# Evolutionary dynamics of temporal niche among tetrapods

**DOI:** 10.64898/2026.03.25.714280

**Authors:** Jacinta Guirguis, Jhoann Canto-Hernández, Catherine Sheard, Daniel Pincheira-Donoso

## Abstract

The diversification of biodiversity progresses as newly evolving species adapt to occupy available niche space – including the temporal dimension. Throughout the history of life, animal species adapted to occupy specific regions, or all regions (cathemerality), of the day-night temporal spectrum. While adaptive radiation theory is predicted to drive much of the global proliferation of biodiversity, the role that the day-night temporal dimension plays in offering ecological opportunity for lineages to diversify remains fundamentally neglected. Using a dataset spanning 19,940 species from across all four main tetrapod lineages (amphibians, squamates (restricted to lizards), mammals and archosaurs), we perform the very first evolutionarily standardised temporal niche diversification analysis and the first to include substantial number of ectotherms. We examine temporal niche as a source of ecological opportunity for adaptive radiation and contrast how lineages have leveraged ecological opportunity spread across 24h time. Findings revealed that tetrapods most frequently transitioned towards diurnality, suggesting a general opening of ecological opportunity in diurnal niche space since the K-Pg mass extinction. Moreover, amphibians are faster and more flexible than amniotes in temporal niche evolution, supported by a relatively higher speed of temporal niche transition and by spending relatively more in time cathemerality, potentially compensating for limitations in geographic dispersal. This interpretation suggests the day-night temporal niche dimension interacts with the geographic dimension, as it exhibits processes which unfold in parallel to (independent of) the geographic dimension as well as processes which unfold in response to what occurs in the geographic dimension.

## Introduction

Adaptive radiation –the evolutionary emergence of ecological diversity from a single ancestor– is widely regarded as a leading driver of biodiversity proliferation (1). This phenomenon is commonly triggered when newly available niche space opens the ‘ecological opportunity’ for evolving lineages to adapt to novel ecological conditions (2,3). The contribution of adaptive radiation to total modern biodiversity has traditionally been estimated through the quantification of ecologically distinct species, or adaptive polymorphisms within species (3), spread over ecological space (e.g., microhabitat types, trophic resources). Surprisingly, however, the role that the temporal dimension of niche space spanning the 24h diel cycle plays in offering ecological opportunity for lineages to diversify remains fundamentally neglected. Over the day-night temporal niche spectrum, temporally divergent biodiversities are exposed to different ecological and climatic challenges (4–8) which shape functional traits specific for their temporal-specific environmental demands on fitness (9). Therefore, just like ecological space, the temporal niche dimension has the potential to contribute to explain a significant amount of biodiversity diversification driven by adaptive radiation.

The interpretation of biodiversity evolution by adaptive radiation from the perspective of both the spatial and the temporal niche dimensions holds a range of important implications. First, and most importantly, the regular incorporation of temporal niche adds a further dimension over which newly arising species can diversify within the same geographic locations and spatial niche components. That is, divergent natural selection has the potential to partition the temporal niche of different species in the same place and adapted to the same microhabitat type, resulting in lineages overlapping in space but segregated in time. Therefore, the spatial and the temporal niche dimensions can be illustrated as being ‘orthogonal’ to one another, amplifying the total amount of niche volume available for natural selection to trigger adaptive radiations. Second, just like in the spatial dimension of niches species can evolve over a spectrum of niche breadth from ecological specialists to generalists, adaptive radiations undergone over the temporal niche dimension can equally drive the evolution of ‘temporal specialists’ that occupy either diurnal or nocturnal niches (i.e. narrow niche breadth), or ‘temporal generalists’ that occupy the whole spectrum (i.e. cathemeral). Finally, the regular incorporation of the temporal niche dimension is expected to shed light on the relative contribution of time and space to the diversification of lineages. In some regions or lineages, and under certain circumstances, it could be possible that diversification is predominantly spread over the spatial niche dimension, while in others, it may be predominantly spread over the temporal dimensions. However, the role that the day-night temporal niche spectrum contributed to the total ecological opportunity available for lineages to radiate remains largely overlooked, thus making it difficult to appreciate the contributions of both the spatial and the temporal dimensions to biodiversity proliferation.

The global biodiversity of tetrapod vertebrates (amphibians, reptiles, mammals and birds) is particularly well suited to address the contribution of temporal niche as a source of ecological opportunity for adaptive radiation. The tetrapod lineage –consisting of ∼38,000 known species– is not only especially well studied, but also, it is the only lineage in nature to have both endothermic (mammals and birds) and ectothermic (amphibians and reptiles) thermoregulators that, therefore, respond in totally contrasting ways to environmental conditions (10–14). For example, as factors such as environmental temperatures experience drastic transitions through space, natural selection theory predicts that lineages will respond by ‘relocating’ newly emerging species across the temporal niche spectrum. The rationale underlying these hypotheses has been reinforced by studies (15,16) that reveal a macroecological gradients whereby ectotherms show an increasing prevalence of nocturnality towards warmer climates.

Here, using a dataset spanning 19,940 tetrapod species, we provide the first evolutionarily standardised (i.e. accounting for differences in evolutionary scale) analysis of lineage diversification associated with the ecological opportunity offered by the day-night temporal niche dimension, and the first empirical analysis aiming to elucidate variation in the macroevolutionary strategies (i.e. relative transition rates) for exploiting temporal ecological opportunity. To this end, using state-dependent speciation and extinction modelling as well as stochastic character mapping, we examine patterns in speciation rates and in the dynamics of temporal niche transitions. We predict that macroevolutionary strategies differ depending on ecophysiology (i.e. ectothermic versus endothermic lineages) and on whether lineages are amniotic or non-amniotic.

## Methods

### Data

We compiled a database of temporal niche spanning 19,940 species from across all four classes of tetrapod vertebrates – amphibians, reptiles (restricted to lizards), archosaurs (which we use to span both birds and crocodiles) and mammals. These data were obtained from a range of sources, including the scientific literature (field guides, journal articles, books), existing class-specific databases (see details below), as well as field and museum records. For temporal niche (defined as the range of time at which a species is active over the 24h diel spectrum) data, we employed the high-level category system (5), where each species was classed as either diurnal (active during the day hours), nocturnal (active during the night hours), or cathemeral (active throughout the 24h of the day). We excluded crepuscular species due to low sample sizes. Temporal niche data were collated from existing databases for mammals (9,17) (we excluded species for which temporal niche data were imputed) and lizards (16,18). For lizards, these sources of data included both cathemeral and crepuscular species within the same category. Therefore, these species were re-classified into the separate categories of cathemeral and crepuscular, but only cathemeral species were retained for analyses. Global-scale databases for archosaurs and amphibians are not currently available, and thus, bird data were obtained primarily from the Handbook of the Birds of the World (19), crocodile data were collected from the literature, while amphibian data were collated as part of the Global Amphibian Biodiversity Project (GABiP, (20,21)).

Finally, the taxonomic nomenclature for amphibians follow Frost (22), for lizards follow Uetz et al. (23), for birds it follows version 8.1 (January 2024) of the HBW Birdlife taxonomic checklist, maintained by Birdlife International, and for mammals it follows version 1.12.1 of the Mammal Diversity Database species checklist (24). Mismatches in names between our data sources and these taxonomies were individually reconciled. All data used in this study are available as supporting information (Dataset 1).

### Phylogenetic trees

For all analyses, we used consensus trees. For amphibians, we used the consensus tree available from Jetz and Pyron (25). For lizards, mammals and birds, to obtain consensus trees we summarised posterior distributions of time-calibrated trees available from Tonini et al. (26), the Phylacine 1.2 database (27), and the Birdtree database (28), respectively, using median node heights with the TreeAnnotator app of Beast 2.5 (29). Our bird phylogeny used the Hackett backbone. For archosaurs, we combined the phylogenies of crocodiles (obtained from TimeTree of Life, version 5, www.timetree.org) (30) with the phylogeny of birds, using an order-level phylogeny of vertebrates (also obtained from TimeTree of Life, version 5) as a back-bone.

### State-dependent diversification analyses

We follow the state-dependent speciation and extinction framework, which improves the estimation of rates of trait evolution by incorporating heterogeneity in diversification rates (speciation − extinction) (31,32), while accounting for the scenario whereby diversification rates may be linked with the evolving trait (33). Under this framework, the speciation rate (*λ_i_*) or extinction rate (*μ_i_*) of a lineage depends on its trait *i*. Using the R package *secsse* (several examined-states dependent speciation and extinction) version 3.1.0 (31,32), we estimate lineage diversification rates and trait transition rates using maximum likelihood optimisation with the function ‘secsse_ml()’, and then infer the temporal niche of ancestors (phylogenetic nodes) using those estimated parameters with the function ‘secsse_loglik()’.

When estimating diversification rates, we accounted for the possibility that they may depend on trait states. Thus, we model scenarios where diversification rates depended on temporal niche (examined-trait dependent models, ETD hereafter) as well as models in which diversification rates depend on a concealed trait that is independent of temporal niche (concealed-trait dependent models, CTD hereafter) (31,32). Because differences in diversification rates could be due to either differential speciation or differential extinction, we ran both ETD and CTD models under two different diversification scenarios. Firstly, in the Varying Speciation, we assume speciation rates differed by each trait state, while extinction rate held constant across all lineages. Secondly, the Varying Speciation and Extinction scenarios, we allowed both speciation and extinction rates to differ by trait state. We did not model scenarios where extinction rates varied with speciation rates held constant due to numerical difficulties and long running times in optimising these models.

When estimating evolutionary (or transition) rates, we ensured all analyses incorporated the same evolutionary model of trait *i*. Thus, we assume the evolution of the trait *i* is bi-directional and passes through an intermediate state. In other words, for all ETD models, we assume that transitions must occur from nocturnality to diurnality via cathemerality, and vice versa (16,34). Similarly, for all CTD models, we assume the same evolutionary transition path, and also assume that the number of states in the concealed trait equals the number of states of our examined trait (three, in our case). In all cases, we assumed all transitions rates differed from one another, and disallowed simultaneous transitions between multiple states as they rarely occur (i.e., at a given point, a temporal niche transition could occur, or a concealed trait transition could occur, but both could not occur simultaneously) (35).

We accounted for unobserved lineages by specifying the sampling fraction using our chosen taxonomies (36). We did not include missing tips encoded as NA in our analyses. This is because *secsse* assigns equal probabilities of being in each state to lineages coded as NA, which, in our case, is biologically unrealistic because temporal niche exhibits phylogenetic signal. Thus, unobserved lineages may be more likely to share the temporal niche of their closest relative included in the analysis than a different temporal niche.

We divided our dataset into amphibians, lizards, mammals and archosaurs. We note that our analyses of lizards do not represent a monophyletic clade as snakes are omitted. Thus, our parameter estimates for this group are biased to varying degrees. This bias is likely strongest at the nodes of the closest relatives of snakes, as ancestral states are heavily influenced by the localised characteristics of the phylogeny (i.e., the branch lengths and topology near the node), which are easily and drastically altered by missing lineages. However, evolutionary rates, which are estimated using information obtained across the entire phylogeny are likely to be relatively less biased.

To ensure the global minima were found during optimisation, we ran each model seven times, with each using a different set of starting points. In the first set, we used estimates of speciation and extinction rates from a birth-death model fit to the branching times of of the phylogeny, with transition rates assumed to be half of the speciation rate. We obtained the second and third sets by multiplying all parameters from the first set by 1.1 and 0.9, respectively. For the fourth set, we multiplied the speciation and extinction rates of the first set by 1.1, while the transition rates were multiplied by 0.9. For the fifth set, the extinction rate was multiplied by 1.1, while the speciation and transition rates of the first set were multiplied by 0.9. For the sixth set, the extinction and transition rates were multiplied by 1.1, while the speciation rate was multiplied by 0.9. Finally, for the seventh set, the speciation and transition rates were multiplied by 1.1, while the extinction rate was multiplied by 0.9. The factors 1.1 and 0.9 were chosen larger perturbations (factors of 2.0 and 0.5) resulted more often in optimisation instability, with the algorithm exploring unrealistic parameter spaces where the likelihood surface was relatively flat, resulting in infeasible computation times. With these starting point sets, only one model (archosaurs, CTD Varying Speciation and Extinction using starting point set 3), was excluded for numerical instability, as invalid mathematical operations were performed by the sampler preventing optimisation.

Using the ‘aicw()’ function from the R package *geiger* **(37)**, we performed model comparison with AIC weights (which penalise models with high numbers of free parameters) to select the best model, and inferred the ancestral states using the function ‘secsse_loglik()’. We only consider nodes that have ‘strong support’ of a particular temporal niche. We define ‘strong support’ as having a probability of at least 0.67, or twice as likely as the second most probably temporal niche (34). Using this threshold, we retained 18,188 out of a total of 19,820 nodes (91.8%) for ancestral state inference.

### Stochastic character mapping

State-dependent models such as *secsse* estimate instantaneous transition rates, which are generally defined as ‘propensities’ for an event to occur, given a specific configuration of the system. In the context of evolution, it is the rate at which evolution could *potentially* progress at, provided certain circumstances (defined by the evolutionary model) were met. More specifically, this means that the propensity to evolve towards another temporal niche (e.g., diurnal) depends on the lineage being in a temporal niche state in which evolution towards that temporal niche is possible (e.g., cathemeral). Critically, this means that transition rates estimated by *secsse* do not represent the realised evolutionary transitions that actually occurred throughout the lineage’s evolutionary history. Consequently, to obtain a comprehensive overview of the macroevolutionary strategies of tetrapods, we additionally estimate the realised evolutionary transition rates by performing stochastic character mapping on the best-supported evolutionary model estimated by *secsse*. We used the function make.simmap() from the R package *phytools* version 2.4.4 (38) to run 1,000 simulations for each tetrapod group, and report the average frequencies of transition, as well as the time spent in each state.

### Standardisation of transition rates

We examine two types of transition rates –instantaneous rates in continuous-time (from *secsse*) and frequencies or discrete-time rates (from *phytools*). To compare transition rates across tetrapods, we standardised them to account for evolutionary scale (continuous-time) and crown age (discrete-time).

For the continuous-time rates, we achieve standardisation by applying equations and terminology from the Gillespie algorithm (i.e., Stochastic Simulation Algorithm) on reaction kinetics (39). This standardisation accounts for differences in evolutionary scale among transition rates (i.e., that baseline rates of evolution may be faster or slower depending on the lineage) thus enabling comparisons of specific transitions between tetrapod groups. To achieve this, we first we calculated the continuous-time rate sum by summing all possible estimated transition rates (nocturnal to cathemeral; cathemeral to diurnal; diurnal to cathemeral; and cathemeral to nocturnal) for each tetrapod group. In the context of our analyses, we define the rate sum as ‘the propensity of a lineage to transition to any temporal niche’. Second, we used the rate sum to calculate the distribution of rates. For this, we divided each individual transition rate by the rate sum. In the context of our analyses, we define the distribution of rates as the ‘relative propensity of lineages to transition to undergo a specific temporal niche transition’. Thus, relative propensities are expressed as proportions which together sum to one for each tetrapod group, forming a categorical probability distribution over all possible transitions, and in which transitions with higher propensities are allocated a larger quantity of the probability space. In this way, the distribution of rates reveals how fast trait evolution occurs in one tetrapod group relative to another tetrapod group, while accounting for differences in their baseline evolutionary rates.

For the discrete-time rates, the standardisation is the same mathematically to the standardisation of the continuous-time rates, but because it is applied to the actual, realised transition rates rather than the propensities, it carries a more intuitive biological meaning. This standardisation accounts for differences in crown age across tetrapod groups. First, we calculated the discrete-time rate to transition to any temporal niche, which we define as the realised count of temporal niche transitions per million years. We did this by taking the mean absolute frequency of temporal niche transition across simulations and divided this by the crown age of each tetrapod group. Second, we calculated the discrete-time distribution of frequencies. The distribution of frequencies in discrete time is the same as in the distribution of rates in continuous time, except that it represents transitions that were realised rather than propensities for a transition to occur. Thus, the distribution of frequencies was calculated by dividing the mean frequency of each possible temporal niche transition by the mean total frequency of temporal niche transition across simulations, to obtain a ‘relative realised transition rate’ for each possible temporal niche transition. In the context of our analyses, we define the ‘relative transition rate’ as the proportion of realised transitions to a given temporal niche in each tetrapod group. Note that the relative propensities to transition and relative realised transition rates are decoupled, in that a temporal niche transition with a high propensity to occur does not necessitate a high realised transition rate.

## Results

Tetrapod lineages accumulate heterogeneously across the day-night temporal niche spectrum (Fig 1), consistent with what we observe in their present-day distribution (Fig 2). Macroevolutionary transitions along the day-night temporal niche dimension vary substantially between amniotes and amphibians, but not between endotherms and ectotherms. We observe that amphibians are faster and more flexible with regards to temporal niche evolution, supported by a relatively higher speed of temporal niche transition and by spending relatively more time as cathemerals (creating more opportunities to transition to either diurnality or nocturnality). Nevertheless, we observe consistency across all tetrapods in other elements of temporal niche diversification (speciation and extinction, instability of cathemerality, relatively recent appearance of diurnality).

**Fig 1.**
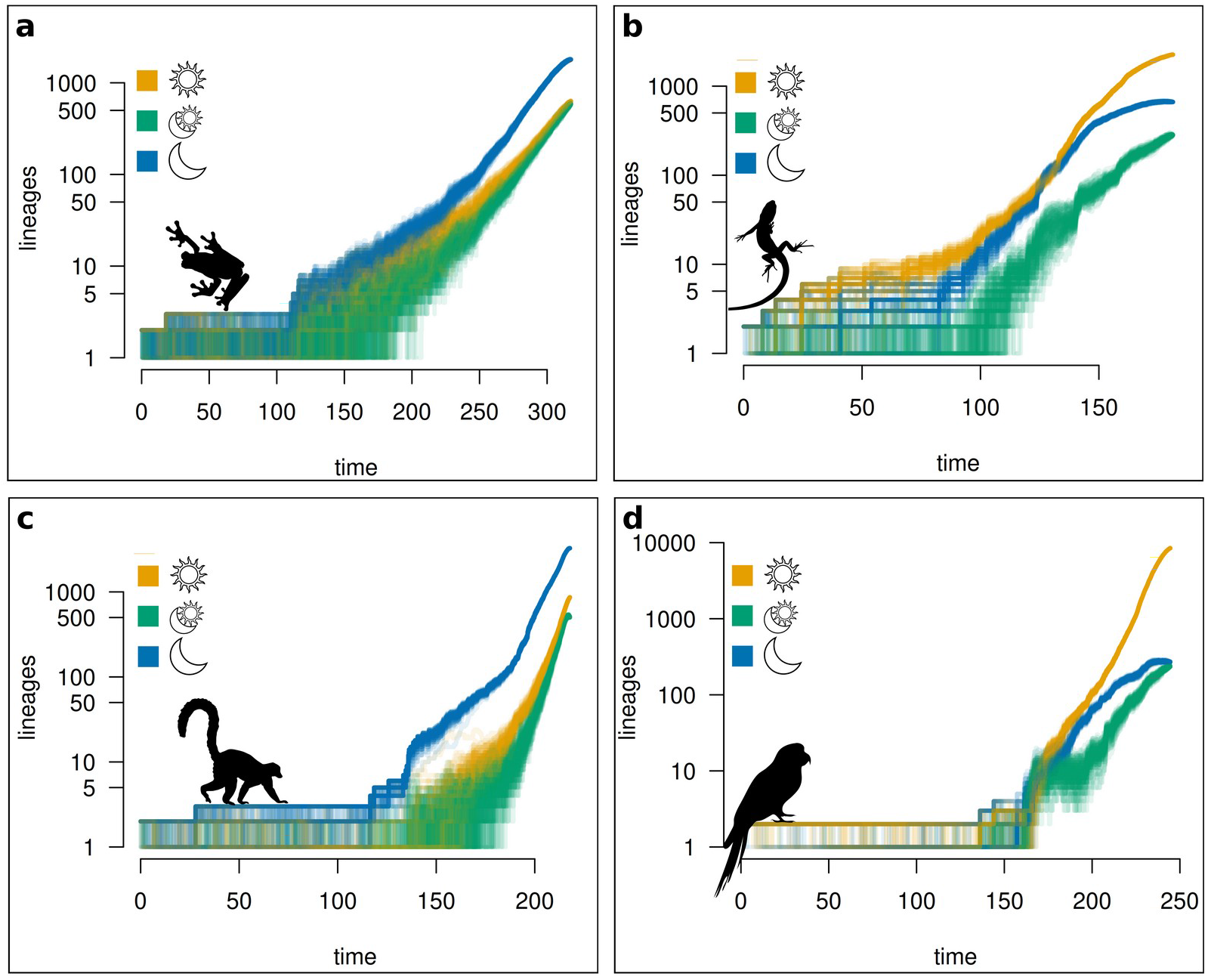
Accumulation of lineages by temporal niche through time. Lineage through time plots showing the number of lineages in each temporal niche (nocturnal, cathemeral and diurnal) from stochastic character maps for (**a**) amphibians, (**b**) lizards, (**c**) mammals and (**d**) archosaurs (birds and crocodiles). Maps were made using the R package *phytool*s on the best supported model derived from the state-dependent speciation and extinction analysis performed using the R package *secsse*, which assumes an ordered model of temporal niche evolution (nocturnal ↔ cathemeral ↔ diurnal). The number of lineages is shown on the *y*-axis while time since the root of each phylogeny is shown on the *x*-axis. For each tetrapod group (**a** – **d**), we generated 1,000 stochastic character maps from which our findings are reported, then randomly selected 100 maps to represent in lineage through time plots. This was to reduce to reduce computational time in making the plots. Silhouette images are of *Agalychnis callidryas* (**a**), *Liolaemus chiliensis* (**b**), *Lemur cata* (**c**) and *Aratinga solstitialis* (**d**), and were obtained from phylopic.org.

**Fig 2.**
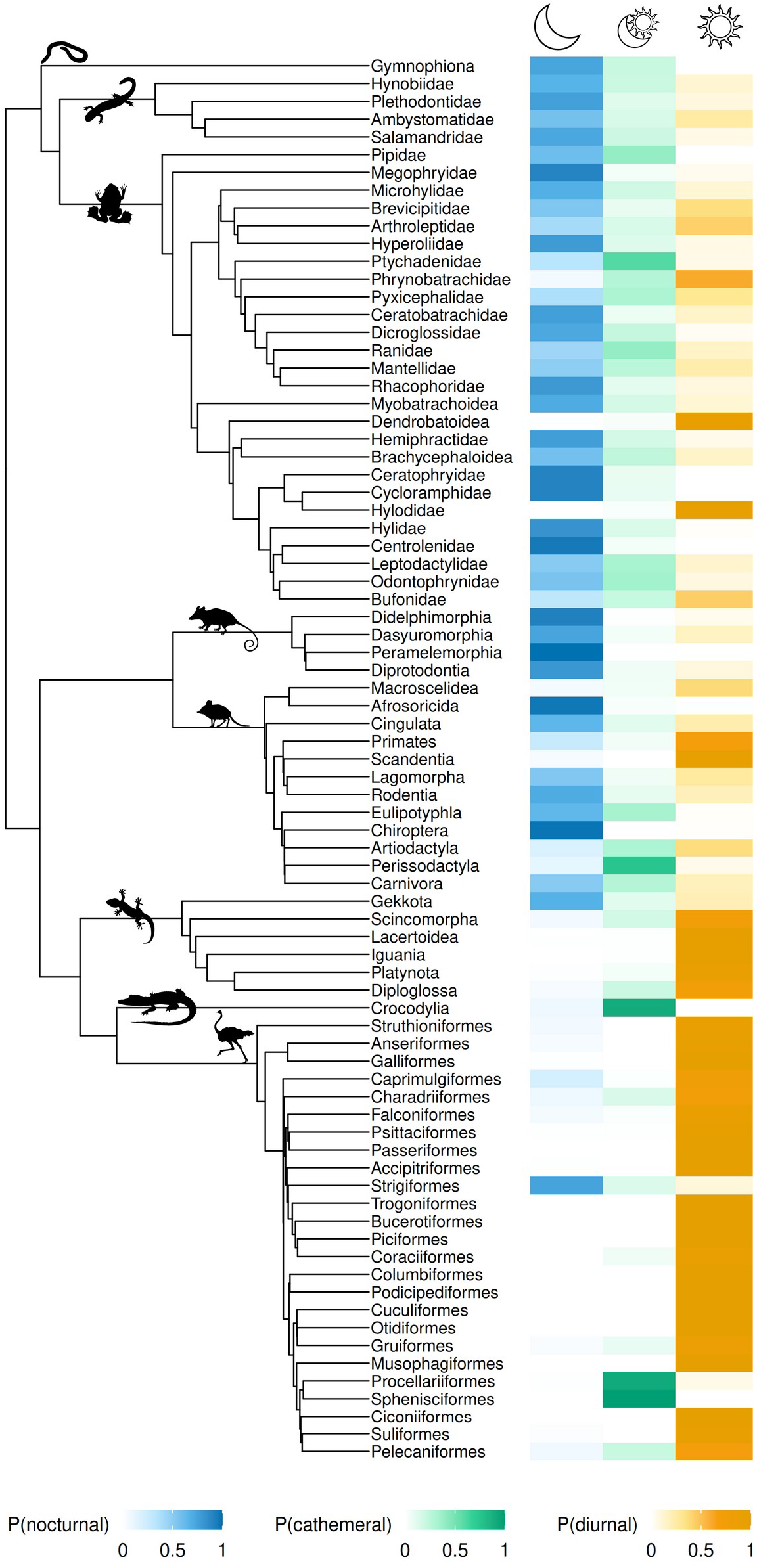
Distribution of temporal niche across the tetrapod tree of life. Phylogram showing the distribution of extant tetrapod lineages among the nocturnal, cathemeral and diurnal temporal niches. Colour bars represent the proportion of extant lineages in each temporal niche. To make the phylogram, consensus trees were obtained (see Methods). Using the R package *ape* and median divergence times obtained from timetree.org, consensus trees were combined into a tetrapod tree, maintaining a single tip per taxonomic group. For amphibians, tip labels primarily refer to family, but my also refer to superfamily (Brachycephaloidea, Dendrobatoidea and Myobatrachoidea) and order (Gymnophiona). For lizards, tip labels refer to superfamily. For mammals and archosaurs (birds and crocodiles), tip labels refer to order. Tips with less than 10 species were removed. Silhouette images shown on root edges of major tetrapod clades represent a species selected from the earliest diverged descendant lineage within that clade. These species are (from the top silhouette image) *Dermophis mexicanus* or the Mexican burrowing caecilian (Gymnophiona), *Onychodactylus fischeri* or Fischer’s clawed salamander (Hynobiidae), *Pipa pipa* or the Suriname toad (Pipidae), *Marmosops* spp or the Neotropical opossums (Didelphimorphia), *Petrodromus tetradactylus* or the four-toed elephant shrew (Macroscelidea), *Gekko gecko* or the tokay gecko (Gekkota), *Crocodylus moreletii* or Morelet’s crocodile (Crocodylia), and *Struthio camelus* or the common ostrich (Struthioniformes). Silhouette images were obtained from phylopic.org.

### Speciation and extinction rates

Our analyses fail to identify a role for temporal niche evolution in influencing rates of lineage diversification among tetrapods, as the CTD Varying Speciation model is always best supported (Table S1). Instead, speciation rates are heterogeneous within each tetrapod group, but in a way which is unlinked to temporal niche, while extinction rates are consistently close to zero (<0.001) (Table S2).

### Relative speed of temporal niche transitions

Tetrapod lineages transition at different rates through the day-night temporal niche, with amphibians transitioning faster than the three amniote groups. This expresses as higher propensity to transition to any temporal niche as well as a higher frequency of realised transitions to any temporal niche. Specifically, the propensity of amphibians to move between temporal niches is 2.52, 1.82 and 2.16 times as high as that of lizards, mammals and archosaurs, respectively (Table 1). Similarly, we found the realised transition rate of amphibians to be 3.75, 3.08 and 7.8 times as high as that of lizards, mammals and archosaurs, respectively (Table 1).

**Table 1.** Standardised temporal niche transition rates of tetrapods. Standardised results from state-dependent speciation and extinction models (SSE) and stochastic character maps. (See Table S2 for raw rates). Table shows results for total transitions (i.e. transitions to any temporal niche), relative propensities (i.e. the propensity of a lineage to transition to a particular temporal niche, if in the state that allows for it), the proportion of realised transitions, as well as the time spent in each temporal niche. Relative propensities are expressed as proportions which together sum to one for each tetrapod group, forming a categorical probability distribution over all possible transitions, and in which transitions with higher propensities are allocated a larger quantity of the probability space. In this way, the distribution of rates reveals how fast trait evolution occurs in one tetrapod group relative to another tetrapod group while accounting for differences in baseline evolutionary rates. ‘Towards diurnality’ refer to the sum of values ‘nocturnal to cathemeral’ and ‘cathemeral to diurnal’. ‘Towards nocturnality’ refer to the sum of values for ‘diurnal to cathemeral’ and ‘cathemeral to nocturnal’. ‘Away from cathemerality’ refer to the sum of values for ‘cathemeral to nocturnal’ and ‘cathemeral to diurnal’.

| Metric | Amphibians | Lizards | Mammals | Archosaurs |
| --- | --- | --- | --- | --- |
| **1. All temporal niche transitions** |  |  |  |  |
| Crown age | 317.6 | 181.3 | 217.8 | 244.5 |
| Total propensity | 0.300 | 0.119 | 0.165 | 0.139 |
| Total transitions | 6207.8 | 945.7 | 1383 | 612.9 |
| Total transitions per million years | 19.543 | 5.217 | 6.349 | 2.507 |
| **2. Diurnality transitions** |  |  |  |  |
| Relative propensity towards diurnality | 0.360 | 0.714 | 0.733 | 0.799 |
| Proportion transitions towards diurnality | 0.516 | 0.605 | 0.558 | 0.703 |
| **3. Nocturnality transitions** |  |  |  |  |
| Relative propensity towards nocturnality | 0.640 | 0.286 | 0.267 | 0.201 |
| Proportion transitions towards nocturnality | 0.484 | 0.395 | 0.442 | 0.297 |
| **4. Cathemerality transitions** |  |  |  |  |
| Relative propensity away from cathemerality | 0.720 | 0.857 | 0.758 | 0.863 |
| Proportion transitions away from cathemerality | 0.493 | 0.516 | 0.527 | 0.594 |
| **5. Time spent in each temporal niche** |  |  |  |  |
| Proportion total time spent in nocturnal | 0.613 | 0.282 | 0.765 | 0.08 |
| Proportion total time spent in cathemeral | 0.178 | 0.077 | 0.092 | 0.043 |
| Proportion total time spent in diurnal | 0.209 | 0.641 | 0.143 | 0.877 |

### Role of cathemerality

Our findings indicate that cathemerality is an unstable state across tetrapods. The propensity to transition away from cathemerality (i.e., the summed propensities to transition from cathemeral to diurnal and cathemeral to nocturnal) is consistently high, with lizards and birds having slightly higher relative propensities (0.86 each) than amphibians and mammals (0.72 and 0.76, respectively) (Table 1). Moreover, in each tetrapod group, approximately half of all realised transitions go away from cathemerality (amphibians 0.49; lizards 0.52; mammals 0.53; archosaurs 0.59) (Table 1). The proportions of realised transitions away from cathemerality are all lower than their respective propensities because time spent in cathemerality is smallest across all tetrapods when compared to time spent in diurnality or nocturnality (Table 1). Thus, tetrapods spend most of their time away from the state of cathemerality, and when they enter it, they leave it relatively rapidly (Table 1).

### Transitions towards diurnality (away from nocturnality)

When considering transitions towards diurnality (or away from nocturnality) (i.e., the summed propensities to transition from nocturnal to cathemeral and from cathemeral to diurnal), two different patterns emerge depending on whether relative propensities or realised transitions are examined. Firstly, our findings reveal that amniotes exhibit propensities to transition towards diurnality (or away from nocturnality) that are roughly twice as high as that of amphibians (Table 1). Secondly, and conversely, the differences observed between amniotes and amphibians are slight when considering the proportions of realised transitions towards diurnality (or away from nocturnality) (Table 1). This discrepancy between the propensities and realised transitions may be attributed to the observation that amphibians spend more time in cathemerality (17.8% of total time) relative to that of amniotes (lizards 7.7%; mammals 9.2%; archosaurs 4.3%) (Table 1). Thus, amphibians compensate for their low propensity to transition towards diurnality by remaining longer in the state in which transition to diurnality is possible.

### Temporal niche of ancestral lineages

Across all taxonomic groups, our findings reveal that diurnality appeared most recently (see below).

Among amphibians, from their nocturnal origins 317 MYA (Pr = 0.768), strong support for cathemerality is first observed 144.3 MYA (Pr = 0.673), while strong support for diurnality is first observed 110.1 MYA (Pr = 0.844). The descendants of the first cathemeral lineage are primarily found in Eurasia and include the primitive frog families Alytidae (painted or midwife frogs, also found in North Africa) and Bombinatoridae (fire-bellied toads and relatives). The descendants of the first diurnal lineage include frog families found throughout tropical Africa and Asia, including Mantellidae (endemic to Madagascar), Nyctibatrachidae (robust frogs, endemic to India), Ranixalidae (leaping frogs, also endemic to India), Rhacophoridea (shrub frogs, found through tropical Africa and Asia), Dicroglossidae (fork-tongued frogs, also found throughout subtropical Africa and Asia), Ceratobatrachidae (ground frogs, found throughout the Malay archipelago), as well as the globally distributed family Ranidae (true frogs).

Among lizards, from their nocturnal origins 181.3 MYA (Pr = 0.997), strong support for cathemerality is first observed 83.8 MYA (Pr = 0.699), while strong support for diurnality was first observed 68.8 MYA (Pr = 0.992). The descendants of the first cathemeral lineage include lacertoidean families primarily found in Central and South America. Specifically, these families are Alopoglossidae (shade lizards), Gymnophthalmidae (spectacled lizards/microteiids), as well as Teiidae (teiids, which include whiptails, tegus, and relatives and are also found in North America). The descendants of the first diurnal lineage are of Phyllodactylidae, a family of geckos (leaf-toed geckos) which are distributed throughout the Americas, Africa, the Middle East and Europe. We note that none of these nodes are among the sister clade of snakes, which are not included in this study.

Among mammals, from their nocturnal origins 217.8 MYA (Pr = 0.988), strong support for cathemerality is first observed 58.8 MYA (Pr = 0.830), while strong support for diurnality is first observed 42.0 MYA (Pr = 0.997). The descendants of the first cathemeral and diurnal lineages include rodents. Specifically, the descendants of the first cathemeral lineage include the families Castoridae (beavers), Geomydae (pocket gophers) and Heteromydiae (kangaroo rats and relatives), while the descendants of the first diurnal lineage include the Central and South American families of Echimyidae (spiny rats), Caviidae (guinea pigs and cavies), Octodontidae (degus and octodonts), Dasyproctidae (agoutis and acouchis), Erethizontidae (New World porcupines, also found in North America), Cuniculidae (pacas), Ctenomyidae (tuco-tucos) and Dinomyidae (the pacarana).

Among archosaurs, from their cathemeral origins 244.5 MYA (Pr = 0.720), we observed apparent evolutionary stasis until after the diversification of birds. From the cathemeral origin of birds 108.4 MYA (Pr = >0.999), strong support for nocturnality is first observed 83.3 MYA (Pr = 0.992), within the Neoaves. The Neoaves include all living birds except Galliformes (gamebirds), Anseriformes (waterfowl) and Struthioniformes (the ratites, which include ostriches, emus, rheas, cassowaries, tinamous and kiwis). Strong support for diurnality is first observed 77.3 MYA (Pr = 0.850), within the Psittacopasserae, which includes both Passeriformes (perching birds) and Psittaciformes (parrots), which we note is a group nested within the Neoaves.

## Discussion

Our study provides the most comprehensive global-scale analysis of evolutionary diversification of tetrapods across the temporal niche dimension (24h diel cycle), which also incorporates the strength of being the first to incorporate a wide proportion of the global diversity of ectotherm lineages. Our study elucidates how tetrapod lineages have undergone lineage diversification from their origins to their present-day by exploiting the ‘temporal ecological opportunity’ that the volume of total niche space (i.e. through ecological space and through the 24h diel cycle) has offered during their phylogenetic histories. We tested the prediction that diversification of temporal niche differed among tetrapods, depending on ecophysiology or whether lineages are amniotic or non-amniotic. Our findings reveal that whether lineages are amniotic or non-amniotic (rather than ecophysiology) plays an important role in influencing the macroevolutionary strategy for diversification throughout the day-night spectrum. Specifically, amphibians are faster and more flexible than amniotes with regards to temporal niche evolution, supported by a relatively higher speed of temporal niche transition and by spending relatively more in time cathemerality.

From across tetrapods, amphibians exhibit the most significant evolutionarily limitations to their ability to disperse due to physiological constraints (40). Our findings suggest that amphibians may compensate for their limitations in geographic dispersal by altering their temporal activity –on ecological timescales through the prevalence of cathemerality throughout their evolutionary history and also on evolutionary timescales through their rapid rates of temporal niche transitions. Our use of relative propensities means that these differences remain even after accounting for the possibility that amphibians may evolve faster in general. Thus, our findings may suggest a hypothesis explaining why the diversification of amphibians throughout the day-night spectrum may have contributed to their persistence as a lineage, and may contribute to their survival throughout the current mass extinction.

### What can be inferred by ancestral reconstructions given the limitations of method and theory?

Early tetrapods are suggested to have undergone an extended period of temporal niche conservatism, with phylogenetic reconstructions indicating restriction to nocturnal niches for more than 300 million years (41). Factors involved in the prolonged period of nocturnality are hypothesised to differ across tetrapod groups. For instance, permeable skin in the earliest tetrapods (amphibians) may have restricted them to nighttime activity due to the risk of desiccation from sun exposure (41,42). In contrast, a classic ‘nocturnal bottleneck’ (34,43,44) hypothesis on the prolonged period of nocturnality among mammals until after the Cretaceous-Palaeogene (K-Pg hereafter) mass extinction posits that such prolonged nocturnality was caused by predation by non-avian dinosaurs (45), which may have dominated diurnal niche space (34,43,44).

Despite their appeal, hypotheses on prolonged periods of nocturnality among early tetrapods present fundamental challenges. The nocturnal bottleneck of mammals pre-dates the K-Pg mass extinction, while nocturnality conservatism among early tetrapods encompasses a time during which three mass extinctions occurred (34,41). Furthermore, while non-avian dinosaurs were thought to be largely diurnal (34), this remains a phenomenon to be supported by a sufficiently substantial body of empirical evidence that at present is still lacking. Thus, while our findings are consistent with these extended periods of temporal niche conservatism, we reiterate previous limitations with absolute inferences of reconstructed trait states of lineages in deep time (46). We do not suggest that diurnality in tetrapods did not exist prior to when we first observed it in our reconstructions, but instead propose that findings on transition rates (which are less sensitive to bias from missing lineages) may more reliably reflect macroevolutionary processes since the K-Pg mass extinction.

## Conclusions

Patterns of diversification are traditionally interpreted across geographic space, yet this perspective overlooks how lineages are partitioned by natural selection into daytime and nighttime assemblages. Our results demonstrate that, at least among tetrapods, the temporal niche dimension constitutes a complementary axis of diversification alongside geographic space. Amphibians show markedly faster realised transitions among temporal niches than amniotes (3-8x higher rates per million years), while also spending disproportionately more time in cathemerality. Combined with well-established constraints on amphibian dispersal (40), this supports a compensatory mechanism whereby rapid temporal niche evolution can offset geographic limitations. In contrast, birds exhibit strong diurnal conservatism, spending nearly 88% of their evolutionary history in diurnality, highlighting divergent macroevolutionary strategies along the day-night spectrum. Rather than viewing space and time as separate arenas, our findings support a model in which temporal and geographic dimensions interact, as spatial constraints are offset by temporal leeway, thus shedding light on the relative contributions of the spatial and temporal dimensions to the diversification of biodiversity. To establish the extent of this interaction, future work should investigate whether the relationship between dispersal ability and temporal niche flexibility extends beyond amphibians to other low-dispersal lineages.

## Acknowledgements

JG is indebted to the UK Natural Environment Research Council DTP Quadrat programme for full PhD funding.

