## Supplementary material for "Evolutionary dynamics of temporal niche among tetrapods": Table S1

**Table S1.** Akaike Information Criterion (*AIC*) of all models fitted using the R package *secsse* version 3.1.0. For each taxonomic group (amphibians, lizards, mammals and archosaurs), modelled scenarios where diversification rates depended on temporal niche (examined-trait dependent models, ETD hereafter) as well as scenarios in which diversification rates depended on a concealed trait that is independent of temporal niche (concealed-trait dependent models, CTD hereafter). Because differences in diversification rates could be due to either differential speciation or differential extinction, we ran both ETD and CTD models under two different diversification scenarios. First, in the Varying Speciation scenario, we assume speciation rates differed by each trait state, while extinction rate was held constant across lineages. Second, the Varying Speciation and Extinction scenarios, we allowed both speciation and extinction rates to differ by trait state. To ensure the global minima was found during the optimisation process, we ran each model seven times, with each using a different set of starting points. In the first set, we used estimates of speciation and extinction rates from a birth-death model fit to the branching times of of the phylogeny, with transition rates assumed to be half of the speciation rate. We obtained the second and third sets by multiplying all parameters from the first set by 1.1 and 0.9, respectively. For the fourth set, we multiplied the speciation and extinction rates of the first set by 1.1, while the transition rates were multiplied by 0.9. For the fifth set, the extinction rate was multiplied by 1.1, while the speciation and transition rates of the first set were multiplied by 0.9. For the sixth set, the extinction and transition rates were multiplied by 1.1, while the speciation rate was multiplied by 0.9. Finally, for the seventh set, the speciation and transition rates were multiplied by 1.1, while the extinction rate was multiplied by 0.9. For each tetrapod group, we selected the best model using the function 'aicw()' from the R package *geiger*, which penalised higher numbers of free parameters. The best-supported model is displayed in the first row of the list for each taxon. Table shows log likelihood (*loglik*), the number of free parameters (*k*), *AIC*, change in *AIC* (*AICΔ*) and *AIC* weights (*AICw).*

| Taxon | Model & Starting Point Set | *loglik* | *k* | *AIC* | *AICΔ* | *AICw* |
|---|---|---|---|---|---|---|
| Amphibians | CTD Varying Speciation 1 | -15,004.204 | 12 | 30032.407 | 0.000 | 0.143 |
| | CTD Varying Speciation 4 | -15,004.204 | 12 | 30032.407 | <0.000 | 0.143 |
| | CTD Varying Speciation 3 | -15,004.204 | 12 | 30032.407 | <0.000 | 0.143 |
| | CTD Varying Speciation 2 | -15,004.204 | 12 | 30032.407 | <0.000 | 0.143 |
| | CTD Varying Speciation 5 | -15,004.204 | 12 | 30032.408 | <0.000 | 0.143 |
| | CTD Varying Speciation 7 | -15,004.204 | 12 | 30032.408 | 0.001 | 0.143 |
| | CTD Varying Speciation & Extinction 7 | -15,004.012 | 14 | 30036.024 | 3.616 | 0.023 |
| | CTD Varying Speciation & Extinction 1 | -15,004.012 | 14 | 30036.024 | 3.617 | 0.023 |
| | CTD Varying Speciation & Extinction 2 | -15,004.013 | 14 | 30036.025 | 3.618 | 0.023 |
| | CTD Varying Speciation & Extinction 5 | -15,004.013 | 14 | 30036.026 | 3.619 | 0.023 |
| | CTD Varying Speciation & Extinction 3 | -15,004.013 | 14 | 30036.026 | 3.619 | 0.023 |
| | CTD Varying Speciation & Extinction 4 | -15,004.013 | 14 | 30036.027 | 3.619 | 0.023 |
| | CTD Varying Speciation 6 | -15,018.729 | 12 | 30061.458 | 29.050 | <0.000 |
| | CTD Varying Speciation & Extinction 6 | -15,040.992 | 14 | 30109.985 | 77.578 | <0.000 |
| | ETD Varying Speciation & Extinction 7 | -15,205.531 | 14 | 30439.062 | 406.655 | <0.000 |
| | ETD Varying Speciation & Extinction 2 | -15,205.531 | 14 | 30439.063 | 406.655 | <0.000 |
| | ETD Varying Speciation 5 | -15,224.912 | 12 | 30473.825 | 441.418 | <0.000 |
| | ETD Varying Speciation 3 | -15,224.913 | 12 | 30473.825 | 441.418 | <0.000 |
| | ETD Varying Speciation 2 | -15,224.913 | 12 | 30473.825 | 441.418 | <0.000 |

| | | | | | | |
|---|---|---|---|---|---|---|
| | ETD Varying Speciation 7 | -15,224.913 | 12 | 30473.825 | 441.418 | <0.000 |
| | ETD Varying Speciation & Extinction 5 | -15,243.787 | 14 | 30515.574 | 483.167 | <0.000 |
| | ETD Varying Speciation & Extinction 3 | -15,243.788 | 14 | 30515.575 | 483.168 | <0.000 |
| | ETD Varying Speciation & Extinction 4 | -15,243.788 | 14 | 30515.575 | 483.168 | <0.000 |
| | ETD Varying Speciation & Extinction 1 | -15,243.795 | 14 | 30515.590 | 483.183 | <0.000 |
| | ETD Varying Speciation & Extinction 6 | -15,243.796 | 14 | 30515.591 | 483.184 | <0.000 |
| | ETD Varying Speciation 1 | -15,290.391 | 12 | 30604.781 | 572.374 | <0.000 |
| | ETD Varying Speciation 6 | -15,290.391 | 12 | 30604.781 | 572.374 | <0.000 |
| | ETD Varying Speciation 4 | -15,290.391 | 12 | 30604.782 | 572.375 | <0.000 |
| Lizards | CTD Varying Speciation 6 | -13,074.338 | 12 | 26172.676 | 0.000 | 0.266 |
| | CTD Varying Speciation 4 | -13,074.338 | 12 | 26172.676 | <0.000 | 0.266 |
| | CTD Varying Speciation 7 | -13,074.338 | 12 | 26172.676 | 0.001 | 0.266 |
| | CTD Varying Speciation & Extinction 6 | -13,073.706 | 14 | 26175.411 | 2.736 | 0.068 |
| | CTD Varying Speciation & Extinction 4 | -13,073.706 | 14 | 26175.411 | 2.736 | 0.068 |
| | CTD Varying Speciation & Extinction 7 | -13,073.706 | 14 | 26175.412 | 2.736 | 0.068 |
| | CTD Varying Speciation 2 | -13,086.808 | 12 | 26197.616 | 24.940 | <0.000 |
| | CTD Varying Speciation 5 | -13,086.809 | 12 | 26197.619 | 24.943 | <0.000 |
| | CTD Varying Speciation 1 | -13,086.810 | 12 | 26197.620 | 24.944 | <0.000 |
| | CTD Varying Speciation & Extinction 5 | -13,086.809 | 14 | 26201.619 | 28.943 | <0.000 |
| | CTD Varying Speciation & Extinction 1 | -13,086.810 | 14 | 26201.619 | 28.944 | <0.000 |
| | CTD Varying Speciation & Extinction 3 | -13,086.814 | 14 | 26201.628 | 28.952 | <0.000 |
| | CTD Varying Speciation 3 | -13,284.976 | 12 | 26593.951 | 421.275 | <0.000 |
| | CTD Varying Speciation & Extinction 2 | -13,284.858 | 14 | 26597.717 | 425.041 | <0.000 |
| | ETD Varying Speciation 5 | -14,453.679 | 12 | 28931.358 | 2758.682 | <0.000 |
| | ETD Varying Speciation 7 | -14,453.679 | 12 | 28931.358 | 2758.682 | <0.000 |
| | ETD Varying Speciation 3 | -14,453.679 | 12 | 28931.358 | 2758.682 | <0.000 |
| | ETD Varying Speciation 2 | -14,453.679 | 12 | 28931.358 | 2758.682 | <0.000 |
| | ETD Varying Speciation 6 | -14,453.679 | 12 | 28931.358 | 2758.682 | <0.000 |
| | ETD Varying Speciation 4 | -14,453.679 | 12 | 28931.358 | 2758.683 | <0.000 |
| | ETD Varying Speciation 1 | -14,453.679 | 12 | 28931.359 | 2758.683 | <0.000 |
| | ETD Varying Speciation & Extinction 1 | -14,453.671 | 14 | 28935.342 | 2762.666 | <0.000 |
| | ETD Varying Speciation & Extinction 7 | -14,453.679 | 14 | 28935.358 | 2762.682 | <0.000 |
| | ETD Varying Speciation & Extinction 2 | -14,453.679 | 14 | 28935.358 | 2762.682 | <0.000 |
| | ETD Varying Speciation & Extinction 3 | -14,453.679 | 14 | 28935.358 | 2762.683 | <0.000 |
| | ETD Varying Speciation & Extinction 5 | -14,453.679 | 14 | 28935.358 | 2762.683 | <0.000 |
| | ETD Varying Speciation & Extinction 4 | -14,453.680 | 14 | 28935.359 | 2762.684 | <0.000 |
| | ETD Varying Speciation & Extinction 6 | -14,453.680 | 14 | 28935.360 | 2762.685 | <0.000 |
| Mammals | CTD Varying Speciation 5 | -15,790.964 | 12 | 31605.928 | 0.000 | 0.288 |
| | CTD Varying Speciation 7 | -15,791.286 | 12 | 31606.572 | 0.644 | 0.209 |
| | CTD Varying Speciation 1 | -15,791.485 | 12 | 31606.969 | 1.042 | 0.171 |
| | CTD Varying Speciation 6 | -15,791.835 | 12 | 31607.669 | 1.742 | 0.121 |
| | CTD Varying Speciation & Extinction 6 | -15,789.921 | 14 | 31607.842 | 1.914 | 0.111 |
| | CTD Varying Speciation 4 | -15,792.444 | 12 | 31608.889 | 2.961 | 0.066 |
| | CTD Varying Speciation & Extinction 5 | -15,791.631 | 14 | 31611.263 | 5.335 | 0.020 |
| | CTD Varying Speciation & Extinction 1 | -15,791.904 | 14 | 31611.807 | 5.880 | 0.015 |
| | CTD Varying Speciation & Extinction 3 | -15,810.897 | 14 | 31649.794 | 43.866 | <0.000 |
| | CTD Varying Speciation 3 | -15,829.464 | 12 | 31682.928 | 77.000 | <0.000 |
| | CTD Varying Speciation 2 | -15,830.209 | 12 | 31684.418 | 78.490 | <0.000 |
| | CTD Varying Speciation & Extinction 7 | -15,851.154 | 14 | 31730.308 | 124.380 | <0.000 |
| | CTD Varying Speciation & Extinction 4 | -15,879.574 | 14 | 31787.149 | 181.221 | <0.000 |
| | CTD Varying Speciation & Extinction 2 | -15,971.869 | 14 | 31971.738 | 365.810 | <0.000 |
| | ETD Varying Speciation 6 | -16,714.662 | 12 | 33453.323 | 1847.395 | <0.000 |
| | ETD Varying Speciation 1 | -16,714.808 | 12 | 33453.617 | 1847.689 | <0.000 |
| | ETD Varying Speciation 7 | -16,715.772 | 12 | 33455.544 | 1849.616 | <0.000 |
| | ETD Varying Speciation 2 | -16,715.925 | 12 | 33455.850 | 1849.922 | <0.000 |
| | ETD Varying Speciation 4 | -16,718.174 | 12 | 33460.348 | 1854.420 | <0.000 |

| | | | | | | |
|---|---|---|---|---|---|---|
| | ETD Varying Speciation 3 | -16,718.585 | 12 | 33461.170 | 1855.243 | <0.000 |
| | ETD Varying Speciation & Extinction 3 | -16,718.544 | 14 | 33465.089 | 1859.161 | <0.000 |
| | ETD Varying Speciation & Extinction 1 | -16,901.612 | 14 | 33831.225 | 2225.297 | <0.000 |
| | ETD Varying Speciation & Extinction 6 | -16,916.160 | 14 | 33860.320 | 2254.392 | <0.000 |
| | ETD Varying Speciation & Extinction 2 | -16,942.908 | 14 | 33913.815 | 2307.887 | <0.000 |
| | ETD Varying Speciation 5 | -16,954.381 | 12 | 33932.762 | 2326.834 | <0.000 |
| | ETD Varying Speciation & Extinction 4 | -16,956.108 | 14 | 33940.215 | 2334.288 | <0.000 |
| | ETD Varying Speciation & Extinction 5 | -16,966.474 | 14 | 33960.949 | 2355.021 | <0.000 |
| | ETD Varying Speciation & Extinction 7 | -16,985.631 | 14 | 33999.262 | 2393.334 | <0.000 |
| Archosaurs | CTD Varying Speciation 4 | -28595.527 | 12 | 57,215.054 | 0.000 | 0.785 |
| | CTD Varying Speciation & Extinction 1 | -28595.511 | 14 | 57,219.023 | 3.968 | 0.108 |
| | CTD Varying Speciation & Extinction 4 | -28595.524 | 14 | 57,219.048 | 3.994 | 0.107 |
| | CTD Varying Speciation 3 | -28637.312 | 12 | 57,298.624 | 83.570 | <0.000 |
| | CTD Varying Speciation 2 | -28637.316 | 12 | 57,298.632 | 83.578 | <0.000 |
| | CTD Varying Speciation 1 | -28637.318 | 12 | 57,298.637 | 83.583 | <0.000 |
| | CTD Varying Speciation 6 | -28637.392 | 12 | 57,298.784 | 83.730 | <0.000 |
| | CTD Varying Speciation & Extinction 6 | -28637.324 | 14 | 57,302.649 | 87.595 | <0.000 |
| | CTD Varying Speciation & Extinction 2 | -28637.333 | 14 | 57,302.665 | 87.611 | <0.000 |
| | CTD Varying Speciation 7 | -28688.643 | 12 | 57,401.285 | 186.231 | <0.000 |
| | CTD Varying Speciation & Extinction 7 | -28687.648 | 14 | 57,403.296 | 188.242 | <0.000 |
| | CTD Varying Speciation 5 | -28798.736 | 12 | 57,621.471 | 406.417 | <0.000 |
| | CTD Varying Speciation & Extinction 5 | -28798.723 | 14 | 57,625.447 | 410.393 | <0.000 |
| | ETD Varying Speciation 6 | -30018.045 | 12 | 60,060.090 | 2,845.036 | <0.000 |
| | ETD Varying Speciation 2 | -30018.049 | 12 | 60,060.098 | 2,845.044 | <0.000 |
| | ETD Varying Speciation 1 | -30018.050 | 12 | 60,060.100 | 2,845.046 | <0.000 |
| | ETD Varying Speciation & Extinction 7 | -30018.050 | 14 | 60,064.100 | 2,849.046 | <0.000 |
| | ETD Varying Speciation 4 | -32040.200 | 12 | 64,104.401 | 6,889.347 | <0.000 |
| | ETD Varying Speciation 5 | -32040.202 | 12 | 64,104.403 | 6,889.349 | <0.000 |
| | ETD Varying Speciation 7 | -32040.203 | 12 | 64,104.406 | 6,889.352 | <0.000 |
| | ETD Varying Speciation 3 | -32040.207 | 12 | 64,104.415 | 6,889.361 | <0.000 |
| | ETD Varying Speciation & Extinction 4 | -32040.202 | 14 | 64,108.404 | 6,893.350 | <0.000 |
| | ETD Varying Speciation & Extinction 2 | -32040.204 | 14 | 64,108.409 | 6,893.355 | <0.000 |
| | ETD Varying Speciation & Extinction 1 | -32040.205 | 14 | 64,108.409 | 6,893.355 | <0.000 |
| | ETD Varying Speciation & Extinction 3 | -32040.206 | 14 | 64,108.412 | 6,893.358 | <0.000 |
| | ETD Varying Speciation & Extinction 5 | -32040.207 | 14 | 64,108.415 | 6,893.361 | <0.000 |
| | ETD Varying Speciation & Extinction 6 | -32040.210 | 14 | 64,108.419 | 6,893.365 | <0.000 |
