## Supplementary material for "Evolutionary dynamics of temporal niche among tetrapods": Table S2

**Table S2.** Estimated parameters for the best-supported model fitted using the R package *secsse* version 3.1.0. For all tetrapod groups (amphibians, lizards, mammals and archosaurs), the 'concealed trait-dependent' (CTD) model with varying speciation only was supported. In this model, rates of diversification varied independently of a lineages' temporal niche. Instead of depending on temporal niche, rates of diversification depended on a concealed trait, which could express as state A, B or C. The speciation rates ($\lambda$) of each concealed state are given below as $\lambda_A$, $\lambda_B$ and $\lambda_C$. The extinction rate ($\mu$) was held constant across all lineages. Lineages evolved from state to state at rate $q$. Our model simultaneously estimated the evolutionary rates of the concealed trait and temporal niche, given below as $q_{A\text{-}B}$, $q_{B\text{-}C}$, $q_{C\text{-}B}$, and $q_{B\text{-}A}$ as well as $q_{Nocturnal\text{-}Cathemeral}$, $q_{Cathemeral\text{-}Diurnal}$, $q_{Diurnal\text{-}Cathemeral}$, and $q_{Cathemeral\text{-}Nocturnal}$, respectively.

| Taxon | $\lambda_A$ | $\lambda_B$ | $\lambda_C$ | $\mu$ | $q_{A\text{-}B}$ | $q_{B\text{-}C}$ | $q_{C\text{-}B}$ | $q_{B\text{-}A}$ | $q_{Nocturnal\text{-}Cathemeral}$ | $q_{Cathemeral\text{-}Diurnal}$ | $q_{Diurnal\text{-}Cathemeral}$ | $q_{Cathemeral\text{-}Nocturnal}$ |
|---|---|---|---|---|---|---|---|---|---|---|---|---|
| Amphibians | 0.111 | 0.052 | 0.014 | <0.001 | 0.041 | 0.013 | 0.003 | 0.003 | 0.019 | 0.017 | 0.009 | 0.055 |
| Lizards | 0.021 | 0.556 | 0.081 | <0.001 | 0.001 | 0.018 | 0.036 | 0.677 | 0.004 | 0.024 | 0.002 | 0.010 |
| Mammals | 0.235 | 0.867 | 0.045 | <0.001 | 0.145 | 1.507 | 0.016 | 0.186 | 0006 | 0.034 | 0.007 | 0.007 |
| Archosaurs | 1.920 | 0.228 | 0.024 | <0.001 | 0.009 | 0.305 | 0.021 | 0.057 | 0.006 | 0.031 | <0.001 | 0.009 |
