## Supplementary material for "Evolutionary dynamics of temporal niche among tetrapods": Dataset 1

| group | scientific_name_updated | phylo_name | activity |
|---|---|---|---|
| amphibians | "Centrolene"_medemi | Centrolene_medemi | nocturnal |
| amphibians | Abavorana_luctuosa | Hylarana_luctuosa | nocturnal |
| amphibians | Acanthixalus_sonjae | Acanthixalus_sonjae | nocturnal |
| amphibians | Acanthixalus_spinosus | Acanthixalus_spinosus | nocturnal |
| amphibians | Acris_blanchardi | Acris_blanchardi | diurnal |
| amphibians | Acris_crepitans | Acris_crepitans | cathemeral |
| amphibians | Acris_gryllus | Acris_gryllus | cathemeral |
| amphibians | Adelastes_hylonomos | Adelastes_hylonomos | nocturnal |
| amphibians | Adelophryne_adiastola | Adelophryne_adiastola | nocturnal |
| amphibians | Adelophryne_baturitensis | Adelophryne_baturitensis | cathemeral |
| amphibians | Adelophryne_gutturosa | Adelophryne_gutturosa | cathemeral |
| amphibians | Adelophryne_maranguapensis | Adelophryne_maranguapensis | diurnal |
| amphibians | Adelophryne_mucronata | Adelophryne_mucronatus | nocturnal |
| amphibians | Adelophryne_pachydactyla | Adelophryne_pachydactyla | nocturnal |
| amphibians | Adelophryne_patamona | Adelophryne_patamona | nocturnal |
| amphibians | Adelphobates_castaneoticus | Adelphobates_castaneoticus | diurnal |
| amphibians | Adelphobates_galactonotus | Adelphobates_galactonotus | diurnal |
| amphibians | Adelphobates_quinquevittatus | Adelphobates_quinquevittatus | diurnal |
| amphibians | Adenomera_ajurauna | Adenomera_ajurauna | nocturnal |
| amphibians | Adenomera_andreae | Adenomera_andreae | cathemeral |
| amphibians | Adenomera_araucaria | Adenomera_araucaria | cathemeral |
| amphibians | Adenomera_bokermanni | Adenomera_bokermanni | nocturnal |
| amphibians | Adenomera_diptyx | Adenomera_diptyx | diurnal |
| amphibians | Adenomera_engelsi | Adenomera_engelsi | cathemeral |
| amphibians | Adenomera_heyeri | Adenomera_heyeri | diurnal |
| amphibians | Adenomera_hylaedactyla | Adenomera_hylaedactyla | cathemeral |
| amphibians | Adenomera_lutzi | Adenomera_lutzi | cathemeral |
| amphibians | Adenomera_marmorata | Adenomera_marmorata | cathemeral |
| amphibians | Adenomera_martinezi | Adenomera_martinezi | diurnal |
| amphibians | Adenomera_nana | Adenomera_nana | nocturnal |
| amphibians | Adenomera_thomei | Adenomera_thomei | nocturnal |
| amphibians | Adenomus_kandianus | Adenomus_kandianus | nocturnal |
| amphibians | Adenomus_kelaartii | Adenomus_kelaartii | cathemeral |

| | | | |
|---|---|---|---|
| amphibians | Afrixalus_aureus | Afrixalus_aureus | cathemeral |
| amphibians | Afrixalus_clarkei | Afrixalus_clarkei | nocturnal |
| amphibians | Afrixalus_delicatus | Afrixalus_delicatus | nocturnal |
| amphibians | Afrixalus_dorsalis | Afrixalus_dorsalis | nocturnal |
| amphibians | Afrixalus_dorsimaculatus | Afrixalus_dorsimaculatus | diurnal |
| amphibians | Afrixalus_equatorialis | Afrixalus_equatorialis | nocturnal |
| amphibians | Afrixalus_fornasini | Afrixalus_fornasini | nocturnal |
| amphibians | Afrixalus_lacteus | Afrixalus_lacteus | nocturnal |
| amphibians | Afrixalus_nigeriensis | Afrixalus_nigeriensis | nocturnal |
| amphibians | Afrixalus_paradorsalis | Afrixalus_paradorsalis | nocturnal |
| amphibians | Afrixalus_quadrivittatus | Afrixalus_quadrivittatus | nocturnal |
| amphibians | Afrixalus_uluguruensis | Afrixalus_uluguruensis | diurnal |
| amphibians | Afrixalus_vibekensis | Afrixalus_vibekensis | nocturnal |
| amphibians | Agalychnis_annae | Agalychnis_annae | nocturnal |
| amphibians | Agalychnis_buckleyi | Hylomantis_buckleyi | nocturnal |
| amphibians | Agalychnis_callidryas | Agalychnis_callidryas | nocturnal |
| amphibians | Agalychnis_dacnicolor | Pachymedusa_dacnicolor | nocturnal |
| amphibians | Agalychnis_lemur | Agalychnis_lemur | nocturnal |
| amphibians | Agalychnis_moreletii | Agalychnis_moreletii | nocturnal |
| amphibians | Agalychnis_psilopygion | Hylomantis_psilopygion | nocturnal |
| amphibians | Agalychnis_saltator | Agalychnis_saltator | cathemeral |
| amphibians | Agalychnis_spurrelli | Agalychnis_spurrelli | nocturnal |
| amphibians | Aglyptodactylus_madagascariensis | Aglyptodactylus_madagascariensis | cathemeral |
| amphibians | Aglyptodactylus_securifer | Aglyptodactylus_securifer | cathemeral |
| amphibians | Allobates_algorei | Allobates_algorei | cathemeral |
| amphibians | Allobates_brunneus | Allobates_brunneus | diurnal |
| amphibians | Allobates_caeruleodactylus | Allobates_caeruleodactylus | crepuscular |
| amphibians | Allobates_chalcopis | Allobates_chalcopis | crepuscular |
| amphibians | Allobates_femoralis | Allobates_femoralis | diurnal |
| amphibians | Allobates_granti | Allobates_granti | diurnal |
| amphibians | Allobates_hodli | Allobates_hodli | nocturnal |
| amphibians | Allobates_insperatus | Allobates_insperatus | diurnal |
| amphibians | Allobates_juanii | Allobates_juanii | diurnal |
| amphibians | Allobates_kingsburyi | Allobates_kingsburyi | diurnal |

| | | | |
|---|---|---|---|
| amphibians | Allobates_marchesianus | Allobates_marchesianus | cathemeral |
| amphibians | Allobates_mcdiarmidi | Allobates_mcdiarmidi | diurnal |
| amphibians | Allobates_melanolaemus | Allobates_melanolaemus | diurnal |
| amphibians | Allobates_myersi | Allobates_myersi | diurnal |
| amphibians | Allobates_nidicola | Allobates_nidicola | crepuscular |
| amphibians | Allobates_niputidea | Allobates_niputidea | diurnal |
| amphibians | Allobates_olfersioides | Allobates_olfersioides | diurnal |
| amphibians | Allobates_ornatus | Allobates_ornatus | diurnal |
| amphibians | Allobates_paleovarzensis | Allobates_paleovarzensis | crepuscular |
| amphibians | Allobates_subfolionidificans | Allobates_subfolionidificans | diurnal |
| amphibians | Allobates_talamancae | Allobates_talamancae | diurnal |
| amphibians | Allobates_trilineatus | Allobates_trilineatus | diurnal |
| amphibians | Allobates_undulatus | Allobates_undulatus | diurnal |
| amphibians | Allobates_wayuu | Allobates_wayuu | crepuscular |
| amphibians | Allobates_zaparo | Allobates_zaparo | diurnal |
| amphibians | Allophryne_relicta | Allophryne_relicta | nocturnal |
| amphibians | Allophryne_ruthveni | Allophryne_ruthveni | cathemeral |
| amphibians | Alsodes_vanzolinii | Alsodes_vanzolinii | nocturnal |
| amphibians | Altiphrynoides_malcolmi | Altiphrynoides_malcolmi | diurnal |
| amphibians | Alytes_cisternasii | Alytes_cisternasii | nocturnal |
| amphibians | Alytes_dickhilleni | Alytes_dickhilleni | nocturnal |
| amphibians | Alytes_maurus | Alytes_maurus | cathemeral |
| amphibians | Alytes_muletensis | Alytes_muletensis | nocturnal |
| amphibians | Alytes_obstetricans | Alytes_obstetricans | nocturnal |
| amphibians | Amazophrynella_minuta | Amazophrynella_minuta | cathemeral |
| amphibians | Ambystoma_annulatum | Ambystoma_annulatum | nocturnal |
| amphibians | Ambystoma_californiense | Ambystoma_californiense | nocturnal |
| amphibians | Ambystoma_cingulatum | Ambystoma_cingulatum | diurnal |
| amphibians | Ambystoma_gracile | Ambystoma_gracile | cathemeral |
| amphibians | Ambystoma_jeffersonianum | Ambystoma_jeffersonianum | diurnal |
| amphibians | Ambystoma_laterale | Ambystoma_laterale | nocturnal |
| amphibians | Ambystoma_mabeei | Ambystoma_mabeei | diurnal |
| amphibians | Ambystoma_macrodactylum | Ambystoma_macrodactylum | nocturnal |
| amphibians | Ambystoma_maculatum | Ambystoma_maculatum | cathemeral |

| | | | |
|---|---|---|---|
| amphibians | Ambystoma_mavortium | Ambystoma_mavortium | nocturnal |
| amphibians | Ambystoma_mexicanum | Ambystoma_mexicanum | diurnal |
| amphibians | Ambystoma_opacum | Ambystoma_opacum | nocturnal |
| amphibians | Ambystoma_ordinarium | Ambystoma_ordinarium | diurnal |
| amphibians | Ambystoma_rosaceum | Ambystoma_rosaceum | nocturnal |
| amphibians | Ambystoma_talpoideum | Ambystoma_talpoideum | cathemeral |
| amphibians | Ambystoma_texanum | Ambystoma_texanum | nocturnal |
| amphibians | Ambystoma_tigrinum | Ambystoma_tigrinum | nocturnal |
| amphibians | Ameerega_altamazonica | Ameerega_altamazonica | crepuscular |
| amphibians | Ameerega_bassleri | Ameerega_bassleri | diurnal |
| amphibians | Ameerega_bilinguis | Ameerega_bilinguis | diurnal |
| amphibians | Ameerega_boliviana | Ameerega_boliviana | diurnal |
| amphibians | Ameerega_flavopicta | Ameerega_flavopicta | diurnal |
| amphibians | Ameerega_hahneli | Ameerega_hahneli | diurnal |
| amphibians | Ameerega_ignipedis | Ameerega_ignipedis | diurnal |
| amphibians | Ameerega_ingeri | Ameerega_ingeri | diurnal |
| amphibians | Ameerega_macero | Ameerega_macero | diurnal |
| amphibians | Ameerega_parvula | Ameerega_parvula | diurnal |
| amphibians | Ameerega_pepperi | Ameerega_pepperi | diurnal |
| amphibians | Ameerega_planipaleae | Ameerega_planipaleae | diurnal |
| amphibians | Ameerega_pongoensis | Ameerega_pongoensis | diurnal |
| amphibians | Ameerega_pulchripecta | Ameerega_pulchripecta | diurnal |
| amphibians | Ameerega_rubriventris | Ameerega_rubriventris | diurnal |
| amphibians | Ameerega_silverstonei | Ameerega_silverstonei | diurnal |
| amphibians | Ameerega_simulans | Ameerega_simulans | diurnal |
| amphibians | Ameerega_trivittata | Ameerega_trivittata | diurnal |
| amphibians | Ameerega_yoshina | Ameerega_yoshina | diurnal |
| amphibians | Amietia_angolensis | Amietia_angolensis | cathemeral |
| amphibians | Amietia_fuscigula | Amietia_fuscigula | diurnal |
| amphibians | Amietia_vertebralis | Amietia_vertebralis | diurnal |
| amphibians | Amietia_wittei | Amietia_wittei | diurnal |
| amphibians | Amolops_bellulus | Amolops_bellulus | nocturnal |
| amphibians | Amolops_cremnobatus | Amolops_cremnobatus | nocturnal |
| amphibians | Amolops_larutensis | Amolops_larutensis | nocturnal |

| | | | |
|---|---|---|---|
| amphibians | Amolops_lifanensis | Amolops_lifanensis | nocturnal |
| amphibians | Amolops_minutus | Amolops_minutus | nocturnal |
| amphibians | Amphiuma_means | Amphiuma_means | nocturnal |
| amphibians | Amphiuma_tridactylum | Amphiuma_tridactylum | crepuscular |
| amphibians | Anaxyrus_americanus | Anaxyrus_americanus | nocturnal |
| amphibians | Anaxyrus_baxteri | Anaxyrus_baxteri | diurnal |
| amphibians | Anaxyrus_boreas | Anaxyrus_boreas | crepuscular |
| amphibians | Anaxyrus_californicus | Anaxyrus_californicus | crepuscular |
| amphibians | Anaxyrus_canorus | Anaxyrus_canorus | diurnal |
| amphibians | Anaxyrus_cognatus | Anaxyrus_cognatus | diurnal |
| amphibians | Anaxyrus_compactilis | Anaxyrus_compactilis | diurnal |
| amphibians | Anaxyrus_debilis | Anaxyrus_debilis | nocturnal |
| amphibians | Anaxyrus_exsul | Anaxyrus_exsul | cathemeral |
| amphibians | Anaxyrus_fowleri | Anaxyrus_fowleri | diurnal |
| amphibians | Anaxyrus_hemiophrys | Anaxyrus_hemiophrys | cathemeral |
| amphibians | Anaxyrus_houstonensis | Anaxyrus_houstonensis | cathemeral |
| amphibians | Anaxyrus_microscaphus | Anaxyrus_microscaphus | nocturnal |
| amphibians | Anaxyrus_nelsoni | Anaxyrus_nelsoni | cathemeral |
| amphibians | Anaxyrus_punctatus | Anaxyrus_punctatus | nocturnal |
| amphibians | Anaxyrus_quercicus | Anaxyrus_quercicus | cathemeral |
| amphibians | Anaxyrus_retiformis | Anaxyrus_retiformis | nocturnal |
| amphibians | Anaxyrus_speciosus | Anaxyrus_speciosus | diurnal |
| amphibians | Anaxyrus_terrestris | Anaxyrus_terrestris | nocturnal |
| amphibians | Anaxyrus_woodhousii | Anaxyrus_woodhousii | nocturnal |
| amphibians | Andinobates_abditus | Andinobates_abditus | diurnal |
| amphibians | Andinobates_altobueyensis | Andinobates_altobueyensis | diurnal |
| amphibians | Andinobates_bombetes | Andinobates_bombetes | diurnal |
| amphibians | Andinobates_cassidyhornae | Andinobates_cassidyhornae | diurnal |
| amphibians | Andinobates_daleswansoni | Andinobates_daleswansoni | diurnal |
| amphibians | Andinobates_dorisswansonae | Andinobates_dorisswansonae | diurnal |
| amphibians | Andinobates_fulguritus | Andinobates_fulguritus | diurnal |
| amphibians | Andinobates_minutus | Andinobates_minutus | diurnal |
| amphibians | Andinobates_opisthomelas | Andinobates_opisthomelas | diurnal |
| amphibians | Andinobates_viridis | Andinobates_viridis | diurnal |

| | | | |
|---|---|---|---|
| amphibians | Andinobates_virolinensis | Andinobates_virolinensis | diurnal |
| amphibians | Andrias_davidianus | Andrias_davidianus | nocturnal |
| amphibians | Andrias_japonicus | Andrias_japonicus | nocturnal |
| amphibians | Aneides_aeneus | Aneides_aeneus | nocturnal |
| amphibians | Aneides_flavipunctatus | Aneides_flavipunctatus | nocturnal |
| amphibians | Aneides_lugubris | Aneides_lugubris | nocturnal |
| amphibians | Aneides_vagrans | Aneides_vagrans | nocturnal |
| amphibians | Anhydrophryne_ngongoniensis | Anhydrophryne_ngongoniensis | cathemeral |
| amphibians | Anilany_helenae | Stumpffia_helenae | diurnal |
| amphibians | Anodonthyla_boulengerii | Anodonthyla_boulengerii | nocturnal |
| amphibians | Anodonthyla_emilei | Anodonthyla_emilei | nocturnal |
| amphibians | Anodonthyla_hutchisoni | Anodonthyla_hutchisoni | nocturnal |
| amphibians | Anodonthyla_jeanbai | Anodonthyla_jeanbai | nocturnal |
| amphibians | Anodonthyla_moramora | Anodonthyla_moramora | nocturnal |
| amphibians | Anodonthyla_pollicaris | Anodonthyla_pollicaris | nocturnal |
| amphibians | Anodonthyla_rouxae | Anodonthyla_rouxae | nocturnal |
| amphibians | Anodonthyla_theoi | Anodonthyla_theoi | nocturnal |
| amphibians | Anodonthyla_vallani | Anodonthyla_vallani | nocturnal |
| amphibians | Anomaloglossus_baeobatrachus | Anomaloglossus_baeobatrachus | diurnal |
| amphibians | Anomaloglossus_degranvillei | Anomaloglossus_degranvillei | diurnal |
| amphibians | Anomaloglossus_guanayensis | Anomaloglossus_guanayensis | diurnal |
| amphibians | Anomaloglossus_kaiei | Anomaloglossus_kaiei | diurnal |
| amphibians | Anomaloglossus_parkerae | Anomaloglossus_parkerae | diurnal |
| amphibians | Anomaloglossus_praderioi | Anomaloglossus_praderioi | diurnal |
| amphibians | Anomaloglossus_roraima | Anomaloglossus_roraima | diurnal |
| amphibians | Anomaloglossus_rufulus | Anomaloglossus_rufulus | diurnal |
| amphibians | Anomaloglossus_stepheni | Anomaloglossus_stepheni | diurnal |
| amphibians | Anomaloglossus_tamacuarensis | Anomaloglossus_tamacuarensis | diurnal |
| amphibians | Anomaloglossus_triunfo | Anomaloglossus_triunfo | diurnal |
| amphibians | Anomaloglossus_verbeeksnyderorum | Anomaloglossus_verbeeksnyderorum | diurnal |
| amphibians | Anomaloglossus_wothuja | Anomaloglossus_wothuja | diurnal |
| amphibians | Ansonia_endauensis | Ansonia_endauensis | nocturnal |
| amphibians | Ansonia_fuliginea | Ansonia_fuliginea | nocturnal |
| amphibians | Ansonia_hanitschi | Ansonia_hanitschi | cathemeral |

| | | | |
|---|---|---|---|
| amphibians | Ansonia_inthanon | Ansonia_inthanon | nocturnal |
| amphibians | Ansonia_jeetsukumarani | Ansonia_jeetsukumarani | nocturnal |
| amphibians | Ansonia_longidigita | Ansonia_longidigita | nocturnal |
| amphibians | Ansonia_platysoma | Ansonia_platysoma | diurnal |
| amphibians | Ansonia_spinulifer | Ansonia_spinulifer | nocturnal |
| amphibians | Ansonia_tiomanica | Ansonia_tiomanica | nocturnal |
| amphibians | Aphantophryne_pansa | Aphantophryne_pansa | diurnal |
| amphibians | Aplastodiscus_albofrenatus | Aplastodiscus_albofrenatus | nocturnal |
| amphibians | Aplastodiscus_arildae | Aplastodiscus_arildae | nocturnal |
| amphibians | Aplastodiscus_cavicola | Aplastodiscus_cavicola | nocturnal |
| amphibians | Aplastodiscus_cochranae | Aplastodiscus_cochranae | nocturnal |
| amphibians | Aplastodiscus_ehrhardti | Aplastodiscus_ehrhardti | nocturnal |
| amphibians | Aplastodiscus_eugenioi | Aplastodiscus_eugenioi | nocturnal |
| amphibians | Aplastodiscus_flumineus | Aplastodiscus_flumineus | nocturnal |
| amphibians | Aplastodiscus_ibirapitanga | Aplastodiscus_ibirapitanga | nocturnal |
| amphibians | Aplastodiscus_leucopygius | Aplastodiscus_leucopygius | nocturnal |
| amphibians | Aplastodiscus_perviridis | Aplastodiscus_perviridis | nocturnal |
| amphibians | Aplastodiscus_sibilatus | Aplastodiscus_sibilatus | nocturnal |
| amphibians | Aplastodiscus_weygoldti | Aplastodiscus_weygoldti | nocturnal |
| amphibians | Aquarana_catesbeiana | Rana_catesbeiana | cathemeral |
| amphibians | Aquarana_clamitans | Rana_clamitans | cathemeral |
| amphibians | Aquarana_grylio | Rana_grylio | nocturnal |
| amphibians | Aquarana_heckscheri | Rana_heckscheri | nocturnal |
| amphibians | Aquarana_septentrionalis | Rana_septentrionalis | cathemeral |
| amphibians | Aquarana_virgatipes | Rana_virgatipes | diurnal |
| amphibians | Arcovomer_passarellii | Arcovomer_passarellii | nocturnal |
| amphibians | Arenophryne_rotunda | Arenophryne_rotunda | nocturnal |
| amphibians | Aromobates_cannatellai | Aromobates_cannatellai | diurnal |
| amphibians | Aromobates_duranti | Aromobates_duranti | diurnal |
| amphibians | Aromobates_ericksonae | Aromobates_ericksonae | diurnal |
| amphibians | Aromobates_mayorgai | Aromobates_mayorgai | diurnal |
| amphibians | Aromobates_molinarii | Aromobates_molinarii | diurnal |
| amphibians | Aromobates_nocturnus | Aromobates_nocturnus | nocturnal |
| amphibians | Aromobates_saltuensis | Aromobates_saltuensis | diurnal |

| | | | |
|---|---|---|---|
| amphibians | Aromobates_serranus | Aromobates_serranus | diurnal |
| amphibians | Arthroleptella_drewesii | Arthroleptella_drewesii | cathemeral |
| amphibians | Arthroleptides_martiensseni | Arthroleptides_martiensseni | diurnal |
| amphibians | Arthroleptis_adelphus | Arthroleptis_adelphus | diurnal |
| amphibians | Arthroleptis_adolfifriederici | Arthroleptis_adolfifriederici | diurnal |
| amphibians | Arthroleptis_bioko | Arthroleptis_bioko | diurnal |
| amphibians | Arthroleptis_crusculum | Arthroleptis_crusculum | diurnal |
| amphibians | Arthroleptis_fichika | Arthroleptis_fichika | diurnal |
| amphibians | Arthroleptis_formosus | Arthroleptis_formosus | diurnal |
| amphibians | Arthroleptis_kidogo | Arthroleptis_kidogo | diurnal |
| amphibians | Arthroleptis_langeri | Arthroleptis_langeri | diurnal |
| amphibians | Arthroleptis_nimbaensis | Arthroleptis_nimbaensis | diurnal |
| amphibians | Arthroleptis_nlonakoensis | Arthroleptis_nlonakoensis | nocturnal |
| amphibians | Arthroleptis_perreti | Arthroleptis_perreti | diurnal |
| amphibians | Arthroleptis_poecilonotus | Arthroleptis_poecilonotus | diurnal |
| amphibians | Arthroleptis_stenodactylus | Arthroleptis_stenodactylus | diurnal |
| amphibians | Arthroleptis_sylvaticus | Arthroleptis_sylvaticus | diurnal |
| amphibians | Arthroleptis_taeniatus | Arthroleptis_taeniatus | diurnal |
| amphibians | Arthroleptis_tanneri | Arthroleptis_tanneri | diurnal |
| amphibians | Arthroleptis_troglodytes | Arthroleptis_troglodytes | nocturnal |
| amphibians | Arthroleptis_tuberosus | Arthroleptis_tuberosus | diurnal |
| amphibians | Arthroleptis_variabilis | Arthroleptis_variabilis | diurnal |
| amphibians | Arthroleptis_wahlbergii | Arthroleptis_wahlbergii | cathemeral |
| amphibians | Arthroleptis_xenodactyloides | Arthroleptis_xenodactyloides | cathemeral |
| amphibians | Arthroleptis_xenodactylus | Arthroleptis_xenodactylus | cathemeral |
| amphibians | Ascaphus_montanus | Ascaphus_montanus | cathemeral |
| amphibians | Ascaphus_truei | Ascaphus_truei | nocturnal |
| amphibians | Assa_darlingtoni | Assa_darlingtoni | cathemeral |
| amphibians | Asterophrys_pullifer | Pseudocallulops_pullifer | nocturnal |
| amphibians | Astylosternus_batesi | Astylosternus_batesi | nocturnal |
| amphibians | Astylosternus_diadematus | Astylosternus_diadematus | nocturnal |
| amphibians | Astylosternus_laticephalus | Astylosternus_laticephalus | nocturnal |
| amphibians | Astylosternus_occidentalis | Astylosternus_occidentalis | nocturnal |
| amphibians | Astylosternus_perreti | Astylosternus_perreti | nocturnal |

| | | | |
|---|---|---|---|
| amphibians | Astylosternus_ranoides | Astylosternus_ranoides | diurnal |
| amphibians | Astylosternus_rheophilus | Astylosternus_rheophilus | cathemeral |
| amphibians | Astylosternus_schioetzi | Astylosternus_schioetzi | nocturnal |
| amphibians | Atelopus_andinus | Atelopus_andinus | diurnal |
| amphibians | Atelopus_angelito | Atelopus_angelito | diurnal |
| amphibians | Atelopus_arsyecue | Atelopus_arsyecue | diurnal |
| amphibians | Atelopus_arthuri | Atelopus_arthuri | diurnal |
| amphibians | Atelopus_balios | Atelopus_balios | diurnal |
| amphibians | Atelopus_bomolochos | Atelopus_bomolochos | diurnal |
| amphibians | Atelopus_boulengeri | Atelopus_boulengeri | nocturnal |
| amphibians | Atelopus_carauta | Atelopus_carauta | diurnal |
| amphibians | Atelopus_carbonerensis | Atelopus_carbonerensis | diurnal |
| amphibians | Atelopus_carrikeri | Atelopus_carrikeri | diurnal |
| amphibians | Atelopus_certus | Atelopus_certus | diurnal |
| amphibians | Atelopus_chiriquiensis | Atelopus_chiriquiensis | diurnal |
| amphibians | Atelopus_chirripoensis | Atelopus_chirripoensis | diurnal |
| amphibians | Atelopus_chocoensis | Atelopus_chocoensis | diurnal |
| amphibians | Atelopus_chrysocorallus | Atelopus_chrysocorallus | diurnal |
| amphibians | Atelopus_coynei | Atelopus_coynei | diurnal |
| amphibians | Atelopus_cruciger | Atelopus_cruciger | diurnal |
| amphibians | Atelopus_dimorphus | Atelopus_dimorphus | diurnal |
| amphibians | Atelopus_ebenoides | Atelopus_ebenoides | diurnal |
| amphibians | Atelopus_elegans | Atelopus_elegans | diurnal |
| amphibians | Atelopus_erythropus | Atelopus_erythropus | diurnal |
| amphibians | Atelopus_eusebianus | Atelopus_eusebianus | diurnal |
| amphibians | Atelopus_exiguus | Atelopus_exiguus | diurnal |
| amphibians | Atelopus_famelicus | Atelopus_famelicus | diurnal |
| amphibians | Atelopus_farci | Atelopus_farci | diurnal |
| amphibians | Atelopus_flavescens | Atelopus_flavescens | diurnal |
| amphibians | Atelopus_franciscus | Atelopus_franciscus | diurnal |
| amphibians | Atelopus_galactogaster | Atelopus_galactogaster | diurnal |
| amphibians | Atelopus_glyphus | Atelopus_glyphus | diurnal |
| amphibians | Atelopus_guanujo | Atelopus_guanujo | diurnal |
| amphibians | Atelopus_guitarraensis | Atelopus_guitarraensis | diurnal |

| | | | |
|---|---|---|---|
| amphibians | Atelopus_halihelos | Atelopus_halihelos | diurnal |
| amphibians | Atelopus_hoogmoedi | Atelopus_hoogmoedi | diurnal |
| amphibians | Atelopus_ignescens | Atelopus_ignescens | diurnal |
| amphibians | Atelopus_laetissimus | Atelopus_laetissimus | diurnal |
| amphibians | Atelopus_limosus | Atelopus_limosus | diurnal |
| amphibians | Atelopus_longibrachius | Atelopus_longibrachius | diurnal |
| amphibians | Atelopus_longirostris | Atelopus_longirostris | cathemeral |
| amphibians | Atelopus_lozanoi | Atelopus_lozanoi | diurnal |
| amphibians | Atelopus_lynchi | Atelopus_lynchi | cathemeral |
| amphibians | Atelopus_mandingues | Atelopus_mandingues | diurnal |
| amphibians | Atelopus_marinkellei | Atelopus_marinkellei | diurnal |
| amphibians | Atelopus_mindoensis | Atelopus_mindoensis | diurnal |
| amphibians | Atelopus_minutulus | Atelopus_minutulus | diurnal |
| amphibians | Atelopus_monohernandezii | Atelopus_monohernandezii | diurnal |
| amphibians | Atelopus_mucubajiensis | Atelopus_mucubajiensis | diurnal |
| amphibians | Atelopus_muisca | Atelopus_muisca | diurnal |
| amphibians | Atelopus_nahumae | Atelopus_nahumae | diurnal |
| amphibians | Atelopus_nanay | Atelopus_nanay | diurnal |
| amphibians | Atelopus_nepiozomus | Atelopus_nepiozomus | diurnal |
| amphibians | Atelopus_nicefori | Atelopus_nicefori | diurnal |
| amphibians | Atelopus_onorei | Atelopus_onorei | diurnal |
| amphibians | Atelopus_orcesi | Atelopus_orcesi | diurnal |
| amphibians | Atelopus_oxapampae | Atelopus_oxapampae | crepuscular |
| amphibians | Atelopus_oxyrhynchus | Atelopus_oxyrhynchus | diurnal |
| amphibians | Atelopus_pachydermus | Atelopus_pachydermus | diurnal |
| amphibians | Atelopus_palmatus | Atelopus_palmatus | cathemeral |
| amphibians | Atelopus_pastuso | Atelopus_pastuso | diurnal |
| amphibians | Atelopus_patazensis | Atelopus_patazensis | diurnal |
| amphibians | Atelopus_pedimarmoratus | Atelopus_pedimarmoratus | diurnal |
| amphibians | Atelopus_peruensis | Atelopus_peruensis | diurnal |
| amphibians | Atelopus_petersi | Atelopus_petersi | diurnal |
| amphibians | Atelopus_pictiventris | Atelopus_pictiventris | diurnal |
| amphibians | Atelopus_pinangoi | Atelopus_pinangoi | diurnal |
| amphibians | Atelopus_planispina | Atelopus_planispina | diurnal |

| | | | |
|---|---|---|---|
| amphibians | Atelopus_podocarpus | Atelopus_podocarpus | diurnal |
| amphibians | Atelopus_pulcher | Atelopus_pulcher | diurnal |
| amphibians | Atelopus_quimbaya | Atelopus_quimbaya | diurnal |
| amphibians | Atelopus_reticulatus | Atelopus_reticulatus | diurnal |
| amphibians | Atelopus_sanjosei | Atelopus_sanjosei | diurnal |
| amphibians | Atelopus_seminiferus | Atelopus_seminiferus | diurnal |
| amphibians | Atelopus_senex | Atelopus_senex | diurnal |
| amphibians | Atelopus_sernai | Atelopus_sernai | diurnal |
| amphibians | Atelopus_simulatus | Atelopus_simulatus | diurnal |
| amphibians | Atelopus_siranus | Atelopus_siranus | diurnal |
| amphibians | Atelopus_sonsonensis | Atelopus_sonsonensis | diurnal |
| amphibians | Atelopus_sorianoi | Atelopus_sorianoi | diurnal |
| amphibians | Atelopus_spumarius | Atelopus_spumarius | diurnal |
| amphibians | Atelopus_spurrelli | Atelopus_spurrelli | diurnal |
| amphibians | Atelopus_subornatus | Atelopus_subornatus | diurnal |
| amphibians | Atelopus_tamaense | Atelopus_tamaensis | diurnal |
| amphibians | Atelopus_tricolor | Atelopus_tricolor | diurnal |
| amphibians | Atelopus_varius | Atelopus_varius | diurnal |
| amphibians | Atelopus_vogli | Atelopus_vogli | diurnal |
| amphibians | Atelopus_walkeri | Atelopus_walkeri | diurnal |
| amphibians | Atelopus_zeteki | Atelopus_zeteki | diurnal |
| amphibians | Atlantihyla_spinipollex | Ptychohyla_spinipollex | nocturnal |
| amphibians | Aubria_masako | Aubria_masako | nocturnal |
| amphibians | Aubria_subsigillata | Aubria_subsigillata | nocturnal |
| amphibians | Austrochaperina_basipalmata | Austrochaperina_basipalmata | diurnal |
| amphibians | Austrochaperina_brevipes | Austrochaperina_brevipes | diurnal |
| amphibians | Austrochaperina_fryi | Austrochaperina_fryi | diurnal |
| amphibians | Austrochaperina_hooglandi | Austrochaperina_hooglandi | diurnal |
| amphibians | Austrochaperina_palmipes | Austrochaperina_palmipes | nocturnal |
| amphibians | Austrochaperina_septentrionalis | Austrochaperina_septentrionalis | diurnal |
| amphibians | Balebreviceps_hillmani | Balebreviceps_hillmani | diurnal |
| amphibians | Barbarophryne_brongersmai | Barbarophryne_brongersmai | nocturnal |
| amphibians | Barygenys_maculata | Barygenys_maculata | diurnal |
| amphibians | Barygenys_parvula | Barygenys_parvula | nocturnal |

| | | | |
|---|---|---|---|
| amphibians | Batrachoseps_attenuatus | Batrachoseps_attenuatus | crepuscular |
| amphibians | Batrachoseps_bramei | Batrachoseps_bramei | nocturnal |
| amphibians | Batrachoseps_campi | Batrachoseps_campi | nocturnal |
| amphibians | Batrachoseps_pacificus | Batrachoseps_pacificus | nocturnal |
| amphibians | Batrachuperus_tibetanus | Batrachuperus_tibetanus | cathemeral |
| amphibians | Batrachyla_taeniata | Batrachyla_taeniata | nocturnal |
| amphibians | Beddomixalus_bijui | Beddomixalus_bijui | nocturnal |
| amphibians | Beduka_amboli | Xanthophryne_tigerina | crepuscular |
| amphibians | Beduka_koynayensis | Xanthophryne_koynayensis | crepuscular |
| amphibians | Blommersia_angolafa | Blommersia_angolafa | cathemeral |
| amphibians | Blommersia_blommersae | Blommersia_blommersae | cathemeral |
| amphibians | Blommersia_dejongi | Blommersia_dejongi | nocturnal |
| amphibians | Blommersia_domerguei | Blommersia_domerguei | cathemeral |
| amphibians | Blommersia_galani | Blommersia_galani | cathemeral |
| amphibians | Blommersia_grandisonae | Blommersia_grandisonae | cathemeral |
| amphibians | Blommersia_kely | Blommersia_kely | diurnal |
| amphibians | Blommersia_sarotra | Blommersia_sarotra | cathemeral |
| amphibians | Blommersia_variabilis | Blommersia_variabilis | cathemeral |
| amphibians | Blommersia_wittei | Blommersia_wittei | cathemeral |
| amphibians | Boana_aguilari | Hypsiboas_aguilari | nocturnal |
| amphibians | Boana_albomarginata | Hypsiboas_albomarginatus | nocturnal |
| amphibians | Boana_albopunctata | Hypsiboas_albopunctatus | nocturnal |
| amphibians | Boana_alfaroi | Hypsiboas_alfaroi | nocturnal |
| amphibians | Boana_almendarizae | Hypsiboas_almendarizae | nocturnal |
| amphibians | Boana_atlantica | Hypsiboas_atlanticus | nocturnal |
| amphibians | Boana_balzani | Hypsiboas_balzani | nocturnal |
| amphibians | Boana_beckeri | Hypsiboas_beckeri | nocturnal |
| amphibians | Boana_bischoffi | Hypsiboas_bischoffi | nocturnal |
| amphibians | Boana_boans | Hypsiboas_boans | cathemeral |
| amphibians | Boana_botumirim | Hypsiboas_botumirim | nocturnal |
| amphibians | Boana_buriti | Hypsiboas_buriti | nocturnal |
| amphibians | Boana_caingua | Hypsiboas_caingua | nocturnal |
| amphibians | Boana_caipora | Hypsiboas_caipora | nocturnal |
| amphibians | Boana_calcarata | Hypsiboas_calcaratus | nocturnal |

| | | | |
|---|---|---|---|
| amphibians | Boana_callipleura | Hypsiboas_callipleura | nocturnal |
| amphibians | Boana_cinerascens | Hypsiboas_cinerascens | nocturnal |
| amphibians | Boana_cipoensis | Hypsiboas_cipoensis | nocturnal |
| amphibians | Boana_crepitans | Hypsiboas_crepitans | nocturnal |
| amphibians | Boana_curupi | Hypsiboas_curupi | nocturnal |
| amphibians | Boana_dentei | Hypsiboas_dentei | nocturnal |
| amphibians | Boana_ericae | Hypsiboas_ericae | nocturnal |
| amphibians | Boana_exastis | Hypsiboas_exastis | nocturnal |
| amphibians | Boana_faber | Hypsiboas_faber | nocturnal |
| amphibians | Boana_fasciata | Hypsiboas_fasciatus | nocturnal |
| amphibians | Boana_freicanecae | Hypsiboas_freicanecae | nocturnal |
| amphibians | Boana_geographica | Hypsiboas_geographicus | nocturnal |
| amphibians | Boana_guentheri | Hypsiboas_guentheri | nocturnal |
| amphibians | Boana_heilprini | Hypsiboas_heilprini | cathemeral |
| amphibians | Boana_hobbsi | Hypsiboas_hobbsi | nocturnal |
| amphibians | Boana_hutchinsi | Hypsiboas_hutchinsi | nocturnal |
| amphibians | Boana_jimenezi | Hypsiboas_jimenezi | nocturnal |
| amphibians | Boana_joaquini | Hypsiboas_joaquini | nocturnal |
| amphibians | Boana_lanciformis | Hypsiboas_lanciformis | nocturnal |
| amphibians | Boana_latistriata | Hypsiboas_latistriatus | nocturnal |
| amphibians | Boana_lemai | Hypsiboas_lemai | nocturnal |
| amphibians | Boana_leptolineata | Hypsiboas_leptolineatus | nocturnal |
| amphibians | Boana_lundii | Hypsiboas_lundii | nocturnal |
| amphibians | Boana_maculateralis | Hypsiboas_maculateralis | nocturnal |
| amphibians | Boana_marginata | Hypsiboas_marginatus | nocturnal |
| amphibians | Boana_marianitae | Hypsiboas_marianitae | nocturnal |
| amphibians | Boana_melanopleura | Hypsiboas_melanopleura | nocturnal |
| amphibians | Boana_microderma | Hypsiboas_microderma | nocturnal |
| amphibians | Boana_multifasciata | Hypsiboas_multifasciatus | nocturnal |
| amphibians | Boana_nympha | Hypsiboas_nympha | nocturnal |
| amphibians | Boana_ornatissima | Hypsiboas_ornatissimus | nocturnal |
| amphibians | Boana_palaestes | Hypsiboas_palaestes | nocturnal |
| amphibians | Boana_paranaiba | Hypsiboas_paranaiba | nocturnal |
| amphibians | Boana_pardalis | Hypsiboas_pardalis | nocturnal |

| | | | |
|---|---|---|---|
| amphibians | Boana_pellucens | Hypsiboas_pellucens | nocturnal |
| amphibians | Boana_picturata | Hypsiboas_picturatus | nocturnal |
| amphibians | Boana_poaju | Hypsiboas_poaju | nocturnal |
| amphibians | Boana_polytaenia | Hypsiboas_polytaenius | nocturnal |
| amphibians | Boana_pombali | Hypsiboas_pombali | nocturnal |
| amphibians | Boana_prasina | Hypsiboas_prasinus | nocturnal |
| amphibians | Boana_pugnax | Hypsiboas_pugnax | nocturnal |
| amphibians | Boana_pulchella | Hypsiboas_pulchellus | nocturnal |
| amphibians | Boana_punctata | Hypsiboas_punctatus | cathemeral |
| amphibians | Boana_raniceps | Hypsiboas_raniceps | cathemeral |
| amphibians | Boana_rhythmica | Hypsiboas_rhythmicus | nocturnal |
| amphibians | Boana_roraima | Hypsiboas_roraima | nocturnal |
| amphibians | Boana_rosenbergi | Hypsiboas_rosenbergi | nocturnal |
| amphibians | Boana_rubracyla | Hypsiboas_rubracylus | nocturnal |
| amphibians | Boana_rufitela | Hypsiboas_rufitelus | nocturnal |
| amphibians | Boana_secedens | Hypsiboas_secedens | nocturnal |
| amphibians | Boana_semiguttata | Hypsiboas_semiguttatus | nocturnal |
| amphibians | Boana_semilineata | Hypsiboas_semilineatus | nocturnal |
| amphibians | Boana_sibleszi | Hypsiboas_sibleszi | nocturnal |
| amphibians | Boana_stellae | Hypsiboas_stellae | nocturnal |
| amphibians | Boana_stenocephala | Hypsiboas_stenocephalus | nocturnal |
| amphibians | Boana_tepuiana | Hypsiboas_tepuianus | nocturnal |
| amphibians | Boana_tetete | Hypsiboas_tetete | nocturnal |
| amphibians | Boehmantis_microtympanum | Boehmantis_microtympanum | nocturnal |
| amphibians | Bokermannohyla_ahenea | Bokermannohyla_ahenea | nocturnal |
| amphibians | Bokermannohyla_alvarengai | Bokermannohyla_alvarengai | nocturnal |
| amphibians | Bokermannohyla_astartea | Bokermannohyla_astartea | nocturnal |
| amphibians | Bokermannohyla_capra | Bokermannohyla_capra | nocturnal |
| amphibians | Bokermannohyla_caramaschii | Bokermannohyla_caramaschii | nocturnal |
| amphibians | Bokermannohyla_carvalhoi | Bokermannohyla_carvalhoi | nocturnal |
| amphibians | Bokermannohyla_circumdata | Bokermannohyla_circumdata | nocturnal |
| amphibians | Bokermannohyla_diamantina | Bokermannohyla_diamantina | nocturnal |
| amphibians | Bokermannohyla_gouveai | Bokermannohyla_gouveai | nocturnal |
| amphibians | Bokermannohyla_hylax | Bokermannohyla_hylax | nocturnal |

| | | | |
|---|---|---|---|
| amphibians | Bokermannohyla_ibitiguara | Bokermannohyla_ibitiguara | nocturnal |
| amphibians | Bokermannohyla_ibitipoca | Bokermannohyla_ibitipoca | nocturnal |
| amphibians | Bokermannohyla_itapoty | Bokermannohyla_itapoty | nocturnal |
| amphibians | Bokermannohyla_izecksohni | Bokermannohyla_izecksohni | nocturnal |
| amphibians | Bokermannohyla_lucianae | Bokermannohyla_lucianae | nocturnal |
| amphibians | Bokermannohyla_luctuosa | Bokermannohyla_luctuosa | nocturnal |
| amphibians | Bokermannohyla_martinsi | Bokermannohyla_martinsi | nocturnal |
| amphibians | Bokermannohyla_nanuzae | Bokermannohyla_nanuzae | nocturnal |
| amphibians | Bokermannohyla_oxente | Bokermannohyla_oxente | nocturnal |
| amphibians | Bokermannohyla_saxicola | Bokermannohyla_saxicola | nocturnal |
| amphibians | Bokermannohyla_vulcaniae | Bokermannohyla_vulcaniae | nocturnal |
| amphibians | Bolitoglossa_adspersa | Bolitoglossa_adspersa | diurnal |
| amphibians | Bolitoglossa_altamazonica | Bolitoglossa_altamazonica | nocturnal |
| amphibians | Bolitoglossa_alvaradoi | Bolitoglossa_alvaradoi | nocturnal |
| amphibians | Bolitoglossa_biseriata | Bolitoglossa_biseriata | nocturnal |
| amphibians | Bolitoglossa_bramei | Bolitoglossa_bramei | nocturnal |
| amphibians | Bolitoglossa_chica | Bolitoglossa_chica | nocturnal |
| amphibians | Bolitoglossa_colonnea | Bolitoglossa_colonnea | nocturnal |
| amphibians | Bolitoglossa_compacta | Bolitoglossa_compacta | nocturnal |
| amphibians | Bolitoglossa_decora | Bolitoglossa_decora | nocturnal |
| amphibians | Bolitoglossa_diaphora | Bolitoglossa_diaphora | nocturnal |
| amphibians | Bolitoglossa_dunni | Bolitoglossa_dunni | nocturnal |
| amphibians | Bolitoglossa_epimela | Bolitoglossa_epimela | nocturnal |
| amphibians | Bolitoglossa_equatoriana | Bolitoglossa_equatoriana | nocturnal |
| amphibians | Bolitoglossa_gomezi | Bolitoglossa_gomezi | nocturnal |
| amphibians | Bolitoglossa_lignicolor | Bolitoglossa_lignicolor | nocturnal |
| amphibians | Bolitoglossa_longissima | Bolitoglossa_longissima | nocturnal |
| amphibians | Bolitoglossa_medemi | Bolitoglossa_medemi | cathemeral |
| amphibians | Bolitoglossa_mexicana | Bolitoglossa_mexicana | nocturnal |
| amphibians | Bolitoglossa_minutula | Bolitoglossa_minutula | nocturnal |
| amphibians | Bolitoglossa_nigrescens | Bolitoglossa_nigrescens | nocturnal |
| amphibians | Bolitoglossa_occidentalis | Bolitoglossa_occidentalis | nocturnal |
| amphibians | Bolitoglossa_odonnelli | Bolitoglossa_odonnelli | nocturnal |
| amphibians | Bolitoglossa_palmata | Bolitoglossa_palmata | nocturnal |

| | | | |
|---|---|---|---|
| amphibians | Bolitoglossa_peruviana | Bolitoglossa_peruviana | cathemeral |
| amphibians | Bolitoglossa_pesrubra | Bolitoglossa_pesrubra | nocturnal |
| amphibians | Bolitoglossa_platydactyla | Bolitoglossa_platydactyla | nocturnal |
| amphibians | Bolitoglossa_porrasorum | Bolitoglossa_porrasorum | nocturnal |
| amphibians | Bolitoglossa_robusta | Bolitoglossa_robusta | nocturnal |
| amphibians | Bolitoglossa_rufescens | Bolitoglossa_rufescens | nocturnal |
| amphibians | Bolitoglossa_schizodactyla | Bolitoglossa_schizodactyla | nocturnal |
| amphibians | Bolitoglossa_sima | Bolitoglossa_sima | nocturnal |
| amphibians | Bolitoglossa_sooyorum | Bolitoglossa_sooyorum | nocturnal |
| amphibians | Bolitoglossa_striatula | Bolitoglossa_striatula | nocturnal |
| amphibians | Bolitoglossa_stuarti | Bolitoglossa_stuarti | nocturnal |
| amphibians | Bolitoglossa_subpalmata | Bolitoglossa_subpalmata | nocturnal |
| amphibians | Bolitoglossa_synoria | Bolitoglossa_synoria | nocturnal |
| amphibians | Bombina_bombina | Bombina_bombina | diurnal |
| amphibians | Bombina_maxima | Bombina_maxima | diurnal |
| amphibians | Bombina_orientalis | Bombina_orientalis | nocturnal |
| amphibians | Boophis_albilabris | Boophis_albilabris | nocturnal |
| amphibians | Boophis_albipunctatus | Boophis_albipunctatus | nocturnal |
| amphibians | Boophis_andohahela | Boophis_andohahela | nocturnal |
| amphibians | Boophis_andreonei | Boophis_andreonei | nocturnal |
| amphibians | Boophis_anjanaharibeensis | Boophis_anjanaharibeensis | nocturnal |
| amphibians | Boophis_ankaratra | Boophis_ankaratra | nocturnal |
| amphibians | Boophis_axelmeyeri | Boophis_axelmeyeri | nocturnal |
| amphibians | Boophis_baetkei | Boophis_baetkei | nocturnal |
| amphibians | Boophis_blommersae | Boophis_blommersae | nocturnal |
| amphibians | Boophis_boehmei | Boophis_boehmei | nocturnal |
| amphibians | Boophis_bottae | Boophis_bottae | nocturnal |
| amphibians | Boophis_brachychir | Boophis_brachychir | nocturnal |
| amphibians | Boophis_calcaratus | Boophis_calcaratus | nocturnal |
| amphibians | Boophis_doulioti | Boophis_doulioti | nocturnal |
| amphibians | Boophis_elenae | Boophis_elenae | nocturnal |
| amphibians | Boophis_englaenderi | Boophis_englaenderi | nocturnal |
| amphibians | Boophis_entingae | Boophis_entingae | nocturnal |
| amphibians | Boophis_erythrodactylus | Boophis_erythrodactylus | nocturnal |

| | | | |
|---|---|---|---|
| amphibians | Boophis_feonnyala | Boophis_feonnyala | nocturnal |
| amphibians | Boophis_goudotii | Boophis_goudotii | nocturnal |
| amphibians | Boophis_guibei | Boophis_guibei | cathemeral |
| amphibians | Boophis_idae | Boophis_idae | nocturnal |
| amphibians | Boophis_jaegeri | Boophis_jaegeri | nocturnal |
| amphibians | Boophis_liami | Boophis_liami | nocturnal |
| amphibians | Boophis_lichenoides | Boophis_lichenoides | nocturnal |
| amphibians | Boophis_lilianae | Boophis_lilianae | nocturnal |
| amphibians | Boophis_luteus | Boophis_luteus | nocturnal |
| amphibians | Boophis_madagascariensis | Boophis_madagascariensis | nocturnal |
| amphibians | Boophis_marojezensis | Boophis_marojezensis | nocturnal |
| amphibians | Boophis_microtympanum | Boophis_microtympanum | cathemeral |
| amphibians | Boophis_miniatus | Boophis_miniatus | nocturnal |
| amphibians | Boophis_occidentalis | Boophis_occidentalis | crepuscular |
| amphibians | Boophis_opisthodon | Boophis_opisthodon | nocturnal |
| amphibians | Boophis_picturatus | Boophis_picturatus | nocturnal |
| amphibians | Boophis_praedictus | Boophis_praedictus | nocturnal |
| amphibians | Boophis_pyrrhus | Boophis_pyrrhus | nocturnal |
| amphibians | Boophis_rappiodes | Boophis_rappiodes | nocturnal |
| amphibians | Boophis_reticulatus | Boophis_reticulatus | nocturnal |
| amphibians | Boophis_rhodoscelis | Boophis_rhodoscelis | nocturnal |
| amphibians | Boophis_roseipalmatus | Boophis_roseipalmatus | nocturnal |
| amphibians | Boophis_rufioculis | Boophis_rufioculis | nocturnal |
| amphibians | Boophis_sambirano | Boophis_sambirano | nocturnal |
| amphibians | Boophis_schuboeae | Boophis_schuboeae | nocturnal |
| amphibians | Boophis_septentrionalis | Boophis_septentrionalis | nocturnal |
| amphibians | Boophis_sibilans | Boophis_sibilans | nocturnal |
| amphibians | Boophis_solomaso | Boophis_solomaso | nocturnal |
| amphibians | Boophis_tampoka | Boophis_tampoka | nocturnal |
| amphibians | Boophis_tasymena | Boophis_tasymena | nocturnal |
| amphibians | Boophis_tephraeomystax | Boophis_tephraeomystax | nocturnal |
| amphibians | Boophis_tsilomaro | Boophis_tsilomaro | nocturnal |
| amphibians | Boophis_ulftunni | Boophis_ulftunni | nocturnal |
| amphibians | Boophis_viridis | Boophis_viridis | nocturnal |

| | | | |
|---|---|---|---|
| amphibians | Boophis_vittatus | Boophis_vittatus | nocturnal |
| amphibians | Boophis_williamsi | Boophis_williamsi | nocturnal |
| amphibians | Boophis_xerophilus | Boophis_xerophilus | nocturnal |
| amphibians | Boreorana_sylvatica | Rana_sylvatica | diurnal |
| amphibians | Boulengerula_taitana | Boulengerula_taitana | cathemeral |
| amphibians | Boulenophrys_baolongensis | Xenophrys_baolongensis | nocturnal |
| amphibians | Boulenophrys_binlingensis | Xenophrys_binlingensis | nocturnal |
| amphibians | Boulenophrys_kuatunensis | Xenophrys_kuatunensis | nocturnal |
| amphibians | Brachycephalus_alipioi | Brachycephalus_alipioi | diurnal |
| amphibians | Brachycephalus_brunneus | Brachycephalus_brunneus | diurnal |
| amphibians | Brachycephalus_didactylus | Brachycephalus_didactylus | diurnal |
| amphibians | Brachycephalus_ephippium | Brachycephalus_ephippium | diurnal |
| amphibians | Brachycephalus_ferruginus | Brachycephalus_ferruginus | diurnal |
| amphibians | Brachycephalus_garbeanus | Brachycephalus_garbeana | diurnal |
| amphibians | Brachycephalus_guarani | Brachycephalus_guarani | diurnal |
| amphibians | Brachycephalus_hermogenesi | Brachycephalus_hermogenesi | diurnal |
| amphibians | Brachycephalus_izecksohni | Brachycephalus_izecksohni | diurnal |
| amphibians | Brachycephalus_margaritatus | Brachycephalus_margaritatus | diurnal |
| amphibians | Brachycephalus_nodoterga | Brachycephalus_nodoterga | diurnal |
| amphibians | Brachycephalus_pernix | Brachycephalus_pernix | diurnal |
| amphibians | Brachycephalus_pitanga | Brachycephalus_pitanga | diurnal |
| amphibians | Brachycephalus_pombali | Brachycephalus_pombali | diurnal |
| amphibians | Brachycephalus_toby | Brachycephalus_toby | diurnal |
| amphibians | Brachycephalus_tridactylus | Brachycephalus_tridactylus | diurnal |
| amphibians | Brachycephalus_vertebralis | Brachycephalus_vertebralis | diurnal |
| amphibians | Brachytarsophrys_platyparietus | Brachytarsophrys_carinense | nocturnal |
| amphibians | Breviceps_adspersus | Breviceps_adspersus | nocturnal |
| amphibians | Breviceps_bagginsi | Breviceps_bagginsi | diurnal |
| amphibians | Breviceps_fichus | Breviceps_fichus | diurnal |
| amphibians | Breviceps_macrops | Breviceps_macrops | nocturnal |
| amphibians | Breviceps_sopranus | Breviceps_sopranus | cathemeral |
| amphibians | Bromeliohyla_bromeliacia | Bromeliohyla_bromeliacia | nocturnal |
| amphibians | Buergeria_buergeri | Buergeria_buergeri | cathemeral |
| amphibians | Bufo_bankorensis | Bufo_bankorensis | nocturnal |

| | | | |
|---|---|---|---|
| amphibians | Bufo_bufo | Bufo_bufo | nocturnal |
| amphibians | Bufo_eichwaldi | Bufo_eichwaldi | nocturnal |
| amphibians | Bufo_gargarizans | Bufo_gargarizans | cathemeral |
| amphibians | Bufo_spinosus | Bufo_spinosus | nocturnal |
| amphibians | Bufo_stejnegeri | Bufo_stejnegeri | nocturnal |
| amphibians | Bufo_torrenticola | Bufo_torrenticola | cathemeral |
| amphibians | Bufo_tuberculatus | Bufo_tuberculatus | nocturnal |
| amphibians | Bufotes_pewzowi | Bufotes_pewzowi | nocturnal |
| amphibians | Bufotes_surdus | Bufotes_surdus | nocturnal |
| amphibians | Cacosternum_capense | Cacosternum_capense | nocturnal |
| amphibians | Cacosternum_namaquense | Cacosternum_namaquense | nocturnal |
| amphibians | Caecilia_volcani | Caecilia_volcani | cathemeral |
| amphibians | Callimedusa_atelopoides | Phyllomedusa_atelopoides | nocturnal |
| amphibians | Callimedusa_baltea | Phyllomedusa_baltea | nocturnal |
| amphibians | Callimedusa_duellmani | Phyllomedusa_duellmani | nocturnal |
| amphibians | Callimedusa_tomopterna | Phyllomedusa_tomopterna | nocturnal |
| amphibians | Callixalus_pictus | Callixalus_pictus | nocturnal |
| amphibians | Callulina_dawida | Callulina_dawida | nocturnal |
| amphibians | Callulina_laphami | Callulina_laphami | nocturnal |
| amphibians | Callulina_shengena | Callulina_shengena | nocturnal |
| amphibians | Callulina_stanleyi | Callulina_stanleyi | nocturnal |
| amphibians | Callulops_doriae | Callulops_doriae | nocturnal |
| amphibians | Calotriton_arnoldi | Calotriton_arnoldi | nocturnal |
| amphibians | Calotriton_asper | Calotriton_asper | nocturnal |
| amphibians | Calyptocephalella_gayi | Calyptocephalella_gayi | cathemeral |
| amphibians | Cardioglossa_alsco | Cardioglossa_alsco | nocturnal |
| amphibians | Cardioglossa_occidentalis | Cardioglossa_occidentalis | nocturnal |
| amphibians | Cardioglossa_trifasciata | Cardioglossa_trifasciata | diurnal |
| amphibians | Celsiella_revocata | Celsiella_revocata | nocturnal |
| amphibians | Centrolene_ballux | Centrolene_ballux | nocturnal |
| amphibians | Centrolene_buckleyi | Centrolene_buckleyi | nocturnal |
| amphibians | Centrolene_condor | Centrolene_condor | nocturnal |
| amphibians | Centrolene_daidalea | Centrolene_daidaleum | nocturnal |
| amphibians | Centrolene_heloderma | Centrolene_heloderma | cathemeral |

| | | | |
|---|---|---|---|
| amphibians | Centrolene_hesperia | Centrolene_hesperium | cathemeral |
| amphibians | Centrolene_huilensis | Centrolene_huilense | nocturnal |
| amphibians | Centrolene_lemniscata | Centrolene_lemniscatum | nocturnal |
| amphibians | Centrolene_muelleri | Centrolene_muelleri | nocturnal |
| amphibians | Centrolene_notosticta | Centrolene_notostictum | nocturnal |
| amphibians | Centrolene_pipilata | Centrolene_pipilatum | nocturnal |
| amphibians | Centrolene_sabini | Centrolene_sabini | nocturnal |
| amphibians | Centrolene_savagei | Centrolene_savagei | nocturnal |
| amphibians | Centrolene_venezuelense | Centrolene_venezuelense | nocturnal |
| amphibians | Ceratophrys_aurita | Ceratophrys_aurita | nocturnal |
| amphibians | Ceratophrys_calcarata | Ceratophrys_calcarata | nocturnal |
| amphibians | Ceratophrys_cornuta | Ceratophrys_cornuta | cathemeral |
| amphibians | Ceratophrys_cranwelli | Ceratophrys_cranwelli | nocturnal |
| amphibians | Ceratophrys_joazeirensis | Ceratophrys_joazeirensis | nocturnal |
| amphibians | Ceratophrys_ornata | Ceratophrys_ornata | nocturnal |
| amphibians | Ceratophrys_stolzmanni | Ceratophrys_stolzmanni | nocturnal |
| amphibians | Ceuthomantis_aracamuni | Ceuthomantis_aracamuni | diurnal |
| amphibians | Ceuthomantis_cavernibardus | Ceuthomantis_cavernibardus | diurnal |
| amphibians | Ceuthomantis_duellmani | Ceuthomantis_duellmani | diurnal |
| amphibians | Ceuthomantis_smaragdinus | Ceuthomantis_smaragdinus | nocturnal |
| amphibians | Chacophrys_pierottii | Chacophrys_pierottii | nocturnal |
| amphibians | Chaperina_fusca | Chaperina_fusca | cathemeral |
| amphibians | Charadrahyla_chaneque | Charadrahyla_chaneque | nocturnal |
| amphibians | Charadrahyla_taeniopus | Charadrahyla_taeniopus | nocturnal |
| amphibians | Charadrahyla_tecuani | Charadrahyla_tecuani | nocturnal |
| amphibians | Chiasmocleis_alagoana | Chiasmocleis_alagoanus | diurnal |
| amphibians | Chiasmocleis_albopunctata | Chiasmocleis_albopunctata | nocturnal |
| amphibians | Chiasmocleis_anatipes | Chiasmocleis_anatipes | nocturnal |
| amphibians | Chiasmocleis_antenori | Syncope_antenori | cathemeral |
| amphibians | Chiasmocleis_atlantica | Chiasmocleis_atlantica | nocturnal |
| amphibians | Chiasmocleis_bassleri | Syncope_bassleri | nocturnal |
| amphibians | Chiasmocleis_capixaba | Chiasmocleis_capixaba | nocturnal |
| amphibians | Chiasmocleis_carvalhoi | Chiasmocleis_carvalhoi | nocturnal |
| amphibians | Chiasmocleis_crucis | Chiasmocleis_crucis | nocturnal |

| | | | |
|---|---|---|---|
| amphibians | Chiasmocleis_gnoma | Chiasmocleis_gnoma | nocturnal |
| amphibians | Chiasmocleis_hudsoni | Syncope_hudsoni | nocturnal |
| amphibians | Chiasmocleis_leucosticta | Chiasmocleis_leucosticta | cathemeral |
| amphibians | Chiasmocleis_mantiqueira | Chiasmocleis_mantiqueira | nocturnal |
| amphibians | Chiasmocleis_schubarti | Chiasmocleis_schubarti | nocturnal |
| amphibians | Chiasmocleis_shudikarensis | Chiasmocleis_shudikarensis | cathemeral |
| amphibians | Chiasmocleis_supercilialba | Syncope_supercilialba | nocturnal |
| amphibians | Chiasmocleis_ventrimaculata | Chiasmocleis_ventrimaculata | nocturnal |
| amphibians | Chimerella_mariaelenae | Chimerella_mariaelenae | nocturnal |
| amphibians | Chioglossa_lusitanica | Chioglossa_lusitanica | nocturnal |
| amphibians | Chirixalus_dudhwaensis | Chiromantis_dudhwaensis | nocturnal |
| amphibians | Chirixalus_nongkhorensis | Chiromantis_nongkhorensis | cathemeral |
| amphibians | Chirixalus_simus | Chiromantis_simus | nocturnal |
| amphibians | Chiromantis_petersii | Chiromantis_petersii | nocturnal |
| amphibians | Chiromantis_xerampelina | Chiromantis_xerampelina | nocturnal |
| amphibians | Choerophryne_arndtorum | Choerophryne_arndtorum | nocturnal |
| amphibians | Choerophryne_burtoni | Choerophryne_burtoni | nocturnal |
| amphibians | Choerophryne_microps | Choerophryne_microps | nocturnal |
| amphibians | Choerophryne_siegfriedi | Albericus_siegfriedi | nocturnal |
| amphibians | Chrysopaa_sternosignata | Chrysopaa_sternosignata | crepuscular |
| amphibians | Chthonerpeton_noctinectes | Chthonerpeton_noctinectes | nocturnal |
| amphibians | Clinotarsus_curtipes | Clinotarsus_curtipes | nocturnal |
| amphibians | Cochranella_euknemos | Cochranella_euknemos | nocturnal |
| amphibians | Cochranella_granulosa | Cochranella_granulosa | nocturnal |
| amphibians | Cochranella_mache | Cochranella_mache | nocturnal |
| amphibians | Colostethus_agilis | Colostethus_agilis | diurnal |
| amphibians | Colostethus_inguinalis | Colostethus_inguinalis | diurnal |
| amphibians | Colostethus_jacobuspetersi | Colostethus_jacobuspetersi | diurnal |
| amphibians | Colostethus_latinasus | Colostethus_latinasus | diurnal |
| amphibians | Colostethus_panamansis | Colostethus_panamansis | diurnal |
| amphibians | Colostethus_pratti | Colostethus_pratti | diurnal |
| amphibians | Colostethus_ucumari | Colostethus_ucumari | cathemeral |
| amphibians | Conraua_alleni | Conraua_alleni | cathemeral |
| amphibians | Conraua_beccarii | Conraua_beccarii | cathemeral |

| | | | |
|---|---|---|---|
| amphibians | Conraua_crassipes | Conraua_crassipes | cathemeral |
| amphibians | Conraua_derooi | Conraua_derooi | cathemeral |
| amphibians | Conraua_goliath | Conraua_goliath | cathemeral |
| amphibians | Conraua_robusta | Conraua_robusta | cathemeral |
| amphibians | Cophixalus_aenigma | Cophixalus_aenigma | nocturnal |
| amphibians | Cophixalus_australis | Cophixalus_australis | nocturnal |
| amphibians | Cophixalus_balbus | Cophixalus_balbus | nocturnal |
| amphibians | Cophixalus_bombiens | Cophixalus_bombiens | nocturnal |
| amphibians | Cophixalus_caverniphilus | Cophixalus_caverniphilus | diurnal |
| amphibians | Cophixalus_cheesmanae | Cophixalus_cheesmanae | nocturnal |
| amphibians | Cophixalus_concinnus | Cophixalus_concinnus | nocturnal |
| amphibians | Cophixalus_cryptotympanum | Cophixalus_cryptotympanum | nocturnal |
| amphibians | Cophixalus_cupricarenus | Cophixalus_cupricarenus | nocturnal |
| amphibians | Cophixalus_desticans | Cophixalus_desticans | nocturnal |
| amphibians | Cophixalus_exiguus | Cophixalus_exiguus | nocturnal |
| amphibians | Cophixalus_hosmeri | Cophixalus_hosmeri | nocturnal |
| amphibians | Cophixalus_infacetus | Cophixalus_infacetus | nocturnal |
| amphibians | Cophixalus_kaindiensis | Cophixalus_kaindiensis | nocturnal |
| amphibians | Cophixalus_mcdonaldi | Cophixalus_mcdonaldi | nocturnal |
| amphibians | Cophixalus_misimae | Cophixalus_misimae | nocturnal |
| amphibians | Cophixalus_monticola | Cophixalus_monticola | nocturnal |
| amphibians | Cophixalus_neglectus | Cophixalus_neglectus | nocturnal |
| amphibians | Cophixalus_ornatus | Cophixalus_ornatus | nocturnal |
| amphibians | Cophixalus_parkeri | Cophixalus_parkeri | nocturnal |
| amphibians | Cophixalus_riparius | Cophixalus_riparius | nocturnal |
| amphibians | Cophixalus_sphagnicola | Cophixalus_sphagnicola | diurnal |
| amphibians | Cophixalus_tetzlaffi | Cophixalus_tetzlaffi | nocturnal |
| amphibians | Cophixalus_timidus | Cophixalus_timidus | cathemeral |
| amphibians | Cophixalus_variabilis | Cophixalus_variabilis | cathemeral |
| amphibians | Cophixalus_verecundus | Cophixalus_verecundus | nocturnal |
| amphibians | Cophyla_berara | Cophyla_berara | nocturnal |
| amphibians | Cophyla_occultans | Cophyla_occultans | nocturnal |
| amphibians | Cophyla_phyllodactyla | Cophyla_phyllodactyla | nocturnal |
| amphibians | Copiula_guttata | Austrochaperina_guttata | diurnal |

| | | | |
|---|---|---|---|
| amphibians | Cornufer_akarithymus | Platymantis_akarithymus | diurnal |
| amphibians | Cornufer_browni | Platymantis_browni | nocturnal |
| amphibians | Cornufer_latro | Platymantis_latro | nocturnal |
| amphibians | Cornufer_magnus | Platymantis_magnus | nocturnal |
| amphibians | Cornufer_nakanaiorum | Platymantis_nakanaiorum | nocturnal |
| amphibians | Cornufer_nexipus | Platymantis_nexipus | nocturnal |
| amphibians | Cornufer_pelewensis | Platymantis_pelewensis | diurnal |
| amphibians | Cornufer_vitianus | Platymantis_vitianus | cathemeral |
| amphibians | Corythomantis_greeningi | Corythomantis_greeningi | nocturnal |
| amphibians | Craugastor_alfredi | Craugastor_alfredi | nocturnal |
| amphibians | Craugastor_anciano | Craugastor_anciano | cathemeral |
| amphibians | Craugastor_andi | Craugastor_andi | nocturnal |
| amphibians | Craugastor_angelicus | Craugastor_angelicus | nocturnal |
| amphibians | Craugastor_augusti | Craugastor_augusti | nocturnal |
| amphibians | Craugastor_aurilegulus | Craugastor_aurilegulus | cathemeral |
| amphibians | Craugastor_bransfordii | Craugastor_bransfordii | cathemeral |
| amphibians | Craugastor_chac | Craugastor_chac | cathemeral |
| amphibians | Craugastor_charadra | Craugastor_charadra | nocturnal |
| amphibians | Craugastor_chrysozetetes | Craugastor_chrysozetetes | cathemeral |
| amphibians | Craugastor_crassidigitus | Craugastor_crassidigitus | cathemeral |
| amphibians | Craugastor_cruzi | Craugastor_cruzi | diurnal |
| amphibians | Craugastor_cuaquero | Craugastor_cuaquero | nocturnal |
| amphibians | Craugastor_emcelae | Craugastor_emcelae | nocturnal |
| amphibians | Craugastor_emleni | Craugastor_emleni | cathemeral |
| amphibians | Craugastor_epochthidius | Craugastor_epochthidius | cathemeral |
| amphibians | Craugastor_escoces | Craugastor_escoces | nocturnal |
| amphibians | Craugastor_fecundus | Craugastor_fecundus | cathemeral |
| amphibians | Craugastor_fitzingeri | Craugastor_fitzingeri | cathemeral |
| amphibians | Craugastor_fleischmanni | Craugastor_fleischmanni | nocturnal |
| amphibians | Craugastor_gollmeri | Craugastor_gollmeri | diurnal |
| amphibians | Craugastor_laevissimus | Craugastor_laevissimus | cathemeral |
| amphibians | Craugastor_laticeps | Craugastor_laticeps | cathemeral |
| amphibians | Craugastor_lauraster | Craugastor_lauraster | cathemeral |
| amphibians | Craugastor_longirostris | Craugastor_longirostris | nocturnal |

| | | | |
|---|---|---|---|
| amphibians | Craugastor_megacephalus | Craugastor_megacephalus | nocturnal |
| amphibians | Craugastor_melanostictus | Craugastor_melanostictus | nocturnal |
| amphibians | Craugastor_merendonensis | Craugastor_merendonensis | nocturnal |
| amphibians | Craugastor_milesi | Craugastor_milesi | cathemeral |
| amphibians | Craugastor_mimus | Craugastor_mimus | cathemeral |
| amphibians | Craugastor_noblei | Craugastor_noblei | cathemeral |
| amphibians | Craugastor_obesus | Craugastor_obesus | nocturnal |
| amphibians | Craugastor_occidentalis | Craugastor_occidentalis | diurnal |
| amphibians | Craugastor_olanchano | Craugastor_olanchano | cathemeral |
| amphibians | Craugastor_omoaensis | Craugastor_omoaensis | diurnal |
| amphibians | Craugastor_pechorum | Craugastor_pechorum | nocturnal |
| amphibians | Craugastor_persimilis | Craugastor_persimilis | diurnal |
| amphibians | Craugastor_polyptychus | Craugastor_polyptychus | diurnal |
| amphibians | Craugastor_psephosypharus | Craugastor_psephosypharus | nocturnal |
| amphibians | Craugastor_punctariolus | Craugastor_punctariolus | nocturnal |
| amphibians | Craugastor_pygmaeus | Craugastor_pygmaeus | diurnal |
| amphibians | Craugastor_ranoides | Craugastor_ranoides | nocturnal |
| amphibians | Craugastor_rayo | Craugastor_rayo | diurnal |
| amphibians | Craugastor_rhodopis | Craugastor_rhodopis | cathemeral |
| amphibians | Craugastor_rostralis | Craugastor_rostralis | cathemeral |
| amphibians | Craugastor_saltuarius | Craugastor_saltuarius | diurnal |
| amphibians | Craugastor_sandersoni | Craugastor_sandersoni | nocturnal |
| amphibians | Craugastor_stadelmani | Craugastor_stadelmani | cathemeral |
| amphibians | Craugastor_stejnegerianus | Craugastor_stejnegerianus | diurnal |
| amphibians | Craugastor_talamancae | Craugastor_talamancae | nocturnal |
| amphibians | Craugastor_taurus | Craugastor_taurus | nocturnal |
| amphibians | Craugastor_underwoodi | Craugastor_underwoodi | diurnal |
| amphibians | Craugastor_yucatanensis | Craugastor_yucatanensis | nocturnal |
| amphibians | Crinia_fimbriata | Crinia_fimbriata | nocturnal |
| amphibians | Crinia_georgiana | Crinia_georgiana | nocturnal |
| amphibians | Crinia_parinsignifera | Crinia_parinsignifera | cathemeral |
| amphibians | Crinia_signifera | Crinia_signifera | nocturnal |
| amphibians | Crossodactylodes_bokermanni | Crossodactylodes_bokermanni | nocturnal |
| amphibians | Crossodactylodes_izecksohni | Crossodactylodes_izecksohni | nocturnal |

| | | | |
|---|---|---|---|
| amphibians | Crossodactylus_caramaschii | Crossodactylus_caramaschii | diurnal |
| amphibians | Crossodactylus_cyclospinus | Crossodactylus_cyclospinus | diurnal |
| amphibians | Crossodactylus_dantei | Crossodactylus_dantei | diurnal |
| amphibians | Crossodactylus_dispar | Crossodactylus_dispar | diurnal |
| amphibians | Crossodactylus_grandis | Crossodactylus_grandis | diurnal |
| amphibians | Crossodactylus_lutzorum | Crossodactylus_lutzorum | diurnal |
| amphibians | Crossodactylus_schmidti | Crossodactylus_schmidti | cathemeral |
| amphibians | Crossodactylus_trachystomus | Crossodactylus_trachystomus | diurnal |
| amphibians | Cruziohyla_calcarifer | Cruziohyla_calcarifer | cathemeral |
| amphibians | Cruziohyla_craspedopus | Cruziohyla_craspedopus | nocturnal |
| amphibians | Cryptothylax_greshoffii | Cryptothylax_greshoffii | nocturnal |
| amphibians | Ctenophryne_aterrima | Ctenophryne_aterrima | nocturnal |
| amphibians | Ctenophryne_geayi | Ctenophryne_geayi | nocturnal |
| amphibians | Cycloramphus_acangatan | Cycloramphus_acangatan | nocturnal |
| amphibians | Cycloramphus_bolitoglossus | Cycloramphus_bolitoglossus | nocturnal |
| amphibians | Cycloramphus_boraceiensis | Cycloramphus_boraceiensis | nocturnal |
| amphibians | Cycloramphus_brasiliensis | Cycloramphus_brasiliensis | nocturnal |
| amphibians | Cycloramphus_catarinensis | Cycloramphus_catarinensis | nocturnal |
| amphibians | Cycloramphus_dubius | Cycloramphus_dubius | nocturnal |
| amphibians | Cycloramphus_eleutherodactylus | Cycloramphus_eleutherodactylus | cathemeral |
| amphibians | Cycloramphus_faustoi | Cycloramphus_faustoi | nocturnal |
| amphibians | Cycloramphus_fuliginosus | Cycloramphus_fuliginosus | nocturnal |
| amphibians | Cycloramphus_izecksohni | Cycloramphus_izecksohni | nocturnal |
| amphibians | Cycloramphus_juimirim | Cycloramphus_juimirim | nocturnal |
| amphibians | Cycloramphus_lithomimeticus | Cycloramphus_lithomimeticus | nocturnal |
| amphibians | Cycloramphus_lutzorum | Cycloramphus_lutzorum | cathemeral |
| amphibians | Cycloramphus_organensis | Cycloramphus_organensis | nocturnal |
| amphibians | Cycloramphus_parvulus | Zachaenus_parvulus | nocturnal |
| amphibians | Cycloramphus_rhyakonastes | Cycloramphus_rhyakonastes | nocturnal |
| amphibians | Cycloramphus_semipalmatus | Cycloramphus_semipalmatus | nocturnal |
| amphibians | Cynops_ensicauda | Cynops_ensicauda | nocturnal |
| amphibians | Cynops_pyrrhogaster | Cynops_pyrrhogaster | nocturnal |
| amphibians | Dasypops_schirchi | Dasypops_schirchi | nocturnal |
| amphibians | Dendrobates_auratus | Dendrobates_auratus | diurnal |

| | | | |
|---|---|---|---|
| amphibians | Dendrobates_leucomelas | Dendrobates_leucomelas | diurnal |
| amphibians | Dendrobates_tinctorius | Dendrobates_tinctorius | diurnal |
| amphibians | Dendrobates_truncatus | Dendrobates_truncatus | diurnal |
| amphibians | Dendrophryniscus_berthalutzae | Dendrophryniscus_berthalutzae | nocturnal |
| amphibians | Dendrophryniscus_brevipollicatus | Dendrophryniscus_brevipollicatus | cathemeral |
| amphibians | Dendrophryniscus_leucomystax | Dendrophryniscus_leucomystax | nocturnal |
| amphibians | Dendrophryniscus_oreites | Dendrophryniscus_oreites | diurnal |
| amphibians | Dendrophryniscus_proboscideus | Dendrophryniscus_proboscideus | nocturnal |
| amphibians | Dendropsophus_anceps | Dendropsophus_anceps | nocturnal |
| amphibians | Dendropsophus_berthalutzae | Dendropsophus_berthalutzae | nocturnal |
| amphibians | Dendropsophus_bifurcus | Dendropsophus_bifurcus | nocturnal |
| amphibians | Dendropsophus_bipunctatus | Dendropsophus_bipunctatus | nocturnal |
| amphibians | Dendropsophus_bokermanni | Dendropsophus_bokermanni | nocturnal |
| amphibians | Dendropsophus_branneri | Dendropsophus_branneri | nocturnal |
| amphibians | Dendropsophus_brevifrons | Dendropsophus_brevifrons | nocturnal |
| amphibians | Dendropsophus_carnifex | Dendropsophus_carnifex | nocturnal |
| amphibians | Dendropsophus_columbianus | Dendropsophus_columbianus | nocturnal |
| amphibians | Dendropsophus_decipiens | Dendropsophus_decipiens | nocturnal |
| amphibians | Dendropsophus_ebraccatus | Dendropsophus_ebraccatus | nocturnal |
| amphibians | Dendropsophus_elegans | Dendropsophus_elegans | nocturnal |
| amphibians | Dendropsophus_elianeae | Dendropsophus_elianeae | nocturnal |
| amphibians | Dendropsophus_garagoensis | Dendropsophus_garagoensis | nocturnal |
| amphibians | Dendropsophus_giesleri | Dendropsophus_giesleri | nocturnal |
| amphibians | Dendropsophus_haddadi | Dendropsophus_haddadi | nocturnal |
| amphibians | Dendropsophus_haraldschultzi | Dendropsophus_haraldschultzi | nocturnal |
| amphibians | Dendropsophus_jimi | Dendropsophus_jimi | nocturnal |
| amphibians | Dendropsophus_joannae | Dendropsophus_joannae | nocturnal |
| amphibians | Dendropsophus_juliani | Dendropsophus_juliani | nocturnal |
| amphibians | Dendropsophus_leali | Dendropsophus_leali | nocturnal |
| amphibians | Dendropsophus_leucophyllatus | Dendropsophus_leucophyllatus | nocturnal |
| amphibians | Dendropsophus_marmoratus | Dendropsophus_marmoratus | nocturnal |
| amphibians | Dendropsophus_mathiassoni | Dendropsophus_mathiassoni | nocturnal |
| amphibians | Dendropsophus_meridensis | Dendropsophus_meridensis | nocturnal |
| amphibians | Dendropsophus_meridianus | Dendropsophus_meridianus | nocturnal |

| | | | |
|---|---|---|---|
| amphibians | Dendropsophus_microcephalus | Dendropsophus_microcephalus | cathemeral |
| amphibians | Dendropsophus_microps | Dendropsophus_microps | nocturnal |
| amphibians | Dendropsophus_minusculus | Dendropsophus_minusculus | nocturnal |
| amphibians | Dendropsophus_minutus | Dendropsophus_minutus | nocturnal |
| amphibians | Dendropsophus_miyatai | Dendropsophus_miyatai | nocturnal |
| amphibians | Dendropsophus_molitor | Dendropsophus_labialis | nocturnal |
| amphibians | Dendropsophus_nahdereri | Dendropsophus_nahdereri | cathemeral |
| amphibians | Dendropsophus_nanus | Dendropsophus_nanus | cathemeral |
| amphibians | Dendropsophus_novaisi | Dendropsophus_novaisi | nocturnal |
| amphibians | Dendropsophus_oliveirai | Dendropsophus_oliveirai | nocturnal |
| amphibians | Dendropsophus_parviceps | Dendropsophus_parviceps | nocturnal |
| amphibians | Dendropsophus_phlebodes | Dendropsophus_phlebodes | nocturnal |
| amphibians | Dendropsophus_praestans | Dendropsophus_praestans | nocturnal |
| amphibians | Dendropsophus_pseudomeridianus | Dendropsophus_pseudomeridianus | nocturnal |
| amphibians | Dendropsophus_reichlei | Dendropsophus_reichlei | nocturnal |
| amphibians | Dendropsophus_rhodopeplus | Dendropsophus_rhodopeplus | nocturnal |
| amphibians | Dendropsophus_riveroi | Dendropsophus_riveroi | nocturnal |
| amphibians | Dendropsophus_robertmertensi | Dendropsophus_robertmertensi | diurnal |
| amphibians | Dendropsophus_rossalleni | Dendropsophus_rossalleni | nocturnal |
| amphibians | Dendropsophus_rubicundulus | Dendropsophus_rubicundulus | nocturnal |
| amphibians | Dendropsophus_ruschii | Dendropsophus_ruschii | nocturnal |
| amphibians | Dendropsophus_salli | Dendropsophus_salli | nocturnal |
| amphibians | Dendropsophus_sanborni | Dendropsophus_sanborni | nocturnal |
| amphibians | Dendropsophus_sarayacuensis | Dendropsophus_sarayacuensis | nocturnal |
| amphibians | Dendropsophus_sartori | Dendropsophus_sartori | nocturnal |
| amphibians | Dendropsophus_schubarti | Dendropsophus_schubarti | nocturnal |
| amphibians | Dendropsophus_seniculus | Dendropsophus_seniculus | nocturnal |
| amphibians | Dendropsophus_soaresi | Dendropsophus_soaresi | nocturnal |
| amphibians | Dendropsophus_studerae | Dendropsophus_studerae | nocturnal |
| amphibians | Dendropsophus_subocularis | Dendropsophus_subocularis | nocturnal |
| amphibians | Dendropsophus_timbeba | Dendropsophus_timbeba | nocturnal |
| amphibians | Dendropsophus_triangulum | Dendropsophus_triangulum | nocturnal |
| amphibians | Dendropsophus_virolinensis | Dendropsophus_virolinensis | nocturnal |
| amphibians | Dendropsophus_werneri | Dendropsophus_werneri | nocturnal |

| | | | |
|---|---|---|---|
| amphibians | Dermatonotus_muelleri | Dermatonotus_muelleri | nocturnal |
| amphibians | Dermophis_costaricense | Dermophis_costaricense | cathemeral |
| amphibians | Dermophis_mexicanus | Dermophis_mexicanus | nocturnal |
| amphibians | Dermophis_oaxacae | Dermophis_oaxacae | nocturnal |
| amphibians | Desmognathus_aeneus | Desmognathus_aeneus | nocturnal |
| amphibians | Desmognathus_apalachicolae | Desmognathus_apalachicolae | cathemeral |
| amphibians | Desmognathus_auriculatus | Desmognathus_auriculatus | diurnal |
| amphibians | Desmognathus_carolinensis | Desmognathus_carolinensis | diurnal |
| amphibians | Desmognathus_conanti | Desmognathus_conanti | cathemeral |
| amphibians | Desmognathus_fuscus | Desmognathus_fuscus | nocturnal |
| amphibians | Desmognathus_imitator | Desmognathus_imitator | cathemeral |
| amphibians | Desmognathus_marmoratus | Desmognathus_marmoratus | nocturnal |
| amphibians | Desmognathus_monticola | Desmognathus_monticola | nocturnal |
| amphibians | Desmognathus_ochrophaeus | Desmognathus_ochrophaeus | nocturnal |
| amphibians | Desmognathus_ocoee | Desmognathus_ocoee | nocturnal |
| amphibians | Desmognathus_orestes | Desmognathus_orestes | diurnal |
| amphibians | Desmognathus_santeetlah | Desmognathus_santeetlah | cathemeral |
| amphibians | Desmognathus_welteri | Desmognathus_welteri | nocturnal |
| amphibians | Diasporus_anthrax | Diasporus_anthrax | cathemeral |
| amphibians | Diasporus_diastema | Diasporus_diastema | cathemeral |
| amphibians | Diasporus_gularis | Diasporus_gularis | nocturnal |
| amphibians | Diasporus_hylaeformis | Diasporus_hylaeformis | nocturnal |
| amphibians | Diasporus_ventrimaculatus | Diasporus_ventrimaculatus | nocturnal |
| amphibians | Diasporus_vocator | Diasporus_vocator | nocturnal |
| amphibians | Dicamptodon_ensatus | Dicamptodon_ensatus | nocturnal |
| amphibians | Dicamptodon_tenebrosus | Dicamptodon_tenebrosus | nocturnal |
| amphibians | Discoglossus_montalentii | Discoglossus_montalentii | cathemeral |
| amphibians | Discoglossus_pictus | Discoglossus_pictus | cathemeral |
| amphibians | Discoglossus_sardus | Discoglossus_sardus | cathemeral |
| amphibians | Discoglossus_scovazzi | Discoglossus_scovazzi | nocturnal |
| amphibians | Dryophytes_andersonii | Hyla_andersonii | diurnal |
| amphibians | Dryophytes_arenicolor | Hyla_arenicolor | nocturnal |
| amphibians | Dryophytes_avivoca | Hyla_avivoca | nocturnal |
| amphibians | Dryophytes_chrysoscelis | Hyla_chrysoscelis | cathemeral |

| | | | |
|---|---|---|---|
| amphibians | Dryophytes_cinereus | Hyla_cinerea | cathemeral |
| amphibians | Dryophytes_eximius | Hyla_eximia | nocturnal |
| amphibians | Dryophytes_femoralis | Hyla_femoralis | cathemeral |
| amphibians | Dryophytes_gratiosus | Hyla_gratiosa | nocturnal |
| amphibians | Dryophytes_squirellus | Hyla_squirella | nocturnal |
| amphibians | Dryophytes_suweonensis | Hyla_suweonensis | cathemeral |
| amphibians | Dryophytes_versicolor | Hyla_versicolor | nocturnal |
| amphibians | Duellmanohyla_rufioculis | Duellmanohyla_rufioculis | nocturnal |
| amphibians | Duellmanohyla_soralia | Duellmanohyla_soralia | nocturnal |
| amphibians | Duellmanohyla_uranochroa | Duellmanohyla_uranochroa | nocturnal |
| amphibians | Duttaphrynus_beddomii | Duttaphrynus_beddomii | diurnal |
| amphibians | Duttaphrynus_himalayanus | Duttaphrynus_himalayanus | nocturnal |
| amphibians | Duttaphrynus_melanostictus | Duttaphrynus_melanostictus | nocturnal |
| amphibians | Duttaphrynus_microtympanum | Duttaphrynus_microtympanum | nocturnal |
| amphibians | Duttaphrynus_parietalis | Duttaphrynus_parietalis | cathemeral |
| amphibians | Duttaphrynus_scaber | Duttaphrynus_scaber | nocturnal |
| amphibians | Dyscophus_antongilii | Dyscophus_antongilii | cathemeral |
| amphibians | Dyscophus_guineti | Dyscophus_guineti | cathemeral |
| amphibians | Dyscophus_insularis | Dyscophus_insularis | cathemeral |
| amphibians | Echinotriton_andersoni | Echinotriton_andersoni | diurnal |
| amphibians | Echinotriton_chinhaiensis | Echinotriton_chinhaiensis | nocturnal |
| amphibians | Ecnomiohyla_fimbrimembra | Ecnomiohyla_fimbrimembra | nocturnal |
| amphibians | Ecnomiohyla_miliaria | Ecnomiohyla_miliaria | nocturnal |
| amphibians | Ecnomiohyla_rabborum | Ecnomiohyla_rabborum | nocturnal |
| amphibians | Ecnomiohyla_salvaje | Ecnomiohyla_salvaje | cathemeral |
| amphibians | Ecnomiohyla_thysanota | Ecnomiohyla_thysanota | nocturnal |
| amphibians | Ecnomiohyla_valancifer | Ecnomiohyla_valancifer | nocturnal |
| amphibians | Ectopoglossus_atopoglossus | Anomaloglossus_atopoglossus | cathemeral |
| amphibians | Ectopoglossus_confusus | Anomaloglossus_confusus | diurnal |
| amphibians | Edalorhina_perezi | Edalorhina_perezi | cathemeral |
| amphibians | Elachistocleis_cesarii | Elachistocleis_cesarii | nocturnal |
| amphibians | Elachistocleis_erythrogaster | Elachistocleis_erythrogaster | cathemeral |
| amphibians | Elachistocleis_piauiensis | Elachistocleis_piauiensis | diurnal |
| amphibians | Elachistocleis_skotogaster | Elachistocleis_skotogaster | nocturnal |

| | | | |
|---|---|---|---|
| amphibians | Eleutherodactylus_abbotti | Eleutherodactylus_abbotti | diurnal |
| amphibians | Eleutherodactylus_adelus | Eleutherodactylus_adelus | diurnal |
| amphibians | Eleutherodactylus_alcoae | Eleutherodactylus_alcoae | nocturnal |
| amphibians | Eleutherodactylus_amadeus | Eleutherodactylus_amadeus | cathemeral |
| amphibians | Eleutherodactylus_amplinympha | Eleutherodactylus_amplinympha | nocturnal |
| amphibians | Eleutherodactylus_angustidigitorum | Eleutherodactylus_angustidigitorum | diurnal |
| amphibians | Eleutherodactylus_antillensis | Eleutherodactylus_antillensis | nocturnal |
| amphibians | Eleutherodactylus_armstrongi | Eleutherodactylus_armstrongi | nocturnal |
| amphibians | Eleutherodactylus_atkinsi | Eleutherodactylus_atkinsi | nocturnal |
| amphibians | Eleutherodactylus_audanti | Eleutherodactylus_audanti | nocturnal |
| amphibians | Eleutherodactylus_auriculatoides | Eleutherodactylus_auriculatoides | nocturnal |
| amphibians | Eleutherodactylus_auriculatus | Eleutherodactylus_auriculatus | cathemeral |
| amphibians | Eleutherodactylus_bakeri | Eleutherodactylus_bakeri | nocturnal |
| amphibians | Eleutherodactylus_barlagnei | Eleutherodactylus_barlagnei | nocturnal |
| amphibians | Eleutherodactylus_bartonsmithi | Eleutherodactylus_bartonsmithi | nocturnal |
| amphibians | Eleutherodactylus_blairhedgesi | Eleutherodactylus_blairhedgesi | nocturnal |
| amphibians | Eleutherodactylus_bresslerae | Eleutherodactylus_bresslerae | nocturnal |
| amphibians | Eleutherodactylus_caribe | Eleutherodactylus_caribe | nocturnal |
| amphibians | Eleutherodactylus_cattus | Eleutherodactylus_cavernicola | nocturnal |
| amphibians | Eleutherodactylus_cochranae | Eleutherodactylus_cochranae | nocturnal |
| amphibians | Eleutherodactylus_cooki | Eleutherodactylus_cooki | nocturnal |
| amphibians | Eleutherodactylus_coqui | Eleutherodactylus_coqui | nocturnal |
| amphibians | Eleutherodactylus_corona | Eleutherodactylus_corona | nocturnal |
| amphibians | Eleutherodactylus_cundalli | Eleutherodactylus_cundalli | nocturnal |
| amphibians | Eleutherodactylus_cuneatus | Eleutherodactylus_cuneatus | cathemeral |
| amphibians | Eleutherodactylus_cystignathoides | Eleutherodactylus_cystignathoides | cathemeral |
| amphibians | Eleutherodactylus_darlingtoni | Eleutherodactylus_darlingtoni | diurnal |
| amphibians | Eleutherodactylus_dilatus | Eleutherodactylus_dilatus | nocturnal |
| amphibians | Eleutherodactylus_dolomedes | Eleutherodactylus_dolomedes | nocturnal |
| amphibians | Eleutherodactylus_eileenae | Eleutherodactylus_eileenae | nocturnal |
| amphibians | Eleutherodactylus_eneidae | Eleutherodactylus_eneidae | nocturnal |
| amphibians | Eleutherodactylus_etheridgei | Eleutherodactylus_etheridgei | nocturnal |
| amphibians | Eleutherodactylus_feichtingeri | Eleutherodactylus_feichtingeri | crepuscular |
| amphibians | Eleutherodactylus_fowleri | Eleutherodactylus_fowleri | nocturnal |

| | | | |
|---|---|---|---|
| amphibians | Eleutherodactylus_glamyrus | Eleutherodactylus_glamyrus | cathemeral |
| amphibians | Eleutherodactylus_glaphycompus | Eleutherodactylus_glaphycompus | cathemeral |
| amphibians | Eleutherodactylus_gossei | Eleutherodactylus_gossei | nocturnal |
| amphibians | Eleutherodactylus_grahami | Eleutherodactylus_grahami | nocturnal |
| amphibians | Eleutherodactylus_griphus | Eleutherodactylus_griphus | nocturnal |
| amphibians | Eleutherodactylus_gryllus | Eleutherodactylus_gryllus | nocturnal |
| amphibians | Eleutherodactylus_guanahacabibes | Eleutherodactylus_guanahacabibes | nocturnal |
| amphibians | Eleutherodactylus_guantanamera | Eleutherodactylus_guantanamera | nocturnal |
| amphibians | Eleutherodactylus_gundlachi | Eleutherodactylus_gundlachi | nocturnal |
| amphibians | Eleutherodactylus_guttilatus | Eleutherodactylus_guttilatus | nocturnal |
| amphibians | Eleutherodactylus_hedricki | Eleutherodactylus_hedricki | nocturnal |
| amphibians | Eleutherodactylus_iberia | Eleutherodactylus_iberia | cathemeral |
| amphibians | Eleutherodactylus_intermedius | Eleutherodactylus_intermedius | nocturnal |
| amphibians | Eleutherodactylus_interorbitalis | Eleutherodactylus_interorbitalis | nocturnal |
| amphibians | Eleutherodactylus_ionthus | Eleutherodactylus_ionthus | nocturnal |
| amphibians | Eleutherodactylus_jamaicensis | Eleutherodactylus_jamaicensis | nocturnal |
| amphibians | Eleutherodactylus_jasperi | Eleutherodactylus_jasperi | nocturnal |
| amphibians | Eleutherodactylus_johnstonei | Eleutherodactylus_johnstonei | nocturnal |
| amphibians | Eleutherodactylus_juanariveroi | Eleutherodactylus_juanariveroi | nocturnal |
| amphibians | Eleutherodactylus_klinikowskii | Eleutherodactylus_klinikowskii | diurnal |
| amphibians | Eleutherodactylus_lamprotes | Eleutherodactylus_lamprotes | diurnal |
| amphibians | Eleutherodactylus_leberi | Eleutherodactylus_leberi | nocturnal |
| amphibians | Eleutherodactylus_lentus | Eleutherodactylus_lentus | nocturnal |
| amphibians | Eleutherodactylus_leoncei | Eleutherodactylus_leoncei | diurnal |
| amphibians | Eleutherodactylus_leprus | Eleutherodactylus_leprus | diurnal |
| amphibians | Eleutherodactylus_limbatus | Eleutherodactylus_limbatus | cathemeral |
| amphibians | Eleutherodactylus_locustus | Eleutherodactylus_locustus | nocturnal |
| amphibians | Eleutherodactylus_mariposa | Eleutherodactylus_mariposa | nocturnal |
| amphibians | Eleutherodactylus_marnockii | Eleutherodactylus_marnockii | nocturnal |
| amphibians | Eleutherodactylus_martinicensis | Eleutherodactylus_martinicensis | nocturnal |
| amphibians | Eleutherodactylus_maurus | Eleutherodactylus_maurus | nocturnal |
| amphibians | Eleutherodactylus_melacara | Eleutherodactylus_melacara | nocturnal |
| amphibians | Eleutherodactylus_monensis | Eleutherodactylus_monensis | cathemeral |
| amphibians | Eleutherodactylus_montanus | Eleutherodactylus_montanus | nocturnal |

| | | | |
|---|---|---|---|
| amphibians | Eleutherodactylus_nortoni | Eleutherodactylus_nortoni | diurnal |
| amphibians | Eleutherodactylus_nubicola | Eleutherodactylus_nubicola | nocturnal |
| amphibians | Eleutherodactylus_orcutti | Eleutherodactylus_orcutti | nocturnal |
| amphibians | Eleutherodactylus_parapelates | Eleutherodactylus_parapelates | diurnal |
| amphibians | Eleutherodactylus_pictissimus | Eleutherodactylus_pictissimus | diurnal |
| amphibians | Eleutherodactylus_pinchoni | Eleutherodactylus_pinchoni | diurnal |
| amphibians | Eleutherodactylus_pipilans | Eleutherodactylus_pipilans | nocturnal |
| amphibians | Eleutherodactylus_planirostris | Eleutherodactylus_planirostris | nocturnal |
| amphibians | Eleutherodactylus_principalis | Eleutherodactylus_principalis | nocturnal |
| amphibians | Eleutherodactylus_richmondi | Eleutherodactylus_richmondi | nocturnal |
| amphibians | Eleutherodactylus_ricordii | Eleutherodactylus_ricordii | nocturnal |
| amphibians | Eleutherodactylus_rivularis | Eleutherodactylus_rivularis | nocturnal |
| amphibians | Eleutherodactylus_ronaldi | Eleutherodactylus_ronaldi | nocturnal |
| amphibians | Eleutherodactylus_ruthae | Eleutherodactylus_ruthae | cathemeral |
| amphibians | Eleutherodactylus_schmidti | Eleutherodactylus_schmidti | nocturnal |
| amphibians | Eleutherodactylus_schwartzi | Eleutherodactylus_schwartzi | nocturnal |
| amphibians | Eleutherodactylus_simulans | Eleutherodactylus_simulans | nocturnal |
| amphibians | Eleutherodactylus_sisyphodemus | Eleutherodactylus_sisyphodemus | nocturnal |
| amphibians | Eleutherodactylus_tetajulia | Eleutherodactylus_tetajulia | cathemeral |
| amphibians | Eleutherodactylus_thomasi | Eleutherodactylus_thomasi | nocturnal |
| amphibians | Eleutherodactylus_toa | Eleutherodactylus_toa | nocturnal |
| amphibians | Eleutherodactylus_tonyi | Eleutherodactylus_tonyi | diurnal |
| amphibians | Eleutherodactylus_turquinensis | Eleutherodactylus_turquinensis | nocturnal |
| amphibians | Eleutherodactylus_tychathrous | Eleutherodactylus_tychathrous | nocturnal |
| amphibians | Eleutherodactylus_unicolor | Eleutherodactylus_unicolor | nocturnal |
| amphibians | Eleutherodactylus_varians | Eleutherodactylus_varians | nocturnal |
| amphibians | Eleutherodactylus_varleyi | Eleutherodactylus_varleyi | nocturnal |
| amphibians | Eleutherodactylus_weinlandi | Eleutherodactylus_weinlandi | nocturnal |
| amphibians | Eleutherodactylus_zugi | Eleutherodactylus_zugi | nocturnal |
| amphibians | Engystomops_coloradorum | Engystomops_coloradorum | nocturnal |
| amphibians | Engystomops_freibergi | Engystomops_freibergi | nocturnal |
| amphibians | Engystomops_montubio | Engystomops_montubio | nocturnal |
| amphibians | Engystomops_petersi | Engystomops_petersi | cathemeral |
| amphibians | Engystomops_pustulatus | Engystomops_pustulatus | nocturnal |

| | | | |
|---|---|---|---|
| amphibians | Engystomops_pustulosus | Engystomops_pustulosus | nocturnal |
| amphibians | Engystomops_randi | Engystomops_randi | nocturnal |
| amphibians | Ensatina_eschscholtzii | Ensatina_eschscholtzii | nocturnal |
| amphibians | Epicrionops_bicolor | Epicrionops_bicolor | nocturnal |
| amphibians | Epidalea_calamita | Epidalea_calamita | nocturnal |
| amphibians | Epipedobates_anthonyi | Epipedobates_anthonyi | diurnal |
| amphibians | Epipedobates_boulengeri | Epipedobates_boulengeri | diurnal |
| amphibians | Epipedobates_darwinwallacei | Epipedobates_darwinwallacei | diurnal |
| amphibians | Epipedobates_espinosai | Epipedobates_espinosai | diurnal |
| amphibians | Epipedobates_machalilla | Epipedobates_machalilla | diurnal |
| amphibians | Epipedobates_narinensis | Epipedobates_narinensis | diurnal |
| amphibians | Epipedobates_tricolor | Epipedobates_tricolor | diurnal |
| amphibians | Espadarana_andina | Espadarana_andina | nocturnal |
| amphibians | Espadarana_callistomma | Espadarana_callistomma | nocturnal |
| amphibians | Espadarana_durrellorum | Centrolene_durrellorum | nocturnal |
| amphibians | Espadarana_prosoblepon | Espadarana_prosoblepon | nocturnal |
| amphibians | Euparkerella_brasiliensis | Euparkerella_brasiliensis | crepuscular |
| amphibians | Euparkerella_cochranae | Euparkerella_cochranae | crepuscular |
| amphibians | Euparkerella_robusta | Euparkerella_robusta | cathemeral |
| amphibians | Euparkerella_tridactyla | Euparkerella_tridactyla | cathemeral |
| amphibians | Euproctus_montanus | Euproctus_montanus | nocturnal |
| amphibians | Euproctus_platycephalus | Euproctus_platycephalus | diurnal |
| amphibians | Eupsophus_altor | Eupsophus_altor | nocturnal |
| amphibians | Eupsophus_contulmoensis | Eupsophus_contulmoensis | nocturnal |
| amphibians | Eupsophus_roseus | Eupsophus_roseus | nocturnal |
| amphibians | Eurycea_bislineata | Eurycea_bislineata | nocturnal |
| amphibians | Eurycea_guttolineata | Eurycea_guttolineata | nocturnal |
| amphibians | Eurycea_longicauda | Eurycea_longicauda | nocturnal |
| amphibians | Eurycea_lucifuga | Eurycea_lucifuga | nocturnal |
| amphibians | Eurycea_nana | Eurycea_nana | diurnal |
| amphibians | Eurycea_quadridigitata | Eurycea_quadridigitata | diurnal |
| amphibians | Eurycea_spelaea | Eurycea_spelaea | nocturnal |
| amphibians | Eurycea_wilderae | Eurycea_wilderae | nocturnal |
| amphibians | Excidobates_captivus | Excidobates_captivus | diurnal |

| | | | |
|---|---|---|---|
| amphibians | Excidobates_condor | Excidobates_condor | diurnal |
| amphibians | Excidobates_mysteriosus | Excidobates_mysteriosus | diurnal |
| amphibians | Exerodonta_bivocata | Exerodonta_bivocata | nocturnal |
| amphibians | Exerodonta_catracha | Exerodonta_catracha | cathemeral |
| amphibians | Exerodonta_melanomma | Exerodonta_melanomma | nocturnal |
| amphibians | Exerodonta_perkinsi | Exerodonta_perkinsi | nocturnal |
| amphibians | Exerodonta_xera | Exerodonta_xera | nocturnal |
| amphibians | Feihyla_kajau | Feihyla_kajau | nocturnal |
| amphibians | Fejervarya_cancrivora | Fejervarya_cancrivora | nocturnal |
| amphibians | Fejervarya_limnocharis | Fejervarya_limnocharis | cathemeral |
| amphibians | Fejervarya_triora | Fejervarya_triora | nocturnal |
| amphibians | Fejervarya_vittigera | Fejervarya_vittigera | nocturnal |
| amphibians | Firouzophrynus_dhufarensis | Duttaphrynus_dhufarensis | nocturnal |
| amphibians | Firouzophrynus_stomaticus | Duttaphrynus_stomaticus | cathemeral |
| amphibians | Flectonotus_fitzgeraldi | Flectonotus_fitzgeraldi | nocturnal |
| amphibians | Flectonotus_pygmaeus | Flectonotus_pygmaeus | nocturnal |
| amphibians | Fritziana_fissilis | Fritziana_fissilis | nocturnal |
| amphibians | Fritziana_goeldii | Fritziana_goeldii | nocturnal |
| amphibians | Fritziana_ohausi | Fritziana_ohausi | nocturnal |
| amphibians | Frostius_erythrophthalmus | Frostius_erythrophthalmus | nocturnal |
| amphibians | Frostius_pernambucensis | Frostius_pernambucensis | nocturnal |
| amphibians | Gastrophryne_carolinensis | Gastrophryne_carolinensis | nocturnal |
| amphibians | Gastrophryne_elegans | Gastrophryne_elegans | nocturnal |
| amphibians | Gastrophryne_olivacea | Gastrophryne_olivacea | cathemeral |
| amphibians | Gastrotheca_albolineata | Gastrotheca_albolineata | nocturnal |
| amphibians | Gastrotheca_andaquiensis | Gastrotheca_andaquiensis | nocturnal |
| amphibians | Gastrotheca_angustifrons | Gastrotheca_angustifrons | nocturnal |
| amphibians | Gastrotheca_antomia | Gastrotheca_antomia | nocturnal |
| amphibians | Gastrotheca_antoniiochoai | Gastrotheca_antoniiochoai | nocturnal |
| amphibians | Gastrotheca_argenteovirens | Gastrotheca_argenteovirens | nocturnal |
| amphibians | Gastrotheca_aureomaculata | Gastrotheca_aureomaculata | cathemeral |
| amphibians | Gastrotheca_christiani | Gastrotheca_christiani | nocturnal |
| amphibians | Gastrotheca_cornuta | Gastrotheca_cornuta | nocturnal |
| amphibians | Gastrotheca_dendronastes | Gastrotheca_dendronastes | nocturnal |

| | | | |
|---|---|---|---|
| amphibians | Gastrotheca_dunni | Gastrotheca_dunni | nocturnal |
| amphibians | Gastrotheca_dysprosita | Gastrotheca_dysprosita | diurnal |
| amphibians | Gastrotheca_ernestoi | Gastrotheca_ernestoi | nocturnal |
| amphibians | Gastrotheca_espeletia | Gastrotheca_espeletia | cathemeral |
| amphibians | Gastrotheca_fissipes | Gastrotheca_fissipes | nocturnal |
| amphibians | Gastrotheca_fulvorufa | Gastrotheca_fulvorufa | nocturnal |
| amphibians | Gastrotheca_galeata | Gastrotheca_galeata | nocturnal |
| amphibians | Gastrotheca_guentheri | Gastrotheca_guentheri | nocturnal |
| amphibians | Gastrotheca_helenae | Gastrotheca_helenae | nocturnal |
| amphibians | Gastrotheca_litonedis | Gastrotheca_litonedis | nocturnal |
| amphibians | Gastrotheca_longipes | Gastrotheca_longipes | nocturnal |
| amphibians | Gastrotheca_marsupiata | Gastrotheca_marsupiata | nocturnal |
| amphibians | Gastrotheca_megacephala | Gastrotheca_megacephala | nocturnal |
| amphibians | Gastrotheca_microdiscus | Gastrotheca_microdiscus | nocturnal |
| amphibians | Gastrotheca_monticola | Gastrotheca_monticola | cathemeral |
| amphibians | Gastrotheca_nicefori | Gastrotheca_nicefori | nocturnal |
| amphibians | Gastrotheca_orophylax | Gastrotheca_orophylax | nocturnal |
| amphibians | Gastrotheca_ossilaginis | Gastrotheca_ossilaginis | nocturnal |
| amphibians | Gastrotheca_ovifera | Gastrotheca_ovifera | cathemeral |
| amphibians | Gastrotheca_pacchamama | Gastrotheca_pacchamama | diurnal |
| amphibians | Gastrotheca_pachachacae | Gastrotheca_pachachacae | nocturnal |
| amphibians | Gastrotheca_phalarosa | Gastrotheca_phalarosa | diurnal |
| amphibians | Gastrotheca_plumbea | Gastrotheca_plumbea | cathemeral |
| amphibians | Gastrotheca_prasina | Gastrotheca_prasina | nocturnal |
| amphibians | Gastrotheca_pseustes | Gastrotheca_pseustes | nocturnal |
| amphibians | Gastrotheca_psychrophila | Gastrotheca_psychrophila | cathemeral |
| amphibians | Gastrotheca_pulchra | Gastrotheca_pulchra | nocturnal |
| amphibians | Gastrotheca_rebeccae | Gastrotheca_rebeccae | nocturnal |
| amphibians | Gastrotheca_recava | Gastrotheca_recava | nocturnal |
| amphibians | Gastrotheca_riobambae | Gastrotheca_riobambae | nocturnal |
| amphibians | Gastrotheca_testudinea | Gastrotheca_testudinea | nocturnal |
| amphibians | Gastrotheca_trachyceps | Gastrotheca_trachyceps | cathemeral |
| amphibians | Gastrotheca_walkeri | Gastrotheca_walkeri | nocturnal |
| amphibians | Gastrotheca_weinlandii | Gastrotheca_weinlandii | nocturnal |

| | | | |
|---|---|---|---|
| amphibians | Gastrotheca_zeugocystis | Gastrotheca_zeugocystis | nocturnal |
| amphibians | Geobatrachus_walkeri | Geobatrachus_walkeri | cathemeral |
| amphibians | Geotrypetes_seraphini | Geotrypetes_seraphini | nocturnal |
| amphibians | Gephyromantis_ambohitra | Gephyromantis_ambohitra | cathemeral |
| amphibians | Gephyromantis_asper | Gephyromantis_asper | cathemeral |
| amphibians | Gephyromantis_blanci | Gephyromantis_blanci | diurnal |
| amphibians | Gephyromantis_boulengeri | Gephyromantis_boulengeri | cathemeral |
| amphibians | Gephyromantis_corvus | Gephyromantis_corvus | cathemeral |
| amphibians | Gephyromantis_decaryi | Gephyromantis_decaryi | cathemeral |
| amphibians | Gephyromantis_eiselti | Gephyromantis_eiselti | diurnal |
| amphibians | Gephyromantis_enki | Gephyromantis_enki | diurnal |
| amphibians | Gephyromantis_granulatus | Gephyromantis_granulatus | nocturnal |
| amphibians | Gephyromantis_horridus | Gephyromantis_horridus | nocturnal |
| amphibians | Gephyromantis_klemmeri | Gephyromantis_klemmeri | diurnal |
| amphibians | Gephyromantis_leucocephalus | Gephyromantis_leucocephalus | diurnal |
| amphibians | Gephyromantis_leucomaculatus | Gephyromantis_leucomaculatus | nocturnal |
| amphibians | Gephyromantis_luteus | Gephyromantis_luteus | cathemeral |
| amphibians | Gephyromantis_moseri | Gephyromantis_moseri | nocturnal |
| amphibians | Gephyromantis_pseudoasper | Gephyromantis_pseudoasper | cathemeral |
| amphibians | Gephyromantis_ranjomavo | Gephyromantis_ranjomavo | nocturnal |
| amphibians | Gephyromantis_redimitus | Gephyromantis_redimitus | cathemeral |
| amphibians | Gephyromantis_rivicola | Gephyromantis_rivicola | diurnal |
| amphibians | Gephyromantis_runewsweeki | Gephyromantis_runewsweeki | diurnal |
| amphibians | Gephyromantis_salegy | Gephyromantis_salegy | nocturnal |
| amphibians | Gephyromantis_schilfi | Gephyromantis_schilfi | nocturnal |
| amphibians | Gephyromantis_silvanus | Gephyromantis_silvanus | cathemeral |
| amphibians | Gephyromantis_spiniferus | Gephyromantis_spiniferus | diurnal |
| amphibians | Gephyromantis_striatus | Gephyromantis_striatus | nocturnal |
| amphibians | Gephyromantis_tahotra | Gephyromantis_tahotra | cathemeral |
| amphibians | Gephyromantis_tandroka | Gephyromantis_tandroka | nocturnal |
| amphibians | Gephyromantis_thelenae | Gephyromantis_thelenae | diurnal |
| amphibians | Gephyromantis_tschenki | Gephyromantis_tschenki | nocturnal |
| amphibians | Gephyromantis_webbi | Gephyromantis_webbi | cathemeral |
| amphibians | Gephyromantis_zavona | Gephyromantis_zavona | nocturnal |

| | | | |
|---|---|---|---|
| amphibians | Glandirana_rugosa | Rugosa_rugosa | nocturnal |
| amphibians | Glyphoglossus_guttulatus | Calluella_guttulata | nocturnal |
| amphibians | Glyphoglossus_molossus | Glyphoglossus_molossus | nocturnal |
| amphibians | Gracixalus_supercornutus | Gracixalus_supercornutus | nocturnal |
| amphibians | Guibemantis_bicalcaratus | Guibemantis_bicalcaratus | diurnal |
| amphibians | Guibemantis_depressiceps | Guibemantis_depressiceps | nocturnal |
| amphibians | Guibemantis_kathrinae | Guibemantis_kathrinae | nocturnal |
| amphibians | Guibemantis_liber | Guibemantis_liber | cathemeral |
| amphibians | Guibemantis_pulcher | Guibemantis_pulcher | cathemeral |
| amphibians | Guibemantis_punctatus | Guibemantis_punctatus | diurnal |
| amphibians | Guibemantis_timidus | Guibemantis_timidus | nocturnal |
| amphibians | Guibemantis_tornieri | Guibemantis_tornieri | nocturnal |
| amphibians | Gymnopis_multiplicata | Gymnopis_multiplicata | cathemeral |
| amphibians | Gyrinophilus_palleucus | Gyrinophilus_palleucus | nocturnal |
| amphibians | Gyrinophilus_porphyriticus | Gyrinophilus_porphyriticus | nocturnal |
| amphibians | Haddadus_aramunha | Haddadus_aramunha | nocturnal |
| amphibians | Haddadus_binotatus | Haddadus_binotatus | cathemeral |
| amphibians | Hadromophryne_natalensis | Hadromophryne_natalensis | nocturnal |
| amphibians | Hamptophryne_boliviana | Hamptophryne_boliviana | cathemeral |
| amphibians | Heleioporus_australiacus | Heleioporus_australiacus | nocturnal |
| amphibians | Heleophryne_orientalis | Heleophryne_orientalis | cathemeral |
| amphibians | Heleophryne_purcelli | Heleophryne_purcelli | diurnal |
| amphibians | Hemidactylium_scutatum | Hemidactylium_scutatum | diurnal |
| amphibians | Hemiphractus_bubalus | Hemiphractus_bubalus | nocturnal |
| amphibians | Hemiphractus_fasciatus | Hemiphractus_fasciatus | cathemeral |
| amphibians | Hemiphractus_helioi | Hemiphractus_helioi | nocturnal |
| amphibians | Hemiphractus_johnsoni | Hemiphractus_johnsoni | nocturnal |
| amphibians | Hemiphractus_proboscideus | Hemiphractus_proboscideus | cathemeral |
| amphibians | Hemiphractus_scutatus | Hemiphractus_scutatus | cathemeral |
| amphibians | Hemisus_guttatus | Hemisus_guttatus | nocturnal |
| amphibians | Hemisus_marmoratus | Hemisus_marmoratus | nocturnal |
| amphibians | Heterixalus_andrakata | Heterixalus_andrakata | nocturnal |
| amphibians | Heterixalus_betsileo | Heterixalus_betsileo | nocturnal |
| amphibians | Heterixalus_boettgeri | Heterixalus_boettgeri | cathemeral |

| | | | |
|---|---|---|---|
| amphibians | Heterixalus_carbonei | Heterixalus_carbonei | nocturnal |
| amphibians | Heterixalus_luteostriatus | Heterixalus_luteostriatus | nocturnal |
| amphibians | Heterixalus_madagascariensis | Heterixalus_madagascariensis | cathemeral |
| amphibians | Heterixalus_punctatus | Heterixalus_punctatus | cathemeral |
| amphibians | Heterixalus_tricolor | Heterixalus_tricolor | cathemeral |
| amphibians | Heterixalus_variabilis | Heterixalus_variabilis | nocturnal |
| amphibians | Hildebrandtia_macrotympanum | Hildebrandtia_macrotympanum | nocturnal |
| amphibians | Hildebrandtia_ornata | Hildebrandtia_ornata | cathemeral |
| amphibians | Holoaden_luederwaldti | Holoaden_luederwaldti | nocturnal |
| amphibians | Holoaden_pholeter | Holoaden_pholeter | nocturnal |
| amphibians | Hoplobatrachus_crassus | Hoplobatrachus_crassus | nocturnal |
| amphibians | Hoplobatrachus_occipitalis | Hoplobatrachus_occipitalis | cathemeral |
| amphibians | Hoplobatrachus_tigerinus | Hoplobatrachus_tigerinus | cathemeral |
| amphibians | Hoplophryne_rogersi | Hoplophryne_rogersi | diurnal |
| amphibians | Huia_cavitympanum | Huia_cavitympanum | nocturnal |
| amphibians | Hyalinobatrachium_aureoguttatum | Hyalinobatrachium_aureoguttatum | nocturnal |
| amphibians | Hyalinobatrachium_cappellei | Hyalinobatrachium_cappellei | nocturnal |
| amphibians | Hyalinobatrachium_carlesvilai | Hyalinobatrachium_carlesvilai | nocturnal |
| amphibians | Hyalinobatrachium_chirripoi | Hyalinobatrachium_chirripoi | nocturnal |
| amphibians | Hyalinobatrachium_colymbiphyllum | Hyalinobatrachium_colymbiphyllum | nocturnal |
| amphibians | Hyalinobatrachium_fleischmanni | Hyalinobatrachium_fleischmanni | nocturnal |
| amphibians | Hyalinobatrachium_guairarepanense | Hyalinobatrachium_guairarepanense | nocturnal |
| amphibians | Hyalinobatrachium_mondolfii | Hyalinobatrachium_mondolfii | cathemeral |
| amphibians | Hyalinobatrachium_orientale | Hyalinobatrachium_orientale | nocturnal |
| amphibians | Hyalinobatrachium_orocostale | Hyalinobatrachium_orocostale | nocturnal |
| amphibians | Hyalinobatrachium_pallidum | Hyalinobatrachium_pallidum | nocturnal |
| amphibians | Hyalinobatrachium_pellucidum | Hyalinobatrachium_pellucidum | nocturnal |
| amphibians | Hyalinobatrachium_talamancae | Hyalinobatrachium_talamancae | nocturnal |
| amphibians | Hyalinobatrachium_tatayoi | Hyalinobatrachium_tatayoi | nocturnal |
| amphibians | Hyalinobatrachium_taylori | Hyalinobatrachium_taylori | nocturnal |
| amphibians | Hyalinobatrachium_tricolor | Hyalinobatrachium_tricolor | nocturnal |
| amphibians | Hyalinobatrachium_valerioi | Hyalinobatrachium_valerioi | nocturnal |
| amphibians | Hydrolaetare_dantasi | Hydrolaetare_dantasi | cathemeral |
| amphibians | Hydrolaetare_schmidti | Hydrolaetare_schmidti | nocturnal |

| | | | |
|---|---|---|---|
| amphibians | Hydromantes_shastae | Hydromantes_shastae | nocturnal |
| amphibians | Hyla_annectans | Hyla_annectans | nocturnal |
| amphibians | Hyla_arborea | Hyla_arborea | cathemeral |
| amphibians | Hyla_chinensis | Hyla_chinensis | nocturnal |
| amphibians | Hyla_felixarabica | Hyla_felixarabica | nocturnal |
| amphibians | Hyla_meridionalis | Hyla_meridionalis | cathemeral |
| amphibians | Hyla_orientalis | Hyla_orientalis | cathemeral |
| amphibians | Hyla_sanchiangensis | Hyla_sanchiangensis | nocturnal |
| amphibians | Hyla_sarda | Hyla_sarda | cathemeral |
| amphibians | Hyla_savignyi | Hyla_savignyi | cathemeral |
| amphibians | Hylarana_aurantiaca | Hylarana_aurantiaca | nocturnal |
| amphibians | Hylarana_chitwanensis | Hylarana_chitwanensis | cathemeral |
| amphibians | Hylarana_erythraea | Hylarana_erythraea | cathemeral |
| amphibians | Hylarana_gracilis | Hylarana_gracilis | cathemeral |
| amphibians | Hylarana_latouchii | Hylarana_latouchii | diurnal |
| amphibians | Hylarana_macrodactyla | Hylarana_macrodactyla | nocturnal |
| amphibians | Hylarana_megalonesa | Hylarana_megalonesa | nocturnal |
| amphibians | Hylarana_nigrovittata | Hylarana_nigrovittata | cathemeral |
| amphibians | Hylarana_taipehensis | Hylarana_taipehensis | cathemeral |
| amphibians | Hylarana_temporalis | Hylarana_temporalis | nocturnal |
| amphibians | Hylodes_amnicola | Hylodes_amnicola | diurnal |
| amphibians | Hylodes_asper | Hylodes_asper | diurnal |
| amphibians | Hylodes_babax | Hylodes_babax | diurnal |
| amphibians | Hylodes_cardosoi | Hylodes_cardosoi | diurnal |
| amphibians | Hylodes_charadranaetes | Hylodes_charadranaetes | diurnal |
| amphibians | Hylodes_dactylocinus | Hylodes_dactylocinus | diurnal |
| amphibians | Hylodes_heyeri | Hylodes_heyeri | diurnal |
| amphibians | Hylodes_lateristrigatus | Hylodes_lateristrigatus | diurnal |
| amphibians | Hylodes_magalhaesi | Hylodes_magalhaesi | diurnal |
| amphibians | Hylodes_meridionalis | Hylodes_meridionalis | diurnal |
| amphibians | Hylodes_nasus | Hylodes_nasus | diurnal |
| amphibians | Hylodes_ornatus | Hylodes_ornatus | diurnal |
| amphibians | Hylodes_otavioi | Hylodes_otavioi | diurnal |
| amphibians | Hylodes_perere | Hylodes_perere | diurnal |

| | | | |
|---|---|---|---|
| amphibians | Hylodes_perplicatus | Hylodes_perplicatus | diurnal |
| amphibians | Hylodes_phyllodes | Hylodes_phyllodes | diurnal |
| amphibians | Hylodes_pipilans | Hylodes_pipilans | diurnal |
| amphibians | Hylodes_regius | Hylodes_regius | diurnal |
| amphibians | Hylodes_sazimai | Hylodes_sazimai | diurnal |
| amphibians | Hylodes_uai | Hylodes_uai | diurnal |
| amphibians | Hylomantis_aspera | Hylomantis_aspera | nocturnal |
| amphibians | Hylomantis_granulosa | Hylomantis_granulosa | nocturnal |
| amphibians | Hylophorbus_rufescens | Hylophorbus_rufescens | nocturnal |
| amphibians | Hylorina_sylvatica | Hylorina_sylvatica | cathemeral |
| amphibians | Hyloscirtus_albopunctulatus | Hyloscirtus_albopunctulatus | nocturnal |
| amphibians | Hyloscirtus_alytolylax | Hyloscirtus_alytolylax | nocturnal |
| amphibians | Hyloscirtus_colymba | Hyloscirtus_colymba | nocturnal |
| amphibians | Hyloscirtus_lascinius | Hyloscirtus_lascinius | nocturnal |
| amphibians | Hyloscirtus_lindae | Hyloscirtus_lindae | nocturnal |
| amphibians | Hyloscirtus_pacha | Hyloscirtus_pacha | nocturnal |
| amphibians | Hyloscirtus_palmeri | Hyloscirtus_palmeri | nocturnal |
| amphibians | Hyloscirtus_pantostictus | Hyloscirtus_pantostictus | nocturnal |
| amphibians | Hyloscirtus_phyllognathus | Hyloscirtus_phyllognathus | nocturnal |
| amphibians | Hyloscirtus_piceigularis | Hyloscirtus_piceigularis | cathemeral |
| amphibians | Hyloscirtus_platydactylus | Hyloscirtus_platydactylus | nocturnal |
| amphibians | Hyloscirtus_princecharlesi | Hyloscirtus_princecharlesi | nocturnal |
| amphibians | Hyloscirtus_psarolaimus | Hyloscirtus_psarolaimus | nocturnal |
| amphibians | Hyloscirtus_ptychodactylus | Hyloscirtus_ptychodactylus | cathemeral |
| amphibians | Hyloscirtus_sarampiona | Hyloscirtus_sarampiona | nocturnal |
| amphibians | Hyloscirtus_staufferorum | Hyloscirtus_staufferorum | nocturnal |
| amphibians | Hyloscirtus_torrenticola | Hyloscirtus_torrenticola | nocturnal |
| amphibians | Hyloxalus_alessandroi | Allobates_alessandroi | diurnal |
| amphibians | Hyloxalus_anthracinus | Hyloxalus_anthracinus | diurnal |
| amphibians | Hyloxalus_awa | Hyloxalus_awa | diurnal |
| amphibians | Hyloxalus_bocagei | Hyloxalus_bocagei | diurnal |
| amphibians | Hyloxalus_breviquartus | Hyloxalus_breviquartus | diurnal |
| amphibians | Hyloxalus_craspedoceps | Hyloxalus_craspedoceps | diurnal |
| amphibians | Hyloxalus_delatorreae | Hyloxalus_delatorreae | diurnal |

| | | | |
|---|---|---|---|
| amphibians | Hyloxalus_elachyhistus | Hyloxalus_elachyhistus | cathemeral |
| amphibians | Hyloxalus_eleutherodactylus | Hyloxalus_eleutherodactylus | diurnal |
| amphibians | Hyloxalus_fascianigrus | Hyloxalus_fascianigrus | diurnal |
| amphibians | Hyloxalus_fuliginosus | Hyloxalus_fuliginosus | diurnal |
| amphibians | Hyloxalus_infraguttatus | Hyloxalus_infraguttatus | diurnal |
| amphibians | Hyloxalus_insulatus | Hyloxalus_insulatus | diurnal |
| amphibians | Hyloxalus_italoi | Hyloxalus_italoi | diurnal |
| amphibians | Hyloxalus_lehmanni | Hyloxalus_lehmanni | diurnal |
| amphibians | Hyloxalus_leucophaeus | Hyloxalus_leucophaeus | diurnal |
| amphibians | Hyloxalus_maculosus | Hyloxalus_maculosus | diurnal |
| amphibians | Hyloxalus_mittermeieri | Hyloxalus_mittermeieri | diurnal |
| amphibians | Hyloxalus_mystax | Hyloxalus_mystax | diurnal |
| amphibians | Hyloxalus_nexipus | Hyloxalus_nexipus | diurnal |
| amphibians | Hyloxalus_patitae | Hyloxalus_patitae | diurnal |
| amphibians | Hyloxalus_picachos | Allobates_picachos | diurnal |
| amphibians | Hyloxalus_pinguis | Hyloxalus_pinguis | diurnal |
| amphibians | Hyloxalus_pulchellus | Hyloxalus_pulchellus | diurnal |
| amphibians | Hyloxalus_pulcherrimus | Hyloxalus_pulcherrimus | diurnal |
| amphibians | Hyloxalus_ramosi | Hyloxalus_ramosi | diurnal |
| amphibians | Hyloxalus_saltuarius | Hyloxalus_saltuarius | diurnal |
| amphibians | Hyloxalus_sauli | Hyloxalus_sauli | diurnal |
| amphibians | Hyloxalus_shuar | Hyloxalus_shuar | cathemeral |
| amphibians | Hyloxalus_sordidatus | Hyloxalus_sordidatus | diurnal |
| amphibians | Hyloxalus_sylvaticus | Hyloxalus_sylvaticus | diurnal |
| amphibians | Hyloxalus_toachi | Hyloxalus_toachi | diurnal |
| amphibians | Hyloxalus_vertebralis | Hyloxalus_vertebralis | diurnal |
| amphibians | Hyloxalus_yasuni | Hyloxalus_yasuni | diurnal |
| amphibians | Hymenochirus_boettgeri | Hymenochirus_boettgeri | nocturnal |
| amphibians | Hynobius_boulengeri | Hynobius_boulengeri | nocturnal |
| amphibians | Hynobius_dunni | Hynobius_dunni | nocturnal |
| amphibians | Hynobius_formosanus | Hynobius_formosanus | nocturnal |
| amphibians | Hynobius_glacialis | Hynobius_glacialis | nocturnal |
| amphibians | Hynobius_kimurae | Hynobius_kimurae | cathemeral |
| amphibians | Hynobius_leechii | Hynobius_leechii | nocturnal |

| | | | |
|---|---|---|---|
| amphibians | Hynobius_naevius | Hynobius_naevius | cathemeral |
| amphibians | Hynobius_nebulosus | Hynobius_nebulosus | nocturnal |
| amphibians | Hynobius_nigrescens | Hynobius_nigrescens | nocturnal |
| amphibians | Hynobius_quelpaertensis | Hynobius_quelpaertensis | diurnal |
| amphibians | Hynobius_retardatus | Hynobius_retardatus | nocturnal |
| amphibians | Hynobius_sonani | Hynobius_sonani | nocturnal |
| amphibians | Hynobius_tokyoensis | Hynobius_tokyoensis | cathemeral |
| amphibians | Hyperolius_castaneus | Hyperolius_castaneus | nocturnal |
| amphibians | Hyperolius_cinereus | Hyperolius_cinereus | diurnal |
| amphibians | Hyperolius_cinnamomeoventris | Hyperolius_cinnamomeoventris | nocturnal |
| amphibians | Hyperolius_cystocandicans | Hyperolius_cystocandicans | nocturnal |
| amphibians | Hyperolius_discodactylus | Hyperolius_discodactylus | diurnal |
| amphibians | Hyperolius_horstockii | Hyperolius_horstockii | diurnal |
| amphibians | Hyperolius_kivuensis | Hyperolius_kivuensis | nocturnal |
| amphibians | Hyperolius_langi | Hyperolius_langi | nocturnal |
| amphibians | Hyperolius_lateralis | Hyperolius_lateralis | nocturnal |
| amphibians | Hyperolius_laurenti | Hyperolius_laurenti | nocturnal |
| amphibians | Hyperolius_montanus | Hyperolius_montanus | nocturnal |
| amphibians | Hyperolius_mosaicus | Hyperolius_mosaicus | nocturnal |
| amphibians | Hyperolius_nasutus | Hyperolius_nasutus | nocturnal |
| amphibians | Hyperolius_nitidulus | Hyperolius_nitidulus | nocturnal |
| amphibians | Hyperolius_pardalis | Hyperolius_pardalis | nocturnal |
| amphibians | Hyperolius_phantasticus | Hyperolius_phantasticus | nocturnal |
| amphibians | Hyperolius_puncticulatus | Hyperolius_puncticulatus | nocturnal |
| amphibians | Hyperolius_pusillus | Hyperolius_pusillus | nocturnal |
| amphibians | Hyperolius_spinigularis | Hyperolius_spinigularis | nocturnal |
| amphibians | Hyperolius_tuberculatus | Hyperolius_tuberculatus | nocturnal |
| amphibians | Hyperolius_tuberilinguis | Hyperolius_tuberilinguis | nocturnal |
| amphibians | Hyperolius_veithi | Hyperolius_veithi | nocturnal |
| amphibians | Hyperolius_viridiflavus | Hyperolius_viridiflavus | cathemeral |
| amphibians | Hypopachus_barberi | Hypopachus_barberi | nocturnal |
| amphibians | Hypopachus_pictiventris | Hypopachus_pictiventris | nocturnal |
| amphibians | Hypopachus_variolosus | Hypopachus_variolosus | nocturnal |
| amphibians | Hypselotriton_fudingensis | Cynops_fudingensis | diurnal |

| | | | |
|---|---|---|---|
| amphibians | Ichthyophis_beddomei | Ichthyophis_beddomei | nocturnal |
| amphibians | Ichthyophis_glutinosus | Ichthyophis_glutinosus | nocturnal |
| amphibians | Ichthyophis_javanicus | Ichthyophis_javanicus | nocturnal |
| amphibians | Ichthyophis_sikkimensis | Ichthyophis_sikkimensis | nocturnal |
| amphibians | Ichthyophis_tricolor | Ichthyophis_tricolor | nocturnal |
| amphibians | Ichthyosaura_alpestris | Ichthyosaura_alpestris | cathemeral |
| amphibians | Incilius_alvarius | Incilius_alvarius | nocturnal |
| amphibians | Incilius_aucoinae | Incilius_aucoinae | nocturnal |
| amphibians | Incilius_campbelli | Incilius_campbelli | diurnal |
| amphibians | Incilius_coccifer | Incilius_coccifer | cathemeral |
| amphibians | Incilius_coniferus | Incilius_coniferus | nocturnal |
| amphibians | Incilius_epioticus | Incilius_epioticus | diurnal |
| amphibians | Incilius_fastidiosus | Incilius_fastidiosus | diurnal |
| amphibians | Incilius_holdridgei | Incilius_holdridgei | diurnal |
| amphibians | Incilius_luetkenii | Incilius_luetkenii | nocturnal |
| amphibians | Incilius_mazatlanensis | Incilius_mazatlanensis | cathemeral |
| amphibians | Incilius_melanochlorus | Incilius_melanochlorus | nocturnal |
| amphibians | Incilius_nebulifer | Incilius_nebulifer | nocturnal |
| amphibians | Incilius_periglenes | Incilius_periglenes | diurnal |
| amphibians | Incilius_valliceps | Incilius_valliceps | cathemeral |
| amphibians | Indirana_brachytarsus | Indirana_brachytarsus | nocturnal |
| amphibians | Indirana_gundia | Indirana_gundia | cathemeral |
| amphibians | Indirana_leithii | Indirana_leithii | cathemeral |
| amphibians | Indirana_semipalmata | Indirana_semipalmata | cathemeral |
| amphibians | Ingerophrynus_biporcatus | Ingerophrynus_biporcatus | nocturnal |
| amphibians | Ingerophrynus_divergens | Ingerophrynus_divergens | nocturnal |
| amphibians | Ingerophrynus_galeatus | Ingerophrynus_galeatus | nocturnal |
| amphibians | Ingerophrynus_ledongensis | Ingerophrynus_ledongensis | diurnal |
| amphibians | Ingerophrynus_parvus | Ingerophrynus_parvus | cathemeral |
| amphibians | Insuetophrynus_acarpicus | Insuetophrynus_acarpicus | nocturnal |
| amphibians | Ischnocnema_bolbodactyla | Ischnocnema_bolbodactyla | cathemeral |
| amphibians | Ischnocnema_concolor | Ischnocnema_concolor | nocturnal |
| amphibians | Ischnocnema_guentheri | Ischnocnema_guentheri | nocturnal |
| amphibians | Ischnocnema_henselii | Ischnocnema_henselii | cathemeral |

| | | | |
|---|---|---|---|
| amphibians | Ischnocnema_holti | Ischnocnema_holti | cathemeral |
| amphibians | Ischnocnema_juipoca | Ischnocnema_juipoca | cathemeral |
| amphibians | Ischnocnema_manezinho | Ischnocnema_manezinho | nocturnal |
| amphibians | Ischnocnema_octavioi | Ischnocnema_octavioi | nocturnal |
| amphibians | Ischnocnema_oea | Ischnocnema_oea | cathemeral |
| amphibians | Ischnocnema_parva | Ischnocnema_parva | cathemeral |
| amphibians | Ischnocnema_penaxavantinho | Ischnocnema_penaxavantinho | crepuscular |
| amphibians | Ischnocnema_randorum | Ischnocnema_randorum | nocturnal |
| amphibians | Ischnocnema_sambaqui | Ischnocnema_sambaqui | nocturnal |
| amphibians | Ischnocnema_spanios | Ischnocnema_spanios | diurnal |
| amphibians | Ischnocnema_verrucosa | Ischnocnema_verrucosa | nocturnal |
| amphibians | Ischnocnema_vizottoi | Ischnocnema_vizottoi | nocturnal |
| amphibians | Isthmohyla_angustilineata | Isthmohyla_angustilineata | nocturnal |
| amphibians | Isthmohyla_calypsa | Isthmohyla_calypsa | nocturnal |
| amphibians | Isthmohyla_infucata | Isthmohyla_infucata | nocturnal |
| amphibians | Isthmohyla_insolita | Isthmohyla_insolita | nocturnal |
| amphibians | Isthmohyla_lancasteri | Isthmohyla_lancasteri | nocturnal |
| amphibians | Isthmohyla_picadoi | Isthmohyla_picadoi | nocturnal |
| amphibians | Isthmohyla_pictipes | Isthmohyla_pictipes | nocturnal |
| amphibians | Isthmohyla_pseudopuma | Isthmohyla_pseudopuma | nocturnal |
| amphibians | Isthmohyla_rivularis | Isthmohyla_rivularis | nocturnal |
| amphibians | Isthmohyla_tica | Isthmohyla_tica | crepuscular |
| amphibians | Isthmohyla_xanthosticta | Isthmohyla_xanthosticta | nocturnal |
| amphibians | Isthmohyla_zeteki | Isthmohyla_zeteki | nocturnal |
| amphibians | Itapotihyla_langsdorffii | Itapotihyla_langsdorffii | cathemeral |
| amphibians | Julianus_uruguayus | Scinax_uruguayus | nocturnal |
| amphibians | Kalophrynus_baluensis | Kalophrynus_baluensis | cathemeral |
| amphibians | Kalophrynus_calciphilus | Kalophrynus_calciphilus | cathemeral |
| amphibians | Kalophrynus_interlineatus | Kalophrynus_interlineatus | nocturnal |
| amphibians | Kalophrynus_nubicola | Kalophrynus_nubicola | diurnal |
| amphibians | Kalophrynus_pleurostigma | Kalophrynus_pleurostigma | cathemeral |
| amphibians | Kalophrynus_yongi | Kalophrynus_yongi | nocturnal |
| amphibians | Kaloula_baleata | Kaloula_baleata | cathemeral |
| amphibians | Kaloula_borealis | Kaloula_borealis | nocturnal |

| | | | |
|---|---|---|---|
| amphibians | Kaloula_conjuncta | Kaloula_conjuncta | nocturnal |
| amphibians | Kaloula_indochinensis | Kaloula_indochinensis | nocturnal |
| amphibians | Kaloula_picta | Kaloula_picta | nocturnal |
| amphibians | Kaloula_pulchra | Kaloula_pulchra | nocturnal |
| amphibians | Kaloula_walteri | Kaloula_walteri | cathemeral |
| amphibians | Kassina_arboricola | Kassina_arboricola | nocturnal |
| amphibians | Kassina_cassinoides | Kassina_cassinoides | nocturnal |
| amphibians | Kassina_cochranae | Kassina_cochranae | nocturnal |
| amphibians | Kassina_decorata | Kassina_decorata | nocturnal |
| amphibians | Kassina_fusca | Kassina_fusca | nocturnal |
| amphibians | Kassina_jozani | Kassina_jozani | nocturnal |
| amphibians | Kassina_kuvangensis | Kassina_kuvangensis | nocturnal |
| amphibians | Kassina_lamottei | Kassina_lamottei | nocturnal |
| amphibians | Kassina_maculifer | Kassina_maculifer | nocturnal |
| amphibians | Kassina_maculosa | Kassina_maculosa | nocturnal |
| amphibians | Kassina_schioetzi | Kassina_schioetzi | nocturnal |
| amphibians | Kassina_senegalensis | Kassina_senegalensis | cathemeral |
| amphibians | Kassina_wazae | Kassina_wazae | nocturnal |
| amphibians | Kurixalus_appendiculatus | Kurixalus_appendiculatus | nocturnal |
| amphibians | Kurixalus_banaensis | Kurixalus_banaensis | nocturnal |
| amphibians | Kurixalus_bisacculus | Kurixalus_bisacculus | nocturnal |
| amphibians | Kurixalus_eiffingeri | Kurixalus_eiffingeri | nocturnal |
| amphibians | Laliostoma_labrosum | Laliostoma_labrosum | nocturnal |
| amphibians | Laotriton_laoensis | Laotriton_laoensis | diurnal |
| amphibians | Leiopelma_archeyi | Leiopelma_archeyi | nocturnal |
| amphibians | Leiopelma_hochstetteri | Leiopelma_hochstetteri | nocturnal |
| amphibians | Lepidobatrachus_asper | Lepidobatrachus_asper | nocturnal |
| amphibians | Lepidobatrachus_laevis | Lepidobatrachus_laevis | nocturnal |
| amphibians | Lepidobatrachus_llanensis | Lepidobatrachus_llanensis | nocturnal |
| amphibians | Leptobrachella_aerea | Leptolalax_aereus | nocturnal |
| amphibians | Leptobrachella_arayai | Leptolalax_arayai | nocturnal |
| amphibians | Leptobrachella_baluensis | Leptobrachella_baluensis | nocturnal |
| amphibians | Leptobrachella_crocea | Leptolalax_croceus | nocturnal |
| amphibians | Leptobrachella_fuliginosa | Leptolalax_fuliginosus | nocturnal |

| | | | |
|---|---|---|---|
| amphibians | Leptobrachella_kajangensis | Leptolalax_kajangensis | nocturnal |
| amphibians | Leptobrachella_kecil | Leptolalax_kecil | nocturnal |
| amphibians | Leptobrachella_liui | Leptolalax_liui | nocturnal |
| amphibians | Leptobrachella_melanoleuca | Leptolalax_melanoleucus | nocturnal |
| amphibians | Leptobrachella_melica | Leptolalax_melicus | nocturnal |
| amphibians | Leptobrachella_sola | Leptolalax_solus | nocturnal |
| amphibians | Leptobrachella_ventripunctata | Leptolalax_ventripunctatus | nocturnal |
| amphibians | Leptobrachium_abbotti | Leptobrachium_abbotti | nocturnal |
| amphibians | Leptobrachium_banae | Leptobrachium_banae | nocturnal |
| amphibians | Leptobrachium_boringii | Leptobrachium_boringii | nocturnal |
| amphibians | Leptobrachium_hainanense | Leptobrachium_hainanense | nocturnal |
| amphibians | Leptobrachium_hasseltii | Leptobrachium_hasseltii | nocturnal |
| amphibians | Leptobrachium_huashen | Leptobrachium_huashen | nocturnal |
| amphibians | Leptobrachium_liui | Leptobrachium_liui | nocturnal |
| amphibians | Leptobrachium_montanum | Leptobrachium_montanum | cathemeral |
| amphibians | Leptobrachium_mouhoti | Leptobrachium_mouhoti | nocturnal |
| amphibians | Leptobrachium_pullum | Leptobrachium_pullum | nocturnal |
| amphibians | Leptobrachium_smithi | Leptobrachium_smithi | nocturnal |
| amphibians | Leptodactylodon_ventrimarmoratus | Leptodactylodon_ventrimarmoratus | diurnal |
| amphibians | Leptodactylus_albilabris | Leptodactylus_albilabris | diurnal |
| amphibians | Leptodactylus_bolivianus | Leptodactylus_bolivianus | cathemeral |
| amphibians | Leptodactylus_bufonius | Leptodactylus_bufonius | nocturnal |
| amphibians | Leptodactylus_caatingae | Leptodactylus_caatingae | cathemeral |
| amphibians | Leptodactylus_camaquara | Leptodactylus_camaquara | diurnal |
| amphibians | Leptodactylus_cunicularius | Leptodactylus_cunicularius | nocturnal |
| amphibians | Leptodactylus_cupreus | Leptodactylus_cupreus | nocturnal |
| amphibians | Leptodactylus_didymus | Leptodactylus_didymus | cathemeral |
| amphibians | Leptodactylus_diedrus | Leptodactylus_diedrus | cathemeral |
| amphibians | Leptodactylus_discodactylus | Leptodactylus_discodactylus | cathemeral |
| amphibians | Leptodactylus_fallax | Leptodactylus_fallax | cathemeral |
| amphibians | Leptodactylus_flavopictus | Leptodactylus_flavopictus | nocturnal |
| amphibians | Leptodactylus_fragilis | Leptodactylus_fragilis | diurnal |
| amphibians | Leptodactylus_furnarius | Leptodactylus_furnarius | cathemeral |
| amphibians | Leptodactylus_fuscus | Leptodactylus_fuscus | cathemeral |

| | | | |
|---|---|---|---|
| amphibians | Leptodactylus_gracilis | Leptodactylus_gracilis | cathemeral |
| amphibians | Leptodactylus_hylodes | Leptodactylus_hylodes | diurnal |
| amphibians | Leptodactylus_insularum | Leptodactylus_insularum | nocturnal |
| amphibians | Leptodactylus_jolyi | Leptodactylus_jolyi | nocturnal |
| amphibians | Leptodactylus_knudseni | Leptodactylus_knudseni | nocturnal |
| amphibians | Leptodactylus_labrosus | Leptodactylus_labrosus | cathemeral |
| amphibians | Leptodactylus_labyrinthicus | Leptodactylus_labyrinthicus | nocturnal |
| amphibians | Leptodactylus_laticeps | Leptodactylus_laticeps | cathemeral |
| amphibians | Leptodactylus_latinasus | Leptodactylus_latinasus | diurnal |
| amphibians | Leptodactylus_latrans | Leptodactylus_latrans | cathemeral |
| amphibians | Leptodactylus_leptodactyloides | Leptodactylus_leptodactyloides | cathemeral |
| amphibians | Leptodactylus_lithonaetes | Leptodactylus_lithonaetes | diurnal |
| amphibians | Leptodactylus_longirostris | Leptodactylus_longirostris | nocturnal |
| amphibians | Leptodactylus_melanonotus | Leptodactylus_melanonotus | cathemeral |
| amphibians | Leptodactylus_myersi | Leptodactylus_myersi | diurnal |
| amphibians | Leptodactylus_mystaceus | Leptodactylus_mystaceus | cathemeral |
| amphibians | Leptodactylus_mystacinus | Leptodactylus_mystacinus | cathemeral |
| amphibians | Leptodactylus_natalensis | Leptodactylus_natalensis | nocturnal |
| amphibians | Leptodactylus_nesiotus | Leptodactylus_nesiotus | crepuscular |
| amphibians | Leptodactylus_notoaktites | Leptodactylus_notoaktites | cathemeral |
| amphibians | Leptodactylus_paraensis | Leptodactylus_paraensis | diurnal |
| amphibians | Leptodactylus_pascoensis | Leptodactylus_pascoensis | diurnal |
| amphibians | Leptodactylus_pentadactylus | Leptodactylus_pentadactylus | nocturnal |
| amphibians | Leptodactylus_peritoaktites | Leptodactylus_peritoaktites | nocturnal |
| amphibians | Leptodactylus_petersii | Leptodactylus_petersii | cathemeral |
| amphibians | Leptodactylus_plaumanni | Leptodactylus_plaumanni | cathemeral |
| amphibians | Leptodactylus_podicipinus | Leptodactylus_podicipinus | nocturnal |
| amphibians | Leptodactylus_poecilochilus | Leptodactylus_poecilochilus | nocturnal |
| amphibians | Leptodactylus_pustulatus | Leptodactylus_pustulatus | diurnal |
| amphibians | Leptodactylus_rhodomerus | Leptodactylus_rhodomerus | nocturnal |
| amphibians | Leptodactylus_rhodomystax | Leptodactylus_rhodomystax | nocturnal |
| amphibians | Leptodactylus_rhodonotus | Leptodactylus_rhodonotus | nocturnal |
| amphibians | Leptodactylus_riveroi | Leptodactylus_riveroi | cathemeral |
| amphibians | Leptodactylus_rugosus | Leptodactylus_rugosus | nocturnal |

| | | | |
|---|---|---|---|
| amphibians | Leptodactylus_savagei | Leptodactylus_savagei | nocturnal |
| amphibians | Leptodactylus_sertanejo | Leptodactylus_sertanejo | cathemeral |
| amphibians | Leptodactylus_silvanimbus | Leptodactylus_silvanimbus | diurnal |
| amphibians | Leptodactylus_spixi | Leptodactylus_spixi | cathemeral |
| amphibians | Leptodactylus_stenodema | Leptodactylus_stenodema | cathemeral |
| amphibians | Leptodactylus_syphax | Leptodactylus_syphax | nocturnal |
| amphibians | Leptodactylus_troglodytes | Leptodactylus_troglodytes | nocturnal |
| amphibians | Leptodactylus_turimiquensis | Leptodactylus_turimiquensis | diurnal |
| amphibians | Leptodactylus_validus | Leptodactylus_validus | diurnal |
| amphibians | Leptodactylus_vastus | Leptodactylus_vastus | cathemeral |
| amphibians | Leptodactylus_ventrimaculatus | Leptodactylus_ventrimaculatus | nocturnal |
| amphibians | Leptodactylus_viridis | Leptodactylus_viridis | nocturnal |
| amphibians | Leptodactylus_wagneri | Leptodactylus_wagneri | nocturnal |
| amphibians | Leptomantis_angulirostris | Rhacophorus_angulirostris | nocturnal |
| amphibians | Leptomantis_cyanopunctatus | Rhacophorus_cyanopunctatus | nocturnal |
| amphibians | Leptomantis_gauni | Rhacophorus_gauni | nocturnal |
| amphibians | Leptopelis_bufonides | Leptopelis_bufonides | nocturnal |
| amphibians | Leptopelis_christyi | Leptopelis_christyi | nocturnal |
| amphibians | Leptopelis_concolor | Leptopelis_concolor | nocturnal |
| amphibians | Leptopelis_flavomaculatus | Leptopelis_flavomaculatus | nocturnal |
| amphibians | Leptopelis_karissimbensis | Leptopelis_karissimbensis | nocturnal |
| amphibians | Leptopelis_ragazzii | Leptopelis_ragazzii | cathemeral |
| amphibians | Leptopelis_spiritusnoctis | Leptopelis_spiritusnoctis | nocturnal |
| amphibians | Leptopelis_susanae | Leptopelis_susanae | cathemeral |
| amphibians | Leptopelis_vannutellii | Leptopelis_vannutellii | cathemeral |
| amphibians | Leptopelis_viridis | Leptopelis_viridis | nocturnal |
| amphibians | Leucostethus_alacris | Colostethus_alacris | diurnal |
| amphibians | Leucostethus_fraterdanieli | Colostethus_fraterdanieli | diurnal |
| amphibians | Leucostethus_ramirezi | Colostethus_ramirezi | diurnal |
| amphibians | Limnodynastes_interioris | Limnodynastes_interioris | nocturnal |
| amphibians | Limnodynastes_peronii | Limnodynastes_peronii | cathemeral |
| amphibians | Limnomedusa_macroglossa | Limnomedusa_macroglossa | nocturnal |
| amphibians | Limnonectes_arathooni | Limnonectes_arathooni | nocturnal |
| amphibians | Limnonectes_blythii | Limnonectes_blythii | nocturnal |

| | | | |
|---|---|---|---|
| amphibians | Limnonectes_dabanus | Limnonectes_dabanus | nocturnal |
| amphibians | Limnonectes_finchi | Limnonectes_finchi | nocturnal |
| amphibians | Limnonectes_hascheanus | Limnonectes_hascheanus | cathemeral |
| amphibians | Limnonectes_ibanorum | Limnonectes_ibanorum | nocturnal |
| amphibians | Limnonectes_kuhlii | Limnonectes_kuhlii | nocturnal |
| amphibians | Limnonectes_leporinus | Limnonectes_leporinus | nocturnal |
| amphibians | Limnonectes_leytensis | Limnonectes_leytensis | nocturnal |
| amphibians | Limnonectes_macrodon | Limnonectes_macrodon | nocturnal |
| amphibians | Limnonectes_malesianus | Limnonectes_malesianus | nocturnal |
| amphibians | Limnonectes_megastomias | Limnonectes_megastomias | cathemeral |
| amphibians | Limnonectes_microtympanum | Limnonectes_microtympanum | nocturnal |
| amphibians | Limnonectes_palavanensis | Limnonectes_palavanensis | nocturnal |
| amphibians | Limnonectes_paramacrodon | Limnonectes_paramacrodon | nocturnal |
| amphibians | Limnonectes_poilani | Limnonectes_poilani | nocturnal |
| amphibians | Limnonectes_shompenorum | Limnonectes_shompenorum | nocturnal |
| amphibians | Limnonectes_timorensis | Limnonectes_timorensis | nocturnal |
| amphibians | Limnonectes_visayanus | Limnonectes_visayanus | nocturnal |
| amphibians | Lissotriton_boscai | Lissotriton_boscai | cathemeral |
| amphibians | Lissotriton_helveticus | Lissotriton_helveticus | nocturnal |
| amphibians | Lissotriton_italicus | Lissotriton_italicus | nocturnal |
| amphibians | Lissotriton_montandoni | Lissotriton_montandoni | cathemeral |
| amphibians | Lissotriton_vulgaris | Lissotriton_vulgaris | cathemeral |
| amphibians | Lithobates_areolatus | Rana_areolata | nocturnal |
| amphibians | Lithobates_berlandieri | Rana_berlandieri | cathemeral |
| amphibians | Lithobates_bwana | Rana_bwana | nocturnal |
| amphibians | Lithobates_capito | Rana_capito | nocturnal |
| amphibians | Lithobates_chiricahuensis | Rana_chiricahuensis | nocturnal |
| amphibians | Lithobates_forreri | Rana_forreri | cathemeral |
| amphibians | Lithobates_juliani | Rana_juliani | cathemeral |
| amphibians | Lithobates_maculatus | Rana_maculata | nocturnal |
| amphibians | Lithobates_palmipes | Rana_palmipes | cathemeral |
| amphibians | Lithobates_palustris | Rana_palustris | nocturnal |
| amphibians | Lithobates_pipiens | Rana_pipiens | nocturnal |
| amphibians | Lithobates_sevosus | Rana_sevosa | diurnal |

| | | | |
|---|---|---|---|
| amphibians | Lithobates_sphenocephalus | Rana_sphenocephala | nocturnal |
| amphibians | Lithobates_taylori | Rana_taylori | cathemeral |
| amphibians | Lithobates_vaillanti | Rana_vaillanti | cathemeral |
| amphibians | Lithobates_vibicarius | Rana_vibicaria | cathemeral |
| amphibians | Lithobates_warszewitschii | Rana_warszewitschii | cathemeral |
| amphibians | Lithodytes_lineatus | Lithodytes_lineatus | cathemeral |
| amphibians | Litoria_aurifera | Litoria_aurifera | cathemeral |
| amphibians | Litoria_bibonius | Litoria_bibonius | nocturnal |
| amphibians | Litoria_bicolor | Litoria_bicolor | diurnal |
| amphibians | Litoria_christianbergmanni | Litoria_christianbergmanni | nocturnal |
| amphibians | Litoria_contrastens | Litoria_contrastens | nocturnal |
| amphibians | Litoria_coplandi | Litoria_coplandi | nocturnal |
| amphibians | Litoria_dentata | Litoria_dentata | nocturnal |
| amphibians | Litoria_ewingii | Litoria_ewingii | diurnal |
| amphibians | Litoria_fallax | Litoria_fallax | cathemeral |
| amphibians | Litoria_flavescens | Litoria_flavescens | nocturnal |
| amphibians | Litoria_gasconi | Litoria_gasconi | nocturnal |
| amphibians | Litoria_hilli | Litoria_hilli | nocturnal |
| amphibians | Litoria_humboldtorum | Litoria_humboldtorum | nocturnal |
| amphibians | Litoria_latopalmata | Litoria_latopalmata | nocturnal |
| amphibians | Litoria_littlejohni | Litoria_littlejohni | nocturnal |
| amphibians | Litoria_longirostris | Litoria_longirostris | nocturnal |
| amphibians | Litoria_megalops | Litoria_megalops | nocturnal |
| amphibians | Litoria_meiriana | Litoria_meiriana | cathemeral |
| amphibians | Litoria_nasuta | Litoria_nasuta | nocturnal |
| amphibians | Litoria_olongburensis | Litoria_olongburensis | nocturnal |
| amphibians | Litoria_pallida | Litoria_pallida | nocturnal |
| amphibians | Litoria_peronii | Litoria_peronii | nocturnal |
| amphibians | Litoria_personata | Litoria_personata | nocturnal |
| amphibians | Litoria_revelata | Litoria_revelata | nocturnal |
| amphibians | Litoria_staccato | Litoria_staccato | nocturnal |
| amphibians | Litoria_verreauxii | Litoria_verreauxii | nocturnal |
| amphibians | Litoria_vocivincens | Litoria_vocivincens | nocturnal |
| amphibians | Liua_tsinpaensis | Liua_tsinpaensis | nocturnal |

| | | | |
|---|---|---|---|
| amphibians | Lyciasalamandra_atifi | Lyciasalamandra_atifi | nocturnal |
| amphibians | Lyciasalamandra_fazilae | Lyciasalamandra_fazilae | nocturnal |
| amphibians | Lyciasalamandra_flavimembris | Lyciasalamandra_flavimembris | nocturnal |
| amphibians | Lyciasalamandra_luschani | Lyciasalamandra_luschani | cathemeral |
| amphibians | Lynchius_nebulanastes | Lynchius_nebulanastes | nocturnal |
| amphibians | Lynchius_simmonsi | Lynchius_simmonsi | cathemeral |
| amphibians | Lysapsus_caraya | Lysapsus_caraya | cathemeral |
| amphibians | Lysapsus_limellum | Lysapsus_limellum | nocturnal |
| amphibians | Macrogenioglottus_alipioi | Macrogenioglottus_alipioi | nocturnal |
| amphibians | Mannophryne_caquetio | Mannophryne_caquetio | diurnal |
| amphibians | Mannophryne_herminae | Mannophryne_herminae | diurnal |
| amphibians | Mannophryne_lamarcai | Mannophryne_lamarcai | diurnal |
| amphibians | Mannophryne_olmonae | Mannophryne_olmonae | diurnal |
| amphibians | Mannophryne_orellana | Mannophryne_orellana | diurnal |
| amphibians | Mannophryne_vulcano | Mannophryne_vulcano | diurnal |
| amphibians | Mantella_aurantiaca | Mantella_aurantiaca | cathemeral |
| amphibians | Mantella_baroni | Mantella_baroni | diurnal |
| amphibians | Mantella_bernhardi | Mantella_bernhardi | diurnal |
| amphibians | Mantella_betsileo | Mantella_betsileo | cathemeral |
| amphibians | Mantella_cowanii | Mantella_cowanii | diurnal |
| amphibians | Mantella_crocea | Mantella_crocea | diurnal |
| amphibians | Mantella_ebenaui | Mantella_ebenaui | diurnal |
| amphibians | Mantella_expectata | Mantella_expectata | diurnal |
| amphibians | Mantella_haraldmeieri | Mantella_haraldmeieri | diurnal |
| amphibians | Mantella_laevigata | Mantella_laevigata | diurnal |
| amphibians | Mantella_madagascariensis | Mantella_madagascariensis | diurnal |
| amphibians | Mantella_manery | Mantella_manery | diurnal |
| amphibians | Mantella_milotympanum | Mantella_milotympanum | diurnal |
| amphibians | Mantella_nigricans | Mantella_nigricans | diurnal |
| amphibians | Mantella_pulchra | Mantella_pulchra | diurnal |
| amphibians | Mantella_viridis | Mantella_viridis | diurnal |
| amphibians | Mantidactylus_aerumnalis | Mantidactylus_aerumnalis | diurnal |
| amphibians | Mantidactylus_albofrenatus | Mantidactylus_albofrenatus | diurnal |
| amphibians | Mantidactylus_alutus | Mantidactylus_alutus | cathemeral |

| | | | |
|---|---|---|---|
| amphibians | Mantidactylus_ambreensis | Mantidactylus_ambreensis | cathemeral |
| amphibians | Mantidactylus_argenteus | Mantidactylus_argenteus | diurnal |
| amphibians | Mantidactylus_bellyi | Mantidactylus_bellyi | cathemeral |
| amphibians | Mantidactylus_betsileanus | Mantidactylus_betsileanus | cathemeral |
| amphibians | Mantidactylus_biporus | Mantidactylus_biporus | diurnal |
| amphibians | Mantidactylus_brevipalmatus | Mantidactylus_brevipalmatus | cathemeral |
| amphibians | Mantidactylus_charlotteae | Mantidactylus_charlotteae | diurnal |
| amphibians | Mantidactylus_delormei | Mantidactylus_delormei | diurnal |
| amphibians | Mantidactylus_femoralis | Mantidactylus_femoralis | nocturnal |
| amphibians | Mantidactylus_grandidieri | Mantidactylus_grandidieri | cathemeral |
| amphibians | Mantidactylus_guttulatus | Mantidactylus_guttulatus | cathemeral |
| amphibians | Mantidactylus_lugubris | Mantidactylus_lugubris | nocturnal |
| amphibians | Mantidactylus_majori | Mantidactylus_majori | cathemeral |
| amphibians | Mantidactylus_melanopleura | Mantidactylus_melanopleura | diurnal |
| amphibians | Mantidactylus_mocquardi | Mantidactylus_mocquardi | cathemeral |
| amphibians | Mantidactylus_opiparis | Mantidactylus_opiparis | cathemeral |
| amphibians | Mantidactylus_pauliani | Mantidactylus_pauliani | diurnal |
| amphibians | Mantidactylus_ulcerosus | Mantidactylus_ulcerosus | cathemeral |
| amphibians | Mantidactylus_zipperi | Mantidactylus_zipperi | cathemeral |
| amphibians | Mantophryne_axanthogaster | Mantophryne_axanthogaster | nocturnal |
| amphibians | Mantophryne_lateralis | Mantophryne_lateralis | nocturnal |
| amphibians | Megaelosia_goeldii | Megaelosia_goeldii | diurnal |
| amphibians | Megastomatohyla_mixe | Megastomatohyla_mixe | nocturnal |
| amphibians | Megastomatohyla_nubicola | Megastomatohyla_nubicola | nocturnal |
| amphibians | Megastomatohyla_pellita | Megastomatohyla_pellita | nocturnal |
| amphibians | Megophrys_montana | Megophrys_montana | nocturnal |
| amphibians | Melanobatrachus_indicus | Melanobatrachus_indicus | nocturnal |
| amphibians | Melanophryniscus_admirabilis | Melanophryniscus_admirabilis | diurnal |
| amphibians | Melanophryniscus_alipioi | Melanophryniscus_alipioi | diurnal |
| amphibians | Melanophryniscus_atroluteus | Melanophryniscus_atroluteus | diurnal |
| amphibians | Melanophryniscus_cambaraensis | Melanophryniscus_cambaraensis | cathemeral |
| amphibians | Melanophryniscus_devincenzii | Melanophryniscus_devincenzii | diurnal |
| amphibians | Melanophryniscus_dorsalis | Melanophryniscus_dorsalis | diurnal |
| amphibians | Melanophryniscus_montevidensis | Melanophryniscus_montevidensis | diurnal |

| | | | |
|---|---|---|---|
| amphibians | Melanophryniscus_moreirae | Melanophryniscus_moreirae | diurnal |
| amphibians | Melanophryniscus_setiba | Melanophryniscus_setiba | diurnal |
| amphibians | Melanophryniscus_simplex | Melanophryniscus_simplex | nocturnal |
| amphibians | Melanophryniscus_spectabilis | Melanophryniscus_spectabilis | diurnal |
| amphibians | Melanophryniscus_stelzneri | Melanophryniscus_stelzneri | diurnal |
| amphibians | Melanophryniscus_tumifrons | Melanophryniscus_tumifrons | diurnal |
| amphibians | Melanophryniscus_vilavelhensis | Melanophryniscus_vilavelhensis | cathemeral |
| amphibians | Mercurana_myristicapalustris | Mercurana_myristicapalustris | nocturnal |
| amphibians | Meristogenys_amoropalamus | Meristogenys_amoropalamus | nocturnal |
| amphibians | Meristogenys_jerboa | Meristogenys_jerboa | cathemeral |
| amphibians | Meristogenys_kinabaluensis | Meristogenys_kinabaluensis | nocturnal |
| amphibians | Meristogenys_maryatiae | Meristogenys_maryatiae | nocturnal |
| amphibians | Meristogenys_orphnocnemis | Meristogenys_orphnocnemis | cathemeral |
| amphibians | Meristogenys_whiteheadi | Meristogenys_whiteheadi | nocturnal |
| amphibians | Mertensiella_caucasica | Mertensiella_caucasica | nocturnal |
| amphibians | Mertensophryne_taitana | Mertensophryne_taitana | cathemeral |
| amphibians | Metaphrynella_sundana | Metaphrynella_sundana | nocturnal |
| amphibians | Micrixalus_elegans | Micrixalus_elegans | diurnal |
| amphibians | Micrixalus_fuscus | Micrixalus_fuscus | diurnal |
| amphibians | Micrixalus_gadgili | Micrixalus_gadgili | diurnal |
| amphibians | Micrixalus_kottigeharensis | Micrixalus_kottigeharensis | diurnal |
| amphibians | Micrixalus_nudis | Micrixalus_nudis | diurnal |
| amphibians | Micrixalus_phyllophilus | Micrixalus_phyllophilus | diurnal |
| amphibians | Micrixalus_saxicola | Micrixalus_saxicola | diurnal |
| amphibians | Micrixalus_thampii | Micrixalus_thampii | diurnal |
| amphibians | Microbatrachella_capensis | Microbatrachella_capensis | nocturnal |
| amphibians | Microhyla_achatina | Microhyla_achatina | nocturnal |
| amphibians | Microhyla_borneensis | Microhyla_borneensis | cathemeral |
| amphibians | Microhyla_fissipes | Microhyla_fissipes | nocturnal |
| amphibians | Microhyla_heymonsi | Microhyla_heymonsi | cathemeral |
| amphibians | Microhyla_mantheyi | Microhyla_mantheyi | nocturnal |
| amphibians | Microhyla_ornata | Microhyla_ornata | nocturnal |
| amphibians | Microhyla_pulchra | Microhyla_pulchra | nocturnal |
| amphibians | Microhyla_rubra | Microhyla_rubra | nocturnal |

| | | | |
|---|---|---|---|
| amphibians | Microkayla_ankohuma | Psychrophrynella_ankohuma | nocturnal |
| amphibians | Microkayla_chacaltaya | Psychrophrynella_chacaltaya | cathemeral |
| amphibians | Microkayla_iatamasi | Psychrophrynella_iatamasi | diurnal |
| amphibians | Microkayla_wettsteini | Psychrophrynella_wettsteini | cathemeral |
| amphibians | Micryletta_erythropoda | Microhyla_erythropoda | nocturnal |
| amphibians | Micryletta_steinegeri | Micryletta_steinegeri | diurnal |
| amphibians | Minervarya_brevipalmata | Fejervarya_brevipalmata | nocturnal |
| amphibians | Minervarya_greenii | Fejervarya_greenii | diurnal |
| amphibians | Minervarya_kirtisinghei | Fejervarya_kirtisinghei | diurnal |
| amphibians | Minervarya_nepalensis | Fejervarya_nepalensis | nocturnal |
| amphibians | Minervarya_nicobariensis | Fejervarya_nicobariensis | cathemeral |
| amphibians | Minervarya_pierrei | Fejervarya_pierrei | nocturnal |
| amphibians | Minervarya_rufescens | Fejervarya_rufescens | nocturnal |
| amphibians | Mixophyes_carbinensis | Mixophyes_carbinensis | nocturnal |
| amphibians | Mixophyes_coggeri | Mixophyes_coggeri | nocturnal |
| amphibians | Mixophyes_fasciolatus | Mixophyes_fasciolatus | nocturnal |
| amphibians | Mixophyes_fleayi | Mixophyes_fleayi | nocturnal |
| amphibians | Mixophyes_iteratus | Mixophyes_iteratus | nocturnal |
| amphibians | Mixophyes_schevilli | Mixophyes_schevilli | nocturnal |
| amphibians | Morerella_cyanophthalma | Morerella_cyanophthalma | nocturnal |
| amphibians | Myersiella_microps | Myersiella_microps | cathemeral |
| amphibians | Myersiohyla_chamaeleo | Myersiohyla_chamaeleo | nocturnal |
| amphibians | Myersiohyla_liliae | Hypsiboas_liliae | nocturnal |
| amphibians | Myersiohyla_loveridgei | Myersiohyla_loveridgei | diurnal |
| amphibians | Myersiohyla_neblinaria | Myersiohyla_neblinaria | nocturnal |
| amphibians | Nannophryne_apolobambica | Nannophryne_apolobambica | cathemeral |
| amphibians | Nannophryne_variegata | Nannophryne_variegata | nocturnal |
| amphibians | Nannophrys_ceylonensis | Nannophrys_ceylonensis | nocturnal |
| amphibians | Nannophrys_naeyakai | Nannophrys_naeyakai | nocturnal |
| amphibians | Nanorana_annandalii | Nanorana_annandalii | nocturnal |
| amphibians | Nanorana_blanfordii | Nanorana_blanfordii | nocturnal |
| amphibians | Nanorana_kangxianensis | Nanorana_kangxianensis | nocturnal |
| amphibians | Nanorana_liebigii | Nanorana_liebigii | nocturnal |
| amphibians | Nanorana_parkeri | Nanorana_parkeri | cathemeral |

| | | | |
|---|---|---|---|
| amphibians | Nanorana_pleskei | Nanorana_pleskei | nocturnal |
| amphibians | Nanorana_polunini | Nanorana_polunini | nocturnal |
| amphibians | Nanorana_rostandi | Nanorana_rostandi | cathemeral |
| amphibians | Nasikabatrachus_sahyadrensis | Nasikabatrachus_sahyadrensis | nocturnal |
| amphibians | Nectophryne_afra | Nectophryne_afra | cathemeral |
| amphibians | Nectophryne_batesii | Nectophryne_batesii | nocturnal |
| amphibians | Nectophrynoides_asperginis | Nectophrynoides_asperginis | nocturnal |
| amphibians | Nectophrynoides_laticeps | Nectophrynoides_laticeps | cathemeral |
| amphibians | Nectophrynoides_paulae | Nectophrynoides_paulae | nocturnal |
| amphibians | Nectophrynoides_poyntoni | Nectophrynoides_poyntoni | diurnal |
| amphibians | Nectophrynoides_pseudotornieri | Nectophrynoides_pseudotornieri | cathemeral |
| amphibians | Necturus_beyeri | Necturus_beyeri | nocturnal |
| amphibians | Necturus_lewisi | Necturus_lewisi | nocturnal |
| amphibians | Necturus_maculosus | Necturus_maculosus | nocturnal |
| amphibians | Necturus_punctatus | Necturus_punctatus | nocturnal |
| amphibians | Neobatrachus_albipes | Neobatrachus_albipes | nocturnal |
| amphibians | Neobatrachus_kunapalari | Neobatrachus_kunapalari | nocturnal |
| amphibians | Neobatrachus_sudelli | Neobatrachus_sudelli | nocturnal |
| amphibians | Neurergus_crocatus | Neurergus_crocatus | nocturnal |
| amphibians | Neurergus_kaiseri | Neurergus_kaiseri | diurnal |
| amphibians | Neurergus_strauchii | Neurergus_strauchii | nocturnal |
| amphibians | Niceforonia_babax | Hypodactylus_babax | nocturnal |
| amphibians | Niceforonia_elassodiscus | Hypodactylus_elassodiscus | diurnal |
| amphibians | Niceforonia_nigrovittata | Hypodactylus_nigrovittatus | cathemeral |
| amphibians | Nidirana_chapaensis | Babina_chapaensis | nocturnal |
| amphibians | Nidirana_daunchina | Babina_daunchina | cathemeral |
| amphibians | Nidirana_pleuraden | Babina_pleuraden | diurnal |
| amphibians | Nimbaphrynoides_occidentalis | Nimbaphrynoides_occidentalis | cathemeral |
| amphibians | Noblella_coloma | Noblella_coloma | cathemeral |
| amphibians | Noblella_duellmani | Noblella_duellmani | nocturnal |
| amphibians | Noblella_lochites | Noblella_lochites | diurnal |
| amphibians | Noblella_myrmecoides | Noblella_myrmecoides | diurnal |
| amphibians | Noblella_personina | Noblella_personina | diurnal |
| amphibians | Notophthalmus_perstriatus | Notophthalmus_perstriatus | nocturnal |

| | | | |
|---|---|---|---|
| amphibians | Notophthalmus_viridescens | Notophthalmus_viridescens | nocturnal |
| amphibians | Nototriton_abscondens | Nototriton_abscondens | nocturnal |
| amphibians | Nototriton_barbouri | Nototriton_barbouri | nocturnal |
| amphibians | Nototriton_brodiei | Nototriton_brodiei | nocturnal |
| amphibians | Nototriton_limnospectator | Nototriton_limnospectator | nocturnal |
| amphibians | Nyctibates_corrugatus | Nyctibates_corrugatus | nocturnal |
| amphibians | Nyctibatrachus_beddomii | Nyctibatrachus_beddomii | cathemeral |
| amphibians | Nyctibatrachus_dattatreyaensis | Nyctibatrachus_dattatreyaensis | nocturnal |
| amphibians | Nyctibatrachus_karnatakaensis | Nyctibatrachus_karnatakaensis | nocturnal |
| amphibians | Nyctibatrachus_kempholeyensis | Nyctibatrachus_kempholeyensis | nocturnal |
| amphibians | Nyctibatrachus_major | Nyctibatrachus_major | nocturnal |
| amphibians | Nyctibatrachus_petraeus | Nyctibatrachus_petraeus | nocturnal |
| amphibians | Nyctimantis_arapapa | Aparasphenodon_arapapa | nocturnal |
| amphibians | Nyctimantis_bokermanni | Aparasphenodon_bokermanni | nocturnal |
| amphibians | Nyctimantis_brunoi | Aparasphenodon_brunoi | nocturnal |
| amphibians | Nyctimantis_rugiceps | Nyctimantis_rugiceps | nocturnal |
| amphibians | Nyctimantis_siemersi | Argenteohyla_siemersi | nocturnal |
| amphibians | Nyctimystes_kuduki | Nyctimystes_kuduki | nocturnal |
| amphibians | Nyctixalus_pictus | Nyctixalus_pictus | nocturnal |
| amphibians | Nyctixalus_spinosus | Nyctixalus_spinosus | nocturnal |
| amphibians | Nymphargus_anomalus | Nymphargus_anomalus | nocturnal |
| amphibians | Nymphargus_balionotus | Cochranella_balionota | nocturnal |
| amphibians | Nymphargus_chancas | Nymphargus_chancas | nocturnal |
| amphibians | Nymphargus_cochranae | Nymphargus_cochranae | nocturnal |
| amphibians | Nymphargus_grandisonae | Nymphargus_grandisonae | nocturnal |
| amphibians | Nymphargus_griffithsi | Nymphargus_griffithsi | nocturnal |
| amphibians | Nymphargus_ignotus | Nymphargus_ignotus | nocturnal |
| amphibians | Nymphargus_lasgralarias | Nymphargus_lasgralarias | nocturnal |
| amphibians | Nymphargus_mariae | Nymphargus_mariae | nocturnal |
| amphibians | Nymphargus_megacheirus | Nymphargus_megacheirus | nocturnal |
| amphibians | Nymphargus_mixomaculatus | Nymphargus_mixomaculatus | nocturnal |
| amphibians | Nymphargus_posadae | Nymphargus_posadae | nocturnal |
| amphibians | Nymphargus_rosada | Nymphargus_rosada | nocturnal |
| amphibians | Nymphargus_sucre | Nymphargus_sucre | nocturnal |

| | | | |
|---|---|---|---|
| amphibians | Nymphargus_vicenteruedai | Nymphargus_vicenteruedai | nocturnal |
| amphibians | Nymphargus_wileyi | Nymphargus_wileyi | nocturnal |
| amphibians | Occidozyga_baluensis | Occidozyga_baluensis | cathemeral |
| amphibians | Occidozyga_laevis | Occidozyga_laevis | cathemeral |
| amphibians | Occidozyga_lima | Occidozyga_lima | cathemeral |
| amphibians | Occidozyga_martensii | Occidozyga_martensii | nocturnal |
| amphibians | Occidozyga_sumatrana | Occidozyga_sumatrana | nocturnal |
| amphibians | Odontobatrachus_natator | Odontobatrachus_natator | nocturnal |
| amphibians | Odontophrynus_carvalhoi | Odontophrynus_carvalhoi | nocturnal |
| amphibians | Odontophrynus_maisuma | Odontophrynus_maisuma | nocturnal |
| amphibians | Odorrana_aureola | Odorrana_aureola | nocturnal |
| amphibians | Odorrana_banaorum | Odorrana_banaorum | nocturnal |
| amphibians | Odorrana_hosii | Odorrana_hosii | cathemeral |
| amphibians | Odorrana_khalam | Odorrana_khalam | nocturnal |
| amphibians | Odorrana_narina | Odorrana_narina | nocturnal |
| amphibians | Odorrana_tormota | Odorrana_tormota | nocturnal |
| amphibians | Odorrana_yentuensis | Odorrana_yentuensis | nocturnal |
| amphibians | Oedipina_collaris | Oedipina_collaris | nocturnal |
| amphibians | Oedipina_cyclocauda | Oedipina_cyclocauda | nocturnal |
| amphibians | Oedipina_gephyra | Oedipina_gephyra | nocturnal |
| amphibians | Oedipina_grandis | Oedipina_grandis | nocturnal |
| amphibians | Oedipina_poelzi | Oedipina_poelzi | nocturnal |
| amphibians | Ololygon_agilis | Scinax_agilis | cathemeral |
| amphibians | Ololygon_albicans | Scinax_albicans | nocturnal |
| amphibians | Ololygon_alcatraz | Scinax_alcatraz | nocturnal |
| amphibians | Ololygon_angrensis | Scinax_angrensis | cathemeral |
| amphibians | Ololygon_arduoa | Scinax_arduous | cathemeral |
| amphibians | Ololygon_argyreornata | Scinax_argyreornatus | cathemeral |
| amphibians | Ololygon_ariadne | Scinax_ariadne | nocturnal |
| amphibians | Ololygon_aromothyella | Scinax_aromothyella | nocturnal |
| amphibians | Ololygon_belloni | Scinax_belloni | nocturnal |
| amphibians | Ololygon_berthae | Scinax_berthae | nocturnal |
| amphibians | Ololygon_brieni | Scinax_brieni | nocturnal |
| amphibians | Ololygon_carnevallii | Scinax_carnevallii | nocturnal |

| | | | |
|---|---|---|---|
| amphibians | Ololygon_catharinae | Scinax_catharinae | nocturnal |
| amphibians | Ololygon_cosenzai | Scinax_cosenzai | nocturnal |
| amphibians | Ololygon_faivovichi | Scinax_faivovichi | nocturnal |
| amphibians | Ololygon_flavoguttata | Scinax_flavoguttatus | nocturnal |
| amphibians | Ololygon_heyeri | Scinax_heyeri | nocturnal |
| amphibians | Ololygon_hiemalis | Scinax_hiemalis | nocturnal |
| amphibians | Ololygon_humilis | Scinax_humilis | nocturnal |
| amphibians | Ololygon_insperata | Scinax_insperatus | nocturnal |
| amphibians | Ololygon_jureia | Scinax_jureia | nocturnal |
| amphibians | Ololygon_littoralis | Scinax_littoralis | nocturnal |
| amphibians | Ololygon_littorea | Scinax_littoreus | nocturnal |
| amphibians | Ololygon_longilinea | Scinax_longilineus | nocturnal |
| amphibians | Ololygon_luizotavioi | Scinax_luizotavioi | nocturnal |
| amphibians | Ololygon_machadoi | Scinax_machadoi | nocturnal |
| amphibians | Ololygon_obtriangulata | Scinax_obtriangulatus | nocturnal |
| amphibians | Ololygon_peixotoi | Scinax_peixotoi | nocturnal |
| amphibians | Ololygon_perpusilla | Scinax_perpusillus | nocturnal |
| amphibians | Ololygon_ranki | Scinax_ranki | nocturnal |
| amphibians | Ololygon_rizibilis | Scinax_rizibilis | cathemeral |
| amphibians | Ololygon_skaios | Scinax_skaios | nocturnal |
| amphibians | Ololygon_strigilata | Scinax_strigilatus | nocturnal |
| amphibians | Ololygon_trapicheiroi | Scinax_trapicheiroi | nocturnal |
| amphibians | Ololygon_v-signata | Scinax_v-signatus | nocturnal |
| amphibians | Ombrana_sikimensis | Ombrana_sikimensis | nocturnal |
| amphibians | Ommatotriton_ophryticus | Ommatotriton_ophryticus | nocturnal |
| amphibians | Ommatotriton_vittatus | Ommatotriton_vittatus | nocturnal |
| amphibians | Onychodactylus_fischeri | Onychodactylus_fischeri | nocturnal |
| amphibians | Onychodactylus_japonicus | Onychodactylus_japonicus | diurnal |
| amphibians | Onychodactylus_nipponoborealis | Onychodactylus_nipponoborealis | nocturnal |
| amphibians | Onychodactylus_zhangyapingi | Onychodactylus_zhangyapingi | nocturnal |
| amphibians | Onychodactylus_zhaoermii | Onychodactylus_zhaoermii | nocturnal |
| amphibians | Oophaga_arborea | Oophaga_arborea | diurnal |
| amphibians | Oophaga_granulifera | Oophaga_granulifera | diurnal |
| amphibians | Oophaga_histrionica | Oophaga_histrionica | diurnal |

| | | | |
|---|---|---|---|
| amphibians | Oophaga_lehmanni | Oophaga_lehmanni | diurnal |
| amphibians | Oophaga_occultator | Oophaga_occultator | diurnal |
| amphibians | Oophaga_pumilio | Oophaga_pumilio | diurnal |
| amphibians | Oophaga_speciosa | Oophaga_speciosa | diurnal |
| amphibians | Oophaga_sylvatica | Oophaga_sylvatica | diurnal |
| amphibians | Oophaga_vicentei | Oophaga_vicentei | diurnal |
| amphibians | Ophryophryne_hansi | Ophryophryne_hansi | nocturnal |
| amphibians | Opisthothylax_immaculatus | Opisthothylax_immaculatus | nocturnal |
| amphibians | Oreobates_amarakaeri | Oreobates_amarakaeri | nocturnal |
| amphibians | Oreobates_ayacucho | Oreobates_ayacucho | diurnal |
| amphibians | Oreobates_barituensis | Oreobates_barituensis | nocturnal |
| amphibians | Oreobates_cruralis | Oreobates_cruralis | nocturnal |
| amphibians | Oreobates_gemcare | Oreobates_gemcare | nocturnal |
| amphibians | Oreobates_machiguenga | Oreobates_machiguenga | nocturnal |
| amphibians | Oreobates_quixensis | Oreobates_quixensis | cathemeral |
| amphibians | Oreobates_sanderi | Oreobates_sanderi | diurnal |
| amphibians | Oreolalax_chuanbeiensis | Oreolalax_chuanbeiensis | nocturnal |
| amphibians | Oreolalax_liangbeiensis | Oreolalax_liangbeiensis | nocturnal |
| amphibians | Oreolalax_omeimontis | Oreolalax_omeimontis | nocturnal |
| amphibians | Oreolalax_popei | Oreolalax_popei | nocturnal |
| amphibians | Oreolalax_xiangchengensis | Oreolalax_xiangchengensis | nocturnal |
| amphibians | Oreophryne_anamiatoi | Oreophryne_anamiatoi | nocturnal |
| amphibians | Oreophryne_atrigularis | Oreophryne_atrigularis | nocturnal |
| amphibians | Oreophryne_clamata | Oreophryne_clamata | nocturnal |
| amphibians | Oreophryne_geislerorum | Oreophryne_geislerorum | nocturnal |
| amphibians | Oreophryne_loriae | Oreophryne_loriae | nocturnal |
| amphibians | Oreophryne_minuta | Oreophryne_minuta | nocturnal |
| amphibians | Oreophryne_notata | Oreophryne_notata | nocturnal |
| amphibians | Oreophryne_oviprotector | Oreophryne_oviprotector | nocturnal |
| amphibians | Oreophryne_sibilans | Oreophryne_sibilans | nocturnal |
| amphibians | Oreophryne_unicolor | Oreophryne_unicolor | nocturnal |
| amphibians | Oreophryne_wapoga | Oreophryne_wapoga | nocturnal |
| amphibians | Oreophrynella_seegobini | Oreophrynella_seegobini | diurnal |
| amphibians | Osornophryne_antisana | Osornophryne_antisana | cathemeral |

| | | | |
|---|---|---|---|
| amphibians | Osornophryne_cofanorum | Osornophryne_cofanorum | nocturnal |
| amphibians | Osornophryne_guacamayo | Osornophryne_guacamayo | cathemeral |
| amphibians | Osornophryne_occidentalis | Osornophryne_occidentalis | cathemeral |
| amphibians | Osornophryne_puruanta | Osornophryne_puruanta | nocturnal |
| amphibians | Osornophryne_simpsoni | Osornophryne_simpsoni | nocturnal |
| amphibians | Osornophryne_sumacoensis | Osornophryne_sumacoensis | cathemeral |
| amphibians | Osteocephalus_alboguttatus | Osteocephalus_alboguttatus | nocturnal |
| amphibians | Osteocephalus_cabrerai | Osteocephalus_cabrerai | nocturnal |
| amphibians | Osteocephalus_deridens | Osteocephalus_deridens | nocturnal |
| amphibians | Osteocephalus_festae | Osteocephalus_festae | nocturnal |
| amphibians | Osteocephalus_fuscifacies | Osteocephalus_fuscifacies | nocturnal |
| amphibians | Osteocephalus_heyeri | Osteocephalus_heyeri | nocturnal |
| amphibians | Osteocephalus_leprieurii | Osteocephalus_leprieurii | nocturnal |
| amphibians | Osteocephalus_oophagus | Osteocephalus_oophagus | crepuscular |
| amphibians | Osteocephalus_planiceps | Osteocephalus_planiceps | nocturnal |
| amphibians | Osteocephalus_subtilis | Osteocephalus_subtilis | nocturnal |
| amphibians | Osteocephalus_taurinus | Osteocephalus_taurinus | nocturnal |
| amphibians | Osteocephalus_yasuni | Osteocephalus_yasuni | cathemeral |
| amphibians | Osteopilus_crucialis | Osteopilus_crucialis | nocturnal |
| amphibians | Osteopilus_dominicensis | Osteopilus_dominicensis | nocturnal |
| amphibians | Osteopilus_ocellatus | Osteopilus_ocellatus | nocturnal |
| amphibians | Osteopilus_septentrionalis | Osteopilus_septentrionalis | nocturnal |
| amphibians | Otophryne_pyburni | Otophryne_pyburni | diurnal |
| amphibians | Otophryne_robusta | Otophryne_robusta | diurnal |
| amphibians | Otophryne_steyermarki | Otophryne_steyermarki | diurnal |
| amphibians | Pachytriton_archospotus | Pachytriton_archospotus | nocturnal |
| amphibians | Pachytriton_brevipes | Pachytriton_brevipes | nocturnal |
| amphibians | Pachytriton_changi | Pachytriton_changi | nocturnal |
| amphibians | Pachytriton_feii | Pachytriton_feii | nocturnal |
| amphibians | Pachytriton_granulosus | Pachytriton_granulosus | nocturnal |
| amphibians | Pachytriton_moi | Pachytriton_moi | nocturnal |
| amphibians | Paedophryne_amauensis | Paedophryne_amauensis | nocturnal |
| amphibians | Paedophryne_verrucosa | Paedophryne_verrucosa | crepuscular |
| amphibians | Paracassina_kounhiensis | Paracassina_kounhiensis | cathemeral |

| | | | |
|---|---|---|---|
| amphibians | Paracassina_obscura | Paracassina_obscura | cathemeral |
| amphibians | Paradoxophyla_palmata | Paradoxophyla_palmata | nocturnal |
| amphibians | Paradoxophyla_tiarano | Paradoxophyla_tiarano | nocturnal |
| amphibians | Paramesotriton_caudopunctatus | Paramesotriton_caudopunctatus | nocturnal |
| amphibians | Paramesotriton_chinensis | Paramesotriton_chinensis | nocturnal |
| amphibians | Paramesotriton_hongkongensis | Paramesotriton_hongkongensis | nocturnal |
| amphibians | Paramesotriton_longliensis | Paramesotriton_longliensis | nocturnal |
| amphibians | Paramesotriton_yunwuensis | Paramesotriton_yunwuensis | cathemeral |
| amphibians | Paratelmatobius_cardosoi | Paratelmatobius_cardosoi | nocturnal |
| amphibians | Paratelmatobius_gaigeae | Paratelmatobius_gaigeae | nocturnal |
| amphibians | Paratelmatobius_lutzii | Paratelmatobius_lutzii | diurnal |
| amphibians | Paratelmatobius_mantiqueira | Paratelmatobius_mantiqueira | nocturnal |
| amphibians | Paratelmatobius_poecilogaster | Paratelmatobius_poecilogaster | nocturnal |
| amphibians | Paratelmatobius_yepiranga | Paratelmatobius_yepiranga | nocturnal |
| amphibians | Paruwrobates_andinus | Ameerega_andina | diurnal |
| amphibians | Paruwrobates_erythromos | Ameerega_erythromos | diurnal |
| amphibians | Pedostibes_tuberculosus | Pedostibes_tuberculosus | nocturnal |
| amphibians | Pelobates_cultripes | Pelobates_cultripes | nocturnal |
| amphibians | Pelobates_fuscus | Pelobates_fuscus | nocturnal |
| amphibians | Pelobates_syriacus | Pelobates_syriacus | cathemeral |
| amphibians | Pelobates_varaldii | Pelobates_varaldii | nocturnal |
| amphibians | Pelodytes_caucasicus | Pelodytes_caucasicus | nocturnal |
| amphibians | Pelodytes_ibericus | Pelodytes_ibericus | cathemeral |
| amphibians | Pelophryne_linanitensis | Pelophryne_linanitensis | nocturnal |
| amphibians | Pelophryne_misera | Pelophryne_misera | diurnal |
| amphibians | Pelophryne_signata | Pelophryne_signata | cathemeral |
| amphibians | Pelophylax_epeiroticus | Pelophylax_epeiroticus | diurnal |
| amphibians | Pelophylax_nigromaculatus | Pelophylax_nigromaculatus | cathemeral |
| amphibians | Pelophylax_perezi | Pelophylax_perezi | cathemeral |
| amphibians | Pelophylax_porosus | Pelophylax_porosus | diurnal |
| amphibians | Pelophylax_ridibundus | Pelophylax_ridibundus | cathemeral |
| amphibians | Pelophylax_saharicus | Pelophylax_saharicus | cathemeral |
| amphibians | Peltophryne_fustiger | Peltophryne_fustiger | nocturnal |
| amphibians | Peltophryne_gundlachi | Peltophryne_gundlachi | nocturnal |

| | | | |
|---|---|---|---|
| amphibians | Peltophryne_lemur | Peltophryne_lemur | nocturnal |
| amphibians | Peltophryne_longinasus | Peltophryne_longinasus | cathemeral |
| amphibians | Peltophryne_peltocephala | Peltophryne_peltocephala | nocturnal |
| amphibians | Peltophryne_taladai | Peltophryne_taladai | nocturnal |
| amphibians | Petropedetes_cameronensis | Petropedetes_cameronensis | nocturnal |
| amphibians | Petropedetes_euskircheni | Petropedetes_euskircheni | nocturnal |
| amphibians | Petropedetes_palmipes | Petropedetes_palmipes | nocturnal |
| amphibians | Petropedetes_perreti | Petropedetes_perreti | nocturnal |
| amphibians | Petropedetes_vulpiae | Petropedetes_vulpiae | cathemeral |
| amphibians | Phaeognathus_hubrichti | Phaeognathus_hubrichti | nocturnal |
| amphibians | Phantasmarana_apuana | Megaelosia_apuana | diurnal |
| amphibians | Phantasmarana_bocainensis | Megaelosia_bocainensis | diurnal |
| amphibians | Phantasmarana_boticariana | Megaelosia_boticariana | diurnal |
| amphibians | Phantasmarana_lutzae | Megaelosia_lutzae | diurnal |
| amphibians | Phantasmarana_massarti | Megaelosia_massarti | diurnal |
| amphibians | Phasmahyla_cochranae | Phasmahyla_cochranae | nocturnal |
| amphibians | Phasmahyla_exilis | Phasmahyla_exilis | nocturnal |
| amphibians | Phasmahyla_guttata | Phasmahyla_guttata | nocturnal |
| amphibians | Phasmahyla_jandaia | Phasmahyla_jandaia | nocturnal |
| amphibians | Phasmahyla_spectabilis | Phasmahyla_spectabilis | nocturnal |
| amphibians | Phasmahyla_timbo | Phasmahyla_timbo | nocturnal |
| amphibians | Philautus_aurantium | Philautus_aurantium | nocturnal |
| amphibians | Philautus_bunitus | Philautus_bunitus | cathemeral |
| amphibians | Philautus_ingeri | Philautus_ingeri | nocturnal |
| amphibians | Philautus_longicrus | Philautus_longicrus | nocturnal |
| amphibians | Philautus_macroscelis | Philautus_macroscelis | nocturnal |
| amphibians | Philautus_mjobergi | Philautus_mjobergi | cathemeral |
| amphibians | Philautus_petersi | Philautus_petersi | nocturnal |
| amphibians | Philautus_saueri | Philautus_saueri | diurnal |
| amphibians | Philoria_frosti | Philoria_frosti | diurnal |
| amphibians | Philoria_kundagungan | Philoria_kundagungan | nocturnal |
| amphibians | Philoria_richmondensis | Philoria_richmondensis | cathemeral |
| amphibians | Philoria_sphagnicola | Philoria_sphagnicolus | diurnal |
| amphibians | Phrynobatrachus_acridoides | Phrynobatrachus_acridoides | cathemeral |

| | | | |
|---|---|---|---|
| amphibians | Phrynobatrachus_africanus | Phrynobatrachus_africanus | diurnal |
| amphibians | Phrynobatrachus_alleni | Phrynobatrachus_alleni | diurnal |
| amphibians | Phrynobatrachus_annulatus | Phrynobatrachus_annulatus | diurnal |
| amphibians | Phrynobatrachus_auritus | Phrynobatrachus_auritus | cathemeral |
| amphibians | Phrynobatrachus_batesii | Phrynobatrachus_batesii | diurnal |
| amphibians | Phrynobatrachus_breviceps | Phrynobatrachus_breviceps | nocturnal |
| amphibians | Phrynobatrachus_bullans | Phrynobatrachus_bullans | cathemeral |
| amphibians | Phrynobatrachus_calcaratus | Phrynobatrachus_calcaratus | cathemeral |
| amphibians | Phrynobatrachus_chukuchuku | Phrynobatrachus_chukuchuku | diurnal |
| amphibians | Phrynobatrachus_cornutus | Phrynobatrachus_cornutus | diurnal |
| amphibians | Phrynobatrachus_danko | Phrynobatrachus_danko | diurnal |
| amphibians | Phrynobatrachus_dispar | Phrynobatrachus_dispar | cathemeral |
| amphibians | Phrynobatrachus_francisci | Phrynobatrachus_francisci | cathemeral |
| amphibians | Phrynobatrachus_fraterculus | Phrynobatrachus_fraterculus | cathemeral |
| amphibians | Phrynobatrachus_ghanensis | Phrynobatrachus_ghanensis | diurnal |
| amphibians | Phrynobatrachus_guineensis | Phrynobatrachus_guineensis | diurnal |
| amphibians | Phrynobatrachus_gutturosus | Phrynobatrachus_gutturosus | cathemeral |
| amphibians | Phrynobatrachus_hylaios | Phrynobatrachus_hylaios | diurnal |
| amphibians | Phrynobatrachus_inexpectatus | Phrynobatrachus_inexpectatus | crepuscular |
| amphibians | Phrynobatrachus_irangi | Phrynobatrachus_irangi | diurnal |
| amphibians | Phrynobatrachus_jimzimkusi | Phrynobatrachus_jimzimkusi | diurnal |
| amphibians | Phrynobatrachus_kakamikro | Phrynobatrachus_kakamikro | nocturnal |
| amphibians | Phrynobatrachus_keniensis | Phrynobatrachus_keniensis | diurnal |
| amphibians | Phrynobatrachus_krefftii | Phrynobatrachus_krefftii | diurnal |
| amphibians | Phrynobatrachus_latifrons | Phrynobatrachus_latifrons | diurnal |
| amphibians | Phrynobatrachus_leveleve | Phrynobatrachus_leveleve | diurnal |
| amphibians | Phrynobatrachus_liberiensis | Phrynobatrachus_liberiensis | diurnal |
| amphibians | Phrynobatrachus_mababiensis | Phrynobatrachus_mababiensis | cathemeral |
| amphibians | Phrynobatrachus_maculiventris | Phrynobatrachus_maculiventris | cathemeral |
| amphibians | Phrynobatrachus_manengoubensis | Phrynobatrachus_manengoubensis | diurnal |
| amphibians | Phrynobatrachus_minutus | Phrynobatrachus_minutus | crepuscular |
| amphibians | Phrynobatrachus_natalensis | Phrynobatrachus_natalensis | cathemeral |
| amphibians | Phrynobatrachus_njiomock | Phrynobatrachus_njiomock | cathemeral |
| amphibians | Phrynobatrachus_ogoensis | Phrynobatrachus_ogoensis | diurnal |

| | | | |
|---|---|---|---|
| amphibians | Phrynobatrachus_parvulus | Phrynobatrachus_parvulus | diurnal |
| amphibians | Phrynobatrachus_phyllophilus | Phrynobatrachus_phyllophilus | diurnal |
| amphibians | Phrynobatrachus_pintoi | Phrynobatrachus_pintoi | diurnal |
| amphibians | Phrynobatrachus_plicatus | Phrynobatrachus_plicatus | diurnal |
| amphibians | Phrynobatrachus_rainerguentheri | Phrynobatrachus_rainerguentheri | diurnal |
| amphibians | Phrynobatrachus_ruthbeateae | Phrynobatrachus_ruthbeateae | diurnal |
| amphibians | Phrynobatrachus_sandersoni | Phrynobatrachus_sandersoni | cathemeral |
| amphibians | Phrynobatrachus_scheffleri | Phrynobatrachus_scheffleri | nocturnal |
| amphibians | Phrynobatrachus_schioetzi | Phrynobatrachus_schioetzi | diurnal |
| amphibians | Phrynobatrachus_steindachneri | Phrynobatrachus_steindachneri | cathemeral |
| amphibians | Phrynobatrachus_stewartae | Phrynobatrachus_stewartae | diurnal |
| amphibians | Phrynobatrachus_tokba | Phrynobatrachus_tokba | diurnal |
| amphibians | Phrynobatrachus_villiersi | Phrynobatrachus_villiersi | diurnal |
| amphibians | Phrynobatrachus_werneri | Phrynobatrachus_werneri | diurnal |
| amphibians | Phrynoidis_asper | Phrynoidis_aspera | cathemeral |
| amphibians | Phrynoidis_juxtasper | Phrynoidis_juxtaspera | cathemeral |
| amphibians | Phrynomantis_bifasciatus | Phrynomantis_bifasciatus | nocturnal |
| amphibians | Phrynomantis_microps | Phrynomantis_microps | nocturnal |
| amphibians | Phrynomedusa_marginata | Phrynomedusa_marginata | nocturnal |
| amphibians | Phrynopus_bracki | Phrynopus_bracki | nocturnal |
| amphibians | Phrynopus_dagmarae | Phrynopus_dagmarae | diurnal |
| amphibians | Phrynopus_horstpauli | Phrynopus_horstpauli | nocturnal |
| amphibians | Phrynopus_kauneorum | Phrynopus_kauneorum | diurnal |
| amphibians | Phrynopus_oblivius | Phrynopus_oblivius | diurnal |
| amphibians | Phyllobates_aurotaenia | Phyllobates_aurotaenia | diurnal |
| amphibians | Phyllobates_bicolor | Phyllobates_bicolor | diurnal |
| amphibians | Phyllobates_lugubris | Phyllobates_lugubris | diurnal |
| amphibians | Phyllobates_terribilis | Phyllobates_terribilis | diurnal |
| amphibians | Phyllobates_vittatus | Phyllobates_vittatus | diurnal |
| amphibians | Phyllodytes_kautskyi | Phyllodytes_kautskyi | cathemeral |
| amphibians | Phyllodytes_luteolus | Phyllodytes_luteolus | cathemeral |
| amphibians | Phyllodytes_melanomystax | Phyllodytes_melanomystax | nocturnal |
| amphibians | Phyllodytes_tuberculosus | Phyllodytes_tuberculosus | nocturnal |
| amphibians | Phyllodytes_wuchereri | Phyllodytes_wuchereri | nocturnal |

| | | | |
|---|---|---|---|
| amphibians | Phyllomedusa_bahiana | Phyllomedusa_bahiana | nocturnal |
| amphibians | Phyllomedusa_bicolor | Phyllomedusa_bicolor | nocturnal |
| amphibians | Phyllomedusa_burmeisteri | Phyllomedusa_burmeisteri | nocturnal |
| amphibians | Phyllomedusa_camba | Phyllomedusa_camba | nocturnal |
| amphibians | Phyllomedusa_coelestis | Phyllomedusa_coelestis | nocturnal |
| amphibians | Phyllomedusa_distincta | Phyllomedusa_distincta | nocturnal |
| amphibians | Phyllomedusa_iheringii | Phyllomedusa_iheringii | nocturnal |
| amphibians | Phyllomedusa_neildi | Phyllomedusa_neildi | nocturnal |
| amphibians | Phyllomedusa_sauvagii | Phyllomedusa_sauvagii | nocturnal |
| amphibians | Phyllomedusa_tarsius | Phyllomedusa_tarsius | nocturnal |
| amphibians | Phyllomedusa_tetraploidea | Phyllomedusa_tetraploidea | nocturnal |
| amphibians | Phyllomedusa_vaillantii | Phyllomedusa_vaillantii | nocturnal |
| amphibians | Physalaemus_aguirrei | Physalaemus_aguirrei | nocturnal |
| amphibians | Physalaemus_albonotatus | Physalaemus_albonotatus | cathemeral |
| amphibians | Physalaemus_angrensis | Physalaemus_angrensis | nocturnal |
| amphibians | Physalaemus_atlanticus | Physalaemus_atlanticus | nocturnal |
| amphibians | Physalaemus_barrioi | Physalaemus_barrioi | nocturnal |
| amphibians | Physalaemus_biligonigerus | Physalaemus_biligonigerus | nocturnal |
| amphibians | Physalaemus_bokermanni | Physalaemus_bokermanni | nocturnal |
| amphibians | Physalaemus_camacan | Physalaemus_camacan | nocturnal |
| amphibians | Physalaemus_centralis | Physalaemus_centralis | nocturnal |
| amphibians | Physalaemus_crombiei | Physalaemus_crombiei | nocturnal |
| amphibians | Physalaemus_cuqui | Physalaemus_cuqui | nocturnal |
| amphibians | Physalaemus_cuvieri | Physalaemus_cuvieri | nocturnal |
| amphibians | Physalaemus_deimaticus | Physalaemus_deimaticus | nocturnal |
| amphibians | Physalaemus_erikae | Physalaemus_erikae | cathemeral |
| amphibians | Physalaemus_erythros | Physalaemus_erythros | nocturnal |
| amphibians | Physalaemus_evangelistai | Physalaemus_evangelistai | nocturnal |
| amphibians | Physalaemus_feioi | Physalaemus_feioi | nocturnal |
| amphibians | Physalaemus_fischeri | Physalaemus_fischeri | nocturnal |
| amphibians | Physalaemus_gracilis | Physalaemus_gracilis | cathemeral |
| amphibians | Physalaemus_jordanensis | Physalaemus_jordanensis | nocturnal |
| amphibians | Physalaemus_kroyeri | Physalaemus_kroyeri | nocturnal |
| amphibians | Physalaemus_lateristriga | Physalaemus_lateristriga | nocturnal |

| | | | |
|---|---|---|---|
| amphibians | Physalaemus_lisei | Physalaemus_lisei | cathemeral |
| amphibians | Physalaemus_maculiventris | Physalaemus_maculiventris | nocturnal |
| amphibians | Physalaemus_marmoratus | Physalaemus_marmoratus | nocturnal |
| amphibians | Physalaemus_maximus | Physalaemus_maximus | nocturnal |
| amphibians | Physalaemus_moreirae | Physalaemus_moreirae | nocturnal |
| amphibians | Physalaemus_nanus | Physalaemus_nanus | nocturnal |
| amphibians | Physalaemus_nattereri | Eupemphix_nattereri | cathemeral |
| amphibians | Physalaemus_obtectus | Physalaemus_obtectus | nocturnal |
| amphibians | Physalaemus_olfersii | Physalaemus_olfersii | nocturnal |
| amphibians | Physalaemus_riograndensis | Physalaemus_riograndensis | cathemeral |
| amphibians | Physalaemus_signifer | Physalaemus_signifer | cathemeral |
| amphibians | Physalaemus_soaresi | Physalaemus_soaresi | nocturnal |
| amphibians | Physalaemus_spiniger | Physalaemus_spiniger | cathemeral |
| amphibians | Phyzelaphryne_miriamae | Phyzelaphryne_miriamae | cathemeral |
| amphibians | Pipa_arrabali | Pipa_arrabali | cathemeral |
| amphibians | Pipa_carvalhoi | Pipa_carvalhoi | cathemeral |
| amphibians | Pipa_pipa | Pipa_pipa | cathemeral |
| amphibians | Pithecopus_ayeaye | Phyllomedusa_ayeaye | nocturnal |
| amphibians | Pithecopus_azureus | Phyllomedusa_azurea | nocturnal |
| amphibians | Pithecopus_hypochondrialis | Phyllomedusa_hypochondrialis | cathemeral |
| amphibians | Pithecopus_megacephalus | Phyllomedusa_megacephala | nocturnal |
| amphibians | Pithecopus_nordestinus | Phyllomedusa_nordestina | nocturnal |
| amphibians | Pithecopus_oreades | Phyllomedusa_oreades | nocturnal |
| amphibians | Pithecopus_palliatus | Phyllomedusa_palliata | nocturnal |
| amphibians | Pithecopus_rohdei | Phyllomedusa_rohdei | nocturnal |
| amphibians | Platymantis_corrugatus | Platymantis_corrugatus | nocturnal |
| amphibians | Platymantis_dorsalis | Platymantis_dorsalis | nocturnal |
| amphibians | Platymantis_negrosensis | Platymantis_negrosensis | nocturnal |
| amphibians | Platymantis_paengi | Platymantis_paengi | nocturnal |
| amphibians | Platymantis_panayensis | Platymantis_panayensis | nocturnal |
| amphibians | Platypelis_cowanii | Platypelis_cowanii | nocturnal |
| amphibians | Platyplectrum_ornatum | Platyplectrum_ornatum | nocturnal |
| amphibians | Platyplectrum_spenceri | Platyplectrum_spenceri | nocturnal |
| amphibians | Plectrohyla_avia | Plectrohyla_avia | crepuscular |

| | | | |
|---|---|---|---|
| amphibians | Plectrohyla_chrysopleura | Plectrohyla_chrysopleura | nocturnal |
| amphibians | Plectrohyla_dasypus | Plectrohyla_dasypus | nocturnal |
| amphibians | Plectrohyla_exquisita | Plectrohyla_exquisita | nocturnal |
| amphibians | Plectrohyla_glandulosa | Plectrohyla_glandulosa | diurnal |
| amphibians | Plectrohyla_guatemalensis | Plectrohyla_guatemalensis | cathemeral |
| amphibians | Plectrohyla_hartwegi | Plectrohyla_hartwegi | nocturnal |
| amphibians | Plectrohyla_ixil | Plectrohyla_ixil | cathemeral |
| amphibians | Plectrohyla_matudai | Plectrohyla_matudai | cathemeral |
| amphibians | Plectrohyla_psiloderma | Plectrohyla_psiloderma | nocturnal |
| amphibians | Plectrohyla_quecchi | Plectrohyla_quecchi | nocturnal |
| amphibians | Plectrohyla_sagorum | Plectrohyla_sagorum | cathemeral |
| amphibians | Plethodon_caddoensis | Plethodon_caddoensis | diurnal |
| amphibians | Plethodon_cinereus | Plethodon_cinereus | nocturnal |
| amphibians | Plethodon_cylindraceus | Plethodon_cylindraceus | diurnal |
| amphibians | Plethodon_dunni | Plethodon_dunni | nocturnal |
| amphibians | Plethodon_elongatus | Plethodon_elongatus | nocturnal |
| amphibians | Plethodon_glutinosus | Plethodon_glutinosus | nocturnal |
| amphibians | Plethodon_hubrichti | Plethodon_hubrichti | nocturnal |
| amphibians | Plethodon_idahoensis | Plethodon_idahoensis | nocturnal |
| amphibians | Plethodon_jordani | Plethodon_jordani | nocturnal |
| amphibians | Plethodon_kentucki | Plethodon_kentucki | nocturnal |
| amphibians | Plethodon_metcalfi | Plethodon_metcalfi | cathemeral |
| amphibians | Plethodon_neomexicanus | Plethodon_neomexicanus | nocturnal |
| amphibians | Plethodon_nettingi | Plethodon_nettingi | nocturnal |
| amphibians | Plethodon_ouachitae | Plethodon_ouachitae | diurnal |
| amphibians | Plethodon_punctatus | Plethodon_punctatus | nocturnal |
| amphibians | Plethodon_richmondi | Plethodon_richmondi | nocturnal |
| amphibians | Plethodon_serratus | Plethodon_serratus | cathemeral |
| amphibians | Plethodon_stormi | Plethodon_stormi | nocturnal |
| amphibians | Plethodon_teyahalee | Plethodon_teyahalee | nocturnal |
| amphibians | Plethodon_vehiculum | Plethodon_vehiculum | nocturnal |
| amphibians | Plethodon_wehrlei | Plethodon_wehrlei | nocturnal |
| amphibians | Plethodon_welleri | Plethodon_welleri | nocturnal |
| amphibians | Plethodon_yonahlossee | Plethodon_yonahlossee | nocturnal |

| | | | |
|---|---|---|---|
| amphibians | Plethodontohyla_alluaudi | Rhombophryne_alluaudi | diurnal |
| amphibians | Plethodontohyla_bipunctata | Plethodontohyla_bipunctata | diurnal |
| amphibians | Plethodontohyla_guentheri | Plethodontohyla_guentheri | cathemeral |
| amphibians | Plethodontohyla_inguinalis | Plethodontohyla_inguinalis | nocturnal |
| amphibians | Plethodontohyla_mihanika | Plethodontohyla_mihanika | nocturnal |
| amphibians | Plethodontohyla_notosticta | Plethodontohyla_notosticta | nocturnal |
| amphibians | Plethodontohyla_ocellata | Plethodontohyla_ocellata | nocturnal |
| amphibians | Plethodontohyla_tuberata | Plethodontohyla_tuberata | nocturnal |
| amphibians | Pleurodeles_nebulosus | Pleurodeles_nebulosus | nocturnal |
| amphibians | Pleurodeles_poireti | Pleurodeles_poireti | nocturnal |
| amphibians | Pleurodeles_waltl | Pleurodeles_waltl | nocturnal |
| amphibians | Pleurodema_brachyops | Pleurodema_brachyops | nocturnal |
| amphibians | Pleurodema_cordobae | Pleurodema_cordobae | nocturnal |
| amphibians | Pleurodema_diplolister | Pleurodema_diplolister | nocturnal |
| amphibians | Pleurodema_kriegi | Pleurodema_kriegi | cathemeral |
| amphibians | Pleurodema_nebulosum | Pleurodema_nebulosum | diurnal |
| amphibians | Pleurodema_thaul | Pleurodema_thaul | nocturnal |
| amphibians | Polypedates_leucomystax | Polypedates_leucomystax | nocturnal |
| amphibians | Polypedates_macrotis | Polypedates_macrotis | nocturnal |
| amphibians | Polypedates_maculatus | Polypedates_maculatus | nocturnal |
| amphibians | Polypedates_megacephalus | Polypedates_megacephalus | nocturnal |
| amphibians | Polypedates_otilophus | Polypedates_otilophus | nocturnal |
| amphibians | Polypedates_pseudocruciger | Polypedates_pseudocruciger | nocturnal |
| amphibians | Polypedates_taeniatus | Polypedates_taeniatus | nocturnal |
| amphibians | Potomotyphlus_kaupii | Potamotyphlus_kaupii | nocturnal |
| amphibians | Poyntonia_paludicola | Poyntonia_paludicola | diurnal |
| amphibians | Poyntonophrynus_damaranus | Poyntonophrynus_damaranus | nocturnal |
| amphibians | Pristimantis_acerus | Pristimantis_acerus | cathemeral |
| amphibians | Pristimantis_achatinus | Pristimantis_achatinus | nocturnal |
| amphibians | Pristimantis_actites | Pristimantis_actites | cathemeral |
| amphibians | Pristimantis_acuminatus | Pristimantis_acuminatus | nocturnal |
| amphibians | Pristimantis_adiastolus | Pristimantis_adiastolus | nocturnal |
| amphibians | Pristimantis_altae | Pristimantis_altae | cathemeral |
| amphibians | Pristimantis_altamazonicus | Pristimantis_altamazonicus | cathemeral |

| | | | |
|---|---|---|---|
| amphibians | Pristimantis_altamnis | Pristimantis_altamnis | nocturnal |
| amphibians | Pristimantis_andinognomus | Pristimantis_andinognomus | nocturnal |
| amphibians | Pristimantis_anemerus | Pristimantis_anemerus | nocturnal |
| amphibians | Pristimantis_aniptopalmatus | Pristimantis_aniptopalmatus | nocturnal |
| amphibians | Pristimantis_anolirex | Pristimantis_anolirex | nocturnal |
| amphibians | Pristimantis_apiculatus | Pristimantis_apiculatus | nocturnal |
| amphibians | Pristimantis_appendiculatus | Pristimantis_appendiculatus | nocturnal |
| amphibians | Pristimantis_aquilonaris | Pristimantis_aquilonaris | cathemeral |
| amphibians | Pristimantis_ardalonychus | Pristimantis_ardalonychus | cathemeral |
| amphibians | Pristimantis_atrabracus | Pristimantis_atrabracus | nocturnal |
| amphibians | Pristimantis_atratus | Pristimantis_atratus | nocturnal |
| amphibians | Pristimantis_aureolineatus | Pristimantis_aureolineatus | cathemeral |
| amphibians | Pristimantis_auricarens | Pristimantis_auricarens | nocturnal |
| amphibians | Pristimantis_avius | Pristimantis_avius | nocturnal |
| amphibians | Pristimantis_balionotus | Pristimantis_balionotus | diurnal |
| amphibians | Pristimantis_bambu | Pristimantis_bambu | nocturnal |
| amphibians | Pristimantis_bellator | Pristimantis_bellator | cathemeral |
| amphibians | Pristimantis_bicantus | Pristimantis_bicantus | nocturnal |
| amphibians | Pristimantis_bicumulus | Pristimantis_bicumulus | nocturnal |
| amphibians | Pristimantis_bipunctatus | Pristimantis_bipunctatus | nocturnal |
| amphibians | Pristimantis_bromeliaceus | Pristimantis_bromeliaceus | nocturnal |
| amphibians | Pristimantis_buccinator | Pristimantis_buccinator | cathemeral |
| amphibians | Pristimantis_caeruleonotus | Pristimantis_caeruleonotus | nocturnal |
| amphibians | Pristimantis_cajamarcensis | Pristimantis_cajamarcensis | cathemeral |
| amphibians | Pristimantis_calcarulatus | Pristimantis_calcarulatus | nocturnal |
| amphibians | Pristimantis_cantitans | Pristimantis_cantitans | cathemeral |
| amphibians | Pristimantis_caprifer | Pristimantis_caprifer | nocturnal |
| amphibians | Pristimantis_carlosceroni | Pristimantis_carlosceroni | nocturnal |
| amphibians | Pristimantis_carvalhoi | Pristimantis_carvalhoi | nocturnal |
| amphibians | Pristimantis_celator | Pristimantis_celator | cathemeral |
| amphibians | Pristimantis_cerasinus | Pristimantis_cerasinus | cathemeral |
| amphibians | Pristimantis_ceuthospilus | Pristimantis_ceuthospilus | cathemeral |
| amphibians | Pristimantis_chalceus | Pristimantis_chalceus | nocturnal |
| amphibians | Pristimantis_chiastonotus | Pristimantis_chiastonotus | diurnal |

| | | | |
|---|---|---|---|
| amphibians | Pristimantis_chloronotus | Pristimantis_chloronotus | cathemeral |
| amphibians | Pristimantis_citriogaster | Pristimantis_citriogaster | nocturnal |
| amphibians | Pristimantis_colodactylus | Pristimantis_colodactylus | diurnal |
| amphibians | Pristimantis_colomai | Pristimantis_colomai | nocturnal |
| amphibians | Pristimantis_colonensis | Pristimantis_colonensis | nocturnal |
| amphibians | Pristimantis_condor | Pristimantis_condor | cathemeral |
| amphibians | Pristimantis_conspicillatus | Pristimantis_conspicillatus | cathemeral |
| amphibians | Pristimantis_coronatus | Pristimantis_coronatus | nocturnal |
| amphibians | Pristimantis_cosnipatae | Pristimantis_cosnipatae | nocturnal |
| amphibians | Pristimantis_cremnobates | Pristimantis_cremnobates | nocturnal |
| amphibians | Pristimantis_crenunguis | Pristimantis_crenunguis | nocturnal |
| amphibians | Pristimantis_croceoinguinis | Pristimantis_croceoinguinis | cathemeral |
| amphibians | Pristimantis_crucifer | Pristimantis_crucifer | nocturnal |
| amphibians | Pristimantis_cruentus | Pristimantis_cruentus | nocturnal |
| amphibians | Pristimantis_cryptomelas | Pristimantis_cryptomelas | nocturnal |
| amphibians | Pristimantis_cuneirostris | Pristimantis_cuneirostris | nocturnal |
| amphibians | Pristimantis_danae | Pristimantis_danae | nocturnal |
| amphibians | Pristimantis_degener | Pristimantis_degener | nocturnal |
| amphibians | Pristimantis_dendrobatoides | Pristimantis_dendrobatoides | nocturnal |
| amphibians | Pristimantis_devillei | Pristimantis_devillei | cathemeral |
| amphibians | Pristimantis_diadematus | Pristimantis_diadematus | cathemeral |
| amphibians | Pristimantis_dissimulatus | Pristimantis_dissimulatus | nocturnal |
| amphibians | Pristimantis_duellmani | Pristimantis_duellmani | nocturnal |
| amphibians | Pristimantis_eriphus | Pristimantis_eriphus | nocturnal |
| amphibians | Pristimantis_esmeraldas | Pristimantis_esmeraldas | nocturnal |
| amphibians | Pristimantis_exoristus | Pristimantis_exoristus | nocturnal |
| amphibians | Pristimantis_fenestratus | Pristimantis_fenestratus | cathemeral |
| amphibians | Pristimantis_festae | Pristimantis_festae | cathemeral |
| amphibians | Pristimantis_gaigei | Pristimantis_gaigei | nocturnal |
| amphibians | Pristimantis_galdi | Pristimantis_galdi | nocturnal |
| amphibians | Pristimantis_ganonotus | Pristimantis_ganonotus | diurnal |
| amphibians | Pristimantis_gentryi | Pristimantis_gentryi | nocturnal |
| amphibians | Pristimantis_gladiator | Pristimantis_gladiator | cathemeral |
| amphibians | Pristimantis_glandulosus | Pristimantis_glandulosus | nocturnal |

| | | | |
|---|---|---|---|
| amphibians | Pristimantis_gryllus | Pristimantis_gryllus | cathemeral |
| amphibians | Pristimantis_guaiquinimensis | Pristimantis_guaiquinimensis | cathemeral |
| amphibians | Pristimantis_gutturalis | Pristimantis_gutturalis | diurnal |
| amphibians | Pristimantis_hectus | Pristimantis_hectus | diurnal |
| amphibians | Pristimantis_ignicolor | Pristimantis_ignicolor | nocturnal |
| amphibians | Pristimantis_illotus | Pristimantis_illotus | nocturnal |
| amphibians | Pristimantis_imitatrix | Pristimantis_imitatrix | cathemeral |
| amphibians | Pristimantis_incomptus | Pristimantis_incomptus | nocturnal |
| amphibians | Pristimantis_infraguttatus | Pristimantis_infraguttatus | nocturnal |
| amphibians | Pristimantis_inusitatus | Pristimantis_inusitatus | nocturnal |
| amphibians | Pristimantis_jorgevelosai | Pristimantis_jorgevelosai | nocturnal |
| amphibians | Pristimantis_katoptroides | Pristimantis_katoptroides | nocturnal |
| amphibians | Pristimantis_kichwarum | Pristimantis_kichwarum | nocturnal |
| amphibians | Pristimantis_labiosus | Pristimantis_labiosus | nocturnal |
| amphibians | Pristimantis_lacrimosus | Pristimantis_lacrimosus | cathemeral |
| amphibians | Pristimantis_lanthanites | Pristimantis_lanthanites | cathemeral |
| amphibians | Pristimantis_laticlavius | Pristimantis_laticlavius | nocturnal |
| amphibians | Pristimantis_latidiscus | Pristimantis_latidiscus | nocturnal |
| amphibians | Pristimantis_leoni | Pristimantis_leoni | cathemeral |
| amphibians | Pristimantis_leucopus | Pristimantis_leucopus | nocturnal |
| amphibians | Pristimantis_leucorrhinus | Pristimantis_leucorrhinus | nocturnal |
| amphibians | Pristimantis_librarius | Pristimantis_librarius | nocturnal |
| amphibians | Pristimantis_lindae | Pristimantis_lindae | nocturnal |
| amphibians | Pristimantis_lirellus | Pristimantis_lirellus | nocturnal |
| amphibians | Pristimantis_lividus | Pristimantis_lividus | nocturnal |
| amphibians | Pristimantis_loustes | Pristimantis_loustes | nocturnal |
| amphibians | Pristimantis_lucasi | Pristimantis_lucasi | nocturnal |
| amphibians | Pristimantis_luscombei | Pristimantis_luscombei | nocturnal |
| amphibians | Pristimantis_luteolateralis | Pristimantis_luteolateralis | nocturnal |
| amphibians | Pristimantis_lymani | Pristimantis_lymani | diurnal |
| amphibians | Pristimantis_lythrodes | Pristimantis_lythrodes | nocturnal |
| amphibians | Pristimantis_malkini | Pristimantis_malkini | nocturnal |
| amphibians | Pristimantis_marmoratus | Pristimantis_marmoratus | cathemeral |
| amphibians | Pristimantis_martiae | Pristimantis_martiae | cathemeral |

| | | | |
|---|---|---|---|
| amphibians | Pristimantis_matidiktyo | Pristimantis_matidiktyo | nocturnal |
| amphibians | Pristimantis_mazar | Pristimantis_mazar | nocturnal |
| amphibians | Pristimantis_memorans | Pristimantis_memorans | cathemeral |
| amphibians | Pristimantis_mendax | Pristimantis_mendax | cathemeral |
| amphibians | Pristimantis_minimus | Pristimantis_minimus | nocturnal |
| amphibians | Pristimantis_minutulus | Pristimantis_minutulus | nocturnal |
| amphibians | Pristimantis_modipeplus | Pristimantis_modipeplus | nocturnal |
| amphibians | Pristimantis_moro | Pristimantis_moro | nocturnal |
| amphibians | Pristimantis_muricatus | Pristimantis_muricatus | nocturnal |
| amphibians | Pristimantis_muscosus | Pristimantis_muscosus | cathemeral |
| amphibians | Pristimantis_museosus | Pristimantis_museosus | nocturnal |
| amphibians | Pristimantis_myersi | Pristimantis_myersi | nocturnal |
| amphibians | Pristimantis_nephophilus | Pristimantis_nephophilus | nocturnal |
| amphibians | Pristimantis_nigrogriseus | Pristimantis_nigrogriseus | nocturnal |
| amphibians | Pristimantis_nyctophylax | Pristimantis_nyctophylax | cathemeral |
| amphibians | Pristimantis_ocellatus | Pristimantis_ocellatus | nocturnal |
| amphibians | Pristimantis_ockendeni | Pristimantis_ockendeni | cathemeral |
| amphibians | Pristimantis_ornatissimus | Pristimantis_ornatissimus | nocturnal |
| amphibians | Pristimantis_ornatus | Pristimantis_ornatus | nocturnal |
| amphibians | Pristimantis_ortizi | Pristimantis_ortizi | nocturnal |
| amphibians | Pristimantis_pardalinus | Pristimantis_pardalinus | diurnal |
| amphibians | Pristimantis_pardalis | Pristimantis_pardalis | nocturnal |
| amphibians | Pristimantis_parvillus | Pristimantis_parvillus | nocturnal |
| amphibians | Pristimantis_paulodutrai | Pristimantis_paulodutrai | nocturnal |
| amphibians | Pristimantis_paululus | Pristimantis_paululus | cathemeral |
| amphibians | Pristimantis_pecki | Pristimantis_pecki | nocturnal |
| amphibians | Pristimantis_percnopterus | Pristimantis_percnopterus | cathemeral |
| amphibians | Pristimantis_percultus | Pristimantis_percultus | cathemeral |
| amphibians | Pristimantis_peruvianus | Pristimantis_peruvianus | cathemeral |
| amphibians | Pristimantis_petersi | Pristimantis_petersi | nocturnal |
| amphibians | Pristimantis_petrobardus | Pristimantis_petrobardus | nocturnal |
| amphibians | Pristimantis_phalaroinguinis | Pristimantis_phalaroinguinis | nocturnal |
| amphibians | Pristimantis_philipi | Pristimantis_philipi | diurnal |
| amphibians | Pristimantis_phoxocephalus | Pristimantis_phoxocephalus | cathemeral |

| | | | |
|---|---|---|---|
| amphibians | Pristimantis_pinguis | Pristimantis_pinguis | diurnal |
| amphibians | Pristimantis_pirrensis | Pristimantis_pirrensis | nocturnal |
| amphibians | Pristimantis_prolatus | Pristimantis_prolatus | nocturnal |
| amphibians | Pristimantis_proserpens | Pristimantis_proserpens | nocturnal |
| amphibians | Pristimantis_pseudoacuminatus | Pristimantis_pseudoacuminatus | cathemeral |
| amphibians | Pristimantis_pteridophilus | Pristimantis_pteridophilus | nocturnal |
| amphibians | Pristimantis_pugnax | Pristimantis_pugnax | cathemeral |
| amphibians | Pristimantis_pycnodermis | Pristimantis_pycnodermis | nocturnal |
| amphibians | Pristimantis_pyrrhomerus | Pristimantis_pyrrhomerus | cathemeral |
| amphibians | Pristimantis_quaquaversus | Pristimantis_quaquaversus | nocturnal |
| amphibians | Pristimantis_quinquagesimus | Pristimantis_quinquagesimus | nocturnal |
| amphibians | Pristimantis_ramagii | Pristimantis_ramagii | nocturnal |
| amphibians | Pristimantis_reichlei | Pristimantis_reichlei | nocturnal |
| amphibians | Pristimantis_reticulatus | Pristimantis_reticulatus | nocturnal |
| amphibians | Pristimantis_rhabdocnemus | Pristimantis_rhabdocnemus | nocturnal |
| amphibians | Pristimantis_rhabdolaemus | Pristimantis_rhabdolaemus | nocturnal |
| amphibians | Pristimantis_rhodoplichus | Pristimantis_rhodoplichus | nocturnal |
| amphibians | Pristimantis_rhodostichus | Pristimantis_rhodostichus | cathemeral |
| amphibians | Pristimantis_ridens | Pristimantis_ridens | nocturnal |
| amphibians | Pristimantis_riveti | Pristimantis_riveti | nocturnal |
| amphibians | Pristimantis_romanorum | Pristimantis_romanorum | nocturnal |
| amphibians | Pristimantis_rosadoi | Pristimantis_rosadoi | nocturnal |
| amphibians | Pristimantis_royi | Pristimantis_royi | nocturnal |
| amphibians | Pristimantis_rubicundus | Pristimantis_rubicundus | nocturnal |
| amphibians | Pristimantis_rufioculis | Pristimantis_rufioculis | nocturnal |
| amphibians | Pristimantis_sagittulus | Pristimantis_sagittulus | nocturnal |
| amphibians | Pristimantis_salaputium | Pristimantis_salaputium | nocturnal |
| amphibians | Pristimantis_sarisarinama | Pristimantis_sarisarinama | crepuscular |
| amphibians | Pristimantis_schultei | Pristimantis_schultei | cathemeral |
| amphibians | Pristimantis_scolodiscus | Pristimantis_scolodiscus | nocturnal |
| amphibians | Pristimantis_serendipitus | Pristimantis_serendipitus | nocturnal |
| amphibians | Pristimantis_simonbolivari | Pristimantis_simonbolivari | cathemeral |
| amphibians | Pristimantis_sirnigeli | Pristimantis_sirnigeli | nocturnal |
| amphibians | Pristimantis_skydmainos | Pristimantis_skydmainos | cathemeral |

| | | | |
|---|---|---|---|
| amphibians | Pristimantis_sobetes | Pristimantis_sobetes | nocturnal |
| amphibians | Pristimantis_spinosus | Pristimantis_spinosus | nocturnal |
| amphibians | Pristimantis_sternothylax | Pristimantis_sternothylax | nocturnal |
| amphibians | Pristimantis_stictogaster | Pristimantis_stictogaster | nocturnal |
| amphibians | Pristimantis_subsigillatus | Pristimantis_subsigillatus | nocturnal |
| amphibians | Pristimantis_sulculus | Pristimantis_sulculus | cathemeral |
| amphibians | Pristimantis_supernatis | Pristimantis_supernatis | nocturnal |
| amphibians | Pristimantis_surdus | Pristimantis_surdus | cathemeral |
| amphibians | Pristimantis_tantanti | Pristimantis_tantanti | diurnal |
| amphibians | Pristimantis_tenebrionis | Pristimantis_tenebrionis | nocturnal |
| amphibians | Pristimantis_thymalopsoides | Pristimantis_thymalopsoides | cathemeral |
| amphibians | Pristimantis_thymelensis | Pristimantis_thymelensis | nocturnal |
| amphibians | Pristimantis_toftae | Pristimantis_toftae | cathemeral |
| amphibians | Pristimantis_truebae | Pristimantis_truebae | diurnal |
| amphibians | Pristimantis_tungurahua | Pristimantis_tungurahua | nocturnal |
| amphibians | Pristimantis_unistrigatus | Pristimantis_unistrigatus | nocturnal |
| amphibians | Pristimantis_uranobates | Pristimantis_uranobates | nocturnal |
| amphibians | Pristimantis_variabilis | Pristimantis_variabilis | cathemeral |
| amphibians | Pristimantis_ventrimarmoratus | Pristimantis_ventrimarmoratus | nocturnal |
| amphibians | Pristimantis_verecundus | Pristimantis_verecundus | nocturnal |
| amphibians | Pristimantis_versicolor | Pristimantis_versicolor | nocturnal |
| amphibians | Pristimantis_vertebralis | Pristimantis_vertebralis | nocturnal |
| amphibians | Pristimantis_vilarsi | Pristimantis_vilarsi | nocturnal |
| amphibians | Pristimantis_w-nigrum | Pristimantis_w-nigrum | cathemeral |
| amphibians | Pristimantis_wagteri | Pristimantis_wagteri | nocturnal |
| amphibians | Pristimantis_walkeri | Pristimantis_walkeri | cathemeral |
| amphibians | Pristimantis_waoranii | Pristimantis_waoranii | diurnal |
| amphibians | Pristimantis_wiensi | Pristimantis_wiensi | nocturnal |
| amphibians | Pristimantis_yumbo | Pristimantis_yumbo | nocturnal |
| amphibians | Pristimantis_zeuctotylus | Pristimantis_zeuctotylus | cathemeral |
| amphibians | Pristimantis_zimmermanae | Pristimantis_zimmermanae | nocturnal |
| amphibians | Proceratophrys_appendiculata | Proceratophrys_appendiculata | cathemeral |
| amphibians | Proceratophrys_avelinoi | Proceratophrys_avelinoi | cathemeral |
| amphibians | Proceratophrys_bigibbosa | Proceratophrys_bigibbosa | cathemeral |

| | | | |
|---|---|---|---|
| amphibians | Proceratophrys_boiei | Proceratophrys_boiei | nocturnal |
| amphibians | Proceratophrys_brauni | Proceratophrys_brauni | diurnal |
| amphibians | Proceratophrys_concavitympanum | Proceratophrys_concavitympanum | nocturnal |
| amphibians | Proceratophrys_cururu | Proceratophrys_cururu | nocturnal |
| amphibians | Proceratophrys_goyana | Proceratophrys_goyana | nocturnal |
| amphibians | Proceratophrys_laticeps | Proceratophrys_laticeps | cathemeral |
| amphibians | Proceratophrys_melanopogon | Proceratophrys_melanopogon | nocturnal |
| amphibians | Proceratophrys_minuta | Proceratophrys_minuta | nocturnal |
| amphibians | Proceratophrys_moehringi | Proceratophrys_moehringi | diurnal |
| amphibians | Proceratophrys_moratoi | Proceratophrys_moratoi | cathemeral |
| amphibians | Proceratophrys_palustris | Proceratophrys_palustris | cathemeral |
| amphibians | Proceratophrys_paviotii | Proceratophrys_paviotii | nocturnal |
| amphibians | Proceratophrys_renalis | Proceratophrys_renalis | nocturnal |
| amphibians | Proceratophrys_sanctaritae | Proceratophrys_sanctaritae | cathemeral |
| amphibians | Proceratophrys_schirchi | Proceratophrys_schirchi | nocturnal |
| amphibians | Proceratophrys_subguttata | Proceratophrys_subguttata | nocturnal |
| amphibians | Proceratophrys_tupinamba | Proceratophrys_tupinambis | cathemeral |
| amphibians | Proteus_anguinus | Proteus_anguinus | cathemeral |
| amphibians | Pseudacris_brachyphona | Pseudacris_brachyphona | nocturnal |
| amphibians | Pseudacris_brimleyi | Pseudacris_brimleyi | cathemeral |
| amphibians | Pseudacris_cadaverina | Pseudacris_cadaverina | cathemeral |
| amphibians | Pseudacris_clarkii | Pseudacris_clarkii | cathemeral |
| amphibians | Pseudacris_crucifer | Pseudacris_crucifer | cathemeral |
| amphibians | Pseudacris_feriarum | Pseudacris_feriarum | cathemeral |
| amphibians | Pseudacris_fouquettei | Pseudacris_fouquettei | cathemeral |
| amphibians | Pseudacris_illinoensis | Pseudacris_illinoensis | nocturnal |
| amphibians | Pseudacris_kalmi | Pseudacris_kalmi | cathemeral |
| amphibians | Pseudacris_maculata | Pseudacris_maculata | cathemeral |
| amphibians | Pseudacris_nigrita | Pseudacris_nigrita | cathemeral |
| amphibians | Pseudacris_ocularis | Pseudacris_ocularis | cathemeral |
| amphibians | Pseudacris_ornata | Pseudacris_ornata | nocturnal |
| amphibians | Pseudacris_streckeri | Pseudacris_streckeri | cathemeral |
| amphibians | Pseudacris_triseriata | Pseudacris_triseriata | nocturnal |
| amphibians | Pseudis_bolbodactyla | Pseudis_bolbodactyla | cathemeral |

| | | | |
|---|---|---|---|
| amphibians | Pseudis_cardosoi | Pseudis_cardosoi | cathemeral |
| amphibians | Pseudis_fusca | Pseudis_fusca | nocturnal |
| amphibians | Pseudis_minuta | Pseudis_minuta | nocturnal |
| amphibians | Pseudis_paradoxa | Pseudis_paradoxa | cathemeral |
| amphibians | Pseudis_tocantins | Pseudis_tocantins | cathemeral |
| amphibians | Pseudobranchus_striatus | Pseudobranchus_striatus | nocturnal |
| amphibians | Pseudoeurycea_brunnata | Pseudoeurycea_brunnata | diurnal |
| amphibians | Pseudohynobius_flavomaculatus | Pseudohynobius_flavomaculatus | nocturnal |
| amphibians | Pseudohynobius_jinfo | Pseudohynobius_jinfo | cathemeral |
| amphibians | Pseudopaludicola_ameghini | Pseudopaludicola_ameghini | diurnal |
| amphibians | Pseudopaludicola_boliviana | Pseudopaludicola_boliviana | cathemeral |
| amphibians | Pseudopaludicola_falcipes | Pseudopaludicola_falcipes | cathemeral |
| amphibians | Pseudopaludicola_murundu | Pseudopaludicola_murundu | nocturnal |
| amphibians | Pseudopaludicola_saltica | Pseudopaludicola_saltica | cathemeral |
| amphibians | Pseudophilautus_alto | Pseudophilautus_alto | cathemeral |
| amphibians | Pseudophilautus_asankai | Pseudophilautus_asankai | cathemeral |
| amphibians | Pseudophilautus_caeruleus | Pseudophilautus_caeruleus | nocturnal |
| amphibians | Pseudophilautus_cuspis | Pseudophilautus_cuspis | nocturnal |
| amphibians | Pseudophilautus_decoris | Pseudophilautus_decoris | nocturnal |
| amphibians | Pseudophilautus_femoralis | Pseudophilautus_femoralis | nocturnal |
| amphibians | Pseudophilautus_fulvus | Pseudophilautus_fulvus | nocturnal |
| amphibians | Pseudophilautus_hoffmanni | Pseudophilautus_hoffmanni | nocturnal |
| amphibians | Pseudophilautus_lunatus | Pseudophilautus_lunatus | nocturnal |
| amphibians | Pseudophilautus_mittermeieri | Pseudophilautus_mittermeieri | nocturnal |
| amphibians | Pseudophilautus_procax | Pseudophilautus_procax | nocturnal |
| amphibians | Pseudophilautus_puranappu | Pseudophilautus_puranappu | cathemeral |
| amphibians | Pseudophilautus_rus | Pseudophilautus_rus | nocturnal |
| amphibians | Pseudophilautus_sarasinorum | Pseudophilautus_sarasinorum | nocturnal |
| amphibians | Pseudophilautus_semiruber | Pseudophilautus_semiruber | diurnal |
| amphibians | Pseudophilautus_silus | Pseudophilautus_silus | nocturnal |
| amphibians | Pseudophilautus_silvaticus | Pseudophilautus_silvaticus | nocturnal |
| amphibians | Pseudophilautus_variabilis | Pseudophilautus_variabilis | nocturnal |
| amphibians | Pseudophilautus_zorro | Pseudophilautus_zorro | diurnal |
| amphibians | Pseudophryne_bibronii | Pseudophryne_bibronii | nocturnal |

| | | | |
|---|---|---|---|
| amphibians | Pseudophryne_dendyi | Pseudophryne_dendyi | nocturnal |
| amphibians | Pseudophryne_pengilleyi | Pseudophryne_pengilleyi | nocturnal |
| amphibians | Pseudophryne_semimarmorata | Pseudophryne_semimarmorata | nocturnal |
| amphibians | Pseudotriton_montanus | Pseudotriton_montanus | nocturnal |
| amphibians | Pseudotriton_ruber | Pseudotriton_ruber | nocturnal |
| amphibians | Psychrophrynella_usurpator | Psychrophrynella_usurpator | diurnal |
| amphibians | Ptychadena_anchietae | Ptychadena_anchietae | cathemeral |
| amphibians | Ptychadena_bibroni | Ptychadena_bibroni | cathemeral |
| amphibians | Ptychadena_erlangeri | Ptychadena_erlangeri | cathemeral |
| amphibians | Ptychadena_mascareniensis | Ptychadena_mascareniensis | cathemeral |
| amphibians | Ptychadena_mossambica | Ptychadena_mossambica | nocturnal |
| amphibians | Ptychadena_neumanni | Ptychadena_neumanni | cathemeral |
| amphibians | Ptychadena_oxyrhynchus | Ptychadena_oxyrhynchus | nocturnal |
| amphibians | Ptychadena_porosissima | Ptychadena_porosissima | nocturnal |
| amphibians | Ptychadena_pumilio | Ptychadena_pumilio | cathemeral |
| amphibians | Ptychadena_schillukorum | Ptychadena_schillukorum | cathemeral |
| amphibians | Ptychadena_stenocephala | Ptychadena_stenocephala | diurnal |
| amphibians | Ptychadena_taenioscelis | Ptychadena_taenioscelis | cathemeral |
| amphibians | Ptychadena_trinodis | Ptychadena_trinodis | nocturnal |
| amphibians | Ptychohyla_hypomykter | Ptychohyla_hypomykter | nocturnal |
| amphibians | Pyxicephalus_adspersus | Pyxicephalus_adspersus | diurnal |
| amphibians | Pyxicephalus_edulis | Pyxicephalus_edulis | cathemeral |
| amphibians | Qosqophryne_flammiventris | Bryophryne_flammiventris | diurnal |
| amphibians | Quasipaa_acanthophora | Quasipaa_acanthophora | nocturnal |
| amphibians | Quasipaa_boulengeri | Quasipaa_boulengeri | nocturnal |
| amphibians | Quasipaa_exilispinosa | Quasipaa_exilispinosa | nocturnal |
| amphibians | Quasipaa_fasciculispina | Quasipaa_fasciculispina | nocturnal |
| amphibians | Quasipaa_spinosa | Quasipaa_spinosa | cathemeral |
| amphibians | Quasipaa_verrucospinosa | Quasipaa_verrucospinosa | nocturnal |
| amphibians | Quilticohyla_acrochorda | Ptychohyla_acrochorda | nocturnal |
| amphibians | Quilticohyla_erythromma | Ptychohyla_erythromma | nocturnal |
| amphibians | Quilticohyla_sanctaecrucis | Ptychohyla_sanctaecrucis | nocturnal |
| amphibians | Rana_amurensis | Rana_amurensis | nocturnal |
| amphibians | Rana_arvalis | Rana_arvalis | cathemeral |

| | | | |
|---|---|---|---|
| amphibians | Rana_chensinensis | Rana_chensinensis | diurnal |
| amphibians | Rana_chevronta | Rana_chevronta | nocturnal |
| amphibians | Rana_coreana | Rana_coreana | diurnal |
| amphibians | Rana_dalmatina | Rana_dalmatina | cathemeral |
| amphibians | Rana_graeca | Rana_graeca | cathemeral |
| amphibians | Rana_iberica | Rana_iberica | cathemeral |
| amphibians | Rana_italica | Rana_italica | cathemeral |
| amphibians | Rana_japonica | Rana_japonica | diurnal |
| amphibians | Rana_kukunoris | Rana_kukunoris | diurnal |
| amphibians | Rana_latastei | Rana_latastei | cathemeral |
| amphibians | Rana_pirica | Rana_pirica | cathemeral |
| amphibians | Rana_pyrenaica | Rana_pyrenaica | cathemeral |
| amphibians | Rana_temporaria | Rana_temporaria | cathemeral |
| amphibians | Ranitomeya_amazonica | Ranitomeya_amazonica | diurnal |
| amphibians | Ranitomeya_benedicta | Ranitomeya_benedicta | diurnal |
| amphibians | Ranitomeya_cyanovittata | Ranitomeya_cyanovittata | diurnal |
| amphibians | Ranitomeya_defleri | Ranitomeya_defleri | diurnal |
| amphibians | Ranitomeya_fantastica | Ranitomeya_fantastica | diurnal |
| amphibians | Ranitomeya_flavovittata | Ranitomeya_flavovittata | diurnal |
| amphibians | Ranitomeya_imitator | Ranitomeya_imitator | diurnal |
| amphibians | Ranitomeya_reticulata | Ranitomeya_reticulata | diurnal |
| amphibians | Ranitomeya_sirensis | Ranitomeya_sirensis | diurnal |
| amphibians | Ranitomeya_summersi | Ranitomeya_summersi | diurnal |
| amphibians | Ranitomeya_toraro | Ranitomeya_toraro | diurnal |
| amphibians | Ranitomeya_uakarii | Ranitomeya_uakarii | diurnal |
| amphibians | Ranitomeya_vanzolinii | Ranitomeya_vanzolinii | diurnal |
| amphibians | Ranitomeya_variabilis | Ranitomeya_variabilis | diurnal |
| amphibians | Ranitomeya_ventrimaculata | Ranitomeya_ventrimaculata | diurnal |
| amphibians | Ranitomeya_yavaricola | Ranitomeya_yavaricola | diurnal |
| amphibians | Ranodon_sibiricus | Ranodon_sibiricus | nocturnal |
| amphibians | Raorchestes_annandalii | Pseudophilautus_annandalii | nocturnal |
| amphibians | Raorchestes_bombayensis | Raorchestes_bombayensis | nocturnal |
| amphibians | Raorchestes_chalazodes | Pseudophilautus_chalazodes | nocturnal |
| amphibians | Raorchestes_flaviventris | Pseudophilautus_flaviventris | diurnal |

| | | | |
|---|---|---|---|
| amphibians | Raorchestes_glandulosus | Raorchestes_glandulosus | nocturnal |
| amphibians | Raorchestes_ponmudi | Raorchestes_ponmudi | cathemeral |
| amphibians | Raorchestes_signatus | Raorchestes_signatus | diurnal |
| amphibians | Rentapia_everetti | Pedostibes_everetti | nocturnal |
| amphibians | Rhacophorus_annamensis | Rhacophorus_annamensis | nocturnal |
| amphibians | Rhacophorus_baluensis | Rhacophorus_baluensis | nocturnal |
| amphibians | Rhacophorus_helenae | Rhacophorus_helenae | nocturnal |
| amphibians | Rhacophorus_kio | Rhacophorus_kio | diurnal |
| amphibians | Rhacophorus_malabaricus | Rhacophorus_malabaricus | nocturnal |
| amphibians | Rhacophorus_pardalis | Rhacophorus_pardalis | nocturnal |
| amphibians | Rhacophorus_pseudomalabaricus | Rhacophorus_pseudomalabaricus | nocturnal |
| amphibians | Rhacophorus_rhodopus | Rhacophorus_rhodopus | nocturnal |
| amphibians | Rhaebo_andinophrynoides | Rhaebo_andinophrynoides | cathemeral |
| amphibians | Rhaebo_blombergi | Rhaebo_blombergi | cathemeral |
| amphibians | Rhaebo_caeruleostictus | Rhaebo_caeruleostictus | diurnal |
| amphibians | Rhaebo_ceratophrys | Rhinella_ceratophrys | cathemeral |
| amphibians | Rhaebo_colomai | Andinophryne_colomai | cathemeral |
| amphibians | Rhaebo_ecuadorensis | Rhaebo_ecuadorensis | nocturnal |
| amphibians | Rhaebo_glaberrimus | Rhaebo_glaberrimus | nocturnal |
| amphibians | Rhaebo_guttatus | Rhaebo_guttatus | nocturnal |
| amphibians | Rhaebo_haematiticus | Rhaebo_haematiticus | cathemeral |
| amphibians | Rhaebo_hypomelas | Rhaebo_hypomelas | diurnal |
| amphibians | Rhaebo_olallai | Andinophryne_olallai | nocturnal |
| amphibians | Rheobates_palmatus | Rheobates_palmatus | diurnal |
| amphibians | Rheobatrachus_silus | Rheobatrachus_silus | nocturnal |
| amphibians | Rheobatrachus_vitellinus | Rheobatrachus_vitellinus | cathemeral |
| amphibians | Rheohyla_miotympanum | Ecnomiohyla_miotympanum | nocturnal |
| amphibians | Rhinatrema_shiv | Rhinatrema_shiv | crepuscular |
| amphibians | Rhinella_achavali | Rhinella_achavali | nocturnal |
| amphibians | Rhinella_alata | Rhinella_alata | diurnal |
| amphibians | Rhinella_amabilis | Rhinella_amabilis | nocturnal |
| amphibians | Rhinella_arborescandens | Rhinella_arborescandens | diurnal |
| amphibians | Rhinella_arenarum | Rhinella_arenarum | nocturnal |
| amphibians | Rhinella_arunco | Rhinella_arunco | cathemeral |

| | | | |
|---|---|---|---|
| amphibians | Rhinella_atacamensis | Rhinella_atacamensis | nocturnal |
| amphibians | Rhinella_chavin | Rhinella_chavin | diurnal |
| amphibians | Rhinella_chrysophora | Rhinella_chrysophora | cathemeral |
| amphibians | Rhinella_crucifer | Rhinella_crucifer | nocturnal |
| amphibians | Rhinella_festae | Rhinella_festae | nocturnal |
| amphibians | Rhinella_granulosa | Rhinella_granulosa | nocturnal |
| amphibians | Rhinella_henseli | Rhinella_henseli | nocturnal |
| amphibians | Rhinella_hoogmoedi | Rhinella_hoogmoedi | nocturnal |
| amphibians | Rhinella_humboldti | Rhinella_humboldti | nocturnal |
| amphibians | Rhinella_icterica | Rhinella_icterica | cathemeral |
| amphibians | Rhinella_justinianoi | Rhinella_justinianoi | diurnal |
| amphibians | Rhinella_lescurei | Rhinella_lescurei | diurnal |
| amphibians | Rhinella_magnussoni | Rhinella_magnussoni | diurnal |
| amphibians | Rhinella_manu | Rhinella_manu | nocturnal |
| amphibians | Rhinella_margaritifera | Rhinella_margaritifera | cathemeral |
| amphibians | Rhinella_marina | Rhinella_marina | cathemeral |
| amphibians | Rhinella_nesiotes | Rhinella_nesiotes | nocturnal |
| amphibians | Rhinella_proboscidea | Rhinella_proboscidea | cathemeral |
| amphibians | Rhinella_pygmaea | Rhinella_pygmaea | nocturnal |
| amphibians | Rhinella_roqueana | Rhinella_roqueana | nocturnal |
| amphibians | Rhinella_rubescens | Rhinella_rubescens | cathemeral |
| amphibians | Rhinella_spinulosa | Rhinella_spinulosa | cathemeral |
| amphibians | Rhinella_yanachaga | Rhinella_yanachaga | nocturnal |
| amphibians | Rhinoderma_darwinii | Rhinoderma_darwinii | cathemeral |
| amphibians | Rhinophrynus_dorsalis | Rhinophrynus_dorsalis | cathemeral |
| amphibians | Rhombophryne_coronata | Rhombophryne_coronata | cathemeral |
| amphibians | Rhombophryne_coudreaui | Rhombophryne_coudreaui | cathemeral |
| amphibians | Rhombophryne_guentherpetersi | Rhombophryne_guentherpetersi | nocturnal |
| amphibians | Rhombophryne_laevipes | Rhombophryne_laevipes | cathemeral |
| amphibians | Rhombophryne_mangabensis | Rhombophryne_mangabensis | nocturnal |
| amphibians | Rhombophryne_matavy | Rhombophryne_matavy | cathemeral |
| amphibians | Rhombophryne_minuta | Rhombophryne_minuta | nocturnal |
| amphibians | Rhombophryne_serratopalpebrosa | Rhombophryne_serratopalpebrosa | nocturnal |
| amphibians | Rhombophryne_testudo | Rhombophryne_testudo | cathemeral |

| | | | |
|---|---|---|---|
| amphibians | Rhyacotriton_variegatus | Rhyacotriton_variegatus | nocturnal |
| amphibians | Rulyrana_mcdiarmidi | Rulyrana_mcdiarmidi | nocturnal |
| amphibians | Rulyrana_spiculata | Rulyrana_spiculata | nocturnal |
| amphibians | Rupirana_cardosoi | Rupirana_cardosoi | diurnal |
| amphibians | Sachatamia_albomaculata | Sachatamia_albomaculata | nocturnal |
| amphibians | Sachatamia_ilex | Sachatamia_ilex | nocturnal |
| amphibians | Sachatamia_orejuela | Sachatamia_orejuela | nocturnal |
| amphibians | Salamandra_algira | Salamandra_algira | nocturnal |
| amphibians | Salamandra_atra | Salamandra_atra | nocturnal |
| amphibians | Salamandra_corsica | Salamandra_corsica | nocturnal |
| amphibians | Salamandra_infraimmaculata | Salamandra_infraimmaculata | nocturnal |
| amphibians | Salamandra_lanzai | Salamandra_lanzai | cathemeral |
| amphibians | Salamandra_salamandra | Salamandra_salamandra | cathemeral |
| amphibians | Salamandrella_keyserlingii | Salamandrella_keyserlingii | diurnal |
| amphibians | Salamandrina_perspicillata | Salamandrina_perspicillata | nocturnal |
| amphibians | Salamandrina_terdigitata | Salamandrina_terdigitata | cathemeral |
| amphibians | Sanguirana_aurantipunctata | Sanguirana_aurantipunctata | nocturnal |
| amphibians | Sarcohyla_arborescandens | Plectrohyla_arborescandens | diurnal |
| amphibians | Sarcohyla_bistincta | Plectrohyla_bistincta | cathemeral |
| amphibians | Sarcohyla_charadricola | Plectrohyla_charadricola | nocturnal |
| amphibians | Sarcohyla_chryses | Plectrohyla_chryses | nocturnal |
| amphibians | Sarcohyla_crassa | Plectrohyla_crassa | nocturnal |
| amphibians | Sarcohyla_pentheter | Plectrohyla_pentheter | cathemeral |
| amphibians | Sarcohyla_robertsorum | Plectrohyla_robertsorum | nocturnal |
| amphibians | Sarcohyla_siopela | Plectrohyla_siopela | cathemeral |
| amphibians | Sarcohyla_thorectes | Plectrohyla_thorectes | nocturnal |
| amphibians | Scaphiophryne_boribory | Scaphiophryne_boribory | nocturnal |
| amphibians | Scaphiophryne_calcarata | Scaphiophryne_calcarata | nocturnal |
| amphibians | Scaphiophryne_gottlebei | Scaphiophryne_gottlebei | cathemeral |
| amphibians | Scaphiophryne_madagascariensis | Scaphiophryne_madagascariensis | nocturnal |
| amphibians | Scaphiophryne_marmorata | Scaphiophryne_marmorata | nocturnal |
| amphibians | Scaphiophryne_menabensis | Scaphiophryne_menabensis | nocturnal |
| amphibians | Scaphiophryne_spinosa | Scaphiophryne_spinosa | cathemeral |
| amphibians | Scaphiopus_couchii | Scaphiopus_couchii | nocturnal |

| | | | |
|---|---|---|---|
| amphibians | Scaphiopus_holbrookii | Scaphiopus_holbrookii | nocturnal |
| amphibians | Scaphiopus_hurterii | Scaphiopus_hurterii | nocturnal |
| amphibians | Scarthyla_goinorum | Scarthyla_goinorum | cathemeral |
| amphibians | Scarthyla_vigilans | Scarthyla_vigilans | cathemeral |
| amphibians | Schismaderma_carens | Schismaderma_carens | cathemeral |
| amphibians | Scinax_acuminatus | Scinax_acuminatus | nocturnal |
| amphibians | Scinax_alter | Scinax_alter | cathemeral |
| amphibians | Scinax_baumgardneri | Scinax_baumgardneri | nocturnal |
| amphibians | Scinax_blairi | Scinax_blairi | nocturnal |
| amphibians | Scinax_boesemani | Scinax_boesemani | nocturnal |
| amphibians | Scinax_boulengeri | Scinax_boulengeri | nocturnal |
| amphibians | Scinax_caldarum | Scinax_caldarum | nocturnal |
| amphibians | Scinax_cretatus | Scinax_cretatus | nocturnal |
| amphibians | Scinax_crospedospilus | Scinax_crospedospilus | nocturnal |
| amphibians | Scinax_cuspidatus | Scinax_cuspidatus | nocturnal |
| amphibians | Scinax_danae | Scinax_danae | nocturnal |
| amphibians | Scinax_duartei | Scinax_duartei | nocturnal |
| amphibians | Scinax_elaeochroa | Scinax_elaeochroa | nocturnal |
| amphibians | Scinax_eurydice | Scinax_eurydice | nocturnal |
| amphibians | Scinax_exiguus | Scinax_exiguus | nocturnal |
| amphibians | Scinax_funereus | Scinax_funereus | nocturnal |
| amphibians | Scinax_fuscomarginatus | Scinax_fuscomarginatus | nocturnal |
| amphibians | Scinax_fuscovarius | Scinax_fuscovarius | nocturnal |
| amphibians | Scinax_garbei | Scinax_garbei | nocturnal |
| amphibians | Scinax_granulatus | Scinax_granulatus | nocturnal |
| amphibians | Scinax_hayii | Scinax_hayii | nocturnal |
| amphibians | Scinax_ictericus | Scinax_ictericus | cathemeral |
| amphibians | Scinax_imbegue | Scinax_imbegue | nocturnal |
| amphibians | Scinax_iquitorum | Scinax_iquitorum | nocturnal |
| amphibians | Scinax_juncae | Scinax_juncae | nocturnal |
| amphibians | Scinax_kennedyi | Scinax_kennedyi | nocturnal |
| amphibians | Scinax_nasicus | Scinax_nasicus | nocturnal |
| amphibians | Scinax_nebulosus | Scinax_nebulosus | nocturnal |
| amphibians | Scinax_pachycrus | Scinax_pachycrus | nocturnal |

| | | | |
|---|---|---|---|
| amphibians | Scinax_perereca | Scinax_perereca | nocturnal |
| amphibians | Scinax_quinquefasciatus | Scinax_quinquefasciatus | cathemeral |
| amphibians | Scinax_rogerioi | Scinax_rogerioi | cathemeral |
| amphibians | Scinax_ruber | Scinax_ruber | cathemeral |
| amphibians | Scinax_similis | Scinax_similis | nocturnal |
| amphibians | Scinax_squalirostris | Scinax_squalirostris | nocturnal |
| amphibians | Scinax_staufferi | Scinax_staufferi | nocturnal |
| amphibians | Scinax_sugillatus | Scinax_sugillatus | nocturnal |
| amphibians | Scinax_tigrinus | Scinax_tigrinus | nocturnal |
| amphibians | Scinax_tymbamirim | Scinax_tymbamirim | nocturnal |
| amphibians | Scinax_wandae | Scinax_wandae | nocturnal |
| amphibians | Scinax_x-signatus | Scinax_x-signatus | nocturnal |
| amphibians | Sclerophrys_channingi | Amietophrynus_channingi | nocturnal |
| amphibians | Sclerophrys_dodsoni | Duttaphrynus_dodsoni | cathemeral |
| amphibians | Sclerophrys_garmani | Amietophrynus_garmani | nocturnal |
| amphibians | Sclerophrys_gracilipes | Amietophrynus_gracilipes | cathemeral |
| amphibians | Sclerophrys_gutturalis | Amietophrynus_gutturalis | cathemeral |
| amphibians | Sclerophrys_hadramautinus | Bufo_arabicus | cathemeral |
| amphibians | Sclerophrys_kassasii | Amietophrynus_kassasii | cathemeral |
| amphibians | Sclerophrys_mauritanica | Amietophrynus_mauritanicus | cathemeral |
| amphibians | Sclerophrys_pantherina | Amietophrynus_pantherinus | nocturnal |
| amphibians | Sclerophrys_pentoni | Bufo_pentoni | cathemeral |
| amphibians | Sclerophrys_perreti | Amietophrynus_perreti | cathemeral |
| amphibians | Sclerophrys_regularis | Amietophrynus_regularis | nocturnal |
| amphibians | Sclerophrys_superciliaris | Amietophrynus_superciliaris | nocturnal |
| amphibians | Sclerophrys_tuberosa | Amietophrynus_tuberosus | diurnal |
| amphibians | Sclerophrys_villiersi | Amietophrynus_villiersi | cathemeral |
| amphibians | Sclerophrys_xeros | Amietophrynus_xeros | nocturnal |
| amphibians | Scotobleps_gabonicus | Scotobleps_gabonicus | nocturnal |
| amphibians | Scutiger_boulengeri | Scutiger_boulengeri | nocturnal |
| amphibians | Scutiger_chintingensis | Scutiger_chintingensis | nocturnal |
| amphibians | Scutiger_liupanensis | Scutiger_liupanensis | nocturnal |
| amphibians | Scutiger_mammatus | Scutiger_mammatus | nocturnal |
| amphibians | Scutiger_muliensis | Scutiger_muliensis | cathemeral |

| | | | |
|---|---|---|---|
| amphibians | Scutiger_pingwuensis | Scutiger_pingwuensis | nocturnal |
| amphibians | Scutiger_wuguanfui | Scutiger_wuguanfui | nocturnal |
| amphibians | Scythrophrys_sawayae | Scythrophrys_sawayae | cathemeral |
| amphibians | Sechellophryne_gardineri | Sechellophryne_gardineri | nocturnal |
| amphibians | Silverstoneia_dalyi | Silverstoneia_dalyi | diurnal |
| amphibians | Silverstoneia_flotator | Silverstoneia_flotator | diurnal |
| amphibians | Silverstoneia_gutturalis | Silverstoneia_gutturalis | diurnal |
| amphibians | Silverstoneia_minima | Silverstoneia_minima | diurnal |
| amphibians | Silverstoneia_minutissima | Silverstoneia_minutissima | diurnal |
| amphibians | Silverstoneia_nubicola | Silverstoneia_nubicola | diurnal |
| amphibians | Silverstoneia_punctiventris | Silverstoneia_punctiventris | diurnal |
| amphibians | Siphonops_annulatus | Siphonops_annulatus | nocturnal |
| amphibians | Siren_intermedia | Siren_intermedia | nocturnal |
| amphibians | Siren_lacertina | Siren_lacertina | nocturnal |
| amphibians | Smilisca_baudinii | Smilisca_baudinii | nocturnal |
| amphibians | Smilisca_cyanosticta | Smilisca_cyanosticta | cathemeral |
| amphibians | Smilisca_dentata | Smilisca_dentata | nocturnal |
| amphibians | Smilisca_fodiens | Smilisca_fodiens | diurnal |
| amphibians | Smilisca_phaeota | Smilisca_phaeota | cathemeral |
| amphibians | Smilisca_puma | Smilisca_puma | nocturnal |
| amphibians | Smilisca_sila | Smilisca_sila | nocturnal |
| amphibians | Smilisca_sordida | Smilisca_sordida | cathemeral |
| amphibians | Sooglossus_thomasseti | Sooglossus_thomasseti | nocturnal |
| amphibians | Spea_bombifrons | Spea_bombifrons | nocturnal |
| amphibians | Spea_hammondii | Spea_hammondii | nocturnal |
| amphibians | Spea_intermontana | Spea_intermontana | nocturnal |
| amphibians | Spea_multiplicata | Spea_multiplicata | nocturnal |
| amphibians | Spelaeophryne_methneri | Spelaeophryne_methneri | diurnal |
| amphibians | Speleomantes_ambrosii | Hydromantes_ambrosii | cathemeral |
| amphibians | Speleomantes_flavus | Hydromantes_flavus | cathemeral |
| amphibians | Speleomantes_genei | Hydromantes_genei | cathemeral |
| amphibians | Speleomantes_imperialis | Hydromantes_imperialis | nocturnal |
| amphibians | Speleomantes_italicus | Hydromantes_italicus | cathemeral |
| amphibians | Speleomantes_sarrabusensis | Hydromantes_sarrabusensis | cathemeral |

| | | | |
|---|---|---|---|
| amphibians | Speleomantes_strinatii | Hydromantes_strinatii | cathemeral |
| amphibians | Speleomantes_supramontis | Hydromantes_supramontis | cathemeral |
| amphibians | Sphaenorhynchus_botocudo | Sphaenorhynchus_botocudo | nocturnal |
| amphibians | Sphaenorhynchus_caramaschii | Sphaenorhynchus_caramaschii | nocturnal |
| amphibians | Sphaenorhynchus_carneus | Sphaenorhynchus_carneus | nocturnal |
| amphibians | Sphaenorhynchus_dorisae | Sphaenorhynchus_dorisae | nocturnal |
| amphibians | Sphaenorhynchus_lacteus | Sphaenorhynchus_lacteus | nocturnal |
| amphibians | Sphaenorhynchus_mirim | Sphaenorhynchus_mirim | nocturnal |
| amphibians | Sphaenorhynchus_palustris | Sphaenorhynchus_palustris | nocturnal |
| amphibians | Sphaenorhynchus_pauloalvini | Sphaenorhynchus_pauloalvini | nocturnal |
| amphibians | Sphaenorhynchus_planicola | Sphaenorhynchus_planicola | nocturnal |
| amphibians | Sphaenorhynchus_prasinus | Sphaenorhynchus_prasinus | nocturnal |
| amphibians | Sphaenorhynchus_surdus | Sphaenorhynchus_surdus | nocturnal |
| amphibians | Sphaerotheca_dobsonii | Sphaerotheca_dobsonii | cathemeral |
| amphibians | Sphaerotheca_rolandae | Sphaerotheca_rolandae | nocturnal |
| amphibians | Sphenophryne_cornuta | Sphenophryne_cornuta | nocturnal |
| amphibians | Sphenophryne_magnitympanum | Liophryne_magnitympanum | nocturnal |
| amphibians | Sphenophryne_schlaginhaufeni | Liophryne_schlaginhaufeni | cathemeral |
| amphibians | Sphenophryne_similis | Liophryne_similis | nocturnal |
| amphibians | Spicospina_flammocaerulea | Spicospina_flammocaerulea | nocturnal |
| amphibians | Spinomantis_aglavei | Spinomantis_aglavei | nocturnal |
| amphibians | Spinomantis_bertini | Spinomantis_bertini | diurnal |
| amphibians | Spinomantis_brunae | Spinomantis_brunae | cathemeral |
| amphibians | Spinomantis_elegans | Spinomantis_elegans | cathemeral |
| amphibians | Spinomantis_fimbriatus | Spinomantis_fimbriatus | nocturnal |
| amphibians | Spinomantis_guibei | Spinomantis_guibei | diurnal |
| amphibians | Spinomantis_massi | Spinomantis_massi | crepuscular |
| amphibians | Spinomantis_microtis | Spinomantis_microtis | diurnal |
| amphibians | Spinomantis_nussbaumi | Spinomantis_nussbaumi | nocturnal |
| amphibians | Spinomantis_peraccae | Spinomantis_peraccae | nocturnal |
| amphibians | Spinomantis_phantasticus | Spinomantis_phantasticus | nocturnal |
| amphibians | Spinomantis_tavaratra | Spinomantis_tavaratra | nocturnal |
| amphibians | Staurois_guttatus | Staurois_guttatus | diurnal |
| amphibians | Staurois_latopalmatus | Staurois_latopalmatus | cathemeral |

| | | | |
|---|---|---|---|
| amphibians | Staurois_natator | Staurois_natator | cathemeral |
| amphibians | Staurois_parvus | Staurois_parvus | diurnal |
| amphibians | Staurois_tuberilinguis | Staurois_tuberilinguis | diurnal |
| amphibians | Stefania_ackawaio | Stefania_ackawaio | nocturnal |
| amphibians | Stefania_ayangannae | Stefania_ayangannae | nocturnal |
| amphibians | Stefania_coxi | Stefania_coxi | nocturnal |
| amphibians | Stefania_evansi | Stefania_evansi | diurnal |
| amphibians | Stefania_neblinae | Stefania_neblinae | cathemeral |
| amphibians | Stefania_riae | Stefania_riae | nocturnal |
| amphibians | Stefania_satelles | Stefania_satelles | nocturnal |
| amphibians | Stefania_scalae | Stefania_scalae | nocturnal |
| amphibians | Stefania_schuberti | Stefania_schuberti | nocturnal |
| amphibians | Stefania_tamacuarina | Stefania_tamacuarina | nocturnal |
| amphibians | Stereocyclops_histrio | Stereocyclops_histrio | nocturnal |
| amphibians | Stereocyclops_incrassatus | Stereocyclops_incrassatus | nocturnal |
| amphibians | Stereocyclops_parkeri | Stereocyclops_parkeri | nocturnal |
| amphibians | Strabomantis_anatipes | Strabomantis_anatipes | nocturnal |
| amphibians | Strabomantis_anomalus | Strabomantis_anomalus | nocturnal |
| amphibians | Strabomantis_biporcatus | Strabomantis_biporcatus | nocturnal |
| amphibians | Strabomantis_bufoniformis | Strabomantis_bufoniformis | nocturnal |
| amphibians | Strabomantis_cerastes | Strabomantis_cerastes | cathemeral |
| amphibians | Strabomantis_cornutus | Strabomantis_cornutus | cathemeral |
| amphibians | Strabomantis_helonotus | Strabomantis_helonotus | nocturnal |
| amphibians | Strabomantis_laticorpus | Strabomantis_laticorpus | diurnal |
| amphibians | Strabomantis_necerus | Strabomantis_necerus | cathemeral |
| amphibians | Strabomantis_sulcatus | Strabomantis_sulcatus | cathemeral |
| amphibians | Strauchbufo_raddei | Strauchbufo_raddei | diurnal |
| amphibians | Strongylopus_fasciatus | Strongylopus_fasciatus | cathemeral |
| amphibians | Stumpffia_analamaina | Stumpffia_analamaina | cathemeral |
| amphibians | Stumpffia_be | Stumpffia_be | nocturnal |
| amphibians | Stumpffia_gimmeli | Stumpffia_gimmeli | nocturnal |
| amphibians | Stumpffia_grandis | Stumpffia_grandis | nocturnal |
| amphibians | Stumpffia_hara | Stumpffia_hara | cathemeral |
| amphibians | Stumpffia_madagascariensis | Stumpffia_madagascariensis | crepuscular |

| | | | |
|---|---|---|---|
| amphibians | Stumpffia_miery | Stumpffia_miery | nocturnal |
| amphibians | Stumpffia_psologlossa | Stumpffia_psologlossa | cathemeral |
| amphibians | Stumpffia_pygmaea | Stumpffia_pygmaea | cathemeral |
| amphibians | Stumpffia_roseifemoralis | Stumpffia_roseifemoralis | nocturnal |
| amphibians | Stumpffia_staffordi | Stumpffia_staffordi | diurnal |
| amphibians | Stumpffia_tetradactyla | Stumpffia_tetradactyla | cathemeral |
| amphibians | Stumpffia_tridactyla | Stumpffia_tridactyla | cathemeral |
| amphibians | Synapturanus_mirandaribeiroi | Synapturanus_mirandaribeiroi | cathemeral |
| amphibians | Synapturanus_rabus | Synapturanus_rabus | nocturnal |
| amphibians | Synapturanus_salseri | Synapturanus_salseri | cathemeral |
| amphibians | Tachycnemis_seychellensis | Tachycnemis_seychellensis | nocturnal |
| amphibians | Taricha_granulosa | Taricha_granulosa | cathemeral |
| amphibians | Taricha_rivularis | Taricha_rivularis | nocturnal |
| amphibians | Taricha_torosa | Taricha_torosa | nocturnal |
| amphibians | Taudactylus_acutirostris | Taudactylus_acutirostris | diurnal |
| amphibians | Taudactylus_diurnus | Taudactylus_diurnus | diurnal |
| amphibians | Taudactylus_eungellensis | Taudactylus_eungellensis | cathemeral |
| amphibians | Taudactylus_pleione | Taudactylus_pleione | diurnal |
| amphibians | Taudactylus_rheophilus | Taudactylus_rheophilus | cathemeral |
| amphibians | Telmatobius_atahualpai | Telmatobius_atahualpai | cathemeral |
| amphibians | Telmatobius_bolivianus | Telmatobius_bolivianus | nocturnal |
| amphibians | Telmatobius_carrillae | Telmatobius_carrillae | nocturnal |
| amphibians | Telmatobius_cirrhacelis | Telmatobius_cirrhacelis | nocturnal |
| amphibians | Telmatobius_culeus | Telmatobius_culeus | nocturnal |
| amphibians | Telmatobius_niger | Telmatobius_niger | diurnal |
| amphibians | Telmatobius_thompsoni | Telmatobius_thompsoni | diurnal |
| amphibians | Telmatobius_truebae | Telmatobius_truebae | nocturnal |
| amphibians | Tepuihyla_edelcae | Tepuihyla_edelcae | nocturnal |
| amphibians | Tepuihyla_exophthalma | Tepuihyla_exophthalma | nocturnal |
| amphibians | Tepuihyla_tuberculosa | Ecnomiohyla_tuberculosa | nocturnal |
| amphibians | Teratohyla_amelie | Teratohyla_amelie | nocturnal |
| amphibians | Teratohyla_midas | Teratohyla_midas | nocturnal |
| amphibians | Teratohyla_pulverata | Teratohyla_pulverata | nocturnal |
| amphibians | Teratohyla_spinosa | Teratohyla_spinosa | nocturnal |

| | | | |
|---|---|---|---|
| amphibians | Theloderma_asperum | Theloderma_asperum | nocturnal |
| amphibians | Theloderma_horridum | Theloderma_horridum | nocturnal |
| amphibians | Theloderma_laeve | Theloderma_laeve | nocturnal |
| amphibians | Thoropa_lutzi | Thoropa_lutzi | nocturnal |
| amphibians | Thoropa_megatympanum | Thoropa_megatympanum | nocturnal |
| amphibians | Thoropa_miliaris | Thoropa_miliaris | nocturnal |
| amphibians | Thoropa_petropolitana | Thoropa_petropolitana | nocturnal |
| amphibians | Thoropa_saxatilis | Thoropa_saxatilis | nocturnal |
| amphibians | Thoropa_taophora | Thoropa_taophora | nocturnal |
| amphibians | Tlalocohyla_loquax | Tlalocohyla_loquax | nocturnal |
| amphibians | Tlalocohyla_picta | Tlalocohyla_picta | nocturnal |
| amphibians | Tlalocohyla_smithii | Tlalocohyla_smithii | nocturnal |
| amphibians | Tomopterna_cryptotis | Tomopterna_cryptotis | nocturnal |
| amphibians | Trachycephalus_atlas | Trachycephalus_atlas | nocturnal |
| amphibians | Trachycephalus_coriaceus | Trachycephalus_coriaceus | nocturnal |
| amphibians | Trachycephalus_cunauaru | Trachycephalus_cunauaru | nocturnal |
| amphibians | Trachycephalus_dibernardoi | Trachycephalus_dibernardoi | nocturnal |
| amphibians | Trachycephalus_imitatrix | Trachycephalus_imitatrix | nocturnal |
| amphibians | Trachycephalus_jordani | Trachycephalus_jordani | nocturnal |
| amphibians | Trachycephalus_lepidus | Trachycephalus_lepidus | nocturnal |
| amphibians | Trachycephalus_mambaiensis | Trachycephalus_mambaiensis | nocturnal |
| amphibians | Trachycephalus_mesophaeus | Trachycephalus_mesophaeus | cathemeral |
| amphibians | Trachycephalus_nigromaculatus | Trachycephalus_nigromaculatus | nocturnal |
| amphibians | Trachycephalus_resinifictrix | Trachycephalus_resinifictrix | nocturnal |
| amphibians | Trachycephalus_typhonius | Trachycephalus_venulosus | cathemeral |
| amphibians | Trachycephalus_venezolanus | Aparasphenodon_venezolanus | nocturnal |
| amphibians | Triprion_petasatus | Triprion_petasatus | cathemeral |
| amphibians | Triprion_spatulatus | Diaglena_spatulata | nocturnal |
| amphibians | Triprion_spinosus | Anotheca_spinosa | cathemeral |
| amphibians | Triturus_carnifex | Triturus_carnifex | nocturnal |
| amphibians | Triturus_cristatus | Triturus_cristatus | cathemeral |
| amphibians | Triturus_dobrogicus | Triturus_dobrogicus | nocturnal |
| amphibians | Triturus_karelinii | Triturus_karelinii | nocturnal |
| amphibians | Triturus_marmoratus | Triturus_marmoratus | cathemeral |

| | | | |
|---|---|---|---|
| amphibians | Triturus_pygmaeus | Triturus_pygmaeus | nocturnal |
| amphibians | Tsingymantis_antitra | Tsingymantis_antitra | nocturnal |
| amphibians | Tylototriton_asperrimus | Tylototriton_asperrimus | nocturnal |
| amphibians | Tylototriton_kweichowensis | Tylototriton_kweichowensis | nocturnal |
| amphibians | Tylototriton_lizhenchangi | Tylototriton_lizhenchangi | nocturnal |
| amphibians | Tylototriton_pseudoverrucosus | Tylototriton_pseudoverrucosus | cathemeral |
| amphibians | Tylototriton_shanjing | Tylototriton_shanjing | nocturnal |
| amphibians | Tylototriton_taliangensis | Tylototriton_taliangensis | nocturnal |
| amphibians | Tylototriton_uyenoi | Tylototriton_uyenoi | nocturnal |
| amphibians | Tylototriton_verrucosus | Tylototriton_verrucosus | cathemeral |
| amphibians | Tylototriton_wenxianensis | Tylototriton_wenxianensis | nocturnal |
| amphibians | Tylototriton_yangi | Tylototriton_yangi | nocturnal |
| amphibians | Typhlonectes_compressicauda | Typhlonectes_compressicauda | nocturnal |
| amphibians | Typhlonectes_natans | Typhlonectes_natans | nocturnal |
| amphibians | Uperodon_globulosus | Uperodon_globulosus | nocturnal |
| amphibians | Uperodon_nagaoi | Ramanella_nagaoi | nocturnal |
| amphibians | Uperodon_systoma | Uperodon_systoma | nocturnal |
| amphibians | Uperodon_taprobanicus | Kaloula_taprobanica | nocturnal |
| amphibians | Uperoleia_aspera | Uperoleia_aspera | nocturnal |
| amphibians | Uperoleia_daviesae | Uperoleia_daviesae | nocturnal |
| amphibians | Uperoleia_laevigata | Uperoleia_laevigata | nocturnal |
| amphibians | Uperoleia_micra | Uperoleia_micra | nocturnal |
| amphibians | Uperoleia_orientalis | Uperoleia_orientalis | nocturnal |
| amphibians | Vitreorana_eurygnatha | Vitreorana_eurygnatha | nocturnal |
| amphibians | Vitreorana_gorzulae | Vitreorana_gorzulae | nocturnal |
| amphibians | Vitreorana_helenae | Vitreorana_helenae | nocturnal |
| amphibians | Vitreorana_ritae | Cochranella_ritae | nocturnal |
| amphibians | Vitreorana_uranoscopa | Vitreorana_uranoscopa | nocturnal |
| amphibians | Wakea_madinika | Wakea_madinika | cathemeral |
| amphibians | Walkerana_diplosticta | Indirana_diplosticta | crepuscular |
| amphibians | Walkerana_leptodactyla | Indirana_leptodactyla | nocturnal |
| amphibians | Walkerana_phrynoderma | Indirana_phrynoderma | diurnal |
| amphibians | Werneria_mertensiana | Werneria_mertensiana | diurnal |
| amphibians | Werneria_preussi | Werneria_preussi | cathemeral |

| | | | |
|---|---|---|---|
| amphibians | Werneria_submontana | Werneria_submontana | diurnal |
| amphibians | Wijayarana_masonii | Huia_masonii | nocturnal |
| amphibians | Wolterstorffina_parvipalmata | Wolterstorffina_parvipalmata | nocturnal |
| amphibians | Xenohyla_truncata | Xenohyla_truncata | nocturnal |
| amphibians | Xenophrys_megacephala | Megophrys_megacephala | diurnal |
| amphibians | Xenophrys_zunhebotoensis | Xenophrys_zunhebotoensis | diurnal |
| amphibians | Xenopus_andrei | Xenopus_andrei | nocturnal |
| amphibians | Xenopus_borealis | Xenopus_borealis | nocturnal |
| amphibians | Xenopus_laevis | Xenopus_laevis | cathemeral |
| amphibians | Xenopus_muelleri | Xenopus_muelleri | nocturnal |
| amphibians | Xenopus_tropicalis | Xenopus_tropicalis | nocturnal |
| amphibians | Xenopus_wittei | Xenopus_wittei | nocturnal |
| amphibians | Xenorhina_arboricola | Xenorhina_arboricola | nocturnal |
| amphibians | Xenorhina_rostrata | Xenorhina_rostrata | diurnal |
| amphibians | Xenorhina_varia | Xenorhina_varia | nocturnal |
| amphibians | Xenorhina_zweifeli | Xenorhina_zweifeli | diurnal |
| amphibians | Yunganastes_bisignatus | Pristimantis_bisignatus | nocturnal |
| amphibians | Yunganastes_mercedesae | Pristimantis_mercedesae | cathemeral |
| amphibians | Yunganastes_pluvicanorus | Pristimantis_pluvicanorus | cathemeral |
| amphibians | Zhangixalus_feae | Rhacophorus_feae | nocturnal |
| amphibians | Zhangixalus_omeimontis | Rhacophorus_omeimontis | nocturnal |
| amphibians | Zhangixalus_schlegelii | Rhacophorus_schlegelii | diurnal |
| lizards | Ablepharus_bivittatus | Ablepharus_bivittatus | diurnal |
| lizards | Ablepharus_budaki | Ablepharus_budaki | diurnal |
| lizards | Ablepharus_chernovi | Ablepharus_chernovi | diurnal |
| lizards | Ablepharus_deserti | Ablepharus_deserti | diurnal |
| lizards | Ablepharus_grayanus | Ablepharus_grayanus | diurnal |
| lizards | Ablepharus_kitaibelii | Ablepharus_kitaibelii | cathemeral |
| lizards | Ablepharus_lindbergi | Ablepharus_lindbergi | diurnal |
| lizards | Ablepharus_pannonicus | Ablepharus_pannonicus | diurnal |
| lizards | Ablepharus_rueppellii | Ablepharus_rueppellii | diurnal |
| lizards | Abronia_aurita | Abronia_aurita | diurnal |
| lizards | Abronia_campbelli | Abronia_campbelli | diurnal |
| lizards | Abronia_deppii | Abronia_deppii | diurnal |

| | | | |
|---|---|---|---|
| lizards | Abronia_fimbriata | Abronia_fimbriata | diurnal |
| lizards | Abronia_graminea | Abronia_graminea | diurnal |
| lizards | Abronia_lythrochila | Abronia_lythrochila | diurnal |
| lizards | Abronia_meledona | Abronia_meledona | diurnal |
| lizards | Abronia_mixteca | Abronia_mixteca | diurnal |
| lizards | Abronia_oaxacae | Abronia_oaxacae | diurnal |
| lizards | Abronia_ochoterenai | Abronia_ochoterenai | diurnal |
| lizards | Abronia_smithi | Abronia_smithi | diurnal |
| lizards | Acanthocercus_adramitanus | Acanthocercus_adramitanus | diurnal |
| lizards | Acanthocercus_atricollis | Acanthocercus_atricollis | diurnal |
| lizards | Acanthocercus_cyanogaster | Acanthocercus_cyanogaster | diurnal |
| lizards | Acanthodactylus_aegyptius | Acanthodactylus_aegyptius | diurnal |
| lizards | Acanthodactylus_bedriagai | Acanthodactylus_bedriagai | diurnal |
| lizards | Acanthodactylus_blanci | Acanthodactylus_blanci | diurnal |
| lizards | Acanthodactylus_blanfordii | Acanthodactylus_blanfordii | diurnal |
| lizards | Acanthodactylus_boskianus | Acanthodactylus_boskianus | diurnal |
| lizards | Acanthodactylus_cantoris | Acanthodactylus_cantoris | diurnal |
| lizards | Acanthodactylus_dumerilii | Acanthodactylus_dumerilii | diurnal |
| lizards | Acanthodactylus_erythrurus | Acanthodactylus_erythrurus | diurnal |
| lizards | Acanthodactylus_hardyi | Acanthodactylus_hardyi | diurnal |
| lizards | Acanthodactylus_longipes | Acanthodactylus_longipes | diurnal |
| lizards | Acanthodactylus_maculatus | Acanthodactylus_maculatus | diurnal |
| lizards | Acanthodactylus_opheodurus | Acanthodactylus_opheodurus | diurnal |
| lizards | Acanthodactylus_pardalis | Acanthodactylus_pardalis | diurnal |
| lizards | Acanthodactylus_savignyi | Acanthodactylus_savignyi | diurnal |
| lizards | Acanthodactylus_schmidti | Acanthodactylus_schmidti | diurnal |
| lizards | Acanthodactylus_schreiberi | Acanthodactylus_schreiberi | diurnal |
| lizards | Acanthodactylus_scutellatus | Acanthodactylus_scutellatus | diurnal |
| lizards | Acanthodactylus_spinicauda | Acanthodactylus_spinicauda | diurnal |
| lizards | Acanthodactylus_tristrami | Acanthodactylus_tristrami | diurnal |
| lizards | Acanthosaura_armata | Acanthosaura_armata | diurnal |
| lizards | Acanthosaura_capra | Acanthosaura_capra | diurnal |
| lizards | Acanthosaura_cardamomensis | Acanthosaura_cardamomensis | cathemeral |
| lizards | Acanthosaura_crucigera | Acanthosaura_crucigera | diurnal |

| | | | |
|---|---|---|---|
| lizards | Acanthosaura_lepidogaster | Acanthosaura_lepidogaster | diurnal |
| lizards | Acanthosaura_nataliae | Acanthosaura_nataliae | diurnal |
| lizards | Acontias_gariepensis | Acontias_gariepensis | nocturnal |
| lizards | Acontias_gracilicauda | Acontias_gracilicauda | cathemeral |
| lizards | Acontias_jappi | Acontias_jappi | cathemeral |
| lizards | Acontias_kgalagadi | Acontias_kgalagadi | nocturnal |
| lizards | Acontias_meleagris | Acontias_meleagris | cathemeral |
| lizards | Acontias_orientalis | Acontias_orientalis | nocturnal |
| lizards | Acontias_percivali | Acontias_percivali | crepuscular |
| lizards | Acontias_plumbeus | Acontias_plumbeus | nocturnal |
| lizards | Acratosaura_mentalis | Acratosaura_mentalis | diurnal |
| lizards | Acritoscincus_duperreyi | Bassiana_duperreyi | diurnal |
| lizards | Acritoscincus_platynotus | Bassiana_platynota | diurnal |
| lizards | Acritoscincus_trilineatus | Bassiana_trilineata | diurnal |
| lizards | Adolfus_alleni | Adolfus_alleni | diurnal |
| lizards | Adolfus_jacksoni | Adolfus_jacksoni | diurnal |
| lizards | Aeluroscalabotes_felinus | Aeluroscalabotes_felinus | nocturnal |
| lizards | Afroedura_africana | Afroedura_africana | diurnal |
| lizards | Afroedura_bogerti | Afroedura_bogerti | nocturnal |
| lizards | Afroedura_broadleyi | Afroedura_broadleyi | nocturnal |
| lizards | Afroedura_halli | Afroedura_halli | nocturnal |
| lizards | Afroedura_hawequensis | Afroedura_hawequensis | diurnal |
| lizards | Afroedura_karroica | Afroedura_karroica | nocturnal |
| lizards | Afroedura_major | Afroedura_major | diurnal |
| lizards | Afroedura_pienaari | Afroedura_pienaari | nocturnal |
| lizards | Afroedura_pondolia | Afroedura_pondolia | diurnal |
| lizards | Afrogecko_porphyreus | Afrogecko_porphyreus | nocturnal |
| lizards | Agama_aculeata | Agama_aculeata | diurnal |
| lizards | Agama_africana | Agama_africana | diurnal |
| lizards | Agama_agama | Agama_agama | diurnal |
| lizards | Agama_anchietae | Agama_anchietae | diurnal |
| lizards | Agama_armata | Agama_armata | diurnal |
| lizards | Agama_atra | Agama_atra | diurnal |
| lizards | Agama_boensis | Agama_boensis | diurnal |

| | | | |
|---|---|---|---|
| lizards | Agama_boueti | Agama_boueti | diurnal |
| lizards | Agama_caudospinosa | Agama_caudospinosa | diurnal |
| lizards | Agama_doriae | Agama_doriae | diurnal |
| lizards | Agama_etoshae | Agama_etoshae | diurnal |
| lizards | Agama_hispida | Agama_hispida | diurnal |
| lizards | Agama_kaimosae | Agama_kaimosae | diurnal |
| lizards | Agama_kirkii | Agama_kirkii | diurnal |
| lizards | Agama_knobeli | Agama_knobeli | diurnal |
| lizards | Agama_lionotus | Agama_lionotus | diurnal |
| lizards | Agama_montana | Agama_montana | diurnal |
| lizards | Agama_mossambica | Agama_mossambica | diurnal |
| lizards | Agama_mwanzae | Agama_mwanzae | diurnal |
| lizards | Agama_planiceps | Agama_planiceps | diurnal |
| lizards | Agama_spinosa | Agama_spinosa | diurnal |
| lizards | Agama_weidholzi | Agama_weidholzi | diurnal |
| lizards | Agamura_persica | Agamura_persica | nocturnal |
| lizards | Ailuronyx_seychellensis | Ailuronyx_seychellensis | nocturnal |
| lizards | Ailuronyx_tachyscopaeus | Ailuronyx_tachyscopaeus | nocturnal |
| lizards | Algyroides_fitzingeri | Algyroides_fitzingeri | diurnal |
| lizards | Algyroides_moreoticus | Algyroides_moreoticus | diurnal |
| lizards | Algyroides_nigropunctatus | Algyroides_nigropunctatus | diurnal |
| lizards | Alopoglossus_angulatus | Alopoglossus_angulatus | diurnal |
| lizards | Alopoglossus_atriventris | Alopoglossus_atriventris | diurnal |
| lizards | Alopoglossus_buckleyi | Alopoglossus_buckleyi | diurnal |
| lizards | Alopoglossus_copii | Alopoglossus_copii | diurnal |
| lizards | Alsophylax_laevis | Alsophylax_laevis | nocturnal |
| lizards | Alsophylax_loricatus | Alsophylax_loricatus | nocturnal |
| lizards | Alsophylax_pipiens | Alsophylax_pipiens | nocturnal |
| lizards | Alsophylax_przewalskii | Alsophylax_przewalskii | nocturnal |
| lizards | Alsophylax_szczerbaki | Alsophylax_szczerbaki | cathemeral |
| lizards | Alsophylax_tadjikiensis | Alsophylax_tadjikiensis | nocturnal |
| lizards | Altiphylax_baturensis | Altiphylax_baturensis | nocturnal |
| lizards | Altiphylax_levitoni | Altiphylax_levitoni | nocturnal |
| lizards | Altiphylax_tokobajevi | Altiphylax_tokobajevi | nocturnal |

| | | | |
|---|---|---|---|
| lizards | Amalosia_jacovae | Amalosia_jacovae | nocturnal |
| lizards | Amalosia_lesueurii | Amalosia_lesueurii | nocturnal |
| lizards | Amalosia_obscura | Amalosia_obscura | nocturnal |
| lizards | Amalosia_rhombifer | Amalosia_rhombifer | nocturnal |
| lizards | Amapasaurus_tetradactylus | Amapasaurus_tetradactylus | diurnal |
| lizards | Amblyrhynchus_cristatus | Amblyrhynchus_cristatus | diurnal |
| lizards | Ameiva_ameiva | Ameiva_ameiva | diurnal |
| lizards | Ameiva_atrigularis | Ameiva_atrigularis | diurnal |
| lizards | Ameiva_bifrontata | Ameiva_bifrontata | diurnal |
| lizards | Ameiva_concolor | Ameiva_concolor | diurnal |
| lizards | Ameiva_parecis | Ameiva_parecis | diurnal |
| lizards | Ameiva_praesignis | Ameiva_praesignis | diurnal |
| lizards | Ameivula_jalapensis | Ameivula_jalapensis | diurnal |
| lizards | Ameivula_mumbuca | Ameivula_mumbuca | diurnal |
| lizards | Ameivula_ocellifera | Ameivula_ocellifera | diurnal |
| lizards | Amphibolurus_burnsi | Lophognathus_burnsi | diurnal |
| lizards | Amphibolurus_muricatus | Amphibolurus_muricatus | diurnal |
| lizards | Amphibolurus_norrisi | Amphibolurus_norrisi | diurnal |
| lizards | Amphiglossus_astrolabi | Amphiglossus_astrolabi | diurnal |
| lizards | Amphiglossus_reticulatus | Amphiglossus_reticulatus | cathemeral |
| lizards | Amphisbaena_alba | Amphisbaena_alba | cathemeral |
| lizards | Amphisbaena_dubia | Amphisbaena_dubia | nocturnal |
| lizards | Anadia_brevifrontalis | Anadia_brevifrontalis | diurnal |
| lizards | Anadia_hobarti | Anadia_hobarti | diurnal |
| lizards | Anadia_rhombifera | Anadia_rhombifera | diurnal |
| lizards | Anatololacerta_anatolica | Anatololacerta_anatolica | diurnal |
| lizards | Anatololacerta_danfordi | Anatololacerta_danfordi | diurnal |
| lizards | Anatololacerta_pelasgiana | Anatololacerta_pelasgiana | diurnal |
| lizards | Andinosaura_oculata | Riama_oculata | diurnal |
| lizards | Andinosaura_vespertina | Riama_vespertina | diurnal |
| lizards | Anepischetosia_maccoyi | Anepischetosia_maccoyi | cathemeral |
| lizards | Anguis_cephallonica | Anguis_cephallonica | diurnal |
| lizards | Anguis_colchica | Anguis_colchica | diurnal |
| lizards | Anguis_fragilis | Anguis_fragilis | diurnal |

| | | | |
|---|---|---|---|
| lizards | Anguis_graeca | Anguis_graeca | diurnal |
| lizards | Anniella_geronimensis | Anniella_geronimensis | diurnal |
| lizards | Anniella_pulchra | Anniella_pulchra | cathemeral |
| lizards | Anolis_acutus | Anolis_acutus | diurnal |
| lizards | Anolis_aeneus | Anolis_aeneus | diurnal |
| lizards | Anolis_aequatorialis | Anolis_aequatorialis | diurnal |
| lizards | Anolis_agassizi | Anolis_agassizi | diurnal |
| lizards | Anolis_aliniger | Anolis_aliniger | diurnal |
| lizards | Anolis_allisoni | Anolis_allisoni | diurnal |
| lizards | Anolis_allogus | Anolis_allogus | diurnal |
| lizards | Anolis_altitudinalis | Anolis_altitudinalis | diurnal |
| lizards | Anolis_alumina | Anolis_alumina | diurnal |
| lizards | Anolis_alutaceus | Anolis_alutaceus | diurnal |
| lizards | Anolis_angusticeps | Anolis_angusticeps | diurnal |
| lizards | Anolis_anisolepis | Anolis_anisolepis | diurnal |
| lizards | Anolis_aquaticus | Anolis_aquaticus | diurnal |
| lizards | Anolis_argenteolus | Anolis_argenteolus | diurnal |
| lizards | Anolis_armouri | Anolis_armouri | diurnal |
| lizards | Anolis_auratus | Anolis_auratus | diurnal |
| lizards | Anolis_bahorucoensis | Anolis_bahorucoensis | diurnal |
| lizards | Anolis_baleatus | Anolis_baleatus | diurnal |
| lizards | Anolis_baracoae | Anolis_baracoae | diurnal |
| lizards | Anolis_barahonae | Anolis_barahonae | diurnal |
| lizards | Anolis_barbatus | Anolis_barbatus | diurnal |
| lizards | Anolis_barbouri | Anolis_barbouri | diurnal |
| lizards | Anolis_barkeri | Anolis_barkeri | diurnal |
| lizards | Anolis_bartschi | Anolis_bartschi | diurnal |
| lizards | Anolis_beckeri | Anolis_beckeri | diurnal |
| lizards | Anolis_bicaorum | Anolis_bicaorum | diurnal |
| lizards | Anolis_bimaculatus | Anolis_bimaculatus | diurnal |
| lizards | Anolis_biporcatus | Anolis_biporcatus | diurnal |
| lizards | Anolis_bitectus | Anolis_bitectus | diurnal |
| lizards | Anolis_blanquillanus | Anolis_blanquillanus | diurnal |
| lizards | Anolis_bombiceps | Anolis_bombiceps | diurnal |

| | | | |
|---|---|---|---|
| lizards | Anolis_bonairensis | Anolis_bonairensis | diurnal |
| lizards | Anolis_boulengerianus | Anolis_boulengerianus | diurnal |
| lizards | Anolis_brasiliensis | Anolis_brasiliensis | diurnal |
| lizards | Anolis_bremeri | Anolis_bremeri | diurnal |
| lizards | Anolis_brevirostris | Anolis_brevirostris | diurnal |
| lizards | Anolis_capito | Anolis_capito | diurnal |
| lizards | Anolis_carlliebi | Anolis_carlliebi | diurnal |
| lizards | Anolis_carolinensis | Anolis_carolinensis | diurnal |
| lizards | Anolis_carpenteri | Anolis_carpenteri | diurnal |
| lizards | Anolis_centralis | Anolis_centralis | diurnal |
| lizards | Anolis_chamaeleonides | Anolis_chamaeleonides | diurnal |
| lizards | Anolis_chloris | Anolis_chloris | diurnal |
| lizards | Anolis_chlorocyanus | Anolis_chlorocyanus | diurnal |
| lizards | Anolis_christophei | Anolis_christophei | diurnal |
| lizards | Anolis_chrysolepis | Anolis_chrysolepis | diurnal |
| lizards | Anolis_chrysops | Anolis_chrysops | diurnal |
| lizards | Anolis_clivicola | Anolis_clivicola | diurnal |
| lizards | Anolis_concolor | Anolis_concolor | diurnal |
| lizards | Anolis_conspersus | Anolis_conspersus | diurnal |
| lizards | Anolis_cooki | Anolis_cooki | diurnal |
| lizards | Anolis_crassulus | Anolis_crassulus | diurnal |
| lizards | Anolis_cristatellus | Anolis_cristatellus | diurnal |
| lizards | Anolis_cupreus | Anolis_cupreus | diurnal |
| lizards | Anolis_cuprinus | Anolis_cuprinus | diurnal |
| lizards | Anolis_cuvieri | Anolis_cuvieri | diurnal |
| lizards | Anolis_cybotes | Anolis_cybotes | diurnal |
| lizards | Anolis_desiradei | Anolis_desiradei | diurnal |
| lizards | Anolis_distichus | Anolis_distichus | diurnal |
| lizards | Anolis_dollfusianus | Anolis_dollfusianus | diurnal |
| lizards | Anolis_duellmani | Anolis_duellmani | diurnal |
| lizards | Anolis_dunni | Anolis_dunni | diurnal |
| lizards | Anolis_equestris | Anolis_equestris | diurnal |
| lizards | Anolis_ernestwilliamsi | Anolis_ernestwilliamsi | diurnal |
| lizards | Anolis_etheridgei | Anolis_etheridgei | diurnal |

| | | | |
|---|---|---|---|
| lizards | Anolis_eugenegrahami | Anolis_eugenegrahami | diurnal |
| lizards | Anolis_evermanni | Anolis_evermanni | diurnal |
| lizards | Anolis_extremus | Anolis_extremus | diurnal |
| lizards | Anolis_fasciatus | Anolis_fasciatus | diurnal |
| lizards | Anolis_festae | Anolis_festae | diurnal |
| lizards | Anolis_fowleri | Anolis_fowleri | diurnal |
| lizards | Anolis_frenatus | Anolis_frenatus | diurnal |
| lizards | Anolis_fuscoauratus | Anolis_fuscoauratus | diurnal |
| lizards | Anolis_gadovii | Anolis_gadovii | diurnal |
| lizards | Anolis_gaigei | Anolis_gaigei | diurnal |
| lizards | Anolis_garmani | Anolis_garmani | diurnal |
| lizards | Anolis_gemmosus | Anolis_gemmosus | diurnal |
| lizards | Anolis_gingivinus | Anolis_gingivinus | diurnal |
| lizards | Anolis_grahami | Anolis_grahami | diurnal |
| lizards | Anolis_griseus | Anolis_griseus | diurnal |
| lizards | Anolis_gundlachi | Anolis_gundlachi | diurnal |
| lizards | Anolis_heterodermus | Anolis_heterodermus | diurnal |
| lizards | Anolis_homolechis | Anolis_homolechis | diurnal |
| lizards | Anolis_humilis | Anolis_humilis | diurnal |
| lizards | Anolis_insolitus | Anolis_insolitus | diurnal |
| lizards | Anolis_isolepis | Anolis_isolepis | diurnal |
| lizards | Anolis_jacare | Anolis_jacare | diurnal |
| lizards | Anolis_johnmeyeri | Anolis_johnmeyeri | diurnal |
| lizards | Anolis_jubar | Anolis_jubar | diurnal |
| lizards | Anolis_kahouannensis | Anolis_kahouannensis | diurnal |
| lizards | Anolis_kemptoni | Anolis_kemptoni | diurnal |
| lizards | Anolis_krugi | Anolis_krugi | diurnal |
| lizards | Anolis_laeviventris | Anolis_laeviventris | diurnal |
| lizards | Anolis_leachii | Anolis_leachii | diurnal |
| lizards | Anolis_lemurinus | Anolis_lemurinus | diurnal |
| lizards | Anolis_limifrons | Anolis_limifrons | diurnal |
| lizards | Anolis_lineatopus | Anolis_lineatopus | diurnal |
| lizards | Anolis_liogaster | Anolis_liogaster | diurnal |
| lizards | Anolis_lionotus | Anolis_lionotus | diurnal |

| | | | |
|---|---|---|---|
| lizards | Anolis_litoralis | Anolis_litoralis | diurnal |
| lizards | Anolis_lividus | Anolis_lividus | diurnal |
| lizards | Anolis_longiceps | Anolis_longiceps | diurnal |
| lizards | Anolis_longitibialis | Anolis_longitibialis | diurnal |
| lizards | Anolis_loveridgei | Anolis_loveridgei | diurnal |
| lizards | Anolis_loysiana | Anolis_loysiana | diurnal |
| lizards | Anolis_luciae | Anolis_luciae | diurnal |
| lizards | Anolis_lucius | Anolis_lucius | diurnal |
| lizards | Anolis_luteogularis | Anolis_luteogularis | diurnal |
| lizards | Anolis_lyra | Anolis_lyra | diurnal |
| lizards | Anolis_macrinii | Anolis_macrinii | diurnal |
| lizards | Anolis_maculiventris | Anolis_maculiventris | diurnal |
| lizards | Anolis_mariarum | Anolis_mariarum | diurnal |
| lizards | Anolis_marmoratus | Anolis_marmoratus | diurnal |
| lizards | Anolis_marron | Anolis_marron | diurnal |
| lizards | Anolis_matudai | Anolis_matudai | cathemeral |
| lizards | Anolis_megapholidotus | Anolis_megapholidotus | diurnal |
| lizards | Anolis_menta | Anolis_menta | diurnal |
| lizards | Anolis_meridionalis | Anolis_meridionalis | diurnal |
| lizards | Anolis_mestrei | Anolis_mestrei | diurnal |
| lizards | Anolis_microlepidotus | Anolis_microlepidotus | diurnal |
| lizards | Anolis_milleri | Anolis_milleri | diurnal |
| lizards | Anolis_naufragus | Anolis_naufragus | diurnal |
| lizards | Anolis_nebuloides | Anolis_nebuloides | diurnal |
| lizards | Anolis_nebulosus | Anolis_nebulosus | diurnal |
| lizards | Anolis_nelsoni | Anolis_nelsoni | diurnal |
| lizards | Anolis_nigrolineatus | Anolis_nigrolineatus | diurnal |
| lizards | Anolis_noblei | Anolis_noblei | diurnal |
| lizards | Anolis_oculatus | Anolis_oculatus | diurnal |
| lizards | Anolis_olssoni | Anolis_olssoni | diurnal |
| lizards | Anolis_omiltemanus | Anolis_omiltemanus | diurnal |
| lizards | Anolis_onca | Anolis_onca | diurnal |
| lizards | Anolis_opalinus | Anolis_opalinus | diurnal |
| lizards | Anolis_ophiolepis | Anolis_ophiolepis | diurnal |

| | | | |
|---|---|---|---|
| lizards | Anolis_ortonii | Anolis_ortonii | diurnal |
| lizards | Anolis_otongae | Anolis_otongae | diurnal |
| lizards | Anolis_oxylophus | Anolis_oxylophus | diurnal |
| lizards | Anolis_pachypus | Anolis_pachypus | diurnal |
| lizards | Anolis_parvicirculatus | Anolis_parvicirculatus | diurnal |
| lizards | Anolis_pentaprion | Anolis_pentaprion | diurnal |
| lizards | Anolis_peraccae | Anolis_peraccae | diurnal |
| lizards | Anolis_petersii | Anolis_petersii | diurnal |
| lizards | Anolis_pigmaequestris | Anolis_pigmaequestris | diurnal |
| lizards | Anolis_placidus | Anolis_placidus | diurnal |
| lizards | Anolis_planiceps | Anolis_planiceps | diurnal |
| lizards | Anolis_podocarpus | Anolis_podocarpus | diurnal |
| lizards | Anolis_poecilopus | Anolis_poecilopus | diurnal |
| lizards | Anolis_pogus | Anolis_pogus | diurnal |
| lizards | Anolis_polylepis | Anolis_polylepis | diurnal |
| lizards | Anolis_porcatus | Anolis_porcatus | diurnal |
| lizards | Anolis_porcus | Anolis_porcus | diurnal |
| lizards | Anolis_proboscis | Anolis_proboscis | diurnal |
| lizards | Anolis_pulchellus | Anolis_pulchellus | diurnal |
| lizards | Anolis_punctatus | Anolis_punctatus | diurnal |
| lizards | Anolis_quaggulus | Anolis_quaggulus | diurnal |
| lizards | Anolis_reconditus | Anolis_reconditus | diurnal |
| lizards | Anolis_rejectus | Anolis_rejectus | diurnal |
| lizards | Anolis_richardii | Anolis_richardii | diurnal |
| lizards | Anolis_rodriguezii | Anolis_rodriguezi | diurnal |
| lizards | Anolis_roquet | Anolis_roquet | diurnal |
| lizards | Anolis_rubribarbaris | Anolis_rubribarbaris | diurnal |
| lizards | Anolis_sabanus | Anolis_sabanus | diurnal |
| lizards | Anolis_sagrei | Anolis_sagrei | diurnal |
| lizards | Anolis_saxatilis | Anolis_whitemani | diurnal |
| lizards | Anolis_schwartzi | Anolis_schwartzi | diurnal |
| lizards | Anolis_scypheus | Anolis_scypheus | diurnal |
| lizards | Anolis_semilineatus | Anolis_semilineatus | diurnal |
| lizards | Anolis_sericeus | Anolis_sericeus | diurnal |

| | | | |
|---|---|---|---|
| lizards | Anolis_sheplani | Anolis_sheplani | diurnal |
| lizards | Anolis_shrevei | Anolis_shrevei | diurnal |
| lizards | Anolis_singularis | Anolis_singularis | diurnal |
| lizards | Anolis_smallwoodi | Anolis_smallwoodi | diurnal |
| lizards | Anolis_smaragdinus | Anolis_smaragdinus | diurnal |
| lizards | Anolis_sminthus | Anolis_sminthus | diurnal |
| lizards | Anolis_spectrum | Anolis_spectrum | diurnal |
| lizards | Anolis_strahmi | Anolis_strahmi | diurnal |
| lizards | Anolis_subocularis | Anolis_subocularis | diurnal |
| lizards | Anolis_tandai | Anolis_tandai | diurnal |
| lizards | Anolis_taylori | Anolis_taylori | diurnal |
| lizards | Anolis_terraealtae | Anolis_terraealtae | diurnal |
| lizards | Anolis_tolimensis | Anolis_tolimensis | diurnal |
| lizards | Anolis_trachyderma | Anolis_trachyderma | diurnal |
| lizards | Anolis_transversalis | Anolis_transversalis | diurnal |
| lizards | Anolis_trinitatis | Anolis_trinitatis | diurnal |
| lizards | Anolis_tropidogaster | Anolis_tropidogaster | diurnal |
| lizards | Anolis_tropidolepis | Anolis_tropidolepis | diurnal |
| lizards | Anolis_tropidonotus | Anolis_tropidonotus | diurnal |
| lizards | Anolis_uniformis | Anolis_uniformis | diurnal |
| lizards | Anolis_utilensis | Anolis_utilensis | diurnal |
| lizards | Anolis_valencienni | Anolis_valencienni | diurnal |
| lizards | Anolis_vermiculatus | Anolis_vermiculatus | diurnal |
| lizards | Anolis_vittigerus | Anolis_vittigerus | diurnal |
| lizards | Anolis_wattsii | Anolis_wattsi | diurnal |
| lizards | Anolis_woodi | Anolis_woodi | diurnal |
| lizards | Anomalopus_leuckartii | Anomalopus_leuckartii | cathemeral |
| lizards | Anomalopus_swansoni | Anomalopus_swansoni | diurnal |
| lizards | Anomalopus_verreauxii | Anomalopus_verreauxi | cathemeral |
| lizards | Anotosaura_vanzolinia | Anotosaura_vanzolinia | diurnal |
| lizards | Apathya_cappadocica | Apathya_cappadocica | diurnal |
| lizards | Aphaniotis_acutirostris | Aphaniotis_acutirostris | diurnal |
| lizards | Aphaniotis_fusca | Aphaniotis_fusca | diurnal |
| lizards | Aphaniotis_ornata | Aphaniotis_ornata | diurnal |

| | | | |
|---|---|---|---|
| lizards | Aprasia_aurita | Aprasia_aurita | cathemeral |
| lizards | Aprasia_haroldi | Aprasia_haroldi | nocturnal |
| lizards | Aprasia_inaurita | Aprasia_inaurita | cathemeral |
| lizards | Aprasia_parapulchella | Aprasia_parapulchella | nocturnal |
| lizards | Aprasia_pseudopulchella | Aprasia_pseudopulchella | nocturnal |
| lizards | Aprasia_pulchella | Aprasia_pulchella | cathemeral |
| lizards | Aprasia_repens | Aprasia_repens | nocturnal |
| lizards | Aprasia_rostrata | Aprasia_rostrata | nocturnal |
| lizards | Aprasia_striolata | Aprasia_striolata | cathemeral |
| lizards | Archaeolacerta_bedriagae | Archaeolacerta_bedriagae | diurnal |
| lizards | Archaius_tigris | Archaius_tigris | diurnal |
| lizards | Aristelliger_barbouri | Aristelliger_barbouri | nocturnal |
| lizards | Aristelliger_cochranae | Aristelliger_cochranae | cathemeral |
| lizards | Aristelliger_expectatus | Aristelliger_expectatus | nocturnal |
| lizards | Aristelliger_georgeensis | Aristelliger_georgeensis | cathemeral |
| lizards | Aristelliger_hechti | Aristelliger_hechti | nocturnal |
| lizards | Aristelliger_lar | Aristelliger_lar | nocturnal |
| lizards | Aristelliger_praesignis | Aristelliger_praesignis | cathemeral |
| lizards | Aristelliger_reyesi | Aristelliger_reyesi | cathemeral |
| lizards | Arthrosaura_kockii | Arthrosaura_kockii | diurnal |
| lizards | Arthrosaura_reticulata | Arthrosaura_reticulata | diurnal |
| lizards | Asaccus_andersoni | Asaccus_andersoni | cathemeral |
| lizards | Asaccus_caudivolvulus | Asaccus_caudivolvulus | nocturnal |
| lizards | Asaccus_elisae | Asaccus_elisae | nocturnal |
| lizards | Asaccus_gallagheri | Asaccus_gallagheri | nocturnal |
| lizards | Asaccus_nasrullahi | Asaccus_nasrullahi | cathemeral |
| lizards | Asaccus_platyrhynchus | Asaccus_platyrhynchus | nocturnal |
| lizards | Asaccus_tangestanensis | Asaccus_tangestanensis | cathemeral |
| lizards | Asaccus_zagrosicus | Asaccus_zagrosicus | cathemeral |
| lizards | Aspidoscelis_burti | Aspidoscelis_burti | diurnal |
| lizards | Aspidoscelis_calidipes | Aspidoscelis_calidipes | diurnal |
| lizards | Aspidoscelis_ceralbensis | Aspidoscelis_ceralbensis | diurnal |
| lizards | Aspidoscelis_communis | Aspidoscelis_communis | diurnal |
| lizards | Aspidoscelis_costatus | Aspidoscelis_costata | diurnal |

| | | | |
|---|---|---|---|
| lizards | Aspidoscelis_cozumela | Aspidoscelis_cozumelae | diurnal |
| lizards | Aspidoscelis_danheimae | Aspidoscelis_danheimae | diurnal |
| lizards | Aspidoscelis_deppii | Aspidoscelis_deppei | diurnal |
| lizards | Aspidoscelis_exsanguis | Aspidoscelis_exsanguis | diurnal |
| lizards | Aspidoscelis_gularis | Aspidoscelis_gularis | diurnal |
| lizards | Aspidoscelis_guttatus | Aspidoscelis_guttata | diurnal |
| lizards | Aspidoscelis_hyperythrus | Aspidoscelis_hyperythra | diurnal |
| lizards | Aspidoscelis_inornatus | Aspidoscelis_inornata | diurnal |
| lizards | Aspidoscelis_labialis | Aspidoscelis_labialis | diurnal |
| lizards | Aspidoscelis_laredoensis | Aspidoscelis_laredoensis | diurnal |
| lizards | Aspidoscelis_lineattissimus | Aspidoscelis_lineattissima | diurnal |
| lizards | Aspidoscelis_marmoratus | Aspidoscelis_marmorata | diurnal |
| lizards | Aspidoscelis_martyris | Cnemidophorus_martyris | diurnal |
| lizards | Aspidoscelis_maximus | Aspidoscelis_maxima | diurnal |
| lizards | Aspidoscelis_motaguae | Aspidoscelis_motaguae | diurnal |
| lizards | Aspidoscelis_neomexicanus | Aspidoscelis_neomexicana | diurnal |
| lizards | Aspidoscelis_neotesselatus | Aspidoscelis_neotesselata | diurnal |
| lizards | Aspidoscelis_opatae | Aspidoscelis_opatae | diurnal |
| lizards | Aspidoscelis_pai | Aspidoscelis_pai | diurnal |
| lizards | Aspidoscelis_parvisocius | Aspidoscelis_parvisocia | diurnal |
| lizards | Aspidoscelis_pictus | Aspidoscelis_picta | diurnal |
| lizards | Aspidoscelis_rodecki | Aspidoscelis_rodecki | diurnal |
| lizards | Aspidoscelis_sackii | Aspidoscelis_sackii | diurnal |
| lizards | Aspidoscelis_scalaris | Aspidoscelis_scalaris | diurnal |
| lizards | Aspidoscelis_sexlineatus | Aspidoscelis_sexlineata | diurnal |
| lizards | Aspidoscelis_sonorae | Aspidoscelis_sonorae | diurnal |
| lizards | Aspidoscelis_stictogrammus | Aspidoscelis_stictogramma | diurnal |
| lizards | Aspidoscelis_tesselatus | Aspidoscelis_tesselata | diurnal |
| lizards | Aspidoscelis_tigris | Aspidoscelis_tigris | diurnal |
| lizards | Aspidoscelis_uniparens | Aspidoscelis_uniparens | diurnal |
| lizards | Aspidoscelis_velox | Aspidoscelis_velox | diurnal |
| lizards | Aspidoscelis_xanthonotus | Aspidoscelis_xanthonota | diurnal |
| lizards | Aspronema_cochabambae | Aspronema_cochabambae | diurnal |
| lizards | Aspronema_dorsivittatum | Aspronema_dorsivittatum | diurnal |

| | | | |
|---|---|---|---|
| lizards | Ateuchosaurus_chinensis | Ateuchosaurus_chinensis | cathemeral |
| lizards | Ateuchosaurus_pellopleurus | Ateuchosaurus_pellopleurus | cathemeral |
| lizards | Atlantolacerta_andreanskyi | Atlantolacerta_andreanskyi | diurnal |
| lizards | Australolacerta_australis | Australolacerta_australis | diurnal |
| lizards | Bachia_bicolor | Bachia_bicolor | diurnal |
| lizards | Bachia_bresslaui | Bachia_bresslaui | diurnal |
| lizards | Bachia_flavescens | Bachia_flavescens | diurnal |
| lizards | Bachia_heteropa | Bachia_heteropa | diurnal |
| lizards | Bachia_scaea | Bachia_scaea | diurnal |
| lizards | Bachia_trisanale | Bachia_trisanale | diurnal |
| lizards | Barisia_ciliaris | Barisia_ciliaris | diurnal |
| lizards | Barisia_imbricata | Barisia_imbricata | diurnal |
| lizards | Basiliscus_basiliscus | Basiliscus_basiliscus | diurnal |
| lizards | Basiliscus_galeritus | Basiliscus_galeritus | diurnal |
| lizards | Basiliscus_plumifrons | Basiliscus_plumifrons | diurnal |
| lizards | Basiliscus_vittatus | Basiliscus_vittatus | diurnal |
| lizards | Bavayia_crassicollis | Bavayia_crassicollis | nocturnal |
| lizards | Bavayia_cyclura | Bavayia_cyclura | nocturnal |
| lizards | Bavayia_exsuccida | Bavayia_exsuccida | nocturnal |
| lizards | Bavayia_geitaina | Bavayia_geitaina | nocturnal |
| lizards | Bavayia_goroensis | Bavayia_goroensis | nocturnal |
| lizards | Bavayia_montana | Bavayia_montana | nocturnal |
| lizards | Bavayia_nubila | Bavayia_nubila | nocturnal |
| lizards | Bavayia_ornata | Bavayia_ornata | nocturnal |
| lizards | Bavayia_pulchella | Bavayia_pulchella | nocturnal |
| lizards | Bavayia_robusta | Bavayia_robusta | nocturnal |
| lizards | Bavayia_sauvagii | Bavayia_sauvagii | nocturnal |
| lizards | Bavayia_septuiclavis | Bavayia_septuiclavis | nocturnal |
| lizards | Bellatorias_frerei | Bellatorias_frerei | cathemeral |
| lizards | Bellatorias_major | Bellatorias_major | diurnal |
| lizards | Bipes_biporus | Bipes_biporus | diurnal |
| lizards | Bipes_canaliculatus | Bipes_canaliculatus | diurnal |
| lizards | Blaesodactylus_ambonihazo | Blaesodactylus_ambonihazo | nocturnal |
| lizards | Blaesodactylus_antongilensis | Blaesodactylus_antongilensis | nocturnal |

| | | | |
|---|---|---|---|
| lizards | Blaesodactylus_boivini | Blaesodactylus_boivini | cathemeral |
| lizards | Blanus_cinereus | Blanus_cinereus | cathemeral |
| lizards | Blanus_mettetali | Blanus_mettetali | nocturnal |
| lizards | Brachylophus_bulabula | Brachylophus_bulabula | diurnal |
| lizards | Brachylophus_fasciatus | Brachylophus_fasciatus | diurnal |
| lizards | Brachylophus_vitiensis | Brachylophus_vitiensis | diurnal |
| lizards | Brachymeles_apus | Brachymeles_apus | cathemeral |
| lizards | Brachymeles_bonitae | Brachymeles_bonitae | diurnal |
| lizards | Brachymeles_boulengeri | Brachymeles_boulengeri | diurnal |
| lizards | Brachymeles_gracilis | Brachymeles_gracilis | diurnal |
| lizards | Brachymeles_orientalis | Brachymeles_orientalis | diurnal |
| lizards | Brachymeles_samarensis | Brachymeles_samarensis | diurnal |
| lizards | Brachymeles_schadenbergi | Brachymeles_schadenbergi | diurnal |
| lizards | Brachymeles_talinis | Brachymeles_talinis | nocturnal |
| lizards | Brachymeles_taylori | Brachymeles_taylori | nocturnal |
| lizards | Brachymeles_tridactylus | Brachymeles_tridactylus | nocturnal |
| lizards | Brachyseps_frontoparietalis | Amphiglossus_frontoparietalis | nocturnal |
| lizards | Brachyseps_gastrostictus | Amphiglossus_macrocercus | diurnal |
| lizards | Brachyseps_punctatus | Amphiglossus_punctatus | diurnal |
| lizards | Bradypodion_damaranum | Bradypodion_damaranum | diurnal |
| lizards | Bradypodion_melanocephalum | Bradypodion_melanocephalum | diurnal |
| lizards | Bradypodion_occidentale | Bradypodion_occidentale | diurnal |
| lizards | Bradypodion_pumilum | Bradypodion_pumilum | diurnal |
| lizards | Bradypodion_setaroi | Bradypodion_setaroi | diurnal |
| lizards | Bradypodion_taeniabronchum | Bradypodion_taeniabronchum | diurnal |
| lizards | Bradypodion_thamnobates | Bradypodion_thamnobates | diurnal |
| lizards | Bradypodion_transvaalense | Bradypodion_transvaalense | diurnal |
| lizards | Bradypodion_ventrale | Bradypodion_ventrale | diurnal |
| lizards | Brasiliscincus_agilis | Brasiliscincus_agilis | diurnal |
| lizards | Brasiliscincus_caissara | Brasiliscincus_caissara | diurnal |
| lizards | Brasiliscincus_heathi | Brasiliscincus_heathi | diurnal |
| lizards | Broadleysaurus_major | Broadleysaurus_major | diurnal |
| lizards | Bronchocela_cristatella | Bronchocela_cristatella | diurnal |
| lizards | Bronchocela_hayeki | Bronchocela_hayeki | diurnal |

| | | | |
|---|---|---|---|
| lizards | Bronchocela_jubata | Bronchocela_jubata | diurnal |
| lizards | Bronchocela_marmorata | Bronchocela_marmorata | diurnal |
| lizards | Bronchocela_smaragdina | Bronchocela_smaragdina | diurnal |
| lizards | Brookesia_antakarana | Brookesia_antakarana | diurnal |
| lizards | Brookesia_brygooi | Brookesia_brygooi | diurnal |
| lizards | Brookesia_decaryi | Brookesia_decaryi | diurnal |
| lizards | Brookesia_desperata | Brookesia_desperata | diurnal |
| lizards | Brookesia_exarmata | Brookesia_exarmata | diurnal |
| lizards | Brookesia_griveaudi | Brookesia_griveaudi | diurnal |
| lizards | Brookesia_minima | Brookesia_minima | diurnal |
| lizards | Brookesia_stumpffi | Brookesia_stumpffi | diurnal |
| lizards | Brookesia_superciliaris | Brookesia_superciliaris | diurnal |
| lizards | Brookesia_thieli | Brookesia_thieli | diurnal |
| lizards | Brookesia_tristis | Brookesia_tristis | diurnal |
| lizards | Bunopus_blanfordii | Bunopus_blanfordii | cathemeral |
| lizards | Bunopus_tuberculatus | Bunopus_tuberculatus | cathemeral |
| lizards | Cachryx_defensor | Ctenosaura_defensor | diurnal |
| lizards | Caesoris_novaecaledoniae | Lioscincus_novaecaledoniae | diurnal |
| lizards | Caledoniscincus_aquilonius | Caledoniscincus_aquilonius | diurnal |
| lizards | Caledoniscincus_atropunctatus | Caledoniscincus_atropunctatus | diurnal |
| lizards | Caledoniscincus_auratus | Caledoniscincus_auratus | diurnal |
| lizards | Caledoniscincus_austrocaledonicus | Caledoniscincus_austrocaledonicus | diurnal |
| lizards | Caledoniscincus_chazeaui | Caledoniscincus_chazeaui | diurnal |
| lizards | Caledoniscincus_festivus | Caledoniscincus_festivus | diurnal |
| lizards | Caledoniscincus_orestes | Caledoniscincus_orestes | diurnal |
| lizards | Caledoniscincus_pelletieri | Caledoniscincus_pelletieri | diurnal |
| lizards | Caledoniscincus_renevieri | Caledoniscincus_renevieri | diurnal |
| lizards | Caledoniscincus_terma | Caledoniscincus_terma | diurnal |
| lizards | Callisaurus_draconoides | Callisaurus_draconoides | diurnal |
| lizards | Callopistes_maculatus | Callopistes_maculatus | diurnal |
| lizards | Calodactylodes_aureus | Calodactylodes_aureus | cathemeral |
| lizards | Calotes_bachae | Calotes_bachae | diurnal |
| lizards | Calotes_calotes | Calotes_calotes | diurnal |
| lizards | Calotes_ceylonensis | Calotes_ceylonensis | diurnal |

| | | | |
|---|---|---|---|
| lizards | Calotes_desilvai | Calotes_desilvai | diurnal |
| lizards | Calotes_emma | Calotes_emma | diurnal |
| lizards | Calotes_grandisquamis | Calotes_grandisquamis | diurnal |
| lizards | Calotes_jerdoni | Calotes_jerdoni | diurnal |
| lizards | Calotes_liocephalus | Calotes_liocephalus | diurnal |
| lizards | Calotes_liolepis | Calotes_liolepis | diurnal |
| lizards | Calotes_minor | Brachysaura_minor | diurnal |
| lizards | Calotes_mystaceus | Calotes_mystaceus | diurnal |
| lizards | Calotes_nemoricola | Calotes_nemoricola | diurnal |
| lizards | Calotes_nigrilabris | Calotes_nigrilabris | diurnal |
| lizards | Calotes_paulus | Oriocalotes_paulus | diurnal |
| lizards | Calotes_pethiyagodai | Calotes_pethiyagodai | diurnal |
| lizards | Calotes_versicolor | Calotes_versicolor | diurnal |
| lizards | Calumma_brevicorne | Calumma_brevicorne | diurnal |
| lizards | Calumma_gehringi | Calumma_guibei | diurnal |
| lizards | Calumma_hilleniusi | Calumma_hilleniusi | diurnal |
| lizards | Calumma_nasutum | Calumma_nasutum | diurnal |
| lizards | Calumma_parsonii | Calumma_parsonii | diurnal |
| lizards | Calyptommatus_leiolepis | Calyptommatus_leiolepis | nocturnal |
| lizards | Calyptommatus_sinebrachiatus | Calyptommatus_sinebrachiatus | nocturnal |
| lizards | Calyptotis_lepidorostrum | Calyptotis_lepidorostrum | cathemeral |
| lizards | Calyptotis_ruficauda | Calyptotis_ruficauda | cathemeral |
| lizards | Calyptotis_scutirostrum | Calyptotis_scutirostrum | cathemeral |
| lizards | Caparaonia_itaiquara | Caparaonia_itaiquara | diurnal |
| lizards | Carinascincus_coventryi | Niveoscincus_coventryi | diurnal |
| lizards | Carinascincus_greeni | Niveoscincus_greeni | diurnal |
| lizards | Carinascincus_metallicus | Niveoscincus_metallicus | diurnal |
| lizards | Carinascincus_microlepidotus | Niveoscincus_microlepidotus | diurnal |
| lizards | Carinascincus_ocellatus | Niveoscincus_ocellatus | diurnal |
| lizards | Carinascincus_orocryptus | Niveoscincus_orocryptus | diurnal |
| lizards | Carinascincus_palfreymani | Niveoscincus_palfreymani | diurnal |
| lizards | Carinascincus_pretiosus | Niveoscincus_pretiosus | diurnal |
| lizards | Carlia_ailanpalai | Carlia_ailanpalai | diurnal |
| lizards | Carlia_amax | Carlia_amax | diurnal |

| | | | |
|---|---|---|---|
| lizards | Carlia_bicarinata | Carlia_bicarinata | diurnal |
| lizards | Carlia_bomberai | Carlia_bomberai | diurnal |
| lizards | Carlia_caesius | Carlia_caesius | diurnal |
| lizards | Carlia_eothen | Carlia_eothen | diurnal |
| lizards | Carlia_gracilis | Carlia_gracilis | diurnal |
| lizards | Carlia_jarnoldae | Carlia_jarnoldae | diurnal |
| lizards | Carlia_longipes | Carlia_longipes | diurnal |
| lizards | Carlia_munda | Carlia_munda | diurnal |
| lizards | Carlia_mysi | Carlia_mysi | diurnal |
| lizards | Carlia_pectoralis | Carlia_pectoralis | diurnal |
| lizards | Carlia_peronii | Carlia_peronii | diurnal |
| lizards | Carlia_rhomboidalis | Carlia_rhomboidalis | diurnal |
| lizards | Carlia_rostralis | Carlia_rostralis | diurnal |
| lizards | Carlia_rubrigularis | Carlia_rubrigularis | diurnal |
| lizards | Carlia_rufilatus | Carlia_rufilatus | diurnal |
| lizards | Carlia_schmeltzii | Carlia_schmeltzii | diurnal |
| lizards | Carlia_storri | Carlia_storri | diurnal |
| lizards | Carlia_tetradactyla | Carlia_tetradactyla | diurnal |
| lizards | Carlia_triacantha | Carlia_triacantha | diurnal |
| lizards | Carlia_tutela | Carlia_tutela | diurnal |
| lizards | Carlia_vivax | Carlia_vivax | diurnal |
| lizards | Carphodactylus_laevis | Carphodactylus_laevis | nocturnal |
| lizards | Celatiscincus_euryotis | Celatiscincus_euryotis | diurnal |
| lizards | Celatiscincus_similis | Celatiscincus_similis | diurnal |
| lizards | Ceratophora_aspera | Ceratophora_aspera | diurnal |
| lizards | Ceratophora_erdeleni | Ceratophora_erdeleni | diurnal |
| lizards | Ceratophora_karu | Ceratophora_karu | diurnal |
| lizards | Ceratophora_stoddartii | Ceratophora_stoddartii | diurnal |
| lizards | Ceratophora_tennentii | Ceratophora_tennentii | diurnal |
| lizards | Cercosaura_argulus | Cercosaura_argulus | diurnal |
| lizards | Cercosaura_eigenmanni | Cercosaura_eigenmanni | diurnal |
| lizards | Cercosaura_manicata | Cercosaura_manicata | diurnal |
| lizards | Cercosaura_ocellata | Cercosaura_ocellata | diurnal |
| lizards | Cercosaura_parkeri | Cercosaura_parkeri | diurnal |

| | | | |
|---|---|---|---|
| lizards | Cercosaura_schreibersii | Cercosaura_schreibersii | diurnal |
| lizards | Chalcides_bedriagai | Chalcides_bedriagai | diurnal |
| lizards | Chalcides_chalcides | Chalcides_chalcides | cathemeral |
| lizards | Chalcides_coeruleopunctatus | Chalcides_coeruleopunctatus | diurnal |
| lizards | Chalcides_colosii | Chalcides_colosii | diurnal |
| lizards | Chalcides_guentheri | Chalcides_guentheri | diurnal |
| lizards | Chalcides_lanzai | Chalcides_lanzai | diurnal |
| lizards | Chalcides_manueli | Chalcides_manueli | diurnal |
| lizards | Chalcides_mertensi | Chalcides_mertensi | diurnal |
| lizards | Chalcides_minutus | Chalcides_minutus | diurnal |
| lizards | Chalcides_mionecton | Chalcides_mionecton | cathemeral |
| lizards | Chalcides_montanus | Chalcides_montanus | diurnal |
| lizards | Chalcides_ocellatus | Chalcides_ocellatus | cathemeral |
| lizards | Chalcides_parallelus | Chalcides_parallelus | diurnal |
| lizards | Chalcides_polylepis | Chalcides_polylepis | cathemeral |
| lizards | Chalcides_pseudostriatus | Chalcides_pseudostriatus | diurnal |
| lizards | Chalcides_ragazzii | Chalcides_ragazzii | diurnal |
| lizards | Chalcides_sepsoides | Chalcides_sepsoides | cathemeral |
| lizards | Chalcides_sexlineatus | Chalcides_sexlineatus | diurnal |
| lizards | Chalcides_striatus | Chalcides_striatus | diurnal |
| lizards | Chalcides_viridanus | Chalcides_viridanus | diurnal |
| lizards | Chalcidoseps_thwaitesi | Chalcidoseps_thwaitesi | diurnal |
| lizards | Chamaeleo_africanus | Chamaeleo_africanus | diurnal |
| lizards | Chamaeleo_calyptratus | Chamaeleo_calyptratus | diurnal |
| lizards | Chamaeleo_chamaeleon | Chamaeleo_chamaeleon | diurnal |
| lizards | Chamaeleo_dilepis | Chamaeleo_dilepis | diurnal |
| lizards | Chamaeleo_gracilis | Chamaeleo_gracilis | diurnal |
| lizards | Chamaeleo_laevigatus | Chamaeleo_laevigatus | diurnal |
| lizards | Chamaeleo_monachus | Chamaeleo_monachus | diurnal |
| lizards | Chamaeleo_namaquensis | Chamaeleo_namaquensis | diurnal |
| lizards | Chamaeleo_senegalensis | Chamaeleo_senegalensis | diurnal |
| lizards | Chamaeleo_zeylanicus | Chamaeleo_zeylanicus | diurnal |
| lizards | Chamaesaura_aenea | Chamaesaura_aenea | diurnal |
| lizards | Chamaesaura_anguina | Chamaesaura_anguina | diurnal |

| | | | |
|---|---|---|---|
| lizards | Chamaesaura_macrolepis | Chamaesaura_macrolepis | diurnal |
| lizards | Chamaesaura_miopropus | Chamaesaura_miopropus | diurnal |
| lizards | Chamaesaura_tenuior | Chamaesaura_tenuior | diurnal |
| lizards | Chatogekko_amazonicus | Chatogekko_amazonicus | diurnal |
| lizards | Chelosania_brunnea | Chelosania_brunnea | diurnal |
| lizards | Chioninia_coctei | Chioninia_coctei | cathemeral |
| lizards | Chioninia_delalandii | Chioninia_delalandii | diurnal |
| lizards | Chioninia_vaillantii | Chioninia_vaillantii | diurnal |
| lizards | Chlamydosaurus_kingii | Chlamydosaurus_kingii | diurnal |
| lizards | Chondrodactylus_angulifer | Chondrodactylus_angulifer | nocturnal |
| lizards | Chondrodactylus_bibronii | Chondrodactylus_bibronii | nocturnal |
| lizards | Chondrodactylus_fitzsimonsi | Chondrodactylus_fitzsimonsi | nocturnal |
| lizards | Chondrodactylus_laevigatus | Pachydactylus_laevigatus | cathemeral |
| lizards | Chondrodactylus_turneri | Chondrodactylus_turneri | nocturnal |
| lizards | Christinus_alexanderi | Christinus_alexanderi | nocturnal |
| lizards | Christinus_guentheri | Christinus_guentheri | nocturnal |
| lizards | Christinus_marmoratus | Christinus_marmoratus | nocturnal |
| lizards | Cnemaspis_affinis | Cnemaspis_affinis | diurnal |
| lizards | Cnemaspis_argus | Cnemaspis_argus | diurnal |
| lizards | Cnemaspis_assamensis | Cnemaspis_assamensis | diurnal |
| lizards | Cnemaspis_aurantiacopes | Cnemaspis_aurantiacopes | nocturnal |
| lizards | Cnemaspis_baueri | Cnemaspis_baueri | cathemeral |
| lizards | Cnemaspis_bayuensis | Cnemaspis_bayuensis | diurnal |
| lizards | Cnemaspis_bidongensis | Cnemaspis_bidongensis | cathemeral |
| lizards | Cnemaspis_biocellata | Cnemaspis_biocellata | cathemeral |
| lizards | Cnemaspis_boulengeri | Cnemaspis_boulengerii | diurnal |
| lizards | Cnemaspis_caudanivea | Cnemaspis_caudanivea | cathemeral |
| lizards | Cnemaspis_chanardi | Cnemaspis_chanardi | diurnal |
| lizards | Cnemaspis_chanthaburiensis | Cnemaspis_chanthaburiensis | cathemeral |
| lizards | Cnemaspis_flavigaster | Cnemaspis_flavigaster | cathemeral |
| lizards | Cnemaspis_flavolineata | Cnemaspis_flavolineata | diurnal |
| lizards | Cnemaspis_gemunu | Cnemaspis_gemunu | diurnal |
| lizards | Cnemaspis_gracilis | Cnemaspis_gracilis | nocturnal |
| lizards | Cnemaspis_grismeri | Cnemaspis_grismeri | diurnal |

| | | | |
|---|---|---|---|
| lizards | Cnemaspis_hangus | Cnemaspis_hangus | diurnal |
| lizards | Cnemaspis_harimau | Cnemaspis_harimau | nocturnal |
| lizards | Cnemaspis_heteropholis | Cnemaspis_heteropholis | cathemeral |
| lizards | Cnemaspis_huaseesom | Cnemaspis_huaseesom | nocturnal |
| lizards | Cnemaspis_jerdonii | Cnemaspis_jerdonii | cathemeral |
| lizards | Cnemaspis_kandiana | Cnemaspis_kandiana | diurnal |
| lizards | Cnemaspis_kendallii | Cnemaspis_kendallii | diurnal |
| lizards | Cnemaspis_kumarasinghei | Cnemaspis_kumarasinghei | nocturnal |
| lizards | Cnemaspis_kumpoli | Cnemaspis_kumpoli | nocturnal |
| lizards | Cnemaspis_limi | Cnemaspis_limi | cathemeral |
| lizards | Cnemaspis_littoralis | Cnemaspis_littoralis | diurnal |
| lizards | Cnemaspis_mcguirei | Cnemaspis_mcguirei | cathemeral |
| lizards | Cnemaspis_monachorum | Cnemaspis_monachorum | diurnal |
| lizards | Cnemaspis_mumpuniae | Cnemaspis_mumpuniae | nocturnal |
| lizards | Cnemaspis_nairi | Cnemaspis_nairi | diurnal |
| lizards | Cnemaspis_nigridia | Cnemaspis_nigridia | diurnal |
| lizards | Cnemaspis_nuicamensis | Cnemaspis_nuicamensis | diurnal |
| lizards | Cnemaspis_omari | Cnemaspis_omari | diurnal |
| lizards | Cnemaspis_ornata | Cnemaspis_ornata | diurnal |
| lizards | Cnemaspis_otai | Cnemaspis_otai | diurnal |
| lizards | Cnemaspis_pemanggilensis | Cnemaspis_pemanggilensis | cathemeral |
| lizards | Cnemaspis_peninsularis | Cnemaspis_peninsularis | diurnal |
| lizards | Cnemaspis_perhentianensis | Cnemaspis_perhentianensis | cathemeral |
| lizards | Cnemaspis_podihuna | Cnemaspis_podihuna | diurnal |
| lizards | Cnemaspis_pseudomcguirei | Cnemaspis_pseudomcguirei | cathemeral |
| lizards | Cnemaspis_psychedelica | Cnemaspis_psychedelica | diurnal |
| lizards | Cnemaspis_punctatonuchalis | Cnemaspis_punctatonuchalis | nocturnal |
| lizards | Cnemaspis_rammalensis | Cnemaspis_rammalensis | diurnal |
| lizards | Cnemaspis_roticanai | Cnemaspis_roticanai | cathemeral |
| lizards | Cnemaspis_scalpensis | Cnemaspis_scalpensis | diurnal |
| lizards | Cnemaspis_selamatkanmerapoh | Cnemaspis_selamatkanmerapoh | cathemeral |
| lizards | Cnemaspis_shahruli | Cnemaspis_shahruli | cathemeral |
| lizards | Cnemaspis_siamensis | Cnemaspis_siamensis | cathemeral |
| lizards | Cnemaspis_silvula | Cnemaspis_silvula | diurnal |

| | | | |
|---|---|---|---|
| lizards | Cnemaspis_stongensis | Cnemaspis_stongensis | nocturnal |
| lizards | Cnemaspis_sundainsula | Cnemaspis_sundainsula | diurnal |
| lizards | Cnemaspis_temiah | Cnemaspis_temiah | diurnal |
| lizards | Cnemaspis_tropidogaster | Cnemaspis_tropidogaster | diurnal |
| lizards | Cnemaspis_tucdupensis | Cnemaspis_tucdupensis | cathemeral |
| lizards | Cnemaspis_yercaudensis | Cnemaspis_yercaudensis | nocturnal |
| lizards | Cnemidophorus_arenivagus | Cnemidophorus_arenivagus | diurnal |
| lizards | Cnemidophorus_arubensis | Cnemidophorus_arubensis | diurnal |
| lizards | Cnemidophorus_cryptus | Cnemidophorus_cryptus | diurnal |
| lizards | Cnemidophorus_gramivagus | Cnemidophorus_gramivagus | diurnal |
| lizards | Cnemidophorus_lemniscatus | Cnemidophorus_lemniscatus | diurnal |
| lizards | Cnemidophorus_murinus | Cnemidophorus_murinus | diurnal |
| lizards | Cnemidophorus_ruatanus | Cnemidophorus_ruatanus | diurnal |
| lizards | Cnemidophorus_ruthveni | Cnemidophorus_ruthveni | diurnal |
| lizards | Coeranoscincus_reticulatus | Coeranoscincus_reticulatus | cathemeral |
| lizards | Coleodactylus_meridionalis | Coleodactylus_meridionalis | diurnal |
| lizards | Coleodactylus_septentrionalis | Coleodactylus_septentrionalis | diurnal |
| lizards | Coleonyx_brevis | Coleonyx_brevis | nocturnal |
| lizards | Coleonyx_elegans | Coleonyx_elegans | nocturnal |
| lizards | Coleonyx_gypsicolus | Coleonyx_gypsicolus | nocturnal |
| lizards | Coleonyx_mitratus | Coleonyx_mitratus | nocturnal |
| lizards | Coleonyx_reticulatus | Coleonyx_reticulatus | nocturnal |
| lizards | Coleonyx_switaki | Coleonyx_switaki | nocturnal |
| lizards | Coleonyx_variegatus | Coleonyx_variegatus | nocturnal |
| lizards | Colobosaura_modesta | Colobosaura_modesta | diurnal |
| lizards | Concinnia_brachysoma | Concinnia_brachyosoma | diurnal |
| lizards | Concinnia_martini | Concinnia_martini | diurnal |
| lizards | Concinnia_sokosoma | Concinnia_sokosoma | diurnal |
| lizards | Concinnia_tenuis | Concinnia_tenuis | cathemeral |
| lizards | Congolacerta_vauereselli | Congolacerta_vauereselli | diurnal |
| lizards | Conolophus_marthae | Conolophus_marthae | diurnal |
| lizards | Conolophus_pallidus | Conolophus_pallidus | diurnal |
| lizards | Conolophus_subcristatus | Conolophus_subcristatus | diurnal |
| lizards | Contomastix_lacertoides | Contomastix_lacertoides | diurnal |

| | | | |
|---|---|---|---|
| lizards | Contomastix_vacariensis | Contomastix_vacariensis | diurnal |
| lizards | Copeoglossum_arajara | Copeoglossum_arajara | diurnal |
| lizards | Copeoglossum_aurae | Copeoglossum_aurae | diurnal |
| lizards | Copeoglossum_nigropunctatum | Copeoglossum_nigropunctatum | diurnal |
| lizards | Cophosaurus_texanus | Cophosaurus_texanus | diurnal |
| lizards | Cophoscincopus_durus | Cophoscincopus_durus | diurnal |
| lizards | Cophoscincopus_greeri | Cophoscincopus_greeri | diurnal |
| lizards | Cophoscincopus_simulans | Cophoscincopus_simulans | diurnal |
| lizards | Cophotis_ceylanica | Cophotis_ceylanica | diurnal |
| lizards | Cordylus_cordylus | Cordylus_cordylus | diurnal |
| lizards | Cordylus_imkeae | Cordylus_imkeae | diurnal |
| lizards | Cordylus_jonesii | Cordylus_jonesii | diurnal |
| lizards | Cordylus_macropholis | Cordylus_macropholis | diurnal |
| lizards | Cordylus_mclachlani | Cordylus_mclachlani | diurnal |
| lizards | Cordylus_meculae | Cordylus_meculae | diurnal |
| lizards | Cordylus_minor | Cordylus_minor | diurnal |
| lizards | Cordylus_niger | Cordylus_niger | diurnal |
| lizards | Cordylus_nyikae | Cordylus_nyikae | diurnal |
| lizards | Cordylus_oelofseni | Cordylus_oelofseni | diurnal |
| lizards | Cordylus_rhodesianus | Cordylus_rhodesianus | diurnal |
| lizards | Cordylus_tropidosternum | Cordylus_tropidosternum | diurnal |
| lizards | Cordylus_vittifer | Cordylus_vittifer | diurnal |
| lizards | Correlophus_belepensis | Correlophus_belepensis | nocturnal |
| lizards | Correlophus_ciliatus | Correlophus_ciliatus | nocturnal |
| lizards | Correlophus_sarasinorum | Correlophus_sarasinorum | nocturnal |
| lizards | Corucia_zebrata | Corucia_zebrata | nocturnal |
| lizards | Coryphophylax_subcristatus | Coryphophylax_subcristatus | diurnal |
| lizards | Corytophanes_cristatus | Corytophanes_cristatus | diurnal |
| lizards | Corytophanes_hernandesii | Corytophanes_hernandesii | diurnal |
| lizards | Corytophanes_percarinatus | Corytophanes_percarinatus | diurnal |
| lizards | Crenadactylus_ocellatus | Crenadactylus_ocellatus | nocturnal |
| lizards | Cricosaura_typica | Cricosaura_typica | cathemeral |
| lizards | Crocodilurus_amazonicus | Crocodilurus_amazonicus | diurnal |
| lizards | Crossobamon_eversmanni | Crossobamon_eversmanni | nocturnal |

| | | | |
|---|---|---|---|
| lizards | Crossobamon_orientalis | Crossobamon_orientalis | nocturnal |
| lizards | Crotaphytus_antiquus | Crotaphytus_antiquus | diurnal |
| lizards | Crotaphytus_bicinctores | Crotaphytus_bicinctores | diurnal |
| lizards | Crotaphytus_collaris | Crotaphytus_collaris | diurnal |
| lizards | Crotaphytus_dickersonae | Crotaphytus_dickersonae | diurnal |
| lizards | Crotaphytus_grismeri | Crotaphytus_grismeri | diurnal |
| lizards | Crotaphytus_insularis | Crotaphytus_insularis | diurnal |
| lizards | Crotaphytus_nebrius | Crotaphytus_nebrius | diurnal |
| lizards | Crotaphytus_reticulatus | Crotaphytus_reticulatus | diurnal |
| lizards | Crotaphytus_vestigium | Crotaphytus_vestigium | diurnal |
| lizards | Cryptactites_peringueyi | Cryptactites_peringueyi | nocturnal |
| lizards | Cryptoblepharus_africanus | Cryptoblepharus_africanus | diurnal |
| lizards | Cryptoblepharus_australis | Cryptoblepharus_australis | diurnal |
| lizards | Cryptoblepharus_bitaeniatus | Cryptoblepharus_bitaeniatus | diurnal |
| lizards | Cryptoblepharus_boutonii | Cryptoblepharus_boutonii | diurnal |
| lizards | Cryptoblepharus_buchananii | Cryptoblepharus_buchananii | diurnal |
| lizards | Cryptoblepharus_burdeni | Cryptoblepharus_burdeni | diurnal |
| lizards | Cryptoblepharus_caudatus | Cryptoblepharus_caudatus | diurnal |
| lizards | Cryptoblepharus_cygnatus | Cryptoblepharus_cygnatus | diurnal |
| lizards | Cryptoblepharus_egeriae | Cryptoblepharus_egeriae | diurnal |
| lizards | Cryptoblepharus_eximius | Cryptoblepharus_eximius | diurnal |
| lizards | Cryptoblepharus_furvus | Cryptoblepharus_furvus | diurnal |
| lizards | Cryptoblepharus_gloriosus | Cryptoblepharus_gloriosus | diurnal |
| lizards | Cryptoblepharus_litoralis | Cryptoblepharus_litoralis | diurnal |
| lizards | Cryptoblepharus_megastictus | Cryptoblepharus_megastictus | diurnal |
| lizards | Cryptoblepharus_metallicus | Cryptoblepharus_metallicus | diurnal |
| lizards | Cryptoblepharus_nigropunctatus | Cryptoblepharus_nigropunctatus | diurnal |
| lizards | Cryptoblepharus_novocaledonicus | Cryptoblepharus_novocaledonicus | diurnal |
| lizards | Cryptoblepharus_novohebridicus | Cryptoblepharus_novohebridicus | diurnal |
| lizards | Cryptoblepharus_pannosus | Cryptoblepharus_pannosus | diurnal |
| lizards | Cryptoblepharus_plagiocephalus | Cryptoblepharus_plagiocephalus | diurnal |
| lizards | Cryptoblepharus_poecilopleurus | Cryptoblepharus_poecilopleurus | diurnal |
| lizards | Cryptoblepharus_pulcher | Cryptoblepharus_pulcher | diurnal |
| lizards | Cryptoblepharus_renschi | Cryptoblepharus_renschi | diurnal |

| | | | |
|---|---|---|---|
| lizards | Cryptoblepharus_ruber | Cryptoblepharus_ruber | diurnal |
| lizards | Cryptoblepharus_virgatus | Cryptoblepharus_virgatus | diurnal |
| lizards | Cryptoblepharus_yulensis | Cryptoblepharus_yulensis | diurnal |
| lizards | Ctenophorus_adelaidensis | Ctenophorus_adelaidensis | diurnal |
| lizards | Ctenophorus_butlerorum | Ctenophorus_butlerorum | diurnal |
| lizards | Ctenophorus_caudicinctus | Ctenophorus_caudicinctus | diurnal |
| lizards | Ctenophorus_clayi | Ctenophorus_clayi | diurnal |
| lizards | Ctenophorus_cristatus | Ctenophorus_cristatus | diurnal |
| lizards | Ctenophorus_decresii | Ctenophorus_decresii | diurnal |
| lizards | Ctenophorus_femoralis | Ctenophorus_femoralis | diurnal |
| lizards | Ctenophorus_fionni | Ctenophorus_fionni | diurnal |
| lizards | Ctenophorus_fordi | Ctenophorus_fordi | diurnal |
| lizards | Ctenophorus_isolepis | Ctenophorus_isolepis | diurnal |
| lizards | Ctenophorus_maculatus | Ctenophorus_maculatus | diurnal |
| lizards | Ctenophorus_maculosus | Ctenophorus_maculosus | diurnal |
| lizards | Ctenophorus_mirrityana | Ctenophorus_mirrityana | diurnal |
| lizards | Ctenophorus_nguyarna | Ctenophorus_nguyarna | diurnal |
| lizards | Ctenophorus_nuchalis | Ctenophorus_nuchalis | diurnal |
| lizards | Ctenophorus_ornatus | Ctenophorus_ornatus | diurnal |
| lizards | Ctenophorus_parviceps | Ctenophorus_parviceps | diurnal |
| lizards | Ctenophorus_pictus | Ctenophorus_pictus | diurnal |
| lizards | Ctenophorus_reticulatus | Ctenophorus_reticulatus | diurnal |
| lizards | Ctenophorus_salinarum | Ctenophorus_salinarum | diurnal |
| lizards | Ctenophorus_scutulatus | Ctenophorus_scutulatus | diurnal |
| lizards | Ctenophorus_vadnappa | Ctenophorus_vadnappa | diurnal |
| lizards | Ctenosaura_acanthura | Ctenosaura_acanthura | diurnal |
| lizards | Ctenosaura_bakeri | Ctenosaura_bakeri | diurnal |
| lizards | Ctenosaura_clarki | Ctenosaura_clarki | diurnal |
| lizards | Ctenosaura_hemilopha | Ctenosaura_hemilopha | diurnal |
| lizards | Ctenosaura_macrolopha | Ctenosaura_macrolopha | diurnal |
| lizards | Ctenosaura_melanosterna | Ctenosaura_melanosterna | diurnal |
| lizards | Ctenosaura_oedirhina | Ctenosaura_oedirhina | diurnal |
| lizards | Ctenosaura_palearis | Ctenosaura_palearis | diurnal |
| lizards | Ctenosaura_pectinata | Ctenosaura_pectinata | diurnal |

| | | | |
|---|---|---|---|
| lizards | Ctenosaura_quinquecarinata | Ctenosaura_quinquecarinata | diurnal |
| lizards | Ctenosaura_similis | Ctenosaura_similis | diurnal |
| lizards | Ctenotus_alacer | Ctenotus_alacer | diurnal |
| lizards | Ctenotus_alleni | Ctenotus_alleni | diurnal |
| lizards | Ctenotus_ariadnae | Ctenotus_ariadnae | diurnal |
| lizards | Ctenotus_arnhemensis | Ctenotus_arnhemensis | diurnal |
| lizards | Ctenotus_atlas | Ctenotus_atlas | diurnal |
| lizards | Ctenotus_australis | Ctenotus_australis | diurnal |
| lizards | Ctenotus_brevipes | Ctenotus_brevipes | diurnal |
| lizards | Ctenotus_brooksi | Ctenotus_brooksi | diurnal |
| lizards | Ctenotus_calurus | Ctenotus_calurus | diurnal |
| lizards | Ctenotus_coggeri | Ctenotus_coggeri | cathemeral |
| lizards | Ctenotus_colletti | Ctenotus_colletti | diurnal |
| lizards | Ctenotus_dux | Ctenotus_dux | diurnal |
| lizards | Ctenotus_essingtonii | Ctenotus_essingtonii | diurnal |
| lizards | Ctenotus_gagudju | Ctenotus_gagudju | diurnal |
| lizards | Ctenotus_gemmula | Ctenotus_gemmula | diurnal |
| lizards | Ctenotus_grandis | Ctenotus_grandis | cathemeral |
| lizards | Ctenotus_hanloni | Ctenotus_hanloni | diurnal |
| lizards | Ctenotus_hebetior | Ctenotus_hebetior | diurnal |
| lizards | Ctenotus_impar | Ctenotus_impar | diurnal |
| lizards | Ctenotus_inornatus | Ctenotus_inornatus | cathemeral |
| lizards | Ctenotus_kurnbudj | Ctenotus_kurnbudj | diurnal |
| lizards | Ctenotus_labillardieri | Ctenotus_labillardieri | diurnal |
| lizards | Ctenotus_lancelini | Ctenotus_lancelini | diurnal |
| lizards | Ctenotus_leae | Ctenotus_leae | diurnal |
| lizards | Ctenotus_leonhardii | Ctenotus_leonhardii | diurnal |
| lizards | Ctenotus_mimetes | Ctenotus_mimetes | diurnal |
| lizards | Ctenotus_nasutus | Ctenotus_nasutus | diurnal |
| lizards | Ctenotus_olympicus | Ctenotus_olympicus | diurnal |
| lizards | Ctenotus_orientalis | Ctenotus_orientalis | diurnal |
| lizards | Ctenotus_pallescens | Ctenotus_pallescens | diurnal |
| lizards | Ctenotus_pantherinus | Ctenotus_pantherinus | cathemeral |
| lizards | Ctenotus_piankai | Ctenotus_piankai | diurnal |

| | | | |
|---|---|---|---|
| lizards | Ctenotus_quattuordecimlineatus | Ctenotus_quattuordecimlineatus | diurnal |
| lizards | Ctenotus_regius | Ctenotus_regius | diurnal |
| lizards | Ctenotus_robustus | Ctenotus_robustus | cathemeral |
| lizards | Ctenotus_rufescens | Ctenotus_rufescens | cathemeral |
| lizards | Ctenotus_rutilans | Ctenotus_rutilans | diurnal |
| lizards | Ctenotus_schomburgkii | Ctenotus_schomburgkii | diurnal |
| lizards | Ctenotus_septenarius | Ctenotus_septenarius | diurnal |
| lizards | Ctenotus_spaldingi | Ctenotus_spaldingi | diurnal |
| lizards | Ctenotus_storri | Ctenotus_storri | diurnal |
| lizards | Ctenotus_strauchii | Ctenotus_strauchii | diurnal |
| lizards | Ctenotus_taeniolatus | Ctenotus_taeniolatus | diurnal |
| lizards | Ctenotus_uber | Ctenotus_uber | diurnal |
| lizards | Ctenotus_vertebralis | Ctenotus_vertebralis | diurnal |
| lizards | Cyclodomorphus_branchialis | Cyclodomorphus_branchialis | cathemeral |
| lizards | Cyclodomorphus_casuarinae | Cyclodomorphus_casuarinae | cathemeral |
| lizards | Cyclodomorphus_celatus | Cyclodomorphus_celatus | nocturnal |
| lizards | Cyclodomorphus_gerrardii | Hemisphaeriodon_gerrardii | cathemeral |
| lizards | Cyclodomorphus_maximus | Cyclodomorphus_maxima | cathemeral |
| lizards | Cyclodomorphus_melanops | Cyclodomorphus_melanops | cathemeral |
| lizards | Cyclodomorphus_michaeli | Cyclodomorphus_michaeli | cathemeral |
| lizards | Cyclodomorphus_praealtus | Cyclodomorphus_praealtus | diurnal |
| lizards | Cyclodomorphus_venustus | Cyclodomorphus_venustus | nocturnal |
| lizards | Cyclura_carinata | Cyclura_carinata | diurnal |
| lizards | Cyclura_collei | Cyclura_collei | diurnal |
| lizards | Cyclura_cornuta | Cyclura_cornuta | diurnal |
| lizards | Cyclura_cychlura | Cyclura_cychlura | diurnal |
| lizards | Cyclura_lewisi | Cyclura_lewisi | diurnal |
| lizards | Cyclura_nubila | Cyclura_nubila | diurnal |
| lizards | Cyclura_pinguis | Cyclura_pinguis | diurnal |
| lizards | Cyclura_ricordi | Cyclura_ricordi | diurnal |
| lizards | Cyclura_rileyi | Cyclura_rileyi | diurnal |
| lizards | Cyrtodactylus_aequalis | Cyrtodactylus_aequalis | nocturnal |
| lizards | Cyrtodactylus_agusanensis | Cyrtodactylus_agusanensis | cathemeral |
| lizards | Cyrtodactylus_albofasciatus | Cyrtodactylus_albofasciatus | nocturnal |

| | | | |
|---|---|---|---|
| lizards | Cyrtodactylus_angularis | Cyrtodactylus_angularis | nocturnal |
| lizards | Cyrtodactylus_annandalei | Cyrtodactylus_annandalei | nocturnal |
| lizards | Cyrtodactylus_astrum | Cyrtodactylus_astrum | nocturnal |
| lizards | Cyrtodactylus_aurensis | Cyrtodactylus_aurensis | nocturnal |
| lizards | Cyrtodactylus_australotitiwangsaensis | Cyrtodactylus_australotitiwangsaensis | nocturnal |
| lizards | Cyrtodactylus_ayeyarwadyensis | Cyrtodactylus_ayeyarwadyensis | nocturnal |
| lizards | Cyrtodactylus_baluensis | Cyrtodactylus_baluensis | nocturnal |
| lizards | Cyrtodactylus_battalensis | Cyrtopodion_battalense | nocturnal |
| lizards | Cyrtodactylus_batucolus | Cyrtodactylus_batucolus | nocturnal |
| lizards | Cyrtodactylus_bintangrendah | Cyrtodactylus_bintangrendah | nocturnal |
| lizards | Cyrtodactylus_bintangtinggi | Cyrtodactylus_bintangtinggi | nocturnal |
| lizards | Cyrtodactylus_biordinis | Cyrtodactylus_biordinis | nocturnal |
| lizards | Cyrtodactylus_boreoclivus | Cyrtodactylus_boreoclivus | nocturnal |
| lizards | Cyrtodactylus_brevidactylus | Cyrtodactylus_brevidactylus | nocturnal |
| lizards | Cyrtodactylus_brevipalmatus | Cyrtodactylus_brevipalmatus | nocturnal |
| lizards | Cyrtodactylus_buchardi | Cyrtodactylus_buchardi | nocturnal |
| lizards | Cyrtodactylus_capreoloides | Cyrtodactylus_capreoloides | nocturnal |
| lizards | Cyrtodactylus_cattienensis | Cyrtodactylus_cattienensis | nocturnal |
| lizards | Cyrtodactylus_cavernicolus | Cyrtodactylus_cavernicolus | nocturnal |
| lizards | Cyrtodactylus_chanhomeae | Cyrtodactylus_chanhomeae | nocturnal |
| lizards | Cyrtodactylus_chrysopylos | Cyrtodactylus_chrysopylos | nocturnal |
| lizards | Cyrtodactylus_collegalensis | Cyrtodactylus_collegalensis | nocturnal |
| lizards | Cyrtodactylus_condorensis | Cyrtodactylus_condorensis | nocturnal |
| lizards | Cyrtodactylus_consobrinus | Cyrtodactylus_consobrinus | nocturnal |
| lizards | Cyrtodactylus_cryptus | Cyrtodactylus_cryptus | nocturnal |
| lizards | Cyrtodactylus_dattanensis | Cyrtopodion_dattanense | nocturnal |
| lizards | Cyrtodactylus_deccanensis | Cyrtodactylus_deccanensis | nocturnal |
| lizards | Cyrtodactylus_dumnuii | Cyrtodactylus_dumnuii | nocturnal |
| lizards | Cyrtodactylus_elok | Cyrtodactylus_elok | nocturnal |
| lizards | Cyrtodactylus_epiroticus | Cyrtodactylus_epiroticus | nocturnal |
| lizards | Cyrtodactylus_erythrops | Cyrtodactylus_erythrops | cathemeral |
| lizards | Cyrtodactylus_fraenatus | Cyrtodactylus_fraenatus | nocturnal |
| lizards | Cyrtodactylus_fumosus | Cyrtodactylus_fumosus | nocturnal |
| lizards | Cyrtodactylus_gansi | Cyrtodactylus_gansi | nocturnal |

| | | | |
|---|---|---|---|
| lizards | Cyrtodactylus_huynhi | Cyrtodactylus_huynhi | nocturnal |
| lizards | Cyrtodactylus_ingeri | Cyrtodactylus_ingeri | nocturnal |
| lizards | Cyrtodactylus_interdigitalis | Cyrtodactylus_interdigitalis | nocturnal |
| lizards | Cyrtodactylus_intermedius | Cyrtodactylus_intermedius | nocturnal |
| lizards | Cyrtodactylus_jarakensis | Cyrtodactylus_jarakensis | nocturnal |
| lizards | Cyrtodactylus_jelawangensis | Cyrtodactylus_jelawangensis | nocturnal |
| lizards | Cyrtodactylus_jellesmae | Cyrtodactylus_jellesmae | nocturnal |
| lizards | Cyrtodactylus_khasiensis | Cyrtodactylus_khasiensis | nocturnal |
| lizards | Cyrtodactylus_laevigatus | Cyrtodactylus_laevigatus | nocturnal |
| lizards | Cyrtodactylus_langkawiensis | Cyrtodactylus_langkawiensis | nocturnal |
| lizards | Cyrtodactylus_lateralis | Cyrtodactylus_lateralis | nocturnal |
| lizards | Cyrtodactylus_leegrismeri | Cyrtodactylus_leegrismeri | nocturnal |
| lizards | Cyrtodactylus_lekaguli | Cyrtodactylus_lekaguli | nocturnal |
| lizards | Cyrtodactylus_loriae | Cyrtodactylus_loriae | nocturnal |
| lizards | Cyrtodactylus_louisiadensis | Cyrtodactylus_louisiadensis | nocturnal |
| lizards | Cyrtodactylus_macrotuberculatus | Cyrtodactylus_macrotuberculatus | nocturnal |
| lizards | Cyrtodactylus_majulah | Cyrtodactylus_majulah | nocturnal |
| lizards | Cyrtodactylus_malayanus | Cyrtodactylus_malayanus | nocturnal |
| lizards | Cyrtodactylus_marmoratus | Cyrtodactylus_marmoratus | nocturnal |
| lizards | Cyrtodactylus_martini | Cyrtodactylus_martini | nocturnal |
| lizards | Cyrtodactylus_martinstolli | Cyrtodactylus_martinstolli | nocturnal |
| lizards | Cyrtodactylus_mcdonaldi | Cyrtodactylus_mcdonaldi | nocturnal |
| lizards | Cyrtodactylus_oldhami | Cyrtodactylus_oldhami | nocturnal |
| lizards | Cyrtodactylus_pageli | Cyrtodactylus_pageli | nocturnal |
| lizards | Cyrtodactylus_pantiensis | Cyrtodactylus_pantiensis | nocturnal |
| lizards | Cyrtodactylus_papilionoides | Cyrtodactylus_papilionoides | nocturnal |
| lizards | Cyrtodactylus_paradoxus | Cyrtodactylus_paradoxus | nocturnal |
| lizards | Cyrtodactylus_payacola | Cyrtodactylus_payacola | nocturnal |
| lizards | Cyrtodactylus_peguensis | Cyrtodactylus_peguensis | nocturnal |
| lizards | Cyrtodactylus_philippinicus | Cyrtodactylus_philippinicus | nocturnal |
| lizards | Cyrtodactylus_phongnhakebangensis | Cyrtodactylus_phongnhakebangensis | nocturnal |
| lizards | Cyrtodactylus_pseudoquadrivirgatus | Cyrtodactylus_pseudoquadrivirgatus | nocturnal |
| lizards | Cyrtodactylus_pubisulcus | Cyrtodactylus_pubisulcus | nocturnal |
| lizards | Cyrtodactylus_pulchellus | Cyrtodactylus_pulchellus | nocturnal |

| | | | |
|---|---|---|---|
| lizards | Cyrtodactylus_redimiculus | Cyrtodactylus_redimiculus | nocturnal |
| lizards | Cyrtodactylus_rubidus | Cyrtodactylus_rubidus | nocturnal |
| lizards | Cyrtodactylus_russelli | Cyrtodactylus_russelli | nocturnal |
| lizards | Cyrtodactylus_sadleiri | Cyrtodactylus_sadleiri | nocturnal |
| lizards | Cyrtodactylus_samroiyot | Cyrtodactylus_samroiyot | nocturnal |
| lizards | Cyrtodactylus_semenanjungensis | Cyrtodactylus_semenanjungensis | nocturnal |
| lizards | Cyrtodactylus_semiadii | Cyrtodactylus_semiadii | nocturnal |
| lizards | Cyrtodactylus_seribuatensis | Cyrtodactylus_seribuatensis | nocturnal |
| lizards | Cyrtodactylus_soba | Cyrtodactylus_soba | nocturnal |
| lizards | Cyrtodactylus_spinosus | Cyrtodactylus_spinosus | nocturnal |
| lizards | Cyrtodactylus_sumonthai | Cyrtodactylus_sumonthai | nocturnal |
| lizards | Cyrtodactylus_sworderi | Cyrtodactylus_sworderi | nocturnal |
| lizards | Cyrtodactylus_takouensis | Cyrtodactylus_takouensis | nocturnal |
| lizards | Cyrtodactylus_tebuensis | Cyrtodactylus_tebuensis | nocturnal |
| lizards | Cyrtodactylus_thochuensis | Cyrtodactylus_thochuensis | nocturnal |
| lizards | Cyrtodactylus_tigroides | Cyrtodactylus_tigroides | nocturnal |
| lizards | Cyrtodactylus_timur | Cyrtodactylus_timur | nocturnal |
| lizards | Cyrtodactylus_tiomanensis | Cyrtodactylus_tiomanensis | nocturnal |
| lizards | Cyrtodactylus_triedrus | Cyrtodactylus_triedrus | nocturnal |
| lizards | Cyrtodactylus_tuberculatus | Cyrtodactylus_tuberculatus | nocturnal |
| lizards | Cyrtodactylus_wakeorum | Cyrtodactylus_wakeorum | nocturnal |
| lizards | Cyrtodactylus_wayakonei | Cyrtodactylus_wayakonei | nocturnal |
| lizards | Cyrtodactylus_yakhuna | Cyrtodactylus_yakhuna | nocturnal |
| lizards | Cyrtodactylus_yangbayensis | Cyrtodactylus_yangbayensis | nocturnal |
| lizards | Cyrtodactylus_yoshii | Cyrtodactylus_yoshii | nocturnal |
| lizards | Cyrtopodion_agamuroides | Cyrtopodion_agamuroides | cathemeral |
| lizards | Cyrtopodion_fortmunroi | Cyrtopodion_fortmunroi | nocturnal |
| lizards | Cyrtopodion_golubevi | Cyrtopodion_golubevi | nocturnal |
| lizards | Cyrtopodion_indusoani | Cyrtopodion_indusoani | cathemeral |
| lizards | Cyrtopodion_kachhense | Cyrtopodion_kachhense | nocturnal |
| lizards | Cyrtopodion_kohsulaimanai | Cyrtopodion_kohsulaimanai | nocturnal |
| lizards | Cyrtopodion_potoharense | Cyrtopodion_potoharense | nocturnal |
| lizards | Cyrtopodion_rohtasfortai | Cyrtopodion_rohtasfortai | nocturnal |
| lizards | Cyrtopodion_scabrum | Cyrtopodion_scabrum | nocturnal |

| | | | |
|---|---|---|---|
| lizards | Cyrtopodion_watsoni | Cyrtopodion_watsoni | cathemeral |
| lizards | Dactylocnemis_pacificus | Dactylocnemis_pacificus | nocturnal |
| lizards | Dalmatolacerta_oxycephala | Dalmatolacerta_oxycephala | diurnal |
| lizards | Darevskia_alpina | Darevskia_alpina | diurnal |
| lizards | Darevskia_armeniaca | Darevskia_armeniaca | diurnal |
| lizards | Darevskia_bendimahiensis | Darevskia_bendimahiensis | diurnal |
| lizards | Darevskia_bithynica | Darevskia_bithynica | diurnal |
| lizards | Darevskia_brauneri | Darevskia_brauneri | diurnal |
| lizards | Darevskia_caucasica | Darevskia_caucasica | diurnal |
| lizards | Darevskia_chlorogaster | Darevskia_chlorogaster | diurnal |
| lizards | Darevskia_daghestanica | Darevskia_daghestanica | diurnal |
| lizards | Darevskia_dahli | Darevskia_dahli | diurnal |
| lizards | Darevskia_defilippii | Darevskia_defilippii | diurnal |
| lizards | Darevskia_derjugini | Darevskia_derjugini | diurnal |
| lizards | Darevskia_lindholmi | Darevskia_lindholmi | diurnal |
| lizards | Darevskia_mixta | Darevskia_mixta | diurnal |
| lizards | Darevskia_parvula | Darevskia_parvula | diurnal |
| lizards | Darevskia_portschinskii | Darevskia_portschinskii | diurnal |
| lizards | Darevskia_praticola | Darevskia_praticola | diurnal |
| lizards | Darevskia_raddei | Darevskia_raddei | diurnal |
| lizards | Darevskia_rostombekowi | Darevskia_rostombekovi | diurnal |
| lizards | Darevskia_rudis | Darevskia_rudis | diurnal |
| lizards | Darevskia_sapphirina | Darevskia_sapphirina | diurnal |
| lizards | Darevskia_saxicola | Darevskia_saxicola | diurnal |
| lizards | Darevskia_unisexualis | Darevskia_unisexualis | diurnal |
| lizards | Darevskia_valentini | Darevskia_valentini | diurnal |
| lizards | Dasia_grisea | Dasia_grisea | diurnal |
| lizards | Dasia_haliana | Dasia_haliana | diurnal |
| lizards | Dasia_olivacea | Dasia_olivacea | diurnal |
| lizards | Dasia_vittata | Dasia_vittata | diurnal |
| lizards | Delma_australis | Delma_australis | diurnal |
| lizards | Delma_borea | Delma_borea | nocturnal |
| lizards | Delma_butleri | Delma_butleri | cathemeral |
| lizards | Delma_concinna | Delma_concinna | cathemeral |

| | | | |
|---|---|---|---|
| lizards | Delma_fraseri | Delma_fraseri | cathemeral |
| lizards | Delma_grayii | Delma_grayii | diurnal |
| lizards | Delma_impar | Delma_impar | diurnal |
| lizards | Delma_inornata | Delma_inornata | cathemeral |
| lizards | Delma_labialis | Delma_labialis | diurnal |
| lizards | Delma_molleri | Delma_molleri | cathemeral |
| lizards | Delma_nasuta | Delma_nasuta | cathemeral |
| lizards | Delma_plebeia | Delma_plebeia | diurnal |
| lizards | Delma_tincta | Delma_tincta | cathemeral |
| lizards | Dendragama_boulengeri | Dendragama_boulengeri | diurnal |
| lizards | Dierogekko_inexpectatus | Dierogekko_inexpectatus | nocturnal |
| lizards | Dierogekko_insularis | Dierogekko_insularis | nocturnal |
| lizards | Dierogekko_kaalaensis | Dierogekko_kaalaensis | nocturnal |
| lizards | Dierogekko_koniambo | Dierogekko_koniambo | nocturnal |
| lizards | Dierogekko_nehoueensis | Dierogekko_nehoueensis | nocturnal |
| lizards | Dierogekko_poumensis | Dierogekko_poumensis | nocturnal |
| lizards | Dierogekko_thomaswhitei | Dierogekko_thomaswhitei | nocturnal |
| lizards | Dierogekko_validiclavis | Dierogekko_validiclavis | nocturnal |
| lizards | Dinarolacerta_mosorensis | Dinarolacerta_mosorensis | diurnal |
| lizards | Diplodactylus_calcicolus | Diplodactylus_calcicolus | nocturnal |
| lizards | Diplodactylus_capensis | Diplodactylus_capensis | nocturnal |
| lizards | Diplodactylus_conspicillatus | Diplodactylus_conspicillatus | nocturnal |
| lizards | Diplodactylus_galeatus | Diplodactylus_galeatus | nocturnal |
| lizards | Diplodactylus_granariensis | Diplodactylus_granariensis | nocturnal |
| lizards | Diplodactylus_klugei | Diplodactylus_klugei | nocturnal |
| lizards | Diplodactylus_lateroides | Diplodactylus_lateroides | nocturnal |
| lizards | Diplodactylus_mitchelli | Diplodactylus_mitchelli | nocturnal |
| lizards | Diplodactylus_ornatus | Diplodactylus_ornatus | nocturnal |
| lizards | Diplodactylus_pulcher | Diplodactylus_pulcher | nocturnal |
| lizards | Diplodactylus_savagei | Diplodactylus_savagei | nocturnal |
| lizards | Diplodactylus_tessellatus | Diplodactylus_tessellatus | nocturnal |
| lizards | Diplodactylus_vittatus | Diplodactylus_vittatus | nocturnal |
| lizards | Diploderma_brevipes | Japalura_brevipes | diurnal |
| lizards | Diploderma_chapaense | Japalura_chapaensis | diurnal |

| | | | |
|---|---|---|---|
| lizards | Diploderma_dymondi | Japalura_dymondi | diurnal |
| lizards | Diploderma_flaviceps | Japalura_flaviceps | diurnal |
| lizards | Diploderma_luei | Japalura_luei | diurnal |
| lizards | Diploderma_makii | Japalura_makii | diurnal |
| lizards | Diploderma_polygonatum | Japalura_polygonata | diurnal |
| lizards | Diploderma_splendidum | Japalura_splendida | diurnal |
| lizards | Diploderma_swinhonis | Japalura_swinhonis | diurnal |
| lizards | Diploderma_varcoae | Japalura_varcoae | diurnal |
| lizards | Diploderma_yunnanense | Japalura_yunnanensis | diurnal |
| lizards | Diploglossus_delasagra | Diploglossus_delasagra | diurnal |
| lizards | Diploglossus_fasciatus | Diploglossus_fasciatus | diurnal |
| lizards | Diploglossus_lessonae | Diploglossus_lessonae | diurnal |
| lizards | Diploglossus_monotropis | Diploglossus_monotropis | diurnal |
| lizards | Diplolaemus_bibronii | Diplolaemus_bibronii | diurnal |
| lizards | Diporiphora_albilabris | Diporiphora_albilabris | diurnal |
| lizards | Diporiphora_amphiboluroides | Diporiphora_amphiboluroides | diurnal |
| lizards | Diporiphora_australis | Diporiphora_australis | diurnal |
| lizards | Diporiphora_bennettii | Diporiphora_bennettii | diurnal |
| lizards | Diporiphora_bilineata | Diporiphora_bilineata | diurnal |
| lizards | Diporiphora_margaretae | Diporiphora_margaretae | diurnal |
| lizards | Diporiphora_nobbi | Diporiphora_nobbi | diurnal |
| lizards | Diporiphora_paraconvergens | Diporiphora_paraconvergens | diurnal |
| lizards | Diporiphora_superba | Diporiphora_superba | diurnal |
| lizards | Diporiphora_winneckei | Diporiphora_winneckei | diurnal |
| lizards | Dipsosaurus_dorsalis | Dipsosaurus_dorsalis | diurnal |
| lizards | Dixonius_hangseesom | Dixonius_hangseesom | nocturnal |
| lizards | Dixonius_siamensis | Dixonius_siamensis | nocturnal |
| lizards | Dixonius_vietnamensis | Dixonius_vietnamensis | nocturnal |
| lizards | Dopasia_gracilis | Dopasia_gracilis | cathemeral |
| lizards | Dopasia_hainanensis | Dopasia_hainanensis | diurnal |
| lizards | Dopasia_harti | Dopasia_harti | diurnal |
| lizards | Dopasia_sokolovi | Dopasia_sokolovi | diurnal |
| lizards | Dracaena_guianensis | Dracaena_guianensis | diurnal |
| lizards | Draco_abbreviatus | Draco_abbreviatus | diurnal |

| | | | |
|---|---|---|---|
| lizards | Draco_beccarii | Draco_beccarii | diurnal |
| lizards | Draco_biaro | Draco_biaro | diurnal |
| lizards | Draco_bimaculatus | Draco_bimaculatus | diurnal |
| lizards | Draco_blanfordii | Draco_blanfordii | diurnal |
| lizards | Draco_cornutus | Draco_cornutus | diurnal |
| lizards | Draco_cyanopterus | Draco_cyanopterus | diurnal |
| lizards | Draco_dussumieri | Draco_dussumieri | diurnal |
| lizards | Draco_fimbriatus | Draco_fimbriatus | diurnal |
| lizards | Draco_formosus | Draco_formosus | diurnal |
| lizards | Draco_haematopogon | Draco_haematopogon | diurnal |
| lizards | Draco_indochinensis | Draco_indochinensis | diurnal |
| lizards | Draco_lineatus | Draco_lineatus | diurnal |
| lizards | Draco_maculatus | Draco_maculatus | diurnal |
| lizards | Draco_maximus | Draco_maximus | diurnal |
| lizards | Draco_melanopogon | Draco_melanopogon | diurnal |
| lizards | Draco_mindanensis | Draco_mindanensis | diurnal |
| lizards | Draco_modiglianii | Draco_modiglianii | diurnal |
| lizards | Draco_obscurus | Draco_obscurus | diurnal |
| lizards | Draco_palawanensis | Draco_palawanensis | diurnal |
| lizards | Draco_quinquefasciatus | Draco_quinquefasciatus | diurnal |
| lizards | Draco_reticulatus | Draco_reticulatus | diurnal |
| lizards | Draco_spilonotus | Draco_spilonotus | diurnal |
| lizards | Draco_spilopterus | Draco_spilopterus | diurnal |
| lizards | Draco_sumatranus | Draco_sumatranus | diurnal |
| lizards | Draco_taeniopterus | Draco_taeniopterus | diurnal |
| lizards | Draco_timoriensis | Draco_timorensis | diurnal |
| lizards | Draco_volans | Draco_volans | diurnal |
| lizards | Dravidogecko_anamallensis | Hemidactylus_anamallensis | nocturnal |
| lizards | Dryadosaura_nordestina | Dryadosaura_nordestina | diurnal |
| lizards | Ebenavia_inunguis | Ebenavia_inunguis | cathemeral |
| lizards | Ebenavia_maintimainty | Ebenavia_maintimainty | nocturnal |
| lizards | Echinosaura_horrida | Echinosaura_horrida | cathemeral |
| lizards | Egernia_cunninghami | Egernia_cunninghami | diurnal |
| lizards | Egernia_depressa | Egernia_depressa | diurnal |

| | | | |
|---|---|---|---|
| lizards | Egernia_douglasi | Egernia_douglasi | cathemeral |
| lizards | Egernia_formosa | Egernia_formosa | diurnal |
| lizards | Egernia_hosmeri | Egernia_hosmeri | diurnal |
| lizards | Egernia_kingii | Egernia_kingii | diurnal |
| lizards | Egernia_mcpheei | Egernia_mcpheei | diurnal |
| lizards | Egernia_napoleonis | Egernia_napoleonis | diurnal |
| lizards | Egernia_rugosa | Egernia_rugosa | diurnal |
| lizards | Egernia_saxatilis | Egernia_saxatilis | diurnal |
| lizards | Egernia_stokesii | Egernia_stokesii | diurnal |
| lizards | Egernia_striolata | Egernia_striolata | diurnal |
| lizards | Elasmodactylus_tetensis | Elasmodactylus_tetensis | nocturnal |
| lizards | Elasmodactylus_tuberculosus | Elasmodactylus_tuberculosus | nocturnal |
| lizards | Elgaria_coerulea | Elgaria_coerulea | diurnal |
| lizards | Elgaria_kingii | Elgaria_kingii | diurnal |
| lizards | Elgaria_multicarinata | Elgaria_multicarinata | cathemeral |
| lizards | Elgaria_panamintina | Elgaria_panamintina | cathemeral |
| lizards | Elgaria_paucicarinata | Elgaria_paucicarinata | diurnal |
| lizards | Emoia_adspersa | Emoia_adspersa | diurnal |
| lizards | Emoia_aenea | Emoia_aenea | diurnal |
| lizards | Emoia_arnoensis | Emoia_arnoensis | diurnal |
| lizards | Emoia_atrocostata | Emoia_atrocostata | diurnal |
| lizards | Emoia_aurulenta | Emoia_aurulenta | diurnal |
| lizards | Emoia_baudini | Emoia_baudini | diurnal |
| lizards | Emoia_bismarckensis | Emoia_bismarckensis | diurnal |
| lizards | Emoia_boettgeri | Emoia_boettgeri | diurnal |
| lizards | Emoia_bogerti | Emoia_bogerti | diurnal |
| lizards | Emoia_brongersmai | Emoia_brongersmai | diurnal |
| lizards | Emoia_caeruleocauda | Emoia_caeruleocauda | diurnal |
| lizards | Emoia_campbelli | Emoia_campbelli | diurnal |
| lizards | Emoia_coggeri | Emoia_coggeri | diurnal |
| lizards | Emoia_concolor | Emoia_concolor | diurnal |
| lizards | Emoia_cyanogaster | Emoia_cyanogaster | diurnal |
| lizards | Emoia_cyanura | Emoia_cyanura | diurnal |
| lizards | Emoia_guttata | Emoia_guttata | diurnal |

| | | | |
|---|---|---|---|
| lizards | Emoia_impar | Emoia_impar | diurnal |
| lizards | Emoia_isolata | Emoia_isolata | diurnal |
| lizards | Emoia_jakati | Emoia_jakati | diurnal |
| lizards | Emoia_jamur | Emoia_jamur | diurnal |
| lizards | Emoia_kordoana | Emoia_kordoana | diurnal |
| lizards | Emoia_longicauda | Emoia_longicauda | diurnal |
| lizards | Emoia_loyaltiensis | Emoia_loyaltiensis | diurnal |
| lizards | Emoia_maculata | Emoia_maculata | diurnal |
| lizards | Emoia_mivarti | Emoia_mivarti | diurnal |
| lizards | Emoia_nativitatis | Emoia_nativittatis | diurnal |
| lizards | Emoia_nigra | Emoia_nigra | diurnal |
| lizards | Emoia_nigromarginata | Emoia_nigromarginata | diurnal |
| lizards | Emoia_obscura | Emoia_obscura | diurnal |
| lizards | Emoia_oriva | Emoia_oriva | diurnal |
| lizards | Emoia_pallidiceps | Emoia_pallidiceps | diurnal |
| lizards | Emoia_parkeri | Emoia_parkeri | diurnal |
| lizards | Emoia_physicae | Emoia_physicae | diurnal |
| lizards | Emoia_physicina | Emoia_physicina | diurnal |
| lizards | Emoia_ponapea | Emoia_ponapea | diurnal |
| lizards | Emoia_popei | Emoia_popei | diurnal |
| lizards | Emoia_pseudocyanura | Emoia_pseudocyanura | diurnal |
| lizards | Emoia_reimschiisseli | Emoia_reimschiisseli | diurnal |
| lizards | Emoia_rennellensis | Emoia_rennellensis | diurnal |
| lizards | Emoia_ruficauda | Emoia_ruficauda | diurnal |
| lizards | Emoia_samoensis | Emoia_samoensis | diurnal |
| lizards | Emoia_sanfordi | Emoia_sanfordi | diurnal |
| lizards | Emoia_schmidti | Emoia_schmidti | diurnal |
| lizards | Emoia_slevini | Emoia_slevini | diurnal |
| lizards | Emoia_submetallica | Emoia_submetallica | diurnal |
| lizards | Emoia_tetrataenia | Emoia_tetrataenia | diurnal |
| lizards | Emoia_tongana | Emoia_tongana | diurnal |
| lizards | Emoia_tropidolepis | Emoia_tropidolepis | diurnal |
| lizards | Emoia_trossula | Emoia_trossula | diurnal |
| lizards | Emoia_veracunda | Emoia_veracunda | diurnal |

| | | | |
|---|---|---|---|
| lizards | Enyalioides_cofanorum | Enyalioides_cofanorum | diurnal |
| lizards | Enyalioides_heterolepis | Enyalioides_heterolepis | diurnal |
| lizards | Enyalioides_laticeps | Enyalioides_laticeps | diurnal |
| lizards | Enyalioides_palpebralis | Enyalioides_palpebralis | diurnal |
| lizards | Enyalioides_praestabilis | Enyalioides_praestabilis | diurnal |
| lizards | Enyalioides_rubrigularis | Enyalioides_rubrigularis | diurnal |
| lizards | Enyalius_bibronii | Enyalius_bibronii | diurnal |
| lizards | Enyalius_bilineatus | Enyalius_bilineatus | diurnal |
| lizards | Enyalius_brasiliensis | Enyalius_brasiliensis | cathemeral |
| lizards | Enyalius_catenatus | Enyalius_catenatus | diurnal |
| lizards | Enyalius_iheringii | Enyalius_iheringii | diurnal |
| lizards | Enyalius_leechii | Enyalius_leechii | diurnal |
| lizards | Enyalius_perditus | Enyalius_perditus | diurnal |
| lizards | Epibator_greeri | Lioscincus_greeri | diurnal |
| lizards | Epibator_nigrofasciolatus | Lioscincus_nigrofasciolatum | diurnal |
| lizards | Eremias_acutirostris | Eremias_acutirostris | diurnal |
| lizards | Eremias_argus | Eremias_argus | diurnal |
| lizards | Eremias_arguta | Eremias_arguta | diurnal |
| lizards | Eremias_brenchleyi | Eremias_brenchleyi | diurnal |
| lizards | Eremias_buechneri | Eremias_buechneri | diurnal |
| lizards | Eremias_grammica | Eremias_grammica | diurnal |
| lizards | Eremias_intermedia | Eremias_intermedia | diurnal |
| lizards | Eremias_lineolata | Eremias_lineolata | diurnal |
| lizards | Eremias_multiocellata | Eremias_multiocellata | diurnal |
| lizards | Eremias_nigrocellata | Eremias_nigrocellata | diurnal |
| lizards | Eremias_nikolskii | Eremias_nikolskii | diurnal |
| lizards | Eremias_persica | Eremias_persica | diurnal |
| lizards | Eremias_pleskei | Eremias_pleskei | diurnal |
| lizards | Eremias_przewalskii | Eremias_przewalskii | diurnal |
| lizards | Eremias_regeli | Eremias_regeli | diurnal |
| lizards | Eremias_scripta | Eremias_scripta | diurnal |
| lizards | Eremias_strauchi | Eremias_strauchi | diurnal |
| lizards | Eremias_stummeri | Eremias_stummeri | diurnal |
| lizards | Eremias_suphani | Eremias_suphani | diurnal |

| | | | |
|---|---|---|---|
| lizards | Eremias_velox | Eremias_velox | diurnal |
| lizards | Eremias_vermiculata | Eremias_vermiculata | diurnal |
| lizards | Eremias_yarkandensis | Eremias_yarkandensis | diurnal |
| lizards | Eremiascincus_douglasi | Eremiascincus_douglasi | cathemeral |
| lizards | Eremiascincus_fasciolatus | Eremiascincus_fasciolatus | nocturnal |
| lizards | Eremiascincus_intermedius | Eremiascincus_intermedius | nocturnal |
| lizards | Eremiascincus_isolepis | Eremiascincus_isolepis | cathemeral |
| lizards | Eremiascincus_musivus | Eremiascincus_musivus | nocturnal |
| lizards | Eremiascincus_pallidus | Eremiascincus_pallidus | nocturnal |
| lizards | Eremiascincus_pardalis | Eremiascincus_pardalis | cathemeral |
| lizards | Eremiascincus_phantasmus | Eremiascincus_phantasmus | nocturnal |
| lizards | Eremiascincus_richardsonii | Eremiascincus_richardsonii | nocturnal |
| lizards | Eroticoscincus_graciloides | Eroticoscincus_graciloides | cathemeral |
| lizards | Eublepharis_angramainyu | Eublepharis_angramainyu | nocturnal |
| lizards | Eublepharis_fuscus | Eublepharis_fuscus | nocturnal |
| lizards | Eublepharis_hardwickii | Eublepharis_hardwickii | nocturnal |
| lizards | Eublepharis_macularius | Eublepharis_macularius | nocturnal |
| lizards | Eublepharis_turcmenicus | Eublepharis_turcmenicus | nocturnal |
| lizards | Eugongylus_albofasciolatus | Eugongylus_albofasciolatus | cathemeral |
| lizards | Eugongylus_rufescens | Eugongylus_rufescens | cathemeral |
| lizards | Eulamprus_heatwolei | Eulamprus_heatwolei | diurnal |
| lizards | Eulamprus_kosciuskoi | Eulamprus_kosciuskoi | diurnal |
| lizards | Eulamprus_leuraensis | Eulamprus_leuraensis | diurnal |
| lizards | Eulamprus_quoyii | Eulamprus_quoyii | diurnal |
| lizards | Eulamprus_tympanum | Eulamprus_tympanum | diurnal |
| lizards | Euleptes_europaea | Euleptes_europaea | nocturnal |
| lizards | Eumeces_algeriensis | Eumeces_algeriensis | diurnal |
| lizards | Eumeces_blythianus | Eumeces_blythianus | diurnal |
| lizards | Eumeces_schneiderii | Eumeces_schneideri | diurnal |
| lizards | Eumecia_anchietae | Eumecia_anchietae | diurnal |
| lizards | Eurolophosaurus_amathites | Eurolophosaurus_amathites | diurnal |
| lizards | Eurolophosaurus_nanuzae | Eurolophosaurus_nanuzae | diurnal |
| lizards | Eurydactylodes_agricolae | Eurydactylodes_agricolae | diurnal |
| lizards | Eurydactylodes_occidentalis | Eurydactylodes_occidentalis | diurnal |

| | | | |
|---|---|---|---|
| lizards | Eurydactylodes_symmetricus | Eurydactylodes_symmetricus | cathemeral |
| lizards | Eurydactylodes_vieillardi | Eurydactylodes_vieillardi | diurnal |
| lizards | Eurylepis_taeniolata | Eurylepis_taeniolata | cathemeral |
| lizards | Euspondylus_acutirostris | Euspondylus_acutirostris | diurnal |
| lizards | Eutropis_beddomei | Eutropis_beddomii | diurnal |
| lizards | Eutropis_carinata | Eutropis_carinata | diurnal |
| lizards | Eutropis_clivicola | Eutropis_clivicola | diurnal |
| lizards | Eutropis_cumingi | Eutropis_cumingi | diurnal |
| lizards | Eutropis_floweri | Eutropis_floweri | diurnal |
| lizards | Eutropis_indeprensa | Eutropis_indeprensa | diurnal |
| lizards | Eutropis_longicaudata | Eutropis_longicaudata | diurnal |
| lizards | Eutropis_macularia | Eutropis_macularia | cathemeral |
| lizards | Eutropis_multicarinata | Eutropis_multicarinata | diurnal |
| lizards | Eutropis_multifasciata | Eutropis_multifasciata | diurnal |
| lizards | Eutropis_quadricarinata | Eutropis_quadricarinata | diurnal |
| lizards | Eutropis_rudis | Eutropis_rudis | diurnal |
| lizards | Eutropis_tammanna | Eutropis_tammanna | diurnal |
| lizards | Feylinia_boulengeri | Chabanaudia_boulengeri | cathemeral |
| lizards | Feylinia_grandisquamis | Feylinia_grandisquamis | crepuscular |
| lizards | Flexiseps_melanurus | Amphiglossus_melanurus | cathemeral |
| lizards | Fojia_bumui | Fojia_bumui | diurnal |
| lizards | Furcifer_antimena | Furcifer_antimena | cathemeral |
| lizards | Furcifer_cephalolepis | Furcifer_cephalolepis | diurnal |
| lizards | Furcifer_labordi | Furcifer_labordi | diurnal |
| lizards | Furcifer_lateralis | Furcifer_lateralis | diurnal |
| lizards | Furcifer_minor | Furcifer_minor | diurnal |
| lizards | Furcifer_oustaleti | Furcifer_oustaleti | diurnal |
| lizards | Furcifer_pardalis | Furcifer_pardalis | diurnal |
| lizards | Furcifer_petteri | Furcifer_petteri | diurnal |
| lizards | Furcifer_timoni | Furcifer_timoni | diurnal |
| lizards | Furcifer_verrucosus | Furcifer_verrucosus | diurnal |
| lizards | Gallotia_atlantica | Gallotia_atlantica | diurnal |
| lizards | Gallotia_auaritae | Gallotia_auaritae | diurnal |
| lizards | Gallotia_bravoana | Gallotia_bravoana | diurnal |

| | | | |
|---|---|---|---|
| lizards | Gallotia_caesaris | Gallotia_caesaris | diurnal |
| lizards | Gallotia_galloti | Gallotia_galloti | diurnal |
| lizards | Gallotia_intermedia | Gallotia_intermedia | diurnal |
| lizards | Gallotia_simonyi | Gallotia_simonyi | diurnal |
| lizards | Gallotia_stehlini | Gallotia_stehlini | diurnal |
| lizards | Gambelia_copeii | Gambelia_copeii | diurnal |
| lizards | Gambelia_sila | Gambelia_sila | diurnal |
| lizards | Gambelia_wislizenii | Gambelia_wislizenii | diurnal |
| lizards | Garthia_gaudichaudii | Garthia_gaudichaudii | nocturnal |
| lizards | Gastropholis_prasina | Gastropholis_prasina | diurnal |
| lizards | Geckolepis_maculata | Geckolepis_maculata | nocturnal |
| lizards | Geckolepis_polylepis | Geckolepis_polylepis | nocturnal |
| lizards | Geckolepis_typica | Geckolepis_typica | nocturnal |
| lizards | Gehyra_australis | Gehyra_australis | nocturnal |
| lizards | Gehyra_baliola | Gehyra_baliola | nocturnal |
| lizards | Gehyra_borroloola | Gehyra_borroloola | nocturnal |
| lizards | Gehyra_catenata | Gehyra_catenata | nocturnal |
| lizards | Gehyra_dubia | Gehyra_dubia | nocturnal |
| lizards | Gehyra_georgpotthasti | Gehyra_georgpotthasti | nocturnal |
| lizards | Gehyra_koira | Gehyra_koira | nocturnal |
| lizards | Gehyra_lacerata | Gehyra_lacerata | nocturnal |
| lizards | Gehyra_minuta | Gehyra_minuta | nocturnal |
| lizards | Gehyra_montium | Gehyra_montium | nocturnal |
| lizards | Gehyra_moritzi | Gehyra_moritzi | nocturnal |
| lizards | Gehyra_mutilata | Gehyra_mutilata | nocturnal |
| lizards | Gehyra_nana | Gehyra_nana | nocturnal |
| lizards | Gehyra_oceanica | Gehyra_oceanica | nocturnal |
| lizards | Gehyra_pilbara | Gehyra_pilbara | nocturnal |
| lizards | Gehyra_punctata | Gehyra_punctata | nocturnal |
| lizards | Gehyra_purpurascens | Gehyra_purpurascens | nocturnal |
| lizards | Gehyra_robusta | Gehyra_robusta | nocturnal |
| lizards | Gehyra_variegata | Gehyra_variegata | nocturnal |
| lizards | Gehyra_versicolor | Gehyra_versicolor | cathemeral |
| lizards | Gehyra_vorax | Gehyra_vorax | cathemeral |

| | | | |
|---|---|---|---|
| lizards | Gehyra_xenopus | Gehyra_xenopus | nocturnal |
| lizards | Gekko_badenii | Gekko_badenii | nocturnal |
| lizards | Gekko_canaensis | Gekko_canaensis | nocturnal |
| lizards | Gekko_chinensis | Gekko_chinensis | nocturnal |
| lizards | Gekko_ernstkelleri | Gekko_ernstkelleri | nocturnal |
| lizards | Gekko_gecko | Gekko_gecko | nocturnal |
| lizards | Gekko_grossmanni | Gekko_grossmanni | nocturnal |
| lizards | Gekko_hokouensis | Gekko_hokouensis | nocturnal |
| lizards | Gekko_horsfieldii | Ptychozoon_horsfieldii | nocturnal |
| lizards | Gekko_intermedium | Ptychozoon_intermedium | nocturnal |
| lizards | Gekko_iskandari | Luperosaurus_iskandari | nocturnal |
| lizards | Gekko_japonicus | Gekko_japonicus | nocturnal |
| lizards | Gekko_kikuchii | Gekko_kikuchii | nocturnal |
| lizards | Gekko_kuhli | Ptychozoon_kuhli | nocturnal |
| lizards | Gekko_lionotum | Ptychozoon_lionotum | nocturnal |
| lizards | Gekko_mindorensis | Gekko_mindorensis | nocturnal |
| lizards | Gekko_monarchus | Gekko_monarchus | nocturnal |
| lizards | Gekko_nicobarensis | Ptychozoon_nicobarensis | nocturnal |
| lizards | Gekko_nutaphandi | Gekko_nutaphandi | nocturnal |
| lizards | Gekko_palmatus | Gekko_palmatus | nocturnal |
| lizards | Gekko_petricolus | Gekko_petricolus | cathemeral |
| lizards | Gekko_rhacophorus | Ptychozoon_rhacophorus | nocturnal |
| lizards | Gekko_russelltraini | Gekko_russelltraini | nocturnal |
| lizards | Gekko_scientiadventura | Gekko_scientiadventura | nocturnal |
| lizards | Gekko_shibatai | Gekko_shibatai | nocturnal |
| lizards | Gekko_siamensis | Gekko_siamensis | nocturnal |
| lizards | Gekko_smithii | Gekko_smithii | nocturnal |
| lizards | Gekko_subpalmatus | Gekko_subpalmatus | nocturnal |
| lizards | Gekko_swinhonis | Gekko_swinhonis | nocturnal |
| lizards | Gekko_takouensis | Gekko_takouensis | nocturnal |
| lizards | Gekko_truongi | Gekko_truongi | nocturnal |
| lizards | Gekko_vittatus | Gekko_vittatus | nocturnal |
| lizards | Gekko_yakuensis | Gekko_yakuensis | nocturnal |
| lizards | Geomyersia_glabra | Geomyersia_glabra | nocturnal |

| | | | |
|---|---|---|---|
| lizards | Gerrhonotus_infernalis | Gerrhonotus_infernalis | cathemeral |
| lizards | Gerrhonotus_liocephalus | Gerrhonotus_liocephalus | diurnal |
| lizards | Gerrhonotus_lugoi | Gerrhonotus_lugoi | nocturnal |
| lizards | Gerrhonotus_ophiurus | Gerrhonotus_ophiurus | diurnal |
| lizards | Gerrhonotus_parvus | Gerrhonotus_parvus | diurnal |
| lizards | Gerrhosaurus_auritus | Gerrhosaurus_auritus | diurnal |
| lizards | Gerrhosaurus_bulsi | Gerrhosaurus_bulsi | diurnal |
| lizards | Gerrhosaurus_flavigularis | Gerrhosaurus_flavigularis | diurnal |
| lizards | Gerrhosaurus_multilineatus | Gerrhosaurus_multilineatus | diurnal |
| lizards | Gerrhosaurus_nigrolineatus | Gerrhosaurus_nigrolineatus | diurnal |
| lizards | Gerrhosaurus_skoogi | Gerrhosaurus_skoogi | diurnal |
| lizards | Gerrhosaurus_typicus | Gerrhosaurus_typicus | cathemeral |
| lizards | Glaphyromorphus_fuscicaudis | Glaphyromorphus_fuscicaudis | cathemeral |
| lizards | Glaphyromorphus_pumilus | Glaphyromorphus_pumilus | cathemeral |
| lizards | Glaphyromorphus_punctulatus | Glaphyromorphus_punctulatus | cathemeral |
| lizards | Glaucomastix_abaetensis | Ameivula_abaetensis | diurnal |
| lizards | Glaucomastix_littoralis | Ameivula_littoralis | diurnal |
| lizards | Goggia_braacki | Goggia_braacki | nocturnal |
| lizards | Goggia_gemmula | Goggia_gemmula | nocturnal |
| lizards | Goggia_hewitti | Goggia_hewitti | nocturnal |
| lizards | Goggia_lineata | Goggia_lineata | nocturnal |
| lizards | Gonatodes_albogularis | Gonatodes_albogularis | diurnal |
| lizards | Gonatodes_alexandermendesi | Gonatodes_alexandermendesi | diurnal |
| lizards | Gonatodes_annularis | Gonatodes_annularis | diurnal |
| lizards | Gonatodes_antillensis | Gonatodes_antillensis | nocturnal |
| lizards | Gonatodes_caudiscutatus | Gonatodes_caudiscutatus | diurnal |
| lizards | Gonatodes_ceciliae | Gonatodes_ceciliae | diurnal |
| lizards | Gonatodes_concinnatus | Gonatodes_concinnatus | diurnal |
| lizards | Gonatodes_daudini | Gonatodes_daudini | diurnal |
| lizards | Gonatodes_hasemani | Gonatodes_hasemani | diurnal |
| lizards | Gonatodes_humeralis | Gonatodes_humeralis | diurnal |
| lizards | Gonatodes_ocellatus | Gonatodes_ocellatus | diurnal |
| lizards | Gonatodes_petersi | Gonatodes_petersi | diurnal |
| lizards | Gonatodes_superciliaris | Gonatodes_superciliaris | diurnal |

| | | | |
|---|---|---|---|
| lizards | Gonatodes_taniae | Gonatodes_taniae | diurnal |
| lizards | Gonatodes_vittatus | Gonatodes_vittatus | diurnal |
| lizards | Gongylomorphus_bojerii | Gongylomorphus_bojerii | diurnal |
| lizards | Goniurosaurus_araneus | Goniurosaurus_araneus | nocturnal |
| lizards | Goniurosaurus_catbaensis | Goniurosaurus_catbaensis | nocturnal |
| lizards | Goniurosaurus_hainanensis | Goniurosaurus_hainanensis | nocturnal |
| lizards | Goniurosaurus_kuroiwae | Goniurosaurus_kuroiwae | nocturnal |
| lizards | Goniurosaurus_luii | Goniurosaurus_luii | nocturnal |
| lizards | Goniurosaurus_orientalis | Goniurosaurus_orientalis | nocturnal |
| lizards | Goniurosaurus_splendens | Goniurosaurus_splendens | nocturnal |
| lizards | Goniurosaurus_toyamai | Goniurosaurus_toyamai | nocturnal |
| lizards | Goniurosaurus_yamashinae | Goniurosaurus_yamashinae | nocturnal |
| lizards | Goniurosaurus_yingdeensis | Goniurosaurus_yingdeensis | nocturnal |
| lizards | Gonocephalus_bellii | Gonocephalus_bellii | diurnal |
| lizards | Gonocephalus_bornensis | Gonocephalus_bornensis | diurnal |
| lizards | Gonocephalus_chamaeleontinus | Gonocephalus_chamaeleontinus | diurnal |
| lizards | Gonocephalus_doriae | Gonocephalus_doriae | diurnal |
| lizards | Gonocephalus_grandis | Gonocephalus_grandis | diurnal |
| lizards | Gonocephalus_kuhlii | Gonocephalus_kuhlii | diurnal |
| lizards | Gonocephalus_lacunosus | Gonocephalus_lacunosus | diurnal |
| lizards | Gonocephalus_liogaster | Gonocephalus_liogaster | diurnal |
| lizards | Gonocephalus_semperi | Gonocephalus_semperi | diurnal |
| lizards | Gonocephalus_sophiae | Gonocephalus_sophiae | diurnal |
| lizards | Gowidon_longirostris | Lophognathus_longirostris | diurnal |
| lizards | Gymnodactylus_amarali | Gymnodactylus_amarali | nocturnal |
| lizards | Gymnodactylus_darwinii | Gymnodactylus_darwinii | nocturnal |
| lizards | Gymnodactylus_geckoides | Gymnodactylus_geckoides | cathemeral |
| lizards | Gymnophthalmus_speciosus | Gymnophthalmus_speciosus | diurnal |
| lizards | Gymnophthalmus_underwoodi | Gymnophthalmus_underwoodi | diurnal |
| lizards | Gymnophthalmus_vanzoi | Gymnophthalmus_vanzoi | diurnal |
| lizards | Haemodracon_riebeckii | Haemodracon_riebeckii | cathemeral |
| lizards | Haemodracon_trachyrhinus | Haemodracon_trachyrhinus | nocturnal |
| lizards | Harpesaurus_borneensis | Harpesaurus_borneensis | diurnal |
| lizards | Harrisoniascincus_zia | Harrisoniascincus_zia | diurnal |

| | | | |
|---|---|---|---|
| lizards | Heliobolus_lugubris | Heliobolus_lugubris | diurnal |
| lizards | Heliobolus_spekii | Heliobolus_spekii | diurnal |
| lizards | Hellenolacerta_graeca | Hellenolacerta_graeca | diurnal |
| lizards | Heloderma_horridum | Heloderma_horridum | cathemeral |
| lizards | Heloderma_suspectum | Heloderma_suspectum | cathemeral |
| lizards | Hemicordylus_capensis | Hemicordylus_capensis | diurnal |
| lizards | Hemicordylus_nebulosus | Hemicordylus_nebulosus | diurnal |
| lizards | Hemidactylus_albofasciatus | Hemidactylus_albofasciatus | nocturnal |
| lizards | Hemidactylus_alkiyumii | Hemidactylus_alkiyumii | nocturnal |
| lizards | Hemidactylus_angulatus | Hemidactylus_angulatus | nocturnal |
| lizards | Hemidactylus_aquilonius | Hemidactylus_aquilonius | nocturnal |
| lizards | Hemidactylus_beninensis | Hemidactylus_beninensis | nocturnal |
| lizards | Hemidactylus_bowringii | Hemidactylus_bowringii | nocturnal |
| lizards | Hemidactylus_brasilianus | Hemidactylus_brasilianus | nocturnal |
| lizards | Hemidactylus_brookii | Hemidactylus_brookii | nocturnal |
| lizards | Hemidactylus_coalescens | Hemidactylus_coalescens | nocturnal |
| lizards | Hemidactylus_craspedotus | Hemidactylus_craspedotus | nocturnal |
| lizards | Hemidactylus_depressus | Hemidactylus_depressus | nocturnal |
| lizards | Hemidactylus_dracaenacolus | Hemidactylus_dracaenacolus | nocturnal |
| lizards | Hemidactylus_echinus | Hemidactylus_echinus | nocturnal |
| lizards | Hemidactylus_fasciatus | Hemidactylus_fasciatus | nocturnal |
| lizards | Hemidactylus_festivus | Hemidactylus_festivus | nocturnal |
| lizards | Hemidactylus_flaviviridis | Hemidactylus_flaviviridis | cathemeral |
| lizards | Hemidactylus_frenatus | Hemidactylus_frenatus | nocturnal |
| lizards | Hemidactylus_garnotii | Hemidactylus_garnotii | nocturnal |
| lizards | Hemidactylus_gleadowi | Hemidactylus_gleadowi | nocturnal |
| lizards | Hemidactylus_hajarensis | Hemidactylus_hajarensis | nocturnal |
| lizards | Hemidactylus_homoeolepis | Hemidactylus_homoeolepis | nocturnal |
| lizards | Hemidactylus_hunae | Hemidactylus_hunae | nocturnal |
| lizards | Hemidactylus_imbricatus | Hemidactylus_imbricatus | nocturnal |
| lizards | Hemidactylus_inexpectatus | Hemidactylus_inexpectatus | nocturnal |
| lizards | Hemidactylus_inintellectus | Hemidactylus_inintellectus | nocturnal |
| lizards | Hemidactylus_isolepis | Hemidactylus_isolepis | nocturnal |
| lizards | Hemidactylus_kamdemtohami | Hemidactylus_kamdemtohami | diurnal |

| | | | |
|---|---|---|---|
| lizards | Hemidactylus_lankae | Hemidactylus_lankae | nocturnal |
| lizards | Hemidactylus_lavadeserticus | Hemidactylus_lavadeserticus | nocturnal |
| lizards | Hemidactylus_leschenaultii | Hemidactylus_leschenaultii | cathemeral |
| lizards | Hemidactylus_longicephalus | Hemidactylus_longicephalus | nocturnal |
| lizards | Hemidactylus_mabouia | Hemidactylus_mabouia | cathemeral |
| lizards | Hemidactylus_macropholis | Hemidactylus_macropholis | nocturnal |
| lizards | Hemidactylus_maculatus | Hemidactylus_maculatus | nocturnal |
| lizards | Hemidactylus_mercatorius | Hemidactylus_mercatorius | nocturnal |
| lizards | Hemidactylus_minutus | Hemidactylus_minutus | nocturnal |
| lizards | Hemidactylus_mrimaensis | Hemidactylus_mrimaensis | cathemeral |
| lizards | Hemidactylus_muriceus | Hemidactylus_muriceus | cathemeral |
| lizards | Hemidactylus_palaichthus | Hemidactylus_palaichthus | nocturnal |
| lizards | Hemidactylus_parvimaculatus | Hemidactylus_parvimaculatus | nocturnal |
| lizards | Hemidactylus_persicus | Hemidactylus_persicus | nocturnal |
| lizards | Hemidactylus_platycephalus | Hemidactylus_platycephalus | nocturnal |
| lizards | Hemidactylus_platyurus | Hemidactylus_platyurus | cathemeral |
| lizards | Hemidactylus_richardsonii | Hemidactylus_richardsonii | nocturnal |
| lizards | Hemidactylus_robustus | Hemidactylus_robustus | nocturnal |
| lizards | Hemidactylus_ruspolii | Hemidactylus_ruspolii | nocturnal |
| lizards | Hemidactylus_sataraensis | Hemidactylus_sataraensis | nocturnal |
| lizards | Hemidactylus_scabriceps | Hemidactylus_scabriceps | nocturnal |
| lizards | Hemidactylus_squamulatus | Hemidactylus_squamulatus | nocturnal |
| lizards | Hemidactylus_stejnegeri | Hemidactylus_stejnegeri | cathemeral |
| lizards | Hemidactylus_tanganicus | Hemidactylus_tanganicus | nocturnal |
| lizards | Hemidactylus_treutleri | Hemidactylus_treutleri | nocturnal |
| lizards | Hemidactylus_triedrus | Hemidactylus_triedrus | nocturnal |
| lizards | Hemidactylus_tropidolepis | Hemidactylus_tropidolepis | nocturnal |
| lizards | Hemidactylus_turcicus | Hemidactylus_turcicus | nocturnal |
| lizards | Hemiergis_decresiensis | Hemiergis_decresiensis | diurnal |
| lizards | Hemiergis_gracilipes | Hemiergis_gracilipes | diurnal |
| lizards | Hemiergis_initialis | Hemiergis_initialis | cathemeral |
| lizards | Hemiergis_millewae | Hemiergis_millewae | cathemeral |
| lizards | Hemiergis_peronii | Hemiergis_peronii | cathemeral |
| lizards | Hemiergis_talbingoensis | Hemiergis_talbingoensis | diurnal |

| | | | |
|---|---|---|---|
| lizards | Hemiphyllodactylus_aurantiacus | Hemiphyllodactylus_aurantiacus | nocturnal |
| lizards | Hemiphyllodactylus_banaensis | Hemiphyllodactylus_banaensis | nocturnal |
| lizards | Hemiphyllodactylus_chiangmaiensis | Hemiphyllodactylus_chiangmaiensis | nocturnal |
| lizards | Hemiphyllodactylus_ganoklonis | Hemiphyllodactylus_ganoklonis | nocturnal |
| lizards | Hemiphyllodactylus_insularis | Hemiphyllodactylus_insularis | nocturnal |
| lizards | Hemiphyllodactylus_kiziriani | Hemiphyllodactylus_kiziriani | nocturnal |
| lizards | Hemiphyllodactylus_tehtarik | Hemiphyllodactylus_tehtarik | nocturnal |
| lizards | Hemiphyllodactylus_typus | Hemiphyllodactylus_typus | nocturnal |
| lizards | Hemitheconyx_caudicinctus | Hemitheconyx_caudicinctus | nocturnal |
| lizards | Hemitheconyx_taylori | Hemitheconyx_taylori | nocturnal |
| lizards | Heremites_auratus | Trachylepis_aurata | diurnal |
| lizards | Heremites_septemtaeniatus | Trachylepis_septemtaeniata | diurnal |
| lizards | Heremites_vittatus | Trachylepis_vittata | cathemeral |
| lizards | Hesperoedura_reticulata | Hesperoedura_reticulata | nocturnal |
| lizards | Heteronotia_atra | Heteronotia_atra | nocturnal |
| lizards | Heteronotia_binoei | Heteronotia_binoei | nocturnal |
| lizards | Heteronotia_planiceps | Heteronotia_planiceps | nocturnal |
| lizards | Heteronotia_spelea | Heteronotia_spelea | nocturnal |
| lizards | Holaspis_guentheri | Holaspis_guentheri | diurnal |
| lizards | Holaspis_laevis | Holaspis_laevis | diurnal |
| lizards | Holbrookia_approximans | Holbrookia_approximans | diurnal |
| lizards | Holbrookia_elegans | Holbrookia_elegans | diurnal |
| lizards | Holbrookia_lacerata | Holbrookia_lacerata | diurnal |
| lizards | Holbrookia_maculata | Holbrookia_maculata | diurnal |
| lizards | Holbrookia_propinqua | Holbrookia_propinqua | diurnal |
| lizards | Holcosus_chaitzami | Holcosus_chaitzami | diurnal |
| lizards | Holcosus_festivus | Holcosus_festivus | diurnal |
| lizards | Holcosus_niceforoi | Holcosus_niceforoi | diurnal |
| lizards | Holcosus_quadrilineatus | Holcosus_quadrilineatus | diurnal |
| lizards | Holcosus_undulatus | Holcosus_undulatus | diurnal |
| lizards | Holodactylus_africanus | Holodactylus_africanus | nocturnal |
| lizards | Homonota_borellii | Homonota_borellii | nocturnal |
| lizards | Homonota_darwinii | Homonota_darwinii | nocturnal |
| lizards | Homonota_taragui | Homonota_taragui | nocturnal |

| | | | |
|---|---|---|---|
| lizards | Homonota_underwoodi | Homonota_underwoodi | nocturnal |
| lizards | Homonota_uruguayensis | Homonota_uruguayensis | cathemeral |
| lizards | Homonota_whitii | Homonota_whitii | nocturnal |
| lizards | Homopholis_fasciata | Homopholis_fasciata | nocturnal |
| lizards | Homopholis_mulleri | Homopholis_mulleri | nocturnal |
| lizards | Homopholis_walbergii | Homopholis_walbergii | cathemeral |
| lizards | Hoplocercus_spinosus | Hoplocercus_spinosus | diurnal |
| lizards | Hoplodactylus_duvaucelii | Hoplodactylus_duvaucelii | nocturnal |
| lizards | Hydrosaurus_amboinensis | Hydrosaurus_amboinensis | diurnal |
| lizards | Hydrosaurus_pustulatus | Hydrosaurus_pustulatus | diurnal |
| lizards | Hypsilurus_godeffroyi | Hypsilurus_godeffroyi | diurnal |
| lizards | Hypsilurus_macrolepis | Hypsilurus_macrolepis | diurnal |
| lizards | Hypsilurus_modestus | Hypsilurus_modestus | diurnal |
| lizards | Hypsilurus_schoedei | Hypsilurus_schoedei | diurnal |
| lizards | Iberolacerta_aranica | Iberolacerta_aranica | diurnal |
| lizards | Iberolacerta_aurelioi | Iberolacerta_aurelioi | diurnal |
| lizards | Iberolacerta_bonnali | Iberolacerta_bonnali | diurnal |
| lizards | Iberolacerta_cyreni | Iberolacerta_cyreni | diurnal |
| lizards | Iberolacerta_galani | Iberolacerta_galani | diurnal |
| lizards | Iberolacerta_horvathi | Iberolacerta_horvathi | diurnal |
| lizards | Iberolacerta_martinezricai | Iberolacerta_martinezricai | diurnal |
| lizards | Iberolacerta_monticola | Iberolacerta_monticola | diurnal |
| lizards | Ichnotropis_capensis | Ichnotropis_capensis | diurnal |
| lizards | Ichnotropis_grandiceps | Ichnotropis_grandiceps | diurnal |
| lizards | Iguana_delicatissima | Iguana_delicatissima | diurnal |
| lizards | Iguana_iguana | Iguana_iguana | diurnal |
| lizards | Insulasaurus_arborens | Insulasaurus_arborens | diurnal |
| lizards | Intellagama_lesueurii | Intellagama_lesueurii | diurnal |
| lizards | Iphisa_elegans | Iphisa_elegans | diurnal |
| lizards | Iranolacerta_brandtii | Iranolacerta_brandtii | diurnal |
| lizards | Isopachys_gyldenstolpei | Isopachys_gyldenstolpei | cathemeral |
| lizards | Janetaescincus_braueri | Janetaescincus_braueri | cathemeral |
| lizards | Japalura_andersoniana | Japalura_andersoniana | diurnal |
| lizards | Japalura_kumaonensis | Japalura_kumaonensis | diurnal |

| | | | |
|---|---|---|---|
| lizards | Japalura_major | Japalura_major | diurnal |
| lizards | Japalura_variegata | Japalura_variegata | diurnal |
| lizards | Kanakysaurus_viviparus | Kanakysaurus_viviparus | cathemeral |
| lizards | Karusasaurus_jordani | Karusasaurus_jordani | diurnal |
| lizards | Karusasaurus_polyzonus | Karusasaurus_polyzonus | diurnal |
| lizards | Kentropyx_altamazonica | Kentropyx_altamazonica | diurnal |
| lizards | Kentropyx_borckiana | Kentropyx_borckiana | diurnal |
| lizards | Kentropyx_calcarata | Kentropyx_calcarata | diurnal |
| lizards | Kentropyx_lagartija | Kentropyx_lagartija | diurnal |
| lizards | Kentropyx_paulensis | Kentropyx_paulensis | diurnal |
| lizards | Kentropyx_pelviceps | Kentropyx_pelviceps | diurnal |
| lizards | Kentropyx_striata | Kentropyx_striata | diurnal |
| lizards | Kentropyx_vanzoi | Kentropyx_vanzoi | diurnal |
| lizards | Kentropyx_viridistriga | Kentropyx_viridistriga | diurnal |
| lizards | Kinyongia_adolfifriderici | Kinyongia_adolfifriderici | diurnal |
| lizards | Kinyongia_boehmei | Kinyongia_boehmei | diurnal |
| lizards | Kinyongia_excubitor | Kinyongia_excubitor | diurnal |
| lizards | Kinyongia_fischeri | Kinyongia_fischeri | diurnal |
| lizards | Kinyongia_oxyrhina | Kinyongia_oxyrhina | diurnal |
| lizards | Kinyongia_tavetana | Kinyongia_tavetana | diurnal |
| lizards | Kinyongia_tenuis | Kinyongia_tenuis | diurnal |
| lizards | Kinyongia_uluguruensis | Kinyongia_uluguruensis | diurnal |
| lizards | Kinyongia_uthmoelleri | Kinyongia_uthmoelleri | diurnal |
| lizards | Lacerta_agilis | Lacerta_agilis | diurnal |
| lizards | Lacerta_bilineata | Lacerta_bilineata | diurnal |
| lizards | Lacerta_media | Lacerta_media | diurnal |
| lizards | Lacerta_pamphylica | Lacerta_pamphylica | diurnal |
| lizards | Lacerta_schreiberi | Lacerta_schreiberi | diurnal |
| lizards | Lacerta_strigata | Lacerta_strigata | diurnal |
| lizards | Lacerta_trilineata | Lacerta_trilineata | diurnal |
| lizards | Lacerta_viridis | Lacerta_viridis | diurnal |
| lizards | Lacertaspis_lepesmei | Lacertaspis_lepesmei | diurnal |
| lizards | Lacertoides_pardalis | Lacertoides_pardalis | diurnal |
| lizards | Laemanctus_longipes | Laemanctus_longipes | diurnal |

| | | | |
|---|---|---|---|
| lizards | Laemanctus_serratus | Laemanctus_serratus | cathemeral |
| lizards | Lamprolepis_smaragdina | Lamprolepis_smaragdina | diurnal |
| lizards | Lampropholis_adonis | Lampropholis_adonis | diurnal |
| lizards | Lampropholis_amicula | Lampropholis_amicula | diurnal |
| lizards | Lampropholis_caligula | Lampropholis_caligula | diurnal |
| lizards | Lampropholis_coggeri | Lampropholis_coggeri | diurnal |
| lizards | Lampropholis_colossus | Lampropholis_colossus | diurnal |
| lizards | Lampropholis_delicata | Lampropholis_delicata | diurnal |
| lizards | Lampropholis_elongata | Lampropholis_elongata | diurnal |
| lizards | Lampropholis_guichenoti | Lampropholis_guichenoti | diurnal |
| lizards | Lampropholis_mirabilis | Lampropholis_mirabilis | diurnal |
| lizards | Lampropholis_robertsi | Lampropholis_robertsi | diurnal |
| lizards | Lankascincus_deignani | Lankascincus_deignani | diurnal |
| lizards | Lankascincus_fallax | Lankascincus_fallax | diurnal |
| lizards | Lankascincus_gansi | Lankascincus_gansi | diurnal |
| lizards | Lankascincus_sripadensis | Lankascincus_sripadensis | diurnal |
| lizards | Lankascincus_taprobanensis | Lankascincus_taprobanensis | diurnal |
| lizards | Lankascincus_taylori | Lankascincus_taylori | diurnal |
| lizards | Lanthanotus_borneensis | Lanthanotus_borneensis | nocturnal |
| lizards | Larutia_larutensis | Larutia_larutensis | diurnal |
| lizards | Larutia_miodactyla | Larutia_miodactyla | diurnal |
| lizards | Larutia_trifasciata | Larutia_trifasciata | diurnal |
| lizards | Latastia_johnstonii | Latastia_johnstonii | diurnal |
| lizards | Latastia_longicaudata | Latastia_longicaudata | diurnal |
| lizards | Laudakia_agrorensis | Laudakia_agrorensis | diurnal |
| lizards | Laudakia_melanura | Laudakia_melanura | diurnal |
| lizards | Laudakia_nupta | Laudakia_nupta | diurnal |
| lizards | Laudakia_tuberculata | Laudakia_tuberculata | diurnal |
| lizards | Leiocephalus_barahonensis | Leiocephalus_barahonensis | diurnal |
| lizards | Leiocephalus_carinatus | Leiocephalus_carinatus | diurnal |
| lizards | Leiocephalus_cubensis | Leiocephalus_cubensis | diurnal |
| lizards | Leiocephalus_inaguae | Leiocephalus_inaguae | diurnal |
| lizards | Leiocephalus_lunatus | Leiocephalus_lunatus | diurnal |
| lizards | Leiocephalus_macropus | Leiocephalus_macropus | diurnal |

| | | | |
|---|---|---|---|
| lizards | Leiocephalus_melanochlorus | Leiocephalus_melanochlorus | diurnal |
| lizards | Leiocephalus_onaneyi | Leiocephalus_onaneyi | diurnal |
| lizards | Leiocephalus_personatus | Leiocephalus_personatus | diurnal |
| lizards | Leiocephalus_psammodromus | Leiocephalus_psammodromus | diurnal |
| lizards | Leiocephalus_raviceps | Leiocephalus_raviceps | diurnal |
| lizards | Leiocephalus_schreibersii | Leiocephalus_schreibersii | diurnal |
| lizards | Leiocephalus_semilineatus | Leiocephalus_semilineatus | diurnal |
| lizards | Leiocephalus_stictigaster | Leiocephalus_stictigaster | diurnal |
| lizards | Leiolepis_belliana | Leiolepis_belliana | diurnal |
| lizards | Leiolepis_guentherpetersi | Leiolepis_guentherpetersi | diurnal |
| lizards | Leiolepis_guttata | Leiolepis_guttata | diurnal |
| lizards | Leiolepis_reevesii | Leiolepis_reevesii | diurnal |
| lizards | Leiolepis_triploida | Leiolepis_triploida | diurnal |
| lizards | Leiosaurus_bellii | Leiosaurus_bellii | diurnal |
| lizards | Leiosaurus_catamarcensis | Leiosaurus_catamarcensis | diurnal |
| lizards | Lepidoblepharis_heyerorum | Lepidoblepharis_heyerorum | cathemeral |
| lizards | Lepidoblepharis_sanctaemartae | Lepidoblepharis_sanctaemartae | diurnal |
| lizards | Lepidoblepharis_xanthostigma | Lepidoblepharis_xanthostigma | diurnal |
| lizards | Lepidodactylus_buleli | Lepidodactylus_buleli | nocturnal |
| lizards | Lepidodactylus_euaensis | Lepidodactylus_euaensis | nocturnal |
| lizards | Lepidodactylus_gardineri | Lepidodactylus_gardineri | nocturnal |
| lizards | Lepidodactylus_guppyi | Lepidodactylus_guppyi | nocturnal |
| lizards | Lepidodactylus_listeri | Lepidodactylus_listeri | nocturnal |
| lizards | Lepidodactylus_lugubris | Lepidodactylus_lugubris | nocturnal |
| lizards | Lepidodactylus_magnus | Lepidodactylus_magnus | nocturnal |
| lizards | Lepidodactylus_manni | Lepidodactylus_manni | nocturnal |
| lizards | Lepidodactylus_moestus | Lepidodactylus_moestus | nocturnal |
| lizards | Lepidodactylus_pumilus | Lepidodactylus_pumilus | nocturnal |
| lizards | Lepidodactylus_ranauensis | Lepidodactylus_ranauensis | nocturnal |
| lizards | Lepidodactylus_vanuatuensis | Lepidodactylus_vanuatuensis | nocturnal |
| lizards | Lepidodactylus_yami | Lepidodactylus_yami | nocturnal |
| lizards | Lepidophyma_flavimaculatum | Lepidophyma_flavimaculatum | cathemeral |
| lizards | Lepidophyma_gaigeae | Lepidophyma_gaigeae | nocturnal |
| lizards | Lepidophyma_micropholis | Lepidophyma_micropholis | cathemeral |

| | | | |
|---|---|---|---|
| lizards | Lepidophyma_pajapanensis | Lepidophyma_pajapanensis | nocturnal |
| lizards | Lepidophyma_smithii | Lepidophyma_smithii | cathemeral |
| lizards | Lepidophyma_tuxtlae | Lepidophyma_tuxtlae | nocturnal |
| lizards | Leposternon_polystegum | Amphisbaena_polystegum | diurnal |
| lizards | Leptosiaphos_blochmanni | Leptosiaphos_blochmanni | diurnal |
| lizards | Leptosiaphos_graueri | Leptosiaphos_graueri | diurnal |
| lizards | Leptosiaphos_kilimensis | Leptosiaphos_kilimensis | cathemeral |
| lizards | Leptosiaphos_meleagris | Leptosiaphos_meleagris | diurnal |
| lizards | Lerista_allanae | Lerista_allanae | nocturnal |
| lizards | Lerista_arenicola | Lerista_arenicola | diurnal |
| lizards | Lerista_baynesi | Lerista_baynesi | cathemeral |
| lizards | Lerista_bipes | Lerista_bipes | cathemeral |
| lizards | Lerista_borealis | Lerista_borealis | cathemeral |
| lizards | Lerista_bougainvillii | Lerista_bougainvillii | cathemeral |
| lizards | Lerista_christinae | Lerista_christinae | cathemeral |
| lizards | Lerista_cinerea | Lerista_cinerea | nocturnal |
| lizards | Lerista_connivens | Lerista_connivens | cathemeral |
| lizards | Lerista_desertorum | Lerista_desertorum | cathemeral |
| lizards | Lerista_distinguenda | Lerista_distinguenda | cathemeral |
| lizards | Lerista_edwardsae | Lerista_edwardsae | cathemeral |
| lizards | Lerista_elegans | Lerista_elegans | cathemeral |
| lizards | Lerista_emmotti | Lerista_emmotti | cathemeral |
| lizards | Lerista_fragilis | Lerista_fragilis | cathemeral |
| lizards | Lerista_frosti | Lerista_frosti | diurnal |
| lizards | Lerista_gerrardii | Lerista_gerrardii | cathemeral |
| lizards | Lerista_griffini | Lerista_griffini | cathemeral |
| lizards | Lerista_humphriesi | Lerista_humphriesi | nocturnal |
| lizards | Lerista_ips | Lerista_ips | cathemeral |
| lizards | Lerista_karlschmidti | Lerista_karlschmidti | nocturnal |
| lizards | Lerista_labialis | Lerista_labialis | cathemeral |
| lizards | Lerista_lineata | Lerista_lineata | cathemeral |
| lizards | Lerista_lineopunctulata | Lerista_lineopunctulata | cathemeral |
| lizards | Lerista_macropisthopus | Lerista_macropisthopus | cathemeral |
| lizards | Lerista_microtis | Lerista_microtis | diurnal |

| | | | |
|---|---|---|---|
| lizards | Lerista_muelleri | Lerista_muelleri | diurnal |
| lizards | Lerista_petersoni | Lerista_petersoni | cathemeral |
| lizards | Lerista_picturata | Lerista_picturata | cathemeral |
| lizards | Lerista_planiventralis | Lerista_planiventralis | cathemeral |
| lizards | Lerista_praepedita | Lerista_praepedita | cathemeral |
| lizards | Lerista_punctatovittata | Lerista_punctatovittata | cathemeral |
| lizards | Lerista_storri | Lerista_storri | nocturnal |
| lizards | Lerista_stylis | Lerista_stylis | nocturnal |
| lizards | Lerista_timida | Lerista_rhodonoides | diurnal |
| lizards | Lerista_uniduo | Lerista_uniduo | cathemeral |
| lizards | Lerista_vittata | Lerista_vittata | nocturnal |
| lizards | Lerista_walkeri | Lerista_walkeri | cathemeral |
| lizards | Lerista_wilkinsi | Lerista_wilkinsi | nocturnal |
| lizards | Lerista_xanthura | Lerista_xanthura | cathemeral |
| lizards | Lialis_burtonis | Lialis_burtonis | cathemeral |
| lizards | Lialis_jicari | Lialis_jicari | diurnal |
| lizards | Liburnascincus_mundivensis | Liburnascincus_mundivensis | diurnal |
| lizards | Liburnascincus_scirtetis | Liburnascincus_scirtetis | diurnal |
| lizards | Liolaemus_abdalai | Liolaemus_abdalai | diurnal |
| lizards | Liolaemus_alticolor | Liolaemus_alticolor | diurnal |
| lizards | Liolaemus_andinus | Liolaemus_andinus | diurnal |
| lizards | Liolaemus_anomalus | Liolaemus_anomalus | diurnal |
| lizards | Liolaemus_arambarensis | Liolaemus_arambarensis | diurnal |
| lizards | Liolaemus_archeforus | Liolaemus_archeforus | diurnal |
| lizards | Liolaemus_atacamensis | Liolaemus_atacamensis | diurnal |
| lizards | Liolaemus_audituvelatus | Liolaemus_audituvelatus | diurnal |
| lizards | Liolaemus_baguali | Liolaemus_baguali | diurnal |
| lizards | Liolaemus_bellii | Liolaemus_bellii | diurnal |
| lizards | Liolaemus_bibronii | Liolaemus_bibronii | diurnal |
| lizards | Liolaemus_bitaeniatus | Liolaemus_bitaeniatus | diurnal |
| lizards | Liolaemus_boulengeri | Liolaemus_boulengeri | diurnal |
| lizards | Liolaemus_burmeisteri | Liolaemus_burmeisteri | diurnal |
| lizards | Liolaemus_cazianiae | Liolaemus_cazianiae | diurnal |
| lizards | Liolaemus_chacoensis | Liolaemus_chacoensis | diurnal |

| | | | |
|---|---|---|---|
| lizards | Liolaemus_chaltin | Liolaemus_chaltin | diurnal |
| lizards | Liolaemus_chavin | Liolaemus_chavin | diurnal |
| lizards | Liolaemus_chiliensis | Liolaemus_chiliensis | diurnal |
| lizards | Liolaemus_chillanensis | Liolaemus_chillanensis | diurnal |
| lizards | Liolaemus_coeruleus | Liolaemus_coeruleus | diurnal |
| lizards | Liolaemus_confusus | Liolaemus_confusus | diurnal |
| lizards | Liolaemus_constanzae | Liolaemus_constanzae | diurnal |
| lizards | Liolaemus_crepuscularis | Liolaemus_crepuscularis | diurnal |
| lizards | Liolaemus_curis | Liolaemus_curis | diurnal |
| lizards | Liolaemus_cyanogaster | Liolaemus_cyanogaster | diurnal |
| lizards | Liolaemus_darwinii | Liolaemus_darwinii | diurnal |
| lizards | Liolaemus_ditadai | Liolaemus_ditadai | diurnal |
| lizards | Liolaemus_dorbignyi | Liolaemus_dorbignyi | diurnal |
| lizards | Liolaemus_elongatus | Liolaemus_elongatus | diurnal |
| lizards | Liolaemus_espinozai | Liolaemus_espinozai | diurnal |
| lizards | Liolaemus_fabiani | Liolaemus_fabiani | diurnal |
| lizards | Liolaemus_fitzgeraldi | Liolaemus_fitzgeraldi | diurnal |
| lizards | Liolaemus_foxi | Liolaemus_foxi | diurnal |
| lizards | Liolaemus_fuscus | Liolaemus_fuscus | diurnal |
| lizards | Liolaemus_gallardoi | Liolaemus_gallardoi | diurnal |
| lizards | Liolaemus_goetschi | Liolaemus_goetschi | diurnal |
| lizards | Liolaemus_gracilis | Liolaemus_gracilis | diurnal |
| lizards | Liolaemus_gravenhorstii | Liolaemus_gravenhorstii | diurnal |
| lizards | Liolaemus_halonastes | Liolaemus_halonastes | diurnal |
| lizards | Liolaemus_hatcheri | Liolaemus_hatcheri | diurnal |
| lizards | Liolaemus_huacahuasicus | Liolaemus_huacahuasicus | diurnal |
| lizards | Liolaemus_incaicus | Liolaemus_incaicus | diurnal |
| lizards | Liolaemus_irregularis | Liolaemus_irregularis | diurnal |
| lizards | Liolaemus_jamesi | Liolaemus_jamesi | diurnal |
| lizards | Liolaemus_kingii | Liolaemus_kingii | diurnal |
| lizards | Liolaemus_kolengh | Liolaemus_kolengh | diurnal |
| lizards | Liolaemus_koslowskyi | Liolaemus_koslowskyi | diurnal |
| lizards | Liolaemus_kriegi | Liolaemus_kriegi | diurnal |
| lizards | Liolaemus_lemniscatus | Liolaemus_lemniscatus | diurnal |

| | | | |
|---|---|---|---|
| lizards | Liolaemus_leopardinus | Liolaemus_leopardinus | diurnal |
| lizards | Liolaemus_lineomaculatus | Liolaemus_lineomaculatus | diurnal |
| lizards | Liolaemus_lutzae | Liolaemus_lutzae | diurnal |
| lizards | Liolaemus_magellanicus | Liolaemus_magellanicus | diurnal |
| lizards | Liolaemus_melanops | Liolaemus_melanops | diurnal |
| lizards | Liolaemus_millcayac | Liolaemus_millcayac | diurnal |
| lizards | Liolaemus_monticola | Liolaemus_monticola | diurnal |
| lizards | Liolaemus_morenoi | Liolaemus_morenoi | diurnal |
| lizards | Liolaemus_multicolor | Liolaemus_multicolor | diurnal |
| lizards | Liolaemus_multimaculatus | Liolaemus_multimaculatus | diurnal |
| lizards | Liolaemus_neuquensis | Liolaemus_neuquensis | diurnal |
| lizards | Liolaemus_nigromaculatus | Liolaemus_nigromaculatus | diurnal |
| lizards | Liolaemus_nigroviridis | Liolaemus_nigroviridis | diurnal |
| lizards | Liolaemus_nitidus | Liolaemus_nitidus | diurnal |
| lizards | Liolaemus_olongasta | Liolaemus_olongasta | diurnal |
| lizards | Liolaemus_ornatus | Liolaemus_ornatus | diurnal |
| lizards | Liolaemus_pachacutec | Liolaemus_pachacutec | diurnal |
| lizards | Liolaemus_pagaburoi | Liolaemus_pagaburoi | diurnal |
| lizards | Liolaemus_parvus | Liolaemus_parvus | diurnal |
| lizards | Liolaemus_patriciaiturrae | Liolaemus_patriciaiturrae | diurnal |
| lizards | Liolaemus_paulinae | Liolaemus_paulinae | diurnal |
| lizards | Liolaemus_petrophilus | Liolaemus_petrophilus | diurnal |
| lizards | Liolaemus_pictus | Liolaemus_pictus | diurnal |
| lizards | Liolaemus_platei | Liolaemus_platei | diurnal |
| lizards | Liolaemus_pleopholis | Liolaemus_pleopholis | diurnal |
| lizards | Liolaemus_poecilochromus | Liolaemus_poecilochromus | diurnal |
| lizards | Liolaemus_pseudoanomalus | Liolaemus_pseudoanomalus | diurnal |
| lizards | Liolaemus_puna | Liolaemus_puna | diurnal |
| lizards | Liolaemus_puritamensis | Liolaemus_puritamensis | diurnal |
| lizards | Liolaemus_ramirezae | Liolaemus_ramirezae | diurnal |
| lizards | Liolaemus_riojanus | Liolaemus_riojanus | diurnal |
| lizards | Liolaemus_robertoi | Liolaemus_robertoi | diurnal |
| lizards | Liolaemus_rosenmanni | Liolaemus_rosenmanni | diurnal |
| lizards | Liolaemus_rothi | Liolaemus_rothi | diurnal |

| | | | |
|---|---|---|---|
| lizards | Liolaemus_ruibali | Liolaemus_ruibali | diurnal |
| lizards | Liolaemus_sagei | Liolaemus_sagei | diurnal |
| lizards | Liolaemus_sanjuanensis | Liolaemus_sanjuanensis | diurnal |
| lizards | Liolaemus_sarmientoi | Liolaemus_sarmientoi | diurnal |
| lizards | Liolaemus_saxatilis | Liolaemus_saxatilis | diurnal |
| lizards | Liolaemus_scapularis | Liolaemus_scapularis | diurnal |
| lizards | Liolaemus_schroederi | Liolaemus_schroederi | diurnal |
| lizards | Liolaemus_septentrionalis | Liolaemus_septentrionalis | diurnal |
| lizards | Liolaemus_signifer | Liolaemus_signifer | diurnal |
| lizards | Liolaemus_silvanae | Liolaemus_silvanae | diurnal |
| lizards | Liolaemus_stolzmanni | Liolaemus_stolzmanni | diurnal |
| lizards | Liolaemus_tandiliensis | Liolaemus_tandiliensis | diurnal |
| lizards | Liolaemus_tari | Liolaemus_tari | diurnal |
| lizards | Liolaemus_tenuis | Liolaemus_tenuis | diurnal |
| lizards | Liolaemus_torresi | Liolaemus_torresi | diurnal |
| lizards | Liolaemus_tristis | Liolaemus_tristis | diurnal |
| lizards | Liolaemus_uspallatensis | Liolaemus_uspallatensis | diurnal |
| lizards | Liolaemus_valdesianus | Liolaemus_valdesianus | diurnal |
| lizards | Liolaemus_vallecurensis | Liolaemus_vallecurensis | diurnal |
| lizards | Liolaemus_velosoi | Liolaemus_velosoi | diurnal |
| lizards | Liolaemus_wari | Liolaemus_wari | diurnal |
| lizards | Liolaemus_wiegmannii | Liolaemus_wiegmannii | diurnal |
| lizards | Liolaemus_xanthoviridis | Liolaemus_xanthoviridis | diurnal |
| lizards | Liolaemus_yanalcu | Liolaemus_yanalcu | diurnal |
| lizards | Liolaemus_zapallarensis | Liolaemus_zapallarensis | diurnal |
| lizards | Liolaemus_zullyae | Liolaemus_zullyae | diurnal |
| lizards | Liopholis_guthega | Liopholis_guthega | diurnal |
| lizards | Liopholis_inornata | Liopholis_inornata | cathemeral |
| lizards | Liopholis_kintorei | Liopholis_kintorei | cathemeral |
| lizards | Liopholis_modesta | Liopholis_modesta | diurnal |
| lizards | Liopholis_montana | Liopholis_montana | diurnal |
| lizards | Liopholis_multiscutata | Liopholis_multiscutata | diurnal |
| lizards | Liopholis_pulchra | Liopholis_pulchra | diurnal |
| lizards | Liopholis_slateri | Liopholis_slateri | cathemeral |

| | | | |
|---|---|---|---|
| lizards | Liopholis_striata | Liopholis_striata | cathemeral |
| lizards | Liopholis_whitii | Liopholis_whitii | diurnal |
| lizards | Lioscincus_steindachneri | Lioscincus_steindachneri | diurnal |
| lizards | Lipinia_auriculata | Lipinia_auriculata | diurnal |
| lizards | Lipinia_inexpectata | Lipinia_inexpectata | cathemeral |
| lizards | Lipinia_leptosoma | Lipinia_leptosoma | diurnal |
| lizards | Lipinia_macrotympanum | Lipinia_macrotympanum | cathemeral |
| lizards | Lipinia_pulchella | Lipinia_pulchella | diurnal |
| lizards | Lipinia_quadrivittata | Lipinia_quadrivittata | diurnal |
| lizards | Lipinia_rabori | Lipinia_rabori | diurnal |
| lizards | Lipinia_rouxi | Lipinia_rouxi | diurnal |
| lizards | Lipinia_vittigera | Lipinia_vittigera | diurnal |
| lizards | Lissolepis_coventryi | Lissolepis_coventryi | diurnal |
| lizards | Lobulia_brongersmai | Lobulia_brongersmai | diurnal |
| lizards | Lobulia_elegans | Lobulia_elegans | diurnal |
| lizards | Lophognathus_gilberti | Lophognathus_gilberti | diurnal |
| lizards | Lophosaurus_boydii | Hypsilurus_boydii | diurnal |
| lizards | Lophosaurus_dilophus | Hypsilurus_dilophus | diurnal |
| lizards | Lophosaurus_spinipes | Hypsilurus_spinipes | diurnal |
| lizards | Loxopholis_guianense | Leposoma_guianense | cathemeral |
| lizards | Loxopholis_hexalepis | Leposoma_hexalepis | diurnal |
| lizards | Loxopholis_osvaldoi | Leposoma_osvaldoi | diurnal |
| lizards | Loxopholis_parietalis | Leposoma_parietale | diurnal |
| lizards | Loxopholis_percarinatum | Leposoma_percarinatum | diurnal |
| lizards | Loxopholis_rugiceps | Leposoma_rugiceps | diurnal |
| lizards | Loxopholis_snethlageae | Leposoma_snethlageae | diurnal |
| lizards | Loxopholis_southi | Leposoma_southi | diurnal |
| lizards | Lubuya_ivensii | Trachylepis_ivensii | diurnal |
| lizards | Lucasium_alboguttatum | Lucasium_alboguttatum | nocturnal |
| lizards | Lucasium_bungabinna | Lucasium_bungabinna | nocturnal |
| lizards | Lucasium_byrnei | Lucasium_byrnei | nocturnal |
| lizards | Lucasium_damaeum | Lucasium_damaeum | nocturnal |
| lizards | Lucasium_maini | Lucasium_maini | nocturnal |
| lizards | Lucasium_occultum | Lucasium_occultum | nocturnal |

| | | | |
|---|---|---|---|
| lizards | Lucasium_squarrosum | Lucasium_squarrosum | nocturnal |
| lizards | Lucasium_steindachneri | Lucasium_steindachneri | nocturnal |
| lizards | Lucasium_wombeyi | Lucasium_wombeyi | nocturnal |
| lizards | Luperosaurus_corfieldi | Luperosaurus_corfieldi | nocturnal |
| lizards | Luperosaurus_cumingii | Luperosaurus_cumingii | nocturnal |
| lizards | Lygisaurus_curtus | Lygisaurus_curtus | diurnal |
| lizards | Lygisaurus_foliorum | Lygisaurus_foliorum | diurnal |
| lizards | Lygisaurus_laevis | Lygisaurus_laevis | diurnal |
| lizards | Lygisaurus_macfarlani | Lygisaurus_macfarlani | diurnal |
| lizards | Lygisaurus_rococo | Lygisaurus_rococo | diurnal |
| lizards | Lygisaurus_zuma | Lygisaurus_zuma | diurnal |
| lizards | Lygodactylus_angolensis | Lygodactylus_angolensis | diurnal |
| lizards | Lygodactylus_angularis | Lygodactylus_angularis | cathemeral |
| lizards | Lygodactylus_bivittis | Lygodactylus_bivittis | diurnal |
| lizards | Lygodactylus_bradfieldi | Lygodactylus_bradfieldi | diurnal |
| lizards | Lygodactylus_capensis | Lygodactylus_capensis | diurnal |
| lizards | Lygodactylus_conraui | Lygodactylus_conraui | diurnal |
| lizards | Lygodactylus_depressus | Lygodactylus_depressus | diurnal |
| lizards | Lygodactylus_gravis | Lygodactylus_gravis | diurnal |
| lizards | Lygodactylus_grotei | Lygodactylus_grotei | diurnal |
| lizards | Lygodactylus_gutturalis | Lygodactylus_gutturalis | diurnal |
| lizards | Lygodactylus_insularis | Lygodactylus_insularis | diurnal |
| lizards | Lygodactylus_keniensis | Lygodactylus_keniensis | diurnal |
| lizards | Lygodactylus_kimhowelli | Lygodactylus_kimhowelli | diurnal |
| lizards | Lygodactylus_klugei | Lygodactylus_klugei | diurnal |
| lizards | Lygodactylus_lawrencei | Lygodactylus_lawrencei | diurnal |
| lizards | Lygodactylus_manni | Lygodactylus_manni | diurnal |
| lizards | Lygodactylus_mirabilis | Lygodactylus_mirabilis | diurnal |
| lizards | Lygodactylus_mombasicus | Lygodactylus_mombasicus | diurnal |
| lizards | Lygodactylus_pauliani | Lygodactylus_pauliani | diurnal |
| lizards | Lygodactylus_picturatus | Lygodactylus_picturatus | diurnal |
| lizards | Lygodactylus_pictus | Lygodactylus_pictus | diurnal |
| lizards | Lygodactylus_scheffleri | Lygodactylus_scheffleri | diurnal |
| lizards | Lygodactylus_scorteccii | Lygodactylus_scorteccii | diurnal |

| | | | |
|---|---|---|---|
| lizards | Lygodactylus_somalicus | Lygodactylus_somalicus | diurnal |
| lizards | Lygodactylus_thomensis | Lygodactylus_thomensis | diurnal |
| lizards | Lygodactylus_tuberosus | Lygodactylus_tuberosus | diurnal |
| lizards | Lygodactylus_verticillatus | Lygodactylus_verticillatus | diurnal |
| lizards | Lygodactylus_williamsi | Lygodactylus_williamsi | diurnal |
| lizards | Lygosoma_angeli | Lygosoma_angeli | nocturnal |
| lizards | Lygosoma_corpulentum | Lygosoma_corpulentum | cathemeral |
| lizards | Lygosoma_koratense | Lygosoma_koratense | nocturnal |
| lizards | Lygosoma_quadrupes | Lygosoma_quadrupes | cathemeral |
| lizards | Lyriocephalus_scutatus | Lyriocephalus_scutatus | diurnal |
| lizards | Mabuya_dominicana | Mabuya_dominicana | diurnal |
| lizards | Mabuya_mabouya | Mabuya_mabouya | diurnal |
| lizards | Macropholidus_annectens | Macropholidus_annectens | diurnal |
| lizards | Macropholidus_ataktolepis | Macropholidus_ataktolepis | diurnal |
| lizards | Macropholidus_ruthveni | Macropholidus_ruthveni | diurnal |
| lizards | Madascincus_melanopleura | Madascincus_melanopleura | diurnal |
| lizards | Madascincus_mouroundavae | Madascincus_mouroundavae | diurnal |
| lizards | Madascincus_stumpffi | Madascincus_stumpffi | nocturnal |
| lizards | Malayodracon_robinsonii | Gonocephalus_robinsonii | diurnal |
| lizards | Manciola_guaporicola | Manciola_guaporicola | diurnal |
| lizards | Mantheyus_phuwuanensis | Mantheyus_phuwuanensis | cathemeral |
| lizards | Maracaiba_meridensis | Maracaiba_meridensis | diurnal |
| lizards | Marisora_alliacea | Marisora_alliacea | diurnal |
| lizards | Marisora_brachypoda | Marisora_brachypoda | diurnal |
| lizards | Marisora_pergravis | Alinea_pergravis | diurnal |
| lizards | Marisora_unimarginata | Marisora_unimarginata | diurnal |
| lizards | Marmorosphax_boulinda | Marmorosphax_boulinda | cathemeral |
| lizards | Marmorosphax_montana | Marmorosphax_montana | cathemeral |
| lizards | Marmorosphax_taom | Marmorosphax_taom | cathemeral |
| lizards | Marmorosphax_tricolor | Marmorosphax_tricolor | cathemeral |
| lizards | Matoatoa_spannringi | Matoatoa_spannringi | nocturnal |
| lizards | Matobosaurus_maltzahni | Matobosaurus_maltzahni | diurnal |
| lizards | Matobosaurus_validus | Matobosaurus_validus | diurnal |
| lizards | Mediodactylus_amictopholis | Mediodactylus_amictopholis | diurnal |

| | | | |
|---|---|---|---|
| lizards | Mediodactylus_brachykolon | Mediodactylus_brachykolon | nocturnal |
| lizards | Mediodactylus_heterocercus | Mediodactylus_heterocercus | nocturnal |
| lizards | Mediodactylus_heteropholis | Mediodactylus_heteropholis | nocturnal |
| lizards | Mediodactylus_kotschyi | Mediodactylus_kotschyi | cathemeral |
| lizards | Mediodactylus_russowii | Mediodactylus_russowii | cathemeral |
| lizards | Mediodactylus_spinicauda | Mediodactylus_spinicauda | nocturnal |
| lizards | Melanoseps_ater | Melanoseps_ater | diurnal |
| lizards | Melanoseps_emmrichi | Melanoseps_emmrichi | cathemeral |
| lizards | Melanoseps_loveridgei | Melanoseps_loveridgei | diurnal |
| lizards | Melanoseps_pygmaeus | Melanoseps_pygmaeus | diurnal |
| lizards | Melanoseps_rondoensis | Melanoseps_rondoensis | diurnal |
| lizards | Menetia_alanae | Menetia_alanae | diurnal |
| lizards | Menetia_greyii | Menetia_greyii | diurnal |
| lizards | Menetia_maini | Menetia_maini | diurnal |
| lizards | Meroles_anchietae | Meroles_anchietae | diurnal |
| lizards | Meroles_ctenodactylus | Meroles_ctenodactylus | diurnal |
| lizards | Meroles_cuneirostris | Meroles_cuneirostris | diurnal |
| lizards | Meroles_knoxii | Meroles_knoxii | diurnal |
| lizards | Meroles_reticulatus | Meroles_reticulatus | diurnal |
| lizards | Meroles_squamulosus | Meroles_squamulosa | diurnal |
| lizards | Meroles_suborbitalis | Meroles_suborbitalis | diurnal |
| lizards | Mesalina_adramitana | Mesalina_adramitana | diurnal |
| lizards | Mesalina_bahaeldini | Mesalina_bahaeldini | diurnal |
| lizards | Mesalina_balfouri | Mesalina_balfouri | diurnal |
| lizards | Mesalina_brevirostris | Mesalina_brevirostris | diurnal |
| lizards | Mesalina_guttulata | Mesalina_guttulata | diurnal |
| lizards | Mesalina_martini | Mesalina_martini | diurnal |
| lizards | Mesalina_olivieri | Mesalina_olivieri | diurnal |
| lizards | Mesalina_pasteuri | Mesalina_pasteuri | diurnal |
| lizards | Mesalina_rubropunctata | Mesalina_rubropunctata | diurnal |
| lizards | Mesalina_simoni | Mesalina_simoni | diurnal |
| lizards | Mesalina_watsonana | Mesalina_watsonana | diurnal |
| lizards | Micrablepharus_atticolus | Micrablepharus_atticolus | diurnal |
| lizards | Micrablepharus_maximiliani | Micrablepharus_maximiliani | diurnal |

| | | | |
|---|---|---|---|
| lizards | Microauris_aurantolabium | Calotes_aurantolabium | diurnal |
| lizards | Microgecko_helenae | Microgecko_helenae | nocturnal |
| lizards | Microgecko_persicus | Microgecko_persicus | nocturnal |
| lizards | Microlophus_albemarlensis | Microlophus_albemarlensis | diurnal |
| lizards | Microlophus_atacamensis | Microlophus_atacamensis | diurnal |
| lizards | Microlophus_delanonis | Microlophus_delanonis | diurnal |
| lizards | Microlophus_habelii | Microlophus_habelii | diurnal |
| lizards | Microlophus_heterolepis | Microlophus_heterolepis | diurnal |
| lizards | Microlophus_occipitalis | Microlophus_occipitalis | diurnal |
| lizards | Microlophus_pacificus | Microlophus_pacificus | diurnal |
| lizards | Microlophus_peruvianus | Microlophus_peruvianus | diurnal |
| lizards | Microlophus_quadrivittatus | Microlophus_quadrivittatus | diurnal |
| lizards | Microlophus_tarapacensis | Microlophus_tarapacensis | diurnal |
| lizards | Microlophus_theresiae | Microlophus_theresiae | diurnal |
| lizards | Microlophus_theresioides | Microlophus_theresioides | diurnal |
| lizards | Microlophus_thoracicus | Microlophus_thoracicus | diurnal |
| lizards | Mniarogekko_chahoua | Mniarogekko_chahoua | nocturnal |
| lizards | Mniarogekko_jalu | Mniarogekko_jalu | nocturnal |
| lizards | Mochlus_brevicaudis | Mochlus_brevicaudis | diurnal |
| lizards | Mochlus_fernandi | Lepidothyris_fernandi | cathemeral |
| lizards | Mochlus_guineensis | Mochlus_guineensis | diurnal |
| lizards | Mochlus_hinkeli | Lepidothyris_hinkeli | diurnal |
| lizards | Mochlus_pembanus | Lygosoma_pembanum | nocturnal |
| lizards | Mochlus_striatus | Lepidothyris_striatus | cathemeral |
| lizards | Mochlus_sundevallii | Mochlus_sundevalli | diurnal |
| lizards | Mokopirirakau_cryptozoicus | Mokopirirakau_cryptozoicus | nocturnal |
| lizards | Mokopirirakau_granulatus | Mokopirirakau_granulatus | cathemeral |
| lizards | Mokopirirakau_kahutarae | Mokopirirakau_kahutarae | nocturnal |
| lizards | Mokopirirakau_nebulosus | Mokopirirakau_nebulosus | nocturnal |
| lizards | Moloch_horridus | Moloch_horridus | diurnal |
| lizards | Monilesaurus_ellioti | Calotes_ellioti | diurnal |
| lizards | Monilesaurus_rouxii | Calotes_rouxii | diurnal |
| lizards | Monopeltis_capensis | Monopeltis_capensis | diurnal |
| lizards | Morethia_adelaidensis | Morethia_adelaidensis | diurnal |

| | | | |
|---|---|---|---|
| lizards | Morethia_boulengeri | Morethia_boulengeri | diurnal |
| lizards | Morethia_butleri | Morethia_butleri | diurnal |
| lizards | Morethia_lineoocellata | Morethia_lineoocellata | diurnal |
| lizards | Morethia_obscura | Morethia_obscura | diurnal |
| lizards | Morethia_ruficauda | Morethia_ruficauda | diurnal |
| lizards | Morethia_storri | Morethia_storri | diurnal |
| lizards | Morethia_taeniopleura | Morethia_taeniopleura | diurnal |
| lizards | Nactus_eboracensis | Nactus_eboracensis | nocturnal |
| lizards | Nactus_multicarinatus | Nactus_multicarinatus | nocturnal |
| lizards | Nactus_pelagicus | Nactus_pelagicus | nocturnal |
| lizards | Nactus_serpensinsula | Nactus_serpensinsula | nocturnal |
| lizards | Nactus_sphaerodactylodes | Nactus_sphaerodactylodes | nocturnal |
| lizards | Namazonurus_campbelli | Namazonurus_campbelli | diurnal |
| lizards | Namazonurus_lawrenci | Namazonurus_lawrenci | diurnal |
| lizards | Namazonurus_namaquensis | Namazonurus_namaquensis | diurnal |
| lizards | Namazonurus_peersi | Namazonurus_peersi | diurnal |
| lizards | Namazonurus_pustulatus | Namazonurus_pustulatus | diurnal |
| lizards | Nannoscincus_garrulus | Nannoscincus_garrulus | cathemeral |
| lizards | Nannoscincus_gracilis | Nannoscincus_gracilis | cathemeral |
| lizards | Nannoscincus_mariei | Nannoscincus_mariei | cathemeral |
| lizards | Narudasia_festiva | Narudasia_festiva | diurnal |
| lizards | Naultinus_elegans | Naultinus_elegans | diurnal |
| lizards | Naultinus_gemmeus | Naultinus_gemmeus | diurnal |
| lizards | Naultinus_grayii | Naultinus_grayii | diurnal |
| lizards | Naultinus_manukanus | Naultinus_manukanus | diurnal |
| lizards | Naultinus_punctatus | Naultinus_punctatus | diurnal |
| lizards | Naultinus_rudis | Naultinus_rudis | diurnal |
| lizards | Naultinus_stellatus | Naultinus_stellatus | diurnal |
| lizards | Naultinus_tuberculatus | Naultinus_tuberculatus | diurnal |
| lizards | Nephrurus_amyae | Nephrurus_amyae | nocturnal |
| lizards | Nephrurus_asper | Nephrurus_asper | nocturnal |
| lizards | Nephrurus_deleani | Nephrurus_deleani | nocturnal |
| lizards | Nephrurus_laevissimus | Nephrurus_laevissimus | nocturnal |
| lizards | Nephrurus_levis | Nephrurus_levis | nocturnal |

| | | | |
|---|---|---|---|
| lizards | Nephrurus_sheai | Nephrurus_sheai | nocturnal |
| lizards | Nephrurus_stellatus | Nephrurus_stellatus | nocturnal |
| lizards | Nephrurus_vertebralis | Nephrurus_vertebralis | nocturnal |
| lizards | Nephrurus_wheeleri | Nephrurus_wheeleri | nocturnal |
| lizards | Nessia_bipes | Nessia_bipes | cathemeral |
| lizards | Nessia_burtonii | Nessia_burtonii | nocturnal |
| lizards | Nessia_layardi | Nessia_layardi | cathemeral |
| lizards | Nessia_monodactyla | Nessia_monodactyla | nocturnal |
| lizards | Neusticurus_bicarinatus | Neusticurus_bicarinatus | diurnal |
| lizards | Neusticurus_rudis | Neusticurus_rudis | diurnal |
| lizards | Neusticurus_tatei | Neusticurus_tatei | diurnal |
| lizards | Ninurta_coeruleopunctatus | Ninurta_coeruleopunctatus | diurnal |
| lizards | Nothobachia_ablephara | Nothobachia_ablephara | cathemeral |
| lizards | Notomabuya_frenata | Notomabuya_frenata | diurnal |
| lizards | Notoscincus_ornatus | Notoscincus_ornatus | diurnal |
| lizards | Nucras_boulengeri | Nucras_boulengeri | diurnal |
| lizards | Nucras_holubi | Nucras_holubi | diurnal |
| lizards | Nucras_intertexta | Nucras_intertexta | diurnal |
| lizards | Nucras_lalandii | Nucras_lalandii | diurnal |
| lizards | Nucras_ornata | Nucras_ornata | diurnal |
| lizards | Nucras_taeniolata | Nucras_taeniolata | diurnal |
| lizards | Nucras_tessellata | Nucras_tessellata | diurnal |
| lizards | Oedodera_marmorata | Oedodera_marmorata | nocturnal |
| lizards | Oedura_castelnaui | Oedura_castelnaui | nocturnal |
| lizards | Oedura_coggeri | Oedura_coggeri | nocturnal |
| lizards | Oedura_filicipoda | Oedura_filicipoda | nocturnal |
| lizards | Oedura_gemmata | Oedura_gemmata | nocturnal |
| lizards | Oedura_gracilis | Oedura_gracilis | nocturnal |
| lizards | Oedura_jowalbinna | Oedura_jowalbinna | nocturnal |
| lizards | Oedura_marmorata | Oedura_marmorata | nocturnal |
| lizards | Oedura_monilis | Oedura_monilis | nocturnal |
| lizards | Oedura_tryoni | Oedura_tryoni | nocturnal |
| lizards | Oligosoma_acrinasum | Oligosoma_acrinasum | diurnal |
| lizards | Oligosoma_aeneum | Oligosoma_aeneum | diurnal |

| | | | |
|---|---|---|---|
| lizards | Oligosoma_alani | Oligosoma_alani | nocturnal |
| lizards | Oligosoma_burganae | Oligosoma_burganae | diurnal |
| lizards | Oligosoma_chloronoton | Oligosoma_chloronoton | diurnal |
| lizards | Oligosoma_fallai | Oligosoma_fallai | cathemeral |
| lizards | Oligosoma_grande | Oligosoma_grande | diurnal |
| lizards | Oligosoma_hardyi | Oligosoma_hardyi | cathemeral |
| lizards | Oligosoma_homalonotum | Oligosoma_homalonotum | diurnal |
| lizards | Oligosoma_inconspicuum | Oligosoma_inconspicuum | diurnal |
| lizards | Oligosoma_infrapunctatum | Oligosoma_infrapunctatum | diurnal |
| lizards | Oligosoma_judgei | Oligosoma_judgei | diurnal |
| lizards | Oligosoma_levidensum | Oligosoma_levidensum | diurnal |
| lizards | Oligosoma_lineoocellatum | Oligosoma_lineoocellatum | diurnal |
| lizards | Oligosoma_longipes | Oligosoma_longipes | diurnal |
| lizards | Oligosoma_maccanni | Oligosoma_maccanni | diurnal |
| lizards | Oligosoma_macgregori | Oligosoma_macgregori | cathemeral |
| lizards | Oligosoma_microlepis | Oligosoma_microlepis | diurnal |
| lizards | Oligosoma_moco | Oligosoma_moco | diurnal |
| lizards | Oligosoma_nigriplantare | Oligosoma_nigriplantare | diurnal |
| lizards | Oligosoma_notosaurus | Oligosoma_notosaurus | diurnal |
| lizards | Oligosoma_oliveri | Oligosoma_oliveri | cathemeral |
| lizards | Oligosoma_ornatum | Oligosoma_ornatum | cathemeral |
| lizards | Oligosoma_otagense | Oligosoma_otagense | diurnal |
| lizards | Oligosoma_polychroma | Oligosoma_polychroma | diurnal |
| lizards | Oligosoma_smithi | Oligosoma_smithi | diurnal |
| lizards | Oligosoma_stenotis | Oligosoma_stenotis | diurnal |
| lizards | Oligosoma_striatum | Oligosoma_striatum | diurnal |
| lizards | Oligosoma_suteri | Oligosoma_suteri | cathemeral |
| lizards | Oligosoma_whitakeri | Oligosoma_whitakeri | cathemeral |
| lizards | Oligosoma_zelandicum | Oligosoma_zelandicum | diurnal |
| lizards | Omanosaura_cyanura | Omanosaura_cyanura | diurnal |
| lizards | Omanosaura_jayakari | Omanosaura_jayakari | diurnal |
| lizards | Ophiodes_intermedius | Ophiodes_intermedius | cathemeral |
| lizards | Ophiodes_striatus | Ophiodes_striatus | diurnal |
| lizards | Ophiodes_vertebralis | Ophiodes_vertebralis | cathemeral |

| | | | |
|---|---|---|---|
| lizards | Ophiomorus_latastii | Ophiomorus_latastii | cathemeral |
| lizards | Ophiomorus_nuchalis | Ophiomorus_nuchalis | nocturnal |
| lizards | Ophiomorus_punctatissimus | Ophiomorus_punctatissimus | diurnal |
| lizards | Ophiomorus_tridactylus | Ophiomorus_tridactylus | cathemeral |
| lizards | Ophioscincus_ophioscincus | Ophioscincus_ophioscincus | cathemeral |
| lizards | Ophioscincus_truncatus | Ophioscincus_truncatus | nocturnal |
| lizards | Ophisaurus_attenuatus | Ophisaurus_attenuatus | diurnal |
| lizards | Ophisaurus_compressus | Ophisaurus_compressus | diurnal |
| lizards | Ophisaurus_mimicus | Ophisaurus_mimicus | cathemeral |
| lizards | Ophisaurus_ventralis | Ophisaurus_ventralis | cathemeral |
| lizards | Ophisops_elegans | Ophisops_elegans | diurnal |
| lizards | Ophisops_jerdonii | Ophisops_jerdonii | diurnal |
| lizards | Ophisops_leschenaultii | Ophisops_leschenaultii | diurnal |
| lizards | Ophisops_microlepis | Ophisops_microlepis | diurnal |
| lizards | Ophisops_occidentalis | Ophisops_occidentalis | diurnal |
| lizards | Oplurus_cuvieri | Oplurus_cuvieri | diurnal |
| lizards | Oplurus_cyclurus | Oplurus_cyclurus | diurnal |
| lizards | Oplurus_fierinensis | Oplurus_fierinensis | diurnal |
| lizards | Oplurus_grandidieri | Oplurus_grandidieri | diurnal |
| lizards | Oplurus_quadrimaculatus | Oplurus_quadrimaculatus | diurnal |
| lizards | Oplurus_saxicola | Oplurus_saxicola | diurnal |
| lizards | Oreosaurus_achlyens | Riama_achlyens | diurnal |
| lizards | Orraya_occultus | Orraya_occultus | nocturnal |
| lizards | Otocryptis_nigristigma | Otocryptis_nigristigma | diurnal |
| lizards | Otocryptis_wiegmanni | Otocryptis_wiegmanni | diurnal |
| lizards | Ouroborus_cataphractus | Ouroborus_cataphractus | cathemeral |
| lizards | Pachydactylus_acuminatus | Pachydactylus_acuminatus | nocturnal |
| lizards | Pachydactylus_atorquatus | Pachydactylus_atorquatus | nocturnal |
| lizards | Pachydactylus_austeni | Pachydactylus_austeni | nocturnal |
| lizards | Pachydactylus_barnardi | Pachydactylus_barnardi | nocturnal |
| lizards | Pachydactylus_bicolor | Pachydactylus_bicolor | nocturnal |
| lizards | Pachydactylus_boehmei | Pachydactylus_boehmei | nocturnal |
| lizards | Pachydactylus_capensis | Pachydactylus_capensis | nocturnal |
| lizards | Pachydactylus_fasciatus | Pachydactylus_fasciatus | nocturnal |

| | | | |
|---|---|---|---|
| lizards | Pachydactylus_geitje | Pachydactylus_geitje | nocturnal |
| lizards | Pachydactylus_haackei | Pachydactylus_haackei | cathemeral |
| lizards | Pachydactylus_kobosensis | Pachydactylus_kobosensis | nocturnal |
| lizards | Pachydactylus_kochii | Colopus_kochii | nocturnal |
| lizards | Pachydactylus_maculatus | Pachydactylus_maculatus | nocturnal |
| lizards | Pachydactylus_maraisi | Pachydactylus_maraisi | nocturnal |
| lizards | Pachydactylus_mariquensis | Pachydactylus_mariquensis | nocturnal |
| lizards | Pachydactylus_namaquensis | Pachydactylus_namaquensis | cathemeral |
| lizards | Pachydactylus_oculatus | Pachydactylus_oculatus | nocturnal |
| lizards | Pachydactylus_parascutatus | Pachydactylus_parascutatus | nocturnal |
| lizards | Pachydactylus_punctatus | Pachydactylus_punctatus | nocturnal |
| lizards | Pachydactylus_purcelli | Pachydactylus_purcelli | nocturnal |
| lizards | Pachydactylus_rangei | Pachydactylus_rangei | nocturnal |
| lizards | Pachydactylus_rugosus | Pachydactylus_rugosus | nocturnal |
| lizards | Pachydactylus_sansteynae | Pachydactylus_sansteynae | nocturnal |
| lizards | Pachydactylus_scutatus | Pachydactylus_scutatus | nocturnal |
| lizards | Pachydactylus_tsodiloensis | Pachydactylus_tsodiloensis | nocturnal |
| lizards | Pachydactylus_vansoni | Pachydactylus_vansoni | nocturnal |
| lizards | Pachydactylus_wahlbergii | Colopus_wahlbergii | nocturnal |
| lizards | Pachydactylus_weberi | Pachydactylus_weberi | nocturnal |
| lizards | Pachydactylus_werneri | Pachydactylus_werneri | nocturnal |
| lizards | Palleon_lolontany | Palleon_lolontany | diurnal |
| lizards | Pamelaescincus_gardineri | Pamelaescincus_gardineri | cathemeral |
| lizards | Panaspis_breviceps | Panaspis_breviceps | diurnal |
| lizards | Panaspis_cabindae | Panaspis_cabindae | cathemeral |
| lizards | Panaspis_maculicollis | Afroablepharus_maculicollis | diurnal |
| lizards | Panaspis_togoensis | Panaspis_togoensis | diurnal |
| lizards | Panaspis_tristaoi | Panaspis_nimbaensis | diurnal |
| lizards | Paniegekko_madjo | Paniegekko_madjo | nocturnal |
| lizards | Papuascincus_morokanus | Papuascincus_morokanus | diurnal |
| lizards | Papuascincus_stanleyanus | Papuascincus_stanleyanus | diurnal |
| lizards | Paracontias_minimus | Paracontias_minimus | diurnal |
| lizards | Paradelma_orientalis | Paradelma_orientalis | cathemeral |
| lizards | Paragehyra_gabriellae | Paragehyra_gabriellae | nocturnal |

| | | | |
|---|---|---|---|
| lizards | Paralaudakia_bochariensis | Paralaudakia_bochariensis | diurnal |
| lizards | Paralaudakia_caucasia | Paralaudakia_caucasia | diurnal |
| lizards | Paralaudakia_erythrogaster | Paralaudakia_erythrogaster | diurnal |
| lizards | Paralaudakia_himalayana | Paralaudakia_himalayana | diurnal |
| lizards | Paralaudakia_lehmanni | Paralaudakia_lehmanni | diurnal |
| lizards | Paralaudakia_stoliczkana | Paralaudakia_stoliczkana | diurnal |
| lizards | Paroedura_androyensis | Paroedura_androyensis | nocturnal |
| lizards | Paroedura_bastardi | Paroedura_bastardi | nocturnal |
| lizards | Paroedura_gracilis | Paroedura_gracilis | nocturnal |
| lizards | Paroedura_ibityensis | Paroedura_ibityensis | cathemeral |
| lizards | Paroedura_karstophila | Paroedura_karstophila | nocturnal |
| lizards | Paroedura_lohatsara | Paroedura_lohatsara | nocturnal |
| lizards | Paroedura_masobe | Paroedura_masobe | nocturnal |
| lizards | Paroedura_oviceps | Paroedura_oviceps | nocturnal |
| lizards | Paroedura_picta | Paroedura_picta | nocturnal |
| lizards | Paroedura_stellata | Paroedura_stellata | nocturnal |
| lizards | Paroedura_stumpffi | Paroedura_stumpffi | nocturnal |
| lizards | Paroedura_tanjaka | Paroedura_tanjaka | cathemeral |
| lizards | Paroedura_vahiny | Paroedura_vahiny | nocturnal |
| lizards | Paroedura_vazimba | Paroedura_vazimba | nocturnal |
| lizards | Parvilacerta_fraasii | Parvilacerta_fraasii | diurnal |
| lizards | Parvilacerta_parva | Parvilacerta_parva | diurnal |
| lizards | Parvoscincus_beyeri | Parvoscincus_beyeri | diurnal |
| lizards | Parvoscincus_boyingi | Parvoscincus_boyingi | diurnal |
| lizards | Parvoscincus_sisoni | Parvoscincus_sisoni | diurnal |
| lizards | Parvoscincus_steerei | Parvoscincus_steerei | diurnal |
| lizards | Parvoscincus_tagapayo | Parvoscincus_tagapayo | diurnal |
| lizards | Pedioplanis_breviceps | Pedioplanis_breviceps | diurnal |
| lizards | Pedioplanis_burchelli | Pedioplanis_burchelli | diurnal |
| lizards | Pedioplanis_husabensis | Pedioplanis_husabensis | diurnal |
| lizards | Pedioplanis_inornata | Pedioplanis_inornata | diurnal |
| lizards | Pedioplanis_lineoocellata | Pedioplanis_lineoocellata | diurnal |
| lizards | Pedioplanis_namaquensis | Pedioplanis_namaquensis | diurnal |
| lizards | Pedioplanis_rubens | Pedioplanis_rubens | diurnal |

| | | | |
|---|---|---|---|
| lizards | Pedioplanis_undata | Pedioplanis_undata | diurnal |
| lizards | Perochirus_ateles | Perochirus_ateles | nocturnal |
| lizards | Perochirus_scutellatus | Perochirus_scutellatus | cathemeral |
| lizards | Petrosaurus_mearnsi | Petrosaurus_mearnsi | diurnal |
| lizards | Petrosaurus_thalassinus | Petrosaurus_thalassinus | diurnal |
| lizards | Phasmasaurus_maruia | Lioscincus_maruia | diurnal |
| lizards | Phasmasaurus_tillieri | Lioscincus_tillieri | diurnal |
| lizards | Phelsuma_abbotti | Phelsuma_abbotti | diurnal |
| lizards | Phelsuma_andamanensis | Phelsuma_andamanense | diurnal |
| lizards | Phelsuma_antanosy | Phelsuma_antanosy | diurnal |
| lizards | Phelsuma_astriata | Phelsuma_astriata | diurnal |
| lizards | Phelsuma_barbouri | Phelsuma_barbouri | diurnal |
| lizards | Phelsuma_berghofi | Phelsuma_berghofi | diurnal |
| lizards | Phelsuma_borbonica | Phelsuma_borbonica | diurnal |
| lizards | Phelsuma_breviceps | Phelsuma_breviceps | diurnal |
| lizards | Phelsuma_cepediana | Phelsuma_cepediana | diurnal |
| lizards | Phelsuma_comorensis | Phelsuma_comorensis | diurnal |
| lizards | Phelsuma_dorsivittata | Phelsuma_dorsivittata | diurnal |
| lizards | Phelsuma_dubia | Phelsuma_dubia | diurnal |
| lizards | Phelsuma_grandis | Phelsuma_grandis | diurnal |
| lizards | Phelsuma_guentheri | Phelsuma_guentheri | cathemeral |
| lizards | Phelsuma_guimbeaui | Phelsuma_guimbeaui | diurnal |
| lizards | Phelsuma_guttata | Phelsuma_guttata | diurnal |
| lizards | Phelsuma_hoeschi | Phelsuma_hoeschi | diurnal |
| lizards | Phelsuma_inexpectata | Phelsuma_inexpectata | diurnal |
| lizards | Phelsuma_kely | Phelsuma_kely | diurnal |
| lizards | Phelsuma_klemmeri | Phelsuma_klemmeri | diurnal |
| lizards | Phelsuma_kochi | Phelsuma_kochi | diurnal |
| lizards | Phelsuma_laticauda | Phelsuma_laticauda | diurnal |
| lizards | Phelsuma_lineata | Phelsuma_lineata | diurnal |
| lizards | Phelsuma_madagascariensis | Phelsuma_madagascariensis | diurnal |
| lizards | Phelsuma_malamakibo | Phelsuma_malamakibo | diurnal |
| lizards | Phelsuma_masohoala | Phelsuma_masohoala | diurnal |
| lizards | Phelsuma_modesta | Phelsuma_modesta | diurnal |

| | | | |
|---|---|---|---|
| lizards | Phelsuma_mutabilis | Phelsuma_mutabilis | diurnal |
| lizards | Phelsuma_nigristriata | Phelsuma_nigristriata | diurnal |
| lizards | Phelsuma_ornata | Phelsuma_ornata | diurnal |
| lizards | Phelsuma_parkeri | Phelsuma_parkeri | diurnal |
| lizards | Phelsuma_parva | Phelsuma_parva | diurnal |
| lizards | Phelsuma_pronki | Phelsuma_pronki | diurnal |
| lizards | Phelsuma_pusilla | Phelsuma_pusilla | diurnal |
| lizards | Phelsuma_quadriocellata | Phelsuma_quadriocellata | diurnal |
| lizards | Phelsuma_robertmertensi | Phelsuma_robertmertensi | diurnal |
| lizards | Phelsuma_rosagularis | Phelsuma_rosagularis | diurnal |
| lizards | Phelsuma_seippi | Phelsuma_seippi | diurnal |
| lizards | Phelsuma_serraticauda | Phelsuma_serraticauda | diurnal |
| lizards | Phelsuma_standingi | Phelsuma_standingi | diurnal |
| lizards | Phelsuma_sundbergi | Phelsuma_sundbergi | diurnal |
| lizards | Phelsuma_v-nigra | Phelsuma_v-nigra | diurnal |
| lizards | Phoenicolacerta_kulzeri | Phoenicolacerta_kulzeri | diurnal |
| lizards | Phoenicolacerta_laevis | Phoenicolacerta_laevis | diurnal |
| lizards | Phoenicolacerta_troodica | Phoenicolacerta_troodica | diurnal |
| lizards | Pholidobolus_affinis | Pholidobolus_affinis | diurnal |
| lizards | Pholidobolus_macbrydei | Pholidobolus_macbrydei | diurnal |
| lizards | Pholidobolus_montium | Pholidobolus_montium | diurnal |
| lizards | Pholidobolus_prefrontalis | Pholidobolus_prefrontalis | diurnal |
| lizards | Pholidobolus_vertebralis | Cercosaura_vertebralis | diurnal |
| lizards | Pholidoscelis_auberi | Ameiva_auberi | diurnal |
| lizards | Pholidoscelis_chrysolaemus | Ameiva_chrysolaema | diurnal |
| lizards | Pholidoscelis_exsul | Ameiva_exsul | diurnal |
| lizards | Pholidoscelis_fuscatus | Ameiva_fuscata | diurnal |
| lizards | Pholidoscelis_plei | Ameiva_plei | diurnal |
| lizards | Pholidoscelis_taeniurus | Ameiva_taeniura | diurnal |
| lizards | Pholidoscelis_wetmorei | Ameiva_wetmorei | diurnal |
| lizards | Phrynocephalus_arabicus | Phrynocephalus_arabicus | diurnal |
| lizards | Phrynocephalus_axillaris | Phrynocephalus_axillaris | diurnal |
| lizards | Phrynocephalus_clarkorum | Phrynocephalus_clarkorum | diurnal |
| lizards | Phrynocephalus_erythrurus | Phrynocephalus_erythrurus | diurnal |

| | | | |
|---|---|---|---|
| lizards | Phrynocephalus_forsythii | Phrynocephalus_forsythii | diurnal |
| lizards | Phrynocephalus_golubewii | Phrynocephalus_golubewii | diurnal |
| lizards | Phrynocephalus_guttatus | Phrynocephalus_guttatus | diurnal |
| lizards | Phrynocephalus_helioscopus | Phrynocephalus_helioscopus | diurnal |
| lizards | Phrynocephalus_interscapularis | Phrynocephalus_interscapularis | diurnal |
| lizards | Phrynocephalus_luteoguttatus | Phrynocephalus_luteoguttatus | diurnal |
| lizards | Phrynocephalus_maculatus | Phrynocephalus_maculatus | diurnal |
| lizards | Phrynocephalus_mystaceus | Phrynocephalus_mystaceus | diurnal |
| lizards | Phrynocephalus_ornatus | Phrynocephalus_ornatus | diurnal |
| lizards | Phrynocephalus_persicus | Phrynocephalus_persicus | diurnal |
| lizards | Phrynocephalus_przewalskii | Phrynocephalus_przewalskii | diurnal |
| lizards | Phrynocephalus_putjatai | Phrynocephalus_putjatai | diurnal |
| lizards | Phrynocephalus_raddei | Phrynocephalus_raddei | diurnal |
| lizards | Phrynocephalus_roborowskii | Phrynocephalus_roborowskii | diurnal |
| lizards | Phrynocephalus_rossikowi | Phrynocephalus_rossikowi | diurnal |
| lizards | Phrynocephalus_scutellatus | Phrynocephalus_scutellatus | diurnal |
| lizards | Phrynocephalus_strauchi | Phrynocephalus_strauchi | diurnal |
| lizards | Phrynocephalus_theobaldi | Phrynocephalus_theobaldi | diurnal |
| lizards | Phrynocephalus_versicolor | Phrynocephalus_versicolor | diurnal |
| lizards | Phrynocephalus_vlangalii | Phrynocephalus_vlangalii | diurnal |
| lizards | Phrynosoma_asio | Phrynosoma_asio | diurnal |
| lizards | Phrynosoma_blainvillii | Phrynosoma_blainvillii | diurnal |
| lizards | Phrynosoma_braconnieri | Phrynosoma_braconnieri | diurnal |
| lizards | Phrynosoma_cornutum | Phrynosoma_cornutum | diurnal |
| lizards | Phrynosoma_coronatum | Phrynosoma_coronatum | diurnal |
| lizards | Phrynosoma_ditmarsi | Phrynosoma_ditmarsi | diurnal |
| lizards | Phrynosoma_douglasii | Phrynosoma_douglasii | diurnal |
| lizards | Phrynosoma_goodei | Phrynosoma_goodei | diurnal |
| lizards | Phrynosoma_hernandesi | Phrynosoma_hernandesi | diurnal |
| lizards | Phrynosoma_mcallii | Phrynosoma_mcallii | diurnal |
| lizards | Phrynosoma_modestum | Phrynosoma_modestum | cathemeral |
| lizards | Phrynosoma_orbiculare | Phrynosoma_orbiculare | diurnal |
| lizards | Phrynosoma_platyrhinos | Phrynosoma_platyrhinos | diurnal |
| lizards | Phrynosoma_solare | Phrynosoma_solare | diurnal |

| | | | |
|---|---|---|---|
| lizards | Phrynosoma_taurus | Phrynosoma_taurus | diurnal |
| lizards | Phyllodactylus_angustidigitus | Phyllodactylus_angustidigitus | nocturnal |
| lizards | Phyllodactylus_baurii | Phyllodactylus_baurii | nocturnal |
| lizards | Phyllodactylus_bugastrolepis | Phyllodactylus_bugastrolepis | nocturnal |
| lizards | Phyllodactylus_delsolari | Phyllodactylus_delsolari | nocturnal |
| lizards | Phyllodactylus_duellmani | Phyllodactylus_duellmani | nocturnal |
| lizards | Phyllodactylus_galapagensis | Phyllodactylus_galapagensis | nocturnal |
| lizards | Phyllodactylus_gerrhopygus | Phyllodactylus_gerrhopygus | cathemeral |
| lizards | Phyllodactylus_insularis | Phyllodactylus_insularis | nocturnal |
| lizards | Phyllodactylus_interandinus | Phyllodactylus_interandinus | nocturnal |
| lizards | Phyllodactylus_johnwrighti | Phyllodactylus_johnwrighti | nocturnal |
| lizards | Phyllodactylus_kofordi | Phyllodactylus_kofordi | nocturnal |
| lizards | Phyllodactylus_lanei | Phyllodactylus_lanei | nocturnal |
| lizards | Phyllodactylus_leei | Phyllodactylus_leei | nocturnal |
| lizards | Phyllodactylus_leoni | Phyllodactylus_leoni | nocturnal |
| lizards | Phyllodactylus_lepidopygus | Phyllodactylus_lepidopygus | nocturnal |
| lizards | Phyllodactylus_microphyllus | Phyllodactylus_microphyllus | nocturnal |
| lizards | Phyllodactylus_nocticolus | Phyllodactylus_nocticolus | nocturnal |
| lizards | Phyllodactylus_palmeus | Phyllodactylus_palmeus | nocturnal |
| lizards | Phyllodactylus_paucituberculatus | Phyllodactylus_paucituberculatus | nocturnal |
| lizards | Phyllodactylus_pulcher | Phyllodactylus_pulcher | nocturnal |
| lizards | Phyllodactylus_reissii | Phyllodactylus_reissii | nocturnal |
| lizards | Phyllodactylus_thompsoni | Phyllodactylus_thompsoni | nocturnal |
| lizards | Phyllodactylus_tuberculosus | Phyllodactylus_tuberculosus | nocturnal |
| lizards | Phyllodactylus_wirshingi | Phyllodactylus_wirshingi | nocturnal |
| lizards | Phyllodactylus_xanti | Phyllodactylus_xanti | nocturnal |
| lizards | Phyllopezus_maranjonensis | Phyllopezus_maranjonensis | nocturnal |
| lizards | Phyllopezus_periosus | Phyllopezus_periosus | nocturnal |
| lizards | Phyllopezus_pollicaris | Phyllopezus_pollicaris | cathemeral |
| lizards | Phyllurus_amnicola | Phyllurus_amnicola | nocturnal |
| lizards | Phyllurus_caudiannulatus | Phyllurus_caudiannulatus | nocturnal |
| lizards | Phyllurus_championae | Phyllurus_championae | nocturnal |
| lizards | Phyllurus_gulbaru | Phyllurus_gulbaru | nocturnal |
| lizards | Phyllurus_nepthys | Phyllurus_nepthys | nocturnal |

| | | | |
|---|---|---|---|
| lizards | Phyllurus_platurus | Phyllurus_platurus | nocturnal |
| lizards | Phymaturus_aguanegra | Phymaturus_aguanegra | diurnal |
| lizards | Phymaturus_aguedae | Phymaturus_aguedae | diurnal |
| lizards | Phymaturus_antofagastensis | Phymaturus_antofagastensis | diurnal |
| lizards | Phymaturus_camilae | Phymaturus_camilae | diurnal |
| lizards | Phymaturus_castillensis | Phymaturus_castillensis | diurnal |
| lizards | Phymaturus_darwini | Phymaturus_darwini | diurnal |
| lizards | Phymaturus_delheyi | Phymaturus_delheyi | diurnal |
| lizards | Phymaturus_palluma | Phymaturus_palluma | diurnal |
| lizards | Phymaturus_patagonicus | Phymaturus_patagonicus | diurnal |
| lizards | Phymaturus_payuniae | Phymaturus_payuniae | diurnal |
| lizards | Phymaturus_punae | Phymaturus_punae | diurnal |
| lizards | Phymaturus_roigorum | Phymaturus_roigorum | diurnal |
| lizards | Phymaturus_sinervoi | Phymaturus_sinervoi | diurnal |
| lizards | Phymaturus_somuncurensis | Phymaturus_somuncurensis | diurnal |
| lizards | Phymaturus_spurcus | Phymaturus_spurcus | diurnal |
| lizards | Phymaturus_tenebrosus | Phymaturus_tenebrosus | diurnal |
| lizards | Phymaturus_videlai | Phymaturus_videlai | diurnal |
| lizards | Phymaturus_vociferator | Phymaturus_vociferator | diurnal |
| lizards | Phymaturus_williamsi | Phymaturus_williamsi | diurnal |
| lizards | Phymaturus_zapalensis | Phymaturus_zapalensis | diurnal |
| lizards | Physignathus_cocincinus | Physignathus_cocincinus | diurnal |
| lizards | Pinoyscincus_coxi | Pinoyscincus_coxi | diurnal |
| lizards | Pinoyscincus_jagori | Pinoyscincus_jagori | diurnal |
| lizards | Pinoyscincus_mindanensis | Pinoyscincus_mindanensis | diurnal |
| lizards | Platysaurus_broadleyi | Platysaurus_broadleyi | diurnal |
| lizards | Platysaurus_capensis | Platysaurus_capensis | diurnal |
| lizards | Platysaurus_imperator | Platysaurus_imperator | diurnal |
| lizards | Platysaurus_intermedius | Platysaurus_intermedius | diurnal |
| lizards | Platysaurus_lebomboensis | Platysaurus_lebomboensis | diurnal |
| lizards | Platysaurus_maculatus | Platysaurus_maculatus | diurnal |
| lizards | Platysaurus_minor | Platysaurus_minor | diurnal |
| lizards | Platysaurus_monotropis | Platysaurus_monotropis | diurnal |
| lizards | Platysaurus_ocellatus | Platysaurus_ocellatus | diurnal |

| | | | |
|---|---|---|---|
| lizards | Platysaurus_orientalis | Platysaurus_orientalis | diurnal |
| lizards | Platysaurus_pungweensis | Platysaurus_pungweensis | diurnal |
| lizards | Plestiodon_anthracinus | Plestiodon_anthracinus | diurnal |
| lizards | Plestiodon_bilineatus | Plestiodon_bilineatus | diurnal |
| lizards | Plestiodon_brevirostris | Plestiodon_brevirostris | diurnal |
| lizards | Plestiodon_callicephalus | Plestiodon_callicephalus | diurnal |
| lizards | Plestiodon_capito | Plestiodon_capito | diurnal |
| lizards | Plestiodon_chinensis | Plestiodon_chinensis | diurnal |
| lizards | Plestiodon_copei | Plestiodon_copei | diurnal |
| lizards | Plestiodon_dugesii | Plestiodon_dugesii | diurnal |
| lizards | Plestiodon_egregius | Plestiodon_egregius | diurnal |
| lizards | Plestiodon_elegans | Plestiodon_elegans | diurnal |
| lizards | Plestiodon_fasciatus | Plestiodon_fasciatus | diurnal |
| lizards | Plestiodon_gilberti | Plestiodon_gilberti | diurnal |
| lizards | Plestiodon_inexpectatus | Plestiodon_inexpectatus | diurnal |
| lizards | Plestiodon_japonicus | Plestiodon_japonicus | diurnal |
| lizards | Plestiodon_kishinouyei | Plestiodon_kishinouyei | diurnal |
| lizards | Plestiodon_lagunensis | Plestiodon_lagunensis | diurnal |
| lizards | Plestiodon_laticeps | Plestiodon_laticeps | diurnal |
| lizards | Plestiodon_latiscutatus | Plestiodon_latiscutatus | diurnal |
| lizards | Plestiodon_longirostris | Plestiodon_longirostris | diurnal |
| lizards | Plestiodon_lynxe | Plestiodon_lynxe | diurnal |
| lizards | Plestiodon_multilineatus | Plestiodon_multilineatus | diurnal |
| lizards | Plestiodon_multivirgatus | Plestiodon_multivirgatus | diurnal |
| lizards | Plestiodon_obsoletus | Plestiodon_obsoletus | cathemeral |
| lizards | Plestiodon_parviauriculatus | Plestiodon_parviauriculatus | diurnal |
| lizards | Plestiodon_quadrilineatus | Plestiodon_quadrilineatus | diurnal |
| lizards | Plestiodon_reynoldsi | Plestiodon_reynoldsi | diurnal |
| lizards | Plestiodon_septentrionalis | Plestiodon_septentrionalis | diurnal |
| lizards | Plestiodon_skiltonianus | Plestiodon_skiltonianus | diurnal |
| lizards | Plestiodon_stimpsonii | Plestiodon_stimpsonii | diurnal |
| lizards | Plestiodon_sumichrasti | Plestiodon_sumichrasti | diurnal |
| lizards | Plestiodon_tetragrammus | Plestiodon_tetragrammus | diurnal |
| lizards | Pletholax_gracilis | Pletholax_gracilis | diurnal |

| | | | |
|---|---|---|---|
| lizards | Plica_caribeana | Plica_caribeana | diurnal |
| lizards | Plica_plica | Plica_plica | diurnal |
| lizards | Plica_umbra | Plica_umbra | diurnal |
| lizards | Podarcis_bocagei | Podarcis_bocagei | diurnal |
| lizards | Podarcis_carbonelli | Podarcis_carbonelli | diurnal |
| lizards | Podarcis_cretensis | Podarcis_cretensis | diurnal |
| lizards | Podarcis_erhardii | Podarcis_erhardii | diurnal |
| lizards | Podarcis_filfolensis | Podarcis_filfolensis | diurnal |
| lizards | Podarcis_gaigeae | Podarcis_gaigeae | diurnal |
| lizards | Podarcis_guadarramae | Podarcis_guadarramae | diurnal |
| lizards | Podarcis_hispanicus | Podarcis_hispanicus | diurnal |
| lizards | Podarcis_lilfordi | Podarcis_lilfordi | diurnal |
| lizards | Podarcis_liolepis | Podarcis_liolepis | diurnal |
| lizards | Podarcis_melisellensis | Podarcis_melisellensis | diurnal |
| lizards | Podarcis_milensis | Podarcis_milensis | diurnal |
| lizards | Podarcis_muralis | Podarcis_muralis | diurnal |
| lizards | Podarcis_peloponnesiacus | Podarcis_peloponnesiacus | diurnal |
| lizards | Podarcis_pityusensis | Podarcis_pityusensis | diurnal |
| lizards | Podarcis_raffonei | Podarcis_raffoneae | diurnal |
| lizards | Podarcis_siculus | Podarcis_siculus | diurnal |
| lizards | Podarcis_tauricus | Podarcis_tauricus | diurnal |
| lizards | Podarcis_tiliguerta | Podarcis_tiliguerta | diurnal |
| lizards | Podarcis_vaucheri | Podarcis_vaucheri | diurnal |
| lizards | Podarcis_waglerianus | Podarcis_waglerianus | diurnal |
| lizards | Pogona_barbata | Pogona_barbata | diurnal |
| lizards | Pogona_henrylawsoni | Pogona_henrylawsoni | diurnal |
| lizards | Pogona_microlepidota | Pogona_microlepidota | diurnal |
| lizards | Pogona_minor | Pogona_minor | diurnal |
| lizards | Pogona_vitticeps | Pogona_vitticeps | diurnal |
| lizards | Polychrus_acutirostris | Polychrus_acutirostris | diurnal |
| lizards | Polychrus_femoralis | Polychrus_femoralis | diurnal |
| lizards | Polychrus_gutturosus | Polychrus_gutturosus | diurnal |
| lizards | Polychrus_jacquelinae | Polychrus_jacquelinae | diurnal |
| lizards | Polychrus_liogaster | Polychrus_liogaster | diurnal |

| | | | |
|---|---|---|---|
| lizards | Polychrus_marmoratus | Polychrus_marmoratus | diurnal |
| lizards | Polychrus_peruvianus | Polychrus_peruvianus | diurnal |
| lizards | Potamites_ecpleopus | Potamites_ecpleopus | diurnal |
| lizards | Potamites_juruazensis | Potamites_juruazensis | diurnal |
| lizards | Prasinohaema_flavipes | Prasinohaema_flavipes | diurnal |
| lizards | Prasinohaema_semoni | Prasinohaema_semoni | diurnal |
| lizards | Prasinohaema_virens | Prasinohaema_virens | diurnal |
| lizards | Pristidactylus_achalensis | Pristidactylus_achalensis | diurnal |
| lizards | Pristidactylus_scapulatus | Pristidactylus_scapulatus | diurnal |
| lizards | Pristidactylus_torquatus | Pristidactylus_torquatus | diurnal |
| lizards | Pristidactylus_volcanensis | Pristidactylus_volcanensis | diurnal |
| lizards | Pristurus_abdelkuri | Pristurus_abdelkuri | diurnal |
| lizards | Pristurus_carteri | Pristurus_carteri | cathemeral |
| lizards | Pristurus_celerrimus | Pristurus_celerrimus | diurnal |
| lizards | Pristurus_collaris | Pristurus_collaris | cathemeral |
| lizards | Pristurus_crucifer | Pristurus_crucifer | diurnal |
| lizards | Pristurus_flavipunctatus | Pristurus_flavipunctatus | diurnal |
| lizards | Pristurus_gallagheri | Pristurus_gallagheri | diurnal |
| lizards | Pristurus_guichardi | Pristurus_guichardi | diurnal |
| lizards | Pristurus_insignis | Pristurus_insignis | diurnal |
| lizards | Pristurus_insignoides | Pristurus_insignoides | diurnal |
| lizards | Pristurus_minimus | Pristurus_minimus | cathemeral |
| lizards | Pristurus_obsti | Pristurus_obsti | diurnal |
| lizards | Pristurus_ornithocephalus | Pristurus_ornithocephalus | diurnal |
| lizards | Pristurus_phillipsii | Pristurus_phillipsii | diurnal |
| lizards | Pristurus_popovi | Pristurus_popovi | diurnal |
| lizards | Pristurus_rupestris | Pristurus_rupestris | diurnal |
| lizards | Pristurus_saada | Pristurus_saada | diurnal |
| lizards | Pristurus_samhaensis | Pristurus_samhaensis | diurnal |
| lizards | Pristurus_sokotranus | Pristurus_sokotranus | diurnal |
| lizards | Pristurus_somalicus | Pristurus_somalicus | diurnal |
| lizards | Proablepharus_reginae | Proablepharus_reginae | diurnal |
| lizards | Proablepharus_tenuis | Proablepharus_tenuis | diurnal |
| lizards | Proctoporus_chasqui | Proctoporus_chasqui | diurnal |

| | | | |
|---|---|---|---|
| lizards | Proctoporus_laudahnae | Riama_laudahnae | diurnal |
| lizards | Proctoporus_oreades | Euspondylus_oreades | diurnal |
| lizards | Proctoporus_sucullucu | Proctoporus_sucullucu | diurnal |
| lizards | Proctoporus_unsaacae | Proctoporus_unsaacae | diurnal |
| lizards | Proscelotes_arnoldi | Proscelotes_arnoldi | diurnal |
| lizards | Proscelotes_eggeli | Proscelotes_eggeli | diurnal |
| lizards | Psammodromus_algirus | Psammodromus_algirus | diurnal |
| lizards | Psammodromus_blanci | Psammodromus_blanci | diurnal |
| lizards | Psammodromus_hispanicus | Psammodromus_hispanicus | diurnal |
| lizards | Psammophilus_blanfordanus | Psammophilus_blanfordanus | diurnal |
| lizards | Psammophilus_dorsalis | Psammophilus_dorsalis | diurnal |
| lizards | Pseudemoia_baudini | Pseudemoia_baudini | diurnal |
| lizards | Pseudemoia_cryodroma | Pseudemoia_cryodroma | diurnal |
| lizards | Pseudemoia_entrecasteauxii | Pseudemoia_entrecasteauxii | diurnal |
| lizards | Pseudemoia_pagenstecheri | Pseudemoia_pagenstecheri | diurnal |
| lizards | Pseudemoia_rawlinsoni | Pseudemoia_rawlinsoni | diurnal |
| lizards | Pseudemoia_spenceri | Pseudemoia_spenceri | diurnal |
| lizards | Pseudocalotes_brevipes | Pseudocalotes_brevipes | diurnal |
| lizards | Pseudocalotes_cybelidermus | Pseudocalotes_cybelidermus | diurnal |
| lizards | Pseudocalotes_flavigula | Pseudocalotes_flavigula | diurnal |
| lizards | Pseudocalotes_floweri | Pseudocalotes_floweri | diurnal |
| lizards | Pseudocalotes_guttalineatus | Pseudocalotes_guttalineatus | diurnal |
| lizards | Pseudocalotes_kakhienensis | Pseudocalotes_kakhienensis | diurnal |
| lizards | Pseudocalotes_larutensis | Pseudocalotes_larutensis | diurnal |
| lizards | Pseudocalotes_rhammanotus | Pseudocalotes_rhammanotus | diurnal |
| lizards | Pseudocalotes_tympanistriga | Pseudocalotes_tympanistriga | diurnal |
| lizards | Pseudoceramodactylus_khobarensis | Pseudoceramodactylus_khobarensis | nocturnal |
| lizards | Pseudocophotis_kontumensis | Pseudocophotis_kontumensis | diurnal |
| lizards | Pseudocordylus_langi | Pseudocordylus_langi | diurnal |
| lizards | Pseudocordylus_melanotus | Pseudocordylus_melanotus | diurnal |
| lizards | Pseudocordylus_microlepidotus | Pseudocordylus_microlepidotus | diurnal |
| lizards | Pseudocordylus_spinosus | Pseudocordylus_spinosus | diurnal |
| lizards | Pseudocordylus_subviridis | Pseudocordylus_subviridis | diurnal |
| lizards | Pseudocordylus_transvaalensis | Pseudocordylus_transvaalensis | diurnal |

| | | | |
|---|---|---|---|
| lizards | Pseudogekko_brevipes | Pseudogekko_brevipes | nocturnal |
| lizards | Pseudogekko_compresicorpus | Pseudogekko_compressicorpus | nocturnal |
| lizards | Pseudogekko_smaragdinus | Pseudogekko_smaragdinus | nocturnal |
| lizards | Pseudogonatodes_guianensis | Pseudogonatodes_guianensis | diurnal |
| lizards | Pseudogonatodes_lunulatus | Pseudogonatodes_lunulatus | cathemeral |
| lizards | Pseudopus_apodus | Pseudopus_apodus | diurnal |
| lizards | Pseudotrapelus_aqabensis | Pseudotrapelus_aqabensis | diurnal |
| lizards | Pseudotrapelus_sinaitus | Pseudotrapelus_sinaitus | diurnal |
| lizards | Psilops_paeminosus | Psilophthalmus_paeminosus | diurnal |
| lizards | Psychosaura_agmosticha | Psychosaura_agmosticha | diurnal |
| lizards | Psychosaura_macrorhyncha | Psychosaura_macrorhyncha | diurnal |
| lizards | Ptenopus_carpi | Ptenopus_carpi | nocturnal |
| lizards | Ptenopus_garrulus | Ptenopus_garrulus | nocturnal |
| lizards | Ptyctolaemus_gularis | Ptyctolaemus_gularis | cathemeral |
| lizards | Ptyodactylus_guttatus | Ptyodactylus_guttatus | cathemeral |
| lizards | Ptyodactylus_hasselquistii | Ptyodactylus_hasselquistii | cathemeral |
| lizards | Ptyodactylus_oudrii | Ptyodactylus_oudrii | nocturnal |
| lizards | Ptyodactylus_puiseuxi | Ptyodactylus_puiseuxi | diurnal |
| lizards | Ptyodactylus_ragazzii | Ptyodactylus_ragazzii | cathemeral |
| lizards | Pygmaeascincus_sadlieri | Pygmaeascincus_sadlieri | diurnal |
| lizards | Pygmaeascincus_timlowi | Pygmaeascincus_timlowi | diurnal |
| lizards | Pygopus_lepidopodus | Pygopus_lepidopodus | cathemeral |
| lizards | Pygopus_nigriceps | Pygopus_nigriceps | nocturnal |
| lizards | Pygopus_schraderi | Pygopus_schraderi | nocturnal |
| lizards | Pygopus_steelescotti | Pygopus_steelescotti | cathemeral |
| lizards | Quedenfeldtia_moerens | Quedenfeldtia_moerens | diurnal |
| lizards | Quedenfeldtia_trachyblepharus | Quedenfeldtia_trachyblepharus | diurnal |
| lizards | Ramigekko_swartbergensis | Ramigekko_swartbergensis | nocturnal |
| lizards | Rankinia_diemensis | Rankinia_diemensis | diurnal |
| lizards | Rhacodactylus_auriculatus | Rhacodactylus_auriculatus | nocturnal |
| lizards | Rhacodactylus_leachianus | Rhacodactylus_leachianus | nocturnal |
| lizards | Rhampholeon_boulengeri | Rhampholeon_boulengeri | diurnal |
| lizards | Rhampholeon_marshalli | Rhampholeon_marshalli | diurnal |
| lizards | Rhampholeon_maspictus | Rhampholeon_maspictus | diurnal |

| | | | |
|---|---|---|---|
| lizards | Rhampholeon_moyeri | Rhampholeon_moyeri | diurnal |
| lizards | Rhampholeon_platyceps | Rhampholeon_platyceps | diurnal |
| lizards | Rhampholeon_spectrum | Rhampholeon_spectrum | diurnal |
| lizards | Rhampholeon_spinosus | Rhampholeon_spinosus | diurnal |
| lizards | Rhampholeon_temporalis | Rhampholeon_temporalis | diurnal |
| lizards | Rhampholeon_tilburyi | Rhampholeon_tilburyi | diurnal |
| lizards | Rhampholeon_uluguruensis | Rhampholeon_uluguruensis | diurnal |
| lizards | Rhoptropella_ocellata | Rhoptropella_ocellata | diurnal |
| lizards | Rhoptropus_afer | Rhoptropus_afer | diurnal |
| lizards | Rhoptropus_barnardi | Rhoptropus_barnardi | diurnal |
| lizards | Rhoptropus_boultoni | Rhoptropus_boultoni | diurnal |
| lizards | Rhoptropus_bradfieldi | Rhoptropus_bradfieldi | diurnal |
| lizards | Rhynchoedura_angusta | Rhynchoedura_angusta | nocturnal |
| lizards | Rhynchoedura_ormsbyi | Rhynchoedura_ormsbyi | nocturnal |
| lizards | Rhynchoedura_ornata | Rhynchoedura_ornata | nocturnal |
| lizards | Rhynchoedura_sexapora | Rhynchoedura_sexapora | nocturnal |
| lizards | Riama_columbiana | Riama_columbiana | diurnal |
| lizards | Rieppeleon_brachyurus | Rieppeleon_brachyurus | diurnal |
| lizards | Rieppeleon_brevicaudatus | Rieppeleon_brevicaudatus | diurnal |
| lizards | Rieppeleon_kerstenii | Rieppeleon_kerstenii | diurnal |
| lizards | Ristella_beddomii | Ristella_beddomii | diurnal |
| lizards | Ristella_rurkii | Ristella_rurkii | diurnal |
| lizards | Ristella_travancorica | Ristella_travancorica | diurnal |
| lizards | Saara_hardwickii | Saara_hardwickii | diurnal |
| lizards | Saara_loricata | Saara_loricata | diurnal |
| lizards | Saiphos_equalis | Saiphos_equalis | cathemeral |
| lizards | Salea_anamallayana | Salea_anamallayana | diurnal |
| lizards | Salea_horsfieldii | Salea_horsfieldii | diurnal |
| lizards | Saltuarius_cornutus | Saltuarius_cornutus | nocturnal |
| lizards | Saltuarius_kateae | Saltuarius_kateae | nocturnal |
| lizards | Saltuarius_moritzi | Saltuarius_moritzi | nocturnal |
| lizards | Saltuarius_salebrosus | Saltuarius_salebrosus | nocturnal |
| lizards | Saltuarius_swaini | Saltuarius_swaini | nocturnal |
| lizards | Saltuarius_wyberba | Saltuarius_wyberba | nocturnal |

| | | | |
|---|---|---|---|
| lizards | Salvator_merianae | Salvator_merianae | diurnal |
| lizards | Salvator_rufescens | Salvator_rufescens | diurnal |
| lizards | Saproscincus_basiliscus | Saproscincus_basiliscus | diurnal |
| lizards | Saproscincus_challengeri | Saproscincus_challengeri | diurnal |
| lizards | Saproscincus_czechurai | Saproscincus_czechurai | diurnal |
| lizards | Saproscincus_eungellensis | Saproscincus_eungellensis | diurnal |
| lizards | Saproscincus_mustelinus | Saproscincus_mustelinus | cathemeral |
| lizards | Saproscincus_oriarus | Saproscincus_oriarus | diurnal |
| lizards | Saproscincus_rosei | Saproscincus_rosei | diurnal |
| lizards | Saproscincus_spectabilis | Saproscincus_spectabilis | diurnal |
| lizards | Saproscincus_tetradactylus | Saproscincus_tetradactylus | diurnal |
| lizards | Saurodactylus_fasciatus | Saurodactylus_fasciatus | nocturnal |
| lizards | Saurodactylus_mauritanicus | Saurodactylus_mauritanicus | diurnal |
| lizards | Sauromalus_ater | Sauromalus_ater | diurnal |
| lizards | Sauromalus_hispidus | Sauromalus_hispidus | diurnal |
| lizards | Sauromalus_varius | Sauromalus_varius | diurnal |
| lizards | Scelarcis_perspicillata | Scelarcis_perspicillata | diurnal |
| lizards | Sceloporus_acanthinus | Sceloporus_acanthinus | diurnal |
| lizards | Sceloporus_adleri | Sceloporus_adleri | diurnal |
| lizards | Sceloporus_aeneus | Sceloporus_aeneus | diurnal |
| lizards | Sceloporus_albiventris | Sceloporus_albiventris | diurnal |
| lizards | Sceloporus_angustus | Sceloporus_angustus | cathemeral |
| lizards | Sceloporus_arenicolus | Sceloporus_arenicolus | diurnal |
| lizards | Sceloporus_becki | Sceloporus_becki | diurnal |
| lizards | Sceloporus_bicanthalis | Sceloporus_bicanthalis | diurnal |
| lizards | Sceloporus_bimaculosus | Sceloporus_bimaculosus | diurnal |
| lizards | Sceloporus_bulleri | Sceloporus_bulleri | diurnal |
| lizards | Sceloporus_caeruleus | Sceloporus_caeruleus | diurnal |
| lizards | Sceloporus_chaneyi | Sceloporus_chaneyi | diurnal |
| lizards | Sceloporus_chrysostictus | Sceloporus_chrysostictus | diurnal |
| lizards | Sceloporus_clarkii | Sceloporus_clarkii | diurnal |
| lizards | Sceloporus_consobrinus | Sceloporus_consobrinus | diurnal |
| lizards | Sceloporus_couchii | Sceloporus_couchii | diurnal |
| lizards | Sceloporus_cowlesi | Sceloporus_cowlesi | diurnal |

| | | | |
|---|---|---|---|
| lizards | Sceloporus_cozumelae | Sceloporus_cozumelae | diurnal |
| lizards | Sceloporus_cryptus | Sceloporus_cryptus | diurnal |
| lizards | Sceloporus_cyanogenys | Sceloporus_cyanogenys | diurnal |
| lizards | Sceloporus_cyanostictus | Sceloporus_cyanostictus | diurnal |
| lizards | Sceloporus_druckercolini | Sceloporus_druckercolini | diurnal |
| lizards | Sceloporus_dugesii | Sceloporus_dugesii | diurnal |
| lizards | Sceloporus_edwardtaylori | Sceloporus_edwardtaylori | diurnal |
| lizards | Sceloporus_exsul | Sceloporus_exsul | diurnal |
| lizards | Sceloporus_formosus | Sceloporus_formosus | diurnal |
| lizards | Sceloporus_gadoviae | Sceloporus_gadoviae | diurnal |
| lizards | Sceloporus_goldmani | Sceloporus_goldmani | diurnal |
| lizards | Sceloporus_graciosus | Sceloporus_graciosus | diurnal |
| lizards | Sceloporus_grammicus | Sceloporus_grammicus | diurnal |
| lizards | Sceloporus_grandaevus | Sceloporus_grandaevus | diurnal |
| lizards | Sceloporus_horridus | Sceloporus_horridus | diurnal |
| lizards | Sceloporus_hunsakeri | Sceloporus_hunsakeri | cathemeral |
| lizards | Sceloporus_internasalis | Sceloporus_internasalis | diurnal |
| lizards | Sceloporus_jalapae | Sceloporus_jalapae | diurnal |
| lizards | Sceloporus_jarrovii | Sceloporus_jarrovii | diurnal |
| lizards | Sceloporus_licki | Sceloporus_licki | diurnal |
| lizards | Sceloporus_macdougalli | Sceloporus_macdougalli | diurnal |
| lizards | Sceloporus_magister | Sceloporus_magister | diurnal |
| lizards | Sceloporus_malachiticus | Sceloporus_malachiticus | diurnal |
| lizards | Sceloporus_megalepidurus | Sceloporus_megalepidurus | diurnal |
| lizards | Sceloporus_melanorhinus | Sceloporus_melanorhinus | diurnal |
| lizards | Sceloporus_merriami | Sceloporus_merriami | diurnal |
| lizards | Sceloporus_minor | Sceloporus_minor | diurnal |
| lizards | Sceloporus_mucronatus | Sceloporus_mucronatus | diurnal |
| lizards | Sceloporus_nelsoni | Sceloporus_nelsoni | diurnal |
| lizards | Sceloporus_occidentalis | Sceloporus_occidentalis | diurnal |
| lizards | Sceloporus_ochoterenae | Sceloporus_ochoterenae | diurnal |
| lizards | Sceloporus_olivaceus | Sceloporus_olivaceus | diurnal |
| lizards | Sceloporus_omiltemanus | Sceloporus_omiltemanus | diurnal |
| lizards | Sceloporus_orcutti | Sceloporus_orcutti | diurnal |

| | | | |
|---|---|---|---|
| lizards | Sceloporus_ornatus | Sceloporus_ornatus | diurnal |
| lizards | Sceloporus_palaciosi | Sceloporus_palaciosi | diurnal |
| lizards | Sceloporus_parvus | Sceloporus_parvus | diurnal |
| lizards | Sceloporus_poinsettii | Sceloporus_poinsettii | diurnal |
| lizards | Sceloporus_pyrocephalus | Sceloporus_pyrocephalus | diurnal |
| lizards | Sceloporus_salvini | Sceloporus_salvini | diurnal |
| lizards | Sceloporus_samcolemani | Sceloporus_samcolemani | diurnal |
| lizards | Sceloporus_scalaris | Sceloporus_scalaris | diurnal |
| lizards | Sceloporus_serrifer | Sceloporus_serrifer | diurnal |
| lizards | Sceloporus_siniferus | Sceloporus_siniferus | diurnal |
| lizards | Sceloporus_slevini | Sceloporus_slevini | diurnal |
| lizards | Sceloporus_smaragdinus | Sceloporus_smaragdinus | diurnal |
| lizards | Sceloporus_smithi | Sceloporus_smithi | diurnal |
| lizards | Sceloporus_spinosus | Sceloporus_spinosus | diurnal |
| lizards | Sceloporus_squamosus | Sceloporus_squamosus | diurnal |
| lizards | Sceloporus_subpictus | Sceloporus_subpictus | diurnal |
| lizards | Sceloporus_sugillatus | Sceloporus_sugillatus | diurnal |
| lizards | Sceloporus_taeniocnemis | Sceloporus_taeniocnemis | diurnal |
| lizards | Sceloporus_teapensis | Sceloporus_teapensis | diurnal |
| lizards | Sceloporus_torquatus | Sceloporus_torquatus | diurnal |
| lizards | Sceloporus_tristichus | Sceloporus_tristichus | diurnal |
| lizards | Sceloporus_undulatus | Sceloporus_undulatus | diurnal |
| lizards | Sceloporus_uniformis | Sceloporus_uniformis | diurnal |
| lizards | Sceloporus_utiformis | Sceloporus_utiformis | diurnal |
| lizards | Sceloporus_variabilis | Sceloporus_variabilis | diurnal |
| lizards | Sceloporus_virgatus | Sceloporus_virgatus | diurnal |
| lizards | Sceloporus_woodi | Sceloporus_woodi | diurnal |
| lizards | Sceloporus_zosteromus | Sceloporus_zosteromus | cathemeral |
| lizards | Scelotes_caffer | Scelotes_caffer | diurnal |
| lizards | Scelotes_gronovii | Scelotes_gronovii | diurnal |
| lizards | Scelotes_limpopoensis | Scelotes_limpopoensis | diurnal |
| lizards | Scincella_assata | Scincella_assatus | diurnal |
| lizards | Scincella_boettgeri | Scincella_boettgeri | cathemeral |
| lizards | Scincella_cherriei | Scincella_cherriei | diurnal |

| | | | |
|---|---|---|---|
| lizards | Scincella_devorator | Scincella_devorator | diurnal |
| lizards | Scincella_gemmingeri | Scincella_gemmingeri | diurnal |
| lizards | Scincella_huanrenensis | Scincella_huanrenensis | diurnal |
| lizards | Scincella_lateralis | Scincella_lateralis | cathemeral |
| lizards | Scincella_melanosticta | Scincella_melanosticta | diurnal |
| lizards | Scincella_modesta | Scincella_modesta | diurnal |
| lizards | Scincella_monticola | Scincella_monticola | cathemeral |
| lizards | Scincella_ochracea | Scincella_ochracea | diurnal |
| lizards | Scincella_reevesii | Scincella_reevesii | cathemeral |
| lizards | Scincella_rufocaudata | Scincella_rufocaudatus | diurnal |
| lizards | Scincella_silvicola | Scincella_silvicola | diurnal |
| lizards | Scincella_tsinlingensis | Scincella_tsinlingensis | diurnal |
| lizards | Scincella_vandenburghi | Scincella_vandenburghi | diurnal |
| lizards | Scincella_victoriana | Scincella_victoriana | diurnal |
| lizards | Scincus_albifasciatus | Scincus_albifasciatus | diurnal |
| lizards | Scincus_mitranus | Scincus_mitranus | diurnal |
| lizards | Scincus_scincus | Scincus_scincus | cathemeral |
| lizards | Sepsina_alberti | Sepsina_alberti | diurnal |
| lizards | Shinisaurus_crocodilurus | Shinisaurus_crocodilurus | diurnal |
| lizards | Sigaloseps_deplanchei | Sigaloseps_deplanchei | cathemeral |
| lizards | Sigaloseps_ruficauda | Sigaloseps_ruficauda | cathemeral |
| lizards | Silvascincus_murrayi | Silvascincus_murrayi | diurnal |
| lizards | Silvascincus_tryoni | Silvascincus_tryoni | diurnal |
| lizards | Simiscincus_aurantiacus | Simiscincus_aurantiacus | nocturnal |
| lizards | Sitana_fusca | Sitana_fusca | diurnal |
| lizards | Sitana_ponticeriana | Sitana_ponticeriana | diurnal |
| lizards | Sitana_sivalensis | Sitana_sivalensis | diurnal |
| lizards | Smaug_barbertonensis | Smaug_barbertonensis | diurnal |
| lizards | Smaug_breyeri | Smaug_breyeri | diurnal |
| lizards | Smaug_depressus | Smaug_depressus | diurnal |
| lizards | Smaug_giganteus | Smaug_giganteus | diurnal |
| lizards | Smaug_mossambicus | Smaug_mossambicus | diurnal |
| lizards | Smaug_regius | Smaug_regius | diurnal |
| lizards | Smaug_vandami | Smaug_vandami | diurnal |

| | | | |
|---|---|---|---|
| lizards | Smaug_warreni | Smaug_warreni | diurnal |
| lizards | Sphaerodactylus_altavelensis | Sphaerodactylus_altavelensis | diurnal |
| lizards | Sphaerodactylus_argus | Sphaerodactylus_argus | diurnal |
| lizards | Sphaerodactylus_cinereus | Sphaerodactylus_cinereus | nocturnal |
| lizards | Sphaerodactylus_clenchi | Sphaerodactylus_clenchi | diurnal |
| lizards | Sphaerodactylus_corticola | Sphaerodactylus_corticola | nocturnal |
| lizards | Sphaerodactylus_darlingtoni | Sphaerodactylus_darlingtoni | diurnal |
| lizards | Sphaerodactylus_difficilis | Sphaerodactylus_difficilis | diurnal |
| lizards | Sphaerodactylus_elegans | Sphaerodactylus_elegans | nocturnal |
| lizards | Sphaerodactylus_fantasticus | Sphaerodactylus_fantasticus | diurnal |
| lizards | Sphaerodactylus_glaucus | Sphaerodactylus_glaucus | cathemeral |
| lizards | Sphaerodactylus_homolepis | Sphaerodactylus_homolepis | diurnal |
| lizards | Sphaerodactylus_inaguae | Sphaerodactylus_inaguae | nocturnal |
| lizards | Sphaerodactylus_kirbyi | Sphaerodactylus_kirbyi | diurnal |
| lizards | Sphaerodactylus_klauberi | Sphaerodactylus_klauberi | diurnal |
| lizards | Sphaerodactylus_leucaster | Sphaerodactylus_leucaster | nocturnal |
| lizards | Sphaerodactylus_levinsi | Sphaerodactylus_levinsi | diurnal |
| lizards | Sphaerodactylus_macrolepis | Sphaerodactylus_macrolepis | diurnal |
| lizards | Sphaerodactylus_millepunctatus | Sphaerodactylus_millepunctatus | cathemeral |
| lizards | Sphaerodactylus_molei | Sphaerodactylus_molei | diurnal |
| lizards | Sphaerodactylus_nicholsi | Sphaerodactylus_nicholsi | cathemeral |
| lizards | Sphaerodactylus_notatus | Sphaerodactylus_notatus | diurnal |
| lizards | Sphaerodactylus_oliveri | Sphaerodactylus_oliveri | diurnal |
| lizards | Sphaerodactylus_pacificus | Sphaerodactylus_pacificus | nocturnal |
| lizards | Sphaerodactylus_parkeri | Sphaerodactylus_parkeri | nocturnal |
| lizards | Sphaerodactylus_parvus | Sphaerodactylus_parvus | nocturnal |
| lizards | Sphaerodactylus_richardsonii | Sphaerodactylus_richardsonii | diurnal |
| lizards | Sphaerodactylus_roosevelti | Sphaerodactylus_roosevelti | nocturnal |
| lizards | Sphaerodactylus_sabanus | Sphaerodactylus_sabanus | cathemeral |
| lizards | Sphaerodactylus_savagei | Sphaerodactylus_savagei | cathemeral |
| lizards | Sphaerodactylus_sputator | Sphaerodactylus_sputator | cathemeral |
| lizards | Sphaerodactylus_torrei | Sphaerodactylus_torrei | diurnal |
| lizards | Sphaerodactylus_townsendi | Sphaerodactylus_townsendi | diurnal |
| lizards | Sphaerodactylus_vincenti | Sphaerodactylus_vincenti | cathemeral |

| | | | |
|---|---|---|---|
| lizards | Sphenomorphus_aignanus | Sphenomorphus_aignanus | diurnal |
| lizards | Sphenomorphus_bacboensis | Sphenomorphus_bacboensis | diurnal |
| lizards | Sphenomorphus_bignelli | Sphenomorphus_bignelli | diurnal |
| lizards | Sphenomorphus_brunneus | Sphenomorphus_brunneus | nocturnal |
| lizards | Sphenomorphus_buenloicus | Sphenomorphus_buenloicus | diurnal |
| lizards | Sphenomorphus_concinnatus | Sphenomorphus_concinnatus | diurnal |
| lizards | Sphenomorphus_cranei | Sphenomorphus_cranei | diurnal |
| lizards | Sphenomorphus_cyanolaemus | Sphenomorphus_cyanolaemus | diurnal |
| lizards | Sphenomorphus_darlingtoni | Sphenomorphus_darlingtoni | nocturnal |
| lizards | Sphenomorphus_dussumieri | Sphenomorphus_dussumieri | diurnal |
| lizards | Sphenomorphus_fasciatus | Sphenomorphus_fasciatus | diurnal |
| lizards | Sphenomorphus_forbesi | Sphenomorphus_forbesi | nocturnal |
| lizards | Sphenomorphus_fragilis | Sphenomorphus_fragilis | nocturnal |
| lizards | Sphenomorphus_granulatus | Sphenomorphus_granulatus | nocturnal |
| lizards | Sphenomorphus_incognitus | Sphenomorphus_incognitus | diurnal |
| lizards | Sphenomorphus_indicus | Sphenomorphus_indicus | diurnal |
| lizards | Sphenomorphus_jobiensis | Sphenomorphus_jobiensis | cathemeral |
| lizards | Sphenomorphus_louisiadensis | Sphenomorphus_louisiadensis | diurnal |
| lizards | Sphenomorphus_maculatus | Sphenomorphus_maculatus | diurnal |
| lizards | Sphenomorphus_maindroni | Sphenomorphus_maindroni | nocturnal |
| lizards | Sphenomorphus_melanopogon | Sphenomorphus_melanopogon | cathemeral |
| lizards | Sphenomorphus_mimicus | Sphenomorphus_mimicus | diurnal |
| lizards | Sphenomorphus_minutus | Sphenomorphus_minutus | nocturnal |
| lizards | Sphenomorphus_multisquamatus | Sphenomorphus_multisquamatus | diurnal |
| lizards | Sphenomorphus_murudensis | Sphenomorphus_murudensis | diurnal |
| lizards | Sphenomorphus_necopinatus | Sphenomorphus_necopinatus | diurnal |
| lizards | Sphenomorphus_nigriventris | Sphenomorphus_nigriventris | nocturnal |
| lizards | Sphenomorphus_nigrolineatus | Sphenomorphus_nigrolineatus | cathemeral |
| lizards | Sphenomorphus_papuae | Sphenomorphus_papuae | nocturnal |
| lizards | Sphenomorphus_praesignis | Sphenomorphus_praesignis | diurnal |
| lizards | Sphenomorphus_pratti | Sphenomorphus_pratti | nocturnal |
| lizards | Sphenomorphus_sabanus | Sphenomorphus_sabanus | diurnal |
| lizards | Sphenomorphus_schlegeli | Sphenomorphus_schlegeli | diurnal |
| lizards | Sphenomorphus_schultzei | Sphenomorphus_schultzei | nocturnal |

| | | | |
|---|---|---|---|
| lizards | Sphenomorphus_scotophilus | Sphenomorphus_scotophilus | diurnal |
| lizards | Sphenomorphus_scutatus | Sphenomorphus_scutatus | diurnal |
| lizards | Sphenomorphus_simus | Sphenomorphus_simus | diurnal |
| lizards | Sphenomorphus_solomonis | Sphenomorphus_solomonis | cathemeral |
| lizards | Sphenomorphus_taiwanensis | Sphenomorphus_taiwanensis | diurnal |
| lizards | Sphenomorphus_undulatus | Sphenomorphus_undulatus | crepuscular |
| lizards | Sphenomorphus_vanheurni | Sphenomorphus_vanheurni | diurnal |
| lizards | Spondylurus_anegadae | Spondylurus_anegadae | diurnal |
| lizards | Spondylurus_caicosae | Spondylurus_caicosae | diurnal |
| lizards | Spondylurus_fulgidus | Spondylurus_fulgidus | diurnal |
| lizards | Spondylurus_monae | Spondylurus_monae | diurnal |
| lizards | Spondylurus_nitidus | Spondylurus_nitidus | diurnal |
| lizards | Spondylurus_powelli | Spondylurus_powelli | diurnal |
| lizards | Spondylurus_sloanii | Spondylurus_sloanii | diurnal |
| lizards | Spondylurus_turksae | Spondylurus_turksae | diurnal |
| lizards | Stenocercus_angel | Stenocercus_angel | diurnal |
| lizards | Stenocercus_azureus | Stenocercus_azureus | diurnal |
| lizards | Stenocercus_bolivarensis | Stenocercus_bolivarensis | diurnal |
| lizards | Stenocercus_cadlei | Stenocercus_cadlei | diurnal |
| lizards | Stenocercus_caducus | Stenocercus_caducus | diurnal |
| lizards | Stenocercus_chinchaoensis | Stenocercus_chinchaoensis | diurnal |
| lizards | Stenocercus_cupreus | Stenocercus_cupreus | diurnal |
| lizards | Stenocercus_dumerilii | Stenocercus_dumerilii | diurnal |
| lizards | Stenocercus_eunetopsis | Stenocercus_eunetopsis | diurnal |
| lizards | Stenocercus_guentheri | Stenocercus_guentheri | diurnal |
| lizards | Stenocercus_huancabambae | Stenocercus_huancabambae | diurnal |
| lizards | Stenocercus_humeralis | Stenocercus_humeralis | diurnal |
| lizards | Stenocercus_percultus | Stenocercus_percultus | diurnal |
| lizards | Stenocercus_prionotus | Stenocercus_prionotus | diurnal |
| lizards | Stenocercus_puyango | Stenocercus_puyango | diurnal |
| lizards | Stenocercus_roseiventris | Stenocercus_roseiventris | diurnal |
| lizards | Stenocercus_varius | Stenocercus_varius | diurnal |
| lizards | Stenodactylus_doriae | Stenodactylus_doriae | nocturnal |
| lizards | Stenodactylus_grandiceps | Stenodactylus_grandiceps | nocturnal |

| | | | |
|---|---|---|---|
| lizards | Stenodactylus_mauritanicus | Stenodactylus_mauritanicus | nocturnal |
| lizards | Stenodactylus_petrii | Stenodactylus_petrii | nocturnal |
| lizards | Stenodactylus_slevini | Stenodactylus_slevini | nocturnal |
| lizards | Strobilurus_torquatus | Strobilurus_torquatus | diurnal |
| lizards | Strophurus_assimilis | Strophurus_assimilis | nocturnal |
| lizards | Strophurus_ciliaris | Strophurus_ciliaris | nocturnal |
| lizards | Strophurus_elderi | Strophurus_elderi | nocturnal |
| lizards | Strophurus_intermedius | Strophurus_intermedius | nocturnal |
| lizards | Strophurus_jeanae | Strophurus_jeanae | nocturnal |
| lizards | Strophurus_krisalys | Strophurus_krisalys | nocturnal |
| lizards | Strophurus_mcmillani | Strophurus_mcmillani | nocturnal |
| lizards | Strophurus_michaelseni | Strophurus_michaelseni | nocturnal |
| lizards | Strophurus_rankini | Strophurus_rankini | nocturnal |
| lizards | Strophurus_spinigerus | Strophurus_spinigerus | nocturnal |
| lizards | Strophurus_strophurus | Strophurus_strophurus | nocturnal |
| lizards | Strophurus_taeniatus | Strophurus_taeniatus | nocturnal |
| lizards | Strophurus_taenicauda | Strophurus_taenicauda | nocturnal |
| lizards | Strophurus_williamsi | Strophurus_williamsi | nocturnal |
| lizards | Strophurus_wilsoni | Strophurus_wilsoni | nocturnal |
| lizards | Takydromus_amurensis | Takydromus_amurensis | diurnal |
| lizards | Takydromus_dorsalis | Takydromus_dorsalis | diurnal |
| lizards | Takydromus_formosanus | Takydromus_formosanus | diurnal |
| lizards | Takydromus_hsuehshanensis | Takydromus_hsuehshanensis | diurnal |
| lizards | Takydromus_khasiensis | Takydromus_khasiensis | diurnal |
| lizards | Takydromus_kuehnei | Takydromus_kuehnei | diurnal |
| lizards | Takydromus_luyeanus | Takydromus_luyeanus | diurnal |
| lizards | Takydromus_sauteri | Takydromus_sauteri | diurnal |
| lizards | Takydromus_septentrionalis | Takydromus_septentrionalis | diurnal |
| lizards | Takydromus_sexlineatus | Takydromus_sexlineatus | diurnal |
| lizards | Takydromus_smaragdinus | Takydromus_smaragdinus | diurnal |
| lizards | Takydromus_stejnegeri | Takydromus_stejnegeri | diurnal |
| lizards | Takydromus_sylvaticus | Takydromus_sylvaticus | diurnal |
| lizards | Takydromus_tachydromoides | Takydromus_tachydromoides | diurnal |
| lizards | Takydromus_toyamai | Takydromus_toyamai | diurnal |

| | | | |
|---|---|---|---|
| lizards | Takydromus_viridipunctatus | Takydromus_viridipunctatus | diurnal |
| lizards | Takydromus_wolteri | Takydromus_wolteri | diurnal |
| lizards | Tarentola_angustimentalis | Tarentola_angustimentalis | nocturnal |
| lizards | Tarentola_annularis | Tarentola_annularis | nocturnal |
| lizards | Tarentola_boehmei | Tarentola_boehmei | nocturnal |
| lizards | Tarentola_boettgeri | Tarentola_boettgeri | nocturnal |
| lizards | Tarentola_chazaliae | Tarentola_chazaliae | nocturnal |
| lizards | Tarentola_crombiei | Tarentola_crombiei | nocturnal |
| lizards | Tarentola_delalandii | Tarentola_delalandii | cathemeral |
| lizards | Tarentola_deserti | Tarentola_deserti | nocturnal |
| lizards | Tarentola_ephippiata | Tarentola_ephippiata | nocturnal |
| lizards | Tarentola_fascicularis | Tarentola_fascicularis | nocturnal |
| lizards | Tarentola_gigas | Tarentola_gigas | nocturnal |
| lizards | Tarentola_gomerensis | Tarentola_gomerensis | nocturnal |
| lizards | Tarentola_mauritanica | Tarentola_mauritanica | cathemeral |
| lizards | Tarentola_mindiae | Tarentola_mindiae | nocturnal |
| lizards | Tarentola_neglecta | Tarentola_neglecta | nocturnal |
| lizards | Techmarscincus_jigurru | Techmarscincus_jigurru | diurnal |
| lizards | Teira_dugesii | Teira_dugesii | diurnal |
| lizards | Teius_oculatus | Teius_oculatus | diurnal |
| lizards | Teius_suquiensis | Teius_suquiensis | diurnal |
| lizards | Teius_teyou | Teius_teyou | diurnal |
| lizards | Tenuidactylus_bogdanovi | Tenuidactylus_bogdanovi | nocturnal |
| lizards | Tenuidactylus_caspius | Tenuidactylus_caspius | cathemeral |
| lizards | Tenuidactylus_dadunensis | Tenuidactylus_dadunensis | nocturnal |
| lizards | Tenuidactylus_elongatus | Tenuidactylus_elongatus | nocturnal |
| lizards | Tenuidactylus_fedtschenkoi | Tenuidactylus_fedtschenkoi | cathemeral |
| lizards | Tenuidactylus_longipes | Tenuidactylus_longipes | cathemeral |
| lizards | Tenuidactylus_turcmenicus | Tenuidactylus_turcmenicus | nocturnal |
| lizards | Teratoscincus_bedriagai | Teratoscincus_bedriagai | nocturnal |
| lizards | Teratoscincus_microlepis | Teratoscincus_microlepis | nocturnal |
| lizards | Teratoscincus_przewalskii | Teratoscincus_przewalskii | nocturnal |
| lizards | Teratoscincus_roborowskii | Teratoscincus_roborowskii | nocturnal |
| lizards | Teratoscincus_scincus | Teratoscincus_scincus | nocturnal |

| | | | |
|---|---|---|---|
| lizards | Tetradactylus_africanus | Tetradactylus_africanus | diurnal |
| lizards | Tetradactylus_breyeri | Tetradactylus_breyeri | diurnal |
| lizards | Tetradactylus_ellenbergeri | Tetradactylus_ellenbergeri | diurnal |
| lizards | Tetradactylus_seps | Tetradactylus_seps | diurnal |
| lizards | Tetradactylus_tetradactylus | Tetradactylus_tetradactylus | diurnal |
| lizards | Tetradactylus_udzungwensis | Tetradactylus_udzungwensis | diurnal |
| lizards | Thecadactylus_oskrobapreinorum | Thecadactylus_oskrobapreinorum | nocturnal |
| lizards | Thecadactylus_rapicauda | Thecadactylus_rapicauda | nocturnal |
| lizards | Thecadactylus_solimoensis | Thecadactylus_solimoensis | nocturnal |
| lizards | Tiliqua_adelaidensis | Tiliqua_adelaidensis | diurnal |
| lizards | Tiliqua_gigas | Tiliqua_gigas | diurnal |
| lizards | Tiliqua_multifasciata | Tiliqua_multifasciata | cathemeral |
| lizards | Tiliqua_nigrolutea | Tiliqua_nigrolutea | diurnal |
| lizards | Tiliqua_occipitalis | Tiliqua_occipitalis | diurnal |
| lizards | Tiliqua_rugosa | Tiliqua_rugosa | diurnal |
| lizards | Tiliqua_scincoides | Tiliqua_scincoides | cathemeral |
| lizards | Timon_lepidus | Timon_lepidus | diurnal |
| lizards | Timon_pater | Timon_pater | diurnal |
| lizards | Timon_princeps | Timon_princeps | diurnal |
| lizards | Timon_tangitanus | Timon_tangitanus | diurnal |
| lizards | Toropuku_stephensi | Toropuku_stephensi | nocturnal |
| lizards | Tracheloptychus_madagascariensis | Tracheloptychus_madagascariensis | diurnal |
| lizards | Trachydactylus_spatalurus | Bunopus_spatalurus | nocturnal |
| lizards | Trachylepis_acutilabris | Trachylepis_acutilabris | diurnal |
| lizards | Trachylepis_affinis | Trachylepis_affinis | diurnal |
| lizards | Trachylepis_albilabris | Trachylepis_albilabris | diurnal |
| lizards | Trachylepis_bayonii | Trachylepis_bayonii | diurnal |
| lizards | Trachylepis_bocagii | Trachylepis_bocagii | diurnal |
| lizards | Trachylepis_boettgeri | Trachylepis_boettgeri | diurnal |
| lizards | Trachylepis_boulengeri | Trachylepis_boulengeri | diurnal |
| lizards | Trachylepis_brauni | Trachylepis_brauni | diurnal |
| lizards | Trachylepis_brevicollis | Trachylepis_brevicollis | cathemeral |
| lizards | Trachylepis_buettneri | Trachylepis_buettneri | diurnal |
| lizards | Trachylepis_capensis | Trachylepis_capensis | diurnal |

| | | | |
|---|---|---|---|
| lizards | Trachylepis_comorensis | Trachylepis_comorensis | diurnal |
| lizards | Trachylepis_depressa | Trachylepis_depressa | diurnal |
| lizards | Trachylepis_dichroma | Trachylepis_dichroma | diurnal |
| lizards | Trachylepis_gravenhorstii | Trachylepis_gravenhorstii | diurnal |
| lizards | Trachylepis_homalocephala | Trachylepis_homalocephala | diurnal |
| lizards | Trachylepis_infralineata | Trachylepis_infralineata | diurnal |
| lizards | Trachylepis_irregularis | Trachylepis_irregularis | diurnal |
| lizards | Trachylepis_lacertiformis | Trachylepis_lacertiformis | crepuscular |
| lizards | Trachylepis_lavarambo | Trachylepis_lavarambo | diurnal |
| lizards | Trachylepis_maculilabris | Trachylepis_maculilabris | diurnal |
| lizards | Trachylepis_madagascariensis | Trachylepis_madagascariensis | diurnal |
| lizards | Trachylepis_margaritifera | Trachylepis_margaritifera | diurnal |
| lizards | Trachylepis_megalura | Trachylepis_megalura | diurnal |
| lizards | Trachylepis_nganghae | Trachylepis_nganghae | diurnal |
| lizards | Trachylepis_occidentalis | Trachylepis_occidentalis | diurnal |
| lizards | Trachylepis_perrotetii | Trachylepis_perrotetii | diurnal |
| lizards | Trachylepis_planifrons | Trachylepis_planifrons | diurnal |
| lizards | Trachylepis_polytropis | Trachylepis_polytropis | diurnal |
| lizards | Trachylepis_pulcherrima | Trachylepis_pulcherrima | diurnal |
| lizards | Trachylepis_punctatissima | Trachylepis_punctatissima | diurnal |
| lizards | Trachylepis_sechellensis | Trachylepis_sechellensis | diurnal |
| lizards | Trachylepis_sparsa | Trachylepis_sparsa | diurnal |
| lizards | Trachylepis_spilogaster | Trachylepis_spilogaster | diurnal |
| lizards | Trachylepis_striata | Trachylepis_striata | diurnal |
| lizards | Trachylepis_sulcata | Trachylepis_sulcata | diurnal |
| lizards | Trachylepis_tavaratra | Trachylepis_tavaratra | diurnal |
| lizards | Trachylepis_tessellata | Trachylepis_tessellata | diurnal |
| lizards | Trachylepis_varia | Trachylepis_varia | diurnal |
| lizards | Trachylepis_variegata | Trachylepis_variegata | diurnal |
| lizards | Trachylepis_vato | Trachylepis_vato | diurnal |
| lizards | Trachylepis_wahlbergii | Trachylepis_angolensis | diurnal |
| lizards | Trachylepis_wrightii | Trachylepis_wrightii | diurnal |
| lizards | Trapelus_agilis | Trapelus_agilis | diurnal |
| lizards | Trapelus_agnetae | Trapelus_agnetae | diurnal |

| | | | |
|---|---|---|---|
| lizards | Trapelus_flavimaculatus | Trapelus_flavimaculatus | diurnal |
| lizards | Trapelus_megalonyx | Trapelus_megalonyx | diurnal |
| lizards | Trapelus_mutabilis | Trapelus_mutabilis | diurnal |
| lizards | Trapelus_rubrigularis | Trapelus_rubrigularis | diurnal |
| lizards | Trapelus_ruderatus | Trapelus_ruderatus | diurnal |
| lizards | Trapelus_sanguinolentus | Trapelus_sanguinolentus | diurnal |
| lizards | Trapelus_savignii | Trapelus_savignii | diurnal |
| lizards | Trapelus_tournevillei | Trapelus_tournevillei | diurnal |
| lizards | Tribolonotus_blanchardi | Tribolonotus_blanchardi | diurnal |
| lizards | Tribolonotus_gracilis | Tribolonotus_gracilis | diurnal |
| lizards | Tribolonotus_novaeguineae | Tribolonotus_novaeguineae | diurnal |
| lizards | Tribolonotus_ponceleti | Tribolonotus_ponceleti | nocturnal |
| lizards | Tribolonotus_pseudoponceleti | Tribolonotus_pseudoponceleti | diurnal |
| lizards | Tribolonotus_schmidti | Tribolonotus_schmidti | diurnal |
| lizards | Trigonodactylus_arabicus | Stenodactylus_arabicus | nocturnal |
| lizards | Trioceros_affinis | Trioceros_affinis | diurnal |
| lizards | Trioceros_balebicornutus | Trioceros_balebicornutus | diurnal |
| lizards | Trioceros_bitaeniatus | Trioceros_bitaeniatus | diurnal |
| lizards | Trioceros_conirostratus | Trioceros_conirostratus | diurnal |
| lizards | Trioceros_cristatus | Trioceros_cristatus | diurnal |
| lizards | Trioceros_ellioti | Trioceros_ellioti | diurnal |
| lizards | Trioceros_hanangensis | Trioceros_hanangensis | diurnal |
| lizards | Trioceros_harennae | Trioceros_harennae | diurnal |
| lizards | Trioceros_hoehnelii | Trioceros_hoehnelii | diurnal |
| lizards | Trioceros_ituriensis | Trioceros_ituriensis | diurnal |
| lizards | Trioceros_jacksonii | Trioceros_jacksonii | diurnal |
| lizards | Trioceros_laterispinis | Trioceros_laterispinis | diurnal |
| lizards | Trioceros_melleri | Trioceros_melleri | diurnal |
| lizards | Trioceros_montium | Trioceros_montium | diurnal |
| lizards | Trioceros_ntunte | Trioceros_ntunte | diurnal |
| lizards | Trioceros_oweni | Trioceros_oweni | diurnal |
| lizards | Trioceros_quadricornis | Trioceros_quadricornis | diurnal |
| lizards | Trioceros_rudis | Trioceros_rudis | diurnal |
| lizards | Trioceros_schubotzi | Trioceros_schubotzi | diurnal |

| | | | |
|---|---|---|---|
| lizards | Trioceros_sternfeldi | Trioceros_sternfeldi | diurnal |
| lizards | Trogonophis_wiegmanni | Trogonophis_wiegmanni | cathemeral |
| lizards | Tropicagama_temporalis | Lophognathus_temporalis | diurnal |
| lizards | Tropidophorus_baviensis | Tropidophorus_baviensis | cathemeral |
| lizards | Tropidophorus_beccarii | Tropidophorus_beccarii | diurnal |
| lizards | Tropidophorus_berdmorei | Tropidophorus_berdmorei | diurnal |
| lizards | Tropidophorus_brookei | Tropidophorus_brookei | diurnal |
| lizards | Tropidophorus_cocincinensis | Tropidophorus_cocincinensis | cathemeral |
| lizards | Tropidophorus_davaoensis | Tropidophorus_davaoensis | diurnal |
| lizards | Tropidophorus_grayi | Tropidophorus_grayi | diurnal |
| lizards | Tropidophorus_hainanus | Tropidophorus_hainanus | diurnal |
| lizards | Tropidophorus_laotus | Tropidophorus_laotus | cathemeral |
| lizards | Tropidophorus_microlepis | Tropidophorus_microlepis | cathemeral |
| lizards | Tropidophorus_micropus | Tropidophorus_micropus | diurnal |
| lizards | Tropidophorus_misaminius | Tropidophorus_misaminius | cathemeral |
| lizards | Tropidophorus_mocquardii | Tropidophorus_mocquardii | diurnal |
| lizards | Tropidophorus_murphyi | Tropidophorus_murphyi | nocturnal |
| lizards | Tropidophorus_noggei | Tropidophorus_noggei | nocturnal |
| lizards | Tropidophorus_partelloi | Tropidophorus_partelloi | cathemeral |
| lizards | Tropidophorus_robinsoni | Tropidophorus_robinsoni | diurnal |
| lizards | Tropidophorus_sinicus | Tropidophorus_sinicus | nocturnal |
| lizards | Tropidosaura_gularis | Tropidosaura_gularis | diurnal |
| lizards | Tropidosaura_montana | Tropidosaura_montana | diurnal |
| lizards | Tropidoscincus_boreus | Tropidoscincus_boreus | diurnal |
| lizards | Tropidoscincus_variabilis | Tropidoscincus_variabilis | diurnal |
| lizards | Tropidurus_bogerti | Tropidurus_bogerti | diurnal |
| lizards | Tropidurus_callathelys | Tropidurus_callathelys | diurnal |
| lizards | Tropidurus_catalanensis | Tropidurus_catalanensis | diurnal |
| lizards | Tropidurus_chromatops | Tropidurus_chromatops | diurnal |
| lizards | Tropidurus_cocorobensis | Tropidurus_cocorobensis | diurnal |
| lizards | Tropidurus_erythrocephalus | Tropidurus_erythrocephalus | diurnal |
| lizards | Tropidurus_etheridgei | Tropidurus_etheridgei | diurnal |
| lizards | Tropidurus_guarani | Tropidurus_guarani | diurnal |
| lizards | Tropidurus_hispidus | Tropidurus_hispidus | diurnal |

| | | | |
|---|---|---|---|
| lizards | Tropidurus_hygomi | Tropidurus_hygomi | diurnal |
| lizards | Tropidurus_itambere | Tropidurus_itambere | diurnal |
| lizards | Tropidurus_jaguaribanus | Tropidurus_jaguaribanus | diurnal |
| lizards | Tropidurus_montanus | Tropidurus_montanus | diurnal |
| lizards | Tropidurus_oreadicus | Tropidurus_oreadicus | diurnal |
| lizards | Tropidurus_pinima | Tropidurus_pinima | diurnal |
| lizards | Tropidurus_psammonastes | Tropidurus_psammonastes | diurnal |
| lizards | Tropidurus_semitaeniatus | Tropidurus_semitaeniatus | diurnal |
| lizards | Tropidurus_spinulosus | Tropidurus_spinulosus | diurnal |
| lizards | Tropidurus_torquatus | Tropidurus_torquatus | diurnal |
| lizards | Tropidurus_xanthochilus | Tropidurus_xanthochilus | diurnal |
| lizards | Tropiocolotes_algericus | Tropiocolotes_algericus | nocturnal |
| lizards | Tropiocolotes_nattereri | Tropiocolotes_nattereri | nocturnal |
| lizards | Tropiocolotes_nubicus | Tropiocolotes_nubicus | nocturnal |
| lizards | Tropiocolotes_somalicus | Tropiocolotes_somalicus | nocturnal |
| lizards | Tropiocolotes_steudneri | Tropiocolotes_steudneri | nocturnal |
| lizards | Tropiocolotes_tripolitanus | Tropiocolotes_tripolitanus | nocturnal |
| lizards | Tukutuku_rakiurae | Tukutuku_rakiurae | cathemeral |
| lizards | Tumbunascincus_luteilateralis | Tumbunascincus_luteilateralis | diurnal |
| lizards | Tupinambis_teguixin | Tupinambis_teguixin | diurnal |
| lizards | Tympanocryptis_centralis | Tympanocryptis_centralis | diurnal |
| lizards | Tympanocryptis_cephalus | Tympanocryptis_cephalus | diurnal |
| lizards | Tympanocryptis_intima | Tympanocryptis_intima | diurnal |
| lizards | Tympanocryptis_lineata | Tympanocryptis_lineata | diurnal |
| lizards | Tympanocryptis_pinguicolla | Tympanocryptis_pinguicolla | diurnal |
| lizards | Tympanocryptis_tetraporophora | Tympanocryptis_tetraporophora | diurnal |
| lizards | Typhlacontias_brevipes | Typhlacontias_brevipes | cathemeral |
| lizards | Typhlacontias_rohani | Typhlacontias_rohani | cathemeral |
| lizards | Tytthoscincus_aesculeticola | Tytthoscincus_aesculeticola | diurnal |
| lizards | Tytthoscincus_bukitensis | Sphenomorphus_bukitensis | diurnal |
| lizards | Tytthoscincus_temmincki | Sphenomorphus_temmincki | diurnal |
| lizards | Uma_cowlesi | Uma_rufopunctata | diurnal |
| lizards | Uma_exsul | Uma_exsul | diurnal |
| lizards | Uma_inornata | Uma_inornata | diurnal |

| | | | |
|---|---|---|---|
| lizards | Uma_notata | Uma_notata | diurnal |
| lizards | Uma_paraphygas | Uma_paraphygas | diurnal |
| lizards | Uma_scoparia | Uma_scoparia | diurnal |
| lizards | Underwoodisaurus_milii | Underwoodisaurus_milii | nocturnal |
| lizards | Underwoodisaurus_seorsus | Underwoodisaurus_seorsus | nocturnal |
| lizards | Uracentron_azureum | Uracentron_azureum | diurnal |
| lizards | Uracentron_flaviceps | Uracentron_flaviceps | diurnal |
| lizards | Uranoscodon_superciliosus | Uranoscodon_superciliosus | diurnal |
| lizards | Urocotyledon_inexpectata | Urocotyledon_inexpectata | nocturnal |
| lizards | Urocotyledon_rasmusseni | Urocotyledon_rasmusseni | nocturnal |
| lizards | Urocotyledon_wolterstorffi | Urocotyledon_wolterstorffi | nocturnal |
| lizards | Uromastyx_acanthinura | Uromastyx_acanthinura | diurnal |
| lizards | Uromastyx_aegyptia | Uromastyx_aegyptia | diurnal |
| lizards | Uromastyx_alfredschmidti | Uromastyx_alfredschmidti | diurnal |
| lizards | Uromastyx_benti | Uromastyx_benti | diurnal |
| lizards | Uromastyx_dispar | Uromastyx_dispar | diurnal |
| lizards | Uromastyx_geyri | Uromastyx_geyri | diurnal |
| lizards | Uromastyx_macfadyeni | Uromastyx_macfadyeni | diurnal |
| lizards | Uromastyx_ocellata | Uromastyx_ocellata | diurnal |
| lizards | Uromastyx_ornata | Uromastyx_ornata | diurnal |
| lizards | Uromastyx_thomasi | Uromastyx_thomasi | diurnal |
| lizards | Uroplatus_alluaudi | Uroplatus_alluaudi | nocturnal |
| lizards | Uroplatus_ebenaui | Uroplatus_ebenaui | nocturnal |
| lizards | Uroplatus_fimbriatus | Uroplatus_fimbriatus | nocturnal |
| lizards | Uroplatus_guentheri | Uroplatus_guentheri | nocturnal |
| lizards | Uroplatus_henkeli | Uroplatus_henkeli | nocturnal |
| lizards | Uroplatus_lineatus | Uroplatus_lineatus | nocturnal |
| lizards | Uroplatus_malahelo | Uroplatus_malahelo | nocturnal |
| lizards | Uroplatus_phantasticus | Uroplatus_phantasticus | nocturnal |
| lizards | Uroplatus_pietschmanni | Uroplatus_pietschmanni | nocturnal |
| lizards | Uroplatus_sameiti | Uroplatus_sameiti | nocturnal |
| lizards | Uroplatus_sikorae | Uroplatus_sikorae | nocturnal |
| lizards | Urosaurus_bicarinatus | Urosaurus_bicarinatus | diurnal |
| lizards | Urosaurus_graciosus | Urosaurus_graciosus | diurnal |

| | | | |
|---|---|---|---|
| lizards | Urosaurus_ornatus | Urosaurus_ornatus | diurnal |
| lizards | Urostrophus_gallardoi | Urostrophus_gallardoi | diurnal |
| lizards | Urostrophus_vautieri | Urostrophus_vautieri | diurnal |
| lizards | Uta_encantadae | Uta_encantadae | diurnal |
| lizards | Uta_nolascensis | Uta_nolascensis | diurnal |
| lizards | Uta_palmeri | Uta_palmeri | diurnal |
| lizards | Uta_squamata | Uta_squamata | diurnal |
| lizards | Uta_stansburiana | Uta_stansburiana | diurnal |
| lizards | Uvidicolus_sphyrurus | Uvidicolus_sphyrurus | nocturnal |
| lizards | Vanzosaura_multiscutata | Vanzosaura_multiscutatus | diurnal |
| lizards | Vanzosaura_rubricauda | Vanzosaura_rubricauda | diurnal |
| lizards | Vanzosaura_savanicola | Vanzosaura_savanicola | diurnal |
| lizards | Varanus_acanthurus | Varanus_acanthurus | diurnal |
| lizards | Varanus_albigularis | Varanus_albigularis | diurnal |
| lizards | Varanus_auffenbergi | Varanus_auffenbergi | diurnal |
| lizards | Varanus_beccarii | Varanus_beccarii | diurnal |
| lizards | Varanus_bengalensis | Varanus_bengalensis | diurnal |
| lizards | Varanus_boehmei | Varanus_boehmei | diurnal |
| lizards | Varanus_bogerti | Varanus_bogerti | diurnal |
| lizards | Varanus_brevicauda | Varanus_brevicauda | diurnal |
| lizards | Varanus_caerulivirens | Varanus_caerulivirens | diurnal |
| lizards | Varanus_caudolineatus | Varanus_caudolineatus | diurnal |
| lizards | Varanus_cumingi | Varanus_cumingi | diurnal |
| lizards | Varanus_doreanus | Varanus_doreanus | diurnal |
| lizards | Varanus_dumerilii | Varanus_dumerilii | cathemeral |
| lizards | Varanus_eremius | Varanus_eremius | diurnal |
| lizards | Varanus_exanthematicus | Varanus_exanthematicus | diurnal |
| lizards | Varanus_finschi | Varanus_finschi | diurnal |
| lizards | Varanus_flavescens | Varanus_flavescens | diurnal |
| lizards | Varanus_giganteus | Varanus_giganteus | diurnal |
| lizards | Varanus_gilleni | Varanus_gilleni | diurnal |
| lizards | Varanus_glauerti | Varanus_glauerti | diurnal |
| lizards | Varanus_glebopalma | Varanus_glebopalma | diurnal |
| lizards | Varanus_gouldii | Varanus_gouldii | diurnal |

| | | | |
|---|---|---|---|
| lizards | Varanus_griseus | Varanus_griseus | diurnal |
| lizards | Varanus_indicus | Varanus_indicus | diurnal |
| lizards | Varanus_jobiensis | Varanus_jobiensis | diurnal |
| lizards | Varanus_juxtindicus | Varanus_juxtindicus | diurnal |
| lizards | Varanus_keithhornei | Varanus_keithhornei | diurnal |
| lizards | Varanus_kingorum | Varanus_kingorum | diurnal |
| lizards | Varanus_komodoensis | Varanus_komodoensis | diurnal |
| lizards | Varanus_kordensis | Varanus_kordensis | diurnal |
| lizards | Varanus_lirungensis | Varanus_lirungensis | diurnal |
| lizards | Varanus_mabitang | Varanus_mabitang | diurnal |
| lizards | Varanus_macraei | Varanus_macraei | diurnal |
| lizards | Varanus_marmoratus | Varanus_marmoratus | diurnal |
| lizards | Varanus_melinus | Varanus_melinus | diurnal |
| lizards | Varanus_mertensi | Varanus_mertensi | diurnal |
| lizards | Varanus_mitchelli | Varanus_mitchelli | diurnal |
| lizards | Varanus_nebulosus | Varanus_nebulosus | diurnal |
| lizards | Varanus_niloticus | Varanus_niloticus | diurnal |
| lizards | Varanus_nuchalis | Varanus_nuchalis | diurnal |
| lizards | Varanus_olivaceus | Varanus_olivaceus | diurnal |
| lizards | Varanus_panoptes | Varanus_panoptes | diurnal |
| lizards | Varanus_pilbarensis | Varanus_pilbarensis | diurnal |
| lizards | Varanus_prasinus | Varanus_prasinus | diurnal |
| lizards | Varanus_primordius | Varanus_primordius | diurnal |
| lizards | Varanus_rosenbergi | Varanus_rosenbergi | diurnal |
| lizards | Varanus_rudicollis | Varanus_rudicollis | diurnal |
| lizards | Varanus_salvadorii | Varanus_salvadorii | diurnal |
| lizards | Varanus_salvator | Varanus_salvator | diurnal |
| lizards | Varanus_scalaris | Varanus_scalaris | diurnal |
| lizards | Varanus_semiremex | Varanus_semiremex | diurnal |
| lizards | Varanus_similis | Varanus_similis | diurnal |
| lizards | Varanus_spenceri | Varanus_spenceri | diurnal |
| lizards | Varanus_spinulosus | Varanus_spinulosus | diurnal |
| lizards | Varanus_storri | Varanus_storri | diurnal |
| lizards | Varanus_timorensis | Varanus_timorensis | diurnal |

| | | | |
|---|---|---|---|
| lizards | Varanus_togianus | Varanus_togianus | diurnal |
| lizards | Varanus_tristis | Varanus_tristis | diurnal |
| lizards | Varanus_varius | Varanus_varius | diurnal |
| lizards | Varanus_yemenensis | Varanus_yemenensis | diurnal |
| lizards | Varzea_bistriata | Varzea_bistriata | diurnal |
| lizards | Vhembelacerta_rupicola | Vhembelacerta_rupicola | diurnal |
| lizards | Woodworthia_brunnea | Woodworthia_brunneus | nocturnal |
| lizards | Woodworthia_chrysosiretica | Woodworthia_chrysosireticus | nocturnal |
| lizards | Woodworthia_maculata | Woodworthia_maculatus | cathemeral |
| lizards | Xantusia_arizonae | Xantusia_arizonae | cathemeral |
| lizards | Xantusia_bezyi | Xantusia_bezyi | diurnal |
| lizards | Xantusia_extorris | Xantusia_extorris | nocturnal |
| lizards | Xantusia_gracilis | Xantusia_gracilis | nocturnal |
| lizards | Xantusia_henshawi | Xantusia_henshawi | cathemeral |
| lizards | Xantusia_riversiana | Xantusia_riversiana | diurnal |
| lizards | Xantusia_sanchezi | Xantusia_sanchezi | cathemeral |
| lizards | Xantusia_vigilis | Xantusia_vigilis | cathemeral |
| lizards | Xenagama_batillifera | Xenagama_batillifera | diurnal |
| lizards | Xenagama_taylori | Xenagama_taylori | diurnal |
| lizards | Xenosaurus_agrenon | Xenosaurus_agrenon | diurnal |
| lizards | Xenosaurus_grandis | Xenosaurus_grandis | diurnal |
| lizards | Xenosaurus_mendozai | Xenosaurus_mendozai | diurnal |
| lizards | Xenosaurus_newmanorum | Xenosaurus_newmanorum | diurnal |
| lizards | Xenosaurus_phalaroanthereon | Xenosaurus_phalaroanthereon | cathemeral |
| lizards | Xenosaurus_platyceps | Xenosaurus_platyceps | diurnal |
| lizards | Xenosaurus_rectocollaris | Xenosaurus_rectocollaris | diurnal |
| lizards | Xenosaurus_tzacualtipantecus | Xenosaurus_tzacualtipantecus | diurnal |
| lizards | Zonosaurus_anelanelany | Zonosaurus_anelanelany | diurnal |
| lizards | Zonosaurus_bemaraha | Zonosaurus_bemaraha | diurnal |
| lizards | Zonosaurus_haraldmeieri | Zonosaurus_haraldmeieri | diurnal |
| lizards | Zonosaurus_karsteni | Zonosaurus_karsteni | diurnal |
| lizards | Zonosaurus_rufipes | Zonosaurus_rufipes | diurnal |
| lizards | Zootoca_vivipara | Zootoca_vivipara | diurnal |
| lizards | Zygaspis_quadrifrons | Zygaspis_quadrifrons | diurnal |

| | | | |
|---|---|---|---|
| mammals | Abditomys_latidens | Abditomys_latidens | nocturnal |
| mammals | Abeomelomys_sevia | Abeomelomys_sevia | nocturnal |
| mammals | Abrawayaomys_ruschii | Abrawayaomys_ruschii | nocturnal |
| mammals | Abrocoma_bennettii | Abrocoma_bennettii | nocturnal |
| mammals | Abrocoma_boliviensis | Abrocoma_boliviensis | diurnal |
| mammals | Abrocoma_budini | Abrocoma_budini | nocturnal |
| mammals | Abrocoma_cinerea | Abrocoma_cinerea | diurnal |
| mammals | Abrocoma_famatina | Abrocoma_famatina | nocturnal |
| mammals | Abrocoma_schistacea | Abrocoma_shistacea | diurnal |
| mammals | Abrocoma_uspallata | Abrocoma_uspallata | diurnal |
| mammals | Abrocoma_vaccarum | Abrocoma_vaccarum | nocturnal |
| mammals | Abrothrix_andina | Abrothrix_andinus | diurnal |
| mammals | Abrothrix_illutea | Abrothrix_illutea | cathemeral |
| mammals | Abrothrix_jelskii | Abrothrix_jelskii | cathemeral |
| mammals | Abrothrix_lanosa | Abrothrix_lanosus | nocturnal |
| mammals | Abrothrix_longipilis | Abrothrix_longipilis | cathemeral |
| mammals | Abrothrix_olivacea | Abrothrix_olivaceus | cathemeral |
| mammals | Abrothrix_sanborni | Abrothrix_sanborni | cathemeral |
| mammals | Acerodon_celebensis | Acerodon_celebensis | nocturnal |
| mammals | Acerodon_humilis | Acerodon_humilis | nocturnal |
| mammals | Acerodon_jubatus | Acerodon_jubatus | diurnal |
| mammals | Acerodon_leucotis | Acerodon_leucotis | nocturnal |
| mammals | Acerodon_macklotii | Acerodon_mackloti | nocturnal |
| mammals | Acinonyx_jubatus | Acinonyx_jubatus | cathemeral |
| mammals | Acomys_cahirinus | Acomys_cahirinus | nocturnal |
| mammals | Acomys_chudeaui | Acomys_airensis | nocturnal |
| mammals | Acomys_cilicicus | Acomys_cilicicus | nocturnal |
| mammals | Acomys_cineraceus | Acomys_cineraceus | nocturnal |
| mammals | Acomys_dimidiatus | Acomys_dimidiatus | nocturnal |
| mammals | Acomys_ignitus | Acomys_ignitus | nocturnal |
| mammals | Acomys_johannis | Acomys_johannis | nocturnal |
| mammals | Acomys_kempi | Acomys_kempi | nocturnal |
| mammals | Acomys_louisae | Acomys_louisae | nocturnal |
| mammals | Acomys_minous | Acomys_minous | nocturnal |

| | | | |
|---|---|---|---|
| mammals | Acomys_mullah | Acomys_mullah | nocturnal |
| mammals | Acomys_nesiotes | Acomys_nesiotes | nocturnal |
| mammals | Acomys_percivali | Acomys_percivali | nocturnal |
| mammals | Acomys_russatus | Acomys_russatus | diurnal |
| mammals | Acomys_seurati | Acomys_seurati | nocturnal |
| mammals | Acomys_spinosissimus | Acomys_spinosissimus | nocturnal |
| mammals | Acomys_subspinosus | Acomys_subspinosus | nocturnal |
| mammals | Acomys_wilsoni | Acomys_wilsoni | nocturnal |
| mammals | Aconaemys_fuscus | Aconaemys_fuscus | nocturnal |
| mammals | Aconaemys_porteri | Aconaemys_porteri | nocturnal |
| mammals | Acrobates_pygmaeus | Acrobates_pygmaeus | nocturnal |
| mammals | Addax_nasomaculatus | Addax_nasomaculatus | diurnal |
| mammals | Aegialomys_xanthaeolus | Aegialomys_xanthaeolus | nocturnal |
| mammals | Aepeomys_lugens | Aepeomys_lugens | nocturnal |
| mammals | Aepeomys_reigi | Aepeomys_reigi | nocturnal |
| mammals | Aepyceros_melampus | Aepyceros_melampus | cathemeral |
| mammals | Aepyprymnus_rufescens | Aepyprymnus_rufescens | nocturnal |
| mammals | Aeretes_melanopterus | Aeretes_melanopterus | nocturnal |
| mammals | Aeromys_tephromelas | Aeromys_tephromelas | nocturnal |
| mammals | Aeromys_thomasi | Aeromys_thomasi | nocturnal |
| mammals | Aethalops_aequalis | Aethalops_aequalis | nocturnal |
| mammals | Aethalops_alecto | Aethalops_alecto | nocturnal |
| mammals | Aethomys_bocagei | Aethomys_bocagei | nocturnal |
| mammals | Aethomys_chrysophilus | Aethomys_chrysophilus | nocturnal |
| mammals | Aethomys_hindei | Aethomys_hindei | cathemeral |
| mammals | Aethomys_ineptus | Aethomys_ineptus | nocturnal |
| mammals | Aethomys_kaiseri | Aethomys_kaiseri | nocturnal |
| mammals | Aethomys_nyikae | Aethomys_nyikae | nocturnal |
| mammals | Aethomys_silindensis | Aethomys_silindensis | nocturnal |
| mammals | Aethomys_stannarius | Aethomys_stannarius | nocturnal |
| mammals | Aethomys_thomasi | Aethomys_thomasi | nocturnal |
| mammals | Afronycteris_helios | Neoromicia_helios | nocturnal |
| mammals | Afronycteris_nanus | Neoromicia_nana | nocturnal |
| mammals | Ailuropoda_melanoleuca | Ailuropoda_melanoleuca | cathemeral |

| | | | |
|---|---|---|---|
| mammals | Ailurops_melanotis | Ailurops_melanotis | nocturnal |
| mammals | Ailurops_ursinus | Ailurops_ursinus | diurnal |
| mammals | Ailurus_fulgens | Ailurus_fulgens | cathemeral |
| mammals | Akodon_aerosus | Akodon_aerosus | nocturnal |
| mammals | Akodon_affinis | Akodon_affinis | diurnal |
| mammals | Akodon_albiventer | Akodon_albiventer | diurnal |
| mammals | Akodon_azarae | Akodon_azarae | cathemeral |
| mammals | Akodon_boliviensis | Akodon_boliviensis | cathemeral |
| mammals | Akodon_budini | Akodon_budini | nocturnal |
| mammals | Akodon_caenosus | Akodon_aliquantulus | cathemeral |
| mammals | Akodon_cursor | Akodon_cursor | nocturnal |
| mammals | Akodon_dayi | Akodon_dayi | cathemeral |
| mammals | Akodon_dolores | Akodon_dolores | nocturnal |
| mammals | Akodon_fumeus | Akodon_fumeus | cathemeral |
| mammals | Akodon_iniscatus | Akodon_iniscatus | nocturnal |
| mammals | Akodon_juninensis | Akodon_juninensis | cathemeral |
| mammals | Akodon_kofordi | Akodon_kofordi | cathemeral |
| mammals | Akodon_lindberghi | Akodon_lindberghi | nocturnal |
| mammals | Akodon_lutescens | Akodon_lutescens | cathemeral |
| mammals | Akodon_mimus | Akodon_mimus | cathemeral |
| mammals | Akodon_mollis | Akodon_mollis | nocturnal |
| mammals | Akodon_montensis | Akodon_montensis | nocturnal |
| mammals | Akodon_mystax | Akodon_mystax | cathemeral |
| mammals | Akodon_orophilus | Akodon_orophilus | nocturnal |
| mammals | Akodon_paranaensis | Akodon_paranaensis | cathemeral |
| mammals | Akodon_pervalens | Akodon_pervalens | cathemeral |
| mammals | Akodon_philipmyersi | Akodon_philipmyersi | nocturnal |
| mammals | Akodon_reigi | Akodon_reigi | nocturnal |
| mammals | Akodon_sanctipaulensis | Akodon_sanctipaulensis | cathemeral |
| mammals | Akodon_siberiae | Akodon_siberiae | cathemeral |
| mammals | Akodon_simulator | Akodon_simulator | nocturnal |
| mammals | Akodon_spegazzinii | Akodon_spegazzinii | nocturnal |
| mammals | Akodon_subfuscus | Akodon_subfuscus | cathemeral |
| mammals | Akodon_surdus | Akodon_surdus | cathemeral |

| | | | |
|---|---|---|---|
| mammals | Akodon_sylvanus | Akodon_sylvanus | cathemeral |
| mammals | Akodon_toba | Akodon_toba | cathemeral |
| mammals | Akodon_torques | Akodon_torques | cathemeral |
| mammals | Akodon_varius | Akodon_varius | cathemeral |
| mammals | Alcelaphus_buselaphus | Alcelaphus_buselaphus | diurnal |
| mammals | Alces_alces | Alces_alces | cathemeral |
| mammals | Alexandromys_evoronensis | Microtus_evoronensis | cathemeral |
| mammals | Alexandromys_fortis | Microtus_fortis | cathemeral |
| mammals | Alexandromys_kikuchii | Microtus_kikuchii | crepuscular |
| mammals | Alexandromys_limnophilus | Microtus_limnophilus | cathemeral |
| mammals | Alexandromys_maximowiczii | Microtus_maximowiczii | cathemeral |
| mammals | Alexandromys_middendorffii | Microtus_middendorffii | cathemeral |
| mammals | Alexandromys_mongolicus | Microtus_mongolicus | cathemeral |
| mammals | Alexandromys_montebelli | Microtus_montebelli | nocturnal |
| mammals | Alexandromys_mujanensis | Microtus_mujanensis | cathemeral |
| mammals | Alexandromys_oeconomus | Microtus_oeconomus | cathemeral |
| mammals | Alexandromys_sachalinensis | Microtus_sachalinensis | cathemeral |
| mammals | Alionycteris_paucidentata | Alionycteris_paucidentata | nocturnal |
| mammals | Allactaga_major | Allactaga_major | nocturnal |
| mammals | Allactaga_severtzovi | Allactaga_severtzovi | nocturnal |
| mammals | Allactodipus_bobrinskii | Allactodipus_bobrinskii | nocturnal |
| mammals | Allenopithecus_nigroviridis | Allenopithecus_nigroviridis | diurnal |
| mammals | Allocebus_trichotis | Allocebus_trichotis | nocturnal |
| mammals | Allochrocebus_lhoesti | Allochrocebus_lhoesti | diurnal |
| mammals | Allochrocebus_preussi | Allochrocebus_preussi | diurnal |
| mammals | Allochrocebus_solatus | Allochrocebus_solatus | diurnal |
| mammals | Allocricetulus_curtatus | Allocricetulus_curtatus | nocturnal |
| mammals | Allocricetulus_eversmanni | Allocricetulus_eversmanni | nocturnal |
| mammals | Alouatta_arctoidea | Alouatta_arctoidea | diurnal |
| mammals | Alouatta_belzebul | Alouatta_belzebul | diurnal |
| mammals | Alouatta_caraya | Alouatta_caraya | diurnal |
| mammals | Alouatta_discolor | Alouatta_discolor | diurnal |
| mammals | Alouatta_guariba | Alouatta_guariba | diurnal |
| mammals | Alouatta_macconnelli | Alouatta_macconnelli | diurnal |

| | | | |
|---|---|---|---|
| mammals | Alouatta_nigerrima | Alouatta_nigerrima | diurnal |
| mammals | Alouatta_palliata | Alouatta_palliata | diurnal |
| mammals | Alouatta_pigra | Alouatta_pigra | diurnal |
| mammals | Alouatta_sara | Alouatta_sara | diurnal |
| mammals | Alticola_albicauda | Alticola_albicaudus | cathemeral |
| mammals | Alticola_argentatus | Alticola_argentatus | diurnal |
| mammals | Alticola_barakshin | Alticola_barakshin | nocturnal |
| mammals | Alticola_lemminus | Alticola_lemminus | cathemeral |
| mammals | Alticola_macrotis | Alticola_macrotis | cathemeral |
| mammals | Alticola_montosus | Alticola_montosa | diurnal |
| mammals | Alticola_olchonensis | Alticola_olchonensis | cathemeral |
| mammals | Alticola_roylei | Alticola_roylei | diurnal |
| mammals | Alticola_semicanus | Alticola_semicanus | nocturnal |
| mammals | Alticola_stoliczkanus | Alticola_stoliczkanus | nocturnal |
| mammals | Alticola_strelzovi | Alticola_strelzowi | cathemeral |
| mammals | Alticola_tuvinicus | Alticola_tuvinicus | cathemeral |
| mammals | Ametrida_centurio | Ametrida_centurio | nocturnal |
| mammals | Ammodillus_imbellis | Ammodillus_imbellis | nocturnal |
| mammals | Ammodorcas_clarkei | Ammodorcas_clarkei | diurnal |
| mammals | Ammospermophilus_harrisii | Ammospermophilus_harrisii | diurnal |
| mammals | Ammospermophilus_interpres | Ammospermophilus_interpres | diurnal |
| mammals | Ammospermophilus_leucurus | Ammospermophilus_leucurus | diurnal |
| mammals | Ammospermophilus_nelsoni | Ammospermophilus_nelsoni | diurnal |
| mammals | Ammotragus_lervia | Ammotragus_lervia | diurnal |
| mammals | Amorphochilus_schnablii | Amorphochilus_schnablii | nocturnal |
| mammals | Anathana_ellioti | Anathana_ellioti | diurnal |
| mammals | Andalgalomys_olrogi | Andalgalomys_olrogi | nocturnal |
| mammals | Andalgalomys_pearsoni | Andalgalomys_pearsoni | nocturnal |
| mammals | Andinomys_edax | Andinomys_edax | nocturnal |
| mammals | Anisomys_imitator | Anisomys_imitator | nocturnal |
| mammals | Anomalurus_beecrofti | Anomalurus_beecrofti | nocturnal |
| mammals | Anomalurus_derbianus | Anomalurus_derbianus | nocturnal |
| mammals | Anomalurus_pelii | Anomalurus_pelii | nocturnal |
| mammals | Anomalurus_pusillus | Anomalurus_pusillus | nocturnal |

| | | | |
|---|---|---|---|
| mammals | Anonymomys_mindorensis | Anonymomys_mindorensis | nocturnal |
| mammals | Anotomys_leander | Anotomys_leander | nocturnal |
| mammals | Anoura_caudifer | Anoura_caudifer | nocturnal |
| mammals | Anoura_cultrata | Anoura_cultrata | nocturnal |
| mammals | Anoura_fistulata | Anoura_fistulata | nocturnal |
| mammals | Anoura_geoffroyi | Anoura_geoffroyi | nocturnal |
| mammals | Anoura_latidens | Anoura_latidens | nocturnal |
| mammals | Anoura_luismanueli | Anoura_luismanueli | nocturnal |
| mammals | Anourosorex_assamensis | Anourosorex_assamensis | nocturnal |
| mammals | Anourosorex_squamipes | Anourosorex_squamipes | nocturnal |
| mammals | Anourosorex_yamashinai | Anourosorex_yamashinai | nocturnal |
| mammals | Antechinomys_laniger | Antechinomys_laniger | nocturnal |
| mammals | Antechinomys_longicaudatus | Sminthopsis_longicaudata | nocturnal |
| mammals | Antechinus_adustus | Antechinus_adustus | diurnal |
| mammals | Antechinus_agilis | Antechinus_agilis | nocturnal |
| mammals | Antechinus_bellus | Antechinus_bellus | nocturnal |
| mammals | Antechinus_flavipes | Antechinus_flavipes | nocturnal |
| mammals | Antechinus_godmani | Antechinus_godmani | diurnal |
| mammals | Antechinus_leo | Antechinus_leo | nocturnal |
| mammals | Antechinus_minimus | Antechinus_minimus | crepuscular |
| mammals | Antechinus_stuartii | Antechinus_stuartii | nocturnal |
| mammals | Antechinus_subtropicus | Antechinus_subtropicus | nocturnal |
| mammals | Antechinus_swainsonii | Antechinus_swainsonii | cathemeral |
| mammals | Anteliomys_chinensis | Eothenomys_chinensis | cathemeral |
| mammals | Anteliomys_custos | Eothenomys_custos | cathemeral |
| mammals | Anteliomys_olitor | Eothenomys_olitor | cathemeral |
| mammals | Anteliomys_proditor | Eothenomys_proditor | cathemeral |
| mammals | Anteliomys_wardi | Eothenomys_wardi | cathemeral |
| mammals | Anthops_ornatus | Anthops_ornatus | nocturnal |
| mammals | Antidorcas_marsupialis | Antidorcas_marsupialis | cathemeral |
| mammals | Antilocapra_americana | Antilocapra_americana | cathemeral |
| mammals | Antilope_cervicapra | Antilope_cervicapra | diurnal |
| mammals | Antrozous_pallidus | Antrozous_pallidus | nocturnal |
| mammals | Aotus_azarae | Aotus_azarae | cathemeral |

| | | | |
|---|---|---|---|
| mammals | Aotus_brumbacki | Aotus_brumbacki | nocturnal |
| mammals | Aotus_griseimembra | Aotus_griseimembra | nocturnal |
| mammals | Aotus_jorgehernandezi | Aotus_jorgehernandezi | nocturnal |
| mammals | Aotus_lemurinus | Aotus_lemurinus | nocturnal |
| mammals | Aotus_miconax | Aotus_miconax | nocturnal |
| mammals | Aotus_nancymai | Aotus_nancymaae | nocturnal |
| mammals | Aotus_nigriceps | Aotus_nigriceps | nocturnal |
| mammals | Aotus_trivirgatus | Aotus_trivirgatus | cathemeral |
| mammals | Aotus_vociferans | Aotus_vociferans | nocturnal |
| mammals | Aotus_zonalis | Aotus_zonalis | nocturnal |
| mammals | Aplodontia_rufa | Aplodontia_rufa | cathemeral |
| mammals | Apodemus_agrarius | Apodemus_agrarius | diurnal |
| mammals | Apodemus_argenteus | Apodemus_argenteus | nocturnal |
| mammals | Apodemus_chevrieri | Apodemus_chevrieri | diurnal |
| mammals | Apodemus_draco | Apodemus_draco | nocturnal |
| mammals | Apodemus_epimelas | Apodemus_epimelas | nocturnal |
| mammals | Apodemus_hyrcanicus | Apodemus_hyrcanicus | nocturnal |
| mammals | Apodemus_latronum | Apodemus_latronum | nocturnal |
| mammals | Apodemus_mystacinus | Apodemus_mystacinus | nocturnal |
| mammals | Apodemus_peninsulae | Apodemus_peninsulae | nocturnal |
| mammals | Apodemus_rusiges | Apodemus_rusiges | nocturnal |
| mammals | Apodemus_semotus | Apodemus_semotus | nocturnal |
| mammals | Apodemus_speciosus | Apodemus_speciosus | nocturnal |
| mammals | Apodemus_witherbyi | Apodemus_witherbyi | nocturnal |
| mammals | Apomys_abrae | Apomys_abrae | nocturnal |
| mammals | Apomys_datae | Apomys_datae | nocturnal |
| mammals | Apomys_gracilirostris | Apomys_gracilirostris | nocturnal |
| mammals | Apomys_hylocoetes | Apomys_hylocetes | nocturnal |
| mammals | Apomys_insignis | Apomys_insignis | nocturnal |
| mammals | Apomys_littoralis | Apomys_littoralis | nocturnal |
| mammals | Apomys_microdon | Apomys_microdon | nocturnal |
| mammals | Apomys_musculus | Apomys_musculus | nocturnal |
| mammals | Apomys_sacobianus | Apomys_sacobianus | nocturnal |
| mammals | Aproteles_bulmerae | Aproteles_bulmerae | nocturnal |

| | | | |
|---|---|---|---|
| mammals | Arabitragus_jayakari | Arabitragus_jayakari | cathemeral |
| mammals | Arborimus_albipes | Arborimus_albipes | nocturnal |
| mammals | Arborimus_longicaudus | Arborimus_longicaudus | nocturnal |
| mammals | Arborimus_pomo | Arborimus_pomo | nocturnal |
| mammals | Archboldomys_luzonensis | Archboldomys_luzonensis | diurnal |
| mammals | Arctictis_binturong | Arctictis_binturong | cathemeral |
| mammals | Arctocebus_aureus | Arctocebus_aureus | nocturnal |
| mammals | Arctocebus_calabarensis | Arctocebus_calabarensis | nocturnal |
| mammals | Arctogalidia_trivirgata | Arctogalidia_trivirgata | nocturnal |
| mammals | Arctonyx_albogularis | Arctonyx_albogularis | cathemeral |
| mammals | Arctonyx_collaris | Arctonyx_collaris | cathemeral |
| mammals | Arctonyx_hoevenii | Arctonyx_hoevenii | cathemeral |
| mammals | Ardops_nichollsi | Ardops_nichollsi | nocturnal |
| mammals | Arielulus_circumdatus | Arielulus_circumdatus | nocturnal |
| mammals | Arielulus_cuprosus | Arielulus_cuprosus | nocturnal |
| mammals | Arielulus_societatis | Arielulus_societatis | nocturnal |
| mammals | Ariteus_flavescens | Ariteus_flavescens | nocturnal |
| mammals | Artibeus_amplus | Artibeus_amplus | nocturnal |
| mammals | Artibeus_concolor | Artibeus_concolor | nocturnal |
| mammals | Artibeus_fimbriatus | Artibeus_fimbriatus | nocturnal |
| mammals | Artibeus_fraterculus | Artibeus_fraterculus | nocturnal |
| mammals | Artibeus_hirsutus | Artibeus_hirsutus | nocturnal |
| mammals | Artibeus_inopinatus | Artibeus_inopinatus | nocturnal |
| mammals | Artibeus_jamaicensis | Artibeus_jamaicensis | nocturnal |
| mammals | Artibeus_lituratus | Artibeus_lituratus | nocturnal |
| mammals | Artibeus_obscurus | Artibeus_obscurus | nocturnal |
| mammals | Artibeus_planirostris | Artibeus_planirostris | nocturnal |
| mammals | Artibeus_schwartzi | Artibeus_schwartzi | nocturnal |
| mammals | Arvicanthis_abyssinicus | Arvicanthis_abyssinicus | diurnal |
| mammals | Arvicanthis_ansorgei | Arvicanthis_ansorgei | diurnal |
| mammals | Arvicanthis_blicki | Arvicanthis_blicki | diurnal |
| mammals | Arvicanthis_nairobae | Arvicanthis_nairobae | diurnal |
| mammals | Arvicanthis_neumanni | Arvicanthis_neumanni | diurnal |
| mammals | Arvicanthis_niloticus | Arvicanthis_niloticus | diurnal |

| | | | |
|---|---|---|---|
| mammals | Arvicanthis_rufinus | Arvicanthis_rufinus | diurnal |
| mammals | Arvicola_amphibius | Arvicola_amphibius | cathemeral |
| mammals | Arvicola_sapidus | Arvicola_sapidus | cathemeral |
| mammals | Asellia_patrizii | Asellia_patrizii | nocturnal |
| mammals | Asellia_tridens | Asellia_tridens | nocturnal |
| mammals | Aselliscus_stoliczkanus | Aselliscus_stoliczkanus | nocturnal |
| mammals | Aselliscus_tricuspidatus | Aselliscus_tricuspidatus | nocturnal |
| mammals | Atelerix_albiventris | Atelerix_albiventris | nocturnal |
| mammals | Atelerix_algirus | Atelerix_algirus | nocturnal |
| mammals | Atelerix_frontalis | Atelerix_frontalis | nocturnal |
| mammals | Atelerix_sclateri | Atelerix_sclateri | nocturnal |
| mammals | Ateles_belzebuth | Ateles_belzebuth | diurnal |
| mammals | Ateles_chamek | Ateles_chamek | diurnal |
| mammals | Ateles_fusciceps | Ateles_fusciceps | diurnal |
| mammals | Ateles_geoffroyi | Ateles_geoffroyi | diurnal |
| mammals | Ateles_hybridus | Ateles_hybridus | diurnal |
| mammals | Ateles_marginatus | Ateles_marginatus | diurnal |
| mammals | Ateles_paniscus | Ateles_paniscus | diurnal |
| mammals | Atelocynus_microtis | Atelocynus_microtis | cathemeral |
| mammals | Atherurus_africanus | Atherurus_africanus | nocturnal |
| mammals | Atherurus_macrourus | Atherurus_macrourus | nocturnal |
| mammals | Atilax_paludinosus | Atilax_paludinosus | nocturnal |
| mammals | Atlantoxerus_getulus | Atlantoxerus_getulus | diurnal |
| mammals | Atopogale_cubana | Solenodon_cubanus | nocturnal |
| mammals | Auliscomys_boliviensis | Auliscomys_boliviensis | diurnal |
| mammals | Auliscomys_pictus | Auliscomys_pictus | cathemeral |
| mammals | Auliscomys_sublimis | Auliscomys_sublimis | nocturnal |
| mammals | Austronomus_australis | Austronomus_australis | nocturnal |
| mammals | Austronomus_kuboriensis | Austronomus_kuboriensis | nocturnal |
| mammals | Avahi_betsileo | Avahi_betsileo | nocturnal |
| mammals | Avahi_cleesei | Avahi_cleesei | nocturnal |
| mammals | Avahi_laniger | Avahi_laniger | nocturnal |
| mammals | Avahi_meridionalis | Avahi_meridionalis | nocturnal |
| mammals | Avahi_mooreorum | Avahi_mooreorum | nocturnal |

| | | | |
|---|---|---|---|
| mammals | Avahi_occidentalis | Avahi_occidentalis | nocturnal |
| mammals | Avahi_peyrierasi | Avahi_peyrierasi | nocturnal |
| mammals | Avahi_ramanantsoavani | Avahi_ramanantsoavanai | nocturnal |
| mammals | Avahi_unicolor | Avahi_unicolor | nocturnal |
| mammals | Axis_axis | Axis_axis | crepuscular |
| mammals | Axis_calamianensis | Axis_calamianensis | diurnal |
| mammals | Axis_kuhlii | Axis_kuhlii | diurnal |
| mammals | Axis_porcinus | Axis_porcinus | crepuscular |
| mammals | Babyrousa_babyrussa | Babyrousa_babyrussa | diurnal |
| mammals | Babyrousa_celebensis | Babyrousa_celebensis | diurnal |
| mammals | Babyrousa_togeanensis | Babyrousa_togeanensis | diurnal |
| mammals | Baeodon_alleni | Rhogeessa_alleni | nocturnal |
| mammals | Baeodon_gracilis | Rhogeessa_gracilis | nocturnal |
| mammals | Baiomys_musculus | Baiomys_musculus | diurnal |
| mammals | Baiomys_taylori | Baiomys_taylori | nocturnal |
| mammals | Baiyankamys_habbema | Baiyankamys_habbema | nocturnal |
| mammals | Baiyankamys_shawmayeri | Baiyankamys_shawmayeri | nocturnal |
| mammals | Balantiopteryx_infusca | Balantiopteryx_infusca | nocturnal |
| mammals | Balantiopteryx_io | Balantiopteryx_io | nocturnal |
| mammals | Balantiopteryx_plicata | Balantiopteryx_plicata | nocturnal |
| mammals | Balionycteris_maculata | Balionycteris_maculata | nocturnal |
| mammals | Bandicota_bengalensis | Bandicota_bengalensis | nocturnal |
| mammals | Barbastella_barbastellus | Barbastella_barbastellus | nocturnal |
| mammals | Barbastella_leucomelas | Barbastella_leucomelas | nocturnal |
| mammals | Bassaricyon_alleni | Bassaricyon_alleni | nocturnal |
| mammals | Bassaricyon_gabbii | Bassaricyon_gabbii | nocturnal |
| mammals | Bassaricyon_medius | Bassaricyon_medius | nocturnal |
| mammals | Bassaricyon_neblina | Bassaricyon_neblina | nocturnal |
| mammals | Bassariscus_astutus | Bassariscus_astutus | nocturnal |
| mammals | Bassariscus_sumichrasti | Bassariscus_sumichrasti | nocturnal |
| mammals | Batomys_dentatus | Batomys_dentatus | nocturnal |
| mammals | Batomys_grantii | Batomys_granti | nocturnal |
| mammals | Batomys_russatus | Batomys_russatus | nocturnal |
| mammals | Batomys_salomonseni | Batomys_salomonseni | nocturnal |

| | | | |
|---|---|---|---|
| mammals | Bauerus_dubiaquercus | Bauerus_dubiaquercus | nocturnal |
| mammals | Bdeogale_crassicauda | Bdeogale_crassicauda | nocturnal |
| mammals | Bdeogale_jacksoni | Bdeogale_jacksoni | nocturnal |
| mammals | Bdeogale_nigripes | Bdeogale_nigripes | nocturnal |
| mammals | Bdeogale_omnivora | Bdeogale_omnivora | nocturnal |
| mammals | Beamys_hindei | Beamys_hindei | nocturnal |
| mammals | Beatragus_hunteri | Beatragus_hunteri | crepuscular |
| mammals | Belomys_pearsonii | Belomys_pearsonii | nocturnal |
| mammals | Berylmys_berdmorei | Berylmys_berdmorei | nocturnal |
| mammals | Berylmys_bowersii | Berylmys_bowersi | nocturnal |
| mammals | Berylmys_manipulus | Berylmys_manipulus | nocturnal |
| mammals | Bettongia_gaimardi | Bettongia_gaimardi | nocturnal |
| mammals | Bettongia_lesueur | Bettongia_lesueur | nocturnal |
| mammals | Bettongia_penicillata | Bettongia_penicillata | nocturnal |
| mammals | Bettongia_tropica | Bettongia_tropica | nocturnal |
| mammals | Bibimys_chacoensis | Bibimys_chacoensis | nocturnal |
| mammals | Bibimys_labiosus | Bibimys_labiosus | nocturnal |
| mammals | Bibimys_torresi | Bibimys_torresi | nocturnal |
| mammals | Biswamoyopterus_biswasi | Biswamoyopterus_biswasi | crepuscular |
| mammals | Blarina_brevicauda | Blarina_brevicauda | nocturnal |
| mammals | Blarina_carolinensis | Blarina_carolinensis | nocturnal |
| mammals | Blarina_hylophaga | Blarina_hylophaga | crepuscular |
| mammals | Blarinella_quadraticauda | Blarinella_quadraticauda | nocturnal |
| mammals | Blarinella_wardi | Blarinella_wardi | nocturnal |
| mammals | Blastocerus_dichotomus | Blastocerus_dichotomus | nocturnal |
| mammals | Boneia_bidens | Rousettus_bidens | nocturnal |
| mammals | Bos_bison | Bison_bison | cathemeral |
| mammals | Bos_bonasus | Bison_bonasus | cathemeral |
| mammals | Bos_gaurus | Bos_gaurus | cathemeral |
| mammals | Bos_javanicus | Bos_javanicus | cathemeral |
| mammals | Bos_mutus | Bos_mutus | cathemeral |
| mammals | Boselaphus_tragocamelus | Boselaphus_tragocamelus | crepuscular |
| mammals | Brachiones_przewalskii | Brachiones_przewalskii | nocturnal |
| mammals | Brachyphylla_cavernarum | Brachyphylla_cavernarum | nocturnal |

| | | | |
|---|---|---|---|
| mammals | Brachyphylla_nana | Brachyphylla_nana | nocturnal |
| mammals | Brachytarsomys_albicauda | Brachytarsomys_albicauda | nocturnal |
| mammals | Brachytarsomys_villosus | Brachytarsomys_villosa | nocturnal |
| mammals | Brachyteles_arachnoides | Brachyteles_arachnoides | diurnal |
| mammals | Brachyteles_hypoxanthus | Brachyteles_hypoxanthus | diurnal |
| mammals | Brachyuromys_betsileoensis | Brachyuromys_betsileoensis | cathemeral |
| mammals | Brachyuromys_ramirohitra | Brachyuromys_ramirohitra | cathemeral |
| mammals | Bradypus_torquatus | Bradypus_torquatus | cathemeral |
| mammals | Bradypus_tridactylus | Bradypus_tridactylus | cathemeral |
| mammals | Bradypus_variegatus | Bradypus_variegatus | cathemeral |
| mammals | Brassomys_albidens | Coccymys_albidens | nocturnal |
| mammals | Brucepattersonius_griserufescens | Brucepattersonius_albinasus | nocturnal |
| mammals | Brucepattersonius_iheringi | Brucepattersonius_iheringi | nocturnal |
| mammals | Bubalus_arnee | Bubalus_arnee | cathemeral |
| mammals | Bubalus_depressicornis | Bubalus_depressicornis | cathemeral |
| mammals | Bubalus_mindorensis | Bubalus_mindorensis | diurnal |
| mammals | Bubalus_quarlesi | Bubalus_quarlesi | cathemeral |
| mammals | Budorcas_taxicolor | Budorcas_taxicolor | diurnal |
| mammals | Bullimus_bagobus | Bullimus_bagobus | nocturnal |
| mammals | Bullimus_gamay | Bullimus_gamay | nocturnal |
| mammals | Bullimus_luzonicus | Bullimus_luzonicus | cathemeral |
| mammals | Bunolagus_monticularis | Bunolagus_monticularis | nocturnal |
| mammals | Bunomys_andrewsi | Bunomys_andrewsi | nocturnal |
| mammals | Bunomys_chrysocomus | Bunomys_chrysocomus | nocturnal |
| mammals | Bunomys_coelestis | Bunomys_coelestis | nocturnal |
| mammals | Bunomys_penitus | Bunomys_penitus | nocturnal |
| mammals | Bunomys_prolatus | Bunomys_prolatus | nocturnal |
| mammals | Burramys_parvus | Burramys_parvus | nocturnal |
| mammals | Cabassous_centralis | Cabassous_centralis | nocturnal |
| mammals | Cabassous_chacoensis | Cabassous_chacoensis | nocturnal |
| mammals | Cabassous_tatouay | Cabassous_tatouay | nocturnal |
| mammals | Cabassous_unicinctus | Cabassous_unicinctus | nocturnal |
| mammals | Cabreramops_aequatorianus | Molossops_aequatorianus | nocturnal |
| mammals | Cacajao_calvus | Cacajao_calvus | diurnal |

| | | | |
|---|---|---|---|
| mammals | Cacajao_hosomi | Cacajao_hosomi | diurnal |
| mammals | Caenolestes_caniventer | Caenolestes_caniventer | nocturnal |
| mammals | Caenolestes_condorensis | Caenolestes_condorensis | nocturnal |
| mammals | Caenolestes_convelatus | Caenolestes_convelatus | nocturnal |
| mammals | Caenolestes_fuliginosus | Caenolestes_fuliginosus | nocturnal |
| mammals | Caenolestes_sangay | Caenolestes_sangay | nocturnal |
| mammals | Callicebus_barbarabrownae | Callicebus_barbarabrownae | diurnal |
| mammals | Callicebus_coimbrai | Callicebus_coimbrai | diurnal |
| mammals | Callicebus_melanochir | Callicebus_melanochir | diurnal |
| mammals | Callicebus_nigrifrons | Callicebus_nigrifrons | diurnal |
| mammals | Callicebus_personatus | Callicebus_personatus | diurnal |
| mammals | Callimico_goeldii | Callimico_goeldii | diurnal |
| mammals | Callistomys_pictus | Callistomys_pictus | nocturnal |
| mammals | Callithrix_aurita | Callithrix_aurita | diurnal |
| mammals | Callithrix_flaviceps | Callithrix_flaviceps | diurnal |
| mammals | Callithrix_geoffroyi | Callithrix_geoffroyi | diurnal |
| mammals | Callithrix_jacchus | Callithrix_jacchus | diurnal |
| mammals | Callithrix_kuhlii | Callithrix_kuhlii | diurnal |
| mammals | Callithrix_penicillata | Callithrix_penicillata | diurnal |
| mammals | Callosciurus_adamsi | Callosciurus_adamsi | diurnal |
| mammals | Callosciurus_baluensis | Callosciurus_baluensis | diurnal |
| mammals | Callosciurus_caniceps | Callosciurus_caniceps | diurnal |
| mammals | Callosciurus_erythraeus | Callosciurus_erythraeus | diurnal |
| mammals | Callosciurus_finlaysonii | Callosciurus_finlaysonii | diurnal |
| mammals | Callosciurus_inornatus | Callosciurus_inornatus | diurnal |
| mammals | Callosciurus_melanogaster | Callosciurus_melanogaster | cathemeral |
| mammals | Callosciurus_nigrovittatus | Callosciurus_nigrovittatus | diurnal |
| mammals | Callosciurus_notatus | Callosciurus_notatus | diurnal |
| mammals | Callosciurus_orestes | Callosciurus_orestes | diurnal |
| mammals | Callosciurus_phayrei | Callosciurus_phayrei | diurnal |
| mammals | Callosciurus_prevostii | Callosciurus_prevostii | diurnal |
| mammals | Callosciurus_pygerythrus | Callosciurus_pygerythrus | diurnal |
| mammals | Callosciurus_quinquestriatus | Callosciurus_quinquestriatus | diurnal |
| mammals | Callospermophilus_lateralis | Callospermophilus_lateralis | diurnal |

| | | | |
|---|---|---|---|
| mammals | Callospermophilus_madrensis | Callospermophilus_madrensis | diurnal |
| mammals | Callospermophilus_saturatus | Callospermophilus_saturatus | diurnal |
| mammals | Calomys_boliviae | Calomys_boliviae | diurnal |
| mammals | Calomys_callidus | Calomys_callidus | nocturnal |
| mammals | Calomys_callosus | Calomys_callosus | nocturnal |
| mammals | Calomys_expulsus | Calomys_expulsus | nocturnal |
| mammals | Calomys_hummelincki | Calomys_hummelincki | nocturnal |
| mammals | Calomys_laucha | Calomys_laucha | nocturnal |
| mammals | Calomys_lepidus | Calomys_lepidus | nocturnal |
| mammals | Calomys_musculinus | Calomys_musculinus | nocturnal |
| mammals | Calomys_sorella | Calomys_sorellus | nocturnal |
| mammals | Calomys_tener | Calomys_tener | nocturnal |
| mammals | Calomys_tocantinsi | Calomys_tocantinsi | nocturnal |
| mammals | Calomys_venustus | Calomys_venustus | crepuscular |
| mammals | Calomyscus_bailwardi | Calomyscus_bailwardi | nocturnal |
| mammals | Calomyscus_baluchi | Calomyscus_baluchi | nocturnal |
| mammals | Calomyscus_elburzensis | Calomyscus_elburzensis | nocturnal |
| mammals | Calomyscus_hotsoni | Calomyscus_hotsoni | nocturnal |
| mammals | Calomyscus_mystax | Calomyscus_mystax | nocturnal |
| mammals | Calomyscus_tsolovi | Calomyscus_tsolovi | nocturnal |
| mammals | Calomyscus_urartensis | Calomyscus_urartensis | nocturnal |
| mammals | Caluromys_derbianus | Caluromys_derbianus | nocturnal |
| mammals | Caluromys_lanatus | Caluromys_lanatus | nocturnal |
| mammals | Caluromys_philander | Caluromys_philander | nocturnal |
| mammals | Caluromysiops_irruptus | Caluromysiops_irrupta | nocturnal |
| mammals | Camelus_ferus | Camelus_ferus | diurnal |
| mammals | Canis_aureus | Canis_aureus | cathemeral |
| mammals | Canis_latrans | Canis_latrans | cathemeral |
| mammals | Canis_lupus | Canis_lupus | cathemeral |
| mammals | Canis_rufus | Canis_rufus | nocturnal |
| mammals | Canis_simensis | Canis_simensis | diurnal |
| mammals | Cannomys_badius | Cannomys_badius | nocturnal |
| mammals | Cansumys_canus | Cansumys_canus | nocturnal |
| mammals | Capra_aegagrus | Capra_aegagrus | diurnal |

| | | | |
|---|---|---|---|
| mammals | Capra_caucasica | Capra_caucasica | diurnal |
| mammals | Capra_cylindricornis | Capra_cylindricornis | diurnal |
| mammals | Capra_falconeri | Capra_falconeri | diurnal |
| mammals | Capra_ibex | Capra_ibex | diurnal |
| mammals | Capra_nubiana | Capra_nubiana | diurnal |
| mammals | Capra_pyrenaica | Capra_pyrenaica | diurnal |
| mammals | Capra_sibirica | Capra_sibirica | diurnal |
| mammals | Capra_walie | Capra_walie | diurnal |
| mammals | Capreolus_capreolus | Capreolus_capreolus | cathemeral |
| mammals | Capreolus_pygargus | Capreolus_pygargus | cathemeral |
| mammals | Capricornis_crispus | Capricornis_crispus | diurnal |
| mammals | Capricornis_rubidus | Capricornis_rubidus | cathemeral |
| mammals | Capricornis_sumatraensis | Capricornis_sumatraensis | cathemeral |
| mammals | Capricornis_swinhoei | Capricornis_swinhoei | diurnal |
| mammals | Caprolagus_hispidus | Caprolagus_hispidus | crepuscular |
| mammals | Capromys_pilorides | Capromys_pilorides | nocturnal |
| mammals | Caracal_aurata | Caracal_aurata | cathemeral |
| mammals | Caracal_caracal | Caracal_caracal | nocturnal |
| mammals | Cardiocranius_paradoxus | Cardiocranius_paradoxus | nocturnal |
| mammals | Cardioderma_cor | Cardioderma_cor | nocturnal |
| mammals | Carlito_syrichta | Tarsius_syrichta | nocturnal |
| mammals | Carollia_brevicauda | Carollia_brevicauda | nocturnal |
| mammals | Carollia_castanea | Carollia_castanea | nocturnal |
| mammals | Carollia_manu | Carollia_manu | nocturnal |
| mammals | Carollia_perspicillata | Carollia_perspicillata | nocturnal |
| mammals | Carollia_sowelli | Carollia_sowelli | nocturnal |
| mammals | Carollia_subrufa | Carollia_subrufa | nocturnal |
| mammals | Carpomys_melanurus | Carpomys_melanurus | nocturnal |
| mammals | Carpomys_phaeurus | Carpomys_phaeurus | nocturnal |
| mammals | Carterodon_sulcidens | Carterodon_sulcidens | nocturnal |
| mammals | Casinycteris_argynnis | Casinycteris_argynnis | nocturnal |
| mammals | Cassistrellus_dimissus | Eptesicus_dimissus | nocturnal |
| mammals | Castor_canadensis | Castor_canadensis | nocturnal |
| mammals | Castor_fiber | Castor_fiber | nocturnal |

| | | | |
|---|---|---|---|
| mammals | Castoria_angustidens | Akodon_serrensis | cathemeral |
| mammals | Catopuma_badia | Catopuma_badia | diurnal |
| mammals | Catopuma_temminckii | Catopuma_temminckii | nocturnal |
| mammals | Cavia_aperea | Cavia_aperea | diurnal |
| mammals | Cavia_fulgida | Cavia_fulgida | diurnal |
| mammals | Cavia_magna | Cavia_magna | diurnal |
| mammals | Cavia_patzelti | Cavia_patzelti | diurnal |
| mammals | Cavia_tschudii | Cavia_tschudii | nocturnal |
| mammals | Cebuella_pygmaea | Cebuella_pygmaea | diurnal |
| mammals | Cebus_albifrons | Cebus_albifrons | diurnal |
| mammals | Cebus_kaapori | Cebus_kaapori | diurnal |
| mammals | Centronycteris_centralis | Centronycteris_centralis | nocturnal |
| mammals | Centronycteris_maximiliani | Centronycteris_maximiliani | nocturnal |
| mammals | Centurio_senex | Centurio_senex | nocturnal |
| mammals | Cephalopachus_bancanus | Tarsius_bancanus | nocturnal |
| mammals | Cephalophorus_callipygus | Cephalophus_callipygus | diurnal |
| mammals | Cephalophorus_leucogaster | Cephalophus_leucogaster | diurnal |
| mammals | Cephalophorus_natalensis | Cephalophus_natalensis | diurnal |
| mammals | Cephalophorus_niger | Cephalophus_niger | cathemeral |
| mammals | Cephalophorus_nigrifrons | Cephalophus_nigrifrons | diurnal |
| mammals | Cephalophorus_ogilbyi | Cephalophus_ogilbyi | diurnal |
| mammals | Cephalophorus_rufilatus | Cephalophus_rufilatus | diurnal |
| mammals | Cephalophorus_weynsi | Cephalophus_weynsi | diurnal |
| mammals | Cephalophula_zebra | Cephalophus_zebra | diurnal |
| mammals | Cephalophus_dorsalis | Cephalophus_dorsalis | diurnal |
| mammals | Cephalophus_jentinki | Cephalophus_jentinki | nocturnal |
| mammals | Cephalophus_silvicultor | Cephalophus_silvicultor | diurnal |
| mammals | Cephalophus_spadix | Cephalophus_spadix | nocturnal |
| mammals | Ceratotherium_simum | Ceratotherium_simum | cathemeral |
| mammals | Cercartetus_caudatus | Cercartetus_caudatus | nocturnal |
| mammals | Cercartetus_concinnus | Cercartetus_concinnus | nocturnal |
| mammals | Cercartetus_lepidus | Cercartetus_lepidus | nocturnal |
| mammals | Cercartetus_nana | Cercartetus_nanus | nocturnal |
| mammals | Cercocebus_agilis | Cercocebus_agilis | diurnal |

| | | | |
|---|---|---|---|
| mammals | Cercocebus_atys | Cercocebus_atys | diurnal |
| mammals | Cercocebus_chrysogaster | Cercocebus_chrysogaster | diurnal |
| mammals | Cercocebus_galeritus | Cercocebus_galeritus | diurnal |
| mammals | Cercocebus_lunulatus | Cercocebus_lunulatus | diurnal |
| mammals | Cercocebus_sanjei | Cercocebus_sanjei | diurnal |
| mammals | Cercocebus_torquatus | Cercocebus_torquatus | diurnal |
| mammals | Cercopithecus_ascanius | Cercopithecus_ascanius | diurnal |
| mammals | Cercopithecus_campbelli | Cercopithecus_campbelli | diurnal |
| mammals | Cercopithecus_cephus | Cercopithecus_cephus | diurnal |
| mammals | Cercopithecus_denti | Cercopithecus_denti | diurnal |
| mammals | Cercopithecus_diana | Cercopithecus_diana | diurnal |
| mammals | Cercopithecus_erythrogaster | Cercopithecus_erythrogaster | diurnal |
| mammals | Cercopithecus_erythrotis | Cercopithecus_erythrotis | diurnal |
| mammals | Cercopithecus_hamlyni | Cercopithecus_hamlyni | diurnal |
| mammals | Cercopithecus_lowei | Cercopithecus_lowei | diurnal |
| mammals | Cercopithecus_mitis | Cercopithecus_mitis | diurnal |
| mammals | Cercopithecus_mona | Cercopithecus_mona | diurnal |
| mammals | Cercopithecus_neglectus | Cercopithecus_neglectus | diurnal |
| mammals | Cercopithecus_nictitans | Cercopithecus_nictitans | diurnal |
| mammals | Cercopithecus_petaurista | Cercopithecus_petaurista | diurnal |
| mammals | Cercopithecus_roloway | Cercopithecus_roloway | diurnal |
| mammals | Cercopithecus_sclateri | Cercopithecus_sclateri | diurnal |
| mammals | Cerdocyon_thous | Cerdocyon_thous | cathemeral |
| mammals | Cerradomys_maracajuensis | Cerradomys_maracajuensis | nocturnal |
| mammals | Cerradomys_marinhus | Cerradomys_marinhus | nocturnal |
| mammals | Cerradomys_scotti | Cerradomys_scotti | nocturnal |
| mammals | Cerradomys_subflavus | Cerradomys_subflavus | nocturnal |
| mammals | Cervus_albirostris | Cervus_albirostris | cathemeral |
| mammals | Cervus_canadensis | Cervus_canadensis | cathemeral |
| mammals | Cervus_elaphus | Cervus_elaphus | cathemeral |
| mammals | Cervus_nippon | Cervus_nippon | crepuscular |
| mammals | Chacodelphys_formosa | Chacodelphys_formosa | nocturnal |
| mammals | Chaetocauda_sichuanensis | Chaetocauda_sichuanensis | nocturnal |
| mammals | Chaetodipus_arenarius | Chaetodipus_arenarius | nocturnal |

| | | | |
|---|---|---|---|
| mammals | Chaetodipus_artus | Chaetodipus_artus | nocturnal |
| mammals | Chaetodipus_baileyi | Chaetodipus_baileyi | nocturnal |
| mammals | Chaetodipus_californicus | Chaetodipus_californicus | nocturnal |
| mammals | Chaetodipus_eremicus | Chaetodipus_eremicus | nocturnal |
| mammals | Chaetodipus_fallax | Chaetodipus_fallax | nocturnal |
| mammals | Chaetodipus_formosus | Chaetodipus_formosus | nocturnal |
| mammals | Chaetodipus_goldmani | Chaetodipus_goldmani | nocturnal |
| mammals | Chaetodipus_hispidus | Chaetodipus_hispidus | nocturnal |
| mammals | Chaetodipus_intermedius | Chaetodipus_intermedius | nocturnal |
| mammals | Chaetodipus_lineatus | Chaetodipus_lineatus | nocturnal |
| mammals | Chaetodipus_nelsoni | Chaetodipus_nelsoni | nocturnal |
| mammals | Chaetodipus_penicillatus | Chaetodipus_penicillatus | nocturnal |
| mammals | Chaetodipus_pernix | Chaetodipus_pernix | nocturnal |
| mammals | Chaetodipus_rudinoris | Chaetodipus_rudinoris | nocturnal |
| mammals | Chaetodipus_spinatus | Chaetodipus_spinatus | nocturnal |
| mammals | Chaetomys_subspinosus | Chaetomys_subspinosus | nocturnal |
| mammals | Chaetophractus_vellerosus | Chaetophractus_vellerosus | diurnal |
| mammals | Chaetophractus_villosus | Chaetophractus_villosus | nocturnal |
| mammals | Chalinolobus_dwyeri | Chalinolobus_dwyeri | nocturnal |
| mammals | Chalinolobus_gouldii | Chalinolobus_gouldii | nocturnal |
| mammals | Chalinolobus_morio | Chalinolobus_morio | nocturnal |
| mammals | Chalinolobus_neocaledonicus | Chalinolobus_neocaledonicus | nocturnal |
| mammals | Chalinolobus_nigrogriseus | Chalinolobus_nigrogriseus | nocturnal |
| mammals | Chalinolobus_picatus | Chalinolobus_picatus | nocturnal |
| mammals | Chalinolobus_tuberculatus | Chalinolobus_tuberculatus | nocturnal |
| mammals | Cheirogaleus_crossleyi | Cheirogaleus_crossleyi | nocturnal |
| mammals | Cheirogaleus_major | Cheirogaleus_major | nocturnal |
| mammals | Cheirogaleus_medius | Cheirogaleus_medius | nocturnal |
| mammals | Cheirogaleus_sibreei | Cheirogaleus_sibreei | nocturnal |
| mammals | Cheiromeles_parvidens | Cheiromeles_parvidens | nocturnal |
| mammals | Cheiromeles_torquatus | Cheiromeles_torquatus | nocturnal |
| mammals | Chelemys_megalonyx | Chelemys_megalonyx | cathemeral |
| mammals | Cheracebus_lucifer | Callicebus_lucifer | diurnal |
| mammals | Cheracebus_lugens | Callicebus_lugens | diurnal |

| | | | |
|---|---|---|---|
| mammals | Cheracebus_medemi | Callicebus_medemi | diurnal |
| mammals | Cheracebus_regulus | Callicebus_regulus | diurnal |
| mammals | Cheracebus_torquatus | Callicebus_torquatus | diurnal |
| mammals | Chibchanomys_trichotis | Chibchanomys_trichotis | nocturnal |
| mammals | Chilomys_instans | Chilomys_instans | nocturnal |
| mammals | Chilonatalus_micropus | Chilonatalus_micropus | nocturnal |
| mammals | Chilonatalus_tumidifrons | Chilonatalus_tumidifrons | nocturnal |
| mammals | Chimarrogale_hantu | Chimarrogale_hantu | diurnal |
| mammals | Chimarrogale_himalayica | Chimarrogale_himalayica | nocturnal |
| mammals | Chimarrogale_platycephala | Chimarrogale_platycephalus | cathemeral |
| mammals | Chimarrogale_styani | Chimarrogale_styani | nocturnal |
| mammals | Chinchilla_chinchilla | Chinchilla_chinchilla | nocturnal |
| mammals | Chinchilla_lanigera | Chinchilla_lanigera | nocturnal |
| mammals | Chinchillula_sahamae | Chinchillula_sahamae | nocturnal |
| mammals | Chionomys_gud | Chionomys_gud | cathemeral |
| mammals | Chionomys_nivalis | Chionomys_nivalis | cathemeral |
| mammals | Chionomys_roberti | Chionomys_roberti | nocturnal |
| mammals | Chiroderma_doriae | Chiroderma_doriae | nocturnal |
| mammals | Chiroderma_improvisum | Chiroderma_improvisum | nocturnal |
| mammals | Chiroderma_salvini | Chiroderma_salvini | nocturnal |
| mammals | Chiroderma_trinitatum | Chiroderma_trinitatum | nocturnal |
| mammals | Chiroderma_villosum | Chiroderma_villosum | nocturnal |
| mammals | Chiromyscus_chiropus | Chiromyscus_chiropus | nocturnal |
| mammals | Chiromyscus_langbianis | Niviventer_langbianis | nocturnal |
| mammals | Chironax_melanocephalus | Chironax_melanocephalus | nocturnal |
| mammals | Chironectes_minimus | Chironectes_minimus | nocturnal |
| mammals | Chiropodomys_calamianensis | Chiropodomys_calamianensis | nocturnal |
| mammals | Chiropodomys_gliroides | Chiropodomys_gliroides | nocturnal |
| mammals | Chiropodomys_karlkoopmani | Chiropodomys_karlkoopmani | nocturnal |
| mammals | Chiropodomys_major | Chiropodomys_major | nocturnal |
| mammals | Chiropodomys_muroides | Chiropodomys_muroides | nocturnal |
| mammals | Chiropodomys_pusillus | Chiropodomys_pusillus | nocturnal |
| mammals | Chiropotes_albinasus | Chiropotes_albinasus | diurnal |
| mammals | Chiropotes_chiropotes | Chiropotes_chiropotes | diurnal |

| | | | |
|---|---|---|---|
| mammals | Chiropotes_satanas | Chiropotes_satanas | diurnal |
| mammals | Chiropotes_utahicki | Chiropotes_utahickae | diurnal |
| mammals | Chiruromys_forbesi | Chiruromys_forbesi | nocturnal |
| mammals | Chiruromys_lamia | Chiruromys_lamia | nocturnal |
| mammals | Chiruromys_vates | Chiruromys_vates | nocturnal |
| mammals | Chlorocebus_aethiops | Chlorocebus_aethiops | diurnal |
| mammals | Chlorocebus_cynosuros | Chlorocebus_cynosuros | diurnal |
| mammals | Chlorocebus_djamdjamensis | Chlorocebus_djamdjamensis | diurnal |
| mammals | Chlorocebus_dryas | Cercopithecus_dryas | diurnal |
| mammals | Chlorocebus_pygerythrus | Chlorocebus_pygerythrus | diurnal |
| mammals | Chlorocebus_sabaeus | Chlorocebus_sabaeus | diurnal |
| mammals | Chlorocebus_tantalus | Chlorocebus_tantalus | diurnal |
| mammals | Chodsigoa_caovansunga | Chodsigoa_caovansunga | nocturnal |
| mammals | Chodsigoa_hypsibia | Chodsigoa_hypsibia | nocturnal |
| mammals | Chodsigoa_parca | Chodsigoa_parca | nocturnal |
| mammals | Chodsigoa_parva | Chodsigoa_parva | cathemeral |
| mammals | Chodsigoa_salenskii | Chodsigoa_salenskii | nocturnal |
| mammals | Chodsigoa_smithii | Chodsigoa_smithii | nocturnal |
| mammals | Chodsigoa_sodalis | Chodsigoa_sodalis | nocturnal |
| mammals | Choeroniscus_godmani | Choeroniscus_godmani | nocturnal |
| mammals | Choeroniscus_minor | Choeroniscus_minor | nocturnal |
| mammals | Choeroniscus_periosus | Choeroniscus_periosus | nocturnal |
| mammals | Choeronycteris_mexicana | Choeronycteris_mexicana | nocturnal |
| mammals | Choeropsis_liberiensis | Choeropsis_liberiensis | nocturnal |
| mammals | Choloepus_didactylus | Choloepus_didactylus | nocturnal |
| mammals | Choloepus_hoffmanni | Choloepus_hoffmanni | nocturnal |
| mammals | Chrotogale_owstoni | Chrotogale_owstoni | nocturnal |
| mammals | Chrotomys_gonzalesi | Chrotomys_gonzalesi | diurnal |
| mammals | Chrotomys_mindorensis | Chrotomys_mindorensis | cathemeral |
| mammals | Chrotomys_silaceus | Chrotomys_silaceus | diurnal |
| mammals | Chrotomys_whiteheadi | Chrotomys_whiteheadi | cathemeral |
| mammals | Chrotopterus_auritus | Chrotopterus_auritus | nocturnal |
| mammals | Chrysocyon_brachyurus | Chrysocyon_brachyurus | cathemeral |
| mammals | Cistugo_lesueuri | Cistugo_lesueuri | nocturnal |

| | | | |
|---|---|---|---|
| mammals | Cistugo_seabrae | Cistugo_seabrae | nocturnal |
| mammals | Civettictis_civetta | Civettictis_civetta | nocturnal |
| mammals | Clethrionomys_californicus | Myodes_californicus | nocturnal |
| mammals | Clethrionomys_centralis | Myodes_centralis | nocturnal |
| mammals | Clethrionomys_gapperi | Myodes_gapperi | cathemeral |
| mammals | Clethrionomys_glareolus | Myodes_glareolus | cathemeral |
| mammals | Clethrionomys_rutilus | Myodes_rutilus | cathemeral |
| mammals | Cloeotis_percivali | Cloeotis_percivali | nocturnal |
| mammals | Clyomys_laticeps | Clyomys_laticeps | diurnal |
| mammals | Cnephaeus_bottae | Eptesicus_bottae | nocturnal |
| mammals | Cnephaeus_floweri | Eptesicus_floweri | nocturnal |
| mammals | Cnephaeus_gobiensis | Eptesicus_gobiensis | nocturnal |
| mammals | Cnephaeus_hottentotus | Eptesicus_hottentotus | nocturnal |
| mammals | Cnephaeus_isabellinus | Eptesicus_isabellinus | nocturnal |
| mammals | Cnephaeus_japonensis | Eptesicus_japonensis | nocturnal |
| mammals | Cnephaeus_nilssonii | Eptesicus_nilssonii | nocturnal |
| mammals | Cnephaeus_pachyotis | Eptesicus_pachyotis | nocturnal |
| mammals | Cnephaeus_platyops | Eptesicus_platyops | nocturnal |
| mammals | Cnephaeus_serotinus | Eptesicus_serotinus | nocturnal |
| mammals | Cnephaeus_tatei | Eptesicus_tatei | nocturnal |
| mammals | Coccymys_ruemmleri | Coccymys_ruemmleri | nocturnal |
| mammals | Coelops_frithii | Coelops_frithii | nocturnal |
| mammals | Coelops_robinsoni | Coelops_robinsoni | nocturnal |
| mammals | Coendou_bicolor | Coendou_bicolor | nocturnal |
| mammals | Coendou_ichillus | Coendou_ichillus | nocturnal |
| mammals | Coendou_insidiosus | Coendou_insidiosus | nocturnal |
| mammals | Coendou_mexicanus | Coendou_mexicanus | nocturnal |
| mammals | Coendou_prehensilis | Coendou_prehensilis | nocturnal |
| mammals | Coendou_quichua | Coendou_quichua | nocturnal |
| mammals | Coendou_roosmalenorum | Coendou_roosmalenorum | nocturnal |
| mammals | Coendou_spinosus | Coendou_spinosus | nocturnal |
| mammals | Coendou_vestitus | Coendou_vestitus | nocturnal |
| mammals | Coleura_afra | Coleura_afra | nocturnal |
| mammals | Coleura_seychellensis | Coleura_seychellensis | nocturnal |

| | | | |
|---|---|---|---|
| mammals | Colobus_angolensis | Colobus_angolensis | diurnal |
| mammals | Colobus_guereza | Colobus_guereza | diurnal |
| mammals | Colobus_polykomos | Colobus_polykomos | diurnal |
| mammals | Colobus_satanas | Colobus_satanas | diurnal |
| mammals | Colobus_vellerosus | Colobus_vellerosus | diurnal |
| mammals | Colomys_goslingi | Colomys_goslingi | nocturnal |
| mammals | Condylura_cristata | Condylura_cristata | cathemeral |
| mammals | Conepatus_chinga | Conepatus_chinga | cathemeral |
| mammals | Conepatus_leuconotus | Conepatus_leuconotus | nocturnal |
| mammals | Conepatus_semistriatus | Conepatus_semistriatus | nocturnal |
| mammals | Congomys_lukolelae | Praomys_lukolelae | nocturnal |
| mammals | Congomys_verschureni | Praomys_verschureni | nocturnal |
| mammals | Congosorex_phillipsorum | Congosorex_phillipsorum | nocturnal |
| mammals | Congosorex_polli | Congosorex_polli | cathemeral |
| mammals | Conilurus_penicillatus | Conilurus_penicillatus | nocturnal |
| mammals | Connochaetes_gnou | Connochaetes_gnou | cathemeral |
| mammals | Connochaetes_taurinus | Connochaetes_taurinus | cathemeral |
| mammals | Cormura_brevirostris | Cormura_brevirostris | nocturnal |
| mammals | Corynorhinus_mexicanus | Corynorhinus_mexicanus | nocturnal |
| mammals | Corynorhinus_rafinesquii | Corynorhinus_rafinesquii | nocturnal |
| mammals | Corynorhinus_townsendii | Corynorhinus_townsendii | nocturnal |
| mammals | Craseomys_andersoni | Myodes_andersoni | nocturnal |
| mammals | Craseomys_regulus | Myodes_regulus | cathemeral |
| mammals | Craseomys_rex | Myodes_rex | cathemeral |
| mammals | Craseomys_rufocanus | Myodes_rufocanus | cathemeral |
| mammals | Craseomys_smithii | Myodes_smithii | cathemeral |
| mammals | Craseonycteris_thonglongyai | Craseonycteris_thonglongyai | nocturnal |
| mammals | Crateromys_heaneyi | Crateromys_heaneyi | nocturnal |
| mammals | Crateromys_paulus | Crateromys_paulus | nocturnal |
| mammals | Crateromys_schadenbergi | Crateromys_schadenbergi | nocturnal |
| mammals | Cremnomys_cutchicus | Cremnomys_cutchicus | nocturnal |
| mammals | Cremnomys_elvira | Cremnomys_elvira | nocturnal |
| mammals | Cricetomys_emini | Cricetomys_emini | nocturnal |
| mammals | Cricetomys_gambianus | Cricetomys_gambianus | nocturnal |

| | | | |
|---|---|---|---|
| mammals | Cricetulus_barabensis | Cricetulus_barabensis | nocturnal |
| mammals | Cricetulus_longicaudatus | Cricetulus_longicaudatus | nocturnal |
| mammals | Cricetulus_sokolovi | Cricetulus_sokolovi | nocturnal |
| mammals | Cricetus_cricetus | Cricetus_cricetus | nocturnal |
| mammals | Crocidura_aleksandrisi | Crocidura_aleksandrisi | nocturnal |
| mammals | Crocidura_allex | Crocidura_allex | nocturnal |
| mammals | Crocidura_andamanensis | Crocidura_andamanensis | nocturnal |
| mammals | Crocidura_ansellorum | Crocidura_ansellorum | nocturnal |
| mammals | Crocidura_arabica | Crocidura_arabica | nocturnal |
| mammals | Crocidura_arispa | Crocidura_arispa | nocturnal |
| mammals | Crocidura_armenica | Crocidura_armenica | nocturnal |
| mammals | Crocidura_attenuata | Crocidura_attenuata | nocturnal |
| mammals | Crocidura_attila | Crocidura_attila | nocturnal |
| mammals | Crocidura_baileyi | Crocidura_baileyi | nocturnal |
| mammals | Crocidura_baluensis | Crocidura_baluensis | nocturnal |
| mammals | Crocidura_beata | Crocidura_beatus | cathemeral |
| mammals | Crocidura_beccarii | Crocidura_beccarii | nocturnal |
| mammals | Crocidura_bottegi | Crocidura_bottegi | nocturnal |
| mammals | Crocidura_bottegoides | Crocidura_bottegoides | nocturnal |
| mammals | Crocidura_brunnea | Crocidura_brunnea | nocturnal |
| mammals | Crocidura_buettikoferi | Crocidura_buettikoferi | nocturnal |
| mammals | Crocidura_caliginea | Crocidura_caliginea | nocturnal |
| mammals | Crocidura_canariensis | Crocidura_canariensis | nocturnal |
| mammals | Crocidura_caspica | Crocidura_caspica | nocturnal |
| mammals | Crocidura_cinderella | Crocidura_cinderella | nocturnal |
| mammals | Crocidura_congobelgica | Crocidura_congobelgica | nocturnal |
| mammals | Crocidura_crenata | Crocidura_crenata | nocturnal |
| mammals | Crocidura_crossei | Crocidura_crossei | nocturnal |
| mammals | Crocidura_cyanea | Crocidura_cyanea | nocturnal |
| mammals | Crocidura_denti | Crocidura_denti | nocturnal |
| mammals | Crocidura_desperata | Crocidura_desperata | nocturnal |
| mammals | Crocidura_dhofarensis | Crocidura_dhofarensis | nocturnal |
| mammals | Crocidura_dolichura | Crocidura_dolichura | nocturnal |
| mammals | Crocidura_douceti | Crocidura_douceti | nocturnal |

| | | | |
|---|---|---|---|
| mammals | Crocidura_dsinezumi | Crocidura_dsinezumi | nocturnal |
| mammals | Crocidura_eisentrauti | Crocidura_eisentrauti | cathemeral |
| mammals | Crocidura_elgonius | Crocidura_elgonius | nocturnal |
| mammals | Crocidura_elongata | Crocidura_elongata | nocturnal |
| mammals | Crocidura_erica | Crocidura_erica | nocturnal |
| mammals | Crocidura_fischeri | Crocidura_fischeri | nocturnal |
| mammals | Crocidura_flavescens | Crocidura_flavescens | nocturnal |
| mammals | Crocidura_floweri | Crocidura_floweri | nocturnal |
| mammals | Crocidura_foxi | Crocidura_foxi | nocturnal |
| mammals | Crocidura_fuliginosa | Crocidura_fuliginosa | nocturnal |
| mammals | Crocidura_fulvastra | Crocidura_fulvastra | nocturnal |
| mammals | Crocidura_fumosa | Crocidura_fumosa | nocturnal |
| mammals | Crocidura_fuscomurina | Crocidura_fuscomurina | nocturnal |
| mammals | Crocidura_glassi | Crocidura_glassi | nocturnal |
| mammals | Crocidura_goliath | Crocidura_goliath | nocturnal |
| mammals | Crocidura_gracilipes | Crocidura_gracilipes | nocturnal |
| mammals | Crocidura_grandiceps | Crocidura_grandiceps | nocturnal |
| mammals | Crocidura_grandis | Crocidura_grandis | nocturnal |
| mammals | Crocidura_grassei | Crocidura_grassei | nocturnal |
| mammals | Crocidura_grayi | Crocidura_grayi | cathemeral |
| mammals | Crocidura_greenwoodi | Crocidura_greenwoodi | nocturnal |
| mammals | Crocidura_harenna | Crocidura_harenna | nocturnal |
| mammals | Crocidura_hildegardeae | Crocidura_hildegardeae | nocturnal |
| mammals | Crocidura_hilliana | Crocidura_hilliana | nocturnal |
| mammals | Crocidura_horsfieldii | Crocidura_horsfieldii | nocturnal |
| mammals | Crocidura_hutanis | Crocidura_hutanis | nocturnal |
| mammals | Crocidura_indochinensis | Crocidura_indochinensis | nocturnal |
| mammals | Crocidura_jacksoni | Crocidura_jacksoni | nocturnal |
| mammals | Crocidura_jenkinsi | Crocidura_jenkinsi | nocturnal |
| mammals | Crocidura_jouvenetae | Crocidura_jouvenetae | nocturnal |
| mammals | Crocidura_katinka | Crocidura_katinka | nocturnal |
| mammals | Crocidura_lamottei | Crocidura_lamottei | nocturnal |
| mammals | Crocidura_lanosa | Crocidura_lanosa | nocturnal |
| mammals | Crocidura_lasiura | Crocidura_lasiura | cathemeral |

| | | | |
|---|---|---|---|
| mammals | Crocidura_latona | Crocidura_latona | nocturnal |
| mammals | Crocidura_lea | Crocidura_lea | nocturnal |
| mammals | Crocidura_lepidura | Crocidura_lepidura | nocturnal |
| mammals | Crocidura_leucodon | Crocidura_leucodon | cathemeral |
| mammals | Crocidura_levicula | Crocidura_levicula | diurnal |
| mammals | Crocidura_littoralis | Crocidura_littoralis | nocturnal |
| mammals | Crocidura_longipes | Crocidura_longipes | nocturnal |
| mammals | Crocidura_lucina | Crocidura_lucina | nocturnal |
| mammals | Crocidura_ludia | Crocidura_ludia | nocturnal |
| mammals | Crocidura_luna | Crocidura_luna | nocturnal |
| mammals | Crocidura_lusitania | Crocidura_lusitania | nocturnal |
| mammals | Crocidura_macarthuri | Crocidura_macarthuri | nocturnal |
| mammals | Crocidura_macmillani | Crocidura_macmillani | nocturnal |
| mammals | Crocidura_macowi | Crocidura_macowi | nocturnal |
| mammals | Crocidura_malayana | Crocidura_malayana | nocturnal |
| mammals | Crocidura_manengubae | Crocidura_manengubae | nocturnal |
| mammals | Crocidura_maquassiensis | Crocidura_maquassiensis | nocturnal |
| mammals | Crocidura_mariquensis | Crocidura_mariquensis | nocturnal |
| mammals | Crocidura_maurisca | Crocidura_maurisca | nocturnal |
| mammals | Crocidura_maxi | Crocidura_maxi | nocturnal |
| mammals | Crocidura_mindorus | Crocidura_mindorus | nocturnal |
| mammals | Crocidura_miya | Crocidura_miya | nocturnal |
| mammals | Crocidura_monax | Crocidura_monax | nocturnal |
| mammals | Crocidura_monticola | Crocidura_monticola | nocturnal |
| mammals | Crocidura_montis | Crocidura_montis | nocturnal |
| mammals | Crocidura_muricauda | Crocidura_muricauda | nocturnal |
| mammals | Crocidura_musseri | Crocidura_musseri | nocturnal |
| mammals | Crocidura_mutesae | Crocidura_mutesae | nocturnal |
| mammals | Crocidura_nana | Crocidura_nana | nocturnal |
| mammals | Crocidura_nanilla | Crocidura_nanilla | nocturnal |
| mammals | Crocidura_negrina | Crocidura_negrina | nocturnal |
| mammals | Crocidura_nicobarica | Crocidura_nicobarica | nocturnal |
| mammals | Crocidura_nigeriae | Crocidura_nigeriae | nocturnal |
| mammals | Crocidura_nigricans | Crocidura_nigricans | nocturnal |

| | | | |
|---|---|---|---|
| mammals | Crocidura_nigripes | Crocidura_nigripes | nocturnal |
| mammals | Crocidura_nigrofusca | Crocidura_nigrofusca | nocturnal |
| mammals | Crocidura_nimbae | Crocidura_nimbae | nocturnal |
| mammals | Crocidura_niobe | Crocidura_niobe | nocturnal |
| mammals | Crocidura_obscurior | Crocidura_obscurior | nocturnal |
| mammals | Crocidura_olivieri | Crocidura_olivieri | nocturnal |
| mammals | Crocidura_orientalis | Crocidura_orientalis | nocturnal |
| mammals | Crocidura_orii | Crocidura_orii | nocturnal |
| mammals | Crocidura_pachyura | Crocidura_pachyura | nocturnal |
| mammals | Crocidura_palawanensis | Crocidura_palawanensis | nocturnal |
| mammals | Crocidura_paradoxura | Crocidura_paradoxura | nocturnal |
| mammals | Crocidura_parvipes | Crocidura_parvipes | nocturnal |
| mammals | Crocidura_pasha | Crocidura_pasha | nocturnal |
| mammals | Crocidura_phaeura | Crocidura_phaeura | nocturnal |
| mammals | Crocidura_picea | Crocidura_picea | nocturnal |
| mammals | Crocidura_pitmani | Crocidura_pitmani | nocturnal |
| mammals | Crocidura_planiceps | Crocidura_planiceps | nocturnal |
| mammals | Crocidura_poensis | Crocidura_poensis | nocturnal |
| mammals | Crocidura_polia | Crocidura_polia | nocturnal |
| mammals | Crocidura_pullata | Crocidura_pullata | nocturnal |
| mammals | Crocidura_raineyi | Crocidura_raineyi | nocturnal |
| mammals | Crocidura_ramona | Crocidura_ramona | nocturnal |
| mammals | Crocidura_rapax | Crocidura_rapax | nocturnal |
| mammals | Crocidura_religiosa | Crocidura_religiosa | nocturnal |
| mammals | Crocidura_rhoditis | Crocidura_rhoditis | nocturnal |
| mammals | Crocidura_roosevelti | Crocidura_roosevelti | nocturnal |
| mammals | Crocidura_russula | Crocidura_russula | cathemeral |
| mammals | Crocidura_selina | Crocidura_selina | nocturnal |
| mammals | Crocidura_serezkyensis | Crocidura_serezkyensis | nocturnal |
| mammals | Crocidura_shantungensis | Crocidura_shantungensis | cathemeral |
| mammals | Crocidura_sibirica | Crocidura_sibirica | nocturnal |
| mammals | Crocidura_sicula | Crocidura_sicula | cathemeral |
| mammals | Crocidura_silacea | Crocidura_silacea | nocturnal |
| mammals | Crocidura_smithii | Crocidura_smithii | nocturnal |

| | | | |
|---|---|---|---|
| mammals | Crocidura_somalica | Crocidura_somalica | nocturnal |
| mammals | Crocidura_stenocephala | Crocidura_stenocephala | nocturnal |
| mammals | Crocidura_suaveolens | Crocidura_suaveolens | nocturnal |
| mammals | Crocidura_susiana | Crocidura_susiana | nocturnal |
| mammals | Crocidura_tanakae | Crocidura_tanakae | nocturnal |
| mammals | Crocidura_tansaniana | Crocidura_tansaniana | nocturnal |
| mammals | Crocidura_tarella | Crocidura_tarella | nocturnal |
| mammals | Crocidura_tarfayaensis | Crocidura_tarfayensis | nocturnal |
| mammals | Crocidura_telfordi | Crocidura_telfordi | cathemeral |
| mammals | Crocidura_tenuis | Crocidura_tenuis | nocturnal |
| mammals | Crocidura_theresae | Crocidura_theresae | nocturnal |
| mammals | Crocidura_turba | Crocidura_turba | nocturnal |
| mammals | Crocidura_ultima | Crocidura_ultima | nocturnal |
| mammals | Crocidura_usambarae | Crocidura_usambarae | nocturnal |
| mammals | Crocidura_viaria | Crocidura_viaria | nocturnal |
| mammals | Crocidura_virgata | Crocidura_virgata | nocturnal |
| mammals | Crocidura_voi | Crocidura_voi | nocturnal |
| mammals | Crocidura_vorax | Crocidura_vorax | nocturnal |
| mammals | Crocidura_vosmaeri | Crocidura_vosmaeri | nocturnal |
| mammals | Crocidura_watasei | Crocidura_watasei | nocturnal |
| mammals | Crocidura_whitakeri | Crocidura_whitakeri | nocturnal |
| mammals | Crocidura_wimmeri | Crocidura_wimmeri | nocturnal |
| mammals | Crocidura_wuchihensis | Crocidura_wuchihensis | nocturnal |
| mammals | Crocidura_xantippe | Crocidura_xantippe | nocturnal |
| mammals | Crocidura_yankariensis | Crocidura_yankariensis | nocturnal |
| mammals | Crocidura_zaphiri | Crocidura_zaphiri | nocturnal |
| mammals | Crocidura_zarudnyi | Crocidura_zarudnyi | nocturnal |
| mammals | Crocidura_zimmeri | Crocidura_zimmeri | nocturnal |
| mammals | Crocidura_zimmermanni | Crocidura_zimmermanni | nocturnal |
| mammals | Crocuta_crocuta | Crocuta_crocuta | nocturnal |
| mammals | Crossarchus_alexandri | Crossarchus_alexandri | diurnal |
| mammals | Crossarchus_ansorgei | Crossarchus_ansorgei | diurnal |
| mammals | Crossarchus_obscurus | Crossarchus_obscurus | diurnal |
| mammals | Crossarchus_platycephalus | Crossarchus_platycephalus | diurnal |

| | | | |
|---|---|---|---|
| mammals | Crossomys_moncktoni | Crossomys_moncktoni | diurnal |
| mammals | Crunomys_celebensis | Crunomys_celebensis | diurnal |
| mammals | Crunomys_fallax | Crunomys_fallax | nocturnal |
| mammals | Crunomys_melanius | Crunomys_melanius | diurnal |
| mammals | Crunomys_suncoides | Crunomys_suncoides | diurnal |
| mammals | Cryptonanus_agricolai | Cryptonanus_agricolai | nocturnal |
| mammals | Cryptonanus_chacoensis | Cryptonanus_chacoensis | nocturnal |
| mammals | Cryptonanus_guahybae | Cryptonanus_guahybae | nocturnal |
| mammals | Cryptonanus_unduaviensis | Cryptonanus_unduaviensis | nocturnal |
| mammals | Cryptoprocta_ferox | Cryptoprocta_ferox | cathemeral |
| mammals | Cryptotis_alticola | Cryptotis_alticola | cathemeral |
| mammals | Cryptotis_endersi | Cryptotis_endersi | cathemeral |
| mammals | Cryptotis_equatoris | Cryptotis_equatoris | cathemeral |
| mammals | Cryptotis_goldmani | Cryptotis_goldmani | cathemeral |
| mammals | Cryptotis_goodwini | Cryptotis_goodwini | nocturnal |
| mammals | Cryptotis_gracilis | Cryptotis_gracilis | nocturnal |
| mammals | Cryptotis_griseoventris | Cryptotis_griseoventris | nocturnal |
| mammals | Cryptotis_magnus | Cryptotis_magna | nocturnal |
| mammals | Cryptotis_mayensis | Cryptotis_mayensis | nocturnal |
| mammals | Cryptotis_medellinius | Cryptotis_medellinia | cathemeral |
| mammals | Cryptotis_meridensis | Cryptotis_meridensis | diurnal |
| mammals | Cryptotis_merriami | Cryptotis_merriami | nocturnal |
| mammals | Cryptotis_merus | Cryptotis_mera | cathemeral |
| mammals | Cryptotis_mexicanus | Cryptotis_mexicana | nocturnal |
| mammals | Cryptotis_montivagus | Cryptotis_montivaga | cathemeral |
| mammals | Cryptotis_orophilus | Cryptotis_orophila | nocturnal |
| mammals | Cryptotis_parvus | Cryptotis_parva | cathemeral |
| mammals | Cryptotis_peregrinus | Cryptotis_peregrina | nocturnal |
| mammals | Cryptotis_peruviensis | Cryptotis_peruviensis | cathemeral |
| mammals | Cryptotis_phillipsii | Cryptotis_phillipsii | nocturnal |
| mammals | Cryptotis_squamipes | Cryptotis_squamipes | cathemeral |
| mammals | Cryptotis_tamensis | Cryptotis_tamensis | cathemeral |
| mammals | Cryptotis_thomasi | Cryptotis_thomasi | cathemeral |
| mammals | Cryptotis_tropicalis | Cryptotis_tropicalis | cathemeral |

| | | | |
|---|---|---|---|
| mammals | Ctenodactylus_gundi | Ctenodactylus_gundi | diurnal |
| mammals | Ctenodactylus_vali | Ctenodactylus_vali | diurnal |
| mammals | Ctenomys_peruanus | Ctenomys_peruanus | diurnal |
| mammals | Cuniculus_paca | Cuniculus_paca | nocturnal |
| mammals | Cuniculus_taczanowskii | Cuniculus_taczanowskii | nocturnal |
| mammals | Cuon_alpinus | Cuon_alpinus | cathemeral |
| mammals | Cuscomys_ashaninka | Cuscomys_ashaninka | nocturnal |
| mammals | Cuscomys_oblativus | Cuscomys_oblativa | nocturnal |
| mammals | Cyclopes_didactylus | Cyclopes_didactylus | nocturnal |
| mammals | Cynictis_penicillata | Cynictis_penicillata | diurnal |
| mammals | Cynocephalus_volans | Cynocephalus_volans | nocturnal |
| mammals | Cynogale_bennettii | Cynogale_bennettii | nocturnal |
| mammals | Cynomops_abrasus | Cynomops_abrasus | nocturnal |
| mammals | Cynomops_greenhalli | Cynomops_greenhalli | nocturnal |
| mammals | Cynomops_mexicanus | Cynomops_mexicanus | nocturnal |
| mammals | Cynomops_milleri | Cynomops_milleri | nocturnal |
| mammals | Cynomops_planirostris | Cynomops_planirostris | nocturnal |
| mammals | Cynomys_gunnisoni | Cynomys_gunnisoni | diurnal |
| mammals | Cynomys_leucurus | Cynomys_leucurus | diurnal |
| mammals | Cynomys_ludovicianus | Cynomys_ludovicianus | diurnal |
| mammals | Cynomys_mexicanus | Cynomys_mexicanus | diurnal |
| mammals | Cynomys_parvidens | Cynomys_parvidens | diurnal |
| mammals | Cynopterus_brachyotis | Cynopterus_brachyotis | nocturnal |
| mammals | Cynopterus_horsfieldii | Cynopterus_horsfieldii | nocturnal |
| mammals | Cynopterus_luzoniensis | Cynopterus_luzoniensis | nocturnal |
| mammals | Cynopterus_minutus | Cynopterus_minutus | nocturnal |
| mammals | Cynopterus_nusatenggara | Cynopterus_nusatenggara | nocturnal |
| mammals | Cynopterus_sphinx | Cynopterus_sphinx | nocturnal |
| mammals | Cynopterus_titthaecheilus | Cynopterus_titthaecheilus | nocturnal |
| mammals | Cyttarops_alecto | Cyttarops_alecto | nocturnal |
| mammals | Dacnomys_millardi | Dacnomys_millardi | nocturnal |
| mammals | Dactylomys_boliviensis | Dactylomys_boliviensis | nocturnal |
| mammals | Dactylomys_dactylinus | Dactylomys_dactylinus | nocturnal |
| mammals | Dactylomys_peruanus | Dactylomys_peruanus | nocturnal |

| | | | |
|---|---|---|---|
| mammals | Dactylopsila_megalura | Dactylopsila_megalura | nocturnal |
| mammals | Dactylopsila_palpator | Dactylonax_palpator | nocturnal |
| mammals | Dactylopsila_tatei | Dactylopsila_tatei | nocturnal |
| mammals | Dactylopsila_trivirgata | Dactylopsila_trivirgata | nocturnal |
| mammals | Dama_dama | Dama_dama | cathemeral |
| mammals | Dama_mesopotamica | Dama_mesopotamica | nocturnal |
| mammals | Damaliscus_lunatus | Damaliscus_lunatus | diurnal |
| mammals | Damaliscus_pygargus | Damaliscus_pygargus | cathemeral |
| mammals | Daptomys_mussoi | Neusticomys_mussoi | nocturnal |
| mammals | Daptomys_oyapocki | Neusticomys_oyapocki | nocturnal |
| mammals | Daptomys_venezuelae | Neusticomys_venezuelae | nocturnal |
| mammals | Dasycercus_blythi | Dasycercus_blythi | nocturnal |
| mammals | Dasycercus_cristicauda | Dasycercus_cristicauda | nocturnal |
| mammals | Dasykaluta_rosamondae | Dasykaluta_rosamondae | diurnal |
| mammals | Dasymys_foxi | Dasymys_foxi | diurnal |
| mammals | Dasymys_incomtus | Dasymys_incomtus | diurnal |
| mammals | Dasymys_montanus | Dasymys_montanus | cathemeral |
| mammals | Dasymys_nudipes | Dasymys_nudipes | diurnal |
| mammals | Dasymys_rufulus | Dasymys_rufulus | nocturnal |
| mammals | Dasyprocta_azarae | Dasyprocta_azarae | diurnal |
| mammals | Dasyprocta_coibae | Dasyprocta_coibae | diurnal |
| mammals | Dasyprocta_fuliginosa | Dasyprocta_fuliginosa | diurnal |
| mammals | Dasyprocta_guamara | Dasyprocta_guamara | diurnal |
| mammals | Dasyprocta_kalinowskii | Dasyprocta_kalinowskii | diurnal |
| mammals | Dasyprocta_leporina | Dasyprocta_leporina | diurnal |
| mammals | Dasyprocta_mexicana | Dasyprocta_mexicana | diurnal |
| mammals | Dasyprocta_prymnolopha | Dasyprocta_prymnolopha | diurnal |
| mammals | Dasyprocta_punctata | Dasyprocta_punctata | diurnal |
| mammals | Dasyprocta_ruatanica | Dasyprocta_ruatanica | diurnal |
| mammals | Dasypus_kappleri | Dasypus_kappleri | nocturnal |
| mammals | Dasypus_mazzai | Dasypus_yepesi | nocturnal |
| mammals | Dasypus_novemcinctus | Dasypus_novemcinctus | cathemeral |
| mammals | Dasypus_pilosus | Dasypus_pilosus | nocturnal |
| mammals | Dasypus_sabanicola | Dasypus_sabanicola | diurnal |

| | | | |
|---|---|---|---|
| mammals | Dasypus_septemcinctus | Dasypus_septemcinctus | cathemeral |
| mammals | Dasyuroides_byrnei | Dasyuroides_byrnei | nocturnal |
| mammals | Dasyurus_albopunctatus | Dasyurus_albopunctatus | diurnal |
| mammals | Dasyurus_geoffroii | Dasyurus_geoffroii | nocturnal |
| mammals | Dasyurus_hallucatus | Dasyurus_hallucatus | nocturnal |
| mammals | Dasyurus_maculatus | Dasyurus_maculatus | cathemeral |
| mammals | Dasyurus_spartacus | Dasyurus_spartacus | nocturnal |
| mammals | Dasyurus_viverrinus | Dasyurus_viverrinus | nocturnal |
| mammals | Daubentonia_madagascariensis | Daubentonia_madagascariensis | nocturnal |
| mammals | Delanymys_brooksi | Delanymys_brooksi | nocturnal |
| mammals | Delomys_dorsalis | Delomys_dorsalis | nocturnal |
| mammals | Delomys_sublineatus | Delomys_sublineatus | nocturnal |
| mammals | Deltamys_kempi | Deltamys_kempi | nocturnal |
| mammals | Dendrogale_melanura | Dendrogale_melanura | diurnal |
| mammals | Dendrogale_murina | Dendrogale_murina | diurnal |
| mammals | Dendrohyrax_arboreus | Dendrohyrax_arboreus | cathemeral |
| mammals | Dendrohyrax_dorsalis | Dendrohyrax_dorsalis | crepuscular |
| mammals | Dendrohyrax_validus | Dendrohyrax_validus | nocturnal |
| mammals | Dendrolagus_bennettianus | Dendrolagus_bennettianus | nocturnal |
| mammals | Dendrolagus_dorianus | Dendrolagus_dorianus | diurnal |
| mammals | Dendrolagus_goodfellowi | Dendrolagus_goodfellowi | diurnal |
| mammals | Dendrolagus_inustus | Dendrolagus_inustus | crepuscular |
| mammals | Dendrolagus_lumholtzi | Dendrolagus_lumholtzi | cathemeral |
| mammals | Dendrolagus_matschiei | Dendrolagus_matschiei | diurnal |
| mammals | Dendrolagus_mbaiso | Dendrolagus_mbaiso | diurnal |
| mammals | Dendrolagus_notatus | Dendrolagus_notatus | diurnal |
| mammals | Dendrolagus_pulcherrimus | Dendrolagus_pulcherrimus | diurnal |
| mammals | Dendrolagus_scottae | Dendrolagus_scottae | diurnal |
| mammals | Dendrolagus_spadix | Dendrolagus_spadix | cathemeral |
| mammals | Dendrolagus_stellarum | Dendrolagus_stellarum | diurnal |
| mammals | Dendrolagus_ursinus | Dendrolagus_ursinus | diurnal |
| mammals | Dendromus_insignis | Dendromus_insignis | nocturnal |
| mammals | Dendromus_kahuziensis | Dendromus_kahuziensis | nocturnal |
| mammals | Dendromus_lovati | Dendromus_lovati | nocturnal |

| | | | |
|---|---|---|---|
| mammals | Dendromus_mesomelas | Dendromus_mesomelas | nocturnal |
| mammals | Dendromus_messorius | Dendromus_messorius | nocturnal |
| mammals | Dendromus_mystacalis | Dendromus_mystacalis | nocturnal |
| mammals | Dendromus_nyasae | Dendromus_kivu | nocturnal |
| mammals | Dendromus_oreas | Dendromus_oreas | nocturnal |
| mammals | Dendromus_vernayi | Dendromus_vernayi | nocturnal |
| mammals | Dendroprionomys_rousseloti | Dendroprionomys_rousseloti | nocturnal |
| mammals | Deomys_ferrugineus | Deomys_ferrugineus | nocturnal |
| mammals | Dephomys_defua | Dephomys_defua | nocturnal |
| mammals | Dermanura_anderseni | Dermanura_anderseni | nocturnal |
| mammals | Dermanura_azteca | Dermanura_azteca | nocturnal |
| mammals | Dermanura_cinerea | Dermanura_cinerea | nocturnal |
| mammals | Dermanura_glauca | Dermanura_glauca | nocturnal |
| mammals | Dermanura_gnoma | Dermanura_gnoma | nocturnal |
| mammals | Dermanura_phaeotis | Dermanura_phaeotis | nocturnal |
| mammals | Dermanura_rosenbergi | Dermanura_rosenbergi | nocturnal |
| mammals | Dermanura_tolteca | Dermanura_tolteca | nocturnal |
| mammals | Dermanura_watsoni | Dermanura_watsoni | nocturnal |
| mammals | Desmalopex_leucoptera | Pteropus_leucopterus | nocturnal |
| mammals | Desmana_moschata | Desmana_moschata | nocturnal |
| mammals | Desmodilliscus_braueri | Desmodilliscus_braueri | nocturnal |
| mammals | Desmodillus_auricularis | Desmodillus_auricularis | nocturnal |
| mammals | Desmodus_rotundus | Desmodus_rotundus | nocturnal |
| mammals | Desmomys_harringtoni | Desmomys_harringtoni | nocturnal |
| mammals | Desmomys_yaldeni | Desmomys_yaldeni | nocturnal |
| mammals | Diaemus_youngii | Diaemus_youngi | nocturnal |
| mammals | Dicerorhinus_sumatrensis | Dicerorhinus_sumatrensis | cathemeral |
| mammals | Diceros_bicornis | Diceros_bicornis | cathemeral |
| mammals | Diclidurus_albus | Diclidurus_albus | nocturnal |
| mammals | Diclidurus_ingens | Diclidurus_ingens | nocturnal |
| mammals | Diclidurus_isabella | Diclidurus_isabella | nocturnal |
| mammals | Diclidurus_scutatus | Diclidurus_scutatus | nocturnal |
| mammals | Dicotyles_tajacu | Pecari_tajacu | cathemeral |
| mammals | Dicrostonyx_hudsonius | Dicrostonyx_hudsonius | cathemeral |

| | | | |
|---|---|---|---|
| mammals | Dicrostonyx_nunatakensis | Dicrostonyx_nunatakensis | cathemeral |
| mammals | Dicrostonyx_unalascensis | Dicrostonyx_unalascensis | cathemeral |
| mammals | Didelphis_albiventris | Didelphis_albiventris | nocturnal |
| mammals | Didelphis_aurita | Didelphis_aurita | nocturnal |
| mammals | Didelphis_imperfecta | Didelphis_imperfecta | nocturnal |
| mammals | Didelphis_marsupialis | Didelphis_marsupialis | nocturnal |
| mammals | Didelphis_pernigra | Didelphis_pernigra | nocturnal |
| mammals | Didelphis_virginiana | Didelphis_virginiana | nocturnal |
| mammals | Dinaromys_bogdanovi | Dinaromys_bogdanovi | nocturnal |
| mammals | Dinomys_branickii | Dinomys_branickii | nocturnal |
| mammals | Diomys_crumpi | Diomys_crumpi | nocturnal |
| mammals | Diphylla_ecaudata | Diphylla_ecaudata | nocturnal |
| mammals | Diplogale_hosei | Diplogale_hosei | nocturnal |
| mammals | Diplomys_caniceps | Diplomys_caniceps | nocturnal |
| mammals | Diplomys_labilis | Diplomys_labilis | nocturnal |
| mammals | Diplothrix_legata | Diplothrix_legata | nocturnal |
| mammals | Dipodomys_agilis | Dipodomys_agilis | nocturnal |
| mammals | Dipodomys_californicus | Dipodomys_californicus | nocturnal |
| mammals | Dipodomys_compactus | Dipodomys_compactus | nocturnal |
| mammals | Dipodomys_deserti | Dipodomys_deserti | nocturnal |
| mammals | Dipodomys_elator | Dipodomys_elator | nocturnal |
| mammals | Dipodomys_gravipes | Dipodomys_gravipes | nocturnal |
| mammals | Dipodomys_heermanni | Dipodomys_heermanni | nocturnal |
| mammals | Dipodomys_ingens | Dipodomys_ingens | nocturnal |
| mammals | Dipodomys_merriami | Dipodomys_merriami | crepuscular |
| mammals | Dipodomys_microps | Dipodomys_microps | cathemeral |
| mammals | Dipodomys_nelsoni | Dipodomys_nelsoni | nocturnal |
| mammals | Dipodomys_nitratoides | Dipodomys_nitratoides | nocturnal |
| mammals | Dipodomys_ordii | Dipodomys_ordii | nocturnal |
| mammals | Dipodomys_panamintinus | Dipodomys_panamintinus | nocturnal |
| mammals | Dipodomys_phillipsii | Dipodomys_phillipsii | nocturnal |
| mammals | Dipodomys_simulans | Dipodomys_simulans | nocturnal |
| mammals | Dipodomys_spectabilis | Dipodomys_spectabilis | cathemeral |
| mammals | Dipodomys_stephensi | Dipodomys_stephensi | nocturnal |

| | | | |
|---|---|---|---|
| mammals | Dipodomys_venustus | Dipodomys_venustus | nocturnal |
| mammals | Dipus_sagitta | Dipus_sagitta | nocturnal |
| mammals | Distoechurus_pennatus | Distoechurus_pennatus | nocturnal |
| mammals | Dobsonia_anderseni | Dobsonia_anderseni | nocturnal |
| mammals | Dobsonia_beauforti | Dobsonia_beauforti | nocturnal |
| mammals | Dobsonia_chapmani | Dobsonia_chapmani | nocturnal |
| mammals | Dobsonia_crenulata | Dobsonia_crenulata | nocturnal |
| mammals | Dobsonia_emersa | Dobsonia_emersa | nocturnal |
| mammals | Dobsonia_exoleta | Dobsonia_exoleta | nocturnal |
| mammals | Dobsonia_inermis | Dobsonia_inermis | nocturnal |
| mammals | Dobsonia_minor | Dobsonia_minor | nocturnal |
| mammals | Dobsonia_moluccensis | Dobsonia_moluccensis | nocturnal |
| mammals | Dobsonia_pannietensis | Dobsonia_pannietensis | nocturnal |
| mammals | Dobsonia_peronii | Dobsonia_peronii | nocturnal |
| mammals | Dobsonia_praedatrix | Dobsonia_praedatrix | nocturnal |
| mammals | Dobsonia_viridis | Dobsonia_viridis | nocturnal |
| mammals | Dolichotis_patagonum | Dolichotis_patagonum | diurnal |
| mammals | Dolichotis_salinicola | Dolichotis_salinicola | diurnal |
| mammals | Dologale_dybowskii | Dologale_dybowskii | nocturnal |
| mammals | Dorcatragus_megalotis | Dorcatragus_megalotis | diurnal |
| mammals | Dorcopsis_atrata | Dorcopsis_atrata | crepuscular |
| mammals | Dorcopsis_hageni | Dorcopsis_hageni | crepuscular |
| mammals | Dorcopsis_luctuosa | Dorcopsis_luctuosa | crepuscular |
| mammals | Dorcopsis_muelleri | Dorcopsis_muelleri | nocturnal |
| mammals | Dorcopsulus_macleayi | Dorcopsulus_macleayi | nocturnal |
| mammals | Dorcopsulus_vanheurni | Dorcopsulus_vanheurni | diurnal |
| mammals | Doryrhina_camerunensis | Hipposideros_camerunensis | nocturnal |
| mammals | Doryrhina_cyclops | Hipposideros_cyclops | nocturnal |
| mammals | Dremomys_gularis | Dremomys_gularis | diurnal |
| mammals | Dremomys_lokriah | Dremomys_lokriah | diurnal |
| mammals | Dremomys_pernyi | Dremomys_pernyi | diurnal |
| mammals | Dremomys_pyrrhomerus | Dremomys_pyrrhomerus | diurnal |
| mammals | Dremomys_rufigenis | Dremomys_rufigenis | diurnal |
| mammals | Dromiciops_gliroides | Dromiciops_gliroides | nocturnal |

| | | | |
|---|---|---|---|
| mammals | Dryomys_laniger | Dryomys_laniger | nocturnal |
| mammals | Dryomys_nitedula | Dryomys_nitedula | nocturnal |
| mammals | Dyacopterus_brooksi | Dyacopterus_brooksi | nocturnal |
| mammals | Dyacopterus_spadiceus | Dyacopterus_spadiceus | nocturnal |
| mammals | Dymecodon_pilirostris | Dymecodon_pilirostris | cathemeral |
| mammals | Echimys_chrysurus | Echimys_chrysurus | nocturnal |
| mammals | Echimys_saturnus | Echimys_saturnus | nocturnal |
| mammals | Echimys_vieirai | Echimys_vieirai | nocturnal |
| mammals | Echinops_telfairi | Echinops_telfairi | nocturnal |
| mammals | Echinosorex_gymnura | Echinosorex_gymnura | nocturnal |
| mammals | Echiothrix_centrosa | Echiothrix_centrosa | nocturnal |
| mammals | Echiothrix_leucura | Echiothrix_leucura | nocturnal |
| mammals | Echymipera_clara | Echymipera_clara | nocturnal |
| mammals | Echymipera_davidi | Echymipera_davidi | nocturnal |
| mammals | Echymipera_echinista | Echymipera_echinista | nocturnal |
| mammals | Echymipera_kalubu | Echymipera_kalubu | nocturnal |
| mammals | Echymipera_rufescens | Echymipera_rufescens | nocturnal |
| mammals | Ectophylla_alba | Ectophylla_alba | nocturnal |
| mammals | Eidolon_dupreanum | Eidolon_dupreanum | nocturnal |
| mammals | Eidolon_helvum | Eidolon_helvum | nocturnal |
| mammals | Eira_barbara | Eira_barbara | diurnal |
| mammals | Elaphodus_cephalophus | Elaphodus_cephalophus | crepuscular |
| mammals | Elephantulus_brachyrhynchus | Elephantulus_brachyrhynchus | crepuscular |
| mammals | Elephantulus_edwardii | Elephantulus_edwardii | crepuscular |
| mammals | Elephantulus_fuscipes | Elephantulus_fuscipes | diurnal |
| mammals | Elephantulus_fuscus | Elephantulus_fuscus | diurnal |
| mammals | Elephantulus_intufi | Elephantulus_intufi | crepuscular |
| mammals | Elephantulus_myurus | Elephantulus_myurus | cathemeral |
| mammals | Elephantulus_rupestris | Elephantulus_rupestris | crepuscular |
| mammals | Elephas_maximus | Elephas_maximus | cathemeral |
| mammals | Eligmodontia_moreni | Eligmodontia_moreni | nocturnal |
| mammals | Eligmodontia_morgani | Eligmodontia_morgani | nocturnal |
| mammals | Eligmodontia_puerulus | Eligmodontia_puerulus | nocturnal |
| mammals | Eligmodontia_typus | Eligmodontia_typus | nocturnal |

| | | | |
|---|---|---|---|
| mammals | Eliomys_melanurus | Eliomys_melanurus | nocturnal |
| mammals | Eliomys_munbyanus | Eliomys_munbyanus | nocturnal |
| mammals | Eliomys_quercinus | Eliomys_quercinus | nocturnal |
| mammals | Eliurus_antsingy | Eliurus_antsingy | nocturnal |
| mammals | Eliurus_carletoni | Eliurus_carletoni | nocturnal |
| mammals | Eliurus_danieli | Eliurus_danieli | nocturnal |
| mammals | Eliurus_grandidieri | Eliurus_grandidieri | nocturnal |
| mammals | Eliurus_majori | Eliurus_majori | nocturnal |
| mammals | Eliurus_minor | Eliurus_minor | nocturnal |
| mammals | Eliurus_myoxinus | Eliurus_myoxinus | nocturnal |
| mammals | Eliurus_penicillatus | Eliurus_penicillatus | nocturnal |
| mammals | Eliurus_petteri | Eliurus_petteri | nocturnal |
| mammals | Eliurus_tanala | Eliurus_tanala | nocturnal |
| mammals | Eliurus_webbi | Eliurus_webbi | nocturnal |
| mammals | Emballonura_alecto | Emballonura_alecto | diurnal |
| mammals | Emballonura_beccarii | Emballonura_beccarii | diurnal |
| mammals | Emballonura_dianae | Emballonura_dianae | diurnal |
| mammals | Emballonura_furax | Emballonura_furax | diurnal |
| mammals | Emballonura_monticola | Emballonura_monticola | diurnal |
| mammals | Emballonura_raffrayana | Emballonura_raffrayana | diurnal |
| mammals | Emballonura_semicaudata | Emballonura_semicaudata | diurnal |
| mammals | Emballonura_serii | Emballonura_serii | diurnal |
| mammals | Enchisthenes_hartii | Enchisthenes_hartii | nocturnal |
| mammals | Eoglaucomys_fimbriatus | Eoglaucomys_fimbriatus | diurnal |
| mammals | Eolagurus_przewalskii | Eolagurus_przewalskii | nocturnal |
| mammals | Eonycteris_major | Eonycteris_major | nocturnal |
| mammals | Eonycteris_robusta | Eonycteris_robusta | nocturnal |
| mammals | Eonycteris_spelaea | Eonycteris_spelaea | nocturnal |
| mammals | Eothenomys_melanogaster | Eothenomys_melanogaster | cathemeral |
| mammals | Eozapus_setchuanus | Eozapus_setchuanus | nocturnal |
| mammals | Episoriculus_caudatus | Episoriculus_caudatus | nocturnal |
| mammals | Episoriculus_leucops | Episoriculus_leucops | nocturnal |
| mammals | Episoriculus_macrurus | Episoriculus_macrurus | nocturnal |
| mammals | Epixerus_ebii | Epixerus_ebii | diurnal |

| | | | |
|---|---|---|---|
| mammals | Epomophorus_angolensis | Epomophorus_angolensis | nocturnal |
| mammals | Epomophorus_anselli | Epomophorus_anselli | nocturnal |
| mammals | Epomophorus_crypturus | Epomophorus_crypturus | nocturnal |
| mammals | Epomophorus_dobsonii | Epomops_dobsonii | nocturnal |
| mammals | Epomophorus_gambianus | Epomophorus_gambianus | nocturnal |
| mammals | Epomophorus_grandis | Epomophorus_grandis | nocturnal |
| mammals | Epomophorus_intermedius | Micropteropus_intermedius | nocturnal |
| mammals | Epomophorus_labiatus | Epomophorus_labiatus | nocturnal |
| mammals | Epomophorus_minor | Epomophorus_minor | nocturnal |
| mammals | Epomophorus_pusillus | Micropteropus_pusillus | nocturnal |
| mammals | Epomophorus_wahlbergi | Epomophorus_wahlbergi | nocturnal |
| mammals | Epomops_buettikoferi | Epomops_buettikoferi | nocturnal |
| mammals | Epomops_franqueti | Epomops_franqueti | nocturnal |
| mammals | Eptesicus_fuscus | Eptesicus_fuscus | nocturnal |
| mammals | Equus_africanus | Equus_africanus | cathemeral |
| mammals | Equus_ferus | Equus_ferus | cathemeral |
| mammals | Equus_grevyi | Equus_grevyi | cathemeral |
| mammals | Equus_hemionus | Equus_hemionus | crepuscular |
| mammals | Equus_kiang | Equus_kiang | cathemeral |
| mammals | Equus_quagga | Equus_quagga | cathemeral |
| mammals | Equus_zebra | Equus_zebra | diurnal |
| mammals | Eremitalpa_granti | Eremitalpa_granti | nocturnal |
| mammals | Eremodipus_lichtensteini | Eremodipus_lichtensteini | nocturnal |
| mammals | Erethizon_dorsatum | Erethizon_dorsatum | nocturnal |
| mammals | Erinaceus_amurensis | Erinaceus_amurensis | nocturnal |
| mammals | Erinaceus_concolor | Erinaceus_concolor | nocturnal |
| mammals | Erinaceus_europaeus | Erinaceus_europaeus | nocturnal |
| mammals | Erinaceus_roumanicus | Erinaceus_roumanicus | nocturnal |
| mammals | Erophylla_bombifrons | Erophylla_bombifrons | nocturnal |
| mammals | Erophylla_sezekorni | Erophylla_sezekorni | nocturnal |
| mammals | Erythrocebus_patas | Erythrocebus_patas | diurnal |
| mammals | Euchoreutes_naso | Euchoreutes_naso | nocturnal |
| mammals | Euderma_maculatum | Euderma_maculatum | nocturnal |
| mammals | Eudiscopus_denticulus | Eudiscopus_denticulus | nocturnal |

| | | | |
|---|---|---|---|
| mammals | Eudorcas_albonotata | Eudorcas_albonotata | diurnal |
| mammals | Eudorcas_rufifrons | Eudorcas_rufifrons | crepuscular |
| mammals | Eudorcas_thomsonii | Eudorcas_thomsonii | cathemeral |
| mammals | Eulemur_albifrons | Eulemur_albifrons | cathemeral |
| mammals | Eulemur_cinereiceps | Eulemur_cinereiceps | cathemeral |
| mammals | Eulemur_collaris | Eulemur_collaris | cathemeral |
| mammals | Eulemur_coronatus | Eulemur_coronatus | cathemeral |
| mammals | Eulemur_flavifrons | Eulemur_flavifrons | cathemeral |
| mammals | Eulemur_fulvus | Eulemur_fulvus | cathemeral |
| mammals | Eulemur_macaco | Eulemur_macaco | cathemeral |
| mammals | Eulemur_mongoz | Eulemur_mongoz | cathemeral |
| mammals | Eulemur_rubriventer | Eulemur_rubriventer | cathemeral |
| mammals | Eulemur_rufifrons | Eulemur_rufifrons | cathemeral |
| mammals | Eulemur_rufus | Eulemur_rufus | cathemeral |
| mammals | Eulemur_sanfordi | Eulemur_sanfordi | cathemeral |
| mammals | Eumops_auripendulus | Eumops_auripendulus | nocturnal |
| mammals | Eumops_bonariensis | Eumops_bonariensis | nocturnal |
| mammals | Eumops_dabbenei | Eumops_dabbenei | nocturnal |
| mammals | Eumops_floridanus | Eumops_floridanus | nocturnal |
| mammals | Eumops_glaucinus | Eumops_glaucinus | nocturnal |
| mammals | Eumops_hansae | Eumops_hansae | nocturnal |
| mammals | Eumops_maurus | Eumops_maurus | nocturnal |
| mammals | Eumops_patagonicus | Eumops_patagonicus | nocturnal |
| mammals | Eumops_perotis | Eumops_perotis | nocturnal |
| mammals | Eumops_trumbulli | Eumops_trumbulli | nocturnal |
| mammals | Eumops_underwoodi | Eumops_underwoodi | nocturnal |
| mammals | Eumops_wilsoni | Eumops_wilsoni | nocturnal |
| mammals | Euneomys_chinchilloides | Euneomys_chinchilloides | nocturnal |
| mammals | Euoticus_elegantulus | Euoticus_elegantulus | nocturnal |
| mammals | Euoticus_pallidus | Euoticus_pallidus | nocturnal |
| mammals | Eupetaurus_cinereus | Eupetaurus_cinereus | nocturnal |
| mammals | Euphractus_sexcinctus | Euphractus_sexcinctus | diurnal |
| mammals | Eupleres_goudotii | Eupleres_goudotii | cathemeral |
| mammals | Euryoryzomys_macconnelli | Euryoryzomys_macconnelli | nocturnal |

| | | | |
|---|---|---|---|
| mammals | Euryoryzomys_nitidus | Euryoryzomys_nitidus | nocturnal |
| mammals | Euryoryzomys_russatus | Euryoryzomys_russatus | nocturnal |
| mammals | Euryzygomatomys_spinosus | Euryzygomatomys_spinosus | nocturnal |
| mammals | Eutamias_sibiricus | Eutamias_sibiricus | diurnal |
| mammals | Euxerus_erythropus | Xerus_erythropus | diurnal |
| mammals | Exilisciurus_concinnus | Exilisciurus_concinnus | diurnal |
| mammals | Exilisciurus_exilis | Exilisciurus_exilis | diurnal |
| mammals | Exilisciurus_whiteheadi | Exilisciurus_whiteheadi | diurnal |
| mammals | Falsistrellus_mackenziei | Falsistrellus_mackenziei | nocturnal |
| mammals | Falsistrellus_tasmaniensis | Falsistrellus_tasmaniensis | nocturnal |
| mammals | Felis_bieti | Felis_bieti | nocturnal |
| mammals | Felis_chaus | Felis_chaus | nocturnal |
| mammals | Felis_margarita | Felis_margarita | nocturnal |
| mammals | Felis_nigripes | Felis_nigripes | nocturnal |
| mammals | Felis_silvestris | Felis_silvestris | nocturnal |
| mammals | Felovia_vae | Felovia_vae | diurnal |
| mammals | Feroculus_feroculus | Feroculus_feroculus | nocturnal |
| mammals | Fossa_fossana | Fossa_fossana | nocturnal |
| mammals | Frateromys_fratrorum | Bunomys_fratrorum | nocturnal |
| mammals | Funambulus_layardi | Funambulus_layardi | diurnal |
| mammals | Funambulus_palmarum | Funambulus_palmarum | diurnal |
| mammals | Funambulus_pennantii | Funambulus_pennantii | diurnal |
| mammals | Funambulus_sublineatus | Funambulus_sublineatus | diurnal |
| mammals | Funambulus_tristriatus | Funambulus_tristriatus | diurnal |
| mammals | Funisciurus_anerythrus | Funisciurus_anerythrus | diurnal |
| mammals | Funisciurus_bayonii | Funisciurus_bayonii | diurnal |
| mammals | Funisciurus_carruthersi | Funisciurus_carruthersi | diurnal |
| mammals | Funisciurus_congicus | Funisciurus_congicus | diurnal |
| mammals | Funisciurus_duchaillui | Funisciurus_duchaillui | diurnal |
| mammals | Funisciurus_isabella | Funisciurus_isabella | diurnal |
| mammals | Funisciurus_lemniscatus | Funisciurus_lemniscatus | diurnal |
| mammals | Funisciurus_leucogenys | Funisciurus_leucogenys | diurnal |
| mammals | Funisciurus_pyrropus | Funisciurus_pyrropus | diurnal |
| mammals | Funisciurus_substriatus | Funisciurus_substriatus | diurnal |

| | | | |
|---|---|---|---|
| mammals | Furipterus_horrens | Furipterus_horrens | nocturnal |
| mammals | Galago_gallarum | Galago_gallarum | nocturnal |
| mammals | Galago_matschiei | Galago_matschiei | nocturnal |
| mammals | Galago_moholi | Galago_moholi | nocturnal |
| mammals | Galago_senegalensis | Galago_senegalensis | nocturnal |
| mammals | Galagoides_demidoff | Galagoides_demidoff | nocturnal |
| mammals | Galagoides_thomasi | Galagoides_thomasi | nocturnal |
| mammals | Galea_flavidens | Galea_flavidens | diurnal |
| mammals | Galea_leucoblephara | Galea_leucoblephara | diurnal |
| mammals | Galea_musteloides | Galea_musteloides | diurnal |
| mammals | Galea_spixii | Galea_spixii | crepuscular |
| mammals | Galegeeska_revoilii | Elephantulus_revoilii | diurnal |
| mammals | Galegeeska_rufescens | Elephantulus_rufescens | crepuscular |
| mammals | Galemys_pyrenaicus | Galemys_pyrenaicus | nocturnal |
| mammals | Galeopterus_variegatus | Galeopterus_variegatus | nocturnal |
| mammals | Galictis_cuja | Galictis_cuja | diurnal |
| mammals | Galictis_vittata | Galictis_vittata | diurnal |
| mammals | Galidia_elegans | Galidia_elegans | diurnal |
| mammals | Galidictis_fasciata | Galidictis_fasciata | nocturnal |
| mammals | Gardnerycteris_crenulata | Mimon_crenulatum | nocturnal |
| mammals | Gardnerycteris_koepckeae | Mimon_koepckeae | nocturnal |
| mammals | Gazella_bennettii | Gazella_bennettii | nocturnal |
| mammals | Gazella_cuvieri | Gazella_cuvieri | nocturnal |
| mammals | Gazella_dorcas | Gazella_dorcas | nocturnal |
| mammals | Gazella_gazella | Gazella_gazella | diurnal |
| mammals | Gazella_leptoceros | Gazella_leptoceros | crepuscular |
| mammals | Gazella_marica | Gazella_marica | crepuscular |
| mammals | Gazella_spekei | Gazella_spekei | crepuscular |
| mammals | Gazella_subgutturosa | Gazella_subgutturosa | diurnal |
| mammals | Genetta_abyssinica | Genetta_abyssinica | nocturnal |
| mammals | Genetta_angolensis | Genetta_angolensis | nocturnal |
| mammals | Genetta_bourloni | Genetta_bourloni | nocturnal |
| mammals | Genetta_cristata | Genetta_cristata | crepuscular |
| mammals | Genetta_fieldiana | Genetta_maculata | nocturnal |

| | | | |
|---|---|---|---|
| mammals | Genetta_genetta | Genetta_genetta | nocturnal |
| mammals | Genetta_johnstoni | Genetta_johnstoni | nocturnal |
| mammals | Genetta_pardina | Genetta_pardina | nocturnal |
| mammals | Genetta_piscivora | Genetta_piscivora | nocturnal |
| mammals | Genetta_poensis | Genetta_poensis | nocturnal |
| mammals | Genetta_servalina | Genetta_servalina | nocturnal |
| mammals | Genetta_thierryi | Genetta_thierryi | nocturnal |
| mammals | Genetta_tigrina | Genetta_tigrina | nocturnal |
| mammals | Genetta_victoriae | Genetta_victoriae | nocturnal |
| mammals | Geocapromys_brownii | Geocapromys_brownii | nocturnal |
| mammals | Geogale_aurita | Geogale_aurita | nocturnal |
| mammals | Geosciurus_inauris | Xerus_inauris | diurnal |
| mammals | Geosciurus_princeps | Xerus_princeps | diurnal |
| mammals | Geoxus_valdivianus | Geoxus_valdivianus | cathemeral |
| mammals | Gerbilliscus_afer | Gerbilliscus_afra | nocturnal |
| mammals | Gerbilliscus_boehmi | Gerbilliscus_boehmi | nocturnal |
| mammals | Gerbilliscus_brantsii | Gerbilliscus_brantsii | nocturnal |
| mammals | Gerbilliscus_gambianus | Gerbilliscus_gambiana | nocturnal |
| mammals | Gerbilliscus_guineae | Gerbilliscus_guineae | nocturnal |
| mammals | Gerbilliscus_inclusus | Gerbilliscus_inclusus | nocturnal |
| mammals | Gerbilliscus_kempii | Gerbilliscus_kempi | nocturnal |
| mammals | Gerbilliscus_leucogaster | Gerbilliscus_leucogaster | nocturnal |
| mammals | Gerbilliscus_nigricaudus | Gerbilliscus_nigricaudus | nocturnal |
| mammals | Gerbilliscus_paeba | Gerbillurus_paeba | nocturnal |
| mammals | Gerbilliscus_phillipsi | Gerbilliscus_phillipsi | nocturnal |
| mammals | Gerbilliscus_robustus | Gerbilliscus_robustus | nocturnal |
| mammals | Gerbilliscus_setzeri | Gerbillurus_setzeri | nocturnal |
| mammals | Gerbilliscus_tytonis | Gerbillurus_tytonis | nocturnal |
| mammals | Gerbilliscus_validus | Gerbilliscus_validus | nocturnal |
| mammals | Gerbilliscus_vallinus | Gerbillurus_vallinus | nocturnal |
| mammals | Gerbillus_acticola | Gerbillus_acticola | nocturnal |
| mammals | Gerbillus_agag | Gerbillus_cosensis | nocturnal |
| mammals | Gerbillus_amoenus | Gerbillus_amoenus | nocturnal |
| mammals | Gerbillus_andersoni | Gerbillus_andersoni | nocturnal |

| | | | |
|---|---|---|---|
| mammals | Gerbillus_aquilus | Gerbillus_aquilus | nocturnal |
| mammals | Gerbillus_bottai | Gerbillus_bottai | nocturnal |
| mammals | Gerbillus_brockmani | Gerbillus_brockmani | nocturnal |
| mammals | Gerbillus_campestris | Gerbillus_campestris | nocturnal |
| mammals | Gerbillus_cheesmani | Gerbillus_cheesmani | nocturnal |
| mammals | Gerbillus_dasyurus | Gerbillus_dasyurus | nocturnal |
| mammals | Gerbillus_dunni | Gerbillus_dunni | nocturnal |
| mammals | Gerbillus_famulus | Gerbillus_famulus | nocturnal |
| mammals | Gerbillus_floweri | Gerbillus_floweri | nocturnal |
| mammals | Gerbillus_gerbillus | Gerbillus_gerbillus | nocturnal |
| mammals | Gerbillus_gleadowi | Gerbillus_gleadowi | nocturnal |
| mammals | Gerbillus_grobbeni | Gerbillus_grobbeni | nocturnal |
| mammals | Gerbillus_harwoodi | Gerbillus_harwoodi | nocturnal |
| mammals | Gerbillus_henleyi | Gerbillus_henleyi | nocturnal |
| mammals | Gerbillus_hesperinus | Gerbillus_hesperinus | nocturnal |
| mammals | Gerbillus_hoogstraali | Gerbillus_hoogstraali | nocturnal |
| mammals | Gerbillus_jamesi | Gerbillus_jamesi | nocturnal |
| mammals | Gerbillus_latastei | Gerbillus_latastei | nocturnal |
| mammals | Gerbillus_lowei | Gerbillus_lowei | nocturnal |
| mammals | Gerbillus_mackilligini | Gerbillus_mackillingini | nocturnal |
| mammals | Gerbillus_maghrebi | Gerbillus_maghrebi | nocturnal |
| mammals | Gerbillus_mesopotamiae | Gerbillus_mesopotamiae | nocturnal |
| mammals | Gerbillus_muriculus | Gerbillus_muriculus | nocturnal |
| mammals | Gerbillus_nancillus | Gerbillus_nancillus | nocturnal |
| mammals | Gerbillus_nanus | Gerbillus_nanus | nocturnal |
| mammals | Gerbillus_nigeriae | Gerbillus_nigeriae | nocturnal |
| mammals | Gerbillus_occiduus | Gerbillus_occiduus | nocturnal |
| mammals | Gerbillus_poecilops | Gerbillus_poecilops | nocturnal |
| mammals | Gerbillus_principulus | Gerbillus_principulus | nocturnal |
| mammals | Gerbillus_pusillus | Gerbillus_pusillus | nocturnal |
| mammals | Gerbillus_pyramidum | Gerbillus_pyramidum | nocturnal |
| mammals | Gerbillus_rosalinda | Gerbillus_rosalinda | nocturnal |
| mammals | Gerbillus_rupicola | Gerbillus_rupicola | nocturnal |
| mammals | Gerbillus_simoni | Gerbillus_simoni | nocturnal |

| | | | |
|---|---|---|---|
| mammals | Gerbillus_somalicus | Gerbillus_somalicus | nocturnal |
| mammals | Gerbillus_stigmonyx | Gerbillus_stigmonyx | nocturnal |
| mammals | Gerbillus_tarabuli | Gerbillus_tarabuli | nocturnal |
| mammals | Gerbillus_watersi | Gerbillus_watersi | nocturnal |
| mammals | Giraffa_camelopardalis | Giraffa_camelopardalis | diurnal |
| mammals | Glaucomys_sabrinus | Glaucomys_sabrinus | nocturnal |
| mammals | Glaucomys_volans | Glaucomys_volans | nocturnal |
| mammals | Glauconycteris_alboguttata | Glauconycteris_alboguttata | nocturnal |
| mammals | Glauconycteris_argentata | Glauconycteris_argentata | nocturnal |
| mammals | Glauconycteris_beatrix | Glauconycteris_beatrix | nocturnal |
| mammals | Glauconycteris_curryae | Glauconycteris_curryae | nocturnal |
| mammals | Glauconycteris_egeria | Glauconycteris_egeria | nocturnal |
| mammals | Glauconycteris_gleni | Glauconycteris_gleni | nocturnal |
| mammals | Glauconycteris_humeralis | Glauconycteris_humeralis | nocturnal |
| mammals | Glauconycteris_kenyacola | Glauconycteris_kenyacola | nocturnal |
| mammals | Glauconycteris_machadoi | Glauconycteris_machadoi | nocturnal |
| mammals | Glauconycteris_poensis | Glauconycteris_poensis | nocturnal |
| mammals | Glauconycteris_superba | Glauconycteris_superba | nocturnal |
| mammals | Glauconycteris_variegata | Glauconycteris_variegata | nocturnal |
| mammals | Glironia_venusta | Glironia_venusta | nocturnal |
| mammals | Glirulus_japonicus | Glirulus_japonicus | nocturnal |
| mammals | Glis_glis | Glis_glis | nocturnal |
| mammals | Glischropus_javanus | Glischropus_javanus | nocturnal |
| mammals | Glischropus_tylopus | Glischropus_tylopus | nocturnal |
| mammals | Glossophaga_commissarisi | Glossophaga_commissarisi | nocturnal |
| mammals | Glossophaga_leachii | Glossophaga_leachii | nocturnal |
| mammals | Glossophaga_longirostris | Glossophaga_longirostris | nocturnal |
| mammals | Glossophaga_morenoi | Glossophaga_morenoi | nocturnal |
| mammals | Glossophaga_soricina | Glossophaga_soricina | nocturnal |
| mammals | Glyphonycteris_behnii | Glyphonycteris_behnii | nocturnal |
| mammals | Glyphonycteris_daviesi | Glyphonycteris_daviesi | nocturnal |
| mammals | Glyphonycteris_sylvestris | Glyphonycteris_sylvestris | nocturnal |
| mammals | Glyphotes_simus | Glyphotes_simus | diurnal |
| mammals | Golunda_ellioti | Golunda_ellioti | cathemeral |

| | | | |
|---|---|---|---|
| mammals | Gorilla_beringei | Gorilla_beringei | diurnal |
| mammals | Gorilla_gorilla | Gorilla_gorilla | diurnal |
| mammals | Gracilinanus_aceramarcae | Gracilinanus_aceramarcae | nocturnal |
| mammals | Gracilinanus_agilis | Gracilinanus_agilis | nocturnal |
| mammals | Gracilinanus_dryas | Gracilinanus_dryas | nocturnal |
| mammals | Gracilinanus_emiliae | Gracilinanus_emiliae | nocturnal |
| mammals | Gracilinanus_marica | Gracilinanus_marica | nocturnal |
| mammals | Gracilinanus_microtarsus | Gracilinanus_microtarsus | nocturnal |
| mammals | Grammomys_aridulus | Grammomys_aridulus | nocturnal |
| mammals | Grammomys_buntingi | Grammomys_buntingi | nocturnal |
| mammals | Grammomys_caniceps | Grammomys_caniceps | nocturnal |
| mammals | Grammomys_cometes | Grammomys_cometes | nocturnal |
| mammals | Grammomys_dolichurus | Grammomys_dolichurus | nocturnal |
| mammals | Grammomys_dryas | Grammomys_dryas | nocturnal |
| mammals | Grammomys_gigas | Grammomys_gigas | nocturnal |
| mammals | Grammomys_ibeanus | Grammomys_ibeanus | nocturnal |
| mammals | Grammomys_macmillani | Grammomys_macmillani | nocturnal |
| mammals | Grammomys_minnae | Grammomys_minnae | nocturnal |
| mammals | Graomys_chacoensis | Graomys_centralis | nocturnal |
| mammals | Graomys_domorum | Graomys_domorum | nocturnal |
| mammals | Graomys_griseoflavus | Graomys_griseoflavus | nocturnal |
| mammals | Graphiurus_christyi | Graphiurus_christyi | nocturnal |
| mammals | Graphiurus_crassicaudatus | Graphiurus_crassicaudatus | nocturnal |
| mammals | Graphiurus_johnstoni | Graphiurus_johnstoni | nocturnal |
| mammals | Graphiurus_kelleni | Graphiurus_kelleni | nocturnal |
| mammals | Graphiurus_lorraineus | Graphiurus_lorraineus | nocturnal |
| mammals | Graphiurus_microtis | Graphiurus_microtis | nocturnal |
| mammals | Graphiurus_monardi | Graphiurus_monardi | nocturnal |
| mammals | Graphiurus_murinus | Graphiurus_murinus | nocturnal |
| mammals | Graphiurus_nagtglasii | Graphiurus_nagtglasii | nocturnal |
| mammals | Graphiurus_ocularis | Graphiurus_ocularis | nocturnal |
| mammals | Graphiurus_platyops | Graphiurus_platyops | nocturnal |
| mammals | Graphiurus_rupicola | Graphiurus_rupicola | nocturnal |
| mammals | Graphiurus_surdus | Graphiurus_surdus | nocturnal |

| | | | |
|---|---|---|---|
| mammals | Gulo_gulo | Gulo_gulo | cathemeral |
| mammals | Gyldenstolpia_fronto | Kunsia_fronto | nocturnal |
| mammals | Gymnobelideus_leadbeateri | Gymnobelideus_leadbeateri | nocturnal |
| mammals | Gymnuromys_roberti | Gymnuromys_roberti | nocturnal |
| mammals | Habromys_chinanteco | Habromys_chinanteco | nocturnal |
| mammals | Habromys_delicatulus | Habromys_delicatulus | nocturnal |
| mammals | Habromys_ixtlani | Habromys_ixtlani | nocturnal |
| mammals | Habromys_lepturus | Habromys_lepturus | nocturnal |
| mammals | Habromys_lophurus | Habromys_lophurus | nocturnal |
| mammals | Habromys_schmidlyi | Habromys_schmidlyi | nocturnal |
| mammals | Habromys_simulatus | Habromys_simulatus | nocturnal |
| mammals | Hadromys_humei | Hadromys_humei | nocturnal |
| mammals | Hadromys_yunnanensis | Hadromys_yunnanensis | nocturnal |
| mammals | Haeromys_margarettae | Haeromys_margarettae | nocturnal |
| mammals | Haeromys_minahassae | Haeromys_minahassae | nocturnal |
| mammals | Haeromys_pusillus | Haeromys_pusillus | nocturnal |
| mammals | Handleyomys_alfaroi | Handleyomys_alfaroi | nocturnal |
| mammals | Handleyomys_chapmani | Handleyomys_chapmani | nocturnal |
| mammals | Handleyomys_fuscatus | Handleyomys_fuscatus | nocturnal |
| mammals | Handleyomys_intectus | Handleyomys_intectus | nocturnal |
| mammals | Handleyomys_melanotis | Handleyomys_melanotis | nocturnal |
| mammals | Handleyomys_rhabdops | Handleyomys_rhabdops | nocturnal |
| mammals | Handleyomys_rostratus | Handleyomys_rostratus | nocturnal |
| mammals | Hapalemur_alaotrensis | Hapalemur_alaotrensis | cathemeral |
| mammals | Hapalemur_aureus | Hapalemur_aureus | cathemeral |
| mammals | Hapalemur_griseus | Hapalemur_griseus | cathemeral |
| mammals | Hapalemur_meridionalis | Hapalemur_meridionalis | cathemeral |
| mammals | Hapalemur_occidentalis | Hapalemur_occidentalis | cathemeral |
| mammals | Hapalomys_delacouri | Hapalomys_delacouri | nocturnal |
| mammals | Hapalomys_longicaudatus | Hapalomys_longicaudatus | nocturnal |
| mammals | Haplonycteris_fischeri | Haplonycteris_fischeri | nocturnal |
| mammals | Harpiocephalus_harpia | Harpiocephalus_harpia | nocturnal |
| mammals | Harpiola_grisea | Harpiola_grisea | nocturnal |
| mammals | Harpiola_isodon | Harpiola_isodon | nocturnal |

| | | | |
|---|---|---|---|
| mammals | Harpyionycteris_celebensis | Harpyionycteris_celebensis | nocturnal |
| mammals | Harpyionycteris_whiteheadi | Harpyionycteris_whiteheadi | nocturnal |
| mammals | Heimyscus_fumosus | Heimyscus_fumosus | nocturnal |
| mammals | Helarctos_malayanus | Helarctos_malayanus | cathemeral |
| mammals | Heliosciurus_gambianus | Heliosciurus_gambianus | diurnal |
| mammals | Heliosciurus_mutabilis | Heliosciurus_mutabilis | diurnal |
| mammals | Heliosciurus_punctatus | Heliosciurus_punctatus | diurnal |
| mammals | Heliosciurus_rufobrachium | Heliosciurus_rufobrachium | diurnal |
| mammals | Heliosciurus_ruwenzorii | Heliosciurus_ruwenzorii | diurnal |
| mammals | Heliosciurus_undulatus | Heliosciurus_undulatus | diurnal |
| mammals | Helogale_hirtula | Helogale_hirtula | diurnal |
| mammals | Helogale_parvula | Helogale_parvula | diurnal |
| mammals | Hemibelideus_lemuroides | Hemibelideus_lemuroides | nocturnal |
| mammals | Hemicentetes_nigriceps | Hemicentetes_nigriceps | nocturnal |
| mammals | Hemicentetes_semispinosus | Hemicentetes_semispinosus | cathemeral |
| mammals | Hemiechinus_auritus | Hemiechinus_auritus | nocturnal |
| mammals | Hemiechinus_collaris | Hemiechinus_collaris | nocturnal |
| mammals | Hemigalus_derbyanus | Hemigalus_derbyanus | nocturnal |
| mammals | Hemitragus_jemlahicus | Hemitragus_jemlahicus | diurnal |
| mammals | Herpailurus_yagouaroundi | Herpailurus_yagouaroundi | diurnal |
| mammals | Herpestes_flavescens | Herpestes_flavescens | diurnal |
| mammals | Herpestes_ichneumon | Herpestes_ichneumon | nocturnal |
| mammals | Herpestes_ochraceus | Herpestes_ochraceus | diurnal |
| mammals | Herpestes_pulverulentus | Herpestes_pulverulentus | diurnal |
| mammals | Herpestes_sanguineus | Herpestes_sanguineus | diurnal |
| mammals | Hesperoptenus_blanfordi | Hesperoptenus_blanfordi | nocturnal |
| mammals | Hesperoptenus_doriae | Hesperoptenus_doriae | nocturnal |
| mammals | Hesperoptenus_tickelli | Hesperoptenus_tickelli | nocturnal |
| mammals | Hesperoptenus_tomesi | Hesperoptenus_tomesi | nocturnal |
| mammals | Heterogeomys_cavator | Orthogeomys_cavator | cathemeral |
| mammals | Heterogeomys_cherriei | Heterogeomys_cherriei | cathemeral |
| mammals | Heterogeomys_dariensis | Heterogeomys_dariensis | cathemeral |
| mammals | Heterogeomys_heterodus | Orthogeomys_heterodus | cathemeral |
| mammals | Heterogeomys_hispidus | Orthogeomys_hispidus | cathemeral |

| | | | |
|---|---|---|---|
| mammals | Heterogeomys_lanius | Orthogeomys_lanius | cathemeral |
| mammals | Heterogeomys_underwoodi | Orthogeomys_underwoodi | cathemeral |
| mammals | Heterohyrax_brucei | Heterohyrax_brucei | diurnal |
| mammals | Heteromys_adspersus | Heteromys_adspersus | nocturnal |
| mammals | Heteromys_anomalus | Heteromys_anomalus | nocturnal |
| mammals | Heteromys_australis | Heteromys_australis | nocturnal |
| mammals | Heteromys_desmarestianus | Heteromys_desmarestianus | nocturnal |
| mammals | Heteromys_gaumeri | Heteromys_gaumeri | nocturnal |
| mammals | Heteromys_irroratus | Heteromys_irroratus | nocturnal |
| mammals | Heteromys_nelsoni | Heteromys_nelsoni | nocturnal |
| mammals | Heteromys_oasicus | Heteromys_oasicus | nocturnal |
| mammals | Heteromys_oresterus | Heteromys_oresterus | nocturnal |
| mammals | Heteromys_pictus | Heteromys_pictus | nocturnal |
| mammals | Heteromys_salvini | Heteromys_salvini | nocturnal |
| mammals | Heteromys_spectabilis | Heteromys_spectabilis | nocturnal |
| mammals | Heteromys_teleus | Heteromys_teleus | nocturnal |
| mammals | Hippocamelus_antisensis | Hippocamelus_antisensis | diurnal |
| mammals | Hippocamelus_bisulcus | Hippocamelus_bisulcus | crepuscular |
| mammals | Hippopotamus_amphibius | Hippopotamus_amphibius | cathemeral |
| mammals | Hipposideros_abae | Hipposideros_abae | nocturnal |
| mammals | Hipposideros_armiger | Hipposideros_armiger | nocturnal |
| mammals | Hipposideros_ater | Hipposideros_ater | nocturnal |
| mammals | Hipposideros_beatus | Hipposideros_beatus | nocturnal |
| mammals | Hipposideros_bicolor | Hipposideros_bicolor | nocturnal |
| mammals | Hipposideros_boeadii | Hipposideros_boeadii | nocturnal |
| mammals | Hipposideros_breviceps | Hipposideros_breviceps | nocturnal |
| mammals | Hipposideros_caffer | Hipposideros_caffer | nocturnal |
| mammals | Hipposideros_calcaratus | Hipposideros_calcaratus | nocturnal |
| mammals | Hipposideros_cervinus | Hipposideros_cervinus | nocturnal |
| mammals | Hipposideros_cineraceus | Hipposideros_cineraceus | nocturnal |
| mammals | Hipposideros_coronatus | Hipposideros_coronatus | nocturnal |
| mammals | Hipposideros_corynophyllus | Hipposideros_corynophyllus | nocturnal |
| mammals | Hipposideros_coxi | Hipposideros_coxi | nocturnal |
| mammals | Hipposideros_crumeniferus | Hipposideros_crumeniferus | nocturnal |

| | | | |
|---|---|---|---|
| mammals | Hipposideros_curtus | Hipposideros_curtus | nocturnal |
| mammals | Hipposideros_demissus | Hipposideros_demissus | nocturnal |
| mammals | Hipposideros_diadema | Hipposideros_diadema | nocturnal |
| mammals | Hipposideros_dinops | Hipposideros_dinops | nocturnal |
| mammals | Hipposideros_doriae | Hipposideros_doriae | nocturnal |
| mammals | Hipposideros_durgadasi | Hipposideros_durgadasi | nocturnal |
| mammals | Hipposideros_dyacorum | Hipposideros_dyacorum | nocturnal |
| mammals | Hipposideros_edwardshilli | Hipposideros_edwardshilli | nocturnal |
| mammals | Hipposideros_fuliginosus | Hipposideros_fuliginosus | nocturnal |
| mammals | Hipposideros_fulvus | Hipposideros_fulvus | nocturnal |
| mammals | Hipposideros_galeritus | Hipposideros_galeritus | nocturnal |
| mammals | Hipposideros_grandis | Hipposideros_grandis | nocturnal |
| mammals | Hipposideros_halophyllus | Hipposideros_halophyllus | nocturnal |
| mammals | Hipposideros_inexpectatus | Hipposideros_inexpectatus | nocturnal |
| mammals | Hipposideros_inornatus | Hipposideros_inornatus | nocturnal |
| mammals | Hipposideros_jonesi | Hipposideros_jonesi | nocturnal |
| mammals | Hipposideros_khaokhouayensis | Hipposideros_khaokhouayensis | nocturnal |
| mammals | Hipposideros_lamottei | Hipposideros_lamottei | nocturnal |
| mammals | Hipposideros_lankadiva | Hipposideros_lankadiva | nocturnal |
| mammals | Hipposideros_larvatus | Hipposideros_larvatus | nocturnal |
| mammals | Hipposideros_lekaguli | Hipposideros_lekaguli | nocturnal |
| mammals | Hipposideros_lylei | Hipposideros_lylei | nocturnal |
| mammals | Hipposideros_macrobullatus | Hipposideros_macrobullatus | nocturnal |
| mammals | Hipposideros_madurae | Hipposideros_madurae | nocturnal |
| mammals | Hipposideros_maggietaylorae | Hipposideros_maggietaylorae | nocturnal |
| mammals | Hipposideros_marisae | Hipposideros_marisae | nocturnal |
| mammals | Hipposideros_megalotis | Hipposideros_megalotis | nocturnal |
| mammals | Hipposideros_muscinus | Hipposideros_muscinus | nocturnal |
| mammals | Hipposideros_nequam | Hipposideros_nequam | nocturnal |
| mammals | Hipposideros_obscurus | Hipposideros_obscurus | nocturnal |
| mammals | Hipposideros_orbiculus | Hipposideros_orbiculus | nocturnal |
| mammals | Hipposideros_papua | Hipposideros_papua | nocturnal |
| mammals | Hipposideros_pelingensis | Hipposideros_pelingensis | nocturnal |
| mammals | Hipposideros_pomona | Hipposideros_pomona | nocturnal |

| | | | |
|---|---|---|---|
| mammals | Hipposideros_pygmaeus | Hipposideros_pygmaeus | nocturnal |
| mammals | Hipposideros_ridleyi | Hipposideros_ridleyi | nocturnal |
| mammals | Hipposideros_rotalis | Hipposideros_rotalis | nocturnal |
| mammals | Hipposideros_ruber | Hipposideros_ruber | cathemeral |
| mammals | Hipposideros_scutinares | Hipposideros_scutinares | nocturnal |
| mammals | Hipposideros_semoni | Hipposideros_semoni | nocturnal |
| mammals | Hipposideros_sorenseni | Hipposideros_sorenseni | nocturnal |
| mammals | Hipposideros_speoris | Hipposideros_speoris | nocturnal |
| mammals | Hipposideros_stenotis | Hipposideros_stenotis | nocturnal |
| mammals | Hipposideros_sumbae | Hipposideros_sumbae | nocturnal |
| mammals | Hipposideros_swinhoii | Hipposideros_pratti | nocturnal |
| mammals | Hipposideros_turpis | Hipposideros_turpis | nocturnal |
| mammals | Hipposideros_wollastoni | Hipposideros_wollastoni | nocturnal |
| mammals | Hippotragus_equinus | Hippotragus_equinus | diurnal |
| mammals | Hippotragus_niger | Hippotragus_niger | cathemeral |
| mammals | Histiotus_alienus | Histiotus_alienus | nocturnal |
| mammals | Histiotus_humboldti | Histiotus_humboldti | nocturnal |
| mammals | Histiotus_laephotis | Histiotus_laephotis | nocturnal |
| mammals | Histiotus_macrotus | Histiotus_macrotus | nocturnal |
| mammals | Histiotus_magellanicus | Histiotus_magellanicus | nocturnal |
| mammals | Histiotus_montanus | Histiotus_montanus | nocturnal |
| mammals | Histiotus_velatus | Histiotus_velatus | nocturnal |
| mammals | Hodomys_alleni | Hodomys_alleni | nocturnal |
| mammals | Holochilus_brasiliensis | Holochilus_brasiliensis | nocturnal |
| mammals | Holochilus_chacarius | Holochilus_chacarius | nocturnal |
| mammals | Holochilus_sciureus | Holochilus_sciureus | nocturnal |
| mammals | Hoolock_hoolock | Hoolock_hoolock | diurnal |
| mammals | Hoolock_leuconedys | Hoolock_leuconedys | diurnal |
| mammals | Hoplomys_gymnurus | Hoplomys_gymnurus | nocturnal |
| mammals | Hsunycteris_thomasi | Lonchophylla_thomasi | nocturnal |
| mammals | Hyaena_hyaena | Hyaena_hyaena | nocturnal |
| mammals | Hybomys_basilii | Hybomys_basilii | nocturnal |
| mammals | Hybomys_lunaris | Hybomys_lunaris | nocturnal |
| mammals | Hybomys_rufocanus | Hybomys_badius | cathemeral |

| | | | |
|---|---|---|---|
| mammals | Hybomys_univittatus | Hybomys_univittatus | cathemeral |
| mammals | Hydrictis_maculicollis | Hydrictis_maculicollis | diurnal |
| mammals | Hydrochoerus_hydrochaeris | Hydrochoerus_hydrochaeris | cathemeral |
| mammals | Hydrochoerus_isthmius | Hydrochoerus_isthmius | nocturnal |
| mammals | Hydromys_chrysogaster | Hydromys_chrysogaster | nocturnal |
| mammals | Hydromys_hussoni | Hydromys_hussoni | nocturnal |
| mammals | Hydromys_neobrittanicus | Hydromys_neobritannicus | nocturnal |
| mammals | Hydromys_ziegleri | Hydromys_ziegleri | nocturnal |
| mammals | Hydropotes_inermis | Hydropotes_inermis | cathemeral |
| mammals | Hyemoschus_aquaticus | Hyemoschus_aquaticus | nocturnal |
| mammals | Hyladelphys_kalinowskii | Hyladelphys_kalinowskii | nocturnal |
| mammals | Hylaeamys_megacephalus | Hylaeamys_megacephalus | nocturnal |
| mammals | Hylaeamys_perenensis | Hylaeamys_perenensis | nocturnal |
| mammals | Hylaeamys_tatei | Hylaeamys_tatei | nocturnal |
| mammals | Hylaeamys_yunganus | Hylaeamys_yunganus | nocturnal |
| mammals | Hylobates_agilis | Hylobates_agilis | diurnal |
| mammals | Hylobates_albibarbis | Hylobates_albibarbis | diurnal |
| mammals | Hylobates_klossii | Hylobates_klossii | diurnal |
| mammals | Hylobates_lar | Hylobates_lar | diurnal |
| mammals | Hylobates_moloch | Hylobates_moloch | diurnal |
| mammals | Hylobates_muelleri | Hylobates_muelleri | diurnal |
| mammals | Hylobates_pileatus | Hylobates_pileatus | diurnal |
| mammals | Hylochoerus_meinertzhageni | Hylochoerus_meinertzhageni | diurnal |
| mammals | Hylomys_parvus | Hylomys_parvus | cathemeral |
| mammals | Hylomys_suillus | Hylomys_suillus | diurnal |
| mammals | Hylomyscus_aeta | Hylomyscus_aeta | nocturnal |
| mammals | Hylomyscus_alleni | Hylomyscus_alleni | nocturnal |
| mammals | Hylomyscus_baeri | Hylomyscus_baeri | nocturnal |
| mammals | Hylomyscus_carillus | Hylomyscus_carillus | nocturnal |
| mammals | Hylomyscus_denniae | Hylomyscus_denniae | nocturnal |
| mammals | Hylomyscus_grandis | Hylomyscus_grandis | nocturnal |
| mammals | Hylomyscus_parvus | Hylomyscus_parvus | nocturnal |
| mammals | Hylomyscus_stella | Hylomyscus_stella | nocturnal |
| mammals | Hylomyscus_walterverheyeni | Hylomyscus_walterverheyeni | nocturnal |

| | | | |
|---|---|---|---|
| mammals | Hylonycteris_underwoodi | Hylonycteris_underwoodi | nocturnal |
| mammals | Hylopetes_alboniger | Hylopetes_alboniger | nocturnal |
| mammals | Hylopetes_bartelsi | Hylopetes_bartelsi | nocturnal |
| mammals | Hylopetes_nigripes | Hylopetes_nigripes | nocturnal |
| mammals | Hylopetes_phayrei | Hylopetes_phayrei | nocturnal |
| mammals | Hylopetes_platyurus | Hylopetes_platyurus | nocturnal |
| mammals | Hylopetes_sagitta | Petinomys_sagitta | nocturnal |
| mammals | Hylopetes_sipora | Hylopetes_sipora | nocturnal |
| mammals | Hylopetes_spadiceus | Hylopetes_spadiceus | nocturnal |
| mammals | Hylopetes_winstoni | Hylopetes_winstoni | nocturnal |
| mammals | Hyomys_dammermani | Hyomys_dammermani | nocturnal |
| mammals | Hyomys_goliath | Hyomys_goliath | nocturnal |
| mammals | Hyosciurus_heinrichi | Hyosciurus_heinrichi | diurnal |
| mammals | Hyosciurus_ileile | Hyosciurus_ileile | diurnal |
| mammals | Hypogeomys_antimena | Hypogeomys_antimena | nocturnal |
| mammals | Hypsignathus_monstrosus | Hypsignathus_monstrosus | nocturnal |
| mammals | Hypsiprymnodon_moschatus | Hypsiprymnodon_moschatus | diurnal |
| mammals | Hypsugo_affinis | Falsistrellus_affinis | nocturnal |
| mammals | Hypsugo_alaschanicus | Pipistrellus_alaschanicus | nocturnal |
| mammals | Hypsugo_arabicus | Pipistrellus_arabicus | nocturnal |
| mammals | Hypsugo_ariel | Pipistrellus_ariel | nocturnal |
| mammals | Hypsugo_cadornae | Pipistrellus_cadornae | nocturnal |
| mammals | Hypsugo_imbricatus | Pipistrellus_imbricatus | nocturnal |
| mammals | Hypsugo_kitcheneri | Hypsugo_kitcheneri | nocturnal |
| mammals | Hypsugo_macrotis | Hypsugo_macrotis | nocturnal |
| mammals | Hypsugo_mordax | Falsistrellus_mordax | nocturnal |
| mammals | Hypsugo_musciculus | Pipistrellus_musciculus | nocturnal |
| mammals | Hypsugo_petersi | Falsistrellus_petersi | nocturnal |
| mammals | Hypsugo_pulveratus | Pipistrellus_pulveratus | nocturnal |
| mammals | Hypsugo_savii | Hypsugo_savii | nocturnal |
| mammals | Hypsugo_vordermanni | Hypsugo_vordermanni | nocturnal |
| mammals | Hystrix_africaeaustralis | Hystrix_africaeaustralis | nocturnal |
| mammals | Hystrix_brachyura | Hystrix_brachyura | nocturnal |
| mammals | Hystrix_crassispinis | Hystrix_crassispinis | nocturnal |

| | | | |
|---|---|---|---|
| mammals | Hystrix_cristata | Hystrix_cristata | nocturnal |
| mammals | Hystrix_indica | Hystrix_indica | nocturnal |
| mammals | Hystrix_javanica | Hystrix_javanica | nocturnal |
| mammals | Hystrix_pumila | Hystrix_pumila | nocturnal |
| mammals | Hystrix_sumatrae | Hystrix_sumatrae | nocturnal |
| mammals | Ia_io | Ia_io | nocturnal |
| mammals | Ichneumia_albicauda | Ichneumia_albicauda | nocturnal |
| mammals | Ichthyomys_hydrobates | Ichthyomys_hydrobates | cathemeral |
| mammals | Ichthyomys_pittieri | Ichthyomys_pittieri | nocturnal |
| mammals | Ichthyomys_stolzmanni | Ichthyomys_stolzmanni | cathemeral |
| mammals | Ichthyomys_tweedii | Ichthyomys_tweedii | cathemeral |
| mammals | Ictidomys_mexicanus | Ictidomys_mexicanus | diurnal |
| mammals | Ictidomys_tridecemlineatus | Ictidomys_tridecemlineatus | diurnal |
| mammals | Ictonyx_striatus | Ictonyx_striatus | nocturnal |
| mammals | Idionycteris_phyllotis | Idionycteris_phyllotis | nocturnal |
| mammals | Idiurus_macrotis | Idiurus_macrotis | nocturnal |
| mammals | Idiurus_zenkeri | Idiurus_zenkeri | nocturnal |
| mammals | Indri_indri | Indri_indri | nocturnal |
| mammals | Iomys_horsfieldii | Iomys_horsfieldii | nocturnal |
| mammals | Iomys_sipora | Iomys_sipora | diurnal |
| mammals | Irenomys_tarsalis | Irenomys_tarsalis | nocturnal |
| mammals | Isoodon_auratus | Isoodon_auratus | nocturnal |
| mammals | Isoodon_macrourus | Isoodon_macrourus | nocturnal |
| mammals | Isoodon_obesulus | Isoodon_obesulus | nocturnal |
| mammals | Isothrix_barbarabrownae | Isothrix_barbarabrownae | nocturnal |
| mammals | Isothrix_bistriata | Isothrix_bistriata | nocturnal |
| mammals | Isothrix_negrensis | Isothrix_negrensis | nocturnal |
| mammals | Isothrix_pagurus | Isothrix_pagurus | nocturnal |
| mammals | Isothrix_sinnamariensis | Isothrix_sinnamariensis | nocturnal |
| mammals | Isthmomys_pirrensis | Isthmomys_pirrensis | nocturnal |
| mammals | Jaculus_blanfordi | Jaculus_blanfordi | nocturnal |
| mammals | Jaculus_jaculus | Jaculus_jaculus | nocturnal |
| mammals | Jaculus_orientalis | Jaculus_orientalis | nocturnal |
| mammals | Juliomys_pictipes | Juliomys_pictipes | nocturnal |

| | | | |
|---|---|---|---|
| mammals | Juliomys_rimofrons | Juliomys_rimofrons | nocturnal |
| mammals | Juscelinomys_huanchacae | Juscelinomys_huanchacae | nocturnal |
| mammals | Kadarsanomys_sodyi | Kadarsanomys_sodyi | nocturnal |
| mammals | Kannabateomys_amblyonyx | Kannabateomys_amblyonyx | nocturnal |
| mammals | Kerivoula_africana | Kerivoula_africana | nocturnal |
| mammals | Kerivoula_agnella | Kerivoula_agnella | nocturnal |
| mammals | Kerivoula_argentata | Kerivoula_argentata | nocturnal |
| mammals | Kerivoula_cuprosa | Kerivoula_cuprosa | nocturnal |
| mammals | Kerivoula_eriophora | Kerivoula_eriophora | nocturnal |
| mammals | Kerivoula_flora | Kerivoula_flora | nocturnal |
| mammals | Kerivoula_hardwickii | Kerivoula_hardwickii | nocturnal |
| mammals | Kerivoula_intermedia | Kerivoula_intermedia | nocturnal |
| mammals | Kerivoula_kachinensis | Kerivoula_kachinensis | nocturnal |
| mammals | Kerivoula_krauensis | Kerivoula_krauensis | nocturnal |
| mammals | Kerivoula_lanosa | Kerivoula_lanosa | nocturnal |
| mammals | Kerivoula_lenis | Kerivoula_lenis | nocturnal |
| mammals | Kerivoula_minuta | Kerivoula_minuta | nocturnal |
| mammals | Kerivoula_muscina | Kerivoula_muscina | nocturnal |
| mammals | Kerivoula_myrella | Kerivoula_myrella | nocturnal |
| mammals | Kerivoula_papillosa | Kerivoula_papillosa | nocturnal |
| mammals | Kerivoula_pellucida | Kerivoula_pellucida | nocturnal |
| mammals | Kerivoula_phalaena | Kerivoula_phalaena | nocturnal |
| mammals | Kerivoula_picta | Kerivoula_picta | nocturnal |
| mammals | Kerivoula_smithii | Kerivoula_smithii | nocturnal |
| mammals | Kerivoula_titania | Kerivoula_titania | nocturnal |
| mammals | Kerivoula_whiteheadi | Kerivoula_whiteheadi | nocturnal |
| mammals | Kerodon_acrobata | Kerodon_acrobata | crepuscular |
| mammals | Kerodon_rupestris | Kerodon_rupestris | crepuscular |
| mammals | Kobus_ellipsiprymnus | Kobus_ellipsiprymnus | crepuscular |
| mammals | Kobus_kob | Kobus_kob | cathemeral |
| mammals | Kobus_leche | Kobus_leche | diurnal |
| mammals | Kobus_megaceros | Kobus_megaceros | diurnal |
| mammals | Kobus_vardonii | Kobus_vardonii | crepuscular |
| mammals | Komodomys_rintjanus | Komodomys_rintjanus | nocturnal |

| | | | |
|---|---|---|---|
| mammals | Kunsia_tomentosus | Kunsia_tomentosus | cathemeral |
| mammals | Laephotis_angolensis | Laephotis_angolensis | nocturnal |
| mammals | Laephotis_capensis | Neoromicia_capensis | nocturnal |
| mammals | Laephotis_malagasyensis | Neoromicia_malagasyensis | nocturnal |
| mammals | Laephotis_matroka | Neoromicia_matroka | nocturnal |
| mammals | Laephotis_namibensis | Laephotis_namibensis | nocturnal |
| mammals | Laephotis_wintoni | Laephotis_wintoni | nocturnal |
| mammals | Lagidium_viscacia | Lagidium_viscacia | cathemeral |
| mammals | Lagidium_wolffsohni | Lagidium_wolffsohni | diurnal |
| mammals | Lagorchestes_conspicillatus | Lagorchestes_conspicillatus | nocturnal |
| mammals | Lagorchestes_hirsutus | Lagorchestes_hirsutus | nocturnal |
| mammals | Lagostomus_maximus | Lagostomus_maximus | nocturnal |
| mammals | Lagostrophus_fasciatus | Lagostrophus_fasciatus | nocturnal |
| mammals | Lagothrix_flavicauda | Oreonax_flavicauda | diurnal |
| mammals | Lagothrix_lagothricha | Lagothrix_lagotricha | diurnal |
| mammals | Lagurus_lagurus | Lagurus_lagurus | cathemeral |
| mammals | Lama_guanicoe | Lama_guanicoe | diurnal |
| mammals | Lama_vicugna | Vicugna_vicugna | diurnal |
| mammals | Lampronycteris_brachyotis | Lampronycteris_brachyotis | nocturnal |
| mammals | Laonastes_aenigmamus | Laonastes_aenigmamus | cathemeral |
| mammals | Lariscus_hosei | Lariscus_hosei | diurnal |
| mammals | Lariscus_insignis | Lariscus_insignis | diurnal |
| mammals | Lariscus_niobe | Lariscus_niobe | diurnal |
| mammals | Lariscus_obscurus | Lariscus_obscurus | diurnal |
| mammals | Lasionycteris_noctivagans | Lasionycteris_noctivagans | nocturnal |
| mammals | Lasiopodomys_brandtii | Lasiopodomys_brandtii | diurnal |
| mammals | Lasiorhinus_krefftii | Lasiorhinus_krefftii | nocturnal |
| mammals | Lasiurus_atratus | Lasiurus_atratus | nocturnal |
| mammals | Lasiurus_blossevillii | Lasiurus_blossevillii | nocturnal |
| mammals | Lasiurus_borealis | Lasiurus_borealis | nocturnal |
| mammals | Lasiurus_castaneus | Lasiurus_castaneus | nocturnal |
| mammals | Lasiurus_cinereus | Lasiurus_cinereus | nocturnal |
| mammals | Lasiurus_degelidus | Lasiurus_degelidus | nocturnal |
| mammals | Lasiurus_ega | Lasiurus_ega | nocturnal |

| | | | |
|---|---|---|---|
| mammals | Lasiurus_egregius | Lasiurus_egregius | nocturnal |
| mammals | Lasiurus_insularis | Lasiurus_insularis | nocturnal |
| mammals | Lasiurus_intermedius | Lasiurus_intermedius | nocturnal |
| mammals | Lasiurus_minor | Lasiurus_minor | nocturnal |
| mammals | Lasiurus_pfeifferi | Lasiurus_pfeifferi | nocturnal |
| mammals | Lasiurus_seminolus | Lasiurus_seminolus | nocturnal |
| mammals | Lasiurus_varius | Lasiurus_varius | nocturnal |
| mammals | Lasiurus_xanthinus | Lasiurus_xanthinus | nocturnal |
| mammals | Latidens_salimalii | Latidens_salimalii | nocturnal |
| mammals | Lavia_frons | Lavia_frons | diurnal |
| mammals | Leggadina_forresti | Leggadina_forresti | nocturnal |
| mammals | Leggadina_lakedownensis | Leggadina_lakedownensis | nocturnal |
| mammals | Leimacomys_buettneri | Leimacomys_buettneri | nocturnal |
| mammals | Leiuromys_occasius | Pattonomys_occasius | nocturnal |
| mammals | Lemmiscus_curtatus | Lemmiscus_curtatus | crepuscular |
| mammals | Lemmus_amurensis | Lemmus_amurensis | cathemeral |
| mammals | Lemmus_lemmus | Lemmus_lemmus | cathemeral |
| mammals | Lemniscomys_barbarus | Lemniscomys_barbarus | nocturnal |
| mammals | Lemniscomys_griselda | Lemniscomys_griselda | nocturnal |
| mammals | Lemniscomys_hoogstraali | Lemniscomys_hoogstraali | nocturnal |
| mammals | Lemniscomys_linulus | Lemniscomys_linulus | nocturnal |
| mammals | Lemniscomys_macculus | Lemniscomys_macculus | cathemeral |
| mammals | Lemniscomys_mittendorfi | Lemniscomys_mittendorfi | nocturnal |
| mammals | Lemniscomys_rosalia | Lemniscomys_rosalia | nocturnal |
| mammals | Lemniscomys_roseveari | Lemniscomys_roseveari | nocturnal |
| mammals | Lemniscomys_striatus | Lemniscomys_striatus | crepuscular |
| mammals | Lemniscomys_zebra | Lemniscomys_zebra | crepuscular |
| mammals | Lemur_catta | Lemur_catta | cathemeral |
| mammals | Lenomys_meyeri | Lenomys_meyeri | nocturnal |
| mammals | Lenothrix_cana | Lenothrix_canus | nocturnal |
| mammals | Leontopithecus_caissara | Leontopithecus_caissara | diurnal |
| mammals | Leontopithecus_chrysomelas | Leontopithecus_chrysomelas | diurnal |
| mammals | Leontopithecus_chrysopygus | Leontopithecus_chrysopygus | diurnal |
| mammals | Leontopithecus_rosalia | Leontopithecus_rosalia | diurnal |

| | | | |
|---|---|---|---|
| mammals | Leopardus_colocola | Leopardus_colocolo | cathemeral |
| mammals | Leopardus_geoffroyi | Leopardus_geoffroyi | cathemeral |
| mammals | Leopardus_guigna | Leopardus_guigna | nocturnal |
| mammals | Leopardus_guttulus | Leopardus_guttulus | cathemeral |
| mammals | Leopardus_jacobita | Leopardus_jacobita | diurnal |
| mammals | Leopardus_pardalis | Leopardus_pardalis | cathemeral |
| mammals | Leopardus_tigrinus | Leopardus_tigrinus | cathemeral |
| mammals | Leopardus_wiedii | Leopardus_wiedii | cathemeral |
| mammals | Leopoldamys_ciliatus | Leopoldamys_ciliatus | nocturnal |
| mammals | Leopoldamys_edwardsi | Leopoldamys_edwardsi | nocturnal |
| mammals | Leopoldamys_milleti | Leopoldamys_milleti | nocturnal |
| mammals | Leopoldamys_neilli | Leopoldamys_neilli | nocturnal |
| mammals | Leopoldamys_sabanus | Leopoldamys_sabanus | nocturnal |
| mammals | Leopoldamys_siporanus | Leopoldamys_siporanus | nocturnal |
| mammals | Lepilemur_aeeclis | Lepilemur_aeeclis | nocturnal |
| mammals | Lepilemur_ahmansoni | Lepilemur_ahmansonorum | nocturnal |
| mammals | Lepilemur_ankaranensis | Lepilemur_ankaranensis | nocturnal |
| mammals | Lepilemur_betsileo | Lepilemur_betsileo | nocturnal |
| mammals | Lepilemur_dorsalis | Lepilemur_dorsalis | nocturnal |
| mammals | Lepilemur_edwardsi | Lepilemur_edwardsi | crepuscular |
| mammals | Lepilemur_fleuretae | Lepilemur_fleuretae | nocturnal |
| mammals | Lepilemur_grewcocki | Lepilemur_grewcockorum | nocturnal |
| mammals | Lepilemur_hollandorum | Lepilemur_hollandorum | nocturnal |
| mammals | Lepilemur_hubbardi | Lepilemur_hubbardorum | nocturnal |
| mammals | Lepilemur_jamesi | Lepilemur_jamesorum | nocturnal |
| mammals | Lepilemur_leucopus | Lepilemur_leucopus | nocturnal |
| mammals | Lepilemur_microdon | Lepilemur_microdon | nocturnal |
| mammals | Lepilemur_milanoii | Lepilemur_milanoii | nocturnal |
| mammals | Lepilemur_mustelinus | Lepilemur_mustelinus | nocturnal |
| mammals | Lepilemur_otto | Lepilemur_otto | nocturnal |
| mammals | Lepilemur_petteri | Lepilemur_petteri | nocturnal |
| mammals | Lepilemur_randrianasoloi | Lepilemur_randrianasoloi | nocturnal |
| mammals | Lepilemur_ruficaudatus | Lepilemur_ruficaudatus | nocturnal |
| mammals | Lepilemur_sahamalaza | Lepilemur_sahamalazensis | nocturnal |

| | | | |
|---|---|---|---|
| mammals | Lepilemur_scottorum | Lepilemur_scottorum | nocturnal |
| mammals | Lepilemur_seali | Lepilemur_seali | nocturnal |
| mammals | Lepilemur_septentrionalis | Lepilemur_septentrionalis | nocturnal |
| mammals | Lepilemur_tymerlachsoni | Lepilemur_tymerlachsoni | nocturnal |
| mammals | Lepilemur_wrighti | Lepilemur_wrightae | nocturnal |
| mammals | Leporillus_conditor | Leporillus_conditor | nocturnal |
| mammals | Leptailurus_serval | Leptailurus_serval | crepuscular |
| mammals | Leptomys_elegans | Leptomys_elegans | nocturnal |
| mammals | Leptomys_ernstmayri | Leptomys_ernstmayri | nocturnal |
| mammals | Leptomys_signatus | Leptomys_signatus | nocturnal |
| mammals | Leptonycteris_curasoae | Leptonycteris_curasoae | nocturnal |
| mammals | Leptonycteris_nivalis | Leptonycteris_nivalis | nocturnal |
| mammals | Leptonycteris_yerbabuenae | Leptonycteris_yerbabuenae | nocturnal |
| mammals | Lepus_alleni | Lepus_alleni | nocturnal |
| mammals | Lepus_americanus | Lepus_americanus | nocturnal |
| mammals | Lepus_arcticus | Lepus_arcticus | nocturnal |
| mammals | Lepus_brachyurus | Lepus_brachyurus | nocturnal |
| mammals | Lepus_californicus | Lepus_californicus | nocturnal |
| mammals | Lepus_callotis | Lepus_callotis | nocturnal |
| mammals | Lepus_capensis | Lepus_capensis | nocturnal |
| mammals | Lepus_castroviejoi | Lepus_castroviejoi | nocturnal |
| mammals | Lepus_comus | Lepus_comus | nocturnal |
| mammals | Lepus_coreanus | Lepus_coreanus | nocturnal |
| mammals | Lepus_corsicanus | Lepus_corsicanus | nocturnal |
| mammals | Lepus_europaeus | Lepus_europaeus | cathemeral |
| mammals | Lepus_fagani | Lepus_fagani | nocturnal |
| mammals | Lepus_flavigularis | Lepus_flavigularis | nocturnal |
| mammals | Lepus_granatensis | Lepus_granatensis | nocturnal |
| mammals | Lepus_habessinicus | Lepus_habessinicus | nocturnal |
| mammals | Lepus_hainanus | Lepus_hainanus | nocturnal |
| mammals | Lepus_mandshuricus | Lepus_mandshuricus | nocturnal |
| mammals | Lepus_microtis | Lepus_victoriae | nocturnal |
| mammals | Lepus_nigricollis | Lepus_nigricollis | nocturnal |
| mammals | Lepus_oiostolus | Lepus_oiostolus | nocturnal |

| | | | |
|---|---|---|---|
| mammals | Lepus_othus | Lepus_othus | nocturnal |
| mammals | Lepus_peguensis | Lepus_peguensis | nocturnal |
| mammals | Lepus_saxatilis | Lepus_saxatilis | nocturnal |
| mammals | Lepus_sinensis | Lepus_sinensis | nocturnal |
| mammals | Lepus_starcki | Lepus_starcki | nocturnal |
| mammals | Lepus_tibetanus | Lepus_tibetanus | crepuscular |
| mammals | Lepus_timidus | Lepus_timidus | nocturnal |
| mammals | Lepus_tolai | Lepus_tolai | nocturnal |
| mammals | Lepus_townsendii | Lepus_townsendii | nocturnal |
| mammals | Lepus_yarkandensis | Lepus_yarkandensis | nocturnal |
| mammals | Lestodelphys_halli | Lestodelphys_halli | nocturnal |
| mammals | Lestoros_inca | Lestoros_inca | nocturnal |
| mammals | Leucocephalophus_adersi | Cephalophus_adersi | diurnal |
| mammals | Liberiictis_kuhni | Liberiictis_kuhni | diurnal |
| mammals | Lichonycteris_obscura | Lichonycteris_obscura | nocturnal |
| mammals | Limnomys_bryophilus | Limnomys_bryophilus | nocturnal |
| mammals | Limnomys_sibuanus | Limnomys_sibuanus | nocturnal |
| mammals | Lionycteris_spurrelli | Lionycteris_spurrelli | nocturnal |
| mammals | Litocranius_walleri | Litocranius_walleri | diurnal |
| mammals | Lonchophylla_bokermanni | Lonchophylla_bokermanni | nocturnal |
| mammals | Lonchophylla_chocoana | Lonchophylla_chocoana | nocturnal |
| mammals | Lonchophylla_concava | Lonchophylla_concava | nocturnal |
| mammals | Lonchophylla_dekeyseri | Lonchophylla_dekeyseri | nocturnal |
| mammals | Lonchophylla_handleyi | Lonchophylla_handleyi | nocturnal |
| mammals | Lonchophylla_hesperia | Lonchophylla_hesperia | nocturnal |
| mammals | Lonchophylla_mordax | Lonchophylla_mordax | nocturnal |
| mammals | Lonchophylla_orcesi | Lonchophylla_orcesi | nocturnal |
| mammals | Lonchophylla_robusta | Lonchophylla_robusta | nocturnal |
| mammals | Lonchorhina_aurita | Lonchorhina_aurita | nocturnal |
| mammals | Lonchorhina_fernandezi | Lonchorhina_fernandezi | nocturnal |
| mammals | Lonchorhina_inusitata | Lonchorhina_inusitata | nocturnal |
| mammals | Lonchorhina_marinkellei | Lonchorhina_marinkellei | nocturnal |
| mammals | Lonchorhina_orinocensis | Lonchorhina_orinocensis | nocturnal |
| mammals | Lonchothrix_emiliae | Lonchothrix_emiliae | nocturnal |

| | | | |
|---|---|---|---|
| mammals | Lontra_canadensis | Lontra_canadensis | nocturnal |
| mammals | Lontra_longicaudis | Lontra_longicaudis | diurnal |
| mammals | Lontra_provocax | Lontra_provocax | nocturnal |
| mammals | Lophiomys_imhausi | Lophiomys_imhausi | nocturnal |
| mammals | Lophocebus_albigena | Lophocebus_albigena | diurnal |
| mammals | Lophocebus_aterrimus | Lophocebus_aterrimus | diurnal |
| mammals | Lophostoma_brasiliense | Lophostoma_brasiliense | nocturnal |
| mammals | Lophostoma_carrikeri | Lophostoma_carrikeri | nocturnal |
| mammals | Lophostoma_evotis | Lophostoma_evotis | nocturnal |
| mammals | Lophostoma_kalkoae | Lophostoma_kalkoae | nocturnal |
| mammals | Lophostoma_occidentale | Lophostoma_occidentalis | nocturnal |
| mammals | Lophostoma_schulzi | Lophostoma_schulzi | nocturnal |
| mammals | Lophostoma_silvicola | Lophostoma_silvicolum | nocturnal |
| mammals | Lophuromys_brevicaudus | Lophuromys_brevicaudus | cathemeral |
| mammals | Lophuromys_chrysopus | Lophuromys_chrysopus | cathemeral |
| mammals | Lophuromys_dieterleni | Lophuromys_dieterleni | nocturnal |
| mammals | Lophuromys_eisentrauti | Lophuromys_eisentrauti | nocturnal |
| mammals | Lophuromys_flavopunctatus | Lophuromys_flavopunctatus | nocturnal |
| mammals | Lophuromys_huttereri | Lophuromys_huttereri | nocturnal |
| mammals | Lophuromys_luteogaster | Lophuromys_luteogaster | nocturnal |
| mammals | Lophuromys_medicaudatus | Lophuromys_medicaudatus | nocturnal |
| mammals | Lophuromys_melanonyx | Lophuromys_melanonyx | diurnal |
| mammals | Lophuromys_nudicaudus | Lophuromys_nudicaudus | diurnal |
| mammals | Lophuromys_rahmi | Lophuromys_rahmi | nocturnal |
| mammals | Lophuromys_roseveari | Lophuromys_roseveari | nocturnal |
| mammals | Lophuromys_sikapusi | Lophuromys_sikapusi | nocturnal |
| mammals | Lophuromys_woosnami | Lophuromys_woosnami | nocturnal |
| mammals | Lorentzimys_nouhuysii | Lorentzimys_nouhuysi | cathemeral |
| mammals | Loris_lydekkerianus | Loris_lydekkerianus | nocturnal |
| mammals | Loris_tardigradus | Loris_tardigradus | nocturnal |
| mammals | Loxodonta_africana | Loxodonta_africana | cathemeral |
| mammals | Loxodontomys_micropus | Loxodontomys_micropus | nocturnal |
| mammals | Lundomys_molitor | Lundomys_molitor | nocturnal |
| mammals | Lupulella_adustus | Canis_adustus | nocturnal |

| | | | |
|---|---|---|---|
| mammals | Lupulella_mesomelas | Canis_mesomelas | cathemeral |
| mammals | Lutra_capensis | Aonyx_capensis | nocturnal |
| mammals | Lutra_cinerea | Aonyx_cinereus | diurnal |
| mammals | Lutra_congica | Aonyx_congicus | nocturnal |
| mammals | Lutra_lutra | Lutra_lutra | nocturnal |
| mammals | Lutra_perspicillata | Lutrogale_perspicillata | diurnal |
| mammals | Lutra_sumatrana | Lutra_sumatrana | cathemeral |
| mammals | Lutreolina_crassicaudata | Lutreolina_crassicaudata | nocturnal |
| mammals | Lutreolina_massoia | Lutreolina_massoia | nocturnal |
| mammals | Lycalopex_culpaeus | Lycalopex_culpaeus | cathemeral |
| mammals | Lycalopex_fulvipes | Lycalopex_fulvipes | diurnal |
| mammals | Lycalopex_grisea | Lycalopex_griseus | cathemeral |
| mammals | Lycalopex_gymnocerca | Lycalopex_gymnocercus | cathemeral |
| mammals | Lycalopex_sechurae | Lycalopex_sechurae | nocturnal |
| mammals | Lycalopex_vetula | Lycalopex_vetulus | nocturnal |
| mammals | Lycaon_pictus | Lycaon_pictus | cathemeral |
| mammals | Lyncodon_patagonicus | Lyncodon_patagonicus | nocturnal |
| mammals | Lynx_canadensis | Lynx_canadensis | cathemeral |
| mammals | Lynx_lynx | Lynx_lynx | cathemeral |
| mammals | Lynx_pardinus | Lynx_pardinus | nocturnal |
| mammals | Lynx_rufus | Lynx_rufus | cathemeral |
| mammals | Lyroderma_lyra | Megaderma_lyra | nocturnal |
| mammals | Macaca_arctoides | Macaca_arctoides | diurnal |
| mammals | Macaca_assamensis | Macaca_assamensis | diurnal |
| mammals | Macaca_cyclopis | Macaca_cyclopis | diurnal |
| mammals | Macaca_fascicularis | Macaca_fascicularis | diurnal |
| mammals | Macaca_fuscata | Macaca_fuscata | diurnal |
| mammals | Macaca_hecki | Macaca_hecki | diurnal |
| mammals | Macaca_leonina | Macaca_leonina | diurnal |
| mammals | Macaca_maura | Macaca_maura | diurnal |
| mammals | Macaca_mulatta | Macaca_mulatta | diurnal |
| mammals | Macaca_nemestrina | Macaca_nemestrina | diurnal |
| mammals | Macaca_nigra | Macaca_nigra | diurnal |
| mammals | Macaca_nigrescens | Macaca_nigrescens | diurnal |

| | | | |
|---|---|---|---|
| mammals | Macaca_ochreata | Macaca_ochreata | diurnal |
| mammals | Macaca_pagensis | Macaca_pagensis | diurnal |
| mammals | Macaca_radiata | Macaca_radiata | diurnal |
| mammals | Macaca_siberu | Macaca_siberu | diurnal |
| mammals | Macaca_silenus | Macaca_silenus | diurnal |
| mammals | Macaca_sinica | Macaca_sinica | diurnal |
| mammals | Macaca_sylvanus | Macaca_sylvanus | diurnal |
| mammals | Macaca_thibetana | Macaca_thibetana | diurnal |
| mammals | Macaca_tonkeana | Macaca_tonkeana | diurnal |
| mammals | Macroderma_gigas | Macroderma_gigas | nocturnal |
| mammals | Macrogalidia_musschenbroekii | Macrogalidia_musschenbroekii | nocturnal |
| mammals | Macroglossus_minimus | Macroglossus_minimus | nocturnal |
| mammals | Macroglossus_sobrinus | Macroglossus_sobrinus | nocturnal |
| mammals | Macronycteris_commersonii | Hipposideros_commersoni | nocturnal |
| mammals | Macronycteris_gigas | Hipposideros_gigas | nocturnal |
| mammals | Macronycteris_thomensis | Hipposideros_thomensis | nocturnal |
| mammals | Macronycteris_vittata | Hipposideros_vittatus | nocturnal |
| mammals | Macrophyllum_macrophyllum | Macrophyllum_macrophyllum | nocturnal |
| mammals | Macropus_fuliginosus | Macropus_fuliginosus | nocturnal |
| mammals | Macropus_giganteus | Macropus_giganteus | cathemeral |
| mammals | Macroscelides_flavicaudatus | Macroscelides_flavicaudatus | crepuscular |
| mammals | Macroscelides_micus | Macroscelides_micus | nocturnal |
| mammals | Macroscelides_proboscideus | Macroscelides_proboscideus | crepuscular |
| mammals | Macrotarsomys_bastardi | Macrotarsomys_bastardi | nocturnal |
| mammals | Macrotarsomys_ingens | Macrotarsomys_ingens | nocturnal |
| mammals | Macrotarsomys_petteri | Macrotarsomys_petteri | nocturnal |
| mammals | Macrotis_lagotis | Macrotis_lagotis | nocturnal |
| mammals | Macrotus_californicus | Macrotus_californicus | nocturnal |
| mammals | Macrotus_waterhousii | Macrotus_waterhousii | nocturnal |
| mammals | Macruromys_major | Macruromys_major | nocturnal |
| mammals | Madoqua_guentheri | Madoqua_guentheri | diurnal |
| mammals | Madoqua_kirkii | Madoqua_kirkii | cathemeral |
| mammals | Madoqua_piacentinii | Madoqua_piacentinii | cathemeral |
| mammals | Madoqua_saltiana | Madoqua_saltiana | nocturnal |

| | | | |
|---|---|---|---|
| mammals | Madromys_blanfordi | Madromys_blanfordi | nocturnal |
| mammals | Makalata_didelphoides | Makalata_didelphoides | nocturnal |
| mammals | Makalata_macrura | Makalata_macrura | nocturnal |
| mammals | Malacomys_cansdalei | Malacomys_cansdalei | nocturnal |
| mammals | Malacomys_edwardsi | Malacomys_edwardsi | nocturnal |
| mammals | Malacomys_longipes | Malacomys_longipes | nocturnal |
| mammals | Malacothrix_typica | Malacothrix_typica | nocturnal |
| mammals | Mallomys_aroaensis | Mallomys_aroaensis | nocturnal |
| mammals | Mallomys_gunung | Mallomys_gunung | nocturnal |
| mammals | Mallomys_istapantap | Mallomys_istapantap | nocturnal |
| mammals | Mallomys_rothschildi | Mallomys_rothschildi | nocturnal |
| mammals | Mammelomys_lanosus | Mammelomys_lanosus | nocturnal |
| mammals | Mammelomys_rattoides | Mammelomys_rattoides | nocturnal |
| mammals | Mandrillus_leucophaeus | Mandrillus_leucophaeus | diurnal |
| mammals | Mandrillus_sphinx | Mandrillus_sphinx | diurnal |
| mammals | Manis_crassicaudata | Manis_crassicaudata | nocturnal |
| mammals | Manis_culionensis | Manis_culionensis | nocturnal |
| mammals | Manis_javanica | Manis_javanica | nocturnal |
| mammals | Manis_pentadactyla | Manis_pentadactyla | nocturnal |
| mammals | Margaretamys_beccarii | Margaretamys_beccarii | nocturnal |
| mammals | Marmosa_alstoni | Marmosa_alstoni | nocturnal |
| mammals | Marmosa_andersoni | Marmosa_andersoni | nocturnal |
| mammals | Marmosa_demerarae | Marmosa_demerarae | nocturnal |
| mammals | Marmosa_lepida | Marmosa_lepida | nocturnal |
| mammals | Marmosa_mexicana | Marmosa_mexicana | nocturnal |
| mammals | Marmosa_murina | Marmosa_murina | nocturnal |
| mammals | Marmosa_paraguayana | Marmosa_paraguayanus | nocturnal |
| mammals | Marmosa_phaea | Marmosa_phaea | nocturnal |
| mammals | Marmosa_robinsoni | Marmosa_robinsoni | nocturnal |
| mammals | Marmosa_rubra | Marmosa_rubra | nocturnal |
| mammals | Marmosa_tyleriana | Marmosa_tyleriana | nocturnal |
| mammals | Marmosa_xerophila | Marmosa_xerophila | nocturnal |
| mammals | Marmosops_caucae | Marmosops_neblina | nocturnal |
| mammals | Marmosops_creightoni | Marmosops_creightoni | nocturnal |

| | | | |
|---|---|---|---|
| mammals | Marmosops_fuscatus | Marmosops_fuscatus | nocturnal |
| mammals | Marmosops_handleyi | Marmosops_handleyi | nocturnal |
| mammals | Marmosops_incanus | Marmosops_incanus | nocturnal |
| mammals | Marmosops_invictus | Marmosops_invictus | nocturnal |
| mammals | Marmosops_noctivagus | Marmosops_noctivagus | nocturnal |
| mammals | Marmosops_ocellatus | Marmosops_ocellatus | nocturnal |
| mammals | Marmosops_parvidens | Marmosops_parvidens | nocturnal |
| mammals | Marmosops_paulensis | Marmosops_paulensis | nocturnal |
| mammals | Marmosops_pinheiroi | Marmosops_pinheiroi | nocturnal |
| mammals | Marmota_baibacina | Marmota_baibacina | diurnal |
| mammals | Marmota_bobak | Marmota_bobak | diurnal |
| mammals | Marmota_broweri | Marmota_broweri | diurnal |
| mammals | Marmota_caligata | Marmota_caligata | diurnal |
| mammals | Marmota_camtschatica | Marmota_camtschatica | diurnal |
| mammals | Marmota_caudata | Marmota_caudata | diurnal |
| mammals | Marmota_flaviventris | Marmota_flaviventris | diurnal |
| mammals | Marmota_himalayana | Marmota_himalayana | diurnal |
| mammals | Marmota_marmota | Marmota_marmota | diurnal |
| mammals | Marmota_menzbieri | Marmota_menzbieri | diurnal |
| mammals | Marmota_monax | Marmota_monax | diurnal |
| mammals | Marmota_olympus | Marmota_olympus | diurnal |
| mammals | Marmota_sibirica | Marmota_sibirica | diurnal |
| mammals | Marmota_vancouverensis | Marmota_vancouverensis | diurnal |
| mammals | Martes_americana | Martes_americana | cathemeral |
| mammals | Martes_flavigula | Martes_flavigula | cathemeral |
| mammals | Martes_foina | Martes_foina | cathemeral |
| mammals | Martes_gwatkinsii | Martes_gwatkinsii | diurnal |
| mammals | Martes_martes | Martes_martes | cathemeral |
| mammals | Martes_melampus | Martes_melampus | cathemeral |
| mammals | Martes_zibellina | Martes_zibellina | cathemeral |
| mammals | Massoutiera_mzabi | Massoutiera_mzabi | diurnal |
| mammals | Mastacomys_fuscus | Mastacomys_fuscus | nocturnal |
| mammals | Mastomys_awashensis | Mastomys_awashensis | nocturnal |
| mammals | Mastomys_coucha | Mastomys_coucha | nocturnal |

| | | | |
|---|---|---|---|
| mammals | Mastomys_erythroleucus | Mastomys_erythroleucus | nocturnal |
| mammals | Mastomys_huberti | Mastomys_huberti | nocturnal |
| mammals | Mastomys_kollmannspergeri | Mastomys_kollmannspergeri | nocturnal |
| mammals | Mastomys_natalensis | Mastomys_natalensis | nocturnal |
| mammals | Mastomys_shortridgei | Mastomys_shortridgei | nocturnal |
| mammals | Maxomys_alticola | Maxomys_alticola | nocturnal |
| mammals | Maxomys_baeodon | Maxomys_baeodon | nocturnal |
| mammals | Maxomys_bartelsii | Maxomys_bartelsii | nocturnal |
| mammals | Maxomys_dollmani | Maxomys_dollmani | nocturnal |
| mammals | Maxomys_hellwaldii | Maxomys_hellwaldii | nocturnal |
| mammals | Maxomys_hylomyoides | Maxomys_hylomyoides | nocturnal |
| mammals | Maxomys_inas | Maxomys_inas | nocturnal |
| mammals | Maxomys_inflatus | Maxomys_inflatus | nocturnal |
| mammals | Maxomys_moi | Maxomys_moi | nocturnal |
| mammals | Maxomys_musschenbroekii | Maxomys_musschenbroekii | nocturnal |
| mammals | Maxomys_ochraceiventer | Maxomys_ochraceiventer | nocturnal |
| mammals | Maxomys_pagensis | Maxomys_pagensis | nocturnal |
| mammals | Maxomys_panglima | Maxomys_panglima | nocturnal |
| mammals | Maxomys_rajah | Maxomys_rajah | nocturnal |
| mammals | Maxomys_surifer | Maxomys_surifer | nocturnal |
| mammals | Maxomys_tajuddinii | Maxomys_tajuddinii | nocturnal |
| mammals | Maxomys_wattsi | Maxomys_wattsi | nocturnal |
| mammals | Maxomys_whiteheadi | Maxomys_whiteheadi | nocturnal |
| mammals | Mazama_americana | Mazama_americana | cathemeral |
| mammals | Mazama_chunyi | Mazama_chunyi | crepuscular |
| mammals | Mazama_jucunda | Mazama_bororo | nocturnal |
| mammals | Mazama_nanus | Mazama_nana | cathemeral |
| mammals | Mazama_rufina | Mazama_rufina | cathemeral |
| mammals | Mazama_temama | Mazama_temama | cathemeral |
| mammals | Megadendromus_nikolausi | Megadendromus_nikolausi | nocturnal |
| mammals | Megaderma_spasma | Megaderma_spasma | nocturnal |
| mammals | Megadontomys_cryophilus | Megadontomys_cryophilus | nocturnal |
| mammals | Megadontomys_nelsoni | Megadontomys_nelsoni | nocturnal |
| mammals | Megadontomys_thomasi | Megadontomys_thomasi | nocturnal |

| | | | |
|---|---|---|---|
| mammals | Megaerops_ecaudatus | Megaerops_ecaudatus | nocturnal |
| mammals | Megaerops_kusnotoi | Megaerops_kusnotoi | nocturnal |
| mammals | Megaerops_niphanae | Megaerops_niphanae | nocturnal |
| mammals | Megaloglossus_azagnyi | Megaloglossus_azagnyi | nocturnal |
| mammals | Megaloglossus_woermanni | Megaloglossus_woermanni | nocturnal |
| mammals | Megasorex_gigas | Megasorex_gigas | cathemeral |
| mammals | Melanomys_caliginosus | Melanomys_caliginosus | diurnal |
| mammals | Melanomys_robustulus | Melanomys_robustulus | nocturnal |
| mammals | Melanomys_zunigae | Melanomys_zunigae | nocturnal |
| mammals | Melasmothrix_naso | Melasmothrix_naso | diurnal |
| mammals | Meles_anakuma | Meles_anakuma | nocturnal |
| mammals | Meles_leucurus | Meles_leucurus | nocturnal |
| mammals | Meles_meles | Meles_meles | nocturnal |
| mammals | Mellivora_capensis | Mellivora_capensis | nocturnal |
| mammals | Melogale_everetti | Melogale_everetti | nocturnal |
| mammals | Melogale_moschata | Melogale_moschata | nocturnal |
| mammals | Melogale_orientalis | Melogale_orientalis | nocturnal |
| mammals | Melogale_personata | Melogale_personata | nocturnal |
| mammals | Melomys_aerosus | Melomys_aerosus | nocturnal |
| mammals | Melomys_arcium | Melomys_arcium | nocturnal |
| mammals | Melomys_bannisteri | Melomys_bannisteri | nocturnal |
| mammals | Melomys_bougainville | Melomys_bougainville | nocturnal |
| mammals | Melomys_burtoni | Melomys_burtoni | nocturnal |
| mammals | Melomys_capensis | Melomys_capensis | nocturnal |
| mammals | Melomys_caurinus | Melomys_caurinus | nocturnal |
| mammals | Melomys_cervinipes | Melomys_cervinipes | nocturnal |
| mammals | Melomys_cooperae | Melomys_cooperae | nocturnal |
| mammals | Melomys_dollmani | Melomys_dollmani | nocturnal |
| mammals | Melomys_fraterculus | Melomys_fraterculus | nocturnal |
| mammals | Melomys_frigicola | Melomys_frigicola | nocturnal |
| mammals | Melomys_fulgens | Melomys_fulgens | nocturnal |
| mammals | Melomys_howi | Melomys_howi | diurnal |
| mammals | Melomys_leucogaster | Melomys_leucogaster | nocturnal |
| mammals | Melomys_lutillus | Melomys_lutillus | nocturnal |

| | | | |
|---|---|---|---|
| mammals | Melomys_matambuai | Melomys_matambuai | nocturnal |
| mammals | Melomys_obiensis | Melomys_obiensis | nocturnal |
| mammals | Melomys_paveli | Melomys_paveli | nocturnal |
| mammals | Melomys_rufescens | Melomys_rufescens | nocturnal |
| mammals | Melomys_talaudium | Melomys_talaudium | nocturnal |
| mammals | Melonycteris_melanops | Melonycteris_melanops | nocturnal |
| mammals | Melursus_ursinus | Melursus_ursinus | cathemeral |
| mammals | Menetes_berdmorei | Menetes_berdmorei | diurnal |
| mammals | Mephitis_macroura | Mephitis_macroura | nocturnal |
| mammals | Mephitis_mephitis | Mephitis_mephitis | nocturnal |
| mammals | Meriones_arimalius | Meriones_arimalius | diurnal |
| mammals | Meriones_crassus | Meriones_crassus | nocturnal |
| mammals | Meriones_grandis | Meriones_grandis | nocturnal |
| mammals | Meriones_hurrianae | Meriones_hurrianae | diurnal |
| mammals | Meriones_libycus | Meriones_libycus | nocturnal |
| mammals | Meriones_meridianus | Meriones_meridianus | nocturnal |
| mammals | Meriones_persicus | Meriones_persicus | nocturnal |
| mammals | Meriones_psammophilus | Meriones_chengi | nocturnal |
| mammals | Meriones_rex | Meriones_rex | nocturnal |
| mammals | Meriones_sacramenti | Meriones_sacramenti | nocturnal |
| mammals | Meriones_shawii | Meriones_shawi | nocturnal |
| mammals | Meriones_tamariscinus | Meriones_tamariscinus | nocturnal |
| mammals | Meriones_tristrami | Meriones_tristrami | nocturnal |
| mammals | Meriones_unguiculatus | Meriones_unguiculatus | nocturnal |
| mammals | Meriones_vinogradovi | Meriones_vinogradovi | cathemeral |
| mammals | Meriones_zarudnyi | Meriones_zarudnyi | nocturnal |
| mammals | Mesechinus_dauuricus | Mesechinus_dauuricus | nocturnal |
| mammals | Mesechinus_hughi | Mesechinus_hughi | nocturnal |
| mammals | Mesembriomys_gouldii | Mesembriomys_gouldii | nocturnal |
| mammals | Mesembriomys_macrurus | Mesembriomys_macrurus | nocturnal |
| mammals | Mesocapromys_auritus | Mesocapromys_auritus | nocturnal |
| mammals | Mesocapromys_melanurus | Mysateles_melanurus | nocturnal |
| mammals | Mesocricetus_auratus | Mesocricetus_auratus | nocturnal |
| mammals | Mesocricetus_brandti | Mesocricetus_brandti | nocturnal |

| | | | |
|---|---|---|---|
| mammals | Mesocricetus_newtoni | Mesocricetus_newtoni | nocturnal |
| mammals | Mesocricetus_raddei | Mesocricetus_raddei | nocturnal |
| mammals | Mesomys_hispidus | Mesomys_hispidus | nocturnal |
| mammals | Mesomys_leniceps | Mesomys_leniceps | nocturnal |
| mammals | Mesomys_occultus | Mesomys_occultus | nocturnal |
| mammals | Mesomys_stimulax | Mesomys_stimulax | nocturnal |
| mammals | Mesophylla_macconnelli | Mesophylla_macconnelli | nocturnal |
| mammals | Metachirus_nudicaudatus | Metachirus_nudicaudatus | nocturnal |
| mammals | Micaelamys_granti | Aethomys_granti | nocturnal |
| mammals | Micaelamys_namaquensis | Aethomys_namaquensis | nocturnal |
| mammals | Mico_acariensis | Mico_acariensis | diurnal |
| mammals | Mico_argentatus | Mico_argentatus | diurnal |
| mammals | Mico_chrysoleucos | Mico_chrysoleucos | diurnal |
| mammals | Mico_emiliae | Mico_emiliae | diurnal |
| mammals | Mico_humeralifer | Mico_humeralifer | diurnal |
| mammals | Mico_humilis | Mico_humilis | diurnal |
| mammals | Mico_intermedius | Mico_intermedius | diurnal |
| mammals | Mico_leucippe | Mico_leucippe | diurnal |
| mammals | Mico_marcai | Mico_marcai | diurnal |
| mammals | Mico_mauesi | Mico_mauesi | diurnal |
| mammals | Mico_melanurus | Mico_melanurus | diurnal |
| mammals | Mico_nigriceps | Mico_nigriceps | diurnal |
| mammals | Mico_rondoni | Mico_rondoni | diurnal |
| mammals | Mico_saterei | Mico_saterei | diurnal |
| mammals | Microcavia_australis | Microcavia_australis | diurnal |
| mammals | Microcavia_niata | Microcavia_niata | diurnal |
| mammals | Microcavia_shiptoni | Microcavia_shiptoni | diurnal |
| mammals | Microcebus_arnholdi | Microcebus_arnholdi | nocturnal |
| mammals | Microcebus_berthae | Microcebus_berthae | nocturnal |
| mammals | Microcebus_bongolavensis | Microcebus_bongolavensis | nocturnal |
| mammals | Microcebus_danfossi | Microcebus_danfossi | nocturnal |
| mammals | Microcebus_gerpi | Microcebus_gerpi | nocturnal |
| mammals | Microcebus_griseorufus | Microcebus_griseorufus | nocturnal |
| mammals | Microcebus_jollyae | Microcebus_jollyae | nocturnal |

| | | | |
|---|---|---|---|
| mammals | Microcebus_lehilahytsara | Microcebus_lehilahytsara | nocturnal |
| mammals | Microcebus_macarthurii | Microcebus_macarthurii | nocturnal |
| mammals | Microcebus_mamiratra | Microcebus_mamiratra | nocturnal |
| mammals | Microcebus_margotmarshae | Microcebus_margotmarshae | nocturnal |
| mammals | Microcebus_marohita | Microcebus_marohita | nocturnal |
| mammals | Microcebus_murinus | Microcebus_murinus | nocturnal |
| mammals | Microcebus_myoxinus | Microcebus_myoxinus | nocturnal |
| mammals | Microcebus_ravelobensis | Microcebus_ravelobensis | nocturnal |
| mammals | Microcebus_rufus | Microcebus_rufus | nocturnal |
| mammals | Microcebus_sambiranensis | Microcebus_sambiranensis | nocturnal |
| mammals | Microcebus_simmonsi | Microcebus_simmonsi | nocturnal |
| mammals | Microcebus_tavaratra | Microcebus_tavaratra | nocturnal |
| mammals | Microdillus_peeli | Microdillus_peeli | nocturnal |
| mammals | Microdipodops_megacephalus | Microdipodops_megacephalus | nocturnal |
| mammals | Microdipodops_pallidus | Microdipodops_pallidus | nocturnal |
| mammals | Microgale_cowani | Microgale_cowani | nocturnal |
| mammals | Microgale_drouhardi | Microgale_drouhardi | nocturnal |
| mammals | Microgale_dryas | Microgale_dryas | nocturnal |
| mammals | Microgale_fotsifotsy | Microgale_fotsifotsy | nocturnal |
| mammals | Microgale_gracilis | Microgale_gracilis | nocturnal |
| mammals | Microgale_gymnorhyncha | Microgale_gymnorhyncha | nocturnal |
| mammals | Microgale_jenkinsae | Microgale_jenkinsae | nocturnal |
| mammals | Microgale_jobihely | Microgale_jobihely | nocturnal |
| mammals | Microgale_longicaudata | Microgale_longicaudata | nocturnal |
| mammals | Microgale_majori | Microgale_majori | nocturnal |
| mammals | Microgale_mergulus | Limnogale_mergulus | nocturnal |
| mammals | Microgale_monticola | Microgale_monticola | nocturnal |
| mammals | Microgale_nasoloi | Microgale_nasoloi | nocturnal |
| mammals | Microgale_parvula | Microgale_parvula | nocturnal |
| mammals | Microgale_principula | Microgale_principula | nocturnal |
| mammals | Microgale_pusilla | Microgale_pusilla | nocturnal |
| mammals | Microgale_soricoides | Microgale_soricoides | nocturnal |
| mammals | Microgale_taiva | Microgale_taiva | nocturnal |
| mammals | Microgale_thomasi | Microgale_thomasi | nocturnal |

| | | | |
|---|---|---|---|
| mammals | Microhydromys_argenteus | Microhydromys_argenteus | nocturnal |
| mammals | Microhydromys_richardsoni | Microhydromys_richardsoni | nocturnal |
| mammals | Micromys_minutus | Micromys_minutus | nocturnal |
| mammals | Micronomus_norfolkensis | Mormopterus_norfolkensis | nocturnal |
| mammals | Micronycteris_brosseti | Micronycteris_brosseti | nocturnal |
| mammals | Micronycteris_giovanniae | Micronycteris_giovanniae | nocturnal |
| mammals | Micronycteris_hirsuta | Micronycteris_hirsuta | nocturnal |
| mammals | Micronycteris_matses | Micronycteris_matses | nocturnal |
| mammals | Micronycteris_megalotis | Micronycteris_megalotis | nocturnal |
| mammals | Micronycteris_microtis | Micronycteris_microtis | nocturnal |
| mammals | Micronycteris_minuta | Micronycteris_minuta | nocturnal |
| mammals | Micronycteris_sanborni | Micronycteris_sanborni | nocturnal |
| mammals | Micronycteris_schmidtorum | Micronycteris_schmidtorum | nocturnal |
| mammals | Microperoryctes_aplini | Microperoryctes_aplini | nocturnal |
| mammals | Microperoryctes_longicauda | Microperoryctes_longicauda | nocturnal |
| mammals | Microperoryctes_murinus | Microperoryctes_murina | nocturnal |
| mammals | Microperoryctes_papuensis | Microperoryctes_papuensis | nocturnal |
| mammals | Micropotamogale_lamottei | Micropotamogale_lamottei | nocturnal |
| mammals | Micropotamogale_ruwenzorii | Micropotamogale_ruwenzorii | nocturnal |
| mammals | Microryzomys_minutus | Microryzomys_minutus | nocturnal |
| mammals | Microsciurus_alfari | Microsciurus_alfari | diurnal |
| mammals | Microsciurus_flaviventer | Microsciurus_flaviventer | diurnal |
| mammals | Microsciurus_mimulus | Microsciurus_mimulus | diurnal |
| mammals | Microsciurus_santanderensis | Microsciurus_santanderensis | diurnal |
| mammals | Microtus_abbreviatus | Microtus_abbreviatus | cathemeral |
| mammals | Microtus_afghanus | Blanfordimys_afghanus | diurnal |
| mammals | Microtus_agrestis | Microtus_agrestis | cathemeral |
| mammals | Microtus_anatolicus | Microtus_anatolicus | cathemeral |
| mammals | Microtus_arvalis | Microtus_arvalis | cathemeral |
| mammals | Microtus_brachycercus | Microtus_brachycercus | nocturnal |
| mammals | Microtus_bucharensis | Blanfordimys_bucharensis | cathemeral |
| mammals | Microtus_cabrerae | Microtus_cabrerae | diurnal |
| mammals | Microtus_canicaudus | Microtus_canicaudus | cathemeral |
| mammals | Microtus_chrotorrhinus | Microtus_chrotorrhinus | diurnal |

| | | | |
|---|---|---|---|
| mammals | Microtus_felteni | Microtus_felteni | nocturnal |
| mammals | Microtus_guatemalensis | Microtus_guatemalensis | diurnal |
| mammals | Microtus_ilaeus | Microtus_ilaeus | crepuscular |
| mammals | Microtus_irani | Microtus_irani | cathemeral |
| mammals | Microtus_longicaudus | Microtus_longicaudus | cathemeral |
| mammals | Microtus_majori | Microtus_majori | nocturnal |
| mammals | Microtus_mexicanus | Microtus_mexicanus | diurnal |
| mammals | Microtus_montanus | Microtus_montanus | cathemeral |
| mammals | Microtus_oaxacensis | Microtus_oaxacensis | cathemeral |
| mammals | Microtus_ochrogaster | Microtus_ochrogaster | cathemeral |
| mammals | Microtus_oregoni | Microtus_oregoni | nocturnal |
| mammals | Microtus_pennsylvanicus | Microtus_pennsylvanicus | cathemeral |
| mammals | Microtus_quasiater | Microtus_quasiater | cathemeral |
| mammals | Microtus_richardsoni | Microtus_richardsoni | nocturnal |
| mammals | Microtus_schelkovnikovi | Microtus_schelkovnikovi | cathemeral |
| mammals | Microtus_townsendii | Microtus_townsendii | cathemeral |
| mammals | Microtus_transcaspicus | Microtus_transcaspicus | cathemeral |
| mammals | Microtus_xanthognathus | Microtus_xanthognathus | cathemeral |
| mammals | Microtus_yuldaschi | Neodon_juldaschi | cathemeral |
| mammals | Mictomys_borealis | Synaptomys_borealis | cathemeral |
| mammals | Mimetillus_moloneyi | Mimetillus_moloneyi | nocturnal |
| mammals | Mimon_bennettii | Mimon_bennettii | nocturnal |
| mammals | Mimon_cozumelae | Mimon_cozumelae | nocturnal |
| mammals | Mindomys_hammondi | Mindomys_hammondi | nocturnal |
| mammals | Miniopterus_australis | Miniopterus_australis | nocturnal |
| mammals | Miniopterus_fraterculus | Miniopterus_fraterculus | nocturnal |
| mammals | Miniopterus_fuscus | Miniopterus_fuscus | nocturnal |
| mammals | Miniopterus_gleni | Miniopterus_gleni | nocturnal |
| mammals | Miniopterus_griveaudi | Miniopterus_griveaudi | nocturnal |
| mammals | Miniopterus_inflatus | Miniopterus_inflatus | nocturnal |
| mammals | Miniopterus_macrocneme | Miniopterus_macrocneme | nocturnal |
| mammals | Miniopterus_magnater | Miniopterus_magnater | nocturnal |
| mammals | Miniopterus_majori | Miniopterus_majori | nocturnal |
| mammals | Miniopterus_manavi | Miniopterus_manavi | nocturnal |

| | | | |
|---|---|---|---|
| mammals | Miniopterus_medius | Miniopterus_medius | nocturnal |
| mammals | Miniopterus_minor | Miniopterus_minor | nocturnal |
| mammals | Miniopterus_natalensis | Miniopterus_natalensis | nocturnal |
| mammals | Miniopterus_newtoni | Miniopterus_newtoni | nocturnal |
| mammals | Miniopterus_paululus | Miniopterus_paululus | nocturnal |
| mammals | Miniopterus_petersoni | Miniopterus_petersoni | nocturnal |
| mammals | Miniopterus_pusillus | Miniopterus_pusillus | nocturnal |
| mammals | Miniopterus_robustior | Miniopterus_robustior | nocturnal |
| mammals | Miniopterus_schreibersii | Miniopterus_schreibersii | nocturnal |
| mammals | Miniopterus_shortridgei | Miniopterus_shortridgei | nocturnal |
| mammals | Miniopterus_sororculus | Miniopterus_sororculus | nocturnal |
| mammals | Miniopterus_tristis | Miniopterus_tristis | nocturnal |
| mammals | Miopithecus_ogouensis | Miopithecus_ogouensis | diurnal |
| mammals | Miopithecus_talapoin | Miopithecus_talapoin | diurnal |
| mammals | Mirimiri_acrodonta | Mirimiri_acrodonta | nocturnal |
| mammals | Mirostrellus_joffrei | Hypsugo_joffrei | nocturnal |
| mammals | Mirza_coquereli | Mirza_coquereli | nocturnal |
| mammals | Mirza_zaza | Mirza_zaza | nocturnal |
| mammals | Molossops_neglectus | Molossops_neglectus | nocturnal |
| mammals | Molossops_temminckii | Molossops_temminckii | nocturnal |
| mammals | Molossus_alvarezi | Molossus_alvarezi | crepuscular |
| mammals | Molossus_aztecus | Molossus_aztecus | nocturnal |
| mammals | Molossus_coibensis | Molossus_coibensis | nocturnal |
| mammals | Molossus_currentium | Molossus_currentium | nocturnal |
| mammals | Molossus_molossus | Molossus_molossus | nocturnal |
| mammals | Molossus_pretiosus | Molossus_pretiosus | nocturnal |
| mammals | Molossus_rufus | Molossus_rufus | nocturnal |
| mammals | Molossus_sinaloae | Molossus_sinaloae | nocturnal |
| mammals | Monodelphis_adusta | Monodelphis_adusta | nocturnal |
| mammals | Monodelphis_americana | Monodelphis_americana | diurnal |
| mammals | Monodelphis_brevicaudata | Monodelphis_brevicaudata | nocturnal |
| mammals | Monodelphis_dimidiata | Monodelphis_dimidiata | diurnal |
| mammals | Monodelphis_domestica | Monodelphis_domestica | nocturnal |
| mammals | Monodelphis_emiliae | Monodelphis_emiliae | nocturnal |

| | | | |
|---|---|---|---|
| mammals | Monodelphis_iheringi | Monodelphis_iheringi | nocturnal |
| mammals | Monodelphis_kunsi | Monodelphis_kunsi | diurnal |
| mammals | Monodelphis_osgoodi | Monodelphis_osgoodi | nocturnal |
| mammals | Monodelphis_palliolata | Monodelphis_palliolata | crepuscular |
| mammals | Monodelphis_reigi | Monodelphis_reigi | nocturnal |
| mammals | Monodelphis_ronaldi | Monodelphis_ronaldi | nocturnal |
| mammals | Monodelphis_scalops | Monodelphis_scalops | diurnal |
| mammals | Monophyllus_plethodon | Monophyllus_plethodon | nocturnal |
| mammals | Monophyllus_redmani | Monophyllus_redmani | nocturnal |
| mammals | Montemys_delectorum | Praomys_delectorum | nocturnal |
| mammals | Monticolomys_koopmani | Monticolomys_koopmani | nocturnal |
| mammals | Mops_aloysiisabaudiae | Chaerephon_aloysiisabaudiae | nocturnal |
| mammals | Mops_ansorgei | Chaerephon_ansorgei | nocturnal |
| mammals | Mops_bemmeleni | Chaerephon_bemmeleni | nocturnal |
| mammals | Mops_bivittatus | Chaerephon_bivittatus | nocturnal |
| mammals | Mops_brachypterus | Mops_brachypterus | nocturnal |
| mammals | Mops_bregullae | Chaerephon_bregullae | nocturnal |
| mammals | Mops_chapini | Chaerephon_chapini | nocturnal |
| mammals | Mops_condylurus | Mops_condylurus | nocturnal |
| mammals | Mops_congicus | Mops_congicus | nocturnal |
| mammals | Mops_demonstrator | Mops_demonstrator | nocturnal |
| mammals | Mops_gallagheri | Chaerephon_gallagheri | nocturnal |
| mammals | Mops_jobensis | Chaerephon_jobensis | nocturnal |
| mammals | Mops_jobimena | Chaerephon_jobimena | nocturnal |
| mammals | Mops_johorensis | Chaerephon_johorensis | nocturnal |
| mammals | Mops_leucostigma | Mops_leucostigma | nocturnal |
| mammals | Mops_major | Chaerephon_major | nocturnal |
| mammals | Mops_midas | Mops_midas | nocturnal |
| mammals | Mops_mops | Mops_mops | nocturnal |
| mammals | Mops_nanulus | Mops_nanulus | nocturnal |
| mammals | Mops_niangarae | Mops_niangarae | nocturnal |
| mammals | Mops_nigeriae | Chaerephon_nigeriae | nocturnal |
| mammals | Mops_niveiventer | Mops_niveiventer | nocturnal |
| mammals | Mops_petersoni | Mops_petersoni | nocturnal |

| | | | |
|---|---|---|---|
| mammals | Mops_plicatus | Chaerephon_plicatus | nocturnal |
| mammals | Mops_pumilus | Chaerephon_pumilus | nocturnal |
| mammals | Mops_russatus | Chaerephon_russatus | nocturnal |
| mammals | Mops_sarasinorum | Mops_sarasinorum | nocturnal |
| mammals | Mops_solomonis | Chaerephon_solomonis | nocturnal |
| mammals | Mops_spurrelli | Mops_spurrelli | nocturnal |
| mammals | Mops_thersites | Mops_thersites | nocturnal |
| mammals | Mops_tomensis | Chaerephon_tomensis | nocturnal |
| mammals | Mops_trevori | Mops_trevori | nocturnal |
| mammals | Mormoops_blainvillei | Mormoops_blainvillei | nocturnal |
| mammals | Mormoops_megalophylla | Mormoops_megalophylla | nocturnal |
| mammals | Mormopterus_acetabulosus | Mormopterus_acetabulosus | nocturnal |
| mammals | Mormopterus_doriae | Mormopterus_doriae | nocturnal |
| mammals | Mormopterus_jugularis | Mormopterus_jugularis | nocturnal |
| mammals | Mormopterus_kalinowskii | Mormopterus_kalinowskii | nocturnal |
| mammals | Mormopterus_minutus | Mormopterus_minutus | nocturnal |
| mammals | Mormopterus_phrudus | Mormopterus_phrudus | nocturnal |
| mammals | Moschiola_indica | Moschiola_indica | nocturnal |
| mammals | Moschiola_meminna | Moschiola_meminna | nocturnal |
| mammals | Moschus_anhuiensis | Moschus_anhuiensis | nocturnal |
| mammals | Moschus_berezovskii | Moschus_berezovskii | nocturnal |
| mammals | Moschus_chrysogaster | Moschus_chrysogaster | nocturnal |
| mammals | Moschus_cupreus | Moschus_cupreus | nocturnal |
| mammals | Moschus_fuscus | Moschus_fuscus | nocturnal |
| mammals | Moschus_leucogaster | Moschus_leucogaster | nocturnal |
| mammals | Moschus_moschiferus | Moschus_moschiferus | crepuscular |
| mammals | Mosia_nigrescens | Mosia_nigrescens | nocturnal |
| mammals | Mungos_gambianus | Mungos_gambianus | diurnal |
| mammals | Mungos_mungo | Mungos_mungo | diurnal |
| mammals | Mungotictis_decemlineata | Mungotictis_decemlineata | diurnal |
| mammals | Muntiacus_atherodes | Muntiacus_atherodes | diurnal |
| mammals | Muntiacus_crinifrons | Muntiacus_crinifrons | cathemeral |
| mammals | Muntiacus_feae | Muntiacus_feae | cathemeral |
| mammals | Muntiacus_gongshanensis | Muntiacus_gongshanensis | crepuscular |

| | | | |
|---|---|---|---|
| mammals | Muntiacus_muntjak | Muntiacus_muntjak | cathemeral |
| mammals | Muntiacus_putaoensis | Muntiacus_putaoensis | cathemeral |
| mammals | Muntiacus_reevesi | Muntiacus_reevesi | cathemeral |
| mammals | Muntiacus_rooseveltorum | Muntiacus_rooseveltorum | cathemeral |
| mammals | Muntiacus_truongsonensis | Muntiacus_truongsonensis | cathemeral |
| mammals | Muntiacus_vaginalis | Muntiacus_vaginalis | cathemeral |
| mammals | Muntiacus_vuquangensis | Muntiacus_vuquangensis | diurnal |
| mammals | Murexia_habbema | Murexia_habbema | nocturnal |
| mammals | Murexia_longicaudata | Murexia_longicaudata | cathemeral |
| mammals | Murexia_melanura | Murexia_melanurus | nocturnal |
| mammals | Murexia_naso | Murexia_naso | crepuscular |
| mammals | Murexia_rothschildi | Murexia_rothschildi | nocturnal |
| mammals | Murina_aenea | Murina_aenea | nocturnal |
| mammals | Murina_aurata | Murina_aurata | nocturnal |
| mammals | Murina_cyclotis | Murina_cyclotis | nocturnal |
| mammals | Murina_florium | Murina_florium | nocturnal |
| mammals | Murina_fusca | Murina_fusca | nocturnal |
| mammals | Murina_harrisoni | Murina_harrisoni | nocturnal |
| mammals | Murina_hilgendorfi | Murina_hilgendorfi | nocturnal |
| mammals | Murina_huttonii | Murina_huttoni | nocturnal |
| mammals | Murina_leucogaster | Murina_leucogaster | nocturnal |
| mammals | Murina_puta | Murina_puta | nocturnal |
| mammals | Murina_rozendaali | Murina_rozendaali | nocturnal |
| mammals | Murina_ryukyuana | Murina_ryukyuana | nocturnal |
| mammals | Murina_suilla | Murina_suilla | nocturnal |
| mammals | Murina_tubinaris | Murina_tubinaris | nocturnal |
| mammals | Murina_ussuriensis | Murina_ussuriensis | nocturnal |
| mammals | Mus_baoulei | Mus_baoulei | nocturnal |
| mammals | Mus_booduga | Mus_booduga | nocturnal |
| mammals | Mus_bufo | Mus_bufo | nocturnal |
| mammals | Mus_callewaerti | Mus_callewaerti | nocturnal |
| mammals | Mus_caroli | Mus_caroli | nocturnal |
| mammals | Mus_cervicolor | Mus_cervicolor | nocturnal |
| mammals | Mus_cookii | Mus_cookii | nocturnal |

| | | | |
|---|---|---|---|
| mammals | Mus_crociduroides | Mus_crociduroides | nocturnal |
| mammals | Mus_cypriacus | Mus_cypriacus | nocturnal |
| mammals | Mus_famulus | Mus_famulus | nocturnal |
| mammals | Mus_fragilicauda | Mus_fragilicauda | nocturnal |
| mammals | Mus_goundae | Mus_goundae | nocturnal |
| mammals | Mus_haussa | Mus_haussa | nocturnal |
| mammals | Mus_imberbis | Muriculus_imberbis | nocturnal |
| mammals | Mus_indutus | Mus_indutus | nocturnal |
| mammals | Mus_macedonicus | Mus_macedonicus | nocturnal |
| mammals | Mus_mahomet | Mus_mahomet | nocturnal |
| mammals | Mus_mattheyi | Mus_mattheyi | nocturnal |
| mammals | Mus_mayori | Mus_mayori | nocturnal |
| mammals | Mus_minutoides | Mus_minutoides | nocturnal |
| mammals | Mus_musculoides | Mus_musculoides | nocturnal |
| mammals | Mus_musculus | Mus_musculus | nocturnal |
| mammals | Mus_neavei | Mus_neavei | nocturnal |
| mammals | Mus_oubanguii | Mus_oubanguii | nocturnal |
| mammals | Mus_pahari | Mus_pahari | nocturnal |
| mammals | Mus_platythrix | Mus_platythrix | nocturnal |
| mammals | Mus_setulosus | Mus_setulosus | nocturnal |
| mammals | Mus_setzeri | Mus_setzeri | nocturnal |
| mammals | Mus_shortridgei | Mus_shortridgei | nocturnal |
| mammals | Mus_sorella | Mus_sorella | nocturnal |
| mammals | Mus_spicilegus | Mus_spicilegus | nocturnal |
| mammals | Mus_spretus | Mus_spretus | nocturnal |
| mammals | Mus_tenellus | Mus_tenellus | nocturnal |
| mammals | Mus_terricolor | Mus_terricolor | nocturnal |
| mammals | Mus_triton | Mus_triton | cathemeral |
| mammals | Mus_vulcani | Mus_vulcani | nocturnal |
| mammals | Muscardinus_avellanarius | Muscardinus_avellanarius | nocturnal |
| mammals | Musonycteris_harrisoni | Musonycteris_harrisoni | nocturnal |
| mammals | Mustela_altaica | Mustela_altaica | nocturnal |
| mammals | Mustela_erminea | Mustela_erminea | cathemeral |
| mammals | Mustela_eversmanii | Mustela_eversmanii | nocturnal |

| | | | |
|---|---|---|---|
| mammals | Mustela_itatsi | Mustela_itatsi | nocturnal |
| mammals | Mustela_kathiah | Mustela_kathiah | cathemeral |
| mammals | Mustela_lutreola | Mustela_lutreola | nocturnal |
| mammals | Mustela_lutreolina | Mustela_lutreolina | nocturnal |
| mammals | Mustela_nigripes | Mustela_nigripes | nocturnal |
| mammals | Mustela_nivalis | Mustela_nivalis | cathemeral |
| mammals | Mustela_nudipes | Mustela_nudipes | cathemeral |
| mammals | Mustela_putorius | Mustela_putorius | cathemeral |
| mammals | Mustela_sibirica | Mustela_sibirica | nocturnal |
| mammals | Mustela_strigidorsa | Mustela_strigidorsa | cathemeral |
| mammals | Mydaus_javanensis | Mydaus_javanensis | nocturnal |
| mammals | Mydaus_marchei | Mydaus_marchei | nocturnal |
| mammals | Mylomys_dybowskii | Mylomys_dybowskii | diurnal |
| mammals | Mylomys_rex | Mylomys_rex | nocturnal |
| mammals | Myocastor_coypus | Myocastor_coypus | nocturnal |
| mammals | Myoictis_leucura | Myoictis_leucura | diurnal |
| mammals | Myoictis_melas | Myoictis_melas | diurnal |
| mammals | Myoictis_wallacii | Myoictis_wallacei | diurnal |
| mammals | Myoictis_wavica | Myoictis_wavicus | diurnal |
| mammals | Myomimus_personatus | Myomimus_personatus | nocturnal |
| mammals | Myomimus_roachi | Myomimus_roachi | nocturnal |
| mammals | Myomimus_setzeri | Myomimus_setzeri | nocturnal |
| mammals | Myomyscus_verreauxi | Myomyscus_verreauxii | nocturnal |
| mammals | Myonycteris_angolensis | Lissonycteris_angolensis | nocturnal |
| mammals | Myonycteris_brachycephala | Myonycteris_brachycephala | nocturnal |
| mammals | Myonycteris_leptodon | Myonycteris_leptodon | nocturnal |
| mammals | Myonycteris_relicta | Myonycteris_relicta | nocturnal |
| mammals | Myonycteris_torquata | Myonycteris_torquata | nocturnal |
| mammals | Myoprocta_acouchy | Myoprocta_acouchy | diurnal |
| mammals | Myoprocta_pratti | Myoprocta_pratti | diurnal |
| mammals | Myopterus_daubentonii | Myopterus_daubentonii | nocturnal |
| mammals | Myopterus_whitleyi | Myopterus_whitleyi | nocturnal |
| mammals | Myopus_schisticolor | Myopus_schisticolor | cathemeral |
| mammals | Myosciurus_pumilio | Myosciurus_pumilio | diurnal |

| | | | |
|---|---|---|---|
| mammals | Myosorex_babaulti | Myosorex_babaulti | cathemeral |
| mammals | Myosorex_blarina | Myosorex_blarina | nocturnal |
| mammals | Myosorex_cafer | Myosorex_cafer | nocturnal |
| mammals | Myosorex_eisentrauti | Myosorex_eisentrauti | cathemeral |
| mammals | Myosorex_geata | Myosorex_geata | cathemeral |
| mammals | Myosorex_kihaulei | Myosorex_kihaulei | nocturnal |
| mammals | Myosorex_longicaudatus | Myosorex_longicaudatus | cathemeral |
| mammals | Myosorex_okuensis | Myosorex_okuensis | cathemeral |
| mammals | Myosorex_rumpii | Myosorex_rumpii | cathemeral |
| mammals | Myosorex_schalleri | Myosorex_schalleri | cathemeral |
| mammals | Myosorex_sclateri | Myosorex_sclateri | cathemeral |
| mammals | Myosorex_tenuis | Myosorex_tenuis | cathemeral |
| mammals | Myosorex_varius | Myosorex_varius | cathemeral |
| mammals | Myotis_adversus | Myotis_adversus | nocturnal |
| mammals | Myotis_albescens | Myotis_albescens | nocturnal |
| mammals | Myotis_alcathoe | Myotis_alcathoe | nocturnal |
| mammals | Myotis_altarium | Myotis_altarium | nocturnal |
| mammals | Myotis_anjouanensis | Myotis_anjouanensis | nocturnal |
| mammals | Myotis_annectans | Myotis_annectans | nocturnal |
| mammals | Myotis_atacamensis | Myotis_atacamensis | nocturnal |
| mammals | Myotis_ater | Myotis_ater | nocturnal |
| mammals | Myotis_auriculus | Myotis_auriculus | nocturnal |
| mammals | Myotis_australis | Myotis_australis | nocturnal |
| mammals | Myotis_austroriparius | Myotis_austroriparius | nocturnal |
| mammals | Myotis_bechsteinii | Myotis_bechsteinii | nocturnal |
| mammals | Myotis_blythii | Myotis_blythii | nocturnal |
| mammals | Myotis_bocagii | Myotis_bocagii | nocturnal |
| mammals | Myotis_bombinus | Myotis_bombinus | nocturnal |
| mammals | Myotis_brandtii | Myotis_brandtii | nocturnal |
| mammals | Myotis_bucharensis | Myotis_bucharensis | nocturnal |
| mammals | Myotis_californicus | Myotis_californicus | nocturnal |
| mammals | Myotis_capaccinii | Myotis_capaccinii | nocturnal |
| mammals | Myotis_chiloensis | Myotis_chiloensis | nocturnal |
| mammals | Myotis_chinensis | Myotis_chinensis | nocturnal |

| | | | |
|---|---|---|---|
| mammals | Myotis_ciliolabrum | Myotis_ciliolabrum | nocturnal |
| mammals | Myotis_cobanensis | Myotis_cobanensis | nocturnal |
| mammals | Myotis_dasycneme | Myotis_dasycneme | nocturnal |
| mammals | Myotis_daubentonii | Myotis_daubentonii | nocturnal |
| mammals | Myotis_davidii | Myotis_davidii | nocturnal |
| mammals | Myotis_dieteri | Myotis_dieteri | nocturnal |
| mammals | Myotis_dinellii | Myotis_dinellii | nocturnal |
| mammals | Myotis_dominicensis | Myotis_dominicensis | nocturnal |
| mammals | Myotis_elegans | Myotis_elegans | nocturnal |
| mammals | Myotis_emarginatus | Myotis_emarginatus | nocturnal |
| mammals | Myotis_evotis | Myotis_evotis | nocturnal |
| mammals | Myotis_fimbriatus | Myotis_fimbriatus | nocturnal |
| mammals | Myotis_findleyi | Myotis_findleyi | nocturnal |
| mammals | Myotis_formosus | Myotis_formosus | nocturnal |
| mammals | Myotis_fortidens | Myotis_fortidens | nocturnal |
| mammals | Myotis_frater | Myotis_frater | nocturnal |
| mammals | Myotis_gomantongensis | Myotis_gomantongensis | nocturnal |
| mammals | Myotis_goudotii | Myotis_goudoti | nocturnal |
| mammals | Myotis_grisescens | Myotis_grisescens | nocturnal |
| mammals | Myotis_hasseltii | Myotis_hasseltii | nocturnal |
| mammals | Myotis_hermani | Myotis_hermani | nocturnal |
| mammals | Myotis_horsfieldii | Myotis_horsfieldii | nocturnal |
| mammals | Myotis_ikonnikovi | Myotis_ikonnikovi | nocturnal |
| mammals | Myotis_insularum | Myotis_insularum | nocturnal |
| mammals | Myotis_keaysi | Myotis_keaysi | nocturnal |
| mammals | Myotis_keenii | Myotis_keenii | nocturnal |
| mammals | Myotis_laniger | Myotis_laniger | nocturnal |
| mammals | Myotis_leibii | Myotis_leibii | nocturnal |
| mammals | Myotis_levis | Myotis_levis | nocturnal |
| mammals | Myotis_longipes | Myotis_longipes | nocturnal |
| mammals | Myotis_lucifugus | Myotis_lucifugus | nocturnal |
| mammals | Myotis_macrodactylus | Myotis_macrodactylus | nocturnal |
| mammals | Myotis_macropus | Myotis_macropus | nocturnal |
| mammals | Myotis_macrotarsus | Myotis_macrotarsus | nocturnal |

| | | | |
|---|---|---|---|
| mammals | Myotis_martiniquensis | Myotis_martiniquensis | nocturnal |
| mammals | Myotis_moluccarum | Myotis_moluccarum | nocturnal |
| mammals | Myotis_montivagus | Myotis_montivagus | nocturnal |
| mammals | Myotis_morrisi | Myotis_morrisi | nocturnal |
| mammals | Myotis_muricola | Myotis_muricola | nocturnal |
| mammals | Myotis_myotis | Myotis_myotis | nocturnal |
| mammals | Myotis_mystacinus | Myotis_mystacinus | nocturnal |
| mammals | Myotis_nattereri | Myotis_nattereri | nocturnal |
| mammals | Myotis_nesopolus | Myotis_nesopolus | nocturnal |
| mammals | Myotis_nigricans | Myotis_nigricans | nocturnal |
| mammals | Myotis_nipalensis | Myotis_nipalensis | nocturnal |
| mammals | Myotis_nyctor | Myotis_nyctor | nocturnal |
| mammals | Myotis_occultus | Myotis_occultus | nocturnal |
| mammals | Myotis_oxyotus | Myotis_oxyotus | nocturnal |
| mammals | Myotis_pequinius | Myotis_pequinius | nocturnal |
| mammals | Myotis_pilosus | Myotis_pilosus | nocturnal |
| mammals | Myotis_planiceps | Myotis_planiceps | nocturnal |
| mammals | Myotis_pruinosus | Myotis_pruinosus | nocturnal |
| mammals | Myotis_punicus | Myotis_punicus | nocturnal |
| mammals | Myotis_ridleyi | Myotis_ridleyi | nocturnal |
| mammals | Myotis_riparius | Myotis_riparius | nocturnal |
| mammals | Myotis_rosseti | Myotis_rosseti | nocturnal |
| mammals | Myotis_ruber | Myotis_ruber | nocturnal |
| mammals | Myotis_rufopictus | Myotis_rufopictus | nocturnal |
| mammals | Myotis_schaubi | Myotis_schaubi | nocturnal |
| mammals | Myotis_scotti | Myotis_scotti | nocturnal |
| mammals | Myotis_septentrionalis | Myotis_septentrionalis | nocturnal |
| mammals | Myotis_sicarius | Myotis_sicarius | nocturnal |
| mammals | Myotis_siligorensis | Myotis_siligorensis | nocturnal |
| mammals | Myotis_simus | Myotis_simus | nocturnal |
| mammals | Myotis_sodalis | Myotis_sodalis | nocturnal |
| mammals | Myotis_stalkeri | Myotis_stalkeri | nocturnal |
| mammals | Myotis_thysanodes | Myotis_thysanodes | nocturnal |
| mammals | Myotis_tricolor | Myotis_tricolor | nocturnal |

| | | | |
|---|---|---|---|
| mammals | Myotis_velifer | Myotis_velifer | nocturnal |
| mammals | Myotis_vivesi | Myotis_vivesi | nocturnal |
| mammals | Myotis_volans | Myotis_volans | nocturnal |
| mammals | Myotis_welwitschii | Myotis_welwitschii | nocturnal |
| mammals | Myotis_yanbarensis | Myotis_yanbarensis | nocturnal |
| mammals | Myotis_yumanensis | Myotis_yumanensis | nocturnal |
| mammals | Myrmecobius_fasciatus | Myrmecobius_fasciatus | diurnal |
| mammals | Myrmecophaga_tridactyla | Myrmecophaga_tridactyla | cathemeral |
| mammals | Mysateles_prehensilis | Mysateles_prehensilis | nocturnal |
| mammals | Mystacina_tuberculata | Mystacina_tuberculata | nocturnal |
| mammals | Mystromys_albicaudatus | Mystromys_albicaudatus | nocturnal |
| mammals | Myzopoda_aurita | Myzopoda_aurita | nocturnal |
| mammals | Myzopoda_schliemanni | Myzopoda_schliemanni | nocturnal |
| mammals | Naemorhedus_baileyi | Naemorhedus_baileyi | diurnal |
| mammals | Naemorhedus_caudatus | Naemorhedus_caudatus | nocturnal |
| mammals | Naemorhedus_goral | Naemorhedus_goral | crepuscular |
| mammals | Nandinia_binotata | Nandinia_binotata | nocturnal |
| mammals | Nanger_dama | Nanger_dama | diurnal |
| mammals | Nanger_granti | Nanger_granti | diurnal |
| mammals | Nanger_soemmerringii | Nanger_soemmerringii | diurnal |
| mammals | Nannosciurus_melanotis | Nannosciurus_melanotis | diurnal |
| mammals | Nanonycteris_veldkampii | Nanonycteris_veldkampii | nocturnal |
| mammals | Napaeozapus_insignis | Napaeozapus_insignis | nocturnal |
| mammals | Nasalis_larvatus | Nasalis_larvatus | diurnal |
| mammals | Nasua_narica | Nasua_narica | diurnal |
| mammals | Nasua_nasua | Nasua_nasua | diurnal |
| mammals | Nasua_olivacea | Nasuella_olivacea | diurnal |
| mammals | Natalus_jamaicensis | Natalus_jamaicensis | nocturnal |
| mammals | Natalus_macrourus | Natalus_espiritosantensis | nocturnal |
| mammals | Natalus_major | Natalus_major | nocturnal |
| mammals | Natalus_mexicanus | Natalus_mexicanus | nocturnal |
| mammals | Natalus_primus | Natalus_primus | nocturnal |
| mammals | Natalus_stramineus | Natalus_stramineus | nocturnal |
| mammals | Natalus_tumidirostris | Natalus_tumidirostris | nocturnal |

| | | | |
|---|---|---|---|
| mammals | Neacomys_guianae | Neacomys_guianae | nocturnal |
| mammals | Neacomys_minutus | Neacomys_minutus | nocturnal |
| mammals | Neacomys_musseri | Neacomys_musseri | nocturnal |
| mammals | Neacomys_pictus | Neacomys_pictus | nocturnal |
| mammals | Neacomys_spinosus | Neacomys_spinosus | nocturnal |
| mammals | Neacomys_tenuipes | Neacomys_tenuipes | nocturnal |
| mammals | Necromys_amoenus | Necromys_amoenus | diurnal |
| mammals | Necromys_lactens | Necromys_lactens | cathemeral |
| mammals | Necromys_lasiurus | Necromys_lasiurus | diurnal |
| mammals | Necromys_lenguarum | Necromys_lenguarum | diurnal |
| mammals | Necromys_obscurus | Necromys_obscurus | diurnal |
| mammals | Necromys_urichi | Necromys_urichi | cathemeral |
| mammals | Nectogale_elegans | Nectogale_elegans | cathemeral |
| mammals | Nectomys_apicalis | Nectomys_apicalis | nocturnal |
| mammals | Nectomys_magdalenae | Nectomys_magdalenae | nocturnal |
| mammals | Nectomys_palmipes | Nectomys_palmipes | nocturnal |
| mammals | Nectomys_rattus | Nectomys_rattus | nocturnal |
| mammals | Nectomys_squamipes | Nectomys_squamipes | nocturnal |
| mammals | Nelsonia_goldmani | Nelsonia_goldmani | nocturnal |
| mammals | Nelsonia_neotomodon | Nelsonia_neotomodon | nocturnal |
| mammals | Neodon_clarkei | Microtus_clarkei | cathemeral |
| mammals | Neodon_fuscus | Lasiopodomys_fuscus | diurnal |
| mammals | Neodon_irene | Neodon_irene | nocturnal |
| mammals | Neodon_leucurus | Phaiomys_leucurus | diurnal |
| mammals | Neoeptesicus_andinus | Eptesicus_andinus | nocturnal |
| mammals | Neoeptesicus_brasiliensis | Eptesicus_brasiliensis | nocturnal |
| mammals | Neoeptesicus_chiriquinus | Eptesicus_chiriquinus | nocturnal |
| mammals | Neoeptesicus_diminutus | Eptesicus_diminutus | nocturnal |
| mammals | Neoeptesicus_furinalis | Eptesicus_furinalis | nocturnal |
| mammals | Neoeptesicus_innoxius | Eptesicus_innoxius | nocturnal |
| mammals | Neofelis_diardi | Neofelis_diardi | cathemeral |
| mammals | Neofelis_nebulosa | Neofelis_nebulosa | nocturnal |
| mammals | Neofiber_alleni | Neofiber_alleni | nocturnal |
| mammals | Neogale_africana | Mustela_africana | diurnal |

| | | | |
|---|---|---|---|
| mammals | Neogale_felipei | Mustela_felipei | cathemeral |
| mammals | Neogale_frenata | Mustela_frenata | cathemeral |
| mammals | Neogale_vison | Neovison_vison | cathemeral |
| mammals | Neomicroxus_bogotensis | Neomicroxus_bogotensis | nocturnal |
| mammals | Neomicroxus_latebricola | Neomicroxus_latebricola | nocturnal |
| mammals | Neomys_anomalus | Neomys_anomalus | cathemeral |
| mammals | Neomys_fodiens | Neomys_fodiens | cathemeral |
| mammals | Neomys_teres | Neomys_teres | nocturnal |
| mammals | Neonycteris_pusilla | Neonycteris_pusilla | nocturnal |
| mammals | Neophascogale_lorentzii | Neophascogale_lorentzii | diurnal |
| mammals | Neoplatymops_mattogrossensis | Molossops_mattogrossensis | nocturnal |
| mammals | Neopteryx_frosti | Neopteryx_frosti | nocturnal |
| mammals | Neoromicia_anchietae | Pipistrellus_anchietae | nocturnal |
| mammals | Neoromicia_guineensis | Neoromicia_guineensis | nocturnal |
| mammals | Neoromicia_somalica | Neoromicia_somalica | nocturnal |
| mammals | Neoromicia_zuluensis | Neoromicia_zuluensis | nocturnal |
| mammals | Neotamias_alpinus | Neotamias_alpinus | diurnal |
| mammals | Neotamias_amoenus | Neotamias_amoenus | diurnal |
| mammals | Neotamias_bulleri | Neotamias_bulleri | diurnal |
| mammals | Neotamias_canipes | Neotamias_canipes | diurnal |
| mammals | Neotamias_cinereicollis | Neotamias_cinereicollis | diurnal |
| mammals | Neotamias_dorsalis | Neotamias_dorsalis | diurnal |
| mammals | Neotamias_durangae | Neotamias_durangae | diurnal |
| mammals | Neotamias_merriami | Neotamias_merriami | diurnal |
| mammals | Neotamias_minimus | Neotamias_minimus | diurnal |
| mammals | Neotamias_obscurus | Neotamias_obscurus | diurnal |
| mammals | Neotamias_ochrogenys | Neotamias_ochrogenys | diurnal |
| mammals | Neotamias_palmeri | Neotamias_palmeri | diurnal |
| mammals | Neotamias_panamintinus | Neotamias_panamintinus | diurnal |
| mammals | Neotamias_quadrimaculatus | Neotamias_quadrimaculatus | diurnal |
| mammals | Neotamias_quadrivittatus | Neotamias_quadrivittatus | diurnal |
| mammals | Neotamias_ruficaudus | Neotamias_ruficaudus | diurnal |
| mammals | Neotamias_rufus | Neotamias_rufus | diurnal |
| mammals | Neotamias_senex | Neotamias_senex | diurnal |

| | | | |
|---|---|---|---|
| mammals | Neotamias_siskiyou | Neotamias_siskiyou | diurnal |
| mammals | Neotamias_sonomae | Neotamias_sonomae | diurnal |
| mammals | Neotamias_speciosus | Neotamias_speciosus | diurnal |
| mammals | Neotamias_townsendii | Neotamias_townsendii | diurnal |
| mammals | Neotamias_umbrinus | Neotamias_umbrinus | diurnal |
| mammals | Neotoma_albigula | Neotoma_albigula | nocturnal |
| mammals | Neotoma_angustapalata | Neotoma_angustapalata | nocturnal |
| mammals | Neotoma_bryanti | Neotoma_bryanti | nocturnal |
| mammals | Neotoma_chrysomelas | Neotoma_chrysomelas | nocturnal |
| mammals | Neotoma_cinerea | Neotoma_cinerea | nocturnal |
| mammals | Neotoma_devia | Neotoma_devia | nocturnal |
| mammals | Neotoma_floridana | Neotoma_floridana | nocturnal |
| mammals | Neotoma_fuscipes | Neotoma_fuscipes | nocturnal |
| mammals | Neotoma_goldmani | Neotoma_goldmani | cathemeral |
| mammals | Neotoma_lepida | Neotoma_lepida | nocturnal |
| mammals | Neotoma_leucodon | Neotoma_leucodon | nocturnal |
| mammals | Neotoma_macrotis | Neotoma_macrotis | nocturnal |
| mammals | Neotoma_magister | Neotoma_magister | nocturnal |
| mammals | Neotoma_mexicana | Neotoma_mexicana | nocturnal |
| mammals | Neotoma_micropus | Neotoma_micropus | nocturnal |
| mammals | Neotoma_nelsoni | Neotoma_nelsoni | nocturnal |
| mammals | Neotoma_palatina | Neotoma_palatina | nocturnal |
| mammals | Neotoma_phenax | Neotoma_phenax | nocturnal |
| mammals | Neotoma_stephensi | Neotoma_stephensi | nocturnal |
| mammals | Neotomodon_alstoni | Neotomodon_alstoni | nocturnal |
| mammals | Neotomys_ebriosus | Neotomys_ebriosus | cathemeral |
| mammals | Neotragus_pygmaeus | Neotragus_pygmaeus | nocturnal |
| mammals | Nephelomys_albigularis | Nephelomys_albigularis | nocturnal |
| mammals | Nephelomys_auriventer | Nephelomys_auriventer | cathemeral |
| mammals | Nephelomys_caracolus | Nephelomys_caracolus | nocturnal |
| mammals | Nephelomys_devius | Nephelomys_devius | nocturnal |
| mammals | Nephelomys_keaysi | Nephelomys_keaysi | nocturnal |
| mammals | Nephelomys_levipes | Nephelomys_levipes | cathemeral |
| mammals | Nephelomys_meridensis | Nephelomys_meridensis | nocturnal |

| | | | |
|---|---|---|---|
| mammals | Nesogale_dobsoni | Microgale_dobsoni | nocturnal |
| mammals | Nesogale_talazaci | Microgale_talazaci | nocturnal |
| mammals | Nesolagus_netscheri | Nesolagus_netscheri | nocturnal |
| mammals | Nesolagus_timminsi | Nesolagus_timminsi | nocturnal |
| mammals | Nesomys_audeberti | Nesomys_audeberti | diurnal |
| mammals | Nesomys_lambertoni | Nesomys_lambertoni | diurnal |
| mammals | Nesomys_rufus | Nesomys_rufus | diurnal |
| mammals | Nesonycteris_fardoulisi | Melonycteris_fardoulisi | nocturnal |
| mammals | Nesonycteris_woodfordi | Melonycteris_woodfordi | nocturnal |
| mammals | Nesoryzomys_fernandinae | Nesoryzomys_fernandinae | nocturnal |
| mammals | Nesoryzomys_narboroughi | Nesoryzomys_narboroughi | nocturnal |
| mammals | Nesoryzomys_swarthi | Nesoryzomys_swarthi | nocturnal |
| mammals | Nesotragus_batesi | Neotragus_batesi | cathemeral |
| mammals | Nesotragus_moschatus | Nesotragus_moschatus | diurnal |
| mammals | Neurotrichus_gibbsii | Neurotrichus_gibbsii | cathemeral |
| mammals | Neusticomys_monticolus | Neusticomys_monticolus | nocturnal |
| mammals | Neusticomys_orcesi | Chibchanomys_orcesi | nocturnal |
| mammals | Nilgiritragus_hylocrius | Nilgiritragus_hylocrius | diurnal |
| mammals | Ningaui_ridei | Ningaui_ridei | nocturnal |
| mammals | Ningaui_timealeyi | Ningaui_timealeyi | nocturnal |
| mammals | Ningaui_yvonneae | Ningaui_yvonneae | nocturnal |
| mammals | Niviventer_andersoni | Niviventer_andersoni | nocturnal |
| mammals | Niviventer_brahma | Niviventer_brahma | nocturnal |
| mammals | Niviventer_cameroni | Niviventer_cameroni | nocturnal |
| mammals | Niviventer_confucianus | Niviventer_confucianus | nocturnal |
| mammals | Niviventer_coninga | Niviventer_coninga | nocturnal |
| mammals | Niviventer_cremoriventer | Niviventer_cremoriventer | nocturnal |
| mammals | Niviventer_culturatus | Niviventer_culturatus | nocturnal |
| mammals | Niviventer_eha | Niviventer_eha | nocturnal |
| mammals | Niviventer_excelsior | Niviventer_excelsior | nocturnal |
| mammals | Niviventer_fraternus | Niviventer_fraternus | nocturnal |
| mammals | Niviventer_fulvescens | Niviventer_fulvescens | nocturnal |
| mammals | Niviventer_hinpoon | Niviventer_hinpoon | nocturnal |
| mammals | Niviventer_lepturus | Niviventer_lepturus | nocturnal |

| | | | |
|---|---|---|---|
| mammals | Niviventer_niviventer | Niviventer_niviventer | nocturnal |
| mammals | Niviventer_rapit | Niviventer_rapit | nocturnal |
| mammals | Niviventer_tenaster | Niviventer_tenaster | nocturnal |
| mammals | Noctilio_albiventris | Noctilio_albiventris | nocturnal |
| mammals | Noctilio_leporinus | Noctilio_leporinus | nocturnal |
| mammals | Nomascus_concolor | Nomascus_concolor | diurnal |
| mammals | Nomascus_gabriellae | Nomascus_gabriellae | diurnal |
| mammals | Nomascus_hainanus | Nomascus_hainanus | diurnal |
| mammals | Nomascus_leucogenys | Nomascus_leucogenys | diurnal |
| mammals | Nomascus_nasutus | Nomascus_nasutus | diurnal |
| mammals | Nomascus_siki | Nomascus_siki | diurnal |
| mammals | Notamacropus_agilis | Macropus_agilis | nocturnal |
| mammals | Notamacropus_dorsalis | Macropus_dorsalis | nocturnal |
| mammals | Notamacropus_eugenii | Macropus_eugenii | nocturnal |
| mammals | Notamacropus_irma | Macropus_irma | diurnal |
| mammals | Notamacropus_parma | Macropus_parma | nocturnal |
| mammals | Notamacropus_parryi | Macropus_parryi | nocturnal |
| mammals | Notamacropus_rufogriseus | Macropus_rufogriseus | nocturnal |
| mammals | Nothocricetulus_migratorius | Cricetulus_migratorius | nocturnal |
| mammals | Notiomys_edwardsii | Notiomys_edwardsii | nocturnal |
| mammals | Notiosorex_cockrumi | Notiosorex_cockrumi | nocturnal |
| mammals | Notiosorex_crawfordi | Notiosorex_crawfordi | cathemeral |
| mammals | Notiosorex_evotis | Notiosorex_evotis | nocturnal |
| mammals | Notocitellus_adocetus | Notocitellus_adocetus | diurnal |
| mammals | Notocitellus_annulatus | Notocitellus_annulatus | diurnal |
| mammals | Notomys_alexis | Notomys_alexis | nocturnal |
| mammals | Notomys_aquilo | Notomys_aquilo | nocturnal |
| mammals | Notomys_cervinus | Notomys_cervinus | nocturnal |
| mammals | Notomys_fuscus | Notomys_fuscus | nocturnal |
| mammals | Notomys_mitchellii | Notomys_mitchellii | nocturnal |
| mammals | Notopteris_macdonaldii | Notopteris_macdonaldi | nocturnal |
| mammals | Notopteris_neocaledonicus | Notopteris_neocaledonica | nocturnal |
| mammals | Nyctalus_aviator | Nyctalus_aviator | nocturnal |
| mammals | Nyctalus_azoreum | Nyctalus_azoreum | nocturnal |

| | | | |
|---|---|---|---|
| mammals | Nyctalus_furvus | Nyctalus_furvus | nocturnal |
| mammals | Nyctalus_lasiopterus | Nyctalus_lasiopterus | nocturnal |
| mammals | Nyctalus_leisleri | Nyctalus_leisleri | nocturnal |
| mammals | Nyctalus_montanus | Nyctalus_montanus | nocturnal |
| mammals | Nyctalus_noctula | Nyctalus_noctula | nocturnal |
| mammals | Nyctalus_plancyi | Nyctalus_plancyi | nocturnal |
| mammals | Nyctereutes_procyonoides | Nyctereutes_procyonoides | cathemeral |
| mammals | Nycteris_arge | Nycteris_arge | nocturnal |
| mammals | Nycteris_aurita | Nycteris_aurita | nocturnal |
| mammals | Nycteris_gambiensis | Nycteris_gambiensis | nocturnal |
| mammals | Nycteris_grandis | Nycteris_grandis | nocturnal |
| mammals | Nycteris_hispida | Nycteris_hispida | nocturnal |
| mammals | Nycteris_intermedia | Nycteris_intermedia | nocturnal |
| mammals | Nycteris_javanica | Nycteris_javanica | nocturnal |
| mammals | Nycteris_macrotis | Nycteris_macrotis | nocturnal |
| mammals | Nycteris_major | Nycteris_major | nocturnal |
| mammals | Nycteris_nana | Nycteris_nana | nocturnal |
| mammals | Nycteris_parisii | Nycteris_parisii | nocturnal |
| mammals | Nycteris_thebaica | Nycteris_thebaica | nocturnal |
| mammals | Nycteris_tragata | Nycteris_tragata | nocturnal |
| mammals | Nycteris_woodi | Nycteris_woodi | nocturnal |
| mammals | Nycticebus_bengalensis | Nycticebus_bengalensis | nocturnal |
| mammals | Nycticebus_coucang | Nycticebus_coucang | nocturnal |
| mammals | Nycticebus_menagensis | Nycticebus_menagensis | nocturnal |
| mammals | Nycticeinops_crassulus | Pipistrellus_crassulus | nocturnal |
| mammals | Nycticeinops_eisentrauti | Pipistrellus_eisentrauti | nocturnal |
| mammals | Nycticeinops_grandidieri | Neoromicia_flavescens | nocturnal |
| mammals | Nycticeinops_schlieffenii | Nycticeinops_schlieffeni | nocturnal |
| mammals | Nycticeius_cubanus | Nycticeius_cubanus | nocturnal |
| mammals | Nycticeius_humeralis | Nycticeius_humeralis | nocturnal |
| mammals | Nyctiellus_lepidus | Nyctiellus_lepidus | nocturnal |
| mammals | Nyctimene_aello | Nyctimene_aello | nocturnal |
| mammals | Nyctimene_albiventer | Nyctimene_albiventer | nocturnal |
| mammals | Nyctimene_cephalotes | Nyctimene_cephalotes | nocturnal |

| | | | |
|---|---|---|---|
| mammals | Nyctimene_certans | Nyctimene_certans | nocturnal |
| mammals | Nyctimene_cyclotis | Nyctimene_cyclotis | nocturnal |
| mammals | Nyctimene_draconilla | Nyctimene_draconilla | nocturnal |
| mammals | Nyctimene_keasti | Nyctimene_keasti | nocturnal |
| mammals | Nyctimene_major | Nyctimene_major | nocturnal |
| mammals | Nyctimene_malaitensis | Nyctimene_malaitensis | nocturnal |
| mammals | Nyctimene_masalai | Nyctimene_masalai | nocturnal |
| mammals | Nyctimene_rabori | Nyctimene_rabori | nocturnal |
| mammals | Nyctimene_robinsoni | Nyctimene_robinsoni | nocturnal |
| mammals | Nyctimene_sanctacrucis | Nyctimene_sanctacrucis | nocturnal |
| mammals | Nyctimene_varia | Nyctimene_minutus | nocturnal |
| mammals | Nyctimene_vizcaccia | Nyctimene_vizcaccia | nocturnal |
| mammals | Nyctinomops_aurispinosus | Nyctinomops_aurispinosus | nocturnal |
| mammals | Nyctinomops_femorosaccus | Nyctinomops_femorosaccus | nocturnal |
| mammals | Nyctinomops_laticaudatus | Nyctinomops_laticaudatus | nocturnal |
| mammals | Nyctinomops_macrotis | Nyctinomops_macrotis | nocturnal |
| mammals | Nyctomys_sumichrasti | Nyctomys_sumichrasti | nocturnal |
| mammals | Nyctophilus_arnhemensis | Nyctophilus_arnhemensis | nocturnal |
| mammals | Nyctophilus_bifax | Nyctophilus_bifax | nocturnal |
| mammals | Nyctophilus_geoffroyi | Nyctophilus_geoffroyi | nocturnal |
| mammals | Nyctophilus_gouldi | Nyctophilus_gouldi | nocturnal |
| mammals | Nyctophilus_heran | Nyctophilus_heran | nocturnal |
| mammals | Nyctophilus_microdon | Nyctophilus_microdon | nocturnal |
| mammals | Nyctophilus_microtis | Nyctophilus_microtis | nocturnal |
| mammals | Nyctophilus_sherrini | Nyctophilus_sherrini | nocturnal |
| mammals | Nyctophilus_walkeri | Nyctophilus_walkeri | nocturnal |
| mammals | Ochotona_alpina | Ochotona_alpina | diurnal |
| mammals | Ochotona_argentata | Ochotona_argentata | crepuscular |
| mammals | Ochotona_cansus | Ochotona_cansus | diurnal |
| mammals | Ochotona_collaris | Ochotona_collaris | diurnal |
| mammals | Ochotona_coreana | Ochotona_coreana | diurnal |
| mammals | Ochotona_curzoniae | Ochotona_curzoniae | diurnal |
| mammals | Ochotona_dauurica | Ochotona_dauurica | diurnal |
| mammals | Ochotona_erythrotis | Ochotona_erythrotis | diurnal |

| | | | |
|---|---|---|---|
| mammals | Ochotona_forresti | Ochotona_forresti | cathemeral |
| mammals | Ochotona_hoffmanni | Ochotona_hoffmanni | diurnal |
| mammals | Ochotona_hyperborea | Ochotona_hyperborea | diurnal |
| mammals | Ochotona_iliensis | Ochotona_iliensis | cathemeral |
| mammals | Ochotona_koslowi | Ochotona_koslowi | diurnal |
| mammals | Ochotona_ladacensis | Ochotona_ladacensis | diurnal |
| mammals | Ochotona_macrotis | Ochotona_macrotis | diurnal |
| mammals | Ochotona_mantchurica | Ochotona_mantchurica | diurnal |
| mammals | Ochotona_nubrica | Ochotona_nubrica | diurnal |
| mammals | Ochotona_opaca | Ochotona_opaca | cathemeral |
| mammals | Ochotona_pallasii | Ochotona_pallasii | diurnal |
| mammals | Ochotona_princeps | Ochotona_princeps | diurnal |
| mammals | Ochotona_pusilla | Ochotona_pusilla | nocturnal |
| mammals | Ochotona_roylii | Ochotona_roylei | diurnal |
| mammals | Ochotona_rufescens | Ochotona_rufescens | diurnal |
| mammals | Ochotona_rutila | Ochotona_rutila | diurnal |
| mammals | Ochotona_thibetana | Ochotona_thibetana | diurnal |
| mammals | Ochotona_thomasi | Ochotona_thomasi | diurnal |
| mammals | Ochotona_turuchanensis | Ochotona_turuchanensis | diurnal |
| mammals | Ochromyscus_brockmani | Myomyscus_brockmani | nocturnal |
| mammals | Ochromyscus_yemeni | Myomyscus_yemeni | nocturnal |
| mammals | Ochrotomys_nuttalli | Ochrotomys_nuttalli | nocturnal |
| mammals | Octodon_bridgesii | Octodon_bridgesi | nocturnal |
| mammals | Octodon_degus | Octodon_degus | diurnal |
| mammals | Octodon_lunatus | Octodon_lunatus | cathemeral |
| mammals | Octodon_pacificus | Octodon_pacificus | diurnal |
| mammals | Octodontomys_gliroides | Octodontomys_gliroides | nocturnal |
| mammals | Octomys_mimax | Octomys_mimax | nocturnal |
| mammals | Odocoileus_hemionus | Odocoileus_hemionus | crepuscular |
| mammals | Odocoileus_pandora | Mazama_pandora | crepuscular |
| mammals | Odocoileus_virginianus | Odocoileus_virginianus | crepuscular |
| mammals | Oecomys_bicolor | Oecomys_bicolor | nocturnal |
| mammals | Oecomys_flavicans | Oecomys_flavicans | nocturnal |
| mammals | Oecomys_mamorae | Oecomys_mamorae | nocturnal |

| | | | |
|---|---|---|---|
| mammals | Oecomys_phaeotis | Oecomys_phaeotis | nocturnal |
| mammals | Oecomys_speciosus | Oecomys_speciosus | nocturnal |
| mammals | Oecomys_superans | Oecomys_superans | nocturnal |
| mammals | Oecomys_trinitatis | Oecomys_trinitatis | nocturnal |
| mammals | Oenomys_hypoxanthus | Oenomys_hypoxanthus | cathemeral |
| mammals | Oenomys_ornatus | Oenomys_ornatus | cathemeral |
| mammals | Okapia_johnstoni | Okapia_johnstoni | diurnal |
| mammals | Olallamys_albicauda | Olallamys_albicauda | nocturnal |
| mammals | Olallamys_edax | Olallamys_edax | nocturnal |
| mammals | Oligoryzomys_andinus | Oligoryzomys_andinus | nocturnal |
| mammals | Oligoryzomys_arenalis | Oligoryzomys_arenalis | nocturnal |
| mammals | Oligoryzomys_brendae | Oligoryzomys_brendae | nocturnal |
| mammals | Oligoryzomys_chacoensis | Oligoryzomys_chacoensis | nocturnal |
| mammals | Oligoryzomys_destructor | Oligoryzomys_destructor | nocturnal |
| mammals | Oligoryzomys_flavescens | Oligoryzomys_flavescens | nocturnal |
| mammals | Oligoryzomys_fulvescens | Oligoryzomys_fulvescens | nocturnal |
| mammals | Oligoryzomys_griseolus | Oligoryzomys_griseolus | nocturnal |
| mammals | Oligoryzomys_longicaudatus | Oligoryzomys_longicaudatus | nocturnal |
| mammals | Oligoryzomys_microtis | Oligoryzomys_microtis | nocturnal |
| mammals | Oligoryzomys_nigripes | Oligoryzomys_nigripes | nocturnal |
| mammals | Oligoryzomys_stramineus | Oligoryzomys_stramineus | nocturnal |
| mammals | Oligoryzomys_vegetus | Oligoryzomys_vegetus | nocturnal |
| mammals | Ondatra_zibethicus | Ondatra_zibethicus | cathemeral |
| mammals | Onychogalea_frenata | Onychogalea_fraenata | nocturnal |
| mammals | Onychogalea_unguifera | Onychogalea_unguifera | nocturnal |
| mammals | Onychomys_arenicola | Onychomys_arenicola | nocturnal |
| mammals | Onychomys_leucogaster | Onychomys_leucogaster | nocturnal |
| mammals | Onychomys_torridus | Onychomys_torridus | nocturnal |
| mammals | Oreamnos_americanus | Oreamnos_americanus | cathemeral |
| mammals | Oreoryzomys_balneator | Oreoryzomys_balneator | nocturnal |
| mammals | Oreoscaptor_mizura | Euroscaptor_mizura | cathemeral |
| mammals | Oreotragus_oreotragus | Oreotragus_oreotragus | diurnal |
| mammals | Orientallactaga_balikunica | Allactaga_balikunica | nocturnal |
| mammals | Orientallactaga_bullata | Allactaga_bullata | nocturnal |

| | | | |
|---|---|---|---|
| mammals | Orientallactaga_sibirica | Allactaga_sibirica | nocturnal |
| mammals | Ornithorhynchus_anatinus | Ornithorhynchus_anatinus | cathemeral |
| mammals | Orthogeomys_grandis | Orthogeomys_cuniculus | cathemeral |
| mammals | Orycteropus_afer | Orycteropus_afer | nocturnal |
| mammals | Oryctolagus_cuniculus | Oryctolagus_cuniculus | cathemeral |
| mammals | Oryx_beisa | Oryx_beisa | cathemeral |
| mammals | Oryx_gazella | Oryx_gazella | cathemeral |
| mammals | Oryx_leucoryx | Oryx_leucoryx | cathemeral |
| mammals | Oryzomys_couesi | Oryzomys_couesi | nocturnal |
| mammals | Oryzomys_dimidiatus | Oryzomys_dimidiatus | cathemeral |
| mammals | Oryzomys_gorgasi | Oryzomys_gorgasi | cathemeral |
| mammals | Oryzomys_palustris | Oryzomys_palustris | nocturnal |
| mammals | Oryzorictes_hova | Oryzorictes_hova | nocturnal |
| mammals | Oryzorictes_tetradactylus | Oryzorictes_tetradactylus | nocturnal |
| mammals | Osgoodomys_banderanus | Osgoodomys_banderanus | nocturnal |
| mammals | Osphranter_antilopinus | Macropus_antilopinus | nocturnal |
| mammals | Osphranter_bernardus | Macropus_bernardus | nocturnal |
| mammals | Osphranter_robustus | Macropus_robustus | nocturnal |
| mammals | Osphranter_rufus | Macropus_rufus | nocturnal |
| mammals | Otocolobus_manul | Otocolobus_manul | cathemeral |
| mammals | Otocyon_megalotis | Otocyon_megalotis | nocturnal |
| mammals | Otohylomys_megalotis | Hylomys_megalotis | cathemeral |
| mammals | Otolemur_crassicaudatus | Otolemur_crassicaudatus | nocturnal |
| mammals | Otolemur_garnettii | Otolemur_garnettii | nocturnal |
| mammals | Otomops_formosus | Otomops_formosus | nocturnal |
| mammals | Otomops_johnstonei | Otomops_johnstonei | nocturnal |
| mammals | Otomops_madagascariensis | Otomops_madagascariensis | nocturnal |
| mammals | Otomops_martiensseni | Otomops_martiensseni | nocturnal |
| mammals | Otomops_papuensis | Otomops_papuensis | nocturnal |
| mammals | Otomops_wroughtoni | Otomops_wroughtoni | nocturnal |
| mammals | Otomys_anchietae | Otomys_anchietae | diurnal |
| mammals | Otomys_angoniensis | Otomys_angoniensis | cathemeral |
| mammals | Otomys_barbouri | Otomys_barbouri | diurnal |
| mammals | Otomys_burtoni | Otomys_burtoni | diurnal |

| | | | |
|---|---|---|---|
| mammals | Otomys_cuanzensis | Otomys_cuanzensis | diurnal |
| mammals | Otomys_denti | Otomys_denti | diurnal |
| mammals | Otomys_irroratus | Otomys_irroratus | diurnal |
| mammals | Otomys_karoensis | Otomys_saundersiae | diurnal |
| mammals | Otomys_lacustris | Otomys_lacustris | diurnal |
| mammals | Otomys_laminatus | Otomys_laminatus | diurnal |
| mammals | Otomys_occidentalis | Otomys_occidentalis | diurnal |
| mammals | Otomys_sloggetti | Otomys_sloggetti | diurnal |
| mammals | Otomys_tropicalis | Otomys_tropicalis | diurnal |
| mammals | Otomys_typus | Otomys_typus | cathemeral |
| mammals | Otomys_unisulcatus | Otomys_unisulcatus | diurnal |
| mammals | Otonycteris_hemprichii | Otonycteris_hemprichii | nocturnal |
| mammals | Otonyctomys_hatti | Otonyctomys_hatti | nocturnal |
| mammals | Otopteropus_cartilagonodus | Otopteropus_cartilagonodus | nocturnal |
| mammals | Otospermophilus_beecheyi | Otospermophilus_beecheyi | diurnal |
| mammals | Otospermophilus_variegatus | Otospermophilus_variegatus | diurnal |
| mammals | Ototylomys_phyllotis | Ototylomys_phyllotis | nocturnal |
| mammals | Ourebia_ourebi | Ourebia_ourebi | cathemeral |
| mammals | Ovibos_moschatus | Ovibos_moschatus | cathemeral |
| mammals | Ovis_ammon | Ovis_ammon | crepuscular |
| mammals | Ovis_canadensis | Ovis_canadensis | crepuscular |
| mammals | Ovis_dalli | Ovis_dalli | diurnal |
| mammals | Ovis_nivicola | Ovis_nivicola | diurnal |
| mammals | Oxymycterus_amazonicus | Oxymycterus_amazonicus | nocturnal |
| mammals | Oxymycterus_caparaoe | Oxymycterus_caparoae | diurnal |
| mammals | Oxymycterus_dasytrichus | Oxymycterus_dasytrichus | cathemeral |
| mammals | Oxymycterus_delator | Oxymycterus_delator | diurnal |
| mammals | Oxymycterus_hiska | Oxymycterus_hiska | cathemeral |
| mammals | Oxymycterus_hucucha | Oxymycterus_hucucha | cathemeral |
| mammals | Oxymycterus_inca | Oxymycterus_inca | diurnal |
| mammals | Oxymycterus_josei | Oxymycterus_josei | diurnal |
| mammals | Oxymycterus_nasutus | Oxymycterus_nasutus | diurnal |
| mammals | Oxymycterus_paramensis | Oxymycterus_paramensis | cathemeral |
| mammals | Oxymycterus_quaestor | Oxymycterus_quaestor | diurnal |

| | | | |
|---|---|---|---|
| mammals | Oxymycterus_rufus | Oxymycterus_rufus | cathemeral |
| mammals | Ozimops_beccarii | Mormopterus_beccarii | nocturnal |
| mammals | Ozimops_loriae | Mormopterus_loriae | nocturnal |
| mammals | Ozimops_planiceps | Mormopterus_planiceps | nocturnal |
| mammals | Ozotoceros_bezoarticus | Ozotoceros_bezoarticus | cathemeral |
| mammals | Pachyuromys_duprasi | Pachyuromys_duprasi | nocturnal |
| mammals | Paguma_larvata | Paguma_larvata | nocturnal |
| mammals | Palawanomys_furvus | Palawanomys_furvus | nocturnal |
| mammals | Palawanosorex_ater | Suncus_ater | nocturnal |
| mammals | Pan_paniscus | Pan_paniscus | diurnal |
| mammals | Pan_troglodytes | Pan_troglodytes | diurnal |
| mammals | Panthera_leo | Panthera_leo | cathemeral |
| mammals | Panthera_onca | Panthera_onca | cathemeral |
| mammals | Panthera_pardus | Panthera_pardus | cathemeral |
| mammals | Panthera_tigris | Panthera_tigris | cathemeral |
| mammals | Panthera_uncia | Panthera_uncia | cathemeral |
| mammals | Pantholops_hodgsonii | Pantholops_hodgsonii | crepuscular |
| mammals | Papagomys_armandvillei | Papagomys_armandvillei | nocturnal |
| mammals | Papio_anubis | Papio_anubis | diurnal |
| mammals | Papio_cynocephalus | Papio_cynocephalus | diurnal |
| mammals | Papio_hamadryas | Papio_hamadryas | diurnal |
| mammals | Papio_kindae | Papio_kindae | diurnal |
| mammals | Papio_papio | Papio_papio | diurnal |
| mammals | Papio_ursinus | Papio_ursinus | diurnal |
| mammals | Parablarinella_griselda | Blarinella_griselda | nocturnal |
| mammals | Parachoerus_wagneri | Catagonus_wagneri | diurnal |
| mammals | Paracrocidura_graueri | Paracrocidura_graueri | nocturnal |
| mammals | Paracrocidura_maxima | Paracrocidura_maxima | nocturnal |
| mammals | Paracrocidura_schoutedeni | Paracrocidura_schoutedeni | nocturnal |
| mammals | Paracynictis_selousi | Paracynictis_selousi | nocturnal |
| mammals | Paradipus_ctenodactylus | Paradipus_ctenodactylus | nocturnal |
| mammals | Paradoxurus_hermaphroditus | Paradoxurus_hermaphroditus | nocturnal |
| mammals | Paradoxurus_jerdoni | Paradoxurus_jerdoni | nocturnal |
| mammals | Paradoxurus_zeylonensis | Paradoxurus_zeylonensis | nocturnal |

| | | | |
|---|---|---|---|
| mammals | Paraechinus_aethiopicus | Paraechinus_aethiopicus | nocturnal |
| mammals | Paraechinus_hypomelas | Paraechinus_hypomelas | nocturnal |
| mammals | Paraechinus_micropus | Paraechinus_micropus | nocturnal |
| mammals | Paraechinus_nudiventris | Paraechinus_nudiventris | nocturnal |
| mammals | Paragalago_cocos | Galagoides_cocos | nocturnal |
| mammals | Paragalago_granti | Galagoides_granti | nocturnal |
| mammals | Paragalago_orinus | Galagoides_orinus | nocturnal |
| mammals | Paragalago_rondoensis | Galagoides_rondoensis | nocturnal |
| mammals | Paragalago_zanzibaricus | Galagoides_zanzibaricus | nocturnal |
| mammals | Parahyaena_brunnea | Parahyaena_brunnea | nocturnal |
| mammals | Paraleptomys_rufilatus | Paraleptomys_rufilatus | nocturnal |
| mammals | Paraleptomys_wilhelmina | Paraleptomys_wilhelmina | nocturnal |
| mammals | Paramelomys_levipes | Paramelomys_levipes | nocturnal |
| mammals | Paramelomys_lorentzii | Paramelomys_lorentzii | nocturnal |
| mammals | Paramelomys_mollis | Paramelomys_mollis | nocturnal |
| mammals | Paramelomys_moncktoni | Paramelomys_moncktoni | nocturnal |
| mammals | Paramelomys_naso | Paramelomys_naso | nocturnal |
| mammals | Paramelomys_platyops | Paramelomys_platyops | nocturnal |
| mammals | Paramelomys_rubex | Paramelomys_rubex | nocturnal |
| mammals | Paramelomys_steini | Paramelomys_steini | nocturnal |
| mammals | Parantechinus_apicalis | Parantechinus_apicalis | nocturnal |
| mammals | Paranyctimene_raptor | Paranyctimene_raptor | nocturnal |
| mammals | Parastrellus_hesperus | Pipistrellus_hesperus | nocturnal |
| mammals | Paratriaenops_auritus | Paratriaenops_auritus | nocturnal |
| mammals | Paratriaenops_furcula | Paratriaenops_furculus | nocturnal |
| mammals | Paraxerus_alexandri | Paraxerus_alexandri | diurnal |
| mammals | Paraxerus_boehmi | Paraxerus_boehmi | diurnal |
| mammals | Paraxerus_cepapi | Paraxerus_cepapi | diurnal |
| mammals | Paraxerus_cooperi | Paraxerus_cooperi | diurnal |
| mammals | Paraxerus_flavovittis | Paraxerus_flavovittis | diurnal |
| mammals | Paraxerus_lucifer | Paraxerus_lucifer | diurnal |
| mammals | Paraxerus_ochraceus | Paraxerus_ochraceus | diurnal |
| mammals | Paraxerus_palliatus | Paraxerus_palliatus | diurnal |
| mammals | Paraxerus_poensis | Paraxerus_poensis | diurnal |

| | | | |
|---|---|---|---|
| mammals | Paraxerus_vexillarius | Paraxerus_vexillarius | diurnal |
| mammals | Paraxerus_vincenti | Paraxerus_vincenti | diurnal |
| mammals | Pardofelis_marmorata | Pardofelis_marmorata | cathemeral |
| mammals | Paremballonura_atrata | Paremballonura_atrata | diurnal |
| mammals | Paremballonura_tiavato | Paremballonura_tiavato | nocturnal |
| mammals | Parotomys_brantsii | Parotomys_brantsii | diurnal |
| mammals | Parotomys_littledalei | Parotomys_littledalei | diurnal |
| mammals | Passalites_nemorivagus | Mazama_nemorivaga | cathemeral |
| mammals | Pattonomys_semivillosus | Pattonomys_semivillosus | nocturnal |
| mammals | Paucidentomys_vermidax | Paucidentomys_vermidax | nocturnal |
| mammals | Paulamys_naso | Paulamys_naso | nocturnal |
| mammals | Paynomys_macronyx | Chelemys_macronyx | nocturnal |
| mammals | Pectinator_spekei | Pectinator_spekei | diurnal |
| mammals | Pedetes_capensis | Pedetes_capensis | nocturnal |
| mammals | Pedetes_surdaster | Pedetes_surdaster | nocturnal |
| mammals | Pekania_pennanti | Martes_pennanti | cathemeral |
| mammals | Pelea_capreolus | Pelea_capreolus | cathemeral |
| mammals | Pelomys_campanae | Pelomys_campanae | nocturnal |
| mammals | Pelomys_fallax | Pelomys_fallax | cathemeral |
| mammals | Pelomys_hopkinsi | Pelomys_hopkinsi | nocturnal |
| mammals | Pelomys_isseli | Pelomys_isseli | nocturnal |
| mammals | Pelomys_minor | Pelomys_minor | nocturnal |
| mammals | Pentalagus_furnessi | Pentalagus_furnessi | nocturnal |
| mammals | Penthetor_lucasii | Penthetor_lucasi | nocturnal |
| mammals | Perameles_bougainville | Perameles_bougainville | nocturnal |
| mammals | Perameles_gunnii | Perameles_gunnii | nocturnal |
| mammals | Perameles_nasuta | Perameles_nasuta | nocturnal |
| mammals | Perimyotis_subflavus | Pipistrellus_subflavus | nocturnal |
| mammals | Perodicticus_edwardsi | Perodicticus_edwardsi | nocturnal |
| mammals | Perodicticus_ibeanus | Perodicticus_ibeanus | nocturnal |
| mammals | Perodicticus_potto | Perodicticus_potto | nocturnal |
| mammals | Perognathus_alticola | Perognathus_alticola | nocturnal |
| mammals | Perognathus_amplus | Perognathus_amplus | nocturnal |
| mammals | Perognathus_fasciatus | Perognathus_fasciatus | nocturnal |

| | | | |
|---|---|---|---|
| mammals | Perognathus_flavescens | Perognathus_flavescens | nocturnal |
| mammals | Perognathus_flavus | Perognathus_flavus | nocturnal |
| mammals | Perognathus_inornatus | Perognathus_inornatus | nocturnal |
| mammals | Perognathus_longimembris | Perognathus_longimembris | nocturnal |
| mammals | Perognathus_merriami | Perognathus_merriami | nocturnal |
| mammals | Perognathus_parvus | Perognathus_parvus | nocturnal |
| mammals | Peromyscus_attwateri | Peromyscus_attwateri | nocturnal |
| mammals | Peromyscus_aztecus | Peromyscus_aztecus | nocturnal |
| mammals | Peromyscus_beatae | Peromyscus_beatae | nocturnal |
| mammals | Peromyscus_boylii | Peromyscus_boylii | nocturnal |
| mammals | Peromyscus_bullatus | Peromyscus_bullatus | nocturnal |
| mammals | Peromyscus_californicus | Peromyscus_californicus | nocturnal |
| mammals | Peromyscus_caniceps | Peromyscus_caniceps | nocturnal |
| mammals | Peromyscus_crinitus | Peromyscus_crinitus | nocturnal |
| mammals | Peromyscus_difficilis | Peromyscus_difficilis | nocturnal |
| mammals | Peromyscus_eremicus | Peromyscus_eremicus | nocturnal |
| mammals | Peromyscus_eva | Peromyscus_eva | nocturnal |
| mammals | Peromyscus_fraterculus | Peromyscus_fraterculus | nocturnal |
| mammals | Peromyscus_furvus | Peromyscus_furvus | nocturnal |
| mammals | Peromyscus_gossypinus | Peromyscus_gossypinus | nocturnal |
| mammals | Peromyscus_grandis | Peromyscus_grandis | nocturnal |
| mammals | Peromyscus_gratus | Peromyscus_gratus | nocturnal |
| mammals | Peromyscus_guatemalensis | Peromyscus_guatemalensis | nocturnal |
| mammals | Peromyscus_gymnotis | Peromyscus_gymnotis | nocturnal |
| mammals | Peromyscus_hooperi | Peromyscus_hooperi | nocturnal |
| mammals | Peromyscus_hylocetes | Peromyscus_hylocetes | nocturnal |
| mammals | Peromyscus_keeni | Peromyscus_keeni | nocturnal |
| mammals | Peromyscus_leucopus | Peromyscus_leucopus | nocturnal |
| mammals | Peromyscus_levipes | Peromyscus_levipes | nocturnal |
| mammals | Peromyscus_madrensis | Peromyscus_madrensis | nocturnal |
| mammals | Peromyscus_maniculatus | Peromyscus_maniculatus | nocturnal |
| mammals | Peromyscus_mayensis | Peromyscus_mayensis | nocturnal |
| mammals | Peromyscus_megalops | Peromyscus_megalops | nocturnal |
| mammals | Peromyscus_mekisturus | Peromyscus_mekisturus | nocturnal |

| | | | |
|---|---|---|---|
| mammals | Peromyscus_melanocarpus | Peromyscus_melanocarpus | nocturnal |
| mammals | Peromyscus_melanophrys | Peromyscus_melanophrys | nocturnal |
| mammals | Peromyscus_melanotis | Peromyscus_melanotis | nocturnal |
| mammals | Peromyscus_melanurus | Peromyscus_melanurus | nocturnal |
| mammals | Peromyscus_merriami | Peromyscus_merriami | nocturnal |
| mammals | Peromyscus_mexicanus | Peromyscus_mexicanus | nocturnal |
| mammals | Peromyscus_nasutus | Peromyscus_nasutus | nocturnal |
| mammals | Peromyscus_ochraventer | Peromyscus_ochraventer | nocturnal |
| mammals | Peromyscus_pectoralis | Peromyscus_pectoralis | nocturnal |
| mammals | Peromyscus_perfulvus | Peromyscus_perfulvus | nocturnal |
| mammals | Peromyscus_polionotus | Peromyscus_polionotus | nocturnal |
| mammals | Peromyscus_polius | Peromyscus_polius | nocturnal |
| mammals | Peromyscus_sagax | Peromyscus_sagax | nocturnal |
| mammals | Peromyscus_schmidlyi | Peromyscus_schmidlyi | nocturnal |
| mammals | Peromyscus_sejugis | Peromyscus_sejugis | nocturnal |
| mammals | Peromyscus_simulus | Peromyscus_simulus | nocturnal |
| mammals | Peromyscus_slevini | Peromyscus_slevini | nocturnal |
| mammals | Peromyscus_spicilegus | Peromyscus_spicilegus | nocturnal |
| mammals | Peromyscus_stephani | Peromyscus_stephani | nocturnal |
| mammals | Peromyscus_stirtoni | Peromyscus_stirtoni | nocturnal |
| mammals | Peromyscus_truei | Peromyscus_truei | nocturnal |
| mammals | Peromyscus_winkelmanni | Peromyscus_winkelmanni | nocturnal |
| mammals | Peromyscus_yucatanicus | Peromyscus_yucatanicus | nocturnal |
| mammals | Peromyscus_zarhynchus | Peromyscus_zarhynchus | nocturnal |
| mammals | Peropteryx_kappleri | Peropteryx_kappleri | nocturnal |
| mammals | Peropteryx_leucoptera | Peropteryx_leucoptera | nocturnal |
| mammals | Peropteryx_macrotis | Peropteryx_macrotis | nocturnal |
| mammals | Peropteryx_pallidoptera | Peropteryx_pallidoptera | nocturnal |
| mammals | Peropteryx_trinitatis | Peropteryx_trinitatis | nocturnal |
| mammals | Peroryctes_broadbenti | Peroryctes_broadbenti | nocturnal |
| mammals | Peroryctes_raffrayanus | Peroryctes_raffrayana | nocturnal |
| mammals | Petaurillus_emiliae | Petaurillus_emiliae | nocturnal |
| mammals | Petaurillus_hosei | Petaurillus_hosei | nocturnal |
| mammals | Petaurillus_kinlochii | Petaurillus_kinlochii | nocturnal |

| | | | |
|---|---|---|---|
| mammals | Petaurista_alborufus | Petaurista_alborufus | nocturnal |
| mammals | Petaurista_elegans | Petaurista_elegans | nocturnal |
| mammals | Petaurista_leucogenys | Petaurista_leucogenys | nocturnal |
| mammals | Petaurista_magnificus | Petaurista_magnificus | nocturnal |
| mammals | Petaurista_nobilis | Petaurista_nobilis | nocturnal |
| mammals | Petaurista_petaurista | Petaurista_petaurista | nocturnal |
| mammals | Petaurista_philippensis | Petaurista_philippensis | nocturnal |
| mammals | Petaurista_xanthotis | Petaurista_xanthotis | nocturnal |
| mammals | Petauroides_volans | Petauroides_volans | nocturnal |
| mammals | Petaurus_abidi | Petaurus_abidi | nocturnal |
| mammals | Petaurus_australis | Petaurus_australis | nocturnal |
| mammals | Petaurus_biacensis | Petaurus_biacensis | nocturnal |
| mammals | Petaurus_breviceps | Petaurus_breviceps | nocturnal |
| mammals | Petaurus_gracilis | Petaurus_gracilis | nocturnal |
| mammals | Petaurus_norfolcensis | Petaurus_norfolcensis | nocturnal |
| mammals | Petinomys_crinitus | Petinomys_crinitus | nocturnal |
| mammals | Petinomys_fuscocapillus | Petinomys_fuscocapillus | nocturnal |
| mammals | Petinomys_genibarbis | Petinomys_genibarbis | nocturnal |
| mammals | Petinomys_hageni | Petinomys_hageni | nocturnal |
| mammals | Petinomys_lugens | Petinomys_lugens | nocturnal |
| mammals | Petinomys_mindanensis | Petinomys_mindanensis | nocturnal |
| mammals | Petinomys_setosus | Petinomys_setosus | nocturnal |
| mammals | Petinomys_vordermanni | Petinomys_vordermanni | nocturnal |
| mammals | Petrodromus_tetradactylus | Petrodromus_tetradactylus | crepuscular |
| mammals | Petrogale_assimilis | Petrogale_assimilis | nocturnal |
| mammals | Petrogale_brachyotis | Petrogale_brachyotis | nocturnal |
| mammals | Petrogale_burbidgei | Petrogale_burbidgei | nocturnal |
| mammals | Petrogale_coenensis | Petrogale_coenensis | nocturnal |
| mammals | Petrogale_concinna | Petrogale_concinna | nocturnal |
| mammals | Petrogale_godmani | Petrogale_godmani | nocturnal |
| mammals | Petrogale_herberti | Petrogale_herberti | nocturnal |
| mammals | Petrogale_inornata | Petrogale_inornata | nocturnal |
| mammals | Petrogale_lateralis | Petrogale_lateralis | nocturnal |
| mammals | Petrogale_mareeba | Petrogale_mareeba | nocturnal |

| | | | |
|---|---|---|---|
| mammals | Petrogale_penicillata | Petrogale_penicillata | nocturnal |
| mammals | Petrogale_persephone | Petrogale_persephone | nocturnal |
| mammals | Petrogale_purpureicollis | Petrogale_purpureicollis | nocturnal |
| mammals | Petrogale_rothschildi | Petrogale_rothschildi | nocturnal |
| mammals | Petrogale_sharmani | Petrogale_sharmani | nocturnal |
| mammals | Petrogale_xanthopus | Petrogale_xanthopus | nocturnal |
| mammals | Petromus_typicus | Petromus_typicus | diurnal |
| mammals | Petromyscus_barbouri | Petromyscus_barbouri | nocturnal |
| mammals | Petromyscus_collinus | Petromyscus_collinus | nocturnal |
| mammals | Petromyscus_monticularis | Petromyscus_monticularis | nocturnal |
| mammals | Petromyscus_shortridgei | Petromyscus_shortridgei | nocturnal |
| mammals | Petrosaltator_rozeti | Elephantulus_rozeti | crepuscular |
| mammals | Phacochoerus_aethiopicus | Phacochoerus_aethiopicus | diurnal |
| mammals | Phacochoerus_africanus | Phacochoerus_africanus | cathemeral |
| mammals | Phaenomys_ferrugineus | Phaenomys_ferrugineus | nocturnal |
| mammals | Phalanger_carmelitae | Phalanger_carmelitae | nocturnal |
| mammals | Phalanger_gymnotis | Phalanger_gymnotis | nocturnal |
| mammals | Phalanger_intercastellanus | Phalanger_intercastellanus | nocturnal |
| mammals | Phalanger_lullulae | Phalanger_lullulae | nocturnal |
| mammals | Phalanger_matanim | Phalanger_matanim | nocturnal |
| mammals | Phalanger_mimicus | Phalanger_mimicus | nocturnal |
| mammals | Phalanger_orientalis | Phalanger_orientalis | nocturnal |
| mammals | Phalanger_ornatus | Phalanger_ornatus | nocturnal |
| mammals | Phalanger_pelengensis | Strigocuscus_pelengensis | nocturnal |
| mammals | Phalanger_rothschildi | Phalanger_rothschildi | nocturnal |
| mammals | Phalanger_sericeus | Phalanger_sericeus | nocturnal |
| mammals | Phalanger_vestitus | Phalanger_vestitus | nocturnal |
| mammals | Phaner_electromontis | Phaner_electromontis | nocturnal |
| mammals | Phaner_furcifer | Phaner_furcifer | nocturnal |
| mammals | Phaner_pallescens | Phaner_pallescens | nocturnal |
| mammals | Phaner_parienti | Phaner_parienti | nocturnal |
| mammals | Pharotis_imogene | Pharotis_imogene | nocturnal |
| mammals | Phascogale_calura | Phascogale_calura | nocturnal |
| mammals | Phascogale_pirata | Phascogale_pirata | nocturnal |

| | | | |
|---|---|---|---|
| mammals | Phascogale_tapoatafa | Phascogale_tapoatafa | nocturnal |
| mammals | Phascolarctos_cinereus | Phascolarctos_cinereus | nocturnal |
| mammals | Phascolosorex_doriae | Phascolosorex_doriae | diurnal |
| mammals | Phascolosorex_dorsalis | Phascolosorex_dorsalis | diurnal |
| mammals | Phataginus_tetradactylus | Phataginus_tetradactyla | diurnal |
| mammals | Phataginus_tricuspis | Phataginus_tricuspis | nocturnal |
| mammals | Phenacomys_intermedius | Phenacomys_intermedius | nocturnal |
| mammals | Phenacomys_ungava | Phenacomys_ungava | nocturnal |
| mammals | Philander_andersoni | Philander_andersoni | nocturnal |
| mammals | Philander_mcilhennyi | Philander_mcilhennyi | cathemeral |
| mammals | Philander_opossum | Philander_opossum | nocturnal |
| mammals | Philander_quica | Philander_frenatus | nocturnal |
| mammals | Philantomba_maxwellii | Philantomba_maxwellii | diurnal |
| mammals | Philantomba_monticola | Philantomba_monticola | diurnal |
| mammals | Philantomba_walteri | Philantomba_walteri | diurnal |
| mammals | Philetor_brachypterus | Philetor_brachypterus | nocturnal |
| mammals | Phloeomys_cumingi | Phloeomys_cumingi | nocturnal |
| mammals | Phloeomys_pallidus | Phloeomys_pallidus | nocturnal |
| mammals | Phodopus_campbelli | Phodopus_campbelli | nocturnal |
| mammals | Phodopus_roborovskii | Phodopus_roborovskii | nocturnal |
| mammals | Phodopus_sungorus | Phodopus_sungorus | nocturnal |
| mammals | Phoniscus_atrox | Phoniscus_atrox | nocturnal |
| mammals | Phoniscus_jagorii | Phoniscus_jagorii | nocturnal |
| mammals | Phoniscus_papuensis | Phoniscus_papuensis | nocturnal |
| mammals | Phylloderma_stenops | Phylloderma_stenops | nocturnal |
| mammals | Phyllomys_blainvilii | Phyllomys_blainvillii | nocturnal |
| mammals | Phyllomys_brasiliensis | Phyllomys_brasiliensis | nocturnal |
| mammals | Phyllomys_dasythrix | Phyllomys_dasythrix | nocturnal |
| mammals | Phyllomys_kerri | Phyllomys_kerri | nocturnal |
| mammals | Phyllomys_lamarum | Phyllomys_lamarum | nocturnal |
| mammals | Phyllomys_lundi | Phyllomys_lundi | nocturnal |
| mammals | Phyllomys_mantiqueirensis | Phyllomys_mantiqueirensis | nocturnal |
| mammals | Phyllomys_medius | Phyllomys_medius | nocturnal |
| mammals | Phyllomys_nigrispinus | Phyllomys_nigrispinus | nocturnal |

| | | | |
|---|---|---|---|
| mammals | Phyllomys_pattoni | Phyllomys_pattoni | nocturnal |
| mammals | Phyllomys_sulinus | Phyllomys_sulinus | nocturnal |
| mammals | Phyllomys_thomasi | Phyllomys_thomasi | nocturnal |
| mammals | Phyllomys_unicolor | Phyllomys_unicolor | nocturnal |
| mammals | Phyllonycteris_aphylla | Phyllonycteris_aphylla | nocturnal |
| mammals | Phyllonycteris_poeyi | Phyllonycteris_poeyi | nocturnal |
| mammals | Phyllops_falcatus | Phyllops_falcatus | nocturnal |
| mammals | Phyllostomus_discolor | Phyllostomus_discolor | nocturnal |
| mammals | Phyllostomus_elongatus | Phyllostomus_elongatus | nocturnal |
| mammals | Phyllostomus_hastatus | Phyllostomus_hastatus | nocturnal |
| mammals | Phyllostomus_latifolius | Phyllostomus_latifolius | nocturnal |
| mammals | Phyllotis_amicus | Phyllotis_amicus | nocturnal |
| mammals | Phyllotis_andium | Phyllotis_andium | nocturnal |
| mammals | Phyllotis_anitae | Phyllotis_anitae | nocturnal |
| mammals | Phyllotis_bonariensis | Phyllotis_bonariensis | nocturnal |
| mammals | Phyllotis_caprinus | Phyllotis_caprinus | cathemeral |
| mammals | Phyllotis_darwinii | Phyllotis_darwini | nocturnal |
| mammals | Phyllotis_definitus | Phyllotis_definitus | cathemeral |
| mammals | Phyllotis_gerbillus | Phyllotis_gerbillus | nocturnal |
| mammals | Phyllotis_haggardi | Phyllotis_haggardi | nocturnal |
| mammals | Phyllotis_limatus | Phyllotis_limatus | nocturnal |
| mammals | Phyllotis_magister | Phyllotis_magister | cathemeral |
| mammals | Phyllotis_osgoodi | Phyllotis_osgoodi | cathemeral |
| mammals | Phyllotis_osilae | Phyllotis_osilae | crepuscular |
| mammals | Phyllotis_xanthopygus | Phyllotis_xanthopygus | nocturnal |
| mammals | Piliocolobus_badius | Piliocolobus_badius | diurnal |
| mammals | Piliocolobus_gordonorum | Piliocolobus_gordonorum | diurnal |
| mammals | Piliocolobus_kirkii | Piliocolobus_kirkii | diurnal |
| mammals | Piliocolobus_oustaleti | Piliocolobus_oustaleti | diurnal |
| mammals | Piliocolobus_pennantii | Piliocolobus_pennantii | diurnal |
| mammals | Piliocolobus_preussi | Piliocolobus_preussi | diurnal |
| mammals | Piliocolobus_rufomitratus | Piliocolobus_rufomitratus | diurnal |
| mammals | Piliocolobus_tephrosceles | Piliocolobus_tephrosceles | diurnal |
| mammals | Pilonycteris_celebensis | Rousettus_celebensis | nocturnal |

| | | | |
|---|---|---|---|
| mammals | Pipanacoctomys_aureus | Tympanoctomys_aureus | nocturnal |
| mammals | Pipistrellus_abramus | Pipistrellus_abramus | nocturnal |
| mammals | Pipistrellus_adamsi | Pipistrellus_adamsi | nocturnal |
| mammals | Pipistrellus_aero | Pipistrellus_aero | nocturnal |
| mammals | Pipistrellus_angulatus | Pipistrellus_angulatus | nocturnal |
| mammals | Pipistrellus_ceylonicus | Pipistrellus_ceylonicus | nocturnal |
| mammals | Pipistrellus_collinus | Pipistrellus_collinus | nocturnal |
| mammals | Pipistrellus_coromandra | Pipistrellus_coromandra | nocturnal |
| mammals | Pipistrellus_endoi | Pipistrellus_endoi | nocturnal |
| mammals | Pipistrellus_hanaki | Pipistrellus_hanaki | nocturnal |
| mammals | Pipistrellus_hesperidus | Pipistrellus_hesperidus | nocturnal |
| mammals | Pipistrellus_inexspectatus | Pipistrellus_inexspectatus | nocturnal |
| mammals | Pipistrellus_javanicus | Pipistrellus_javanicus | nocturnal |
| mammals | Pipistrellus_kuhlii | Pipistrellus_kuhlii | nocturnal |
| mammals | Pipistrellus_maderensis | Pipistrellus_maderensis | nocturnal |
| mammals | Pipistrellus_minahassae | Pipistrellus_minahassae | nocturnal |
| mammals | Pipistrellus_nanulus | Pipistrellus_nanulus | nocturnal |
| mammals | Pipistrellus_nathusii | Pipistrellus_nathusii | nocturnal |
| mammals | Pipistrellus_papuanus | Pipistrellus_papuanus | nocturnal |
| mammals | Pipistrellus_paterculus | Pipistrellus_paterculus | nocturnal |
| mammals | Pipistrellus_permixtus | Pipistrellus_permixtus | nocturnal |
| mammals | Pipistrellus_pipistrellus | Pipistrellus_pipistrellus | nocturnal |
| mammals | Pipistrellus_pygmaeus | Pipistrellus_pygmaeus | nocturnal |
| mammals | Pipistrellus_raceyi | Pipistrellus_raceyi | nocturnal |
| mammals | Pipistrellus_rusticus | Pipistrellus_rusticus | nocturnal |
| mammals | Pipistrellus_stenopterus | Pipistrellus_stenopterus | nocturnal |
| mammals | Pipistrellus_wattsi | Pipistrellus_wattsi | nocturnal |
| mammals | Pipistrellus_westralis | Pipistrellus_westralis | nocturnal |
| mammals | Pithecheir_melanurus | Pithecheir_melanurus | nocturnal |
| mammals | Pithecheir_parvus | Pithecheir_parvus | nocturnal |
| mammals | Pithecia_aequatorialis | Pithecia_aequatorialis | diurnal |
| mammals | Pithecia_albicans | Pithecia_albicans | diurnal |
| mammals | Pithecia_chrysocephala | Pithecia_chrysocephala | diurnal |
| mammals | Pithecia_milleri | Pithecia_milleri | diurnal |

| | | | |
|---|---|---|---|
| mammals | Pithecia_pithecia | Pithecia_pithecia | diurnal |
| mammals | Pithecia_vanzolinii | Pithecia_vanzolinii | diurnal |
| mammals | Plagiodontia_aedium | Plagiodontia_aedium | nocturnal |
| mammals | Planigale_gilesi | Planigale_gilesi | crepuscular |
| mammals | Planigale_ingrami | Planigale_ingrami | nocturnal |
| mammals | Planigale_maculata | Planigale_maculata | nocturnal |
| mammals | Planigale_novaeguineae | Planigale_novaeguineae | nocturnal |
| mammals | Planigale_tenuirostris | Planigale_tenuirostris | nocturnal |
| mammals | Platacanthomys_lasiurus | Platacanthomys_lasiurus | nocturnal |
| mammals | Platalina_genovensium | Platalina_genovensium | nocturnal |
| mammals | Platymops_setiger | Platymops_setiger | nocturnal |
| mammals | Platyrrhinus_albericoi | Platyrrhinus_albericoi | nocturnal |
| mammals | Platyrrhinus_aquilus | Platyrrhinus_aquilus | nocturnal |
| mammals | Platyrrhinus_aurarius | Platyrrhinus_aurarius | nocturnal |
| mammals | Platyrrhinus_brachycephalus | Platyrrhinus_brachycephalus | nocturnal |
| mammals | Platyrrhinus_dorsalis | Platyrrhinus_dorsalis | nocturnal |
| mammals | Platyrrhinus_helleri | Platyrrhinus_helleri | nocturnal |
| mammals | Platyrrhinus_infuscus | Platyrrhinus_infuscus | nocturnal |
| mammals | Platyrrhinus_ismaeli | Platyrrhinus_ismaeli | nocturnal |
| mammals | Platyrrhinus_lineatus | Platyrrhinus_lineatus | nocturnal |
| mammals | Platyrrhinus_masu | Platyrrhinus_masu | nocturnal |
| mammals | Platyrrhinus_matapalensis | Platyrrhinus_matapalensis | nocturnal |
| mammals | Platyrrhinus_recifinus | Platyrrhinus_recifinus | nocturnal |
| mammals | Platyrrhinus_umbratus | Platyrrhinus_umbratus | nocturnal |
| mammals | Platyrrhinus_vittatus | Platyrrhinus_vittatus | nocturnal |
| mammals | Plecotus_auritus | Plecotus_auritus | nocturnal |
| mammals | Plecotus_austriacus | Plecotus_austriacus | nocturnal |
| mammals | Plecotus_balensis | Plecotus_balensis | nocturnal |
| mammals | Plecotus_christii | Plecotus_christii | nocturnal |
| mammals | Plecotus_kolombatovici | Plecotus_kolombatovici | nocturnal |
| mammals | Plecotus_macrobullaris | Plecotus_macrobullaris | nocturnal |
| mammals | Plecotus_ognevi | Plecotus_ognevi | nocturnal |
| mammals | Plecotus_sacrimontis | Plecotus_sacrimontis | nocturnal |
| mammals | Plecotus_sardus | Plecotus_sardus | nocturnal |

| | | | |
|---|---|---|---|
| mammals | Plecotus_taivanus | Plecotus_taivanus | nocturnal |
| mammals | Plecotus_teneriffae | Plecotus_teneriffae | nocturnal |
| mammals | Plecturocebus_aureipalatii | Callicebus_aureipalatii | diurnal |
| mammals | Plecturocebus_baptista | Callicebus_baptista | diurnal |
| mammals | Plecturocebus_bernhardi | Callicebus_bernhardi | diurnal |
| mammals | Plecturocebus_brunneus | Callicebus_brunneus | diurnal |
| mammals | Plecturocebus_caligatus | Callicebus_caligatus | diurnal |
| mammals | Plecturocebus_cinerascens | Callicebus_cinerascens | diurnal |
| mammals | Plecturocebus_cupreus | Callicebus_cupreus | diurnal |
| mammals | Plecturocebus_discolor | Callicebus_discolor | diurnal |
| mammals | Plecturocebus_donacophilus | Callicebus_donacophilus | diurnal |
| mammals | Plecturocebus_hoffmannsi | Callicebus_hoffmannsi | diurnal |
| mammals | Plecturocebus_modestus | Callicebus_modestus | diurnal |
| mammals | Plecturocebus_moloch | Callicebus_moloch | diurnal |
| mammals | Plecturocebus_oenanthe | Callicebus_oenanthe | diurnal |
| mammals | Plecturocebus_olallae | Callicebus_olallae | diurnal |
| mammals | Plecturocebus_ornatus | Callicebus_ornatus | diurnal |
| mammals | Plecturocebus_pallescens | Callicebus_pallescens | diurnal |
| mammals | Plecturocebus_stephennashi | Callicebus_stephennashi | diurnal |
| mammals | Plerotes_anchietae | Plerotes_anchietae | nocturnal |
| mammals | Podogymnura_aureospinula | Podogymnura_aureospinula | nocturnal |
| mammals | Podogymnura_truei | Podogymnura_truei | nocturnal |
| mammals | Podomys_floridanus | Podomys_floridanus | nocturnal |
| mammals | Podoxymys_roraimae | Podoxymys_roraimae | nocturnal |
| mammals | Poecilictis_libyca | Ictonyx_libycus | nocturnal |
| mammals | Poecilogale_albinucha | Poecilogale_albinucha | nocturnal |
| mammals | Poelagus_marjorita | Poelagus_marjorita | nocturnal |
| mammals | Poemys_melanotis | Dendromus_melanotis | nocturnal |
| mammals | Poemys_nyikae | Dendromus_nyikae | nocturnal |
| mammals | Pogonomelomys_bruijnii | Pogonomelomys_bruijnii | nocturnal |
| mammals | Pogonomelomys_mayeri | Pogonomelomys_mayeri | nocturnal |
| mammals | Pogonomys_championi | Pogonomys_championi | nocturnal |
| mammals | Pogonomys_fergussoniensis | Pogonomys_fergussoniensis | nocturnal |
| mammals | Pogonomys_loriae | Pogonomys_loriae | nocturnal |

| | | | |
|---|---|---|---|
| mammals | Pogonomys_macrourus | Pogonomys_macrourus | nocturnal |
| mammals | Pogonomys_sylvestris | Pogonomys_sylvestris | nocturnal |
| mammals | Poiana_leightoni | Poiana_leightoni | nocturnal |
| mammals | Poiana_richardsonii | Poiana_richardsonii | nocturnal |
| mammals | Poliocitellus_franklinii | Poliocitellus_franklinii | diurnal |
| mammals | Pongo_abelii | Pongo_abelii | diurnal |
| mammals | Pongo_pygmaeus | Pongo_pygmaeus | diurnal |
| mammals | Porcula_salvania | Porcula_salvania | diurnal |
| mammals | Potamochoerus_larvatus | Potamochoerus_larvatus | nocturnal |
| mammals | Potamochoerus_porcus | Potamochoerus_porcus | nocturnal |
| mammals | Potamogale_velox | Potamogale_velox | nocturnal |
| mammals | Potorous_longipes | Potorous_longipes | nocturnal |
| mammals | Potorous_tridactylus | Potorous_tridactylus | nocturnal |
| mammals | Potos_flavus | Potos_flavus | nocturnal |
| mammals | Praomys_daltoni | Praomys_daltoni | nocturnal |
| mammals | Praomys_degraaffi | Praomys_degraaffi | nocturnal |
| mammals | Praomys_hartwigi | Praomys_hartwigi | nocturnal |
| mammals | Praomys_jacksoni | Praomys_jacksoni | nocturnal |
| mammals | Praomys_minor | Praomys_minor | nocturnal |
| mammals | Praomys_misonnei | Praomys_misonnei | nocturnal |
| mammals | Praomys_morio | Praomys_morio | nocturnal |
| mammals | Praomys_mutoni | Praomys_mutoni | nocturnal |
| mammals | Praomys_obscurus | Praomys_obscurus | nocturnal |
| mammals | Praomys_petteri | Praomys_petteri | nocturnal |
| mammals | Praomys_rostratus | Praomys_rostratus | nocturnal |
| mammals | Praomys_tullbergi | Praomys_tullbergi | nocturnal |
| mammals | Presbytis_chrysomelas | Presbytis_chrysomelas | diurnal |
| mammals | Presbytis_comata | Presbytis_comata | diurnal |
| mammals | Presbytis_femoralis | Presbytis_femoralis | diurnal |
| mammals | Presbytis_frontata | Presbytis_frontata | diurnal |
| mammals | Presbytis_hosei | Presbytis_hosei | diurnal |
| mammals | Presbytis_melalophos | Presbytis_melalophos | diurnal |
| mammals | Presbytis_natunae | Presbytis_natunae | diurnal |
| mammals | Presbytis_potenziani | Presbytis_potenziani | diurnal |

| | | | |
|---|---|---|---|
| mammals | Presbytis_rubicunda | Presbytis_rubicunda | diurnal |
| mammals | Presbytis_siamensis | Presbytis_siamensis | diurnal |
| mammals | Presbytis_thomasi | Presbytis_thomasi | diurnal |
| mammals | Priodontes_maximus | Priodontes_maximus | nocturnal |
| mammals | Prionailurus_bengalensis | Prionailurus_bengalensis | cathemeral |
| mammals | Prionailurus_planiceps | Prionailurus_planiceps | nocturnal |
| mammals | Prionailurus_rubiginosus | Prionailurus_rubiginosus | nocturnal |
| mammals | Prionailurus_viverrinus | Prionailurus_viverrinus | nocturnal |
| mammals | Prionodon_linsang | Prionodon_linsang | nocturnal |
| mammals | Prionodon_pardicolor | Prionodon_pardicolor | nocturnal |
| mammals | Prionomys_batesi | Prionomys_batesi | nocturnal |
| mammals | Procapra_gutturosa | Procapra_gutturosa | diurnal |
| mammals | Procapra_picticaudata | Procapra_picticaudata | crepuscular |
| mammals | Procapra_przewalskii | Procapra_przewalskii | diurnal |
| mammals | Procavia_capensis | Procavia_capensis | diurnal |
| mammals | Procolobus_verus | Procolobus_verus | diurnal |
| mammals | Procyon_cancrivorus | Procyon_cancrivorus | nocturnal |
| mammals | Procyon_lotor | Procyon_lotor | cathemeral |
| mammals | Procyon_pygmaeus | Procyon_pygmaeus | nocturnal |
| mammals | Proechimys_brevicauda | Proechimys_brevicauda | nocturnal |
| mammals | Proechimys_canicollis | Proechimys_canicollis | nocturnal |
| mammals | Proechimys_chrysaeolus | Proechimys_chrysaeolus | nocturnal |
| mammals | Proechimys_cuvieri | Proechimys_cuvieri | nocturnal |
| mammals | Proechimys_decumanus | Proechimys_decumanus | nocturnal |
| mammals | Proechimys_echinothrix | Proechimys_echinothrix | nocturnal |
| mammals | Proechimys_gardneri | Proechimys_gardneri | nocturnal |
| mammals | Proechimys_goeldii | Proechimys_goeldii | nocturnal |
| mammals | Proechimys_guairae | Proechimys_guairae | nocturnal |
| mammals | Proechimys_guyannensis | Proechimys_guyannensis | nocturnal |
| mammals | Proechimys_hoplomyoides | Proechimys_hoplomyoides | nocturnal |
| mammals | Proechimys_kulinae | Proechimys_kulinae | nocturnal |
| mammals | Proechimys_longicaudatus | Proechimys_longicaudatus | nocturnal |
| mammals | Proechimys_mincae | Proechimys_mincae | nocturnal |
| mammals | Proechimys_oconnelli | Proechimys_oconnelli | nocturnal |

| | | | |
|---|---|---|---|
| mammals | Proechimys_pattoni | Proechimys_pattoni | nocturnal |
| mammals | Proechimys_quadruplicatus | Proechimys_quadruplicatus | nocturnal |
| mammals | Proechimys_roberti | Proechimys_roberti | nocturnal |
| mammals | Proechimys_semispinosus | Proechimys_semispinosus | nocturnal |
| mammals | Proechimys_simonsi | Proechimys_simonsi | nocturnal |
| mammals | Proechimys_steerei | Proechimys_steerei | nocturnal |
| mammals | Proechimys_trinitatis | Proechimys_trinitatis | nocturnal |
| mammals | Prolemur_simus | Prolemur_simus | cathemeral |
| mammals | Promops_centralis | Promops_centralis | nocturnal |
| mammals | Promops_davisoni | Promops_davisoni | nocturnal |
| mammals | Promops_nasutus | Promops_nasutus | nocturnal |
| mammals | Pronolagus_crassicaudatus | Pronolagus_crassicaudatus | nocturnal |
| mammals | Pronolagus_randensis | Pronolagus_randensis | nocturnal |
| mammals | Pronolagus_rupestris | Pronolagus_rupestris | nocturnal |
| mammals | Propithecus_candidus | Propithecus_candidus | diurnal |
| mammals | Propithecus_coquereli | Propithecus_coquereli | diurnal |
| mammals | Propithecus_coronatus | Propithecus_coronatus | diurnal |
| mammals | Propithecus_deckenii | Propithecus_deckenii | diurnal |
| mammals | Propithecus_diadema | Propithecus_diadema | diurnal |
| mammals | Propithecus_edwardsi | Propithecus_edwardsi | diurnal |
| mammals | Propithecus_perrieri | Propithecus_perrieri | diurnal |
| mammals | Propithecus_tattersalli | Propithecus_tattersalli | diurnal |
| mammals | Propithecus_verreauxi | Propithecus_verreauxi | diurnal |
| mammals | Prosciurillus_abstrusus | Prosciurillus_abstrusus | diurnal |
| mammals | Prosciurillus_leucomus | Prosciurillus_leucomus | diurnal |
| mammals | Prosciurillus_murinus | Prosciurillus_murinus | diurnal |
| mammals | Prosciurillus_rosenbergii | Prosciurillus_rosenbergii | diurnal |
| mammals | Prosciurillus_weberi | Prosciurillus_weberi | diurnal |
| mammals | Proteles_cristatus | Proteles_cristata | nocturnal |
| mammals | Protoxerus_aubinnii | Protoxerus_aubinnii | diurnal |
| mammals | Protoxerus_stangeri | Protoxerus_stangeri | diurnal |
| mammals | Psammomys_obesus | Psammomys_obesus | cathemeral |
| mammals | Pseudantechinus_bilarni | Pseudantechinus_bilarni | nocturnal |
| mammals | Pseudantechinus_macdonnellensis | Pseudantechinus_macdonnellensis | nocturnal |

| | | | |
|---|---|---|---|
| mammals | Pseudantechinus_mimulus | Pseudantechinus_mimulus | nocturnal |
| mammals | Pseudantechinus_ningbing | Pseudantechinus_ningbing | nocturnal |
| mammals | Pseudantechinus_woolleyae | Pseudantechinus_woolleyae | nocturnal |
| mammals | Pseudocheirus_occidentalis | Pseudocheirus_occidentalis | nocturnal |
| mammals | Pseudocheirus_peregrinus | Pseudocheirus_peregrinus | nocturnal |
| mammals | Pseudochirops_albertisii | Pseudochirops_albertisii | nocturnal |
| mammals | Pseudochirops_archeri | Pseudochirops_archeri | nocturnal |
| mammals | Pseudochirops_corinnae | Pseudochirops_corinnae | nocturnal |
| mammals | Pseudochirops_coronatus | Pseudochirops_coronatus | nocturnal |
| mammals | Pseudochirops_cupreus | Pseudochirops_cupreus | nocturnal |
| mammals | Pseudochirops_dahlii | Petropseudes_dahli | nocturnal |
| mammals | Pseudochirulus_canescens | Pseudochirulus_canescens | nocturnal |
| mammals | Pseudochirulus_caroli | Pseudochirulus_caroli | nocturnal |
| mammals | Pseudochirulus_cinereus | Pseudochirulus_cinereus | nocturnal |
| mammals | Pseudochirulus_forbesi | Pseudochirulus_forbesi | nocturnal |
| mammals | Pseudochirulus_herbertensis | Pseudochirulus_herbertensis | nocturnal |
| mammals | Pseudochirulus_larvatus | Pseudochirulus_larvatus | nocturnal |
| mammals | Pseudochirulus_mayeri | Pseudochirulus_mayeri | nocturnal |
| mammals | Pseudochirulus_schlegelii | Pseudochirulus_schlegeli | nocturnal |
| mammals | Pseudohydromys_ellermani | Pseudohydromys_ellermani | nocturnal |
| mammals | Pseudohydromys_fuscus | Pseudohydromys_fuscus | nocturnal |
| mammals | Pseudois_nayaur | Pseudois_nayaur | diurnal |
| mammals | Pseudomys_albocinereus | Pseudomys_albocinereus | nocturnal |
| mammals | Pseudomys_apodemoides | Pseudomys_apodemoides | nocturnal |
| mammals | Pseudomys_australis | Pseudomys_australis | nocturnal |
| mammals | Pseudomys_bolami | Pseudomys_bolami | nocturnal |
| mammals | Pseudomys_calabyi | Pseudomys_calabyi | nocturnal |
| mammals | Pseudomys_chapmani | Pseudomys_chapmani | nocturnal |
| mammals | Pseudomys_delicatulus | Pseudomys_delicatulus | nocturnal |
| mammals | Pseudomys_desertor | Pseudomys_desertor | diurnal |
| mammals | Pseudomys_fumeus | Pseudomys_fumeus | nocturnal |
| mammals | Pseudomys_gouldii | Pseudomys_fieldi | nocturnal |
| mammals | Pseudomys_gracilicaudatus | Pseudomys_gracilicaudatus | nocturnal |
| mammals | Pseudomys_hermannsburgensis | Pseudomys_hermannsburgensis | nocturnal |

| | | | |
|---|---|---|---|
| mammals | Pseudomys_higginsi | Pseudomys_higginsi | nocturnal |
| mammals | Pseudomys_johnsoni | Pseudomys_johnsoni | nocturnal |
| mammals | Pseudomys_nanus | Pseudomys_nanus | nocturnal |
| mammals | Pseudomys_novaehollandiae | Pseudomys_novaehollandiae | nocturnal |
| mammals | Pseudomys_occidentalis | Pseudomys_occidentalis | nocturnal |
| mammals | Pseudomys_oralis | Pseudomys_oralis | nocturnal |
| mammals | Pseudomys_patrius | Pseudomys_patrius | nocturnal |
| mammals | Pseudomys_shortridgei | Pseudomys_shortridgei | diurnal |
| mammals | Pseudoromicia_brunnea | Neoromicia_brunnea | nocturnal |
| mammals | Pseudoromicia_rendalli | Neoromicia_rendalli | nocturnal |
| mammals | Pseudoromicia_tenuipinnis | Neoromicia_tenuipinnis | nocturnal |
| mammals | Pseudoryx_nghetinhensis | Pseudoryx_nghetinhensis | diurnal |
| mammals | Pseudoryzomys_simplex | Pseudoryzomys_simplex | nocturnal |
| mammals | Pseudosoriculus_fumidus | Episoriculus_fumidus | cathemeral |
| mammals | Ptenochirus_jagorii | Ptenochirus_jagori | nocturnal |
| mammals | Ptenochirus_minor | Ptenochirus_minor | nocturnal |
| mammals | Ptenochirus_wetmorei | Megaerops_wetmorei | nocturnal |
| mammals | Pteralopex_anceps | Pteralopex_anceps | nocturnal |
| mammals | Pteralopex_atrata | Pteralopex_atrata | nocturnal |
| mammals | Pteralopex_flanneryi | Pteralopex_flanneryi | nocturnal |
| mammals | Pteralopex_taki | Pteralopex_taki | nocturnal |
| mammals | Pteromys_momonga | Pteromys_momonga | nocturnal |
| mammals | Pteromys_volans | Pteromys_volans | nocturnal |
| mammals | Pteromyscus_pulverulentus | Pteromyscus_pulverulentus | nocturnal |
| mammals | Pteronotus_davyi | Pteronotus_davyi | nocturnal |
| mammals | Pteronotus_gymnonotus | Pteronotus_gymnonotus | nocturnal |
| mammals | Pteronotus_macleayii | Pteronotus_macleayii | nocturnal |
| mammals | Pteronotus_mesoamericanus | Pteronotus_mesoamericanus | nocturnal |
| mammals | Pteronotus_paraguanensis | Pteronotus_paraguanensis | nocturnal |
| mammals | Pteronotus_parnellii | Pteronotus_parnellii | nocturnal |
| mammals | Pteronotus_personatus | Pteronotus_personatus | nocturnal |
| mammals | Pteronotus_quadridens | Pteronotus_quadridens | nocturnal |
| mammals | Pteronotus_rubiginosus | Pteronotus_rubiginosus | nocturnal |
| mammals | Pteronura_brasiliensis | Pteronura_brasiliensis | diurnal |

| | | | |
|---|---|---|---|
| mammals | Pteropus_admiralitatum | Pteropus_admiralitatum | nocturnal |
| mammals | Pteropus_aldabrensis | Pteropus_aldabrensis | nocturnal |
| mammals | Pteropus_alecto | Pteropus_alecto | nocturnal |
| mammals | Pteropus_anetianus | Pteropus_anetianus | nocturnal |
| mammals | Pteropus_aruensis | Pteropus_aruensis | nocturnal |
| mammals | Pteropus_caniceps | Pteropus_caniceps | nocturnal |
| mammals | Pteropus_capistratus | Pteropus_capistratus | nocturnal |
| mammals | Pteropus_chrysoproctus | Pteropus_chrysoproctus | nocturnal |
| mammals | Pteropus_cognatus | Pteropus_cognatus | nocturnal |
| mammals | Pteropus_conspicillatus | Pteropus_conspicillatus | nocturnal |
| mammals | Pteropus_dasymallus | Pteropus_dasymallus | nocturnal |
| mammals | Pteropus_faunulus | Pteropus_faunulus | nocturnal |
| mammals | Pteropus_fundatus | Pteropus_fundatus | nocturnal |
| mammals | Pteropus_gilliardi | Pteropus_gilliardorum | nocturnal |
| mammals | Pteropus_griseus | Pteropus_griseus | nocturnal |
| mammals | Pteropus_hypomelanus | Pteropus_hypomelanus | nocturnal |
| mammals | Pteropus_intermedius | Pteropus_intermedius | nocturnal |
| mammals | Pteropus_keyensis | Pteropus_keyensis | nocturnal |
| mammals | Pteropus_livingstonii | Pteropus_livingstonii | nocturnal |
| mammals | Pteropus_lombocensis | Pteropus_lombocensis | nocturnal |
| mammals | Pteropus_lylei | Pteropus_lylei | nocturnal |
| mammals | Pteropus_macrotis | Pteropus_macrotis | nocturnal |
| mammals | Pteropus_mahaganus | Pteropus_mahaganus | nocturnal |
| mammals | Pteropus_mariannus | Pteropus_mariannus | nocturnal |
| mammals | Pteropus_melanopogon | Pteropus_melanopogon | nocturnal |
| mammals | Pteropus_melanotus | Pteropus_melanotus | diurnal |
| mammals | Pteropus_molossinus | Pteropus_molossinus | diurnal |
| mammals | Pteropus_neohibernicus | Pteropus_neohibernicus | nocturnal |
| mammals | Pteropus_niger | Pteropus_niger | nocturnal |
| mammals | Pteropus_nitendiensis | Pteropus_nitendiensis | nocturnal |
| mammals | Pteropus_ocularis | Pteropus_ocularis | nocturnal |
| mammals | Pteropus_ornatus | Pteropus_ornatus | nocturnal |
| mammals | Pteropus_personatus | Pteropus_personatus | nocturnal |
| mammals | Pteropus_pohlei | Pteropus_pohlei | nocturnal |

| | | | |
|---|---|---|---|
| mammals | Pteropus_poliocephalus | Pteropus_poliocephalus | nocturnal |
| mammals | Pteropus_pselaphon | Pteropus_pselaphon | nocturnal |
| mammals | Pteropus_pumilus | Pteropus_pumilus | nocturnal |
| mammals | Pteropus_rayneri | Pteropus_rayneri | nocturnal |
| mammals | Pteropus_rennelli | Pteropus_rennelli | nocturnal |
| mammals | Pteropus_rodricensis | Pteropus_rodricensis | nocturnal |
| mammals | Pteropus_rufus | Pteropus_rufus | nocturnal |
| mammals | Pteropus_samoensis | Pteropus_samoensis | cathemeral |
| mammals | Pteropus_scapulatus | Pteropus_scapulatus | nocturnal |
| mammals | Pteropus_seychellensis | Pteropus_seychellensis | nocturnal |
| mammals | Pteropus_speciosus | Pteropus_speciosus | nocturnal |
| mammals | Pteropus_temminckii | Pteropus_temminckii | nocturnal |
| mammals | Pteropus_tonganus | Pteropus_tonganus | nocturnal |
| mammals | Pteropus_tuberculatus | Pteropus_tuberculatus | nocturnal |
| mammals | Pteropus_ualanus | Pteropus_ualanus | nocturnal |
| mammals | Pteropus_vampyrus | Pteropus_vampyrus | nocturnal |
| mammals | Pteropus_vetulus | Pteropus_vetulus | nocturnal |
| mammals | Pteropus_voeltzkowi | Pteropus_voeltzkowi | nocturnal |
| mammals | Pteropus_woodfordi | Pteropus_woodfordi | nocturnal |
| mammals | Ptilocercus_lowii | Ptilocercus_lowii | nocturnal |
| mammals | Pudu_mephistophiles | Pudu_mephistophiles | nocturnal |
| mammals | Pudu_puda | Pudu_puda | crepuscular |
| mammals | Puma_concolor | Puma_concolor | cathemeral |
| mammals | Punomys_kofordi | Punomys_kofordi | cathemeral |
| mammals | Punomys_lemminus | Punomys_lemminus | diurnal |
| mammals | Pygathrix_cinerea | Pygathrix_cinerea | diurnal |
| mammals | Pygathrix_nemaeus | Pygathrix_nemaeus | diurnal |
| mammals | Pygathrix_nigripes | Pygathrix_nigripes | diurnal |
| mammals | Pygeretmus_platyurus | Pygeretmus_platyurus | nocturnal |
| mammals | Pygeretmus_pumilio | Pygeretmus_pumilio | nocturnal |
| mammals | Pygeretmus_shitkovi | Pygeretmus_zhitkovi | nocturnal |
| mammals | Pygoderma_bilabiatum | Pygoderma_bilabiatum | nocturnal |
| mammals | Rangifer_tarandus | Rangifer_tarandus | cathemeral |
| mammals | Raphicerus_campestris | Raphicerus_campestris | cathemeral |

| | | | |
|---|---|---|---|
| mammals | Raphicerus_melanotis | Raphicerus_melanotis | nocturnal |
| mammals | Raphicerus_sharpei | Raphicerus_sharpei | nocturnal |
| mammals | Rattus_adustus | Rattus_adustus | nocturnal |
| mammals | Rattus_andamanensis | Rattus_andamanensis | nocturnal |
| mammals | Rattus_argentiventer | Rattus_argentiventer | nocturnal |
| mammals | Rattus_baluensis | Rattus_baluensis | nocturnal |
| mammals | Rattus_blangorum | Rattus_blangorum | nocturnal |
| mammals | Rattus_bontanus | Rattus_bontanus | nocturnal |
| mammals | Rattus_ceramicus | Nesoromys_ceramicus | nocturnal |
| mammals | Rattus_colletti | Rattus_colletti | nocturnal |
| mammals | Rattus_elaphinus | Rattus_elaphinus | nocturnal |
| mammals | Rattus_enganus | Rattus_enganus | nocturnal |
| mammals | Rattus_everetti | Rattus_everetti | nocturnal |
| mammals | Rattus_exulans | Rattus_exulans | nocturnal |
| mammals | Rattus_feliceus | Rattus_feliceus | nocturnal |
| mammals | Rattus_fuscipes | Rattus_fuscipes | nocturnal |
| mammals | Rattus_giluwensis | Rattus_giluwensis | nocturnal |
| mammals | Rattus_hainaldi | Rattus_hainaldi | nocturnal |
| mammals | Rattus_hoffmanni | Rattus_hoffmanni | nocturnal |
| mammals | Rattus_hoogerwerfi | Rattus_hoogerwerfi | nocturnal |
| mammals | Rattus_jobiensis | Rattus_jobiensis | nocturnal |
| mammals | Rattus_korinchi | Rattus_korinchi | nocturnal |
| mammals | Rattus_leucopus | Rattus_leucopus | nocturnal |
| mammals | Rattus_losea | Rattus_losea | nocturnal |
| mammals | Rattus_lugens | Rattus_lugens | nocturnal |
| mammals | Rattus_lutreola | Rattus_lutreolus | cathemeral |
| mammals | Rattus_marmosurus | Rattus_marmosurus | nocturnal |
| mammals | Rattus_mindorensis | Rattus_mindorensis | nocturnal |
| mammals | Rattus_montanus | Rattus_montanus | diurnal |
| mammals | Rattus_mordax | Rattus_mordax | nocturnal |
| mammals | Rattus_morotaiensis | Rattus_morotaiensis | nocturnal |
| mammals | Rattus_niobe | Rattus_niobe | nocturnal |
| mammals | Rattus_nitidus | Rattus_nitidus | cathemeral |
| mammals | Rattus_norvegicus | Rattus_norvegicus | nocturnal |

| | | | |
|---|---|---|---|
| mammals | Rattus_novaeguineae | Rattus_novaeguineae | nocturnal |
| mammals | Rattus_omichlodes | Rattus_omichlodes | nocturnal |
| mammals | Rattus_osgoodi | Rattus_osgoodi | nocturnal |
| mammals | Rattus_palmarum | Rattus_palmarum | nocturnal |
| mammals | Rattus_pelurus | Rattus_pelurus | nocturnal |
| mammals | Rattus_praetor | Rattus_praetor | nocturnal |
| mammals | Rattus_pyctoris | Rattus_pyctoris | nocturnal |
| mammals | Rattus_rattus | Rattus_rattus | nocturnal |
| mammals | Rattus_richardsoni | Rattus_richardsoni | nocturnal |
| mammals | Rattus_salocco | Rattus_salocco | nocturnal |
| mammals | Rattus_satarae | Rattus_satarae | nocturnal |
| mammals | Rattus_sordidus | Rattus_sordidus | nocturnal |
| mammals | Rattus_steini | Rattus_steini | nocturnal |
| mammals | Rattus_stoicus | Rattus_stoicus | nocturnal |
| mammals | Rattus_tanezumi | Rattus_tanezumi | nocturnal |
| mammals | Rattus_tawitawiensis | Rattus_tawitawiensis | nocturnal |
| mammals | Rattus_timorensis | Rattus_timorensis | nocturnal |
| mammals | Rattus_tiomanicus | Rattus_tiomanicus | nocturnal |
| mammals | Rattus_tunneyi | Rattus_tunneyi | nocturnal |
| mammals | Rattus_vandeuseni | Rattus_vandeuseni | nocturnal |
| mammals | Rattus_verecundus | Rattus_verecundus | nocturnal |
| mammals | Rattus_villosissimus | Rattus_villosissimus | nocturnal |
| mammals | Rattus_xanthurus | Rattus_xanthurus | nocturnal |
| mammals | Ratufa_affinis | Ratufa_affinis | diurnal |
| mammals | Ratufa_bicolor | Ratufa_bicolor | diurnal |
| mammals | Ratufa_indica | Ratufa_indica | diurnal |
| mammals | Ratufa_macroura | Ratufa_macroura | diurnal |
| mammals | Redunca_arundinum | Redunca_arundinum | cathemeral |
| mammals | Redunca_fulvorufula | Redunca_fulvorufula | cathemeral |
| mammals | Redunca_redunca | Redunca_redunca | cathemeral |
| mammals | Reithrodon_auritus | Reithrodon_auritus | nocturnal |
| mammals | Reithrodon_typicus | Reithrodon_typicus | nocturnal |
| mammals | Reithrodontomys_brevirostris | Reithrodontomys_brevirostris | nocturnal |
| mammals | Reithrodontomys_burti | Reithrodontomys_burti | nocturnal |

| | | | |
|---|---|---|---|
| mammals | Reithrodontomys_chrysopsis | Reithrodontomys_chrysopsis | nocturnal |
| mammals | Reithrodontomys_creper | Reithrodontomys_creper | nocturnal |
| mammals | Reithrodontomys_darienensis | Reithrodontomys_darienensis | nocturnal |
| mammals | Reithrodontomys_fulvescens | Reithrodontomys_fulvescens | nocturnal |
| mammals | Reithrodontomys_gracilis | Reithrodontomys_gracilis | nocturnal |
| mammals | Reithrodontomys_hirsutus | Reithrodontomys_hirsutus | nocturnal |
| mammals | Reithrodontomys_humulis | Reithrodontomys_humulis | nocturnal |
| mammals | Reithrodontomys_megalotis | Reithrodontomys_megalotis | nocturnal |
| mammals | Reithrodontomys_mexicanus | Reithrodontomys_mexicanus | nocturnal |
| mammals | Reithrodontomys_microdon | Reithrodontomys_microdon | nocturnal |
| mammals | Reithrodontomys_montanus | Reithrodontomys_montanus | nocturnal |
| mammals | Reithrodontomys_paradoxus | Reithrodontomys_paradoxus | nocturnal |
| mammals | Reithrodontomys_raviventris | Reithrodontomys_raviventris | nocturnal |
| mammals | Reithrodontomys_rodriguezi | Reithrodontomys_rodriguezi | nocturnal |
| mammals | Reithrodontomys_spectabilis | Reithrodontomys_spectabilis | nocturnal |
| mammals | Reithrodontomys_sumichrasti | Reithrodontomys_sumichrasti | nocturnal |
| mammals | Reithrodontomys_tenuirostris | Reithrodontomys_tenuirostris | nocturnal |
| mammals | Reithrodontomys_zacatecae | Reithrodontomys_zacatecae | nocturnal |
| mammals | Rhabdomys_pumilio | Rhabdomys_pumilio | diurnal |
| mammals | Rhagomys_longilingua | Rhagomys_longilingua | nocturnal |
| mammals | Rheithrosciurus_macrotis | Rheithrosciurus_macrotis | diurnal |
| mammals | Rheomys_mexicanus | Rheomys_mexicanus | cathemeral |
| mammals | Rheomys_raptor | Rheomys_raptor | cathemeral |
| mammals | Rheomys_thomasi | Rheomys_thomasi | cathemeral |
| mammals | Rheomys_underwoodi | Rheomys_underwoodi | cathemeral |
| mammals | Rhinoceros_sondaicus | Rhinoceros_sondaicus | cathemeral |
| mammals | Rhinoceros_unicornis | Rhinoceros_unicornis | cathemeral |
| mammals | Rhinolophus_acuminatus | Rhinolophus_acuminatus | nocturnal |
| mammals | Rhinolophus_adami | Rhinolophus_adami | nocturnal |
| mammals | Rhinolophus_affinis | Rhinolophus_affinis | nocturnal |
| mammals | Rhinolophus_alcyone | Rhinolophus_alcyone | nocturnal |
| mammals | Rhinolophus_arcuatus | Rhinolophus_arcuatus | nocturnal |
| mammals | Rhinolophus_beddomei | Rhinolophus_beddomei | nocturnal |
| mammals | Rhinolophus_blasii | Rhinolophus_blasii | nocturnal |

| | | | |
|---|---|---|---|
| mammals | Rhinolophus_bocharicus | Rhinolophus_bocharicus | nocturnal |
| mammals | Rhinolophus_borneensis | Rhinolophus_borneensis | nocturnal |
| mammals | Rhinolophus_canuti | Rhinolophus_canuti | nocturnal |
| mammals | Rhinolophus_capensis | Rhinolophus_capensis | nocturnal |
| mammals | Rhinolophus_celebensis | Rhinolophus_celebensis | nocturnal |
| mammals | Rhinolophus_clivosus | Rhinolophus_clivosus | nocturnal |
| mammals | Rhinolophus_coelophyllus | Rhinolophus_coelophyllus | nocturnal |
| mammals | Rhinolophus_cognatus | Rhinolophus_cognatus | nocturnal |
| mammals | Rhinolophus_convexus | Rhinolophus_convexus | nocturnal |
| mammals | Rhinolophus_creaghi | Rhinolophus_creaghi | nocturnal |
| mammals | Rhinolophus_darlingi | Rhinolophus_darlingi | nocturnal |
| mammals | Rhinolophus_deckenii | Rhinolophus_deckenii | nocturnal |
| mammals | Rhinolophus_denti | Rhinolophus_denti | nocturnal |
| mammals | Rhinolophus_eloquens | Rhinolophus_eloquens | nocturnal |
| mammals | Rhinolophus_euryale | Rhinolophus_euryale | nocturnal |
| mammals | Rhinolophus_euryotis | Rhinolophus_euryotis | nocturnal |
| mammals | Rhinolophus_ferrumequinum | Rhinolophus_ferrumequinum | nocturnal |
| mammals | Rhinolophus_formosae | Rhinolophus_formosae | nocturnal |
| mammals | Rhinolophus_fumigatus | Rhinolophus_fumigatus | nocturnal |
| mammals | Rhinolophus_guineensis | Rhinolophus_guineensis | nocturnal |
| mammals | Rhinolophus_hildebrandtii | Rhinolophus_hildebrandtii | nocturnal |
| mammals | Rhinolophus_hilli | Rhinolophus_hilli | nocturnal |
| mammals | Rhinolophus_hillorum | Rhinolophus_hillorum | nocturnal |
| mammals | Rhinolophus_hipposideros | Rhinolophus_hipposideros | nocturnal |
| mammals | Rhinolophus_inops | Rhinolophus_inops | nocturnal |
| mammals | Rhinolophus_keyensis | Rhinolophus_keyensis | nocturnal |
| mammals | Rhinolophus_landeri | Rhinolophus_landeri | nocturnal |
| mammals | Rhinolophus_lepidus | Rhinolophus_lepidus | nocturnal |
| mammals | Rhinolophus_luctus | Rhinolophus_luctus | nocturnal |
| mammals | Rhinolophus_maclaudi | Rhinolophus_maclaudi | nocturnal |
| mammals | Rhinolophus_macrotis | Rhinolophus_macrotis | nocturnal |
| mammals | Rhinolophus_madurensis | Rhinolophus_madurensis | nocturnal |
| mammals | Rhinolophus_maendeleo | Rhinolophus_maendeleo | nocturnal |
| mammals | Rhinolophus_malayanus | Rhinolophus_malayanus | nocturnal |

| | | | |
|---|---|---|---|
| mammals | Rhinolophus_marshalli | Rhinolophus_marshalli | nocturnal |
| mammals | Rhinolophus_megaphyllus | Rhinolophus_megaphyllus | nocturnal |
| mammals | Rhinolophus_mehelyi | Rhinolophus_mehelyi | nocturnal |
| mammals | Rhinolophus_mitratus | Rhinolophus_mitratus | nocturnal |
| mammals | Rhinolophus_montanus | Rhinolophus_montanus | nocturnal |
| mammals | Rhinolophus_nereis | Rhinolophus_nereis | nocturnal |
| mammals | Rhinolophus_osgoodi | Rhinolophus_osgoodi | nocturnal |
| mammals | Rhinolophus_pearsonii | Rhinolophus_pearsonii | nocturnal |
| mammals | Rhinolophus_philippinensis | Rhinolophus_philippinensis | nocturnal |
| mammals | Rhinolophus_pusillus | Rhinolophus_pusillus | nocturnal |
| mammals | Rhinolophus_rex | Rhinolophus_rex | nocturnal |
| mammals | Rhinolophus_robinsoni | Rhinolophus_robinsoni | nocturnal |
| mammals | Rhinolophus_rouxii | Rhinolophus_rouxii | nocturnal |
| mammals | Rhinolophus_rufus | Rhinolophus_rufus | nocturnal |
| mammals | Rhinolophus_ruwenzorii | Rhinolophus_ruwenzorii | nocturnal |
| mammals | Rhinolophus_sakejiensis | Rhinolophus_sakejiensis | nocturnal |
| mammals | Rhinolophus_sedulus | Rhinolophus_sedulus | nocturnal |
| mammals | Rhinolophus_shameli | Rhinolophus_shameli | nocturnal |
| mammals | Rhinolophus_shortridgei | Rhinolophus_shortridgei | nocturnal |
| mammals | Rhinolophus_siamensis | Rhinolophus_siamensis | nocturnal |
| mammals | Rhinolophus_silvestris | Rhinolophus_silvestris | nocturnal |
| mammals | Rhinolophus_simulator | Rhinolophus_simulator | nocturnal |
| mammals | Rhinolophus_sinicus | Rhinolophus_sinicus | nocturnal |
| mammals | Rhinolophus_stheno | Rhinolophus_stheno | nocturnal |
| mammals | Rhinolophus_subbadius | Rhinolophus_subbadius | nocturnal |
| mammals | Rhinolophus_subrufus | Rhinolophus_subrufus | nocturnal |
| mammals | Rhinolophus_swinnyi | Rhinolophus_swinnyi | nocturnal |
| mammals | Rhinolophus_thomasi | Rhinolophus_thomasi | nocturnal |
| mammals | Rhinolophus_trifoliatus | Rhinolophus_trifoliatus | nocturnal |
| mammals | Rhinolophus_virgo | Rhinolophus_virgo | nocturnal |
| mammals | Rhinolophus_yunanensis | Rhinolophus_yunanensis | nocturnal |
| mammals | Rhinolophus_ziama | Rhinolophus_ziama | nocturnal |
| mammals | Rhinonicteris_aurantia | Rhinonicteris_aurantia | nocturnal |
| mammals | Rhinophylla_alethina | Rhinophylla_alethina | nocturnal |

| | | | |
|---|---|---|---|
| mammals | Rhinophylla_fischerae | Rhinophylla_fischerae | nocturnal |
| mammals | Rhinophylla_pumilio | Rhinophylla_pumilio | nocturnal |
| mammals | Rhinopithecus_avunculus | Rhinopithecus_avunculus | diurnal |
| mammals | Rhinopithecus_bieti | Rhinopithecus_bieti | diurnal |
| mammals | Rhinopithecus_brelichi | Rhinopithecus_brelichi | diurnal |
| mammals | Rhinopithecus_roxellana | Rhinopithecus_roxellana | diurnal |
| mammals | Rhinopithecus_strykeri | Rhinopithecus_strykeri | diurnal |
| mammals | Rhinopoma_hardwickii | Rhinopoma_hardwickii | nocturnal |
| mammals | Rhinopoma_macinnesi | Rhinopoma_macinnesi | nocturnal |
| mammals | Rhinopoma_microphyllum | Rhinopoma_microphyllum | nocturnal |
| mammals | Rhinopoma_muscatellum | Rhinopoma_muscatellum | nocturnal |
| mammals | Rhinosciurus_laticaudatus | Rhinosciurus_laticaudatus | diurnal |
| mammals | Rhipidomys_austrinus | Rhipidomys_austrinus | nocturnal |
| mammals | Rhipidomys_caucensis | Rhipidomys_caucensis | nocturnal |
| mammals | Rhipidomys_couesi | Rhipidomys_couesi | nocturnal |
| mammals | Rhipidomys_emiliae | Rhipidomys_emiliae | nocturnal |
| mammals | Rhipidomys_fulviventer | Rhipidomys_fulviventer | nocturnal |
| mammals | Rhipidomys_gardneri | Rhipidomys_gardneri | nocturnal |
| mammals | Rhipidomys_latimanus | Rhipidomys_latimanus | nocturnal |
| mammals | Rhipidomys_leucodactylus | Rhipidomys_leucodactylus | nocturnal |
| mammals | Rhipidomys_macconnelli | Rhipidomys_macconnelli | nocturnal |
| mammals | Rhipidomys_macrurus | Rhipidomys_macrurus | nocturnal |
| mammals | Rhipidomys_mastacalis | Rhipidomys_mastacalis | nocturnal |
| mammals | Rhipidomys_nitela | Rhipidomys_nitela | nocturnal |
| mammals | Rhipidomys_ochrogaster | Rhipidomys_ochrogaster | nocturnal |
| mammals | Rhipidomys_venezuelae | Rhipidomys_venezuelae | nocturnal |
| mammals | Rhipidomys_venustus | Rhipidomys_venustus | nocturnal |
| mammals | Rhipidomys_wetzeli | Rhipidomys_wetzeli | nocturnal |
| mammals | Rhizomys_sinensis | Rhizomys_sinensis | nocturnal |
| mammals | Rhizomys_sumatrensis | Rhizomys_sumatrensis | nocturnal |
| mammals | Rhogeessa_aenea | Rhogeessa_aeneus | nocturnal |
| mammals | Rhogeessa_genowaysi | Rhogeessa_genowaysi | nocturnal |
| mammals | Rhogeessa_hussoni | Rhogeessa_hussoni | nocturnal |
| mammals | Rhogeessa_io | Rhogeessa_io | nocturnal |

| | | | |
|---|---|---|---|
| mammals | Rhogeessa_minutilla | Rhogeessa_minutilla | nocturnal |
| mammals | Rhogeessa_mira | Rhogeessa_mira | nocturnal |
| mammals | Rhogeessa_parvula | Rhogeessa_parvula | nocturnal |
| mammals | Rhogeessa_tumida | Rhogeessa_tumida | nocturnal |
| mammals | Rhombomys_opimus | Rhombomys_opimus | diurnal |
| mammals | Rhynchocyon_chrysopygus | Rhynchocyon_chrysopygus | diurnal |
| mammals | Rhynchocyon_cirnei | Rhynchocyon_cirnei | diurnal |
| mammals | Rhynchocyon_petersi | Rhynchocyon_petersi | diurnal |
| mammals | Rhynchocyon_udzungwensis | Rhynchocyon_udzungwensis | diurnal |
| mammals | Rhynchogale_melleri | Rhynchogale_melleri | nocturnal |
| mammals | Rhyncholestes_raphanurus | Rhyncholestes_raphanurus | nocturnal |
| mammals | Rhynchomys_banahao | Rhynchomys_banahao | nocturnal |
| mammals | Rhynchomys_isarogensis | Rhynchomys_isarogensis | nocturnal |
| mammals | Rhynchomys_soricoides | Rhynchomys_soricoides | cathemeral |
| mammals | Rhynchomys_tapulao | Rhynchomys_tapulao | nocturnal |
| mammals | Rhynchonycteris_naso | Rhynchonycteris_naso | nocturnal |
| mammals | Rhyneptesicus_nasutus | Eptesicus_nasutus | nocturnal |
| mammals | Romerolagus_diazi | Romerolagus_diazi | crepuscular |
| mammals | Rousettus_aegyptiacus | Rousettus_aegyptiacus | nocturnal |
| mammals | Rousettus_amplexicaudatus | Rousettus_amplexicaudatus | nocturnal |
| mammals | Rousettus_leschenaultii | Rousettus_leschenaultii | nocturnal |
| mammals | Rousettus_linduensis | Rousettus_linduensis | nocturnal |
| mammals | Rousettus_madagascariensis | Rousettus_madagascariensis | nocturnal |
| mammals | Rousettus_obliviosus | Rousettus_obliviosus | nocturnal |
| mammals | Rousettus_spinalatus | Rousettus_spinalatus | nocturnal |
| mammals | Rubrisciurus_rubriventer | Rubrisciurus_rubriventer | diurnal |
| mammals | Rucervus_duvaucelii | Rucervus_duvaucelii | diurnal |
| mammals | Rucervus_eldii | Rucervus_eldii | crepuscular |
| mammals | Rungwecebus_kipunji | Rungwecebus_kipunji | diurnal |
| mammals | Rupicapra_pyrenaica | Rupicapra_pyrenaica | diurnal |
| mammals | Rupicapra_rupicapra | Rupicapra_rupicapra | cathemeral |
| mammals | Rusa_alfredi | Rusa_alfredi | nocturnal |
| mammals | Rusa_marianna | Rusa_marianna | nocturnal |
| mammals | Rusa_timorensis | Rusa_timorensis | diurnal |

| | | | |
|---|---|---|---|
| mammals | Rusa_unicolor | Rusa_unicolor | cathemeral |
| mammals | Ruwenzorisorex_suncoides | Ruwenzorisorex_suncoides | nocturnal |
| mammals | Saccolaimus_flaviventris | Saccolaimus_flaviventris | nocturnal |
| mammals | Saccolaimus_mixtus | Saccolaimus_mixtus | nocturnal |
| mammals | Saccolaimus_peli | Saccolaimus_peli | nocturnal |
| mammals | Saccolaimus_saccolaimus | Saccolaimus_saccolaimus | nocturnal |
| mammals | Saccopteryx_antioquensis | Saccopteryx_antioquensis | nocturnal |
| mammals | Saccopteryx_bilineata | Saccopteryx_bilineata | nocturnal |
| mammals | Saccopteryx_canescens | Saccopteryx_canescens | nocturnal |
| mammals | Saccopteryx_gymnura | Saccopteryx_gymnura | nocturnal |
| mammals | Saccopteryx_leptura | Saccopteryx_leptura | nocturnal |
| mammals | Saccostomus_campestris | Saccostomus_campestris | nocturnal |
| mammals | Saccostomus_mearnsi | Saccostomus_mearnsi | nocturnal |
| mammals | Saguinus_bicolor | Saguinus_bicolor | diurnal |
| mammals | Saguinus_fuscicollis | Saguinus_fuscicollis | diurnal |
| mammals | Saguinus_geoffroyi | Saguinus_geoffroyi | diurnal |
| mammals | Saguinus_imperator | Saguinus_imperator | diurnal |
| mammals | Saguinus_inustus | Saguinus_inustus | diurnal |
| mammals | Saguinus_labiatus | Saguinus_labiatus | diurnal |
| mammals | Saguinus_leucopus | Saguinus_leucopus | diurnal |
| mammals | Saguinus_martinsi | Saguinus_martinsi | diurnal |
| mammals | Saguinus_midas | Saguinus_midas | diurnal |
| mammals | Saguinus_mystax | Saguinus_mystax | diurnal |
| mammals | Saguinus_niger | Saguinus_niger | diurnal |
| mammals | Saguinus_nigricollis | Saguinus_nigricollis | diurnal |
| mammals | Saguinus_oedipus | Saguinus_oedipus | diurnal |
| mammals | Saguinus_tripartitus | Saguinus_tripartitus | diurnal |
| mammals | Saguinus_weddelli | Saguinus_melanoleucus | diurnal |
| mammals | Saiga_tatarica | Saiga_tatarica | diurnal |
| mammals | Saimiri_boliviensis | Saimiri_boliviensis | diurnal |
| mammals | Saimiri_oerstedii | Saimiri_oerstedii | diurnal |
| mammals | Saimiri_sciureus | Saimiri_sciureus | diurnal |
| mammals | Saimiri_ustus | Saimiri_ustus | diurnal |
| mammals | Saimiri_vanzolinii | Saimiri_vanzolinii | diurnal |

| | | | |
|---|---|---|---|
| mammals | Salanoia_concolor | Salanoia_concolor | diurnal |
| mammals | Salinomys_delicatus | Salinomys_delicatus | nocturnal |
| mammals | Salpingotus_crassicauda | Salpingotus_crassicauda | nocturnal |
| mammals | Salpingotus_heptneri | Salpingotus_heptneri | nocturnal |
| mammals | Salpingotus_kozlovi | Salpingotus_kozlovi | nocturnal |
| mammals | Salpingotus_michaelis | Salpingotulus_michaelis | nocturnal |
| mammals | Salpingotus_pallidus | Salpingotus_pallidus | nocturnal |
| mammals | Santamartamys_rufodorsalis | Santamartamys_rufodorsalis | nocturnal |
| mammals | Sapajus_apella | Sapajus_apella | diurnal |
| mammals | Sapajus_flavius | Sapajus_flavius | diurnal |
| mammals | Sapajus_libidinosus | Sapajus_libidinosus | diurnal |
| mammals | Sapajus_xanthosternos | Sapajus_xanthosternos | diurnal |
| mammals | Sarcophilus_harrisii | Sarcophilus_harrisii | nocturnal |
| mammals | Sauromys_petrophilus | Sauromys_petrophilus | nocturnal |
| mammals | Saxatilomys_paulinae | Saxatilomys_paulinae | nocturnal |
| mammals | Scapteromys_aquaticus | Scapteromys_aquaticus | nocturnal |
| mammals | Scapteromys_tumidus | Scapteromys_tumidus | nocturnal |
| mammals | Scarturus_elater | Allactaga_elater | nocturnal |
| mammals | Scarturus_euphraticus | Allactaga_euphratica | nocturnal |
| mammals | Scarturus_hotsoni | Allactaga_hotsoni | nocturnal |
| mammals | Scarturus_tetradactylus | Allactaga_tetradactyla | nocturnal |
| mammals | Scarturus_vinogradovi | Allactaga_vinogradovi | nocturnal |
| mammals | Scarturus_williamsi | Allactaga_williamsi | nocturnal |
| mammals | Sciurillus_pusillus | Sciurillus_pusillus | diurnal |
| mammals | Sciurocheirus_alleni | Sciurocheirus_alleni | nocturnal |
| mammals | Sciurocheirus_gabonensis | Sciurocheirus_gabonensis | nocturnal |
| mammals | Sciurotamias_davidianus | Sciurotamias_davidianus | diurnal |
| mammals | Sciurotamias_forresti | Sciurotamias_forresti | diurnal |
| mammals | Sciurus_aberti | Sciurus_aberti | diurnal |
| mammals | Sciurus_aestuans | Sciurus_aestuans | diurnal |
| mammals | Sciurus_alleni | Sciurus_alleni | diurnal |
| mammals | Sciurus_anomalus | Sciurus_anomalus | diurnal |
| mammals | Sciurus_arizonensis | Sciurus_arizonensis | diurnal |
| mammals | Sciurus_aureogaster | Sciurus_aureogaster | diurnal |

| | | | |
|---|---|---|---|
| mammals | Sciurus_carolinensis | Sciurus_carolinensis | diurnal |
| mammals | Sciurus_colliaei | Sciurus_colliaei | diurnal |
| mammals | Sciurus_deppei | Sciurus_deppei | diurnal |
| mammals | Sciurus_granatensis | Sciurus_granatensis | diurnal |
| mammals | Sciurus_griseus | Sciurus_griseus | diurnal |
| mammals | Sciurus_ignitus | Sciurus_ignitus | diurnal |
| mammals | Sciurus_igniventris | Sciurus_igniventris | diurnal |
| mammals | Sciurus_lis | Sciurus_lis | diurnal |
| mammals | Sciurus_nayaritensis | Sciurus_nayaritensis | diurnal |
| mammals | Sciurus_niger | Sciurus_niger | diurnal |
| mammals | Sciurus_oculatus | Sciurus_oculatus | diurnal |
| mammals | Sciurus_pucheranii | Sciurus_pucheranii | diurnal |
| mammals | Sciurus_pyrrhinus | Sciurus_pyrrhinus | diurnal |
| mammals | Sciurus_richmondi | Sciurus_richmondi | diurnal |
| mammals | Sciurus_spadiceus | Sciurus_spadiceus | diurnal |
| mammals | Sciurus_stramineus | Sciurus_stramineus | diurnal |
| mammals | Sciurus_variegatoides | Sciurus_variegatoides | diurnal |
| mammals | Sciurus_vulgaris | Sciurus_vulgaris | diurnal |
| mammals | Sciurus_yucatanensis | Sciurus_yucatanensis | diurnal |
| mammals | Scleronycteris_ega | Scleronycteris_ega | nocturnal |
| mammals | Scolomys_melanops | Scolomys_melanops | nocturnal |
| mammals | Scolomys_ucayalensis | Scolomys_ucayalensis | nocturnal |
| mammals | Scoteanax_rueppellii | Scoteanax_rueppellii | nocturnal |
| mammals | Scotinomys_teguina | Scotinomys_teguina | diurnal |
| mammals | Scotinomys_xerampelinus | Scotinomys_xerampelinus | diurnal |
| mammals | Scotoecus_albofuscus | Scotoecus_albofuscus | nocturnal |
| mammals | Scotoecus_hirundo | Scotoecus_hirundo | nocturnal |
| mammals | Scotoecus_pallidus | Scotoecus_pallidus | nocturnal |
| mammals | Scotomanes_ornatus | Scotomanes_ornatus | nocturnal |
| mammals | Scotonycteris_zenkeri | Scotonycteris_zenkeri | nocturnal |
| mammals | Scotophilus_collinus | Scotophilus_collinus | nocturnal |
| mammals | Scotophilus_dinganii | Scotophilus_dinganii | nocturnal |
| mammals | Scotophilus_heathii | Scotophilus_heathii | nocturnal |
| mammals | Scotophilus_kuhlii | Scotophilus_kuhlii | nocturnal |

| | | | |
|---|---|---|---|
| mammals | Scotophilus_leucogaster | Scotophilus_leucogaster | nocturnal |
| mammals | Scotophilus_marovaza | Scotophilus_marovaza | nocturnal |
| mammals | Scotophilus_nigrita | Scotophilus_nigrita | nocturnal |
| mammals | Scotophilus_nucella | Scotophilus_nucella | nocturnal |
| mammals | Scotophilus_nux | Scotophilus_nux | nocturnal |
| mammals | Scotophilus_robustus | Scotophilus_robustus | nocturnal |
| mammals | Scotophilus_tandrefana | Scotophilus_tandrefana | nocturnal |
| mammals | Scotophilus_viridis | Scotophilus_viridis | nocturnal |
| mammals | Scotorepens_balstoni | Scotorepens_balstoni | nocturnal |
| mammals | Scotorepens_greyii | Scotorepens_greyii | nocturnal |
| mammals | Scotorepens_orion | Scotorepens_orion | nocturnal |
| mammals | Scotorepens_sanborni | Scotorepens_sanborni | nocturnal |
| mammals | Scotozous_dormeri | Scotozous_dormeri | nocturnal |
| mammals | Scutisorex_somereni | Scutisorex_somereni | cathemeral |
| mammals | Sekeetamys_calurus | Sekeetamys_calurus | cathemeral |
| mammals | Selevinia_betpakdalaensis | Selevinia_betpakdalaensis | nocturnal |
| mammals | Semnopithecus_ajax | Semnopithecus_ajax | diurnal |
| mammals | Semnopithecus_entellus | Semnopithecus_entellus | diurnal |
| mammals | Semnopithecus_hector | Semnopithecus_hector | diurnal |
| mammals | Semnopithecus_hypoleucos | Semnopithecus_hypoleucos | diurnal |
| mammals | Semnopithecus_johnii | Trachypithecus_johnii | diurnal |
| mammals | Semnopithecus_priam | Semnopithecus_priam | diurnal |
| mammals | Semnopithecus_schistaceus | Semnopithecus_schistaceus | diurnal |
| mammals | Semnopithecus_vetulus | Trachypithecus_vetulus | diurnal |
| mammals | Serengetimys_pernanus | Mastomys_pernanus | nocturnal |
| mammals | Setifer_setosus | Setifer_setosus | nocturnal |
| mammals | Setonix_brachyurus | Setonix_brachyurus | nocturnal |
| mammals | Sicista_betulina | Sicista_betulina | nocturnal |
| mammals | Sicista_caucasica | Sicista_caucasica | nocturnal |
| mammals | Sicista_caudata | Sicista_caudata | nocturnal |
| mammals | Sicista_concolor | Sicista_concolor | nocturnal |
| mammals | Sicista_kazbegica | Sicista_kazbegica | nocturnal |
| mammals | Sicista_kluchorica | Sicista_kluchorica | nocturnal |
| mammals | Sicista_napaea | Sicista_napaea | nocturnal |

| | | | |
|---|---|---|---|
| mammals | Sicista_pseudonapaea | Sicista_pseudonapaea | nocturnal |
| mammals | Sicista_severtzovi | Sicista_severtzovi | nocturnal |
| mammals | Sicista_strandi | Sicista_strandi | nocturnal |
| mammals | Sicista_subtilis | Sicista_subtilis | nocturnal |
| mammals | Sicista_tianschanica | Sicista_tianshanica | nocturnal |
| mammals | Sigmodon_alleni | Sigmodon_alleni | cathemeral |
| mammals | Sigmodon_alstoni | Sigmodon_alstoni | cathemeral |
| mammals | Sigmodon_arizonae | Sigmodon_arizonae | cathemeral |
| mammals | Sigmodon_fulviventer | Sigmodon_fulviventer | cathemeral |
| mammals | Sigmodon_hirsutus | Sigmodon_hirsutus | cathemeral |
| mammals | Sigmodon_hispidus | Sigmodon_hispidus | cathemeral |
| mammals | Sigmodon_inopinatus | Sigmodon_inopinatus | cathemeral |
| mammals | Sigmodon_leucotis | Sigmodon_leucotis | diurnal |
| mammals | Sigmodon_mascotensis | Sigmodon_mascotensis | cathemeral |
| mammals | Sigmodon_ochrognathus | Sigmodon_ochrognathus | diurnal |
| mammals | Sigmodon_peruanus | Sigmodon_peruanus | cathemeral |
| mammals | Sigmodon_toltecus | Sigmodon_toltecus | diurnal |
| mammals | Sigmodontomys_alfari | Sigmodontomys_alfari | nocturnal |
| mammals | Simias_concolor | Simias_concolor | diurnal |
| mammals | Sminthopsis_archeri | Sminthopsis_archeri | nocturnal |
| mammals | Sminthopsis_bindi | Sminthopsis_bindi | nocturnal |
| mammals | Sminthopsis_butleri | Sminthopsis_butleri | nocturnal |
| mammals | Sminthopsis_crassicaudata | Sminthopsis_crassicaudata | nocturnal |
| mammals | Sminthopsis_dolichura | Sminthopsis_dolichura | nocturnal |
| mammals | Sminthopsis_douglasi | Sminthopsis_douglasi | nocturnal |
| mammals | Sminthopsis_fuliginosa | Sminthopsis_griseoventer | nocturnal |
| mammals | Sminthopsis_gilberti | Sminthopsis_gilberti | nocturnal |
| mammals | Sminthopsis_granulipes | Sminthopsis_granulipes | nocturnal |
| mammals | Sminthopsis_hirtipes | Sminthopsis_hirtipes | nocturnal |
| mammals | Sminthopsis_leucopus | Sminthopsis_leucopus | nocturnal |
| mammals | Sminthopsis_macroura | Sminthopsis_macroura | nocturnal |
| mammals | Sminthopsis_murina | Sminthopsis_murina | nocturnal |
| mammals | Sminthopsis_ooldea | Sminthopsis_ooldea | nocturnal |
| mammals | Sminthopsis_psammophila | Sminthopsis_psammophila | nocturnal |

| | | | |
|---|---|---|---|
| mammals | Sminthopsis_virginiae | Sminthopsis_virginiae | nocturnal |
| mammals | Sminthopsis_youngsoni | Sminthopsis_youngsoni | nocturnal |
| mammals | Smutsia_gigantea | Smutsia_gigantea | nocturnal |
| mammals | Smutsia_temminckii | Smutsia_temminckii | cathemeral |
| mammals | Solenodon_paradoxus | Solenodon_paradoxus | nocturnal |
| mammals | Solisorex_pearsoni | Solisorex_pearsoni | nocturnal |
| mammals | Solomys_ponceleti | Solomys_ponceleti | nocturnal |
| mammals | Solomys_salebrosus | Solomys_salebrosus | nocturnal |
| mammals | Solomys_sapientis | Solomys_sapientis | nocturnal |
| mammals | Sommeromys_macrorhinos | Sommeromys_macrorhinos | nocturnal |
| mammals | Sooretamys_angouya | Sooretamys_angouya | nocturnal |
| mammals | Sorex_alpinus | Sorex_alpinus | cathemeral |
| mammals | Sorex_antinorii | Sorex_antinorii | cathemeral |
| mammals | Sorex_araneus | Sorex_araneus | cathemeral |
| mammals | Sorex_arcticus | Sorex_arcticus | cathemeral |
| mammals | Sorex_arizonae | Sorex_arizonae | cathemeral |
| mammals | Sorex_asper | Sorex_asper | cathemeral |
| mammals | Sorex_bedfordiae | Sorex_bedfordiae | nocturnal |
| mammals | Sorex_bendirii | Sorex_bendirii | diurnal |
| mammals | Sorex_buchariensis | Sorex_buchariensis | cathemeral |
| mammals | Sorex_caecutiens | Sorex_caecutiens | cathemeral |
| mammals | Sorex_camtschaticus | Sorex_camtschatica | cathemeral |
| mammals | Sorex_cansulus | Sorex_cansulus | cathemeral |
| mammals | Sorex_coronatus | Sorex_coronatus | cathemeral |
| mammals | Sorex_cylindricauda | Sorex_cylindricauda | cathemeral |
| mammals | Sorex_daphaenodon | Sorex_daphaenodon | cathemeral |
| mammals | Sorex_emarginatus | Sorex_emarginatus | cathemeral |
| mammals | Sorex_excelsus | Sorex_excelsus | nocturnal |
| mammals | Sorex_fumeus | Sorex_fumeus | cathemeral |
| mammals | Sorex_gracillimus | Sorex_gracillimus | nocturnal |
| mammals | Sorex_granarius | Sorex_granarius | cathemeral |
| mammals | Sorex_haydeni | Sorex_haydeni | diurnal |
| mammals | Sorex_hosonoi | Sorex_hosonoi | cathemeral |
| mammals | Sorex_isodon | Sorex_isodon | cathemeral |

| | | | |
|---|---|---|---|
| mammals | Sorex_ixtlanensis | Sorex_ixtlanensis | cathemeral |
| mammals | Sorex_jacksoni | Sorex_jacksoni | cathemeral |
| mammals | Sorex_kozlovi | Sorex_kozlovi | cathemeral |
| mammals | Sorex_leucogaster | Sorex_leucogaster | cathemeral |
| mammals | Sorex_longirostris | Sorex_longirostris | cathemeral |
| mammals | Sorex_lyelli | Sorex_lyelli | cathemeral |
| mammals | Sorex_maritimensis | Sorex_maritimensis | cathemeral |
| mammals | Sorex_mediopua | Sorex_mediopua | cathemeral |
| mammals | Sorex_merriami | Sorex_merriami | cathemeral |
| mammals | Sorex_milleri | Sorex_milleri | cathemeral |
| mammals | Sorex_minutissimus | Sorex_minutissimus | cathemeral |
| mammals | Sorex_minutus | Sorex_minutus | crepuscular |
| mammals | Sorex_monticolus | Sorex_monticolus | cathemeral |
| mammals | Sorex_nanus | Sorex_nanus | cathemeral |
| mammals | Sorex_navigator | Sorex_alaskanus | cathemeral |
| mammals | Sorex_oreopolus | Sorex_oreopolus | cathemeral |
| mammals | Sorex_orizabae | Sorex_orizabae | cathemeral |
| mammals | Sorex_pacificus | Sorex_pacificus | cathemeral |
| mammals | Sorex_palustris | Sorex_palustris | cathemeral |
| mammals | Sorex_planiceps | Sorex_planiceps | cathemeral |
| mammals | Sorex_portenkoi | Sorex_portenkoi | cathemeral |
| mammals | Sorex_preblei | Sorex_preblei | cathemeral |
| mammals | Sorex_pribilofensis | Sorex_pribilofensis | cathemeral |
| mammals | Sorex_raddei | Sorex_raddei | nocturnal |
| mammals | Sorex_roboratus | Sorex_roboratus | cathemeral |
| mammals | Sorex_rohweri | Sorex_rohweri | cathemeral |
| mammals | Sorex_samniticus | Sorex_samniticus | cathemeral |
| mammals | Sorex_satunini | Sorex_satunini | nocturnal |
| mammals | Sorex_saussurei | Sorex_saussurei | cathemeral |
| mammals | Sorex_sclateri | Sorex_sclateri | cathemeral |
| mammals | Sorex_shinto | Sorex_shinto | nocturnal |
| mammals | Sorex_sinalis | Sorex_sinalis | cathemeral |
| mammals | Sorex_sonomae | Sorex_sonomae | nocturnal |
| mammals | Sorex_tenellus | Sorex_tenellus | cathemeral |

| | | | |
|---|---|---|---|
| mammals | Sorex_thibetanus | Sorex_thibetanus | cathemeral |
| mammals | Sorex_trowbridgii | Sorex_trowbridgii | cathemeral |
| mammals | Sorex_tundrensis | Sorex_tundrensis | cathemeral |
| mammals | Sorex_ugyunak | Sorex_ugyunak | cathemeral |
| mammals | Sorex_unguiculatus | Sorex_unguiculatus | cathemeral |
| mammals | Sorex_vagrans | Sorex_vagrans | cathemeral |
| mammals | Sorex_ventralis | Sorex_ventralis | cathemeral |
| mammals | Sorex_veraecrucis | Sorex_veraecrucis | cathemeral |
| mammals | Sorex_veraepacis | Sorex_veraepacis | cathemeral |
| mammals | Sorex_volnuchini | Sorex_volnuchini | cathemeral |
| mammals | Soricomys_kalinga | Soricomys_kalinga | diurnal |
| mammals | Soricomys_musseri | Soricomys_musseri | diurnal |
| mammals | Soriculus_nigrescens | Soriculus_nigrescens | nocturnal |
| mammals | Spalacopus_cyanus | Spalacopus_cyanus | diurnal |
| mammals | Speothos_venaticus | Speothos_venaticus | diurnal |
| mammals | Spermophilopsis_leptodactyla | Spermophilopsis_leptodactylus | diurnal |
| mammals | Spermophilus_alaschanicus | Spermophilus_alashanicus | diurnal |
| mammals | Spermophilus_brevicauda | Spermophilus_brevicauda | diurnal |
| mammals | Spermophilus_citellus | Spermophilus_citellus | diurnal |
| mammals | Spermophilus_dauricus | Spermophilus_dauricus | diurnal |
| mammals | Spermophilus_erythrogenys | Spermophilus_erythrogenys | diurnal |
| mammals | Spermophilus_fulvus | Spermophilus_fulvus | diurnal |
| mammals | Spermophilus_major | Spermophilus_major | diurnal |
| mammals | Spermophilus_musicus | Spermophilus_musicus | diurnal |
| mammals | Spermophilus_nilkaensis | Spermophilus_ralli | diurnal |
| mammals | Spermophilus_pallidicauda | Spermophilus_pallidicauda | diurnal |
| mammals | Spermophilus_pygmaeus | Spermophilus_pygmaeus | diurnal |
| mammals | Spermophilus_relictus | Spermophilus_relictus | diurnal |
| mammals | Spermophilus_suslicus | Spermophilus_suslicus | diurnal |
| mammals | Spermophilus_taurensis | Spermophilus_taurensis | diurnal |
| mammals | Spermophilus_xanthoprymnus | Spermophilus_xanthoprymnus | diurnal |
| mammals | Sphaerias_blanfordi | Sphaerias_blanfordi | nocturnal |
| mammals | Sphaeronycteris_toxophyllum | Sphaeronycteris_toxophyllum | nocturnal |
| mammals | Spilocuscus_kraemeri | Spilocuscus_kraemeri | nocturnal |

| | | | |
|---|---|---|---|
| mammals | Spilocuscus_maculatus | Spilocuscus_maculatus | nocturnal |
| mammals | Spilocuscus_papuensis | Spilocuscus_papuensis | cathemeral |
| mammals | Spilocuscus_rufoniger | Spilocuscus_rufoniger | cathemeral |
| mammals | Spilocuscus_wilsoni | Spilocuscus_wilsoni | cathemeral |
| mammals | Spilogale_angustifrons | Spilogale_angustifrons | nocturnal |
| mammals | Spilogale_gracilis | Spilogale_gracilis | nocturnal |
| mammals | Spilogale_putorius | Spilogale_putorius | nocturnal |
| mammals | Spilogale_pygmaea | Spilogale_pygmaea | nocturnal |
| mammals | Srilankamys_ohiensis | Srilankamys_ohiensis | nocturnal |
| mammals | Steatomys_caurinus | Steatomys_caurinus | nocturnal |
| mammals | Steatomys_cuppedius | Steatomys_cuppedius | nocturnal |
| mammals | Steatomys_jacksoni | Steatomys_jacksoni | nocturnal |
| mammals | Steatomys_krebsii | Steatomys_krebsii | nocturnal |
| mammals | Steatomys_opimus | Steatomys_opimus | nocturnal |
| mammals | Steatomys_parvus | Steatomys_parvus | nocturnal |
| mammals | Steatomys_pratensis | Steatomys_pratensis | nocturnal |
| mammals | Stenocephalemys_albipes | Stenocephalemys_albipes | nocturnal |
| mammals | Stenocephalemys_albocaudatus | Stenocephalemys_albocaudata | nocturnal |
| mammals | Stenocephalemys_griseicauda | Stenocephalemys_griseicauda | nocturnal |
| mammals | Stenocephalemys_ruppi | Stenocephalemys_ruppi | nocturnal |
| mammals | Stenocranius_gregalis | Microtus_gregalis | cathemeral |
| mammals | Stenoderma_rufum | Stenoderma_rufum | nocturnal |
| mammals | Stenonycteris_lanosa | Rousettus_lanosus | nocturnal |
| mammals | Stochomys_longicaudatus | Stochomys_longicaudatus | nocturnal |
| mammals | Strigocuscus_celebensis | Strigocuscus_celebensis | nocturnal |
| mammals | Sturnira_aratathomasi | Sturnira_aratathomasi | nocturnal |
| mammals | Sturnira_bidens | Sturnira_bidens | nocturnal |
| mammals | Sturnira_bogotensis | Sturnira_bogotensis | nocturnal |
| mammals | Sturnira_erythromos | Sturnira_erythromos | nocturnal |
| mammals | Sturnira_koopmanhilli | Sturnira_koopmanhilli | nocturnal |
| mammals | Sturnira_lilium | Sturnira_lilium | nocturnal |
| mammals | Sturnira_ludovici | Sturnira_ludovici | nocturnal |
| mammals | Sturnira_luisi | Sturnira_luisi | nocturnal |
| mammals | Sturnira_magna | Sturnira_magna | nocturnal |

| | | | |
|---|---|---|---|
| mammals | Sturnira_mistratensis | Sturnira_mistratensis | nocturnal |
| mammals | Sturnira_mordax | Sturnira_mordax | nocturnal |
| mammals | Sturnira_nana | Sturnira_nana | nocturnal |
| mammals | Sturnira_oporaphilum | Sturnira_oporaphilum | nocturnal |
| mammals | Sturnira_perla | Sturnira_perla | nocturnal |
| mammals | Sturnira_sorianoi | Sturnira_sorianoi | nocturnal |
| mammals | Sturnira_tildae | Sturnira_tildae | nocturnal |
| mammals | Styloctenium_mindorense | Styloctenium_mindorensis | nocturnal |
| mammals | Styloctenium_wallacei | Styloctenium_wallacei | nocturnal |
| mammals | Stylodipus_andrewsi | Stylodipus_andrewsi | nocturnal |
| mammals | Stylodipus_sungorus | Stylodipus_sungorus | nocturnal |
| mammals | Stylodipus_telum | Stylodipus_telum | nocturnal |
| mammals | Subulo_gouazoubira | Mazama_gouazoubira | cathemeral |
| mammals | Suncus_aequatorius | Suncus_aequatorius | nocturnal |
| mammals | Suncus_dayi | Suncus_dayi | nocturnal |
| mammals | Suncus_etruscus | Suncus_etruscus | nocturnal |
| mammals | Suncus_hosei | Suncus_hosei | cathemeral |
| mammals | Suncus_hututsi | Suncus_hututsi | nocturnal |
| mammals | Suncus_infinitesimus | Suncus_infinitesimus | cathemeral |
| mammals | Suncus_lixa | Suncus_lixus | nocturnal |
| mammals | Suncus_malayanus | Suncus_malayanus | nocturnal |
| mammals | Suncus_megalurus | Suncus_megalura | nocturnal |
| mammals | Suncus_mertensi | Suncus_mertensi | nocturnal |
| mammals | Suncus_montanus | Suncus_montanus | nocturnal |
| mammals | Suncus_murinus | Suncus_murinus | nocturnal |
| mammals | Suncus_remyi | Suncus_remyi | nocturnal |
| mammals | Suncus_stoliczkanus | Suncus_stoliczkanus | nocturnal |
| mammals | Suncus_varilla | Suncus_varilla | nocturnal |
| mammals | Suncus_zeylanicus | Suncus_zeylanicus | nocturnal |
| mammals | Sundamys_annandalei | Rattus_annandalei | nocturnal |
| mammals | Sundamys_infraluteus | Sundamys_infraluteus | nocturnal |
| mammals | Sundamys_maxi | Sundamys_maxi | nocturnal |
| mammals | Sundamys_muelleri | Sundamys_muelleri | nocturnal |
| mammals | Sundasciurus_brookei | Sundasciurus_brookei | diurnal |

| | | | |
|---|---|---|---|
| mammals | Sundasciurus_davensis | Sundasciurus_davensis | diurnal |
| mammals | Sundasciurus_everetti | Dremomys_everetti | diurnal |
| mammals | Sundasciurus_fraterculus | Sundasciurus_fraterculus | diurnal |
| mammals | Sundasciurus_hippurus | Sundasciurus_hippurus | diurnal |
| mammals | Sundasciurus_hoogstraali | Sundasciurus_hoogstraali | diurnal |
| mammals | Sundasciurus_jentinki | Sundasciurus_jentinki | diurnal |
| mammals | Sundasciurus_juvencus | Sundasciurus_juvencus | diurnal |
| mammals | Sundasciurus_lowii | Sundasciurus_lowii | diurnal |
| mammals | Sundasciurus_mindanensis | Sundasciurus_mindanensis | diurnal |
| mammals | Sundasciurus_moellendorffi | Sundasciurus_moellendorffi | diurnal |
| mammals | Sundasciurus_philippinensis | Sundasciurus_philippinensis | diurnal |
| mammals | Sundasciurus_rabori | Sundasciurus_rabori | diurnal |
| mammals | Sundasciurus_samarensis | Sundasciurus_samarensis | diurnal |
| mammals | Sundasciurus_steerii | Sundasciurus_steerii | diurnal |
| mammals | Sundasciurus_tenuis | Sundasciurus_tenuis | diurnal |
| mammals | Surdisorex_norae | Surdisorex_norae | cathemeral |
| mammals | Surdisorex_polulus | Surdisorex_polulus | cathemeral |
| mammals | Suricata_suricatta | Suricata_suricatta | diurnal |
| mammals | Sus_ahoenobarbus | Sus_ahoenobarbus | diurnal |
| mammals | Sus_barbatus | Sus_barbatus | diurnal |
| mammals | Sus_cebifrons | Sus_cebifrons | nocturnal |
| mammals | Sus_celebensis | Sus_celebensis | diurnal |
| mammals | Sus_oliveri | Sus_oliveri | nocturnal |
| mammals | Sus_philippensis | Sus_philippensis | nocturnal |
| mammals | Sus_scrofa | Sus_scrofa | cathemeral |
| mammals | Sus_verrucosus | Sus_verrucosus | nocturnal |
| mammals | Syconycteris_australis | Syconycteris_australis | nocturnal |
| mammals | Syconycteris_carolinae | Syconycteris_carolinae | nocturnal |
| mammals | Syconycteris_hobbit | Syconycteris_hobbit | nocturnal |
| mammals | Sylvicapra_grimmia | Sylvicapra_grimmia | diurnal |
| mammals | Sylvilagus_aquaticus | Sylvilagus_aquaticus | nocturnal |
| mammals | Sylvilagus_audubonii | Sylvilagus_audubonii | crepuscular |
| mammals | Sylvilagus_bachmani | Sylvilagus_bachmani | crepuscular |
| mammals | Sylvilagus_brasiliensis | Sylvilagus_brasiliensis | nocturnal |

| | | | |
|---|---|---|---|
| mammals | Sylvilagus_cunicularius | Sylvilagus_cunicularius | crepuscular |
| mammals | Sylvilagus_dicei | Sylvilagus_dicei | nocturnal |
| mammals | Sylvilagus_floridanus | Sylvilagus_floridanus | crepuscular |
| mammals | Sylvilagus_graysoni | Sylvilagus_graysoni | nocturnal |
| mammals | Sylvilagus_holzneri | Sylvilagus_robustus | nocturnal |
| mammals | Sylvilagus_idahoensis | Brachylagus_idahoensis | crepuscular |
| mammals | Sylvilagus_insonus | Sylvilagus_insonus | nocturnal |
| mammals | Sylvilagus_nuttallii | Sylvilagus_nuttallii | crepuscular |
| mammals | Sylvilagus_obscurus | Sylvilagus_obscurus | diurnal |
| mammals | Sylvilagus_palustris | Sylvilagus_palustris | nocturnal |
| mammals | Sylvilagus_transitionalis | Sylvilagus_transitionalis | crepuscular |
| mammals | Sylvisorex_camerunensis | Sylvisorex_camerunensis | cathemeral |
| mammals | Sylvisorex_granti | Sylvisorex_granti | cathemeral |
| mammals | Sylvisorex_howelli | Sylvisorex_howelli | nocturnal |
| mammals | Sylvisorex_isabellae | Sylvisorex_isabellae | cathemeral |
| mammals | Sylvisorex_johnstoni | Sylvisorex_johnstoni | cathemeral |
| mammals | Sylvisorex_konganensis | Sylvisorex_konganensis | cathemeral |
| mammals | Sylvisorex_lunaris | Sylvisorex_lunaris | cathemeral |
| mammals | Sylvisorex_morio | Sylvisorex_morio | cathemeral |
| mammals | Sylvisorex_ollula | Sylvisorex_ollula | cathemeral |
| mammals | Sylvisorex_oriundus | Sylvisorex_oriundus | cathemeral |
| mammals | Sylvisorex_pluvialis | Sylvisorex_pluvialis | cathemeral |
| mammals | Sylvisorex_vulcanorum | Sylvisorex_vulcanorum | cathemeral |
| mammals | Symphalangus_syndactylus | Symphalangus_syndactylus | diurnal |
| mammals | Synaptomys_cooperi | Synaptomys_cooperi | cathemeral |
| mammals | Syncerus_caffer | Syncerus_caffer | cathemeral |
| mammals | Syntheosciurus_brochus | Syntheosciurus_brochus | diurnal |
| mammals | Tachyglossus_aculeatus | Tachyglossus_aculeatus | nocturnal |
| mammals | Tadarida_aegyptiaca | Tadarida_aegyptiaca | nocturnal |
| mammals | Tadarida_brasiliensis | Tadarida_brasiliensis | nocturnal |
| mammals | Tadarida_fulminans | Tadarida_fulminans | nocturnal |
| mammals | Tadarida_insignis | Tadarida_insignis | nocturnal |
| mammals | Tadarida_latouchei | Tadarida_latouchei | nocturnal |
| mammals | Tadarida_lobata | Tadarida_lobata | nocturnal |

| | | | |
|---|---|---|---|
| mammals | Tadarida_teniotis | Tadarida_teniotis | nocturnal |
| mammals | Tadarida_ventralis | Tadarida_ventralis | nocturnal |
| mammals | Taeromys_callitrichus | Taeromys_callitrichus | nocturnal |
| mammals | Taeromys_celebensis | Taeromys_celebensis | nocturnal |
| mammals | Taeromys_dominator | Paruromys_dominator | nocturnal |
| mammals | Taeromys_hamatus | Taeromys_hamatus | nocturnal |
| mammals | Taeromys_punicans | Taeromys_punicans | nocturnal |
| mammals | Taeromys_taerae | Taeromys_taerae | nocturnal |
| mammals | Tamandua_mexicana | Tamandua_mexicana | cathemeral |
| mammals | Tamandua_tetradactyla | Tamandua_tetradactyla | diurnal |
| mammals | Tamias_striatus | Tamias_striatus | diurnal |
| mammals | Tamiasciurus_douglasii | Tamiasciurus_douglasii | diurnal |
| mammals | Tamiasciurus_hudsonicus | Tamiasciurus_hudsonicus | diurnal |
| mammals | Tamiops_maritimus | Tamiops_maritimus | diurnal |
| mammals | Tamiops_mcclellandii | Tamiops_macclellandii | diurnal |
| mammals | Tamiops_rodolphii | Tamiops_rodolphii | diurnal |
| mammals | Tamiops_swinhoei | Tamiops_swinhoei | diurnal |
| mammals | Tapecomys_primus | Tapecomys_primus | nocturnal |
| mammals | Tapecomys_wolffsohni | Phyllotis_wolffsohni | cathemeral |
| mammals | Taphozous_achates | Taphozous_achates | nocturnal |
| mammals | Taphozous_australis | Taphozous_australis | nocturnal |
| mammals | Taphozous_georgianus | Taphozous_georgianus | crepuscular |
| mammals | Taphozous_hamiltoni | Taphozous_hamiltoni | nocturnal |
| mammals | Taphozous_hildegardeae | Taphozous_hildegardeae | nocturnal |
| mammals | Taphozous_hilli | Taphozous_hilli | nocturnal |
| mammals | Taphozous_kapalgensis | Taphozous_kapalgensis | nocturnal |
| mammals | Taphozous_longimanus | Taphozous_longimanus | nocturnal |
| mammals | Taphozous_mauritianus | Taphozous_mauritianus | nocturnal |
| mammals | Taphozous_melanopogon | Taphozous_melanopogon | nocturnal |
| mammals | Taphozous_nudiventris | Taphozous_nudiventris | nocturnal |
| mammals | Taphozous_perforatus | Taphozous_perforatus | nocturnal |
| mammals | Taphozous_theobaldi | Taphozous_theobaldi | nocturnal |
| mammals | Taphozous_troughtoni | Taphozous_troughtoni | nocturnal |
| mammals | Tapirus_bairdii | Tapirus_bairdii | nocturnal |

| | | | |
|---|---|---|---|
| mammals | Tapirus_indicus | Tapirus_indicus | nocturnal |
| mammals | Tapirus_pinchaque | Tapirus_pinchaque | cathemeral |
| mammals | Tapirus_terrestris | Tapirus_terrestris | cathemeral |
| mammals | Tarsipes_rostratus | Tarsipes_rostratus | nocturnal |
| mammals | Tarsius_dentatus | Tarsius_dentatus | nocturnal |
| mammals | Tarsius_lariang | Tarsius_lariang | nocturnal |
| mammals | Tarsius_pelengensis | Tarsius_pelengensis | nocturnal |
| mammals | Tarsius_pumilus | Tarsius_pumilus | nocturnal |
| mammals | Tarsius_sangirensis | Tarsius_sangirensis | nocturnal |
| mammals | Tarsius_tarsier | Tarsius_tarsier | nocturnal |
| mammals | Tarsius_tumpara | Tarsius_tumpara | nocturnal |
| mammals | Tarsius_wallacei | Tarsius_wallacei | nocturnal |
| mammals | Tarsomys_apoensis | Tarsomys_apoensis | cathemeral |
| mammals | Tateomys_macrocercus | Tateomys_macrocercus | nocturnal |
| mammals | Tateomys_rhinogradoides | Tateomys_rhinogradoides | nocturnal |
| mammals | Tatera_indica | Tatera_indica | nocturnal |
| mammals | Taterillus_arenarius | Taterillus_arenarius | nocturnal |
| mammals | Taterillus_congicus | Taterillus_congicus | nocturnal |
| mammals | Taterillus_emini | Taterillus_emini | nocturnal |
| mammals | Taterillus_gracilis | Taterillus_gracilis | nocturnal |
| mammals | Taterillus_lacustris | Taterillus_lacustris | nocturnal |
| mammals | Taterillus_petteri | Taterillus_petteri | nocturnal |
| mammals | Taterillus_pygargus | Taterillus_pygargus | nocturnal |
| mammals | Taterillus_tranieri | Taterillus_tranieri | nocturnal |
| mammals | Taxidea_taxus | Taxidea_taxus | nocturnal |
| mammals | Tayassu_pecari | Tayassu_pecari | cathemeral |
| mammals | Tenrec_ecaudatus | Tenrec_ecaudatus | nocturnal |
| mammals | Tetracerus_quadricornis | Tetracerus_quadricornis | crepuscular |
| mammals | Thainycteris_aureocollaris | Arielulus_aureocollaris | nocturnal |
| mammals | Thainycteris_torquata | Arielulus_torquatus | nocturnal |
| mammals | Thallomys_loringi | Thallomys_loringi | nocturnal |
| mammals | Thallomys_nigricauda | Thallomys_nigricauda | nocturnal |
| mammals | Thallomys_paedulcus | Thallomys_paedulcus | nocturnal |
| mammals | Thallomys_shortridgei | Thallomys_shortridgei | nocturnal |

| | | | |
|---|---|---|---|
| mammals | Thalpomys_cerradensis | Thalpomys_cerradensis | nocturnal |
| mammals | Thalpomys_lasiotis | Thalpomys_lasiotis | nocturnal |
| mammals | Thamnomys_kempi | Thamnomys_kempi | nocturnal |
| mammals | Thamnomys_kuru | Grammomys_kuru | nocturnal |
| mammals | Thamnomys_schoutedeni | Thamnomys_schoutedeni | nocturnal |
| mammals | Thamnomys_venustus | Thamnomys_venustus | nocturnal |
| mammals | Thaptomys_nigrita | Thaptomys_nigrita | diurnal |
| mammals | Theropithecus_gelada | Theropithecus_gelada | diurnal |
| mammals | Thomasomys_aureus | Thomasomys_aureus | nocturnal |
| mammals | Thomasomys_baeops | Thomasomys_baeops | nocturnal |
| mammals | Thomasomys_bombycinus | Thomasomys_bombycinus | nocturnal |
| mammals | Thomasomys_caudivarius | Thomasomys_caudivarius | nocturnal |
| mammals | Thomasomys_cinereus | Thomasomys_cinereus | nocturnal |
| mammals | Thomasomys_cinnameus | Thomasomys_cinnameus | nocturnal |
| mammals | Thomasomys_erro | Thomasomys_erro | nocturnal |
| mammals | Thomasomys_hudsoni | Thomasomys_hudsoni | nocturnal |
| mammals | Thomasomys_incanus | Thomasomys_incanus | nocturnal |
| mammals | Thomasomys_kalinowskii | Thomasomys_kalinowskii | nocturnal |
| mammals | Thomasomys_paramorum | Thomasomys_paramorum | nocturnal |
| mammals | Thomasomys_praetor | Thomasomys_praetor | nocturnal |
| mammals | Thomasomys_pyrrhonotus | Thomasomys_pyrrhonotus | nocturnal |
| mammals | Thomasomys_taczanowskii | Thomasomys_taczanowskii | nocturnal |
| mammals | Thomasomys_ucucha | Thomasomys_ucucha | nocturnal |
| mammals | Thomasomys_vestitus | Thomasomys_vestitus | nocturnal |
| mammals | Thomasomys_vulcani | Thomasomys_vulcani | nocturnal |
| mammals | Thoopterus_nigrescens | Thoopterus_nigrescens | nocturnal |
| mammals | Thrichomys_apereoides | Thrichomys_apereoides | nocturnal |
| mammals | Thrichomys_inermis | Thrichomys_inermis | crepuscular |
| mammals | Thrichomys_laurentius | Thrichomys_laurentius | crepuscular |
| mammals | Thrichomys_pachyurus | Thrichomys_pachyurus | crepuscular |
| mammals | Thryonomys_gregorianus | Thryonomys_gregorianus | nocturnal |
| mammals | Thryonomys_swinderianus | Thryonomys_swinderianus | nocturnal |
| mammals | Thylamys_elegans | Thylamys_elegans | nocturnal |
| mammals | Thylamys_karimii | Thylamys_karimii | nocturnal |

| | | | |
|---|---|---|---|
| mammals | Thylamys_macrurus | Thylamys_macrurus | nocturnal |
| mammals | Thylamys_pallidior | Thylamys_pallidior | nocturnal |
| mammals | Thylamys_pusillus | Thylamys_pusillus | nocturnal |
| mammals | Thylamys_tatei | Thylamys_tatei | nocturnal |
| mammals | Thylamys_velutinus | Thylamys_velutinus | nocturnal |
| mammals | Thylamys_venustus | Thylamys_venustus | nocturnal |
| mammals | Thylogale_billardierii | Thylogale_billardierii | nocturnal |
| mammals | Thylogale_brownii | Thylogale_browni | nocturnal |
| mammals | Thylogale_brunii | Thylogale_brunii | nocturnal |
| mammals | Thylogale_calabyi | Thylogale_calabyi | nocturnal |
| mammals | Thylogale_stigmatica | Thylogale_stigmatica | cathemeral |
| mammals | Thylogale_thetis | Thylogale_thetis | cathemeral |
| mammals | Thyroptera_devivoi | Thyroptera_devivoi | nocturnal |
| mammals | Thyroptera_discifera | Thyroptera_discifera | nocturnal |
| mammals | Thyroptera_lavali | Thyroptera_lavali | nocturnal |
| mammals | Thyroptera_tricolor | Thyroptera_tricolor | nocturnal |
| mammals | Tlacuatzin_canescens | Tlacuatzin_canescens | nocturnal |
| mammals | Tokudaia_muenninki | Tokudaia_muenninki | nocturnal |
| mammals | Tokudaia_tokunoshimensis | Tokudaia_tokunoshimensis | nocturnal |
| mammals | Tolypeutes_matacus | Tolypeutes_matacus | nocturnal |
| mammals | Tolypeutes_tricinctus | Tolypeutes_tricinctus | nocturnal |
| mammals | Tomopeas_ravum | Tomopeas_ravus | nocturnal |
| mammals | Tonatia_bidens | Tonatia_bidens | nocturnal |
| mammals | Tonatia_saurophila | Tonatia_saurophila | nocturnal |
| mammals | Tonkinomys_daovantieni | Tonkinomys_daovantieni | nocturnal |
| mammals | Toromys_grandis | Toromys_grandis | nocturnal |
| mammals | Trachops_cirrhosus | Trachops_cirrhosus | nocturnal |
| mammals | Trachypithecus_auratus | Trachypithecus_auratus | diurnal |
| mammals | Trachypithecus_barbei | Trachypithecus_barbei | diurnal |
| mammals | Trachypithecus_cristatus | Trachypithecus_cristatus | diurnal |
| mammals | Trachypithecus_delacouri | Trachypithecus_delacouri | diurnal |
| mammals | Trachypithecus_francoisi | Trachypithecus_francoisi | diurnal |
| mammals | Trachypithecus_geei | Trachypithecus_geei | diurnal |
| mammals | Trachypithecus_germaini | Trachypithecus_germaini | diurnal |

| | | | |
|---|---|---|---|
| mammals | Trachypithecus_hatinhensis | Trachypithecus_hatinhensis | diurnal |
| mammals | Trachypithecus_laotum | Trachypithecus_laotum | diurnal |
| mammals | Trachypithecus_obscurus | Trachypithecus_obscurus | diurnal |
| mammals | Trachypithecus_phayrei | Trachypithecus_phayrei | diurnal |
| mammals | Trachypithecus_pileatus | Trachypithecus_pileatus | diurnal |
| mammals | Trachypithecus_poliocephalus | Trachypithecus_poliocephalus | diurnal |
| mammals | Trachypithecus_shortridgei | Trachypithecus_shortridgei | diurnal |
| mammals | Tragelaphus_angasii | Tragelaphus_angasii | diurnal |
| mammals | Tragelaphus_buxtoni | Tragelaphus_buxtoni | nocturnal |
| mammals | Tragelaphus_derbianus | Tragelaphus_derbianus | crepuscular |
| mammals | Tragelaphus_eurycerus | Tragelaphus_eurycerus | nocturnal |
| mammals | Tragelaphus_imberbis | Tragelaphus_imberbis | nocturnal |
| mammals | Tragelaphus_oryx | Tragelaphus_oryx | nocturnal |
| mammals | Tragelaphus_scriptus | Tragelaphus_scriptus | nocturnal |
| mammals | Tragelaphus_spekii | Tragelaphus_spekii | diurnal |
| mammals | Tragelaphus_strepsiceros | Tragelaphus_strepsiceros | cathemeral |
| mammals | Tragulus_javanicus | Tragulus_javanicus | nocturnal |
| mammals | Tragulus_kanchil | Tragulus_kanchil | cathemeral |
| mammals | Tragulus_napu | Tragulus_napu | cathemeral |
| mammals | Tragulus_nigricans | Tragulus_nigricans | nocturnal |
| mammals | Tragulus_versicolor | Tragulus_versicolor | nocturnal |
| mammals | Tragulus_williamsoni | Tragulus_williamsoni | nocturnal |
| mammals | Transandinomys_bolivaris | Transandinomys_bolivaris | nocturnal |
| mammals | Transandinomys_talamancae | Transandinomys_talamancae | nocturnal |
| mammals | Tremarctos_ornatus | Tremarctos_ornatus | diurnal |
| mammals | Triaenops_menamena | Triaenops_rufus | nocturnal |
| mammals | Triaenops_persicus | Triaenops_persicus | nocturnal |
| mammals | Trichosurus_caninus | Trichosurus_caninus | nocturnal |
| mammals | Trichosurus_cunninghami | Trichosurus_cunninghami | nocturnal |
| mammals | Trichosurus_vulpecula | Trichosurus_vulpecula | nocturnal |
| mammals | Trichys_fasciculata | Trichys_fasciculata | nocturnal |
| mammals | Trinomys_albispinus | Trinomys_albispinus | nocturnal |
| mammals | Trinomys_dimidiatus | Trinomys_dimidiatus | nocturnal |
| mammals | Trinomys_eliasi | Trinomys_eliasi | crepuscular |

| | | | |
|---|---|---|---|
| mammals | Trinomys_iheringi | Trinomys_iheringi | nocturnal |
| mammals | Trinomys_mirapitanga | Trinomys_mirapitanga | nocturnal |
| mammals | Trinomys_moojeni | Trinomys_moojeni | nocturnal |
| mammals | Trinomys_paratus | Trinomys_paratus | nocturnal |
| mammals | Trinomys_setosus | Trinomys_setosus | nocturnal |
| mammals | Trinomys_yonenagae | Trinomys_yonenagae | nocturnal |
| mammals | Trinycteris_nicefori | Trinycteris_nicefori | nocturnal |
| mammals | Trogopterus_xanthipes | Trogopterus_xanthipes | nocturnal |
| mammals | Tryphomys_adustus | Tryphomys_adustus | nocturnal |
| mammals | Tscherskia_triton | Tscherskia_triton | nocturnal |
| mammals | Tupaia_belangeri | Tupaia_belangeri | diurnal |
| mammals | Tupaia_chrysogaster | Tupaia_chrysogaster | diurnal |
| mammals | Tupaia_dorsalis | Tupaia_dorsalis | diurnal |
| mammals | Tupaia_everetti | Urogale_everetti | diurnal |
| mammals | Tupaia_glis | Tupaia_glis | diurnal |
| mammals | Tupaia_gracilis | Tupaia_gracilis | diurnal |
| mammals | Tupaia_javanica | Tupaia_javanica | diurnal |
| mammals | Tupaia_longipes | Tupaia_longipes | diurnal |
| mammals | Tupaia_minor | Tupaia_minor | diurnal |
| mammals | Tupaia_montana | Tupaia_montana | diurnal |
| mammals | Tupaia_nicobarica | Tupaia_nicobarica | diurnal |
| mammals | Tupaia_palawanensis | Tupaia_palawanensis | diurnal |
| mammals | Tupaia_picta | Tupaia_picta | diurnal |
| mammals | Tupaia_splendidula | Tupaia_splendidula | diurnal |
| mammals | Tupaia_tana | Tupaia_tana | diurnal |
| mammals | Tylomys_bullaris | Tylomys_bullaris | nocturnal |
| mammals | Tylomys_mirae | Tylomys_mirae | nocturnal |
| mammals | Tylomys_nudicaudus | Tylomys_nudicaudus | nocturnal |
| mammals | Tylomys_panamensis | Tylomys_panamensis | nocturnal |
| mammals | Tylonycteris_pachypus | Tylonycteris_pachypus | nocturnal |
| mammals | Tylonycteris_robustula | Tylonycteris_robustula | nocturnal |
| mammals | Tympanoctomys_barrerae | Tympanoctomys_barrerae | nocturnal |
| mammals | Tympanoctomys_kirchnerorum | Tympanoctomys_kirchnerorum | nocturnal |
| mammals | Typhlomys_cinereus | Typhlomys_cinereus | nocturnal |

| | | | |
|---|---|---|---|
| mammals | Typomys_planifrons | Hybomys_planifrons | diurnal |
| mammals | Typomys_trivirgatus | Hybomys_trivirgatus | cathemeral |
| mammals | Uranomys_ruddi | Uranomys_ruddi | nocturnal |
| mammals | Urocitellus_armatus | Urocitellus_armatus | diurnal |
| mammals | Urocitellus_beldingi | Urocitellus_beldingi | diurnal |
| mammals | Urocitellus_brunneus | Urocitellus_brunneus | diurnal |
| mammals | Urocitellus_canus | Urocitellus_canus | diurnal |
| mammals | Urocitellus_columbianus | Urocitellus_columbianus | diurnal |
| mammals | Urocitellus_elegans | Urocitellus_elegans | diurnal |
| mammals | Urocitellus_mollis | Urocitellus_mollis | diurnal |
| mammals | Urocitellus_parryii | Urocitellus_parryii | diurnal |
| mammals | Urocitellus_richardsonii | Urocitellus_richardsonii | diurnal |
| mammals | Urocitellus_townsendii | Urocitellus_townsendii | diurnal |
| mammals | Urocitellus_undulatus | Urocitellus_undulatus | diurnal |
| mammals | Urocitellus_washingtoni | Urocitellus_washingtoni | diurnal |
| mammals | Urocricetus_alticola | Cricetulus_alticola | nocturnal |
| mammals | Urocricetus_kamensis | Cricetulus_kamensis | cathemeral |
| mammals | Urocyon_cinereoargenteus | Urocyon_cinereoargenteus | nocturnal |
| mammals | Urocyon_littoralis | Urocyon_littoralis | nocturnal |
| mammals | Uroderma_bilobatum | Uroderma_bilobatum | nocturnal |
| mammals | Uroderma_magnirostrum | Uroderma_magnirostrum | nocturnal |
| mammals | Uromys_anak | Uromys_anak | nocturnal |
| mammals | Uromys_caudimaculatus | Uromys_caudimaculatus | nocturnal |
| mammals | Uromys_hadrourus | Uromys_hadrourus | nocturnal |
| mammals | Uromys_neobrittanicus | Uromys_neobritannicus | nocturnal |
| mammals | Uromys_rex | Uromys_rex | nocturnal |
| mammals | Uropsilus_andersoni | Uropsilus_andersoni | cathemeral |
| mammals | Uropsilus_gracilis | Uropsilus_gracilis | cathemeral |
| mammals | Uropsilus_investigator | Uropsilus_investigator | cathemeral |
| mammals | Uropsilus_soricipes | Uropsilus_soricipes | cathemeral |
| mammals | Urotrichus_talpoides | Urotrichus_talpoides | cathemeral |
| mammals | Ursus_americanus | Ursus_americanus | cathemeral |
| mammals | Ursus_arctos | Ursus_arctos | cathemeral |
| mammals | Ursus_maritimus | Ursus_maritimus | cathemeral |

| | | | |
|---|---|---|---|
| mammals | Ursus_thibetanus | Ursus_thibetanus | cathemeral |
| mammals | Urva_auropunctata | Herpestes_auropunctatus | diurnal |
| mammals | Urva_brachyura | Herpestes_brachyurus | diurnal |
| mammals | Urva_edwardsii | Herpestes_edwardsii | diurnal |
| mammals | Urva_fusca | Herpestes_fuscus | nocturnal |
| mammals | Urva_javanica | Herpestes_javanicus | nocturnal |
| mammals | Urva_semitorquata | Herpestes_semitorquatus | diurnal |
| mammals | Urva_smithii | Herpestes_smithii | diurnal |
| mammals | Urva_urva | Herpestes_urva | diurnal |
| mammals | Urva_vitticollis | Herpestes_vitticollis | diurnal |
| mammals | Vampyressa_melissa | Vampyressa_melissa | nocturnal |
| mammals | Vampyressa_pusilla | Vampyressa_pusilla | nocturnal |
| mammals | Vampyressa_thyone | Vampyressa_thyone | nocturnal |
| mammals | Vampyriscus_bidens | Vampyriscus_bidens | nocturnal |
| mammals | Vampyriscus_brocki | Vampyriscus_brocki | nocturnal |
| mammals | Vampyriscus_nymphaeus | Vampyriscus_nymphaea | nocturnal |
| mammals | Vampyrodes_caraccioli | Vampyrodes_caraccioli | nocturnal |
| mammals | Vampyrodes_major | Vampyrodes_major | nocturnal |
| mammals | Vampyrum_spectrum | Vampyrum_spectrum | nocturnal |
| mammals | Vandeleuria_nolthenii | Vandeleuria_nolthenii | nocturnal |
| mammals | Vandeleuria_oleraceus | Vandeleuria_oleracea | nocturnal |
| mammals | Vansonia_rueppellii | Pipistrellus_rueppellii | nocturnal |
| mammals | Varecia_rubra | Varecia_rubra | diurnal |
| mammals | Varecia_variegata | Varecia_variegata | diurnal |
| mammals | Vernaya_fulva | Vernaya_fulva | nocturnal |
| mammals | Vespadelus_baverstocki | Vespadelus_baverstocki | nocturnal |
| mammals | Vespadelus_caurinus | Vespadelus_caurinus | nocturnal |
| mammals | Vespadelus_darlingtoni | Vespadelus_darlingtoni | nocturnal |
| mammals | Vespadelus_douglasorum | Vespadelus_douglasorum | nocturnal |
| mammals | Vespadelus_finlaysoni | Vespadelus_finlaysoni | nocturnal |
| mammals | Vespadelus_pumilus | Vespadelus_pumilus | nocturnal |
| mammals | Vespadelus_regulus | Vespadelus_regulus | nocturnal |
| mammals | Vespadelus_troughtoni | Vespadelus_troughtoni | nocturnal |
| mammals | Vespadelus_vulturnus | Vespadelus_vulturnus | nocturnal |

| | | | |
|---|---|---|---|
| mammals | Vespertilio_murinus | Vespertilio_murinus | nocturnal |
| mammals | Vespertilio_sinensis | Vespertilio_sinensis | nocturnal |
| mammals | Viverra_megaspila | Viverra_megaspila | nocturnal |
| mammals | Viverra_tangalunga | Viverra_tangalunga | nocturnal |
| mammals | Viverra_zibetha | Viverra_zibetha | nocturnal |
| mammals | Viverricula_indica | Viverricula_indica | nocturnal |
| mammals | Voalavo_antsahabensis | Voalavo_antsahabensis | nocturnal |
| mammals | Voalavo_gymnocaudus | Voalavo_gymnocaudus | nocturnal |
| mammals | Volemys_musseri | Volemys_musseri | cathemeral |
| mammals | Vombatus_ursinus | Vombatus_ursinus | nocturnal |
| mammals | Vormela_peregusna | Vormela_peregusna | nocturnal |
| mammals | Vulpes_bengalensis | Vulpes_bengalensis | nocturnal |
| mammals | Vulpes_cana | Vulpes_cana | nocturnal |
| mammals | Vulpes_chama | Vulpes_chama | nocturnal |
| mammals | Vulpes_corsac | Vulpes_corsac | nocturnal |
| mammals | Vulpes_ferrilata | Vulpes_ferrilata | nocturnal |
| mammals | Vulpes_lagopus | Vulpes_lagopus | nocturnal |
| mammals | Vulpes_macrotis | Vulpes_macrotis | nocturnal |
| mammals | Vulpes_pallida | Vulpes_pallida | nocturnal |
| mammals | Vulpes_rueppellii | Vulpes_rueppellii | nocturnal |
| mammals | Vulpes_velox | Vulpes_velox | nocturnal |
| mammals | Vulpes_vulpes | Vulpes_vulpes | cathemeral |
| mammals | Vulpes_zerda | Vulpes_zerda | nocturnal |
| mammals | Wallabia_bicolor | Wallabia_bicolor | nocturnal |
| mammals | Wiedomys_cerradensis | Wiedomys_cerradensis | nocturnal |
| mammals | Wiedomys_pyrrhorinos | Wiedomys_pyrrhorhinos | nocturnal |
| mammals | Wilfredomys_oenax | Wilfredomys_oenax | nocturnal |
| mammals | Wyulda_squamicaudata | Wyulda_squamicaudata | nocturnal |
| mammals | Xanthonycticebus_pygmaeus | Nycticebus_pygmaeus | nocturnal |
| mammals | Xenogale_naso | Herpestes_naso | diurnal |
| mammals | Xenomys_nelsoni | Xenomys_nelsoni | nocturnal |
| mammals | Xenuromys_barbatus | Xenuromys_barbatus | cathemeral |
| mammals | Xeromys_myoides | Xeromys_myoides | nocturnal |
| mammals | Xeronycteris_vieirai | Xeronycteris_vieirai | nocturnal |

| | | | |
|---|---|---|---|
| mammals | Xerospermophilus_mohavensis | Xerospermophilus_mohavensis | diurnal |
| mammals | Xerospermophilus_perotensis | Xerospermophilus_perotensis | diurnal |
| mammals | Xerospermophilus_spilosoma | Xerospermophilus_spilosoma | diurnal |
| mammals | Xerospermophilus_tereticaudus | Xerospermophilus_tereticaudus | diurnal |
| mammals | Xerus_rutilus | Xerus_rutilus | diurnal |
| mammals | Zaedyus_pichiy | Zaedyus_pichiy | diurnal |
| mammals | Zaglossus_attenboroughi | Zaglossus_attenboroughi | nocturnal |
| mammals | Zaglossus_bartoni | Zaglossus_bartoni | nocturnal |
| mammals | Zaglossus_bruijnii | Zaglossus_bruijnii | nocturnal |
| mammals | Zapus_hudsonius | Zapus_hudsonius | nocturnal |
| mammals | Zapus_princeps | Zapus_princeps | nocturnal |
| mammals | Zapus_trinotatus | Zapus_trinotatus | nocturnal |
| mammals | Zelotomys_hildegardeae | Zelotomys_hildegardeae | nocturnal |
| mammals | Zelotomys_woosnami | Zelotomys_woosnami | nocturnal |
| mammals | Zenkerella_insignis | Zenkerella_insignis | cathemeral |
| mammals | Zygodontomys_brevicauda | Zygodontomys_brevicauda | nocturnal |
| mammals | Zygodontomys_brunneus | Zygodontomys_brunneus | nocturnal |
| mammals | Zyzomys_argurus | Zyzomys_argurus | nocturnal |
| mammals | Zyzomys_maini | Zyzomys_maini | nocturnal |
| mammals | Zyzomys_palatalis | Zyzomys_palatalis | nocturnal |
| mammals | Zyzomys_pedunculatus | Zyzomys_pedunculatus | nocturnal |
| mammals | Zyzomys_woodwardi | Zyzomys_woodwardi | nocturnal |
| birds | Abeillia_abeillei | Abeillia_abeillei | diurnal |
| birds | Abroscopus_albogularis | Abroscopus_albogularis | diurnal |
| birds | Abroscopus_schisticeps | Abroscopus_schisticeps | diurnal |
| birds | Abroscopus_superciliaris | Abroscopus_superciliaris | diurnal |
| birds | Aburria_aburri | Aburria_aburri | diurnal |
| birds | Acanthagenys_rufogularis | Acanthagenys_rufogularis | diurnal |
| birds | Acanthidops_bairdi | Acanthidops_bairdii | diurnal |
| birds | Acanthis_flammea | Carduelis_flammea | diurnal |
| birds | Acanthisitta_chloris | Acanthisitta_chloris | diurnal |
| birds | Acanthiza_chrysorrhoa | Acanthiza_chrysorrhoa | diurnal |
| birds | Acanthiza_cinerea | Gerygone_cinerea | diurnal |
| birds | Acanthiza_ewingii | Acanthiza_ewingii | diurnal |

| | | | |
|---|---|---|---|
| birds | Acanthiza_inornata | Acanthiza_inornata | diurnal |
| birds | Acanthiza_iredalei | Acanthiza_iredalei | diurnal |
| birds | Acanthiza_katherina | Acanthiza_katherina | diurnal |
| birds | Acanthiza_lineata | Acanthiza_lineata | diurnal |
| birds | Acanthiza_murina | Acanthiza_murina | diurnal |
| birds | Acanthiza_nana | Acanthiza_nana | diurnal |
| birds | Acanthiza_pusilla | Acanthiza_pusilla | diurnal |
| birds | Acanthiza_reguloides | Acanthiza_reguloides | diurnal |
| birds | Acanthiza_robustirostris | Acanthiza_robustirostris | diurnal |
| birds | Acanthiza_uropygialis | Acanthiza_uropygialis | diurnal |
| birds | Acanthorhynchus_superciliosus | Acanthorhynchus_superciliosus | diurnal |
| birds | Acanthorhynchus_tenuirostris | Acanthorhynchus_tenuirostris | diurnal |
| birds | Acanthornis_magna | Acanthornis_magna | diurnal |
| birds | Accipiter_albogularis | Accipiter_albogularis | diurnal |
| birds | Accipiter_badius | Accipiter_badius | diurnal |
| birds | Accipiter_bicolor | Accipiter_bicolor | diurnal |
| birds | Accipiter_brachyurus | Accipiter_brachyurus | diurnal |
| birds | Accipiter_brevipes | Accipiter_brevipes | diurnal |
| birds | Accipiter_butleri | Accipiter_butleri | diurnal |
| birds | Accipiter_castanilius | Accipiter_castanilius | diurnal |
| birds | Accipiter_cirrocephalus | Accipiter_cirrocephalus | diurnal |
| birds | Accipiter_collaris | Accipiter_collaris | diurnal |
| birds | Accipiter_cooperii | Accipiter_cooperii | diurnal |
| birds | Accipiter_erythrauchen | Accipiter_erythrauchen | diurnal |
| birds | Accipiter_erythropus | Accipiter_erythropus | diurnal |
| birds | Accipiter_fasciatus | Accipiter_fasciatus | diurnal |
| birds | Accipiter_francesiae | Accipiter_francesiae | diurnal |
| birds | Accipiter_gentilis | Accipiter_gentilis | diurnal |
| birds | Accipiter_griseiceps | Accipiter_griseiceps | diurnal |
| birds | Accipiter_gularis | Accipiter_gularis | diurnal |
| birds | Accipiter_gundlachi | Accipiter_gundlachi | diurnal |
| birds | Accipiter_haplochrous | Accipiter_haplochrous | diurnal |
| birds | Accipiter_henstii | Accipiter_henstii | diurnal |
| birds | Accipiter_imitator | Accipiter_imitator | diurnal |

| | | | |
|---|---|---|---|
| birds | Accipiter_luteoschistaceus | Accipiter_luteoschistaceus | diurnal |
| birds | Accipiter_madagascariensis | Accipiter_madagascariensis | diurnal |
| birds | Accipiter_melanochlamys | Accipiter_melanochlamys | diurnal |
| birds | Accipiter_melanoleucus | Accipiter_melanoleucus | diurnal |
| birds | Accipiter_meyerianus | Accipiter_meyerianus | diurnal |
| birds | Accipiter_minullus | Accipiter_minullus | diurnal |
| birds | Accipiter_nisus | Accipiter_nisus | diurnal |
| birds | Accipiter_novaehollandiae | Accipiter_novaehollandiae | diurnal |
| birds | Accipiter_ovampensis | Accipiter_ovampensis | diurnal |
| birds | Accipiter_poliocephalus | Accipiter_poliocephalus | diurnal |
| birds | Accipiter_poliogaster | Accipiter_poliogaster | diurnal |
| birds | Accipiter_rhodogaster | Accipiter_rhodogaster | diurnal |
| birds | Accipiter_rufitorques | Accipiter_rufitorques | diurnal |
| birds | Accipiter_rufiventris | Accipiter_rufiventris | diurnal |
| birds | Accipiter_soloensis | Accipiter_soloensis | diurnal |
| birds | Accipiter_striatus | Accipiter_striatus | diurnal |
| birds | Accipiter_superciliosus | Accipiter_superciliosus | diurnal |
| birds | Accipiter_trinotatus | Accipiter_trinotatus | diurnal |
| birds | Accipiter_trivirgatus | Accipiter_trivirgatus | diurnal |
| birds | Accipiter_virgatus | Accipiter_virgatus | diurnal |
| birds | Aceros_nipalensis | Aceros_nipalensis | diurnal |
| birds | Achaetops_pycnopygius | Achaetops_pycnopygius | diurnal |
| birds | Acridotheres_cristatellus | Acridotheres_cristatellus | diurnal |
| birds | Acridotheres_fuscus | Acridotheres_fuscus | diurnal |
| birds | Acridotheres_ginginianus | Acridotheres_ginginianus | diurnal |
| birds | Acridotheres_grandis | Acridotheres_grandis | diurnal |
| birds | Acridotheres_melanopterus | Sturnus_melanopterus | diurnal |
| birds | Acridotheres_tristis | Acridotheres_tristis | diurnal |
| birds | Acritillas_indica | Iole_indica | diurnal |
| birds | Acrobatornis_fonsecai | Acrobatornis_fonsecai | diurnal |
| birds | Acrocephalus_aequinoctialis | Acrocephalus_aequinoctialis | diurnal |
| birds | Acrocephalus_agricola | Acrocephalus_agricola | diurnal |
| birds | Acrocephalus_arundinaceus | Acrocephalus_arundinaceus | diurnal |
| birds | Acrocephalus_atyphus | Acrocephalus_atyphus | diurnal |

| | | | |
|---|---|---|---|
| birds | Acrocephalus_australis | Acrocephalus_australis | diurnal |
| birds | Acrocephalus_bistrigiceps | Acrocephalus_bistrigiceps | diurnal |
| birds | Acrocephalus_brevipennis | Acrocephalus_brevipennis | diurnal |
| birds | Acrocephalus_caffer | Acrocephalus_caffer | diurnal |
| birds | Acrocephalus_concinens | Acrocephalus_concinens | diurnal |
| birds | Acrocephalus_dumetorum | Acrocephalus_dumetorum | diurnal |
| birds | Acrocephalus_familiaris | Acrocephalus_familiaris | diurnal |
| birds | Acrocephalus_gracilirostris | Acrocephalus_gracilirostris | diurnal |
| birds | Acrocephalus_griseldis | Acrocephalus_griseldis | diurnal |
| birds | Acrocephalus_kerearako | Acrocephalus_kerearako | diurnal |
| birds | Acrocephalus_melanopogon | Acrocephalus_melanopogon | diurnal |
| birds | Acrocephalus_mendanae | Acrocephalus_mendanae | diurnal |
| birds | Acrocephalus_newtoni | Acrocephalus_newtoni | diurnal |
| birds | Acrocephalus_paludicola | Acrocephalus_paludicola | diurnal |
| birds | Acrocephalus_palustris | Acrocephalus_palustris | diurnal |
| birds | Acrocephalus_rehsei | Acrocephalus_rehsei | diurnal |
| birds | Acrocephalus_rimitarae | Acrocephalus_rimatarae | diurnal |
| birds | Acrocephalus_rodericanus | Acrocephalus_rodericanus | diurnal |
| birds | Acrocephalus_rufescens | Acrocephalus_rufescens | diurnal |
| birds | Acrocephalus_schoenobaenus | Acrocephalus_schoenobaenus | diurnal |
| birds | Acrocephalus_scirpaceus | Acrocephalus_scirpaceus | diurnal |
| birds | Acrocephalus_sechellensis | Acrocephalus_sechellensis | diurnal |
| birds | Acrocephalus_sorghophilus | Acrocephalus_sorghophilus | diurnal |
| birds | Acrocephalus_stentoreus | Acrocephalus_stentoreus | diurnal |
| birds | Acrocephalus_syrinx | Acrocephalus_syrinx | diurnal |
| birds | Acrocephalus_taiti | Acrocephalus_taiti | diurnal |
| birds | Acrocephalus_tangorum | Acrocephalus_tangorum | diurnal |
| birds | Acrocephalus_vaughani | Acrocephalus_vaughani | diurnal |
| birds | Acropternis_orthonyx | Acropternis_orthonyx | diurnal |
| birds | Acryllium_vulturinum | Acryllium_vulturinum | diurnal |
| birds | Actenoides_bougainvillei | Actenoides_bougainvillei | diurnal |
| birds | Actenoides_concretus | Actenoides_concretus | diurnal |
| birds | Actenoides_hombroni | Actenoides_hombroni | diurnal |
| birds | Actenoides_lindsayi | Actenoides_lindsayi | diurnal |

| | | | |
|---|---|---|---|
| birds | Actenoides_monachus | Actenoides_monachus | diurnal |
| birds | Actenoides_princeps | Actenoides_princeps | diurnal |
| birds | Actinodura_egertoni | Actinodura_egertoni | diurnal |
| birds | Actinodura_ramsayi | Actinodura_ramsayi | diurnal |
| birds | Actitis_hypoleucos | Actitis_hypoleucos | diurnal |
| birds | Actitis_macularius | Actitis_macularius | diurnal |
| birds | Actophilornis_africanus | Actophilornis_africanus | diurnal |
| birds | Actophilornis_albinucha | Actophilornis_albinucha | diurnal |
| birds | Adelomyia_melanogenys | Adelomyia_melanogenys | diurnal |
| birds | Aechmophorus_clarkii | Aechmophorus_clarkii | diurnal |
| birds | Aechmophorus_occidentalis | Aechmophorus_occidentalis | diurnal |
| birds | Aegithalos_caudatus | Aegithalos_caudatus | diurnal |
| birds | Aegithalos_concinnus | Aegithalos_concinnus | diurnal |
| birds | Aegithalos_fuliginosus | Aegithalos_fuliginosus | diurnal |
| birds | Aegithalos_iouschistos | Aegithalos_iouschistos | diurnal |
| birds | Aegithalos_leucogenys | Aegithalos_leucogenys | diurnal |
| birds | Aegithalos_niveogularis | Aegithalos_niveogularis | diurnal |
| birds | Aegithina_lafresnayei | Aegithina_lafresnayei | diurnal |
| birds | Aegithina_nigrolutea | Aegithina_nigrolutea | diurnal |
| birds | Aegithina_tiphia | Aegithina_tiphia | diurnal |
| birds | Aegithina_viridissima | Aegithina_viridissima | diurnal |
| birds | Aegolius_acadicus | Aegolius_acadicus | nocturnal |
| birds | Aegolius_funereus | Aegolius_funereus | cathemeral |
| birds | Aegolius_harrisii | Aegolius_harrisii | nocturnal |
| birds | Aegolius_ridgwayi | Aegolius_ridgwayi | nocturnal |
| birds | Aegotheles_albertisi | Aegotheles_albertisi | nocturnal |
| birds | Aegotheles_archboldi | Aegotheles_archboldi | nocturnal |
| birds | Aegotheles_bennettii | Aegotheles_bennettii | nocturnal |
| birds | Aegotheles_crinifrons | Aegotheles_crinifrons | nocturnal |
| birds | Aegotheles_cristatus | Aegotheles_cristatus | nocturnal |
| birds | Aegotheles_insignis | Aegotheles_insignis | nocturnal |
| birds | Aegotheles_tatei | Aegotheles_tatei | nocturnal |
| birds | Aegotheles_wallacii | Aegotheles_wallacii | nocturnal |
| birds | Aegypius_monachus | Aegypius_monachus | diurnal |

| | | | |
|---|---|---|---|
| birds | Aepypodius_arfakianus | Aepypodius_arfakianus | diurnal |
| birds | Aepypodius_bruijnii | Aepypodius_bruijnii | diurnal |
| birds | Aerodramus_elaphrus | Collocalia_elaphra | diurnal |
| birds | Aerodramus_francicus | Collocalia_francica | diurnal |
| birds | Aerodramus_hirundinaceus | Collocalia_hirundinacea | diurnal |
| birds | Aerodramus_infuscatus | Collocalia_infuscata | diurnal |
| birds | Aerodramus_inquietus | Collocalia_inquieta | diurnal |
| birds | Aerodramus_leucophaeus | Collocalia_leucophaea | diurnal |
| birds | Aerodramus_maximus | Collocalia_maxima | nocturnal |
| birds | Aerodramus_mearnsi | Collocalia_mearnsi | diurnal |
| birds | Aerodramus_papuensis | Collocalia_papuensis | nocturnal |
| birds | Aerodramus_spodiopygius | Collocalia_spodiopygia | diurnal |
| birds | Aerodramus_terraereginae | Collocalia_terraereginae | diurnal |
| birds | Aerodramus_unicolor | Collocalia_unicolor | cathemeral |
| birds | Aerodramus_vanikorensis | Collocalia_vanikorensis | diurnal |
| birds | Aeronautes_andecolus | Aeronautes_andecolus | diurnal |
| birds | Aeronautes_montivagus | Aeronautes_montivagus | diurnal |
| birds | Aeronautes_saxatalis | Aeronautes_saxatalis | diurnal |
| birds | Aethia_cristatella | Aethia_cristatella | diurnal |
| birds | Aethia_psittacula | Aethia_psittacula | diurnal |
| birds | Aethia_pusilla | Aethia_pusilla | diurnal |
| birds | Aethia_pygmaea | Aethia_pygmaea | diurnal |
| birds | Aethomyias_arfakianus | Sericornis_arfakianus | diurnal |
| birds | Aethomyias_nigrorufus | Crateroscelis_nigrorufa | diurnal |
| birds | Aethomyias_papuensis | Sericornis_papuensis | diurnal |
| birds | Aethomyias_perspicillatus | Sericornis_perspicillatus | diurnal |
| birds | Aethomyias_rufescens | Sericornis_rufescens | diurnal |
| birds | Aethomyias_spilodera | Sericornis_spilodera | diurnal |
| birds | Aethopyga_boltoni | Aethopyga_boltoni | diurnal |
| birds | Aethopyga_christinae | Aethopyga_christinae | diurnal |
| birds | Aethopyga_duyvenbodei | Aethopyga_duyvenbodei | diurnal |
| birds | Aethopyga_flagrans | Aethopyga_flagrans | diurnal |
| birds | Aethopyga_gouldiae | Aethopyga_gouldiae | diurnal |
| birds | Aethopyga_ignicauda | Aethopyga_ignicauda | diurnal |

| | | | |
|---|---|---|---|
| birds | Aethopyga_mystacalis | Aethopyga_mystacalis | diurnal |
| birds | Aethopyga_nipalensis | Aethopyga_nipalensis | diurnal |
| birds | Aethopyga_primigenia | Aethopyga_primigenia | diurnal |
| birds | Aethopyga_pulcherrima | Aethopyga_pulcherrima | diurnal |
| birds | Aethopyga_saturata | Aethopyga_saturata | diurnal |
| birds | Aethopyga_shelleyi | Aethopyga_shelleyi | diurnal |
| birds | Aethopyga_siparaja | Aethopyga_siparaja | diurnal |
| birds | Afropavo_congensis | Afropavo_congensis | diurnal |
| birds | Afrotis_afra | Eupodotis_afra | diurnal |
| birds | Afrotis_afraoides | Eupodotis_afraoides | diurnal |
| birds | Agamia_agami | Agamia_agami | diurnal |
| birds | Agapornis_canus | Agapornis_canus | diurnal |
| birds | Agapornis_fischeri | Agapornis_fischeri | diurnal |
| birds | Agapornis_lilianae | Agapornis_lilianae | diurnal |
| birds | Agapornis_nigrigenis | Agapornis_nigrigenis | diurnal |
| birds | Agapornis_personatus | Agapornis_personatus | diurnal |
| birds | Agapornis_pullarius | Agapornis_pullarius | diurnal |
| birds | Agapornis_roseicollis | Agapornis_roseicollis | diurnal |
| birds | Agapornis_swindernianus | Agapornis_swindernianus | diurnal |
| birds | Agapornis_taranta | Agapornis_taranta | diurnal |
| birds | Agelaioides_badius | Agelaioides_badius | diurnal |
| birds | Agelaius_assimilis | Agelaius_assimilis | diurnal |
| birds | Agelaius_humeralis | Agelaius_humeralis | diurnal |
| birds | Agelaius_tricolor | Agelaius_tricolor | diurnal |
| birds | Agelaius_xanthomus | Agelaius_xanthomus | diurnal |
| birds | Agelastes_meleagrides | Agelastes_meleagrides | diurnal |
| birds | Agelastes_niger | Agelastes_niger | diurnal |
| birds | Agelasticus_cyanopus | Agelasticus_cyanopus | diurnal |
| birds | Agelasticus_thilius | Agelasticus_thilius | diurnal |
| birds | Agelasticus_xanthophthalmus | Agelasticus_xanthophthalmus | diurnal |
| birds | Aglaeactis_aliciae | Aglaeactis_aliciae | diurnal |
| birds | Aglaeactis_castelnaudii | Aglaeactis_castelnaudii | diurnal |
| birds | Aglaeactis_cupripennis | Aglaeactis_cupripennis | diurnal |
| birds | Aglaeactis_pamela | Aglaeactis_pamela | diurnal |

| | | | |
|---|---|---|---|
| birds | Aglaiocercus_berlepschi | Aglaiocercus_berlepschi | diurnal |
| birds | Aglaiocercus_coelestis | Aglaiocercus_coelestis | diurnal |
| birds | Aglaiocercus_kingii | Aglaiocercus_kingi | diurnal |
| birds | Agricola_infuscatus | Bradornis_infuscatus | diurnal |
| birds | Agricola_pallidus | Bradornis_pallidus | diurnal |
| birds | Agriornis_albicauda | Agriornis_albicauda | diurnal |
| birds | Agriornis_lividus | Agriornis_lividus | diurnal |
| birds | Agriornis_micropterus | Agriornis_micropterus | diurnal |
| birds | Agriornis_montanus | Agriornis_montanus | diurnal |
| birds | Agriornis_murinus | Agriornis_murinus | diurnal |
| birds | Agropsar_philippensis | Sturnus_philippensis | diurnal |
| birds | Agropsar_sturninus | Sturnus_sturninus | diurnal |
| birds | Ailuroedus_buccoides | Ailuroedus_buccoides | diurnal |
| birds | Ailuroedus_crassirostris | Ailuroedus_crassirostris | diurnal |
| birds | Ailuroedus_melanotis | Ailuroedus_melanotis | diurnal |
| birds | Aimophila_notosticta | Aimophila_notosticta | diurnal |
| birds | Aimophila_rufescens | Aimophila_rufescens | diurnal |
| birds | Aimophila_ruficeps | Aimophila_ruficeps | diurnal |
| birds | Aix_galericulata | Aix_galericulata | diurnal |
| birds | Aix_sponsa | Aix_sponsa | diurnal |
| birds | Akletos_goeldii | Myrmeciza_goeldii | diurnal |
| birds | Akletos_melanoceps | Myrmeciza_melanoceps | diurnal |
| birds | Alaemon_alaudipes | Alaemon_alaudipes | diurnal |
| birds | Alauda_arvensis | Alauda_arvensis | diurnal |
| birds | Alauda_gulgula | Alauda_gulgula | diurnal |
| birds | Alauda_leucoptera | Melanocorypha_leucoptera | diurnal |
| birds | Alauda_razae | Alauda_razae | diurnal |
| birds | Alaudala_raytal | Calandrella_raytal | diurnal |
| birds | Alaudala_rufescens | Calandrella_cheleensis | diurnal |
| birds | Alca_torda | Alca_torda | diurnal |
| birds | Alcedo_atthis | Alcedo_atthis | cathemeral |
| birds | Alcedo_euryzona | Alcedo_euryzona | diurnal |
| birds | Alcedo_hercules | Alcedo_hercules | diurnal |
| birds | Alcedo_meninting | Alcedo_meninting | diurnal |

| | | | |
|---|---|---|---|
| birds | Alcedo_quadribrachys | Alcedo_quadribrachys | diurnal |
| birds | Alcedo_semitorquata | Alcedo_semitorquata | diurnal |
| birds | Alcippe_brunneicauda | Alcippe_brunneicauda | diurnal |
| birds | Alcippe_morrisonia | Alcippe_morrisonia | diurnal |
| birds | Alcippe_nipalensis | Alcippe_nipalensis | diurnal |
| birds | Alcippe_peracensis | Alcippe_peracensis | diurnal |
| birds | Alcippe_poioicephala | Alcippe_poioicephala | diurnal |
| birds | Alcippe_pyrrhoptera | Alcippe_pyrrhoptera | diurnal |
| birds | Alcurus_striatus | Pycnonotus_striatus | diurnal |
| birds | Aleadryas_rufinucha | Aleadryas_rufinucha | diurnal |
| birds | Alectoris_barbara | Alectoris_barbara | diurnal |
| birds | Alectoris_chukar | Alectoris_chukar | diurnal |
| birds | Alectoris_graeca | Alectoris_graeca | diurnal |
| birds | Alectoris_magna | Alectoris_magna | diurnal |
| birds | Alectoris_philbyi | Alectoris_philbyi | diurnal |
| birds | Alectoris_rufa | Alectoris_rufa | diurnal |
| birds | Alectroenas_madagascariensis | Alectroenas_madagascariensis | diurnal |
| birds | Alectroenas_pulcherrimus | Alectroenas_pulcherrima | diurnal |
| birds | Alectroenas_sganzini | Alectroenas_sganzini | diurnal |
| birds | Alectrurus_risora | Alectrurus_risora | diurnal |
| birds | Alectrurus_tricolor | Alectrurus_tricolor | diurnal |
| birds | Alectura_lathami | Alectura_lathami | diurnal |
| birds | Alethe_diademata | Alethe_diademata | diurnal |
| birds | Alexandrinus_eques | Psittacula_eques | diurnal |
| birds | Alexandrinus_krameri | Psittacula_krameri | diurnal |
| birds | Alisterus_amboinensis | Alisterus_amboinensis | diurnal |
| birds | Alisterus_chloropterus | Alisterus_chloropterus | diurnal |
| birds | Alisterus_scapularis | Alisterus_scapularis | diurnal |
| birds | Alle_alle | Alle_alle | diurnal |
| birds | Allenia_fusca | Margarops_fuscus | diurnal |
| birds | Alophoixus_bres | Alophoixus_bres | diurnal |
| birds | Alophoixus_flaveolus | Alophoixus_flaveolus | diurnal |
| birds | Alophoixus_ochraceus | Alophoixus_ochraceus | diurnal |
| birds | Alophoixus_pallidus | Alophoixus_pallidus | diurnal |

| | | | |
|---|---|---|---|
| birds | Alophoixus_phaeocephalus | Alophoixus_phaeocephalus | diurnal |
| birds | Alopochelidon_fucata | Alopochelidon_fucata | diurnal |
| birds | Alopochen_aegyptiaca | Alopochen_aegyptiaca | diurnal |
| birds | Amadina_erythrocephala | Amadina_erythrocephala | diurnal |
| birds | Amadina_fasciata | Amadina_fasciata | diurnal |
| birds | Amalocichla_incerta | Amalocichla_incerta | diurnal |
| birds | Amalocichla_sclateriana | Amalocichla_sclateriana | diurnal |
| birds | Amandava_amandava | Amandava_amandava | diurnal |
| birds | Amandava_formosa | Amandava_formosa | diurnal |
| birds | Amandava_subflava | Amandava_subflava | diurnal |
| birds | Amaurocichla_bocagii | Amaurocichla_bocagei | diurnal |
| birds | Amaurolimnas_concolor | Amaurolimnas_concolor | diurnal |
| birds | Amaurornis_cinerea | Porzana_cinerea | diurnal |
| birds | Amaurornis_moluccana | Amaurornis_moluccana | diurnal |
| birds | Amaurornis_olivacea | Amaurornis_olivacea | diurnal |
| birds | Amaurornis_phoenicurus | Amaurornis_phoenicurus | diurnal |
| birds | Amaurospiza_moesta | Amaurospiza_moesta | diurnal |
| birds | Amaurospiza_relicta | Amaurospiza_concolor | diurnal |
| birds | Amazilia_boucardi | Amazilia_boucardi | diurnal |
| birds | Amazilia_cyanifrons | Amazilia_cyanifrons | diurnal |
| birds | Amazilia_franciae | Amazilia_franciae | diurnal |
| birds | Amazilia_rutila | Amazilia_rutila | diurnal |
| birds | Amazilia_tzacatl | Amazilia_tzacatl | diurnal |
| birds | Amazilia_yucatanensis | Amazilia_yucatanensis | diurnal |
| birds | Amazilis_amazilia | Amazilia_amazilia | diurnal |
| birds | Amazona_aestiva | Amazona_aestiva | diurnal |
| birds | Amazona_agilis | Amazona_agilis | diurnal |
| birds | Amazona_albifrons | Amazona_albifrons | diurnal |
| birds | Amazona_amazonica | Amazona_amazonica | diurnal |
| birds | Amazona_arausiaca | Amazona_arausiaca | diurnal |
| birds | Amazona_autumnalis | Amazona_autumnalis | diurnal |
| birds | Amazona_barbadensis | Amazona_barbadensis | diurnal |
| birds | Amazona_brasiliensis | Amazona_brasiliensis | diurnal |
| birds | Amazona_collaria | Amazona_collaria | diurnal |

| | | | |
|---|---|---|---|
| birds | Amazona_dufresniana | Amazona_dufresniana | diurnal |
| birds | Amazona_farinosa | Amazona_farinosa | diurnal |
| birds | Amazona_festiva | Amazona_festiva | diurnal |
| birds | Amazona_finschi | Amazona_finschi | diurnal |
| birds | Amazona_guildingii | Amazona_guildingii | diurnal |
| birds | Amazona_imperialis | Amazona_imperialis | diurnal |
| birds | Amazona_leucocephala | Amazona_leucocephala | diurnal |
| birds | Amazona_mercenarius | Amazona_mercenaria | diurnal |
| birds | Amazona_ochrocephala | Amazona_ochrocephala | diurnal |
| birds | Amazona_pretrei | Amazona_pretrei | diurnal |
| birds | Amazona_rhodocorytha | Amazona_rhodocorytha | diurnal |
| birds | Amazona_tucumana | Amazona_tucumana | diurnal |
| birds | Amazona_ventralis | Amazona_ventralis | diurnal |
| birds | Amazona_versicolor | Amazona_versicolor | diurnal |
| birds | Amazona_vinacea | Amazona_vinacea | diurnal |
| birds | Amazona_viridigenalis | Amazona_viridigenalis | diurnal |
| birds | Amazona_vittata | Amazona_vittata | diurnal |
| birds | Amazona_xantholora | Amazona_xantholora | diurnal |
| birds | Amazonetta_brasiliensis | Amazonetta_brasiliensis | diurnal |
| birds | Amblycercus_holosericeus | Amblycercus_holosericeus | diurnal |
| birds | Amblyornis_flavifrons | Amblyornis_flavifrons | diurnal |
| birds | Amblyornis_inornata | Amblyornis_inornata | diurnal |
| birds | Amblyornis_macgregoriae | Amblyornis_macgregoriae | diurnal |
| birds | Amblyornis_subalaris | Amblyornis_subalaris | diurnal |
| birds | Amblyospiza_albifrons | Amblyospiza_albifrons | diurnal |
| birds | Amblyramphus_holosericeus | Amblyramphus_holosericeus | diurnal |
| birds | Ammodramus_aurifrons | Ammodramus_aurifrons | diurnal |
| birds | Ammodramus_humeralis | Ammodramus_humeralis | diurnal |
| birds | Ammodramus_savannarum | Ammodramus_savannarum | diurnal |
| birds | Ammomanes_cinctura | Ammomanes_cinctura | diurnal |
| birds | Ammomanes_deserti | Ammomanes_deserti | diurnal |
| birds | Ammomanes_phoenicura | Ammomanes_phoenicura | diurnal |
| birds | Ammonastes_pelzelni | Myrmeciza_pelzelni | diurnal |
| birds | Ammoperdix_griseogularis | Ammoperdix_griseogularis | diurnal |

| | | | |
|---|---|---|---|
| birds | Ammoperdix_heyi | Ammoperdix_heyi | diurnal |
| birds | Ammospiza_caudacuta | Ammodramus_caudacutus | diurnal |
| birds | Ammospiza_leconteii | Ammodramus_leconteii | diurnal |
| birds | Ammospiza_maritima | Ammodramus_maritimus | diurnal |
| birds | Ammospiza_nelsoni | Ammodramus_nelsoni | diurnal |
| birds | Ampeliceps_coronatus | Ampeliceps_coronatus | diurnal |
| birds | Ampelioides_tschudii | Ampelioides_tschudii | diurnal |
| birds | Ampelion_rubrocristatus | Ampelion_rubrocristatus | diurnal |
| birds | Ampelion_rufaxilla | Ampelion_rufaxilla | diurnal |
| birds | Ampelornis_griseiceps | Myrmeciza_griseiceps | diurnal |
| birds | Amphispiza_bilineata | Amphispiza_bilineata | diurnal |
| birds | Amphispiza_quinquestriata | Aimophila_quinquestriata | diurnal |
| birds | Amytornis_barbatus | Amytornis_barbatus | diurnal |
| birds | Amytornis_dorotheae | Amytornis_dorotheae | diurnal |
| birds | Amytornis_goyderi | Amytornis_goyderi | diurnal |
| birds | Amytornis_housei | Amytornis_housei | diurnal |
| birds | Amytornis_purnelli | Amytornis_purnelli | diurnal |
| birds | Amytornis_striatus | Amytornis_striatus | diurnal |
| birds | Amytornis_textilis | Amytornis_textilis | diurnal |
| birds | Amytornis_woodwardi | Amytornis_woodwardi | diurnal |
| birds | Anabacerthia_amaurotis | Anabacerthia_amaurotis | diurnal |
| birds | Anabacerthia_lichtensteini | Philydor_lichtensteini | diurnal |
| birds | Anabacerthia_ruficaudata | Philydor_ruficaudatum | diurnal |
| birds | Anabacerthia_striaticollis | Anabacerthia_striaticollis | diurnal |
| birds | Anabacerthia_variegaticeps | Anabacerthia_variegaticeps | diurnal |
| birds | Anabathmis_hartlaubii | Nectarinia_hartlaubii | diurnal |
| birds | Anabathmis_newtonii | Nectarinia_newtonii | diurnal |
| birds | Anabathmis_reichenbachii | Nectarinia_reichenbachii | diurnal |
| birds | Anabazenops_dorsalis | Anabazenops_dorsalis | diurnal |
| birds | Anabazenops_fuscus | Anabazenops_fuscus | diurnal |
| birds | Anairetes_alpinus | Anairetes_alpinus | diurnal |
| birds | Anairetes_fernandezianus | Anairetes_fernandezianus | diurnal |
| birds | Anairetes_flavirostris | Anairetes_flavirostris | diurnal |
| birds | Anairetes_nigrocristatus | Anairetes_nigrocristatus | diurnal |

| | | | |
|---|---|---|---|
| birds | Anairetes_parulus | Anairetes_parulus | diurnal |
| birds | Anairetes_reguloides | Anairetes_reguloides | diurnal |
| birds | Anaplectes_rubriceps | Anaplectes_rubriceps | diurnal |
| birds | Anarhynchus_frontalis | Anarhynchus_frontalis | cathemeral |
| birds | Anas_acuta | Anas_acuta | diurnal |
| birds | Anas_aucklandica | Anas_aucklandica | nocturnal |
| birds | Anas_bahamensis | Anas_bahamensis | diurnal |
| birds | Anas_bernieri | Anas_bernieri | diurnal |
| birds | Anas_capensis | Anas_capensis | diurnal |
| birds | Anas_castanea | Anas_castanea | diurnal |
| birds | Anas_crecca | Anas_crecca | diurnal |
| birds | Anas_erythrorhyncha | Anas_erythrorhyncha | diurnal |
| birds | Anas_flavirostris | Anas_flavirostris | diurnal |
| birds | Anas_georgica | Anas_georgica | diurnal |
| birds | Anas_gibberifrons | Anas_gibberifrons | diurnal |
| birds | Anas_luzonica | Anas_luzonica | diurnal |
| birds | Anas_melleri | Anas_melleri | diurnal |
| birds | Anas_platyrhynchos | Anas_platyrhynchos | diurnal |
| birds | Anas_poecilorhyncha | Anas_poecilorhyncha | diurnal |
| birds | Anas_rubripes | Anas_rubripes | diurnal |
| birds | Anas_sparsa | Anas_sparsa | diurnal |
| birds | Anas_superciliosa | Anas_superciliosa | diurnal |
| birds | Anas_undulata | Anas_undulata | diurnal |
| birds | Anastomus_lamelligerus | Anastomus_lamelligerus | diurnal |
| birds | Anastomus_oscitans | Anastomus_oscitans | diurnal |
| birds | Ancistrops_strigilatus | Ancistrops_strigilatus | diurnal |
| birds | Andigena_cucullata | Andigena_cucullata | diurnal |
| birds | Andigena_hypoglauca | Andigena_hypoglauca | diurnal |
| birds | Andigena_laminirostris | Andigena_laminirostris | diurnal |
| birds | Andigena_nigrirostris | Andigena_nigrirostris | diurnal |
| birds | Androdon_aequatorialis | Androdon_aequatorialis | diurnal |
| birds | Andropadus_importunus | Andropadus_importunus | diurnal |
| birds | Androphobus_viridis | Androphobus_viridis | diurnal |
| birds | Anhima_cornuta | Anhima_cornuta | diurnal |

| | | | |
|---|---|---|---|
| birds | Anhinga_anhinga | Anhinga_anhinga | diurnal |
| birds | Anhinga_melanogaster | Anhinga_melanogaster | diurnal |
| birds | Anisognathus_igniventris | Anisognathus_igniventris | diurnal |
| birds | Anisognathus_lacrymosus | Anisognathus_lacrymosus | diurnal |
| birds | Anisognathus_melanogenys | Anisognathus_melanogenys | diurnal |
| birds | Anisognathus_notabilis | Anisognathus_notabilis | diurnal |
| birds | Anisognathus_somptuosus | Anisognathus_somptuosus | diurnal |
| birds | Anodorhynchus_hyacinthinus | Anodorhynchus_hyacinthinus | diurnal |
| birds | Anodorhynchus_leari | Anodorhynchus_leari | diurnal |
| birds | Anomalospiza_imberbis | Anomalospiza_imberbis | diurnal |
| birds | Anopetia_gounellei | Anopetia_gounellei | diurnal |
| birds | Anorrhinus_austeni | Anorrhinus_austeni | diurnal |
| birds | Anorrhinus_galeritus | Anorrhinus_galeritus | diurnal |
| birds | Anorrhinus_tickelli | Anorrhinus_tickelli | diurnal |
| birds | Anous_albivittus | Procelsterna_albivitta | diurnal |
| birds | Anous_ceruleus | Procelsterna_cerulea | diurnal |
| birds | Anous_minutus | Anous_minutus | diurnal |
| birds | Anous_stolidus | Anous_stolidus | diurnal |
| birds | Anous_tenuirostris | Anous_tenuirostris | diurnal |
| birds | Anser_albifrons | Anser_albifrons | diurnal |
| birds | Anser_anser | Anser_anser | diurnal |
| birds | Anser_brachyrhynchus | Anser_brachyrhynchus | diurnal |
| birds | Anser_caerulescens | Chen_caerulescens | diurnal |
| birds | Anser_canagicus | Chen_canagica | diurnal |
| birds | Anser_cygnoid | Anser_cygnoides | diurnal |
| birds | Anser_fabalis | Anser_fabalis | diurnal |
| birds | Anser_indicus | Anser_indicus | diurnal |
| birds | Anser_rossii | Chen_rossii | diurnal |
| birds | Anseranas_semipalmata | Anseranas_semipalmata | diurnal |
| birds | Anthipes_monileger | Ficedula_monileger | diurnal |
| birds | Anthipes_solitaris | Ficedula_solitaris | diurnal |
| birds | Anthobaphes_violacea | Nectarinia_violacea | diurnal |
| birds | Anthocephala_floriceps | Anthocephala_floriceps | diurnal |
| birds | Anthochaera_carunculata | Anthochaera_carunculata | diurnal |

| | | | |
|---|---|---|---|
| birds | Anthochaera_chrysoptera | Anthochaera_chrysoptera | diurnal |
| birds | Anthochaera_paradoxa | Anthochaera_paradoxa | diurnal |
| birds | Anthochaera_phrygia | Xanthomyza_phrygia | diurnal |
| birds | Anthornis_melanura | Anthornis_melanura | diurnal |
| birds | Anthoscopus_caroli | Anthoscopus_caroli | diurnal |
| birds | Anthoscopus_flavifrons | Anthoscopus_flavifrons | diurnal |
| birds | Anthoscopus_minutus | Anthoscopus_minutus | diurnal |
| birds | Anthoscopus_musculus | Anthoscopus_musculus | diurnal |
| birds | Anthoscopus_parvulus | Anthoscopus_parvulus | diurnal |
| birds | Anthoscopus_punctifrons | Anthoscopus_punctifrons | diurnal |
| birds | Anthracoceros_albirostris | Anthracoceros_albirostris | diurnal |
| birds | Anthracoceros_coronatus | Anthracoceros_coronatus | diurnal |
| birds | Anthracoceros_malayanus | Anthracoceros_malayanus | diurnal |
| birds | Anthracoceros_marchei | Anthracoceros_marchei | diurnal |
| birds | Anthracoceros_montani | Anthracoceros_montani | diurnal |
| birds | Anthracothorax_dominicus | Anthracothorax_dominicus | diurnal |
| birds | Anthracothorax_mango | Anthracothorax_mango | diurnal |
| birds | Anthracothorax_nigricollis | Anthracothorax_nigricollis | diurnal |
| birds | Anthracothorax_veraguensis | Anthracothorax_veraguensis | diurnal |
| birds | Anthracothorax_viridigula | Anthracothorax_viridigula | diurnal |
| birds | Anthracothorax_viridis | Anthracothorax_viridis | diurnal |
| birds | Anthreptes_anchietae | Anthreptes_anchietae | diurnal |
| birds | Anthreptes_aurantius | Anthreptes_aurantium | diurnal |
| birds | Anthreptes_longuemarei | Anthreptes_longuemarei | diurnal |
| birds | Anthreptes_malacensis | Anthreptes_malacensis | diurnal |
| birds | Anthreptes_neglectus | Anthreptes_neglectus | diurnal |
| birds | Anthreptes_orientalis | Anthreptes_orientalis | diurnal |
| birds | Anthreptes_rectirostris | Anthreptes_rectirostris | diurnal |
| birds | Anthreptes_reichenowi | Anthreptes_reichenowi | diurnal |
| birds | Anthreptes_rhodolaemus | Anthreptes_rhodolaemus | diurnal |
| birds | Anthreptes_rubritorques | Anthreptes_rubritorques | diurnal |
| birds | Anthreptes_seimundi | Nectarinia_seimundi | diurnal |
| birds | Anthreptes_simplex | Anthreptes_simplex | diurnal |
| birds | Anthropoides_paradiseus | Grus_paradisea | diurnal |

| | | | |
|---|---|---|---|
| birds | Anthropoides_virgo | Grus_virgo | diurnal |
| birds | Anthus_antarcticus | Anthus_antarcticus | diurnal |
| birds | Anthus_berthelotii | Anthus_berthelotii | diurnal |
| birds | Anthus_bogotensis | Anthus_bogotensis | diurnal |
| birds | Anthus_brachyurus | Anthus_brachyurus | diurnal |
| birds | Anthus_caffer | Anthus_caffer | diurnal |
| birds | Anthus_campestris | Anthus_campestris | diurnal |
| birds | Anthus_cervinus | Anthus_cervinus | diurnal |
| birds | Anthus_correndera | Anthus_correndera | diurnal |
| birds | Anthus_crenatus | Anthus_crenatus | diurnal |
| birds | Anthus_furcatus | Anthus_furcatus | diurnal |
| birds | Anthus_godlewskii | Anthus_godlewskii | diurnal |
| birds | Anthus_gustavi | Anthus_gustavi | diurnal |
| birds | Anthus_gutturalis | Anthus_gutturalis | diurnal |
| birds | Anthus_hellmayri | Anthus_hellmayri | diurnal |
| birds | Anthus_hodgsoni | Anthus_hodgsoni | diurnal |
| birds | Anthus_hoeschi | Anthus_hoeschi | diurnal |
| birds | Anthus_leucophrys | Anthus_leucophrys | diurnal |
| birds | Anthus_lineiventris | Anthus_lineiventris | diurnal |
| birds | Anthus_lutescens | Anthus_lutescens | diurnal |
| birds | Anthus_melindae | Anthus_melindae | diurnal |
| birds | Anthus_nattereri | Anthus_nattereri | diurnal |
| birds | Anthus_novaeseelandiae | Anthus_novaeseelandiae | diurnal |
| birds | Anthus_pallidiventris | Anthus_pallidiventris | diurnal |
| birds | Anthus_petrosus | Anthus_petrosus | diurnal |
| birds | Anthus_pratensis | Anthus_pratensis | diurnal |
| birds | Anthus_richardi | Anthus_richardi | diurnal |
| birds | Anthus_roseatus | Anthus_roseatus | diurnal |
| birds | Anthus_rubescens | Anthus_rubescens | diurnal |
| birds | Anthus_rufulus | Anthus_rufulus | diurnal |
| birds | Anthus_similis | Anthus_similis | diurnal |
| birds | Anthus_sokokensis | Anthus_sokokensis | diurnal |
| birds | Anthus_spinoletta | Anthus_spinoletta | diurnal |
| birds | Anthus_sylvanus | Anthus_sylvanus | diurnal |

| | | | |
|---|---|---|---|
| birds | Anthus_trivialis | Anthus_trivialis | diurnal |
| birds | Anthus_vaalensis | Anthus_vaalensis | diurnal |
| birds | Antilophia_galeata | Antilophia_galeata | diurnal |
| birds | Antrostomus_badius | Caprimulgus_badius | nocturnal |
| birds | Antrostomus_carolinensis | Caprimulgus_carolinensis | nocturnal |
| birds | Antrostomus_cubanensis | Caprimulgus_cubanensis | nocturnal |
| birds | Antrostomus_ekmani | Caprimulgus_ekmani | nocturnal |
| birds | Antrostomus_noctitherus | Caprimulgus_noctitherus | nocturnal |
| birds | Antrostomus_ridgwayi | Caprimulgus_ridgwayi | nocturnal |
| birds | Antrostomus_rufus | Caprimulgus_rufus | nocturnal |
| birds | Antrostomus_salvini | Caprimulgus_salvini | nocturnal |
| birds | Antrostomus_saturatus | Caprimulgus_saturatus | nocturnal |
| birds | Antrostomus_sericocaudatus | Caprimulgus_sericocaudatus | nocturnal |
| birds | Antrostomus_vociferus | Caprimulgus_vociferus | nocturnal |
| birds | Anumbius_annumbi | Anumbius_annumbi | diurnal |
| birds | Anurophasis_monorthonyx | Anurophasis_monorthonyx | diurnal |
| birds | Apalharpactes_mackloti | Apalharpactes_mackloti | diurnal |
| birds | Apalis_alticola | Apalis_alticola | diurnal |
| birds | Apalis_bamendae | Apalis_bamendae | diurnal |
| birds | Apalis_binotata | Apalis_binotata | diurnal |
| birds | Apalis_chapini | Apalis_chapini | diurnal |
| birds | Apalis_chariessa | Apalis_chariessa | diurnal |
| birds | Apalis_chirindensis | Apalis_chirindensis | diurnal |
| birds | Apalis_cinerea | Apalis_cinerea | diurnal |
| birds | Apalis_flavida | Apalis_flavida | diurnal |
| birds | Apalis_jacksoni | Apalis_jacksoni | diurnal |
| birds | Apalis_karamojae | Apalis_karamojae | diurnal |
| birds | Apalis_melanocephala | Apalis_melanocephala | diurnal |
| birds | Apalis_nigriceps | Apalis_nigriceps | diurnal |
| birds | Apalis_personata | Apalis_personata | diurnal |
| birds | Apalis_porphyrolaema | Apalis_porphyrolaema | diurnal |
| birds | Apalis_ruddi | Apalis_ruddi | diurnal |
| birds | Apalis_rufogularis | Apalis_rufogularis | diurnal |
| birds | Apalis_sharpii | Apalis_sharpii | diurnal |

| | | | |
|---|---|---|---|
| birds | Apalis_thoracica | Apalis_thoracica | diurnal |
| birds | Apaloderma_narina | Apaloderma_narina | diurnal |
| birds | Apaloderma_vittatum | Apaloderma_vittatum | diurnal |
| birds | Apalopteron_familiare | Apalopteron_familiare | diurnal |
| birds | Aphanotriccus_audax | Aphanotriccus_audax | diurnal |
| birds | Aphanotriccus_capitalis | Aphanotriccus_capitalis | diurnal |
| birds | Aphelocephala_leucopsis | Aphelocephala_leucopsis | diurnal |
| birds | Aphelocephala_nigricincta | Aphelocephala_nigricincta | diurnal |
| birds | Aphelocephala_pectoralis | Aphelocephala_pectoralis | diurnal |
| birds | Aphelocoma_californica | Aphelocoma_californica | diurnal |
| birds | Aphelocoma_coerulescens | Aphelocoma_coerulescens | diurnal |
| birds | Aphelocoma_ultramarina | Aphelocoma_ultramarina | diurnal |
| birds | Aphelocoma_unicolor | Aphelocoma_unicolor | diurnal |
| birds | Aphrastura_masafuerae | Aphrastura_masafuerae | diurnal |
| birds | Aphrastura_spinicauda | Aphrastura_spinicauda | diurnal |
| birds | Aphrodroma_brevirostris | Lugensa_brevirostris | cathemeral |
| birds | Aplonis_atrifusca | Aplonis_atrifusca | diurnal |
| birds | Aplonis_brunneicapillus | Aplonis_brunneicapillus | diurnal |
| birds | Aplonis_cantoroides | Aplonis_cantoroides | diurnal |
| birds | Aplonis_cinerascens | Aplonis_cinerascens | diurnal |
| birds | Aplonis_crassa | Aplonis_crassa | diurnal |
| birds | Aplonis_dichroa | Aplonis_dichroa | diurnal |
| birds | Aplonis_feadensis | Aplonis_feadensis | diurnal |
| birds | Aplonis_grandis | Aplonis_grandis | diurnal |
| birds | Aplonis_insularis | Aplonis_insularis | diurnal |
| birds | Aplonis_magna | Aplonis_magna | diurnal |
| birds | Aplonis_metallica | Aplonis_metallica | diurnal |
| birds | Aplonis_minor | Aplonis_minor | diurnal |
| birds | Aplonis_mysolensis | Aplonis_mysolensis | diurnal |
| birds | Aplonis_mystacea | Aplonis_mystacea | diurnal |
| birds | Aplonis_opaca | Aplonis_opaca | diurnal |
| birds | Aplonis_panayensis | Aplonis_panayensis | diurnal |
| birds | Aplonis_pelzelni | Aplonis_pelzelni | diurnal |
| birds | Aplonis_santovestris | Aplonis_santovestris | diurnal |

| | | | |
|---|---|---|---|
| birds | Aplonis_striata | Aplonis_striata | diurnal |
| birds | Aplonis_tabuensis | Aplonis_tabuensis | diurnal |
| birds | Aplonis_zelandica | Aplonis_zelandica | diurnal |
| birds | Aplopelia_larvata | Aplopelia_larvata | diurnal |
| birds | Aprositornis_disjuncta | Myrmeciza_disjuncta | diurnal |
| birds | Aprosmictus_erythropterus | Aprosmictus_erythropterus | diurnal |
| birds | Aprosmictus_jonquillaceus | Aprosmictus_jonquillaceus | diurnal |
| birds | Aptenodytes_forsteri | Aptenodytes_forsteri | cathemeral |
| birds | Aptenodytes_patagonicus | Aptenodytes_patagonicus | cathemeral |
| birds | Apteryx_australis | Apteryx_australis | nocturnal |
| birds | Apteryx_haastii | Apteryx_haastii | nocturnal |
| birds | Apteryx_owenii | Apteryx_owenii | nocturnal |
| birds | Apus_affinis | Apus_affinis | diurnal |
| birds | Apus_apus | Apus_apus | diurnal |
| birds | Apus_barbatus | Apus_barbatus | diurnal |
| birds | Apus_berliozi | Apus_berliozi | diurnal |
| birds | Apus_bradfieldi | Apus_bradfieldi | diurnal |
| birds | Apus_caffer | Apus_caffer | diurnal |
| birds | Apus_horus | Apus_horus | diurnal |
| birds | Apus_niansae | Apus_niansae | diurnal |
| birds | Apus_nipalensis | Apus_nipalensis | diurnal |
| birds | Apus_pacificus | Apus_pacificus | diurnal |
| birds | Apus_pallidus | Apus_pallidus | diurnal |
| birds | Apus_unicolor | Apus_unicolor | diurnal |
| birds | Aquila_adalberti | Aquila_adalberti | diurnal |
| birds | Aquila_africana | Aquila_africanus | diurnal |
| birds | Aquila_audax | Aquila_audax | diurnal |
| birds | Aquila_chrysaetos | Aquila_chrysaetos | diurnal |
| birds | Aquila_fasciata | Aquila_fasciatus | diurnal |
| birds | Aquila_gurneyi | Aquila_gurneyi | diurnal |
| birds | Aquila_heliaca | Aquila_heliaca | diurnal |
| birds | Aquila_rapax | Aquila_rapax | diurnal |
| birds | Aquila_spilogaster | Hieraaetus_spilogaster | diurnal |
| birds | Aquila_verreauxii | Aquila_verreauxii | diurnal |

| | | | |
|---|---|---|---|
| birds | Ara_ambiguus | Ara_ambiguus | diurnal |
| birds | Ara_ararauna | Ara_ararauna | diurnal |
| birds | Ara_chloropterus | Ara_chloropterus | diurnal |
| birds | Ara_glaucogularis | Ara_glaucogularis | diurnal |
| birds | Ara_macao | Ara_macao | diurnal |
| birds | Ara_rubrogenys | Ara_rubrogenys | diurnal |
| birds | Ara_severus | Ara_severus | diurnal |
| birds | Arachnothera_affinis | Arachnothera_affinis | diurnal |
| birds | Arachnothera_chrysogenys | Arachnothera_chrysogenys | diurnal |
| birds | Arachnothera_clarae | Arachnothera_clarae | diurnal |
| birds | Arachnothera_crassirostris | Arachnothera_crassirostris | diurnal |
| birds | Arachnothera_flavigaster | Arachnothera_flavigaster | diurnal |
| birds | Arachnothera_longirostra | Arachnothera_longirostra | diurnal |
| birds | Arachnothera_magna | Arachnothera_magna | diurnal |
| birds | Arachnothera_robusta | Arachnothera_robusta | diurnal |
| birds | Aramides_cajaneus | Aramides_cajanea | nocturnal |
| birds | Aramus_guarauna | Aramus_guarauna | cathemeral |
| birds | Aratinga_auricapillus | Aratinga_auricapillus | diurnal |
| birds | Aratinga_jandaya | Aratinga_jandaya | diurnal |
| birds | Aratinga_nenday | Nandayus_nenday | diurnal |
| birds | Aratinga_solstitialis | Aratinga_solstitialis | diurnal |
| birds | Aratinga_weddellii | Aratinga_weddellii | diurnal |
| birds | Arborophila_ardens | Arborophila_ardens | diurnal |
| birds | Arborophila_atrogularis | Arborophila_atrogularis | diurnal |
| birds | Arborophila_brunneopectus | Arborophila_brunneopectus | diurnal |
| birds | Arborophila_cambodiana | Arborophila_cambodiana | diurnal |
| birds | Arborophila_crudigularis | Arborophila_crudigularis | diurnal |
| birds | Arborophila_davidi | Arborophila_davidi | diurnal |
| birds | Arborophila_gingica | Arborophila_gingica | diurnal |
| birds | Arborophila_hyperythra | Arborophila_hyperythra | diurnal |
| birds | Arborophila_javanica | Arborophila_javanica | diurnal |
| birds | Arborophila_mandellii | Arborophila_mandellii | diurnal |
| birds | Arborophila_orientalis | Arborophila_orientalis | diurnal |
| birds | Arborophila_rubrirostris | Arborophila_rubrirostris | diurnal |

| | | | |
|---|---|---|---|
| birds | Arborophila_rufipectus | Arborophila_rufipectus | diurnal |
| birds | Arborophila_rufogularis | Arborophila_rufogularis | diurnal |
| birds | Arborophila_torqueola | Arborophila_torqueola | diurnal |
| birds | Arcanator_orostruthus | Modulatrix_orostruthus | diurnal |
| birds | Archboldia_papuensis | Archboldia_papuensis | diurnal |
| birds | Archilochus_alexandri | Archilochus_alexandri | diurnal |
| birds | Archilochus_colubris | Archilochus_colubris | diurnal |
| birds | Ardea_alba | Casmerodius_albus | diurnal |
| birds | Ardea_cinerea | Ardea_cinerea | cathemeral |
| birds | Ardea_cocoi | Ardea_cocoi | cathemeral |
| birds | Ardea_goliath | Ardea_goliath | diurnal |
| birds | Ardea_intermedia | Mesophoyx_intermedia | diurnal |
| birds | Ardea_melanocephala | Ardea_melanocephala | cathemeral |
| birds | Ardea_pacifica | Ardea_pacifica | diurnal |
| birds | Ardea_purpurea | Ardea_purpurea | diurnal |
| birds | Ardea_sumatrana | Ardea_sumatrana | diurnal |
| birds | Ardenna_bulleri | Puffinus_bulleri | cathemeral |
| birds | Ardenna_carneipes | Puffinus_carneipes | cathemeral |
| birds | Ardenna_creatopus | Puffinus_creatopus | cathemeral |
| birds | Ardenna_gravis | Puffinus_gravis | cathemeral |
| birds | Ardenna_grisea | Puffinus_griseus | cathemeral |
| birds | Ardenna_pacifica | Puffinus_pacificus | cathemeral |
| birds | Ardenna_tenuirostris | Puffinus_tenuirostris | cathemeral |
| birds | Ardeola_bacchus | Ardeola_bacchus | diurnal |
| birds | Ardeola_grayii | Ardeola_grayii | cathemeral |
| birds | Ardeola_ralloides | Ardeola_ralloides | diurnal |
| birds | Ardeotis_arabs | Ardeotis_arabs | diurnal |
| birds | Ardeotis_australis | Ardeotis_australis | diurnal |
| birds | Ardeotis_kori | Ardeotis_kori | diurnal |
| birds | Ardeotis_nigriceps | Ardeotis_nigriceps | diurnal |
| birds | Arenaria_interpres | Arenaria_interpres | diurnal |
| birds | Arenaria_melanocephala | Arenaria_melanocephala | diurnal |
| birds | Argusianus_argus | Argusianus_argus | diurnal |
| birds | Argya_affinis | Turdoides_affinis | diurnal |

| | | | |
|---|---|---|---|
| birds | Argya_altirostris | Turdoides_altirostris | diurnal |
| birds | Argya_aylmeri | Turdoides_aylmeri | diurnal |
| birds | Argya_caudata | Turdoides_caudata | diurnal |
| birds | Argya_cinereifrons | Garrulax_cinereifrons | diurnal |
| birds | Argya_earlei | Turdoides_earlei | diurnal |
| birds | Argya_fulva | Turdoides_fulva | diurnal |
| birds | Argya_gularis | Turdoides_gularis | diurnal |
| birds | Argya_longirostris | Turdoides_longirostris | diurnal |
| birds | Argya_malcolmi | Turdoides_malcolmi | diurnal |
| birds | Argya_rubiginosa | Turdoides_rubiginosa | diurnal |
| birds | Argya_rufescens | Turdoides_rufescens | diurnal |
| birds | Argya_squamiceps | Turdoides_squamiceps | diurnal |
| birds | Argya_subrufa | Turdoides_subrufa | diurnal |
| birds | Arizelocichla_masukuensis | Andropadus_masukuensis | diurnal |
| birds | Arizelocichla_milanjensis | Andropadus_milanjensis | diurnal |
| birds | Arizelocichla_montana | Andropadus_montanus | diurnal |
| birds | Arizelocichla_tephrolaema | Andropadus_tephrolaemus | diurnal |
| birds | Arremon_abeillei | Arremon_abeillei | diurnal |
| birds | Arremon_aurantiirostris | Arremon_aurantiirostris | diurnal |
| birds | Arremon_brunneinucha | Arremon_brunneinucha | diurnal |
| birds | Arremon_castaneiceps | Arremon_castaneiceps | diurnal |
| birds | Arremon_crassirostris | Arremon_crassirostris | diurnal |
| birds | Arremon_flavirostris | Arremon_flavirostris | diurnal |
| birds | Arremon_franciscanus | Arremon_franciscanus | diurnal |
| birds | Arremon_schlegeli | Arremon_schlegeli | diurnal |
| birds | Arremon_semitorquatus | Arremon_semitorquatus | diurnal |
| birds | Arremon_taciturnus | Arremon_taciturnus | diurnal |
| birds | Arremon_torquatus | Arremon_torquatus | diurnal |
| birds | Arremon_virenticeps | Arremon_virenticeps | diurnal |
| birds | Arremonops_chloronotus | Arremonops_chloronotus | diurnal |
| birds | Arremonops_conirostris | Arremonops_conirostris | diurnal |
| birds | Arremonops_rufivirgatus | Arremonops_rufivirgatus | diurnal |
| birds | Arremonops_tocuyensis | Arremonops_tocuyensis | diurnal |
| birds | Arses_insularis | Arses_insularis | diurnal |

| | | | |
|---|---|---|---|
| birds | Arses_kaupi | Arses_kaupi | diurnal |
| birds | Arses_telescopthalmus | Arses_telescophthalmus | diurnal |
| birds | Artamella_viridis | Artamella_viridis | diurnal |
| birds | Artamus_cinereus | Artamus_cinereus | diurnal |
| birds | Artamus_cyanopterus | Artamus_cyanopterus | diurnal |
| birds | Artamus_fuscus | Artamus_fuscus | diurnal |
| birds | Artamus_insignis | Artamus_insignis | diurnal |
| birds | Artamus_leucoryn | Artamus_leucorynchus | diurnal |
| birds | Artamus_maximus | Artamus_maximus | diurnal |
| birds | Artamus_mentalis | Artamus_mentalis | diurnal |
| birds | Artamus_minor | Artamus_minor | diurnal |
| birds | Artamus_personatus | Artamus_personatus | diurnal |
| birds | Artamus_superciliosus | Artamus_superciliosus | diurnal |
| birds | Artemisiospiza_belli | Amphispiza_belli | diurnal |
| birds | Artisornis_metopias | Artisornis_metopias | diurnal |
| birds | Artisornis_moreaui | Artisornis_moreaui | diurnal |
| birds | Artomyias_fuliginosa | Muscicapa_infuscata | diurnal |
| birds | Artomyias_ussheri | Muscicapa_ussheri | diurnal |
| birds | Arundinax_aedon | Acrocephalus_aedon | diurnal |
| birds | Arundinicola_leucocephala | Arundinicola_leucocephala | diurnal |
| birds | Asarcornis_scutulata | Cairina_scutulata | nocturnal |
| birds | Asemospiza_fuliginosa | Tiaris_fuliginosus | diurnal |
| birds | Asemospiza_obscura | Tiaris_obscurus | diurnal |
| birds | Ashbyia_lovensis | Ashbyia_lovensis | diurnal |
| birds | Asio_abyssinicus | Asio_abyssinicus | nocturnal |
| birds | Asio_capensis | Asio_capensis | nocturnal |
| birds | Asio_clamator | Pseudoscops_clamator | nocturnal |
| birds | Asio_flammeus | Asio_flammeus | cathemeral |
| birds | Asio_otus | Asio_otus | nocturnal |
| birds | Asio_stygius | Asio_stygius | nocturnal |
| birds | Aspatha_gularis | Aspatha_gularis | diurnal |
| birds | Asthenes_anthoides | Asthenes_anthoides | diurnal |
| birds | Asthenes_arequipae | Asthenes_arequipae | diurnal |
| birds | Asthenes_baeri | Asthenes_baeri | diurnal |

| | | | |
|---|---|---|---|
| birds | Asthenes_berlepschi | Asthenes_berlepschi | diurnal |
| birds | Asthenes_coryi | Schizoeaca_coryi | diurnal |
| birds | Asthenes_dorbignyi | Asthenes_dorbignyi | diurnal |
| birds | Asthenes_flammulata | Asthenes_flammulata | diurnal |
| birds | Asthenes_fuliginosa | Schizoeaca_fuliginosa | diurnal |
| birds | Asthenes_griseomurina | Schizoeaca_griseomurina | diurnal |
| birds | Asthenes_harterti | Schizoeaca_harterti | diurnal |
| birds | Asthenes_helleri | Schizoeaca_helleri | diurnal |
| birds | Asthenes_heterura | Asthenes_heterura | diurnal |
| birds | Asthenes_huancavelicae | Asthenes_huancavelicae | diurnal |
| birds | Asthenes_hudsoni | Asthenes_hudsoni | diurnal |
| birds | Asthenes_humilis | Asthenes_humilis | diurnal |
| birds | Asthenes_luizae | Asthenes_luizae | diurnal |
| birds | Asthenes_maculicauda | Asthenes_maculicauda | diurnal |
| birds | Asthenes_modesta | Asthenes_modesta | diurnal |
| birds | Asthenes_moreirae | Oreophylax_moreirae | diurnal |
| birds | Asthenes_ottonis | Asthenes_ottonis | diurnal |
| birds | Asthenes_palpebralis | Schizoeaca_palpebralis | diurnal |
| birds | Asthenes_perijana | Schizoeaca_perijana | diurnal |
| birds | Asthenes_pudibunda | Asthenes_pudibunda | diurnal |
| birds | Asthenes_pyrrholeuca | Asthenes_pyrrholeuca | diurnal |
| birds | Asthenes_urubambensis | Asthenes_urubambensis | diurnal |
| birds | Asthenes_vilcabambae | Schizoeaca_vilcabambae | diurnal |
| birds | Asthenes_virgata | Asthenes_virgata | diurnal |
| birds | Asthenes_wyatti | Asthenes_wyatti | diurnal |
| birds | Astrapia_mayeri | Astrapia_mayeri | diurnal |
| birds | Astrapia_nigra | Astrapia_nigra | diurnal |
| birds | Astrapia_rothschildi | Astrapia_rothschildi | diurnal |
| birds | Astrapia_splendidissima | Astrapia_splendidissima | diurnal |
| birds | Astrapia_stephaniae | Astrapia_stephaniae | diurnal |
| birds | Atalotriccus_pilaris | Atalotriccus_pilaris | diurnal |
| birds | Atelornis_crossleyi | Atelornis_crossleyi | cathemeral |
| birds | Atelornis_pittoides | Atelornis_pittoides | cathemeral |
| birds | Athene_blewitti | Heteroglaux_blewitti | diurnal |

| | | | |
|---|---|---|---|
| birds | Athene_brama | Athene_brama | nocturnal |
| birds | Athene_noctua | Athene_noctua | nocturnal |
| birds | Athene_superciliaris | Ninox_superciliaris | nocturnal |
| birds | Atimastillas_flavicollis | Chlorocichla_flavicollis | diurnal |
| birds | Atlapetes_albiceps | Atlapetes_albiceps | diurnal |
| birds | Atlapetes_albinucha | Atlapetes_albinucha | diurnal |
| birds | Atlapetes_albofrenatus | Atlapetes_albofrenatus | diurnal |
| birds | Atlapetes_canigenis | Atlapetes_canigenis | diurnal |
| birds | Atlapetes_citrinellus | Atlapetes_citrinellus | diurnal |
| birds | Atlapetes_flaviceps | Atlapetes_flaviceps | diurnal |
| birds | Atlapetes_forbesi | Atlapetes_forbesi | diurnal |
| birds | Atlapetes_fulviceps | Atlapetes_fulviceps | diurnal |
| birds | Atlapetes_fuscoolivaceus | Atlapetes_fuscoolivaceus | diurnal |
| birds | Atlapetes_latinuchus | Atlapetes_latinuchus | diurnal |
| birds | Atlapetes_leucopis | Atlapetes_leucopis | diurnal |
| birds | Atlapetes_leucopterus | Atlapetes_leucopterus | diurnal |
| birds | Atlapetes_melanocephalus | Atlapetes_melanocephalus | diurnal |
| birds | Atlapetes_melanolaemus | Atlapetes_melanolaemus | diurnal |
| birds | Atlapetes_melanopsis | Atlapetes_melanopsis | diurnal |
| birds | Atlapetes_nationi | Atlapetes_nationi | diurnal |
| birds | Atlapetes_pallidiceps | Atlapetes_pallidiceps | diurnal |
| birds | Atlapetes_pallidinucha | Atlapetes_pallidinucha | diurnal |
| birds | Atlapetes_personatus | Atlapetes_personatus | diurnal |
| birds | Atlapetes_pileatus | Atlapetes_pileatus | diurnal |
| birds | Atlapetes_rufigenis | Atlapetes_rufigenis | diurnal |
| birds | Atlapetes_rufinucha | Atlapetes_rufinucha | diurnal |
| birds | Atlapetes_schistaceus | Atlapetes_schistaceus | diurnal |
| birds | Atlapetes_seebohmi | Atlapetes_seebohmi | diurnal |
| birds | Atlapetes_semirufus | Atlapetes_semirufus | diurnal |
| birds | Atlapetes_tricolor | Atlapetes_tricolor | diurnal |
| birds | Atrichornis_clamosus | Atrichornis_clamosus | diurnal |
| birds | Atrichornis_rufescens | Atrichornis_rufescens | diurnal |
| birds | Attagis_gayi | Attagis_gayi | diurnal |
| birds | Attagis_malouinus | Attagis_malouinus | diurnal |

| | | | |
|---|---|---|---|
| birds | Atticora_fasciata | Atticora_fasciata | diurnal |
| birds | Atticora_pileata | Notiochelidon_pileata | diurnal |
| birds | Atticora_tibialis | Neochelidon_tibialis | diurnal |
| birds | Attila_bolivianus | Attila_bolivianus | diurnal |
| birds | Attila_cinnamomeus | Attila_cinnamomeus | diurnal |
| birds | Attila_citriniventris | Attila_citriniventris | diurnal |
| birds | Attila_phoenicurus | Attila_phoenicurus | diurnal |
| birds | Attila_rufus | Attila_rufus | diurnal |
| birds | Attila_spadiceus | Attila_spadiceus | diurnal |
| birds | Attila_torridus | Attila_torridus | diurnal |
| birds | Augastes_lumachella | Augastes_lumachella | diurnal |
| birds | Augastes_scutatus | Augastes_scutatus | diurnal |
| birds | Aulacorhynchus_coeruleicinctis | Aulacorhynchus_coeruleicinctis | diurnal |
| birds | Aulacorhynchus_derbianus | Aulacorhynchus_derbianus | diurnal |
| birds | Aulacorhynchus_haematopygus | Aulacorhynchus_haematopygus | diurnal |
| birds | Aulacorhynchus_huallagae | Aulacorhynchus_huallagae | diurnal |
| birds | Aulacorhynchus_prasinus | Aulacorhynchus_prasinus | diurnal |
| birds | Aulacorhynchus_sulcatus | Aulacorhynchus_sulcatus | diurnal |
| birds | Auriparus_flaviceps | Auriparus_flaviceps | diurnal |
| birds | Automolus_infuscatus | Automolus_infuscatus | diurnal |
| birds | Automolus_leucophthalmus | Automolus_leucophthalmus | diurnal |
| birds | Automolus_melanopezus | Automolus_melanopezus | diurnal |
| birds | Automolus_ochrolaemus | Automolus_ochrolaemus | diurnal |
| birds | Automolus_rufipileatus | Automolus_rufipileatus | diurnal |
| birds | Automolus_subulatus | Hyloctistes_subulatus | diurnal |
| birds | Aviceda_cuculoides | Aviceda_cuculoides | diurnal |
| birds | Aviceda_jerdoni | Aviceda_jerdoni | diurnal |
| birds | Aviceda_leuphotes | Aviceda_leuphotes | diurnal |
| birds | Aviceda_subcristata | Aviceda_subcristata | diurnal |
| birds | Avocettula_recurvirostris | Avocettula_recurvirostris | diurnal |
| birds | Aythya_affinis | Aythya_affinis | diurnal |
| birds | Aythya_americana | Aythya_americana | diurnal |
| birds | Aythya_australis | Aythya_australis | diurnal |
| birds | Aythya_baeri | Aythya_baeri | diurnal |

| | | | |
|---|---|---|---|
| birds | Aythya_collaris | Aythya_collaris | diurnal |
| birds | Aythya_ferina | Aythya_ferina | diurnal |
| birds | Aythya_innotata | Aythya_innotata | diurnal |
| birds | Aythya_marila | Aythya_marila | diurnal |
| birds | Aythya_novaeseelandiae | Aythya_novaeseelandiae | diurnal |
| birds | Aythya_nyroca | Aythya_nyroca | diurnal |
| birds | Aythya_valisineria | Aythya_valisineria | diurnal |
| birds | Baeolophus_atricristatus | Baeolophus_atricristatus | diurnal |
| birds | Baeolophus_bicolor | Baeolophus_bicolor | diurnal |
| birds | Baeolophus_inornatus | Baeolophus_inornatus | diurnal |
| birds | Baeolophus_ridgwayi | Baeolophus_ridgwayi | diurnal |
| birds | Baeolophus_wollweberi | Baeolophus_wollweberi | diurnal |
| birds | Baeopogon_indicator | Baeopogon_indicator | diurnal |
| birds | Balaeniceps_rex | Balaeniceps_rex | diurnal |
| birds | Balearica_pavonina | Balearica_pavonina | diurnal |
| birds | Balearica_regulorum | Balearica_regulorum | diurnal |
| birds | Bambusicola_fytchii | Bambusicola_fytchii | diurnal |
| birds | Bambusicola_thoracicus | Bambusicola_thoracicus | diurnal |
| birds | Bangsia_arcaei | Bangsia_arcaei | diurnal |
| birds | Bangsia_aureocincta | Bangsia_aureocincta | diurnal |
| birds | Bangsia_edwardsi | Bangsia_edwardsi | diurnal |
| birds | Bangsia_flavovirens | Chlorospingus_flavovirens | diurnal |
| birds | Bangsia_melanochlamys | Bangsia_melanochlamys | diurnal |
| birds | Bangsia_rothschildi | Bangsia_rothschildi | diurnal |
| birds | Barnardius_zonarius | Barnardius_zonarius | diurnal |
| birds | Bartramia_longicauda | Bartramia_longicauda | diurnal |
| birds | Baryphthengus_martii | Baryphthengus_martii | diurnal |
| birds | Baryphthengus_ruficapillus | Baryphthengus_ruficapillus | diurnal |
| birds | Basileuterus_belli | Basileuterus_belli | diurnal |
| birds | Basileuterus_culicivorus | Basileuterus_culicivorus | diurnal |
| birds | Basileuterus_griseiceps | Basileuterus_griseiceps | diurnal |
| birds | Basileuterus_hypoleucus | Basileuterus_hypoleucus | diurnal |
| birds | Basileuterus_ignotus | Basileuterus_ignotus | diurnal |
| birds | Basileuterus_lachrymosus | Euthlypis_lachrymosa | diurnal |

| | | | |
|---|---|---|---|
| birds | Basileuterus_melanogenys | Basileuterus_melanogenys | diurnal |
| birds | Basileuterus_rufifrons | Basileuterus_rufifrons | diurnal |
| birds | Basileuterus_trifasciatus | Basileuterus_trifasciatus | diurnal |
| birds | Basileuterus_tristriatus | Basileuterus_tristriatus | diurnal |
| birds | Basilinna_leucotis | Hylocharis_leucotis | diurnal |
| birds | Basilinna_xantusii | Hylocharis_xantusii | diurnal |
| birds | Basilornis_corythaix | Basilornis_corythaix | diurnal |
| birds | Batara_cinerea | Batara_cinerea | diurnal |
| birds | Bathmocercus_cerviniventris | Bathmocercus_cerviniventris | diurnal |
| birds | Bathmocercus_rufus | Bathmocercus_rufus | diurnal |
| birds | Batis_capensis | Batis_capensis | diurnal |
| birds | Batis_diops | Batis_diops | diurnal |
| birds | Batis_fratrum | Batis_fratrum | diurnal |
| birds | Batis_ituriensis | Batis_ituriensis | diurnal |
| birds | Batis_margaritae | Batis_margaritae | diurnal |
| birds | Batis_minima | Batis_minima | diurnal |
| birds | Batis_minor | Batis_minor | diurnal |
| birds | Batis_minulla | Batis_minulla | diurnal |
| birds | Batis_mixta | Batis_mixta | diurnal |
| birds | Batis_molitor | Batis_molitor | diurnal |
| birds | Batis_orientalis | Batis_orientalis | diurnal |
| birds | Batis_perkeo | Batis_perkeo | diurnal |
| birds | Batis_poensis | Batis_poensis | diurnal |
| birds | Batis_pririt | Batis_pririt | diurnal |
| birds | Batis_senegalensis | Batis_senegalensis | diurnal |
| birds | Batis_soror | Batis_soror | diurnal |
| birds | Batrachostomus_auritus | Batrachostomus_auritus | nocturnal |
| birds | Batrachostomus_hodgsoni | Batrachostomus_hodgsoni | nocturnal |
| birds | Batrachostomus_javensis | Batrachostomus_javensis | nocturnal |
| birds | Batrachostomus_moniliger | Batrachostomus_moniliger | nocturnal |
| birds | Batrachostomus_septimus | Batrachostomus_septimus | nocturnal |
| birds | Batrachostomus_stellatus | Batrachostomus_stellatus | nocturnal |
| birds | Belocercus_longicaudus | Psittacula_longicauda | diurnal |
| birds | Berenicornis_comatus | Aceros_comatus | diurnal |

| | | | |
|---|---|---|---|
| birds | Berlepschia_rikeri | Berlepschia_rikeri | diurnal |
| birds | Bernieria_madagascariensis | Bernieria_madagascariensis | diurnal |
| birds | Bias_musicus | Bias_musicus | diurnal |
| birds | Biatas_nigropectus | Biatas_nigropectus | diurnal |
| birds | Biziura_lobata | Biziura_lobata | diurnal |
| birds | Bleda_canicapillus | Bleda_canicapillus | diurnal |
| birds | Bleda_eximius | Bleda_eximius | diurnal |
| birds | Bleda_syndactylus | Bleda_syndactylus | diurnal |
| birds | Blythipicus_pyrrhotis | Blythipicus_pyrrhotis | diurnal |
| birds | Blythipicus_rubiginosus | Blythipicus_rubiginosus | diurnal |
| birds | Bocagia_minuta | Tchagra_minutus | diurnal |
| birds | Boissonneaua_flavescens | Boissonneaua_flavescens | diurnal |
| birds | Boissonneaua_jardini | Boissonneaua_jardini | diurnal |
| birds | Boissonneaua_matthewsii | Boissonneaua_matthewsii | diurnal |
| birds | Bolbopsittacus_lunulatus | Bolbopsittacus_lunulatus | diurnal |
| birds | Bolborhynchus_ferrugineifrons | Bolborhynchus_ferrugineifrons | diurnal |
| birds | Bolborhynchus_lineola | Bolborhynchus_lineola | diurnal |
| birds | Bolborhynchus_orbygnesius | Bolborhynchus_orbygnesius | diurnal |
| birds | Bolemoreus_frenatus | Lichenostomus_frenatus | diurnal |
| birds | Bolemoreus_hindwoodi | Lichenostomus_hindwoodi | diurnal |
| birds | Bombycilla_cedrorum | Bombycilla_cedrorum | diurnal |
| birds | Bombycilla_garrulus | Bombycilla_garrulus | diurnal |
| birds | Bombycilla_japonica | Bombycilla_japonica | diurnal |
| birds | Bonasa_umbellus | Bonasa_umbellus | diurnal |
| birds | Bostrychia_carunculata | Bostrychia_carunculata | diurnal |
| birds | Bostrychia_hagedash | Bostrychia_hagedash | diurnal |
| birds | Bostrychia_rara | Bostrychia_rara | diurnal |
| birds | Botaurus_lentiginosus | Botaurus_lentiginosus | nocturnal |
| birds | Botaurus_pinnatus | Botaurus_pinnatus | nocturnal |
| birds | Botaurus_poiciloptilus | Botaurus_poiciloptilus | cathemeral |
| birds | Botaurus_stellaris | Botaurus_stellaris | cathemeral |
| birds | Brachycope_anomala | Brachycope_anomala | diurnal |
| birds | Brachygalba_albogularis | Brachygalba_albogularis | diurnal |
| birds | Brachygalba_goeringi | Brachygalba_goeringi | diurnal |

| | | | |
|---|---|---|---|
| birds | Brachygalba_lugubris | Brachygalba_lugubris | diurnal |
| birds | Brachygalba_salmoni | Brachygalba_salmoni | diurnal |
| birds | Brachypodius_atriceps | Pycnonotus_atriceps | diurnal |
| birds | Brachypteracias_leptosomus | Brachypteracias_leptosomus | diurnal |
| birds | Brachypteryx_leucophris | Brachypteryx_leucophrys | diurnal |
| birds | Brachypteryx_montana | Brachypteryx_montana | diurnal |
| birds | Brachyramphus_brevirostris | Brachyramphus_brevirostris | diurnal |
| birds | Brachyramphus_marmoratus | Brachyramphus_marmoratus | diurnal |
| birds | Bradornis_comitatus | Muscicapa_comitata | diurnal |
| birds | Bradornis_mariquensis | Bradornis_mariquensis | diurnal |
| birds | Bradornis_microrhynchus | Bradornis_microrhynchus | diurnal |
| birds | Bradypterus_baboecala | Bradypterus_baboecala | diurnal |
| birds | Bradypterus_barratti | Bradypterus_barratti | diurnal |
| birds | Bradypterus_brunneus | Dromaeocercus_brunneus | diurnal |
| birds | Bradypterus_carpalis | Bradypterus_carpalis | diurnal |
| birds | Bradypterus_cinnamomeus | Bradypterus_cinnamomeus | diurnal |
| birds | Bradypterus_grandis | Bradypterus_grandis | diurnal |
| birds | Bradypterus_graueri | Bradypterus_graueri | diurnal |
| birds | Bradypterus_lopezi | Bradypterus_lopezi | diurnal |
| birds | Bradypterus_seebohmi | Amphilais_seebohmi | diurnal |
| birds | Bradypterus_sylvaticus | Bradypterus_sylvaticus | diurnal |
| birds | Branta_bernicla | Branta_bernicla | diurnal |
| birds | Branta_canadensis | Branta_canadensis | diurnal |
| birds | Branta_leucopsis | Branta_leucopsis | diurnal |
| birds | Branta_ruficollis | Branta_ruficollis | diurnal |
| birds | Branta_sandvicensis | Branta_sandvicensis | diurnal |
| birds | Brotogeris_chiriri | Brotogeris_chiriri | diurnal |
| birds | Brotogeris_chrysoptera | Brotogeris_chrysoptera | diurnal |
| birds | Brotogeris_cyanoptera | Brotogeris_cyanoptera | diurnal |
| birds | Brotogeris_jugularis | Brotogeris_jugularis | diurnal |
| birds | Brotogeris_pyrrhoptera | Brotogeris_pyrrhoptera | diurnal |
| birds | Brotogeris_sanctithomae | Brotogeris_sanctithomae | diurnal |
| birds | Brotogeris_tirica | Brotogeris_tirica | diurnal |
| birds | Brotogeris_versicolurus | Brotogeris_versicolurus | diurnal |

| | | | |
|---|---|---|---|
| birds | Brunhilda_charmosyna | Estrilda_charmosyna | diurnal |
| birds | Brunhilda_erythronotos | Estrilda_erythronotos | diurnal |
| birds | Bubalornis_albirostris | Bubalornis_albirostris | diurnal |
| birds | Bubalornis_niger | Bubalornis_niger | diurnal |
| birds | Bubo_africanus | Bubo_africanus | nocturnal |
| birds | Bubo_ascalaphus | Bubo_ascalaphus | nocturnal |
| birds | Bubo_bengalensis | Bubo_bengalensis | nocturnal |
| birds | Bubo_blakistoni | Ketupa_blakistoni | cathemeral |
| birds | Bubo_bubo | Bubo_bubo | cathemeral |
| birds | Bubo_capensis | Bubo_capensis | nocturnal |
| birds | Bubo_cinerascens | Bubo_cinerascens | nocturnal |
| birds | Bubo_coromandus | Bubo_coromandus | diurnal |
| birds | Bubo_lacteus | Bubo_lacteus | nocturnal |
| birds | Bubo_leucostictus | Bubo_leucostictus | nocturnal |
| birds | Bubo_magellanicus | Bubo_magellanicus | diurnal |
| birds | Bubo_nipalensis | Bubo_nipalensis | cathemeral |
| birds | Bubo_poensis | Bubo_poensis | nocturnal |
| birds | Bubo_sumatranus | Bubo_sumatranus | nocturnal |
| birds | Bubo_virginianus | Bubo_virginianus | nocturnal |
| birds | Bubulcus_ibis | Bubulcus_ibis | diurnal |
| birds | Bucanetes_githagineus | Bucanetes_githagineus | diurnal |
| birds | Bucanetes_mongolicus | Eremopsaltria_mongolicus | diurnal |
| birds | Buccanodon_duchaillui | Buccanodon_duchaillui | diurnal |
| birds | Bucco_capensis | Bucco_capensis | diurnal |
| birds | Bucephala_albeola | Bucephala_albeola | diurnal |
| birds | Bucephala_clangula | Bucephala_clangula | diurnal |
| birds | Bucephala_islandica | Bucephala_islandica | diurnal |
| birds | Buceros_bicornis | Buceros_bicornis | diurnal |
| birds | Buceros_hydrocorax | Buceros_hydrocorax | diurnal |
| birds | Buceros_rhinoceros | Buceros_rhinoceros | diurnal |
| birds | Bucorvus_abyssinicus | Bucorvus_abyssinicus | diurnal |
| birds | Bucorvus_leadbeateri | Bucorvus_cafer | diurnal |
| birds | Bugeranus_carunculatus | Grus_carunculatus | diurnal |
| birds | Bulweria_bulwerii | Bulweria_bulwerii | cathemeral |

| | | | |
|---|---|---|---|
| birds | Buphagus_africanus | Buphagus_africanus | diurnal |
| birds | Buphagus_erythrorynchus | Buphagus_erythrorhynchus | diurnal |
| birds | Burhinus_bistriatus | Burhinus_bistriatus | nocturnal |
| birds | Burhinus_capensis | Burhinus_capensis | nocturnal |
| birds | Burhinus_grallarius | Burhinus_grallarius | nocturnal |
| birds | Burhinus_oedicnemus | Burhinus_oedicnemus | nocturnal |
| birds | Burhinus_senegalensis | Burhinus_senegalensis | nocturnal |
| birds | Burhinus_superciliaris | Burhinus_superciliaris | nocturnal |
| birds | Burhinus_vermiculatus | Burhinus_vermiculatus | nocturnal |
| birds | Busarellus_nigricollis | Busarellus_nigricollis | diurnal |
| birds | Butastur_indicus | Butastur_indicus | diurnal |
| birds | Butastur_liventer | Butastur_liventer | diurnal |
| birds | Butastur_rufipennis | Butastur_rufipennis | diurnal |
| birds | Butastur_teesa | Butastur_teesa | diurnal |
| birds | Buteo_albigula | Buteo_albigula | diurnal |
| birds | Buteo_albonotatus | Buteo_albonotatus | diurnal |
| birds | Buteo_augur | Buteo_augur | diurnal |
| birds | Buteo_auguralis | Buteo_auguralis | diurnal |
| birds | Buteo_brachypterus | Buteo_brachypterus | diurnal |
| birds | Buteo_brachyurus | Buteo_brachyurus | diurnal |
| birds | Buteo_buteo | Buteo_buteo | diurnal |
| birds | Buteo_galapagoensis | Buteo_galapagoensis | diurnal |
| birds | Buteo_hemilasius | Buteo_hemilasius | diurnal |
| birds | Buteo_jamaicensis | Buteo_jamaicensis | diurnal |
| birds | Buteo_lagopus | Buteo_lagopus | diurnal |
| birds | Buteo_lineatus | Buteo_lineatus | diurnal |
| birds | Buteo_nitidus | Buteo_nitidus | diurnal |
| birds | Buteo_oreophilus | Buteo_oreophilus | diurnal |
| birds | Buteo_platypterus | Buteo_platypterus | diurnal |
| birds | Buteo_regalis | Buteo_regalis | diurnal |
| birds | Buteo_ridgwayi | Buteo_ridgwayi | diurnal |
| birds | Buteo_rufinus | Buteo_rufinus | diurnal |
| birds | Buteo_rufofuscus | Buteo_rufofuscus | diurnal |
| birds | Buteo_solitarius | Buteo_solitarius | diurnal |

| | | | |
|---|---|---|---|
| birds | Buteo_swainsoni | Buteo_swainsoni | diurnal |
| birds | Buteo_ventralis | Buteo_ventralis | diurnal |
| birds | Buteogallus_aequinoctialis | Buteogallus_aequinoctialis | diurnal |
| birds | Buteogallus_anthracinus | Buteogallus_anthracinus | diurnal |
| birds | Buteogallus_coronatus | Harpyhaliaetus_coronatus | diurnal |
| birds | Buteogallus_meridionalis | Buteogallus_meridionalis | diurnal |
| birds | Buteogallus_schistaceus | Leucopternis_schistaceus | diurnal |
| birds | Buteogallus_solitarius | Harpyhaliaetus_solitarius | diurnal |
| birds | Buteogallus_urubitinga | Buteogallus_urubitinga | diurnal |
| birds | Buthraupis_montana | Buthraupis_montana | diurnal |
| birds | Butorides_striata | Butorides_striata | cathemeral |
| birds | Bycanistes_brevis | Bycanistes_brevis | diurnal |
| birds | Bycanistes_bucinator | Bycanistes_bucinator | diurnal |
| birds | Bycanistes_cylindricus | Bycanistes_cylindricus | diurnal |
| birds | Bycanistes_fistulator | Bycanistes_fistulator | diurnal |
| birds | Bycanistes_subcylindricus | Bycanistes_subcylindricus | diurnal |
| birds | Cacatua_alba | Cacatua_alba | diurnal |
| birds | Cacatua_ducorpsii | Cacatua_ducorpsii | diurnal |
| birds | Cacatua_galerita | Cacatua_galerita | diurnal |
| birds | Cacatua_goffiniana | Cacatua_goffiniana | diurnal |
| birds | Cacatua_haematuropygia | Cacatua_haematuropygia | diurnal |
| birds | Cacatua_leadbeateri | Cacatua_leadbeateri | diurnal |
| birds | Cacatua_moluccensis | Cacatua_moluccensis | diurnal |
| birds | Cacatua_ophthalmica | Cacatua_ophthalmica | diurnal |
| birds | Cacatua_pastinator | Cacatua_pastinator | diurnal |
| birds | Cacatua_sanguinea | Cacatua_sanguinea | diurnal |
| birds | Cacatua_sulphurea | Cacatua_sulphurea | diurnal |
| birds | Cacatua_tenuirostris | Cacatua_tenuirostris | diurnal |
| birds | Cacicus_cela | Cacicus_cela | diurnal |
| birds | Cacicus_chrysonotus | Cacicus_chrysonotus | diurnal |
| birds | Cacicus_chrysopterus | Cacicus_chrysopterus | diurnal |
| birds | Cacicus_haemorrhous | Cacicus_haemorrhous | diurnal |
| birds | Cacicus_koepckeae | Cacicus_koepckeae | diurnal |
| birds | Cacicus_latirostris | Ocyalus_latirostris | diurnal |

| | | | |
|---|---|---|---|
| birds | Cacicus_microrhynchus | Cacicus_microrhynchus | diurnal |
| birds | Cacicus_oseryi | Clypicterus_oseryi | diurnal |
| birds | Cacicus_sclateri | Cacicus_sclateri | diurnal |
| birds | Cacicus_solitarius | Cacicus_solitarius | diurnal |
| birds | Cacicus_uropygialis | Cacicus_uropygialis | diurnal |
| birds | Cacomantis_castaneiventris | Cacomantis_castaneiventris | diurnal |
| birds | Cacomantis_flabelliformis | Cacomantis_flabelliformis | diurnal |
| birds | Cacomantis_merulinus | Cacomantis_merulinus | diurnal |
| birds | Cacomantis_sonneratii | Cacomantis_sonneratii | diurnal |
| birds | Cacomantis_variolosus | Cacomantis_variolosus | diurnal |
| birds | Cairina_moschata | Cairina_moschata | diurnal |
| birds | Calamanthus_campestris | Calamanthus_campestris | diurnal |
| birds | Calamanthus_fuliginosus | Calamanthus_fuliginosus | diurnal |
| birds | Calamonastes_fasciolatus | Camaroptera_fasciolata | diurnal |
| birds | Calamonastes_simplex | Camaroptera_simplex | diurnal |
| birds | Calamonastes_undosus | Camaroptera_undosa | diurnal |
| birds | Calamonastides_gracilirostris | Chloropeta_gracilirostris | diurnal |
| birds | Calamospiza_melanocorys | Calamospiza_melanocorys | diurnal |
| birds | Calandrella_acutirostris | Calandrella_acutirostris | diurnal |
| birds | Calandrella_brachydactyla | Calandrella_brachydactyla | diurnal |
| birds | Calandrella_cinerea | Calandrella_cinerea | diurnal |
| birds | Calcarius_lapponicus | Calcarius_lapponicus | diurnal |
| birds | Calcarius_ornatus | Calcarius_ornatus | diurnal |
| birds | Calcarius_pictus | Calcarius_pictus | diurnal |
| birds | Calendulauda_africanoides | Mirafra_africanoides | diurnal |
| birds | Calendulauda_albescens | Certhilauda_albescens | diurnal |
| birds | Calendulauda_burra | Certhilauda_burra | diurnal |
| birds | Calendulauda_poecilosterna | Mirafra_poecilosterna | diurnal |
| birds | Calendulauda_rufa | Mirafra_rufa | diurnal |
| birds | Calendulauda_sabota | Mirafra_sabota | diurnal |
| birds | Calherodius_leuconotus | Gorsachius_leuconotus | nocturnal |
| birds | Calicalicus_madagascariensis | Calicalicus_madagascariensis | diurnal |
| birds | Calidris_acuminata | Calidris_acuminata | diurnal |
| birds | Calidris_alba | Calidris_alba | cathemeral |

| | | | |
|---|---|---|---|
| birds | Calidris_alpina | Calidris_alpina | cathemeral |
| birds | Calidris_bairdii | Calidris_bairdii | diurnal |
| birds | Calidris_canutus | Calidris_canutus | cathemeral |
| birds | Calidris_falcinellus | Limicola_falcinellus | diurnal |
| birds | Calidris_ferruginea | Calidris_ferruginea | cathemeral |
| birds | Calidris_fuscicollis | Calidris_fuscicollis | diurnal |
| birds | Calidris_himantopus | Calidris_himantopus | diurnal |
| birds | Calidris_maritima | Calidris_maritima | diurnal |
| birds | Calidris_mauri | Calidris_mauri | diurnal |
| birds | Calidris_melanotos | Calidris_melanotos | diurnal |
| birds | Calidris_minuta | Calidris_minuta | diurnal |
| birds | Calidris_minutilla | Calidris_minutilla | cathemeral |
| birds | Calidris_ptilocnemis | Calidris_ptilocnemis | diurnal |
| birds | Calidris_pugnax | Philomachus_pugnax | cathemeral |
| birds | Calidris_pusilla | Calidris_pusilla | diurnal |
| birds | Calidris_pygmaea | Eurynorhynchus_pygmeus | diurnal |
| birds | Calidris_ruficollis | Calidris_ruficollis | diurnal |
| birds | Calidris_subminuta | Calidris_subminuta | diurnal |
| birds | Calidris_subruficollis | Tryngites_subruficollis | diurnal |
| birds | Calidris_temminckii | Calidris_temminckii | diurnal |
| birds | Calidris_tenuirostris | Calidris_tenuirostris | cathemeral |
| birds | Calidris_virgata | Aphriza_virgata | diurnal |
| birds | Caliechthrus_leucolophus | Caliechthrus_leucolophus | diurnal |
| birds | Caligavis_chrysops | Lichenostomus_chrysops | diurnal |
| birds | Caligavis_obscura | Lichenostomus_obscurus | diurnal |
| birds | Caligavis_subfrenata | Lichenostomus_subfrenatus | diurnal |
| birds | Callaeas_cinereus | Callaeas_cinereus | diurnal |
| birds | Calliope_calliope | Luscinia_calliope | diurnal |
| birds | Calliope_obscura | Luscinia_obscura | diurnal |
| birds | Calliope_pectardens | Luscinia_pectardens | diurnal |
| birds | Calliope_pectoralis | Luscinia_pectoralis | diurnal |
| birds | Callipepla_californica | Callipepla_californica | diurnal |
| birds | Callipepla_douglasii | Callipepla_douglasii | diurnal |
| birds | Callipepla_gambelii | Callipepla_gambelii | diurnal |

| | | | |
|---|---|---|---|
| birds | Callipepla_squamata | Callipepla_squamata | diurnal |
| birds | Calliphlox_amethystina | Calliphlox_amethystina | diurnal |
| birds | Callocephalon_fimbriatum | Callocephalon_fimbriatum | diurnal |
| birds | Callonetta_leucophrys | Callonetta_leucophrys | diurnal |
| birds | Calochaetes_coccineus | Calochaetes_coccineus | diurnal |
| birds | Calonectris_diomedea | Calonectris_diomedea | cathemeral |
| birds | Calonectris_leucomelas | Calonectris_leucomelas | cathemeral |
| birds | Caloperdix_oculeus | Caloperdix_oculeus | diurnal |
| birds | Caloramphus_fuliginosus | Calorhamphus_fuliginosus | diurnal |
| birds | Calothorax_lucifer | Calothorax_lucifer | diurnal |
| birds | Calothorax_pulcher | Calothorax_pulcher | diurnal |
| birds | Calypte_anna | Calypte_anna | diurnal |
| birds | Calypte_costae | Calypte_costae | diurnal |
| birds | Calyptocichla_serinus | Calyptocichla_serina | diurnal |
| birds | Calyptomena_hosii | Calyptomena_hosii | diurnal |
| birds | Calyptomena_viridis | Calyptomena_viridis | diurnal |
| birds | Calyptomena_whiteheadi | Calyptomena_whiteheadi | diurnal |
| birds | Calyptophilus_frugivorus | Calyptophilus_frugivorus | diurnal |
| birds | Calyptophilus_tertius | Calyptophilus_tertius | diurnal |
| birds | Calyptorhynchus_banksii | Calyptorhynchus_banksii | diurnal |
| birds | Calyptorhynchus_lathami | Calyptorhynchus_lathami | diurnal |
| birds | Calyptura_cristata | Calyptura_cristata | diurnal |
| birds | Camaroptera_brachyura | Camaroptera_brachyura | diurnal |
| birds | Camaroptera_chloronota | Camaroptera_chloronota | diurnal |
| birds | Camaroptera_superciliaris | Camaroptera_superciliaris | diurnal |
| birds | Campephaga_flava | Campephaga_flava | diurnal |
| birds | Campephaga_petiti | Campephaga_petiti | diurnal |
| birds | Campephaga_phoenicea | Campephaga_phoenicea | diurnal |
| birds | Campephaga_quiscalina | Campephaga_quiscalina | diurnal |
| birds | Campephilus_gayaquilensis | Campephilus_gayaquilensis | diurnal |
| birds | Campephilus_guatemalensis | Campephilus_guatemalensis | diurnal |
| birds | Campephilus_haematogaster | Campephilus_haematogaster | diurnal |
| birds | Campephilus_imperialis | Campephilus_imperialis | diurnal |
| birds | Campephilus_leucopogon | Campephilus_leucopogon | diurnal |

| | | | |
|---|---|---|---|
| birds | Campephilus_magellanicus | Campephilus_magellanicus | diurnal |
| birds | Campephilus_melanoleucos | Campephilus_melanoleucos | diurnal |
| birds | Campephilus_pollens | Campephilus_pollens | diurnal |
| birds | Campephilus_principalis | Campephilus_principalis | diurnal |
| birds | Campephilus_robustus | Campephilus_robustus | diurnal |
| birds | Campephilus_rubricollis | Campephilus_rubricollis | diurnal |
| birds | Campethera_abingoni | Campethera_abingoni | diurnal |
| birds | Campethera_bennettii | Campethera_bennettii | diurnal |
| birds | Campethera_cailliautii | Campethera_cailliautii | diurnal |
| birds | Campethera_maculosa | Campethera_maculosa | diurnal |
| birds | Campethera_mombassica | Campethera_mombassica | diurnal |
| birds | Campethera_notata | Campethera_notata | diurnal |
| birds | Campethera_nubica | Campethera_nubica | diurnal |
| birds | Campethera_punctuligera | Campethera_punctuligera | diurnal |
| birds | Campethera_tullbergi | Campethera_tullbergi | diurnal |
| birds | Campicoloides_bifasciatus | Oenanthe_bifasciata | diurnal |
| birds | Campochaera_sloetii | Campochaera_sloetii | diurnal |
| birds | Campocolinus_albogularis | Francolinus_albogularis | diurnal |
| birds | Campocolinus_coqui | Francolinus_coqui | diurnal |
| birds | Campocolinus_schlegelii | Francolinus_schlegelii | diurnal |
| birds | Camptostoma_imberbe | Camptostoma_imberbe | diurnal |
| birds | Camptostoma_obsoletum | Camptostoma_obsoletum | diurnal |
| birds | Campylopterus_duidae | Campylopterus_duidae | diurnal |
| birds | Campylopterus_ensipennis | Campylopterus_ensipennis | diurnal |
| birds | Campylopterus_falcatus | Campylopterus_falcatus | diurnal |
| birds | Campylopterus_hemileucurus | Campylopterus_hemileucurus | diurnal |
| birds | Campylopterus_hyperythrus | Campylopterus_hyperythrus | diurnal |
| birds | Campylopterus_largipennis | Campylopterus_largipennis | diurnal |
| birds | Campylopterus_phainopeplus | Campylopterus_phainopeplus | diurnal |
| birds | Campylopterus_villaviscensio | Campylopterus_villaviscensio | diurnal |
| birds | Campylorhamphus_falcularius | Campylorhamphus_falcularius | diurnal |
| birds | Campylorhamphus_procurvoides | Campylorhamphus_procurvoides | diurnal |
| birds | Campylorhamphus_pusillus | Campylorhamphus_pusillus | diurnal |
| birds | Campylorhamphus_trochilirostris | Campylorhamphus_trochilirostris | diurnal |

| | | | |
|---|---|---|---|
| birds | Campylorhynchus_albobrunneus | Campylorhynchus_albobrunneus | diurnal |
| birds | Campylorhynchus_brunneicapillus | Campylorhynchus_brunneicapillus | diurnal |
| birds | Campylorhynchus_chiapensis | Campylorhynchus_chiapensis | diurnal |
| birds | Campylorhynchus_fasciatus | Campylorhynchus_fasciatus | diurnal |
| birds | Campylorhynchus_griseus | Campylorhynchus_griseus | diurnal |
| birds | Campylorhynchus_gularis | Campylorhynchus_gularis | diurnal |
| birds | Campylorhynchus_jocosus | Campylorhynchus_jocosus | diurnal |
| birds | Campylorhynchus_megalopterus | Campylorhynchus_megalopterus | diurnal |
| birds | Campylorhynchus_nuchalis | Campylorhynchus_nuchalis | diurnal |
| birds | Campylorhynchus_rufinucha | Campylorhynchus_rufinucha | diurnal |
| birds | Campylorhynchus_turdinus | Campylorhynchus_turdinus | diurnal |
| birds | Campylorhynchus_yucatanicus | Campylorhynchus_yucatanicus | diurnal |
| birds | Campylorhynchus_zonatus | Campylorhynchus_zonatus | diurnal |
| birds | Canachites_canadensis | Dendragapus_canadensis | diurnal |
| birds | Canirallus_oculeus | Canirallus_oculeus | diurnal |
| birds | Cantorchilus_griseus | Thryothorus_griseus | diurnal |
| birds | Cantorchilus_guarayanus | Thryothorus_guarayanus | diurnal |
| birds | Cantorchilus_leucopogon | Thryothorus_leucopogon | diurnal |
| birds | Cantorchilus_leucotis | Thryothorus_leucotis | diurnal |
| birds | Cantorchilus_longirostris | Thryothorus_longirostris | diurnal |
| birds | Cantorchilus_modestus | Thryothorus_modestus | diurnal |
| birds | Cantorchilus_nigricapillus | Thryothorus_nigricapillus | diurnal |
| birds | Cantorchilus_semibadius | Thryothorus_semibadius | diurnal |
| birds | Cantorchilus_superciliaris | Thryothorus_superciliaris | diurnal |
| birds | Cantorchilus_thoracicus | Thryothorus_thoracicus | diurnal |
| birds | Capito_aurovirens | Capito_aurovirens | diurnal |
| birds | Capito_brunneipectus | Capito_brunneipectus | diurnal |
| birds | Capito_dayi | Capito_dayi | diurnal |
| birds | Capito_hypoleucus | Capito_hypoleucus | diurnal |
| birds | Capito_maculicoronatus | Capito_maculicoronatus | diurnal |
| birds | Capito_niger | Capito_niger | diurnal |
| birds | Capito_quinticolor | Capito_quinticolor | diurnal |
| birds | Capito_squamatus | Capito_squamatus | diurnal |
| birds | Capito_wallacei | Capito_wallacei | diurnal |

| | | | |
|---|---|---|---|
| birds | Caprimulgus_aegyptius | Caprimulgus_aegyptius | nocturnal |
| birds | Caprimulgus_affinis | Caprimulgus_affinis | nocturnal |
| birds | Caprimulgus_asiaticus | Caprimulgus_asiaticus | nocturnal |
| birds | Caprimulgus_atripennis | Caprimulgus_atripennis | nocturnal |
| birds | Caprimulgus_batesi | Caprimulgus_batesi | nocturnal |
| birds | Caprimulgus_celebensis | Caprimulgus_celebensis | nocturnal |
| birds | Caprimulgus_clarus | Caprimulgus_clarus | nocturnal |
| birds | Caprimulgus_climacurus | Caprimulgus_climacurus | nocturnal |
| birds | Caprimulgus_concretus | Caprimulgus_concretus | nocturnal |
| birds | Caprimulgus_donaldsoni | Caprimulgus_donaldsoni | nocturnal |
| birds | Caprimulgus_europaeus | Caprimulgus_europaeus | cathemeral |
| birds | Caprimulgus_fossii | Caprimulgus_fossii | nocturnal |
| birds | Caprimulgus_fraenatus | Caprimulgus_fraenatus | nocturnal |
| birds | Caprimulgus_indicus | Caprimulgus_indicus | nocturnal |
| birds | Caprimulgus_inornatus | Caprimulgus_inornatus | nocturnal |
| birds | Caprimulgus_longipennis | Macrodipteryx_longipennis | nocturnal |
| birds | Caprimulgus_macrurus | Caprimulgus_macrurus | nocturnal |
| birds | Caprimulgus_madagascariensis | Caprimulgus_madagascariensis | nocturnal |
| birds | Caprimulgus_mahrattensis | Caprimulgus_mahrattensis | nocturnal |
| birds | Caprimulgus_manillensis | Caprimulgus_manillensis | nocturnal |
| birds | Caprimulgus_natalensis | Caprimulgus_natalensis | nocturnal |
| birds | Caprimulgus_nubicus | Caprimulgus_nubicus | nocturnal |
| birds | Caprimulgus_pectoralis | Caprimulgus_pectoralis | nocturnal |
| birds | Caprimulgus_poliocephalus | Caprimulgus_poliocephalus | nocturnal |
| birds | Caprimulgus_pulchellus | Caprimulgus_pulchellus | nocturnal |
| birds | Caprimulgus_ruficollis | Caprimulgus_ruficollis | nocturnal |
| birds | Caprimulgus_rufigena | Caprimulgus_rufigena | nocturnal |
| birds | Caprimulgus_tristigma | Caprimulgus_tristigma | nocturnal |
| birds | Caprimulgus_vexillarius | Macrodipteryx_vexillarius | cathemeral |
| birds | Capsiempis_flaveola | Capsiempis_flaveola | diurnal |
| birds | Caracara_plancus | Caracara_plancus | diurnal |
| birds | Cardellina_canadensis | Wilsonia_canadensis | diurnal |
| birds | Cardellina_pusilla | Wilsonia_pusilla | diurnal |
| birds | Cardellina_rubra | Ergaticus_ruber | diurnal |

| | | | |
|---|---|---|---|
| birds | Cardellina_rubrifrons | Cardellina_rubrifrons | diurnal |
| birds | Cardellina_versicolor | Ergaticus_versicolor | diurnal |
| birds | Cardinalis_cardinalis | Cardinalis_cardinalis | diurnal |
| birds | Cardinalis_phoeniceus | Cardinalis_phoeniceus | diurnal |
| birds | Cardinalis_sinuatus | Cardinalis_sinuatus | diurnal |
| birds | Carduelis_carduelis | Carduelis_carduelis | diurnal |
| birds | Carduelis_citrinella | Carduelis_citrinella | diurnal |
| birds | Cariama_cristata | Cariama_cristata | diurnal |
| birds | Caridonax_fulgidus | Caridonax_fulgidus | diurnal |
| birds | Carpococcyx_radiceus | Carpococcyx_radiatus | diurnal |
| birds | Carpococcyx_renauldi | Carpococcyx_renauldi | diurnal |
| birds | Carpodacus_edwardsii | Carpodacus_edwardsii | diurnal |
| birds | Carpodacus_erythrinus | Carpodacus_erythrinus | diurnal |
| birds | Carpodacus_pulcherrimus | Carpodacus_pulcherrimus | diurnal |
| birds | Carpodacus_puniceus | Carpodacus_puniceus | diurnal |
| birds | Carpodacus_rhodochlamys | Carpodacus_rhodochlamys | diurnal |
| birds | Carpodacus_rodochroa | Carpodacus_rodochroa | diurnal |
| birds | Carpodacus_rodopeplus | Carpodacus_rodopeplus | diurnal |
| birds | Carpodacus_roseus | Carpodacus_roseus | diurnal |
| birds | Carpodacus_rubicilla | Carpodacus_rubicilla | diurnal |
| birds | Carpodacus_rubicilloides | Carpodacus_rubicilloides | diurnal |
| birds | Carpodacus_sibiricus | Uragus_sibiricus | diurnal |
| birds | Carpodacus_sipahi | Haematospiza_sipahi | diurnal |
| birds | Carpodacus_subhimachalus | Pinicola_subhimachala | diurnal |
| birds | Carpodacus_synoicus | Carpodacus_synoicus | diurnal |
| birds | Carpodacus_thura | Carpodacus_thura | diurnal |
| birds | Carpodacus_vinaceus | Carpodacus_vinaceus | diurnal |
| birds | Carpodacus_waltoni | Carpodacus_eos | diurnal |
| birds | Carpodectes_antoniae | Carpodectes_antoniae | diurnal |
| birds | Carpodectes_hopkei | Carpodectes_hopkei | diurnal |
| birds | Carpodectes_nitidus | Carpodectes_nitidus | diurnal |
| birds | Carpornis_cucullata | Carpornis_cucullata | diurnal |
| birds | Carpornis_melanocephala | Carpornis_melanocephala | diurnal |
| birds | Carpospiza_brachydactyla | Petronia_brachydactyla | diurnal |

| | | | |
|---|---|---|---|
| birds | Carterornis_chrysomela | Monarcha_chrysomela | diurnal |
| birds | Carterornis_leucotis | Monarcha_leucotis | diurnal |
| birds | Carterornis_pileatus | Monarcha_pileatus | diurnal |
| birds | Caryothraustes_canadensis | Caryothraustes_canadensis | diurnal |
| birds | Caryothraustes_celaeno | Rhodothraupis_celaeno | diurnal |
| birds | Caryothraustes_erythromelas | Periporphyrus_erythromelas | diurnal |
| birds | Caryothraustes_poliogaster | Caryothraustes_poliogaster | diurnal |
| birds | Casiornis_fuscus | Casiornis_fuscus | diurnal |
| birds | Casiornis_rufus | Casiornis_rufus | diurnal |
| birds | Cassiculus_melanicterus | Cacicus_melanicterus | diurnal |
| birds | Castanozoster_thoracicus | Poospiza_thoracica | diurnal |
| birds | Casuarius_bennetti | Casuarius_bennetti | diurnal |
| birds | Casuarius_casuarius | Casuarius_casuarius | diurnal |
| birds | Casuarius_unappendiculatus | Casuarius_unappendiculatus | diurnal |
| birds | Catamblyrhynchus_diadema | Catamblyrhynchus_diadema | diurnal |
| birds | Catamenia_analis | Catamenia_analis | diurnal |
| birds | Catamenia_homochroa | Catamenia_homochroa | diurnal |
| birds | Catamenia_inornata | Catamenia_inornata | diurnal |
| birds | Cataponera_turdoides | Cataponera_turdoides | diurnal |
| birds | Catharacta_antarctica | Catharacta_antarctica | nocturnal |
| birds | Catharacta_chilensis | Catharacta_chilensis | diurnal |
| birds | Catharacta_maccormicki | Catharacta_maccormicki | diurnal |
| birds | Catharacta_skua | Catharacta_skua | diurnal |
| birds | Catharopeza_bishopi | Catharopeza_bishopi | diurnal |
| birds | Cathartes_aura | Cathartes_aura | diurnal |
| birds | Cathartes_burrovianus | Cathartes_burrovianus | diurnal |
| birds | Cathartes_melambrotus | Cathartes_melambrotus | diurnal |
| birds | Catharus_aurantiirostris | Catharus_aurantiirostris | diurnal |
| birds | Catharus_bicknelli | Catharus_bicknelli | diurnal |
| birds | Catharus_dryas | Catharus_dryas | diurnal |
| birds | Catharus_frantzii | Catharus_frantzii | diurnal |
| birds | Catharus_fuscater | Catharus_fuscater | diurnal |
| birds | Catharus_fuscescens | Catharus_fuscescens | diurnal |
| birds | Catharus_gracilirostris | Catharus_gracilirostris | diurnal |

| | | | |
|---|---|---|---|
| birds | Catharus_guttatus | Catharus_guttatus | diurnal |
| birds | Catharus_mexicanus | Catharus_mexicanus | diurnal |
| birds | Catharus_minimus | Catharus_minimus | diurnal |
| birds | Catharus_occidentalis | Catharus_occidentalis | diurnal |
| birds | Catharus_ustulatus | Catharus_ustulatus | diurnal |
| birds | Catherpes_mexicanus | Catherpes_mexicanus | diurnal |
| birds | Catreus_wallichii | Catreus_wallichi | diurnal |
| birds | Catriscus_brevirostris | Schoenicola_brevirostris | diurnal |
| birds | Ceblepyris_caesius | Coracina_caesia | diurnal |
| birds | Ceblepyris_cinereus | Coracina_cinerea | diurnal |
| birds | Ceblepyris_pectoralis | Coracina_pectoralis | diurnal |
| birds | Cecropis_abyssinica | Hirundo_abyssinica | diurnal |
| birds | Cecropis_cucullata | Hirundo_cucullata | diurnal |
| birds | Cecropis_daurica | Hirundo_daurica | diurnal |
| birds | Cecropis_semirufa | Hirundo_semirufa | diurnal |
| birds | Cecropis_senegalensis | Hirundo_senegalensis | diurnal |
| birds | Celebesica_abbotti | Coracina_abbotti | diurnal |
| birds | Celeus_castaneus | Celeus_castaneus | diurnal |
| birds | Celeus_elegans | Celeus_elegans | diurnal |
| birds | Celeus_flavescens | Celeus_flavescens | diurnal |
| birds | Celeus_flavus | Celeus_flavus | diurnal |
| birds | Celeus_galeatus | Dryocopus_galeatus | diurnal |
| birds | Celeus_loricatus | Celeus_loricatus | diurnal |
| birds | Celeus_lugubris | Celeus_lugubris | diurnal |
| birds | Celeus_spectabilis | Celeus_spectabilis | diurnal |
| birds | Celeus_torquatus | Celeus_torquatus | diurnal |
| birds | Celeus_undatus | Celeus_undatus | diurnal |
| birds | Centrocercus_urophasianus | Centrocercus_urophasianus | diurnal |
| birds | Centropus_andamanensis | Centropus_andamanensis | diurnal |
| birds | Centropus_anselli | Centropus_anselli | diurnal |
| birds | Centropus_ateralbus | Centropus_ateralbus | diurnal |
| birds | Centropus_bengalensis | Centropus_bengalensis | diurnal |
| birds | Centropus_bernsteini | Centropus_bernsteini | diurnal |
| birds | Centropus_celebensis | Centropus_celebensis | diurnal |

| | | | |
|---|---|---|---|
| birds | Centropus_chalybeus | Centropus_chalybeus | diurnal |
| birds | Centropus_chlororhynchos | Centropus_chlororhynchus | diurnal |
| birds | Centropus_cupreicaudus | Centropus_cupreicaudus | diurnal |
| birds | Centropus_goliath | Centropus_goliath | diurnal |
| birds | Centropus_grillii | Centropus_grillii | diurnal |
| birds | Centropus_leucogaster | Centropus_leucogaster | diurnal |
| birds | Centropus_melanops | Centropus_melanops | diurnal |
| birds | Centropus_menbeki | Centropus_menbeki | diurnal |
| birds | Centropus_milo | Centropus_milo | diurnal |
| birds | Centropus_monachus | Centropus_monachus | diurnal |
| birds | Centropus_nigrorufus | Centropus_nigrorufus | diurnal |
| birds | Centropus_phasianinus | Centropus_phasianinus | diurnal |
| birds | Centropus_rectunguis | Centropus_rectunguis | diurnal |
| birds | Centropus_senegalensis | Centropus_senegalensis | diurnal |
| birds | Centropus_sinensis | Centropus_sinensis | diurnal |
| birds | Centropus_spilopterus | Centropus_spilopterus | diurnal |
| birds | Centropus_steerii | Centropus_steerii | diurnal |
| birds | Centropus_superciliosus | Centropus_superciliosus | diurnal |
| birds | Centropus_toulou | Centropus_toulou | diurnal |
| birds | Centropus_unirufus | Centropus_unirufus | diurnal |
| birds | Centropus_violaceus | Centropus_violaceus | diurnal |
| birds | Centropus_viridis | Centropus_viridis | diurnal |
| birds | Cephalopterus_glabricollis | Cephalopterus_glabricollis | diurnal |
| birds | Cephalopterus_ornatus | Cephalopterus_ornatus | diurnal |
| birds | Cephalopterus_penduliger | Cephalopterus_penduliger | diurnal |
| birds | Cephalopyrus_flammiceps | Cephalopyrus_flammiceps | diurnal |
| birds | Cepphus_carbo | Cepphus_carbo | diurnal |
| birds | Cepphus_columba | Cepphus_columba | diurnal |
| birds | Cepphus_grylle | Cepphus_grylle | diurnal |
| birds | Ceratogymna_atrata | Ceratogymna_atrata | diurnal |
| birds | Ceratogymna_elata | Ceratogymna_elata | diurnal |
| birds | Ceratopipra_chloromeros | Pipra_chloromeros | diurnal |
| birds | Ceratopipra_cornuta | Pipra_cornuta | diurnal |
| birds | Ceratopipra_erythrocephala | Pipra_erythrocephala | diurnal |

| | | | |
|---|---|---|---|
| birds | Ceratopipra_mentalis | Pipra_mentalis | diurnal |
| birds | Ceratopipra_rubrocapilla | Pipra_rubrocapilla | diurnal |
| birds | Cercibis_oxycerca | Cercibis_oxycerca | diurnal |
| birds | Cercococcyx_mechowi | Cercococcyx_mechowi | diurnal |
| birds | Cercococcyx_montanus | Cercococcyx_montanus | diurnal |
| birds | Cercococcyx_olivinus | Cercococcyx_olivinus | diurnal |
| birds | Cercomacra_brasiliana | Cercomacra_brasiliana | diurnal |
| birds | Cercomacra_carbonaria | Cercomacra_carbonaria | diurnal |
| birds | Cercomacra_cinerascens | Cercomacra_cinerascens | diurnal |
| birds | Cercomacra_ferdinandi | Cercomacra_ferdinandi | diurnal |
| birds | Cercomacra_manu | Cercomacra_manu | diurnal |
| birds | Cercomacra_melanaria | Cercomacra_melanaria | diurnal |
| birds | Cercomacra_nigricans | Cercomacra_nigricans | diurnal |
| birds | Cercomacroides_laeta | Cercomacra_laeta | diurnal |
| birds | Cercomacroides_nigrescens | Cercomacra_nigrescens | diurnal |
| birds | Cercomacroides_parkeri | Cercomacra_parkeri | diurnal |
| birds | Cercomacroides_serva | Cercomacra_serva | diurnal |
| birds | Cercomacroides_tyrannina | Cercomacra_tyrannina | diurnal |
| birds | Cercotrichas_galactotes | Erythropygia_galactotes | diurnal |
| birds | Cercotrichas_hartlaubi | Erythropygia_hartlaubi | diurnal |
| birds | Cercotrichas_leucophrys | Erythropygia_leucophrys | diurnal |
| birds | Cercotrichas_paena | Erythropygia_paena | diurnal |
| birds | Cercotrichas_podobe | Cercotrichas_podobe | diurnal |
| birds | Cereopsis_novaehollandiae | Cereopsis_novaehollandiae | diurnal |
| birds | Cerorhinca_monocerata | Cerorhinca_monocerata | diurnal |
| birds | Certhia_americana | Certhia_americana | diurnal |
| birds | Certhia_brachydactyla | Certhia_brachydactyla | diurnal |
| birds | Certhia_discolor | Certhia_discolor | diurnal |
| birds | Certhia_familiaris | Certhia_familiaris | diurnal |
| birds | Certhia_himalayana | Certhia_himalayana | diurnal |
| birds | Certhia_nipalensis | Certhia_nipalensis | diurnal |
| birds | Certhiasomus_stictolaemus | Deconychura_stictolaema | diurnal |
| birds | Certhiaxis_cinnamomeus | Certhiaxis_cinnamomeus | diurnal |
| birds | Certhiaxis_mustelinus | Certhiaxis_mustelinus | diurnal |

| | | | |
|---|---|---|---|
| birds | Certhidea_olivacea | Certhidea_olivacea | diurnal |
| birds | Certhilauda_chuana | Certhilauda_chuana | diurnal |
| birds | Certhilauda_curvirostris | Certhilauda_curvirostris | diurnal |
| birds | Certhionyx_variegatus | Certhionyx_variegatus | diurnal |
| birds | Cettia_brunnifrons | Cettia_brunnifrons | diurnal |
| birds | Cettia_castaneocoronata | Tesia_castaneocoronata | diurnal |
| birds | Cettia_cetti | Cettia_cetti | diurnal |
| birds | Cettia_major | Cettia_major | diurnal |
| birds | Ceuthmochares_aereus | Ceuthmochares_aereus | diurnal |
| birds | Ceyx_argentatus | Alcedo_argentata | diurnal |
| birds | Ceyx_azureus | Alcedo_azurea | diurnal |
| birds | Ceyx_cyanopectus | Alcedo_cyanopectus | diurnal |
| birds | Ceyx_erithaca | Ceyx_erithaca | diurnal |
| birds | Ceyx_fallax | Ceyx_fallax | diurnal |
| birds | Ceyx_lepidus | Ceyx_lepidus | diurnal |
| birds | Ceyx_melanurus | Ceyx_melanurus | diurnal |
| birds | Ceyx_pusillus | Alcedo_pusilla | diurnal |
| birds | Ceyx_websteri | Alcedo_websteri | diurnal |
| birds | Chaetocercus_berlepschi | Chaetocercus_berlepschi | diurnal |
| birds | Chaetocercus_bombus | Chaetocercus_bombus | diurnal |
| birds | Chaetocercus_heliodor | Chaetocercus_heliodor | diurnal |
| birds | Chaetocercus_jourdanii | Chaetocercus_jourdanii | diurnal |
| birds | Chaetocercus_mulsant | Chaetocercus_mulsant | diurnal |
| birds | Chaetops_frenatus | Chaetops_frenatus | diurnal |
| birds | Chaetorhynchus_papuensis | Chaetorhynchus_papuensis | diurnal |
| birds | Chaetura_brachyura | Chaetura_brachyura | diurnal |
| birds | Chaetura_chapmani | Chaetura_chapmani | diurnal |
| birds | Chaetura_cinereiventris | Chaetura_cinereiventris | diurnal |
| birds | Chaetura_egregia | Chaetura_egregia | diurnal |
| birds | Chaetura_martinica | Chaetura_martinica | diurnal |
| birds | Chaetura_pelagica | Chaetura_pelagica | diurnal |
| birds | Chaetura_vauxi | Chaetura_vauxi | diurnal |
| birds | Chalcites_basalis | Chrysococcyx_basalis | diurnal |
| birds | Chalcites_lucidus | Chrysococcyx_lucidus | diurnal |

| | | | |
|---|---|---|---|
| birds | Chalcites_megarhynchus | Rhamphomantis_megarhynchus | diurnal |
| birds | Chalcites_meyerii | Chrysococcyx_meyeri | diurnal |
| birds | Chalcites_minutillus | Chrysococcyx_minutillus | diurnal |
| birds | Chalcites_osculans | Chrysococcyx_osculans | diurnal |
| birds | Chalcites_ruficollis | Chrysococcyx_ruficollis | diurnal |
| birds | Chalcomitra_adelberti | Nectarinia_adelberti | diurnal |
| birds | Chalcomitra_amethystina | Nectarinia_amethystina | diurnal |
| birds | Chalcomitra_balfouri | Nectarinia_balfouri | diurnal |
| birds | Chalcomitra_fuliginosa | Nectarinia_fuliginosa | diurnal |
| birds | Chalcomitra_hunteri | Nectarinia_hunteri | diurnal |
| birds | Chalcomitra_rubescens | Nectarinia_rubescens | diurnal |
| birds | Chalcomitra_senegalensis | Nectarinia_senegalensis | diurnal |
| birds | Chalcoparia_singalensis | Anthreptes_singalensis | diurnal |
| birds | Chalcophaps_indica | Chalcophaps_indica | diurnal |
| birds | Chalcophaps_stephani | Chalcophaps_stephani | diurnal |
| birds | Chalcopsitta_atra | Chalcopsitta_atra | diurnal |
| birds | Chalcopsitta_cardinalis | Chalcopsitta_cardinalis | diurnal |
| birds | Chalcopsitta_duivenbodei | Chalcopsitta_duivenbodei | diurnal |
| birds | Chalcopsitta_scintillata | Chalcopsitta_sintillata | diurnal |
| birds | Chalcostigma_herrani | Chalcostigma_herrani | diurnal |
| birds | Chalcostigma_heteropogon | Chalcostigma_heteropogon | diurnal |
| birds | Chalcostigma_olivaceum | Chalcostigma_olivaceum | diurnal |
| birds | Chalcostigma_ruficeps | Chalcostigma_ruficeps | diurnal |
| birds | Chalcostigma_stanleyi | Chalcostigma_stanleyi | diurnal |
| birds | Chalybura_buffonii | Chalybura_buffonii | diurnal |
| birds | Chalybura_urochrysia | Chalybura_urochrysia | diurnal |
| birds | Chamaea_fasciata | Chamaea_fasciata | diurnal |
| birds | Chamaepetes_unicolor | Chamaepetes_unicolor | diurnal |
| birds | Chamaetylas_choloensis | Alethe_choloensis | diurnal |
| birds | Chamaetylas_fuelleborni | Alethe_fuelleborni | diurnal |
| birds | Chamaetylas_poliophrys | Alethe_poliophrys | diurnal |
| birds | Chamaeza_campanisona | Chamaeza_campanisona | diurnal |
| birds | Chamaeza_meruloides | Chamaeza_meruloides | diurnal |
| birds | Chamaeza_mollissima | Chamaeza_mollissima | diurnal |

| | | | |
|---|---|---|---|
| birds | Chamaeza_nobilis | Chamaeza_nobilis | diurnal |
| birds | Chamaeza_ruficauda | Chamaeza_ruficauda | diurnal |
| birds | Chamaeza_turdina | Chamaeza_turdina | diurnal |
| birds | Charadrius_alexandrinus | Charadrius_alexandrinus | diurnal |
| birds | Charadrius_alticola | Charadrius_alticola | diurnal |
| birds | Charadrius_asiaticus | Charadrius_asiaticus | diurnal |
| birds | Charadrius_bicinctus | Charadrius_bicinctus | cathemeral |
| birds | Charadrius_collaris | Charadrius_collaris | diurnal |
| birds | Charadrius_dubius | Charadrius_dubius | diurnal |
| birds | Charadrius_falklandicus | Charadrius_falklandicus | diurnal |
| birds | Charadrius_hiaticula | Charadrius_hiaticula | cathemeral |
| birds | Charadrius_leschenaultii | Charadrius_leschenaultii | diurnal |
| birds | Charadrius_marginatus | Charadrius_marginatus | cathemeral |
| birds | Charadrius_melodus | Charadrius_melodus | cathemeral |
| birds | Charadrius_modestus | Charadrius_modestus | diurnal |
| birds | Charadrius_montanus | Charadrius_montanus | diurnal |
| birds | Charadrius_obscurus | Charadrius_obscurus | cathemeral |
| birds | Charadrius_pecuarius | Charadrius_pecuarius | cathemeral |
| birds | Charadrius_peronii | Charadrius_peronii | diurnal |
| birds | Charadrius_placidus | Charadrius_placidus | diurnal |
| birds | Charadrius_ruficapillus | Charadrius_ruficapillus | diurnal |
| birds | Charadrius_sanctaehelenae | Charadrius_sanctaehelenae | diurnal |
| birds | Charadrius_semipalmatus | Charadrius_semipalmatus | cathemeral |
| birds | Charadrius_thoracicus | Charadrius_thoracicus | diurnal |
| birds | Charadrius_tricollaris | Charadrius_tricollaris | cathemeral |
| birds | Charadrius_veredus | Charadrius_veredus | cathemeral |
| birds | Charadrius_vociferus | Charadrius_vociferus | cathemeral |
| birds | Charadrius_wilsonia | Charadrius_wilsonia | cathemeral |
| birds | Charitospiza_eucosma | Charitospiza_eucosma | diurnal |
| birds | Charmosyna_amabilis | Charmosyna_amabilis | diurnal |
| birds | Charmosyna_diadema | Charmosyna_diadema | diurnal |
| birds | Charmosyna_josefinae | Charmosyna_josefinae | diurnal |
| birds | Charmosyna_margarethae | Charmosyna_margarethae | diurnal |
| birds | Charmosyna_meeki | Charmosyna_meeki | diurnal |

| | | | |
|---|---|---|---|
| birds | Charmosyna_multistriata | Charmosyna_multistriata | diurnal |
| birds | Charmosyna_palmarum | Charmosyna_palmarum | diurnal |
| birds | Charmosyna_papou | Charmosyna_papou | diurnal |
| birds | Charmosyna_placentis | Charmosyna_placentis | diurnal |
| birds | Charmosyna_pulchella | Charmosyna_pulchella | diurnal |
| birds | Charmosyna_rubrigularis | Charmosyna_rubrigularis | diurnal |
| birds | Charmosyna_rubronotata | Charmosyna_rubronotata | diurnal |
| birds | Charmosyna_toxopei | Charmosyna_toxopei | diurnal |
| birds | Charmosyna_wilhelminae | Charmosyna_wilhelminae | diurnal |
| birds | Chasiempis_sandwichensis | Chasiempis_sandwichensis | diurnal |
| birds | Chauna_chavaria | Chauna_chavaria | diurnal |
| birds | Chauna_torquata | Chauna_torquata | diurnal |
| birds | Chelictinia_riocourii | Chelictinia_riocourii | diurnal |
| birds | Chelidoptera_tenebrosa | Chelidoptera_tenebrosa | diurnal |
| birds | Chelidorhynx_hypoxanthus | Chelidorhynx_hypoxantha | diurnal |
| birds | Chenonetta_jubata | Chenonetta_jubata | diurnal |
| birds | Chenorhamphus_grayi | Malurus_grayi | diurnal |
| birds | Cheramoeca_leucosterna | Cheramoeca_leucosterna | diurnal |
| birds | Chersophilus_duponti | Chersophilus_duponti | diurnal |
| birds | Chionis_albus | Chionis_albus | diurnal |
| birds | Chionis_minor | Chionis_minor | diurnal |
| birds | Chionomesa_fimbriata | Amazilia_fimbriata | diurnal |
| birds | Chiroxiphia_boliviana | Chiroxiphia_boliviana | diurnal |
| birds | Chiroxiphia_caudata | Chiroxiphia_caudata | diurnal |
| birds | Chiroxiphia_lanceolata | Chiroxiphia_lanceolata | diurnal |
| birds | Chiroxiphia_linearis | Chiroxiphia_linearis | diurnal |
| birds | Chiroxiphia_pareola | Chiroxiphia_pareola | diurnal |
| birds | Chlamydera_cerviniventris | Chlamydera_cerviniventris | diurnal |
| birds | Chlamydera_guttata | Chlamydera_guttata | diurnal |
| birds | Chlamydera_lauterbachi | Chlamydera_lauterbachi | diurnal |
| birds | Chlamydera_maculata | Chlamydera_maculata | diurnal |
| birds | Chlamydera_nuchalis | Chlamydera_nuchalis | diurnal |
| birds | Chlamydochaera_jefferyi | Chlamydochaera_jefferyi | diurnal |
| birds | Chlamydotis_undulata | Chlamydotis_undulata | diurnal |

| | | | |
|---|---|---|---|
| birds | Chlidonias_albostriatus | Sterna_albostriata | diurnal |
| birds | Chlidonias_hybrida | Chlidonias_hybrida | diurnal |
| birds | Chlidonias_leucopterus | Chlidonias_leucopterus | diurnal |
| birds | Chlidonias_niger | Chlidonias_niger | diurnal |
| birds | Chloebia_gouldiae | Erythrura_gouldiae | diurnal |
| birds | Chloephaga_hybrida | Chloephaga_hybrida | diurnal |
| birds | Chloephaga_melanoptera | Chloephaga_melanoptera | diurnal |
| birds | Chloephaga_picta | Chloephaga_picta | diurnal |
| birds | Chloephaga_poliocephala | Chloephaga_poliocephala | diurnal |
| birds | Chloephaga_rubidiceps | Chloephaga_rubidiceps | diurnal |
| birds | Chlorestes_candida | Amazilia_candida | diurnal |
| birds | Chlorestes_cyanus | Hylocharis_cyanus | diurnal |
| birds | Chlorestes_eliciae | Hylocharis_eliciae | diurnal |
| birds | Chlorestes_julie | Damophila_julie | diurnal |
| birds | Chlorestes_notata | Chlorestes_notata | diurnal |
| birds | Chloris_chloris | Carduelis_chloris | diurnal |
| birds | Chloris_monguilloti | Carduelis_monguilloti | diurnal |
| birds | Chloris_sinica | Carduelis_sinica | diurnal |
| birds | Chloris_spinoides | Carduelis_spinoides | diurnal |
| birds | Chloroceryle_aenea | Chloroceryle_aenea | diurnal |
| birds | Chloroceryle_americana | Chloroceryle_americana | diurnal |
| birds | Chloroceryle_inda | Chloroceryle_inda | diurnal |
| birds | Chlorochrysa_calliparaea | Chlorochrysa_calliparaea | diurnal |
| birds | Chlorochrysa_nitidissima | Chlorochrysa_nitidissima | diurnal |
| birds | Chlorochrysa_phoenicotis | Chlorochrysa_phoenicotis | diurnal |
| birds | Chlorocichla_falkensteini | Chlorocichla_falkensteini | diurnal |
| birds | Chlorocichla_flaviventris | Chlorocichla_flaviventris | diurnal |
| birds | Chlorocichla_laetissima | Chlorocichla_laetissima | diurnal |
| birds | Chlorocichla_prigoginei | Chlorocichla_prigoginei | diurnal |
| birds | Chlorocichla_simplex | Chlorocichla_simplex | diurnal |
| birds | Chlorodrepanis_flava | Hemignathus_flavus | diurnal |
| birds | Chlorodrepanis_stejnegeri | Hemignathus_kauaiensis | diurnal |
| birds | Chlorodrepanis_virens | Hemignathus_virens | diurnal |
| birds | Chlorophanes_spiza | Chlorophanes_spiza | diurnal |

| | | | |
|---|---|---|---|
| birds | Chlorophoneus_bocagei | Telophorus_bocagei | diurnal |
| birds | Chlorophoneus_multicolor | Telophorus_multicolor | diurnal |
| birds | Chlorophoneus_nigrifrons | Telophorus_nigrifrons | diurnal |
| birds | Chlorophoneus_olivaceus | Telophorus_olivaceus | diurnal |
| birds | Chlorophoneus_sulfureopectus | Telophorus_sulfureopectus | diurnal |
| birds | Chlorophonia_callophrys | Chlorophonia_callophrys | diurnal |
| birds | Chlorophonia_cyanea | Chlorophonia_cyanea | diurnal |
| birds | Chlorophonia_flavirostris | Chlorophonia_flavirostris | diurnal |
| birds | Chlorophonia_occipitalis | Chlorophonia_occipitalis | diurnal |
| birds | Chlorophonia_pyrrhophrys | Chlorophonia_pyrrhophrys | diurnal |
| birds | Chloropicoides_rafflesii | Dinopium_rafflesii | diurnal |
| birds | Chloropipo_flavicapilla | Xenopipo_flavicapilla | diurnal |
| birds | Chloropipo_unicolor | Xenopipo_unicolor | diurnal |
| birds | Chloropsis_aurifrons | Chloropsis_aurifrons | diurnal |
| birds | Chloropsis_cyanopogon | Chloropsis_cyanopogon | diurnal |
| birds | Chloropsis_flavipennis | Chloropsis_flavipennis | diurnal |
| birds | Chloropsis_hardwickii | Chloropsis_hardwickii | diurnal |
| birds | Chloropsis_palawanensis | Chloropsis_palawanensis | diurnal |
| birds | Chloropsis_sonnerati | Chloropsis_sonnerati | diurnal |
| birds | Chlorornis_riefferii | Chlorornis_riefferii | diurnal |
| birds | Chlorospingus_canigularis | Chlorospingus_canigularis | diurnal |
| birds | Chlorospingus_flavigularis | Chlorospingus_flavigularis | diurnal |
| birds | Chlorospingus_flavopectus | Chlorospingus_ophthalmicus | diurnal |
| birds | Chlorospingus_inornatus | Chlorospingus_inornatus | diurnal |
| birds | Chlorospingus_parvirostris | Chlorospingus_parvirostris | diurnal |
| birds | Chlorospingus_pileatus | Chlorospingus_pileatus | diurnal |
| birds | Chlorospingus_semifuscus | Chlorospingus_semifuscus | diurnal |
| birds | Chlorospingus_tacarcunae | Chlorospingus_tacarcunae | diurnal |
| birds | Chlorostilbon_lucidus | Chlorostilbon_lucidus | diurnal |
| birds | Chlorostilbon_mellisugus | Chlorostilbon_mellisugus | diurnal |
| birds | Chlorostilbon_olivaresi | Chlorostilbon_olivaresi | diurnal |
| birds | Chlorostilbon_poortmani | Chlorostilbon_poortmani | diurnal |
| birds | Chlorostilbon_russatus | Chlorostilbon_russatus | diurnal |
| birds | Chlorostilbon_stenurus | Chlorostilbon_stenurus | diurnal |

| | | | |
|---|---|---|---|
| birds | Chondestes_grammacus | Chondestes_grammacus | diurnal |
| birds | Chondrohierax_uncinatus | Chondrohierax_uncinatus | diurnal |
| birds | Chordeiles_acutipennis | Chordeiles_acutipennis | cathemeral |
| birds | Chordeiles_gundlachii | Chordeiles_gundlachii | cathemeral |
| birds | Chordeiles_minor | Chordeiles_minor | nocturnal |
| birds | Chordeiles_nacunda | Podager_nacunda | nocturnal |
| birds | Chordeiles_pusillus | Chordeiles_pusillus | nocturnal |
| birds | Chordeiles_rupestris | Chordeiles_rupestris | nocturnal |
| birds | Chrysococcyx_caprius | Chrysococcyx_caprius | diurnal |
| birds | Chrysococcyx_cupreus | Chrysococcyx_cupreus | diurnal |
| birds | Chrysococcyx_flavigularis | Chrysococcyx_flavigularis | diurnal |
| birds | Chrysococcyx_klaas | Chrysococcyx_klaas | diurnal |
| birds | Chrysococcyx_maculatus | Chrysococcyx_maculatus | diurnal |
| birds | Chrysococcyx_xanthorhynchus | Chrysococcyx_xanthorhynchus | diurnal |
| birds | Chrysocolaptes_festivus | Chrysocolaptes_festivus | diurnal |
| birds | Chrysocolaptes_lucidus | Chrysocolaptes_lucidus | diurnal |
| birds | Chrysocolaptes_validus | Reinwardtipicus_validus | diurnal |
| birds | Chrysocorypha_delatrii | Tachyphonus_delatrii | diurnal |
| birds | Chrysoena_luteovirens | Ptilinopus_luteovirens | diurnal |
| birds | Chrysoena_victor | Ptilinopus_victor | diurnal |
| birds | Chrysoena_viridis | Ptilinopus_layardi | diurnal |
| birds | Chrysolampis_mosquitus | Chrysolampis_mosquitus | diurnal |
| birds | Chrysolophus_amherstiae | Chrysolophus_amherstiae | diurnal |
| birds | Chrysolophus_pictus | Chrysolophus_pictus | diurnal |
| birds | Chrysominla_strigula | Minla_strigula | diurnal |
| birds | Chrysomma_altirostre | Chrysomma_altirostre | diurnal |
| birds | Chrysomma_sinense | Chrysomma_sinense | diurnal |
| birds | Chrysomus_icterocephalus | Chrysomus_icterocephalus | diurnal |
| birds | Chrysophlegma_flavinucha | Picus_flavinucha | diurnal |
| birds | Chrysophlegma_mentale | Picus_mentalis | diurnal |
| birds | Chrysophlegma_miniaceum | Picus_mineaceus | diurnal |
| birds | Chrysothlypis_chrysomelas | Chrysothlypis_chrysomelas | diurnal |
| birds | Chrysothlypis_salmoni | Chrysothlypis_salmoni | diurnal |
| birds | Chrysuronia_brevirostris | Amazilia_brevirostris | diurnal |

| | | | |
|---|---|---|---|
| birds | Chrysuronia_coeruleogularis | Lepidopyga_coeruleogularis | diurnal |
| birds | Chrysuronia_goudoti | Lepidopyga_goudoti | diurnal |
| birds | Chrysuronia_grayi | Hylocharis_grayi | diurnal |
| birds | Chrysuronia_leucogaster | Amazilia_leucogaster | diurnal |
| birds | Chrysuronia_lilliae | Lepidopyga_lilliae | diurnal |
| birds | Chrysuronia_oenone | Chrysuronia_oenone | diurnal |
| birds | Chrysuronia_versicolor | Amazilia_versicolor | diurnal |
| birds | Chunga_burmeisteri | Chunga_burmeisteri | diurnal |
| birds | Ciccaba_albitarsis | Strix_albitarsis | nocturnal |
| birds | Ciccaba_huhula | Strix_huhula | nocturnal |
| birds | Ciccaba_nigrolineata | Strix_nigrolineata | nocturnal |
| birds | Ciccaba_virgata | Strix_virgata | nocturnal |
| birds | Cichladusa_arquata | Cichladusa_arquata | diurnal |
| birds | Cichladusa_guttata | Cichladusa_guttata | diurnal |
| birds | Cichladusa_ruficauda | Cichladusa_ruficauda | diurnal |
| birds | Cichlocolaptes_leucophrus | Cichlocolaptes_leucophrus | diurnal |
| birds | Cichlopsis_leucogenys | Cichlopsis_leucogenys | diurnal |
| birds | Cicinnurus_magnificus | Cicinnurus_magnificus | diurnal |
| birds | Cicinnurus_regius | Cicinnurus_regius | diurnal |
| birds | Cicinnurus_respublica | Cicinnurus_respublica | diurnal |
| birds | Ciconia_abdimii | Ciconia_abdimii | diurnal |
| birds | Ciconia_boyciana | Ciconia_boyciana | diurnal |
| birds | Ciconia_ciconia | Ciconia_ciconia | diurnal |
| birds | Ciconia_episcopus | Ciconia_episcopus | diurnal |
| birds | Ciconia_maguari | Ciconia_maguari | diurnal |
| birds | Ciconia_nigra | Ciconia_nigra | diurnal |
| birds | Cinclidium_frontale | Cinclidium_frontale | diurnal |
| birds | Cinclocerthia_gutturalis | Cinclocerthia_gutturalis | diurnal |
| birds | Cinclocerthia_ruficauda | Cinclocerthia_ruficauda | diurnal |
| birds | Cinclodes_antarcticus | Cinclodes_antarcticus | diurnal |
| birds | Cinclodes_aricomae | Cinclodes_aricomae | diurnal |
| birds | Cinclodes_atacamensis | Cinclodes_atacamensis | diurnal |
| birds | Cinclodes_comechingonus | Cinclodes_comechingonus | diurnal |
| birds | Cinclodes_excelsior | Cinclodes_excelsior | diurnal |

| | | | |
|---|---|---|---|
| birds | Cinclodes_nigrofumosus | Cinclodes_nigrofumosus | diurnal |
| birds | Cinclodes_oustaleti | Cinclodes_oustaleti | diurnal |
| birds | Cinclodes_pabsti | Cinclodes_pabsti | diurnal |
| birds | Cinclodes_palliatus | Cinclodes_palliatus | diurnal |
| birds | Cinclodes_patagonicus | Cinclodes_patagonicus | diurnal |
| birds | Cinclodes_taczanowskii | Cinclodes_taczanowskii | diurnal |
| birds | Cincloramphus_bivittatus | Buettikoferella_bivittata | diurnal |
| birds | Cincloramphus_cruralis | Cincloramphus_cruralis | diurnal |
| birds | Cincloramphus_grosvenori | Megalurulus_grosvenori | diurnal |
| birds | Cincloramphus_llaneae | Megalurulus_llaneae | diurnal |
| birds | Cincloramphus_mariae | Megalurulus_mariei | diurnal |
| birds | Cincloramphus_mathewsi | Cincloramphus_mathewsi | diurnal |
| birds | Cincloramphus_rubiginosus | Megalurulus_rubiginosus | diurnal |
| birds | Cincloramphus_rufus | Trichocichla_rufa | diurnal |
| birds | Cincloramphus_timoriensis | Megalurus_timoriensis | diurnal |
| birds | Cinclosoma_ajax | Cinclosoma_ajax | diurnal |
| birds | Cinclosoma_castaneothorax | Cinclosoma_castaneothorax | diurnal |
| birds | Cinclosoma_castanotum | Cinclosoma_castanotum | diurnal |
| birds | Cinclosoma_cinnamomeum | Cinclosoma_cinnamomeum | diurnal |
| birds | Cinclosoma_punctatum | Cinclosoma_punctatum | diurnal |
| birds | Cinclus_cinclus | Cinclus_cinclus | diurnal |
| birds | Cinclus_leucocephalus | Cinclus_leucocephalus | diurnal |
| birds | Cinclus_mexicanus | Cinclus_mexicanus | diurnal |
| birds | Cinclus_pallasii | Cinclus_pallasii | diurnal |
| birds | Cinclus_schulzii | Cinclus_schulzi | diurnal |
| birds | Cinnycerthia_fulva | Cinnycerthia_fulva | diurnal |
| birds | Cinnycerthia_olivascens | Cinnycerthia_olivascens | diurnal |
| birds | Cinnycerthia_peruana | Cinnycerthia_peruana | diurnal |
| birds | Cinnycerthia_unirufa | Cinnycerthia_unirufa | diurnal |
| birds | Cinnyricinclus_leucogaster | Cinnyricinclus_leucogaster | diurnal |
| birds | Cinnyris_afer | Nectarinia_afra | diurnal |
| birds | Cinnyris_asiaticus | Nectarinia_asiatica | diurnal |
| birds | Cinnyris_batesi | Nectarinia_batesi | diurnal |
| birds | Cinnyris_bifasciatus | Nectarinia_bifasciata | diurnal |

| | | | |
|---|---|---|---|
| birds | Cinnyris_bouvieri | Nectarinia_bouvieri | diurnal |
| birds | Cinnyris_chalcomelas | Nectarinia_chalcomelas | diurnal |
| birds | Cinnyris_chalybeus | Nectarinia_chalybea | diurnal |
| birds | Cinnyris_chloropygius | Nectarinia_chloropygia | diurnal |
| birds | Cinnyris_coccinigastrus | Nectarinia_coccinigaster | diurnal |
| birds | Cinnyris_comorensis | Nectarinia_comorensis | diurnal |
| birds | Cinnyris_congensis | Nectarinia_congensis | diurnal |
| birds | Cinnyris_coquerellii | Nectarinia_coquerellii | diurnal |
| birds | Cinnyris_cupreus | Nectarinia_cuprea | diurnal |
| birds | Cinnyris_dussumieri | Nectarinia_dussumieri | diurnal |
| birds | Cinnyris_erythrocercus | Nectarinia_erythrocerca | diurnal |
| birds | Cinnyris_fuscus | Nectarinia_fusca | diurnal |
| birds | Cinnyris_habessinicus | Nectarinia_habessinica | diurnal |
| birds | Cinnyris_humbloti | Nectarinia_humbloti | diurnal |
| birds | Cinnyris_johannae | Nectarinia_johannae | diurnal |
| birds | Cinnyris_jugularis | Nectarinia_jugularis | diurnal |
| birds | Cinnyris_lotenius | Nectarinia_lotenia | diurnal |
| birds | Cinnyris_loveridgei | Nectarinia_loveridgei | diurnal |
| birds | Cinnyris_ludovicensis | Nectarinia_ludovicensis | diurnal |
| birds | Cinnyris_manoensis | Nectarinia_manoensis | diurnal |
| birds | Cinnyris_mariquensis | Nectarinia_mariquensis | diurnal |
| birds | Cinnyris_mediocris | Nectarinia_mediocris | diurnal |
| birds | Cinnyris_minullus | Nectarinia_minulla | diurnal |
| birds | Cinnyris_moreaui | Nectarinia_moreaui | diurnal |
| birds | Cinnyris_nectarinioides | Nectarinia_nectarinioides | diurnal |
| birds | Cinnyris_neergaardi | Nectarinia_neergardi | diurnal |
| birds | Cinnyris_notatus | Nectarinia_notata | diurnal |
| birds | Cinnyris_osea | Nectarinia_osea | diurnal |
| birds | Cinnyris_pulchellus | Nectarinia_pulchella | diurnal |
| birds | Cinnyris_regius | Nectarinia_regia | diurnal |
| birds | Cinnyris_reichenowi | Nectarinia_preussi | diurnal |
| birds | Cinnyris_rockefelleri | Nectarinia_rockefelleri | diurnal |
| birds | Cinnyris_rufipennis | Nectarinia_rufipennis | diurnal |
| birds | Cinnyris_shelleyi | Nectarinia_shelleyi | diurnal |

| | | | |
|---|---|---|---|
| birds | Cinnyris_sovimanga | Nectarinia_sovimanga | diurnal |
| birds | Cinnyris_stuhlmanni | Nectarinia_stuhlmanni | diurnal |
| birds | Cinnyris_superbus | Nectarinia_superba | diurnal |
| birds | Cinnyris_talatala | Nectarinia_talatala | diurnal |
| birds | Cinnyris_tsavoensis | Nectarinia_tsavoensis | diurnal |
| birds | Cinnyris_ursulae | Nectarinia_ursulae | diurnal |
| birds | Cinnyris_venustus | Nectarinia_venusta | diurnal |
| birds | Circaetus_cinerascens | Circaetus_cinerascens | diurnal |
| birds | Circaetus_cinereus | Circaetus_cinereus | diurnal |
| birds | Circaetus_fasciolatus | Circaetus_fasciolatus | diurnal |
| birds | Circaetus_gallicus | Circaetus_gallicus | diurnal |
| birds | Circaetus_pectoralis | Circaetus_pectoralis | diurnal |
| birds | Circus_aeruginosus | Circus_aeruginosus | diurnal |
| birds | Circus_approximans | Circus_approximans | diurnal |
| birds | Circus_assimilis | Circus_assimilis | diurnal |
| birds | Circus_buffoni | Circus_buffoni | diurnal |
| birds | Circus_cinereus | Circus_cinereus | diurnal |
| birds | Circus_cyaneus | Circus_cyaneus | diurnal |
| birds | Circus_macrourus | Circus_macrourus | diurnal |
| birds | Circus_maillardi | Circus_maillardi | diurnal |
| birds | Circus_maurus | Circus_maurus | diurnal |
| birds | Circus_melanoleucos | Circus_melanoleucos | diurnal |
| birds | Circus_pygargus | Circus_pygargus | diurnal |
| birds | Circus_ranivorus | Circus_ranivorus | diurnal |
| birds | Circus_spilonotus | Circus_spilonotus | diurnal |
| birds | Cissa_chinensis | Cissa_chinensis | diurnal |
| birds | Cissa_hypoleuca | Cissa_hypoleuca | diurnal |
| birds | Cissomela_pectoralis | Certhionyx_pectoralis | diurnal |
| birds | Cissopis_leverianus | Cissopis_leverianus | diurnal |
| birds | Cisticola_aberdare | Cisticola_aberdare | diurnal |
| birds | Cisticola_aberrans | Cisticola_aberrans | diurnal |
| birds | Cisticola_anonymus | Cisticola_anonymus | diurnal |
| birds | Cisticola_aridulus | Cisticola_aridulus | diurnal |
| birds | Cisticola_ayresii | Cisticola_ayresii | diurnal |

| | | | |
|---|---|---|---|
| birds | Cisticola_bodessa | Cisticola_bodessa | diurnal |
| birds | Cisticola_brachypterus | Cisticola_brachypterus | diurnal |
| birds | Cisticola_brunnescens | Cisticola_brunnescens | diurnal |
| birds | Cisticola_bulliens | Cisticola_bulliens | diurnal |
| birds | Cisticola_cantans | Cisticola_cantans | diurnal |
| birds | Cisticola_carruthersi | Cisticola_carruthersi | diurnal |
| birds | Cisticola_cherina | Cisticola_cherina | diurnal |
| birds | Cisticola_chiniana | Cisticola_chiniana | diurnal |
| birds | Cisticola_chubbi | Cisticola_chubbi | diurnal |
| birds | Cisticola_cinereolus | Cisticola_cinereolus | diurnal |
| birds | Cisticola_dambo | Cisticola_dambo | diurnal |
| birds | Cisticola_erythrops | Cisticola_erythrops | diurnal |
| birds | Cisticola_exilis | Cisticola_exilis | diurnal |
| birds | Cisticola_eximius | Cisticola_eximius | diurnal |
| birds | Cisticola_fulvicapilla | Cisticola_fulvicapilla | diurnal |
| birds | Cisticola_galactotes | Cisticola_galactotes | diurnal |
| birds | Cisticola_guinea | Cisticola_guinea | diurnal |
| birds | Cisticola_haesitatus | Cisticola_haesitatus | diurnal |
| birds | Cisticola_hunteri | Cisticola_hunteri | diurnal |
| birds | Cisticola_juncidis | Cisticola_juncidis | diurnal |
| birds | Cisticola_lais | Cisticola_lais | diurnal |
| birds | Cisticola_lateralis | Cisticola_lateralis | diurnal |
| birds | Cisticola_melanurus | Cisticola_melanurus | diurnal |
| birds | Cisticola_nana | Cisticola_nanus | diurnal |
| birds | Cisticola_natalensis | Cisticola_natalensis | diurnal |
| birds | Cisticola_nigriloris | Cisticola_nigriloris | diurnal |
| birds | Cisticola_njombe | Cisticola_njombe | diurnal |
| birds | Cisticola_pipiens | Cisticola_pipiens | diurnal |
| birds | Cisticola_restrictus | Cisticola_restrictus | diurnal |
| birds | Cisticola_robustus | Cisticola_robustus | diurnal |
| birds | Cisticola_ruficeps | Cisticola_ruficeps | diurnal |
| birds | Cisticola_rufilatus | Cisticola_rufilatus | diurnal |
| birds | Cisticola_rufus | Cisticola_rufus | diurnal |
| birds | Cisticola_subruficapilla | Cisticola_subruficapilla | diurnal |

| | | | |
|---|---|---|---|
| birds | Cisticola_textrix | Cisticola_textrix | diurnal |
| birds | Cisticola_tinniens | Cisticola_tinniens | diurnal |
| birds | Cisticola_woosnami | Cisticola_woosnami | diurnal |
| birds | Cistothorus_apolinari | Cistothorus_apolinari | diurnal |
| birds | Cistothorus_meridae | Cistothorus_meridae | diurnal |
| birds | Cistothorus_palustris | Cistothorus_palustris | diurnal |
| birds | Cistothorus_platensis | Cistothorus_platensis | diurnal |
| birds | Cittura_cyanotis | Cittura_cyanotis | diurnal |
| birds | Cladorhynchus_leucocephalus | Cladorhynchus_leucocephalus | diurnal |
| birds | Clamator_coromandus | Clamator_coromandus | diurnal |
| birds | Clamator_glandarius | Clamator_glandarius | diurnal |
| birds | Clamator_jacobinus | Clamator_jacobinus | diurnal |
| birds | Clamator_levaillantii | Clamator_levaillantii | diurnal |
| birds | Clanga_clanga | Aquila_clanga | diurnal |
| birds | Clanga_pomarina | Aquila_pomarina | diurnal |
| birds | Clangula_hyemalis | Clangula_hyemalis | diurnal |
| birds | Claravis_pretiosa | Claravis_pretiosa | diurnal |
| birds | Cleptornis_marchei | Cleptornis_marchei | diurnal |
| birds | Clibanornis_dendrocolaptoides | Clibanornis_dendrocolaptoides | diurnal |
| birds | Clibanornis_erythrocephalus | Hylocryptus_erythrocephalus | diurnal |
| birds | Clibanornis_rectirostris | Hylocryptus_rectirostris | diurnal |
| birds | Clibanornis_rubiginosus | Automolus_rubiginosus | diurnal |
| birds | Climacteris_affinis | Climacteris_affinis | diurnal |
| birds | Climacteris_erythrops | Climacteris_erythrops | diurnal |
| birds | Climacteris_melanurus | Climacteris_melanurus | diurnal |
| birds | Climacteris_picumnus | Climacteris_picumnus | diurnal |
| birds | Climacteris_rufus | Climacteris_rufus | diurnal |
| birds | Clytoctantes_alixii | Clytoctantes_alixii | diurnal |
| birds | Clytoctantes_atrogularis | Clytoctantes_atrogularis | diurnal |
| birds | Clytolaema_rubricauda | Clytolaema_rubricauda | diurnal |
| birds | Clytomyias_insignis | Clytomyias_insignis | diurnal |
| birds | Clytorhynchus_hamlini | Clytorhynchus_hamlini | diurnal |
| birds | Clytorhynchus_nigrogularis | Clytorhynchus_nigrogularis | diurnal |
| birds | Clytorhynchus_pachycephaloides | Clytorhynchus_pachycephaloides | diurnal |

| | | | |
|---|---|---|---|
| birds | Clytorhynchus_vitiensis | Clytorhynchus_vitiensis | diurnal |
| birds | Clytospiza_monteiri | Clytospiza_monteiri | diurnal |
| birds | Cnemarchus_erythropygius | Cnemarchus_erythropygius | diurnal |
| birds | Cnemathraupis_aureodorsalis | Buthraupis_aureodorsalis | diurnal |
| birds | Cnemathraupis_eximia | Buthraupis_eximia | diurnal |
| birds | Cnemophilus_loriae | Cnemophilus_loriae | diurnal |
| birds | Cnemophilus_macgregorii | Cnemophilus_macgregorii | diurnal |
| birds | Cnemoscopus_rubrirostris | Cnemoscopus_rubrirostris | diurnal |
| birds | Cnemotriccus_fuscatus | Cnemotriccus_fuscatus | diurnal |
| birds | Cnipodectes_subbrunneus | Cnipodectes_subbrunneus | diurnal |
| birds | Coccopygia_melanotis | Estrilda_melanotis | diurnal |
| birds | Coccopygia_quartinia | Estrilda_quartinia | diurnal |
| birds | Coccothraustes_coccothraustes | Coccothraustes_coccothraustes | diurnal |
| birds | Coccycua_cinerea | Coccycua_cinerea | diurnal |
| birds | Coccycua_minuta | Coccycua_minuta | diurnal |
| birds | Coccycua_pumila | Coccycua_pumila | diurnal |
| birds | Coccyzus_americanus | Coccyzus_americanus | diurnal |
| birds | Coccyzus_erythropthalmus | Coccyzus_erythropthalmus | diurnal |
| birds | Coccyzus_euleri | Coccyzus_euleri | diurnal |
| birds | Coccyzus_ferrugineus | Coccyzus_ferrugineus | diurnal |
| birds | Coccyzus_lansbergi | Coccyzus_lansbergi | diurnal |
| birds | Coccyzus_longirostris | Coccyzus_longirostris | diurnal |
| birds | Coccyzus_melacoryphus | Coccyzus_melacoryphus | diurnal |
| birds | Coccyzus_merlini | Coccyzus_merlini | diurnal |
| birds | Coccyzus_minor | Coccyzus_minor | diurnal |
| birds | Coccyzus_pluvialis | Coccyzus_pluvialis | diurnal |
| birds | Coccyzus_rufigularis | Coccyzus_rufigularis | diurnal |
| birds | Coccyzus_vetula | Coccyzus_vetula | diurnal |
| birds | Coccyzus_vieilloti | Coccyzus_vieilloti | diurnal |
| birds | Cochlearius_cochlearius | Cochlearius_cochlearius | cathemeral |
| birds | Cochoa_azurea | Cochoa_azurea | diurnal |
| birds | Cochoa_beccarii | Cochoa_beccarii | diurnal |
| birds | Cochoa_purpurea | Cochoa_purpurea | diurnal |
| birds | Cochoa_viridis | Cochoa_viridis | diurnal |

| | | | |
|---|---|---|---|
| birds | Coeligena_bonapartei | Coeligena_bonapartei | diurnal |
| birds | Coeligena_coeligena | Coeligena_coeligena | diurnal |
| birds | Coeligena_helianthea | Coeligena_helianthea | diurnal |
| birds | Coeligena_iris | Coeligena_iris | diurnal |
| birds | Coeligena_lutetiae | Coeligena_lutetiae | diurnal |
| birds | Coeligena_phalerata | Coeligena_phalerata | diurnal |
| birds | Coeligena_prunellei | Coeligena_prunellei | diurnal |
| birds | Coeligena_torquata | Coeligena_torquata | diurnal |
| birds | Coeligena_violifer | Coeligena_violifer | diurnal |
| birds | Coeligena_wilsoni | Coeligena_wilsoni | diurnal |
| birds | Coenocorypha_aucklandica | Coenocorypha_aucklandica | cathemeral |
| birds | Coenocorypha_pusilla | Coenocorypha_pusilla | diurnal |
| birds | Coereba_flaveola | Coereba_flaveola | diurnal |
| birds | Colaptes_atricollis | Colaptes_atricollis | diurnal |
| birds | Colaptes_auratus | Colaptes_auratus | diurnal |
| birds | Colaptes_auricularis | Colaptes_auricularis | diurnal |
| birds | Colaptes_campestris | Colaptes_campestris | diurnal |
| birds | Colaptes_chrysoides | Colaptes_chrysoides | diurnal |
| birds | Colaptes_fernandinae | Colaptes_fernandinae | diurnal |
| birds | Colaptes_melanochloros | Colaptes_melanochloros | diurnal |
| birds | Colaptes_pitius | Colaptes_pitius | diurnal |
| birds | Colaptes_punctigula | Colaptes_punctigula | diurnal |
| birds | Colaptes_rivolii | Colaptes_rivolii | diurnal |
| birds | Colaptes_rubiginosus | Colaptes_rubiginosus | diurnal |
| birds | Colaptes_rupicola | Colaptes_rupicola | diurnal |
| birds | Colibri_coruscans | Colibri_coruscans | diurnal |
| birds | Colibri_delphinae | Colibri_delphinae | diurnal |
| birds | Colibri_serrirostris | Colibri_serrirostris | diurnal |
| birds | Colibri_thalassinus | Colibri_thalassinus | diurnal |
| birds | Colinus_cristatus | Colinus_cristatus | diurnal |
| birds | Colinus_leucopogon | Colinus_leucopogon | diurnal |
| birds | Colinus_nigrogularis | Colinus_nigrogularis | diurnal |
| birds | Colinus_virginianus | Colinus_virginianus | diurnal |
| birds | Colius_castanotus | Colius_castanotus | diurnal |

| | | | |
|---|---|---|---|
| birds | Colius_colius | Colius_colius | diurnal |
| birds | Colius_leucocephalus | Colius_leucocephalus | diurnal |
| birds | Colius_striatus | Colius_striatus | diurnal |
| birds | Collocalia_esculenta | Collocalia_esculenta | diurnal |
| birds | Collocalia_linchi | Collocalia_linchi | diurnal |
| birds | Collocalia_troglodytes | Collocalia_troglodytes | diurnal |
| birds | Colluricincla_boweri | Colluricincla_boweri | diurnal |
| birds | Colluricincla_harmonica | Colluricincla_harmonica | diurnal |
| birds | Colluricincla_megarhyncha | Colluricincla_megarhyncha | diurnal |
| birds | Colluricincla_tenebrosa | Colluricincla_umbrina | diurnal |
| birds | Colluricincla_woodwardi | Colluricincla_woodwardi | diurnal |
| birds | Colonia_colonus | Colonia_colonus | diurnal |
| birds | Colorhamphus_parvirostris | Colorhamphus_parvirostris | diurnal |
| birds | Columba_albinucha | Columba_albinucha | diurnal |
| birds | Columba_albitorques | Columba_albitorques | diurnal |
| birds | Columba_arquatrix | Columba_arquatrix | diurnal |
| birds | Columba_bollii | Columba_bollii | diurnal |
| birds | Columba_delegorguei | Columba_delegorguei | diurnal |
| birds | Columba_elphinstonii | Columba_elphinstonii | diurnal |
| birds | Columba_eversmanni | Columba_eversmanni | diurnal |
| birds | Columba_guinea | Columba_guinea | diurnal |
| birds | Columba_hodgsonii | Columba_hodgsonii | diurnal |
| birds | Columba_iriditorques | Columba_iriditorques | diurnal |
| birds | Columba_janthina | Columba_janthina | diurnal |
| birds | Columba_leucomela | Columba_leucomela | diurnal |
| birds | Columba_leuconota | Columba_leuconota | diurnal |
| birds | Columba_livia | Columba_livia | diurnal |
| birds | Columba_malherbii | Columba_malherbii | diurnal |
| birds | Columba_oenas | Columba_oenas | diurnal |
| birds | Columba_oliviae | Columba_oliviae | diurnal |
| birds | Columba_pallidiceps | Columba_pallidiceps | diurnal |
| birds | Columba_palumboides | Columba_palumboides | diurnal |
| birds | Columba_pollenii | Columba_pollenii | diurnal |
| birds | Columba_pulchricollis | Columba_pulchricollis | diurnal |

| | | | |
|---|---|---|---|
| birds | Columba_punicea | Columba_punicea | diurnal |
| birds | Columba_rupestris | Columba_rupestris | diurnal |
| birds | Columba_sjostedti | Columba_sjostedti | diurnal |
| birds | Columba_thomensis | Columba_thomensis | diurnal |
| birds | Columba_torringtoniae | Columba_torringtoniae | diurnal |
| birds | Columba_trocaz | Columba_trocaz | diurnal |
| birds | Columba_unicincta | Columba_unicincta | diurnal |
| birds | Columba_vitiensis | Columba_vitiensis | diurnal |
| birds | Columbina_buckleyi | Columbina_buckleyi | diurnal |
| birds | Columbina_cruziana | Columbina_cruziana | diurnal |
| birds | Columbina_cyanopis | Columbina_cyanopis | diurnal |
| birds | Columbina_inca | Columbina_inca | diurnal |
| birds | Columbina_minuta | Columbina_minuta | diurnal |
| birds | Columbina_passerina | Columbina_passerina | diurnal |
| birds | Columbina_picui | Columbina_picui | diurnal |
| birds | Columbina_squammata | Columbina_squammata | diurnal |
| birds | Columbina_talpacoti | Columbina_talpacoti | diurnal |
| birds | Compsothraupis_loricata | Compsothraupis_loricata | diurnal |
| birds | Conioptilon_mcilhennyi | Conioptilon_mcilhennyi | diurnal |
| birds | Conirostrum_albifrons | Conirostrum_albifrons | diurnal |
| birds | Conirostrum_bicolor | Conirostrum_bicolor | diurnal |
| birds | Conirostrum_binghami | Oreomanes_fraseri | diurnal |
| birds | Conirostrum_cinereum | Conirostrum_cinereum | diurnal |
| birds | Conirostrum_ferrugineiventre | Conirostrum_ferrugineiventre | diurnal |
| birds | Conirostrum_leucogenys | Conirostrum_leucogenys | diurnal |
| birds | Conirostrum_margaritae | Conirostrum_margaritae | diurnal |
| birds | Conirostrum_rufum | Conirostrum_rufum | diurnal |
| birds | Conirostrum_sitticolor | Conirostrum_sitticolor | diurnal |
| birds | Conirostrum_speciosum | Conirostrum_speciosum | diurnal |
| birds | Conirostrum_tamarugense | Conirostrum_tamarugense | diurnal |
| birds | Conopias_albovittatus | Conopias_albovittatus | diurnal |
| birds | Conopias_cinchoneti | Conopias_cinchoneti | diurnal |
| birds | Conopias_parvus | Conopias_parvus | diurnal |
| birds | Conopophaga_ardesiaca | Conopophaga_ardesiaca | diurnal |

| | | | |
|---|---|---|---|
| birds | Conopophaga_aurita | Conopophaga_aurita | diurnal |
| birds | Conopophaga_castaneiceps | Conopophaga_castaneiceps | diurnal |
| birds | Conopophaga_cearae | Conopophaga_cearae | diurnal |
| birds | Conopophaga_lineata | Conopophaga_lineata | diurnal |
| birds | Conopophaga_melanogaster | Conopophaga_melanogaster | diurnal |
| birds | Conopophaga_melanops | Conopophaga_melanops | diurnal |
| birds | Conopophaga_peruviana | Conopophaga_peruviana | diurnal |
| birds | Conopophaga_roberti | Conopophaga_roberti | diurnal |
| birds | Conopophila_albogularis | Conopophila_albogularis | diurnal |
| birds | Conopophila_rufogularis | Conopophila_rufogularis | diurnal |
| birds | Conopophila_whitei | Conopophila_whitei | diurnal |
| birds | Conothraupis_mesoleuca | Conothraupis_mesoleuca | diurnal |
| birds | Conothraupis_speculigera | Conothraupis_speculigera | diurnal |
| birds | Contopus_albogularis | Contopus_albogularis | diurnal |
| birds | Contopus_caribaeus | Contopus_caribaeus | diurnal |
| birds | Contopus_cinereus | Contopus_cinereus | diurnal |
| birds | Contopus_cooperi | Contopus_cooperi | diurnal |
| birds | Contopus_fumigatus | Contopus_fumigatus | diurnal |
| birds | Contopus_hispaniolensis | Contopus_hispaniolensis | diurnal |
| birds | Contopus_latirostris | Contopus_latirostris | diurnal |
| birds | Contopus_lugubris | Contopus_lugubris | diurnal |
| birds | Contopus_nigrescens | Contopus_nigrescens | diurnal |
| birds | Contopus_ochraceus | Contopus_ochraceus | diurnal |
| birds | Contopus_pallidus | Contopus_pallidus | diurnal |
| birds | Contopus_pertinax | Contopus_pertinax | diurnal |
| birds | Contopus_punensis | Contopus_punensis | diurnal |
| birds | Contopus_sordidulus | Contopus_sordidulus | diurnal |
| birds | Contopus_virens | Contopus_virens | diurnal |
| birds | Copsychus_albospecularis | Copsychus_albospecularis | diurnal |
| birds | Copsychus_fulicatus | Saxicoloides_fulicatus | diurnal |
| birds | Copsychus_pyrropygus | Trichixos_pyrropygus | diurnal |
| birds | Copsychus_saularis | Copsychus_saularis | diurnal |
| birds | Copsychus_sechellarum | Copsychus_sechellarum | diurnal |
| birds | Coracias_abyssinicus | Coracias_abyssinicus | diurnal |

| | | | |
|---|---|---|---|
| birds | Coracias_benghalensis | Coracias_benghalensis | cathemeral |
| birds | Coracias_caudatus | Coracias_caudatus | diurnal |
| birds | Coracias_cyanogaster | Coracias_cyanogaster | diurnal |
| birds | Coracias_garrulus | Coracias_garrulus | diurnal |
| birds | Coracias_naevius | Coracias_naevia | diurnal |
| birds | Coracias_spatulatus | Coracias_spatulatus | diurnal |
| birds | Coracias_temminckii | Coracias_temminckii | diurnal |
| birds | Coracina_atriceps | Coracina_atriceps | diurnal |
| birds | Coracina_boyeri | Coracina_boyeri | diurnal |
| birds | Coracina_caeruleogrisea | Coracina_caeruleogrisea | diurnal |
| birds | Coracina_caledonica | Coracina_caledonica | diurnal |
| birds | Coracina_javensis | Coracina_javensis | diurnal |
| birds | Coracina_lineata | Coracina_lineata | diurnal |
| birds | Coracina_longicauda | Coracina_longicauda | diurnal |
| birds | Coracina_macei | Coracina_macei | diurnal |
| birds | Coracina_maxima | Coracina_maxima | diurnal |
| birds | Coracina_novaehollandiae | Coracina_novaehollandiae | diurnal |
| birds | Coracina_papuensis | Coracina_papuensis | diurnal |
| birds | Coracina_parvula | Coracina_parvula | diurnal |
| birds | Coracina_striata | Coracina_striata | diurnal |
| birds | Coracopsis_nigra | Coracopsis_nigra | diurnal |
| birds | Coracopsis_vasa | Coracopsis_vasa | diurnal |
| birds | Coracornis_raveni | Coracornis_raveni | diurnal |
| birds | Coragyps_atratus | Coragyps_atratus | diurnal |
| birds | Corapipo_gutturalis | Corapipo_gutturalis | diurnal |
| birds | Corapipo_leucorrhoa | Corapipo_leucorrhoa | diurnal |
| birds | Corcorax_melanorhamphos | Corcorax_melanorhamphos | diurnal |
| birds | Cormobates_leucophaea | Cormobates_leucophaea | diurnal |
| birds | Cormobates_placens | Cormobates_placens | diurnal |
| birds | Corthylio_calendula | Regulus_calendula | diurnal |
| birds | Corvus_albicollis | Corvus_albicollis | diurnal |
| birds | Corvus_albus | Corvus_albus | diurnal |
| birds | Corvus_bennetti | Corvus_bennetti | diurnal |
| birds | Corvus_brachyrhynchos | Corvus_brachyrhynchos | diurnal |

| | | | |
|---|---|---|---|
| birds | Corvus_capensis | Corvus_capensis | diurnal |
| birds | Corvus_caurinus | Corvus_caurinus | diurnal |
| birds | Corvus_corax | Corvus_corax | diurnal |
| birds | Corvus_corone | Corvus_corone | diurnal |
| birds | Corvus_coronoides | Corvus_coronoides | diurnal |
| birds | Corvus_crassirostris | Corvus_crassirostris | diurnal |
| birds | Corvus_cryptoleucus | Corvus_cryptoleucus | diurnal |
| birds | Corvus_dauuricus | Corvus_dauuricus | diurnal |
| birds | Corvus_enca | Corvus_enca | diurnal |
| birds | Corvus_florensis | Corvus_florensis | diurnal |
| birds | Corvus_frugilegus | Corvus_frugilegus | diurnal |
| birds | Corvus_hawaiiensis | Corvus_hawaiiensis | diurnal |
| birds | Corvus_imparatus | Corvus_imparatus | diurnal |
| birds | Corvus_jamaicensis | Corvus_jamaicensis | diurnal |
| birds | Corvus_kubaryi | Corvus_kubaryi | diurnal |
| birds | Corvus_macrorhynchos | Corvus_macrorhynchos | diurnal |
| birds | Corvus_mellori | Corvus_mellori | diurnal |
| birds | Corvus_monedula | Corvus_monedula | diurnal |
| birds | Corvus_moneduloides | Corvus_moneduloides | diurnal |
| birds | Corvus_nasicus | Corvus_nasicus | diurnal |
| birds | Corvus_orru | Corvus_orru | diurnal |
| birds | Corvus_ossifragus | Corvus_ossifragus | diurnal |
| birds | Corvus_palmarum | Corvus_palmarum | diurnal |
| birds | Corvus_pectoralis | Corvus_torquatus | diurnal |
| birds | Corvus_rhipidurus | Corvus_rhipidurus | diurnal |
| birds | Corvus_ruficollis | Corvus_ruficollis | diurnal |
| birds | Corvus_sinaloae | Corvus_sinaloae | diurnal |
| birds | Corvus_splendens | Corvus_splendens | diurnal |
| birds | Corvus_tasmanicus | Corvus_tasmanicus | diurnal |
| birds | Corvus_tristis | Corvus_tristis | diurnal |
| birds | Corvus_typicus | Corvus_typicus | diurnal |
| birds | Corvus_unicolor | Corvus_unicolor | diurnal |
| birds | Corvus_validus | Corvus_validus | diurnal |
| birds | Corvus_woodfordi | Corvus_woodfordi | diurnal |

| | | | |
|---|---|---|---|
| birds | Corydon_sumatranus | Corydon_sumatranus | diurnal |
| birds | Corydospiza_alaudina | Phrygilus_alaudinus | diurnal |
| birds | Corydospiza_carbonaria | Phrygilus_carbonarius | diurnal |
| birds | Coryphaspiza_melanotis | Coryphaspiza_melanotis | diurnal |
| birds | Coryphistera_alaudina | Coryphistera_alaudina | diurnal |
| birds | Coryphospingus_cucullatus | Coryphospingus_cucullatus | diurnal |
| birds | Coryphospingus_pileatus | Coryphospingus_pileatus | diurnal |
| birds | Corythaeola_cristata | Corythaeola_cristata | diurnal |
| birds | Corythaixoides_concolor | Corythaixoides_concolor | diurnal |
| birds | Corythaixoides_personatus | Corythaixoides_personatus | diurnal |
| birds | Corythopis_delalandi | Corythopis_delalandi | diurnal |
| birds | Corythopis_torquatus | Corythopis_torquatus | diurnal |
| birds | Corythornis_cristatus | Alcedo_cristata | diurnal |
| birds | Corythornis_leucogaster | Alcedo_leucogaster | diurnal |
| birds | Corythornis_madagascariensis | Ceyx_madagascariensis | diurnal |
| birds | Corythornis_vintsioides | Alcedo_vintsioides | diurnal |
| birds | Coscoroba_coscoroba | Coscoroba_coscoroba | diurnal |
| birds | Cossypha_albicapillus | Cossypha_albicapilla | diurnal |
| birds | Cossypha_cyanocampter | Cossypha_cyanocampter | diurnal |
| birds | Cossypha_heinrichi | Cossypha_heinrichi | diurnal |
| birds | Cossypha_heuglini | Cossypha_heuglini | diurnal |
| birds | Cossypha_natalensis | Cossypha_natalensis | diurnal |
| birds | Cossypha_niveicapilla | Cossypha_niveicapilla | diurnal |
| birds | Cossypha_semirufa | Cossypha_semirufa | diurnal |
| birds | Cossyphicula_isabellae | Cossypha_isabellae | diurnal |
| birds | Cossyphicula_roberti | Cossyphicula_roberti | diurnal |
| birds | Cotinga_amabilis | Cotinga_amabilis | diurnal |
| birds | Cotinga_cayana | Cotinga_cayana | diurnal |
| birds | Cotinga_cotinga | Cotinga_cotinga | diurnal |
| birds | Cotinga_maculata | Cotinga_maculata | diurnal |
| birds | Cotinga_maynana | Cotinga_maynana | diurnal |
| birds | Cotinga_ridgwayi | Cotinga_ridgwayi | diurnal |
| birds | Coturnicops_notatus | Coturnicops_notatus | diurnal |
| birds | Coturnix_coromandelica | Coturnix_coromandelica | diurnal |

| | | | |
|---|---|---|---|
| birds | Coturnix_coturnix | Coturnix_coturnix | diurnal |
| birds | Coturnix_delegorguei | Coturnix_delegorguei | diurnal |
| birds | Coturnix_japonica | Coturnix_japonica | diurnal |
| birds | Coturnix_pectoralis | Coturnix_pectoralis | diurnal |
| birds | Coua_caerulea | Coua_caerulea | diurnal |
| birds | Coua_coquereli | Coua_coquereli | diurnal |
| birds | Coua_cristata | Coua_cristata | diurnal |
| birds | Coua_cursor | Coua_cursor | diurnal |
| birds | Coua_gigas | Coua_gigas | diurnal |
| birds | Coua_reynaudii | Coua_reynaudii | diurnal |
| birds | Coua_ruficeps | Coua_ruficeps | diurnal |
| birds | Coua_serriana | Coua_serriana | diurnal |
| birds | Coua_verreauxi | Coua_verreauxi | diurnal |
| birds | Cracticus_cassicus | Cracticus_cassicus | diurnal |
| birds | Cracticus_mentalis | Cracticus_mentalis | diurnal |
| birds | Cracticus_nigrogularis | Cracticus_nigrogularis | diurnal |
| birds | Cracticus_torquatus | Cracticus_torquatus | diurnal |
| birds | Cranioleuca_albicapilla | Cranioleuca_albicapilla | diurnal |
| birds | Cranioleuca_albiceps | Cranioleuca_albiceps | diurnal |
| birds | Cranioleuca_antisiensis | Cranioleuca_antisiensis | diurnal |
| birds | Cranioleuca_berlepschi | Thripophaga_berlepschi | diurnal |
| birds | Cranioleuca_curtata | Cranioleuca_curtata | diurnal |
| birds | Cranioleuca_demissa | Cranioleuca_demissa | diurnal |
| birds | Cranioleuca_erythrops | Cranioleuca_erythrops | diurnal |
| birds | Cranioleuca_hellmayri | Cranioleuca_hellmayri | diurnal |
| birds | Cranioleuca_marcapatae | Cranioleuca_marcapatae | diurnal |
| birds | Cranioleuca_muelleri | Cranioleuca_muelleri | diurnal |
| birds | Cranioleuca_obsoleta | Cranioleuca_obsoleta | diurnal |
| birds | Cranioleuca_pallida | Cranioleuca_pallida | diurnal |
| birds | Cranioleuca_pyrrhophia | Cranioleuca_pyrrhophia | diurnal |
| birds | Cranioleuca_semicinerea | Cranioleuca_semicinerea | diurnal |
| birds | Cranioleuca_subcristata | Cranioleuca_subcristata | diurnal |
| birds | Cranioleuca_vulpecula | Cranioleuca_vulpecula | diurnal |
| birds | Cranioleuca_vulpina | Cranioleuca_vulpina | diurnal |

| | | | |
|---|---|---|---|
| birds | Crax_alector | Crax_alector | diurnal |
| birds | Crax_blumenbachii | Crax_blumenbachii | diurnal |
| birds | Crax_daubentoni | Crax_daubentoni | diurnal |
| birds | Crax_globulosa | Crax_globulosa | diurnal |
| birds | Crax_rubra | Crax_rubra | diurnal |
| birds | Creagrus_furcatus | Creagrus_furcatus | diurnal |
| birds | Creatophora_cinerea | Creatophora_cinerea | diurnal |
| birds | Creurgops_dentatus | Creurgops_dentatus | diurnal |
| birds | Creurgops_verticalis | Creurgops_verticalis | diurnal |
| birds | Crinifer_piscator | Crinifer_piscator | diurnal |
| birds | Crinifer_zonurus | Crinifer_zonurus | diurnal |
| birds | Criniferoides_leucogaster | Corythaixoides_leucogaster | diurnal |
| birds | Criniger_barbatus | Criniger_barbatus | diurnal |
| birds | Criniger_calurus | Criniger_calurus | diurnal |
| birds | Criniger_chloronotus | Criniger_chloronotus | diurnal |
| birds | Criniger_ndussumensis | Criniger_ndussumensis | diurnal |
| birds | Criniger_olivaceus | Criniger_olivaceus | diurnal |
| birds | Crithagra_albogularis | Serinus_albogularis | diurnal |
| birds | Crithagra_ankoberensis | Serinus_ankoberensis | diurnal |
| birds | Crithagra_atrogularis | Serinus_atrogularis | diurnal |
| birds | Crithagra_burtoni | Serinus_burtoni | diurnal |
| birds | Crithagra_citrinelloides | Serinus_citrinelloides | diurnal |
| birds | Crithagra_citrinipectus | Serinus_citrinipectus | diurnal |
| birds | Crithagra_concolor | Neospiza_concolor | diurnal |
| birds | Crithagra_donaldsoni | Serinus_donaldsoni | diurnal |
| birds | Crithagra_dorsostriata | Serinus_dorsostriatus | diurnal |
| birds | Crithagra_flaviventris | Serinus_flaviventris | diurnal |
| birds | Crithagra_frontalis | Serinus_frontalis | diurnal |
| birds | Crithagra_gularis | Serinus_gularis | diurnal |
| birds | Crithagra_hyposticta | Serinus_hypostictus | diurnal |
| birds | Crithagra_koliensis | Serinus_koliensis | diurnal |
| birds | Crithagra_leucoptera | Serinus_leucopterus | diurnal |
| birds | Crithagra_leucopygia | Serinus_leucopygius | diurnal |
| birds | Crithagra_mennelli | Serinus_mennelli | diurnal |

| | | | |
|---|---|---|---|
| birds | Crithagra_mozambica | Serinus_mozambicus | diurnal |
| birds | Crithagra_reichardi | Serinus_reichardi | diurnal |
| birds | Crithagra_rothschildi | Serinus_rothschildi | diurnal |
| birds | Crithagra_rufobrunnea | Serinus_rufobrunneus | diurnal |
| birds | Crithagra_scotops | Serinus_scotops | diurnal |
| birds | Crithagra_striolata | Serinus_striolatus | diurnal |
| birds | Crithagra_sulphurata | Serinus_sulphuratus | diurnal |
| birds | Crithagra_symonsi | Serinus_symonsi | diurnal |
| birds | Crithagra_totta | Serinus_totta | diurnal |
| birds | Crithagra_tristriata | Serinus_tristriatus | diurnal |
| birds | Crithagra_xantholaema | Serinus_xantholaemus | diurnal |
| birds | Crithagra_xanthopygia | Serinus_xanthopygius | diurnal |
| birds | Crossleyia_tenebrosa | Bernieria_tenebrosa | diurnal |
| birds | Crossleyia_xanthophrys | Crossleyia_xanthophrys | diurnal |
| birds | Crossoptilon_auritum | Crossoptilon_auritum | diurnal |
| birds | Crossoptilon_crossoptilon | Crossoptilon_crossoptilon | diurnal |
| birds | Crossoptilon_mantchuricum | Crossoptilon_mantchuricum | diurnal |
| birds | Crotophaga_ani | Crotophaga_ani | diurnal |
| birds | Crotophaga_major | Crotophaga_major | diurnal |
| birds | Crotophaga_sulcirostris | Crotophaga_sulcirostris | diurnal |
| birds | Crypsirina_cucullata | Crypsirina_cucullata | diurnal |
| birds | Crypsirina_temia | Crypsirina_temia | diurnal |
| birds | Cryptillas_victorini | Bradypterus_victorini | diurnal |
| birds | Cryptoleucopteryx_plumbea | Leucopternis_plumbeus | diurnal |
| birds | Cryptolybia_olivacea | Stactolaema_olivacea | diurnal |
| birds | Cryptomicroeca_flaviventris | Eopsaltria_flaviventris | diurnal |
| birds | Cryptopezus_nattereri | Hylopezus_nattereri | diurnal |
| birds | Cryptophaps_poecilorrhoa | Cryptophaps_poecilorrhoa | diurnal |
| birds | Cryptopipo_holochlora | Xenopipo_holochlora | diurnal |
| birds | Cryptospiza_jacksoni | Cryptospiza_jacksoni | diurnal |
| birds | Cryptospiza_reichenovii | Cryptospiza_reichenovii | diurnal |
| birds | Cryptospiza_salvadorii | Cryptospiza_salvadorii | diurnal |
| birds | Cryptospiza_shelleyi | Cryptospiza_shelleyi | diurnal |
| birds | Cryptosylvicola_randrianasoloi | Cryptosylvicola_randrianasoloi | diurnal |

| | | | |
|---|---|---|---|
| birds | Crypturellus_bartletti | Crypturellus_bartletti | diurnal |
| birds | Crypturellus_boucardi | Crypturellus_boucardi | diurnal |
| birds | Crypturellus_brevirostris | Crypturellus_brevirostris | diurnal |
| birds | Crypturellus_casiquiare | Crypturellus_casiquiare | diurnal |
| birds | Crypturellus_cinereus | Crypturellus_cinereus | diurnal |
| birds | Crypturellus_cinnamomeus | Crypturellus_cinnamomeus | diurnal |
| birds | Crypturellus_noctivagus | Crypturellus_noctivagus | diurnal |
| birds | Crypturellus_obsoletus | Crypturellus_obsoletus | diurnal |
| birds | Crypturellus_parvirostris | Crypturellus_parvirostris | diurnal |
| birds | Crypturellus_soui | Crypturellus_soui | diurnal |
| birds | Crypturellus_tataupa | Crypturellus_tataupa | diurnal |
| birds | Crypturellus_undulatus | Crypturellus_undulatus | diurnal |
| birds | Crypturellus_variegatus | Crypturellus_variegatus | diurnal |
| birds | Cuculus_canorus | Cuculus_canorus | diurnal |
| birds | Cuculus_clamosus | Cuculus_clamosus | diurnal |
| birds | Cuculus_gularis | Cuculus_gularis | diurnal |
| birds | Cuculus_poliocephalus | Cuculus_poliocephalus | diurnal |
| birds | Cuculus_rochii | Cuculus_rochii | diurnal |
| birds | Cuculus_saturatus | Cuculus_saturatus | diurnal |
| birds | Cuculus_solitarius | Cuculus_solitarius | diurnal |
| birds | Culicicapa_ceylonensis | Culicicapa_ceylonensis | diurnal |
| birds | Culicicapa_helianthea | Culicicapa_helianthea | diurnal |
| birds | Culicivora_caudacuta | Culicivora_caudacuta | diurnal |
| birds | Curaeus_curaeus | Curaeus_curaeus | diurnal |
| birds | Curruca_boehmi | Sylvia_boehmi | diurnal |
| birds | Curruca_buryi | Sylvia_buryi | diurnal |
| birds | Curruca_communis | Sylvia_communis | diurnal |
| birds | Curruca_conspicillata | Sylvia_conspicillata | diurnal |
| birds | Curruca_deserticola | Sylvia_deserticola | diurnal |
| birds | Curruca_layardi | Sylvia_layardi | diurnal |
| birds | Curruca_leucomelaena | Sylvia_leucomelaena | diurnal |
| birds | Curruca_lugens | Sylvia_lugens | diurnal |
| birds | Curruca_melanocephala | Sylvia_melanocephala | diurnal |
| birds | Curruca_melanothorax | Sylvia_melanothorax | diurnal |

| | | | |
|---|---|---|---|
| birds | Curruca_mystacea | Sylvia_mystacea | diurnal |
| birds | Curruca_nisoria | Sylvia_nisoria | diurnal |
| birds | Curruca_ruppeli | Sylvia_rueppelli | diurnal |
| birds | Curruca_subcoerulea | Sylvia_subcaerulea | diurnal |
| birds | Curruca_undata | Sylvia_undata | diurnal |
| birds | Cursorius_cursor | Cursorius_cursor | diurnal |
| birds | Cursorius_rufus | Cursorius_rufus | diurnal |
| birds | Cutia_nipalensis | Cutia_nipalensis | diurnal |
| birds | Cyanerpes_caeruleus | Cyanerpes_caeruleus | diurnal |
| birds | Cyanerpes_cyaneus | Cyanerpes_cyaneus | diurnal |
| birds | Cyanerpes_lucidus | Cyanerpes_lucidus | diurnal |
| birds | Cyanerpes_nitidus | Cyanerpes_nitidus | diurnal |
| birds | Cyanicterus_cyanicterus | Cyanicterus_cyanicterus | diurnal |
| birds | Cyanistes_caeruleus | Parus_caeruleus | diurnal |
| birds | Cyanistes_cyanus | Parus_cyanus | diurnal |
| birds | Cyanistes_teneriffae | Parus_teneriffae | diurnal |
| birds | Cyanochen_cyanoptera | Cyanochen_cyanoptera | diurnal |
| birds | Cyanocitta_cristata | Cyanocitta_cristata | diurnal |
| birds | Cyanocitta_stelleri | Cyanocitta_stelleri | diurnal |
| birds | Cyanocompsa_parellina | Cyanocompsa_parellina | diurnal |
| birds | Cyanocorax_affinis | Cyanocorax_affinis | diurnal |
| birds | Cyanocorax_beecheii | Cyanocorax_beecheii | diurnal |
| birds | Cyanocorax_cayanus | Cyanocorax_cayanus | diurnal |
| birds | Cyanocorax_chrysops | Cyanocorax_chrysops | diurnal |
| birds | Cyanocorax_coeruleus | Cyanocorax_caeruleus | diurnal |
| birds | Cyanocorax_colliei | Calocitta_colliei | diurnal |
| birds | Cyanocorax_cristatellus | Cyanocorax_cristatellus | diurnal |
| birds | Cyanocorax_cyanomelas | Cyanocorax_cyanomelas | diurnal |
| birds | Cyanocorax_cyanopogon | Cyanocorax_cyanopogon | diurnal |
| birds | Cyanocorax_dickeyi | Cyanocorax_dickeyi | diurnal |
| birds | Cyanocorax_formosus | Calocitta_formosa | diurnal |
| birds | Cyanocorax_heilprini | Cyanocorax_heilprini | diurnal |
| birds | Cyanocorax_melanocyaneus | Cyanocorax_melanocyaneus | diurnal |
| birds | Cyanocorax_morio | Cyanocorax_morio | diurnal |

| | | | |
|---|---|---|---|
| birds | Cyanocorax_mystacalis | Cyanocorax_mystacalis | diurnal |
| birds | Cyanocorax_sanblasianus | Cyanocorax_sanblasianus | diurnal |
| birds | Cyanocorax_violaceus | Cyanocorax_violaceus | diurnal |
| birds | Cyanocorax_yncas | Cyanocorax_yncas | diurnal |
| birds | Cyanocorax_yucatanicus | Cyanocorax_yucatanicus | diurnal |
| birds | Cyanoderma_chrysaeum | Stachyris_chrysaea | diurnal |
| birds | Cyanoderma_erythropterum | Stachyris_erythroptera | diurnal |
| birds | Cyanoderma_melanothorax | Stachyris_melanothorax | diurnal |
| birds | Cyanoderma_pyrrhops | Stachyris_pyrrhops | diurnal |
| birds | Cyanoderma_ruficeps | Stachyris_ruficeps | diurnal |
| birds | Cyanoderma_rufifrons | Stachyris_rufifrons | diurnal |
| birds | Cyanograucalus_azureus | Coracina_azurea | diurnal |
| birds | Cyanolanius_madagascarinus | Cyanolanius_madagascarinus | diurnal |
| birds | Cyanoliseus_patagonus | Cyanoliseus_patagonus | diurnal |
| birds | Cyanoloxia_brissonii | Cyanocompsa_brissonii | diurnal |
| birds | Cyanoloxia_cyanoides | Cyanocompsa_cyanoides | diurnal |
| birds | Cyanoloxia_glaucocaerulea | Cyanoloxia_glaucocaerulea | diurnal |
| birds | Cyanolyca_argentigula | Cyanolyca_argentigula | diurnal |
| birds | Cyanolyca_armillata | Cyanolyca_armillata | diurnal |
| birds | Cyanolyca_cucullata | Cyanolyca_cucullata | diurnal |
| birds | Cyanolyca_mirabilis | Cyanolyca_mirabilis | diurnal |
| birds | Cyanolyca_nanus | Cyanolyca_nana | diurnal |
| birds | Cyanolyca_pumilo | Cyanolyca_pumilo | diurnal |
| birds | Cyanolyca_turcosa | Cyanolyca_turcosa | diurnal |
| birds | Cyanolyca_viridicyanus | Cyanolyca_viridicyanus | diurnal |
| birds | Cyanomitra_alinae | Nectarinia_alinae | diurnal |
| birds | Cyanomitra_bannermani | Nectarinia_bannermani | diurnal |
| birds | Cyanomitra_cyanolaema | Nectarinia_cyanolaema | diurnal |
| birds | Cyanomitra_olivacea | Nectarinia_olivacea | diurnal |
| birds | Cyanomitra_oritis | Nectarinia_oritis | diurnal |
| birds | Cyanomitra_verreauxii | Nectarinia_veroxii | diurnal |
| birds | Cyanomitra_verticalis | Nectarinia_verticalis | diurnal |
| birds | Cyanopica_cyanus | Cyanopica_cyanus | diurnal |
| birds | Cyanopsitta_spixii | Cyanopsitta_spixii | diurnal |

| | | | |
|---|---|---|---|
| birds | Cyanoptila_cyanomelana | Cyanoptila_cyanomelana | diurnal |
| birds | Cyanoramphus_auriceps | Cyanoramphus_auriceps | diurnal |
| birds | Cyanoramphus_forbesi | Cyanoramphus_forbesi | diurnal |
| birds | Cyanoramphus_malherbi | Cyanoramphus_malherbi | diurnal |
| birds | Cyanoramphus_novaezelandiae | Cyanoramphus_novaezelandiae | diurnal |
| birds | Cyanoramphus_unicolor | Cyanoramphus_unicolor | diurnal |
| birds | Cyclarhis_gujanensis | Cyclarhis_gujanensis | diurnal |
| birds | Cyclarhis_nigrirostris | Cyclarhis_nigrirostris | diurnal |
| birds | Cyclopsitta_diophthalma | Cyclopsitta_diophthalma | diurnal |
| birds | Cyclopsitta_gulielmitertii | Cyclopsitta_gulielmitertii | diurnal |
| birds | Cygnus_atratus | Cygnus_atratus | diurnal |
| birds | Cygnus_buccinator | Cygnus_buccinator | diurnal |
| birds | Cygnus_columbianus | Cygnus_columbianus | diurnal |
| birds | Cygnus_cygnus | Cygnus_cygnus | diurnal |
| birds | Cygnus_melancoryphus | Cygnus_melancoryphus | diurnal |
| birds | Cygnus_olor | Cygnus_olor | diurnal |
| birds | Cymbilaimus_lineatus | Cymbilaimus_lineatus | diurnal |
| birds | Cymbilaimus_sanctaemariae | Cymbilaimus_sanctaemariae | diurnal |
| birds | Cymbirhynchus_macrorhynchos | Cymbirhynchus_macrorhynchos | diurnal |
| birds | Cynanthus_latirostris | Cynanthus_latirostris | diurnal |
| birds | Cyornis_brunneatus | Rhinomyias_brunneatus | diurnal |
| birds | Cyornis_caerulatus | Cyornis_caerulatus | diurnal |
| birds | Cyornis_colonus | Rhinomyias_colonus | diurnal |
| birds | Cyornis_concretus | Cyornis_concretus | diurnal |
| birds | Cyornis_hainanus | Cyornis_hainanus | diurnal |
| birds | Cyornis_herioti | Cyornis_herioti | diurnal |
| birds | Cyornis_lemprieri | Cyornis_lemprieri | diurnal |
| birds | Cyornis_olivaceus | Rhinomyias_olivaceus | diurnal |
| birds | Cyornis_omissus | Cyornis_omissus | diurnal |
| birds | Cyornis_pallidipes | Cyornis_pallipes | diurnal |
| birds | Cyornis_rubeculoides | Cyornis_rubeculoides | diurnal |
| birds | Cyornis_ruficauda | Rhinomyias_ruficauda | diurnal |
| birds | Cyornis_rufigastra | Cyornis_rufigastra | diurnal |
| birds | Cyornis_tickelliae | Cyornis_tickelliae | diurnal |

| | | | |
|---|---|---|---|
| birds | Cyornis_umbratilis | Rhinomyias_umbratilis | diurnal |
| birds | Cyornis_unicolor | Cyornis_unicolor | diurnal |
| birds | Cyphorhinus_arada | Cyphorhinus_arada | diurnal |
| birds | Cyphorhinus_phaeocephalus | Cyphorhinus_phaeocephalus | diurnal |
| birds | Cyphorhinus_thoracicus | Cyphorhinus_thoracicus | diurnal |
| birds | Cyphos_macrodactylus | Bucco_macrodactylus | diurnal |
| birds | Cypseloides_cherriei | Cypseloides_cherriei | cathemeral |
| birds | Cypseloides_cryptus | Cypseloides_cryptus | diurnal |
| birds | Cypseloides_fumigatus | Cypseloides_fumigatus | diurnal |
| birds | Cypseloides_lemosi | Cypseloides_lemosi | diurnal |
| birds | Cypseloides_niger | Cypseloides_niger | diurnal |
| birds | Cypseloides_senex | Cypseloides_senex | diurnal |
| birds | Cypsnagra_hirundinacea | Cypsnagra_hirundinacea | diurnal |
| birds | Cyrtonyx_montezumae | Cyrtonyx_montezumae | diurnal |
| birds | Cyrtonyx_ocellatus | Cyrtonyx_ocellatus | diurnal |
| birds | Dacelo_gaudichaud | Dacelo_gaudichaud | diurnal |
| birds | Dacelo_leachii | Dacelo_leachii | diurnal |
| birds | Dacelo_novaeguineae | Dacelo_novaeguineae | diurnal |
| birds | Dacelo_rex | Clytoceyx_rex | cathemeral |
| birds | Dacelo_tyro | Dacelo_tyro | diurnal |
| birds | Dacnis_albiventris | Dacnis_albiventris | diurnal |
| birds | Dacnis_berlepschi | Dacnis_berlepschi | diurnal |
| birds | Dacnis_cayana | Dacnis_cayana | diurnal |
| birds | Dacnis_flaviventer | Dacnis_flaviventer | diurnal |
| birds | Dacnis_lineata | Dacnis_lineata | diurnal |
| birds | Dacnis_nigripes | Dacnis_nigripes | diurnal |
| birds | Dacnis_venusta | Dacnis_venusta | diurnal |
| birds | Dactylortyx_thoracicus | Dactylortyx_thoracicus | diurnal |
| birds | Daphoenositta_chrysoptera | Daphoenositta_chrysoptera | diurnal |
| birds | Daphoenositta_miranda | Daphoenositta_miranda | diurnal |
| birds | Daption_capense | Daption_capense | cathemeral |
| birds | Daptrius_ater | Daptrius_ater | diurnal |
| birds | Dasycrotapha_plateni | Stachyris_plateni | diurnal |
| birds | Dasycrotapha_speciosa | Dasycrotapha_speciosa | diurnal |

| | | | |
|---|---|---|---|
| birds | Dasylophus_superciliosus | Phaenicophaeus_superciliosus | diurnal |
| birds | Dasyornis_brachypterus | Dasyornis_brachypterus | diurnal |
| birds | Dasyornis_broadbenti | Dasyornis_broadbenti | diurnal |
| birds | Dasyornis_longirostris | Dasyornis_longirostris | diurnal |
| birds | Deconychura_longicauda | Deconychura_longicauda | diurnal |
| birds | Delacourella_capistrata | Nesocharis_capistrata | diurnal |
| birds | Deleornis_fraseri | Anthreptes_fraseri | diurnal |
| birds | Delichon_dasypus | Delichon_dasypus | diurnal |
| birds | Delichon_nipalense | Delichon_nipalense | diurnal |
| birds | Delichon_urbicum | Delichon_urbicum | diurnal |
| birds | Deltarhynchus_flammulatus | Deltarhynchus_flammulatus | diurnal |
| birds | Dendrexetastes_rufigula | Dendrexetastes_rufigula | diurnal |
| birds | Dendrocincla_anabatina | Dendrocincla_anabatina | diurnal |
| birds | Dendrocincla_fuliginosa | Dendrocincla_fuliginosa | diurnal |
| birds | Dendrocincla_homochroa | Dendrocincla_homochroa | diurnal |
| birds | Dendrocincla_merula | Dendrocincla_merula | diurnal |
| birds | Dendrocincla_turdina | Dendrocincla_turdina | diurnal |
| birds | Dendrocincla_tyrannina | Dendrocincla_tyrannina | diurnal |
| birds | Dendrocitta_bayleii | Dendrocitta_bayleyi | diurnal |
| birds | Dendrocitta_formosae | Dendrocitta_formosae | diurnal |
| birds | Dendrocitta_frontalis | Dendrocitta_frontalis | diurnal |
| birds | Dendrocitta_leucogastra | Dendrocitta_leucogastra | diurnal |
| birds | Dendrocitta_occipitalis | Dendrocitta_occipitalis | diurnal |
| birds | Dendrocitta_vagabunda | Dendrocitta_vagabunda | diurnal |
| birds | Dendrocolaptes_certhia | Dendrocolaptes_certhia | diurnal |
| birds | Dendrocolaptes_hoffmannsi | Dendrocolaptes_hoffmannsi | diurnal |
| birds | Dendrocolaptes_picumnus | Dendrocolaptes_picumnus | diurnal |
| birds | Dendrocolaptes_platyrostris | Dendrocolaptes_platyrostris | diurnal |
| birds | Dendrocolaptes_sanctithomae | Dendrocolaptes_sanctithomae | diurnal |
| birds | Dendrocopos_assimilis | Dendrocopos_assimilis | diurnal |
| birds | Dendrocopos_atratus | Dendrocopos_atratus | diurnal |
| birds | Dendrocopos_darjellensis | Dendrocopos_darjellensis | diurnal |
| birds | Dendrocopos_himalayensis | Dendrocopos_himalayensis | diurnal |
| birds | Dendrocopos_hyperythrus | Dendrocopos_hyperythrus | diurnal |

| | | | |
|---|---|---|---|
| birds | Dendrocopos_leucopterus | Dendrocopos_leucopterus | diurnal |
| birds | Dendrocopos_leucotos | Dendrocopos_leucotos | diurnal |
| birds | Dendrocopos_macei | Dendrocopos_macei | diurnal |
| birds | Dendrocopos_major | Dendrocopos_major | diurnal |
| birds | Dendrocopos_noguchii | Dendrocopos_noguchii | diurnal |
| birds | Dendrocopos_syriacus | Dendrocopos_syriacus | diurnal |
| birds | Dendrocygna_arborea | Dendrocygna_arborea | nocturnal |
| birds | Dendrocygna_arcuata | Dendrocygna_arcuata | diurnal |
| birds | Dendrocygna_autumnalis | Dendrocygna_autumnalis | diurnal |
| birds | Dendrocygna_bicolor | Dendrocygna_bicolor | nocturnal |
| birds | Dendrocygna_eytoni | Dendrocygna_eytoni | nocturnal |
| birds | Dendrocygna_guttata | Dendrocygna_guttata | nocturnal |
| birds | Dendrocygna_javanica | Dendrocygna_javanica | diurnal |
| birds | Dendrocygna_viduata | Dendrocygna_viduata | nocturnal |
| birds | Dendroma_erythroptera | Philydor_erythropterum | diurnal |
| birds | Dendroma_rufa | Philydor_rufum | diurnal |
| birds | Dendronanthus_indicus | Dendronanthus_indicus | diurnal |
| birds | Dendropicos_abyssinicus | Dendropicos_abyssinicus | diurnal |
| birds | Dendropicos_dorae | Dendrocopos_dorae | diurnal |
| birds | Dendropicos_elachus | Dendropicos_elachus | diurnal |
| birds | Dendropicos_elliotii | Mesopicos_elliotii | diurnal |
| birds | Dendropicos_fuscescens | Dendropicos_fuscescens | diurnal |
| birds | Dendropicos_gabonensis | Dendropicos_gabonensis | diurnal |
| birds | Dendropicos_goertae | Mesopicos_goertae | diurnal |
| birds | Dendropicos_griseocephalus | Mesopicos_griseocephalus | diurnal |
| birds | Dendropicos_namaquus | Thripias_namaquus | diurnal |
| birds | Dendropicos_obsoletus | Dendrocopos_obsoletus | diurnal |
| birds | Dendropicos_poecilolaemus | Dendropicos_poecilolaemus | diurnal |
| birds | Dendropicos_pyrrhogaster | Thripias_pyrrhogaster | diurnal |
| birds | Dendropicos_spodocephalus | Dendropicos_spodocephalus | diurnal |
| birds | Dendropicos_stierlingi | Dendropicos_stierlingi | diurnal |
| birds | Dendropicos_xantholophus | Thripias_xantholophus | diurnal |
| birds | Dendroplex_kienerii | Dendroplex_kienerii | diurnal |
| birds | Dendroplex_picus | Dendroplex_picus | diurnal |

| | | | |
|---|---|---|---|
| birds | Dendrortyx_barbatus | Dendrortyx_barbatus | diurnal |
| birds | Dendrortyx_leucophrys | Dendrortyx_leucophrys | diurnal |
| birds | Dendrortyx_macroura | Dendrortyx_macroura | diurnal |
| birds | Deroptyus_accipitrinus | Deroptyus_accipitrinus | diurnal |
| birds | Dessonornis_anomalus | Cossypha_anomala | diurnal |
| birds | Dessonornis_archeri | Cossypha_archeri | diurnal |
| birds | Dessonornis_caffer | Cossypha_caffra | diurnal |
| birds | Dessonornis_humeralis | Cossypha_humeralis | diurnal |
| birds | Devioeca_papuana | Microeca_papuana | diurnal |
| birds | Dicaeum_aeneum | Dicaeum_aeneum | diurnal |
| birds | Dicaeum_agile | Dicaeum_agile | diurnal |
| birds | Dicaeum_annae | Dicaeum_annae | diurnal |
| birds | Dicaeum_anthonyi | Dicaeum_anthonyi | diurnal |
| birds | Dicaeum_aureolimbatum | Dicaeum_aureolimbatum | diurnal |
| birds | Dicaeum_australe | Dicaeum_australe | diurnal |
| birds | Dicaeum_bicolor | Dicaeum_bicolor | diurnal |
| birds | Dicaeum_celebicum | Dicaeum_celebicum | diurnal |
| birds | Dicaeum_chrysorrheum | Dicaeum_chrysorrheum | diurnal |
| birds | Dicaeum_concolor | Dicaeum_concolor | diurnal |
| birds | Dicaeum_cruentatum | Dicaeum_cruentatum | diurnal |
| birds | Dicaeum_erythrorhynchos | Dicaeum_erythrorhynchos | diurnal |
| birds | Dicaeum_erythrothorax | Dicaeum_erythrothorax | diurnal |
| birds | Dicaeum_everetti | Dicaeum_everetti | diurnal |
| birds | Dicaeum_eximium | Dicaeum_eximium | diurnal |
| birds | Dicaeum_geelvinkianum | Dicaeum_geelvinkianum | diurnal |
| birds | Dicaeum_haematostictum | Dicaeum_haematostictum | diurnal |
| birds | Dicaeum_hirundinaceum | Dicaeum_hirundinaceum | diurnal |
| birds | Dicaeum_hypoleucum | Dicaeum_hypoleucum | diurnal |
| birds | Dicaeum_igniferum | Dicaeum_igniferum | diurnal |
| birds | Dicaeum_ignipectus | Dicaeum_ignipectus | diurnal |
| birds | Dicaeum_maugei | Dicaeum_maugei | diurnal |
| birds | Dicaeum_melanozanthum | Dicaeum_melanoxanthum | diurnal |
| birds | Dicaeum_monticolum | Dicaeum_monticolum | diurnal |
| birds | Dicaeum_nehrkorni | Dicaeum_nehrkorni | diurnal |

| | | | |
|---|---|---|---|
| birds | Dicaeum_nigrilore | Dicaeum_nigrilore | diurnal |
| birds | Dicaeum_nitidum | Dicaeum_nitidum | diurnal |
| birds | Dicaeum_pectorale | Dicaeum_pectorale | diurnal |
| birds | Dicaeum_proprium | Dicaeum_proprium | diurnal |
| birds | Dicaeum_pygmaeum | Dicaeum_pygmaeum | diurnal |
| birds | Dicaeum_quadricolor | Dicaeum_quadricolor | diurnal |
| birds | Dicaeum_retrocinctum | Dicaeum_retrocinctum | diurnal |
| birds | Dicaeum_sanguinolentum | Dicaeum_sanguinolentum | diurnal |
| birds | Dicaeum_trigonostigma | Dicaeum_trigonostigma | diurnal |
| birds | Dicaeum_tristrami | Dicaeum_tristrami | diurnal |
| birds | Dicaeum_trochileum | Dicaeum_trochileum | diurnal |
| birds | Dicaeum_vincens | Dicaeum_vincens | diurnal |
| birds | Dicaeum_vulneratum | Dicaeum_vulneratum | diurnal |
| birds | Dichrozona_cincta | Dichrozona_cincta | diurnal |
| birds | Dicrurus_adsimilis | Dicrurus_adsimilis | diurnal |
| birds | Dicrurus_aeneus | Dicrurus_aeneus | diurnal |
| birds | Dicrurus_aldabranus | Dicrurus_aldabranus | nocturnal |
| birds | Dicrurus_andamanensis | Dicrurus_andamanensis | diurnal |
| birds | Dicrurus_annectens | Dicrurus_annectans | diurnal |
| birds | Dicrurus_atripennis | Dicrurus_atripennis | diurnal |
| birds | Dicrurus_balicassius | Dicrurus_balicassius | diurnal |
| birds | Dicrurus_bracteatus | Dicrurus_bracteatus | diurnal |
| birds | Dicrurus_caerulescens | Dicrurus_caerulescens | diurnal |
| birds | Dicrurus_forficatus | Dicrurus_forficatus | diurnal |
| birds | Dicrurus_fuscipennis | Dicrurus_fuscipennis | diurnal |
| birds | Dicrurus_hottentottus | Dicrurus_hottentottus | diurnal |
| birds | Dicrurus_leucophaeus | Dicrurus_leucophaeus | diurnal |
| birds | Dicrurus_ludwigii | Dicrurus_ludwigii | diurnal |
| birds | Dicrurus_megarhynchus | Dicrurus_megarhynchus | diurnal |
| birds | Dicrurus_modestus | Dicrurus_modestus | nocturnal |
| birds | Dicrurus_montanus | Dicrurus_montanus | diurnal |
| birds | Dicrurus_paradiseus | Dicrurus_paradiseus | diurnal |
| birds | Dicrurus_remifer | Dicrurus_remifer | diurnal |
| birds | Dicrurus_sumatranus | Dicrurus_sumatranus | diurnal |

| | | | |
|---|---|---|---|
| birds | Dicrurus_waldenii | Dicrurus_waldenii | diurnal |
| birds | Didunculus_strigirostris | Didunculus_strigirostris | diurnal |
| birds | Diglossa_albilatera | Diglossa_albilatera | diurnal |
| birds | Diglossa_baritula | Diglossa_baritula | diurnal |
| birds | Diglossa_brunneiventris | Diglossa_brunneiventris | diurnal |
| birds | Diglossa_caerulescens | Diglossa_caerulescens | diurnal |
| birds | Diglossa_carbonaria | Diglossa_carbonaria | diurnal |
| birds | Diglossa_cyanea | Diglossa_cyanea | diurnal |
| birds | Diglossa_duidae | Diglossa_duidae | diurnal |
| birds | Diglossa_glauca | Diglossa_glauca | diurnal |
| birds | Diglossa_gloriosa | Diglossa_gloriosa | diurnal |
| birds | Diglossa_gloriosissima | Diglossa_gloriosissima | diurnal |
| birds | Diglossa_humeralis | Diglossa_humeralis | diurnal |
| birds | Diglossa_indigotica | Diglossa_indigotica | diurnal |
| birds | Diglossa_lafresnayii | Diglossa_lafresnayii | diurnal |
| birds | Diglossa_major | Diglossa_major | diurnal |
| birds | Diglossa_mystacalis | Diglossa_mystacalis | diurnal |
| birds | Diglossa_plumbea | Diglossa_plumbea | diurnal |
| birds | Diglossa_sittoides | Diglossa_sittoides | diurnal |
| birds | Diglossa_venezuelensis | Diglossa_venezuelensis | diurnal |
| birds | Dinemellia_dinemelli | Dinemellia_dinemelli | diurnal |
| birds | Dinopium_benghalense | Dinopium_benghalense | diurnal |
| birds | Dinopium_javanense | Dinopium_javanense | diurnal |
| birds | Dinopium_shorii | Dinopium_shorii | diurnal |
| birds | Diomedea_amsterdamensis | Diomedea_amsterdamensis | cathemeral |
| birds | Diomedea_epomophora | Diomedea_epomophora | cathemeral |
| birds | Diomedea_exulans | Diomedea_exulans | cathemeral |
| birds | Diopsittaca_nobilis | Diopsittaca_nobilis | diurnal |
| birds | Discosura_conversii | Discosura_conversii | diurnal |
| birds | Discosura_langsdorffi | Discosura_langsdorffi | diurnal |
| birds | Discosura_letitiae | Discosura_letitiae | diurnal |
| birds | Discosura_longicaudus | Discosura_longicaudus | diurnal |
| birds | Discosura_popelairii | Discosura_popelairii | diurnal |
| birds | Diuca_diuca | Diuca_diuca | diurnal |

| | | | |
|---|---|---|---|
| birds | Dives_dives | Dives_dives | diurnal |
| birds | Dives_warczewiczi | Dives_warszewiczi | diurnal |
| birds | Dolichonyx_oryzivorus | Dolichonyx_oryzivorus | diurnal |
| birds | Doliornis_remseni | Doliornis_remseni | diurnal |
| birds | Doliornis_sclateri | Doliornis_sclateri | diurnal |
| birds | Donacobius_atricapilla | Donacobius_atricapilla | diurnal |
| birds | Donacospiza_albifrons | Donacospiza_albifrons | diurnal |
| birds | Doricha_eliza | Doricha_eliza | diurnal |
| birds | Doricha_enicura | Doricha_enicura | diurnal |
| birds | Doryfera_johannae | Doryfera_johannae | diurnal |
| birds | Doryfera_ludovicae | Doryfera_ludovicae | diurnal |
| birds | Drepanis_coccinea | Vestiaria_coccinea | diurnal |
| birds | Drepanoptila_holosericea | Drepanoptila_holosericea | diurnal |
| birds | Drepanorhynchus_reichenowi | Nectarinia_reichenowi | diurnal |
| birds | Drepanornis_albertisi | Epimachus_albertisi | diurnal |
| birds | Drepanornis_bruijnii | Epimachus_bruijnii | diurnal |
| birds | Dreptes_thomensis | Nectarinia_thomensis | diurnal |
| birds | Dromaius_novaehollandiae | Dromaius_novaehollandiae | diurnal |
| birds | Dromas_ardeola | Dromas_ardeola | nocturnal |
| birds | Dromococcyx_pavoninus | Dromococcyx_pavoninus | diurnal |
| birds | Dromococcyx_phasianellus | Dromococcyx_phasianellus | diurnal |
| birds | Drymocichla_incana | Drymocichla_incana | diurnal |
| birds | Drymodes_brunneopygia | Drymodes_brunneopygia | diurnal |
| birds | Drymodes_superciliaris | Drymodes_superciliaris | diurnal |
| birds | Drymophila_caudata | Drymophila_caudata | diurnal |
| birds | Drymophila_devillei | Drymophila_devillei | diurnal |
| birds | Drymophila_ferruginea | Drymophila_ferruginea | diurnal |
| birds | Drymophila_genei | Drymophila_genei | diurnal |
| birds | Drymophila_malura | Drymophila_malura | diurnal |
| birds | Drymophila_ochropyga | Drymophila_ochropyga | diurnal |
| birds | Drymophila_rubricollis | Drymophila_rubricollis | diurnal |
| birds | Drymophila_squamata | Drymophila_squamata | diurnal |
| birds | Drymornis_bridgesii | Drymornis_bridgesii | diurnal |
| birds | Drymotoxeres_pucheranii | Campylorhamphus_pucherani | diurnal |

| | | | |
|---|---|---|---|
| birds | Dryobates_cathpharius | Dendrocopos_cathpharius | diurnal |
| birds | Dryobates_minor | Dendrocopos_minor | diurnal |
| birds | Dryobates_nuttallii | Picoides_nuttallii | diurnal |
| birds | Dryobates_pubescens | Picoides_pubescens | diurnal |
| birds | Dryobates_scalaris | Picoides_scalaris | diurnal |
| birds | Dryocopus_hodgei | Dryocopus_hodgei | diurnal |
| birds | Dryocopus_javensis | Dryocopus_javensis | diurnal |
| birds | Dryocopus_martius | Dryocopus_martius | diurnal |
| birds | Dryolimnas_cuvieri | Dryolimnas_cuvieri | diurnal |
| birds | Dryoscopus_angolensis | Dryoscopus_angolensis | diurnal |
| birds | Dryoscopus_cubla | Dryoscopus_cubla | diurnal |
| birds | Dryoscopus_gambensis | Dryoscopus_gambensis | diurnal |
| birds | Dryoscopus_pringlii | Dryoscopus_pringlii | diurnal |
| birds | Dryoscopus_sabini | Dryoscopus_sabini | diurnal |
| birds | Dryoscopus_senegalensis | Dryoscopus_senegalensis | diurnal |
| birds | Dryotriorchis_spectabilis | Dryotriorchis_spectabilis | diurnal |
| birds | Dubusia_castaneoventris | Delothraupis_castaneoventris | diurnal |
| birds | Dubusia_taeniata | Dubusia_taeniata | diurnal |
| birds | Ducula_aenea | Ducula_aenea | diurnal |
| birds | Ducula_aurorae | Ducula_aurorae | diurnal |
| birds | Ducula_badia | Ducula_badia | diurnal |
| birds | Ducula_bakeri | Ducula_bakeri | diurnal |
| birds | Ducula_basilica | Ducula_basilica | diurnal |
| birds | Ducula_bicolor | Ducula_bicolor | diurnal |
| birds | Ducula_brenchleyi | Ducula_brenchleyi | diurnal |
| birds | Ducula_carola | Ducula_carola | diurnal |
| birds | Ducula_chalconota | Ducula_chalconota | diurnal |
| birds | Ducula_cineracea | Ducula_cineracea | diurnal |
| birds | Ducula_concinna | Ducula_concinna | diurnal |
| birds | Ducula_finschii | Ducula_finschii | diurnal |
| birds | Ducula_forsteni | Ducula_forsteni | diurnal |
| birds | Ducula_galeata | Ducula_galeata | diurnal |
| birds | Ducula_goliath | Ducula_goliath | diurnal |
| birds | Ducula_lacernulata | Ducula_lacernulata | diurnal |

| | | | |
|---|---|---|---|
| birds | Ducula_latrans | Ducula_latrans | diurnal |
| birds | Ducula_luctuosa | Ducula_luctuosa | diurnal |
| birds | Ducula_melanochroa | Ducula_melanochroa | diurnal |
| birds | Ducula_mindorensis | Ducula_mindorensis | diurnal |
| birds | Ducula_mullerii | Ducula_mullerii | diurnal |
| birds | Ducula_myristicivora | Ducula_myristicivora | diurnal |
| birds | Ducula_oceanica | Ducula_oceanica | diurnal |
| birds | Ducula_pacifica | Ducula_pacifica | diurnal |
| birds | Ducula_perspicillata | Ducula_perspicillata | diurnal |
| birds | Ducula_pickeringii | Ducula_pickeringii | diurnal |
| birds | Ducula_pinon | Ducula_pinon | diurnal |
| birds | Ducula_pistrinaria | Ducula_pistrinaria | diurnal |
| birds | Ducula_poliocephala | Ducula_poliocephala | diurnal |
| birds | Ducula_radiata | Ducula_radiata | diurnal |
| birds | Ducula_rosacea | Ducula_rosacea | diurnal |
| birds | Ducula_rubricera | Ducula_rubricera | diurnal |
| birds | Ducula_rufigaster | Ducula_rufigaster | diurnal |
| birds | Ducula_spilorrhoa | Ducula_spilorrhoa | diurnal |
| birds | Ducula_whartoni | Ducula_whartoni | diurnal |
| birds | Ducula_zoeae | Ducula_zoeae | diurnal |
| birds | Dulus_dominicus | Dulus_dominicus | diurnal |
| birds | Dumetella_carolinensis | Dumetella_carolinensis | diurnal |
| birds | Dumetia_atriceps | Rhopocichla_atriceps | diurnal |
| birds | Dumetia_hyperythra | Dumetia_hyperythra | diurnal |
| birds | Dyaphorophyia_blissetti | Platysteira_blissetti | diurnal |
| birds | Dyaphorophyia_castanea | Platysteira_castanea | diurnal |
| birds | Dyaphorophyia_concreta | Platysteira_concreta | diurnal |
| birds | Dyaphorophyia_tonsa | Platysteira_tonsa | diurnal |
| birds | Dysithamnus_leucostictus | Dysithamnus_leucostictus | diurnal |
| birds | Dysithamnus_mentalis | Dysithamnus_mentalis | diurnal |
| birds | Dysithamnus_occidentalis | Dysithamnus_occidentalis | diurnal |
| birds | Dysithamnus_plumbeus | Dysithamnus_plumbeus | diurnal |
| birds | Dysithamnus_puncticeps | Dysithamnus_puncticeps | diurnal |
| birds | Dysithamnus_stictothorax | Dysithamnus_stictothorax | diurnal |

| | | | |
|---|---|---|---|
| birds | Dysithamnus_striaticeps | Dysithamnus_striaticeps | diurnal |
| birds | Dysithamnus_xanthopterus | Dysithamnus_xanthopterus | diurnal |
| birds | Eclectus_roratus | Eclectus_roratus | nocturnal |
| birds | Edolisoma_anale | Coracina_analis | diurnal |
| birds | Edolisoma_ceramense | Coracina_ceramensis | diurnal |
| birds | Edolisoma_coerulescens | Coracina_coerulescens | diurnal |
| birds | Edolisoma_dispar | Coracina_dispar | diurnal |
| birds | Edolisoma_dohertyi | Coracina_dohertyi | diurnal |
| birds | Edolisoma_holopolium | Coracina_holopolia | diurnal |
| birds | Edolisoma_incertum | Coracina_incerta | diurnal |
| birds | Edolisoma_melas | Coracina_melas | diurnal |
| birds | Edolisoma_mindanense | Coracina_mindanensis | diurnal |
| birds | Edolisoma_montanum | Coracina_montana | diurnal |
| birds | Edolisoma_morio | Coracina_morio | diurnal |
| birds | Edolisoma_ostentum | Coracina_ostenta | diurnal |
| birds | Edolisoma_schisticeps | Coracina_schisticeps | diurnal |
| birds | Edolisoma_sula | Coracina_sula | diurnal |
| birds | Edolisoma_tenuirostre | Coracina_tenuirostris | diurnal |
| birds | Egretta_ardesiaca | Egretta_ardesiaca | diurnal |
| birds | Egretta_caerulea | Egretta_caerulea | diurnal |
| birds | Egretta_eulophotes | Egretta_eulophotes | diurnal |
| birds | Egretta_novaehollandiae | Egretta_novaehollandiae | diurnal |
| birds | Egretta_picata | Ardea_picata | diurnal |
| birds | Egretta_rufescens | Egretta_rufescens | diurnal |
| birds | Egretta_sacra | Egretta_sacra | cathemeral |
| birds | Egretta_thula | Egretta_thula | diurnal |
| birds | Egretta_tricolor | Egretta_tricolor | diurnal |
| birds | Egretta_vinaceigula | Egretta_vinaceigula | diurnal |
| birds | Elachura_formosa | Spelaeornis_formosus | diurnal |
| birds | Elaenia_albiceps | Elaenia_albiceps | diurnal |
| birds | Elaenia_chiriquensis | Elaenia_chiriquensis | diurnal |
| birds | Elaenia_cristata | Elaenia_cristata | diurnal |
| birds | Elaenia_dayi | Elaenia_dayi | diurnal |
| birds | Elaenia_fallax | Elaenia_fallax | diurnal |

| | | | |
|---|---|---|---|
| birds | Elaenia_flavogaster | Elaenia_flavogaster | diurnal |
| birds | Elaenia_frantzii | Elaenia_frantzii | diurnal |
| birds | Elaenia_gigas | Elaenia_gigas | diurnal |
| birds | Elaenia_martinica | Elaenia_martinica | diurnal |
| birds | Elaenia_mesoleuca | Elaenia_mesoleuca | diurnal |
| birds | Elaenia_obscura | Elaenia_obscura | diurnal |
| birds | Elaenia_pallatangae | Elaenia_pallatangae | diurnal |
| birds | Elaenia_parvirostris | Elaenia_parvirostris | diurnal |
| birds | Elaenia_ridleyana | Elaenia_ridleyana | diurnal |
| birds | Elaenia_ruficeps | Elaenia_ruficeps | diurnal |
| birds | Elaenia_spectabilis | Elaenia_spectabilis | diurnal |
| birds | Elaenia_strepera | Elaenia_strepera | diurnal |
| birds | Elanoides_forficatus | Elanoides_forficatus | diurnal |
| birds | Elanus_axillaris | Elanus_axillaris | cathemeral |
| birds | Elanus_caeruleus | Elanus_caeruleus | diurnal |
| birds | Elanus_leucurus | Elanus_leucurus | diurnal |
| birds | Elanus_scriptus | Elanus_scriptus | nocturnal |
| birds | Elaphrornis_palliseri | Bradypterus_palliseri | diurnal |
| birds | Electron_carinatum | Electron_carinatum | diurnal |
| birds | Electron_platyrhynchum | Electron_platyrhynchum | diurnal |
| birds | Eleoscytalopus_indigoticus | Eleoscytalopus_indigoticus | diurnal |
| birds | Eleoscytalopus_psychopompus | Eleoscytalopus_psychopompus | diurnal |
| birds | Eleothreptus_anomalus | Eleothreptus_anomalus | nocturnal |
| birds | Eleothreptus_candicans | Eleothreptus_candicans | nocturnal |
| birds | Elliotomyia_chionogaster | Amazilia_chionogaster | diurnal |
| birds | Elliotomyia_viridicauda | Amazilia_viridicauda | diurnal |
| birds | Elminia_albicauda | Elminia_albicauda | diurnal |
| birds | Elminia_albiventris | Elminia_albiventris | diurnal |
| birds | Elminia_albonotata | Elminia_albonotata | diurnal |
| birds | Elminia_longicauda | Elminia_longicauda | diurnal |
| birds | Elminia_nigromitrata | Elminia_nigromitrata | diurnal |
| birds | Elseyornis_melanops | Elseyornis_melanops | diurnal |
| birds | Emarginata_schlegelii | Cercomela_schlegelii | diurnal |
| birds | Emarginata_sinuata | Cercomela_sinuata | diurnal |

| | | | |
|---|---|---|---|
| birds | Emarginata_tractrac | Cercomela_tractrac | diurnal |
| birds | Emberiza_affinis | Emberiza_affinis | diurnal |
| birds | Emberiza_aureola | Emberiza_aureola | diurnal |
| birds | Emberiza_bruniceps | Emberiza_bruniceps | diurnal |
| birds | Emberiza_buchanani | Emberiza_buchanani | diurnal |
| birds | Emberiza_cabanisi | Emberiza_cabanisi | diurnal |
| birds | Emberiza_caesia | Emberiza_caesia | diurnal |
| birds | Emberiza_calandra | Miliaria_calandra | diurnal |
| birds | Emberiza_capensis | Emberiza_capensis | diurnal |
| birds | Emberiza_chrysophrys | Emberiza_chrysophrys | diurnal |
| birds | Emberiza_cia | Emberiza_cia | diurnal |
| birds | Emberiza_cineracea | Emberiza_cineracea | diurnal |
| birds | Emberiza_cioides | Emberiza_cioides | diurnal |
| birds | Emberiza_cirlus | Emberiza_cirlus | diurnal |
| birds | Emberiza_citrinella | Emberiza_citrinella | diurnal |
| birds | Emberiza_elegans | Emberiza_elegans | diurnal |
| birds | Emberiza_flaviventris | Emberiza_flaviventris | diurnal |
| birds | Emberiza_fucata | Emberiza_fucata | diurnal |
| birds | Emberiza_godlewskii | Emberiza_godlewskii | diurnal |
| birds | Emberiza_hortulana | Emberiza_hortulana | diurnal |
| birds | Emberiza_impetuani | Emberiza_impetuani | diurnal |
| birds | Emberiza_jankowskii | Emberiza_jankowskii | diurnal |
| birds | Emberiza_koslowi | Emberiza_koslowi | diurnal |
| birds | Emberiza_lathami | Melophus_lathami | diurnal |
| birds | Emberiza_leucocephalos | Emberiza_leucocephalos | diurnal |
| birds | Emberiza_melanocephala | Emberiza_melanocephala | diurnal |
| birds | Emberiza_pallasi | Emberiza_pallasi | diurnal |
| birds | Emberiza_poliopleura | Emberiza_poliopleura | diurnal |
| birds | Emberiza_pusilla | Emberiza_pusilla | diurnal |
| birds | Emberiza_rustica | Emberiza_rustica | diurnal |
| birds | Emberiza_rutila | Emberiza_rutila | diurnal |
| birds | Emberiza_schoeniclus | Emberiza_schoeniclus | diurnal |
| birds | Emberiza_siemsseni | Latoucheornis_siemsseni | diurnal |
| birds | Emberiza_socotrana | Emberiza_socotrana | diurnal |

| | | | |
|---|---|---|---|
| birds | Emberiza_spodocephala | Emberiza_spodocephala | diurnal |
| birds | Emberiza_stewarti | Emberiza_stewarti | diurnal |
| birds | Emberiza_striolata | Emberiza_striolata | diurnal |
| birds | Emberiza_sulphurata | Emberiza_sulphurata | diurnal |
| birds | Emberiza_tahapisi | Emberiza_tahapisi | diurnal |
| birds | Emberiza_tristrami | Emberiza_tristrami | diurnal |
| birds | Emberiza_variabilis | Emberiza_variabilis | diurnal |
| birds | Emberiza_yessoensis | Emberiza_yessoensis | diurnal |
| birds | Emberizoides_herbicola | Emberizoides_herbicola | diurnal |
| birds | Emberizoides_ypiranganus | Emberizoides_ypiranganus | diurnal |
| birds | Embernagra_longicauda | Embernagra_longicauda | diurnal |
| birds | Embernagra_platensis | Embernagra_platensis | diurnal |
| birds | Emblema_pictum | Emblema_pictum | diurnal |
| birds | Eminia_lepida | Eminia_lepida | diurnal |
| birds | Empidonax_affinis | Empidonax_affinis | diurnal |
| birds | Empidonax_albigularis | Empidonax_albigularis | diurnal |
| birds | Empidonax_alnorum | Empidonax_alnorum | diurnal |
| birds | Empidonax_atriceps | Empidonax_atriceps | diurnal |
| birds | Empidonax_difficilis | Empidonax_difficilis | diurnal |
| birds | Empidonax_flavescens | Empidonax_flavescens | diurnal |
| birds | Empidonax_flaviventris | Empidonax_flaviventris | diurnal |
| birds | Empidonax_fulvifrons | Empidonax_fulvifrons | diurnal |
| birds | Empidonax_minimus | Empidonax_minimus | diurnal |
| birds | Empidonax_oberholseri | Empidonax_oberholseri | diurnal |
| birds | Empidonax_occidentalis | Empidonax_occidentalis | diurnal |
| birds | Empidonax_traillii | Empidonax_traillii | diurnal |
| birds | Empidonax_virescens | Empidonax_virescens | diurnal |
| birds | Empidonax_wrightii | Empidonax_wrightii | diurnal |
| birds | Empidonomus_varius | Empidonomus_varius | diurnal |
| birds | Empidornis_semipartitus | Empidornis_semipartitus | diurnal |
| birds | Enicognathus_ferrugineus | Enicognathus_ferrugineus | diurnal |
| birds | Enicognathus_leptorhynchus | Enicognathus_leptorhynchus | diurnal |
| birds | Enicurus_immaculatus | Enicurus_immaculatus | diurnal |
| birds | Enicurus_leschenaulti | Enicurus_leschenaulti | diurnal |

| | | | |
|---|---|---|---|
| birds | Enicurus_maculatus | Enicurus_maculatus | diurnal |
| birds | Enicurus_ruficapillus | Enicurus_ruficapillus | diurnal |
| birds | Enicurus_schistaceus | Enicurus_schistaceus | diurnal |
| birds | Enicurus_scouleri | Enicurus_scouleri | diurnal |
| birds | Enicurus_velatus | Enicurus_velatus | diurnal |
| birds | Enodes_erythrophris | Enodes_erythrophris | diurnal |
| birds | Ensifera_ensifera | Ensifera_ensifera | diurnal |
| birds | Entomodestes_coracinus | Entomodestes_coracinus | diurnal |
| birds | Entomodestes_leucotis | Entomodestes_leucotis | diurnal |
| birds | Entomyzon_cyanotis | Entomyzon_cyanotis | diurnal |
| birds | Eolophus_roseicapilla | Cacatua_roseicapilla | diurnal |
| birds | Eophona_migratoria | Eophona_migratoria | diurnal |
| birds | Eophona_personata | Eophona_personata | diurnal |
| birds | Eopsaltria_australis | Eopsaltria_australis | diurnal |
| birds | Eopsaltria_griseogularis | Eopsaltria_griseogularis | diurnal |
| birds | Eos_bornea | Eos_bornea | diurnal |
| birds | Eos_cyanogenia | Eos_cyanogenia | diurnal |
| birds | Eos_histrio | Eos_histrio | diurnal |
| birds | Eos_reticulata | Eos_reticulata | diurnal |
| birds | Eos_semilarvata | Eos_semilarvata | diurnal |
| birds | Eos_squamata | Eos_squamata | diurnal |
| birds | Ephippiorhynchus_asiaticus | Ephippiorhynchus_asiaticus | diurnal |
| birds | Ephippiorhynchus_senegalensis | Ephippiorhynchus_senegalensis | diurnal |
| birds | Ephippiospingus_dorsalis | Phrygilus_dorsalis | diurnal |
| birds | Ephippiospingus_erythronotus | Phrygilus_erythronotus | diurnal |
| birds | Epimachus_fastosus | Epimachus_fastuosus | diurnal |
| birds | Epimachus_meyeri | Epimachus_meyeri | diurnal |
| birds | Epinecrophylla_erythrura | Epinecrophylla_erythrura | diurnal |
| birds | Epinecrophylla_fulviventris | Epinecrophylla_fulviventris | diurnal |
| birds | Epinecrophylla_gutturalis | Myrmotherula_gutturalis | diurnal |
| birds | Epinecrophylla_haematonota | Epinecrophylla_haematonota | diurnal |
| birds | Epinecrophylla_leucophthalma | Epinecrophylla_leucophthalma | diurnal |
| birds | Epinecrophylla_ornata | Epinecrophylla_ornata | diurnal |
| birds | Epinecrophylla_spodionota | Epinecrophylla_spodionota | diurnal |

| | | | |
|---|---|---|---|
| birds | Epthianura_albifrons | Epthianura_albifrons | diurnal |
| birds | Epthianura_aurifrons | Epthianura_aurifrons | diurnal |
| birds | Epthianura_crocea | Epthianura_crocea | diurnal |
| birds | Epthianura_tricolor | Epthianura_tricolor | diurnal |
| birds | Eremalauda_dunni | Eremalauda_dunni | diurnal |
| birds | Eremomela_atricollis | Eremomela_atricollis | diurnal |
| birds | Eremomela_badiceps | Eremomela_badiceps | diurnal |
| birds | Eremomela_canescens | Eremomela_canescens | diurnal |
| birds | Eremomela_flavicrissalis | Eremomela_flavicrissalis | diurnal |
| birds | Eremomela_gregalis | Eremomela_gregalis | diurnal |
| birds | Eremomela_icteropygialis | Eremomela_icteropygialis | diurnal |
| birds | Eremomela_pusilla | Eremomela_pusilla | diurnal |
| birds | Eremomela_scotops | Eremomela_scotops | diurnal |
| birds | Eremomela_turneri | Eremomela_turneri | diurnal |
| birds | Eremomela_usticollis | Eremomela_usticollis | diurnal |
| birds | Eremophila_alpestris | Eremophila_alpestris | diurnal |
| birds | Eremophila_bilopha | Eremophila_bilopha | diurnal |
| birds | Eremopterix_australis | Eremopterix_australis | diurnal |
| birds | Eremopterix_griseus | Eremopterix_griseus | diurnal |
| birds | Eremopterix_hova | Mirafra_hova | diurnal |
| birds | Eremopterix_leucopareia | Eremopterix_leucopareia | diurnal |
| birds | Eremopterix_leucotis | Eremopterix_leucotis | diurnal |
| birds | Eremopterix_signatus | Eremopterix_signatus | diurnal |
| birds | Eriocnemis_aline | Eriocnemis_alinae | diurnal |
| birds | Eriocnemis_cupreoventris | Eriocnemis_cupreoventris | diurnal |
| birds | Eriocnemis_derbyi | Eriocnemis_derbyi | diurnal |
| birds | Eriocnemis_glaucopoides | Eriocnemis_glaucopoides | diurnal |
| birds | Eriocnemis_godini | Eriocnemis_godini | diurnal |
| birds | Eriocnemis_luciani | Eriocnemis_luciani | diurnal |
| birds | Eriocnemis_mirabilis | Eriocnemis_mirabilis | diurnal |
| birds | Eriocnemis_mosquera | Eriocnemis_mosquera | diurnal |
| birds | Eriocnemis_nigrivestis | Eriocnemis_nigrivestis | diurnal |
| birds | Eriocnemis_vestita | Eriocnemis_vestita | diurnal |
| birds | Erithacus_rubecula | Erithacus_rubecula | diurnal |

| | | | |
|---|---|---|---|
| birds | Erpornis_zantholeuca | Erpornis_zantholeuca | diurnal |
| birds | Erythrocercus_holochlorus | Erythrocercus_holochlorus | diurnal |
| birds | Erythrocercus_livingstonei | Erythrocercus_livingstonei | diurnal |
| birds | Erythrocercus_mccallii | Erythrocercus_mccallii | diurnal |
| birds | Erythrogenys_erythrocnemis | Pomatorhinus_erythrocnemis | diurnal |
| birds | Erythrogenys_erythrogenys | Pomatorhinus_erythrogenys | diurnal |
| birds | Erythrogenys_hypoleucos | Pomatorhinus_hypoleucos | diurnal |
| birds | Erythrogonys_cinctus | Erythrogonys_cinctus | diurnal |
| birds | Erythropitta_arquata | Pitta_arcuata | diurnal |
| birds | Erythropitta_erythrogaster | Pitta_erythrogaster | diurnal |
| birds | Erythropitta_granatina | Pitta_granatina | diurnal |
| birds | Erythropitta_kochi | Pitta_kochi | diurnal |
| birds | Erythropitta_ussheri | Pitta_ussheri | diurnal |
| birds | Erythrotriorchis_buergersi | Erythrotriorchis_buergersi | diurnal |
| birds | Erythrotriorchis_radiatus | Erythrotriorchis_radiatus | diurnal |
| birds | Erythrura_coloria | Erythrura_coloria | diurnal |
| birds | Erythrura_cyaneovirens | Erythrura_cyaneovirens | diurnal |
| birds | Erythrura_hyperythra | Erythrura_hyperythra | diurnal |
| birds | Erythrura_kleinschmidti | Erythrura_kleinschmidti | diurnal |
| birds | Erythrura_papuana | Erythrura_papuana | diurnal |
| birds | Erythrura_pealii | Erythrura_pealii | diurnal |
| birds | Erythrura_prasina | Erythrura_prasina | diurnal |
| birds | Erythrura_psittacea | Erythrura_psittacea | diurnal |
| birds | Erythrura_trichroa | Erythrura_trichroa | diurnal |
| birds | Erythrura_tricolor | Erythrura_tricolor | diurnal |
| birds | Erythrura_viridifacies | Erythrura_viridifacies | diurnal |
| birds | Esacus_magnirostris | Esacus_giganteus | diurnal |
| birds | Esacus_recurvirostris | Esacus_recurvirostris | nocturnal |
| birds | Estrilda_astrild | Estrilda_astrild | diurnal |
| birds | Estrilda_atricapilla | Estrilda_atricapilla | diurnal |
| birds | Estrilda_kandti | Estrilda_kandti | diurnal |
| birds | Estrilda_melpoda | Estrilda_melpoda | diurnal |
| birds | Estrilda_nigriloris | Estrilda_nigriloris | diurnal |
| birds | Estrilda_nonnula | Estrilda_nonnula | diurnal |

| | | | |
|---|---|---|---|
| birds | Estrilda_paludicola | Estrilda_paludicola | diurnal |
| birds | Estrilda_poliopareia | Estrilda_poliopareia | diurnal |
| birds | Estrilda_rhodopyga | Estrilda_rhodopyga | diurnal |
| birds | Estrilda_rufibarba | Estrilda_rufibarba | diurnal |
| birds | Estrilda_troglodytes | Estrilda_troglodytes | diurnal |
| birds | Eubucco_bourcierii | Eubucco_bourcierii | diurnal |
| birds | Eubucco_richardsoni | Eubucco_richardsoni | diurnal |
| birds | Eubucco_tucinkae | Eubucco_tucinkae | diurnal |
| birds | Eubucco_versicolor | Eubucco_versicolor | diurnal |
| birds | Euchrepomis_callinota | Terenura_callinota | diurnal |
| birds | Euchrepomis_humeralis | Terenura_humeralis | diurnal |
| birds | Euchrepomis_sharpei | Terenura_sharpei | diurnal |
| birds | Euchrepomis_spodioptila | Terenura_spodioptila | diurnal |
| birds | Eucometis_penicillata | Eucometis_penicillata | diurnal |
| birds | Eudocimus_albus | Eudocimus_albus | diurnal |
| birds | Eudocimus_ruber | Eudocimus_ruber | diurnal |
| birds | Eudromia_elegans | Eudromia_elegans | diurnal |
| birds | Eudromia_formosa | Eudromia_formosa | diurnal |
| birds | Eudromias_morinellus | Eudromias_morinellus | cathemeral |
| birds | Eudynamys_scolopaceus | Eudynamys_scolopaceus | diurnal |
| birds | Eudyptes_chrysocome | Eudyptes_chrysocome | cathemeral |
| birds | Eudyptes_chrysolophus | Eudyptes_chrysolophus | cathemeral |
| birds | Eudyptes_pachyrhynchus | Eudyptes_pachyrhynchus | cathemeral |
| birds | Eudyptes_robustus | Eudyptes_robustus | cathemeral |
| birds | Eudyptes_schlegeli | Eudyptes_schlegeli | cathemeral |
| birds | Eudyptes_sclateri | Eudyptes_sclateri | cathemeral |
| birds | Eudyptula_minor | Eudyptula_minor | cathemeral |
| birds | Eugenes_fulgens | Eugenes_fulgens | diurnal |
| birds | Eugerygone_rubra | Eugerygone_rubra | diurnal |
| birds | Eugralla_paradoxa | Eugralla_paradoxa | diurnal |
| birds | Eulabeornis_castaneoventris | Eulabeornis_castaneoventris | cathemeral |
| birds | Eulacestoma_nigropectus | Eulacestoma_nigropectus | diurnal |
| birds | Eulampis_holosericeus | Eulampis_holosericeus | diurnal |
| birds | Eulampis_jugularis | Eulampis_jugularis | diurnal |

| | | | |
|---|---|---|---|
| birds | Eulidia_yarrellii | Eulidia_yarrellii | diurnal |
| birds | Eulipoa_wallacei | Eulipoa_wallacei | diurnal |
| birds | Eumomota_superciliosa | Eumomota_superciliosa | diurnal |
| birds | Eumyias_additus | Rhinomyias_additus | diurnal |
| birds | Eumyias_albicaudatus | Eumyias_albicaudatus | diurnal |
| birds | Eumyias_hyacinthinus | Cyornis_hyacinthinus | diurnal |
| birds | Eumyias_indigo | Eumyias_indigo | diurnal |
| birds | Eumyias_panayensis | Eumyias_panayensis | diurnal |
| birds | Eumyias_sordidus | Eumyias_sordidus | diurnal |
| birds | Eumyias_thalassinus | Eumyias_thalassinus | diurnal |
| birds | Euneornis_campestris | Euneornis_campestris | diurnal |
| birds | Eunymphicus_cornutus | Eunymphicus_cornutus | diurnal |
| birds | Euodice_cantans | Lonchura_cantans | diurnal |
| birds | Euodice_malabarica | Lonchura_malabarica | diurnal |
| birds | Eupetes_macrocerus | Eupetes_macrocerus | diurnal |
| birds | Eupetomena_cirrochloris | Aphantochroa_cirrochloris | diurnal |
| birds | Eupetomena_macroura | Eupetomena_macroura | diurnal |
| birds | Euphagus_carolinus | Euphagus_carolinus | diurnal |
| birds | Euphagus_cyanocephalus | Euphagus_cyanocephalus | diurnal |
| birds | Eupherusa_cyanophrys | Eupherusa_cyanophrys | diurnal |
| birds | Eupherusa_eximia | Eupherusa_eximia | diurnal |
| birds | Eupherusa_nigriventris | Eupherusa_nigriventris | diurnal |
| birds | Eupherusa_poliocerca | Eupherusa_poliocerca | diurnal |
| birds | Eupherusa_ridgwayi | Thalurania_ridgwayi | diurnal |
| birds | Euphonia_affinis | Euphonia_affinis | diurnal |
| birds | Euphonia_anneae | Euphonia_anneae | diurnal |
| birds | Euphonia_cayennensis | Euphonia_cayennensis | diurnal |
| birds | Euphonia_chalybea | Euphonia_chalybea | diurnal |
| birds | Euphonia_chlorotica | Euphonia_chlorotica | diurnal |
| birds | Euphonia_chrysopasta | Euphonia_chrysopasta | diurnal |
| birds | Euphonia_concinna | Euphonia_concinna | diurnal |
| birds | Euphonia_cyanocephala | Euphonia_cyanocephala | diurnal |
| birds | Euphonia_elegantissima | Euphonia_elegantissima | diurnal |
| birds | Euphonia_finschi | Euphonia_finschi | diurnal |

| | | | |
|---|---|---|---|
| birds | Euphonia_fulvicrissa | Euphonia_fulvicrissa | diurnal |
| birds | Euphonia_gouldi | Euphonia_gouldi | diurnal |
| birds | Euphonia_hirundinacea | Euphonia_hirundinacea | diurnal |
| birds | Euphonia_imitans | Euphonia_imitans | diurnal |
| birds | Euphonia_jamaica | Euphonia_jamaica | diurnal |
| birds | Euphonia_laniirostris | Euphonia_laniirostris | diurnal |
| birds | Euphonia_luteicapilla | Euphonia_luteicapilla | diurnal |
| birds | Euphonia_mesochrysa | Euphonia_mesochrysa | diurnal |
| birds | Euphonia_minuta | Euphonia_minuta | diurnal |
| birds | Euphonia_musica | Euphonia_musica | diurnal |
| birds | Euphonia_pectoralis | Euphonia_pectoralis | diurnal |
| birds | Euphonia_plumbea | Euphonia_plumbea | diurnal |
| birds | Euphonia_rufiventris | Euphonia_rufiventris | diurnal |
| birds | Euphonia_saturata | Euphonia_saturata | diurnal |
| birds | Euphonia_trinitatis | Euphonia_trinitatis | diurnal |
| birds | Euphonia_violacea | Euphonia_violacea | diurnal |
| birds | Euphonia_xanthogaster | Euphonia_xanthogaster | diurnal |
| birds | Euplectes_afer | Euplectes_afer | diurnal |
| birds | Euplectes_albonotatus | Euplectes_albonotatus | diurnal |
| birds | Euplectes_ardens | Euplectes_ardens | diurnal |
| birds | Euplectes_aureus | Euplectes_aureus | diurnal |
| birds | Euplectes_axillaris | Euplectes_axillaris | diurnal |
| birds | Euplectes_capensis | Euplectes_capensis | diurnal |
| birds | Euplectes_diadematus | Euplectes_diadematus | diurnal |
| birds | Euplectes_franciscanus | Euplectes_franciscanus | diurnal |
| birds | Euplectes_gierowii | Euplectes_gierowii | diurnal |
| birds | Euplectes_hartlaubi | Euplectes_hartlaubi | diurnal |
| birds | Euplectes_hordeaceus | Euplectes_hordeaceus | diurnal |
| birds | Euplectes_jacksoni | Euplectes_jacksoni | diurnal |
| birds | Euplectes_macroura | Euplectes_macroura | diurnal |
| birds | Euplectes_nigroventris | Euplectes_nigroventris | diurnal |
| birds | Euplectes_orix | Euplectes_orix | diurnal |
| birds | Euplectes_progne | Euplectes_progne | diurnal |
| birds | Euplectes_psammacromius | Euplectes_psammocromius | diurnal |

| | | | |
|---|---|---|---|
| birds | Eupodotis_caerulescens | Eupodotis_caerulescens | diurnal |
| birds | Eupodotis_senegalensis | Eupodotis_senegalensis | diurnal |
| birds | Eupsittula_aurea | Aratinga_aurea | diurnal |
| birds | Eupsittula_cactorum | Aratinga_cactorum | diurnal |
| birds | Eupsittula_canicularis | Aratinga_canicularis | diurnal |
| birds | Eupsittula_nana | Aratinga_nana | diurnal |
| birds | Eupsittula_pertinax | Aratinga_pertinax | diurnal |
| birds | Euptilotis_neoxenus | Euptilotis_neoxenus | diurnal |
| birds | Euptilotus_eutilotus | Pycnonotus_eutilotus | diurnal |
| birds | Eurillas_ansorgei | Andropadus_ansorgei | diurnal |
| birds | Eurillas_curvirostris | Andropadus_curvirostris | diurnal |
| birds | Eurillas_gracilis | Andropadus_gracilis | diurnal |
| birds | Eurillas_latirostris | Andropadus_latirostris | diurnal |
| birds | Eurillas_virens | Andropadus_virens | diurnal |
| birds | Eurocephalus_anguitimens | Eurocephalus_anguitimens | diurnal |
| birds | Eurocephalus_ruppelli | Eurocephalus_rueppelli | diurnal |
| birds | Eurochelidon_sirintarae | Eurochelidon_sirintarae | diurnal |
| birds | Eurostopodus_archboldi | Eurostopodus_archboldi | nocturnal |
| birds | Eurostopodus_argus | Eurostopodus_argus | nocturnal |
| birds | Eurostopodus_diabolicus | Eurostopodus_diabolicus | nocturnal |
| birds | Eurostopodus_mystacalis | Eurostopodus_mystacalis | nocturnal |
| birds | Eurostopodus_papuensis | Eurostopodus_papuensis | nocturnal |
| birds | Euryceros_prevostii | Euryceros_prevostii | diurnal |
| birds | Eurylaimus_javanicus | Eurylaimus_javanicus | diurnal |
| birds | Eurylaimus_ochromalus | Eurylaimus_ochromalus | diurnal |
| birds | Euryptila_subcinnamomea | Euryptila_subcinnamomea | diurnal |
| birds | Eurypyga_helias | Eurypyga_helias | diurnal |
| birds | Eurystomus_gularis | Eurystomus_gularis | diurnal |
| birds | Euscarthmus_meloryphus | Euscarthmus_meloryphus | diurnal |
| birds | Euscarthmus_rufomarginatus | Euscarthmus_rufomarginatus | diurnal |
| birds | Euschistospiza_cinereovinacea | Euschistospiza_cinereovinacea | diurnal |
| birds | Euschistospiza_dybowskii | Euschistospiza_dybowskii | diurnal |
| birds | Eutoxeres_aquila | Eutoxeres_aquila | diurnal |
| birds | Eutoxeres_condamini | Eutoxeres_condamini | diurnal |

| | | | |
|---|---|---|---|
| birds | Eutrichomyias_rowleyi | Eutrichomyias_rowleyi | diurnal |
| birds | Eutriorchis_astur | Eutriorchis_astur | diurnal |
| birds | Falcipennis_falcipennis | Dendragapus_falcipennis | diurnal |
| birds | Falco_alopex | Falco_alopex | diurnal |
| birds | Falco_araeus | Falco_araea | diurnal |
| birds | Falco_ardosiaceus | Falco_ardosiaceus | diurnal |
| birds | Falco_berigora | Falco_berigora | diurnal |
| birds | Falco_biarmicus | Falco_biarmicus | nocturnal |
| birds | Falco_cenchroides | Falco_cenchroides | diurnal |
| birds | Falco_cherrug | Falco_cherrug | diurnal |
| birds | Falco_chicquera | Falco_chicquera | diurnal |
| birds | Falco_columbarius | Falco_columbarius | diurnal |
| birds | Falco_deiroleucus | Falco_deiroleucus | diurnal |
| birds | Falco_dickinsoni | Falco_dickinsoni | diurnal |
| birds | Falco_eleonorae | Falco_eleonorae | nocturnal |
| birds | Falco_fasciinucha | Falco_fasciinucha | diurnal |
| birds | Falco_femoralis | Falco_femoralis | diurnal |
| birds | Falco_hypoleucos | Falco_hypoleucos | diurnal |
| birds | Falco_jugger | Falco_jugger | diurnal |
| birds | Falco_longipennis | Falco_longipennis | cathemeral |
| birds | Falco_mexicanus | Falco_mexicanus | diurnal |
| birds | Falco_moluccensis | Falco_moluccensis | diurnal |
| birds | Falco_naumanni | Falco_naumanni | diurnal |
| birds | Falco_newtoni | Falco_newtoni | diurnal |
| birds | Falco_novaeseelandiae | Falco_novaeseelandiae | diurnal |
| birds | Falco_peregrinus | Falco_peregrinus | diurnal |
| birds | Falco_punctatus | Falco_punctatus | diurnal |
| birds | Falco_rufigularis | Falco_rufigularis | nocturnal |
| birds | Falco_rupicoloides | Falco_rupicoloides | diurnal |
| birds | Falco_rusticolus | Falco_rusticolus | diurnal |
| birds | Falco_sparverius | Falco_sparverius | diurnal |
| birds | Falco_subniger | Falco_subniger | diurnal |
| birds | Falco_tinnunculus | Falco_tinnunculus | diurnal |
| birds | Falco_zoniventris | Falco_zoniventris | diurnal |

| | | | |
|---|---|---|---|
| birds | Falculea_palliata | Falculea_palliata | diurnal |
| birds | Falcunculus_frontatus | Falcunculus_frontatus | diurnal |
| birds | Ferminia_cerverai | Ferminia_cerverai | diurnal |
| birds | Ficedula_albicollis | Ficedula_albicollis | diurnal |
| birds | Ficedula_basilanica | Ficedula_basilanica | diurnal |
| birds | Ficedula_buruensis | Ficedula_buruensis | diurnal |
| birds | Ficedula_crypta | Ficedula_crypta | diurnal |
| birds | Ficedula_dumetoria | Ficedula_dumetoria | diurnal |
| birds | Ficedula_erithacus | Ficedula_hodgsonii | diurnal |
| birds | Ficedula_hodgsoni | Muscicapella_hodgsoni | diurnal |
| birds | Ficedula_hyperythra | Ficedula_hyperythra | diurnal |
| birds | Ficedula_hypoleuca | Ficedula_hypoleuca | diurnal |
| birds | Ficedula_mugimaki | Ficedula_mugimaki | diurnal |
| birds | Ficedula_narcissina | Ficedula_narcissina | diurnal |
| birds | Ficedula_nigrorufa | Ficedula_nigrorufa | diurnal |
| birds | Ficedula_parva | Ficedula_parva | diurnal |
| birds | Ficedula_platenae | Ficedula_platenae | diurnal |
| birds | Ficedula_ruficauda | Muscicapa_ruficauda | diurnal |
| birds | Ficedula_rufigula | Ficedula_rufigula | diurnal |
| birds | Ficedula_sapphira | Ficedula_sapphira | diurnal |
| birds | Ficedula_semitorquata | Ficedula_semitorquata | diurnal |
| birds | Ficedula_strophiata | Ficedula_strophiata | diurnal |
| birds | Ficedula_subrubra | Ficedula_subrubra | diurnal |
| birds | Ficedula_superciliaris | Ficedula_superciliaris | diurnal |
| birds | Ficedula_tricolor | Ficedula_tricolor | diurnal |
| birds | Ficedula_westermanni | Ficedula_westermanni | diurnal |
| birds | Ficedula_zanthopygia | Ficedula_zanthopygia | diurnal |
| birds | Florisuga_fusca | Florisuga_fusca | diurnal |
| birds | Florisuga_mellivora | Florisuga_mellivora | diurnal |
| birds | Fluvicola_albiventer | Fluvicola_albiventer | diurnal |
| birds | Fluvicola_nengeta | Fluvicola_nengeta | diurnal |
| birds | Fluvicola_pica | Fluvicola_pica | diurnal |
| birds | Formicarius_analis | Formicarius_analis | diurnal |
| birds | Formicarius_colma | Formicarius_colma | diurnal |

| | | | |
|---|---|---|---|
| birds | Formicarius_moniliger | Formicarius_moniliger | diurnal |
| birds | Formicarius_nigricapillus | Formicarius_nigricapillus | diurnal |
| birds | Formicarius_rufifrons | Formicarius_rufifrons | diurnal |
| birds | Formicarius_rufipectus | Formicarius_rufipectus | diurnal |
| birds | Formicivora_acutirostris | Stymphalornis_acutirostris | diurnal |
| birds | Formicivora_erythronotos | Formicivora_erythronotos | diurnal |
| birds | Formicivora_grisea | Formicivora_grisea | diurnal |
| birds | Formicivora_iheringi | Formicivora_iheringi | diurnal |
| birds | Formicivora_intermedia | Formicivora_intermedia | diurnal |
| birds | Formicivora_melanogaster | Formicivora_melanogaster | diurnal |
| birds | Formicivora_rufa | Formicivora_rufa | diurnal |
| birds | Formicivora_serrana | Formicivora_serrana | diurnal |
| birds | Forpus_coelestis | Forpus_coelestis | diurnal |
| birds | Forpus_conspicillatus | Forpus_conspicillatus | diurnal |
| birds | Forpus_cyanopygius | Forpus_cyanopygius | diurnal |
| birds | Forpus_modestus | Forpus_modestus | diurnal |
| birds | Forpus_passerinus | Forpus_passerinus | diurnal |
| birds | Forpus_xanthops | Forpus_xanthops | diurnal |
| birds | Forpus_xanthopterygius | Forpus_xanthopterygius | diurnal |
| birds | Foudia_eminentissima | Foudia_eminentissima | diurnal |
| birds | Foudia_flavicans | Foudia_flavicans | diurnal |
| birds | Foudia_madagascariensis | Foudia_madagascariensis | diurnal |
| birds | Foudia_omissa | Foudia_omissa | diurnal |
| birds | Foudia_rubra | Foudia_rubra | diurnal |
| birds | Foudia_sechellarum | Foudia_sechellarum | diurnal |
| birds | Foulehaio_carunculatus | Foulehaio_carunculatus | diurnal |
| birds | Francolinus_francolinus | Francolinus_francolinus | diurnal |
| birds | Francolinus_pictus | Francolinus_pictus | diurnal |
| birds | Francolinus_pintadeanus | Francolinus_pintadeanus | diurnal |
| birds | Fraseria_caerulescens | Muscicapa_caerulescens | diurnal |
| birds | Fraseria_cinerascens | Fraseria_cinerascens | diurnal |
| birds | Fraseria_griseigularis | Myioparus_griseigularis | diurnal |
| birds | Fraseria_lendu | Muscicapa_lendu | diurnal |
| birds | Fraseria_ocreata | Fraseria_ocreata | diurnal |

| | | | |
|---|---|---|---|
| birds | Fraseria_olivascens | Muscicapa_olivascens | diurnal |
| birds | Fraseria_plumbea | Myioparus_plumbeus | diurnal |
| birds | Fratercula_arctica | Fratercula_arctica | diurnal |
| birds | Fratercula_cirrhata | Fratercula_cirrhata | diurnal |
| birds | Fratercula_corniculata | Fratercula_corniculata | diurnal |
| birds | Frederickena_unduliger | Frederickena_unduligera | diurnal |
| birds | Frederickena_viridis | Frederickena_viridis | diurnal |
| birds | Fregata_andrewsi | Fregata_andrewsi | diurnal |
| birds | Fregata_aquila | Fregata_aquila | diurnal |
| birds | Fregata_ariel | Fregata_ariel | diurnal |
| birds | Fregata_magnificens | Fregata_magnificens | diurnal |
| birds | Fregata_minor | Fregata_minor | diurnal |
| birds | Fregetta_grallaria | Fregetta_grallaria | cathemeral |
| birds | Fregetta_tropica | Fregetta_tropica | cathemeral |
| birds | Fringilla_coelebs | Fringilla_coelebs | diurnal |
| birds | Fringilla_montifringilla | Fringilla_montifringilla | diurnal |
| birds | Fringilla_teydea | Fringilla_teydea | diurnal |
| birds | Fulica_alai | Fulica_alai | diurnal |
| birds | Fulica_ardesiaca | Fulica_ardesiaca | diurnal |
| birds | Fulica_armillata | Fulica_armillata | diurnal |
| birds | Fulica_atra | Fulica_atra | diurnal |
| birds | Fulica_cornuta | Fulica_cornuta | diurnal |
| birds | Fulica_cristata | Fulica_cristata | diurnal |
| birds | Fulica_gigantea | Fulica_gigantea | diurnal |
| birds | Fulica_leucoptera | Fulica_leucoptera | diurnal |
| birds | Fulica_rufifrons | Fulica_rufifrons | diurnal |
| birds | Fulmarus_glacialis | Fulmarus_glacialis | cathemeral |
| birds | Fulmarus_glacialoides | Fulmarus_glacialoides | cathemeral |
| birds | Fulvetta_cinereiceps | Alcippe_cinereiceps | diurnal |
| birds | Fulvetta_ludlowi | Alcippe_ludlowi | diurnal |
| birds | Fulvetta_ruficapilla | Alcippe_ruficapilla | diurnal |
| birds | Fulvetta_striaticollis | Alcippe_striaticollis | diurnal |
| birds | Fulvetta_vinipectus | Alcippe_vinipectus | diurnal |
| birds | Furnarius_cinnamomeus | Furnarius_cinnamomeus | diurnal |

| | | | |
|---|---|---|---|
| birds | Furnarius_cristatus | Furnarius_cristatus | diurnal |
| birds | Furnarius_figulus | Furnarius_figulus | diurnal |
| birds | Furnarius_leucopus | Furnarius_leucopus | diurnal |
| birds | Furnarius_longirostris | Furnarius_longirostris | diurnal |
| birds | Furnarius_minor | Furnarius_minor | diurnal |
| birds | Furnarius_rufus | Furnarius_rufus | diurnal |
| birds | Furnarius_torridus | Furnarius_torridus | diurnal |
| birds | Gactornis_enarratus | Caprimulgus_enarratus | nocturnal |
| birds | Galbalcyrhynchus_leucotis | Galbalcyrhynchus_leucotis | diurnal |
| birds | Galbalcyrhynchus_purusianus | Galbalcyrhynchus_purusianus | diurnal |
| birds | Galbula_albirostris | Galbula_albirostris | diurnal |
| birds | Galbula_chalcothorax | Galbula_chalcothorax | diurnal |
| birds | Galbula_cyanescens | Galbula_cyanescens | diurnal |
| birds | Galbula_cyanicollis | Galbula_cyanicollis | diurnal |
| birds | Galbula_dea | Galbula_dea | diurnal |
| birds | Galbula_galbula | Galbula_galbula | diurnal |
| birds | Galbula_leucogastra | Galbula_leucogastra | diurnal |
| birds | Galbula_pastazae | Galbula_pastazae | diurnal |
| birds | Galbula_ruficauda | Galbula_ruficauda | diurnal |
| birds | Galbula_tombacea | Galbula_tombacea | diurnal |
| birds | Galerida_cristata | Galerida_cristata | diurnal |
| birds | Galerida_deva | Galerida_deva | diurnal |
| birds | Galerida_magnirostris | Galerida_magnirostris | diurnal |
| birds | Galerida_modesta | Galerida_modesta | diurnal |
| birds | Galerida_theklae | Galerida_theklae | diurnal |
| birds | Gallicolumba_keayi | Gallicolumba_keayi | diurnal |
| birds | Gallicolumba_luzonica | Gallicolumba_luzonica | diurnal |
| birds | Gallicolumba_menagei | Gallicolumba_menagei | diurnal |
| birds | Gallicolumba_platenae | Gallicolumba_platenae | diurnal |
| birds | Gallicolumba_rufigula | Gallicolumba_rufigula | diurnal |
| birds | Gallicolumba_tristigmata | Gallicolumba_tristigmata | diurnal |
| birds | Gallinago_macrodactyla | Gallinago_macrodactyla | diurnal |
| birds | Gallinago_media | Gallinago_media | nocturnal |
| birds | Gallinago_megala | Gallinago_megala | nocturnal |

| | | | |
|---|---|---|---|
| birds | Gallinago_nemoricola | Gallinago_nemoricola | diurnal |
| birds | Gallinago_nigripennis | Gallinago_nigripennis | nocturnal |
| birds | Gallinago_paraguaiae | Gallinago_paraguaiae | diurnal |
| birds | Gallinago_solitaria | Gallinago_solitaria | diurnal |
| birds | Gallinago_stenura | Gallinago_stenura | cathemeral |
| birds | Gallinago_stricklandii | Gallinago_stricklandii | nocturnal |
| birds | Gallinago_undulata | Gallinago_undulata | nocturnal |
| birds | Gallinula_chloropus | Gallinula_chloropus | diurnal |
| birds | Gallinula_tenebrosa | Gallinula_tenebrosa | diurnal |
| birds | Gallirallus_australis | Gallirallus_australis | diurnal |
| birds | Gallirex_johnstoni | Ruwenzorornis_johnstoni | diurnal |
| birds | Gallirex_porphyreolophus | Tauraco_porphyreolophus | diurnal |
| birds | Galloperdix_bicalcarata | Galloperdix_bicalcarata | diurnal |
| birds | Galloperdix_lunulata | Galloperdix_lunulata | diurnal |
| birds | Galloperdix_spadicea | Galloperdix_spadicea | diurnal |
| birds | Gallus_gallus | Gallus_gallus | diurnal |
| birds | Gallus_lafayettii | Gallus_lafayetii | diurnal |
| birds | Gallus_sonneratii | Gallus_sonneratii | diurnal |
| birds | Gallus_varius | Gallus_varius | diurnal |
| birds | Gampsonyx_swainsonii | Gampsonyx_swainsonii | diurnal |
| birds | Gampsorhynchus_rufulus | Gampsorhynchus_rufulus | diurnal |
| birds | Garritornis_isidorei | Pomatostomus_isidorei | diurnal |
| birds | Garrodia_nereis | Garrodia_nereis | diurnal |
| birds | Garrulax_canorus | Garrulax_canorus | diurnal |
| birds | Garrulax_leucolophus | Garrulax_leucolophus | diurnal |
| birds | Garrulax_maesi | Garrulax_maesi | diurnal |
| birds | Garrulax_merulinus | Garrulax_merulinus | diurnal |
| birds | Garrulax_monileger | Garrulax_monileger | diurnal |
| birds | Garrulax_rufifrons | Garrulax_rufifrons | diurnal |
| birds | Garrulax_strepitans | Garrulax_strepitans | diurnal |
| birds | Garrulus_glandarius | Garrulus_glandarius | diurnal |
| birds | Garrulus_lanceolatus | Garrulus_lanceolatus | diurnal |
| birds | Garrulus_lidthi | Garrulus_lidthi | diurnal |
| birds | Gavia_adamsii | Gavia_adamsii | diurnal |

| | | | |
|---|---|---|---|
| birds | Gavia_arctica | Gavia_arctica | diurnal |
| birds | Gavia_immer | Gavia_immer | diurnal |
| birds | Gavia_stellata | Gavia_stellata | diurnal |
| birds | Gavicalis_fasciogularis | Lichenostomus_fasciogularis | diurnal |
| birds | Gavicalis_versicolor | Lichenostomus_versicolor | diurnal |
| birds | Gavicalis_virescens | Lichenostomus_virescens | diurnal |
| birds | Gecinulus_grantia | Gecinulus_grantia | diurnal |
| birds | Gecinulus_viridis | Gecinulus_viridis | diurnal |
| birds | Gelochelidon_nilotica | Sterna_nilotica | diurnal |
| birds | Gennaeodryas_placens | Poecilodryas_placens | diurnal |
| birds | Geobiastes_squamiger | Brachypteracias_squamiger | cathemeral |
| birds | Geocerthia_serrana | Upucerthia_serrana | diurnal |
| birds | Geococcyx_californianus | Geococcyx_californianus | diurnal |
| birds | Geococcyx_velox | Geococcyx_velox | diurnal |
| birds | Geoffroyus_geoffroyi | Geoffroyus_geoffroyi | diurnal |
| birds | Geoffroyus_heteroclitus | Geoffroyus_heteroclitus | diurnal |
| birds | Geoffroyus_simplex | Geoffroyus_simplex | diurnal |
| birds | Geokichla_camaronensis | Zoothera_camaronensis | diurnal |
| birds | Geokichla_cinerea | Zoothera_cinerea | diurnal |
| birds | Geokichla_citrina | Zoothera_citrina | diurnal |
| birds | Geokichla_crossleyi | Zoothera_crossleyi | diurnal |
| birds | Geokichla_dumasi | Zoothera_dumasi | diurnal |
| birds | Geokichla_erythronota | Zoothera_erythronota | diurnal |
| birds | Geokichla_gurneyi | Zoothera_gurneyi | diurnal |
| birds | Geokichla_guttata | Zoothera_guttata | diurnal |
| birds | Geokichla_interpres | Zoothera_interpres | diurnal |
| birds | Geokichla_oberlaenderi | Zoothera_oberlaenderi | diurnal |
| birds | Geokichla_piaggiae | Zoothera_piaggiae | diurnal |
| birds | Geokichla_princei | Zoothera_princei | diurnal |
| birds | Geokichla_sibirica | Zoothera_sibirica | diurnal |
| birds | Geokichla_spiloptera | Zoothera_spiloptera | diurnal |
| birds | Geokichla_wardii | Zoothera_wardii | diurnal |
| birds | Geopelia_cuneata | Geopelia_cuneata | diurnal |
| birds | Geopelia_humeralis | Geopelia_humeralis | diurnal |

| | | | |
|---|---|---|---|
| birds | Geopelia_placida | Geopelia_placida | diurnal |
| birds | Geopelia_striata | Geopelia_striata | diurnal |
| birds | Geophaps_plumifera | Geophaps_plumifera | diurnal |
| birds | Geophaps_scripta | Geophaps_scripta | diurnal |
| birds | Geophaps_smithii | Geophaps_smithii | diurnal |
| birds | Geositta_antarctica | Geositta_antarctica | diurnal |
| birds | Geositta_crassirostris | Geositta_crassirostris | diurnal |
| birds | Geositta_cunicularia | Geositta_cunicularia | diurnal |
| birds | Geositta_isabellina | Geositta_isabellina | diurnal |
| birds | Geositta_maritima | Geositta_maritima | diurnal |
| birds | Geositta_peruviana | Geositta_peruviana | diurnal |
| birds | Geositta_poeciloptera | Geositta_poeciloptera | diurnal |
| birds | Geositta_punensis | Geositta_punensis | diurnal |
| birds | Geositta_rufipennis | Geositta_rufipennis | diurnal |
| birds | Geositta_saxicolina | Geositta_saxicolina | diurnal |
| birds | Geositta_tenuirostris | Geositta_tenuirostris | diurnal |
| birds | Geospiza_conirostris | Geospiza_conirostris | diurnal |
| birds | Geospiza_difficilis | Geospiza_difficilis | diurnal |
| birds | Geospiza_fortis | Geospiza_fortis | diurnal |
| birds | Geospiza_fuliginosa | Geospiza_fuliginosa | diurnal |
| birds | Geospiza_heliobates | Camarhynchus_heliobates | diurnal |
| birds | Geospiza_magnirostris | Geospiza_magnirostris | diurnal |
| birds | Geospiza_pallida | Camarhynchus_pallidus | diurnal |
| birds | Geospiza_parvula | Camarhynchus_parvulus | diurnal |
| birds | Geospiza_pauper | Camarhynchus_pauper | diurnal |
| birds | Geospiza_psittacula | Camarhynchus_psittacula | diurnal |
| birds | Geospiza_scandens | Geospiza_scandens | diurnal |
| birds | Geospizopsis_plebejus | Phrygilus_plebejus | diurnal |
| birds | Geospizopsis_unicolor | Phrygilus_unicolor | diurnal |
| birds | Geothlypis_aequinoctialis | Geothlypis_aequinoctialis | diurnal |
| birds | Geothlypis_auricularis | Geothlypis_auricularis | diurnal |
| birds | Geothlypis_beldingi | Geothlypis_beldingi | diurnal |
| birds | Geothlypis_chiriquensis | Geothlypis_chiriquensis | diurnal |
| birds | Geothlypis_flavovelata | Geothlypis_flavovelata | diurnal |

| | | | |
|---|---|---|---|
| birds | Geothlypis_formosa | Oporornis_formosus | diurnal |
| birds | Geothlypis_nelsoni | Geothlypis_nelsoni | diurnal |
| birds | Geothlypis_philadelphia | Oporornis_philadelphia | diurnal |
| birds | Geothlypis_poliocephala | Geothlypis_poliocephala | diurnal |
| birds | Geothlypis_rostrata | Geothlypis_rostrata | diurnal |
| birds | Geothlypis_semiflava | Geothlypis_semiflava | diurnal |
| birds | Geothlypis_speciosa | Geothlypis_speciosa | diurnal |
| birds | Geothlypis_tolmiei | Oporornis_tolmiei | diurnal |
| birds | Geothlypis_trichas | Geothlypis_trichas | diurnal |
| birds | Geotrygon_caniceps | Geotrygon_caniceps | diurnal |
| birds | Geotrygon_chrysia | Geotrygon_chrysia | diurnal |
| birds | Geotrygon_montana | Geotrygon_montana | diurnal |
| birds | Geotrygon_mystacea | Geotrygon_mystacea | diurnal |
| birds | Geotrygon_saphirina | Geotrygon_saphirina | diurnal |
| birds | Geotrygon_versicolor | Geotrygon_versicolor | diurnal |
| birds | Geotrygon_violacea | Geotrygon_violacea | diurnal |
| birds | Geranoaetus_albicaudatus | Buteo_albicaudatus | diurnal |
| birds | Geranoaetus_melanoleucus | Geranoaetus_melanoleucus | diurnal |
| birds | Geranoaetus_polyosoma | Buteo_polyosoma | diurnal |
| birds | Geranospiza_caerulescens | Geranospiza_caerulescens | diurnal |
| birds | Geronticus_eremita | Geronticus_eremita | diurnal |
| birds | Gerygone_albofrontata | Gerygone_albofrontata | diurnal |
| birds | Gerygone_chloronota | Gerygone_chloronota | diurnal |
| birds | Gerygone_chrysogaster | Gerygone_chrysogaster | diurnal |
| birds | Gerygone_dorsalis | Gerygone_dorsalis | diurnal |
| birds | Gerygone_flavolateralis | Gerygone_flavolateralis | diurnal |
| birds | Gerygone_fusca | Gerygone_fusca | diurnal |
| birds | Gerygone_hypoxantha | Gerygone_hypoxantha | diurnal |
| birds | Gerygone_igata | Gerygone_igata | diurnal |
| birds | Gerygone_inornata | Gerygone_inornata | diurnal |
| birds | Gerygone_levigaster | Gerygone_levigaster | diurnal |
| birds | Gerygone_magnirostris | Gerygone_magnirostris | diurnal |
| birds | Gerygone_modesta | Gerygone_modesta | diurnal |
| birds | Gerygone_mouki | Gerygone_mouki | diurnal |

| | | | |
|---|---|---|---|
| birds | Gerygone_olivacea | Gerygone_olivacea | diurnal |
| birds | Gerygone_palpebrosa | Gerygone_palpebrosa | diurnal |
| birds | Gerygone_ruficollis | Gerygone_ruficollis | diurnal |
| birds | Gerygone_sulphurea | Gerygone_sulphurea | diurnal |
| birds | Gerygone_tenebrosa | Gerygone_tenebrosa | diurnal |
| birds | Glaucestrilda_coerulescens | Estrilda_caerulescens | diurnal |
| birds | Glaucestrilda_perreini | Estrilda_perreini | diurnal |
| birds | Glaucestrilda_thomensis | Estrilda_thomensis | diurnal |
| birds | Glaucidium_albertinum | Glaucidium_albertinum | nocturnal |
| birds | Glaucidium_bolivianum | Glaucidium_bolivianum | cathemeral |
| birds | Glaucidium_brasilianum | Glaucidium_brasilianum | diurnal |
| birds | Glaucidium_brodiei | Glaucidium_brodiei | diurnal |
| birds | Glaucidium_capense | Glaucidium_capense | diurnal |
| birds | Glaucidium_castaneum | Glaucidium_castaneum | diurnal |
| birds | Glaucidium_castanotum | Glaucidium_castanonotum | diurnal |
| birds | Glaucidium_cuculoides | Glaucidium_cuculoides | diurnal |
| birds | Glaucidium_gnoma | Glaucidium_gnoma | diurnal |
| birds | Glaucidium_griseiceps | Glaucidium_griseiceps | diurnal |
| birds | Glaucidium_hardyi | Glaucidium_hardyi | diurnal |
| birds | Glaucidium_jardinii | Glaucidium_jardinii | cathemeral |
| birds | Glaucidium_minutissimum | Glaucidium_minutissimum | cathemeral |
| birds | Glaucidium_nana | Glaucidium_nanum | cathemeral |
| birds | Glaucidium_perlatum | Glaucidium_perlatum | diurnal |
| birds | Glaucidium_sjostedti | Glaucidium_sjostedti | nocturnal |
| birds | Glaucidium_tephronotum | Glaucidium_tephronotum | nocturnal |
| birds | Glaucis_aeneus | Glaucis_aeneus | diurnal |
| birds | Glaucis_dohrnii | Glaucis_dohrnii | diurnal |
| birds | Glaucis_hirsutus | Glaucis_hirsutus | diurnal |
| birds | Gliciphila_melanops | Phylidonyris_melanops | diurnal |
| birds | Gliciphila_notabilis | Phylidonyris_notabilis | diurnal |
| birds | Gliciphila_undulata | Phylidonyris_undulatus | diurnal |
| birds | Glossopsitta_concinna | Glossopsitta_concinna | diurnal |
| birds | Glossopsitta_porphyrocephala | Glossopsitta_porphyrocephala | diurnal |
| birds | Glossopsitta_pusilla | Glossopsitta_pusilla | diurnal |

| | | | |
|---|---|---|---|
| birds | Glycichaera_fallax | Glycichaera_fallax | diurnal |
| birds | Glyphorynchus_spirurus | Glyphorynchus_spirurus | diurnal |
| birds | Gnorimopsar_chopi | Gnorimopsar_chopi | diurnal |
| birds | Goldmania_bella | Goethalsia_bella | diurnal |
| birds | Goldmania_violiceps | Goldmania_violiceps | diurnal |
| birds | Goodfellowia_miranda | Basilornis_mirandus | diurnal |
| birds | Gorsachius_goisagi | Gorsachius_goisagi | cathemeral |
| birds | Gorsachius_melanolophus | Gorsachius_melanolophus | cathemeral |
| birds | Goura_cristata | Goura_cristata | diurnal |
| birds | Goura_scheepmakeri | Goura_scheepmakeri | diurnal |
| birds | Goura_victoria | Goura_victoria | diurnal |
| birds | Gracula_ptilogenys | Gracula_ptilogenys | diurnal |
| birds | Gracula_religiosa | Gracula_religiosa | diurnal |
| birds | Gracupica_contra | Sturnus_contra | diurnal |
| birds | Gracupica_nigricollis | Sturnus_nigricollis | diurnal |
| birds | Grafisia_torquata | Grafisia_torquata | diurnal |
| birds | Grallaria_albigula | Grallaria_albigula | diurnal |
| birds | Grallaria_alleni | Grallaria_alleni | diurnal |
| birds | Grallaria_andicolus | Grallaria_andicolus | diurnal |
| birds | Grallaria_bangsi | Grallaria_bangsi | diurnal |
| birds | Grallaria_blakei | Grallaria_blakei | diurnal |
| birds | Grallaria_capitalis | Grallaria_capitalis | diurnal |
| birds | Grallaria_carrikeri | Grallaria_carrikeri | diurnal |
| birds | Grallaria_dignissima | Grallaria_dignissima | diurnal |
| birds | Grallaria_eludens | Grallaria_eludens | diurnal |
| birds | Grallaria_erythroleuca | Grallaria_erythroleuca | diurnal |
| birds | Grallaria_erythrotis | Grallaria_erythrotis | diurnal |
| birds | Grallaria_excelsa | Grallaria_excelsa | diurnal |
| birds | Grallaria_flavotincta | Grallaria_flavotincta | diurnal |
| birds | Grallaria_gigantea | Grallaria_gigantea | diurnal |
| birds | Grallaria_guatimalensis | Grallaria_guatimalensis | diurnal |
| birds | Grallaria_haplonota | Grallaria_haplonota | diurnal |
| birds | Grallaria_hypoleuca | Grallaria_hypoleuca | diurnal |
| birds | Grallaria_kaestneri | Grallaria_kaestneri | diurnal |

| | | | |
|---|---|---|---|
| birds | Grallaria_milleri | Grallaria_milleri | diurnal |
| birds | Grallaria_nuchalis | Grallaria_nuchalis | diurnal |
| birds | Grallaria_przewalskii | Grallaria_przewalskii | diurnal |
| birds | Grallaria_quitensis | Grallaria_quitensis | diurnal |
| birds | Grallaria_ruficapilla | Grallaria_ruficapilla | diurnal |
| birds | Grallaria_rufocinerea | Grallaria_rufocinerea | diurnal |
| birds | Grallaria_rufula | Grallaria_rufula | diurnal |
| birds | Grallaria_squamigera | Grallaria_squamigera | diurnal |
| birds | Grallaria_varia | Grallaria_varia | diurnal |
| birds | Grallaria_watkinsi | Grallaria_watkinsi | diurnal |
| birds | Grallaricula_cucullata | Grallaricula_cucullata | diurnal |
| birds | Grallaricula_ferrugineipectus | Grallaricula_ferrugineipectus | diurnal |
| birds | Grallaricula_flavirostris | Grallaricula_flavirostris | diurnal |
| birds | Grallaricula_lineifrons | Grallaricula_lineifrons | diurnal |
| birds | Grallaricula_nana | Grallaricula_nana | diurnal |
| birds | Grallaricula_ochraceifrons | Grallaricula_ochraceifrons | diurnal |
| birds | Grallaricula_peruviana | Grallaricula_peruviana | diurnal |
| birds | Grallina_bruijnii | Grallina_bruijni | diurnal |
| birds | Grallina_cyanoleuca | Grallina_cyanoleuca | diurnal |
| birds | Graminicola_bengalensis | Graminicola_bengalensis | diurnal |
| birds | Grammatoptila_striata | Garrulax_striatus | diurnal |
| birds | Granatellus_pelzelni | Granatellus_pelzelni | diurnal |
| birds | Granatellus_sallaei | Granatellus_sallaei | diurnal |
| birds | Granatellus_venustus | Granatellus_venustus | diurnal |
| birds | Granatina_granatina | Uraeginthus_granatinus | diurnal |
| birds | Granatina_ianthinogaster | Uraeginthus_ianthinogaster | diurnal |
| birds | Grandala_coelicolor | Grandala_coelicolor | diurnal |
| birds | Grantiella_picta | Grantiella_picta | diurnal |
| birds | Graueria_vittata | Graueria_vittata | diurnal |
| birds | Graydidascalus_brachyurus | Graydidascalus_brachyurus | diurnal |
| birds | Griseotyrannus_aurantioatrocristatus | Empidonomus_aurantioatrocristatus | diurnal |
| birds | Grus_americana | Grus_americana | diurnal |
| birds | Grus_antigone | Grus_antigone | diurnal |
| birds | Grus_canadensis | Grus_canadensis | diurnal |

| | | | |
|---|---|---|---|
| birds | Grus_grus | Grus_grus | diurnal |
| birds | Grus_japonensis | Grus_japonensis | diurnal |
| birds | Grus_monacha | Grus_monacha | diurnal |
| birds | Grus_nigricollis | Grus_nigricollis | diurnal |
| birds | Grus_rubicunda | Grus_rubicunda | diurnal |
| birds | Grus_vipio | Grus_vipio | diurnal |
| birds | Guadalcanaria_inexpectata | Guadalcanaria_inexpectata | diurnal |
| birds | Guaruba_guarouba | Guaruba_guarouba | diurnal |
| birds | Gubernatrix_cristata | Gubernatrix_cristata | diurnal |
| birds | Gubernetes_yetapa | Gubernetes_yetapa | diurnal |
| birds | Guira_guira | Guira_guira | diurnal |
| birds | Gulosus_aristotelis | Phalacrocorax_aristotelis | diurnal |
| birds | Guttera_plumifera | Guttera_plumifera | diurnal |
| birds | Guttera_pucherani | Guttera_pucherani | diurnal |
| birds | Guyramemua_affinis | Suiriri_islerorum | diurnal |
| birds | Gygis_alba | Gygis_alba | diurnal |
| birds | Gymnobucco_bonapartei | Gymnobucco_bonapartei | diurnal |
| birds | Gymnobucco_calvus | Gymnobucco_calvus | diurnal |
| birds | Gymnobucco_peli | Gymnobucco_peli | diurnal |
| birds | Gymnobucco_sladeni | Gymnobucco_sladeni | diurnal |
| birds | Gymnocichla_nudiceps | Gymnocichla_nudiceps | diurnal |
| birds | Gymnocrex_plumbeiventris | Gymnocrex_plumbeiventris | diurnal |
| birds | Gymnoderus_foetidus | Gymnoderus_foetidus | diurnal |
| birds | Gymnogyps_californianus | Gymnogyps_californianus | diurnal |
| birds | Gymnomystax_mexicanus | Gymnomystax_mexicanus | diurnal |
| birds | Gymnomyza_aubryana | Gymnomyza_aubryana | diurnal |
| birds | Gymnomyza_samoensis | Gymnomyza_samoensis | diurnal |
| birds | Gymnomyza_viridis | Gymnomyza_viridis | diurnal |
| birds | Gymnophaps_albertisii | Gymnophaps_albertisii | diurnal |
| birds | Gymnophaps_mada | Gymnophaps_mada | diurnal |
| birds | Gymnophaps_solomonensis | Gymnophaps_solomonensis | diurnal |
| birds | Gymnopithys_leucaspis | Gymnopithys_leucaspis | diurnal |
| birds | Gymnopithys_rufigula | Gymnopithys_rufigula | diurnal |
| birds | Gymnorhina_tibicen | Gymnorhina_tibicen | diurnal |

| | | | |
|---|---|---|---|
| birds | Gymnorhinus_cyanocephalus | Gymnorhinus_cyanocephalus | diurnal |
| birds | Gymnoris_dentata | Petronia_dentata | diurnal |
| birds | Gymnoris_pyrgita | Petronia_pyrgita | diurnal |
| birds | Gymnoris_superciliaris | Petronia_superciliaris | diurnal |
| birds | Gymnoris_xanthocollis | Petronia_xanthocollis | diurnal |
| birds | Gypaetus_barbatus | Gypaetus_barbatus | diurnal |
| birds | Gypohierax_angolensis | Gypohierax_angolensis | diurnal |
| birds | Gyps_africanus | Gyps_africanus | diurnal |
| birds | Gyps_bengalensis | Gyps_bengalensis | diurnal |
| birds | Gyps_coprotheres | Gyps_coprotheres | diurnal |
| birds | Gyps_fulvus | Gyps_fulvus | diurnal |
| birds | Gyps_himalayensis | Gyps_himalayensis | diurnal |
| birds | Gyps_indicus | Gyps_indicus | diurnal |
| birds | Gyps_rueppelli | Gyps_rueppellii | diurnal |
| birds | Gypsophila_brevicaudata | Napothera_brevicaudata | diurnal |
| birds | Gypsophila_crassa | Napothera_crassa | diurnal |
| birds | Gypsophila_rufipecta | Turdinus_rufipectus | diurnal |
| birds | Habia_atrimaxillaris | Habia_atrimaxillaris | diurnal |
| birds | Habia_carmioli | Chlorothraupis_carmioli | diurnal |
| birds | Habia_cristata | Habia_cristata | diurnal |
| birds | Habia_frenata | Chlorothraupis_frenata | diurnal |
| birds | Habia_fuscicauda | Habia_fuscicauda | diurnal |
| birds | Habia_gutturalis | Habia_gutturalis | diurnal |
| birds | Habia_olivacea | Chlorothraupis_olivacea | diurnal |
| birds | Habia_rubica | Habia_rubica | diurnal |
| birds | Habia_stolzmanni | Chlorothraupis_stolzmanni | diurnal |
| birds | Haematoderus_militaris | Haematoderus_militaris | diurnal |
| birds | Haematopus_ater | Haematopus_ater | diurnal |
| birds | Haematopus_chathamensis | Haematopus_chathamensis | diurnal |
| birds | Haematopus_fuliginosus | Haematopus_fuliginosus | diurnal |
| birds | Haematopus_leucopodus | Haematopus_leucopodus | diurnal |
| birds | Haematopus_longirostris | Haematopus_longirostris | diurnal |
| birds | Haematopus_moquini | Haematopus_moquini | diurnal |
| birds | Haematopus_ostralegus | Haematopus_ostralegus | diurnal |

| | | | |
|---|---|---|---|
| birds | Haematopus_palliatus | Haematopus_palliatus | diurnal |
| birds | Haematopus_unicolor | Haematopus_unicolor | diurnal |
| birds | Haematortyx_sanguiniceps | Haematortyx_sanguiniceps | diurnal |
| birds | Haemorhous_cassinii | Carpodacus_cassinii | diurnal |
| birds | Haemorhous_mexicanus | Carpodacus_mexicanus | diurnal |
| birds | Haemorhous_purpureus | Carpodacus_purpureus | diurnal |
| birds | Hafferia_fortis | Myrmeciza_fortis | diurnal |
| birds | Hafferia_immaculata | Myrmeciza_immaculata | diurnal |
| birds | Halcyon_albiventris | Halcyon_albiventris | diurnal |
| birds | Halcyon_badia | Halcyon_badia | diurnal |
| birds | Halcyon_chelicuti | Halcyon_chelicuti | diurnal |
| birds | Halcyon_coromanda | Halcyon_coromanda | diurnal |
| birds | Halcyon_cyanoventris | Halcyon_cyanoventris | diurnal |
| birds | Halcyon_leucocephala | Halcyon_leucocephala | diurnal |
| birds | Halcyon_malimbica | Halcyon_malimbica | diurnal |
| birds | Halcyon_pileata | Halcyon_pileata | diurnal |
| birds | Halcyon_senegalensis | Halcyon_senegalensis | diurnal |
| birds | Halcyon_senegaloides | Halcyon_senegaloides | diurnal |
| birds | Halcyon_smyrnensis | Halcyon_smyrnensis | diurnal |
| birds | Haliaeetus_albicilla | Haliaeetus_albicilla | diurnal |
| birds | Haliaeetus_leucocephalus | Haliaeetus_leucocephalus | diurnal |
| birds | Haliaeetus_leucogaster | Haliaeetus_leucogaster | diurnal |
| birds | Haliaeetus_leucoryphus | Haliaeetus_leucoryphus | diurnal |
| birds | Haliaeetus_pelagicus | Haliaeetus_pelagicus | diurnal |
| birds | Haliaeetus_sanfordi | Haliaeetus_sanfordi | diurnal |
| birds | Haliaeetus_vocifer | Haliaeetus_vocifer | diurnal |
| birds | Haliaeetus_vociferoides | Haliaeetus_vociferoides | diurnal |
| birds | Haliastur_indus | Haliastur_indus | diurnal |
| birds | Haliastur_sphenurus | Haliastur_sphenurus | diurnal |
| birds | Halobaena_caerulea | Halobaena_caerulea | cathemeral |
| birds | Hamirostra_melanosternon | Hamirostra_melanosternon | diurnal |
| birds | Hapalopsittaca_amazonina | Hapalopsittaca_amazonina | diurnal |
| birds | Hapalopsittaca_fuertesi | Hapalopsittaca_fuertesi | diurnal |
| birds | Hapalopsittaca_melanotis | Hapalopsittaca_melanotis | diurnal |

| | | | |
|---|---|---|---|
| birds | Hapalopsittaca_pyrrhops | Hapalopsittaca_pyrrhops | diurnal |
| birds | Hapaloptila_castanea | Hapaloptila_castanea | diurnal |
| birds | Haplophaedia_assimilis | Haplophaedia_assimilis | diurnal |
| birds | Haplophaedia_aureliae | Haplophaedia_aureliae | diurnal |
| birds | Haplophaedia_lugens | Haplophaedia_lugens | diurnal |
| birds | Haplospiza_unicolor | Haplospiza_unicolor | diurnal |
| birds | Harpactes_ardens | Harpactes_ardens | diurnal |
| birds | Harpactes_diardii | Harpactes_diardii | diurnal |
| birds | Harpactes_duvaucelii | Harpactes_duvaucelii | diurnal |
| birds | Harpactes_erythrocephalus | Harpactes_erythrocephalus | diurnal |
| birds | Harpactes_kasumba | Harpactes_kasumba | diurnal |
| birds | Harpactes_oreskios | Harpactes_oreskios | diurnal |
| birds | Harpactes_orrhophaeus | Harpactes_orrhophaeus | diurnal |
| birds | Harpactes_wardi | Harpactes_wardi | diurnal |
| birds | Harpactes_whiteheadi | Harpactes_whiteheadi | diurnal |
| birds | Harpagus_bidentatus | Harpagus_bidentatus | diurnal |
| birds | Harpagus_diodon | Harpagus_diodon | diurnal |
| birds | Harpia_harpyja | Harpia_harpyja | diurnal |
| birds | Harpyopsis_novaeguineae | Harpyopsis_novaeguineae | diurnal |
| birds | Hartertula_flavoviridis | Neomixis_flavoviridis | diurnal |
| birds | Hartlaubius_auratus | Saroglossa_aurata | diurnal |
| birds | Hedydipna_collaris | Anthreptes_collaris | diurnal |
| birds | Hedydipna_metallica | Anthreptes_metallicus | diurnal |
| birds | Hedydipna_pallidigaster | Anthreptes_pallidigaster | diurnal |
| birds | Hedydipna_platura | Anthreptes_platurus | diurnal |
| birds | Heleia_crassirostris | Heleia_crassirostris | diurnal |
| birds | Heleia_goodfellowi | Lophozosterops_goodfellowi | diurnal |
| birds | Heleia_muelleri | Heleia_muelleri | diurnal |
| birds | Heleia_pinaiae | Lophozosterops_pinaiae | diurnal |
| birds | Heleia_squamiceps | Lophozosterops_squamiceps | diurnal |
| birds | Heleia_squamifrons | Oculocincta_squamifrons | diurnal |
| birds | Heleia_wallacei | Zosterops_wallacei | diurnal |
| birds | Heliactin_bilophus | Heliactin_bilophus | diurnal |
| birds | Heliangelus_amethysticollis | Heliangelus_amethysticollis | diurnal |

| | | | |
|---|---|---|---|
| birds | Heliangelus_exortis | Heliangelus_exortis | diurnal |
| birds | Heliangelus_mavors | Heliangelus_mavors | diurnal |
| birds | Heliangelus_micraster | Heliangelus_micraster | diurnal |
| birds | Heliangelus_regalis | Heliangelus_regalis | diurnal |
| birds | Heliangelus_strophianus | Heliangelus_strophianus | diurnal |
| birds | Heliangelus_viola | Heliangelus_viola | diurnal |
| birds | Helicolestes_hamatus | Helicolestes_hamatus | diurnal |
| birds | Heliobletus_contaminatus | Heliobletus_contaminatus | diurnal |
| birds | Heliodoxa_aurescens | Heliodoxa_aurescens | diurnal |
| birds | Heliodoxa_branickii | Heliodoxa_branickii | diurnal |
| birds | Heliodoxa_gularis | Heliodoxa_gularis | diurnal |
| birds | Heliodoxa_imperatrix | Heliodoxa_imperatrix | diurnal |
| birds | Heliodoxa_jacula | Heliodoxa_jacula | diurnal |
| birds | Heliodoxa_leadbeateri | Heliodoxa_leadbeateri | diurnal |
| birds | Heliodoxa_rubinoides | Heliodoxa_rubinoides | diurnal |
| birds | Heliodoxa_schreibersii | Heliodoxa_schreibersii | diurnal |
| birds | Heliodoxa_xanthogonys | Heliodoxa_xanthogonys | diurnal |
| birds | Heliomaster_constantii | Heliomaster_constantii | diurnal |
| birds | Heliomaster_furcifer | Heliomaster_furcifer | diurnal |
| birds | Heliomaster_longirostris | Heliomaster_longirostris | diurnal |
| birds | Heliomaster_squamosus | Heliomaster_squamosus | diurnal |
| birds | Heliopais_personatus | Heliopais_personatus | diurnal |
| birds | Heliornis_fulica | Heliornis_fulica | diurnal |
| birds | Heliothryx_auritus | Heliothryx_auritus | diurnal |
| birds | Heliothryx_barroti | Heliothryx_barroti | diurnal |
| birds | Hellmayrea_gularis | Hellmayrea_gularis | diurnal |
| birds | Helmitheros_vermivorum | Helmitheros_vermivorum | diurnal |
| birds | Helopsaltes_certhiola | Locustella_certhiola | diurnal |
| birds | Helopsaltes_ochotensis | Locustella_ochotensis | diurnal |
| birds | Helopsaltes_pleskei | Locustella_pleskei | diurnal |
| birds | Helopsaltes_pryeri | Locustella_pryeri | diurnal |
| birds | Hemicircus_canente | Hemicircus_canente | diurnal |
| birds | Hemicircus_concretus | Hemicircus_concretus | diurnal |
| birds | Hemignathus_wilsoni | Hemignathus_munroi | diurnal |

| | | | |
|---|---|---|---|
| birds | Hemimacronyx_chloris | Anthus_chloris | diurnal |
| birds | Hemiphaga_novaeseelandiae | Hemiphaga_novaeseelandiae | diurnal |
| birds | Hemiprocne_comata | Hemiprocne_comata | diurnal |
| birds | Hemiprocne_coronata | Hemiprocne_coronata | diurnal |
| birds | Hemiprocne_longipennis | Hemiprocne_longipennis | diurnal |
| birds | Hemiprocne_mystacea | Hemiprocne_mystacea | nocturnal |
| birds | Hemipus_hirundinaceus | Hemipus_hirundinaceus | diurnal |
| birds | Hemipus_picatus | Hemipus_picatus | diurnal |
| birds | Hemitesia_neumanni | Hemitesia_neumanni | diurnal |
| birds | Hemitesia_pallidipes | Cettia_pallidipes | diurnal |
| birds | Hemithraupis_flavicollis | Hemithraupis_flavicollis | diurnal |
| birds | Hemithraupis_guira | Hemithraupis_guira | diurnal |
| birds | Hemithraupis_ruficapilla | Hemithraupis_ruficapilla | diurnal |
| birds | Hemitriccus_cinnamomeipectus | Hemitriccus_cinnamomeipectus | diurnal |
| birds | Hemitriccus_diops | Hemitriccus_diops | diurnal |
| birds | Hemitriccus_flammulatus | Hemitriccus_flammulatus | diurnal |
| birds | Hemitriccus_granadensis | Hemitriccus_granadensis | diurnal |
| birds | Hemitriccus_griseipectus | Hemitriccus_griseipectus | diurnal |
| birds | Hemitriccus_inornatus | Hemitriccus_inornatus | diurnal |
| birds | Hemitriccus_iohannis | Hemitriccus_iohannis | diurnal |
| birds | Hemitriccus_josephinae | Hemitriccus_josephinae | diurnal |
| birds | Hemitriccus_kaempferi | Hemitriccus_kaempferi | diurnal |
| birds | Hemitriccus_margaritaceiventer | Hemitriccus_margaritaceiventer | diurnal |
| birds | Hemitriccus_minimus | Hemitriccus_minimus | diurnal |
| birds | Hemitriccus_minor | Hemitriccus_minor | diurnal |
| birds | Hemitriccus_nidipendulus | Hemitriccus_nidipendulus | diurnal |
| birds | Hemitriccus_obsoletus | Hemitriccus_obsoletus | diurnal |
| birds | Hemitriccus_orbitatus | Hemitriccus_orbitatus | diurnal |
| birds | Hemitriccus_rufigularis | Hemitriccus_rufigularis | diurnal |
| birds | Hemitriccus_spodiops | Hemitriccus_spodiops | diurnal |
| birds | Hemitriccus_striaticollis | Hemitriccus_striaticollis | diurnal |
| birds | Hemitriccus_zosterops | Hemitriccus_zosterops | diurnal |
| birds | Hemixos_castanonotus | Hemixos_castanonotus | diurnal |
| birds | Hemixos_flavala | Hemixos_flavala | diurnal |

| | | | |
|---|---|---|---|
| birds | Hemixos_leucogrammicus | Pycnonotus_leucogrammicus | diurnal |
| birds | Henicopernis_longicauda | Henicopernis_longicauda | diurnal |
| birds | Henicophaps_albifrons | Henicophaps_albifrons | diurnal |
| birds | Henicophaps_foersteri | Henicophaps_foersteri | diurnal |
| birds | Henicorhina_leucophrys | Henicorhina_leucophrys | diurnal |
| birds | Henicorhina_leucoptera | Henicorhina_leucoptera | diurnal |
| birds | Henicorhina_leucosticta | Henicorhina_leucosticta | diurnal |
| birds | Herpetotheres_cachinnans | Herpetotheres_cachinnans | diurnal |
| birds | Herpsilochmus_atricapillus | Herpsilochmus_atricapillus | diurnal |
| birds | Herpsilochmus_axillaris | Herpsilochmus_axillaris | diurnal |
| birds | Herpsilochmus_dorsimaculatus | Herpsilochmus_dorsimaculatus | diurnal |
| birds | Herpsilochmus_dugandi | Herpsilochmus_dugandi | diurnal |
| birds | Herpsilochmus_gentryi | Herpsilochmus_gentryi | diurnal |
| birds | Herpsilochmus_longirostris | Herpsilochmus_longirostris | diurnal |
| birds | Herpsilochmus_motacilloides | Herpsilochmus_motacilloides | diurnal |
| birds | Herpsilochmus_parkeri | Herpsilochmus_parkeri | diurnal |
| birds | Herpsilochmus_pectoralis | Herpsilochmus_pectoralis | diurnal |
| birds | Herpsilochmus_pileatus | Herpsilochmus_pileatus | diurnal |
| birds | Herpsilochmus_roraimae | Herpsilochmus_roraimae | diurnal |
| birds | Herpsilochmus_rufimarginatus | Herpsilochmus_rufimarginatus | diurnal |
| birds | Herpsilochmus_sellowi | Herpsilochmus_sellowi | diurnal |
| birds | Herpsilochmus_stictocephalus | Herpsilochmus_stictocephalus | diurnal |
| birds | Herpsilochmus_sticturus | Herpsilochmus_sticturus | diurnal |
| birds | Hesperiphona_abeillei | Coccothraustes_abeillei | diurnal |
| birds | Hesperiphona_vespertina | Coccothraustes_vespertinus | diurnal |
| birds | Heterocercus_aurantiivertex | Heterocercus_aurantiivertex | diurnal |
| birds | Heterocercus_flavivertex | Heterocercus_flavivertex | diurnal |
| birds | Heterocercus_linteatus | Heterocercus_linteatus | diurnal |
| birds | Heteromirafra_archeri | Heteromirafra_archeri | diurnal |
| birds | Heteromirafra_ruddi | Heteromirafra_ruddi | diurnal |
| birds | Heteromunia_pectoralis | Heteromunia_pectoralis | diurnal |
| birds | Heteromyias_albispecularis | Heteromyias_albispecularis | diurnal |
| birds | Heteronetta_atricapilla | Heteronetta_atricapilla | diurnal |
| birds | Heterophasia_auricularis | Heterophasia_auricularis | diurnal |

| | | | |
|---|---|---|---|
| birds | Heterophasia_capistrata | Heterophasia_capistrata | diurnal |
| birds | Heterophasia_gracilis | Heterophasia_gracilis | diurnal |
| birds | Heterophasia_melanoleuca | Heterophasia_melanoleuca | diurnal |
| birds | Heterophasia_picaoides | Heterophasia_picaoides | diurnal |
| birds | Heterophasia_pulchella | Heterophasia_pulchella | diurnal |
| birds | Heteroscenes_pallidus | Cuculus_pallidus | diurnal |
| birds | Heterospingus_rubrifrons | Heterospingus_rubrifrons | diurnal |
| birds | Heterospingus_xanthopygius | Heterospingus_xanthopygius | diurnal |
| birds | Heterotetrax_humilis | Eupodotis_humilis | diurnal |
| birds | Heterotetrax_rueppelii | Eupodotis_rueppellii | diurnal |
| birds | Heterotetrax_vigorsii | Eupodotis_vigorsii | diurnal |
| birds | Heteroxenicus_stellatus | Brachypteryx_stellata | diurnal |
| birds | Hieraaetus_ayresii | Hieraaetus_ayresii | diurnal |
| birds | Hieraaetus_morphnoides | Hieraaetus_morphnoides | diurnal |
| birds | Hieraaetus_pennatus | Hieraaetus_pennatus | diurnal |
| birds | Hieraaetus_wahlbergi | Aquila_wahlbergi | diurnal |
| birds | Hierococcyx_fugax | Cuculus_fugax | diurnal |
| birds | Hierococcyx_sparverioides | Cuculus_sparverioides | diurnal |
| birds | Hierococcyx_vagans | Cuculus_vagans | diurnal |
| birds | Hierococcyx_varius | Cuculus_varius | diurnal |
| birds | Himalayapsitta_cyanocephala | Psittacula_cyanocephala | diurnal |
| birds | Himalayapsitta_finschii | Psittacula_finschii | diurnal |
| birds | Himalayapsitta_himalayana | Psittacula_himalayana | diurnal |
| birds | Himantopus_himantopus | Himantopus_himantopus | cathemeral |
| birds | Himantopus_novaezelandiae | Himantopus_novaezelandiae | diurnal |
| birds | Himantornis_haematopus | Himantornis_haematopus | diurnal |
| birds | Himatione_sanguinea | Himatione_sanguinea | diurnal |
| birds | Hippolais_icterina | Hippolais_icterina | diurnal |
| birds | Hippolais_languida | Hippolais_languida | diurnal |
| birds | Hippolais_olivetorum | Hippolais_olivetorum | diurnal |
| birds | Hippolais_polyglotta | Hippolais_polyglotta | diurnal |
| birds | Hirundapus_caudacutus | Hirundapus_caudacutus | diurnal |
| birds | Hirundapus_celebensis | Hirundapus_celebensis | diurnal |
| birds | Hirundapus_cochinchinensis | Hirundapus_cochinchinensis | diurnal |

| | | | |
|---|---|---|---|
| birds | Hirundapus_giganteus | Hirundapus_giganteus | diurnal |
| birds | Hirundinea_ferruginea | Hirundinea_ferruginea | diurnal |
| birds | Hirundo_aethiopica | Hirundo_aethiopica | diurnal |
| birds | Hirundo_albigularis | Hirundo_albigularis | diurnal |
| birds | Hirundo_angolensis | Hirundo_angolensis | diurnal |
| birds | Hirundo_atrocaerulea | Hirundo_atrocaerulea | diurnal |
| birds | Hirundo_dimidiata | Hirundo_dimidiata | diurnal |
| birds | Hirundo_javanica | Hirundo_domicola | diurnal |
| birds | Hirundo_leucosoma | Hirundo_leucosoma | diurnal |
| birds | Hirundo_lucida | Hirundo_lucida | diurnal |
| birds | Hirundo_megaensis | Hirundo_megaensis | diurnal |
| birds | Hirundo_neoxena | Hirundo_neoxena | cathemeral |
| birds | Hirundo_nigrita | Hirundo_nigrita | diurnal |
| birds | Hirundo_nigrorufa | Hirundo_nigrorufa | diurnal |
| birds | Hirundo_rustica | Hirundo_rustica | diurnal |
| birds | Hirundo_smithii | Hirundo_smithii | diurnal |
| birds | Hirundo_tahitica | Hirundo_tahitica | diurnal |
| birds | Histrionicus_histrionicus | Histrionicus_histrionicus | diurnal |
| birds | Histurgops_ruficauda | Histurgops_ruficaudus | diurnal |
| birds | Hoploxypterus_cayanus | Vanellus_cayanus | diurnal |
| birds | Horizocerus_albocristatus | Tropicranus_albocristatus | diurnal |
| birds | Horizocerus_hartlaubi | Tockus_hartlaubi | diurnal |
| birds | Horornis_acanthizoides | Cettia_acanthizoides | diurnal |
| birds | Horornis_annae | Cettia_annae | diurnal |
| birds | Horornis_canturians | Cettia_canturians | diurnal |
| birds | Horornis_carolinae | Cettia_carolinae | diurnal |
| birds | Horornis_diphone | Cettia_diphone | diurnal |
| birds | Horornis_flavolivaceus | Cettia_flavolivacea | diurnal |
| birds | Horornis_fortipes | Cettia_fortipes | diurnal |
| birds | Horornis_parens | Cettia_parens | diurnal |
| birds | Horornis_ruficapilla | Cettia_ruficapilla | diurnal |
| birds | Horornis_seebohmi | Cettia_seebohmi | diurnal |
| birds | Houbaropsis_bengalensis | Houbaropsis_bengalensis | diurnal |
| birds | Hydrobates_castro | Oceanodroma_castro | cathemeral |

| | | | |
|---|---|---|---|
| birds | Hydrobates_furcatus | Oceanodroma_furcata | cathemeral |
| birds | Hydrobates_homochroa | Oceanodroma_homochroa | diurnal |
| birds | Hydrobates_leucorhous | Oceanodroma_leucorhoa | cathemeral |
| birds | Hydrobates_matsudairae | Oceanodroma_matsudairae | diurnal |
| birds | Hydrobates_melania | Oceanodroma_melania | diurnal |
| birds | Hydrobates_microsoma | Halocyptena_microsoma | cathemeral |
| birds | Hydrobates_pelagicus | Hydrobates_pelagicus | cathemeral |
| birds | Hydrobates_tethys | Oceanodroma_tethys | cathemeral |
| birds | Hydrobates_tristrami | Oceanodroma_tristrami | cathemeral |
| birds | Hydrocoloeus_minutus | Larus_minutus | diurnal |
| birds | Hydrophasianus_chirurgus | Hydrophasianus_chirurgus | diurnal |
| birds | Hydroprogne_caspia | Sterna_caspia | diurnal |
| birds | Hydropsalis_cayennensis | Caprimulgus_cayennensis | nocturnal |
| birds | Hydropsalis_climacocerca | Hydropsalis_climacocerca | nocturnal |
| birds | Hydropsalis_maculicaudus | Caprimulgus_maculicaudus | nocturnal |
| birds | Hydropsalis_torquata | Hydropsalis_torquata | nocturnal |
| birds | Hydrornis_baudii | Pitta_baudii | diurnal |
| birds | Hydrornis_caeruleus | Pitta_caerulea | diurnal |
| birds | Hydrornis_cyaneus | Pitta_cyanea | diurnal |
| birds | Hydrornis_elliotii | Pitta_elliotii | diurnal |
| birds | Hydrornis_guajanus | Pitta_guajana | diurnal |
| birds | Hydrornis_gurneyi | Pitta_gurneyi | diurnal |
| birds | Hydrornis_nipalensis | Pitta_nipalensis | diurnal |
| birds | Hydrornis_oatesi | Pitta_oatesi | diurnal |
| birds | Hydrornis_phayrei | Pitta_phayrei | diurnal |
| birds | Hydrornis_soror | Pitta_soror | diurnal |
| birds | Hylacola_cauta | Hylacola_cauta | diurnal |
| birds | Hylacola_pyrrhopygia | Hylacola_pyrrhopygia | diurnal |
| birds | Hylatomus_lineatus | Dryocopus_lineatus | diurnal |
| birds | Hylatomus_pileatus | Dryocopus_pileatus | diurnal |
| birds | Hylatomus_schulzii | Dryocopus_schulzi | diurnal |
| birds | Hylexetastes_perrotii | Hylexetastes_perrotii | diurnal |
| birds | Hylexetastes_stresemanni | Hylexetastes_stresemanni | diurnal |
| birds | Hylia_prasina | Hylia_prasina | diurnal |

| | | | |
|---|---|---|---|
| birds | Hyliota_australis | Hyliota_australis | diurnal |
| birds | Hyliota_flavigaster | Hyliota_flavigaster | diurnal |
| birds | Hyliota_violacea | Hyliota_violacea | diurnal |
| birds | Hylocharis_chrysura | Hylocharis_chrysura | diurnal |
| birds | Hylocharis_sapphirina | Hylocharis_sapphirina | diurnal |
| birds | Hylocichla_mustelina | Hylocichla_mustelina | diurnal |
| birds | Hylocitrea_bonensis | Hylocitrea_bonensis | diurnal |
| birds | Hylomanes_momotula | Hylomanes_momotula | diurnal |
| birds | Hylonympha_macrocerca | Hylonympha_macrocerca | diurnal |
| birds | Hylopezus_auricularis | Hylopezus_auricularis | diurnal |
| birds | Hylopezus_macularius | Hylopezus_macularius | diurnal |
| birds | Hylopezus_ochroleucus | Hylopezus_ochroleucus | diurnal |
| birds | Hylopezus_perspicillatus | Hylopezus_perspicillatus | diurnal |
| birds | Hylophilus_amaurocephalus | Hylophilus_amaurocephalus | diurnal |
| birds | Hylophilus_brunneiceps | Hylophilus_brunneiceps | diurnal |
| birds | Hylophilus_flavipes | Hylophilus_flavipes | diurnal |
| birds | Hylophilus_olivaceus | Hylophilus_olivaceus | diurnal |
| birds | Hylophilus_pectoralis | Hylophilus_pectoralis | diurnal |
| birds | Hylophilus_poicilotis | Hylophilus_poicilotis | diurnal |
| birds | Hylophilus_semicinereus | Hylophilus_semicinereus | diurnal |
| birds | Hylophilus_thoracicus | Hylophilus_thoracicus | diurnal |
| birds | Hylophylax_naevioides | Hylophylax_naevioides | diurnal |
| birds | Hylophylax_naevius | Hylophylax_naevius | diurnal |
| birds | Hylophylax_punctulatus | Hylophylax_punctulatus | diurnal |
| birds | Hylopsar_cupreocauda | Lamprotornis_cupreocauda | diurnal |
| birds | Hylopsar_purpureiceps | Lamprotornis_purpureiceps | diurnal |
| birds | Hylorchilus_navai | Hylorchilus_navai | diurnal |
| birds | Hylorchilus_sumichrasti | Hylorchilus_sumichrasti | diurnal |
| birds | Hymenolaimus_malacorhynchos | Hymenolaimus_malacorhynchos | diurnal |
| birds | Hymenops_perspicillatus | Hymenops_perspicillatus | diurnal |
| birds | Hypargos_margaritatus | Hypargos_margaritatus | diurnal |
| birds | Hypargos_niveoguttatus | Hypargos_niveoguttatus | diurnal |
| birds | Hypergerus_atriceps | Hypergerus_atriceps | diurnal |
| birds | Hypnelus_ruficollis | Hypnelus_ruficollis | diurnal |

| | | | |
|---|---|---|---|
| birds | Hypocnemis_cantator | Hypocnemis_cantator | diurnal |
| birds | Hypocnemis_hypoxantha | Hypocnemis_hypoxantha | diurnal |
| birds | Hypocnemoides_maculicauda | Hypocnemoides_maculicauda | diurnal |
| birds | Hypocnemoides_melanopogon | Hypocnemoides_melanopogon | diurnal |
| birds | Hypocolius_ampelinus | Hypocolius_ampelinus | diurnal |
| birds | Hypocryptadius_cinnamomeus | Hypocryptadius_cinnamomeus | diurnal |
| birds | Hypoedaleus_guttatus | Hypoedaleus_guttatus | diurnal |
| birds | Hypopyrrhus_pyrohypogaster | Hypopyrrhus_pyrohypogaster | diurnal |
| birds | Hypositta_corallirostris | Hypositta_corallirostris | diurnal |
| birds | Hypotaenidia_insignis | Gallirallus_insignis | diurnal |
| birds | Hypotaenidia_okinawae | Gallirallus_okinawae | diurnal |
| birds | Hypotaenidia_owstoni | Gallirallus_owstoni | nocturnal |
| birds | Hypotaenidia_philippensis | Gallirallus_philippensis | cathemeral |
| birds | Hypotaenidia_rovianae | Gallirallus_rovianae | diurnal |
| birds | Hypotaenidia_sylvestris | Gallirallus_sylvestris | diurnal |
| birds | Hypotaenidia_torquata | Gallirallus_torquatus | diurnal |
| birds | Hypotaenidia_woodfordi | Nesoclopeus_woodfordi | diurnal |
| birds | Hypothymis_azurea | Hypothymis_azurea | diurnal |
| birds | Hypothymis_coelestis | Hypothymis_coelestis | diurnal |
| birds | Hypothymis_helenae | Hypothymis_helenae | diurnal |
| birds | Hypsipetes_affinis | Alophoixus_affinis | diurnal |
| birds | Hypsipetes_amaurotis | Ixos_amaurotis | diurnal |
| birds | Hypsipetes_borbonicus | Hypsipetes_borbonicus | diurnal |
| birds | Hypsipetes_crassirostris | Hypsipetes_crassirostris | diurnal |
| birds | Hypsipetes_everetti | Ixos_everetti | diurnal |
| birds | Hypsipetes_leucocephalus | Hypsipetes_leucocephalus | diurnal |
| birds | Hypsipetes_madagascariensis | Hypsipetes_madagascariensis | diurnal |
| birds | Hypsipetes_olivaceus | Hypsipetes_olivaceus | diurnal |
| birds | Hypsipetes_parvirostris | Hypsipetes_parvirostris | diurnal |
| birds | Hypsipetes_philippinus | Ixos_philippinus | diurnal |
| birds | Hypsipetes_rufigularis | Ixos_rufigularis | diurnal |
| birds | Hypsipetes_thompsoni | Hypsipetes_thompsoni | diurnal |
| birds | Ianthocincla_bieti | Garrulax_bieti | diurnal |
| birds | Ianthocincla_lunulata | Garrulax_lunulatus | diurnal |

| | | | |
|---|---|---|---|
| birds | Ianthocincla_maxima | Garrulax_maximus | diurnal |
| birds | Ianthocincla_ocellata | Garrulax_ocellatus | diurnal |
| birds | Ianthocincla_rufogularis | Garrulax_rufogularis | diurnal |
| birds | Ianthocincla_sukatschewi | Garrulax_sukatschewi | diurnal |
| birds | Ibidorhyncha_struthersii | Ibidorhyncha_struthersii | diurnal |
| birds | Ibycter_americanus | Ibycter_americanus | diurnal |
| birds | Icteria_virens | Icteria_virens | diurnal |
| birds | Icterus_abeillei | Icterus_abeillei | diurnal |
| birds | Icterus_auratus | Icterus_auratus | diurnal |
| birds | Icterus_auricapillus | Icterus_auricapillus | diurnal |
| birds | Icterus_bonana | Icterus_bonana | diurnal |
| birds | Icterus_bullockiorum | Icterus_bullockii | diurnal |
| birds | Icterus_cayanensis | Icterus_cayanensis | diurnal |
| birds | Icterus_chrysater | Icterus_chrysater | diurnal |
| birds | Icterus_croconotus | Icterus_croconotus | diurnal |
| birds | Icterus_cucullatus | Icterus_cucullatus | diurnal |
| birds | Icterus_dominicensis | Icterus_dominicensis | diurnal |
| birds | Icterus_galbula | Icterus_galbula | diurnal |
| birds | Icterus_graceannae | Icterus_graceannae | diurnal |
| birds | Icterus_graduacauda | Icterus_graduacauda | diurnal |
| birds | Icterus_gularis | Icterus_gularis | diurnal |
| birds | Icterus_icterus | Icterus_icterus | diurnal |
| birds | Icterus_jamacaii | Icterus_jamacaii | diurnal |
| birds | Icterus_laudabilis | Icterus_laudabilis | diurnal |
| birds | Icterus_leucopteryx | Icterus_leucopteryx | diurnal |
| birds | Icterus_maculialatus | Icterus_maculialatus | diurnal |
| birds | Icterus_mesomelas | Icterus_mesomelas | diurnal |
| birds | Icterus_nigrogularis | Icterus_nigrogularis | diurnal |
| birds | Icterus_oberi | Icterus_oberi | diurnal |
| birds | Icterus_parisorum | Icterus_parisorum | diurnal |
| birds | Icterus_pectoralis | Icterus_pectoralis | diurnal |
| birds | Icterus_prosthemelas | Icterus_prosthemelas | diurnal |
| birds | Icterus_pustulatus | Icterus_pustulatus | diurnal |
| birds | Icterus_spurius | Icterus_spurius | diurnal |

| | | | |
|---|---|---|---|
| birds | Icterus_wagleri | Icterus_wagleri | diurnal |
| birds | Icthyophaga_humilis | Ichthyophaga_humilis | diurnal |
| birds | Icthyophaga_ichthyaetus | Ichthyophaga_ichthyaetus | diurnal |
| birds | Ictinaetus_malaiensis | Ictinaetus_malayensis | diurnal |
| birds | Ictinia_mississippiensis | Ictinia_mississippiensis | diurnal |
| birds | Ictinia_plumbea | Ictinia_plumbea | diurnal |
| birds | Idiopsar_brachyurus | Idiopsar_brachyurus | diurnal |
| birds | Iduna_caligata | Hippolais_caligata | diurnal |
| birds | Iduna_natalensis | Chloropeta_natalensis | diurnal |
| birds | Iduna_pallida | Hippolais_pallida | diurnal |
| birds | Iduna_rama | Hippolais_rama | diurnal |
| birds | Iduna_similis | Chloropeta_similis | diurnal |
| birds | Ifrita_kowaldi | Ifrita_kowaldi | diurnal |
| birds | Ilicura_militaris | Ilicura_militaris | diurnal |
| birds | Illadopsis_albipectus | Illadopsis_albipectus | diurnal |
| birds | Illadopsis_cleaveri | Illadopsis_cleaveri | diurnal |
| birds | Illadopsis_fulvescens | Illadopsis_fulvescens | diurnal |
| birds | Illadopsis_puveli | Illadopsis_puveli | diurnal |
| birds | Illadopsis_pyrrhoptera | Illadopsis_pyrrhoptera | diurnal |
| birds | Illadopsis_rufescens | Illadopsis_rufescens | diurnal |
| birds | Illadopsis_rufipennis | Illadopsis_rufipennis | diurnal |
| birds | Illadopsis_turdina | Ptyrticus_turdinus | diurnal |
| birds | Incana_incana | Incana_incana | diurnal |
| birds | Incaspiza_laeta | Incaspiza_laeta | diurnal |
| birds | Incaspiza_ortizi | Incaspiza_ortizi | diurnal |
| birds | Incaspiza_personata | Incaspiza_personata | diurnal |
| birds | Incaspiza_pulchra | Incaspiza_pulchra | diurnal |
| birds | Incaspiza_watkinsi | Incaspiza_watkinsi | diurnal |
| birds | Indicator_archipelagicus | Indicator_archipelagicus | diurnal |
| birds | Indicator_exilis | Indicator_exilis | diurnal |
| birds | Indicator_indicator | Indicator_indicator | diurnal |
| birds | Indicator_maculatus | Indicator_maculatus | diurnal |
| birds | Indicator_meliphilus | Indicator_meliphilus | diurnal |
| birds | Indicator_minor | Indicator_minor | diurnal |

| | | | |
|---|---|---|---|
| birds | Indicator_pumilio | Indicator_pumilio | diurnal |
| birds | Indicator_variegatus | Indicator_variegatus | diurnal |
| birds | Indicator_willcocksi | Indicator_willcocksi | diurnal |
| birds | Indicator_xanthonotus | Indicator_xanthonotus | diurnal |
| birds | Inezia_caudata | Inezia_caudata | diurnal |
| birds | Inezia_inornata | Inezia_inornata | diurnal |
| birds | Inezia_subflava | Inezia_subflava | diurnal |
| birds | Inezia_tenuirostris | Inezia_tenuirostris | diurnal |
| birds | Iodopleura_fusca | Iodopleura_fusca | diurnal |
| birds | Iodopleura_isabellae | Iodopleura_isabellae | diurnal |
| birds | Iodopleura_pipra | Iodopleura_pipra | diurnal |
| birds | Iole_finschii | Alophoixus_finschii | diurnal |
| birds | Iole_palawanensis | Ixos_palawanensis | diurnal |
| birds | Iole_propinqua | Iole_propinqua | diurnal |
| birds | Irania_gutturalis | Irania_gutturalis | diurnal |
| birds | Irediparra_gallinacea | Irediparra_gallinacea | diurnal |
| birds | Irena_cyanogastra | Irena_cyanogastra | diurnal |
| birds | Irena_puella | Irena_puella | diurnal |
| birds | Iridophanes_pulcherrimus | Iridophanes_pulcherrimus | diurnal |
| birds | Iridosornis_analis | Iridosornis_analis | diurnal |
| birds | Iridosornis_jelskii | Iridosornis_jelskii | diurnal |
| birds | Iridosornis_porphyrocephalus | Iridosornis_porphyrocephalus | diurnal |
| birds | Iridosornis_reinhardti | Iridosornis_reinhardti | diurnal |
| birds | Iridosornis_rufivertex | Iridosornis_rufivertex | diurnal |
| birds | Isleria_guttata | Myrmotherula_guttata | diurnal |
| birds | Isleria_hauxwelli | Myrmotherula_hauxwelli | diurnal |
| birds | Islerothraupis_cristata | Tachyphonus_cristatus | diurnal |
| birds | Islerothraupis_luctuosa | Tachyphonus_luctuosus | diurnal |
| birds | Islerothraupis_rufiventer | Tachyphonus_rufiventer | diurnal |
| birds | Ispidina_lecontei | Ceyx_lecontei | diurnal |
| birds | Ispidina_picta | Ceyx_pictus | diurnal |
| birds | Ithaginis_cruentus | Ithaginis_cruentus | diurnal |
| birds | Ixidia_cyaniventris | Pycnonotus_cyaniventris | diurnal |
| birds | Ixidia_erythropthalmos | Pycnonotus_erythropthalmos | diurnal |

| | | | |
|---|---|---|---|
| birds | Ixidia_squamata | Pycnonotus_squamatus | diurnal |
| birds | Ixobrychus_cinnamomeus | Ixobrychus_cinnamomeus | diurnal |
| birds | Ixobrychus_eurhythmus | Ixobrychus_eurhythmus | diurnal |
| birds | Ixobrychus_flavicollis | Ixobrychus_flavicollis | cathemeral |
| birds | Ixobrychus_involucris | Ixobrychus_involucris | nocturnal |
| birds | Ixobrychus_minutus | Ixobrychus_minutus | diurnal |
| birds | Ixobrychus_sinensis | Ixobrychus_sinensis | cathemeral |
| birds | Ixobrychus_sturmii | Ixobrychus_sturmii | cathemeral |
| birds | Ixonotus_guttatus | Ixonotus_guttatus | diurnal |
| birds | Ixoreus_naevius | Zoothera_naevia | diurnal |
| birds | Ixos_malaccensis | Ixos_malaccensis | diurnal |
| birds | Ixos_mcclellandii | Hypsipetes_mcclellandii | diurnal |
| birds | Ixos_nicobariensis | Hypsipetes_nicobariensis | diurnal |
| birds | Jabiru_mycteria | Jabiru_mycteria | diurnal |
| birds | Jacamaralcyon_tridactyla | Jacamaralcyon_tridactyla | diurnal |
| birds | Jacamerops_aureus | Jacamerops_aureus | diurnal |
| birds | Jacana_jacana | Jacana_jacana | diurnal |
| birds | Jacana_spinosa | Jacana_spinosa | diurnal |
| birds | Jubula_lettii | Jubula_lettii | nocturnal |
| birds | Junco_hyemalis | Junco_hyemalis | diurnal |
| birds | Junco_insularis | Junco_insularis | diurnal |
| birds | Junco_phaeonotus | Junco_phaeonotus | diurnal |
| birds | Junco_vulcani | Junco_vulcani | diurnal |
| birds | Jynx_ruficollis | Jynx_ruficollis | diurnal |
| birds | Jynx_torquilla | Jynx_torquilla | diurnal |
| birds | Kakamega_poliothorax | Kakamega_poliothorax | diurnal |
| birds | Kaupifalco_monogrammicus | Kaupifalco_monogrammicus | diurnal |
| birds | Kempiella_flavovirescens | Microeca_flavovirescens | diurnal |
| birds | Kempiella_griseoceps | Microeca_griseoceps | diurnal |
| birds | Kenopia_striata | Kenopia_striata | diurnal |
| birds | Ketupa_flavipes | Ketupa_flavipes | cathemeral |
| birds | Ketupa_ketupu | Ketupa_ketupu | nocturnal |
| birds | Ketupa_zeylonensis | Ketupa_zeylonensis | cathemeral |
| birds | Klais_guimeti | Klais_guimeti | diurnal |

| | | | |
|---|---|---|---|
| birds | Kleinothraupis_atropileus | Hemispingus_atropileus | diurnal |
| birds | Kleinothraupis_auricularis | Hemispingus_auricularis | diurnal |
| birds | Kleinothraupis_calophrys | Hemispingus_calophrys | diurnal |
| birds | Kleinothraupis_parodii | Hemispingus_parodii | diurnal |
| birds | Kleinothraupis_reyi | Hemispingus_reyi | diurnal |
| birds | Knipolegus_aterrimus | Knipolegus_aterrimus | diurnal |
| birds | Knipolegus_cyanirostris | Knipolegus_cyanirostris | diurnal |
| birds | Knipolegus_franciscanus | Knipolegus_franciscanus | diurnal |
| birds | Knipolegus_hudsoni | Knipolegus_hudsoni | diurnal |
| birds | Knipolegus_lophotes | Knipolegus_lophotes | diurnal |
| birds | Knipolegus_nigerrimus | Knipolegus_nigerrimus | diurnal |
| birds | Knipolegus_orenocensis | Knipolegus_orenocensis | diurnal |
| birds | Knipolegus_poecilocercus | Knipolegus_poecilocercus | diurnal |
| birds | Knipolegus_poecilurus | Knipolegus_poecilurus | diurnal |
| birds | Knipolegus_signatus | Knipolegus_signatus | diurnal |
| birds | Knipolegus_striaticeps | Knipolegus_striaticeps | diurnal |
| birds | Kupeornis_chapini | Kupeornis_chapini | diurnal |
| birds | Kupeornis_gilberti | Kupeornis_gilberti | diurnal |
| birds | Kupeornis_rufocinctus | Kupeornis_rufocinctus | diurnal |
| birds | Kurochkinegramma_hypogrammica | Hypogramma_hypogrammicum | diurnal |
| birds | Lacedo_pulchella | Lacedo_pulchella | diurnal |
| birds | Lafresnaya_lafresnayi | Lafresnaya_lafresnayi | diurnal |
| birds | Lagonosticta_larvata | Lagonosticta_larvata | diurnal |
| birds | Lagonosticta_nitidula | Lagonosticta_nitidula | diurnal |
| birds | Lagonosticta_rara | Lagonosticta_rara | diurnal |
| birds | Lagonosticta_rhodopareia | Lagonosticta_rhodopareia | diurnal |
| birds | Lagonosticta_rubricata | Lagonosticta_rubricata | diurnal |
| birds | Lagonosticta_rufopicta | Lagonosticta_rufopicta | diurnal |
| birds | Lagonosticta_sanguinodorsalis | Lagonosticta_sanguinodorsalis | diurnal |
| birds | Lagonosticta_senegala | Lagonosticta_senegala | diurnal |
| birds | Lagonosticta_umbrinodorsalis | Lagonosticta_umbrinodorsalis | diurnal |
| birds | Lagonosticta_virata | Lagonosticta_virata | diurnal |
| birds | Lagopus_lagopus | Lagopus_lagopus | diurnal |
| birds | Lagopus_leucura | Lagopus_leucura | diurnal |

| | | | |
|---|---|---|---|
| birds | Lagopus_muta | Lagopus_muta | diurnal |
| birds | Lalage_atrovirens | Lalage_atrovirens | diurnal |
| birds | Lalage_aurea | Lalage_aurea | diurnal |
| birds | Lalage_fimbriata | Coracina_fimbriata | diurnal |
| birds | Lalage_leucomela | Lalage_leucomela | diurnal |
| birds | Lalage_leucopyga | Lalage_leucopyga | diurnal |
| birds | Lalage_leucopygialis | Lalage_leucopygialis | diurnal |
| birds | Lalage_maculosa | Lalage_maculosa | diurnal |
| birds | Lalage_melanoleuca | Lalage_melanoleuca | diurnal |
| birds | Lalage_melanoptera | Coracina_melanoptera | diurnal |
| birds | Lalage_melaschistos | Coracina_melaschistos | diurnal |
| birds | Lalage_moesta | Lalage_moesta | diurnal |
| birds | Lalage_nigra | Lalage_nigra | diurnal |
| birds | Lalage_polioptera | Coracina_polioptera | diurnal |
| birds | Lalage_sharpei | Lalage_sharpei | diurnal |
| birds | Lalage_sueurii | Lalage_sueurii | diurnal |
| birds | Lalage_tricolor | Lalage_tricolor | diurnal |
| birds | Lalage_typica | Coracina_typica | diurnal |
| birds | Lampornis_amethystinus | Lampornis_amethystinus | diurnal |
| birds | Lampornis_castaneoventris | Lampornis_castaneoventris | diurnal |
| birds | Lampornis_clemenciae | Lampornis_clemenciae | diurnal |
| birds | Lampornis_hemileucus | Lampornis_hemileucus | diurnal |
| birds | Lampornis_sybillae | Lampornis_sybillae | diurnal |
| birds | Lampornis_viridipallens | Lampornis_viridipallens | diurnal |
| birds | Lamprolaima_rhami | Lamprolaima_rhami | diurnal |
| birds | Lamprolia_victoriae | Lamprolia_victoriae | diurnal |
| birds | Lampropsar_tanagrinus | Lampropsar_tanagrinus | diurnal |
| birds | Lamprospiza_melanoleuca | Lamprospiza_melanoleuca | diurnal |
| birds | Lamprotornis_acuticaudus | Lamprotornis_acuticaudus | diurnal |
| birds | Lamprotornis_albicapillus | Spreo_albicapillus | diurnal |
| birds | Lamprotornis_australis | Lamprotornis_australis | diurnal |
| birds | Lamprotornis_bicolor | Spreo_bicolor | diurnal |
| birds | Lamprotornis_caudatus | Lamprotornis_caudatus | diurnal |
| birds | Lamprotornis_chalcurus | Lamprotornis_chalcurus | diurnal |

| | | | |
|---|---|---|---|
| birds | Lamprotornis_chalybaeus | Lamprotornis_chalybaeus | diurnal |
| birds | Lamprotornis_chloropterus | Lamprotornis_chloropterus | diurnal |
| birds | Lamprotornis_fischeri | Spreo_fischeri | diurnal |
| birds | Lamprotornis_hildebrandti | Lamprotornis_hildebrandti | diurnal |
| birds | Lamprotornis_iris | Coccycolius_iris | diurnal |
| birds | Lamprotornis_mevesii | Lamprotornis_mevesii | diurnal |
| birds | Lamprotornis_nitens | Lamprotornis_nitens | diurnal |
| birds | Lamprotornis_ornatus | Lamprotornis_ornatus | diurnal |
| birds | Lamprotornis_pulcher | Lamprotornis_pulcher | diurnal |
| birds | Lamprotornis_purpureus | Lamprotornis_purpureus | diurnal |
| birds | Lamprotornis_purpuroptera | Lamprotornis_purpuroptera | diurnal |
| birds | Lamprotornis_regius | Cosmopsarus_regius | diurnal |
| birds | Lamprotornis_shelleyi | Lamprotornis_shelleyi | diurnal |
| birds | Lamprotornis_splendidus | Lamprotornis_splendidus | diurnal |
| birds | Lamprotornis_superbus | Lamprotornis_superbus | diurnal |
| birds | Lamprotornis_unicolor | Cosmopsarus_unicolor | diurnal |
| birds | Laniarius_aethiopicus | Laniarius_aethiopicus | diurnal |
| birds | Laniarius_amboimensis | Laniarius_amboimensis | diurnal |
| birds | Laniarius_atrococcineus | Laniarius_atrococcineus | diurnal |
| birds | Laniarius_atroflavus | Laniarius_atroflavus | diurnal |
| birds | Laniarius_barbarus | Laniarius_barbarus | diurnal |
| birds | Laniarius_bicolor | Laniarius_bicolor | diurnal |
| birds | Laniarius_brauni | Laniarius_brauni | diurnal |
| birds | Laniarius_erythrogaster | Laniarius_erythrogaster | diurnal |
| birds | Laniarius_ferrugineus | Laniarius_ferrugineus | diurnal |
| birds | Laniarius_fuelleborni | Laniarius_fuelleborni | diurnal |
| birds | Laniarius_funebris | Laniarius_funebris | diurnal |
| birds | Laniarius_leucorhynchus | Laniarius_leucorhynchus | diurnal |
| birds | Laniarius_luehderi | Laniarius_luehderi | diurnal |
| birds | Laniarius_mufumbiri | Laniarius_mufumbiri | diurnal |
| birds | Laniarius_poensis | Laniarius_poensis | diurnal |
| birds | Laniarius_ruficeps | Laniarius_ruficeps | diurnal |
| birds | Laniarius_turatii | Laniarius_turatii | diurnal |
| birds | Laniellus_albonotatus | Crocias_albonotatus | diurnal |

| | | | |
|---|---|---|---|
| birds | Laniellus_langbianis | Crocias_langbianis | diurnal |
| birds | Laniisoma_buckleyi | Laniisoma_buckleyi | diurnal |
| birds | Laniisoma_elegans | Laniisoma_elegans | diurnal |
| birds | Lanio_aurantius | Lanio_aurantius | diurnal |
| birds | Lanio_fulvus | Lanio_fulvus | diurnal |
| birds | Lanio_leucothorax | Lanio_leucothorax | diurnal |
| birds | Lanio_versicolor | Lanio_versicolor | diurnal |
| birds | Laniocera_hypopyrra | Laniocera_hypopyrra | diurnal |
| birds | Laniocera_rufescens | Laniocera_rufescens | diurnal |
| birds | Lanioturdus_torquatus | Lanioturdus_torquatus | diurnal |
| birds | Lanius_bucephalus | Lanius_bucephalus | diurnal |
| birds | Lanius_cabanisi | Lanius_cabanisi | diurnal |
| birds | Lanius_collaris | Lanius_collaris | diurnal |
| birds | Lanius_collurio | Lanius_collurio | diurnal |
| birds | Lanius_collurioides | Lanius_collurioides | diurnal |
| birds | Lanius_corvinus | Corvinella_corvina | diurnal |
| birds | Lanius_cristatus | Lanius_cristatus | diurnal |
| birds | Lanius_dorsalis | Lanius_dorsalis | diurnal |
| birds | Lanius_excubitor | Lanius_excubitor | diurnal |
| birds | Lanius_excubitoroides | Lanius_excubitoroides | diurnal |
| birds | Lanius_gubernator | Lanius_gubernator | diurnal |
| birds | Lanius_isabellinus | Lanius_isabellinus | diurnal |
| birds | Lanius_ludovicianus | Lanius_ludovicianus | diurnal |
| birds | Lanius_mackinnoni | Lanius_mackinnoni | diurnal |
| birds | Lanius_melanoleucus | Urolestes_melanoleucus | diurnal |
| birds | Lanius_meridionalis | Lanius_meridionalis | diurnal |
| birds | Lanius_minor | Lanius_minor | diurnal |
| birds | Lanius_newtoni | Lanius_newtoni | diurnal |
| birds | Lanius_nubicus | Lanius_nubicus | diurnal |
| birds | Lanius_schach | Lanius_schach | diurnal |
| birds | Lanius_senator | Lanius_senator | diurnal |
| birds | Lanius_somalicus | Lanius_somalicus | diurnal |
| birds | Lanius_souzae | Lanius_souzae | diurnal |
| birds | Lanius_sphenocercus | Lanius_sphenocercus | diurnal |

| | | | |
|---|---|---|---|
| birds | Lanius_tephronotus | Lanius_tephronotus | diurnal |
| birds | Lanius_tigrinus | Lanius_tigrinus | diurnal |
| birds | Lanius_validirostris | Lanius_validirostris | diurnal |
| birds | Lanius_vittatus | Lanius_vittatus | diurnal |
| birds | Larosterna_inca | Larosterna_inca | diurnal |
| birds | Larus_argentatus | Larus_argentatus | diurnal |
| birds | Larus_armenicus | Larus_armenicus | diurnal |
| birds | Larus_atlanticus | Larus_atlanticus | diurnal |
| birds | Larus_atricilla | Larus_atricilla | diurnal |
| birds | Larus_audouinii | Larus_audouinii | diurnal |
| birds | Larus_belcheri | Larus_belcheri | diurnal |
| birds | Larus_brunnicephalus | Larus_brunnicephalus | diurnal |
| birds | Larus_bulleri | Larus_bulleri | diurnal |
| birds | Larus_cachinnans | Larus_cachinnans | diurnal |
| birds | Larus_californicus | Larus_californicus | diurnal |
| birds | Larus_cirrocephalus | Larus_cirrocephalus | diurnal |
| birds | Larus_crassirostris | Larus_crassirostris | diurnal |
| birds | Larus_delawarensis | Larus_delawarensis | diurnal |
| birds | Larus_dominicanus | Larus_dominicanus | diurnal |
| birds | Larus_fuliginosus | Larus_fuliginosus | diurnal |
| birds | Larus_fuscus | Larus_fuscus | diurnal |
| birds | Larus_genei | Larus_genei | diurnal |
| birds | Larus_glaucescens | Larus_glaucescens | diurnal |
| birds | Larus_glaucoides | Larus_glaucoides | diurnal |
| birds | Larus_hartlaubii | Larus_hartlaubii | diurnal |
| birds | Larus_heermanni | Larus_heermanni | diurnal |
| birds | Larus_hemprichii | Larus_hemprichii | diurnal |
| birds | Larus_hyperboreus | Larus_hyperboreus | diurnal |
| birds | Larus_ichthyaetus | Larus_ichthyaetus | diurnal |
| birds | Larus_leucophthalmus | Larus_leucophthalmus | diurnal |
| birds | Larus_livens | Larus_livens | diurnal |
| birds | Larus_maculipennis | Larus_maculipennis | diurnal |
| birds | Larus_marinus | Larus_marinus | diurnal |
| birds | Larus_melanocephalus | Larus_melanocephalus | diurnal |

| | | | |
|---|---|---|---|
| birds | Larus_modestus | Larus_modestus | cathemeral |
| birds | Larus_novaehollandiae | Larus_novaehollandiae | diurnal |
| birds | Larus_occidentalis | Larus_occidentalis | diurnal |
| birds | Larus_pacificus | Larus_pacificus | diurnal |
| birds | Larus_philadelphia | Larus_philadelphia | diurnal |
| birds | Larus_pipixcan | Larus_pipixcan | diurnal |
| birds | Larus_relictus | Larus_relictus | diurnal |
| birds | Larus_ridibundus | Larus_ridibundus | diurnal |
| birds | Larus_schistisagus | Larus_schistisagus | diurnal |
| birds | Larus_scoresbii | Leucophaeus_scoresbii | diurnal |
| birds | Larus_serranus | Larus_serranus | diurnal |
| birds | Larvivora_akahige | Erithacus_akahige | diurnal |
| birds | Larvivora_brunnea | Luscinia_brunnea | diurnal |
| birds | Larvivora_cyane | Luscinia_cyane | diurnal |
| birds | Larvivora_komadori | Erithacus_komadori | diurnal |
| birds | Larvivora_ruficeps | Luscinia_ruficeps | diurnal |
| birds | Larvivora_sibilans | Luscinia_sibilans | diurnal |
| birds | Laterallus_albigularis | Laterallus_albigularis | diurnal |
| birds | Laterallus_exilis | Laterallus_exilis | diurnal |
| birds | Laterallus_flaviventer | Porzana_flaviventer | diurnal |
| birds | Laterallus_jamaicensis | Laterallus_jamaicensis | diurnal |
| birds | Laterallus_leucopyrrhus | Laterallus_leucopyrrhus | diurnal |
| birds | Laterallus_levraudi | Laterallus_levraudi | diurnal |
| birds | Laterallus_melanophaius | Laterallus_melanophaius | diurnal |
| birds | Laterallus_rogersi | Atlantisia_rogersi | cathemeral |
| birds | Laterallus_ruber | Laterallus_ruber | diurnal |
| birds | Laterallus_spilonota | Laterallus_spilonotus | diurnal |
| birds | Laterallus_xenopterus | Laterallus_xenopterus | diurnal |
| birds | Lathamus_discolor | Lathamus_discolor | diurnal |
| birds | Lathrotriccus_euleri | Lathrotriccus_euleri | diurnal |
| birds | Lathrotriccus_griseipectus | Lathrotriccus_griseipectus | diurnal |
| birds | Laticilla_burnesii | Prinia_burnesii | diurnal |
| birds | Legatus_leucophaius | Legatus_leucophaius | diurnal |
| birds | Leiopicus_auriceps | Dendrocopos_auriceps | diurnal |

| | | | |
|---|---|---|---|
| birds | Leiopicus_mahrattensis | Dendrocopos_mahrattensis | diurnal |
| birds | Leiopicus_medius | Dendrocopos_medius | diurnal |
| birds | Leioptila_annectens | Heterophasia_annectens | diurnal |
| birds | Leiothlypis_celata | Vermivora_celata | diurnal |
| birds | Leiothlypis_crissalis | Vermivora_crissalis | diurnal |
| birds | Leiothlypis_luciae | Vermivora_luciae | diurnal |
| birds | Leiothlypis_peregrina | Vermivora_peregrina | diurnal |
| birds | Leiothlypis_ruficapilla | Vermivora_ruficapilla | diurnal |
| birds | Leiothlypis_virginiae | Vermivora_virginiae | diurnal |
| birds | Leiothrix_argentauris | Leiothrix_argentauris | diurnal |
| birds | Leiothrix_lutea | Leiothrix_lutea | diurnal |
| birds | Leipoa_ocellata | Leipoa_ocellata | diurnal |
| birds | Leistes_bellicosus | Sturnella_bellicosa | diurnal |
| birds | Leistes_loyca | Sturnella_loyca | diurnal |
| birds | Leistes_militaris | Sturnella_militaris | diurnal |
| birds | Leistes_superciliaris | Sturnella_superciliaris | diurnal |
| birds | Leonardina_woodi | Leonardina_woodi | diurnal |
| birds | Lepidocolaptes_affinis | Lepidocolaptes_affinis | diurnal |
| birds | Lepidocolaptes_albolineatus | Lepidocolaptes_albolineatus | diurnal |
| birds | Lepidocolaptes_angustirostris | Lepidocolaptes_angustirostris | diurnal |
| birds | Lepidocolaptes_falcinellus | Lepidocolaptes_falcinellus | diurnal |
| birds | Lepidocolaptes_lacrymiger | Lepidocolaptes_lacrymiger | diurnal |
| birds | Lepidocolaptes_leucogaster | Lepidocolaptes_leucogaster | diurnal |
| birds | Lepidocolaptes_souleyetii | Lepidocolaptes_souleyetii | diurnal |
| birds | Lepidocolaptes_squamatus | Lepidocolaptes_squamatus | diurnal |
| birds | Lepidogrammus_cumingi | Phaenicophaeus_cumingi | diurnal |
| birds | Lepidopygia_nana | Lonchura_nana | diurnal |
| birds | Lepidothrix_coeruleocapilla | Lepidothrix_coeruleocapilla | diurnal |
| birds | Lepidothrix_coronata | Lepidothrix_coronata | diurnal |
| birds | Lepidothrix_iris | Lepidothrix_iris | diurnal |
| birds | Lepidothrix_isidorei | Lepidothrix_isidorei | diurnal |
| birds | Lepidothrix_nattereri | Lepidothrix_nattereri | diurnal |
| birds | Lepidothrix_serena | Lepidothrix_serena | diurnal |
| birds | Lepidothrix_suavissima | Lepidothrix_suavissima | diurnal |

| | | | |
|---|---|---|---|
| birds | Leptasthenura_aegithaloides | Leptasthenura_aegithaloides | diurnal |
| birds | Leptasthenura_andicola | Leptasthenura_andicola | diurnal |
| birds | Leptasthenura_fuliginiceps | Leptasthenura_fuliginiceps | diurnal |
| birds | Leptasthenura_pileata | Leptasthenura_pileata | diurnal |
| birds | Leptasthenura_platensis | Leptasthenura_platensis | diurnal |
| birds | Leptasthenura_setaria | Leptasthenura_setaria | diurnal |
| birds | Leptasthenura_striata | Leptasthenura_striata | diurnal |
| birds | Leptasthenura_striolata | Leptasthenura_striolata | diurnal |
| birds | Leptasthenura_xenothorax | Leptasthenura_xenothorax | diurnal |
| birds | Leptocoma_aspasia | Nectarinia_aspasia | diurnal |
| birds | Leptocoma_calcostetha | Nectarinia_calcostetha | diurnal |
| birds | Leptocoma_minima | Nectarinia_minima | diurnal |
| birds | Leptocoma_sperata | Nectarinia_sperata | diurnal |
| birds | Leptocoma_zeylonica | Nectarinia_zeylonica | diurnal |
| birds | Leptodon_cayanensis | Leptodon_cayanensis | diurnal |
| birds | Leptopoecile_elegans | Leptopoecile_elegans | diurnal |
| birds | Leptopoecile_sophiae | Leptopoecile_sophiae | diurnal |
| birds | Leptopogon_amaurocephalus | Leptopogon_amaurocephalus | diurnal |
| birds | Leptopogon_rufipectus | Leptopogon_rufipectus | diurnal |
| birds | Leptopogon_superciliaris | Leptopogon_superciliaris | diurnal |
| birds | Leptopogon_taczanowskii | Leptopogon_taczanowskii | diurnal |
| birds | Leptopterus_chabert | Leptopterus_chabert | diurnal |
| birds | Leptoptilos_crumenifer | Leptoptilos_crumeniferus | diurnal |
| birds | Leptoptilos_dubius | Leptoptilos_dubius | diurnal |
| birds | Leptoptilos_javanicus | Leptoptilos_javanicus | diurnal |
| birds | Leptosittaca_branickii | Leptosittaca_branickii | diurnal |
| birds | Leptosomus_discolor | Leptosomus_discolor | diurnal |
| birds | Leptotila_battyi | Leptotila_battyi | diurnal |
| birds | Leptotila_cassinii | Leptotila_cassini | diurnal |
| birds | Leptotila_conoveri | Leptotila_conoveri | diurnal |
| birds | Leptotila_jamaicensis | Leptotila_jamaicensis | diurnal |
| birds | Leptotila_megalura | Leptotila_megalura | diurnal |
| birds | Leptotila_ochraceiventris | Leptotila_ochraceiventris | diurnal |
| birds | Leptotila_pallida | Leptotila_pallida | diurnal |

| | | | |
|---|---|---|---|
| birds | Leptotila_plumbeiceps | Leptotila_plumbeiceps | diurnal |
| birds | Leptotila_rufaxilla | Leptotila_rufaxilla | diurnal |
| birds | Leptotila_verreauxi | Leptotila_verreauxi | diurnal |
| birds | Leptotila_wellsi | Leptotila_wellsi | diurnal |
| birds | Leptotrygon_veraguensis | Geotrygon_veraguensis | diurnal |
| birds | Lerwa_lerwa | Lerwa_lerwa | diurnal |
| birds | Lesbia_nuna | Lesbia_nuna | diurnal |
| birds | Lesbia_victoriae | Lesbia_victoriae | diurnal |
| birds | Lessonia_oreas | Lessonia_oreas | diurnal |
| birds | Lessonia_rufa | Lessonia_rufa | diurnal |
| birds | Leucippus_fallax | Leucippus_fallax | diurnal |
| birds | Leucocarbo_atriceps | Phalacrocorax_atriceps | diurnal |
| birds | Leucocarbo_bougainvilliorum | Phalacrocorax_bougainvillii | diurnal |
| birds | Leucocarbo_campbelli | Phalacrocorax_campbelli | diurnal |
| birds | Leucocarbo_carunculatus | Phalacrocorax_carunculatus | diurnal |
| birds | Leucocarbo_chalconotus | Phalacrocorax_chalconotus | diurnal |
| birds | Leucocarbo_colensoi | Phalacrocorax_colensoi | diurnal |
| birds | Leucocarbo_magellanicus | Phalacrocorax_magellanicus | diurnal |
| birds | Leucocarbo_onslowi | Phalacrocorax_onslowi | diurnal |
| birds | Leucocarbo_ranfurlyi | Phalacrocorax_ranfurlyi | diurnal |
| birds | Leucochloris_albicollis | Leucochloris_albicollis | diurnal |
| birds | Leucogeranus_leucogeranus | Grus_leucogeranus | diurnal |
| birds | Leucolia_violiceps | Amazilia_violiceps | diurnal |
| birds | Leuconotopicus_albolarvatus | Picoides_albolarvatus | diurnal |
| birds | Leuconotopicus_borealis | Picoides_borealis | diurnal |
| birds | Leuconotopicus_fumigatus | Picoides_fumigatus | diurnal |
| birds | Leuconotopicus_stricklandi | Picoides_stricklandi | diurnal |
| birds | Leuconotopicus_villosus | Picoides_villosus | diurnal |
| birds | Leucopeza_semperi | Leucopeza_semperi | diurnal |
| birds | Leucopternis_kuhli | Leucopternis_kuhli | diurnal |
| birds | Leucopternis_melanops | Leucopternis_melanops | diurnal |
| birds | Leucopternis_semiplumbeus | Leucopternis_semiplumbeus | diurnal |
| birds | Leucosarcia_melanoleuca | Leucosarcia_melanoleuca | diurnal |
| birds | Leucosticte_arctoa | Leucosticte_arctoa | diurnal |

| | | | |
|---|---|---|---|
| birds | Leucosticte_atrata | Leucosticte_atrata | diurnal |
| birds | Leucosticte_australis | Leucosticte_australis | diurnal |
| birds | Leucosticte_brandti | Leucosticte_brandti | diurnal |
| birds | Leucosticte_nemoricola | Leucosticte_nemoricola | diurnal |
| birds | Leucosticte_tephrocotis | Leucosticte_tephrocotis | diurnal |
| birds | Lewinia_mirifica | Lewinia_mirifica | diurnal |
| birds | Lewinia_pectoralis | Lewinia_pectoralis | diurnal |
| birds | Lewinia_striata | Gallirallus_striatus | diurnal |
| birds | Lichenostomus_cratitius | Lichenostomus_cratitius | diurnal |
| birds | Lichenostomus_melanops | Lichenostomus_melanops | diurnal |
| birds | Lichmera_alboauricularis | Lichmera_alboauricularis | diurnal |
| birds | Lichmera_argentauris | Lichmera_argentauris | diurnal |
| birds | Lichmera_deningeri | Lichmera_deningeri | diurnal |
| birds | Lichmera_flavicans | Lichmera_flavicans | diurnal |
| birds | Lichmera_incana | Lichmera_incana | diurnal |
| birds | Lichmera_indistincta | Lichmera_indistincta | diurnal |
| birds | Lichmera_lombokia | Lichmera_lombokia | diurnal |
| birds | Lichmera_monticola | Lichmera_monticola | diurnal |
| birds | Lichmera_notabilis | Lichmera_notabilis | diurnal |
| birds | Lichmera_squamata | Lichmera_squamata | diurnal |
| birds | Limnoctites_rectirostris | Limnoctites_rectirostris | diurnal |
| birds | Limnoctites_sulphuriferus | Cranioleuca_sulphurifera | diurnal |
| birds | Limnodromus_griseus | Limnodromus_griseus | cathemeral |
| birds | Limnodromus_scolopaceus | Limnodromus_scolopaceus | diurnal |
| birds | Limnodromus_semipalmatus | Limnodromus_semipalmatus | diurnal |
| birds | Limnornis_curvirostris | Limnornis_curvirostris | diurnal |
| birds | Limnothlypis_swainsonii | Limnothlypis_swainsonii | diurnal |
| birds | Limosa_fedoa | Limosa_fedoa | diurnal |
| birds | Limosa_haemastica | Limosa_haemastica | diurnal |
| birds | Limosa_lapponica | Limosa_lapponica | diurnal |
| birds | Limosa_limosa | Limosa_limosa | diurnal |
| birds | Linaria_cannabina | Carduelis_cannabina | diurnal |
| birds | Linaria_flavirostris | Carduelis_flavirostris | diurnal |
| birds | Linaria_johannis | Carduelis_johannis | diurnal |

| | | | |
|---|---|---|---|
| birds | Linaria_yemenensis | Carduelis_yemenensis | diurnal |
| birds | Linurgus_olivaceus | Linurgus_olivaceus | diurnal |
| birds | Liocichla_omeiensis | Liocichla_omeiensis | diurnal |
| birds | Liocichla_phoenicea | Liocichla_phoenicea | diurnal |
| birds | Lioparus_chrysotis | Alcippe_chrysotis | diurnal |
| birds | Liosceles_thoracicus | Liosceles_thoracicus | diurnal |
| birds | Lipaugus_ater | Tijuca_atra | diurnal |
| birds | Lipaugus_conditus | Tijuca_condita | diurnal |
| birds | Lipaugus_fuscocinereus | Lipaugus_fuscocinereus | diurnal |
| birds | Lipaugus_lanioides | Lipaugus_lanioides | diurnal |
| birds | Lipaugus_streptophorus | Lipaugus_streptophorus | diurnal |
| birds | Lipaugus_unirufus | Lipaugus_unirufus | diurnal |
| birds | Lipaugus_uropygialis | Lipaugus_uropygialis | diurnal |
| birds | Lipaugus_vociferans | Lipaugus_vociferans | diurnal |
| birds | Lissotis_hartlaubii | Eupodotis_hartlaubii | diurnal |
| birds | Lissotis_melanogaster | Eupodotis_melanogaster | diurnal |
| birds | Loboparadisea_sericea | Loboparadisea_sericea | diurnal |
| birds | Lobotos_lobatus | Campephaga_lobata | diurnal |
| birds | Lochmias_nematura | Lochmias_nematura | diurnal |
| birds | Locustella_alfredi | Bradypterus_alfredi | diurnal |
| birds | Locustella_castanea | Bradypterus_castaneus | diurnal |
| birds | Locustella_caudata | Bradypterus_caudatus | diurnal |
| birds | Locustella_fluviatilis | Locustella_fluviatilis | diurnal |
| birds | Locustella_lanceolata | Locustella_lanceolata | diurnal |
| birds | Locustella_luscinioides | Locustella_luscinioides | diurnal |
| birds | Locustella_luteoventris | Bradypterus_luteoventris | diurnal |
| birds | Locustella_naevia | Locustella_naevia | diurnal |
| birds | Locustella_seebohmi | Bradypterus_seebohmi | diurnal |
| birds | Locustella_tacsanowskia | Bradypterus_tacsanowskius | diurnal |
| birds | Locustella_thoracica | Bradypterus_thoracicus | diurnal |
| birds | Loddigesia_mirabilis | Loddigesia_mirabilis | diurnal |
| birds | Lonchura_caniceps | Lonchura_caniceps | diurnal |
| birds | Lonchura_castaneothorax | Lonchura_castaneothorax | diurnal |
| birds | Lonchura_ferruginosa | Lonchura_ferruginosa | diurnal |

| | | | |
|---|---|---|---|
| birds | Lonchura_flaviprymna | Lonchura_flaviprymna | diurnal |
| birds | Lonchura_forbesi | Lonchura_forbesi | diurnal |
| birds | Lonchura_fuscans | Lonchura_fuscans | diurnal |
| birds | Lonchura_grandis | Lonchura_grandis | diurnal |
| birds | Lonchura_hunsteini | Lonchura_hunsteini | diurnal |
| birds | Lonchura_kelaarti | Lonchura_kelaarti | diurnal |
| birds | Lonchura_leucogastra | Lonchura_leucogastra | diurnal |
| birds | Lonchura_leucogastroides | Lonchura_leucogastroides | diurnal |
| birds | Lonchura_leucosticta | Lonchura_leucosticta | diurnal |
| birds | Lonchura_maja | Lonchura_maja | diurnal |
| birds | Lonchura_malacca | Lonchura_malacca | diurnal |
| birds | Lonchura_melaena | Lonchura_melaena | diurnal |
| birds | Lonchura_molucca | Lonchura_molucca | diurnal |
| birds | Lonchura_montana | Lonchura_montana | diurnal |
| birds | Lonchura_monticola | Lonchura_monticola | diurnal |
| birds | Lonchura_nevermanni | Lonchura_nevermanni | diurnal |
| birds | Lonchura_pallida | Lonchura_pallida | diurnal |
| birds | Lonchura_punctulata | Lonchura_punctulata | diurnal |
| birds | Lonchura_quinticolor | Lonchura_quinticolor | diurnal |
| birds | Lonchura_spectabilis | Lonchura_spectabilis | diurnal |
| birds | Lonchura_striata | Lonchura_striata | diurnal |
| birds | Lonchura_stygia | Lonchura_stygia | diurnal |
| birds | Lonchura_teerinki | Lonchura_teerinki | diurnal |
| birds | Lonchura_tristissima | Lonchura_tristissima | diurnal |
| birds | Lonchura_vana | Lonchura_vana | diurnal |
| birds | Lophaetus_occipitalis | Lophaetus_occipitalis | diurnal |
| birds | Lophoceros_alboterminatus | Tockus_alboterminatus | diurnal |
| birds | Lophoceros_bradfieldi | Tockus_bradfieldi | diurnal |
| birds | Lophoceros_camurus | Tockus_camurus | diurnal |
| birds | Lophoceros_fasciatus | Tockus_fasciatus | diurnal |
| birds | Lophoceros_hemprichii | Tockus_hemprichii | diurnal |
| birds | Lophoceros_nasutus | Tockus_nasutus | diurnal |
| birds | Lophoceros_pallidirostris | Tockus_pallidirostris | diurnal |
| birds | Lophoictinia_isura | Lophoictinia_isura | diurnal |

| | | | |
|---|---|---|---|
| birds | Lopholaimus_antarcticus | Lopholaimus_antarcticus | diurnal |
| birds | Lophonetta_specularioides | Lophonetta_specularioides | diurnal |
| birds | Lophophanes_cristatus | Parus_cristatus | diurnal |
| birds | Lophophanes_dichrous | Parus_dichrous | diurnal |
| birds | Lophophorus_impejanus | Lophophorus_impejanus | diurnal |
| birds | Lophophorus_lhuysii | Lophophorus_lhuysii | diurnal |
| birds | Lophophorus_sclateri | Lophophorus_sclateri | diurnal |
| birds | Lophorina_intercedens | Ptiloris_intercedens | diurnal |
| birds | Lophorina_magnifica | Ptiloris_magnificus | diurnal |
| birds | Lophorina_paradisea | Ptiloris_paradiseus | diurnal |
| birds | Lophorina_superba | Lophorina_superba | diurnal |
| birds | Lophorina_victoriae | Ptiloris_victoriae | diurnal |
| birds | Lophornis_adorabilis | Lophornis_adorabilis | diurnal |
| birds | Lophornis_brachylophus | Lophornis_brachylophus | diurnal |
| birds | Lophornis_chalybeus | Lophornis_chalybeus | diurnal |
| birds | Lophornis_delattrei | Lophornis_delattrei | diurnal |
| birds | Lophornis_gouldii | Lophornis_gouldii | diurnal |
| birds | Lophornis_helenae | Lophornis_helenae | diurnal |
| birds | Lophornis_magnificus | Lophornis_magnificus | diurnal |
| birds | Lophornis_ornatus | Lophornis_ornatus | diurnal |
| birds | Lophornis_pavoninus | Lophornis_pavoninus | diurnal |
| birds | Lophornis_stictolophus | Lophornis_stictolophus | diurnal |
| birds | Lophospingus_griseocristatus | Lophospingus_griseocristatus | diurnal |
| birds | Lophospingus_pusillus | Lophospingus_pusillus | diurnal |
| birds | Lophostrix_cristata | Lophostrix_cristata | nocturnal |
| birds | Lophotibis_cristata | Lophotibis_cristata | diurnal |
| birds | Lophotis_gindiana | Eupodotis_gindiana | diurnal |
| birds | Lophotis_ruficrista | Eupodotis_ruficrista | diurnal |
| birds | Lophotriccus_eulophotes | Lophotriccus_eulophotes | diurnal |
| birds | Lophotriccus_galeatus | Lophotriccus_galeatus | diurnal |
| birds | Lophotriccus_pileatus | Lophotriccus_pileatus | diurnal |
| birds | Lophotriccus_vitiosus | Lophotriccus_vitiosus | diurnal |
| birds | Lophotriorchis_kienerii | Lophotriorchis_kienerii | diurnal |
| birds | Lophura_bulweri | Lophura_bulweri | diurnal |

| | | | |
|---|---|---|---|
| birds | Lophura_diardi | Lophura_diardi | diurnal |
| birds | Lophura_edwardsi | Lophura_edwardsi | diurnal |
| birds | Lophura_erythrophthalma | Lophura_erythrophthalma | diurnal |
| birds | Lophura_ignita | Lophura_ignita | diurnal |
| birds | Lophura_inornata | Lophura_inornata | diurnal |
| birds | Lophura_leucomelanos | Lophura_leucomelanos | diurnal |
| birds | Lophura_nycthemera | Lophura_nycthemera | diurnal |
| birds | Loriculus_aurantiifrons | Loriculus_aurantiifrons | diurnal |
| birds | Loriculus_beryllinus | Loriculus_beryllinus | diurnal |
| birds | Loriculus_catamene | Loriculus_catamene | diurnal |
| birds | Loriculus_exilis | Loriculus_exilis | diurnal |
| birds | Loriculus_flosculus | Loriculus_flosculus | diurnal |
| birds | Loriculus_galgulus | Loriculus_galgulus | diurnal |
| birds | Loriculus_philippensis | Loriculus_philippensis | diurnal |
| birds | Loriculus_sclateri | Loriculus_sclateri | diurnal |
| birds | Loriculus_tener | Loriculus_tener | diurnal |
| birds | Loriculus_vernalis | Loriculus_vernalis | diurnal |
| birds | Lorius_albidinucha | Lorius_albidinucha | diurnal |
| birds | Lorius_chlorocercus | Lorius_chlorocercus | diurnal |
| birds | Lorius_domicella | Lorius_domicella | diurnal |
| birds | Lorius_garrulus | Lorius_garrulus | diurnal |
| birds | Lorius_hypoinochrous | Lorius_hypoinochrous | diurnal |
| birds | Lorius_lory | Lorius_lory | diurnal |
| birds | Loxia_curvirostra | Loxia_curvirostra | diurnal |
| birds | Loxia_leucoptera | Loxia_leucoptera | diurnal |
| birds | Loxia_pytyopsittacus | Loxia_pytyopsittacus | diurnal |
| birds | Loxia_scotica | Loxia_scotica | diurnal |
| birds | Loxigilla_noctis | Loxigilla_noctis | diurnal |
| birds | Loxioides_bailleui | Loxioides_bailleui | diurnal |
| birds | Loxipasser_anoxanthus | Loxipasser_anoxanthus | diurnal |
| birds | Loxops_caeruleirostris | Loxops_caeruleirostris | diurnal |
| birds | Loxops_coccineus | Loxops_coccineus | diurnal |
| birds | Lullula_arborea | Lullula_arborea | diurnal |
| birds | Lurocalis_rufiventris | Lurocalis_rufiventris | nocturnal |

| | | | |
|---|---|---|---|
| birds | Lurocalis_semitorquatus | Lurocalis_semitorquatus | nocturnal |
| birds | Luscinia_luscinia | Luscinia_luscinia | diurnal |
| birds | Luscinia_megarhynchos | Luscinia_megarhynchos | diurnal |
| birds | Luscinia_phaenicuroides | Hodgsonius_phaenicuroides | diurnal |
| birds | Luscinia_svecica | Luscinia_svecica | diurnal |
| birds | Lybius_chaplini | Lybius_chaplini | diurnal |
| birds | Lybius_guifsobalito | Lybius_guifsobalito | diurnal |
| birds | Lybius_leucocephalus | Lybius_leucocephalus | diurnal |
| birds | Lybius_rubrifacies | Lybius_rubrifacies | diurnal |
| birds | Lybius_torquatus | Lybius_torquatus | diurnal |
| birds | Lybius_undatus | Lybius_undatus | diurnal |
| birds | Lybius_vieilloti | Lybius_vieilloti | diurnal |
| birds | Lycocorax_pyrrhopterus | Lycocorax_pyrrhopterus | diurnal |
| birds | Lymnocryptes_minimus | Lymnocryptes_minimus | nocturnal |
| birds | Lyncornis_macrotis | Eurostopodus_macrotis | nocturnal |
| birds | Lyncornis_temminckii | Eurostopodus_temminckii | nocturnal |
| birds | Lyrurus_tetrix | Tetrao_tetrix | diurnal |
| birds | Macgregoria_pulchra | Macgregoria_pulchra | diurnal |
| birds | Machaerirhynchus_flaviventer | Machaerirhynchus_flaviventer | diurnal |
| birds | Machaerirhynchus_nigripectus | Machaerirhynchus_nigripectus | diurnal |
| birds | Machaeropterus_deliciosus | Machaeropterus_deliciosus | diurnal |
| birds | Machaeropterus_pyrocephalus | Machaeropterus_pyrocephalus | diurnal |
| birds | Machaeropterus_regulus | Machaeropterus_regulus | diurnal |
| birds | Macheiramphus_alcinus | Macheiramphus_alcinus | diurnal |
| birds | Machetornis_rixosa | Machetornis_rixosa | diurnal |
| birds | Machlolophus_holsti | Parus_holsti | diurnal |
| birds | Machlolophus_nuchalis | Parus_nuchalis | diurnal |
| birds | Machlolophus_spilonotus | Parus_spilonotus | diurnal |
| birds | Machlolophus_xanthogenys | Parus_xanthogenys | diurnal |
| birds | Mackenziaena_leachii | Mackenziaena_leachii | diurnal |
| birds | Mackenziaena_severa | Mackenziaena_severa | diurnal |
| birds | Macroagelaius_imthurni | Macroagelaius_imthurni | diurnal |
| birds | Macroagelaius_subalaris | Macroagelaius_subalaris | diurnal |
| birds | Macrocephalon_maleo | Macrocephalon_maleo | diurnal |

| | | | |
|---|---|---|---|
| birds | Macronectes_giganteus | Macronectes_giganteus | cathemeral |
| birds | Macronectes_halli | Macronectes_halli | cathemeral |
| birds | Macronus_ptilosus | Macronous_ptilosus | diurnal |
| birds | Macronus_striaticeps | Macronous_striaticeps | diurnal |
| birds | Macronyx_ameliae | Macronyx_ameliae | diurnal |
| birds | Macronyx_aurantiigula | Macronyx_aurantiigula | diurnal |
| birds | Macronyx_capensis | Macronyx_capensis | diurnal |
| birds | Macronyx_croceus | Macronyx_croceus | diurnal |
| birds | Macronyx_flavicollis | Macronyx_flavicollis | diurnal |
| birds | Macronyx_fuelleborni | Macronyx_fuellebornii | diurnal |
| birds | Macronyx_grimwoodi | Macronyx_grimwoodi | diurnal |
| birds | Macronyx_sharpei | Macronyx_sharpei | diurnal |
| birds | Macropsalis_forcipata | Macropsalis_forcipata | nocturnal |
| birds | Macropygia_amboinensis | Macropygia_amboinensis | diurnal |
| birds | Macropygia_emiliana | Macropygia_emiliana | diurnal |
| birds | Macropygia_mackinlayi | Macropygia_mackinlayi | diurnal |
| birds | Macropygia_nigrirostris | Macropygia_nigrirostris | diurnal |
| birds | Macropygia_phasianella | Macropygia_phasianella | diurnal |
| birds | Macropygia_ruficeps | Macropygia_ruficeps | diurnal |
| birds | Macropygia_rufipennis | Macropygia_rufipennis | diurnal |
| birds | Macropygia_tenuirostris | Macropygia_tenuirostris | diurnal |
| birds | Macropygia_unchall | Macropygia_unchall | diurnal |
| birds | Macrosphenus_concolor | Macrosphenus_concolor | diurnal |
| birds | Macrosphenus_flavicans | Macrosphenus_flavicans | diurnal |
| birds | Macrosphenus_kempi | Macrosphenus_kempi | diurnal |
| birds | Macrosphenus_kretschmeri | Macrosphenus_kretschmeri | diurnal |
| birds | Macrosphenus_pulitzeri | Macrosphenus_pulitzeri | diurnal |
| birds | Madanga_ruficollis | Madanga_ruficollis | diurnal |
| birds | Magumma_parva | Hemignathus_parvus | diurnal |
| birds | Malacocincla_abbotti | Malacocincla_abbotti | diurnal |
| birds | Malacocincla_sepiaria | Malacocincla_sepiaria | diurnal |
| birds | Malaconotus_blanchoti | Malaconotus_blanchoti | diurnal |
| birds | Malaconotus_gladiator | Malaconotus_gladiator | diurnal |
| birds | Malaconotus_lagdeni | Malaconotus_lagdeni | diurnal |

| | | | |
|---|---|---|---|
| birds | Malaconotus_monteiri | Malaconotus_monteiri | diurnal |
| birds | Malacopteron_affine | Malacopteron_affine | diurnal |
| birds | Malacopteron_albogulare | Malacopteron_albogulare | diurnal |
| birds | Malacopteron_cinereum | Malacopteron_cinereum | diurnal |
| birds | Malacopteron_magnirostre | Malacopteron_magnirostre | diurnal |
| birds | Malacopteron_magnum | Malacopteron_magnum | diurnal |
| birds | Malacopteron_palawanense | Malacopteron_palawanense | diurnal |
| birds | Malacoptila_fulvogularis | Malacoptila_fulvogularis | diurnal |
| birds | Malacoptila_fusca | Malacoptila_fusca | diurnal |
| birds | Malacoptila_mystacalis | Malacoptila_mystacalis | diurnal |
| birds | Malacoptila_panamensis | Malacoptila_panamensis | diurnal |
| birds | Malacoptila_rufa | Malacoptila_rufa | diurnal |
| birds | Malacoptila_semicincta | Malacoptila_semicincta | diurnal |
| birds | Malacoptila_striata | Malacoptila_striata | diurnal |
| birds | Malacorhynchus_membranaceus | Malacorhynchus_membranaceus | diurnal |
| birds | Malcorus_pectoralis | Malcorus_pectoralis | diurnal |
| birds | Malia_grata | Malia_grata | diurnal |
| birds | Malimbus_ballmanni | Malimbus_ballmanni | diurnal |
| birds | Malimbus_cassini | Malimbus_cassini | diurnal |
| birds | Malimbus_coronatus | Malimbus_coronatus | diurnal |
| birds | Malimbus_erythrogaster | Malimbus_erythrogaster | diurnal |
| birds | Malimbus_ibadanensis | Malimbus_ibadanensis | diurnal |
| birds | Malimbus_malimbicus | Malimbus_malimbicus | diurnal |
| birds | Malimbus_nitens | Malimbus_nitens | diurnal |
| birds | Malimbus_racheliae | Malimbus_racheliae | diurnal |
| birds | Malimbus_rubricollis | Malimbus_rubricollis | diurnal |
| birds | Malimbus_scutatus | Malimbus_scutatus | diurnal |
| birds | Malindangia_mcgregori | Coracina_mcgregori | diurnal |
| birds | Malurus_alboscapulatus | Malurus_alboscapulatus | diurnal |
| birds | Malurus_amabilis | Malurus_amabilis | diurnal |
| birds | Malurus_coronatus | Malurus_coronatus | diurnal |
| birds | Malurus_cyaneus | Malurus_cyaneus | diurnal |
| birds | Malurus_cyanocephalus | Malurus_cyanocephalus | diurnal |
| birds | Malurus_elegans | Malurus_elegans | diurnal |

| | | | |
|---|---|---|---|
| birds | Malurus_lamberti | Malurus_lamberti | diurnal |
| birds | Malurus_leucopterus | Malurus_leucopterus | diurnal |
| birds | Malurus_melanocephalus | Malurus_melanocephalus | diurnal |
| birds | Malurus_pulcherrimus | Malurus_pulcherrimus | diurnal |
| birds | Malurus_splendens | Malurus_splendens | diurnal |
| birds | Manacus_aurantiacus | Manacus_aurantiacus | diurnal |
| birds | Manacus_candei | Manacus_candei | diurnal |
| birds | Manacus_manacus | Manacus_manacus | diurnal |
| birds | Manacus_vitellinus | Manacus_vitellinus | diurnal |
| birds | Mandingoa_nitidula | Mandingoa_nitidula | diurnal |
| birds | Manorina_flavigula | Manorina_flavigula | diurnal |
| birds | Manorina_melanocephala | Manorina_melanocephala | diurnal |
| birds | Manorina_melanophrys | Manorina_melanophrys | diurnal |
| birds | Manorina_melanotis | Manorina_melanotis | diurnal |
| birds | Manucerthia_mana | Oreomystis_mana | diurnal |
| birds | Manucodia_ater | Manucodia_ater | diurnal |
| birds | Manucodia_chalybatus | Manucodia_chalybatus | diurnal |
| birds | Manucodia_comrii | Manucodia_comrii | diurnal |
| birds | Manucodia_jobiensis | Manucodia_jobiensis | diurnal |
| birds | Mareca_americana | Anas_americana | diurnal |
| birds | Mareca_falcata | Anas_falcata | diurnal |
| birds | Mareca_penelope | Anas_penelope | diurnal |
| birds | Mareca_sibilatrix | Anas_sibilatrix | diurnal |
| birds | Mareca_strepera | Anas_strepera | diurnal |
| birds | Margaroperdix_madagarensis | Margaroperdix_madagascariensis | diurnal |
| birds | Margarops_fuscatus | Margarops_fuscatus | diurnal |
| birds | Margarornis_bellulus | Margarornis_bellulus | diurnal |
| birds | Margarornis_rubiginosus | Margarornis_rubiginosus | diurnal |
| birds | Margarornis_squamiger | Margarornis_squamiger | diurnal |
| birds | Margarornis_stellatus | Margarornis_stellatus | diurnal |
| birds | Marmaronetta_angustirostris | Marmaronetta_angustirostris | diurnal |
| birds | Maschalethraupis_surinama | Tachyphonus_surinamus | diurnal |
| birds | Masius_chrysopterus | Masius_chrysopterus | diurnal |
| birds | Mayrornis_lessoni | Mayrornis_lessoni | diurnal |

| | | | |
|---|---|---|---|
| birds | Mayrornis_versicolor | Mayrornis_versicolor | diurnal |
| birds | Mazaria_propinqua | Synallaxis_propinqua | diurnal |
| birds | Mearnsia_picina | Mearnsia_picina | diurnal |
| birds | Mecocerculus_calopterus | Mecocerculus_calopterus | diurnal |
| birds | Mecocerculus_hellmayri | Mecocerculus_hellmayri | diurnal |
| birds | Mecocerculus_leucophrys | Mecocerculus_leucophrys | diurnal |
| birds | Mecocerculus_minor | Mecocerculus_minor | diurnal |
| birds | Mecocerculus_poecilocercus | Mecocerculus_poecilocercus | diurnal |
| birds | Mecocerculus_stictopterus | Mecocerculus_stictopterus | diurnal |
| birds | Megabyas_flammulatus | Megabyas_flammulatus | diurnal |
| birds | Megaceryle_alcyon | Megaceryle_alcyon | diurnal |
| birds | Megaceryle_lugubris | Megaceryle_lugubris | diurnal |
| birds | Megaceryle_maxima | Megaceryle_maxima | diurnal |
| birds | Megaceryle_torquata | Megaceryle_torquata | diurnal |
| birds | Megadyptes_antipodes | Megadyptes_antipodes | cathemeral |
| birds | Megalampitta_gigantea | Melampitta_gigantea | diurnal |
| birds | Megaloprepia_formosa | Ptilinopus_bernsteinii | diurnal |
| birds | Megaloprepia_magnifica | Ptilinopus_magnificus | diurnal |
| birds | Megalurus_palustris | Megalurus_palustris | diurnal |
| birds | Megapodius_bernsteinii | Megapodius_bernsteinii | diurnal |
| birds | Megapodius_cumingii | Megapodius_cumingii | diurnal |
| birds | Megapodius_decollatus | Megapodius_affinis | diurnal |
| birds | Megapodius_eremita | Megapodius_eremita | diurnal |
| birds | Megapodius_freycinet | Megapodius_freycinet | diurnal |
| birds | Megapodius_laperouse | Megapodius_laperouse | diurnal |
| birds | Megapodius_layardi | Megapodius_layardi | diurnal |
| birds | Megapodius_nicobariensis | Megapodius_nicobariensis | diurnal |
| birds | Megapodius_pritchardii | Megapodius_pritchardii | diurnal |
| birds | Megapodius_reinwardt | Megapodius_reinwardt | diurnal |
| birds | Megarynchus_pitangua | Megarynchus_pitangua | diurnal |
| birds | Megascops_albogularis | Megascops_albogularis | nocturnal |
| birds | Megascops_asio | Megascops_asio | cathemeral |
| birds | Megascops_atricapilla | Megascops_atricapilla | nocturnal |
| birds | Megascops_choliba | Megascops_choliba | nocturnal |

| | | | |
|---|---|---|---|
| birds | Megascops_clarkii | Megascops_clarkii | nocturnal |
| birds | Megascops_cooperi | Megascops_cooperi | nocturnal |
| birds | Megascops_guatemalae | Megascops_guatemalae | nocturnal |
| birds | Megascops_ingens | Megascops_ingens | nocturnal |
| birds | Megascops_kennicottii | Megascops_kennicottii | nocturnal |
| birds | Megascops_marshalli | Megascops_marshalli | nocturnal |
| birds | Megascops_nudipes | Megascops_nudipes | nocturnal |
| birds | Megascops_roboratus | Megascops_roboratus | nocturnal |
| birds | Megascops_sanctaecatarinae | Megascops_sanctaecatarinae | nocturnal |
| birds | Megascops_trichopsis | Megascops_trichopsis | nocturnal |
| birds | Megastictus_margaritatus | Megastictus_margaritatus | diurnal |
| birds | Megatriorchis_doriae | Megatriorchis_doriae | diurnal |
| birds | Megaxenops_parnaguae | Megaxenops_parnaguae | diurnal |
| birds | Megazosterops_palauensis | Megazosterops_palauensis | diurnal |
| birds | Meiglyptes_jugularis | Meiglyptes_jugularis | diurnal |
| birds | Meiglyptes_tristis | Meiglyptes_tristis | diurnal |
| birds | Meiglyptes_tukki | Meiglyptes_tukki | diurnal |
| birds | Melaenornis_annamarulae | Melaenornis_annamarulae | diurnal |
| birds | Melaenornis_ardesiacus | Melaenornis_ardesiacus | diurnal |
| birds | Melaenornis_brunneus | Dioptrornis_brunneus | diurnal |
| birds | Melaenornis_chocolatinus | Dioptrornis_chocolatinus | diurnal |
| birds | Melaenornis_edolioides | Melaenornis_edolioides | diurnal |
| birds | Melaenornis_pammelaina | Melaenornis_pammelaina | diurnal |
| birds | Melampitta_lugubris | Melampitta_lugubris | diurnal |
| birds | Melanerpes_aurifrons | Melanerpes_aurifrons | diurnal |
| birds | Melanerpes_cactorum | Melanerpes_cactorum | diurnal |
| birds | Melanerpes_candidus | Melanerpes_candidus | diurnal |
| birds | Melanerpes_carolinus | Melanerpes_carolinus | diurnal |
| birds | Melanerpes_chrysauchen | Melanerpes_chrysauchen | diurnal |
| birds | Melanerpes_chrysogenys | Melanerpes_chrysogenys | diurnal |
| birds | Melanerpes_cruentatus | Melanerpes_cruentatus | diurnal |
| birds | Melanerpes_erythrocephalus | Melanerpes_erythrocephalus | diurnal |
| birds | Melanerpes_flavifrons | Melanerpes_flavifrons | diurnal |
| birds | Melanerpes_formicivorus | Melanerpes_formicivorus | diurnal |

| | | | |
|---|---|---|---|
| birds | Melanerpes_herminieri | Melanerpes_herminieri | diurnal |
| birds | Melanerpes_hoffmannii | Melanerpes_hoffmannii | diurnal |
| birds | Melanerpes_hypopolius | Melanerpes_hypopolius | diurnal |
| birds | Melanerpes_lewis | Melanerpes_lewis | diurnal |
| birds | Melanerpes_portoricensis | Melanerpes_portoricensis | diurnal |
| birds | Melanerpes_pucherani | Melanerpes_pucherani | diurnal |
| birds | Melanerpes_pygmaeus | Melanerpes_pygmaeus | diurnal |
| birds | Melanerpes_radiolatus | Melanerpes_radiolatus | diurnal |
| birds | Melanerpes_rubricapillus | Melanerpes_rubricapillus | diurnal |
| birds | Melanerpes_striatus | Melanerpes_striatus | diurnal |
| birds | Melanerpes_superciliaris | Melanerpes_superciliaris | diurnal |
| birds | Melanerpes_uropygialis | Melanerpes_uropygialis | diurnal |
| birds | Melaniparus_afer | Parus_afer | diurnal |
| birds | Melaniparus_albiventris | Parus_albiventris | diurnal |
| birds | Melaniparus_carpi | Parus_carpi | diurnal |
| birds | Melaniparus_cinerascens | Parus_cinerascens | diurnal |
| birds | Melaniparus_fasciiventer | Parus_fasciiventer | diurnal |
| birds | Melaniparus_fringillinus | Parus_fringillinus | diurnal |
| birds | Melaniparus_funereus | Parus_funereus | diurnal |
| birds | Melaniparus_griseiventris | Parus_griseiventris | diurnal |
| birds | Melaniparus_leucomelas | Parus_leucomelas | diurnal |
| birds | Melaniparus_leuconotus | Parus_leuconotus | diurnal |
| birds | Melaniparus_niger | Parus_niger | diurnal |
| birds | Melaniparus_rufiventris | Parus_rufiventris | diurnal |
| birds | Melaniparus_thruppi | Parus_thruppi | diurnal |
| birds | Melanitta_fusca | Melanitta_fusca | diurnal |
| birds | Melanitta_nigra | Melanitta_nigra | diurnal |
| birds | Melanitta_perspicillata | Melanitta_perspicillata | diurnal |
| birds | Melanocharis_arfakiana | Melanocharis_arfakiana | diurnal |
| birds | Melanocharis_longicauda | Melanocharis_longicauda | diurnal |
| birds | Melanocharis_nigra | Melanocharis_nigra | diurnal |
| birds | Melanocharis_striativentris | Melanocharis_striativentris | diurnal |
| birds | Melanocharis_versteri | Melanocharis_versteri | diurnal |
| birds | Melanochlora_sultanea | Melanochlora_sultanea | diurnal |

| | | | |
|---|---|---|---|
| birds | Melanocichla_calva | Garrulax_calvus | diurnal |
| birds | Melanocichla_lugubris | Garrulax_lugubris | diurnal |
| birds | Melanocorypha_bimaculata | Melanocorypha_bimaculata | diurnal |
| birds | Melanocorypha_calandra | Melanocorypha_calandra | diurnal |
| birds | Melanocorypha_maxima | Melanocorypha_maxima | diurnal |
| birds | Melanocorypha_mongolica | Melanocorypha_mongolica | diurnal |
| birds | Melanocorypha_yeltoniensis | Melanocorypha_yeltoniensis | diurnal |
| birds | Melanodera_melanodera | Melanodera_melanodera | diurnal |
| birds | Melanodera_xanthogramma | Melanodera_xanthogramma | diurnal |
| birds | Melanodryas_cucullata | Melanodryas_cucullata | diurnal |
| birds | Melanodryas_vittata | Melanodryas_vittata | diurnal |
| birds | Melanopareia_elegans | Melanopareia_elegans | diurnal |
| birds | Melanopareia_maranonica | Melanopareia_maranonica | diurnal |
| birds | Melanopareia_maximiliani | Melanopareia_maximiliani | diurnal |
| birds | Melanopareia_torquata | Melanopareia_torquata | diurnal |
| birds | Melanoperdix_niger | Melanoperdix_niger | diurnal |
| birds | Melanoptila_glabrirostris | Melanoptila_glabrirostris | diurnal |
| birds | Melanorectes_nigrescens | Pitohui_nigrescens | diurnal |
| birds | Melanospiza_bicolor | Tiaris_bicolor | diurnal |
| birds | Melanospiza_richardsoni | Melanospiza_richardsoni | diurnal |
| birds | Melanotis_caerulescens | Melanotis_caerulescens | diurnal |
| birds | Melanotis_hypoleucus | Melanotis_hypoleucus | diurnal |
| birds | Meleagris_gallopavo | Meleagris_gallopavo | diurnal |
| birds | Meleagris_ocellata | Meleagris_ocellata | diurnal |
| birds | Meliarchus_sclateri | Melidectes_sclateri | diurnal |
| birds | Melichneutes_robustus | Melichneutes_robustus | diurnal |
| birds | Melidectes_belfordi | Melidectes_belfordi | diurnal |
| birds | Melidectes_foersteri | Melidectes_foersteri | diurnal |
| birds | Melidectes_leucostephes | Melidectes_leucostephes | diurnal |
| birds | Melidectes_ochromelas | Melidectes_ochromelas | diurnal |
| birds | Melidectes_rufocrissalis | Melidectes_rufocrissalis | diurnal |
| birds | Melidectes_torquatus | Melidectes_torquatus | diurnal |
| birds | Melidora_macrorrhina | Melidora_macrorrhina | cathemeral |
| birds | Melierax_canorus | Melierax_canorus | diurnal |

| | | | |
|---|---|---|---|
| birds | Melierax_metabates | Melierax_metabates | diurnal |
| birds | Melierax_poliopterus | Melierax_poliopterus | diurnal |
| birds | Melignomon_eisentrauti | Melignomon_eisentrauti | diurnal |
| birds | Melignomon_zenkeri | Melignomon_zenkeri | diurnal |
| birds | Melilestes_megarhynchus | Melilestes_megarhynchus | diurnal |
| birds | Melionyx_fuscus | Melidectes_fuscus | diurnal |
| birds | Melionyx_nouhuysi | Melidectes_nouhuysi | diurnal |
| birds | Melionyx_princeps | Melidectes_princeps | diurnal |
| birds | Meliphacator_provocator | Xanthotis_provocator | diurnal |
| birds | Meliphaga_aruensis | Meliphaga_aruensis | diurnal |
| birds | Meliphaga_lewinii | Meliphaga_lewinii | diurnal |
| birds | Meliphaga_notata | Meliphaga_notata | diurnal |
| birds | Melipotes_ater | Melipotes_ater | diurnal |
| birds | Melipotes_fumigatus | Melipotes_fumigatus | diurnal |
| birds | Melipotes_gymnops | Melipotes_gymnops | diurnal |
| birds | Melithreptus_affinis | Melithreptus_affinis | diurnal |
| birds | Melithreptus_albogularis | Melithreptus_albogularis | diurnal |
| birds | Melithreptus_brevirostris | Melithreptus_brevirostris | diurnal |
| birds | Melithreptus_gularis | Melithreptus_gularis | diurnal |
| birds | Melithreptus_lunatus | Melithreptus_lunatus | diurnal |
| birds | Melithreptus_validirostris | Melithreptus_validirostris | diurnal |
| birds | Melitograis_gilolensis | Melitograis_gilolensis | diurnal |
| birds | Mellisuga_helenae | Mellisuga_helenae | diurnal |
| birds | Mellisuga_minima | Mellisuga_minima | diurnal |
| birds | Melloria_quoyi | Cracticus_quoyi | diurnal |
| birds | Melocichla_mentalis | Melocichla_mentalis | diurnal |
| birds | Melopsittacus_undulatus | Melopsittacus_undulatus | diurnal |
| birds | Melopyrrha_portoricensis | Loxigilla_portoricensis | diurnal |
| birds | Melopyrrha_violacea | Loxigilla_violacea | diurnal |
| birds | Melospiza_georgiana | Melospiza_georgiana | diurnal |
| birds | Melospiza_lincolnii | Melospiza_lincolnii | diurnal |
| birds | Melospiza_melodia | Melospiza_melodia | diurnal |
| birds | Melozone_aberti | Pipilo_aberti | diurnal |
| birds | Melozone_albicollis | Pipilo_albicollis | diurnal |

| | | | |
|---|---|---|---|
| birds | Melozone_biarcuata | Melozone_biarcuata | diurnal |
| birds | Melozone_cabanisi | Melozone_cabanisi | diurnal |
| birds | Melozone_crissalis | Pipilo_crissalis | diurnal |
| birds | Melozone_fusca | Pipilo_fuscus | diurnal |
| birds | Melozone_kieneri | Melozone_kieneri | diurnal |
| birds | Melozone_leucotis | Melozone_leucotis | diurnal |
| birds | Mentocrex_kioloides | Canirallus_kioloides | diurnal |
| birds | Menura_alberti | Menura_alberti | diurnal |
| birds | Menura_novaehollandiae | Menura_novaehollandiae | diurnal |
| birds | Merganetta_armata | Merganetta_armata | diurnal |
| birds | Mergellus_albellus | Mergellus_albellus | diurnal |
| birds | Mergus_merganser | Mergus_merganser | diurnal |
| birds | Mergus_octosetaceus | Mergus_octosetaceus | diurnal |
| birds | Mergus_serrator | Mergus_serrator | diurnal |
| birds | Mergus_squamatus | Mergus_squamatus | diurnal |
| birds | Meropogon_forsteni | Meropogon_forsteni | diurnal |
| birds | Merops_albicollis | Merops_albicollis | diurnal |
| birds | Merops_apiaster | Merops_apiaster | diurnal |
| birds | Merops_boehmi | Merops_boehmi | diurnal |
| birds | Merops_breweri | Merops_breweri | diurnal |
| birds | Merops_bullockoides | Merops_bullockoides | diurnal |
| birds | Merops_bulocki | Merops_bulocki | diurnal |
| birds | Merops_gularis | Merops_gularis | diurnal |
| birds | Merops_hirundineus | Merops_hirundineus | diurnal |
| birds | Merops_leschenaulti | Merops_leschenaulti | diurnal |
| birds | Merops_malimbicus | Merops_malimbicus | diurnal |
| birds | Merops_muelleri | Merops_muelleri | diurnal |
| birds | Merops_nubicus | Merops_nubicus | diurnal |
| birds | Merops_oreobates | Merops_oreobates | diurnal |
| birds | Merops_orientalis | Merops_orientalis | diurnal |
| birds | Merops_ornatus | Merops_ornatus | diurnal |
| birds | Merops_persicus | Merops_persicus | diurnal |
| birds | Merops_philippinus | Merops_philippinus | diurnal |
| birds | Merops_pusillus | Merops_pusillus | diurnal |

| | | | |
|---|---|---|---|
| birds | Merops_revoilii | Merops_revoilii | diurnal |
| birds | Merops_superciliosus | Merops_superciliosus | diurnal |
| birds | Merops_variegatus | Merops_variegatus | diurnal |
| birds | Merops_viridis | Merops_viridis | diurnal |
| birds | Merulaxis_ater | Merulaxis_ater | diurnal |
| birds | Mesembrinibis_cayennensis | Mesembrinibis_cayennensis | diurnal |
| birds | Mesitornis_unicolor | Mesitornis_unicolor | diurnal |
| birds | Mesitornis_variegatus | Mesitornis_variegatus | diurnal |
| birds | Metallura_aeneocauda | Metallura_aeneocauda | diurnal |
| birds | Metallura_baroni | Metallura_baroni | diurnal |
| birds | Metallura_eupogon | Metallura_eupogon | diurnal |
| birds | Metallura_iracunda | Metallura_iracunda | diurnal |
| birds | Metallura_odomae | Metallura_odomae | diurnal |
| birds | Metallura_phoebe | Metallura_phoebe | diurnal |
| birds | Metallura_theresiae | Metallura_theresiae | diurnal |
| birds | Metallura_tyrianthina | Metallura_tyrianthina | diurnal |
| birds | Metallura_williami | Metallura_williami | diurnal |
| birds | Metopidius_indicus | Metopidius_indicus | diurnal |
| birds | Metopothrix_aurantiaca | Metopothrix_aurantiaca | diurnal |
| birds | Metriopelia_aymara | Metriopelia_aymara | diurnal |
| birds | Metriopelia_ceciliae | Metriopelia_ceciliae | diurnal |
| birds | Metriopelia_melanoptera | Metriopelia_melanoptera | diurnal |
| birds | Metriopelia_morenoi | Metriopelia_morenoi | diurnal |
| birds | Micrastur_buckleyi | Micrastur_buckleyi | diurnal |
| birds | Micrastur_gilvicollis | Micrastur_gilvicollis | diurnal |
| birds | Micrastur_mirandollei | Micrastur_mirandollei | diurnal |
| birds | Micrastur_plumbeus | Micrastur_plumbeus | diurnal |
| birds | Micrastur_ruficollis | Micrastur_ruficollis | diurnal |
| birds | Micrastur_semitorquatus | Micrastur_semitorquatus | diurnal |
| birds | Microbates_cinereiventris | Microbates_cinereiventris | diurnal |
| birds | Microbates_collaris | Microbates_collaris | diurnal |
| birds | Microcarbo_africanus | Phalacrocorax_africanus | diurnal |
| birds | Microcarbo_coronatus | Phalacrocorax_coronatus | diurnal |
| birds | Microcarbo_melanoleucos | Phalacrocorax_melanoleucos | diurnal |

| | | | |
|---|---|---|---|
| birds | Microcarbo_niger | Phalacrocorax_niger | diurnal |
| birds | Microcarbo_pygmaeus | Phalacrocorax_pygmeus | diurnal |
| birds | Microcerculus_bambla | Microcerculus_bambla | diurnal |
| birds | Microcerculus_marginatus | Microcerculus_marginatus | diurnal |
| birds | Microcerculus_philomela | Microcerculus_philomela | diurnal |
| birds | Microcerculus_ustulatus | Microcerculus_ustulatus | diurnal |
| birds | Microchera_albocoronata | Microchera_albocoronata | diurnal |
| birds | Microchera_chionura | Elvira_chionura | diurnal |
| birds | Microchera_cupreiceps | Elvira_cupreiceps | diurnal |
| birds | Microdynamis_parva | Microdynamis_parva | diurnal |
| birds | Microeca_fascinans | Microeca_fascinans | diurnal |
| birds | Microeca_flavigaster | Microeca_flavigaster | diurnal |
| birds | Microeca_hemixantha | Microeca_hemixantha | diurnal |
| birds | Microhierax_caerulescens | Microhierax_caerulescens | diurnal |
| birds | Microhierax_erythrogenys | Microhierax_erythrogenys | diurnal |
| birds | Microhierax_fringillarius | Microhierax_fringillarius | diurnal |
| birds | Microhierax_latifrons | Microhierax_latifrons | diurnal |
| birds | Microhierax_melanoleucos | Microhierax_melanoleucos | diurnal |
| birds | Microligea_palustris | Microligea_palustris | diurnal |
| birds | Micromacronus_leytensis | Micromacronus_leytensis | diurnal |
| birds | Micromonacha_lanceolata | Micromonacha_lanceolata | diurnal |
| birds | Micronisus_gabar | Melierax_gabar | diurnal |
| birds | Microparra_capensis | Microparra_capensis | diurnal |
| birds | Micropsitta_bruijnii | Micropsitta_bruijnii | diurnal |
| birds | Micropsitta_finschii | Micropsitta_finschii | diurnal |
| birds | Micropsitta_geelvinkiana | Micropsitta_geelvinkiana | diurnal |
| birds | Micropsitta_keiensis | Micropsitta_keiensis | diurnal |
| birds | Micropsitta_meeki | Micropsitta_meeki | diurnal |
| birds | Micropsitta_pusio | Micropsitta_pusio | diurnal |
| birds | Micropternus_brachyurus | Celeus_brachyurus | diurnal |
| birds | Microptilotis_albilineatus | Meliphaga_albilineata | diurnal |
| birds | Microptilotis_albonotatus | Meliphaga_albonotata | diurnal |
| birds | Microptilotis_analogus | Meliphaga_analoga | diurnal |
| birds | Microptilotis_flavirictus | Meliphaga_flavirictus | diurnal |

| | | | |
|---|---|---|---|
| birds | Microptilotis_gracilis | Meliphaga_gracilis | diurnal |
| birds | Microptilotis_mimikae | Meliphaga_mimikae | diurnal |
| birds | Microptilotis_montanus | Meliphaga_montana | diurnal |
| birds | Microptilotis_orientalis | Meliphaga_orientalis | diurnal |
| birds | Microptilotis_reticulatus | Meliphaga_reticulata | diurnal |
| birds | Microptilotis_vicina | Meliphaga_vicina | diurnal |
| birds | Micropygia_schomburgkii | Micropygia_schomburgkii | diurnal |
| birds | Microrhopias_quixensis | Microrhopias_quixensis | diurnal |
| birds | Microspingus_alticola | Poospiza_alticola | diurnal |
| birds | Microspingus_cinereus | Poospiza_cinerea | diurnal |
| birds | Microspingus_erythrophrys | Poospiza_erythrophrys | diurnal |
| birds | Microspingus_lateralis | Poospiza_lateralis | diurnal |
| birds | Microspingus_melanoleucus | Poospiza_melanoleuca | diurnal |
| birds | Microspingus_torquatus | Poospiza_torquata | diurnal |
| birds | Microspingus_trifasciatus | Hemispingus_trifasciatus | diurnal |
| birds | Microstilbon_burmeisteri | Microstilbon_burmeisteri | diurnal |
| birds | Microtarsus_melanoleucos | Pycnonotus_melanoleucos | diurnal |
| birds | Microxenops_milleri | Xenops_milleri | diurnal |
| birds | Milvago_chimachima | Milvago_chimachima | diurnal |
| birds | Milvus_migrans | Milvus_migrans | diurnal |
| birds | Milvus_milvus | Milvus_milvus | diurnal |
| birds | Mimus_dorsalis | Mimus_dorsalis | diurnal |
| birds | Mimus_gilvus | Mimus_gilvus | diurnal |
| birds | Mimus_graysoni | Mimus_graysoni | diurnal |
| birds | Mimus_gundlachii | Mimus_gundlachii | diurnal |
| birds | Mimus_longicaudatus | Mimus_longicaudatus | diurnal |
| birds | Mimus_macdonaldi | Mimus_macdonaldi | diurnal |
| birds | Mimus_melanotis | Mimus_melanotis | diurnal |
| birds | Mimus_parvulus | Mimus_parvulus | diurnal |
| birds | Mimus_patagonicus | Mimus_patagonicus | diurnal |
| birds | Mimus_polyglottos | Mimus_polyglottos | diurnal |
| birds | Mimus_saturninus | Mimus_saturninus | diurnal |
| birds | Mimus_thenca | Mimus_thenca | diurnal |
| birds | Mimus_trifasciatus | Mimus_trifasciatus | diurnal |

| | | | |
|---|---|---|---|
| birds | Mimus_triurus | Mimus_triurus | diurnal |
| birds | Minla_ignotincta | Minla_ignotincta | diurnal |
| birds | Mino_anais | Mino_anais | diurnal |
| birds | Mino_dumontii | Mino_dumontii | diurnal |
| birds | Mionectes_macconnelli | Mionectes_macconnelli | diurnal |
| birds | Mionectes_oleagineus | Mionectes_oleagineus | diurnal |
| birds | Mionectes_olivaceus | Mionectes_olivaceus | diurnal |
| birds | Mionectes_rufiventris | Mionectes_rufiventris | diurnal |
| birds | Mionectes_striaticollis | Mionectes_striaticollis | diurnal |
| birds | Mirafra_africana | Mirafra_africana | diurnal |
| birds | Mirafra_albicauda | Mirafra_albicauda | diurnal |
| birds | Mirafra_angolensis | Mirafra_angolensis | diurnal |
| birds | Mirafra_apiata | Mirafra_apiata | diurnal |
| birds | Mirafra_assamica | Mirafra_assamica | diurnal |
| birds | Mirafra_cheniana | Mirafra_cheniana | diurnal |
| birds | Mirafra_collaris | Mirafra_collaris | diurnal |
| birds | Mirafra_cordofanica | Mirafra_cordofanica | diurnal |
| birds | Mirafra_erythroptera | Mirafra_erythroptera | diurnal |
| birds | Mirafra_hypermetra | Mirafra_hypermetra | diurnal |
| birds | Mirafra_javanica | Mirafra_javanica | diurnal |
| birds | Mirafra_passerina | Mirafra_passerina | diurnal |
| birds | Mirafra_pulpa | Mirafra_pulpa | diurnal |
| birds | Mirafra_rufocinnamomea | Mirafra_rufocinnamomea | diurnal |
| birds | Mirafra_somalica | Mirafra_somalica | diurnal |
| birds | Mirafra_williamsi | Mirafra_williamsi | diurnal |
| birds | Mitrephanes_olivaceus | Mitrephanes_olivaceus | diurnal |
| birds | Mitrephanes_phaeocercus | Mitrephanes_phaeocercus | diurnal |
| birds | Mitrospingus_cassinii | Mitrospingus_cassinii | diurnal |
| birds | Mitrospingus_oleagineus | Mitrospingus_oleagineus | diurnal |
| birds | Mitu_mitu | Mitu_mitu | diurnal |
| birds | Mitu_salvini | Mitu_salvini | diurnal |
| birds | Mitu_tomentosum | Mitu_tomentosum | diurnal |
| birds | Mitu_tuberosum | Mitu_tuberosum | diurnal |
| birds | Mixornis_flavicollis | Macronous_flavicollis | diurnal |

| | | | |
|---|---|---|---|
| birds | Mixornis_gularis | Macronous_gularis | diurnal |
| birds | Mniotilta_varia | Mniotilta_varia | diurnal |
| birds | Modulatrix_stictigula | Modulatrix_stictigula | diurnal |
| birds | Mohoua_albicilla | Mohoua_albicilla | diurnal |
| birds | Mohoua_novaeseelandiae | Mohoua_novaeseelandiae | diurnal |
| birds | Mohoua_ochrocephala | Mohoua_ochrocephala | diurnal |
| birds | Molothrus_aeneus | Molothrus_aeneus | diurnal |
| birds | Molothrus_armenti | Molothrus_armenti | diurnal |
| birds | Molothrus_ater | Molothrus_ater | diurnal |
| birds | Molothrus_bonariensis | Molothrus_bonariensis | diurnal |
| birds | Molothrus_oryzivorus | Molothrus_oryzivorus | diurnal |
| birds | Molothrus_rufoaxillaris | Molothrus_rufoaxillaris | diurnal |
| birds | Momotus_aequatorialis | Momotus_aequatorialis | diurnal |
| birds | Momotus_mexicanus | Momotus_mexicanus | diurnal |
| birds | Momotus_momota | Momotus_momota | diurnal |
| birds | Monachella_muelleriana | Monachella_muelleriana | diurnal |
| birds | Monarcha_castaneiventris | Monarcha_castaneiventris | diurnal |
| birds | Monarcha_cinerascens | Monarcha_cinerascens | diurnal |
| birds | Monarcha_frater | Monarcha_frater | diurnal |
| birds | Monarcha_melanopsis | Monarcha_melanopsis | diurnal |
| birds | Monasa_atra | Monasa_atra | diurnal |
| birds | Monasa_flavirostris | Monasa_flavirostris | diurnal |
| birds | Monasa_morphoeus | Monasa_morphoeus | diurnal |
| birds | Monasa_nigrifrons | Monasa_nigrifrons | diurnal |
| birds | Monias_benschi | Monias_benschi | diurnal |
| birds | Montecincla_cachinnans | Strophocincla_cachinnans | diurnal |
| birds | Monticola_angolensis | Monticola_angolensis | diurnal |
| birds | Monticola_brevipes | Monticola_brevipes | diurnal |
| birds | Monticola_cinclorhyncha | Monticola_cinclorhynchus | diurnal |
| birds | Monticola_explorator | Monticola_explorator | diurnal |
| birds | Monticola_gularis | Monticola_gularis | diurnal |
| birds | Monticola_imerina | Monticola_imerinus | diurnal |
| birds | Monticola_rufiventris | Monticola_rufiventris | diurnal |
| birds | Monticola_rupestris | Monticola_rupestris | diurnal |

| | | | |
|---|---|---|---|
| birds | Monticola_saxatilis | Monticola_saxatilis | diurnal |
| birds | Monticola_semirufus | Myrmecocichla_semirufa | diurnal |
| birds | Monticola_sharpei | Monticola_sharpei | diurnal |
| birds | Monticola_solitarius | Monticola_solitarius | diurnal |
| birds | Montifringilla_adamsi | Montifringilla_adamsi | diurnal |
| birds | Montifringilla_nivalis | Montifringilla_nivalis | diurnal |
| birds | Morococcyx_erythropygus | Morococcyx_erythropygus | diurnal |
| birds | Morphnarchus_princeps | Leucopternis_princeps | diurnal |
| birds | Morphnus_guianensis | Morphnus_guianensis | diurnal |
| birds | Morus_bassanus | Morus_bassanus | diurnal |
| birds | Morus_capensis | Morus_capensis | diurnal |
| birds | Morus_serrator | Morus_serrator | diurnal |
| birds | Motacilla_aguimp | Motacilla_aguimp | diurnal |
| birds | Motacilla_alba | Motacilla_alba | diurnal |
| birds | Motacilla_capensis | Motacilla_capensis | diurnal |
| birds | Motacilla_cinerea | Motacilla_cinerea | diurnal |
| birds | Motacilla_citreola | Motacilla_citreola | diurnal |
| birds | Motacilla_clara | Motacilla_clara | diurnal |
| birds | Motacilla_flava | Motacilla_flava | diurnal |
| birds | Motacilla_flaviventris | Motacilla_flaviventris | diurnal |
| birds | Motacilla_grandis | Motacilla_grandis | diurnal |
| birds | Motacilla_maderaspatensis | Motacilla_madaraspatensis | diurnal |
| birds | Moupinia_poecilotis | Chrysomma_poecilotis | diurnal |
| birds | Mulleripicus_fulvus | Mulleripicus_fulvus | diurnal |
| birds | Mulleripicus_funebris | Mulleripicus_funebris | diurnal |
| birds | Mulleripicus_pulverulentus | Mulleripicus_pulverulentus | diurnal |
| birds | Muscicapa_adusta | Muscicapa_adusta | diurnal |
| birds | Muscicapa_aquatica | Muscicapa_aquatica | diurnal |
| birds | Muscicapa_cassini | Muscicapa_cassini | diurnal |
| birds | Muscicapa_dauurica | Muscicapa_dauurica | diurnal |
| birds | Muscicapa_epulata | Muscicapa_epulata | diurnal |
| birds | Muscicapa_ferruginea | Muscicapa_ferruginea | diurnal |
| birds | Muscicapa_gambagae | Muscicapa_gambagae | diurnal |
| birds | Muscicapa_griseisticta | Muscicapa_griseisticta | diurnal |

| | | | |
|---|---|---|---|
| birds | Muscicapa_muttui | Muscicapa_muttui | diurnal |
| birds | Muscicapa_randi | Muscicapa_randi | diurnal |
| birds | Muscicapa_sethsmithi | Muscicapa_sethsmithi | diurnal |
| birds | Muscicapa_sibirica | Muscicapa_sibirica | diurnal |
| birds | Muscicapa_striata | Muscicapa_striata | diurnal |
| birds | Muscigralla_brevicauda | Muscigralla_brevicauda | diurnal |
| birds | Muscipipra_vetula | Muscipipra_vetula | diurnal |
| birds | Muscisaxicola_albifrons | Muscisaxicola_albifrons | diurnal |
| birds | Muscisaxicola_albilora | Muscisaxicola_albilora | diurnal |
| birds | Muscisaxicola_alpinus | Muscisaxicola_alpinus | diurnal |
| birds | Muscisaxicola_capistratus | Muscisaxicola_capistratus | diurnal |
| birds | Muscisaxicola_cinereus | Muscisaxicola_cinereus | diurnal |
| birds | Muscisaxicola_flavinucha | Muscisaxicola_flavinucha | diurnal |
| birds | Muscisaxicola_fluviatilis | Muscisaxicola_fluviatilis | diurnal |
| birds | Muscisaxicola_frontalis | Muscisaxicola_frontalis | diurnal |
| birds | Muscisaxicola_griseus | Muscisaxicola_griseus | diurnal |
| birds | Muscisaxicola_juninensis | Muscisaxicola_juninensis | diurnal |
| birds | Muscisaxicola_maclovianus | Muscisaxicola_maclovianus | diurnal |
| birds | Muscisaxicola_maculirostris | Muscisaxicola_maculirostris | diurnal |
| birds | Muscisaxicola_rufivertex | Muscisaxicola_rufivertex | diurnal |
| birds | Musophaga_rossae | Musophaga_rossae | diurnal |
| birds | Musophaga_violacea | Musophaga_violacea | diurnal |
| birds | Myadestes_coloratus | Myadestes_coloratus | diurnal |
| birds | Myadestes_elisabeth | Myadestes_elisabeth | diurnal |
| birds | Myadestes_genibarbis | Myadestes_genibarbis | diurnal |
| birds | Myadestes_lanaiensis | Myadestes_lanaiensis | diurnal |
| birds | Myadestes_melanops | Myadestes_melanops | diurnal |
| birds | Myadestes_obscurus | Myadestes_obscurus | diurnal |
| birds | Myadestes_occidentalis | Myadestes_occidentalis | diurnal |
| birds | Myadestes_palmeri | Myadestes_palmeri | diurnal |
| birds | Myadestes_ralloides | Myadestes_ralloides | diurnal |
| birds | Myadestes_townsendi | Myadestes_townsendi | diurnal |
| birds | Myadestes_unicolor | Myadestes_unicolor | diurnal |
| birds | Mycerobas_affinis | Mycerobas_affinis | diurnal |

| | | | |
|---|---|---|---|
| birds | Mycerobas_carnipes | Mycerobas_carnipes | diurnal |
| birds | Mycerobas_icterioides | Mycerobas_icterioides | diurnal |
| birds | Mycerobas_melanozanthos | Mycerobas_melanozanthos | diurnal |
| birds | Mycteria_americana | Mycteria_americana | diurnal |
| birds | Mycteria_cinerea | Mycteria_cinerea | diurnal |
| birds | Mycteria_ibis | Mycteria_ibis | diurnal |
| birds | Mycteria_leucocephala | Mycteria_leucocephala | diurnal |
| birds | Myiagra_alecto | Myiagra_alecto | diurnal |
| birds | Myiagra_azureocapilla | Myiagra_azureocapilla | diurnal |
| birds | Myiagra_caledonica | Myiagra_caledonica | diurnal |
| birds | Myiagra_cervinicauda | Myiagra_cervinicauda | diurnal |
| birds | Myiagra_cyanoleuca | Myiagra_cyanoleuca | diurnal |
| birds | Myiagra_ferrocyanea | Myiagra_ferrocyanea | diurnal |
| birds | Myiagra_galeata | Myiagra_galeata | diurnal |
| birds | Myiagra_hebetior | Myiagra_hebetior | diurnal |
| birds | Myiagra_inquieta | Myiagra_inquieta | diurnal |
| birds | Myiagra_rubecula | Myiagra_rubecula | diurnal |
| birds | Myiagra_ruficollis | Myiagra_ruficollis | diurnal |
| birds | Myiagra_vanikorensis | Myiagra_vanikorensis | diurnal |
| birds | Myiarchus_antillarum | Myiarchus_antillarum | diurnal |
| birds | Myiarchus_apicalis | Myiarchus_apicalis | diurnal |
| birds | Myiarchus_barbirostris | Myiarchus_barbirostris | diurnal |
| birds | Myiarchus_cephalotes | Myiarchus_cephalotes | diurnal |
| birds | Myiarchus_cinerascens | Myiarchus_cinerascens | diurnal |
| birds | Myiarchus_crinitus | Myiarchus_crinitus | diurnal |
| birds | Myiarchus_ferox | Myiarchus_ferox | diurnal |
| birds | Myiarchus_magnirostris | Myiarchus_magnirostris | diurnal |
| birds | Myiarchus_nugator | Myiarchus_nugator | diurnal |
| birds | Myiarchus_nuttingi | Myiarchus_nuttingi | diurnal |
| birds | Myiarchus_oberi | Myiarchus_oberi | diurnal |
| birds | Myiarchus_panamensis | Myiarchus_panamensis | diurnal |
| birds | Myiarchus_phaeocephalus | Myiarchus_phaeocephalus | diurnal |
| birds | Myiarchus_sagrae | Myiarchus_sagrae | diurnal |
| birds | Myiarchus_semirufus | Myiarchus_semirufus | diurnal |

| | | | |
|---|---|---|---|
| birds | Myiarchus_stolidus | Myiarchus_stolidus | diurnal |
| birds | Myiarchus_swainsoni | Myiarchus_swainsoni | diurnal |
| birds | Myiarchus_tuberculifer | Myiarchus_tuberculifer | diurnal |
| birds | Myiarchus_tyrannulus | Myiarchus_tyrannulus | diurnal |
| birds | Myiarchus_validus | Myiarchus_validus | diurnal |
| birds | Myiarchus_venezuelensis | Myiarchus_venezuelensis | diurnal |
| birds | Myiarchus_yucatanensis | Myiarchus_yucatanensis | diurnal |
| birds | Myiobius_atricaudus | Myiobius_atricaudus | diurnal |
| birds | Myiobius_barbatus | Myiobius_barbatus | diurnal |
| birds | Myiobius_sulphureipygius | Myiobius_sulphureipygius | diurnal |
| birds | Myiobius_villosus | Myiobius_villosus | diurnal |
| birds | Myioborus_albifacies | Myioborus_albifacies | diurnal |
| birds | Myioborus_brunniceps | Myioborus_brunniceps | diurnal |
| birds | Myioborus_castaneocapilla | Myioborus_castaneocapilla | diurnal |
| birds | Myioborus_flavivertex | Myioborus_flavivertex | diurnal |
| birds | Myioborus_melanocephalus | Myioborus_melanocephalus | diurnal |
| birds | Myioborus_miniatus | Myioborus_miniatus | diurnal |
| birds | Myioborus_ornatus | Myioborus_ornatus | diurnal |
| birds | Myioborus_pictus | Myioborus_pictus | diurnal |
| birds | Myioborus_torquatus | Myioborus_torquatus | diurnal |
| birds | Myiodynastes_bairdii | Myiodynastes_bairdii | diurnal |
| birds | Myiodynastes_chrysocephalus | Myiodynastes_chrysocephalus | diurnal |
| birds | Myiodynastes_hemichrysus | Myiodynastes_hemichrysus | diurnal |
| birds | Myiodynastes_luteiventris | Myiodynastes_luteiventris | diurnal |
| birds | Myiodynastes_maculatus | Myiodynastes_maculatus | diurnal |
| birds | Myiomela_diana | Cinclidium_diana | diurnal |
| birds | Myiomela_leucura | Cinclidium_leucurum | diurnal |
| birds | Myiopagis_caniceps | Myiopagis_caniceps | diurnal |
| birds | Myiopagis_cotta | Myiopagis_cotta | diurnal |
| birds | Myiopagis_flavivertex | Myiopagis_flavivertex | diurnal |
| birds | Myiopagis_gaimardii | Myiopagis_gaimardii | diurnal |
| birds | Myiopagis_olallai | Myiopagis_olallai | diurnal |
| birds | Myiopagis_subplacens | Myiopagis_subplacens | diurnal |
| birds | Myiopagis_viridicata | Myiopagis_viridicata | diurnal |

| | | | |
|---|---|---|---|
| birds | Myiophobus_cryptoxanthus | Myiophobus_cryptoxanthus | diurnal |
| birds | Myiophobus_fasciatus | Myiophobus_fasciatus | diurnal |
| birds | Myiophobus_flavicans | Myiophobus_flavicans | diurnal |
| birds | Myiophobus_inornatus | Myiophobus_inornatus | diurnal |
| birds | Myiophobus_phoenicomitra | Myiophobus_phoenicomitra | diurnal |
| birds | Myiophobus_roraimae | Myiophobus_roraimae | diurnal |
| birds | Myiopsitta_monachus | Myiopsitta_monachus | diurnal |
| birds | Myiornis_albiventris | Myiornis_albiventris | diurnal |
| birds | Myiornis_atricapillus | Myiornis_atricapillus | diurnal |
| birds | Myiornis_auricularis | Myiornis_auricularis | diurnal |
| birds | Myiornis_ecaudatus | Myiornis_ecaudatus | diurnal |
| birds | Myiotheretes_fumigatus | Myiotheretes_fumigatus | diurnal |
| birds | Myiotheretes_fuscorufus | Myiotheretes_fuscorufus | diurnal |
| birds | Myiotheretes_pernix | Myiotheretes_pernix | diurnal |
| birds | Myiotheretes_striaticollis | Myiotheretes_striaticollis | diurnal |
| birds | Myiothlypis_basilica | Basileuterus_basilicus | diurnal |
| birds | Myiothlypis_bivittata | Basileuterus_bivittatus | diurnal |
| birds | Myiothlypis_chlorophrys | Basileuterus_chlorophrys | diurnal |
| birds | Myiothlypis_chrysogaster | Basileuterus_chrysogaster | diurnal |
| birds | Myiothlypis_cinereicollis | Basileuterus_cinereicollis | diurnal |
| birds | Myiothlypis_conspicillata | Basileuterus_conspicillatus | diurnal |
| birds | Myiothlypis_coronata | Basileuterus_coronatus | diurnal |
| birds | Myiothlypis_flaveola | Basileuterus_flaveolus | diurnal |
| birds | Myiothlypis_fraseri | Basileuterus_fraseri | diurnal |
| birds | Myiothlypis_fulvicauda | Phaeothlypis_fulvicauda | diurnal |
| birds | Myiothlypis_leucoblephara | Basileuterus_leucoblepharus | diurnal |
| birds | Myiothlypis_leucophrys | Basileuterus_leucophrys | diurnal |
| birds | Myiothlypis_luteoviridis | Basileuterus_luteoviridis | diurnal |
| birds | Myiothlypis_nigrocristata | Basileuterus_nigrocristatus | diurnal |
| birds | Myiothlypis_rivularis | Phaeothlypis_rivularis | diurnal |
| birds | Myiothlypis_signata | Basileuterus_signatus | diurnal |
| birds | Myiotriccus_ornatus | Myiotriccus_ornatus | diurnal |
| birds | Myiozetetes_cayanensis | Myiozetetes_cayanensis | diurnal |
| birds | Myiozetetes_granadensis | Myiozetetes_granadensis | diurnal |

| | | | |
|---|---|---|---|
| birds | Myiozetetes_luteiventris | Myiozetetes_luteiventris | diurnal |
| birds | Myiozetetes_similis | Myiozetetes_similis | diurnal |
| birds | Myophonus_caeruleus | Myophonus_caeruleus | diurnal |
| birds | Myophonus_glaucinus | Myophonus_glaucinus | diurnal |
| birds | Myophonus_horsfieldii | Myophonus_horsfieldii | diurnal |
| birds | Myophonus_melanurus | Myophonus_melanurus | diurnal |
| birds | Myophonus_robinsoni | Myophonus_robinsoni | diurnal |
| birds | Myopornis_boehmi | Muscicapa_boehmi | diurnal |
| birds | Myornis_senilis | Myornis_senilis | diurnal |
| birds | Myrmeciza_longipes | Myrmeciza_longipes | diurnal |
| birds | Myrmecocichla_aethiops | Myrmecocichla_aethiops | diurnal |
| birds | Myrmecocichla_arnotti | Myrmecocichla_arnoti | diurnal |
| birds | Myrmecocichla_formicivora | Myrmecocichla_formicivora | diurnal |
| birds | Myrmecocichla_monticola | Oenanthe_monticola | diurnal |
| birds | Myrmecocichla_nigra | Myrmecocichla_nigra | diurnal |
| birds | Myrmelastes_caurensis | Schistocichla_caurensis | diurnal |
| birds | Myrmelastes_hyperythrus | Myrmeciza_hyperythra | diurnal |
| birds | Myrmelastes_leucostigma | Schistocichla_leucostigma | diurnal |
| birds | Myrmelastes_schistaceus | Schistocichla_schistacea | diurnal |
| birds | Myrmia_micrura | Myrmia_micrura | diurnal |
| birds | Myrmoborus_leucophrys | Myrmoborus_leucophrys | diurnal |
| birds | Myrmoborus_lophotes | Percnostola_lophotes | diurnal |
| birds | Myrmoborus_lugubris | Myrmoborus_lugubris | diurnal |
| birds | Myrmoborus_melanurus | Myrmoborus_melanurus | diurnal |
| birds | Myrmoborus_myotherinus | Myrmoborus_myotherinus | diurnal |
| birds | Myrmochanes_hemileucus | Myrmochanes_hemileucus | diurnal |
| birds | Myrmoderus_ferrugineus | Myrmeciza_ferruginea | diurnal |
| birds | Myrmoderus_loricatus | Myrmeciza_loricata | diurnal |
| birds | Myrmoderus_ruficauda | Myrmeciza_ruficauda | diurnal |
| birds | Myrmoderus_squamosus | Myrmeciza_squamosa | diurnal |
| birds | Myrmophylax_atrothorax | Myrmeciza_atrothorax | diurnal |
| birds | Myrmorchilus_strigilatus | Myrmorchilus_strigilatus | diurnal |
| birds | Myrmornis_torquata | Myrmornis_torquata | diurnal |
| birds | Myrmothera_berlepschi | Hylopezus_berlepschi | diurnal |

| | | | |
|---|---|---|---|
| birds | Myrmothera_campanisona | Myrmothera_campanisona | diurnal |
| birds | Myrmothera_dives | Hylopezus_dives | diurnal |
| birds | Myrmothera_fulviventris | Hylopezus_fulviventris | diurnal |
| birds | Myrmothera_simplex | Myrmothera_simplex | diurnal |
| birds | Myrmotherula_ambigua | Myrmotherula_ambigua | diurnal |
| birds | Myrmotherula_assimilis | Myrmotherula_assimilis | diurnal |
| birds | Myrmotherula_axillaris | Myrmotherula_axillaris | diurnal |
| birds | Myrmotherula_behni | Myrmotherula_behni | diurnal |
| birds | Myrmotherula_brachyura | Myrmotherula_brachyura | diurnal |
| birds | Myrmotherula_cherriei | Myrmotherula_cherriei | diurnal |
| birds | Myrmotherula_grisea | Myrmotherula_grisea | diurnal |
| birds | Myrmotherula_ignota | Myrmotherula_ignota | diurnal |
| birds | Myrmotherula_iheringi | Myrmotherula_iheringi | diurnal |
| birds | Myrmotherula_klagesi | Myrmotherula_klagesi | diurnal |
| birds | Myrmotherula_longicauda | Myrmotherula_longicauda | diurnal |
| birds | Myrmotherula_longipennis | Myrmotherula_longipennis | diurnal |
| birds | Myrmotherula_luctuosa | Myrmotherula_luctuosa | diurnal |
| birds | Myrmotherula_menetriesii | Myrmotherula_menetriesii | diurnal |
| birds | Myrmotherula_minor | Myrmotherula_minor | diurnal |
| birds | Myrmotherula_multostriata | Myrmotherula_multostriata | diurnal |
| birds | Myrmotherula_pacifica | Myrmotherula_pacifica | diurnal |
| birds | Myrmotherula_schisticolor | Myrmotherula_schisticolor | diurnal |
| birds | Myrmotherula_sclateri | Myrmotherula_sclateri | diurnal |
| birds | Myrmotherula_snowi | Myrmotherula_snowi | diurnal |
| birds | Myrmotherula_sunensis | Myrmotherula_sunensis | diurnal |
| birds | Myrmotherula_surinamensis | Myrmotherula_surinamensis | diurnal |
| birds | Myrmotherula_unicolor | Myrmotherula_unicolor | diurnal |
| birds | Myrmotherula_urosticta | Myrmotherula_urosticta | diurnal |
| birds | Myrtis_fanny | Myrtis_fanny | diurnal |
| birds | Mystacornis_crossleyi | Mystacornis_crossleyi | diurnal |
| birds | Myza_celebensis | Myza_celebensis | diurnal |
| birds | Myza_sarasinorum | Myza_sarasinorum | diurnal |
| birds | Myzomela_adolphinae | Myzomela_adolphinae | diurnal |
| birds | Myzomela_albigula | Myzomela_albigula | diurnal |

| | | | |
|---|---|---|---|
| birds | Myzomela_blasii | Myzomela_blasii | diurnal |
| birds | Myzomela_boiei | Myzomela_boiei | diurnal |
| birds | Myzomela_caledonica | Myzomela_caledonica | diurnal |
| birds | Myzomela_cardinalis | Myzomela_cardinalis | diurnal |
| birds | Myzomela_chermesina | Myzomela_chermesina | diurnal |
| birds | Myzomela_chloroptera | Myzomela_chloroptera | diurnal |
| birds | Myzomela_cineracea | Myzomela_cineracea | diurnal |
| birds | Myzomela_cruentata | Myzomela_cruentata | diurnal |
| birds | Myzomela_eichhorni | Myzomela_eichhorni | diurnal |
| birds | Myzomela_eques | Myzomela_eques | diurnal |
| birds | Myzomela_erythrocephala | Myzomela_erythrocephala | diurnal |
| birds | Myzomela_erythromelas | Myzomela_erythromelas | diurnal |
| birds | Myzomela_jugularis | Myzomela_jugularis | diurnal |
| birds | Myzomela_kuehni | Myzomela_kuehni | diurnal |
| birds | Myzomela_lafargei | Myzomela_lafargei | diurnal |
| birds | Myzomela_malaitae | Myzomela_malaitae | diurnal |
| birds | Myzomela_melanocephala | Myzomela_melanocephala | diurnal |
| birds | Myzomela_nigrita | Myzomela_nigrita | diurnal |
| birds | Myzomela_obscura | Myzomela_obscura | diurnal |
| birds | Myzomela_pammelaena | Myzomela_pammelaena | diurnal |
| birds | Myzomela_pulchella | Myzomela_pulchella | diurnal |
| birds | Myzomela_rosenbergii | Myzomela_rosenbergii | diurnal |
| birds | Myzomela_rubratra | Myzomela_rubratra | diurnal |
| birds | Myzomela_sanguinolenta | Myzomela_sanguinolenta | diurnal |
| birds | Myzomela_sclateri | Myzomela_sclateri | diurnal |
| birds | Myzomela_tristrami | Myzomela_tristrami | diurnal |
| birds | Myzomela_vulnerata | Myzomela_vulnerata | diurnal |
| birds | Myzomela_wakoloensis | Myzomela_wakoloensis | diurnal |
| birds | Myzornis_pyrrhoura | Myzornis_pyrrhoura | diurnal |
| birds | Nannopsittaca_dachilleae | Nannopsittaca_dachilleae | diurnal |
| birds | Nannopsittaca_panychlora | Nannopsittaca_panychlora | diurnal |
| birds | Nannopterum_auritum | Phalacrocorax_auritus | diurnal |
| birds | Nannopterum_brasilianum | Phalacrocorax_brasilianus | diurnal |
| birds | Nannopterum_harrisi | Phalacrocorax_harrisi | diurnal |

| | | | |
|---|---|---|---|
| birds | Napothera_epilepidota | Napothera_epilepidota | diurnal |
| birds | Nasica_longirostris | Nasica_longirostris | diurnal |
| birds | Neafrapus_boehmi | Neafrapus_boehmi | diurnal |
| birds | Neafrapus_cassini | Neafrapus_cassini | diurnal |
| birds | Necrosyrtes_monachus | Necrosyrtes_monachus | diurnal |
| birds | Nectarinia_bocagii | Nectarinia_bocagii | diurnal |
| birds | Nectarinia_famosa | Nectarinia_famosa | diurnal |
| birds | Nectarinia_johnstoni | Nectarinia_johnstoni | diurnal |
| birds | Nectarinia_kilimensis | Nectarinia_kilimensis | diurnal |
| birds | Nectarinia_purpureiventris | Nectarinia_purpureiventris | diurnal |
| birds | Nectarinia_tacazze | Nectarinia_tacazze | diurnal |
| birds | Nemosia_pileata | Nemosia_pileata | diurnal |
| birds | Nemosia_rourei | Nemosia_rourei | diurnal |
| birds | Neochen_jubata | Neochen_jubata | diurnal |
| birds | Neochmia_modesta | Neochmia_modesta | diurnal |
| birds | Neochmia_phaeton | Neochmia_phaeton | diurnal |
| birds | Neochmia_ruficauda | Neochmia_ruficauda | diurnal |
| birds | Neochmia_temporalis | Neochmia_temporalis | diurnal |
| birds | Neocichla_gutturalis | Neocichla_gutturalis | diurnal |
| birds | Neocossyphus_poensis | Neocossyphus_poensis | diurnal |
| birds | Neocossyphus_rufus | Neocossyphus_rufus | diurnal |
| birds | Neoctantes_niger | Neoctantes_niger | diurnal |
| birds | Neodrepanis_coruscans | Neodrepanis_coruscans | diurnal |
| birds | Neodrepanis_hypoxantha | Neodrepanis_hypoxantha | diurnal |
| birds | Neohierax_insignis | Polihierax_insignis | diurnal |
| birds | Neolalage_banksiana | Neolalage_banksiana | diurnal |
| birds | Neolestes_torquatus | Neolestes_torquatus | diurnal |
| birds | Neomixis_striatigula | Neomixis_striatigula | diurnal |
| birds | Neomixis_tenella | Neomixis_tenella | diurnal |
| birds | Neomixis_viridis | Neomixis_viridis | diurnal |
| birds | Neomorphus_geoffroyi | Neomorphus_geoffroyi | diurnal |
| birds | Neomorphus_pucheranii | Neomorphus_pucheranii | diurnal |
| birds | Neomorphus_radiolosus | Neomorphus_radiolosus | diurnal |
| birds | Neomorphus_rufipennis | Neomorphus_rufipennis | diurnal |

| | | | |
|---|---|---|---|
| birds | Neopelma_aurifrons | Neopelma_aurifrons | diurnal |
| birds | Neopelma_chrysocephalum | Neopelma_chrysocephalum | diurnal |
| birds | Neopelma_chrysolophum | Neopelma_chrysolophum | diurnal |
| birds | Neopelma_pallescens | Neopelma_pallescens | diurnal |
| birds | Neopelma_sulphureiventer | Neopelma_sulphureiventer | diurnal |
| birds | Neophedina_cincta | Riparia_cincta | diurnal |
| birds | Neophema_chrysogaster | Neophema_chrysogaster | diurnal |
| birds | Neophema_chrysostoma | Neophema_chrysostoma | diurnal |
| birds | Neophema_elegans | Neophema_elegans | diurnal |
| birds | Neophema_petrophila | Neophema_petrophila | diurnal |
| birds | Neophema_pulchella | Neophema_pulchella | diurnal |
| birds | Neophema_splendida | Neophema_splendida | diurnal |
| birds | Neophron_percnopterus | Neophron_percnopterus | diurnal |
| birds | Neopipo_cinnamomea | Neopipo_cinnamomea | diurnal |
| birds | Neopsephotus_bourkii | Neopsephotus_bourkii | diurnal |
| birds | Neopsittacus_musschenbroekii | Neopsittacus_musschenbroekii | diurnal |
| birds | Neopsittacus_pullicauda | Neopsittacus_pullicauda | diurnal |
| birds | Neosericornis_citreogularis | Sericornis_citreogularis | diurnal |
| birds | Neothraupis_fasciata | Neothraupis_fasciata | diurnal |
| birds | Neotis_denhami | Neotis_denhami | diurnal |
| birds | Neotis_heuglinii | Neotis_heuglinii | diurnal |
| birds | Neotis_ludwigii | Neotis_ludwigii | diurnal |
| birds | Neotis_nuba | Neotis_nuba | diurnal |
| birds | Neoxolmis_rufiventris | Neoxolmis_rufiventris | diurnal |
| birds | Nephelomyias_lintoni | Myiophobus_lintoni | diurnal |
| birds | Nephelomyias_ochraceiventris | Myiophobus_ochraceiventris | diurnal |
| birds | Nephelomyias_pulcher | Myiophobus_pulcher | diurnal |
| birds | Nephelornis_oneilli | Nephelornis_oneilli | diurnal |
| birds | Nesillas_brevicaudata | Nesillas_brevicaudata | diurnal |
| birds | Nesillas_mariae | Nesillas_mariae | diurnal |
| birds | Nesillas_typica | Nesillas_typica | diurnal |
| birds | Nesocharis_ansorgei | Nesocharis_ansorgei | diurnal |
| birds | Nesocharis_shelleyi | Nesocharis_shelleyi | diurnal |
| birds | Nesoctites_micromegas | Nesoctites_micromegas | diurnal |

| | | | |
|---|---|---|---|
| birds | Nesoenas_mayeri | Nesoenas_mayeri | diurnal |
| birds | Nesoenas_picturatus | Nesoenas_picturata | diurnal |
| birds | Nesofregetta_fuliginosa | Nesofregetta_fuliginosa | cathemeral |
| birds | Nesophlox_evelynae | Calliphlox_evelynae | diurnal |
| birds | Nesopsar_nigerrimus | Nesopsar_nigerrimus | diurnal |
| birds | Nesoptilotis_flavicollis | Lichenostomus_flavicollis | diurnal |
| birds | Nesoptilotis_leucotis | Lichenostomus_leucotis | diurnal |
| birds | Nesospingus_speculiferus | Nesospingus_speculiferus | diurnal |
| birds | Nesospiza_acunhae | Nesospiza_acunhae | diurnal |
| birds | Nesospiza_wilkinsi | Nesospiza_wilkinsi | diurnal |
| birds | Nesotriccus_ridgwayi | Nesotriccus_ridgwayi | diurnal |
| birds | Nestor_meridionalis | Nestor_meridionalis | diurnal |
| birds | Nestor_notabilis | Nestor_notabilis | diurnal |
| birds | Netta_erythrophthalma | Netta_erythrophthalma | diurnal |
| birds | Netta_peposaca | Netta_peposaca | diurnal |
| birds | Netta_rufina | Netta_rufina | diurnal |
| birds | Nettapus_auritus | Nettapus_auritus | diurnal |
| birds | Nettapus_coromandelianus | Nettapus_coromandelianus | diurnal |
| birds | Nettapus_pulchellus | Nettapus_pulchellus | diurnal |
| birds | Newtonia_amphichroa | Newtonia_amphichroa | diurnal |
| birds | Newtonia_archboldi | Newtonia_archboldi | diurnal |
| birds | Newtonia_brunneicauda | Newtonia_brunneicauda | diurnal |
| birds | Newtonia_fanovanae | Newtonia_fanovanae | diurnal |
| birds | Nicator_chloris | Nicator_chloris | diurnal |
| birds | Nicator_gularis | Nicator_gularis | diurnal |
| birds | Nicator_vireo | Nicator_vireo | diurnal |
| birds | Nicopsitta_columboides | Psittacula_columboides | diurnal |
| birds | Nigrita_bicolor | Nigrita_bicolor | diurnal |
| birds | Nigrita_canicapillus | Nigrita_canicapillus | diurnal |
| birds | Nigrita_fusconotus | Nigrita_fusconotus | diurnal |
| birds | Nigrita_luteifrons | Nigrita_luteifrons | diurnal |
| birds | Nilaus_afer | Nilaus_afer | diurnal |
| birds | Niltava_davidi | Niltava_davidi | diurnal |
| birds | Niltava_grandis | Niltava_grandis | diurnal |

| | | | |
|---|---|---|---|
| birds | Niltava_sumatrana | Niltava_sumatrana | diurnal |
| birds | Niltava_sundara | Niltava_sundara | diurnal |
| birds | Niltava_vivida | Niltava_vivida | diurnal |
| birds | Ninox_affinis | Ninox_affinis | nocturnal |
| birds | Ninox_boobook | Ninox_boobook | nocturnal |
| birds | Ninox_connivens | Ninox_connivens | nocturnal |
| birds | Ninox_jacquinoti | Ninox_jacquinoti | nocturnal |
| birds | Ninox_natalis | Ninox_natalis | nocturnal |
| birds | Ninox_novaeseelandiae | Ninox_novaeseelandiae | nocturnal |
| birds | Ninox_odiosa | Ninox_odiosa | nocturnal |
| birds | Ninox_philippensis | Ninox_philippensis | nocturnal |
| birds | Ninox_punctulata | Ninox_punctulata | nocturnal |
| birds | Ninox_rudolfi | Ninox_rudolfi | nocturnal |
| birds | Ninox_rufa | Ninox_rufa | nocturnal |
| birds | Ninox_scutulata | Ninox_scutulata | nocturnal |
| birds | Ninox_strenua | Ninox_strenua | nocturnal |
| birds | Ninox_theomacha | Ninox_theomacha | nocturnal |
| birds | Nipponia_nippon | Nipponia_nippon | diurnal |
| birds | Nisaetus_alboniger | Nisaetus_alboniger | diurnal |
| birds | Nisaetus_cirrhatus | Nisaetus_cirrhatus | diurnal |
| birds | Nisaetus_nanus | Nisaetus_nanus | diurnal |
| birds | Nisaetus_nipalensis | Nisaetus_nipalensis | diurnal |
| birds | Nisaetus_philippensis | Nisaetus_philippensis | diurnal |
| birds | Nomonyx_dominicus | Nomonyx_dominicus | diurnal |
| birds | Nonnula_amaurocephala | Nonnula_amaurocephala | diurnal |
| birds | Nonnula_brunnea | Nonnula_brunnea | diurnal |
| birds | Nonnula_frontalis | Nonnula_frontalis | diurnal |
| birds | Nonnula_rubecula | Nonnula_rubecula | diurnal |
| birds | Nonnula_ruficapilla | Nonnula_ruficapilla | diurnal |
| birds | Nonnula_sclateri | Nonnula_sclateri | diurnal |
| birds | Northiella_haematogaster | Northiella_haematogaster | diurnal |
| birds | Notharchus_macrorhynchos | Notharchus_macrorhynchos | diurnal |
| birds | Notharchus_ordii | Notharchus_ordii | diurnal |
| birds | Notharchus_pectoralis | Notharchus_pectoralis | diurnal |

| | | | |
|---|---|---|---|
| birds | Notharchus_swainsoni | Notharchus_swainsoni | diurnal |
| birds | Notharchus_tectus | Notharchus_tectus | diurnal |
| birds | Nothocercus_bonapartei | Nothocercus_bonapartei | diurnal |
| birds | Nothocercus_nigrocapillus | Nothocercus_nigrocapillus | diurnal |
| birds | Nothocrax_urumutum | Nothocrax_urumutum | diurnal |
| birds | Nothoprocta_cinerascens | Nothoprocta_cinerascens | diurnal |
| birds | Nothoprocta_curvirostris | Nothoprocta_curvirostris | diurnal |
| birds | Nothoprocta_ornata | Nothoprocta_ornata | diurnal |
| birds | Nothoprocta_pentlandii | Nothoprocta_pentlandii | diurnal |
| birds | Nothoprocta_perdicaria | Nothoprocta_perdicaria | diurnal |
| birds | Nothura_boraquira | Nothura_boraquira | diurnal |
| birds | Nothura_darwinii | Nothura_darwinii | diurnal |
| birds | Nothura_maculosa | Nothura_maculosa | diurnal |
| birds | Nothura_minor | Nothura_minor | diurnal |
| birds | Notiomystis_cincta | Notiomystis_cincta | diurnal |
| birds | Notopholia_corusca | Lamprotornis_corruscus | diurnal |
| birds | Nucifraga_caryocatactes | Nucifraga_caryocatactes | diurnal |
| birds | Nucifraga_columbiana | Nucifraga_columbiana | diurnal |
| birds | Numenius_americanus | Numenius_americanus | diurnal |
| birds | Numenius_arquata | Numenius_arquata | diurnal |
| birds | Numenius_borealis | Numenius_borealis | diurnal |
| birds | Numenius_madagascariensis | Numenius_madagascariensis | cathemeral |
| birds | Numenius_phaeopus | Numenius_phaeopus | diurnal |
| birds | Numenius_tahitiensis | Numenius_tahitiensis | diurnal |
| birds | Numenius_tenuirostris | Numenius_tenuirostris | diurnal |
| birds | Numida_meleagris | Numida_meleagris | diurnal |
| birds | Nyctanassa_violacea | Nyctanassa_violacea | cathemeral |
| birds | Nyctibius_aethereus | Nyctibius_aethereus | nocturnal |
| birds | Nyctibius_grandis | Nyctibius_grandis | nocturnal |
| birds | Nyctibius_griseus | Nyctibius_griseus | nocturnal |
| birds | Nyctibius_jamaicensis | Nyctibius_jamaicensis | nocturnal |
| birds | Nyctibius_leucopterus | Nyctibius_leucopterus | nocturnal |
| birds | Nyctibius_maculosus | Nyctibius_maculosus | nocturnal |
| birds | Nycticorax_caledonicus | Nycticorax_caledonicus | cathemeral |

| | | | |
|---|---|---|---|
| birds | Nycticorax_nycticorax | Nycticorax_nycticorax | cathemeral |
| birds | Nyctidromus_albicollis | Nyctidromus_albicollis | nocturnal |
| birds | Nyctidromus_anthonyi | Caprimulgus_anthonyi | nocturnal |
| birds | Nyctiphrynus_mcleodii | Nyctiphrynus_mcleodii | nocturnal |
| birds | Nyctiphrynus_ocellatus | Nyctiphrynus_ocellatus | nocturnal |
| birds | Nyctiphrynus_rosenbergi | Nyctiphrynus_rosenbergi | nocturnal |
| birds | Nyctiphrynus_yucatanicus | Nyctiphrynus_yucatanicus | nocturnal |
| birds | Nyctipolus_hirundinaceus | Caprimulgus_hirundinaceus | nocturnal |
| birds | Nyctipolus_nigrescens | Caprimulgus_nigrescens | nocturnal |
| birds | Nyctiprogne_leucopyga | Nyctiprogne_leucopyga | nocturnal |
| birds | Nyctyornis_amictus | Nyctyornis_amictus | diurnal |
| birds | Nyctyornis_athertoni | Nyctyornis_athertoni | diurnal |
| birds | Nymphicus_hollandicus | Nymphicus_hollandicus | diurnal |
| birds | Nystactes_tamatia | Bucco_tamatia | diurnal |
| birds | Nystalus_chacuru | Nystalus_chacuru | diurnal |
| birds | Nystalus_maculatus | Nystalus_maculatus | diurnal |
| birds | Nystalus_radiatus | Nystalus_radiatus | diurnal |
| birds | Nystalus_striolatus | Nystalus_striolatus | diurnal |
| birds | Oceanites_gracilis | Oceanites_gracilis | diurnal |
| birds | Oceanites_oceanicus | Oceanites_oceanicus | diurnal |
| birds | Ochetorhynchus_andaecola | Ochetorhynchus_andaecola | diurnal |
| birds | Ochetorhynchus_melanurus | Ochetorhynchus_melanura | diurnal |
| birds | Ochetorhynchus_phoenicurus | Ochetorhynchus_phoenicurus | diurnal |
| birds | Ochetorhynchus_ruficaudus | Ochetorhynchus_ruficaudus | diurnal |
| birds | Ochthoeca_cinnamomeiventris | Ochthoeca_cinnamomeiventris | diurnal |
| birds | Ochthoeca_fumicolor | Ochthoeca_fumicolor | diurnal |
| birds | Ochthoeca_leucophrys | Ochthoeca_leucophrys | diurnal |
| birds | Ochthoeca_nigrita | Ochthoeca_nigrita | diurnal |
| birds | Ochthoeca_oenanthoides | Ochthoeca_oenanthoides | diurnal |
| birds | Ochthoeca_piurae | Ochthoeca_piurae | diurnal |
| birds | Ochthoeca_rufipectoralis | Ochthoeca_rufipectoralis | diurnal |
| birds | Ochthoeca_salvini | Tumbezia_salvini | diurnal |
| birds | Ochthoeca_thoracica | Ochthoeca_thoracica | diurnal |
| birds | Ochthornis_littoralis | Ochthornis_littoralis | diurnal |

| | | | |
|---|---|---|---|
| birds | Ocreatus_underwoodii | Ocreatus_underwoodii | diurnal |
| birds | Ocyceros_birostris | Ocyceros_birostris | diurnal |
| birds | Ocyceros_gingalensis | Ocyceros_gingalensis | diurnal |
| birds | Ocyceros_griseus | Ocyceros_griseus | diurnal |
| birds | Ocyphaps_lophotes | Ocyphaps_lophotes | diurnal |
| birds | Odontophorus_atrifrons | Odontophorus_atrifrons | diurnal |
| birds | Odontophorus_balliviani | Odontophorus_balliviani | diurnal |
| birds | Odontophorus_capueira | Odontophorus_capueira | diurnal |
| birds | Odontophorus_columbianus | Odontophorus_columbianus | diurnal |
| birds | Odontophorus_dialeucos | Odontophorus_dialeucos | diurnal |
| birds | Odontophorus_erythrops | Odontophorus_erythrops | diurnal |
| birds | Odontophorus_gujanensis | Odontophorus_gujanensis | diurnal |
| birds | Odontophorus_guttatus | Odontophorus_guttatus | diurnal |
| birds | Odontophorus_hyperythrus | Odontophorus_hyperythrus | diurnal |
| birds | Odontophorus_leucolaemus | Odontophorus_leucolaemus | diurnal |
| birds | Odontophorus_melanonotus | Odontophorus_melanonotus | diurnal |
| birds | Odontophorus_melanotis | Odontophorus_melanotis | diurnal |
| birds | Odontophorus_speciosus | Odontophorus_speciosus | diurnal |
| birds | Odontophorus_stellatus | Odontophorus_stellatus | diurnal |
| birds | Odontophorus_strophium | Odontophorus_strophium | diurnal |
| birds | Odontorchilus_branickii | Odontorchilus_branickii | diurnal |
| birds | Odontorchilus_cinereus | Odontorchilus_cinereus | diurnal |
| birds | Oedistoma_iliolophus | Toxorhamphus_iliolophus | diurnal |
| birds | Oedistoma_pygmaeum | Oedistoma_pygmaeum | diurnal |
| birds | Oena_capensis | Oena_capensis | diurnal |
| birds | Oenanthe_albifrons | Myrmecocichla_albifrons | diurnal |
| birds | Oenanthe_albonigra | Oenanthe_albonigra | diurnal |
| birds | Oenanthe_bottae | Oenanthe_bottae | diurnal |
| birds | Oenanthe_cypriaca | Oenanthe_cypriaca | diurnal |
| birds | Oenanthe_deserti | Oenanthe_deserti | diurnal |
| birds | Oenanthe_familiaris | Cercomela_familiaris | diurnal |
| birds | Oenanthe_finschii | Oenanthe_finschii | diurnal |
| birds | Oenanthe_fusca | Cercomela_fusca | diurnal |
| birds | Oenanthe_hispanica | Oenanthe_hispanica | diurnal |

| | | | |
|---|---|---|---|
| birds | Oenanthe_isabellina | Oenanthe_isabellina | diurnal |
| birds | Oenanthe_leucopyga | Oenanthe_leucopyga | diurnal |
| birds | Oenanthe_leucura | Oenanthe_leucura | diurnal |
| birds | Oenanthe_lugens | Oenanthe_lugens | diurnal |
| birds | Oenanthe_melanura | Cercomela_melanura | diurnal |
| birds | Oenanthe_moesta | Oenanthe_moesta | diurnal |
| birds | Oenanthe_monacha | Oenanthe_monacha | diurnal |
| birds | Oenanthe_oenanthe | Oenanthe_oenanthe | diurnal |
| birds | Oenanthe_phillipsi | Oenanthe_phillipsi | diurnal |
| birds | Oenanthe_picata | Oenanthe_picata | diurnal |
| birds | Oenanthe_pileata | Oenanthe_pileata | diurnal |
| birds | Oenanthe_pleschanka | Oenanthe_pleschanka | diurnal |
| birds | Oenanthe_scotocerca | Cercomela_scotocerca | diurnal |
| birds | Oenanthe_xanthoprymna | Oenanthe_xanthoprymna | diurnal |
| birds | Ognorhynchus_icterotis | Ognorhynchus_icterotis | diurnal |
| birds | Oncostoma_cinereigulare | Oncostoma_cinereigulare | diurnal |
| birds | Oncostoma_olivaceum | Oncostoma_olivaceum | diurnal |
| birds | Oneillornis_lunulatus | Gymnopithys_lunulatus | diurnal |
| birds | Oneillornis_salvini | Gymnopithys_salvini | diurnal |
| birds | Onychognathus_albirostris | Onychognathus_albirostris | diurnal |
| birds | Onychognathus_blythii | Onychognathus_blythii | diurnal |
| birds | Onychognathus_frater | Onychognathus_frater | diurnal |
| birds | Onychognathus_fulgidus | Onychognathus_fulgidus | diurnal |
| birds | Onychognathus_morio | Onychognathus_morio | cathemeral |
| birds | Onychognathus_nabouroup | Onychognathus_nabouroup | diurnal |
| birds | Onychognathus_neumanni | Onychognathus_neumanni | diurnal |
| birds | Onychognathus_salvadorii | Onychognathus_salvadorii | diurnal |
| birds | Onychognathus_tenuirostris | Onychognathus_tenuirostris | diurnal |
| birds | Onychognathus_tristramii | Onychognathus_tristramii | diurnal |
| birds | Onychognathus_walleri | Onychognathus_walleri | diurnal |
| birds | Onychoprion_aleuticus | Sterna_aleutica | diurnal |
| birds | Onychoprion_anaethetus | Sterna_anaethetus | diurnal |
| birds | Onychoprion_fuscatus | Sterna_fuscata | diurnal |
| birds | Onychoprion_lunatus | Sterna_lunata | diurnal |

| | | | |
|---|---|---|---|
| birds | Onychorhynchus_coronatus | Onychorhynchus_coronatus | diurnal |
| birds | Onychorhynchus_mexicanus | Onychorhynchus_mexicanus | diurnal |
| birds | Onychorhynchus_occidentalis | Onychorhynchus_occidentalis | diurnal |
| birds | Onychorhynchus_swainsoni | Onychorhynchus_swainsoni | diurnal |
| birds | Onychostruthus_taczanowskii | Montifringilla_taczanowskii | diurnal |
| birds | Opisthocomus_hoazin | Opisthocomus_hoazin | nocturnal |
| birds | Opisthoprora_euryptera | Opisthoprora_euryptera | diurnal |
| birds | Oporornis_agilis | Oporornis_agilis | diurnal |
| birds | Orchesticus_abeillei | Orchesticus_abeillei | diurnal |
| birds | Oreocharis_arfaki | Oreocharis_arfaki | diurnal |
| birds | Oreoica_gutturalis | Oreoica_gutturalis | diurnal |
| birds | Oreolais_pulcher | Oreolais_pulchra | diurnal |
| birds | Oreolais_ruwenzorii | Oreolais_ruwenzorii | diurnal |
| birds | Oreomystis_bairdi | Oreomystis_bairdi | diurnal |
| birds | Oreonympha_nobilis | Oreonympha_nobilis | diurnal |
| birds | Oreophasis_derbianus | Oreophasis_derbianus | diurnal |
| birds | Oreophilais_robertsi | Prinia_robertsi | diurnal |
| birds | Oreopholus_ruficollis | Oreopholus_ruficollis | diurnal |
| birds | Oreopsar_bolivianus | Oreopsar_bolivianus | diurnal |
| birds | Oreopsittacus_arfaki | Oreopsittacus_arfaki | diurnal |
| birds | Oreornis_chrysogenys | Oreornis_chrysogenys | diurnal |
| birds | Oreortyx_pictus | Oreortyx_pictus | diurnal |
| birds | Oreoscoptes_montanus | Oreoscoptes_montanus | diurnal |
| birds | Oreoscopus_gutturalis | Oreoscopus_gutturalis | diurnal |
| birds | Oreostruthus_fuliginosus | Oreostruthus_fuliginosus | diurnal |
| birds | Oreothlypis_gutturalis | Parula_gutturalis | diurnal |
| birds | Oreothlypis_superciliosa | Parula_superciliosa | diurnal |
| birds | Oreothraupis_arremonops | Oreothraupis_arremonops | diurnal |
| birds | Oreotrochilus_adela | Oreotrochilus_adela | diurnal |
| birds | Oreotrochilus_chimborazo | Oreotrochilus_chimborazo | diurnal |
| birds | Oreotrochilus_estella | Oreotrochilus_estella | diurnal |
| birds | Oreotrochilus_leucopleurus | Oreotrochilus_leucopleurus | diurnal |
| birds | Oreotrochilus_melanogaster | Oreotrochilus_melanogaster | diurnal |
| birds | Origma_murina | Crateroscelis_murina | diurnal |

| | | | |
|---|---|---|---|
| birds | Origma_robusta | Crateroscelis_robusta | diurnal |
| birds | Origma_solitaria | Origma_solitaria | diurnal |
| birds | Oriolia_bernieri | Oriolia_bernieri | diurnal |
| birds | Oriolus_albiloris | Oriolus_albiloris | diurnal |
| birds | Oriolus_auratus | Oriolus_auratus | diurnal |
| birds | Oriolus_brachyrynchus | Oriolus_brachyrhynchus | diurnal |
| birds | Oriolus_chinensis | Oriolus_chinensis | diurnal |
| birds | Oriolus_crassirostris | Oriolus_crassirostris | diurnal |
| birds | Oriolus_cruentus | Oriolus_cruentus | diurnal |
| birds | Oriolus_flavocinctus | Oriolus_flavocinctus | diurnal |
| birds | Oriolus_forsteni | Oriolus_forsteni | diurnal |
| birds | Oriolus_isabellae | Oriolus_isabellae | diurnal |
| birds | Oriolus_larvatus | Oriolus_larvatus | diurnal |
| birds | Oriolus_mellianus | Oriolus_mellianus | diurnal |
| birds | Oriolus_monacha | Oriolus_monacha | diurnal |
| birds | Oriolus_nigripennis | Oriolus_nigripennis | diurnal |
| birds | Oriolus_oriolus | Oriolus_oriolus | diurnal |
| birds | Oriolus_percivali | Oriolus_percivali | diurnal |
| birds | Oriolus_sagittatus | Oriolus_sagittatus | diurnal |
| birds | Oriolus_steerii | Oriolus_steerii | diurnal |
| birds | Oriolus_szalayi | Oriolus_szalayi | diurnal |
| birds | Oriolus_tenuirostris | Oriolus_tenuirostris | diurnal |
| birds | Oriolus_traillii | Oriolus_traillii | diurnal |
| birds | Oriolus_xanthonotus | Oriolus_xanthonotus | diurnal |
| birds | Oriolus_xanthornus | Oriolus_xanthornus | diurnal |
| birds | Oriturus_superciliosus | Oriturus_superciliosus | diurnal |
| birds | Ornithion_brunneicapillus | Ornithion_brunneicapillus | diurnal |
| birds | Ornithion_inerme | Ornithion_inerme | diurnal |
| birds | Ornithion_semiflavum | Ornithion_semiflavum | diurnal |
| birds | Ornorectes_cristatus | Pitohui_cristatus | diurnal |
| birds | Oroanassa_magnifica | Gorsachius_magnificus | nocturnal |
| birds | Orochelidon_andecola | Haplochelidon_andecola | diurnal |
| birds | Orochelidon_flavipes | Notiochelidon_flavipes | diurnal |
| birds | Orochelidon_murina | Notiochelidon_murina | diurnal |

| | | | |
|---|---|---|---|
| birds | Ortalis_canicollis | Ortalis_canicollis | diurnal |
| birds | Ortalis_cinereiceps | Ortalis_cinereiceps | diurnal |
| birds | Ortalis_erythroptera | Ortalis_erythroptera | diurnal |
| birds | Ortalis_garrula | Ortalis_garrula | diurnal |
| birds | Ortalis_guttata | Ortalis_guttata | diurnal |
| birds | Ortalis_leucogastra | Ortalis_leucogastra | diurnal |
| birds | Ortalis_motmot | Ortalis_motmot | diurnal |
| birds | Ortalis_poliocephala | Ortalis_poliocephala | diurnal |
| birds | Ortalis_ruficauda | Ortalis_ruficauda | diurnal |
| birds | Ortalis_vetula | Ortalis_vetula | diurnal |
| birds | Ortalis_wagleri | Ortalis_wagleri | diurnal |
| birds | Orthogonys_chloricterus | Orthogonys_chloricterus | diurnal |
| birds | Orthonyx_spaldingii | Orthonyx_spaldingii | diurnal |
| birds | Orthonyx_temminckii | Orthonyx_temminckii | diurnal |
| birds | Orthopsittaca_manilatus | Orthopsittaca_manilata | diurnal |
| birds | Orthorhyncus_cristatus | Orthorhyncus_cristatus | diurnal |
| birds | Orthotomus_atrogularis | Orthotomus_atrogularis | diurnal |
| birds | Orthotomus_castaneiceps | Orthotomus_castaneiceps | diurnal |
| birds | Orthotomus_cinereiceps | Orthotomus_cinereiceps | diurnal |
| birds | Orthotomus_derbianus | Orthotomus_derbianus | diurnal |
| birds | Orthotomus_frontalis | Orthotomus_frontalis | diurnal |
| birds | Orthotomus_nigriceps | Orthotomus_nigriceps | diurnal |
| birds | Orthotomus_ruficeps | Orthotomus_ruficeps | diurnal |
| birds | Orthotomus_samarensis | Orthotomus_samarensis | diurnal |
| birds | Orthotomus_sepium | Orthotomus_sepium | diurnal |
| birds | Orthotomus_sericeus | Orthotomus_sericeus | diurnal |
| birds | Orthotomus_sutorius | Orthotomus_sutorius | diurnal |
| birds | Ortygornis_pondicerianus | Francolinus_pondicerianus | diurnal |
| birds | Ortygornis_sephaena | Francolinus_sephaena | diurnal |
| birds | Ortygospiza_atricollis | Ortygospiza_atricollis | diurnal |
| birds | Otidiphaps_nobilis | Otidiphaps_nobilis | diurnal |
| birds | Otis_tarda | Otis_tarda | diurnal |
| birds | Otocichla_mupinensis | Turdus_mupinensis | diurnal |
| birds | Otus_angelinae | Otus_angelinae | nocturnal |

| | | | |
|---|---|---|---|
| birds | Otus_bakkamoena | Otus_bakkamoena | nocturnal |
| birds | Otus_beccarii | Otus_beccarii | nocturnal |
| birds | Otus_brucei | Otus_brucei | cathemeral |
| birds | Otus_elegans | Otus_elegans | nocturnal |
| birds | Otus_hartlaubi | Otus_hartlaubi | diurnal |
| birds | Otus_insularis | Otus_insularis | nocturnal |
| birds | Otus_ireneae | Otus_ireneae | nocturnal |
| birds | Otus_lempiji | Otus_lempiji | nocturnal |
| birds | Otus_lettia | Otus_lettia | nocturnal |
| birds | Otus_magicus | Otus_magicus | nocturnal |
| birds | Otus_manadensis | Otus_manadensis | nocturnal |
| birds | Otus_sagittatus | Otus_sagittatus | nocturnal |
| birds | Otus_scops | Otus_scops | cathemeral |
| birds | Otus_semitorques | Otus_semitorques | nocturnal |
| birds | Otus_senegalensis | Otus_senegalensis | nocturnal |
| birds | Otus_spilocephalus | Otus_spilocephalus | nocturnal |
| birds | Otus_sunia | Otus_sunia | nocturnal |
| birds | Otus_umbra | Otus_umbra | nocturnal |
| birds | Oxylabes_madagascariensis | Oxylabes_madagascariensis | diurnal |
| birds | Oxypogon_guerinii | Oxypogon_guerinii | diurnal |
| birds | Oxyruncus_cristatus | Oxyruncus_cristatus | diurnal |
| birds | Oxyura_australis | Oxyura_australis | diurnal |
| birds | Oxyura_jamaicensis | Oxyura_jamaicensis | diurnal |
| birds | Oxyura_leucocephala | Oxyura_leucocephala | diurnal |
| birds | Oxyura_maccoa | Oxyura_maccoa | diurnal |
| birds | Oxyura_vittata | Oxyura_vittata | diurnal |
| birds | Pachycare_flavogriseum | Pachycare_flavogriseum | diurnal |
| birds | Pachycephala_albiventris | Pachycephala_albiventris | diurnal |
| birds | Pachycephala_caledonica | Pachycephala_caledonica | diurnal |
| birds | Pachycephala_cinerea | Pachycephala_grisola | diurnal |
| birds | Pachycephala_flavifrons | Pachycephala_flavifrons | diurnal |
| birds | Pachycephala_griseonota | Pachycephala_griseonota | diurnal |
| birds | Pachycephala_homeyeri | Pachycephala_homeyeri | diurnal |
| birds | Pachycephala_hyperythra | Pachycephala_hyperythra | diurnal |

| | | | |
|---|---|---|---|
| birds | Pachycephala_implicata | Pachycephala_implicata | diurnal |
| birds | Pachycephala_inornata | Pachycephala_inornata | diurnal |
| birds | Pachycephala_jacquinoti | Pachycephala_jacquinoti | diurnal |
| birds | Pachycephala_lanioides | Pachycephala_lanioides | diurnal |
| birds | Pachycephala_leucogastra | Pachycephala_leucogastra | diurnal |
| birds | Pachycephala_lorentzi | Pachycephala_lorentzi | diurnal |
| birds | Pachycephala_melanura | Pachycephala_melanura | diurnal |
| birds | Pachycephala_meyeri | Pachycephala_meyeri | diurnal |
| birds | Pachycephala_modesta | Pachycephala_modesta | diurnal |
| birds | Pachycephala_monacha | Pachycephala_monacha | diurnal |
| birds | Pachycephala_olivacea | Pachycephala_olivacea | diurnal |
| birds | Pachycephala_orpheus | Pachycephala_orpheus | diurnal |
| birds | Pachycephala_pectoralis | Pachycephala_pectoralis | diurnal |
| birds | Pachycephala_phaionota | Pachycephala_phaionota | diurnal |
| birds | Pachycephala_philippinensis | Pachycephala_philippinensis | diurnal |
| birds | Pachycephala_rufiventris | Pachycephala_rufiventris | diurnal |
| birds | Pachycephala_rufogularis | Pachycephala_rufogularis | diurnal |
| birds | Pachycephala_schlegelii | Pachycephala_schlegelii | diurnal |
| birds | Pachycephala_simplex | Pachycephala_simplex | diurnal |
| birds | Pachycephala_soror | Pachycephala_soror | diurnal |
| birds | Pachycephala_sulfuriventer | Pachycephala_sulfuriventer | diurnal |
| birds | Pachycephala_tenebrosa | Colluricincla_tenebrosa | diurnal |
| birds | Pachycephalopsis_hattamensis | Pachycephalopsis_hattamensis | diurnal |
| birds | Pachycephalopsis_poliosoma | Pachycephalopsis_poliosoma | diurnal |
| birds | Pachycoccyx_audeberti | Pachycoccyx_audeberti | diurnal |
| birds | Pachyptila_belcheri | Pachyptila_belcheri | cathemeral |
| birds | Pachyptila_crassirostris | Pachyptila_crassirostris | cathemeral |
| birds | Pachyptila_desolata | Pachyptila_desolata | cathemeral |
| birds | Pachyptila_salvini | Pachyptila_salvini | cathemeral |
| birds | Pachyptila_turtur | Pachyptila_turtur | cathemeral |
| birds | Pachyptila_vittata | Pachyptila_vittata | cathemeral |
| birds | Pachyramphus_aglaiae | Pachyramphus_aglaiae | diurnal |
| birds | Pachyramphus_albogriseus | Pachyramphus_albogriseus | diurnal |
| birds | Pachyramphus_castaneus | Pachyramphus_castaneus | diurnal |

| | | | |
|---|---|---|---|
| birds | Pachyramphus_cinnamomeus | Pachyramphus_cinnamomeus | diurnal |
| birds | Pachyramphus_homochrous | Pachyramphus_homochrous | diurnal |
| birds | Pachyramphus_major | Pachyramphus_major | diurnal |
| birds | Pachyramphus_marginatus | Pachyramphus_marginatus | diurnal |
| birds | Pachyramphus_minor | Pachyramphus_minor | diurnal |
| birds | Pachyramphus_niger | Pachyramphus_niger | diurnal |
| birds | Pachyramphus_polychopterus | Pachyramphus_polychopterus | diurnal |
| birds | Pachyramphus_rufus | Pachyramphus_rufus | diurnal |
| birds | Pachyramphus_spodiurus | Pachyramphus_spodiurus | diurnal |
| birds | Pachyramphus_surinamus | Pachyramphus_surinamus | diurnal |
| birds | Pachyramphus_validus | Pachyramphus_validus | diurnal |
| birds | Pachyramphus_versicolor | Pachyramphus_versicolor | diurnal |
| birds | Pachyramphus_viridis | Pachyramphus_viridis | diurnal |
| birds | Pachysylvia_aurantiifrons | Hylophilus_aurantiifrons | diurnal |
| birds | Pachysylvia_decurtata | Hylophilus_decurtatus | diurnal |
| birds | Pachysylvia_hypochrysea | Vireo_hypochryseus | diurnal |
| birds | Pachysylvia_hypoxantha | Hylophilus_hypoxanthus | diurnal |
| birds | Pachysylvia_muscicapina | Hylophilus_muscicapinus | diurnal |
| birds | Pachysylvia_semibrunnea | Hylophilus_semibrunneus | diurnal |
| birds | Padda_fuscata | Padda_fuscata | diurnal |
| birds | Padda_oryzivora | Padda_oryzivora | diurnal |
| birds | Pagodroma_nivea | Pagodroma_nivea | cathemeral |
| birds | Pagophila_eburnea | Pagophila_eburnea | diurnal |
| birds | Palaeornis_eupatria | Psittacula_eupatria | diurnal |
| birds | Palmeria_dolei | Palmeria_dolei | diurnal |
| birds | Pampa_rufa | Campylopterus_rufus | diurnal |
| birds | Pampusana_beccarii | Gallicolumba_beccarii | diurnal |
| birds | Pampusana_erythroptera | Gallicolumba_erythroptera | diurnal |
| birds | Pampusana_jobiensis | Gallicolumba_jobiensis | diurnal |
| birds | Pampusana_kubaryi | Gallicolumba_kubaryi | diurnal |
| birds | Pampusana_rubescens | Gallicolumba_rubescens | diurnal |
| birds | Pampusana_stairi | Gallicolumba_stairi | diurnal |
| birds | Pampusana_xanthonura | Gallicolumba_xanthonura | diurnal |
| birds | Pandion_haliaetus | Pandion_haliaetus | diurnal |

| | | | |
|---|---|---|---|
| birds | Panterpe_insignis | Panterpe_insignis | diurnal |
| birds | Panurus_biarmicus | Panurus_biarmicus | diurnal |
| birds | Panyptila_cayennensis | Panyptila_cayennensis | diurnal |
| birds | Panyptila_sanctihieronymi | Panyptila_sanctihieronymi | diurnal |
| birds | Papasula_abbotti | Papasula_abbotti | diurnal |
| birds | Parabuteo_leucorrhous | Buteo_leucorrhous | diurnal |
| birds | Parabuteo_unicinctus | Parabuteo_unicinctus | diurnal |
| birds | Paraclaravis_geoffroyi | Claravis_godefrida | diurnal |
| birds | Paraclaravis_mondetoura | Claravis_mondetoura | diurnal |
| birds | Paradigalla_brevicauda | Paradigalla_brevicauda | diurnal |
| birds | Paradigalla_carunculata | Paradigalla_carunculata | diurnal |
| birds | Paradisaea_apoda | Paradisaea_apoda | diurnal |
| birds | Paradisaea_decora | Paradisaea_decora | diurnal |
| birds | Paradisaea_guilielmi | Paradisaea_guilielmi | diurnal |
| birds | Paradisaea_minor | Paradisaea_minor | diurnal |
| birds | Paradisaea_raggiana | Paradisaea_raggiana | diurnal |
| birds | Paradisaea_rubra | Paradisaea_rubra | diurnal |
| birds | Paradisornis_rudolphi | Paradisaea_rudolphi | diurnal |
| birds | Paradoxornis_aemodium | Conostoma_oemodium | diurnal |
| birds | Paradoxornis_flavirostris | Paradoxornis_flavirostris | diurnal |
| birds | Paradoxornis_gularis | Paradoxornis_gularis | diurnal |
| birds | Paradoxornis_guttaticollis | Paradoxornis_guttaticollis | diurnal |
| birds | Paradoxornis_heudei | Paradoxornis_heudei | diurnal |
| birds | Paradoxornis_paradoxus | Paradoxornis_paradoxus | diurnal |
| birds | Paradoxornis_unicolor | Paradoxornis_unicolor | diurnal |
| birds | Paragallinula_angulata | Gallinula_angulata | diurnal |
| birds | Paramythia_montium | Paramythia_montium | diurnal |
| birds | Parayuhina_diademata | Yuhina_diademata | diurnal |
| birds | Pardaliparus_amabilis | Parus_amabilis | diurnal |
| birds | Pardaliparus_elegans | Parus_elegans | diurnal |
| birds | Pardaliparus_venustulus | Parus_venustulus | diurnal |
| birds | Pardalotus_punctatus | Pardalotus_punctatus | diurnal |
| birds | Pardalotus_quadragintus | Pardalotus_quadragintus | diurnal |
| birds | Pardalotus_rubricatus | Pardalotus_rubricatus | diurnal |

| | | | |
|---|---|---|---|
| birds | Pardipicus_caroli | Campethera_caroli | diurnal |
| birds | Pardipicus_nivosus | Campethera_nivosa | diurnal |
| birds | Pardirallus_nigricans | Pardirallus_nigricans | diurnal |
| birds | Pardirallus_sanguinolentus | Pardirallus_sanguinolentus | cathemeral |
| birds | Pareudiastes_silvestris | Gallinula_silvestris | diurnal |
| birds | Parkerthraustes_humeralis | Parkerthraustes_humeralis | diurnal |
| birds | Parkesia_motacilla | Seiurus_motacilla | diurnal |
| birds | Parkesia_noveboracensis | Seiurus_noveboracensis | diurnal |
| birds | Parmoptila_jamesoni | Parmoptila_jamesoni | diurnal |
| birds | Parmoptila_rubrifrons | Parmoptila_rubrifrons | diurnal |
| birds | Parmoptila_woodhousei | Parmoptila_woodhousei | diurnal |
| birds | Paroaria_baeri | Paroaria_baeri | diurnal |
| birds | Paroaria_capitata | Paroaria_capitata | diurnal |
| birds | Paroaria_coronata | Paroaria_coronata | diurnal |
| birds | Paroaria_dominicana | Paroaria_dominicana | diurnal |
| birds | Paroaria_gularis | Paroaria_gularis | diurnal |
| birds | Paroreomyza_maculata | Paroreomyza_maculata | diurnal |
| birds | Paroreomyza_montana | Paroreomyza_montana | diurnal |
| birds | Parotia_carolae | Parotia_carolae | diurnal |
| birds | Parotia_lawesii | Parotia_lawesii | diurnal |
| birds | Parotia_sefilata | Parotia_sefilata | diurnal |
| birds | Parotia_wahnesi | Parotia_wahnesi | diurnal |
| birds | Parus_major | Parus_major | diurnal |
| birds | Parus_monticolus | Parus_monticolus | diurnal |
| birds | Passer_ammodendri | Passer_ammodendri | diurnal |
| birds | Passer_castanopterus | Passer_castanopterus | diurnal |
| birds | Passer_cinnamomeus | Passer_rutilans | diurnal |
| birds | Passer_cordofanicus | Passer_cordofanicus | diurnal |
| birds | Passer_diffusus | Passer_diffusus | diurnal |
| birds | Passer_domesticus | Passer_domesticus | diurnal |
| birds | Passer_eminibey | Passer_eminibey | diurnal |
| birds | Passer_euchlorus | Passer_euchlorus | diurnal |
| birds | Passer_flaveolus | Passer_flaveolus | diurnal |
| birds | Passer_gongonensis | Passer_gongonensis | diurnal |

| | | | |
|---|---|---|---|
| birds | Passer_griseus | Passer_griseus | diurnal |
| birds | Passer_hispaniolensis | Passer_hispaniolensis | diurnal |
| birds | Passer_iagoensis | Passer_iagoensis | diurnal |
| birds | Passer_insularis | Passer_insularis | diurnal |
| birds | Passer_luteus | Passer_luteus | diurnal |
| birds | Passer_melanurus | Passer_melanurus | diurnal |
| birds | Passer_moabiticus | Passer_moabiticus | diurnal |
| birds | Passer_montanus | Passer_montanus | diurnal |
| birds | Passer_motitensis | Passer_motitensis | diurnal |
| birds | Passer_pyrrhonotus | Passer_pyrrhonotus | diurnal |
| birds | Passer_rufocinctus | Passer_rufocinctus | diurnal |
| birds | Passer_shelleyi | Passer_shelleyi | diurnal |
| birds | Passer_simplex | Passer_simplex | diurnal |
| birds | Passer_suahelicus | Passer_suahelicus | diurnal |
| birds | Passer_swainsonii | Passer_swainsonii | diurnal |
| birds | Passerculus_bairdii | Ammodramus_bairdii | diurnal |
| birds | Passerculus_henslowii | Ammodramus_henslowii | diurnal |
| birds | Passerculus_sandwichensis | Passerculus_sandwichensis | diurnal |
| birds | Passerella_arborea | Spizella_arborea | diurnal |
| birds | Passerella_iliaca | Passerella_iliaca | diurnal |
| birds | Passerina_amoena | Passerina_amoena | diurnal |
| birds | Passerina_caerulea | Passerina_caerulea | diurnal |
| birds | Passerina_ciris | Passerina_ciris | diurnal |
| birds | Passerina_cyanea | Passerina_cyanea | diurnal |
| birds | Passerina_leclancherii | Passerina_leclancherii | diurnal |
| birds | Passerina_rositae | Passerina_rositae | diurnal |
| birds | Passerina_versicolor | Passerina_versicolor | diurnal |
| birds | Pastor_roseus | Sturnus_roseus | diurnal |
| birds | Patagioenas_araucana | Patagioenas_araucana | diurnal |
| birds | Patagioenas_caribaea | Patagioenas_caribaea | diurnal |
| birds | Patagioenas_cayennensis | Patagioenas_cayennensis | diurnal |
| birds | Patagioenas_corensis | Patagioenas_corensis | diurnal |
| birds | Patagioenas_fasciata | Patagioenas_fasciata | diurnal |
| birds | Patagioenas_flavirostris | Patagioenas_flavirostris | diurnal |

| | | | |
|---|---|---|---|
| birds | Patagioenas_inornata | Patagioenas_inornata | diurnal |
| birds | Patagioenas_leucocephala | Patagioenas_leucocephala | diurnal |
| birds | Patagioenas_maculosa | Patagioenas_maculosa | diurnal |
| birds | Patagioenas_nigrirostris | Patagioenas_nigrirostris | diurnal |
| birds | Patagioenas_oenops | Patagioenas_oenops | diurnal |
| birds | Patagioenas_picazuro | Patagioenas_picazuro | diurnal |
| birds | Patagioenas_plumbea | Patagioenas_plumbea | diurnal |
| birds | Patagioenas_speciosa | Patagioenas_speciosa | diurnal |
| birds | Patagioenas_squamosa | Patagioenas_squamosa | diurnal |
| birds | Patagioenas_subvinacea | Patagioenas_subvinacea | diurnal |
| birds | Patagona_gigas | Patagona_gigas | diurnal |
| birds | Pauxi_pauxi | Pauxi_pauxi | diurnal |
| birds | Pauxi_unicornis | Pauxi_unicornis | nocturnal |
| birds | Pavo_cristatus | Pavo_cristatus | diurnal |
| birds | Pavo_muticus | Pavo_muticus | diurnal |
| birds | Pedionomus_torquatus | Pedionomus_torquatus | diurnal |
| birds | Pelargopsis_amauroptera | Pelargopsis_amauroptera | diurnal |
| birds | Pelargopsis_capensis | Pelargopsis_capensis | diurnal |
| birds | Pelargopsis_melanorhyncha | Pelargopsis_melanorhyncha | diurnal |
| birds | Pelecanoides_garnotii | Pelecanoides_garnotii | cathemeral |
| birds | Pelecanoides_georgicus | Pelecanoides_georgicus | cathemeral |
| birds | Pelecanoides_magellani | Pelecanoides_magellani | cathemeral |
| birds | Pelecanoides_urinatrix | Pelecanoides_urinatrix | cathemeral |
| birds | Pelecanus_conspicillatus | Pelecanus_conspicillatus | diurnal |
| birds | Pelecanus_crispus | Pelecanus_crispus | diurnal |
| birds | Pelecanus_erythrorhynchos | Pelecanus_erythrorhynchos | diurnal |
| birds | Pelecanus_occidentalis | Pelecanus_occidentalis | diurnal |
| birds | Pelecanus_onocrotalus | Pelecanus_onocrotalus | diurnal |
| birds | Pelecanus_philippensis | Pelecanus_philippensis | diurnal |
| birds | Pelecanus_rufescens | Pelecanus_rufescens | diurnal |
| birds | Peliperdix_lathami | Francolinus_lathami | diurnal |
| birds | Pellorneum_albiventre | Pellorneum_albiventre | diurnal |
| birds | Pellorneum_bicolor | Trichastoma_bicolor | diurnal |
| birds | Pellorneum_capistratum | Pellorneum_capistratum | diurnal |

| | | | |
|---|---|---|---|
| birds | Pellorneum_celebense | Trichastoma_celebense | diurnal |
| birds | Pellorneum_cinereiceps | Malacocincla_cinereiceps | diurnal |
| birds | Pellorneum_fuscocapillus | Pellorneum_fuscocapillus | diurnal |
| birds | Pellorneum_malaccense | Malacocincla_malaccensis | diurnal |
| birds | Pellorneum_pyrrogenys | Trichastoma_pyrrogenys | diurnal |
| birds | Pellorneum_rostratum | Trichastoma_rostratum | diurnal |
| birds | Pellorneum_ruficeps | Pellorneum_ruficeps | diurnal |
| birds | Pellorneum_tickelli | Trichastoma_tickelli | diurnal |
| birds | Peltohyas_australis | Charadrius_australis | nocturnal |
| birds | Peltops_blainvillii | Peltops_blainvillii | diurnal |
| birds | Peltops_montanus | Peltops_montanus | diurnal |
| birds | Penelope_albipennis | Penelope_albipennis | diurnal |
| birds | Penelope_argyrotis | Penelope_argyrotis | diurnal |
| birds | Penelope_barbata | Penelope_barbata | diurnal |
| birds | Penelope_dabbenei | Penelope_dabbenei | diurnal |
| birds | Penelope_jacquacu | Penelope_jacquacu | diurnal |
| birds | Penelope_jacucaca | Penelope_jacucaca | diurnal |
| birds | Penelope_marail | Penelope_marail | diurnal |
| birds | Penelope_montagnii | Penelope_montagnii | diurnal |
| birds | Penelope_obscura | Penelope_obscura | diurnal |
| birds | Penelope_ortoni | Penelope_ortoni | diurnal |
| birds | Penelope_perspicax | Penelope_perspicax | diurnal |
| birds | Penelope_pileata | Penelope_pileata | diurnal |
| birds | Penelope_purpurascens | Penelope_purpurascens | diurnal |
| birds | Penelope_superciliaris | Penelope_superciliaris | diurnal |
| birds | Penelopides_affinis | Penelopides_affinis | diurnal |
| birds | Penelopides_manillae | Penelopides_manillae | diurnal |
| birds | Penelopides_mindorensis | Penelopides_mindorensis | diurnal |
| birds | Penelopides_panini | Penelopides_panini | diurnal |
| birds | Penelopina_nigra | Penelopina_nigra | diurnal |
| birds | Peneoenanthe_pulverulenta | Eopsaltria_pulverulenta | diurnal |
| birds | Peneothello_bimaculata | Peneothello_bimaculata | diurnal |
| birds | Peneothello_cryptoleuca | Peneothello_cryptoleuca | diurnal |
| birds | Peneothello_cyanus | Peneothello_cyanus | diurnal |

| | | | |
|---|---|---|---|
| birds | Peneothello_sigillata | Peneothello_sigillatus | diurnal |
| birds | Percnostola_rufifrons | Percnostola_rufifrons | diurnal |
| birds | Perdicula_argoondah | Perdicula_argoondah | diurnal |
| birds | Perdicula_asiatica | Perdicula_asiatica | diurnal |
| birds | Perdicula_erythrorhyncha | Perdicula_erythrorhyncha | diurnal |
| birds | Perdicula_manipurensis | Perdicula_manipurensis | diurnal |
| birds | Perdix_dauurica | Perdix_dauurica | diurnal |
| birds | Perdix_hodgsoniae | Perdix_hodgsoniae | diurnal |
| birds | Perdix_perdix | Perdix_perdix | diurnal |
| birds | Pericrocotus_brevirostris | Pericrocotus_brevirostris | diurnal |
| birds | Pericrocotus_cantonensis | Pericrocotus_cantonensis | diurnal |
| birds | Pericrocotus_cinnamomeus | Pericrocotus_cinnamomeus | diurnal |
| birds | Pericrocotus_divaricatus | Pericrocotus_divaricatus | diurnal |
| birds | Pericrocotus_erythropygius | Pericrocotus_erythropygius | diurnal |
| birds | Pericrocotus_ethologus | Pericrocotus_ethologus | diurnal |
| birds | Pericrocotus_flammeus | Pericrocotus_flammeus | diurnal |
| birds | Pericrocotus_igneus | Pericrocotus_igneus | diurnal |
| birds | Pericrocotus_lansbergei | Pericrocotus_lansbergei | diurnal |
| birds | Pericrocotus_miniatus | Pericrocotus_miniatus | diurnal |
| birds | Pericrocotus_roseus | Pericrocotus_roseus | diurnal |
| birds | Pericrocotus_solaris | Pericrocotus_solaris | diurnal |
| birds | Pericrocotus_tegimae | Pericrocotus_tegimae | diurnal |
| birds | Periparus_ater | Parus_ater | diurnal |
| birds | Periparus_rubidiventris | Parus_rubidiventris | diurnal |
| birds | Periparus_rufonuchalis | Parus_rufonuchalis | diurnal |
| birds | Perisoreus_canadensis | Perisoreus_canadensis | diurnal |
| birds | Perisoreus_infaustus | Perisoreus_infaustus | diurnal |
| birds | Perisoreus_internigrans | Perisoreus_internigrans | diurnal |
| birds | Perissocephalus_tricolor | Perissocephalus_tricolor | diurnal |
| birds | Pernis_apivorus | Pernis_apivorus | diurnal |
| birds | Pernis_celebensis | Pernis_celebensis | diurnal |
| birds | Pernis_ptilorhynchus | Pernis_ptilorhyncus | diurnal |
| birds | Petrochelidon_ariel | Hirundo_ariel | diurnal |
| birds | Petrochelidon_fluvicola | Hirundo_fluvicola | diurnal |

| | | | |
|---|---|---|---|
| birds | Petrochelidon_fulva | Petrochelidon_fulva | diurnal |
| birds | Petrochelidon_nigricans | Hirundo_nigricans | diurnal |
| birds | Petrochelidon_preussi | Hirundo_preussi | diurnal |
| birds | Petrochelidon_pyrrhonota | Petrochelidon_pyrrhonota | diurnal |
| birds | Petrochelidon_rufigula | Hirundo_rufigula | diurnal |
| birds | Petrochelidon_rufocollaris | Petrochelidon_rufocollaris | diurnal |
| birds | Petrochelidon_spilodera | Hirundo_spilodera | diurnal |
| birds | Petroica_archboldi | Petroica_archboldi | diurnal |
| birds | Petroica_australis | Petroica_australis | diurnal |
| birds | Petroica_bivittata | Petroica_bivittata | diurnal |
| birds | Petroica_goodenovii | Petroica_goodenovii | diurnal |
| birds | Petroica_macrocephala | Petroica_macrocephala | diurnal |
| birds | Petroica_multicolor | Petroica_multicolor | diurnal |
| birds | Petroica_phoenicea | Petroica_phoenicea | diurnal |
| birds | Petroica_rodinogaster | Petroica_rodinogaster | diurnal |
| birds | Petroica_rosea | Petroica_rosea | diurnal |
| birds | Petroica_traversi | Petroica_traversi | diurnal |
| birds | Petronia_petronia | Petronia_petronia | diurnal |
| birds | Petrophassa_albipennis | Petrophassa_albipennis | diurnal |
| birds | Petrophassa_rufipennis | Petrophassa_rufipennis | diurnal |
| birds | Peucaea_aestivalis | Aimophila_aestivalis | diurnal |
| birds | Peucaea_botterii | Aimophila_botterii | diurnal |
| birds | Peucaea_carpalis | Aimophila_carpalis | diurnal |
| birds | Peucaea_cassinii | Aimophila_cassinii | diurnal |
| birds | Peucaea_humeralis | Aimophila_humeralis | diurnal |
| birds | Peucaea_mystacalis | Aimophila_mystacalis | diurnal |
| birds | Peucaea_ruficauda | Aimophila_ruficauda | diurnal |
| birds | Peucaea_sumichrasti | Aimophila_sumichrasti | diurnal |
| birds | Peucedramus_taeniatus | Peucedramus_taeniatus | diurnal |
| birds | Pezopetes_capitalis | Pezopetes_capitalis | diurnal |
| birds | Pezoporus_occidentalis | Pezoporus_occidentalis | diurnal |
| birds | Pezoporus_wallicus | Pezoporus_wallicus | diurnal |
| birds | Phacellodomus_dorsalis | Phacellodomus_dorsalis | diurnal |
| birds | Phacellodomus_erythrophthalmus | Phacellodomus_erythrophthalmus | diurnal |

| | | | |
|---|---|---|---|
| birds | Phacellodomus_inornatus | Phacellodomus_inornatus | diurnal |
| birds | Phacellodomus_maculipectus | Phacellodomus_maculipectus | diurnal |
| birds | Phacellodomus_ruber | Phacellodomus_ruber | diurnal |
| birds | Phacellodomus_rufifrons | Phacellodomus_rufifrons | diurnal |
| birds | Phacellodomus_sibilatrix | Phacellodomus_sibilatrix | diurnal |
| birds | Phacellodomus_striaticeps | Phacellodomus_striaticeps | diurnal |
| birds | Phacellodomus_striaticollis | Phacellodomus_striaticollis | diurnal |
| birds | Phaenicophaeus_curvirostris | Phaenicophaeus_curvirostris | diurnal |
| birds | Phaenicophaeus_diardi | Phaenicophaeus_diardi | diurnal |
| birds | Phaenicophaeus_sumatranus | Phaenicophaeus_sumatranus | diurnal |
| birds | Phaenicophaeus_tristis | Phaenicophaeus_tristis | diurnal |
| birds | Phaenicophaeus_viridirostris | Phaenicophaeus_viridirostris | diurnal |
| birds | Phaenicophilus_palmarum | Phaenicophilus_palmarum | diurnal |
| birds | Phaenicophilus_poliocephalus | Phaenicophilus_poliocephalus | diurnal |
| birds | Phaenostictus_mcleannani | Phaenostictus_mcleannani | diurnal |
| birds | Phaeochroa_cuvierii | Phaeochroa_cuvierii | diurnal |
| birds | Phaeomyias_murina | Phaeomyias_murina | diurnal |
| birds | Phaeomyias_tumbezana | Phaeomyias_tumbezana | diurnal |
| birds | Phaeoptila_sordida | Cynanthus_sordidus | diurnal |
| birds | Phaethornis_anthophilus | Phaethornis_anthophilus | diurnal |
| birds | Phaethornis_atrimentalis | Phaethornis_atrimentalis | diurnal |
| birds | Phaethornis_augusti | Phaethornis_augusti | diurnal |
| birds | Phaethornis_bourcieri | Phaethornis_bourcieri | diurnal |
| birds | Phaethornis_eurynome | Phaethornis_eurynome | diurnal |
| birds | Phaethornis_griseogularis | Phaethornis_griseogularis | diurnal |
| birds | Phaethornis_guy | Phaethornis_guy | diurnal |
| birds | Phaethornis_hispidus | Phaethornis_hispidus | diurnal |
| birds | Phaethornis_idaliae | Phaethornis_idaliae | diurnal |
| birds | Phaethornis_koepckeae | Phaethornis_koepckeae | diurnal |
| birds | Phaethornis_longirostris | Phaethornis_longirostris | diurnal |
| birds | Phaethornis_longuemareus | Phaethornis_longuemareus | diurnal |
| birds | Phaethornis_malaris | Phaethornis_malaris | diurnal |
| birds | Phaethornis_nattereri | Phaethornis_nattereri | diurnal |
| birds | Phaethornis_philippii | Phaethornis_philippii | diurnal |

| | | | |
|---|---|---|---|
| birds | Phaethornis_pretrei | Phaethornis_pretrei | diurnal |
| birds | Phaethornis_ruber | Phaethornis_ruber | diurnal |
| birds | Phaethornis_rupurumii | Phaethornis_rupurumii | diurnal |
| birds | Phaethornis_squalidus | Phaethornis_squalidus | diurnal |
| birds | Phaethornis_striigularis | Phaethornis_striigularis | diurnal |
| birds | Phaethornis_stuarti | Phaethornis_stuarti | diurnal |
| birds | Phaethornis_subochraceus | Phaethornis_subochraceus | diurnal |
| birds | Phaethornis_superciliosus | Phaethornis_superciliosus | diurnal |
| birds | Phaethornis_syrmatophorus | Phaethornis_syrmatophorus | diurnal |
| birds | Phaethornis_yaruqui | Phaethornis_yaruqui | diurnal |
| birds | Phaetusa_simplex | Phaetusa_simplex | diurnal |
| birds | Phainopepla_nitens | Phainopepla_nitens | diurnal |
| birds | Phainoptila_melanoxantha | Phainoptila_melanoxantha | diurnal |
| birds | Phalacrocorax_capensis | Phalacrocorax_capensis | diurnal |
| birds | Phalacrocorax_capillatus | Phalacrocorax_capillatus | diurnal |
| birds | Phalacrocorax_carbo | Phalacrocorax_carbo | diurnal |
| birds | Phalacrocorax_featherstoni | Phalacrocorax_featherstoni | diurnal |
| birds | Phalacrocorax_fuscescens | Phalacrocorax_fuscescens | diurnal |
| birds | Phalacrocorax_fuscicollis | Phalacrocorax_fuscicollis | diurnal |
| birds | Phalacrocorax_neglectus | Phalacrocorax_neglectus | diurnal |
| birds | Phalacrocorax_nigrogularis | Phalacrocorax_nigrogularis | diurnal |
| birds | Phalacrocorax_punctatus | Phalacrocorax_punctatus | diurnal |
| birds | Phalacrocorax_sulcirostris | Phalacrocorax_sulcirostris | diurnal |
| birds | Phalacrocorax_varius | Phalacrocorax_varius | diurnal |
| birds | Phalaenoptilus_nuttallii | Phalaenoptilus_nuttallii | nocturnal |
| birds | Phalaropus_fulicarius | Phalaropus_fulicarius | diurnal |
| birds | Phalaropus_lobatus | Phalaropus_lobatus | diurnal |
| birds | Phalcoboenus_albogularis | Phalcoboenus_albogularis | diurnal |
| birds | Phalcoboenus_australis | Phalcoboenus_australis | diurnal |
| birds | Phalcoboenus_chimango | Milvago_chimango | diurnal |
| birds | Phalcoboenus_megalopterus | Phalcoboenus_megalopterus | diurnal |
| birds | Phapitreron_amethystinus | Phapitreron_amethystinus | diurnal |
| birds | Phapitreron_cinereiceps | Phapitreron_cinereiceps | diurnal |
| birds | Phapitreron_leucotis | Phapitreron_leucotis | diurnal |

| | | | |
|---|---|---|---|
| birds | Phaps_chalcoptera | Phaps_chalcoptera | diurnal |
| birds | Phaps_elegans | Phaps_elegans | diurnal |
| birds | Phaps_histrionica | Phaps_histrionica | diurnal |
| birds | Pharomachrus_antisianus | Pharomachrus_antisianus | diurnal |
| birds | Pharomachrus_auriceps | Pharomachrus_auriceps | diurnal |
| birds | Pharomachrus_fulgidus | Pharomachrus_fulgidus | diurnal |
| birds | Pharomachrus_mocinno | Pharomachrus_mocinno | diurnal |
| birds | Pharomachrus_pavoninus | Pharomachrus_pavoninus | diurnal |
| birds | Phasianus_colchicus | Phasianus_colchicus | diurnal |
| birds | Phasianus_versicolor | Phasianus_versicolor | diurnal |
| birds | Phedina_borbonica | Phedina_borbonica | diurnal |
| birds | Phedinopsis_brazzae | Phedina_brazzae | diurnal |
| birds | Phegornis_mitchellii | Phegornis_mitchellii | diurnal |
| birds | Phelpsia_inornata | Phelpsia_inornatus | diurnal |
| birds | Pheucticus_aureoventris | Pheucticus_aureoventris | diurnal |
| birds | Pheucticus_chrysogaster | Pheucticus_chrysogaster | diurnal |
| birds | Pheucticus_chrysopeplus | Pheucticus_chrysopeplus | diurnal |
| birds | Pheucticus_ludovicianus | Pheucticus_ludovicianus | diurnal |
| birds | Pheucticus_melanocephalus | Pheucticus_melanocephalus | diurnal |
| birds | Pheucticus_tibialis | Pheucticus_tibialis | diurnal |
| birds | Pheugopedius_atrogularis | Thryothorus_atrogularis | diurnal |
| birds | Pheugopedius_coraya | Thryothorus_coraya | diurnal |
| birds | Pheugopedius_eisenmanni | Thryothorus_eisenmanni | diurnal |
| birds | Pheugopedius_euophrys | Thryothorus_euophrys | diurnal |
| birds | Pheugopedius_fasciatoventris | Thryothorus_fasciatoventris | diurnal |
| birds | Pheugopedius_felix | Thryothorus_felix | diurnal |
| birds | Pheugopedius_genibarbis | Thryothorus_genibarbis | diurnal |
| birds | Pheugopedius_maculipectus | Thryothorus_maculipectus | diurnal |
| birds | Pheugopedius_mystacalis | Thryothorus_mystacalis | diurnal |
| birds | Pheugopedius_rutilus | Thryothorus_rutilus | diurnal |
| birds | Pheugopedius_sclateri | Thryothorus_sclateri | diurnal |
| birds | Pheugopedius_spadix | Thryothorus_spadix | diurnal |
| birds | Phibalura_flavirostris | Phibalura_flavirostris | diurnal |
| birds | Phigys_solitarius | Phigys_solitarius | diurnal |

| | | | |
|---|---|---|---|
| birds | Philemon_albitorques | Philemon_albitorques | diurnal |
| birds | Philemon_argenticeps | Philemon_argenticeps | diurnal |
| birds | Philemon_brassi | Philemon_brassi | diurnal |
| birds | Philemon_buceroides | Philemon_buceroides | diurnal |
| birds | Philemon_citreogularis | Philemon_citreogularis | diurnal |
| birds | Philemon_cockerelli | Philemon_cockerelli | diurnal |
| birds | Philemon_corniculatus | Philemon_corniculatus | diurnal |
| birds | Philemon_diemenensis | Philemon_diemenensis | diurnal |
| birds | Philemon_eichhorni | Philemon_eichhorni | diurnal |
| birds | Philemon_fuscicapillus | Philemon_fuscicapillus | diurnal |
| birds | Philemon_inornatus | Philemon_inornatus | diurnal |
| birds | Philemon_kisserensis | Philemon_kisserensis | diurnal |
| birds | Philemon_meyeri | Philemon_meyeri | diurnal |
| birds | Philemon_moluccensis | Philemon_moluccensis | diurnal |
| birds | Philemon_subcorniculatus | Philemon_subcorniculatus | diurnal |
| birds | Philentoma_pyrhoptera | Philentoma_pyrhoptera | diurnal |
| birds | Philentoma_velata | Philentoma_velata | diurnal |
| birds | Philepitta_castanea | Philepitta_castanea | diurnal |
| birds | Philepitta_schlegeli | Philepitta_schlegeli | diurnal |
| birds | Philesturnus_carunculatus | Philesturnus_carunculatus | diurnal |
| birds | Philetairus_socius | Philetairus_socius | diurnal |
| birds | Philodice_bryantae | Calliphlox_bryantae | diurnal |
| birds | Philodice_mitchellii | Calliphlox_mitchellii | diurnal |
| birds | Philohydor_lictor | Pitangus_lictor | diurnal |
| birds | Philortyx_fasciatus | Philortyx_fasciatus | diurnal |
| birds | Philydor_atricapillus | Philydor_atricapillus | diurnal |
| birds | Philydor_erythrocercum | Philydor_erythrocercum | diurnal |
| birds | Philydor_fuscipenne | Philydor_fuscipenne | diurnal |
| birds | Philydor_pyrrhodes | Philydor_pyrrhodes | diurnal |
| birds | Phimosus_infuscatus | Phimosus_infuscatus | diurnal |
| birds | Phlegopsis_borbae | Skutchia_borbae | diurnal |
| birds | Phlegopsis_erythroptera | Phlegopsis_erythroptera | diurnal |
| birds | Phlegopsis_nigromaculata | Phlegopsis_nigromaculata | diurnal |
| birds | Phleocryptes_melanops | Phleocryptes_melanops | diurnal |

| | | | |
|---|---|---|---|
| birds | Phlogophilus_harterti | Phlogophilus_harterti | diurnal |
| birds | Phlogophilus_hemileucurus | Phlogophilus_hemileucurus | diurnal |
| birds | Phodilus_badius | Phodilus_badius | nocturnal |
| birds | Phoebastria_albatrus | Phoebastria_albatrus | cathemeral |
| birds | Phoebastria_immutabilis | Phoebastria_immutabilis | cathemeral |
| birds | Phoebastria_irrorata | Phoebastria_irrorata | cathemeral |
| birds | Phoebastria_nigripes | Phoebastria_nigripes | cathemeral |
| birds | Phoebetria_fusca | Phoebetria_fusca | cathemeral |
| birds | Phoebetria_palpebrata | Phoebetria_palpebrata | cathemeral |
| birds | Phoenicircus_carnifex | Phoenicircus_carnifex | diurnal |
| birds | Phoenicircus_nigricollis | Phoenicircus_nigricollis | diurnal |
| birds | Phoeniconaias_minor | Phoeniconaias_minor | diurnal |
| birds | Phoenicoparrus_andinus | Phoenicoparrus_andinus | diurnal |
| birds | Phoenicoparrus_jamesi | Phoenicoparrus_jamesi | diurnal |
| birds | Phoenicopterus_chilensis | Phoenicopterus_chilensis | diurnal |
| birds | Phoenicopterus_ruber | Phoenicopterus_ruber | diurnal |
| birds | Phoeniculus_bollei | Phoeniculus_bollei | diurnal |
| birds | Phoeniculus_damarensis | Phoeniculus_damarensis | diurnal |
| birds | Phoeniculus_purpureus | Phoeniculus_purpureus | diurnal |
| birds | Phoeniculus_somaliensis | Phoeniculus_somaliensis | diurnal |
| birds | Phoenicurus_alaschanicus | Phoenicurus_alaschanicus | diurnal |
| birds | Phoenicurus_auroreus | Phoenicurus_auroreus | diurnal |
| birds | Phoenicurus_bicolor | Rhyacornis_bicolor | diurnal |
| birds | Phoenicurus_coeruleocephala | Phoenicurus_caeruleocephala | diurnal |
| birds | Phoenicurus_erythrogastrus | Phoenicurus_erythrogastrus | diurnal |
| birds | Phoenicurus_erythronotus | Phoenicurus_erythronotus | diurnal |
| birds | Phoenicurus_frontalis | Phoenicurus_frontalis | diurnal |
| birds | Phoenicurus_fuliginosus | Rhyacornis_fuliginosa | diurnal |
| birds | Phoenicurus_hodgsoni | Phoenicurus_hodgsoni | diurnal |
| birds | Phoenicurus_leucocephalus | Chaimarrornis_leucocephalus | diurnal |
| birds | Phoenicurus_moussieri | Phoenicurus_moussieri | diurnal |
| birds | Phoenicurus_ochruros | Phoenicurus_ochruros | diurnal |
| birds | Phoenicurus_phoenicurus | Phoenicurus_phoenicurus | diurnal |
| birds | Phoenicurus_schisticeps | Phoenicurus_schisticeps | diurnal |

| | | | |
|---|---|---|---|
| birds | Pholia_sharpii | Cinnyricinclus_sharpii | diurnal |
| birds | Pholidornis_rushiae | Pholidornis_rushiae | diurnal |
| birds | Phonipara_canora | Tiaris_canorus | diurnal |
| birds | Phonygammus_keraudrenii | Manucodia_keraudrenii | diurnal |
| birds | Phragmacia_substriata | Phragmacia_substriata | diurnal |
| birds | Phrygilus_atriceps | Phrygilus_atriceps | diurnal |
| birds | Phrygilus_gayi | Phrygilus_gayi | diurnal |
| birds | Phrygilus_patagonicus | Phrygilus_patagonicus | diurnal |
| birds | Phrygilus_punensis | Phrygilus_punensis | diurnal |
| birds | Phylidonyris_niger | Phylidonyris_niger | diurnal |
| birds | Phylidonyris_novaehollandiae | Phylidonyris_novaehollandiae | diurnal |
| birds | Phylidonyris_pyrrhopterus | Phylidonyris_pyrrhopterus | diurnal |
| birds | Phyllaemulor_bracteatus | Nyctibius_bracteatus | nocturnal |
| birds | Phyllanthus_atripennis | Phyllanthus_atripennis | diurnal |
| birds | Phyllastrephus_albigularis | Phyllastrephus_albigularis | diurnal |
| birds | Phyllastrephus_baumanni | Phyllastrephus_baumanni | diurnal |
| birds | Phyllastrephus_cabanisi | Phyllastrephus_cabanisi | diurnal |
| birds | Phyllastrephus_cerviniventris | Phyllastrephus_cerviniventris | diurnal |
| birds | Phyllastrephus_debilis | Phyllastrephus_debilis | diurnal |
| birds | Phyllastrephus_fischeri | Phyllastrephus_fischeri | diurnal |
| birds | Phyllastrephus_flavostriatus | Phyllastrephus_flavostriatus | diurnal |
| birds | Phyllastrephus_fulviventris | Phyllastrephus_fulviventris | diurnal |
| birds | Phyllastrephus_hypochloris | Phyllastrephus_hypochloris | diurnal |
| birds | Phyllastrephus_icterinus | Phyllastrephus_icterinus | diurnal |
| birds | Phyllastrephus_lorenzi | Phyllastrephus_lorenzi | diurnal |
| birds | Phyllastrephus_poensis | Phyllastrephus_poensis | diurnal |
| birds | Phyllastrephus_poliocephalus | Phyllastrephus_poliocephalus | diurnal |
| birds | Phyllastrephus_strepitans | Phyllastrephus_strepitans | diurnal |
| birds | Phyllastrephus_terrestris | Phyllastrephus_terrestris | diurnal |
| birds | Phyllastrephus_xavieri | Phyllastrephus_xavieri | diurnal |
| birds | Phyllergates_cucullatus | Orthotomus_cuculatus | diurnal |
| birds | Phyllergates_heterolaemus | Orthotomus_heterolaemus | diurnal |
| birds | Phyllolais_pulchella | Phyllolais_pulchella | diurnal |
| birds | Phyllomyias_burmeisteri | Phyllomyias_burmeisteri | diurnal |

| | | | |
|---|---|---|---|
| birds | Phyllomyias_cinereiceps | Phyllomyias_cinereiceps | diurnal |
| birds | Phyllomyias_fasciatus | Phyllomyias_fasciatus | diurnal |
| birds | Phyllomyias_griseiceps | Phyllomyias_griseiceps | diurnal |
| birds | Phyllomyias_griseocapilla | Phyllomyias_griseocapilla | diurnal |
| birds | Phyllomyias_nigrocapillus | Phyllomyias_nigrocapillus | diurnal |
| birds | Phyllomyias_plumbeiceps | Phyllomyias_plumbeiceps | diurnal |
| birds | Phyllomyias_reiseri | Phyllomyias_reiseri | diurnal |
| birds | Phyllomyias_sclateri | Phyllomyias_sclateri | diurnal |
| birds | Phyllomyias_urichi | Phyllomyias_urichi | diurnal |
| birds | Phyllomyias_uropygialis | Phyllomyias_uropygialis | diurnal |
| birds | Phyllomyias_virescens | Phyllomyias_virescens | diurnal |
| birds | Phyllomyias_zeledoni | Phyllomyias_zeledoni | diurnal |
| birds | Phylloscartes_beckeri | Phylloscartes_beckeri | diurnal |
| birds | Phylloscartes_ceciliae | Phylloscartes_ceciliae | diurnal |
| birds | Phylloscartes_difficilis | Phylloscartes_difficilis | diurnal |
| birds | Phylloscartes_flaviventris | Phylloscartes_flaviventris | diurnal |
| birds | Phylloscartes_flavovirens | Phylloscartes_flavovirens | diurnal |
| birds | Phylloscartes_gualaquizae | Phylloscartes_gualaquizae | diurnal |
| birds | Phylloscartes_kronei | Phylloscartes_kronei | diurnal |
| birds | Phylloscartes_nigrifrons | Phylloscartes_nigrifrons | diurnal |
| birds | Phylloscartes_oustaleti | Phylloscartes_oustaleti | diurnal |
| birds | Phylloscartes_parkeri | Phylloscartes_parkeri | diurnal |
| birds | Phylloscartes_paulista | Phylloscartes_paulista | diurnal |
| birds | Phylloscartes_roquettei | Phylloscartes_roquettei | diurnal |
| birds | Phylloscartes_superciliaris | Phylloscartes_superciliaris | diurnal |
| birds | Phylloscartes_sylviolus | Phylloscartes_sylviolus | diurnal |
| birds | Phylloscartes_ventralis | Phylloscartes_ventralis | diurnal |
| birds | Phylloscartes_virescens | Phylloscartes_virescens | diurnal |
| birds | Phylloscopus_affinis | Phylloscopus_affinis | diurnal |
| birds | Phylloscopus_amoenus | Phylloscopus_amoenus | diurnal |
| birds | Phylloscopus_armandii | Phylloscopus_armandii | diurnal |
| birds | Phylloscopus_bonelli | Phylloscopus_bonelli | diurnal |
| birds | Phylloscopus_borealis | Phylloscopus_borealis | diurnal |
| birds | Phylloscopus_borealoides | Phylloscopus_borealoides | diurnal |

| | | | |
|---|---|---|---|
| birds | Phylloscopus_budongoensis | Phylloscopus_budongoensis | diurnal |
| birds | Phylloscopus_burkii | Seicercus_burkii | diurnal |
| birds | Phylloscopus_cantator | Phylloscopus_cantator | diurnal |
| birds | Phylloscopus_castaniceps | Seicercus_castaniceps | diurnal |
| birds | Phylloscopus_cebuensis | Phylloscopus_cebuensis | diurnal |
| birds | Phylloscopus_chloronotus | Phylloscopus_chloronotus | diurnal |
| birds | Phylloscopus_collybita | Phylloscopus_collybita | diurnal |
| birds | Phylloscopus_coronatus | Phylloscopus_coronatus | diurnal |
| birds | Phylloscopus_emeiensis | Phylloscopus_emeiensis | diurnal |
| birds | Phylloscopus_fuligiventer | Phylloscopus_fuligiventer | diurnal |
| birds | Phylloscopus_fuscatus | Phylloscopus_fuscatus | diurnal |
| birds | Phylloscopus_grammiceps | Seicercus_grammiceps | diurnal |
| birds | Phylloscopus_griseolus | Phylloscopus_griseolus | diurnal |
| birds | Phylloscopus_hainanus | Phylloscopus_hainanus | diurnal |
| birds | Phylloscopus_herberti | Phylloscopus_herberti | diurnal |
| birds | Phylloscopus_humei | Phylloscopus_humei | diurnal |
| birds | Phylloscopus_ijimae | Phylloscopus_ijimae | diurnal |
| birds | Phylloscopus_inornatus | Phylloscopus_inornatus | diurnal |
| birds | Phylloscopus_intensior | Phylloscopus_davisoni | diurnal |
| birds | Phylloscopus_intermedius | Seicercus_affinis | diurnal |
| birds | Phylloscopus_kansuensis | Phylloscopus_kansuensis | diurnal |
| birds | Phylloscopus_laetus | Phylloscopus_laetus | diurnal |
| birds | Phylloscopus_laurae | Phylloscopus_laurae | diurnal |
| birds | Phylloscopus_maculipennis | Phylloscopus_maculipennis | diurnal |
| birds | Phylloscopus_magnirostris | Phylloscopus_magnirostris | diurnal |
| birds | Phylloscopus_montis | Seicercus_montis | diurnal |
| birds | Phylloscopus_neglectus | Phylloscopus_neglectus | diurnal |
| birds | Phylloscopus_nitidus | Phylloscopus_nitidus | diurnal |
| birds | Phylloscopus_occipitalis | Phylloscopus_occipitalis | diurnal |
| birds | Phylloscopus_olivaceus | Phylloscopus_olivaceus | diurnal |
| birds | Phylloscopus_orientalis | Phylloscopus_orientalis | diurnal |
| birds | Phylloscopus_poliocephalus | Phylloscopus_poliocephalus | diurnal |
| birds | Phylloscopus_poliogenys | Seicercus_poliogenys | diurnal |
| birds | Phylloscopus_presbytes | Phylloscopus_presbytes | diurnal |

| | | | |
|---|---|---|---|
| birds | Phylloscopus_proregulus | Phylloscopus_proregulus | diurnal |
| birds | Phylloscopus_pulcher | Phylloscopus_pulcher | diurnal |
| birds | Phylloscopus_reguloides | Phylloscopus_reguloides | diurnal |
| birds | Phylloscopus_ricketti | Phylloscopus_ricketti | diurnal |
| birds | Phylloscopus_ruficapilla | Phylloscopus_ruficapilla | diurnal |
| birds | Phylloscopus_sarasinorum | Phylloscopus_sarasinorum | diurnal |
| birds | Phylloscopus_schwarzi | Phylloscopus_schwarzi | diurnal |
| birds | Phylloscopus_sibilatrix | Phylloscopus_sibilatrix | diurnal |
| birds | Phylloscopus_sindianus | Phylloscopus_sindianus | diurnal |
| birds | Phylloscopus_subaffinis | Phylloscopus_subaffinis | diurnal |
| birds | Phylloscopus_subviridis | Phylloscopus_subviridis | diurnal |
| birds | Phylloscopus_tenellipes | Phylloscopus_tenellipes | diurnal |
| birds | Phylloscopus_trivirgatus | Phylloscopus_trivirgatus | diurnal |
| birds | Phylloscopus_trochiloides | Phylloscopus_trochiloides | diurnal |
| birds | Phylloscopus_trochilus | Phylloscopus_trochilus | diurnal |
| birds | Phylloscopus_tytleri | Phylloscopus_tytleri | diurnal |
| birds | Phylloscopus_umbrovirens | Phylloscopus_umbrovirens | diurnal |
| birds | Phylloscopus_xanthoschistos | Phylloscopus_xanthoschistos | diurnal |
| birds | Phylloscopus_yunnanensis | Phylloscopus_yunnanensis | diurnal |
| birds | Phytotoma_raimondii | Phytotoma_raimondii | diurnal |
| birds | Phytotoma_rara | Phytotoma_rara | diurnal |
| birds | Phytotoma_rutila | Phytotoma_rutila | diurnal |
| birds | Piaya_cayana | Piaya_cayana | diurnal |
| birds | Piaya_melanogaster | Piaya_melanogaster | diurnal |
| birds | Pica_hudsonia | Pica_hudsonia | diurnal |
| birds | Pica_nutalli | Pica_nuttalli | diurnal |
| birds | Pica_pica | Pica_pica | diurnal |
| birds | Picathartes_gymnocephalus | Picathartes_gymnocephalus | diurnal |
| birds | Picathartes_oreas | Picathartes_oreas | diurnal |
| birds | Picoides_arcticus | Picoides_arcticus | diurnal |
| birds | Picoides_canicapillus | Dendrocopos_canicapillus | diurnal |
| birds | Picoides_kizuki | Dendrocopos_kizuki | diurnal |
| birds | Picoides_maculatus | Dendrocopos_maculatus | diurnal |
| birds | Picoides_moluccensis | Dendrocopos_moluccensis | diurnal |

| | | | |
|---|---|---|---|
| birds | Picoides_temminckii | Dendrocopos_temminckii | diurnal |
| birds | Picoides_tridactylus | Picoides_tridactylus | diurnal |
| birds | Piculus_aurulentus | Piculus_aurulentus | diurnal |
| birds | Piculus_callopterus | Piculus_callopterus | diurnal |
| birds | Piculus_chrysochloros | Piculus_chrysochloros | diurnal |
| birds | Piculus_flavigula | Piculus_flavigula | diurnal |
| birds | Piculus_leucolaemus | Piculus_leucolaemus | diurnal |
| birds | Piculus_simplex | Piculus_simplex | diurnal |
| birds | Picumnus_albosquamatus | Picumnus_albosquamatus | diurnal |
| birds | Picumnus_aurifrons | Picumnus_aurifrons | diurnal |
| birds | Picumnus_castelnau | Picumnus_castelnau | diurnal |
| birds | Picumnus_cinnamomeus | Picumnus_cinnamomeus | diurnal |
| birds | Picumnus_cirratus | Picumnus_cirratus | diurnal |
| birds | Picumnus_dorbignyanus | Picumnus_dorbignyanus | diurnal |
| birds | Picumnus_exilis | Picumnus_exilis | diurnal |
| birds | Picumnus_fulvescens | Picumnus_fulvescens | diurnal |
| birds | Picumnus_fuscus | Picumnus_fuscus | diurnal |
| birds | Picumnus_granadensis | Picumnus_granadensis | diurnal |
| birds | Picumnus_innominatus | Picumnus_innominatus | diurnal |
| birds | Picumnus_lafresnayi | Picumnus_lafresnayi | diurnal |
| birds | Picumnus_limae | Picumnus_limae | diurnal |
| birds | Picumnus_minutissimus | Picumnus_minutissimus | diurnal |
| birds | Picumnus_nebulosus | Picumnus_nebulosus | diurnal |
| birds | Picumnus_nigropunctatus | Picumnus_nigropunctatus | diurnal |
| birds | Picumnus_olivaceus | Picumnus_olivaceus | diurnal |
| birds | Picumnus_pumilus | Picumnus_pumilus | diurnal |
| birds | Picumnus_pygmaeus | Picumnus_pygmaeus | diurnal |
| birds | Picumnus_rufiventris | Picumnus_rufiventris | diurnal |
| birds | Picumnus_sclateri | Picumnus_sclateri | diurnal |
| birds | Picumnus_spilogaster | Picumnus_spilogaster | diurnal |
| birds | Picumnus_squamulatus | Picumnus_squamulatus | diurnal |
| birds | Picumnus_steindachneri | Picumnus_steindachneri | diurnal |
| birds | Picumnus_subtilis | Picumnus_subtilis | diurnal |
| birds | Picumnus_temminckii | Picumnus_temminckii | diurnal |

| | | | |
|---|---|---|---|
| birds | Picumnus_varzeae | Picumnus_varzeae | diurnal |
| birds | Picus_awokera | Picus_awokera | diurnal |
| birds | Picus_canus | Picus_canus | diurnal |
| birds | Picus_chlorolophus | Picus_chlorolophus | diurnal |
| birds | Picus_erythropygius | Picus_erythropygius | diurnal |
| birds | Picus_puniceus | Picus_puniceus | diurnal |
| birds | Picus_squamatus | Picus_squamatus | diurnal |
| birds | Picus_viridanus | Picus_viridanus | diurnal |
| birds | Picus_viridis | Picus_viridis | diurnal |
| birds | Picus_vittatus | Picus_vittatus | diurnal |
| birds | Picus_xanthopygaeus | Picus_xanthopygaeus | diurnal |
| birds | Piezorina_cinerea | Piezorhina_cinerea | diurnal |
| birds | Pilherodius_pileatus | Pilherodius_pileatus | diurnal |
| birds | Pinarochroa_sordida | Cercomela_sordida | diurnal |
| birds | Pinarocorys_erythropygia | Pinarocorys_erythropygia | diurnal |
| birds | Pinarocorys_nigricans | Pinarocorys_nigricans | diurnal |
| birds | Pinaroloxias_inornata | Pinaroloxias_inornata | diurnal |
| birds | Pinarornis_plumosus | Pinarornis_plumosus | diurnal |
| birds | Pinicola_enucleator | Pinicola_enucleator | diurnal |
| birds | Pionites_leucogaster | Pionites_leucogaster | diurnal |
| birds | Pionites_melanocephalus | Pionites_melanocephalus | diurnal |
| birds | Pionopsitta_pileata | Pionopsitta_pileata | diurnal |
| birds | Pionus_chalcopterus | Pionus_chalcopterus | diurnal |
| birds | Pionus_fuscus | Pionus_fuscus | diurnal |
| birds | Pionus_maximiliani | Pionus_maximiliani | diurnal |
| birds | Pionus_menstruus | Pionus_menstruus | diurnal |
| birds | Pionus_senilis | Pionus_senilis | diurnal |
| birds | Pionus_sordidus | Pionus_sordidus | diurnal |
| birds | Pionus_tumultuosus | Pionus_tumultuosus | diurnal |
| birds | Pipile_cujubi | Pipile_cujubi | diurnal |
| birds | Pipile_cumanensis | Pipile_cumanensis | diurnal |
| birds | Pipile_jacutinga | Pipile_jacutinga | diurnal |
| birds | Pipile_pipile | Pipile_pipile | nocturnal |
| birds | Pipilo_chlorurus | Pipilo_chlorurus | diurnal |

| | | | |
|---|---|---|---|
| birds | Pipilo_erythrophthalmus | Pipilo_erythrophthalmus | diurnal |
| birds | Pipilo_maculatus | Pipilo_maculatus | diurnal |
| birds | Pipilo_ocai | Pipilo_ocai | diurnal |
| birds | Pipra_aureola | Pipra_aureola | diurnal |
| birds | Pipra_fasciicauda | Pipra_fasciicauda | diurnal |
| birds | Pipra_filicauda | Pipra_filicauda | diurnal |
| birds | Pipraeidea_bonariensis | Thraupis_bonariensis | diurnal |
| birds | Pipraeidea_melanonota | Pipraeidea_melanonota | diurnal |
| birds | Pipreola_arcuata | Pipreola_arcuata | diurnal |
| birds | Pipreola_aureopectus | Pipreola_aureopectus | diurnal |
| birds | Pipreola_chlorolepidota | Pipreola_chlorolepidota | diurnal |
| birds | Pipreola_formosa | Pipreola_formosa | diurnal |
| birds | Pipreola_frontalis | Pipreola_frontalis | diurnal |
| birds | Pipreola_intermedia | Pipreola_intermedia | diurnal |
| birds | Pipreola_lubomirskii | Pipreola_lubomirskii | diurnal |
| birds | Pipreola_pulchra | Pipreola_pulchra | diurnal |
| birds | Pipreola_riefferii | Pipreola_riefferii | diurnal |
| birds | Pipreola_whitelyi | Pipreola_whitelyi | diurnal |
| birds | Piprites_chloris | Piprites_chloris | diurnal |
| birds | Piprites_griseiceps | Piprites_griseiceps | diurnal |
| birds | Piprites_pileata | Piprites_pileata | diurnal |
| birds | Piranga_bidentata | Piranga_bidentata | diurnal |
| birds | Piranga_erythrocephala | Piranga_erythrocephala | diurnal |
| birds | Piranga_flava | Piranga_flava | diurnal |
| birds | Piranga_hepatica | Piranga_hepatica | diurnal |
| birds | Piranga_leucoptera | Piranga_leucoptera | diurnal |
| birds | Piranga_ludoviciana | Piranga_ludoviciana | diurnal |
| birds | Piranga_olivacea | Piranga_olivacea | diurnal |
| birds | Piranga_roseogularis | Piranga_roseogularis | diurnal |
| birds | Piranga_rubra | Piranga_rubra | diurnal |
| birds | Piranga_rubriceps | Piranga_rubriceps | diurnal |
| birds | Pitangus_sulphuratus | Pitangus_sulphuratus | diurnal |
| birds | Pithecophaga_jefferyi | Pithecophaga_jefferyi | diurnal |
| birds | Pithys_albifrons | Pithys_albifrons | diurnal |

| | | | |
|---|---|---|---|
| birds | Pithys_castaneus | Pithys_castaneus | diurnal |
| birds | Pitohui_dichrous | Pitohui_dichrous | diurnal |
| birds | Pitohui_kirhocephalus | Pitohui_kirhocephalus | diurnal |
| birds | Pitta_anerythra | Pitta_anerythra | diurnal |
| birds | Pitta_angolensis | Pitta_angolensis | diurnal |
| birds | Pitta_brachyura | Pitta_brachyura | diurnal |
| birds | Pitta_elegans | Pitta_elegans | diurnal |
| birds | Pitta_iris | Pitta_iris | diurnal |
| birds | Pitta_maxima | Pitta_maxima | diurnal |
| birds | Pitta_megarhyncha | Pitta_megarhyncha | diurnal |
| birds | Pitta_moluccensis | Pitta_moluccensis | diurnal |
| birds | Pitta_nympha | Pitta_nympha | diurnal |
| birds | Pitta_sordida | Pitta_sordida | diurnal |
| birds | Pitta_steerii | Pitta_steerii | diurnal |
| birds | Pitta_superba | Pitta_superba | diurnal |
| birds | Pitta_versicolor | Pitta_versicolor | diurnal |
| birds | Pittasoma_michleri | Pittasoma_michleri | diurnal |
| birds | Pittasoma_rufopileatum | Pittasoma_rufopileatum | diurnal |
| birds | Pityriasis_gymnocephala | Pityriasis_gymnocephala | diurnal |
| birds | Platalea_ajaja | Platalea_ajaja | diurnal |
| birds | Platalea_alba | Platalea_alba | diurnal |
| birds | Platalea_flavipes | Platalea_flavipes | diurnal |
| birds | Platalea_minor | Platalea_minor | diurnal |
| birds | Platalea_regia | Platalea_regia | cathemeral |
| birds | Platycercus_adscitus | Platycercus_adscitus | diurnal |
| birds | Platycercus_caledonicus | Platycercus_caledonicus | diurnal |
| birds | Platycercus_elegans | Platycercus_elegans | diurnal |
| birds | Platycercus_eximius | Platycercus_eximius | diurnal |
| birds | Platycercus_icterotis | Platycercus_icterotis | diurnal |
| birds | Platycercus_venustus | Platycercus_venustus | diurnal |
| birds | Platylophus_galericulatus | Platylophus_galericulatus | diurnal |
| birds | Platyrinchus_cancrominus | Platyrinchus_cancrominus | diurnal |
| birds | Platyrinchus_coronatus | Platyrinchus_coronatus | diurnal |
| birds | Platyrinchus_flavigularis | Platyrinchus_flavigularis | diurnal |

| | | | |
|---|---|---|---|
| birds | Platyrinchus_leucoryphus | Platyrinchus_leucoryphus | diurnal |
| birds | Platyrinchus_mystaceus | Platyrinchus_mystaceus | diurnal |
| birds | Platyrinchus_platyrhynchos | Platyrinchus_platyrhynchos | diurnal |
| birds | Platyrinchus_saturatus | Platyrinchus_saturatus | diurnal |
| birds | Platysmurus_leucopterus | Platysmurus_leucopterus | diurnal |
| birds | Platyspiza_crassirostris | Platyspiza_crassirostris | diurnal |
| birds | Platysteira_albifrons | Platysteira_albifrons | diurnal |
| birds | Platysteira_cyanea | Platysteira_cyanea | diurnal |
| birds | Platysteira_peltata | Platysteira_peltata | diurnal |
| birds | Plectorhyncha_lanceolata | Plectorhyncha_lanceolata | diurnal |
| birds | Plectrophenax_hyperboreus | Plectrophenax_hyperboreus | diurnal |
| birds | Plectrophenax_nivalis | Plectrophenax_nivalis | diurnal |
| birds | Plectropterus_gambensis | Plectropterus_gambensis | diurnal |
| birds | Plegadis_chihi | Plegadis_chihi | diurnal |
| birds | Plegadis_falcinellus | Plegadis_falcinellus | diurnal |
| birds | Plegadis_ridgwayi | Plegadis_ridgwayi | diurnal |
| birds | Plesiodryas_albonotata | Poecilodryas_albonotata | diurnal |
| birds | Plocepasser_donaldsoni | Plocepasser_donaldsoni | diurnal |
| birds | Plocepasser_mahali | Plocepasser_mahali | diurnal |
| birds | Plocepasser_rufoscapulatus | Plocepasser_rufoscapulatus | diurnal |
| birds | Plocepasser_superciliosus | Plocepasser_superciliosus | diurnal |
| birds | Ploceus_albinucha | Ploceus_albinucha | diurnal |
| birds | Ploceus_alienus | Ploceus_alienus | diurnal |
| birds | Ploceus_angolensis | Ploceus_angolensis | diurnal |
| birds | Ploceus_aurantius | Ploceus_aurantius | diurnal |
| birds | Ploceus_aureonucha | Ploceus_aureonucha | diurnal |
| birds | Ploceus_badius | Ploceus_badius | diurnal |
| birds | Ploceus_baglafecht | Ploceus_baglafecht | diurnal |
| birds | Ploceus_bannermani | Ploceus_bannermani | diurnal |
| birds | Ploceus_batesi | Ploceus_batesi | diurnal |
| birds | Ploceus_benghalensis | Ploceus_benghalensis | diurnal |
| birds | Ploceus_bertrandi | Ploceus_bertrandi | diurnal |
| birds | Ploceus_bicolor | Ploceus_bicolor | diurnal |
| birds | Ploceus_bojeri | Ploceus_bojeri | diurnal |

| | | | |
|---|---|---|---|
| birds | Ploceus_burnieri | Ploceus_burnieri | diurnal |
| birds | Ploceus_capensis | Ploceus_capensis | diurnal |
| birds | Ploceus_castaneiceps | Ploceus_castaneiceps | diurnal |
| birds | Ploceus_castanops | Ploceus_castanops | diurnal |
| birds | Ploceus_cucullatus | Ploceus_cucullatus | diurnal |
| birds | Ploceus_dichrocephalus | Ploceus_dichrocephalus | diurnal |
| birds | Ploceus_dorsomaculatus | Ploceus_dorsomaculatus | diurnal |
| birds | Ploceus_flavipes | Ploceus_flavipes | diurnal |
| birds | Ploceus_galbula | Ploceus_galbula | diurnal |
| birds | Ploceus_golandi | Ploceus_golandi | diurnal |
| birds | Ploceus_grandis | Ploceus_grandis | diurnal |
| birds | Ploceus_heuglini | Ploceus_heuglini | diurnal |
| birds | Ploceus_hypoxanthus | Ploceus_hypoxanthus | diurnal |
| birds | Ploceus_insignis | Ploceus_insignis | diurnal |
| birds | Ploceus_intermedius | Ploceus_intermedius | diurnal |
| birds | Ploceus_jacksoni | Ploceus_jacksoni | diurnal |
| birds | Ploceus_katangae | Ploceus_katangae | diurnal |
| birds | Ploceus_luteolus | Ploceus_luteolus | diurnal |
| birds | Ploceus_manyar | Ploceus_manyar | diurnal |
| birds | Ploceus_megarhynchus | Ploceus_megarhynchus | diurnal |
| birds | Ploceus_melanocephalus | Ploceus_melanocephalus | diurnal |
| birds | Ploceus_melanogaster | Ploceus_melanogaster | diurnal |
| birds | Ploceus_nelicourvi | Ploceus_nelicourvi | diurnal |
| birds | Ploceus_nicolli | Ploceus_nicolli | diurnal |
| birds | Ploceus_nigerrimus | Ploceus_nigerrimus | diurnal |
| birds | Ploceus_nigricollis | Ploceus_nigricollis | diurnal |
| birds | Ploceus_nigrimentus | Ploceus_nigrimentus | diurnal |
| birds | Ploceus_ocularis | Ploceus_ocularis | diurnal |
| birds | Ploceus_olivaceiceps | Ploceus_olivaceiceps | diurnal |
| birds | Ploceus_pelzelni | Ploceus_pelzelni | diurnal |
| birds | Ploceus_philippinus | Ploceus_philippinus | diurnal |
| birds | Ploceus_preussi | Ploceus_preussi | diurnal |
| birds | Ploceus_princeps | Ploceus_princeps | diurnal |
| birds | Ploceus_reichardi | Ploceus_reichardi | diurnal |

| | | | |
|---|---|---|---|
| birds | Ploceus_rubiginosus | Ploceus_rubiginosus | diurnal |
| birds | Ploceus_ruweti | Ploceus_ruweti | diurnal |
| birds | Ploceus_sakalava | Ploceus_sakalava | diurnal |
| birds | Ploceus_sanctithomae | Ploceus_sanctithomae | diurnal |
| birds | Ploceus_spekei | Ploceus_spekei | diurnal |
| birds | Ploceus_spekeoides | Ploceus_spekeoides | diurnal |
| birds | Ploceus_subaureus | Ploceus_subaureus | diurnal |
| birds | Ploceus_subpersonatus | Ploceus_subpersonatus | diurnal |
| birds | Ploceus_superciliosus | Ploceus_superciliosus | diurnal |
| birds | Ploceus_taeniopterus | Ploceus_taeniopterus | diurnal |
| birds | Ploceus_temporalis | Ploceus_temporalis | diurnal |
| birds | Ploceus_tricolor | Ploceus_tricolor | diurnal |
| birds | Ploceus_velatus | Ploceus_velatus | diurnal |
| birds | Ploceus_vitellinus | Ploceus_vitellinus | diurnal |
| birds | Ploceus_weynsi | Ploceus_weynsi | diurnal |
| birds | Ploceus_xanthops | Ploceus_xanthops | diurnal |
| birds | Ploceus_xanthopterus | Ploceus_xanthopterus | diurnal |
| birds | Pluvialis_apricaria | Pluvialis_apricaria | cathemeral |
| birds | Pluvialis_dominica | Pluvialis_dominica | diurnal |
| birds | Pluvialis_fulva | Pluvialis_fulva | diurnal |
| birds | Pluvialis_squatarola | Pluvialis_squatarola | cathemeral |
| birds | Pluvianellus_socialis | Pluvianellus_socialis | diurnal |
| birds | Pluvianus_aegyptius | Pluvianus_aegyptius | diurnal |
| birds | Pnoepyga_albiventer | Pnoepyga_albiventer | diurnal |
| birds | Pnoepyga_pusilla | Pnoepyga_pusilla | diurnal |
| birds | Podargus_ocellatus | Podargus_ocellatus | nocturnal |
| birds | Podargus_papuensis | Podargus_papuensis | cathemeral |
| birds | Podargus_strigoides | Podargus_strigoides | nocturnal |
| birds | Podica_senegalensis | Podica_senegalensis | diurnal |
| birds | Podiceps_auritus | Podiceps_auritus | diurnal |
| birds | Podiceps_cristatus | Podiceps_cristatus | diurnal |
| birds | Podiceps_gallardoi | Podiceps_gallardoi | diurnal |
| birds | Podiceps_major | Podiceps_major | diurnal |
| birds | Podiceps_nigricollis | Podiceps_nigricollis | diurnal |

| | | | |
|---|---|---|---|
| birds | Podiceps_occipitalis | Podiceps_occipitalis | diurnal |
| birds | Podiceps_taczanowskii | Podiceps_taczanowskii | diurnal |
| birds | Podilymbus_podiceps | Podilymbus_podiceps | diurnal |
| birds | Podoces_biddulphi | Podoces_biddulphi | diurnal |
| birds | Podoces_hendersoni | Podoces_hendersoni | diurnal |
| birds | Podoces_panderi | Podoces_panderi | diurnal |
| birds | Podoces_pleskei | Podoces_pleskei | diurnal |
| birds | Poecile_atricapillus | Parus_atricapillus | diurnal |
| birds | Poecile_carolinensis | Parus_carolinensis | diurnal |
| birds | Poecile_cinctus | Parus_cinctus | diurnal |
| birds | Poecile_davidi | Parus_davidi | diurnal |
| birds | Poecile_gambeli | Parus_gambeli | diurnal |
| birds | Poecile_hudsonicus | Parus_hudsonicus | diurnal |
| birds | Poecile_lugubris | Parus_lugubris | diurnal |
| birds | Poecile_montanus | Parus_montanus | diurnal |
| birds | Poecile_palustris | Parus_palustris | diurnal |
| birds | Poecile_rufescens | Parus_rufescens | diurnal |
| birds | Poecile_sclateri | Parus_sclateri | diurnal |
| birds | Poecile_superciliosus | Parus_superciliosus | diurnal |
| birds | Poecilodryas_brachyura | Poecilodryas_brachyura | diurnal |
| birds | Poecilodryas_hypoleuca | Poecilodryas_hypoleuca | diurnal |
| birds | Poecilodryas_superciliosa | Poecilodryas_superciliosa | diurnal |
| birds | Poecilotriccus_albifacies | Poecilotriccus_albifacies | diurnal |
| birds | Poecilotriccus_calopterus | Poecilotriccus_calopterus | diurnal |
| birds | Poecilotriccus_capitalis | Poecilotriccus_capitalis | diurnal |
| birds | Poecilotriccus_fumifrons | Poecilotriccus_fumifrons | diurnal |
| birds | Poecilotriccus_latirostris | Poecilotriccus_latirostris | diurnal |
| birds | Poecilotriccus_luluae | Poecilotriccus_luluae | diurnal |
| birds | Poecilotriccus_plumbeiceps | Poecilotriccus_plumbeiceps | diurnal |
| birds | Poecilotriccus_pulchellus | Poecilotriccus_pulchellus | diurnal |
| birds | Poecilotriccus_ruficeps | Poecilotriccus_ruficeps | diurnal |
| birds | Poecilotriccus_russatus | Poecilotriccus_russatus | diurnal |
| birds | Poecilotriccus_sylvia | Poecilotriccus_sylvia | diurnal |
| birds | Poeoptera_kenricki | Poeoptera_kenricki | diurnal |

| | | | |
|---|---|---|---|
| birds | Poeoptera_lugubris | Poeoptera_lugubris | diurnal |
| birds | Poeoptera_stuhlmanni | Poeoptera_stuhlmanni | diurnal |
| birds | Poephila_acuticauda | Poephila_acuticauda | diurnal |
| birds | Poephila_cincta | Poephila_cincta | diurnal |
| birds | Poephila_personata | Poephila_personata | diurnal |
| birds | Pogoniulus_atroflavus | Pogoniulus_atroflavus | diurnal |
| birds | Pogoniulus_bilineatus | Pogoniulus_bilineatus | diurnal |
| birds | Pogoniulus_chrysoconus | Pogoniulus_chrysoconus | diurnal |
| birds | Pogoniulus_coryphaea | Pogoniulus_coryphaeus | diurnal |
| birds | Pogoniulus_leucomystax | Pogoniulus_leucomystax | diurnal |
| birds | Pogoniulus_pusillus | Pogoniulus_pusillus | diurnal |
| birds | Pogoniulus_scolopaceus | Pogoniulus_scolopaceus | diurnal |
| birds | Pogoniulus_simplex | Pogoniulus_simplex | diurnal |
| birds | Pogoniulus_subsulphureus | Pogoniulus_subsulphureus | diurnal |
| birds | Pogonocichla_stellata | Pogonocichla_stellata | diurnal |
| birds | Pogonornis_bidentatus | Lybius_bidentatus | diurnal |
| birds | Pogonornis_dubius | Lybius_dubius | diurnal |
| birds | Pogonornis_melanopterus | Lybius_melanopterus | diurnal |
| birds | Pogonornis_minor | Lybius_minor | diurnal |
| birds | Pogonornis_rolleti | Lybius_rolleti | diurnal |
| birds | Pogonotriccus_chapmani | Phylloscartes_chapmani | diurnal |
| birds | Pogonotriccus_eximius | Phylloscartes_eximius | diurnal |
| birds | Pogonotriccus_lanyoni | Phylloscartes_lanyoni | diurnal |
| birds | Pogonotriccus_ophthalmicus | Phylloscartes_ophthalmicus | diurnal |
| birds | Pogonotriccus_orbitalis | Phylloscartes_orbitalis | diurnal |
| birds | Pogonotriccus_poecilotis | Phylloscartes_poecilotis | diurnal |
| birds | Pogonotriccus_venezuelanus | Phylloscartes_venezuelanus | diurnal |
| birds | Poicephalus_cryptoxanthus | Poicephalus_cryptoxanthus | diurnal |
| birds | Poicephalus_flavifrons | Poicephalus_flavifrons | diurnal |
| birds | Poicephalus_gulielmi | Poicephalus_gulielmi | diurnal |
| birds | Poicephalus_meyeri | Poicephalus_meyeri | diurnal |
| birds | Poicephalus_robustus | Poicephalus_robustus | diurnal |
| birds | Poicephalus_rueppellii | Poicephalus_rueppellii | diurnal |
| birds | Poicephalus_rufiventris | Poicephalus_rufiventris | diurnal |

| | | | |
|---|---|---|---|
| birds | Poicephalus_senegalus | Poicephalus_senegalus | diurnal |
| birds | Poikilocarbo_gaimardi | Phalacrocorax_gaimardi | diurnal |
| birds | Polemaetus_bellicosus | Polemaetus_bellicosus | diurnal |
| birds | Polihierax_semitorquatus | Polihierax_semitorquatus | diurnal |
| birds | Poliocephalus_poliocephalus | Poliocephalus_poliocephalus | diurnal |
| birds | Poliocephalus_rufopectus | Poliocephalus_rufopectus | diurnal |
| birds | Poliocrania_exsul | Myrmeciza_exsul | diurnal |
| birds | Poliolais_lopezi | Poliolais_lopezi | diurnal |
| birds | Poliolophus_urostictus | Pycnonotus_urostictus | diurnal |
| birds | Polioptila_albiloris | Polioptila_albiloris | diurnal |
| birds | Polioptila_caerulea | Polioptila_caerulea | diurnal |
| birds | Polioptila_californica | Polioptila_californica | diurnal |
| birds | Polioptila_dumicola | Polioptila_dumicola | diurnal |
| birds | Polioptila_guianensis | Polioptila_guianensis | diurnal |
| birds | Polioptila_lactea | Polioptila_lactea | diurnal |
| birds | Polioptila_lembeyei | Polioptila_lembeyei | diurnal |
| birds | Polioptila_nigriceps | Polioptila_nigriceps | diurnal |
| birds | Polioptila_plumbea | Polioptila_plumbea | diurnal |
| birds | Polioptila_schistaceigula | Polioptila_schistaceigula | diurnal |
| birds | Polioxolmis_rufipennis | Polioxolmis_rufipennis | diurnal |
| birds | Polyboroides_typus | Polyboroides_typus | diurnal |
| birds | Polyerata_amabilis | Amazilia_amabilis | diurnal |
| birds | Polyerata_rosenbergi | Amazilia_rosenbergi | diurnal |
| birds | Polyonymus_caroli | Polyonymus_caroli | diurnal |
| birds | Polyplectron_bicalcaratum | Polyplectron_bicalcaratum | diurnal |
| birds | Polyplectron_chalcurum | Polyplectron_chalcurum | diurnal |
| birds | Polyplectron_germaini | Polyplectron_germaini | diurnal |
| birds | Polyplectron_inopinatum | Polyplectron_inopinatum | diurnal |
| birds | Polyplectron_malacense | Polyplectron_malacense | diurnal |
| birds | Polyplectron_napoleonis | Polyplectron_napoleonis | diurnal |
| birds | Polysticta_stelleri | Polysticta_stelleri | diurnal |
| birds | Polystictus_pectoralis | Polystictus_pectoralis | diurnal |
| birds | Polystictus_superciliaris | Polystictus_superciliaris | diurnal |
| birds | Polytelis_alexandrae | Polytelis_alexandrae | diurnal |

| | | | |
|---|---|---|---|
| birds | Polytelis_anthopeplus | Polytelis_anthopeplus | diurnal |
| birds | Polytelis_swainsonii | Polytelis_swainsonii | diurnal |
| birds | Polytmus_guainumbi | Polytmus_guainumbi | diurnal |
| birds | Polytmus_milleri | Polytmus_milleri | diurnal |
| birds | Polytmus_theresiae | Polytmus_theresiae | diurnal |
| birds | Pomarea_dimidiata | Pomarea_dimidiata | diurnal |
| birds | Pomarea_iphis | Pomarea_iphis | diurnal |
| birds | Pomarea_mendozae | Pomarea_mendozae | diurnal |
| birds | Pomarea_nigra | Pomarea_nigra | diurnal |
| birds | Pomarea_whitneyi | Pomarea_whitneyi | diurnal |
| birds | Pomatorhinus_ferruginosus | Pomatorhinus_ferruginosus | diurnal |
| birds | Pomatorhinus_horsfieldii | Pomatorhinus_horsfieldii | diurnal |
| birds | Pomatorhinus_montanus | Pomatorhinus_montanus | diurnal |
| birds | Pomatorhinus_ochraceiceps | Pomatorhinus_ochraceiceps | diurnal |
| birds | Pomatorhinus_ruficollis | Pomatorhinus_ruficollis | diurnal |
| birds | Pomatorhinus_schisticeps | Pomatorhinus_schisticeps | diurnal |
| birds | Pomatorhinus_superciliaris | Xiphirhynchus_superciliaris | diurnal |
| birds | Pomatostomus_halli | Pomatostomus_halli | diurnal |
| birds | Pomatostomus_ruficeps | Pomatostomus_ruficeps | diurnal |
| birds | Pomatostomus_superciliosus | Pomatostomus_superciliosus | diurnal |
| birds | Pomatostomus_temporalis | Pomatostomus_temporalis | diurnal |
| birds | Poodytes_albolimbatus | Megalurus_albolimbatus | diurnal |
| birds | Poodytes_carteri | Eremiornis_carteri | diurnal |
| birds | Poodytes_gramineus | Megalurus_gramineus | diurnal |
| birds | Poodytes_punctatus | Bowdleria_punctata | diurnal |
| birds | Pooecetes_gramineus | Pooecetes_gramineus | diurnal |
| birds | Poospiza_baeri | Compsospiza_baeri | diurnal |
| birds | Poospiza_boliviana | Poospiza_boliviana | diurnal |
| birds | Poospiza_goeringi | Hemispingus_goeringi | diurnal |
| birds | Poospiza_hispaniolensis | Poospiza_hispaniolensis | diurnal |
| birds | Poospiza_nigrorufa | Poospiza_nigrorufa | diurnal |
| birds | Poospiza_ornata | Poospiza_ornata | diurnal |
| birds | Poospiza_rubecula | Poospiza_rubecula | diurnal |
| birds | Poospiza_rufosuperciliaris | Hemispingus_rufosuperciliaris | diurnal |

| | | | |
|---|---|---|---|
| birds | Poospiza_whitii | Poospiza_whitii | diurnal |
| birds | Poospizopsis_caesar | Poospiza_caesar | diurnal |
| birds | Poospizopsis_hypocondria | Poospiza_hypochondria | diurnal |
| birds | Porphyrio_alleni | Porphyrio_alleni | cathemeral |
| birds | Porphyrio_flavirostris | Porphyrio_flavirostris | diurnal |
| birds | Porphyrio_martinicus | Porphyrio_martinica | diurnal |
| birds | Porphyrio_porphyrio | Porphyrio_porphyrio | nocturnal |
| birds | Porphyriops_melanops | Gallinula_melanops | diurnal |
| birds | Porphyrolaema_porphyrolaema | Porphyrolaema_porphyrolaema | diurnal |
| birds | Porphyrospiza_caerulescens | Porphyrospiza_caerulescens | diurnal |
| birds | Porzana_albicollis | Porzana_albicollis | diurnal |
| birds | Porzana_carolina | Porzana_carolina | diurnal |
| birds | Porzana_fasciata | Anurolimnas_fasciatus | diurnal |
| birds | Porzana_fluminea | Porzana_fluminea | diurnal |
| birds | Premnoplex_brunnescens | Premnoplex_brunnescens | diurnal |
| birds | Premnornis_guttuliger | Premnornis_guttuligera | diurnal |
| birds | Primolius_auricollis | Primolius_auricollis | diurnal |
| birds | Primolius_couloni | Primolius_couloni | diurnal |
| birds | Primolius_maracana | Primolius_maracana | diurnal |
| birds | Prinia_atrogularis | Prinia_atrogularis | diurnal |
| birds | Prinia_bairdii | Prinia_bairdii | diurnal |
| birds | Prinia_buchanani | Prinia_buchanani | diurnal |
| birds | Prinia_cinereocapilla | Prinia_cinereocapilla | diurnal |
| birds | Prinia_crinigera | Prinia_crinigera | diurnal |
| birds | Prinia_erythroptera | Heliolais_erythropterus | diurnal |
| birds | Prinia_familiaris | Prinia_familiaris | diurnal |
| birds | Prinia_flavicans | Prinia_flavicans | diurnal |
| birds | Prinia_flaviventris | Prinia_flaviventris | diurnal |
| birds | Prinia_fluviatilis | Prinia_fluviatilis | diurnal |
| birds | Prinia_gracilis | Prinia_gracilis | diurnal |
| birds | Prinia_hodgsonii | Prinia_hodgsonii | diurnal |
| birds | Prinia_inornata | Prinia_inornata | diurnal |
| birds | Prinia_maculosa | Prinia_maculosa | diurnal |
| birds | Prinia_molleri | Prinia_molleri | diurnal |

| | | | |
|---|---|---|---|
| birds | Prinia_polychroa | Prinia_polychroa | diurnal |
| birds | Prinia_rufescens | Prinia_rufescens | diurnal |
| birds | Prinia_rufifrons | Spiloptila_rufifrons | diurnal |
| birds | Prinia_socialis | Prinia_socialis | diurnal |
| birds | Prinia_somalica | Prinia_somalica | diurnal |
| birds | Prinia_subflava | Prinia_subflava | diurnal |
| birds | Prinia_sylvatica | Prinia_sylvatica | diurnal |
| birds | Prioniturus_discurus | Prioniturus_discurus | diurnal |
| birds | Prioniturus_flavicans | Prioniturus_flavicans | diurnal |
| birds | Prioniturus_luconensis | Prioniturus_luconensis | diurnal |
| birds | Prioniturus_mada | Prioniturus_mada | diurnal |
| birds | Prioniturus_montanus | Prioniturus_montanus | diurnal |
| birds | Prioniturus_platenae | Prioniturus_platenae | diurnal |
| birds | Prioniturus_platurus | Prioniturus_platurus | cathemeral |
| birds | Prioniturus_verticalis | Prioniturus_verticalis | diurnal |
| birds | Prioniturus_waterstradti | Prioniturus_waterstradti | diurnal |
| birds | Prionochilus_maculatus | Prionochilus_maculatus | diurnal |
| birds | Prionochilus_olivaceus | Prionochilus_olivaceus | diurnal |
| birds | Prionochilus_percussus | Prionochilus_percussus | diurnal |
| birds | Prionochilus_plateni | Prionochilus_plateni | diurnal |
| birds | Prionochilus_thoracicus | Prionochilus_thoracicus | diurnal |
| birds | Prionochilus_xanthopygius | Prionochilus_xanthopygius | diurnal |
| birds | Prionodura_newtoniana | Prionodura_newtoniana | diurnal |
| birds | Prionops_alberti | Prionops_alberti | diurnal |
| birds | Prionops_caniceps | Prionops_caniceps | diurnal |
| birds | Prionops_plumatus | Prionops_plumatus | diurnal |
| birds | Prionops_poliolophus | Prionops_poliolophus | diurnal |
| birds | Prionops_retzii | Prionops_retzii | diurnal |
| birds | Prionops_rufiventris | Prionops_rufiventris | diurnal |
| birds | Prionops_scopifrons | Prionops_scopifrons | diurnal |
| birds | Priotelus_temnurus | Priotelus_temnurus | diurnal |
| birds | Probosciger_aterrimus | Probosciger_aterrimus | diurnal |
| birds | Procarduelis_nipalensis | Carpodacus_nipalensis | diurnal |
| birds | Procellaria_aequinoctialis | Procellaria_aequinoctialis | cathemeral |

| | | | |
|---|---|---|---|
| birds | Procellaria_cinerea | Procellaria_cinerea | cathemeral |
| birds | Procellaria_parkinsoni | Procellaria_parkinsoni | cathemeral |
| birds | Procellaria_westlandica | Procellaria_westlandica | cathemeral |
| birds | Procnias_albus | Procnias_albus | diurnal |
| birds | Procnias_averano | Procnias_averano | diurnal |
| birds | Procnias_nudicollis | Procnias_nudicollis | diurnal |
| birds | Procnias_tricarunculatus | Procnias_tricarunculatus | diurnal |
| birds | Prodotiscus_insignis | Prodotiscus_insignis | diurnal |
| birds | Prodotiscus_regulus | Prodotiscus_regulus | diurnal |
| birds | Prodotiscus_zambesiae | Prodotiscus_zambesiae | diurnal |
| birds | Progne_chalybea | Progne_chalybea | diurnal |
| birds | Progne_cryptoleuca | Progne_cryptoleuca | diurnal |
| birds | Progne_dominicensis | Progne_dominicensis | diurnal |
| birds | Progne_elegans | Progne_elegans | diurnal |
| birds | Progne_murphyi | Progne_murphyi | diurnal |
| birds | Progne_sinaloae | Progne_sinaloae | diurnal |
| birds | Progne_subis | Progne_subis | diurnal |
| birds | Progne_tapera | Progne_tapera | diurnal |
| birds | Promerops_cafer | Promerops_cafer | diurnal |
| birds | Promerops_gurneyi | Promerops_gurneyi | diurnal |
| birds | Prosopeia_personata | Prosopeia_personata | diurnal |
| birds | Prosopeia_tabuensis | Prosopeia_tabuensis | diurnal |
| birds | Prosthemadera_novaeseelandiae | Prosthemadera_novaeseelandiae | diurnal |
| birds | Protonotaria_citrea | Protonotaria_citrea | diurnal |
| birds | Prunella_atrogularis | Prunella_atrogularis | diurnal |
| birds | Prunella_collaris | Prunella_collaris | diurnal |
| birds | Prunella_fulvescens | Prunella_fulvescens | diurnal |
| birds | Prunella_himalayana | Prunella_himalayana | diurnal |
| birds | Prunella_immaculata | Prunella_immaculata | diurnal |
| birds | Prunella_koslowi | Prunella_koslowi | diurnal |
| birds | Prunella_modularis | Prunella_modularis | diurnal |
| birds | Prunella_montanella | Prunella_montanella | diurnal |
| birds | Prunella_ocularis | Prunella_ocularis | diurnal |
| birds | Prunella_rubeculoides | Prunella_rubeculoides | diurnal |

| | | | |
|---|---|---|---|
| birds | Prunella_rubida | Prunella_rubida | diurnal |
| birds | Prunella_strophiata | Prunella_strophiata | diurnal |
| birds | Psalidoprocne_albiceps | Psalidoprocne_albiceps | diurnal |
| birds | Psalidoprocne_fuliginosa | Psalidoprocne_fuliginosa | diurnal |
| birds | Psalidoprocne_nitens | Psalidoprocne_nitens | diurnal |
| birds | Psalidoprocne_obscura | Psalidoprocne_obscura | diurnal |
| birds | Psaltria_exilis | Psaltria_exilis | diurnal |
| birds | Psaltriparus_minimus | Psaltriparus_minimus | diurnal |
| birds | Psarisomus_dalhousiae | Psarisomus_dalhousiae | diurnal |
| birds | Psarocolius_angustifrons | Psarocolius_angustifrons | diurnal |
| birds | Psarocolius_atrovirens | Psarocolius_atrovirens | diurnal |
| birds | Psarocolius_bifasciatus | Psarocolius_bifasciatus | diurnal |
| birds | Psarocolius_cassini | Psarocolius_cassini | diurnal |
| birds | Psarocolius_decumanus | Psarocolius_decumanus | diurnal |
| birds | Psarocolius_guatimozinus | Psarocolius_guatimozinus | diurnal |
| birds | Psarocolius_montezuma | Psarocolius_montezuma | diurnal |
| birds | Psarocolius_viridis | Psarocolius_viridis | diurnal |
| birds | Psarocolius_wagleri | Psarocolius_wagleri | diurnal |
| birds | Pselliophorus_tibialis | Pselliophorus_tibialis | diurnal |
| birds | Psephotellus_chrysopterygius | Psephotus_chrysopterygius | diurnal |
| birds | Psephotellus_dissimilis | Psephotus_dissimilis | diurnal |
| birds | Psephotellus_varius | Psephotus_varius | diurnal |
| birds | Psephotus_haematonotus | Psephotus_haematonotus | diurnal |
| birds | Pseudasthenes_cactorum | Asthenes_cactorum | diurnal |
| birds | Pseudasthenes_humicola | Asthenes_humicola | diurnal |
| birds | Pseudasthenes_patagonica | Asthenes_patagonica | diurnal |
| birds | Pseudasthenes_steinbachi | Asthenes_steinbachi | diurnal |
| birds | Pseudastur_albicollis | Leucopternis_albicollis | diurnal |
| birds | Pseudastur_occidentalis | Leucopternis_occidentalis | diurnal |
| birds | Pseudelaenia_leucospodia | Pseudelaenia_leucospodia | diurnal |
| birds | Pseudeos_fuscata | Pseudeos_fuscata | diurnal |
| birds | Pseudhirundo_griseopyga | Pseudhirundo_griseopyga | diurnal |
| birds | Pseudibis_davisoni | Pseudibis_davisoni | diurnal |
| birds | Pseudibis_papillosa | Pseudibis_papillosa | diurnal |

| | | | |
|---|---|---|---|
| birds | Pseudobias_wardi | Pseudobias_wardi | diurnal |
| birds | Pseudocalyptomena_graueri | Pseudocalyptomena_graueri | diurnal |
| birds | Pseudochelidon_eurystomina | Pseudochelidon_eurystomina | diurnal |
| birds | Pseudocolaptes_boissonneauii | Pseudocolaptes_boissonneautii | diurnal |
| birds | Pseudocolaptes_lawrencii | Pseudocolaptes_lawrencii | diurnal |
| birds | Pseudocolopteryx_acutipennis | Pseudocolopteryx_acutipennis | diurnal |
| birds | Pseudocolopteryx_flaviventris | Pseudocolopteryx_flaviventris | diurnal |
| birds | Pseudocolopteryx_sclateri | Pseudocolopteryx_sclateri | diurnal |
| birds | Pseudoleistes_guirahuro | Pseudoleistes_guirahuro | diurnal |
| birds | Pseudoleistes_virescens | Pseudoleistes_virescens | diurnal |
| birds | Pseudonestor_xanthophrys | Pseudonestor_xanthophrys | diurnal |
| birds | Pseudonigrita_arnaudi | Pseudonigrita_arnaudi | diurnal |
| birds | Pseudonigrita_cabanisi | Pseudonigrita_cabanisi | diurnal |
| birds | Pseudopipra_pipra | Pipra_pipra | diurnal |
| birds | Pseudopodoces_humilis | Pseudopodoces_humilis | diurnal |
| birds | Pseudorectes_ferrugineus | Pitohui_ferrugineus | diurnal |
| birds | Pseudorectes_incertus | Pitohui_incertus | diurnal |
| birds | Pseudosaltator_rufiventris | Saltator_rufiventris | diurnal |
| birds | Pseudoscops_grammicus | Pseudoscops_grammicus | nocturnal |
| birds | Pseudoseisura_cristata | Pseudoseisura_cristata | diurnal |
| birds | Pseudoseisura_gutturalis | Pseudoseisura_gutturalis | diurnal |
| birds | Pseudoseisura_lophotes | Pseudoseisura_lophotes | diurnal |
| birds | Pseudoseisura_unirufa | Pseudoseisura_unirufa | diurnal |
| birds | Pseudospingus_verticalis | Hemispingus_verticalis | diurnal |
| birds | Pseudospingus_xanthophthalmus | Hemispingus_xanthophthalmus | diurnal |
| birds | Pseudotriccus_pelzelni | Pseudotriccus_pelzelni | diurnal |
| birds | Pseudotriccus_ruficeps | Pseudotriccus_ruficeps | diurnal |
| birds | Pseudotriccus_simplex | Pseudotriccus_simplex | diurnal |
| birds | Psilopogon_armillaris | Megalaima_armillaris | diurnal |
| birds | Psilopogon_asiaticus | Megalaima_asiatica | diurnal |
| birds | Psilopogon_australis | Megalaima_australis | diurnal |
| birds | Psilopogon_chrysopogon | Megalaima_chrysopogon | diurnal |
| birds | Psilopogon_corvinus | Megalaima_corvina | diurnal |
| birds | Psilopogon_faiostrictus | Megalaima_faiostricta | diurnal |

| | | | |
|---|---|---|---|
| birds | Psilopogon_flavifrons | Megalaima_flavifrons | diurnal |
| birds | Psilopogon_franklinii | Megalaima_franklinii | diurnal |
| birds | Psilopogon_haemacephalus | Megalaima_haemacephala | diurnal |
| birds | Psilopogon_henricii | Megalaima_henricii | diurnal |
| birds | Psilopogon_incognitus | Megalaima_incognita | diurnal |
| birds | Psilopogon_javensis | Megalaima_javensis | diurnal |
| birds | Psilopogon_lagrandieri | Megalaima_lagrandieri | diurnal |
| birds | Psilopogon_lineatus | Megalaima_lineata | diurnal |
| birds | Psilopogon_monticola | Megalaima_monticola | diurnal |
| birds | Psilopogon_mystacophanos | Megalaima_mystacophanos | diurnal |
| birds | Psilopogon_oorti | Megalaima_oorti | diurnal |
| birds | Psilopogon_pulcherrimus | Megalaima_pulcherrima | diurnal |
| birds | Psilopogon_pyrolophus | Psilopogon_pyrolophus | diurnal |
| birds | Psilopogon_rafflesii | Megalaima_rafflesii | diurnal |
| birds | Psilopogon_virens | Megalaima_virens | diurnal |
| birds | Psilopogon_viridis | Megalaima_viridis | diurnal |
| birds | Psilopogon_zeylanicus | Megalaima_zeylanica | diurnal |
| birds | Psilopsiagon_aurifrons | Psilopsiagon_aurifrons | diurnal |
| birds | Psilopsiagon_aymara | Psilopsiagon_aymara | diurnal |
| birds | Psilorhamphus_guttatus | Psilorhamphus_guttatus | diurnal |
| birds | Psiloscops_flammeolus | Otus_flammeolus | nocturnal |
| birds | Psittacara_acuticaudatus | Aratinga_acuticaudata | diurnal |
| birds | Psittacara_chloropterus | Aratinga_chloroptera | diurnal |
| birds | Psittacara_erythrogenys | Aratinga_erythrogenys | diurnal |
| birds | Psittacara_euops | Aratinga_euops | diurnal |
| birds | Psittacara_finschi | Aratinga_finschi | diurnal |
| birds | Psittacara_holochlorus | Aratinga_holochlora | diurnal |
| birds | Psittacara_leucophthalmus | Aratinga_leucophthalma | diurnal |
| birds | Psittacara_mitratus | Aratinga_mitrata | diurnal |
| birds | Psittacara_rubritorquis | Aratinga_rubritorquis | diurnal |
| birds | Psittacara_wagleri | Aratinga_wagleri | diurnal |
| birds | Psittacella_brehmii | Psittacella_brehmii | diurnal |
| birds | Psittacella_madaraszi | Psittacella_madaraszi | diurnal |
| birds | Psittacella_modesta | Psittacella_modesta | diurnal |

| | | | |
|---|---|---|---|
| birds | Psittacella_picta | Psittacella_picta | diurnal |
| birds | Psittacula_alexandri | Psittacula_alexandri | diurnal |
| birds | Psittacula_caniceps | Psittacula_caniceps | diurnal |
| birds | Psittacula_derbiana | Psittacula_derbiana | diurnal |
| birds | Psittaculirostris_desmarestii | Psittaculirostris_desmarestii | diurnal |
| birds | Psittaculirostris_edwardsii | Psittaculirostris_edwardsii | diurnal |
| birds | Psittaculirostris_salvadorii | Psittaculirostris_salvadorii | diurnal |
| birds | Psittacus_erithacus | Psittacus_erithacus | diurnal |
| birds | Psitteuteles_goldiei | Psitteuteles_goldiei | diurnal |
| birds | Psitteuteles_versicolor | Psitteuteles_versicolor | diurnal |
| birds | Psittinus_cyanurus | Psittinus_cyanurus | diurnal |
| birds | Psittirostra_psittacea | Psittirostra_psittacea | diurnal |
| birds | Psittrichas_fulgidus | Psittrichas_fulgidus | diurnal |
| birds | Psophia_crepitans | Psophia_crepitans | diurnal |
| birds | Psophia_leucoptera | Psophia_leucoptera | diurnal |
| birds | Psophia_viridis | Psophia_viridis | diurnal |
| birds | Psophocichla_litsitsirupa | Psophocichla_litsitsirupa | diurnal |
| birds | Psophodes_cristatus | Psophodes_cristatus | diurnal |
| birds | Psophodes_nigrogularis | Psophodes_nigrogularis | diurnal |
| birds | Psophodes_occidentalis | Psophodes_occidentalis | diurnal |
| birds | Psophodes_olivaceus | Psophodes_olivaceus | diurnal |
| birds | Pteridophora_alberti | Pteridophora_alberti | diurnal |
| birds | Pternistis_adspersus | Francolinus_adspersus | diurnal |
| birds | Pternistis_afer | Francolinus_afer | diurnal |
| birds | Pternistis_ahantensis | Francolinus_ahantensis | diurnal |
| birds | Pternistis_bicalcaratus | Francolinus_bicalcaratus | diurnal |
| birds | Pternistis_camerunensis | Francolinus_camerunensis | diurnal |
| birds | Pternistis_capensis | Francolinus_capensis | diurnal |
| birds | Pternistis_castaneicollis | Francolinus_castaneicollis | diurnal |
| birds | Pternistis_clappertoni | Francolinus_clappertoni | diurnal |
| birds | Pternistis_erckelii | Francolinus_erckelii | diurnal |
| birds | Pternistis_hartlaubi | Francolinus_hartlaubi | diurnal |
| birds | Pternistis_harwoodi | Francolinus_harwoodi | diurnal |
| birds | Pternistis_hildebrandti | Francolinus_hildebrandti | diurnal |

| | | | |
|---|---|---|---|
| birds | Pternistis_icterorhynchus | Francolinus_icterorhynchus | diurnal |
| birds | Pternistis_jacksoni | Francolinus_jacksoni | diurnal |
| birds | Pternistis_leucoscepus | Francolinus_leucoscepus | diurnal |
| birds | Pternistis_natalensis | Francolinus_natalensis | diurnal |
| birds | Pternistis_ochropectus | Francolinus_ochropectus | diurnal |
| birds | Pternistis_rufopictus | Francolinus_rufopictus | diurnal |
| birds | Pternistis_squamatus | Francolinus_squamatus | diurnal |
| birds | Pternistis_swierstrai | Francolinus_swierstrai | diurnal |
| birds | Pterocles_bicinctus | Pterocles_bicinctus | nocturnal |
| birds | Pterocles_lichtensteinii | Pterocles_lichtensteinii | nocturnal |
| birds | Pterocles_quadricinctus | Pterocles_quadricinctus | nocturnal |
| birds | Pterodroma_alba | Pterodroma_alba | cathemeral |
| birds | Pterodroma_arminjoniana | Pterodroma_arminjoniana | cathemeral |
| birds | Pterodroma_axillaris | Pterodroma_axillaris | cathemeral |
| birds | Pterodroma_cahow | Pterodroma_cahow | cathemeral |
| birds | Pterodroma_cervicalis | Pterodroma_cervicalis | cathemeral |
| birds | Pterodroma_cookii | Pterodroma_cookii | nocturnal |
| birds | Pterodroma_defilippiana | Pterodroma_defilippiana | cathemeral |
| birds | Pterodroma_externa | Pterodroma_externa | cathemeral |
| birds | Pterodroma_feae | Pterodroma_feae | cathemeral |
| birds | Pterodroma_hasitata | Pterodroma_hasitata | cathemeral |
| birds | Pterodroma_hypoleuca | Pterodroma_hypoleuca | cathemeral |
| birds | Pterodroma_inexpectata | Pterodroma_inexpectata | cathemeral |
| birds | Pterodroma_lessonii | Pterodroma_lessonii | cathemeral |
| birds | Pterodroma_leucoptera | Pterodroma_leucoptera | cathemeral |
| birds | Pterodroma_macroptera | Pterodroma_macroptera | cathemeral |
| birds | Pterodroma_mollis | Pterodroma_mollis | cathemeral |
| birds | Pterodroma_neglecta | Pterodroma_neglecta | cathemeral |
| birds | Pterodroma_nigripennis | Pterodroma_nigripennis | cathemeral |
| birds | Pterodroma_phaeopygia | Pterodroma_phaeopygia | cathemeral |
| birds | Pterodroma_pycrofti | Pterodroma_pycrofti | cathemeral |
| birds | Pterodroma_solandri | Pterodroma_solandri | cathemeral |
| birds | Pterodroma_ultima | Pterodroma_ultima | cathemeral |
| birds | Pteroglossus_aracari | Pteroglossus_aracari | diurnal |

| | | | |
|---|---|---|---|
| birds | Pteroglossus_azara | Pteroglossus_azara | diurnal |
| birds | Pteroglossus_bailloni | Pteroglossus_bailloni | diurnal |
| birds | Pteroglossus_beauharnaisii | Pteroglossus_beauharnaesii | diurnal |
| birds | Pteroglossus_bitorquatus | Pteroglossus_bitorquatus | diurnal |
| birds | Pteroglossus_castanotis | Pteroglossus_castanotis | diurnal |
| birds | Pteroglossus_frantzii | Pteroglossus_frantzii | diurnal |
| birds | Pteroglossus_inscriptus | Pteroglossus_inscriptus | diurnal |
| birds | Pteroglossus_pluricinctus | Pteroglossus_pluricinctus | diurnal |
| birds | Pteroglossus_torquatus | Pteroglossus_torquatus | diurnal |
| birds | Pteroglossus_viridis | Pteroglossus_viridis | diurnal |
| birds | Pteronetta_hartlaubii | Pteronetta_hartlaubii | diurnal |
| birds | Pterophanes_cyanopterus | Pterophanes_cyanopterus | diurnal |
| birds | Pteroptochos_castaneus | Pteroptochos_castaneus | diurnal |
| birds | Pteroptochos_megapodius | Pteroptochos_megapodius | diurnal |
| birds | Pteroptochos_tarnii | Pteroptochos_tarnii | diurnal |
| birds | Pterorhinus_caerulatus | Garrulax_caerulatus | diurnal |
| birds | Pterorhinus_chinensis | Garrulax_chinensis | diurnal |
| birds | Pterorhinus_davidi | Garrulax_davidi | diurnal |
| birds | Pterorhinus_delesserti | Garrulax_delesserti | diurnal |
| birds | Pterorhinus_gularis | Garrulax_gularis | diurnal |
| birds | Pterorhinus_koslowi | Babax_koslowi | diurnal |
| birds | Pterorhinus_lanceolatus | Babax_lanceolatus | diurnal |
| birds | Pterorhinus_nuchalis | Garrulax_nuchalis | diurnal |
| birds | Pterorhinus_pectoralis | Garrulax_pectoralis | diurnal |
| birds | Pterorhinus_perspicillatus | Garrulax_perspicillatus | diurnal |
| birds | Pterorhinus_ruficollis | Garrulax_ruficollis | diurnal |
| birds | Pterorhinus_sannio | Garrulax_sannio | diurnal |
| birds | Pterorhinus_waddelli | Babax_waddelli | diurnal |
| birds | Pteruthius_aenobarbus | Pteruthius_aenobarbus | diurnal |
| birds | Pteruthius_flaviscapis | Pteruthius_flaviscapis | diurnal |
| birds | Pteruthius_melanotis | Pteruthius_melanotis | diurnal |
| birds | Pteruthius_rufiventer | Pteruthius_rufiventer | diurnal |
| birds | Pteruthius_xanthochlorus | Pteruthius_xanthochlorus | diurnal |
| birds | Ptilinopus_aurantiifrons | Ptilinopus_aurantiifrons | diurnal |

| | | | |
|---|---|---|---|
| birds | Ptilinopus_cinctus | Ptilinopus_cinctus | diurnal |
| birds | Ptilinopus_coralensis | Ptilinopus_coralensis | diurnal |
| birds | Ptilinopus_coronulatus | Ptilinopus_coronulatus | diurnal |
| birds | Ptilinopus_dupetithouarsii | Ptilinopus_dupetithouarsii | diurnal |
| birds | Ptilinopus_eugeniae | Ptilinopus_eugeniae | diurnal |
| birds | Ptilinopus_greyi | Ptilinopus_greyii | diurnal |
| birds | Ptilinopus_huttoni | Ptilinopus_huttoni | diurnal |
| birds | Ptilinopus_insolitus | Ptilinopus_insolitus | diurnal |
| birds | Ptilinopus_insularis | Ptilinopus_insularis | diurnal |
| birds | Ptilinopus_iozonus | Ptilinopus_iozonus | diurnal |
| birds | Ptilinopus_melanospilus | Ptilinopus_melanospilus | diurnal |
| birds | Ptilinopus_monacha | Ptilinopus_monacha | diurnal |
| birds | Ptilinopus_nainus | Ptilinopus_naina | diurnal |
| birds | Ptilinopus_ornatus | Ptilinopus_ornatus | diurnal |
| birds | Ptilinopus_pelewensis | Ptilinopus_pelewensis | diurnal |
| birds | Ptilinopus_perlatus | Ptilinopus_perlatus | diurnal |
| birds | Ptilinopus_perousii | Ptilinopus_perousii | diurnal |
| birds | Ptilinopus_porphyraceus | Ptilinopus_porphyraceus | diurnal |
| birds | Ptilinopus_porphyreus | Ptilinopus_porphyreus | diurnal |
| birds | Ptilinopus_pulchellus | Ptilinopus_pulchellus | diurnal |
| birds | Ptilinopus_purpuratus | Ptilinopus_purpuratus | diurnal |
| birds | Ptilinopus_regina | Ptilinopus_regina | diurnal |
| birds | Ptilinopus_richardsii | Ptilinopus_richardsii | diurnal |
| birds | Ptilinopus_rivoli | Ptilinopus_rivoli | diurnal |
| birds | Ptilinopus_roseicapilla | Ptilinopus_roseicapilla | diurnal |
| birds | Ptilinopus_solomonensis | Ptilinopus_solomonensis | diurnal |
| birds | Ptilinopus_superbus | Ptilinopus_superbus | diurnal |
| birds | Ptilinopus_tannensis | Ptilinopus_tannensis | diurnal |
| birds | Ptilinopus_viridis | Ptilinopus_viridis | diurnal |
| birds | Ptiliogonys_caudatus | Ptilogonys_caudatus | diurnal |
| birds | Ptiliogonys_cinereus | Ptilogonys_cinereus | diurnal |
| birds | Ptilocichla_leucogrammica | Ptilocichla_leucogrammica | diurnal |
| birds | Ptilocichla_mindanensis | Ptilocichla_mindanensis | diurnal |
| birds | Ptilonorhynchus_violaceus | Ptilonorhynchus_violaceus | diurnal |

| | | | |
|---|---|---|---|
| birds | Ptilopachus_nahani | Francolinus_nahani | diurnal |
| birds | Ptilopachus_petrosus | Ptilopachus_petrosus | diurnal |
| birds | Ptiloprora_erythropleura | Ptiloprora_erythropleura | diurnal |
| birds | Ptiloprora_guisei | Ptiloprora_guisei | diurnal |
| birds | Ptiloprora_mayri | Ptiloprora_mayri | diurnal |
| birds | Ptiloprora_meekiana | Ptiloprora_meekiana | diurnal |
| birds | Ptiloprora_perstriata | Ptiloprora_perstriata | diurnal |
| birds | Ptiloprora_plumbea | Ptiloprora_plumbea | diurnal |
| birds | Ptilopsis_granti | Ptilopsis_granti | nocturnal |
| birds | Ptilopsis_leucotis | Otus_leucotis | nocturnal |
| birds | Ptilorrhoa_caerulescens | Ptilorrhoa_caerulescens | diurnal |
| birds | Ptilorrhoa_castanonota | Ptilorrhoa_castanonota | diurnal |
| birds | Ptilorrhoa_leucosticta | Ptilorrhoa_leucosticta | diurnal |
| birds | Ptilotula_flavescens | Lichenostomus_flavescens | diurnal |
| birds | Ptilotula_fusca | Lichenostomus_fuscus | diurnal |
| birds | Ptilotula_keartlandi | Lichenostomus_keartlandi | diurnal |
| birds | Ptilotula_ornata | Lichenostomus_ornatus | diurnal |
| birds | Ptilotula_penicillata | Lichenostomus_penicillatus | diurnal |
| birds | Ptilotula_plumula | Lichenostomus_plumulus | diurnal |
| birds | Ptiloxena_atroviolacea | Dives_atroviolaceus | diurnal |
| birds | Ptychoramphus_aleuticus | Ptychoramphus_aleuticus | cathemeral |
| birds | Ptyonoprogne_concolor | Hirundo_concolor | diurnal |
| birds | Ptyonoprogne_obsoleta | Hirundo_obsoleta | diurnal |
| birds | Pucrasia_macrolopha | Pucrasia_macrolopha | diurnal |
| birds | Puffinus_gavia | Puffinus_gavia | cathemeral |
| birds | Puffinus_huttoni | Puffinus_huttoni | cathemeral |
| birds | Puffinus_lherminieri | Puffinus_lherminieri | cathemeral |
| birds | Puffinus_nativitatis | Puffinus_nativitatis | cathemeral |
| birds | Puffinus_puffinus | Puffinus_puffinus | cathemeral |
| birds | Pulsatrix_koeniswaldiana | Pulsatrix_koeniswaldiana | nocturnal |
| birds | Pulsatrix_melanota | Pulsatrix_melanota | nocturnal |
| birds | Pulsatrix_perspicillata | Pulsatrix_perspicillata | nocturnal |
| birds | Purnella_albifrons | Phylidonyris_albifrons | diurnal |
| birds | Purpureicephalus_spurius | Purpureicephalus_spurius | diurnal |

| | | | |
|---|---|---|---|
| birds | Pycnonotus_aurigaster | Pycnonotus_aurigaster | diurnal |
| birds | Pycnonotus_barbatus | Pycnonotus_barbatus | diurnal |
| birds | Pycnonotus_blanfordi | Pycnonotus_blanfordi | diurnal |
| birds | Pycnonotus_brunneus | Pycnonotus_brunneus | diurnal |
| birds | Pycnonotus_cafer | Pycnonotus_cafer | diurnal |
| birds | Pycnonotus_capensis | Pycnonotus_capensis | diurnal |
| birds | Pycnonotus_finlaysoni | Pycnonotus_finlaysoni | diurnal |
| birds | Pycnonotus_flavescens | Pycnonotus_flavescens | diurnal |
| birds | Pycnonotus_goiavier | Pycnonotus_goiavier | diurnal |
| birds | Pycnonotus_jocosus | Pycnonotus_jocosus | diurnal |
| birds | Pycnonotus_leucogenys | Pycnonotus_leucogenys | diurnal |
| birds | Pycnonotus_leucotis | Pycnonotus_leucotis | diurnal |
| birds | Pycnonotus_luteolus | Pycnonotus_luteolus | diurnal |
| birds | Pycnonotus_nigricans | Pycnonotus_nigricans | diurnal |
| birds | Pycnonotus_penicillatus | Pycnonotus_penicillatus | diurnal |
| birds | Pycnonotus_plumosus | Pycnonotus_plumosus | diurnal |
| birds | Pycnonotus_simplex | Pycnonotus_simplex | diurnal |
| birds | Pycnonotus_sinensis | Pycnonotus_sinensis | diurnal |
| birds | Pycnonotus_taivanus | Pycnonotus_taivanus | diurnal |
| birds | Pycnonotus_tympanistrigus | Pycnonotus_tympanistrigus | diurnal |
| birds | Pycnonotus_xantholaemus | Pycnonotus_xantholaemus | diurnal |
| birds | Pycnonotus_xanthopygos | Pycnonotus_xanthopygos | diurnal |
| birds | Pycnonotus_xanthorrhous | Pycnonotus_xanthorrhous | diurnal |
| birds | Pycnoptilus_floccosus | Pycnoptilus_floccosus | diurnal |
| birds | Pycnopygius_cinereus | Pycnopygius_cinereus | diurnal |
| birds | Pycnopygius_ixoides | Pycnopygius_ixoides | diurnal |
| birds | Pycnopygius_stictocephalus | Pycnopygius_stictocephalus | diurnal |
| birds | Pygarrhichas_albogularis | Pygarrhichas_albogularis | diurnal |
| birds | Pygiptila_stellaris | Pygiptila_stellaris | diurnal |
| birds | Pygochelidon_cyanoleuca | Pygochelidon_cyanoleuca | diurnal |
| birds | Pygochelidon_melanoleuca | Atticora_melanoleuca | diurnal |
| birds | Pygoscelis_adeliae | Pygoscelis_adeliae | cathemeral |
| birds | Pygoscelis_antarcticus | Pygoscelis_antarcticus | cathemeral |
| birds | Pygoscelis_papua | Pygoscelis_papua | cathemeral |

| | | | |
|---|---|---|---|
| birds | Pyrenestes_minor | Pyrenestes_minor | diurnal |
| birds | Pyrenestes_ostrinus | Pyrenestes_ostrinus | diurnal |
| birds | Pyrenestes_sanguineus | Pyrenestes_sanguineus | diurnal |
| birds | Pyrgilauda_davidiana | Montifringilla_davidiana | diurnal |
| birds | Pyrgilauda_ruficollis | Montifringilla_ruficollis | diurnal |
| birds | Pyrgilauda_theresae | Montifringilla_theresae | diurnal |
| birds | Pyriglena_atra | Pyriglena_atra | diurnal |
| birds | Pyriglena_leuconota | Pyriglena_leuconota | diurnal |
| birds | Pyriglena_leucoptera | Pyriglena_leucoptera | diurnal |
| birds | Pyrilia_barrabandi | Pyrilia_barrabandi | diurnal |
| birds | Pyrilia_caica | Pyrilia_caica | diurnal |
| birds | Pyrilia_haematotis | Pyrilia_haematotis | diurnal |
| birds | Pyrilia_pulchra | Pyrilia_pulchra | diurnal |
| birds | Pyrilia_vulturina | Pyrilia_vulturina | diurnal |
| birds | Pyrocephalus_rubinus | Pyrocephalus_rubinus | diurnal |
| birds | Pyroderus_scutatus | Pyroderus_scutatus | diurnal |
| birds | Pyrrhocorax_graculus | Pyrrhocorax_graculus | diurnal |
| birds | Pyrrhocorax_pyrrhocorax | Pyrrhocorax_pyrrhocorax | diurnal |
| birds | Pyrrholaemus_brunneus | Pyrrholaemus_brunneus | diurnal |
| birds | Pyrrholaemus_sagittatus | Chthonicola_sagittatus | diurnal |
| birds | Pyrrhomyias_cinnamomeus | Pyrrhomyias_cinnamomeus | diurnal |
| birds | Pyrrhoplectes_epauletta | Pyrrhoplectes_epauletta | diurnal |
| birds | Pyrrhula_aurantiaca | Pyrrhula_aurantiaca | diurnal |
| birds | Pyrrhula_erythaca | Pyrrhula_erythaca | diurnal |
| birds | Pyrrhula_erythrocephala | Pyrrhula_erythrocephala | diurnal |
| birds | Pyrrhula_leucogenis | Pyrrhula_leucogenis | diurnal |
| birds | Pyrrhula_murina | Pyrrhula_murina | diurnal |
| birds | Pyrrhula_nipalensis | Pyrrhula_nipalensis | diurnal |
| birds | Pyrrhula_pyrrhula | Pyrrhula_pyrrhula | diurnal |
| birds | Pyrrhura_albipectus | Pyrrhura_albipectus | diurnal |
| birds | Pyrrhura_cruentata | Pyrrhura_cruentata | diurnal |
| birds | Pyrrhura_egregia | Pyrrhura_egregia | diurnal |
| birds | Pyrrhura_frontalis | Pyrrhura_frontalis | diurnal |
| birds | Pyrrhura_hoematotis | Pyrrhura_hoematotis | diurnal |

| | | | |
|---|---|---|---|
| birds | Pyrrhura_hoffmanni | Pyrrhura_hoffmanni | diurnal |
| birds | Pyrrhura_lepida | Pyrrhura_lepida | diurnal |
| birds | Pyrrhura_leucotis | Pyrrhura_leucotis | diurnal |
| birds | Pyrrhura_melanura | Pyrrhura_melanura | diurnal |
| birds | Pyrrhura_molinae | Pyrrhura_molinae | diurnal |
| birds | Pyrrhura_orcesi | Pyrrhura_orcesi | diurnal |
| birds | Pyrrhura_perlata | Pyrrhura_perlata | diurnal |
| birds | Pyrrhura_picta | Pyrrhura_picta | diurnal |
| birds | Pyrrhura_rupicola | Pyrrhura_rupicola | diurnal |
| birds | Pyrrhura_viridicata | Pyrrhura_viridicata | diurnal |
| birds | Pyrrhurus_scandens | Pyrrhurus_scandens | diurnal |
| birds | Pytilia_afra | Pytilia_afra | diurnal |
| birds | Pytilia_hypogrammica | Pytilia_hypogrammica | diurnal |
| birds | Pytilia_lineata | Pytilia_lineata | diurnal |
| birds | Pytilia_melba | Pytilia_melba | diurnal |
| birds | Pytilia_phoenicoptera | Pytilia_phoenicoptera | diurnal |
| birds | Quelea_cardinalis | Quelea_cardinalis | diurnal |
| birds | Quelea_erythrops | Quelea_erythrops | diurnal |
| birds | Quelea_quelea | Quelea_quelea | diurnal |
| birds | Querula_purpurata | Querula_purpurata | diurnal |
| birds | Quiscalus_lugubris | Quiscalus_lugubris | diurnal |
| birds | Quiscalus_major | Quiscalus_major | diurnal |
| birds | Quiscalus_mexicanus | Quiscalus_mexicanus | diurnal |
| birds | Quiscalus_nicaraguensis | Quiscalus_nicaraguensis | diurnal |
| birds | Quiscalus_niger | Quiscalus_niger | diurnal |
| birds | Quiscalus_quiscula | Quiscalus_quiscula | diurnal |
| birds | Quoyornis_georgianus | Eopsaltria_georgiana | diurnal |
| birds | Radjah_radjah | Tadorna_radjah | diurnal |
| birds | Rallicula_forbesi | Rallina_forbesi | diurnal |
| birds | Rallicula_leucospila | Rallina_leucospila | diurnal |
| birds | Rallicula_mayri | Rallina_mayri | diurnal |
| birds | Rallicula_rubra | Rallina_rubra | diurnal |
| birds | Rallina_eurizonoides | Rallina_eurizonoides | cathemeral |
| birds | Rallina_fasciata | Rallina_fasciata | diurnal |

| | | | |
|---|---|---|---|
| birds | Rallina_tricolor | Rallina_tricolor | cathemeral |
| birds | Rallus_antarcticus | Rallus_antarcticus | cathemeral |
| birds | Rallus_aquaticus | Rallus_aquaticus | diurnal |
| birds | Rallus_elegans | Rallus_elegans | diurnal |
| birds | Rallus_longirostris | Rallus_longirostris | cathemeral |
| birds | Rallus_madagascariensis | Rallus_madagascariensis | diurnal |
| birds | Ramphastos_ambiguus | Ramphastos_ambiguus | diurnal |
| birds | Ramphastos_brevis | Ramphastos_brevis | diurnal |
| birds | Ramphastos_dicolorus | Ramphastos_dicolorus | diurnal |
| birds | Ramphastos_sulfuratus | Ramphastos_sulfuratus | diurnal |
| birds | Ramphastos_toco | Ramphastos_toco | diurnal |
| birds | Ramphastos_tucanus | Ramphastos_tucanus | diurnal |
| birds | Ramphastos_vitellinus | Ramphastos_vitellinus | diurnal |
| birds | Ramphiculus_fischeri | Ptilinopus_fischeri | diurnal |
| birds | Ramphiculus_jambu | Ptilinopus_jambu | diurnal |
| birds | Ramphiculus_leclancheri | Ptilinopus_leclancheri | diurnal |
| birds | Ramphiculus_marchei | Ptilinopus_marchei | diurnal |
| birds | Ramphiculus_merrilli | Ptilinopus_merrilli | diurnal |
| birds | Ramphiculus_occipitalis | Ptilinopus_occipitalis | diurnal |
| birds | Ramphiculus_subgularis | Ptilinopus_subgularis | diurnal |
| birds | Ramphocaenus_melanurus | Ramphocaenus_melanurus | diurnal |
| birds | Ramphocelus_bresilius | Ramphocelus_bresilius | diurnal |
| birds | Ramphocelus_carbo | Ramphocelus_carbo | diurnal |
| birds | Ramphocelus_dimidiatus | Ramphocelus_dimidiatus | diurnal |
| birds | Ramphocelus_flammigerus | Ramphocelus_flammigerus | diurnal |
| birds | Ramphocelus_melanogaster | Ramphocelus_melanogaster | diurnal |
| birds | Ramphocelus_nigrogularis | Ramphocelus_nigrogularis | diurnal |
| birds | Ramphocelus_passerinii | Ramphocelus_passerinii | diurnal |
| birds | Ramphocelus_sanguinolentus | Ramphocelus_sanguinolentus | diurnal |
| birds | Ramphocinclus_brachyurus | Ramphocinclus_brachyurus | diurnal |
| birds | Ramphocoris_clotbey | Rhamphocoris_clotbey | diurnal |
| birds | Ramphodon_naevius | Ramphodon_naevius | diurnal |
| birds | Ramphomicron_dorsale | Ramphomicron_dorsale | diurnal |
| birds | Ramphomicron_microrhynchum | Ramphomicron_microrhynchum | diurnal |

| | | | |
|---|---|---|---|
| birds | Ramphotrigon_fuscicauda | Ramphotrigon_fuscicauda | diurnal |
| birds | Ramphotrigon_megacephalum | Ramphotrigon_megacephalum | diurnal |
| birds | Ramphotrigon_ruficauda | Ramphotrigon_ruficauda | diurnal |
| birds | Ramsayornis_fasciatus | Ramsayornis_fasciatus | diurnal |
| birds | Ramsayornis_modestus | Ramsayornis_modestus | diurnal |
| birds | Randia_pseudozosterops | Randia_pseudozosterops | diurnal |
| birds | Recurvirostra_americana | Recurvirostra_americana | diurnal |
| birds | Recurvirostra_andina | Recurvirostra_andina | diurnal |
| birds | Recurvirostra_avosetta | Recurvirostra_avosetta | diurnal |
| birds | Recurvirostra_novaehollandiae | Recurvirostra_novaehollandiae | diurnal |
| birds | Regulus_goodfellowi | Regulus_goodfellowi | diurnal |
| birds | Regulus_regulus | Regulus_regulus | diurnal |
| birds | Regulus_satrapa | Regulus_satrapa | diurnal |
| birds | Reinwardtoena_browni | Reinwardtoena_browni | diurnal |
| birds | Reinwardtoena_crassirostris | Reinwardtoena_crassirostris | diurnal |
| birds | Reinwardtoena_reinwardti | Reinwardtoena_reinwardtsi | diurnal |
| birds | Remiz_consobrinus | Remiz_consobrinus | diurnal |
| birds | Remiz_coronatus | Remiz_coronatus | diurnal |
| birds | Remiz_pendulinus | Remiz_pendulinus | diurnal |
| birds | Rhabdornis_grandis | Rhabdornis_grandis | diurnal |
| birds | Rhabdornis_inornatus | Rhabdornis_inornatus | diurnal |
| birds | Rhabdornis_mystacalis | Rhabdornis_mystacalis | diurnal |
| birds | Rhabdotorrhinus_corrugatus | Aceros_corrugatus | diurnal |
| birds | Rhabdotorrhinus_exarhatus | Penelopides_exarhatus | diurnal |
| birds | Rhabdotorrhinus_leucocephalus | Aceros_leucocephalus | diurnal |
| birds | Rhagologus_leucostigma | Rhagologus_leucostigma | diurnal |
| birds | Rhamphocharis_crassirostris | Melanocharis_crassirostris | diurnal |
| birds | Rhaphidura_leucopygialis | Rhaphidura_leucopygialis | diurnal |
| birds | Rhaphidura_sabini | Rhaphidura_sabini | diurnal |
| birds | Rhea_americana | Rhea_americana | diurnal |
| birds | Rhea_pennata | Rhea_pennata | diurnal |
| birds | Rhegmatorhina_berlepschi | Rhegmatorhina_berlepschi | diurnal |
| birds | Rhegmatorhina_cristata | Rhegmatorhina_cristata | diurnal |
| birds | Rhegmatorhina_gymnops | Rhegmatorhina_gymnops | diurnal |

| | | | |
|---|---|---|---|
| birds | Rhegmatorhina_hoffmannsi | Rhegmatorhina_hoffmannsi | diurnal |
| birds | Rhegmatorhina_melanosticta | Rhegmatorhina_melanosticta | diurnal |
| birds | Rheinardia_ocellata | Rheinardia_ocellata | diurnal |
| birds | Rhinocrypta_lanceolata | Rhinocrypta_lanceolata | diurnal |
| birds | Rhinopomastus_aterrimus | Rhinopomastus_aterrimus | diurnal |
| birds | Rhinopomastus_castaneiceps | Phoeniculus_castaneiceps | diurnal |
| birds | Rhinopomastus_cyanomelas | Rhinopomastus_cyanomelas | diurnal |
| birds | Rhinopomastus_minor | Rhinopomastus_minor | diurnal |
| birds | Rhinoptilus_chalcopterus | Rhinoptilus_chalcopterus | nocturnal |
| birds | Rhinoptilus_cinctus | Rhinoptilus_cinctus | nocturnal |
| birds | Rhinortha_chlorophaea | Phaenicophaeus_chlorophaeus | diurnal |
| birds | Rhipidura_albicollis | Rhipidura_albicollis | diurnal |
| birds | Rhipidura_albolimbata | Rhipidura_albolimbata | diurnal |
| birds | Rhipidura_atra | Rhipidura_atra | diurnal |
| birds | Rhipidura_aureola | Rhipidura_aureola | diurnal |
| birds | Rhipidura_brachyrhyncha | Rhipidura_brachyrhyncha | diurnal |
| birds | Rhipidura_cockerelli | Rhipidura_cockerelli | diurnal |
| birds | Rhipidura_cyaniceps | Rhipidura_cyaniceps | diurnal |
| birds | Rhipidura_dahli | Rhipidura_dahli | diurnal |
| birds | Rhipidura_dedemi | Rhipidura_dedemi | diurnal |
| birds | Rhipidura_diluta | Rhipidura_diluta | diurnal |
| birds | Rhipidura_drownei | Rhipidura_drownei | diurnal |
| birds | Rhipidura_euryura | Rhipidura_euryura | diurnal |
| birds | Rhipidura_fuliginosa | Rhipidura_fuliginosa | diurnal |
| birds | Rhipidura_fuscorufa | Rhipidura_fuscorufa | diurnal |
| birds | Rhipidura_hyperythra | Rhipidura_hyperythra | diurnal |
| birds | Rhipidura_javanica | Rhipidura_javanica | diurnal |
| birds | Rhipidura_kubaryi | Rhipidura_kubaryi | diurnal |
| birds | Rhipidura_lepida | Rhipidura_lepida | diurnal |
| birds | Rhipidura_leucophrys | Rhipidura_leucophrys | diurnal |
| birds | Rhipidura_leucothorax | Rhipidura_leucothorax | diurnal |
| birds | Rhipidura_maculipectus | Rhipidura_maculipectus | diurnal |
| birds | Rhipidura_malaitae | Rhipidura_malaitae | diurnal |
| birds | Rhipidura_matthiae | Rhipidura_matthiae | diurnal |

| | | | |
|---|---|---|---|
| birds | Rhipidura_nebulosa | Rhipidura_nebulosa | diurnal |
| birds | Rhipidura_nigrocinnamomea | Rhipidura_nigrocinnamomea | diurnal |
| birds | Rhipidura_opistherythra | Rhipidura_opistherythra | diurnal |
| birds | Rhipidura_perlata | Rhipidura_perlata | diurnal |
| birds | Rhipidura_personata | Rhipidura_personata | diurnal |
| birds | Rhipidura_phasiana | Rhipidura_phasiana | diurnal |
| birds | Rhipidura_phoenicura | Rhipidura_phoenicura | diurnal |
| birds | Rhipidura_rennelliana | Rhipidura_rennelliana | diurnal |
| birds | Rhipidura_rufidorsa | Rhipidura_rufidorsa | diurnal |
| birds | Rhipidura_rufifrons | Rhipidura_rufifrons | diurnal |
| birds | Rhipidura_rufiventris | Rhipidura_rufiventris | diurnal |
| birds | Rhipidura_semirubra | Rhipidura_semirubra | diurnal |
| birds | Rhipidura_spilodera | Rhipidura_verreauxi | diurnal |
| birds | Rhipidura_superciliaris | Rhipidura_superciliaris | diurnal |
| birds | Rhipidura_superflua | Rhipidura_superflua | diurnal |
| birds | Rhipidura_tenebrosa | Rhipidura_tenebrosa | diurnal |
| birds | Rhipidura_teysmanni | Rhipidura_teysmanni | diurnal |
| birds | Rhipidura_threnothorax | Rhipidura_threnothorax | diurnal |
| birds | Rhizothera_longirostris | Rhizothera_longirostris | diurnal |
| birds | Rhodinocichla_rosea | Rhodinocichla_rosea | diurnal |
| birds | Rhodopechys_sanguineus | Rhodopechys_sanguineus | diurnal |
| birds | Rhodophoneus_cruentus | Rhodophoneus_cruentus | diurnal |
| birds | Rhodopis_vesper | Rhodopis_vesper | diurnal |
| birds | Rhodospingus_cruentus | Rhodospingus_cruentus | diurnal |
| birds | Rhodospiza_obsoleta | Rhodopechys_obsoletus | diurnal |
| birds | Rhodostethia_rosea | Rhodostethia_rosea | diurnal |
| birds | Rhopias_gularis | Myrmotherula_gularis | diurnal |
| birds | Rhopophilus_pekinensis | Rhopophilus_pekinensis | diurnal |
| birds | Rhopornis_ardesiacus | Rhopornis_ardesiacus | diurnal |
| birds | Rhopospina_fruticeti | Phrygilus_fruticeti | diurnal |
| birds | Rhynchocyclus_brevirostris | Rhynchocyclus_brevirostris | diurnal |
| birds | Rhynchocyclus_fulvipectus | Rhynchocyclus_fulvipectus | diurnal |
| birds | Rhynchocyclus_olivaceus | Rhynchocyclus_olivaceus | diurnal |
| birds | Rhynchocyclus_pacificus | Rhynchocyclus_pacificus | diurnal |

| | | | |
|---|---|---|---|
| birds | Rhynchophanes_mccownii | Calcarius_mccownii | diurnal |
| birds | Rhynchopsitta_pachyrhyncha | Rhynchopsitta_pachyrhyncha | diurnal |
| birds | Rhynchopsitta_terrisi | Rhynchopsitta_terrisi | diurnal |
| birds | Rhynchortyx_cinctus | Rhynchortyx_cinctus | diurnal |
| birds | Rhynchospiza_stolzmanni | Aimophila_stolzmanni | diurnal |
| birds | Rhynchospiza_strigiceps | Aimophila_strigiceps | diurnal |
| birds | Rhynchostruthus_socotranus | Rhynchostruthus_socotranus | diurnal |
| birds | Rhynchotus_rufescens | Rhynchotus_rufescens | diurnal |
| birds | Rhynochetos_jubatus | Rhynochetos_jubatus | diurnal |
| birds | Rhyticeros_cassidix | Aceros_cassidix | diurnal |
| birds | Rhyticeros_narcondami | Aceros_narcondami | diurnal |
| birds | Rhyticeros_plicatus | Aceros_plicatus | diurnal |
| birds | Rhyticeros_subruficollis | Aceros_subruficollis | diurnal |
| birds | Rhyticeros_undulatus | Aceros_undulatus | diurnal |
| birds | Rhytipterna_holerythra | Rhytipterna_holerythra | diurnal |
| birds | Rhytipterna_immunda | Rhytipterna_immunda | diurnal |
| birds | Rhytipterna_simplex | Rhytipterna_simplex | diurnal |
| birds | Riccordia_bicolor | Cyanophaia_bicolor | diurnal |
| birds | Riccordia_maugaeus | Chlorostilbon_maugaeus | diurnal |
| birds | Riccordia_ricordii | Chlorostilbon_ricordii | diurnal |
| birds | Riccordia_swainsonii | Chlorostilbon_swainsonii | diurnal |
| birds | Ridgwayia_pinicola | Zoothera_pinicola | diurnal |
| birds | Riparia_congica | Riparia_congica | diurnal |
| birds | Riparia_riparia | Riparia_riparia | diurnal |
| birds | Rissa_brevirostris | Rissa_brevirostris | diurnal |
| birds | Rissa_tridactyla | Rissa_tridactyla | diurnal |
| birds | Robsonius_rabori | Robsonius_rabori | diurnal |
| birds | Rollandia_microptera | Rollandia_microptera | diurnal |
| birds | Rollandia_rolland | Rollandia_rolland | diurnal |
| birds | Rollulus_rouloul | Rollulus_rouloul | diurnal |
| birds | Roraimia_adusta | Roraimia_adusta | diurnal |
| birds | Rostrhamus_sociabilis | Rostrhamus_sociabilis | diurnal |
| birds | Rougetius_rougetii | Rougetius_rougetii | diurnal |
| birds | Rowettia_goughensis | Rowettia_goughensis | diurnal |

| | | | |
|---|---|---|---|
| birds | Rubigula_melanictera | Pycnonotus_melanicterus | diurnal |
| birds | Rufirallus_castaneiceps | Anurolimnas_castaneiceps | diurnal |
| birds | Rufirallus_viridis | Anurolimnas_viridis | diurnal |
| birds | Rukia_longirostra | Rukia_longirostra | diurnal |
| birds | Rukia_ruki | Rukia_ruki | diurnal |
| birds | Rupicola_peruvianus | Rupicola_peruvianus | diurnal |
| birds | Rupicola_rupicola | Rupicola_rupicola | diurnal |
| birds | Rupornis_magnirostris | Buteo_magnirostris | diurnal |
| birds | Rynchops_flavirostris | Rynchops_flavirostris | cathemeral |
| birds | Rynchops_niger | Rynchops_niger | cathemeral |
| birds | Sagittarius_serpentarius | Sagittarius_serpentarius | diurnal |
| birds | Sakesphorus_canadensis | Sakesphorus_canadensis | diurnal |
| birds | Sakesphorus_cristatus | Sakesphorus_cristatus | diurnal |
| birds | Sakesphorus_luctuosus | Sakesphorus_luctuosus | diurnal |
| birds | Salpinctes_obsoletus | Salpinctes_obsoletus | diurnal |
| birds | Salpornis_spilonota | Salpornis_spilonotus | diurnal |
| birds | Saltator_albicollis | Saltator_albicollis | diurnal |
| birds | Saltator_atriceps | Saltator_atriceps | diurnal |
| birds | Saltator_atricollis | Saltator_atricollis | diurnal |
| birds | Saltator_atripennis | Saltator_atripennis | diurnal |
| birds | Saltator_aurantiirostris | Saltator_aurantiirostris | diurnal |
| birds | Saltator_cinctus | Saltator_cinctus | diurnal |
| birds | Saltator_coerulescens | Saltator_coerulescens | diurnal |
| birds | Saltator_fuliginosus | Saltator_fuliginosus | diurnal |
| birds | Saltator_grossus | Saltator_grossus | diurnal |
| birds | Saltator_maxillosus | Saltator_maxillosus | diurnal |
| birds | Saltator_maximus | Saltator_maximus | diurnal |
| birds | Saltator_multicolor | Saltatricula_multicolor | diurnal |
| birds | Saltator_nigriceps | Saltator_nigriceps | diurnal |
| birds | Saltator_orenocensis | Saltator_orenocensis | diurnal |
| birds | Saltator_similis | Saltator_similis | diurnal |
| birds | Saltator_striatipectus | Saltator_striatipectus | diurnal |
| birds | Salvadorina_waigiuensis | Salvadorina_waigiuensis | diurnal |
| birds | Sapayoa_aenigma | Sapayoa_aenigma | diurnal |

| | | | |
|---|---|---|---|
| birds | Sappho_sparganurus | Sappho_sparganura | diurnal |
| birds | Sarcogyps_calvus | Sarcogyps_calvus | diurnal |
| birds | Sarcophanops_samarensis | Eurylaimus_samarensis | diurnal |
| birds | Sarcophanops_steerii | Eurylaimus_steerii | diurnal |
| birds | Sarcops_calvus | Sarcops_calvus | diurnal |
| birds | Sarcoramphus_papa | Sarcoramphus_papa | diurnal |
| birds | Sarkidiornis_melanotos | Sarkidiornis_melanotos | diurnal |
| birds | Saroglossa_spilopterus | Saroglossa_spiloptera | diurnal |
| birds | Sarothrura_affinis | Sarothrura_affinis | diurnal |
| birds | Sarothrura_ayresi | Sarothrura_ayresi | diurnal |
| birds | Sarothrura_boehmi | Sarothrura_boehmi | diurnal |
| birds | Sarothrura_elegans | Sarothrura_elegans | diurnal |
| birds | Sarothrura_insularis | Sarothrura_insularis | diurnal |
| birds | Sarothrura_lugens | Sarothrura_lugens | diurnal |
| birds | Sarothrura_pulchra | Sarothrura_pulchra | diurnal |
| birds | Sarothrura_rufa | Sarothrura_rufa | diurnal |
| birds | Sarothrura_watersi | Sarothrura_watersi | diurnal |
| birds | Sasia_abnormis | Sasia_abnormis | diurnal |
| birds | Sasia_ochracea | Sasia_ochracea | diurnal |
| birds | Satrapa_icterophrys | Satrapa_icterophrys | diurnal |
| birds | Saucerottia_beryllina | Amazilia_beryllina | diurnal |
| birds | Saucerottia_castaneiventris | Amazilia_castaneiventris | diurnal |
| birds | Saucerottia_cyanocephala | Amazilia_cyanocephala | diurnal |
| birds | Saucerottia_cyanura | Amazilia_cyanura | diurnal |
| birds | Saucerottia_edward | Amazilia_edward | diurnal |
| birds | Saucerottia_saucerottei | Amazilia_saucerrottei | diurnal |
| birds | Saucerottia_tobaci | Amazilia_tobaci | diurnal |
| birds | Saundersilarus_saundersi | Larus_saundersi | diurnal |
| birds | Saxicola_caprata | Saxicola_caprata | diurnal |
| birds | Saxicola_dacotiae | Saxicola_dacotiae | diurnal |
| birds | Saxicola_ferreus | Saxicola_ferreus | diurnal |
| birds | Saxicola_gutturalis | Saxicola_gutturalis | diurnal |
| birds | Saxicola_insignis | Saxicola_insignis | diurnal |
| birds | Saxicola_jerdoni | Saxicola_jerdoni | diurnal |

| | | | |
|---|---|---|---|
| birds | Saxicola_macrorhynchus | Saxicola_macrorhynchus | diurnal |
| birds | Saxicola_rubetra | Saxicola_rubetra | diurnal |
| birds | Saxicola_tectes | Saxicola_tectes | diurnal |
| birds | Saxicola_torquatus | Saxicola_torquatus | diurnal |
| birds | Sayornis_nigricans | Sayornis_nigricans | diurnal |
| birds | Sayornis_phoebe | Sayornis_phoebe | diurnal |
| birds | Sayornis_saya | Sayornis_saya | diurnal |
| birds | Scelorchilus_albicollis | Scelorchilus_albicollis | diurnal |
| birds | Scelorchilus_rubecula | Scelorchilus_rubecula | diurnal |
| birds | Scenopoeetes_dentirostris | Scenopoeetes_dentirostris | diurnal |
| birds | Scepomycter_winifredae | Bathmocercus_winifredae | diurnal |
| birds | Schetba_rufa | Schetba_rufa | diurnal |
| birds | Schiffornis_major | Schiffornis_major | diurnal |
| birds | Schiffornis_turdina | Schiffornis_turdina | diurnal |
| birds | Schiffornis_virescens | Schiffornis_virescens | diurnal |
| birds | Schistes_geoffroyi | Schistes_geoffroyi | diurnal |
| birds | Schistochlamys_melanopis | Schistochlamys_melanopis | diurnal |
| birds | Schistochlamys_ruficapillus | Schistochlamys_ruficapillus | diurnal |
| birds | Schistolais_leontica | Prinia_leontica | diurnal |
| birds | Schistolais_leucopogon | Prinia_leucopogon | diurnal |
| birds | Schoenicola_platyurus | Schoenicola_platyurus | diurnal |
| birds | Schoenicola_striatus | Chaetornis_striata | diurnal |
| birds | Schoeniophylax_phryganophilus | Schoeniophylax_phryganophilus | diurnal |
| birds | Schoeniparus_brunneus | Alcippe_brunnea | diurnal |
| birds | Schoeniparus_castaneceps | Alcippe_castaneceps | diurnal |
| birds | Schoeniparus_cinereus | Alcippe_cinerea | diurnal |
| birds | Schoeniparus_dubius | Alcippe_dubia | diurnal |
| birds | Schoeniparus_rufogularis | Alcippe_rufogularis | diurnal |
| birds | Schoeniparus_variegaticeps | Alcippe_variegaticeps | diurnal |
| birds | Schoutedenapus_myoptilus | Schoutedenapus_myoptilus | diurnal |
| birds | Sciaphylax_hemimelaena | Myrmeciza_hemimelaena | diurnal |
| birds | Scissirostrum_dubium | Scissirostrum_dubium | diurnal |
| birds | Sclateria_naevia | Sclateria_naevia | diurnal |
| birds | Scleroptila_finschi | Francolinus_finschi | diurnal |

| | | | |
|---|---|---|---|
| birds | Scleroptila_gutturalis | Francolinus_levaillantoides | diurnal |
| birds | Scleroptila_levaillantii | Francolinus_levaillantii | diurnal |
| birds | Scleroptila_psilolaema | Francolinus_psilolaemus | diurnal |
| birds | Scleroptila_shelleyi | Francolinus_shelleyi | diurnal |
| birds | Scleroptila_streptophora | Francolinus_streptophorus | diurnal |
| birds | Sclerurus_albigularis | Sclerurus_albigularis | diurnal |
| birds | Sclerurus_caudacutus | Sclerurus_caudacutus | diurnal |
| birds | Sclerurus_guatemalensis | Sclerurus_guatemalensis | diurnal |
| birds | Sclerurus_mexicanus | Sclerurus_mexicanus | diurnal |
| birds | Sclerurus_rufigularis | Sclerurus_rufigularis | diurnal |
| birds | Sclerurus_scansor | Sclerurus_scansor | diurnal |
| birds | Scolopax_minor | Scolopax_minor | diurnal |
| birds | Scolopax_rusticola | Scolopax_rusticola | nocturnal |
| birds | Scolopax_saturata | Scolopax_saturata | nocturnal |
| birds | Scopus_umbretta | Scopus_umbretta | diurnal |
| birds | Scotocerca_inquieta | Scotocerca_inquieta | diurnal |
| birds | Scotopelia_bouvieri | Scotopelia_bouvieri | nocturnal |
| birds | Scotopelia_peli | Scotopelia_peli | nocturnal |
| birds | Scotopelia_ussheri | Scotopelia_ussheri | nocturnal |
| birds | Scytalopus_acutirostris | Scytalopus_acutirostris | diurnal |
| birds | Scytalopus_affinis | Scytalopus_affinis | diurnal |
| birds | Scytalopus_altirostris | Scytalopus_altirostris | diurnal |
| birds | Scytalopus_argentifrons | Scytalopus_argentifrons | diurnal |
| birds | Scytalopus_atratus | Scytalopus_atratus | diurnal |
| birds | Scytalopus_bolivianus | Scytalopus_bolivianus | diurnal |
| birds | Scytalopus_canus | Scytalopus_canus | diurnal |
| birds | Scytalopus_caracae | Scytalopus_caracae | diurnal |
| birds | Scytalopus_chocoensis | Scytalopus_chocoensis | diurnal |
| birds | Scytalopus_femoralis | Scytalopus_femoralis | diurnal |
| birds | Scytalopus_griseicollis | Scytalopus_griseicollis | diurnal |
| birds | Scytalopus_iraiensis | Scytalopus_iraiensis | diurnal |
| birds | Scytalopus_latrans | Scytalopus_latrans | diurnal |
| birds | Scytalopus_macropus | Scytalopus_macropus | diurnal |
| birds | Scytalopus_magellanicus | Scytalopus_magellanicus | diurnal |

| | | | |
|---|---|---|---|
| birds | Scytalopus_meridanus | Scytalopus_meridanus | diurnal |
| birds | Scytalopus_micropterus | Scytalopus_micropterus | diurnal |
| birds | Scytalopus_novacapitalis | Scytalopus_novacapitalis | diurnal |
| birds | Scytalopus_parkeri | Scytalopus_parkeri | diurnal |
| birds | Scytalopus_parvirostris | Scytalopus_parvirostris | diurnal |
| birds | Scytalopus_robbinsi | Scytalopus_robbinsi | diurnal |
| birds | Scytalopus_schulenbergi | Scytalopus_schulenbergi | diurnal |
| birds | Scytalopus_simonsi | Scytalopus_simonsi | diurnal |
| birds | Scytalopus_speluncae | Scytalopus_speluncae | diurnal |
| birds | Scytalopus_spillmanni | Scytalopus_spillmanni | diurnal |
| birds | Scytalopus_superciliaris | Scytalopus_superciliaris | diurnal |
| birds | Scytalopus_unicolor | Scytalopus_unicolor | diurnal |
| birds | Scytalopus_urubambae | Scytalopus_urubambae | diurnal |
| birds | Scytalopus_vicinior | Scytalopus_vicinior | diurnal |
| birds | Scytalopus_zimmeri | Scytalopus_zimmeri | diurnal |
| birds | Scythrops_novaehollandiae | Scythrops_novaehollandiae | diurnal |
| birds | Seiurus_aurocapilla | Seiurus_aurocapilla | diurnal |
| birds | Selasphorus_ardens | Selasphorus_ardens | diurnal |
| birds | Selasphorus_calliope | Stellula_calliope | diurnal |
| birds | Selasphorus_ellioti | Atthis_ellioti | diurnal |
| birds | Selasphorus_flammula | Selasphorus_flammula | diurnal |
| birds | Selasphorus_heloisa | Atthis_heloisa | diurnal |
| birds | Selasphorus_platycercus | Selasphorus_platycercus | diurnal |
| birds | Selasphorus_rufus | Selasphorus_rufus | diurnal |
| birds | Selasphorus_sasin | Selasphorus_sasin | diurnal |
| birds | Selasphorus_scintilla | Selasphorus_scintilla | diurnal |
| birds | Selenidera_gouldii | Selenidera_gouldii | diurnal |
| birds | Selenidera_maculirostris | Selenidera_maculirostris | diurnal |
| birds | Selenidera_nattereri | Selenidera_nattereri | diurnal |
| birds | Selenidera_piperivora | Selenidera_piperivora | diurnal |
| birds | Selenidera_reinwardtii | Selenidera_reinwardtii | diurnal |
| birds | Selenidera_spectabilis | Selenidera_spectabilis | diurnal |
| birds | Seleucidis_melanoleucus | Seleucidis_melanoleucus | diurnal |
| birds | Semioptera_wallacii | Semioptera_wallacii | diurnal |

| | | | |
|---|---|---|---|
| birds | Semnornis_frantzii | Semnornis_frantzii | diurnal |
| birds | Semnornis_ramphastinus | Semnornis_ramphastinus | diurnal |
| birds | Sephanoides_fernandensis | Sephanoides_fernandensis | diurnal |
| birds | Sephanoides_sephaniodes | Sephanoides_sephaniodes | diurnal |
| birds | Sericornis_beccarii | Sericornis_beccarii | diurnal |
| birds | Sericornis_frontalis | Sericornis_frontalis | diurnal |
| birds | Sericornis_humilis | Sericornis_humilis | diurnal |
| birds | Sericornis_keri | Sericornis_keri | diurnal |
| birds | Sericornis_magnirostra | Sericornis_magnirostra | diurnal |
| birds | Sericornis_nouhuysi | Sericornis_nouhuysi | diurnal |
| birds | Sericossypha_albocristata | Sericossypha_albocristata | diurnal |
| birds | Sericulus_aureus | Sericulus_aureus | diurnal |
| birds | Sericulus_bakeri | Sericulus_bakeri | diurnal |
| birds | Sericulus_chrysocephalus | Sericulus_chrysocephalus | diurnal |
| birds | Serilophus_lunatus | Serilophus_lunatus | diurnal |
| birds | Serinus_alario | Serinus_alario | diurnal |
| birds | Serinus_canaria | Serinus_canaria | diurnal |
| birds | Serinus_canicollis | Serinus_canicollis | diurnal |
| birds | Serinus_flavivertex | Serinus_flavivertex | diurnal |
| birds | Serinus_nigriceps | Serinus_nigriceps | diurnal |
| birds | Serinus_pusillus | Serinus_pusillus | diurnal |
| birds | Serinus_serinus | Serinus_serinus | diurnal |
| birds | Serinus_syriacus | Serinus_syriacus | diurnal |
| birds | Serpophaga_cinerea | Serpophaga_cinerea | diurnal |
| birds | Serpophaga_hypoleuca | Serpophaga_hypoleuca | diurnal |
| birds | Serpophaga_nigricans | Serpophaga_nigricans | diurnal |
| birds | Serpophaga_subcristata | Serpophaga_subcristata | diurnal |
| birds | Setopagis_parvula | Caprimulgus_parvulus | nocturnal |
| birds | Setopagis_whitelyi | Caprimulgus_whitelyi | nocturnal |
| birds | Setophaga_adelaidae | Dendroica_adelaidae | diurnal |
| birds | Setophaga_americana | Parula_americana | diurnal |
| birds | Setophaga_angelae | Dendroica_angelae | diurnal |
| birds | Setophaga_caerulescens | Dendroica_caerulescens | diurnal |
| birds | Setophaga_castanea | Dendroica_castanea | diurnal |

| | | | |
|---|---|---|---|
| birds | Setophaga_cerulea | Dendroica_cerulea | diurnal |
| birds | Setophaga_chrysoparia | Dendroica_chrysoparia | diurnal |
| birds | Setophaga_citrina | Wilsonia_citrina | diurnal |
| birds | Setophaga_coronata | Dendroica_coronata | diurnal |
| birds | Setophaga_delicata | Dendroica_delicata | diurnal |
| birds | Setophaga_discolor | Dendroica_discolor | diurnal |
| birds | Setophaga_dominica | Dendroica_dominica | diurnal |
| birds | Setophaga_fusca | Dendroica_fusca | diurnal |
| birds | Setophaga_graciae | Dendroica_graciae | diurnal |
| birds | Setophaga_kirtlandii | Dendroica_kirtlandii | diurnal |
| birds | Setophaga_magnolia | Dendroica_magnolia | diurnal |
| birds | Setophaga_nigrescens | Dendroica_nigrescens | diurnal |
| birds | Setophaga_occidentalis | Dendroica_occidentalis | diurnal |
| birds | Setophaga_palmarum | Dendroica_palmarum | diurnal |
| birds | Setophaga_pensylvanica | Dendroica_pensylvanica | diurnal |
| birds | Setophaga_petechia | Dendroica_petechia | diurnal |
| birds | Setophaga_pharetra | Dendroica_pharetra | diurnal |
| birds | Setophaga_pinus | Dendroica_pinus | diurnal |
| birds | Setophaga_pitiayumi | Parula_pitiayumi | diurnal |
| birds | Setophaga_pityophila | Dendroica_pityophila | diurnal |
| birds | Setophaga_plumbea | Dendroica_plumbea | diurnal |
| birds | Setophaga_ruticilla | Setophaga_ruticilla | diurnal |
| birds | Setophaga_striata | Dendroica_striata | diurnal |
| birds | Setophaga_subita | Dendroica_subita | diurnal |
| birds | Setophaga_tigrina | Dendroica_tigrina | diurnal |
| birds | Setophaga_townsendi | Dendroica_townsendi | diurnal |
| birds | Setophaga_virens | Dendroica_virens | diurnal |
| birds | Setophaga_vitellina | Dendroica_vitellina | diurnal |
| birds | Setornis_criniger | Setornis_criniger | diurnal |
| birds | Sheppardia_aequatorialis | Sheppardia_aequatorialis | diurnal |
| birds | Sheppardia_bocagei | Sheppardia_bocagei | diurnal |
| birds | Sheppardia_cyornithopsis | Sheppardia_cyornithopsis | diurnal |
| birds | Sheppardia_gabela | Sheppardia_gabela | diurnal |
| birds | Sheppardia_gunningi | Sheppardia_gunningi | diurnal |

| | | | |
|---|---|---|---|
| birds | Sheppardia_lowei | Sheppardia_lowei | diurnal |
| birds | Sheppardia_montana | Sheppardia_montana | diurnal |
| birds | Sheppardia_polioptera | Cossypha_polioptera | diurnal |
| birds | Sheppardia_sharpei | Sheppardia_sharpei | diurnal |
| birds | Sholicola_major | Myiomela_major | diurnal |
| birds | Sialia_currucoides | Sialia_currucoides | diurnal |
| birds | Sialia_mexicana | Sialia_mexicana | diurnal |
| birds | Sialia_sialis | Sialia_sialis | diurnal |
| birds | Sibia_morrisoniana | Actinodura_morrisoniana | diurnal |
| birds | Sibia_nipalensis | Actinodura_nipalensis | diurnal |
| birds | Sibia_souliei | Actinodura_souliei | diurnal |
| birds | Sibia_waldeni | Actinodura_waldeni | diurnal |
| birds | Sibirionetta_formosa | Anas_formosa | diurnal |
| birds | Sicalis_auriventris | Sicalis_auriventris | diurnal |
| birds | Sicalis_citrina | Sicalis_citrina | diurnal |
| birds | Sicalis_columbiana | Sicalis_columbiana | diurnal |
| birds | Sicalis_flaveola | Sicalis_flaveola | diurnal |
| birds | Sicalis_lebruni | Sicalis_lebruni | diurnal |
| birds | Sicalis_lutea | Sicalis_lutea | diurnal |
| birds | Sicalis_luteocephala | Sicalis_luteocephala | diurnal |
| birds | Sicalis_luteola | Sicalis_luteola | diurnal |
| birds | Sicalis_olivascens | Sicalis_olivascens | diurnal |
| birds | Sicalis_raimondii | Sicalis_raimondii | diurnal |
| birds | Sicalis_taczanowskii | Sicalis_taczanowskii | diurnal |
| birds | Sicalis_uropigyalis | Sicalis_uropygialis | diurnal |
| birds | Sigelus_silens | Sigelus_silens | diurnal |
| birds | Silvicultrix_diadema | Ochthoeca_diadema | diurnal |
| birds | Silvicultrix_frontalis | Ochthoeca_frontalis | diurnal |
| birds | Silvicultrix_jelskii | Ochthoeca_jelskii | diurnal |
| birds | Silvicultrix_pulchella | Ochthoeca_pulchella | diurnal |
| birds | Siphonorhis_brewsteri | Siphonorhis_brewsteri | nocturnal |
| birds | Sipia_berlepschi | Myrmeciza_berlepschi | diurnal |
| birds | Sipia_laemosticta | Myrmeciza_laemosticta | diurnal |
| birds | Sipia_nigricauda | Myrmeciza_nigricauda | diurnal |

| | | | |
|---|---|---|---|
| birds | Sipodotus_wallacii | Sipodotus_wallacii | diurnal |
| birds | Siptornis_striaticollis | Siptornis_striaticollis | diurnal |
| birds | Sirystes_albogriseus | Sirystes_albogriseus | diurnal |
| birds | Sirystes_sibilator | Sirystes_sibilator | diurnal |
| birds | Sitta_azurea | Sitta_azurea | diurnal |
| birds | Sitta_canadensis | Sitta_canadensis | diurnal |
| birds | Sitta_cashmirensis | Sitta_cashmirensis | diurnal |
| birds | Sitta_castanea | Sitta_castanea | diurnal |
| birds | Sitta_europaea | Sitta_europaea | diurnal |
| birds | Sitta_frontalis | Sitta_frontalis | diurnal |
| birds | Sitta_himalayensis | Sitta_himalayensis | diurnal |
| birds | Sitta_krueperi | Sitta_krueperi | diurnal |
| birds | Sitta_ledanti | Sitta_ledanti | diurnal |
| birds | Sitta_leucopsis | Sitta_leucopsis | diurnal |
| birds | Sitta_magna | Sitta_magna | diurnal |
| birds | Sitta_nagaensis | Sitta_nagaensis | diurnal |
| birds | Sitta_neumayer | Sitta_neumayer | diurnal |
| birds | Sitta_oenochlamys | Sitta_oenochlamys | diurnal |
| birds | Sitta_pusilla | Sitta_pusilla | diurnal |
| birds | Sitta_pygmaea | Sitta_pygmaea | diurnal |
| birds | Sitta_solangiae | Sitta_solangiae | diurnal |
| birds | Sitta_tephronota | Sitta_tephronota | diurnal |
| birds | Sitta_villosa | Sitta_villosa | diurnal |
| birds | Sitta_whiteheadi | Sitta_whiteheadi | diurnal |
| birds | Sitta_yunnanensis | Sitta_yunnanensis | diurnal |
| birds | Sittasomus_griseicapillus | Sittasomus_griseicapillus | diurnal |
| birds | Sittiparus_semilarvatus | Parus_semilarvatus | diurnal |
| birds | Sittiparus_varius | Parus_varius | diurnal |
| birds | Siva_cyanouroptera | Minla_cyanouroptera | diurnal |
| birds | Smicrornis_brevirostris | Smicrornis_brevirostris | diurnal |
| birds | Smithornis_capensis | Smithornis_capensis | diurnal |
| birds | Smithornis_rufolateralis | Smithornis_rufolateralis | diurnal |
| birds | Smithornis_sharpei | Smithornis_sharpei | diurnal |
| birds | Smutsornis_africanus | Rhinoptilus_africanus | nocturnal |

| | | | |
|---|---|---|---|
| birds | Snowornis_cryptolophus | Snowornis_cryptolophus | diurnal |
| birds | Snowornis_subalaris | Snowornis_subalaris | diurnal |
| birds | Somateria_fischeri | Somateria_fischeri | diurnal |
| birds | Somateria_mollissima | Somateria_mollissima | diurnal |
| birds | Somateria_spectabilis | Somateria_spectabilis | diurnal |
| birds | Spartonoica_maluroides | Spartonoica_maluroides | diurnal |
| birds | Spatula_clypeata | Anas_clypeata | diurnal |
| birds | Spatula_cyanoptera | Anas_cyanoptera | diurnal |
| birds | Spatula_discors | Anas_discors | diurnal |
| birds | Spatula_hottentota | Anas_hottentota | diurnal |
| birds | Spatula_platalea | Anas_platalea | diurnal |
| birds | Spatula_querquedula | Anas_querquedula | diurnal |
| birds | Spatula_rhynchotis | Anas_rhynchotis | diurnal |
| birds | Spatula_smithii | Anas_smithii | diurnal |
| birds | Spatula_versicolor | Anas_versicolor | diurnal |
| birds | Speculanas_specularis | Speculanas_specularis | diurnal |
| birds | Speculipastor_bicolor | Speculipastor_bicolor | diurnal |
| birds | Spelaeornis_badeigularis | Spelaeornis_badeigularis | diurnal |
| birds | Spelaeornis_caudatus | Spelaeornis_caudatus | diurnal |
| birds | Spelaeornis_chocolatinus | Spelaeornis_chocolatinus | diurnal |
| birds | Spelaeornis_longicaudatus | Spelaeornis_longicaudatus | diurnal |
| birds | Spelaeornis_troglodytoides | Spelaeornis_troglodytoides | diurnal |
| birds | Spermestes_bicolor | Lonchura_bicolor | diurnal |
| birds | Spermestes_cucullata | Lonchura_cucullata | diurnal |
| birds | Spermestes_fringilloides | Lonchura_fringilloides | diurnal |
| birds | Spermestes_griseicapilla | Lonchura_griseicapilla | diurnal |
| birds | Spermophaga_haematina | Spermophaga_haematina | diurnal |
| birds | Spermophaga_poliogenys | Spermophaga_poliogenys | diurnal |
| birds | Spermophaga_ruficapilla | Spermophaga_ruficapilla | diurnal |
| birds | Sphecotheres_hypoleucus | Sphecotheres_hypoleucus | diurnal |
| birds | Sphecotheres_vieilloti | Sphecotheres_vieilloti | diurnal |
| birds | Sphecotheres_viridis | Sphecotheres_viridis | diurnal |
| birds | Spheniscus_demersus | Spheniscus_demersus | cathemeral |
| birds | Spheniscus_humboldti | Spheniscus_humboldti | cathemeral |

| | | | |
|---|---|---|---|
| birds | Spheniscus_magellanicus | Spheniscus_magellanicus | cathemeral |
| birds | Spheniscus_mendiculus | Spheniscus_mendiculus | cathemeral |
| birds | Sphenoeacus_afer | Sphenoeacus_afer | diurnal |
| birds | Sphenopsis_frontalis | Hemispingus_frontalis | diurnal |
| birds | Sphenopsis_melanotis | Hemispingus_melanotis | diurnal |
| birds | Sphyrapicus_nuchalis | Sphyrapicus_nuchalis | diurnal |
| birds | Sphyrapicus_ruber | Sphyrapicus_ruber | diurnal |
| birds | Sphyrapicus_thyroideus | Sphyrapicus_thyroideus | diurnal |
| birds | Sphyrapicus_varius | Sphyrapicus_varius | diurnal |
| birds | Spilopelia_chinensis | Stigmatopelia_chinensis | diurnal |
| birds | Spilopelia_senegalensis | Stigmatopelia_senegalensis | diurnal |
| birds | Spiloptila_clamans | Spiloptila_clamans | diurnal |
| birds | Spilornis_cheela | Spilornis_cheela | diurnal |
| birds | Spilornis_elgini | Spilornis_elgini | diurnal |
| birds | Spilornis_holospilus | Spilornis_holospilus | diurnal |
| birds | Spindalis_dominicensis | Spindalis_dominicensis | diurnal |
| birds | Spindalis_nigricephala | Spindalis_nigricephala | diurnal |
| birds | Spindalis_portoricensis | Spindalis_portoricensis | diurnal |
| birds | Spindalis_zena | Spindalis_zena | diurnal |
| birds | Spinus_atratus | Carduelis_atrata | diurnal |
| birds | Spinus_atriceps | Carduelis_atriceps | diurnal |
| birds | Spinus_barbatus | Carduelis_barbata | diurnal |
| birds | Spinus_crassirostris | Carduelis_crassirostris | diurnal |
| birds | Spinus_cucullatus | Carduelis_cucullata | diurnal |
| birds | Spinus_dominicensis | Carduelis_dominicensis | diurnal |
| birds | Spinus_lawrencei | Carduelis_lawrencei | diurnal |
| birds | Spinus_magellanicus | Carduelis_magellanica | diurnal |
| birds | Spinus_notatus | Carduelis_notata | diurnal |
| birds | Spinus_pinus | Carduelis_pinus | diurnal |
| birds | Spinus_psaltria | Carduelis_psaltria | diurnal |
| birds | Spinus_spinescens | Carduelis_spinescens | diurnal |
| birds | Spinus_spinus | Carduelis_spinus | diurnal |
| birds | Spinus_thibetanus | Serinus_thibetanus | diurnal |
| birds | Spinus_tristis | Carduelis_tristis | diurnal |

| | | | |
|---|---|---|---|
| birds | Spinus_uropygialis | Carduelis_uropygialis | diurnal |
| birds | Spinus_xanthogastrus | Carduelis_xanthogastra | diurnal |
| birds | Spinus_yarrellii | Carduelis_yarrellii | diurnal |
| birds | Spiza_americana | Spiza_americana | diurnal |
| birds | Spizaetus_melanoleucus | Spizaetus_melanoleucus | diurnal |
| birds | Spizaetus_ornatus | Spizaetus_ornatus | diurnal |
| birds | Spizaetus_tyrannus | Spizaetus_tyrannus | diurnal |
| birds | Spizella_atrogularis | Spizella_atrogularis | diurnal |
| birds | Spizella_breweri | Spizella_breweri | diurnal |
| birds | Spizella_pallida | Spizella_pallida | diurnal |
| birds | Spizella_passerina | Spizella_passerina | diurnal |
| birds | Spizella_pusilla | Spizella_pusilla | diurnal |
| birds | Spizella_wortheni | Spizella_wortheni | diurnal |
| birds | Spiziapteryx_circumcincta | Spiziapteryx_circumcincta | diurnal |
| birds | Spizixos_canifrons | Spizixos_canifrons | diurnal |
| birds | Spizixos_semitorques | Spizixos_semitorques | diurnal |
| birds | Spizocorys_conirostris | Spizocorys_conirostris | diurnal |
| birds | Spizocorys_fringillaris | Spizocorys_fringillaris | diurnal |
| birds | Spizocorys_obbiensis | Spizocorys_obbiensis | diurnal |
| birds | Spizocorys_personata | Spizocorys_personata | diurnal |
| birds | Spizocorys_sclateri | Spizocorys_sclateri | diurnal |
| birds | Spodiopsar_cineraceus | Sturnus_cineraceus | diurnal |
| birds | Spodiopsar_sericeus | Sturnus_sericeus | diurnal |
| birds | Spodiornis_rusticus | Haplospiza_rustica | diurnal |
| birds | Sporathraupis_cyanocephala | Thraupis_cyanocephala | diurnal |
| birds | Sporophila_albogularis | Sporophila_albogularis | diurnal |
| birds | Sporophila_americana | Sporophila_americana | diurnal |
| birds | Sporophila_angolensis | Oryzoborus_angolensis | diurnal |
| birds | Sporophila_atrirostris | Oryzoborus_atrirostris | diurnal |
| birds | Sporophila_bouvreuil | Sporophila_bouvreuil | diurnal |
| birds | Sporophila_bouvronides | Sporophila_bouvronides | diurnal |
| birds | Sporophila_caerulescens | Sporophila_caerulescens | diurnal |
| birds | Sporophila_castaneiventris | Sporophila_castaneiventris | diurnal |
| birds | Sporophila_cinnamomea | Sporophila_cinnamomea | diurnal |

| | | | |
|---|---|---|---|
| birds | Sporophila_collaris | Sporophila_collaris | diurnal |
| birds | Sporophila_corvina | Sporophila_corvina | diurnal |
| birds | Sporophila_crassirostris | Oryzoborus_crassirostris | diurnal |
| birds | Sporophila_falcirostris | Sporophila_falcirostris | diurnal |
| birds | Sporophila_fringilloides | Dolospingus_fringilloides | diurnal |
| birds | Sporophila_frontalis | Sporophila_frontalis | diurnal |
| birds | Sporophila_hypochroma | Sporophila_hypochroma | diurnal |
| birds | Sporophila_hypoxantha | Sporophila_hypoxantha | diurnal |
| birds | Sporophila_intermedia | Sporophila_intermedia | diurnal |
| birds | Sporophila_leucoptera | Sporophila_leucoptera | diurnal |
| birds | Sporophila_lineola | Sporophila_lineola | diurnal |
| birds | Sporophila_luctuosa | Sporophila_luctuosa | diurnal |
| birds | Sporophila_maximiliani | Oryzoborus_maximiliani | diurnal |
| birds | Sporophila_melanogaster | Sporophila_melanogaster | diurnal |
| birds | Sporophila_minuta | Sporophila_minuta | diurnal |
| birds | Sporophila_murallae | Sporophila_murallae | diurnal |
| birds | Sporophila_nigricollis | Sporophila_nigricollis | diurnal |
| birds | Sporophila_nigrorufa | Sporophila_nigrorufa | diurnal |
| birds | Sporophila_nuttingi | Oryzoborus_nuttingi | diurnal |
| birds | Sporophila_palustris | Sporophila_palustris | diurnal |
| birds | Sporophila_peruviana | Sporophila_peruviana | diurnal |
| birds | Sporophila_plumbea | Sporophila_plumbea | diurnal |
| birds | Sporophila_ruficollis | Sporophila_ruficollis | diurnal |
| birds | Sporophila_schistacea | Sporophila_schistacea | diurnal |
| birds | Sporophila_simplex | Sporophila_simplex | diurnal |
| birds | Sporophila_telasco | Sporophila_telasco | diurnal |
| birds | Sporophila_torqueola | Sporophila_torqueola | diurnal |
| birds | Sporopipes_frontalis | Sporopipes_frontalis | diurnal |
| birds | Sporopipes_squamifrons | Sporopipes_squamifrons | diurnal |
| birds | Stachyris_grammiceps | Stachyris_grammiceps | diurnal |
| birds | Stachyris_herberti | Stachyris_herberti | diurnal |
| birds | Stachyris_humei | Sphenocichla_humei | diurnal |
| birds | Stachyris_leucotis | Stachyris_leucotis | diurnal |
| birds | Stachyris_maculata | Stachyris_maculata | diurnal |

| | | | |
|---|---|---|---|
| birds | Stachyris_nigriceps | Stachyris_nigriceps | diurnal |
| birds | Stachyris_nigricollis | Stachyris_nigricollis | diurnal |
| birds | Stachyris_oglei | Stachyris_oglei | diurnal |
| birds | Stachyris_poliocephala | Stachyris_poliocephala | diurnal |
| birds | Stachyris_strialata | Stachyris_striolata | diurnal |
| birds | Stachyris_thoracica | Stachyris_thoracica | diurnal |
| birds | Stactolaema_anchietae | Stactolaema_anchietae | diurnal |
| birds | Stactolaema_leucotis | Stactolaema_leucotis | diurnal |
| birds | Stactolaema_whytii | Stactolaema_whytii | diurnal |
| birds | Stagonopleura_bella | Stagonopleura_bella | diurnal |
| birds | Stagonopleura_oculata | Stagonopleura_oculata | diurnal |
| birds | Staphida_everetti | Yuhina_everetti | diurnal |
| birds | Starnoenas_cyanocephala | Starnoenas_cyanocephala | diurnal |
| birds | Steatornis_caripensis | Steatornis_caripensis | nocturnal |
| birds | Steganopus_tricolor | Steganopus_tricolor | diurnal |
| birds | Stelgidillas_gracilirostris | Andropadus_gracilirostris | diurnal |
| birds | Stelgidopteryx_ruficollis | Stelgidopteryx_ruficollis | diurnal |
| birds | Stelgidopteryx_serripennis | Stelgidopteryx_serripennis | diurnal |
| birds | Stenostira_scita | Stenostira_scita | diurnal |
| birds | Stephanoaetus_coronatus | Stephanoaetus_coronatus | diurnal |
| birds | Stephanophorus_diadematus | Stephanophorus_diadematus | diurnal |
| birds | Stephanoxis_lalandi | Stephanoxis_lalandi | diurnal |
| birds | Stercorarius_longicaudus | Stercorarius_longicaudus | diurnal |
| birds | Stercorarius_parasiticus | Stercorarius_parasiticus | diurnal |
| birds | Stercorarius_pomarinus | Stercorarius_pomarinus | diurnal |
| birds | Sterna_forsteri | Sterna_forsteri | diurnal |
| birds | Sterna_hirundinacea | Sterna_hirundinacea | diurnal |
| birds | Sterna_hirundo | Sterna_hirundo | diurnal |
| birds | Sterna_paradisaea | Sterna_paradisaea | diurnal |
| birds | Sterna_repressa | Sterna_repressa | diurnal |
| birds | Sterna_striata | Sterna_striata | diurnal |
| birds | Sterna_trudeaui | Sterna_trudeaui | diurnal |
| birds | Sterna_virgata | Sterna_virgata | diurnal |
| birds | Sterna_vittata | Sterna_vittata | diurnal |

| | | | |
|---|---|---|---|
| birds | Sternoclyta_cyanopectus | Sternoclyta_cyanopectus | diurnal |
| birds | Sternula_albifrons | Sterna_albifrons | diurnal |
| birds | Sternula_antillarum | Sterna_antillarum | diurnal |
| birds | Sternula_balaenarum | Sterna_balaenarum | diurnal |
| birds | Sternula_lorata | Sterna_lorata | diurnal |
| birds | Sternula_nereis | Sterna_nereis | diurnal |
| birds | Sternula_saundersi | Sterna_saundersi | diurnal |
| birds | Sternula_superciliaris | Sterna_superciliaris | diurnal |
| birds | Sterrhoptilus_capitalis | Stachyris_capitalis | diurnal |
| birds | Sterrhoptilus_dennistouni | Stachyris_dennistouni | diurnal |
| birds | Sterrhoptilus_nigrocapitatus | Stachyris_nigrocapitata | diurnal |
| birds | Stictonetta_naevosa | Stictonetta_naevosa | diurnal |
| birds | Stigmatura_budytoides | Stigmatura_budytoides | diurnal |
| birds | Stigmatura_napensis | Stigmatura_napensis | diurnal |
| birds | Stiltia_isabella | Stiltia_isabella | diurnal |
| birds | Stiphrornis_erythrothorax | Stiphrornis_erythrothorax | diurnal |
| birds | Stipiturus_malachurus | Stipiturus_malachurus | diurnal |
| birds | Stipiturus_mallee | Stipiturus_mallee | diurnal |
| birds | Stipiturus_ruficeps | Stipiturus_ruficeps | diurnal |
| birds | Stizorhina_finschi | Stizorhina_finschi | diurnal |
| birds | Stizorhina_fraseri | Stizorhina_fraseri | diurnal |
| birds | Stomiopera_flava | Lichenostomus_flavus | diurnal |
| birds | Stomiopera_unicolor | Lichenostomus_unicolor | diurnal |
| birds | Strepera_fuliginosa | Strepera_fuliginosa | diurnal |
| birds | Strepera_graculina | Strepera_graculina | diurnal |
| birds | Strepera_versicolor | Strepera_versicolor | diurnal |
| birds | Streptocitta_albertinae | Streptocitta_albertinae | diurnal |
| birds | Streptocitta_albicollis | Streptocitta_albicollis | diurnal |
| birds | Streptopelia_bitorquata | Streptopelia_bitorquata | diurnal |
| birds | Streptopelia_capicola | Streptopelia_capicola | diurnal |
| birds | Streptopelia_decaocto | Streptopelia_decaocto | diurnal |
| birds | Streptopelia_decipiens | Streptopelia_decipiens | diurnal |
| birds | Streptopelia_hypopyrrha | Streptopelia_hypopyrrha | diurnal |
| birds | Streptopelia_lugens | Streptopelia_lugens | diurnal |

| | | | |
|---|---|---|---|
| birds | Streptopelia_orientalis | Streptopelia_orientalis | diurnal |
| birds | Streptopelia_reichenowi | Streptopelia_reichenowi | diurnal |
| birds | Streptopelia_roseogrisea | Streptopelia_roseogrisea | diurnal |
| birds | Streptopelia_semitorquata | Streptopelia_semitorquata | diurnal |
| birds | Streptopelia_tranquebarica | Streptopelia_tranquebarica | diurnal |
| birds | Streptopelia_turtur | Streptopelia_turtur | diurnal |
| birds | Streptopelia_vinacea | Streptopelia_vinacea | diurnal |
| birds | Streptoprocne_biscutata | Streptoprocne_biscutata | diurnal |
| birds | Streptoprocne_rutila | Streptoprocne_rutila | diurnal |
| birds | Streptoprocne_semicollaris | Streptoprocne_semicollaris | diurnal |
| birds | Streptoprocne_zonaris | Streptoprocne_zonaris | diurnal |
| birds | Stresemannia_bougainvillei | Stresemannia_bougainvillei | diurnal |
| birds | Strigops_habroptila | Strigops_habroptila | nocturnal |
| birds | Strix_aluco | Strix_aluco | cathemeral |
| birds | Strix_butleri | Strix_butleri | nocturnal |
| birds | Strix_chacoensis | Strix_chacoensis | nocturnal |
| birds | Strix_hylophila | Strix_hylophila | nocturnal |
| birds | Strix_leptogrammica | Strix_leptogrammica | nocturnal |
| birds | Strix_nebulosa | Strix_nebulosa | cathemeral |
| birds | Strix_occidentalis | Strix_occidentalis | nocturnal |
| birds | Strix_rufipes | Strix_rufipes | nocturnal |
| birds | Strix_seloputo | Strix_seloputo | nocturnal |
| birds | Strix_uralensis | Strix_uralensis | cathemeral |
| birds | Strix_varia | Strix_varia | nocturnal |
| birds | Strix_woodfordii | Strix_woodfordii | nocturnal |
| birds | Struthidea_cinerea | Struthidea_cinerea | diurnal |
| birds | Struthio_camelus | Struthio_camelus | diurnal |
| birds | Sturnella_magna | Sturnella_magna | diurnal |
| birds | Sturnella_neglecta | Sturnella_neglecta | diurnal |
| birds | Sturnia_malabarica | Sturnus_malabaricus | diurnal |
| birds | Sturnia_pagodarum | Sturnus_pagodarum | diurnal |
| birds | Sturnia_sinensis | Sturnus_sinensis | diurnal |
| birds | Sturnus_unicolor | Sturnus_unicolor | diurnal |
| birds | Sturnus_vulgaris | Sturnus_vulgaris | diurnal |

| | | | |
|---|---|---|---|
| birds | Sublegatus_arenarum | Sublegatus_arenarum | diurnal |
| birds | Sublegatus_modestus | Sublegatus_modestus | diurnal |
| birds | Sublegatus_obscurior | Sublegatus_obscurior | diurnal |
| birds | Sugomel_nigrum | Certhionyx_niger | diurnal |
| birds | Suiriri_suiriri | Suiriri_suiriri | diurnal |
| birds | Sula_dactylatra | Sula_dactylatra | diurnal |
| birds | Sula_leucogaster | Sula_leucogaster | diurnal |
| birds | Sula_nebouxii | Sula_nebouxii | diurnal |
| birds | Sula_sula | Sula_sula | nocturnal |
| birds | Sula_variegata | Sula_variegata | diurnal |
| birds | Surnia_ulula | Surnia_ulula | cathemeral |
| birds | Surniculus_lugubris | Surniculus_lugubris | diurnal |
| birds | Surniculus_velutinus | Surniculus_velutinus | diurnal |
| birds | Suthora_alphonsiana | Paradoxornis_alphonsianus | diurnal |
| birds | Suthora_atrosuperciliaris | Paradoxornis_atrosuperciliaris | diurnal |
| birds | Suthora_conspicillata | Paradoxornis_conspicillatus | diurnal |
| birds | Suthora_davidiana | Paradoxornis_davidianus | diurnal |
| birds | Suthora_fulvifrons | Paradoxornis_fulvifrons | diurnal |
| birds | Suthora_nipalensis | Paradoxornis_nipalensis | diurnal |
| birds | Suthora_verreauxi | Paradoxornis_verreauxi | diurnal |
| birds | Suthora_webbiana | Paradoxornis_webbianus | diurnal |
| birds | Suthora_zappeyi | Paradoxornis_zappeyi | diurnal |
| birds | Swynnertonia_swynnertoni | Swynnertonia_swynnertoni | diurnal |
| birds | Sylvia_abyssinica | Pseudoalcippe_abyssinica | diurnal |
| birds | Sylvia_atricapilla | Sylvia_atricapilla | diurnal |
| birds | Sylvia_borin | Sylvia_borin | diurnal |
| birds | Sylvia_dohrni | Horizorhinus_dohrni | diurnal |
| birds | Sylvia_galinieri | Parophasma_galinieri | diurnal |
| birds | Sylvia_nigricapillus | Lioptilus_nigricapillus | diurnal |
| birds | Sylvietta_brachyura | Sylvietta_brachyura | diurnal |
| birds | Sylvietta_denti | Sylvietta_denti | diurnal |
| birds | Sylvietta_isabellina | Sylvietta_isabellina | diurnal |
| birds | Sylvietta_leucophrys | Sylvietta_leucophrys | diurnal |
| birds | Sylvietta_philippae | Sylvietta_philippae | diurnal |

| | | | |
|---|---|---|---|
| birds | Sylvietta_rufescens | Sylvietta_rufescens | diurnal |
| birds | Sylvietta_ruficapilla | Sylvietta_ruficapilla | diurnal |
| birds | Sylvietta_virens | Sylvietta_virens | diurnal |
| birds | Sylvietta_whytii | Sylvietta_whytii | diurnal |
| birds | Sylviorthorhynchus_desmurii | Sylviorthorhynchus_desmursii | diurnal |
| birds | Sylviorthorhynchus_yanacensis | Leptasthenura_yanacensis | diurnal |
| birds | Sylviparus_modestus | Sylviparus_modestus | diurnal |
| birds | Syma_megarhyncha | Syma_megarhyncha | diurnal |
| birds | Syma_torotoro | Syma_torotoro | diurnal |
| birds | Symposiachrus_axillaris | Monarcha_axillaris | diurnal |
| birds | Symposiachrus_barbatus | Monarcha_barbatus | diurnal |
| birds | Symposiachrus_boanensis | Monarcha_boanensis | diurnal |
| birds | Symposiachrus_brehmii | Monarcha_brehmii | diurnal |
| birds | Symposiachrus_guttula | Monarcha_guttula | diurnal |
| birds | Symposiachrus_manadensis | Monarcha_manadensis | diurnal |
| birds | Symposiachrus_sacerdotum | Monarcha_sacerdotum | diurnal |
| birds | Symposiachrus_trivirgatus | Monarcha_trivirgatus | diurnal |
| birds | Symposiachrus_verticalis | Monarcha_verticalis | diurnal |
| birds | Symposiachrus_vidua | Monarcha_viduus | diurnal |
| birds | Synallaxis_albescens | Synallaxis_albescens | diurnal |
| birds | Synallaxis_albigularis | Synallaxis_albigularis | diurnal |
| birds | Synallaxis_albilora | Synallaxis_albilora | diurnal |
| birds | Synallaxis_azarae | Synallaxis_azarae | diurnal |
| birds | Synallaxis_brachyura | Synallaxis_brachyura | diurnal |
| birds | Synallaxis_cabanisi | Synallaxis_cabanisi | diurnal |
| birds | Synallaxis_candei | Synallaxis_candei | diurnal |
| birds | Synallaxis_castanea | Synallaxis_castanea | diurnal |
| birds | Synallaxis_cherriei | Synallaxis_cherriei | diurnal |
| birds | Synallaxis_cinerascens | Synallaxis_cinerascens | diurnal |
| birds | Synallaxis_cinerea | Synallaxis_whitneyi | diurnal |
| birds | Synallaxis_cinnamomea | Synallaxis_cinnamomea | diurnal |
| birds | Synallaxis_courseni | Synallaxis_courseni | diurnal |
| birds | Synallaxis_erythrothorax | Synallaxis_erythrothorax | diurnal |
| birds | Synallaxis_frontalis | Synallaxis_frontalis | diurnal |

| | | | |
|---|---|---|---|
| birds | Synallaxis_fuscorufa | Synallaxis_fuscorufa | diurnal |
| birds | Synallaxis_gujanensis | Synallaxis_gujanensis | diurnal |
| birds | Synallaxis_hellmayri | Gyalophylax_hellmayri | diurnal |
| birds | Synallaxis_hypochondriaca | Siptornopsis_hypochondriaca | diurnal |
| birds | Synallaxis_hypospodia | Synallaxis_hypospodia | diurnal |
| birds | Synallaxis_infuscata | Synallaxis_infuscata | diurnal |
| birds | Synallaxis_kollari | Synallaxis_kollari | diurnal |
| birds | Synallaxis_macconnelli | Synallaxis_macconnelli | diurnal |
| birds | Synallaxis_maranonica | Synallaxis_maranonica | diurnal |
| birds | Synallaxis_moesta | Synallaxis_moesta | diurnal |
| birds | Synallaxis_ruficapilla | Synallaxis_ruficapilla | diurnal |
| birds | Synallaxis_rutilans | Synallaxis_rutilans | diurnal |
| birds | Synallaxis_scutata | Synallaxis_scutata | diurnal |
| birds | Synallaxis_spixi | Synallaxis_spixi | diurnal |
| birds | Synallaxis_stictothorax | Synallaxis_stictothorax | diurnal |
| birds | Synallaxis_tithys | Synallaxis_tithys | diurnal |
| birds | Synallaxis_unirufa | Synallaxis_unirufa | diurnal |
| birds | Synallaxis_zimmeri | Synallaxis_zimmeri | diurnal |
| birds | Syndactyla_dimidiata | Syndactyla_dimidiata | diurnal |
| birds | Syndactyla_guttulata | Syndactyla_guttulata | diurnal |
| birds | Syndactyla_roraimae | Syndactyla_roraimae | diurnal |
| birds | Syndactyla_ruficollis | Syndactyla_ruficollis | diurnal |
| birds | Syndactyla_rufosuperciliata | Syndactyla_rufosuperciliata | diurnal |
| birds | Syndactyla_striata | Simoxenops_striatus | diurnal |
| birds | Syndactyla_subalaris | Syndactyla_subalaris | diurnal |
| birds | Syndactyla_ucayalae | Simoxenops_ucayalae | diurnal |
| birds | Synoicus_chinensis | Coturnix_chinensis | diurnal |
| birds | Synoicus_ypsilophorus | Coturnix_ypsilophora | diurnal |
| birds | Synthliboramphus_antiquus | Synthliboramphus_antiquus | diurnal |
| birds | Synthliboramphus_craveri | Synthliboramphus_craveri | diurnal |
| birds | Synthliboramphus_hypoleucus | Synthliboramphus_hypoleucus | diurnal |
| birds | Synthliboramphus_wumizusume | Synthliboramphus_wumizusume | diurnal |
| birds | Sypheotides_indicus | Sypheotides_indicus | diurnal |
| birds | Syrigma_sibilatrix | Syrigma_sibilatrix | diurnal |

| | | | |
|---|---|---|---|
| birds | Syrmaticus_ellioti | Syrmaticus_ellioti | diurnal |
| birds | Syrmaticus_humiae | Syrmaticus_humiae | diurnal |
| birds | Syrmaticus_mikado | Syrmaticus_mikado | diurnal |
| birds | Syrmaticus_reevesii | Syrmaticus_reevesii | diurnal |
| birds | Syrmaticus_soemmerringii | Syrmaticus_soemmerringii | diurnal |
| birds | Syrrhaptes_tibetanus | Syrrhaptes_tibetanus | diurnal |
| birds | Systellura_longirostris | Caprimulgus_longirostris | nocturnal |
| birds | Taccocua_leschenaultii | Phaenicophaeus_leschenaultii | diurnal |
| birds | Tachornis_furcata | Tachornis_furcata | diurnal |
| birds | Tachornis_phoenicobia | Tachornis_phoenicobia | diurnal |
| birds | Tachornis_squamata | Tachornis_squamata | diurnal |
| birds | Tachuris_rubrigastra | Tachuris_rubrigastra | diurnal |
| birds | Tachybaptus_dominicus | Tachybaptus_dominicus | diurnal |
| birds | Tachybaptus_novaehollandiae | Tachybaptus_novaehollandiae | diurnal |
| birds | Tachybaptus_pelzelnii | Tachybaptus_pelzelnii | diurnal |
| birds | Tachybaptus_ruficollis | Tachybaptus_ruficollis | diurnal |
| birds | Tachycineta_albiventer | Tachycineta_albiventer | diurnal |
| birds | Tachycineta_bicolor | Tachycineta_bicolor | diurnal |
| birds | Tachycineta_cyaneoviridis | Tachycineta_cyaneoviridis | diurnal |
| birds | Tachycineta_euchrysea | Tachycineta_euchrysea | diurnal |
| birds | Tachycineta_leucorrhoa | Tachycineta_leucorrhoa | diurnal |
| birds | Tachycineta_meyeni | Tachycineta_meyeni | diurnal |
| birds | Tachycineta_stolzmanni | Tachycineta_stolzmanni | diurnal |
| birds | Tachycineta_thalassina | Tachycineta_thalassina | diurnal |
| birds | Tachyeres_brachypterus | Tachyeres_brachypterus | diurnal |
| birds | Tachyeres_leucocephalus | Tachyeres_leucocephalus | diurnal |
| birds | Tachyeres_patachonicus | Tachyeres_patachonicus | diurnal |
| birds | Tachyeres_pteneres | Tachyeres_pteneres | diurnal |
| birds | Tachymarptis_aequatorialis | Tachymarptis_aequatorialis | diurnal |
| birds | Tachymarptis_melba | Tachymarptis_melba | nocturnal |
| birds | Tachyphonus_coronatus | Tachyphonus_coronatus | diurnal |
| birds | Tachyphonus_phoenicius | Tachyphonus_phoenicius | diurnal |
| birds | Tachyphonus_rufus | Tachyphonus_rufus | diurnal |
| birds | Tadorna_cana | Tadorna_cana | diurnal |

| | | | |
|---|---|---|---|
| birds | Tadorna_ferruginea | Tadorna_ferruginea | diurnal |
| birds | Tadorna_tadorna | Tadorna_tadorna | diurnal |
| birds | Tadorna_tadornoides | Tadorna_tadornoides | diurnal |
| birds | Tadorna_variegata | Tadorna_variegata | diurnal |
| birds | Taeniopygia_bichenovii | Taeniopygia_bichenovii | diurnal |
| birds | Taeniopygia_guttata | Taeniopygia_guttata | diurnal |
| birds | Taeniotriccus_andrei | Taeniotriccus_andrei | diurnal |
| birds | Talaphorus_chlorocercus | Leucippus_chlorocercus | diurnal |
| birds | Talegalla_cuvieri | Talegalla_cuvieri | diurnal |
| birds | Talegalla_fuscirostris | Talegalla_fuscirostris | diurnal |
| birds | Talegalla_jobiensis | Talegalla_jobiensis | diurnal |
| birds | Tangara_abbas | Thraupis_abbas | diurnal |
| birds | Tangara_argentea | Tangara_cyanoptera | diurnal |
| birds | Tangara_argyrofenges | Tangara_argyrofenges | diurnal |
| birds | Tangara_arthus | Tangara_arthus | diurnal |
| birds | Tangara_cabanisi | Tangara_cabanisi | diurnal |
| birds | Tangara_callophrys | Tangara_callophrys | diurnal |
| birds | Tangara_cayana | Tangara_cayana | diurnal |
| birds | Tangara_chilensis | Tangara_chilensis | diurnal |
| birds | Tangara_chrysotis | Tangara_chrysotis | diurnal |
| birds | Tangara_cucullata | Tangara_cucullata | diurnal |
| birds | Tangara_cyanicollis | Tangara_cyanicollis | diurnal |
| birds | Tangara_cyanocephala | Tangara_cyanocephala | diurnal |
| birds | Tangara_cyanoptera | Thraupis_cyanoptera | diurnal |
| birds | Tangara_cyanotis | Tangara_cyanotis | diurnal |
| birds | Tangara_cyanoventris | Tangara_cyanoventris | diurnal |
| birds | Tangara_desmaresti | Tangara_desmaresti | diurnal |
| birds | Tangara_dowii | Tangara_dowii | diurnal |
| birds | Tangara_episcopus | Thraupis_episcopus | diurnal |
| birds | Tangara_fastuosa | Tangara_fastuosa | diurnal |
| birds | Tangara_florida | Tangara_florida | diurnal |
| birds | Tangara_fucosa | Tangara_fucosa | diurnal |
| birds | Tangara_glaucocolpa | Thraupis_glaucocolpa | diurnal |
| birds | Tangara_guttata | Tangara_guttata | diurnal |

| | | | |
|---|---|---|---|
| birds | Tangara_gyrola | Tangara_gyrola | diurnal |
| birds | Tangara_heinei | Tangara_heinei | diurnal |
| birds | Tangara_icterocephala | Tangara_icterocephala | diurnal |
| birds | Tangara_inornata | Tangara_inornata | diurnal |
| birds | Tangara_johannae | Tangara_johannae | diurnal |
| birds | Tangara_labradorides | Tangara_labradorides | diurnal |
| birds | Tangara_larvata | Tangara_larvata | diurnal |
| birds | Tangara_lavinia | Tangara_lavinia | diurnal |
| birds | Tangara_mexicana | Tangara_mexicana | diurnal |
| birds | Tangara_meyerdeschauenseei | Tangara_meyerdeschauenseei | diurnal |
| birds | Tangara_nigrocincta | Tangara_nigrocincta | diurnal |
| birds | Tangara_nigroviridis | Tangara_nigroviridis | diurnal |
| birds | Tangara_ornata | Thraupis_ornata | diurnal |
| birds | Tangara_palmarum | Thraupis_palmarum | diurnal |
| birds | Tangara_palmeri | Tangara_palmeri | diurnal |
| birds | Tangara_parzudakii | Tangara_parzudakii | diurnal |
| birds | Tangara_peruviana | Tangara_peruviana | diurnal |
| birds | Tangara_phillipsi | Tangara_phillipsi | diurnal |
| birds | Tangara_preciosa | Tangara_preciosa | diurnal |
| birds | Tangara_punctata | Tangara_punctata | diurnal |
| birds | Tangara_ruficervix | Tangara_ruficervix | diurnal |
| birds | Tangara_rufigenis | Tangara_rufigenis | diurnal |
| birds | Tangara_rufigula | Tangara_rufigula | diurnal |
| birds | Tangara_sayaca | Thraupis_sayaca | diurnal |
| birds | Tangara_schrankii | Tangara_schrankii | diurnal |
| birds | Tangara_seledon | Tangara_seledon | diurnal |
| birds | Tangara_varia | Tangara_varia | diurnal |
| birds | Tangara_vassorii | Tangara_vassorii | diurnal |
| birds | Tangara_velia | Tangara_velia | diurnal |
| birds | Tangara_viridicollis | Tangara_viridicollis | diurnal |
| birds | Tangara_vitriolina | Tangara_vitriolina | diurnal |
| birds | Tangara_xanthocephala | Tangara_xanthocephala | diurnal |
| birds | Tangara_xanthogastra | Tangara_xanthogastra | diurnal |
| birds | Tanygnathus_gramineus | Tanygnathus_gramineus | diurnal |

| | | | |
|---|---|---|---|
| birds | Tanygnathus_lucionensis | Tanygnathus_lucionensis | diurnal |
| birds | Tanygnathus_megalorynchos | Tanygnathus_megalorynchos | diurnal |
| birds | Tanygnathus_sumatranus | Tanygnathus_sumatranus | cathemeral |
| birds | Tanysiptera_carolinae | Tanysiptera_carolinae | diurnal |
| birds | Tanysiptera_danae | Tanysiptera_danae | diurnal |
| birds | Tanysiptera_ellioti | Tanysiptera_ellioti | diurnal |
| birds | Tanysiptera_galatea | Tanysiptera_galatea | diurnal |
| birds | Tanysiptera_hydrocharis | Tanysiptera_hydrocharis | diurnal |
| birds | Tanysiptera_nympha | Tanysiptera_nympha | diurnal |
| birds | Tanysiptera_riedelii | Tanysiptera_riedelii | diurnal |
| birds | Tanysiptera_sylvia | Tanysiptera_sylvia | diurnal |
| birds | Taoniscus_nanus | Taoniscus_nanus | diurnal |
| birds | Tapera_naevia | Tapera_naevia | diurnal |
| birds | Taphrolesbia_griseiventris | Taphrolesbia_griseiventris | diurnal |
| birds | Taphrospilus_hypostictus | Taphrospilus_hypostictus | diurnal |
| birds | Taraba_major | Taraba_major | diurnal |
| birds | Tarphonomus_certhioides | Tarphonomus_certhioides | diurnal |
| birds | Tarphonomus_harterti | Tarphonomus_harterti | diurnal |
| birds | Tarsiger_chrysaeus | Tarsiger_chrysaeus | diurnal |
| birds | Tarsiger_cyanurus | Tarsiger_cyanurus | diurnal |
| birds | Tarsiger_hyperythrus | Tarsiger_hyperythrus | diurnal |
| birds | Tarsiger_indicus | Tarsiger_indicus | diurnal |
| birds | Tarsiger_johnstoniae | Tarsiger_johnstoniae | diurnal |
| birds | Tauraco_bannermani | Tauraco_bannermani | diurnal |
| birds | Tauraco_corythaix | Tauraco_corythaix | diurnal |
| birds | Tauraco_erythrolophus | Tauraco_erythrolophus | diurnal |
| birds | Tauraco_fischeri | Tauraco_fischeri | diurnal |
| birds | Tauraco_hartlaubi | Tauraco_hartlaubi | diurnal |
| birds | Tauraco_leucolophus | Tauraco_leucolophus | diurnal |
| birds | Tauraco_leucotis | Tauraco_leucotis | diurnal |
| birds | Tauraco_livingstonii | Tauraco_livingstonii | diurnal |
| birds | Tauraco_macrorhynchus | Tauraco_macrorhynchus | diurnal |
| birds | Tauraco_persa | Tauraco_persa | diurnal |
| birds | Tauraco_ruspolii | Tauraco_ruspolii | diurnal |

| | | | |
|---|---|---|---|
| birds | Tauraco_schalowi | Tauraco_schalowi | diurnal |
| birds | Tauraco_schuettii | Tauraco_schuetti | diurnal |
| birds | Tchagra_australis | Tchagra_australis | diurnal |
| birds | Tchagra_jamesi | Tchagra_jamesi | diurnal |
| birds | Tchagra_senegalus | Tchagra_senegalus | diurnal |
| birds | Tchagra_tchagra | Tchagra_tchagra | diurnal |
| birds | Telacanthura_melanopygia | Telacanthura_melanopygia | diurnal |
| birds | Telacanthura_ussheri | Telacanthura_ussheri | diurnal |
| birds | Teledromas_fuscus | Teledromas_fuscus | diurnal |
| birds | Telespiza_cantans | Telespiza_cantans | diurnal |
| birds | Telespiza_ultima | Telespiza_ultima | diurnal |
| birds | Telophorus_dohertyi | Telophorus_dohertyi | diurnal |
| birds | Telophorus_viridis | Telophorus_viridis | diurnal |
| birds | Telophorus_zeylonus | Telophorus_zeylonus | diurnal |
| birds | Temnotrogon_roseigaster | Priotelus_roseigaster | diurnal |
| birds | Temnurus_temnurus | Temnurus_temnurus | diurnal |
| birds | Tephrodornis_pondicerianus | Tephrodornis_pondicerianus | diurnal |
| birds | Tephrodornis_sylvicola | Tephrodornis_gularis | diurnal |
| birds | Tephrophilus_wetmorei | Buthraupis_wetmorei | diurnal |
| birds | Tephrozosterops_stalkeri | Tephrozosterops_stalkeri | diurnal |
| birds | Terathopius_ecaudatus | Terathopius_ecaudatus | diurnal |
| birds | Terenotriccus_erythrurus | Terenotriccus_erythrurus | diurnal |
| birds | Terenura_maculata | Terenura_maculata | diurnal |
| birds | Terenura_sicki | Terenura_sicki | diurnal |
| birds | Teretistris_fernandinae | Teretistris_fernandinae | diurnal |
| birds | Teretistris_fornsi | Teretistris_fornsi | diurnal |
| birds | Terpsiphone_atrocaudata | Terpsiphone_atrocaudata | diurnal |
| birds | Terpsiphone_atrochalybeia | Terpsiphone_atrochalybeia | diurnal |
| birds | Terpsiphone_bedfordi | Terpsiphone_bedfordi | diurnal |
| birds | Terpsiphone_bourbonnensis | Terpsiphone_bourbonnensis | diurnal |
| birds | Terpsiphone_cinnamomea | Terpsiphone_cinnamomea | diurnal |
| birds | Terpsiphone_corvina | Terpsiphone_corvina | diurnal |
| birds | Terpsiphone_cyanescens | Terpsiphone_cyanescens | diurnal |
| birds | Terpsiphone_mutata | Terpsiphone_mutata | diurnal |

| | | | |
|---|---|---|---|
| birds | Terpsiphone_paradisi | Terpsiphone_paradisi | diurnal |
| birds | Terpsiphone_rufiventer | Terpsiphone_rufiventer | diurnal |
| birds | Terpsiphone_rufocinerea | Terpsiphone_rufocinerea | diurnal |
| birds | Terpsiphone_viridis | Terpsiphone_viridis | diurnal |
| birds | Tersina_viridis | Tersina_viridis | diurnal |
| birds | Tesia_cyaniventer | Tesia_cyaniventer | diurnal |
| birds | Tesia_everetti | Tesia_everetti | diurnal |
| birds | Tesia_olivea | Tesia_olivea | diurnal |
| birds | Tetrao_urogalloides | Tetrao_parvirostris | diurnal |
| birds | Tetrao_urogallus | Tetrao_urogallus | diurnal |
| birds | Tetraogallus_altaicus | Tetraogallus_altaicus | diurnal |
| birds | Tetraogallus_caspius | Tetraogallus_caspius | diurnal |
| birds | Tetraogallus_caucasicus | Tetraogallus_caucasicus | diurnal |
| birds | Tetraogallus_himalayensis | Tetraogallus_himalayensis | diurnal |
| birds | Tetraogallus_tibetanus | Tetraogallus_tibetanus | diurnal |
| birds | Tetraophasis_obscurus | Tetraophasis_obscurus | diurnal |
| birds | Tetraophasis_szechenyii | Tetraophasis_szechenyii | diurnal |
| birds | Tetrastes_bonasia | Bonasa_bonasia | diurnal |
| birds | Tetrastes_sewerzowi | Bonasa_sewerzowi | diurnal |
| birds | Tetrax_tetrax | Tetrax_tetrax | diurnal |
| birds | Thalassarche_bulleri | Thalassarche_bulleri | cathemeral |
| birds | Thalassarche_cauta | Thalassarche_cauta | cathemeral |
| birds | Thalassarche_chlororhynchos | Thalassarche_chlororhynchos | cathemeral |
| birds | Thalassarche_chrysostoma | Thalassarche_chrysostoma | cathemeral |
| birds | Thalassarche_melanophris | Thalassarche_melanophrys | cathemeral |
| birds | Thalasseus_bengalensis | Sterna_bengalensis | diurnal |
| birds | Thalasseus_bergii | Sterna_bergii | diurnal |
| birds | Thalasseus_bernsteini | Sterna_bernsteini | diurnal |
| birds | Thalasseus_elegans | Sterna_elegans | diurnal |
| birds | Thalasseus_sandvicensis | Sterna_sandvicensis | diurnal |
| birds | Thalassoica_antarctica | Thalassoica_antarctica | cathemeral |
| birds | Thalassornis_leuconotus | Thalassornis_leuconotus | diurnal |
| birds | Thalurania_colombica | Thalurania_colombica | diurnal |
| birds | Thalurania_furcata | Thalurania_furcata | diurnal |

| | | | |
|---|---|---|---|
| birds | Thalurania_glaucopis | Thalurania_glaucopis | diurnal |
| birds | Thalurania_watertonii | Thalurania_watertonii | diurnal |
| birds | Thamnistes_anabatinus | Thamnistes_anabatinus | diurnal |
| birds | Thamnolaea_cinnamomeiventris | Myrmecocichla_cinnamomeiventris | diurnal |
| birds | Thamnomanes_ardesiacus | Thamnomanes_ardesiacus | diurnal |
| birds | Thamnomanes_caesius | Thamnomanes_caesius | diurnal |
| birds | Thamnomanes_saturninus | Thamnomanes_saturninus | diurnal |
| birds | Thamnomanes_schistogynus | Thamnomanes_schistogynus | diurnal |
| birds | Thamnophilus_aethiops | Thamnophilus_aethiops | diurnal |
| birds | Thamnophilus_amazonicus | Thamnophilus_amazonicus | diurnal |
| birds | Thamnophilus_ambiguus | Thamnophilus_ambiguus | diurnal |
| birds | Thamnophilus_aroyae | Thamnophilus_aroyae | diurnal |
| birds | Thamnophilus_atrinucha | Thamnophilus_atrinucha | diurnal |
| birds | Thamnophilus_bernardi | Sakesphorus_bernardi | diurnal |
| birds | Thamnophilus_bridgesi | Thamnophilus_bridgesi | diurnal |
| birds | Thamnophilus_caerulescens | Thamnophilus_caerulescens | diurnal |
| birds | Thamnophilus_cryptoleucus | Thamnophilus_cryptoleucus | diurnal |
| birds | Thamnophilus_doliatus | Thamnophilus_doliatus | diurnal |
| birds | Thamnophilus_insignis | Thamnophilus_insignis | diurnal |
| birds | Thamnophilus_melanonotus | Sakesphorus_melanonotus | diurnal |
| birds | Thamnophilus_melanothorax | Sakesphorus_melanothorax | diurnal |
| birds | Thamnophilus_multistriatus | Thamnophilus_multistriatus | diurnal |
| birds | Thamnophilus_murinus | Thamnophilus_murinus | diurnal |
| birds | Thamnophilus_nigriceps | Thamnophilus_nigriceps | diurnal |
| birds | Thamnophilus_nigrocinereus | Thamnophilus_nigrocinereus | diurnal |
| birds | Thamnophilus_palliatus | Thamnophilus_palliatus | diurnal |
| birds | Thamnophilus_pelzelni | Thamnophilus_pelzelni | diurnal |
| birds | Thamnophilus_praecox | Thamnophilus_praecox | diurnal |
| birds | Thamnophilus_punctatus | Thamnophilus_punctatus | diurnal |
| birds | Thamnophilus_ruficapillus | Thamnophilus_ruficapillus | diurnal |
| birds | Thamnophilus_schistaceus | Thamnophilus_schistaceus | diurnal |
| birds | Thamnophilus_stictocephalus | Thamnophilus_stictocephalus | diurnal |
| birds | Thamnophilus_sticturus | Thamnophilus_sticturus | diurnal |
| birds | Thamnophilus_tenuepunctatus | Thamnophilus_tenuepunctatus | diurnal |

| | | | |
|---|---|---|---|
| birds | Thamnophilus_torquatus | Thamnophilus_torquatus | diurnal |
| birds | Thamnophilus_unicolor | Thamnophilus_unicolor | diurnal |
| birds | Thamnophilus_zarumae | Thamnophilus_zarumae | diurnal |
| birds | Thamnornis_chloropetoides | Thamnornis_chloropetoides | diurnal |
| birds | Thaumasius_baeri | Leucippus_baeri | diurnal |
| birds | Thaumasius_taczanowskii | Leucippus_taczanowskii | diurnal |
| birds | Thaumastura_cora | Thaumastura_cora | diurnal |
| birds | Thaumatibis_gigantea | Thaumatibis_gigantea | diurnal |
| birds | Theristicus_caerulescens | Theristicus_caerulescens | diurnal |
| birds | Theristicus_caudatus | Theristicus_caudatus | diurnal |
| birds | Theristicus_melanopis | Theristicus_melanopis | diurnal |
| birds | Thescelocichla_leucopleura | Thescelocichla_leucopleura | diurnal |
| birds | Thinocorus_orbignyianus | Thinocorus_orbignyianus | diurnal |
| birds | Thinocorus_rumicivorus | Thinocorus_rumicivorus | diurnal |
| birds | Thinornis_cucullatus | Thinornis_rubricollis | cathemeral |
| birds | Thinornis_novaeseelandiae | Thinornis_novaeseelandiae | cathemeral |
| birds | Thlypopsis_fulviceps | Thlypopsis_fulviceps | diurnal |
| birds | Thlypopsis_inornata | Thlypopsis_inornata | diurnal |
| birds | Thlypopsis_ornata | Thlypopsis_ornata | diurnal |
| birds | Thlypopsis_pectoralis | Thlypopsis_pectoralis | diurnal |
| birds | Thlypopsis_pyrrhocoma | Pyrrhocoma_ruficeps | diurnal |
| birds | Thlypopsis_ruficeps | Thlypopsis_ruficeps | diurnal |
| birds | Thlypopsis_sordida | Thlypopsis_sordida | diurnal |
| birds | Thlypopsis_superciliaris | Hemispingus_superciliaris | diurnal |
| birds | Threnetes_niger | Threnetes_niger | diurnal |
| birds | Threnetes_ruckeri | Threnetes_ruckeri | diurnal |
| birds | Threskiornis_aethiopicus | Threskiornis_aethiopicus | diurnal |
| birds | Threskiornis_melanocephalus | Threskiornis_melanocephalus | diurnal |
| birds | Threskiornis_moluccus | Threskiornis_molucca | diurnal |
| birds | Threskiornis_spinicollis | Threskiornis_spinicollis | diurnal |
| birds | Thripadectes_flammulatus | Thripadectes_flammulatus | diurnal |
| birds | Thripadectes_holostictus | Thripadectes_holostictus | diurnal |
| birds | Thripadectes_ignobilis | Thripadectes_ignobilis | diurnal |
| birds | Thripadectes_melanorhynchus | Thripadectes_melanorhynchus | diurnal |

| | | | |
|---|---|---|---|
| birds | Thripadectes_rufobrunneus | Thripadectes_rufobrunneus | diurnal |
| birds | Thripadectes_scrutator | Thripadectes_scrutator | diurnal |
| birds | Thripadectes_virgaticeps | Thripadectes_virgaticeps | diurnal |
| birds | Thripophaga_cherriei | Thripophaga_cherriei | diurnal |
| birds | Thripophaga_fusciceps | Thripophaga_fusciceps | diurnal |
| birds | Thripophaga_gutturata | Cranioleuca_gutturata | diurnal |
| birds | Thryomanes_bewickii | Thryomanes_bewickii | diurnal |
| birds | Thryophilus_pleurostictus | Thryothorus_pleurostictus | diurnal |
| birds | Thryophilus_rufalbus | Thryothorus_rufalbus | diurnal |
| birds | Thryophilus_sinaloa | Thryothorus_sinaloa | diurnal |
| birds | Thryorchilus_browni | Thryorchilus_browni | diurnal |
| birds | Thryothorus_ludovicianus | Thryothorus_ludovicianus | diurnal |
| birds | Tiaris_olivaceus | Tiaris_olivaceus | diurnal |
| birds | Tichodroma_muraria | Tichodroma_muraria | diurnal |
| birds | Tickellia_hodgsoni | Tickellia_hodgsoni | diurnal |
| birds | Tigrisoma_fasciatum | Tigrisoma_fasciatum | nocturnal |
| birds | Tigrisoma_lineatum | Tigrisoma_lineatum | cathemeral |
| birds | Tigrisoma_mexicanum | Tigrisoma_mexicanum | nocturnal |
| birds | Tilmatura_dupontii | Tilmatura_dupontii | diurnal |
| birds | Timalia_pileata | Timalia_pileata | diurnal |
| birds | Timeliopsis_fulvigula | Timeliopsis_fulvigula | diurnal |
| birds | Timeliopsis_griseigula | Timeliopsis_griseigula | diurnal |
| birds | Tinamotis_ingoufi | Tinamotis_ingoufi | diurnal |
| birds | Tinamotis_pentlandii | Tinamotis_pentlandii | diurnal |
| birds | Tinamus_guttatus | Tinamus_guttatus | diurnal |
| birds | Tinamus_major | Tinamus_major | diurnal |
| birds | Tinamus_solitarius | Tinamus_solitarius | diurnal |
| birds | Tinamus_tao | Tinamus_tao | diurnal |
| birds | Tityra_cayana | Tityra_cayana | diurnal |
| birds | Tityra_inquisitor | Tityra_inquisitor | diurnal |
| birds | Tityra_semifasciata | Tityra_semifasciata | diurnal |
| birds | Tmetothylacus_tenellus | Tmetothylacus_tenellus | diurnal |
| birds | Tockus_deckeni | Tockus_deckeni | diurnal |
| birds | Tockus_erythrorhynchus | Tockus_erythrorhynchus | diurnal |

| | | | |
|---|---|---|---|
| birds | Tockus_flavirostris | Tockus_flavirostris | diurnal |
| birds | Tockus_leucomelas | Tockus_leucomelas | diurnal |
| birds | Tockus_monteiri | Tockus_monteiri | diurnal |
| birds | Todiramphus_albonotatus | Todiramphus_albonotatus | diurnal |
| birds | Todiramphus_australasia | Todiramphus_australasia | diurnal |
| birds | Todiramphus_chloris | Todiramphus_chloris | diurnal |
| birds | Todiramphus_cinnamominus | Todiramphus_cinnamominus | diurnal |
| birds | Todiramphus_diops | Todiramphus_diops | diurnal |
| birds | Todiramphus_farquhari | Todiramphus_farquhari | diurnal |
| birds | Todiramphus_gambieri | Todiramphus_gambieri | diurnal |
| birds | Todiramphus_godeffroyi | Todiramphus_godeffroyi | diurnal |
| birds | Todiramphus_leucopygius | Todiramphus_leucopygius | diurnal |
| birds | Todiramphus_macleayii | Todiramphus_macleayii | diurnal |
| birds | Todiramphus_nigrocyaneus | Todiramphus_nigrocyaneus | diurnal |
| birds | Todiramphus_pyrrhopygius | Todiramphus_pyrrhopygius | diurnal |
| birds | Todiramphus_sanctus | Todiramphus_sanctus | diurnal |
| birds | Todiramphus_saurophagus | Todiramphus_saurophaga | diurnal |
| birds | Todiramphus_tutus | Todiramphus_tutus | diurnal |
| birds | Todiramphus_veneratus | Todiramphus_veneratus | diurnal |
| birds | Todiramphus_winchelli | Todiramphus_winchelli | diurnal |
| birds | Todirostrum_chrysocrotaphum | Todirostrum_chrysocrotaphum | diurnal |
| birds | Todirostrum_cinereum | Todirostrum_cinereum | diurnal |
| birds | Todirostrum_maculatum | Todirostrum_maculatum | diurnal |
| birds | Todirostrum_nigriceps | Todirostrum_nigriceps | diurnal |
| birds | Todirostrum_pictum | Todirostrum_pictum | diurnal |
| birds | Todirostrum_poliocephalum | Todirostrum_poliocephalum | diurnal |
| birds | Todirostrum_viridanum | Todirostrum_viridanum | diurnal |
| birds | Todus_angustirostris | Todus_angustirostris | diurnal |
| birds | Todus_mexicanus | Todus_mexicanus | diurnal |
| birds | Todus_multicolor | Todus_multicolor | diurnal |
| birds | Todus_subulatus | Todus_subulatus | diurnal |
| birds | Todus_todus | Todus_todus | diurnal |
| birds | Tolmomyias_assimilis | Tolmomyias_assimilis | diurnal |
| birds | Tolmomyias_flaviventris | Tolmomyias_flaviventris | diurnal |

| | | | |
|---|---|---|---|
| birds | Tolmomyias_flavotectus | Tolmomyias_flavotectus | diurnal |
| birds | Tolmomyias_poliocephalus | Tolmomyias_poliocephalus | diurnal |
| birds | Tolmomyias_sulphurescens | Tolmomyias_sulphurescens | diurnal |
| birds | Tolmomyias_traylori | Tolmomyias_traylori | diurnal |
| birds | Tolmomyias_viridiceps | Tolmomyias_viridiceps | diurnal |
| birds | Topaza_pella | Topaza_pella | diurnal |
| birds | Torgos_tracheliotos | Torgos_tracheliotos | diurnal |
| birds | Torreornis_inexpectata | Torreornis_inexpectata | diurnal |
| birds | Touit_costaricensis | Touit_costaricensis | diurnal |
| birds | Touit_dilectissimus | Touit_dilectissimus | diurnal |
| birds | Touit_huetii | Touit_huetii | diurnal |
| birds | Touit_melanonotus | Touit_melanonotus | diurnal |
| birds | Touit_purpuratus | Touit_purpuratus | diurnal |
| birds | Touit_stictopterus | Touit_stictopterus | diurnal |
| birds | Touit_surdus | Touit_surdus | diurnal |
| birds | Toxorhamphus_novaeguineae | Toxorhamphus_novaeguineae | diurnal |
| birds | Toxorhamphus_poliopterus | Toxorhamphus_poliopterus | diurnal |
| birds | Toxostoma_bendirei | Toxostoma_bendirei | diurnal |
| birds | Toxostoma_cinereum | Toxostoma_cinereum | diurnal |
| birds | Toxostoma_crissale | Toxostoma_crissale | diurnal |
| birds | Toxostoma_curvirostre | Toxostoma_curvirostre | diurnal |
| birds | Toxostoma_guttatum | Toxostoma_guttatum | diurnal |
| birds | Toxostoma_lecontei | Toxostoma_lecontei | diurnal |
| birds | Toxostoma_longirostre | Toxostoma_longirostre | diurnal |
| birds | Toxostoma_ocellatum | Toxostoma_ocellatum | diurnal |
| birds | Toxostoma_redivivum | Toxostoma_redivivum | diurnal |
| birds | Toxostoma_rufum | Toxostoma_rufum | diurnal |
| birds | Trachylaemus_purpuratus | Trachyphonus_purpuratus | diurnal |
| birds | Trachyphonus_darnaudii | Trachyphonus_darnaudii | diurnal |
| birds | Trachyphonus_erythrocephalus | Trachyphonus_erythrocephalus | diurnal |
| birds | Trachyphonus_margaritatus | Trachyphonus_margaritatus | diurnal |
| birds | Trachyphonus_vaillantii | Trachyphonus_vaillantii | diurnal |
| birds | Tragopan_blythii | Tragopan_blythii | diurnal |
| birds | Tragopan_caboti | Tragopan_caboti | diurnal |

| | | | |
|---|---|---|---|
| birds | Tragopan_melanocephalus | Tragopan_melanocephalus | diurnal |
| birds | Tragopan_satyra | Tragopan_satyra | diurnal |
| birds | Tragopan_temminckii | Tragopan_temminckii | diurnal |
| birds | Tregellasia_capito | Tregellasia_capito | diurnal |
| birds | Tregellasia_leucops | Tregellasia_leucops | diurnal |
| birds | Treron_apicauda | Treron_apicauda | diurnal |
| birds | Treron_australis | Treron_australis | diurnal |
| birds | Treron_bicinctus | Treron_bicinctus | diurnal |
| birds | Treron_calvus | Treron_calvus | diurnal |
| birds | Treron_capellei | Treron_capellei | diurnal |
| birds | Treron_curvirostra | Treron_curvirostra | diurnal |
| birds | Treron_floris | Treron_floris | diurnal |
| birds | Treron_formosae | Treron_formosae | diurnal |
| birds | Treron_fulvicollis | Treron_fulvicollis | diurnal |
| birds | Treron_griseicauda | Treron_griseicauda | diurnal |
| birds | Treron_olax | Treron_olax | diurnal |
| birds | Treron_oxyurus | Treron_oxyurus | diurnal |
| birds | Treron_pembaensis | Treron_pembaensis | diurnal |
| birds | Treron_phoenicopterus | Treron_phoenicopterus | diurnal |
| birds | Treron_pompadora | Treron_pompadora | diurnal |
| birds | Treron_psittaceus | Treron_psittaceus | diurnal |
| birds | Treron_sanctithomae | Treron_sanctithomae | diurnal |
| birds | Treron_seimundi | Treron_seimundi | diurnal |
| birds | Treron_sieboldii | Treron_sieboldii | diurnal |
| birds | Treron_sphenurus | Treron_sphenurus | diurnal |
| birds | Treron_teysmannii | Treron_teysmannii | diurnal |
| birds | Treron_vernans | Treron_vernans | diurnal |
| birds | Treron_waalia | Treron_waalia | diurnal |
| birds | Tribonyx_mortierii | Gallinula_mortierii | diurnal |
| birds | Tribonyx_ventralis | Gallinula_ventralis | diurnal |
| birds | Trichodere_cockerelli | Trichodere_cockerelli | diurnal |
| birds | Trichoglossus_chlorolepidotus | Trichoglossus_chlorolepidotus | diurnal |
| birds | Trichoglossus_euteles | Trichoglossus_euteles | diurnal |
| birds | Trichoglossus_flavoviridis | Trichoglossus_flavoviridis | diurnal |

| | | | |
|---|---|---|---|
| birds | Trichoglossus_haematodus | Trichoglossus_haematodus | diurnal |
| birds | Trichoglossus_johnstoniae | Trichoglossus_johnstoniae | diurnal |
| birds | Trichoglossus_ornatus | Trichoglossus_ornatus | diurnal |
| birds | Trichoglossus_rubiginosus | Trichoglossus_rubiginosus | diurnal |
| birds | Tricholaema_diademata | Tricholaema_diademata | diurnal |
| birds | Tricholaema_frontata | Tricholaema_frontata | diurnal |
| birds | Tricholaema_hirsuta | Tricholaema_hirsuta | diurnal |
| birds | Tricholaema_lacrymosa | Tricholaema_lacrymosa | diurnal |
| birds | Tricholaema_leucomelas | Tricholaema_leucomelas | diurnal |
| birds | Tricholaema_melanocephala | Tricholaema_melanocephala | diurnal |
| birds | Tricholestes_criniger | Tricholestes_criniger | diurnal |
| birds | Trichothraupis_melanops | Trichothraupis_melanops | diurnal |
| birds | Triclaria_malachitacea | Triclaria_malachitacea | diurnal |
| birds | Trigonoceps_occipitalis | Trigonoceps_occipitalis | diurnal |
| birds | Tringa_brevipes | Heteroscelus_brevipes | diurnal |
| birds | Tringa_erythropus | Tringa_erythropus | cathemeral |
| birds | Tringa_flavipes | Tringa_flavipes | cathemeral |
| birds | Tringa_glareola | Tringa_glareola | diurnal |
| birds | Tringa_guttifer | Tringa_guttifer | diurnal |
| birds | Tringa_incana | Heteroscelus_incanus | diurnal |
| birds | Tringa_melanoleuca | Tringa_melanoleuca | cathemeral |
| birds | Tringa_nebularia | Tringa_nebularia | cathemeral |
| birds | Tringa_ochropus | Tringa_ochropus | diurnal |
| birds | Tringa_semipalmata | Catoptrophorus_semipalmatus | cathemeral |
| birds | Tringa_solitaria | Tringa_solitaria | diurnal |
| birds | Tringa_stagnatilis | Tringa_stagnatilis | diurnal |
| birds | Tringa_totanus | Tringa_totanus | cathemeral |
| birds | Trochalopteron_affine | Garrulax_affinis | diurnal |
| birds | Trochalopteron_austeni | Garrulax_austeni | diurnal |
| birds | Trochalopteron_elliotii | Garrulax_elliotii | diurnal |
| birds | Trochalopteron_erythrocephalum | Garrulax_erythrocephalus | diurnal |
| birds | Trochalopteron_formosum | Garrulax_formosus | diurnal |
| birds | Trochalopteron_henrici | Garrulax_henrici | diurnal |
| birds | Trochalopteron_lineatum | Garrulax_lineatus | diurnal |

| | | | |
|---|---|---|---|
| birds | Trochalopteron_milnei | Garrulax_milnei | diurnal |
| birds | Trochalopteron_morrisonianum | Garrulax_morrisonianus | diurnal |
| birds | Trochalopteron_squamatum | Garrulax_squamatus | diurnal |
| birds | Trochalopteron_subunicolor | Garrulax_subunicolor | diurnal |
| birds | Trochalopteron_variegatum | Garrulax_variegatus | diurnal |
| birds | Trochalopteron_virgatum | Garrulax_virgatus | diurnal |
| birds | Trochalopteron_yersini | Garrulax_yersini | diurnal |
| birds | Trochilus_polytmus | Trochilus_polytmus | diurnal |
| birds | Trochilus_scitulus | Trochilus_scitulus | diurnal |
| birds | Trochocercus_cyanomelas | Trochocercus_cyanomelas | diurnal |
| birds | Trochocercus_nitens | Trochocercus_nitens | diurnal |
| birds | Troglodytes_aedon | Troglodytes_aedon | diurnal |
| birds | Troglodytes_cobbi | Troglodytes_cobbi | diurnal |
| birds | Troglodytes_monticola | Troglodytes_monticola | diurnal |
| birds | Troglodytes_ochraceus | Troglodytes_ochraceus | diurnal |
| birds | Troglodytes_rufociliatus | Troglodytes_rufociliatus | diurnal |
| birds | Troglodytes_rufulus | Troglodytes_rufulus | diurnal |
| birds | Troglodytes_sissonii | Troglodytes_sissonii | diurnal |
| birds | Troglodytes_solstitialis | Troglodytes_solstitialis | diurnal |
| birds | Troglodytes_tanneri | Troglodytes_tanneri | diurnal |
| birds | Troglodytes_troglodytes | Troglodytes_troglodytes | diurnal |
| birds | Trogon_bairdii | Trogon_bairdii | diurnal |
| birds | Trogon_citreolus | Trogon_citreolus | diurnal |
| birds | Trogon_clathratus | Trogon_clathratus | diurnal |
| birds | Trogon_collaris | Trogon_collaris | diurnal |
| birds | Trogon_comptus | Trogon_comptus | diurnal |
| birds | Trogon_curucui | Trogon_curucui | diurnal |
| birds | Trogon_elegans | Trogon_elegans | diurnal |
| birds | Trogon_massena | Trogon_massena | diurnal |
| birds | Trogon_melanocephalus | Trogon_melanocephalus | diurnal |
| birds | Trogon_melanurus | Trogon_melanurus | diurnal |
| birds | Trogon_mexicanus | Trogon_mexicanus | diurnal |
| birds | Trogon_personatus | Trogon_personatus | diurnal |
| birds | Trogon_rufus | Trogon_rufus | diurnal |

| | | | |
|---|---|---|---|
| birds | Trogon_surrucura | Trogon_surrucura | diurnal |
| birds | Trogon_violaceus | Trogon_violaceus | diurnal |
| birds | Trogon_viridis | Trogon_viridis | diurnal |
| birds | Trugon_terrestris | Trugon_terrestris | diurnal |
| birds | Tunchiornis_ochraceiceps | Hylophilus_ochraceiceps | diurnal |
| birds | Turacoena_manadensis | Turacoena_manadensis | diurnal |
| birds | Turacoena_modesta | Turacoena_modesta | diurnal |
| birds | Turdinus_atrigularis | Turdinus_atrigularis | diurnal |
| birds | Turdinus_macrodactylus | Turdinus_macrodactylus | diurnal |
| birds | Turdinus_marmoratus | Turdinus_marmorata | diurnal |
| birds | Turdoides_bicolor | Turdoides_bicolor | diurnal |
| birds | Turdoides_gymnogenys | Turdoides_gymnogenys | diurnal |
| birds | Turdoides_hartlaubii | Turdoides_hartlaubii | diurnal |
| birds | Turdoides_hindei | Turdoides_hindei | diurnal |
| birds | Turdoides_hypoleuca | Turdoides_hypoleuca | diurnal |
| birds | Turdoides_jardineii | Turdoides_jardineii | diurnal |
| birds | Turdoides_leucocephala | Turdoides_leucocephala | diurnal |
| birds | Turdoides_leucopygia | Turdoides_leucopygia | diurnal |
| birds | Turdoides_melanops | Turdoides_melanops | diurnal |
| birds | Turdoides_plebejus | Turdoides_plebejus | diurnal |
| birds | Turdoides_reinwardtii | Turdoides_reinwardii | diurnal |
| birds | Turdoides_sharpei | Turdoides_sharpei | diurnal |
| birds | Turdoides_squamulata | Turdoides_squamulata | diurnal |
| birds | Turdoides_tenebrosa | Turdoides_tenebrosa | diurnal |
| birds | Turdus_albicollis | Turdus_albicollis | diurnal |
| birds | Turdus_albocinctus | Turdus_albocinctus | diurnal |
| birds | Turdus_amaurochalinus | Turdus_amaurochalinus | diurnal |
| birds | Turdus_assimilis | Turdus_assimilis | diurnal |
| birds | Turdus_aurantius | Turdus_aurantius | diurnal |
| birds | Turdus_bewsheri | Turdus_bewsheri | diurnal |
| birds | Turdus_boulboul | Turdus_boulboul | diurnal |
| birds | Turdus_cardis | Turdus_cardis | diurnal |
| birds | Turdus_celaenops | Turdus_celaenops | diurnal |
| birds | Turdus_chiguanco | Turdus_chiguanco | diurnal |

| | | | |
|---|---|---|---|
| birds | Turdus_chrysolaus | Turdus_chrysolaus | diurnal |
| birds | Turdus_dissimilis | Turdus_dissimilis | diurnal |
| birds | Turdus_eremita | Nesocichla_eremita | diurnal |
| birds | Turdus_falcklandii | Turdus_falcklandii | diurnal |
| birds | Turdus_feae | Turdus_feae | diurnal |
| birds | Turdus_flavipes | Turdus_flavipes | diurnal |
| birds | Turdus_fulviventris | Turdus_fulviventris | diurnal |
| birds | Turdus_fumigatus | Turdus_fumigatus | diurnal |
| birds | Turdus_fuscater | Turdus_fuscater | diurnal |
| birds | Turdus_grayi | Turdus_grayi | diurnal |
| birds | Turdus_haplochrous | Turdus_haplochrous | diurnal |
| birds | Turdus_hauxwelli | Turdus_hauxwelli | diurnal |
| birds | Turdus_helleri | Turdus_helleri | diurnal |
| birds | Turdus_hortulorum | Turdus_hortulorum | diurnal |
| birds | Turdus_ignobilis | Turdus_ignobilis | diurnal |
| birds | Turdus_infuscatus | Turdus_infuscatus | diurnal |
| birds | Turdus_jamaicensis | Turdus_jamaicensis | diurnal |
| birds | Turdus_kessleri | Turdus_kessleri | diurnal |
| birds | Turdus_lawrencii | Turdus_lawrencii | diurnal |
| birds | Turdus_leucomelas | Turdus_leucomelas | diurnal |
| birds | Turdus_leucops | Turdus_leucops | diurnal |
| birds | Turdus_lherminieri | Turdus_lherminieri | diurnal |
| birds | Turdus_libonyana | Turdus_libonyanus | diurnal |
| birds | Turdus_ludoviciae | Turdus_ludoviciae | diurnal |
| birds | Turdus_maculirostris | Turdus_maculirostris | diurnal |
| birds | Turdus_maranonicus | Turdus_maranonicus | diurnal |
| birds | Turdus_menachensis | Turdus_menachensis | diurnal |
| birds | Turdus_merula | Turdus_merula | diurnal |
| birds | Turdus_migratorius | Turdus_migratorius | diurnal |
| birds | Turdus_naumanni | Turdus_naumanni | diurnal |
| birds | Turdus_nigrescens | Turdus_nigrescens | diurnal |
| birds | Turdus_nigriceps | Turdus_nigriceps | diurnal |
| birds | Turdus_nudigenis | Turdus_nudigenis | diurnal |
| birds | Turdus_obscurus | Turdus_obscurus | diurnal |

| | | | |
|---|---|---|---|
| birds | Turdus_obsoletus | Turdus_obsoletus | diurnal |
| birds | Turdus_olivaceofuscus | Turdus_olivaceofuscus | diurnal |
| birds | Turdus_olivaceus | Turdus_olivaceus | diurnal |
| birds | Turdus_olivater | Turdus_olivater | diurnal |
| birds | Turdus_pallidus | Turdus_pallidus | diurnal |
| birds | Turdus_philomelos | Turdus_philomelos | diurnal |
| birds | Turdus_pilaris | Turdus_pilaris | diurnal |
| birds | Turdus_plebejus | Turdus_plebejus | diurnal |
| birds | Turdus_plumbeus | Turdus_plumbeus | diurnal |
| birds | Turdus_poliocephalus | Turdus_poliocephalus | diurnal |
| birds | Turdus_reevei | Turdus_reevei | diurnal |
| birds | Turdus_rubrocanus | Turdus_rubrocanus | diurnal |
| birds | Turdus_ruficollis | Turdus_ruficollis | diurnal |
| birds | Turdus_rufitorques | Turdus_rufitorques | diurnal |
| birds | Turdus_rufiventris | Turdus_rufiventris | diurnal |
| birds | Turdus_rufopalliatus | Turdus_rufopalliatus | diurnal |
| birds | Turdus_serranus | Turdus_serranus | diurnal |
| birds | Turdus_subalaris | Turdus_subalaris | diurnal |
| birds | Turdus_swalesi | Turdus_swalesi | diurnal |
| birds | Turdus_tephronotus | Turdus_tephronotus | diurnal |
| birds | Turdus_torquatus | Turdus_torquatus | diurnal |
| birds | Turdus_unicolor | Turdus_unicolor | diurnal |
| birds | Turdus_viscivorus | Turdus_viscivorus | diurnal |
| birds | Turnix_castanotus | Turnix_castanotus | nocturnal |
| birds | Turnix_hottentottus | Turnix_hottentottus | diurnal |
| birds | Turnix_maculosus | Turnix_maculosus | nocturnal |
| birds | Turnix_melanogaster | Turnix_melanogaster | cathemeral |
| birds | Turnix_nigricollis | Turnix_nigricollis | diurnal |
| birds | Turnix_olivii | Turnix_olivii | diurnal |
| birds | Turnix_pyrrhothorax | Turnix_pyrrhothorax | cathemeral |
| birds | Turnix_suscitator | Turnix_suscitator | diurnal |
| birds | Turnix_sylvaticus | Turnix_sylvaticus | nocturnal |
| birds | Turnix_tanki | Turnix_tanki | diurnal |
| birds | Turnix_varius | Turnix_varius | cathemeral |

| | | | |
|---|---|---|---|
| birds | Turnix_velox | Turnix_velox | cathemeral |
| birds | Turtur_abyssinicus | Turtur_abyssinicus | diurnal |
| birds | Turtur_afer | Turtur_afer | diurnal |
| birds | Turtur_brehmeri | Turtur_brehmeri | diurnal |
| birds | Turtur_chalcospilos | Turtur_chalcospilos | diurnal |
| birds | Turtur_tympanistria | Turtur_tympanistria | diurnal |
| birds | Tychaedon_barbata | Erythropygia_barbata | diurnal |
| birds | Tychaedon_coryphoeus | Erythropygia_coryphaeus | diurnal |
| birds | Tychaedon_leucosticta | Erythropygia_leucosticta | diurnal |
| birds | Tychaedon_quadrivirgata | Erythropygia_quadrivirgata | diurnal |
| birds | Tychaedon_signata | Erythropygia_signata | diurnal |
| birds | Tylas_eduardi | Tylas_eduardi | diurnal |
| birds | Tympanuchus_pallidicinctus | Tympanuchus_pallidicinctus | diurnal |
| birds | Tympanuchus_phasianellus | Tympanuchus_phasianellus | diurnal |
| birds | Tyranneutes_stolzmanni | Tyranneutes_stolzmanni | diurnal |
| birds | Tyranneutes_virescens | Tyranneutes_virescens | diurnal |
| birds | Tyrannopsis_sulphurea | Tyrannopsis_sulphurea | diurnal |
| birds | Tyrannulus_elatus | Tyrannulus_elatus | diurnal |
| birds | Tyrannus_albogularis | Tyrannus_albogularis | diurnal |
| birds | Tyrannus_caudifasciatus | Tyrannus_caudifasciatus | diurnal |
| birds | Tyrannus_couchii | Tyrannus_couchii | diurnal |
| birds | Tyrannus_crassirostris | Tyrannus_crassirostris | diurnal |
| birds | Tyrannus_cubensis | Tyrannus_cubensis | diurnal |
| birds | Tyrannus_dominicensis | Tyrannus_dominicensis | diurnal |
| birds | Tyrannus_forficatus | Tyrannus_forficatus | diurnal |
| birds | Tyrannus_melancholicus | Tyrannus_melancholicus | diurnal |
| birds | Tyrannus_niveigularis | Tyrannus_niveigularis | diurnal |
| birds | Tyrannus_savana | Tyrannus_savana | diurnal |
| birds | Tyrannus_tyrannus | Tyrannus_tyrannus | diurnal |
| birds | Tyrannus_verticalis | Tyrannus_verticalis | diurnal |
| birds | Tyrannus_vociferans | Tyrannus_vociferans | diurnal |
| birds | Tyto_alba | Tyto_alba | cathemeral |
| birds | Tyto_capensis | Tyto_capensis | nocturnal |
| birds | Tyto_glaucops | Tyto_glaucops | nocturnal |

| | | | |
|---|---|---|---|
| birds | Tyto_longimembris | Tyto_longimembris | nocturnal |
| birds | Tyto_novaehollandiae | Tyto_novaehollandiae | nocturnal |
| birds | Tyto_soumagnei | Tyto_soumagnei | nocturnal |
| birds | Tyto_tenebricosa | Tyto_tenebricosa | nocturnal |
| birds | Upucerthia_albigula | Upucerthia_albigula | diurnal |
| birds | Upucerthia_dumetaria | Upucerthia_dumetaria | diurnal |
| birds | Upucerthia_validirostris | Upucerthia_validirostris | diurnal |
| birds | Upupa_epops | Upupa_epops | diurnal |
| birds | Uraeginthus_angolensis | Uraeginthus_angolensis | diurnal |
| birds | Uraeginthus_bengalus | Uraeginthus_bengalus | diurnal |
| birds | Uraeginthus_cyanocephalus | Uraeginthus_cyanocephalus | diurnal |
| birds | Uratelornis_chimaera | Uratelornis_chimaera | cathemeral |
| birds | Uria_aalge | Uria_aalge | diurnal |
| birds | Uria_lomvia | Uria_lomvia | diurnal |
| birds | Urile_penicillatus | Phalacrocorax_penicillatus | diurnal |
| birds | Urile_urile | Phalacrocorax_urile | diurnal |
| birds | Urochroa_bougueri | Urochroa_bougueri | diurnal |
| birds | Urocissa_caerulea | Urocissa_caerulea | diurnal |
| birds | Urocissa_erythroryncha | Urocissa_erythrorhyncha | diurnal |
| birds | Urocissa_flavirostris | Urocissa_flavirostris | diurnal |
| birds | Urocissa_ornata | Urocissa_ornata | diurnal |
| birds | Urocissa_whiteheadi | Urocissa_whiteheadi | diurnal |
| birds | Urocolius_indicus | Urocolius_indicus | diurnal |
| birds | Urocolius_macrourus | Urocolius_macrourus | diurnal |
| birds | Urocynchramus_pylzowi | Urocynchramus_pylzowi | diurnal |
| birds | Urodynamis_taitensis | Eudynamys_taitensis | diurnal |
| birds | Urolais_epichlorus | Urolais_epichlorus | diurnal |
| birds | Uromyias_agilis | Anairetes_agilis | diurnal |
| birds | Uromyias_agraphia | Anairetes_agraphia | diurnal |
| birds | Uropelia_campestris | Uropelia_campestris | diurnal |
| birds | Uropsalis_lyra | Uropsalis_lyra | nocturnal |
| birds | Uropsalis_segmentata | Uropsalis_segmentata | nocturnal |
| birds | Uropsila_leucogastra | Uropsila_leucogastra | diurnal |
| birds | Urosphena_squameiceps | Urosphena_squameiceps | diurnal |

| | | | |
|---|---|---|---|
| birds | Urosphena_subulata | Urosphena_subulata | diurnal |
| birds | Urosphena_whiteheadi | Urosphena_whiteheadi | diurnal |
| birds | Urosticte_benjamini | Urosticte_benjamini | diurnal |
| birds | Urosticte_ruficrissa | Urosticte_ruficrissa | diurnal |
| birds | Urothraupis_stolzmanni | Urothraupis_stolzmanni | diurnal |
| birds | Urotriorchis_macrourus | Urotriorchis_macrourus | diurnal |
| birds | Vanellus_albiceps | Vanellus_albiceps | diurnal |
| birds | Vanellus_armatus | Vanellus_armatus | diurnal |
| birds | Vanellus_chilensis | Vanellus_chilensis | diurnal |
| birds | Vanellus_cinereus | Vanellus_cinereus | diurnal |
| birds | Vanellus_coronatus | Vanellus_coronatus | nocturnal |
| birds | Vanellus_crassirostris | Vanellus_crassirostris | diurnal |
| birds | Vanellus_duvaucelii | Vanellus_duvaucelii | diurnal |
| birds | Vanellus_gregarius | Vanellus_gregarius | diurnal |
| birds | Vanellus_leucurus | Vanellus_leucurus | diurnal |
| birds | Vanellus_lugubris | Vanellus_lugubris | cathemeral |
| birds | Vanellus_malabaricus | Vanellus_malarbaricus | nocturnal |
| birds | Vanellus_melanocephalus | Vanellus_melanocephalus | diurnal |
| birds | Vanellus_melanopterus | Vanellus_melanopterus | diurnal |
| birds | Vanellus_miles | Vanellus_miles | cathemeral |
| birds | Vanellus_senegallus | Vanellus_senegallus | diurnal |
| birds | Vanellus_spinosus | Vanellus_spinosus | diurnal |
| birds | Vanellus_superciliosus | Vanellus_superciliosus | cathemeral |
| birds | Vanellus_tectus | Vanellus_tectus | nocturnal |
| birds | Vanellus_vanellus | Vanellus_vanellus | cathemeral |
| birds | Vanga_curvirostris | Vanga_curvirostris | diurnal |
| birds | Vauriella_albigularis | Rhinomyias_albigularis | diurnal |
| birds | Vauriella_goodfellowi | Rhinomyias_goodfellowi | diurnal |
| birds | Vauriella_gularis | Rhinomyias_gularis | diurnal |
| birds | Vauriella_insignis | Rhinomyias_insignis | diurnal |
| birds | Veles_binotatus | Caprimulgus_binotatus | nocturnal |
| birds | Veniliornis_affinis | Veniliornis_affinis | diurnal |
| birds | Veniliornis_callonotus | Veniliornis_callonotus | diurnal |
| birds | Veniliornis_cassini | Veniliornis_cassini | diurnal |

| | | | |
|---|---|---|---|
| birds | Veniliornis_chocoensis | Veniliornis_chocoensis | diurnal |
| birds | Veniliornis_dignus | Veniliornis_dignus | diurnal |
| birds | Veniliornis_frontalis | Veniliornis_frontalis | diurnal |
| birds | Veniliornis_kirkii | Veniliornis_kirkii | diurnal |
| birds | Veniliornis_lignarius | Veniliornis_lignarius | diurnal |
| birds | Veniliornis_maculifrons | Veniliornis_maculifrons | diurnal |
| birds | Veniliornis_mixtus | Veniliornis_mixtus | diurnal |
| birds | Veniliornis_nigriceps | Veniliornis_nigriceps | diurnal |
| birds | Veniliornis_passerinus | Veniliornis_passerinus | diurnal |
| birds | Veniliornis_sanguineus | Veniliornis_sanguineus | diurnal |
| birds | Veniliornis_spilogaster | Veniliornis_spilogaster | diurnal |
| birds | Vermivora_chrysoptera | Vermivora_chrysoptera | diurnal |
| birds | Vermivora_cyanoptera | Vermivora_pinus | diurnal |
| birds | Verreauxia_africana | Sasia_africana | diurnal |
| birds | Vidua_camerunensis | Vidua_camerunensis | diurnal |
| birds | Vidua_chalybeata | Vidua_chalybeata | diurnal |
| birds | Vidua_codringtoni | Vidua_codringtoni | diurnal |
| birds | Vidua_fischeri | Vidua_fischeri | diurnal |
| birds | Vidua_funerea | Vidua_funerea | diurnal |
| birds | Vidua_hypocherina | Vidua_hypocherina | diurnal |
| birds | Vidua_interjecta | Vidua_interjecta | diurnal |
| birds | Vidua_larvaticola | Vidua_larvaticola | diurnal |
| birds | Vidua_macroura | Vidua_macroura | diurnal |
| birds | Vidua_maryae | Vidua_maryae | diurnal |
| birds | Vidua_nigeriae | Vidua_nigeriae | diurnal |
| birds | Vidua_obtusa | Vidua_obtusa | diurnal |
| birds | Vidua_orientalis | Vidua_orientalis | diurnal |
| birds | Vidua_paradisaea | Vidua_paradisaea | diurnal |
| birds | Vidua_purpurascens | Vidua_purpurascens | diurnal |
| birds | Vidua_raricola | Vidua_raricola | diurnal |
| birds | Vidua_regia | Vidua_regia | diurnal |
| birds | Vidua_togoensis | Vidua_togoensis | diurnal |
| birds | Vidua_wilsoni | Vidua_wilsoni | diurnal |
| birds | Vini_australis | Vini_australis | diurnal |

| | | | |
|---|---|---|---|
| birds | Vini_kuhlii | Vini_kuhlii | diurnal |
| birds | Vini_peruviana | Vini_peruviana | diurnal |
| birds | Vini_stepheni | Vini_stepheni | diurnal |
| birds | Vireo_altiloquus | Vireo_altiloquus | diurnal |
| birds | Vireo_atricapilla | Vireo_atricapilla | diurnal |
| birds | Vireo_bairdi | Vireo_bairdi | diurnal |
| birds | Vireo_bellii | Vireo_bellii | diurnal |
| birds | Vireo_brevipennis | Vireo_brevipennis | diurnal |
| birds | Vireo_caribaeus | Vireo_caribaeus | diurnal |
| birds | Vireo_carmioli | Vireo_carmioli | diurnal |
| birds | Vireo_cassinii | Vireo_cassinii | diurnal |
| birds | Vireo_crassirostris | Vireo_crassirostris | diurnal |
| birds | Vireo_flavifrons | Vireo_flavifrons | diurnal |
| birds | Vireo_flavoviridis | Vireo_flavoviridis | diurnal |
| birds | Vireo_gilvus | Vireo_gilvus | diurnal |
| birds | Vireo_gracilirostris | Vireo_gracilirostris | diurnal |
| birds | Vireo_griseus | Vireo_griseus | diurnal |
| birds | Vireo_gundlachii | Vireo_gundlachii | diurnal |
| birds | Vireo_huttoni | Vireo_huttoni | diurnal |
| birds | Vireo_latimeri | Vireo_latimeri | diurnal |
| birds | Vireo_leucophrys | Vireo_leucophrys | diurnal |
| birds | Vireo_magister | Vireo_magister | diurnal |
| birds | Vireo_masteri | Vireo_masteri | diurnal |
| birds | Vireo_modestus | Vireo_modestus | diurnal |
| birds | Vireo_nanus | Vireo_nanus | diurnal |
| birds | Vireo_nelsoni | Vireo_nelsoni | diurnal |
| birds | Vireo_olivaceus | Vireo_olivaceus | diurnal |
| birds | Vireo_osburni | Vireo_osburni | diurnal |
| birds | Vireo_pallens | Vireo_pallens | diurnal |
| birds | Vireo_philadelphicus | Vireo_philadelphicus | diurnal |
| birds | Vireo_plumbeus | Vireo_plumbeus | diurnal |
| birds | Vireo_sclateri | Hylophilus_sclateri | diurnal |
| birds | Vireo_solitarius | Vireo_solitarius | diurnal |
| birds | Vireo_vicinior | Vireo_vicinior | diurnal |

| | | | |
|---|---|---|---|
| birds | Vireolanius_leucotis | Vireolanius_leucotis | diurnal |
| birds | Vireolanius_melitophrys | Vireolanius_melitophrys | diurnal |
| birds | Vireolanius_pulchellus | Vireolanius_pulchellus | diurnal |
| birds | Volatinia_jacarina | Volatinia_jacarina | diurnal |
| birds | Vosea_whitemanensis | Melidectes_whitemanensis | diurnal |
| birds | Vultur_gryphus | Vultur_gryphus | diurnal |
| birds | Wetmorethraupis_sterrhopteron | Wetmorethraupis_sterrhopteron | diurnal |
| birds | Willisornis_poecilinotus | Willisornis_poecilinotus | diurnal |
| birds | Xanthocephalus_xanthocephalus | Xanthocephalus_xanthocephalus | diurnal |
| birds | Xanthomixis_apperti | Bernieria_apperti | diurnal |
| birds | Xanthomixis_cinereiceps | Bernieria_cinereiceps | diurnal |
| birds | Xanthomixis_zosterops | Bernieria_zosterops | diurnal |
| birds | Xanthopsar_flavus | Xanthopsar_flavus | diurnal |
| birds | Xanthotis_flaviventer | Xanthotis_flaviventer | diurnal |
| birds | Xanthotis_macleayanus | Xanthotis_macleayanus | diurnal |
| birds | Xanthotis_polygrammus | Xanthotis_polygrammus | diurnal |
| birds | Xema_sabini | Xema_sabini | diurnal |
| birds | Xenerpestes_minlosi | Xenerpestes_minlosi | diurnal |
| birds | Xenerpestes_singularis | Xenerpestes_singularis | diurnal |
| birds | Xenicus_gilviventris | Xenicus_gilviventris | diurnal |
| birds | Xenodacnis_parina | Xenodacnis_parina | diurnal |
| birds | Xenoligea_montana | Xenoligea_montana | diurnal |
| birds | Xenoperdix_udzungwensis | Xenoperdix_udzungwensis | diurnal |
| birds | Xenopipo_atronitens | Xenopipo_atronitens | diurnal |
| birds | Xenopipo_uniformis | Xenopipo_uniformis | diurnal |
| birds | Xenopirostris_polleni | Xenopirostris_polleni | diurnal |
| birds | Xenopirostris_xenopirostris | Xenopirostris_xenopirostris | diurnal |
| birds | Xenops_minutus | Xenops_minutus | diurnal |
| birds | Xenops_rutilus | Xenops_rutilans | diurnal |
| birds | Xenops_tenuirostris | Xenops_tenuirostris | diurnal |
| birds | Xenopsaris_albinucha | Xenopsaris_albinucha | diurnal |
| birds | Xenornis_setifrons | Xenornis_setifrons | diurnal |
| birds | Xenospingus_concolor | Xenospingus_concolor | diurnal |
| birds | Xenospiza_baileyi | Xenospiza_baileyi | diurnal |

| | | | |
|---|---|---|---|
| birds | Xenotriccus_callizonus | Xenotriccus_callizonus | diurnal |
| birds | Xenotriccus_mexicanus | Xenotriccus_mexicanus | diurnal |
| birds | Xenus_cinereus | Xenus_cinereus | diurnal |
| birds | Xiphidiopicus_percussus | Xiphidiopicus_percussus | diurnal |
| birds | Xiphocolaptes_albicollis | Xiphocolaptes_albicollis | diurnal |
| birds | Xiphocolaptes_falcirostris | Xiphocolaptes_falcirostris | diurnal |
| birds | Xiphocolaptes_major | Xiphocolaptes_major | diurnal |
| birds | Xiphocolaptes_promeropirhynchus | Xiphocolaptes_promeropirhynchus | diurnal |
| birds | Xipholena_atropurpurea | Xipholena_atropurpurea | diurnal |
| birds | Xipholena_lamellipennis | Xipholena_lamellipennis | diurnal |
| birds | Xipholena_punicea | Xipholena_punicea | diurnal |
| birds | Xiphorhynchus_erythropygius | Xiphorhynchus_erythropygius | diurnal |
| birds | Xiphorhynchus_flavigaster | Xiphorhynchus_flavigaster | diurnal |
| birds | Xiphorhynchus_fuscus | Xiphorhynchus_fuscus | diurnal |
| birds | Xiphorhynchus_guttatus | Xiphorhynchus_guttatus | diurnal |
| birds | Xiphorhynchus_lachrymosus | Xiphorhynchus_lachrymosus | diurnal |
| birds | Xiphorhynchus_obsoletus | Xiphorhynchus_obsoletus | diurnal |
| birds | Xiphorhynchus_ocellatus | Xiphorhynchus_ocellatus | diurnal |
| birds | Xiphorhynchus_pardalotus | Xiphorhynchus_pardalotus | diurnal |
| birds | Xiphorhynchus_spixii | Xiphorhynchus_spixii | diurnal |
| birds | Xiphorhynchus_susurrans | Xiphorhynchus_susurrans | diurnal |
| birds | Xiphorhynchus_triangularis | Xiphorhynchus_triangularis | diurnal |
| birds | Xolmis_cinereus | Xolmis_cinereus | diurnal |
| birds | Xolmis_coronatus | Xolmis_coronatus | diurnal |
| birds | Xolmis_dominicanus | Xolmis_dominicanus | diurnal |
| birds | Xolmis_irupero | Xolmis_irupero | diurnal |
| birds | Xolmis_pyrope | Xolmis_pyrope | diurnal |
| birds | Xolmis_salinarum | Xolmis_salinarum | diurnal |
| birds | Yuhina_bakeri | Yuhina_bakeri | diurnal |
| birds | Yuhina_brunneiceps | Yuhina_brunneiceps | diurnal |
| birds | Yuhina_flavicollis | Yuhina_flavicollis | diurnal |
| birds | Yuhina_gularis | Yuhina_gularis | diurnal |
| birds | Yuhina_humilis | Yuhina_humilis | diurnal |
| birds | Yuhina_nigrimenta | Yuhina_nigrimenta | diurnal |

| | | | |
|---|---|---|---|
| birds | Yuhina_occipitalis | Yuhina_occipitalis | diurnal |
| birds | Zanclostomus_javanicus | Phaenicophaeus_javanicus | diurnal |
| birds | Zanda_baudinii | Calyptorhynchus_baudinii | diurnal |
| birds | Zanda_funerea | Calyptorhynchus_funereus | diurnal |
| birds | Zanda_latirostris | Calyptorhynchus_latirostris | diurnal |
| birds | Zapornia_atra | Porzana_atra | diurnal |
| birds | Zapornia_flavirostra | Amaurornis_flavirostra | diurnal |
| birds | Zapornia_parva | Porzana_parva | diurnal |
| birds | Zapornia_paykullii | Porzana_paykullii | nocturnal |
| birds | Zapornia_tabuensis | Porzana_tabuensis | diurnal |
| birds | Zaratornis_stresemanni | Zaratornis_stresemanni | diurnal |
| birds | Zavattariornis_stresemanni | Zavattariornis_stresemanni | diurnal |
| birds | Zebrilus_undulatus | Zebrilus_undulatus | diurnal |
| birds | Zeledonia_coronata | Zeledonia_coronata | diurnal |
| birds | Zenaida_asiatica | Zenaida_asiatica | diurnal |
| birds | Zenaida_auriculata | Zenaida_auriculata | diurnal |
| birds | Zenaida_aurita | Zenaida_aurita | diurnal |
| birds | Zenaida_galapagoensis | Zenaida_galapagoensis | diurnal |
| birds | Zenaida_graysoni | Zenaida_graysoni | diurnal |
| birds | Zenaida_macroura | Zenaida_macroura | diurnal |
| birds | Zenaida_meloda | Zenaida_meloda | diurnal |
| birds | Zentrygon_albifacies | Geotrygon_albifacies | diurnal |
| birds | Zentrygon_carrikeri | Geotrygon_carrikeri | diurnal |
| birds | Zentrygon_chiriquensis | Geotrygon_chiriquensis | diurnal |
| birds | Zentrygon_costaricensis | Geotrygon_costaricensis | diurnal |
| birds | Zentrygon_frenata | Geotrygon_frenata | diurnal |
| birds | Zentrygon_goldmani | Geotrygon_goldmani | diurnal |
| birds | Zentrygon_lawrencii | Geotrygon_lawrencii | diurnal |
| birds | Zentrygon_linearis | Geotrygon_linearis | diurnal |
| birds | Zimmerius_bolivianus | Zimmerius_bolivianus | diurnal |
| birds | Zimmerius_chrysops | Zimmerius_chrysops | diurnal |
| birds | Zimmerius_cinereicapilla | Zimmerius_cinereicapilla | diurnal |
| birds | Zimmerius_gracilipes | Zimmerius_gracilipes | diurnal |
| birds | Zimmerius_improbus | Zimmerius_improbus | diurnal |

| | | | |
|---|---|---|---|
| birds | Zimmerius_vilissimus | Zimmerius_vilissimus | diurnal |
| birds | Zimmerius_viridiflavus | Zimmerius_viridiflavus | diurnal |
| birds | Zonotrichia_albicollis | Zonotrichia_albicollis | diurnal |
| birds | Zonotrichia_atricapilla | Zonotrichia_atricapilla | diurnal |
| birds | Zonotrichia_capensis | Zonotrichia_capensis | diurnal |
| birds | Zonotrichia_leucophrys | Zonotrichia_leucophrys | diurnal |
| birds | Zonotrichia_querula | Zonotrichia_querula | diurnal |
| birds | Zoonavena_thomensis | Zoonavena_thomensis | diurnal |
| birds | Zoothera_andromedae | Zoothera_andromedae | diurnal |
| birds | Zoothera_dixoni | Zoothera_dixoni | diurnal |
| birds | Zoothera_everetti | Zoothera_everetti | diurnal |
| birds | Zoothera_heinei | Zoothera_heinei | diurnal |
| birds | Zoothera_heinrichi | Geomalia_heinrichi | diurnal |
| birds | Zoothera_lunulata | Zoothera_lunulata | diurnal |
| birds | Zoothera_machiki | Zoothera_machiki | diurnal |
| birds | Zoothera_major | Zoothera_major | diurnal |
| birds | Zoothera_margaretae | Zoothera_margaretae | diurnal |
| birds | Zoothera_marginata | Zoothera_marginata | diurnal |
| birds | Zoothera_mollissima | Zoothera_mollissima | diurnal |
| birds | Zoothera_monticola | Zoothera_monticola | diurnal |
| birds | Zosterops_abyssinicus | Zosterops_abyssinicus | diurnal |
| birds | Zosterops_albogularis | Zosterops_albogularis | diurnal |
| birds | Zosterops_anomalus | Zosterops_anomalus | diurnal |
| birds | Zosterops_atricapilla | Zosterops_atricapilla | diurnal |
| birds | Zosterops_atriceps | Zosterops_atriceps | diurnal |
| birds | Zosterops_atrifrons | Zosterops_atrifrons | diurnal |
| birds | Zosterops_borbonicus | Zosterops_borbonicus | diurnal |
| birds | Zosterops_brunneus | Speirops_brunneus | diurnal |
| birds | Zosterops_buruensis | Zosterops_buruensis | diurnal |
| birds | Zosterops_ceylonensis | Zosterops_ceylonensis | diurnal |
| birds | Zosterops_chloris | Zosterops_chloris | diurnal |
| birds | Zosterops_chloronothos | Zosterops_chloronothus | diurnal |
| birds | Zosterops_cinereus | Zosterops_cinereus | diurnal |
| birds | Zosterops_citrinella | Zosterops_citrinella | diurnal |

| | | | |
|---|---|---|---|
| birds | Zosterops_consobrinorum | Zosterops_consobrinorum | diurnal |
| birds | Zosterops_conspicillatus | Zosterops_conspicillatus | diurnal |
| birds | Zosterops_emiliae | Chlorocharis_emiliae | diurnal |
| birds | Zosterops_erythropleurus | Zosterops_erythropleurus | diurnal |
| birds | Zosterops_everetti | Zosterops_everetti | diurnal |
| birds | Zosterops_explorator | Zosterops_explorator | diurnal |
| birds | Zosterops_ficedulinus | Zosterops_ficedulinus | diurnal |
| birds | Zosterops_finschii | Zosterops_finschii | diurnal |
| birds | Zosterops_flavifrons | Zosterops_flavifrons | diurnal |
| birds | Zosterops_fuscicapilla | Zosterops_fuscicapilla | diurnal |
| birds | Zosterops_grayi | Zosterops_grayi | diurnal |
| birds | Zosterops_griseotinctus | Zosterops_griseotinctus | diurnal |
| birds | Zosterops_griseovirescens | Zosterops_griseovirescens | diurnal |
| birds | Zosterops_hypolais | Zosterops_hypolais | diurnal |
| birds | Zosterops_hypoxanthus | Zosterops_hypoxanthus | diurnal |
| birds | Zosterops_inornatus | Zosterops_inornatus | diurnal |
| birds | Zosterops_japonicus | Zosterops_japonicus | diurnal |
| birds | Zosterops_kuehni | Zosterops_kuehni | diurnal |
| birds | Zosterops_kulambangrae | Zosterops_kulambangrae | diurnal |
| birds | Zosterops_lacertosus | Woodfordia_lacertosa | diurnal |
| birds | Zosterops_lateralis | Zosterops_lateralis | diurnal |
| birds | Zosterops_leucophaeus | Speirops_leucophoeus | diurnal |
| birds | Zosterops_lugubris | Speirops_lugubris | diurnal |
| birds | Zosterops_luteirostris | Zosterops_luteirostris | diurnal |
| birds | Zosterops_luteus | Zosterops_luteus | diurnal |
| birds | Zosterops_maderaspatanus | Zosterops_maderaspatanus | diurnal |
| birds | Zosterops_mayottensis | Zosterops_mayottensis | diurnal |
| birds | Zosterops_meeki | Zosterops_meeki | diurnal |
| birds | Zosterops_melanocephalus | Speirops_melanocephalus | diurnal |
| birds | Zosterops_metcalfii | Zosterops_metcalfii | diurnal |
| birds | Zosterops_meyeni | Zosterops_meyeni | diurnal |
| birds | Zosterops_minor | Zosterops_minor | diurnal |
| birds | Zosterops_minutus | Zosterops_minutus | diurnal |
| birds | Zosterops_modestus | Zosterops_modestus | diurnal |

| | | | |
|---|---|---|---|
| birds | Zosterops_mouroniensis | Zosterops_mouroniensis | diurnal |
| birds | Zosterops_murphyi | Zosterops_murphyi | diurnal |
| birds | Zosterops_mysorensis | Zosterops_mysorensis | diurnal |
| birds | Zosterops_natalis | Zosterops_natalis | diurnal |
| birds | Zosterops_nigrorum | Zosterops_nigrorum | diurnal |
| birds | Zosterops_novaeguineae | Zosterops_novaeguineae | diurnal |
| birds | Zosterops_oleagineus | Zosterops_oleagineus | diurnal |
| birds | Zosterops_olivaceus | Zosterops_olivaceus | diurnal |
| birds | Zosterops_pallidus | Zosterops_pallidus | diurnal |
| birds | Zosterops_palpebrosus | Zosterops_palpebrosus | diurnal |
| birds | Zosterops_poliogastrus | Zosterops_poliogastrus | diurnal |
| birds | Zosterops_rendovae | Zosterops_rendovae | diurnal |
| birds | Zosterops_rennellianus | Zosterops_rennellianus | diurnal |
| birds | Zosterops_rotensis | Zosterops_rotensis | diurnal |
| birds | Zosterops_samoensis | Zosterops_samoensis | diurnal |
| birds | Zosterops_sanctaecrucis | Zosterops_sanctaecrucis | diurnal |
| birds | Zosterops_semperi | Zosterops_semperi | diurnal |
| birds | Zosterops_senegalensis | Zosterops_senegalensis | diurnal |
| birds | Zosterops_splendidus | Zosterops_splendidus | diurnal |
| birds | Zosterops_stresemanni | Zosterops_stresemanni | diurnal |
| birds | Zosterops_superciliosus | Woodfordia_superciliosa | diurnal |
| birds | Zosterops_tenuirostris | Zosterops_tenuirostris | diurnal |
| birds | Zosterops_uropygialis | Zosterops_uropygialis | diurnal |
| birds | Zosterops_vellalavella | Zosterops_vellalavella | diurnal |
| birds | Zosterops_xanthochroa | Zosterops_xanthochroa | diurnal |
| birds | Zosterornis_latistriatus | Stachyris_latistriata | diurnal |
| birds | Zosterornis_nigrorum | Stachyris_nigrorum | diurnal |
| birds | Zosterornis_striatus | Stachyris_striata | diurnal |
| birds | Zosterornis_whiteheadi | Stachyris_whiteheadi | diurnal |
| crocodiles | Alligator_mississippiensis | Alligator_mississippiensis | cathemeral |
| crocodiles | Alligator_sinensis | Alligator_sinensis | cathemeral |
| crocodiles | Caiman_crocodilus | Caiman_crocodilus | cathemeral |
| crocodiles | Caiman_latirostris | Caiman_latirostris | cathemeral |
| crocodiles | Caiman_yacare | Caiman_yacare | cathemeral |

| | | | |
|---|---|---|---|
| crocodiles | Crocodylus_acutus | Crocodylus_acutus | cathemeral |
| crocodiles | Crocodylus_intermedius | Crocodylus_intermedius | cathemeral |
| crocodiles | Crocodylus_mindorensis | Crocodylus_mindorensis | cathemeral |
| crocodiles | Crocodylus_moreletii | Crocodylus_moreletii | cathemeral |
| crocodiles | Crocodylus_niloticus | Crocodylus_niloticus | cathemeral |
| crocodiles | Crocodylus_novaeguineae | Crocodylus_novaeguineae | cathemeral |
| crocodiles | Crocodylus_palustris | Crocodylus_palustris | cathemeral |
| crocodiles | Crocodylus_porosus | Crocodylus_porosus | cathemeral |
| crocodiles | Crocodylus_rhombifer | Crocodylus_rhombifer | cathemeral |
| crocodiles | Crocodylus_siamensis | Crocodylus_siamensis | cathemeral |
| crocodiles | Gavialis_gangeticus | Gavialis_gangeticus | cathemeral |
| crocodiles | Mecistops_cataphractus | Mecistops_cataphractus | cathemeral |
| crocodiles | Melanosuchus_niger | Melanosuchus_niger | cathemeral |
| crocodiles | Osteolaemus_tetraspis | Osteolaemus_tetraspis | cathemeral |
| crocodiles | Paleosuchus_palpebrosus | Paleosuchus_palpebrosus | nocturnal |
| crocodiles | Paleosuchus_trigonatus | Paleosuchus_trigonatus | nocturnal |
| crocodiles | Tomistoma_schlegelii | Tomistoma_schlegelii | cathemeral |
